# Haplotype-aware single-cell multiomics uncovers functional effects of somatic structural variation

**DOI:** 10.1101/2021.11.11.468039

**Authors:** Hyobin Jeong, Karen Grimes, Peter-Martin Bruch, Tobias Rausch, Patrick Hasenfeld, Radhakrishnan Sabarinathan, David Porubsky, Sophie A. Herbst, Büşra Erarslan-Uysal, Johann-Christoph Jann, Tobias Marschall, Daniel Nowak, Jean-Pierre Bourquin, Andreas E. Kulozik, Sascha Dietrich, Beat Bornhauser, Ashley D. Sanders, Jan O. Korbel

## Abstract

Somatic structural variants (SVs) are widespread in cancer genomes, however, their impact on tumorigenesis and intra-tumour heterogeneity is incompletely understood, since methods to functionally characterize the broad spectrum of SVs arising in cancerous single-cells are lacking. We present a computational method, scNOVA, that couples SV discovery with nucleosome occupancy analysis by haplotype-resolved single-cell sequencing, to systematically uncover SV effects on *cis*-regulatory elements and gene activity. Application to leukemias and cell lines uncovered SV outcomes at several loci, including dysregulated cancer-related pathways and mono-allelic oncogene expression near SV breakpoints. At the intra-patient level, we identified different yet overlapping subclonal SVs that converge on aberrant Wnt signaling. We also deconvoluted the effects of catastrophic chromosomal rearrangements resulting in oncogenic transcription factor dysregulation. scNOVA directly links SVs to their functional consequences, opening the door for single-cell multiomics of SVs in heterogeneous cell populations.

## Introduction

The mutational landscapes of numerous cancers have recently been catalogued in pan-cancer surveys^1,2^, revealing that ~55% of somatic driver mutations represent SVs, which outnumber base-pair substitutions as cancer drivers^2^. Somatic SVs drive tumor development, progression and therapy resistance, and cause inter- and intra-tumor heterogeneity^3–8^. Somatic DNA rearrangement processes generate a broad spectrum of SV classes, which range from simple SVs such as deletions and inversions to catastrophic rearrangement classes resulting from chromothripsis, chromoplexy and breakage-fusion bridge (BFB) cycles^9–13^. However, with the exception of focal gene deletions and amplifications, the functional effects of SVs are understudied^1,2,6,14,15^. Indeed, the impact of SVs often escapes attention as gene panels, exomes and genomes sequenced with moderate coverage suffer from low SV discovery sensitivity, especially for subclonal SVs^1,2^, which restrains applications in precision oncology. Computational methods integrating bulk-cell RNA-seq and cancer genome sequencing data have been employed to infer the functional outcomes of a wider spectrum of SVs^14–17^. These methods, however, have several limitations: They require generation of large cohorts with paired RNA-seq and genomic data to analyse recurrent SVs, which limits their routine use, or they are restricted to studying SVs that result in allele-specific outlier expression in a tumour sample^14–17^. Furthermore, intra-tumour heterogeneity is not taken into account by these methods, which are therefore essentially restricted to the main clone.

Unambiguous inference that a somatic mutation mediates an effect can be drawn by measuring both genotypes and molecular phenotypes in the same cell, a feat realizable by single-cell multiomics methods^18–22^. These methods allow investigating the functional impact of mutations in genetically heterogeneous contexts, for example, by coupling single-cell DNA and RNA sequencing. Several methods have been developed^18–21^, but all present methods cover only a small subset of the spectrum of SVs arising in cancer: chromosome-arm level changes, and interstitial as well as terminal somatic copy-number alterations (SCNAs) greater than 10 Megabase (Mb) in size, which jointly represent only 37% of SV drivers seen in a typical cancer genome^2^. SCNAs smaller than 10Mb, as well as translocations, balanced inversions, and complex rearrangement classes^9,10,12,23^ – which together account for the majority of driver SVs^2^ – escape detection with these methods.

Here, we describe scNOVA (for <u>s</u>ingle-<u>c</u>ell <u>N</u>ucleosome <u>O</u>ccupancy and Genetic <u>V</u>ariation <u>A</u>nalysis), a method to dissect the full spectrum of somatic SVs ≥200kb in size by inferring their functional conequences using single-cell multiomics principles. scNOVA leverages data from template-strand sequencing (Strand-seq^24^), a haplotype-resolved single-cell technique, in two orthogonal ways: [*i*] it uses the specific DNA fragmentation pattern resulting from Micrococcal nuclease (MNase) digestion of Strand-seq libraries to directly measure nucleosome occupancy (NO) and indirectly infer *cis*-regulatory element (CRE) accessibility^25–27^ in each cell, and [*ii*] it couples this ‘molecular phenotype proxy’ with SV discovery by single-cell tri-channel processing (scTRIP), via joint modeling of read-orientation, read-depth, and haplotype-phase^28^ – in the same individual cell. MNase digests linker DNA between nucleosomes, while nucleosome-protected DNA (or DNA protected by some other strongly bound proteins) remains intact, allowing inference of NO along the genome by interpreting sequence read counts^25–27,29^. Prior work has shown how nucleosome positioning and occupancy reflects gene regulation, with active enhancers exhibiting decreased NO, and with actively transcribed genes showing reduced NO at the transcriptional start and termination sites, as well as reduced nucleosome packing within the gene body^25–27,29–31^. However, the relationships between NO and somatic mutational landscapes in cancer cells are unexplored. scNOVA addresses this by integrating somatic SVs and NO measurements in the same cell – in a haplotype-resolved manner – to dissect the functional impact of somatic SVs in genetically heterogeneous samples. Using scNOVA, we functionally deconvolute a wide variety of SVs in single cells providing insights into their functional outcomes, including complex DNA rearrangements that previously resisted comprehensive characterization.

## Results

### Haplotype-aware NO reveals changes in CREs, cell types, and differential gene activity in Strand-seq data

#### Resemblance of Strand-seq and MNase-based NO tracks

Using Strand-seq, a technique in which non-template strands are labeled during DNA replication using Bromodeoxyuridine (BrdU)^24^, we recently reported comprehensive SV discovery in single cells at 200kb resolution^28^. We next sought to investigate the functional outcomes of these SVs in individual cellular genomes by single-cell multiomics. We hypothesized that NO patterns derived from MNase cuts made during Strand-seq library preparation^24^ would represent a promising readout, allowing to couple SV discovery with a data type equivalent to single-cell MNase-seq data^26^ – to link somatic SVs and NO profiles in the same cell (**Fig. 1a, Fig. S1**). To address our hypothesis, we first assessed the suitability of Strand-seq for revealing nucleosome locations, by comparing Strand-seq to bulk MNase-seq data generated for the NA12878 lymphoblastoid cell line (LCL). Each high-quality Strand-seq library (*N*=95 single cells) was sequenced to a median of 540,379 mapped non-duplicate read pairs^32^, which amounts to ~0.018x coverage per cell (**Table S1**). To directly compare Strand-seq with bulk-cell MNase-seq data^33^, we pooled these Strand-seq data into “pseudo-bulk” tracks, and then measured NO along the genome (**Methods**). We found Strand-seq and MNase-seq were highly concordant in terms of uniformity of coverage and inferred nucleosome positions at DNase-I hypersensitive sites (Spearman’s *r*=0.68) (**Fig. 1b,c**). Measuring NO genome-wide at the binding sites of CTCF^34^ (a key chromatin organizer) revealed a nucleosome-depleted region at the center of the binding site in the Strand-seq track, as previously reported for MNase-seq^29^ (**Fig. S2)**, and nucleosome positioning near CTCF sites^26,29^ closely matched between Strand-seq and MNase-seq (**Fig. 1d**). Nucleosome repeat length^29^ estimates were consistent between independent Strand-seq experiments (195.4 ±0.4bp) and concordant with bulk MNase-seq (193.7 ±0.6bp) (**Fig. S2**), showing these patterns are reproducible. In addition, both assays measured NO in all fifteen chromatin states identified by the Roadmap Epigenome Consortium^35^. Among these chromatin states, Strand-seq and MNase-seq revealed the highest NO signals on average for the polycomb-repressed state and the bivalent enhancer state, whereas the lowest average NO signals were consistently seen for the active transcription start site (TSS) state (**Fig. S3**). This indicates that Strand-seq based NO tracks closely resemble those derived using classical MNase-seq^26,29^, and we thus concluded that Strand-seq data enable direct measurement of NO in single cells.

**Fig. 1.**
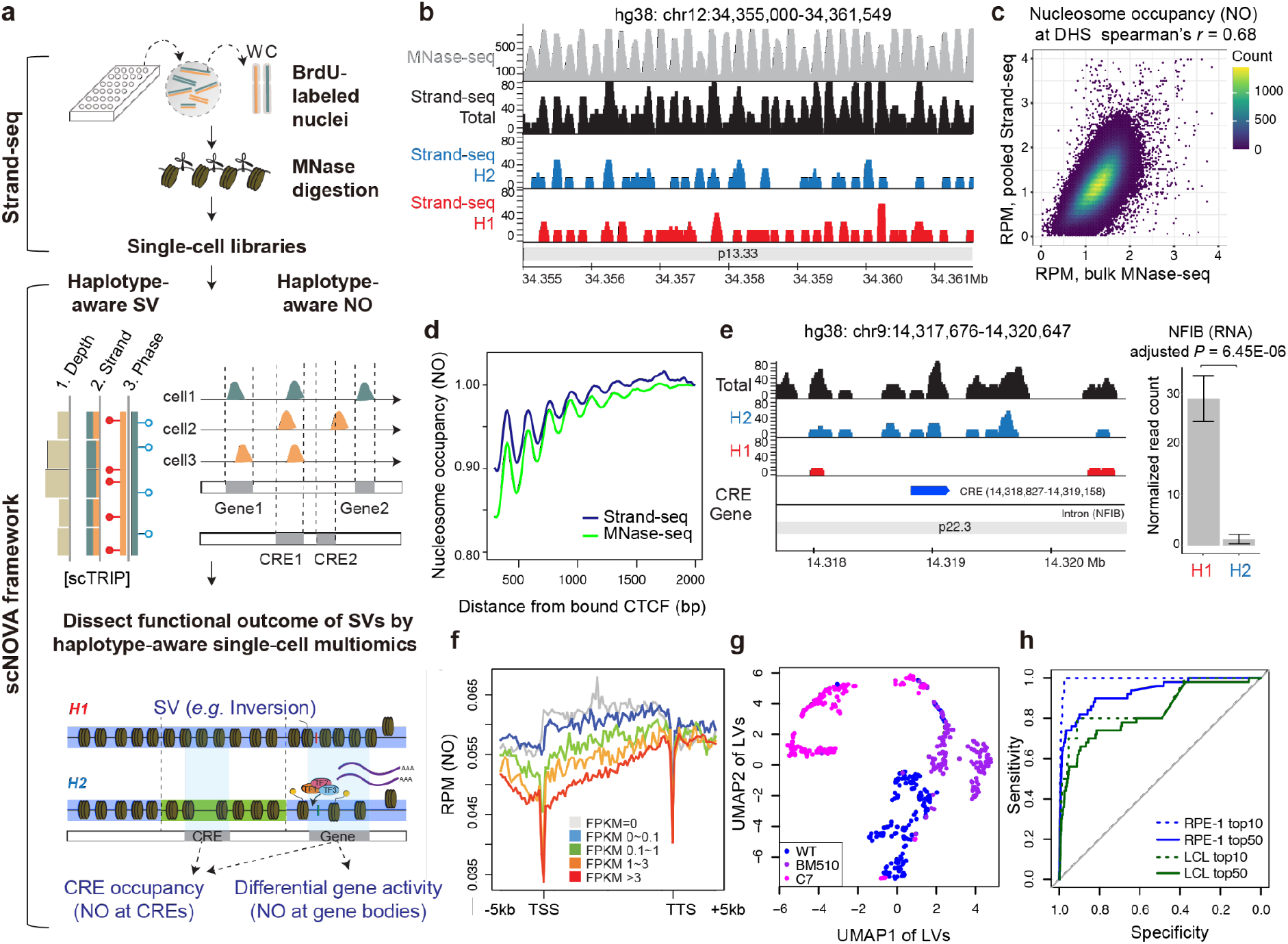
scNOVA: Haplotype-aware single-cell multiomics to functionally characterize SVs. (**a**) Leveraging Strand-seq, the scNOVA framework discovers haplotype-phased SVs by performing single-cell tri-channel processing (scTRIP)^28^, and then generates, in the same single cell, phased nucleosome occupancy (NO) tracks to allow SV functional characterization. Orange: Strand-seq reads mapped to the W (Watson) strand; green: C (Crick) strand. (**b**) Strand-seq based NO tracks in NA12878 reveal nucleosome positions highly concordant with high-coverage bulk MNase-seq, depicted for a chromosome 12 locus with regular nucleosome positioning^113^. Red: NO tracks mapping to haplotype 1 (H1); blue: H2; black: NO measured by combining phased and unphased Strand-seq reads; grey: MNase-seq. The y-axis depicts read counts at each base pair in 10bp bins. (**c**) Correlated NO at consensus DNase I hypersensitive sites^35^, shown for NA12878. (**d**) Averaged nucleosome patterns at bound CTCF binding sites^34^ in NA12878, using pseudo-bulk Strand-seq and bulk MNase-seq. (**e**) Pseudo-bulk haplotype-phased NO tracks based on Strand-seq, depicting a previously defined CRE^36^ in NA12878 with haplotype-specific absence of NO on H1 (10% FDR). Bar chart on the right shows the allele-specific expression on H1 of *NFIB* – the inferred target gene of this CRE. Total: aggregated phased and unphased Strand-seq reads. (**f**) Inverse correlation of NO at gene bodies and bulk RNA-seq expression values. NO is based on pseudo-bulk Strand-seq libraries from RPE-1. RPM: reads per million. TTS: transcription termination site. To facilitate visualisation, gene bodies were scaled to the same length. (**g**) Cell type classification based on NO at gene bodies (AUC=0.96). Cell line codes: Blue: RPE-1. Purple: BM510. Magenta: C7. LV: latent variable. (**h**) Receiver operating characteristics for inferring altered gene activity based on analyzing NO at gene bodies, using pseudo-bulk Strand-seq libraries from in *silico cell* mixing. Ground truth sets of differentially active genes are based on bulk RNA-seq; performance was estimated for the top 10 and top 50 most differentially active genes, using scNOVA. The computed mean AUC was 0.93 for the top 10 genes (0.88 for the top 50 genes).

#### scNOVA reveals haplotype-specific CRE accessibility

Encouraged by these observations, we developed scNOVA, a computational framework that harnesses Strand-seq data to measure NO genome-wide within a cell, and couples this readout with SVs discovered in the same cell to functionally characterize SV landscapes (**Fig. 1a**). Given that Strand-seq data are haplotype-resolved by nature^32^, we hypothesized we could identify haplotype-specific differences in nucleosome occupancy (in short: haplotype-specific NO) – which may, for example, arise as a result of a germline genetic variant or a somatic mutation present on one of the chromosomal homologues. To address this, we performed analyses of haplotype-specific NO at 66,254 CREs previously defined^36^ in NA12878. First, we phased 24,652,658/49,205,197 (50.1%) of the Strand-seq read fragments onto chromosomal haplotypes, and pooled these reads to generate phased pseudo-bulk NO tracks for each of the two haplotypes (denoted ‘H1’ and ‘H2’; **Fig. 1b**). Using these phased NO tracks, we then employed scNOVA to test for haplotype-specific NO within a 1 kilobase (kb) region centred around each CRE (**Fig. S4**). This analysis showed haplotype-specific NO at 727 out of 66,254 CREs (Exact test) when controlling the false discovery rate (FDR) at 10%. These 727 CREs were 1.4–fold enriched on chromosome X (*P*=0.019; hypergeometric test), presumably due to the X-inactivation process^37^ operating in the female-derived NA12878 cell line (**Fig. S5**). We next investigated whether haplotype-specific NO at CREs reflects allele-specific gene expression, by using bulk RNA-seq data^34^ that we phased using the Strand-seq haplotypes as a guide (**Methods**). After assigning putative target genes for each CRE based on the nearest gene approach^38^, genes targeted by CREs with haplotype-specific NO showed a significant proclivity to be allele-specifically expressed (**Fig. S4**; *P*=0.0021; hypergeometric test; 1.5–fold enrichment). A CRE inferred to target the *NFIB* gene, for example, exhibited a clear pattern of haplotype-specific NO: in this CRE, phased NO reads were exclusively assigned to H2, indicating increased CRE accessibility on H1 (**Fig. 1e**; Exact test; 10% FDR), and this was coupled with mono-allelic *NFIB* expression from the H1 haplotype (**Fig. 1e**; FDR-adjusted *P*<2.3E-10; likelihood ratio test). Likewise, a CRE targeting *WWC3*, a gene subject to X-inactivation, demonstrated absence of NO on H2 (10% FDR) – the active chromosome X – coupled with mono-allelic *WWC3* expression from H2 (**Fig. S5**; FDR-adjusted *P*<2.5E-40). These data suggest that scNOVA enables haplotype-phased NO measurements, with haplotype-specific NO patterns at CREs reflecting target gene expression patterns on individual homologs.

#### Cell-type classification by NO profiling

Since NO within gene bodies reflects gene activity^29^, we further hypothesized that we could use NO tracks derived from Strand-seq to infer cell types based on interpreting the activity of lineage-specific genes. In order to address this, we first tested whether Strand-seq based NO profiles at gene bodies reflect global gene expression patterns of a cell. We pooled 33 Million fragments (including phased and unphased reads) from 79 Strand-seq libraries generated for the retinal pigment epithelial-1 (RPE-1) cell line^28^, and then used bulk RNA-seq data for the same line to measure NO at actively expressed versus silent gene bodies. Analysis of these pseudo-bulk NO data revealed a strong inverse correlation between NO at gene bodies and bulk gene expression (*P*<2.2e-16; Spearman’s *r* of up to −0.24; **Fig. 1f**, **Fig. S6**), where highly expressed genes showed lower NO within their gene bodies (and *vice versa)*, consistent with prior MNase-seq based findings^29^. We next explored the utility of NO for cell type classification, by implementing a multivariate dimensionality-reduction framework into scNOVA. We performed *in silico* mixing of Strand-seq libraries generated for different LCLs and RPE cell lines, and built a classifier (**Methods**) that separates distinct cell types by partial least squares discriminant analysis (**Fig. S7**). We first used a training set of 179 mixed libraries, and initially considered 19,629 features, which reflects ENSEMBL^39^ gene bodies with detectable Strand-seq reads to allow for NO measurements. After feature selection, 1,738 features were retained in the final model. We then used a non-overlapping set of 123 single cells to assess performance, all of which scNOVA classified accurately (area under the curve (AUC)=1; **Fig. S7**). We also successfully discriminated between cells from three highly-related RPE cell lines (RPE-1, BM510 and C7), which all originate from the same donor, but which underwent distinct genome instability and selection trajectories^28,40^ (AUC=0.96; **Fig. 1g, Fig. S7**). Thus, scNOVA enables accurate cell type classification based on NO, even amongst closely related cells, indicating that it can be applied to heterogeneous cell samples.

#### Inferring gene activity changes in pseudo-clones

Having confirmed that genome-wide NO measurements can be used to distinguish cell types, we next tested whether NO at gene bodies is also suited to infer changes in the activity of individual genes. Such functionality would be potentially highly relevant for applying scNOVA in cancers that show dysregulated cancer-related gene expression in distinct subclones^17^. Integrating deep convolutional neural networks (CNNs) and negative binomial generalized linear models (**Fig. S8, S9**), we hence devised an scNOVA module to predict gene activity changes between defined cell populations (*i.e*. subclones) using NO tracks. To define the ground truth of gene expression, we used bulk RNA-seq data generated for distinct cell lines. Then, to parameterize scNOVA, we mixed different numbers of Strand-seq libraries from these cell lines *in silico* to create “pseudo-clones”, and performed benchmarking to assess the ability of scNOVA to predict changes in gene activity between pseudo-clones (each composed of cells from a single cell line) by analyzing NO at gene bodies (**Fig. S10, S11**). Indeed, scNOVA’s differential gene activity score (**Methods**) was highly predictive of the 10 most differentially expressed genes, where analyses of pseudo-clones comprising 156 RPE-1 and 46 HG01573 (LCL) libraries revealed an AUC of 0.93 (AUC=0.88 for the 50 most differentially expressed genes; see **Fig. 1h**). Gene activity changes inferred included well-known markers of epithelial (*e.g. EGFR*, *VCAN*) and lymphoid (*e.g. CD74*, *CD100*) cell lineages (**Table S2**). When controlling the FDR at 10%, 10/10 (100%) of the most overexpressed genes in RPE-1, as well as 7/10 (70%) of the most overexpressed genes in HG01573 were correctly captured by scNOVA. scNOVA made informative predictions even when we performed *in silico* cell mixing to simulate minor subclones present with a very low cell fraction (CF) of only 30 cells (CF=20%), 8 cells (CF=5%), and 2 cells (CF=1.3%), respectively, resulting in AUCs of 0.92, 0.79, 0.68. When cell fractions were smaller than 10%, we found that an alternative mode of scNOVA (**Methods)** that uses partial least squares discriminant analysis (PLS-DA) to infer gene activity changes provides slightly improved performances (with AUCs reaching 0.90, 0.82, 0.70 for CFs of 20%, 5%, and 1.3%, respectively; **Fig S11**). We also created pseudo-clones from RPE cells, represented by 156 RPE-1 cells (the original hTERT-immortalised cell line) and 154 C7 cells (which underwent transformation)^40^, where we measured an AUC of 0.73 (**Fig. S10**). Comparing these related lines, scNOVA inferred 615 genes to increase in activity in C7, which included several cancer-related genes (*e.g. CDK1*, *EEF1A2*), and amongst those 615 genes “carcinoma” represented the most enriched functional category (**Table S2**) – in line with the transformed status of C7. We concluded that scNOVA enables accurate inference of gene activity changes based on analyzing Strand-seq-derived NO tracks. Using these inferred gene activity changes as a proxy for molecular phenotypes, together with single-cell SV discovery, we next sought to investigate the functional outcomes of somatic SV landscapes.

### Benchmarking scNOVA: functional outcomes of somatic SV landscapes in lymphoblastoid cell lines

#### Benchmarking in cell lines

Before turning to patient samples, we first benchmarked the coupling of SV calling and NO analysis – in the same individual cell – using cell lines. We used *N*=25 LCLs generated from diverse 1000 Genomes Project (1KGP) samples, which were recently subjected to Strand-seq by the Human Genome Structural Variation Consortium to construct a haplotype-resolved germline SV resource^41^. Single-cell SV discovery in 1,372 Strand-seq libraries (**Table S1**) revealed extensive intra-sample genetic heterogeneity in these widely used cell lines: We detected 205 somatic SVs overall, with 24/25 (96%) LCLs showing at least one somatic SV – a 7-fold increase in the proportion of LCLs exhibiting somatic heterogeneity compared to a prior report^42^ (**Table S3, Supplementary Data**). A significant subset of these somatic SVs were present at high clonality (with >10% CF), affecting 13/25 LCLs. These high-clonality SVs, unexpectedly, included novel subclonal SV in NA12878, arguably the most widely sequenced human cell line^34,43,44^ – which showed a previously not described 500kb somatic 19q13.12 deletion seen with CF=21%, along with two mutually exclusive 22q11.2 somatic deletions (with CFs of 21% and 57%, respectively; **Fig. S12, S13**). The 22q11.2 events map to the known site of IGL recombination, a somatic rearrangement occurring during normal B cell development and thought to have little transcriptional consequences^45^. Thus, we focused on the 19q13.12 event, which led to the loss of a copy of *ZNF382* – a known tumor suppressor and repressor of c-Myc^46^. Application of scNOVA revealed significantly increased activity of *ERCC6*, notably a target of c-Myc/Max dimers^47^, in cells harboring the somatic deletion (10% FDR; **Table S2**).

We sought to validate this finding using orthogonal datasets, by reanalyzing Fluidigm and Smart-seq single-cell RNA-seq (scRNA-seq) data previously generated in NA12878^48,49^. We subjected these datasets to an array of computational tools for SCNA discovery from scRNA-seq data (InferCNV^50^, HoneyBADGER^51^ and CONICSmat^52^), none of which identified SCNAs affecting 19q13.12 or 22q11.2 (**Table S4**). This might not be surprising, as these somatic deletions comprised only 11 and 12 expressed genes, respectively – fewer than the recommended number of genes (*i.e*. 100) for confident SCNA calling using scRNA-seq^52^. We therefore attempted to detect these somatic deletions in a targeted manner (by “genotyping”), via inputting the high-resolution SV breakpoint coordinates from our framework into CONICSmat. We tested different CONICSmat posterior probability cutoffs^52^ to explore SCNA genotyping in the context of a low number of expressed genes (**Fig. S13**). This analysis inferred the presence of the 19q13.12 somatic deletion in NA12878 across a wide range of CONICSmat posterior probability cutoffs (**Fig. S13**). We next performed differential expression analysis between subclones composed of cells with high-confidence deletion genotypes, which revealed that *ERCC6* is over-expressed in cells harboring the 19q13.12 somatic deletion, thus verifying the prediction made by scNOVA (10% FDR, **Fig. S13**). Additionally, cells harboring the 19q13.12 deletion showed several dysregulated signaling pathways compared to cells not exhibiting this event, which included the dysregulation of c-Myc/Max targets and aberrant MAPK signaling (10% FDR, **Fig. S13**). These results suggest that scNOVA is able to leverage SVs inaccessible to scRNA-seq based discovery to dissect the functional effects of these variants – either directly using NO based inference, or by leveraging high-resolution breakpoint assignments from our framework for targeted identification of SCNAs (that would otherwise escape detection) in scRNA-seq data.

#### Complex subclonal rearrangements resulting in dysregulated MAPK signaling

Encouraged by these somatic SV data in NA12878, we next turned our attention to NA20509, the LCL with the most abundant SV subclone (85% CF). SVs discovered in this subclone included complex rearrangements that appeared to affect two chromosomes, providing a relevant use case since Strand-seq enables the identification of subclonal complex and inter-chromosomal SVs inaccessible to discovery by other methods^28^. Analyses of these somatic SVs revealed a 49Mb dispersed duplication on 5q, and a 2.5Mb inverted duplication on 17p with an adjacent terminal deletion on the same haplotype – a signature characteristic of the single cell footprint defined^28^ for BFB rearrangements (**Fig. 2a**). We next pursued a translocation analysis^28^, which revealed that the rearranged 5q and 17p segments were fused into a ~115Mb derivative chromosome (**Fig. S14**), reflecting a common mechanism through which BFBs can be stabilised. Bulk whole genome paired-end sequencing (WGS) validated these subclonal somatic rearrangements in a separate NA20509 cell stock^53^, with ~30% CF (**Fig. S14**). Using scNOVA, we inferred that 18 genes were dysregulated in the BFB subclone (**Fig. 2b**). None of these genes reside in the genomic segments affected by the somatic DNA rearrangements, suggesting that the gene activity changes arose from gene regulatory effects in *trans*. To further characterize the functional consequences of these *trans* effects, we devised an scNOVA module to test for the overrepresentation of functionally-related gene sets^54^, such as transcription factor (TF) targets (**Methods**). Using this module, we observed that the dysregulated genes inferred in the BFB subclone were enriched for the downstream targets of c-Myc and Max (10% FDR, **Fig. 2c**). These two TFs form heterodimers to regulate cell proliferation in the MAPK signaling cascade^55,56^, which is the same pathway we previously associated with the somatic 19q13.12 deletion in NA12878. Indeed, we observed that *MAP2K3*, a MAPK signaling pathway member that activates c-Myc/Max via p38^57,58^, is located in the center of the BFB-mediated inverted duplication on the derivative chromosome (**Fig. 2a**), likely resulting in *MAP2K3* dosage increase. Consistent with these observations, analysis of bulk RNA-seq data available for 24 of these LCLs^41^ showed that NA20509 exhibits the highest *MAP2K3* expression, and the highest c-Myc/Max target expression, amongst the entire LCL panel (**Fig. S15, Fig. 2d**). Additionally, haplotype-resolved RNA-seq analysis in NA20509 (**Methods**) revealed mono-allelic expression of *MAP2K3* restricted to the complex derivative chromosome haplotype (*P*=1.8e-73; FDR-adjusted likelihood ratio test; **Fig. 2e**), indicating that only the somatically rearranged haplotype exhibits *MAP2K3* activity. Collectively, these data orthogonally support the predictions of our scNOVA framework that a complex BFB-mediated rearrangement in NA20509 leads to activation of the c-Myc/Max pathway (model shown in **Fig. 2c**). These findings showcase scNOVA’s ability to predict global gene activity outcomes associated with a subclonal SV, allowing us to now functionally characterize somatic SV landscapes in patient samples.

**Fig. 2.**
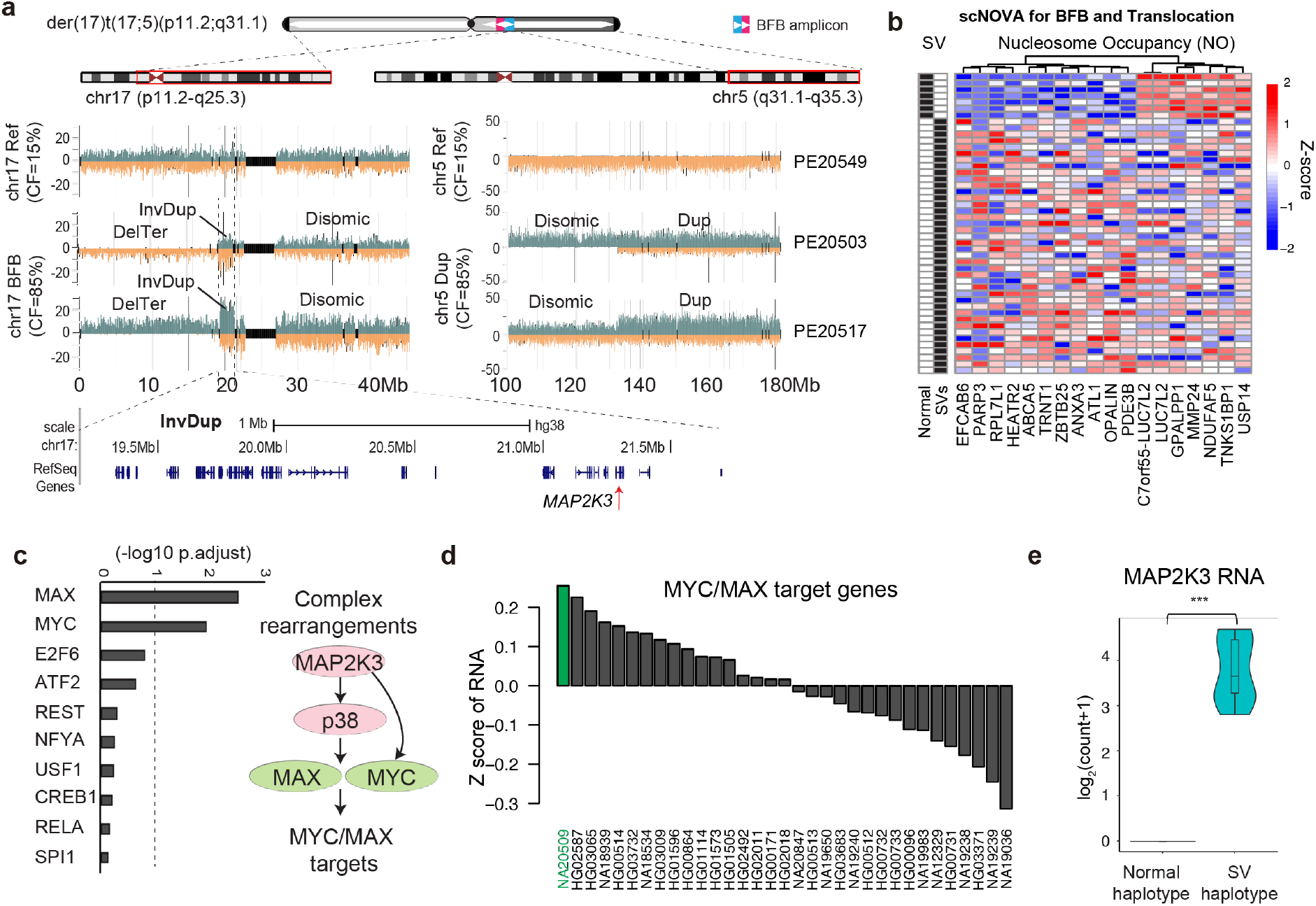
Functional outcome of subclonal SV heterogeneity in lymphoblastoid cell lines (LCLs). (**a**) Complex, BFB-mediated rearrangements in NA20509 (17p) and terminal dispersed duplication (5q) present with CF=85%, here shown for representative single cells. Ref: normal chromosome. InvDup: inverted duplication. DelTer: terminal deletion. Reads denoting somatic SVs, discovered using scTRIP^28^, map to the W (Watson, orange) or C (Crick, green) strand. Grey: single cell IDs. (**b**) Heatmap of 18 genes with inferred altered gene activity amongst subclones, using scNOVA (cells denoted ‘chr5, 17 SVs’ refers to the subclone bearing the complex SVs). (**c**) Gene set overrepresentation analysis for TF target genes showing significant enrichment of c-Myc and Max targets in the subclone bearing complex SVs. Right panel: Model for c-Myc/Max target activation in NA20509 based on scNOVA, combined with prior knowledge. (**d**) Mean RNA expression Z-scores of c-Myc/Max target genes across 33 LCLs. **(e)** Haplotype resolved RNA-seq read counts at heterozygous SNP sites within the *MAP2K3* gene locus, revealing monoallelic expression from the rearranged (BFB) haplotype (FDR-adjusted *P*=1.1e-73; likelihood ratio test).

### Haplotype-specific functional outcomes of somatic DNA rearrangements in leukemia

#### Subclonal SVs in a chronic lymphocytic leukemia (CLL) primary patient sample

Encouraged by this benchmarking exercise, we next applied scNOVA to functionally dissect leukemia samples. We first analysed primary B-cells from a 61-year-old CLL patient (CLL_24). Routine clinical cytogenetics did not show any prototypical genomic alteration^59^ in CLL_24. By contrast, our single-cell analysis of 86 Strand-seq libraries revealed extensive cellular heterogeneity; in this single leukemia sample, we identified 11 different karyotypes represented by 13 SVs occurring at CFs between 1-5% (**Table S3**). While these CF levels are readily amenable to Strand-seq, they are considered to be below the detection limit of WGS^28^. Consequently, our approach detected a higher number of somatic subclones for CLL_24 than for any CLL patient sample subjected to WGS by the International Cancer Genome Consortium (ICGC)^3^. Chromosome 10q24 showed especially pronounced subclonal heterogeneity; here, 7 independent and partially overlapping hemizygous deletions were found, ranging from 2-31Mb in size, and clustering proximal to the chromosome fragile site *FRA10B^60^* (**Fig. 3a, Fig. S16**). Overlap analysis of the deletions revealed a 1.4 Mb ‘minimal segment’ at 10q24.32 that was lost in 11 different cells, and occurred independently on both haplotypes (**Fig. 3b**), confirming somatic 10q24 deletions arose multiple times in this patient. Prior studies reported somatic 10q24 deletions in 1-4% of CLLs^61–63^, which showed enrichment in relapsed/refractory and high-risk cases^64^, suggesting this SV represents a somatic driver. We furthermore verified this deletion in two independent cohorts, where we detected somatic 10q24.32 deletions in 6/306 (2%) CLLs from the ICGC cohort analyzed by SNP arrays^62^ and 4/96 (4%) CLLs from the Pan-Cancer Analysis of Whole Genomes (PCAWG) cohort analyzed by WGS2 (**Fig. 3b, Fig. S19**). It has been proposed that loss of *NFKB2*, present in the minimal segment, may be responsible for the driver phenotype^65^, but a molecular analysis of the consequences of this somatic SV has been lacking. We therefore sought to characterise the functional consequences of 10q24.32 deletions arising in CLL_24.

**Fig. 3.**
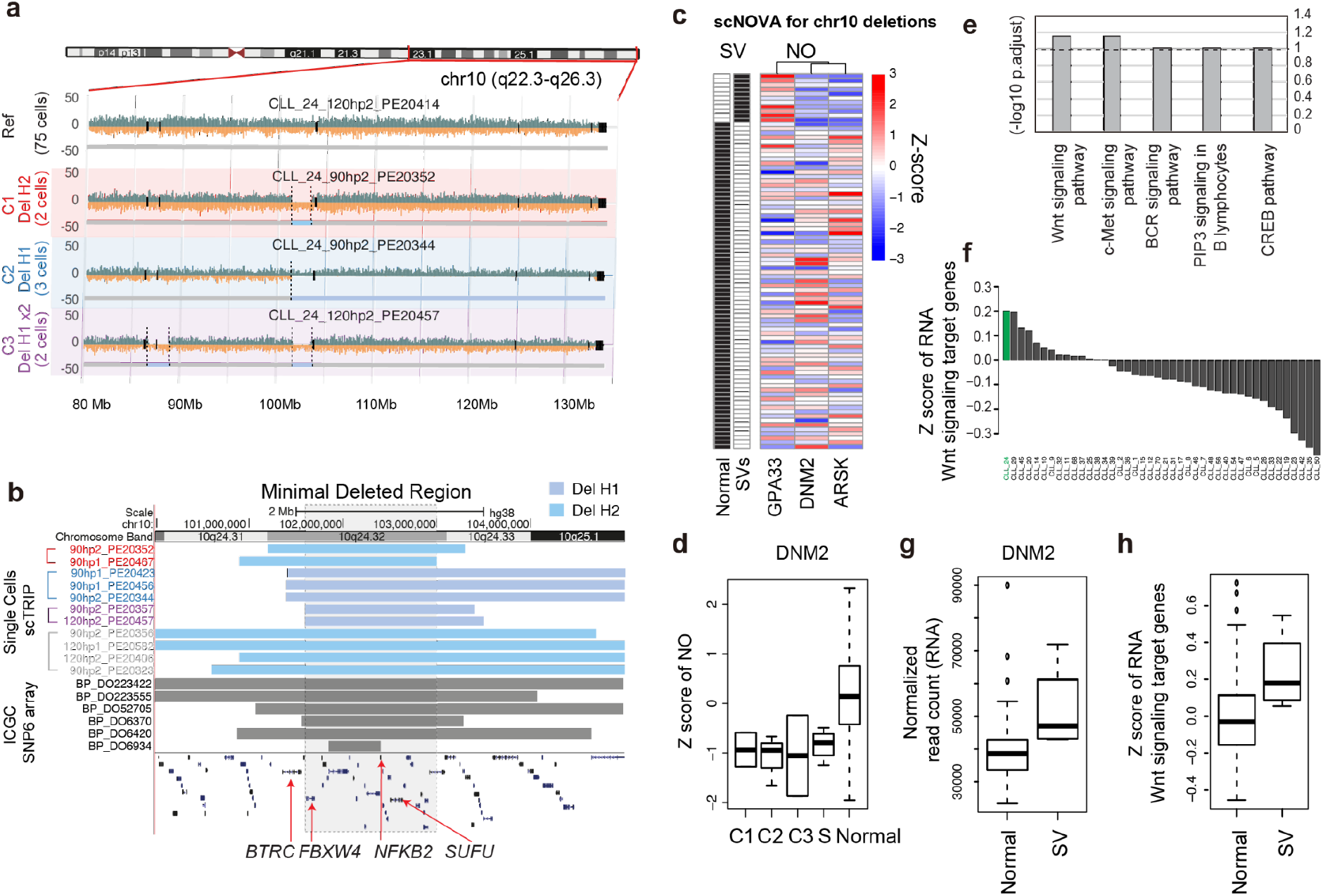
Subclonal SVs in chronic lymphocytic leukemia (CLL) associated with aberrant Wnt signaling. (**a**) Single-cell SV discovery in CLL_24, revealing a diversity of subclonal deletions at chromosome 10q24.32 (all cells exhibiting deletions shown in **Fig. S16**). (**b**) Single cell breakpoint analysis reveals a minimally deleted segment (chr10:101615000-103028000; hg38), which displays recurring deletions in a separate cohort of CLLs (ICGC samples). (**c**) Heatmap of genes altered in activity in cells bearing 10q24.32 deletions, using scNOVA (10% FDR). **(d)** Inference of subclonal gene activity changes in CLL_24 for *DNM2*. ‘Normal’ indicates normal karyotype. C1, C2 and C3 denote subclones harboring deletions (Del) in the ‘minimal region’ at 10q24.32. S combines four single cells that exhibit individual deletions in the same minimal region (seen in *N*=1 cell each). (**e**) Pathway modules with differential activity, in cells exhibiting 10q24.32 deletions versus cells bearing a normal 10q (10% FDR). (**f**) Bulk RNA-seq analysis in 42 CLLs. Mean expression Z-scores for canonical Wnt signaling target genes are shown for each donor (green: CLL_24). (**g-h**) Minimal segment deletion-bearing (SV) CLL samples from the ICGC demonstrate overexpression of *DNM2* and Wnt signaling target genes compared to CLLs not bearing such SV (Normal) (*P*=0.0098; likelihood ratio test).

#### Sub-clonal functional consequences of different 10q24.32 deletions

Having learned that the recurrent 10q24.32 deletion is subclonal in CLL_24 now provides an opportunity to directly compare the phenotypes of cells with and without the deletion and explore the molecular consequences of the somatic SV in (nearly) isogenic subclones. We applied scNOVA and identified three differentially active genes: *GPA33*, *ARSK* and *DNM2*, which we found dysregulated in a joint comparison of the eleven 10q24.32 deletion-bearing cells with the remaining 75 cells (10% FDR; **Fig**. **3c****, Fig. S17**). Notably, all of these genes reside on other chromosomes. For instance, in cells harboring the deletion we observed reduced NO at the *DNM2* locus on chromosome 19, indicating increased activity of Dynamin 2, a GTP-binding protein previously shown to be overexpressed^66^ in other leukemias (**Fig. 3d**). Given these observed *trans* effects, we next performed a molecular phenotyping analysis similar to the NA20509 case, with the aim to measure global expression outcomes associated with the 10q24 deletion. Since three differentially active genes (**Fig**. **3c**) are too few for a gene set overrepresentation analysis, we instead used scNOVA to jointly model NO at gene bodies across gene sets from MSigDB^67^, an approach that is more sensitive since single genes not reaching significance on their own can contribute to the joint model as a group (**Methods**). This analysis showed increased activity of the Wnt signaling pathway in cells carrying a somatic deletion at 10q24.32 (10% FDR; **Fig. 3e**). We also identified c-Met signaling, which is promoted by the Wnt pathway^68^, to be significantly more active in the SV bearing cells (FDR 10%; second rank after Wnt signaling). In support of this prediction, we performed bulk RNA-seq for 42 CLLs, which notably revealed CLL_24 as an outlier demonstrating the highest Wnt signaling target gene expression amongst all CLLs examined (**Fig. 3f**). Moreover, the availability of bulk RNA-seq data for 178 CLL patient samples from ICGC^62^, allowed us to attempt further validation of our predictions. We indeed observed significant overexpression of *DNM2*, as well as significant upregulation of Wnt and c-Met signalling in CLL samples carrying somatic 10q24.32 deletions (10% FDR; CLLs with the deletion: *N*=4; no 10q24.32 SV: *N*=174; **Fig. 3g-h**, **Fig. S20**), which orthogonally supports the predictions made by scNOVA. Aberrant activation of Wnt signaling is crucial for CLL pathogenesis^69^. To date, few genetic lesions have been linked with aberrant Wnt signalling in CLL^69^, and our findings suggest this pathway is associated with recurrent somatic loss of 10q24.32.

While these results suggest the Wnt pathway is activated in cells containing any deletion spanning the “minimal segment”, we also tested the activity of each independent subclone represented by distinct 10q24 somatic deletions (**Fig. S16**). Given the low CF of these subclones (each ≤3.5%), we used the alternative PLS-DA mode of scNOVA to infer gene activity for low frequency subclones represented by at least two cells. This clone-by-clone analysis suggested that the Wnt signaling pathway activation was concentrated within just two of the subclones, with subclone ‘C2’ showing the highest pathway activation and subclone ‘C1’ showing no evidence of Wnt signaling activation (**Fig. S17**). Together, these analyses suggest that scNOVA can uncover sub-clonal functional outcomes of driver SVs, and disentangle the relative contribution of low-abundant subclones to aberrant pathway activity in primary patient samples. Studies to date have not yet identified which genes specifically promote aberrant Wnt pathway activity at 10q24.32, a locus comprising several genes that are known to, or have been suspected to, negatively regulate Wnt signaling (**Fig. 3b, Fig. S17, Supplementary Discussions**). This includes genes directly located in the 10q24.32 minimal segment, such as: *NFKB2* (encoding Nuclear Factor Kappa B Subunit 2), and *FBXW4* (encoding F-box/WD repeat-containing protein 4)^70^, as well as genes very close to the minimal segment, such as: *BTRC* (F-box/WD repeat-containing protein 1A)^71^, which resides just 58kb upstream from the minimal region boundary and was found deleted in 9/11 cells in CLL_24. Notably, when analysing published ICGC transcriptomic data^62^, *NFKB2* did not show significant changes in gene expression (FDR-adjusted *P*=0.507; Wald’s test), whereas both *FBXW4* and *BTRC* were clearly down-regulated in CLL samples exhibiting 10q24.32 deletions (FDR-adjusted *P*-values 0.00478 and 0.000646, respectively); **Fig. S17**). This suggests that *NFKB2* hemizygous loss is unlikely to be responsible for Wnt activation in 10q24.32 deletion bearing cells, whereas *FBXW4* and *BTRC* represent promising targets for future investigation.

#### Haplotype-specific NO patterns in RUNX1-RUNX1T1 driven acute myeloid leukaemia (AML)

Having gained new insight into recurrent SVs in CLL, we next applied scNOVA to a primary AML patient sample (32-year-old male; patient ID=AML_1). We sorted CD34+ primary cells from AML_1 (**Fig. S21**), and produced 42 Strand-seq libraries. SV discovery revealed a 46,XY,t(8;21)(q22;q22) karyotype consistent with clinical diagnostics, with the t(8;21) translocation, detected based on template strand co-segregation^28^, resulting in the well-known *RUNX1-RUNX1T1^72^* gene fusion (**Fig. 4a, Fig. S22**). We pooled the single cells into phased pseudo-bulk NO tracks to map the breakpoints of this SV to intron 1 of *RUNX1T1* and intron 5 of *RUNX1*, consistent with prior reports^73^. We did not detect subclonal heterogeneity in AML_1, indicating a stable karyotype, which precluded the use of scNOVA to assign subclones to dysregulated pathways. Instead, we explored genome-wide patterns of gene dysregulation in AML_1 by analysing haplotype differences in NO at gene bodies. Since cancer-specific alterations are often localized to a single homolog^17^, we reasoned that haplotype-specific NO analysis could be informative to predict cancer-related gene dysregulation (**Methods**). Using the phased pseudo-bulk NO tracks, scNOVA revealed 11 genes with significant haplotype-specific NO at 10% FDR (**Table S2**). This included the *RUNX1T1* gene body, which was more accessible on the derivative (translocation) haplotype in line with its haplotype-specific activity (**Fig. 4b**), suggesting that haplotype-specific NO reflects the aberrant gene activity at this oncogenic locus^73^. The additional genes showing haplotype-specific NO included *ARFGEF1* and *NPR2*, which were previously reported to be dysregulated in cells bearing the *RUNX1*-*RUNX1T1* gene fusion^74^, and CD98 (*SLC3A2*), which promotes AML lethality^75^ and represents a putative therapeutic target (**Fig. 4b, Fig. S23**).

**Fig. 4.**
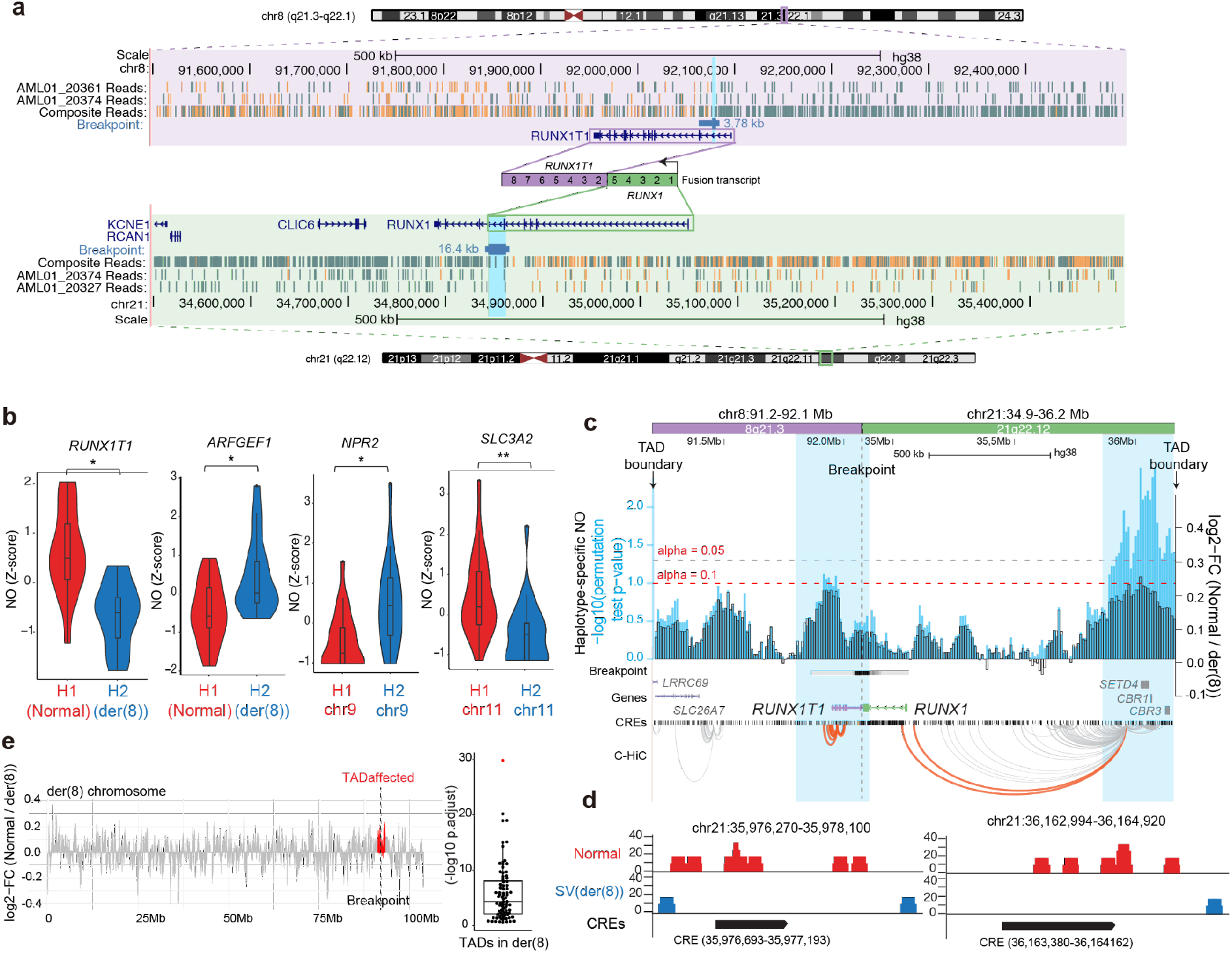
Haplotype-specific NO patterns in *RUNX1-RUNX1T1* driven acute myeloid leukaemia (AML). (**a**) Balanced t(8;21) translocation in AML_1, discovered based on strand co-segregation (*P-*value for translocation discovery using strand co-segregation^28^: *P*=0.00003, FDR-adjusted Fisher’s exact test, **Fig. S22**). SV breakpoints were fine-mapped to the region highlighted in light blue. (**b**) A violin plot demonstrates haplotype-specific NO at the *RUNX1T1* gene body (10% FDR), consistent with aberrant activity of the locus on der(8). Haplotype-specific NO additionally found at the *ARFGEF1*, *NPR2*, and *SLC3A2* was shown next to it (10% FDR). (**c**) Haplotype-specific NO around the SV breakpoint. Fold changes of haplotype-specific NO, measured between the *RUNX1-RUNX1T1* containing derivative chromosome (der(8)) and corresponding regions on the unaffected homologue (Normal), are shown in black, and –log10(*P*-values) in light blue. Enhancer-target gene physical interactions based on chromatin conformation capture^76,114^ are depicted in orange (interactions involving *RUNX1* and *RUNX1T1*) and grey (involving other loci). (**d**) Significant CREs located within the distal peak region, demonstrating haplotype-specific absence of NO on der(8) at 10% FDR, indicative for increased CRE accessibility on der(8). (**e**) Haplotype-specific NO measured between der(8) and corresponding regions of the unaffected homologue. Red: regions corresponding to the fused TAD. A beeswarm plot shows that the fused TAD (red) is an outlier in terms of haplotype-specific NO on der(8) (*P*-values based on KS tests).

These analyses notably suggest the ability of scNOVA to uncover cancer-related changes in gene activity based only on analyzing genome-wide haplotype-specific NO, an approach with potential for discovering cancer drivers and therapy targets in genetically heterogeneous samples.

Next, we considered that fine-grained analysis of haplotype-resolved NO profiles may allow us to dissect the consequences of this balanced translocation on the local *cis*-regulatory landscape. We used the phased pseudo-bulk NO tracks to analyse haplotype-specific NO on either side of the SV breakpoint with scNOVA, by using a sliding 300kb window with a 10kb offset (**Methods)**. This analysis revealed widespread haplotype-specific NO around the SV breakpoint: particularly, we observed broadly decreased NO (suggestive of increased chromatin accessibility) on the SV-affected when compared to the unaffected homolog (**Fig. 4c**). At finer-scale, analysis of haplotype-specific NO showed a pronounced decrease in NO in a large distal region 0.8 to 1.1Mb upstream of *RUNX1* (*P*<0.003; likelihood ratio test, adjusted using permutations; **Fig. 4c**). This region was previously shown to interact physically with the *RUNX1* promoter in normal CD34+ cells^76^. Application of scNOVA revealed two significant CREs in this distal segment showing absence of NO on the derivative chromosome at FDR 10% (Exact test), indicating the CREs are more accessible on the derivative chromosome (**Fig. 4d, Table S5**). The depletion of nucleosomes at these two CREs in physical proximity to the *RUNX1* promoter^76^ suggests that they may foster or support aberrant *RUNX1-RUNX1T1* expression in AML_1. Furthermore, chromosome-wide analyses indicated that the haplotype-imbalanced NO was restricted to the two topologically associating domains (TAD) that were predicted to be fused as a result of the translocation (**Fig. 4e**). These analyses suggest the potential of scNOVA for dissecting changes in local *cis*-regulatory landscapes resulting from somatic driver SVs, and reveal epigenetic remodelling of regulatory regions in association with the t(8;21) translocation in AML_1 likely contributing to aberrant *RUNX1-RUNX1T1* fusion gene expression.

### Haplotype-specific functional outcomes of complex DNA rearrangement processes in leukemia

#### Deconvoluting the outcomes of subclonal chromothripsis with scNOVA

We previously reported a subclonal chromothripsis event with unclear functional outcome in a T-ALL patient-derived xenograft (PDX) sample (TALL-P1)^28^. While catastrophic mutation processes such as chromothripsis play key roles in cancer development, deciphering their functional effects remains an essentially unsolved challenge^9,12,77^. This motivated us to use scNOVA to deconvolute the outcomes of this subclonal DNA rearrangement. TALL-P1, in particular, harbors two subclones^28^: the dominant (CF=70%) subclone lacks complex SVs, whereas the minor subclone (CF=30%) exhibits a chromothripsis event rearranging most of chromosome 6q (depicted in **Fig. 5a**). Analysis using scNOVA identified 12 genes with differential NO between the two subclones (for simplicity, denoted the ‘chromothripsis-associated gene activity signature’ below; 10% FDR; **Fig. 5b; Table S2**). Almost all (10/12 (83%)) of these genes reside on different chromosomes other than chromosome 6, leading us to search for *trans* regulatory effects arising from the chromothripsis event. Closer analysis of the ~90Mb-sized rearrangement showed that 27 TFs reside in the chromothriptic region (**Fig. 5a**). We performed gene-set overrepresentation testing using the target genes of these 27 TFs with scNOVA, which notably revealed only one TF – c-Myb, the product of the *MYB* oncogene – as significantly enriched (FDR-adjusted *P*-value=0.00015; **Fig. 5c**). In fact, half (6/12) of the genes which scNOVA inferred to be altered in activity represent known c-Myb targets (**Fig. 5b-c; Table S6**). 5 out of these 6 genes were inferred to be upregulated in the chromothriptic subclone, which included: *RHOH -* a Rho GTPase frequently overexpressed in T-ALL^78^, *NOTCH1* - a TF and prototypical oncogene frequently exhibiting activating point mutations in T-ALL^79^, and *SLC9A7 (NHE7)* - a membrane protein previously shown to enhance breast tumor formation^80^. The only gene inferred to be downregulated was *PRKCB* - a PKC kinase shown to display tumour suppressive functions in a colon cancer model^81^. Taken together, scNOVA nominates *MYB* as a candidate driver gene dysregulated as a consequence of chromothripsis, by revealing the aberrant activity of several *MYB* target genes.

**Fig. 5.**
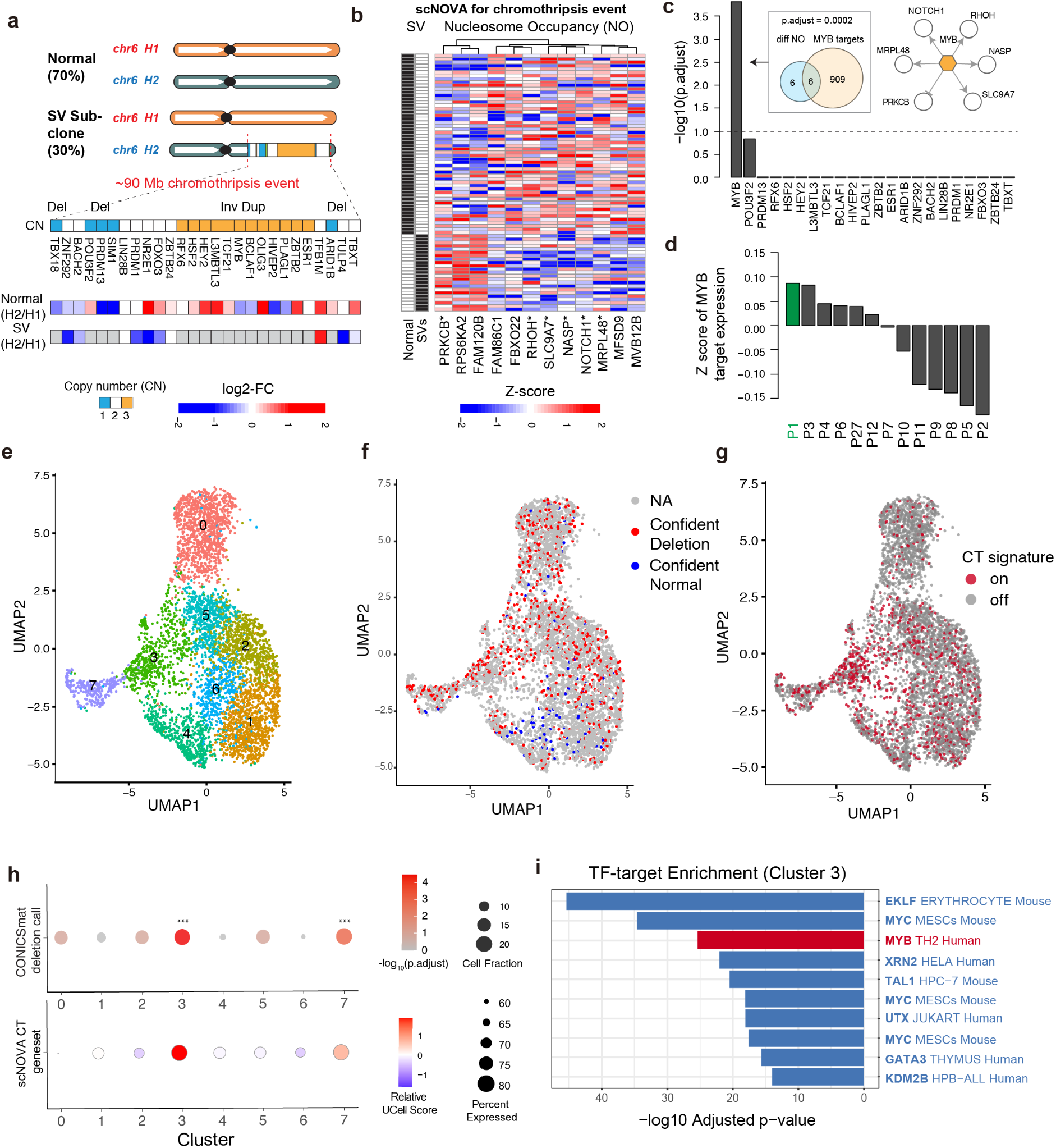
Dissecting a complex karyotype in a T-ALL patient-derived sample using scNOVA. (**a**) 27 TF genes located in a segment that underwent chromothripsis^28^ on 6q in TALL-P1 are depicted along with haplotype-specific NO measurements in each subclone, which scNOVA generated for CREs assigned to the nearest genes. FC: fold-change of normalized haplotype-specific NO (shown for each subclone). Grey: haplotype-specific not measured (for segments deleted or invertedly duplicated). ‘Normal’ denotes the subclone bearing a normal (not rearranged) chromosome 6. (**b**) Heatmap of 12 genes with differential activity between subclones in TALL-P1, based on scNOVA. ‘SVs’: cells bearing the chromothripsis event. ‘Normal’: normal chromosome. Asterisks denote TF targets highlighted in (c). (**c**) TF target enrichment analyses based on the 12 genes from (b), revealing c-Myb as the only significant hit. Venn diagram depicts enrichment of c-Myb targets among the 12 genes (P-value based on an FDR-adjusted hypergeometric test). Upper right: network with c-Myb and its target genes based on scNOVA, combined with prior knowledge. (**d**) Mean Z-scores of c-Myb target gene expression measured by bulk RNA-seq in a panel of 13 T-ALL-derived samples. TALL-P1 (P1) exhibited the overall highest expression of c-Myb targets. (**e**) UMAP of scRNA-seq data showing eight unsupervised clusters in T-ALL-P1. (**f**) Single-cells confidently inferred to exhibit chromosome 6 SCNAs (shown in red) or cells confidently called ‘normal’ (shown in blue) using scRNA-seq data were projected to the UMAP plot. (**g**) Single-cells inferred to be active for the signature genes of the chromothriptic (CT) subclone were shown in red. (**h**) Upper panel: Dot plot shows the significance of over-representation of scRNA-seq based SCNA calls in each of the clusters. Dot color indicates the significance score (-log_10_ adjusted P) from Fisher’s exact test, and the dot sizes denote the cell fraction (CF) of having SCNA calls in each cluster. Cluster 3 and Cluster 7 were significantly overrepresented for scRNA-seq based SCNA calls (*P*-values based on FDR-adjusted Fisher exact tests). Lower panel: Dot plot for the scRNA-seq based gene set level expression summary for the 12 genes from panel (b). Dot color shows the relative UCell score per cluster, and dot size the percent of cells with scorable expression of the geneset. Gene set level expression was derived using UCell^82^ with the directionality of expression changes taken into account. (**i**) Bar graph showing *P*-values for the top 10 significant TFs from the TF-target over-representation analysis of differentially expressed genes for cluster 3. Red color denotes *MYB*, blue color denotes TFs which interact with/are transactivated by *MYB*. (For the corresponding *P*-values in cluster 7, where *MYB* was also significant, see **Table S10**).

#### Orthogonal support from bulk RNA-seq

To corroborate these predictions, we first performed bulk RNA-seq in a panel of 13 T-ALL samples, including TALL-P1. Leveraging the whole-chromosome haplotypes from Strand-seq, we performed allele-specific expression analysis resolved by haplotype in TALL-P1, which revealed aberrant expression of *MYB* from the SV-affected chromosome (*P*=0.0317; likelihood ratio test, 1.4-fold increase in homolog-resolved gene expression; **Fig. S25**) – consistent with *MYB* dysregulation emerging as a direct consequence of chromothripsis. We next compared gene expression measurements for known c-Myb targets (**Table S7**) among all 13 T-ALL samples, which showed that TALL-P1 exhibited the highest expression of c-Myb targets amongst all samples tested, including high outlier expression of *RHOH* (**Fig. 5d, Fig. S26**) – thus further corroborating the inferences made with scNOVA.

#### Verification and refinement of the chromothripsis-associated gene activity signature via scRNA-seq

Second, to provide a more detailed analysis of the chromothripsis subclone we generated an orthogonal scRNA-seq (10X Genomics) dataset comprising 5,504 TALL-P1 cells (**Fig. 5e**). As seen earlier, when applying several current methods for inferring SCNAs from scRNA-seq data^50–52^, we were unable to discover any somatic SVs on chromosome 6 based on these single cell expression data (**Table S4**) – in spite of the fact that the chromotriptic region collectively comprises 227 expressed genes. However, when using the high-resolution breakpoints identified by our framework as an input to CONICSmat, we successfully produced targeted “genotype” calls in a subset of the single cell data. CONICSmat confidently genotyped 15% of the scRNA-seq dataset for which 729 cells were predicted to harbor the chromothripsis rearrangement, and 109 cells were called “normal” [disomic] across the entire chromosome 6 (for the remainder of cells no confident assignment could be made; **Fig. 5f**). Those single cells inferred to contain the chromothripsis event notably showed significant enrichment in two clusters, denoted cluster 3 and cluster 7, which emerged from unsupervised clustering analysis (adjusted *P*=3.43e-5 and 1.15e-3 respectively; FDR-adjusted Fisher’s exact test; **Fig. 5h**) – whereas confident normal cells were enriched in clusters 4 and 6 (**Fig. S28**). We next tested whether the chromothripsis-associated gene activity signature inferred by scNOVA could be used to independently identify cells harbouring the sub-clonal SV. Using UCell^82^, we found both cluster 3 and cluster 7 were significantly enriched for cells comprising high expression scores for the 12 genes contained in this gene activity signature (**Fig. 5h, g**; *P*=3.39e-38 and *P*=2.15e-4, respectively; FDR-adjusted Fisher’s exact test). These complementary analyses therefore suggested that the chromothriptic clone is enriched in scRNA-seq clusters 3 and 7. Notably, differentially expressed genes of both clusters 3 and 7 were significantly enriched with c-Myb target genes (adjusted *P*=4.25e-26, and *P*=1.88e-26, respectively; **Fig. 5i**, **Table S10**), which orthogonally validates the aberrant c-Myb activity initially predicted by scNOVA. In addition to c-Myb, cluster 3 was enriched for *EKLF* and *MYC* targets, which act downstream of c-Myb^83,84^. Finally, we performed trajectory analysis on the scRNA-seq data^85^, which suggested a more primitive state in clusters 3 and 7 compared to other clusters (**Fig. S29**). This is perhaps explained by a differentiation block at the progenitor stage, which has been previously associated with c-Myb hyperactivity in leukemia^86^. The agreement between genotype (a somatic chromothripsis event) and molecular phenotype (gene activity dysregulation of c-Myb targets) in these orthogonal data sets directly supports the ability of scNOVA to infer functional consequences of complex SVs in the same leukemic single cells. Collectively, our results indicate that allele-specific c-Myb dysregulation resulted from a one-off catastrophic rearrangement event in TALL-P1, leading to aberrant expression of c-Myb targets and a more primitive cell state. To our knowledge, this is the first example of how single-cell multiomics can be used to decipher the functional effects of chromothripsis – emphasising the potential of scNOVA to dissect the functional consequences of catastrophic mutational processes in other cancers.

## Discussion

### A novel single cell multiomic framework to functionally characterise somatic SVs

Functional characterization of somatic SVs is of critical importance for the interpretation of cancer genomes because SVs outnumber somatic point mutations as drivers^1,2^. The scNOVA single-cell multiomic framework directly links somatic SVs to their functional consequences by integrating genetic and molecular readouts from the same cell. Using scNOVA we can now: dissect intra-sample genetic heterogeneity at single-cell resolution, measure the haplotype-specific impact of SVs, explore global gene dysregulation in SV-containing cells, discriminate genetically-distinct subclones, and identify unique and shared functional consequences of heterogeneous SVs affecting the same genomic interval. Strand-seq accesses the full spectrum of SV classes ≥200kb in size^28^, which scNOVA now couples to NO measurements to enable their simultaneous functional characterization within the same single-cell sequencing experiment. According to the PCAWG cancer genomic resource, 83% of cancer SV drivers fall within the SV spectrum2 accessible to Strand-seq. This means that scNOVA can interrogate the gene and pathway activity changes resulting from a considerably wider spectrum of SVs compared to prior single-cell multiomics methods, which are largely restricted to arm-level or very large interstitial SCNAs >10Mb in size (see also **Supplementary Discussion**). Despite employing several state-of-the-art methods for inferring SCNA in paired scRNAseq datasets, we were unable to discover the somatic SVs predicted with Strand-seq data when using scRNAseq, unless we first defined the high-resolution breakpoints for targeted SCNA detection. This is consistent with the resolution of our framework being significantly higher compared to the discovery of SCNA based on scRNA-seq alone. On the other hand, using this targeted genotyping approach, we found that Strand-seq and scRNA-seq can be successfully combined to more completely deconvolute patterns of subclonality and intra-tumor heterogeneity. This study altogether analysed 2,178 single-cell genomes and 12,911 single cell transcriptomes (**Table S1, Table S4**), and using these we showed that scNOVA infers changes in gene activity that are generally well-recapitulated by expression data. Our study linked SVs representing copy-imbalanced, copy-neutral and complex rearrangement classes with a functional outcome. Since copy-neutral SVs are captured by our framework, largely euploid (copy-number stable) malignancies previously considered inaccessible^21^ can now be studied by single-cell multinomics using our framework.

### Linking intra-sample genetic complexity to expression dysregulation

Due to the widened spectrum of SVs accessible to our framework, scNOVA enables a more comprehensive dissection of subclonal genetic complexity. We report a >7-fold increase in somatic SVs compared with prior studies in LCLs, and implicate novel somatic SVs in LCLs with gene activity changes. For instance, we found a somatic deletion at 19q13.12 in the widely-used NA12878 cell line, and linked this subclonal SV to *ERCC6* over-expression and dysregulation of cancer-related pathways, such as MAPK signaling and c-Myc/Max target gene activation (**Fig. S13**). We also observed unusual karyotypic diversity in a primary CLL sample, which harbored distinct deletions affecting the 10q24.32 locus. By measuring the subclonal consequences of distinct SV events in the same patient, scNOVA linked this previously described putative driver to aberrant activation of the Wnt signaling pathway. Whether the *FRA10B* fragile site is involved in the formation of these somatic deletions remains to be seen, and will necessitate analyses of larger cohorts. However, we note that CLL_24 exhibits a SNP (rs118137427; 3.7% allele frequency in Europeans) which tags expanded repeat sequences at *FRA10B* that were associated with the acquisition of terminal 10q mosaic deletions in normal blood^87^. Based on the PCAWG resource comprising 94 CLLs^2^, rs118137427 is seen in 2/4 (50%) CLLs with 10q24.32 deletions, but in only 6/90 (6.7%) CLLs with an intact 10q (*P*=0.035; Fisher’s exact test), suggesting a possible link between germline genetic variants at *FRA10B* and somatic SVs in CLL that future studies may explore.

### Functional dissection of complex karyotypes

SVs functionally characterized by scNOVA include those arising from catastrophic rearrangement processes. While complex SV classes have been associated with aggressive tumor phenotypes^2,77^, their functional impact has remained largely unclear, with functional interpretation typically limited to gene loci that are highly amplified in copy-number^2,23,77^. We showcase the high resolution of scNOVA by functionally dissecting a BFB-mediated complex karyotype in NA20509, resulting in monoallelic *MAP2K3* expression and c-Myc/Max target dysregulation. In addition we dissect a subclonal chromothripsis event in a T-ALL-derived sample, directly linking chromothripsis to c-Myb target deregulation by single-cell multiomics, to report a previously undescribed mechanism for *MYB* oncogene activation in T-ALL. Pursuing paired scRNA-seq for the same patient sample enabled us to locate the chromothriptic clone in the clustered expression data, which in turn allowed us to comprehensively characterize the differential gene activity changes and primitive cell state of the somatic subclone. The agreement we found between these orthogonal datasets verified the predictions made by scNOVA, illustrating how we can now comprehensively infer the functional consequences of complex subclonal SVs in the same leukemic single cells. These results highlight significant potential of scNOVA as a framework to study the impact and dynamics of complex rearrangements in cancer.

### Technical considerations and remaining limitations

scNOVA considers three separate genomic readouts: somatic SVs, gene activity, and CRE accessibility around the SV breakpoints – all obtained from only one single-cell assay: Strand-seq. Since our workflow does not require coupling distinct experimental modalities in each individual cell, well-known challenges of single-cell multiomics methods^21^ such as increased costs (*e.g*. from pursuing paired RNA and DNA sequencing from the same cells) and data sparseness (*e.g*. loss of data from any one of the paired experimental modalities^88^) are overcome. One could consider also integrating ATAC-seq into the Strand-seq assay, potentially enabling enhanced analysis of chromatin accessibility; however, this would come at the price of significantly lower resolution of SV calls, given the less uniform coverage of transposase- compared to MNase-mediated cuts^89^. The relatively high coverage achieved by Strand-seq enables the analysis of haplotype-specific NO along whole chromosome homologs (**Fig. S30**), allowing dissection of SV functional outcomes *in cis*. This provides certain advantages over classical allele-specific analyses, which are restricted to phased sequences containing heterozygous SNPs typically reflecting up to 15% of the genome^17^.

Nevertheless, key challenges in single-cell sequencing remain, and the full spectrum of somatic mutations arising in an individual cell is likely to remain inaccessible to studies in the foreseeable future. Because scNOVA harnesses the increasingly used Strand-seq technique^41,90,91^, it captures the whole spectrum of SVs ≥200kb, but misses smaller ones such as mobile element insertions (albeit, this SV class more rarely acts as cancer driver^2^). While revealing differentially active genes, scNOVA does not span the dynamic expression range accessible to RNA-seq nor provides quantitative gene expression measurements of individual genes. Finally, Strand-seq requires dividing cells for BrdU labeling (**Fig. 1a**), and is hence not applicable for non-dividing cells and formalin-fixed samples. However, it can be utilized for dividing cells in cell lines, PDX models, organoids, primary fresh frozen progenitor cells, cells in regenerating tissues, leukaemia cells, and cancer samples amenable to culture. As customary when developing new methods, we used cell lines for benchmarking followed by proof-of-principle application in patient-derived samples. Generalisation of these studies in larger patient cohorts will in the future allow elucidating relevant roles of subclonal SVs in leukaemogenesis.

### Potential future applications: non-genetic determinants of cancer, pre-cancer studies, translational medicine

Technical challenges and limitations notwithstanding, we foresee a wide variety of potential future applications using scNOVA. Our framework offers potential for studies on the determinants and consequences of genomic instability in cancer, and may promote research into the interplay of genetic and non-genetic cancer determinants, including epigenetic plasticity implicated in metastatic potential^21^. It likewise could be used to advance surveys of precancerous lesions, in which roles of SV are understudied^21,92^. Robust determination of SVs in single cells is a known challenge^92^, and scNOVA can increase confidence in single-cell SV discovery by allowing to identify haplotype-resolved SVs and their functional effects in the same cell. This is exemplified by the *RUNX1-RUNX1T1* oncogenic fusion in AML, which in addition to yielding a rearranged DNA sequence results in a discernible pattern of haplotype-specific NO at CREs around the rearrangement breakpoint. In addition, scNOVA might offer future value in precision oncology by exposing subclonal driver alterations along with their targetable functional outcomes, to potentially allow therapeutic targeting of cancer subclones. Furthermore, SVs can accidentally arise in key model cell lines, and scNOVA’s features are ideally suited to functionally characterise unwanted heterogeneity in such samples. As a proof-of-principle, we demonstrated dissection of the outcomes of undesired SVs in widely used LCLs, which includes NA12878, key reference cell line for 1KGP, ENCODE, and NIST^34,43,44^. Indeed, the extent of somatic SVs that our framework uncovered in LCLs indicates that human model cell lines, used for fundamental discoveries in translational medicine and tumour biology, may show more widespread and functionally impactful genetic heterogeneity than currently anticipated. Unwanted somatic SVs also arise as a by-product of CRISPR-Cas9 genome editing, which generates micronuclei and chromosome bridges in human primary cells, structures that initiate the formation of chromothripsis^93^. scNOVA could promote the safety of therapeutically relevant genome editing in the future, by enabling the simultaneous detection and functional characterization of such potentially pathogenic editing outcomes in single cells.

In summary, scNOVA moves directly from SV landscapes to their functional consequences in heterogeneous cell populations. By making a broad spectrum of somatic SVs accessible for functional characterisation genome-wide, this single-cell multiomic framework serves as the foundation for deciphering the impact of somatic rearrangement processes in cancer.

## Acknowledgements

We thank Arnaud Krebs, Judith Zaugg, Karsten Rippe and Isidro Cortés-Ciriano for providing thoughtful feedback on the development of scNOVA. We also thank Malte Paulsen (Flow Cytometry Core Facility) for assistance in cell sorting, Benjamin Raeder for assisting in Strand-seq library preparation, and the EMBL Genomics Core Facility for assisting in single-cell automation (Jürgen Zimmermann and Vladimir Benes) and scRNA-seq library preparation (Laura Villacorta). Finally, we thank Wolfram Höps for assistance with single-cell analysis, as well as Margit Happich and Paulina Richter-Pechanska for assistance with RNA-seq analysis.

## Author contributions

Study design (including conceptualisation of haplotype-specific NO analysis, cell-type classification, and altered gene activity using Strand-seq data): H.J., K.G., A.D.S., J.O.K.; development of scNOVA computational method: H.J., K.G., A.D.S., J.O.K.; single-cell SV discovery; H.J., K.G., A.D.S., J.O.K.; LCL Strand-seq experiments: A.D.S., P.H.; CLL Strand-seq experiments: K.G., P.B., S.D.; AML Strand-seq experiments: K.G., J.-C.J., D.N.; WGS-based somatic SV discovery and verification: T.R.; whole chromosome haplotype-phasing: H.J., D.P., T.M.; LCL RNA-seq analysis: H.J.; LCL scRNA-seq analysis: H.J.; CLL RNA-seq analysis: H.J., S.H., P.B., S.D.; T-ALL RNA-seq analysis: H.J., B.E.-U., A.E.K.; PCAWG SV driver spectrum analysis: R.S., J.O.K.; Joint first authors: H.J., K.G. Joint senior and corresponding authors: A.D.S., J.O.K. The manuscript was written by H.J., K.G., A.D.S. and J.O.K., with additional contributions from all authors.

## Data and materials availability

The code of scNOVA is available open source at https://github.com/jeongdo801/scNOVA, with no restrictions on reuse. LCL data are available under the following accessions: Strand-seq (PRJEB39750); RNA-seq (ERP123231); WGS (PRJEB37677). Leukemia patient data were deposited in the European Genome-phenome Archive (EGA), under the following accession numbers: T-ALL Strand-seq and scRNA-seq (EGAS00001003365), CLL Strand-seq (EGAS00001004925), AML Strand-seq (EGAS00001004903), T-ALL bulk RNA-seq (EGAS00001003248), CLL bulk RNA-seq (accession in progress).

## Methods

### Ethics statement

The protocols used in this study received approval from the relevant institutional review boards and local ethics committees. Written informed consent was obtained from patients, and all experiments were consistent with current bioethical policies. T-ALL experiments were approved by the ethics commission of the Kanton Zurich (approval number 2014-0383).

### Strand-seq in primary leukemia cells

Peripheral blood mononuclear cells of a previously untreated female CLL patient (routine diagnostics: *IGHV* unmutated, no *TP53* mutation, no detected alteration in 6q21, 8q24, 11q22.3, 12q13, 13q14 und 17p13) were isolated after obtaining informed consent. Cells were isolated and cultured using previously established protocols^94^. CLL cells were cultured at 1×10^6^ cells/ml in Roswell Park Memorial Institute (RPMI) medium (Gibco by Life technologies), supplemented with 10 % human serum (PAN BIOTECH), 1 % Pen/Strep (GIBCO by Life Technologies) and 1 % Glutamine (GIBCO by Life Technologies). Cells were stimulated with 1 μg/ml Resiquimod (Enzo) and 50 ng/ml IL-2 (Sigma). BrdU (40 μM; Sigma) was incorporated for 90 h and 120 h, respectively, to perform non-template strand labeling. Single nuclei from each timepoint were sorted into 96-well plates using a BD FACSMelody cell sorter, followed by Strand-seq library preparation (described below).

In the case of the AML sample, frozen primary mononuclear cells from a bone marrow aspirate were thawed and stained with CD34-APC (clone 581; Biolegend), CD38-PeCy7 (clone HB7; eBioscience), CD45Ra-FITC (clone HI100; eBioscience), CD90-PE (clone 5E10; eBioscience), and LIVE/DEAD™ Fixable Near-IR Dead Cell Stain (Thermofisher). Single, viable, CD34+ cells (**Fig. S21**) were sorted using a BD FACSAria™ Fusion Cell Sorter into ice-cold Serum-Free Expansion Medium (SFEM) supplemented with 100 ng/ml SCF and Flt3 (Stem Cell Technologies), 20 ng/ml IL-3, IL-6, G-CSF and TPO (Stem Cell Technologies). Cells were plated in Corning Costar Ultra-Low Attachment 96-well flat-bottom plates (Sigma) at 1×10^5^ cells/ml in warm medium as above. 24 h after culture, 40 μM BrdU was added. Nuclei were isolated after 43 h total culture time, and BrdU-incorporating nuclei sorted into 96-well plates followed by Strand-seq library preparation.

Strand-seq libraries were prepared using a Biomek FXP liquid handling robotic system, as described previously^24,95^. Libraries were sequenced on an Illumina NextSeq500 sequencing platform (MID-mode, 75 bp paired-end sequencing protocol) to a median depth of 551,831 (CLL) and 371,159 (AML) mapped non-duplicate fragments per cell, respectively. Strand-seq data generated with this protocol^24^, involving MNase digestion, largely comprise mononucleosome-sized (140-180bp) fragments (**Table S1**).

### Strand-seq data preprocessing and library selection in leukemia-derived single cells

Reads from Strand-seq (fastq) libraries were aligned to the hg38 assembly using bwa^96^, as previously described^28^. Sequence reads with low quality (MAPQ<10), supplementary reads, and duplicated reads were removed. Single cell library selection was performed as described previously^28^. The single-cell footprints of different SV classes were discovered using the principle of single cell tri-channel processing (scTRIP) of Strand-seq data, using the MosaiCatcher computational pipeline with default settings^28^, as described in the scNOVA workflow further below. In the AML sample, a reciprocal translocation involving chromosomes 8 and 21 resulting in a *RUNX1-RUNX1T1* fusion gene was inferred based on template strand co-segregation^28^, using a MosaiCatcher FDR-adjusted *P*-value of 0.000026 (event seen in 37/37 cells; 5 cells with sister chromatid exchange (SCEs) affecting the respective translocation partners were excluded from the FDR-adjusted *P*-value computation using the MosaiCatcher pipeline^28^). As we identified extensive subclonal heterogeneity in the CLL sample, we used the ‘lenient SV calling parameterization’ available using the MosaiCatcher pipeline^28^ to allow for sensitive detection of SVs at CFs from 1 to 5%, including in individual cells.

### scNOVA workflow: coupling NO measurements and SV discovery in the same cell

We developed scNOVA as a computational tool for coupling discovered somatic SVs with analyses of NO profiles – in the same cell. The scNOVA workflow covers a set of different operations from single-cell SV discovery (using the previously described scTRIP method^28^) to NO profiling at CREs, and gene as well as pathway dysregulation inference based on NO at gene bodies, and can be used in a haplotype-aware or -unaware manner. To maximize reusability, interoperability and reproducibility we combined all of scNOVA’s modules into a coherent workflow using snakemake. Alternatively these modules can be executed individually.

#### Nucleosome occupancy (NO): data analysis and operational definition utilised

We operationally defined nucleosome occupancy (NO) closely following definitions from a prior study^29^: NO maps were calculated by counting how many reads from the Strand-seq libraries (which typically comprise mono-nucleosomal fragments ~140-180 base pairs in size; see **Table S1**, **Fig. S2**) covered a given base pair based on aligning reads to the GRCh38 (hg38) genome assembly with BWA^96^. Genomic regions with unusual (such as artificially high) coverage were considered artifacts, and were automatically excluded (“blacklisted”) by our Strand-seq analysis workflow as previously described^28^. No further peak calling or smoothing was conducted, and no assumptions on the length of the nucleosomal DNA were made to derive NO maps, as nucleosome boundaries were determined on both sides of the nucleosome by paired-end sequencing^29^. For the calculation of NO around bound CTCF binding sites (downloaded from ENCODE^34^), the averaged profile was scaled^29^ to yield an NO equal to 1 at position −2000bp from the center of the bound CTCF site.

#### Cell type classification

To initially assess the utility of Strand-seq derived NO patterns for cell type classification, we generated feature sets from the NO at the body of genes (defined as the region from the TSS to the transcription termination site (TTS), which includes exons and introns) at the single-cell level. NO in gene-body regions was normalized by segmental copy number status, and by library size to obtain reads per million (RPM), which we transformed into log2 scale. This feature set was used for the unsupervised dimension reduction plot (**Fig. S7**) and for training of a supervised classification model based on partial least squares discriminant analysis (PLS-DA)^97^. To train the supervised cell type classifier based on gene-body NO, we used 179 single-cell Strand-seq libraries generated from two diploid LCLs (50 cells from HG02018, 50 cells from NA19036), as well as from a replicate of the near-diploid RPE-1 cell line (79 cells; ‘replicate 1’ ^28^). 19,629 ENSEMBL genes with at least one read detected at the respective gene-bodies were considered as initial input feature sets. An X matrix [179 cells-by-features] and a Y matrix [179 cells-by-two cell type] were prepared for PLS-DA, in order to find latent variables which can explain the variability in Y using linear combinations of features in the X matrix. Using three latent variables which explain 98.48% of the variance of the Y variable, variable importance of projection (VIP) for each feature was calculated. Highly informative features (VIP>90% of null distribution) were selected to build the final classifier. Classification performance of the final classifier was evaluated by leave-one-out cross validation using the 179 cells, which yielded 100% accuracy of classification (are under the curve (AUC)=1) with six latent variables. Finally, we applied this model to an independent validation using a different LCL (46 cells from HG01573) and the same RPE-1 epithelial cell line, albeit using a different replicate (77 cells; ‘replicate 2’), verifying 100% classification accuracy with independently generated data (AUC=1). To generate the UMAP shown in **Fig. S7**, 302 cells including those from the training set (179 cells) and the independent validation set (123 cells) were projected onto the final classification model. The resulting prediction score of six latent variables for each cell was used to perform UMAP^98^ for dimensionality reduction.

#### Haplotype-phasing of single-cell NO tracks

As previously described, Strand-seq directly resolves its underlying sequence reads onto haplotypes ranging from telomere to telomere^32^ (chromosome-length haplotyping). scNOVA phases NO profiles onto a chromosomal homolog using the StrandPhaseR algorithm^32^, which is employed wherever the template strand segregation pattern of a chromosome enables unambiguous haplotype-phasing – that is, for Watson/Crick (WC) or Crick/Watson (CW) template state configurations in Strand-seq libraries^32,95^. Haplotype-specific analyses pursued by scNOVA employ phased reads (normalized by locus copy-number), whereas the inference of gene activity changes uses both phased reads (from chromosomes with a WC or CW configuration) and unphased reads (from chromosomes with a CC or WW configuration^32,95^).

#### Inference of haplotype-specific NO at cis-regulatory elements (CREs)

To allow inference of haplotype-specific NO at CREs, scNOVA requires the location of CREs as another input in bed format (REs_hg38.bed). CREs can be defined using DNase I hypersensitive sites (DHSs), or alternatively based on accessible chromatin segments obtained using ATAC-seq^89^. As a default functionality, scNOVA considers DHSs provided from 127 epigenomes^35^ from the Roadmap Epigenomics Consortium and ENCODE (covering a variety of human tissue types). scNOVA also provides the option to use CREs from user-defined DNA accessibility profiling experiments (provided in ‘bed’ format). After aggregating haplotype-phased single-cell NO tracks into pseudo-bulk tracks, NO is measured based on assessing the read depth at defined CREs, using haplotype-resolved reads. Haplotype-specific NO is measured using the Exact Test followed by controlling the False Discovery Rate^99^ (FDR), using EdgeR software^100^. CREs are assigned to their likely target genes using a nearest gene approach using the prioritisation rules described in^38^.

#### Inference of genome-wide changes in gene activity

This haplotype-unaware module of scNOVA considers all reads – whether they are phased or not – to infer gene activity alterations via analysis of differential patterns of NO along gene bodies. scNOVA obtains gene loci from ENSEMBL (GRCh38.81), converted into bed format (Genebody_hg38.81.bed). Strand-seq reads falling within the start and end position of genes (Genebody_hg38.81.bed) were identified with the Deeptool multiBamSummary function^101^, using the following parameters: multiBamSummary BED-file --BED Genebody_hg38.81.bed --bamfiles Input.bam --extendReads -- outRawCounts output.tab -out output.npz scNOVA’s gene dysregulation inference module contains two steps. *Step 1* filters out genes unlikely to be expressed (‘not expressed’, NEs). *Step 2* infers dysregulated (*i.e*. differentially expressed) genes between subclones using a generalized linear model.

In *Step 1*, scNOVA first aims to infer gene expression ‘On’ and ‘Off’ states^102^ from NO, by analysing NO as well as gene context-specific sequence features along gene bodies using deep convolutional neural networks^103^ (CNNs).

Both phased and unphased single-cell reads were used to generate NO profiles. Feature sets were incorporated into one-dimensional CNNs. To define the feature sets for each gene, we considered genomic regions spanning the body of genes, which we – for the purpose of the CNN – expanded from 5kb upstream of the TSS until 5kb downstream of the TTS, to include 5kb of flanking non-transcribed sequences on each flank which appeared informative as well (**Fig. 1f**). Each gene was divided into 150 bins, whereby we considered genomic annotations of the start and end coordinates of genes provided via ENSEMBL’s GTF file, as follows: 50 bins for the region −5kb to the TSS, 50 bins for the gene-body, and 50 bins for the region from the TTS to +5kb. Five layers of feature sets were derived for those 150 bins: NO, single cell variance of NO, GC content, CpG content, and replication timing, which we included in the CNNs to assist bin stratification. To compute NO for *Step 1*, the read depth within each of the 150 bins was first normalized by bin length using smooth spline fitting in R, and then normalized by library size to obtain read per million (RPM) measurements, which subsequently were scaled by the locus copy-number status. To compute the single cell variance of NO, coefficients of variation (CV = standard deviation / mean) for the single-cell read depth were calculated for each bin. Systematic effects of mean on the CV were regressed out using smooth spline fitting. GC and CpG content were computed based on the nucleotide content of each bin, using the Homer annotate peak tool^104^. For replication timing, we used a pre-processed signal track (hg19) from the UCSC Genome Browser database^105^ (UW Repli-seq track), which we mapped to the hg38 genome using the UCSC LiftOver tool. scNOVA can also consider CNNs for inferring expression in a single cell, which use four rather than five layers of feature sets (excluding single-cell variance of NO, which cannot be computed for a single cell).

To define ground-truth labels of not expressed genes (NEs) and expressed genes (EGs), we used bulk RNA-seq data from three RPE cell lines (RPE-1, BM510, and C7)^28^. Reads were aligned onto hg38 with STAR aligner (v2.5.3)^106^, using gene annotations from ENSEMBL GTF (GRCh38.81). FPKM values were obtained with Alfred^107^; genes with FPKM>1 were labeled as EGs, and all remaining as NEs. We used the following numbers of EGs and NEs for training: RPE-1: 10413 EGs, 9131 NEs; BM510: 10339 EGs, 9205 NEs; C7: 10486 EGs, 9058 NEs. We used the hyperopt package^108^ to search for optimal hyperparameters for the CNNs (**Fig. S8**).

In leave-one-chromosome-out cross-validation^103^ experiments (where we trained a model leaving out a certain chromosome, and then applied the model to the chromosome previously left out), the CNNs outperformed random forest and SVM based models with the same set of features (**Fig. S9**). To assess model performance for different number of aggregated cells (clones of different sizes), we pooled Strand-seq data to generate randomized pseudo-bulk datasets for 80, 40, 20, 5, and 1 cell (s), respectively, and evaluated CNN performance using leave-one-chromosome out cross-validation. Trained models for each chromosome, and for different clone set sizes, are made available along with the code of scNOVA to facilitate application to new data sets. By default, scNOVA operates with the model trained with a pseudo-bulk of 80 cells, to estimate the probability of each gene to represent an NE in each clone. Genes likely to be unexpressed (NE status probability≥0.9) across clones are filtered out in *Step 1*, and all remaining genes used in *Step 2*.

In *Step 2*, scNOVA by default employs negative binomial generalized linear models, available in the DESeq2 algorithm^109^, to infer genes with differential activity between individual cells or clones. As an input, scNOVA computes single-cell count tables of gene-body NO. When running this step with subclones, all individual cells of the subclone are considered ‘replicates’ in DESeq2 terminology^109^. Subclones (or cells) are compared in a pairwise manner using a two-sided Wald test to infer genome-wide alterations in gene activity. Based on this, we defined the differential gene activity score as the sign of the fold change in NO at gene bodies, multiplied by -log10 p-values. Genes with significantly altered activity were identified using a 10% FDR threshold. Additionally, to facilitate the analysis of small CF subclones, scNOVA provides an alternative mode which employs partial least squares discriminant analysis (PLS-DA)^97^ to identify discriminatory feature sets as gene sets showing altered activity. To do this, scNOVA builds a PLS-DA^97^ discriminant model to classify cells in a given subclone 1 and subclone 2 based on single-cell count tables of gene-body NO as feature sets. This model provides a variable importance of projection (VIP) and significance compared to a null distribution in the form of a *P*-value for each gene analyzed. Similar to the default setting, genes with altered activity were identified using a 10% FDR cutoff when using PLS-DA for inferring changes in gene activity between subclones. Benchmarking both modes (see **Fig. S11)** suggested that whereas both DESeq2 and PLS-DA offer acceptable performance, the alternative mode (PLS-DA) outperforms the default setting when the subclonal CF is below 10%, whereas the default mode (DESeq2) generated superior results for CF values of 10% or greater.

#### Molecular phenotype analysis in gene-sets

This module of scNOVA uses defined gene-sets, obtained from public resources, to identify overrepresented sets of functionally related genes changing in activity between subclones (or individual cells). Two types of analyses are enabled by this module: (1) gene-set overrepresentation analysis, which, for example, can be used to investigate the enrichment of targets of a major transcription factor (TF) among genes showing a change in activity according to gene-body analysis of NO; (2) joint modeling of NO across predefined gene-sets, using pathway definitions from MSigDB^67^.

In the case of gene-set overrepresentation analysis, we collected TF target genes from database entries (EnrichR^54^) as well as by reviewing the literature. When reviewing the literature, we created curated lists of target genes for TFs based on published genome-wide studies using the following strict criteria: (*i*) target genes show evidence of binding of the TF of interest by ChIP-seq; (*ii*) the same genes must additionally show differential expression when the TF of interest is experimentally silenced (our curated target gene lists are available in **Table S7**). For each TF, the significance of overlap between its target gene set and genes exhibiting differential NO was computed using hypergeometric tests, followed by controlling the FDR at 10%.

To jointly model differential NO across all genes of predefined pathways, scNOVA first generates a single-cell gene-body NO table using Strand-seq read count data, with these read counts then being normalized using the median-of-ratios method from DESeq2^109^. For each member in the biological pathway gene sets from MSigDB^67^, scNOVA then computes mean normalized NO values, in each single-cell, as a proxy for pathway-level NO. Lowly variable genes (standard deviation <80%) are removed. Pathway-level NO is compared between cells with and without SVs using linear mixed model fitting followed by likelihood ratio testing, and controlling the FDR at 10%. For linear mixed model fitting, SV status is defined as a fixed effect and different Strand-seq library batches are defined as random effects, by scNOVA.

#### Inference of haplotype-specific NO at the body of genes

To measure haplotype-specific NO at gene bodies, scNOVA measures read depth along gene bodies using haplotype-phased single-cell NO tracks. For each gene, gene-body NO measurements from both haplotypes are converted into log2-scale and compared using a generalized linear model likelihood ratio test. To infer haplotype-specific gene deregulation in an AML_1 based on NO, we first filtered out genes inferred to be unexpressed (NE status probability≥0.9) using scNOVA’s CNN. For the remaining genes we computed gene-body NO resolved by haplotype. For *RUNX1* and *RUNX1T1*, we computed haplotype-aware NO for the partial gene-body regions participating in the gene fusion (**Fig. S23**). For each gene, single-cell gene-body NO from two haplotypes was converted into log2-scale and compared using a generalized linear model likelihood ratio test, controlled using an FDR of 10%.

#### Haplotype-resolved SV discovery in single cells

The scNOVA computational framework utilizes the previously described scTRIP method for haplotype-aware SV discovery of the full spectrum of somatic SVs ≥200kb in size in Strand-seq data, by executing the MosaiCatcher computational pipeline^28^. In brief, this pipeline integrates three ‘channels’ – template strand, read depth and haplotype-phase – to discover deletions, duplications, balanced inversions, inverted duplications, balanced translocations, unbalanced translocations and diverse classes of complex SV including BFBs and chromothripsis events in single cells, and it maps these SVs to a defined chromosomal homolog. All single cells are subjected to SV discovery, regardless of chromosomal template strand configuration^28^ (such as Watson/Crick (WC), Crick/Crick (CC), or Watson/Watson (WW)), and joint modeling of the data is pursued which increases the detection sensitivity for SVs present in more than one single cell^28^. By default, scNOVA employs the ‘strict’ scTRIP SV caller, which has been optimized for detecting SVs with CF≥5% ^28^. SV discovery can be bypassed in the scNOVA framework, to focus downstream functional investigation to user-defined somatic SVs.

#### Single-cell RNA sequencing and data processing

Primary human T-ALL cells were recovered from cryopreserved bone marrow aspirates of patients enrolled in the ALL-BFM 2009 study. Patient-derived xenografts (PDX) were generated as previously described by intrafemoral injection of 1 Million viable primary ALL cells in NSG mice^110^ PDX-derived (P1)^28^ cells were frozen until processing. For scRNA-seq library preparation, cryopreserved cells were thawed rapidly at 37 °C and resuspended in 10 ml warm Roswell Park Memorial Institute (RPMI) medium with 100 μg/ml Dnase I. Cells were centrifuged for 5 mins at 300 *g*, and resuspended in ice-cold phosphate buffered saline (PBS) with 2% foetal bovine serum (FBS) and 5mM EDTA. Cells were stained on ice with anti-murine-CD45-PE (mCD45)(clone 30-F11; BioLegend; 1:20) in the dark for 30 mins. 1:100 DAPI was added and incubated in the dark for 5 mins before sorting. Triple negative cells (DAPI-mCD45-GFP-) were sorted (Fig. S27) using a BD FACSAria™ Fusion Cell Sorter into ice cold 0.03% bovine serum albumin (BSA) in PBS. All isolated cells were immediately used for scRNA-seq libraries, which were generated as per the standard 10x Genomics Chromium 3′ (v.3.1 Chemistry) protocol. Completed libraries were sequenced on a NextSeq5000 sequencer (HIGH-mode, 75 bp paired-end).

Sequenced transcripts were aligned to both human and mouse genomes (GRCh38 and mm10) and quantified into count matrices using Cellranger mkfastq and count workflows (10X Genomics, V 3.1.0, default parameters). The R package Seurat^111^ (V 4.0.3) was used for QC of single cells and unsupervised clustering of the data. Briefly, human cells were separated from multiplets/mouse contamination based on >97 % of their reads aligning to GRCh38. Further filtering for high quality cells accepted only those with >200 but <20,000 total RNA counts and a percentage of mitochondrial reads <10%. Finally, remaining mouse transcripts were removed prior to further analysis. Normalisation, scaling and regression of mitochondrial read percentage was carried out using the scTransform package^112^. Dimensionality reduction and differential expression analysis of identified clusters was performed using Seurat.

#### Single-cell gene signature scoring using UCell

The activity of the scNOVA-identified gene set from TALL-P1 in scRNA-seq data was profiled using the UCell package ^82^. Briefly, signature genes considered were those with either increased (implying decreased expression) or decreased (implying increased expression) nucleosome occupancy (see Fig. 5b), or genes encoding TFs whose targets showed differential nucleosome occupancy (see Fig. 5c). The following gene set was used for T-ALL-P1: “PRKCB-”, “RPS6KA2-”, “FAM120B-”, “FAM86C1+”, “FBXO22+”, “RHOH+”, “SLC9A7+”, “NASP+”, “NOTCH1+”, “MRPL48+”, “MFSD9+”, “MVB12B+”, “MYB+” (with “+” for upregulated, and “−” for downregulated). The score per single cell for the entire directional gene set was calculated using the AddModuleScore_UCell() function. Cells were considered to be ‘active’ for the signature genes if their respective UCell score was greater than or equal to the median UCell score of the entire dataset, plus the standard deviation.

