## Supplementary Materials for "Haplotype-aware single-cell multiomics uncovers functional effects of somatic structural variation"

**Index:**

This Supplement Information is divided into Figures, Tables, Data, Supplementary Notes for Methodological Details, Supplementary Discussion and References.


**Supplementary Figures**

**Supplementary Tables**

**Supplementary Data**

**Supplementary Notes for Methodological Details**

*1*. Estimating genome-wide coverage

*2*. Analysis and comparison of NO profiles derived from Strand-seq and MNase-seq

*3*. Visualization of NO at gene bodies for genes stratified by their expression level

*4*. Analysis of previously reported scMNase-seq data

*5*. Identifying optimal parameters for inferring changes in gene activity using NO

*6*. NO-based inference of altered gene activity by scNOVA

*7.* Molecular phenotype analysis in gene-sets in a cell lines and leukemia samples

*8.* Analysis of haplotype-specific chromatin accessibility near rearrangement breakpoints in an AML patient

*9.* Analysis of haplotype-specific chromatin accessibility near rearrangement breakpoints in T-ALL P1

*10.* Bulk-cell RNA-seq data processing and allele-specific expression analysis

*11*. Bulk RNA-seq analysis in thirteen T-ALL patient-derived samples

*12.* Bulk RNA-seq analysis in 42 CLLs

*13*. Haplotype-resolved bulk RNA-seq analysis in LCLs from HGSVC consortium

*14*. Clinical diagnostic information for CLL_24

*15*. Clinical diagnostic information for AML_1

*16*. 10q deletion discovery in CLL samples from PCAWG

*17*. Strand-seq in a panel of lymphoblastoid cell lines (LCLs)

*18*. WGS-based subclonal SV analysis in NA20509

*19*. Manual curation of somatic SVs in LCLs to achieve a high-quality callset

*20*. scRNA-seq data analysis for inferring somatic copy-number alterations (SCNAs)

*21*. Pseudotime/cell-type analysis of scRNA-seq data

**Supplementary Discussion**

*I*. Scope of scNOVA compared to single-cell multiomics methods focusing on SCNAs

*II*. Known and suspected Wnt signaling regulators near 10q24.32

*III*. Nucleosome repeat length measurements: considerations for future users

*IV*. Further details of functional outcomes of somatic rearrangement landscapes in lymphoblastoid cell lines

*V*. Regulatory landscape changes mediated by a 14q32 inversion in T-cell acute lymphoblastic leukemia (T-ALL)

### Supplementary Figures

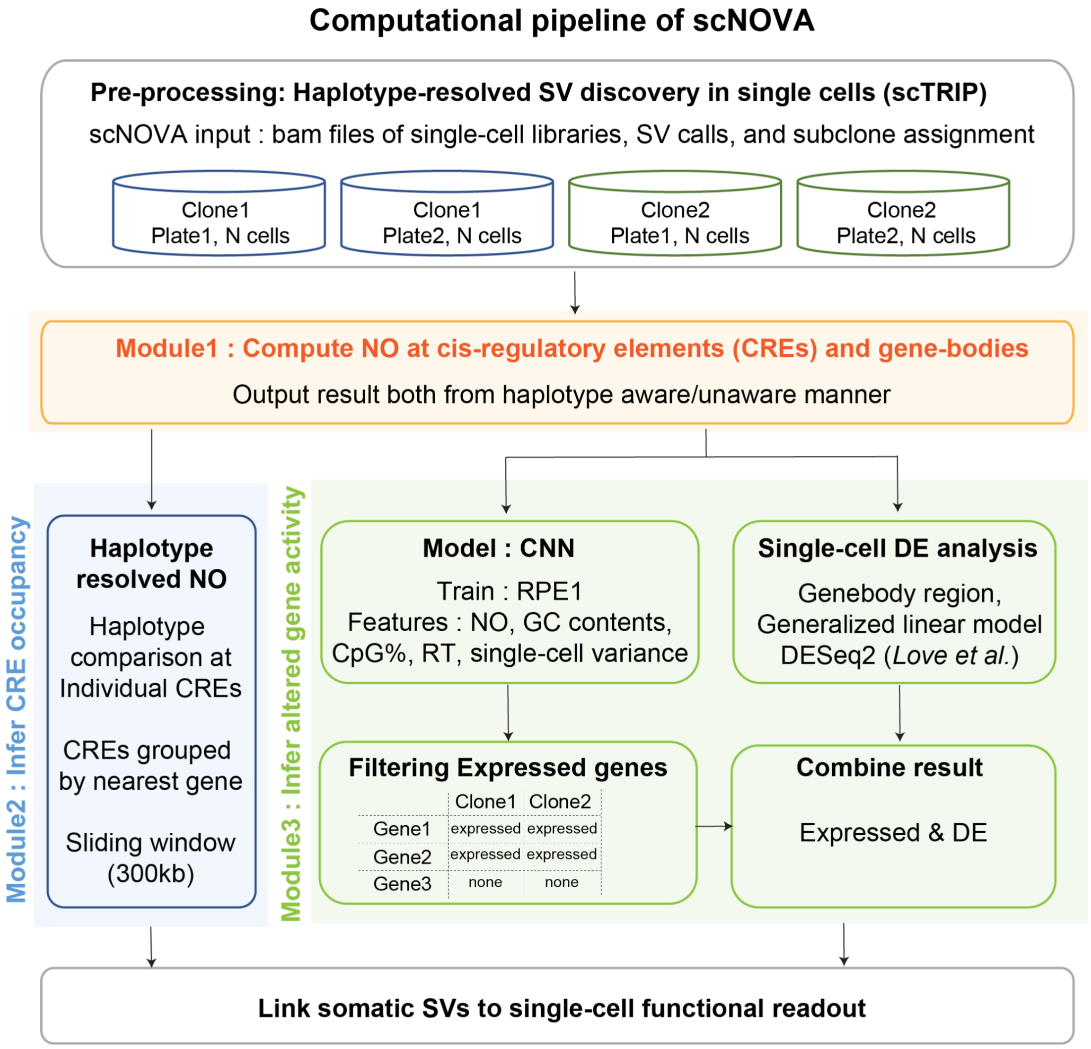


**Figure S1. Overview of components of the scNOVA computational workflow.** scNOVA employs single cell tri-channel processing (scTRIP) as realised in the MosaiCatcher pipeline to perform haplotype-aware somatic SV discovery[1]. Modules of scNOVA enable single-cell mulitomics of these somatic SVs, including investigation of CRE occupancy in *cis*, and inference of altered gene/pathway activity *in trans*. To infer alterations in gene activity, scNOVA integrates deep convolutional neural network (CNN) based machine learning, and negative binomial generalized linear models. The framework dissects intra-sample genetic heterogeneity at single-cell resolution, measures the haplotype-specific impact of somatic SVs, can be used to explore global gene dysregulation in SV-containing cells, can discriminate between genetically-distinct subclones, and can uncover shared functional consequences of heterogeneous SVs affecting the same chromosomal interval.

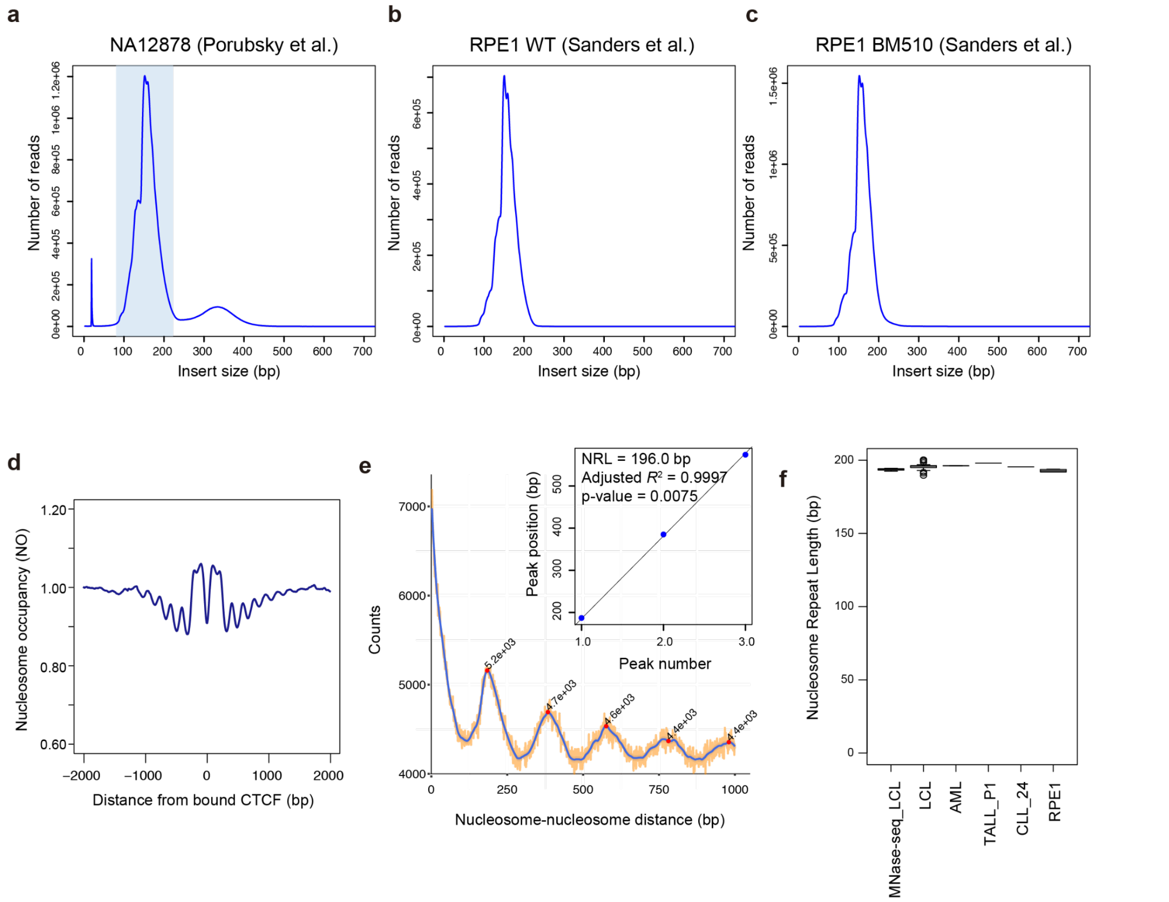


**Figure S2. Strand-seq reflects the characteristics of MNase-seq profiles. (a-c)** Fragment size distribution calculated from the distance between paired-end read alignment positions represent nucleosomal fragments in Strand-seq data from multiple independent experiments. **(a)** Strand-seq libraries from NA12878[2] show a bimodal read length (insert size) distribution implying existence of both mono-nucleosomal and di-nucleosomal sized fragments (see **Supplementary Discussion III**). NA12878 (also known as GM12878, or CEPH1463) represents a widely used human reference and benchmarking sample. **(b)** By comparison, Strand-seq libraries for RPE-1 (here denoted RPE1 WT) and **(c)** the RPE-1 derived BM510 cell line were generated using stringent size selection[1] at 250-350bp (a size representing mononucleosome fragments that still include the sequencing adapters). These data are unimodal reflecting the specific enrichment of mono-nucleosomal sized fragments. Strand-seq libraries newly generated in this study (**Table S1**) follow the strict size selection in [1], and select the 250-350bp (unimodal) sized fragments. For all down-stream analyses we performed an *in-silico* size selection of NA12878 for fragments between 80-220 base pairs in size (after trimming adapter sequences) to enrich for mono-nucleosomal sized fragments (indicated by the first peak in blue background), to be consistent with other Strand-seq libraries investigated in this study. **(d)** Genome-wide averaged nucleosome patterns at CTCF binding sites, based on pooled Strand-seq libraries generated for NA12878. CTCF binding sites for NA12878 were obtained from ENCODE[3]. **(e)** Representative histogram of distances between nucleosomes calculated from pooled Strand-seq HG00096 (N=69 single cell libraries) using NucTools[4]. Plot for chromosome 19 shown as an example. Peak positions represent the distances between the nearest-neighbor[5], followed by the 2nd, 3rd, etc. Inlet figure shows the scatter plot of peak position versus the peak number considering 1st~3rd neighboring nucleosomes, using the peaks in the histogram in (d). Nucleosome repeat length[5] was estimated based on slope values derived using linear regression. **(f)** Nucleosome repeat length[5] estimates for pooled Strand-seq libraries shown for different cell types (LCL, AML, T-ALL (TALL_P1), CLL (CLL_24), and RPE-1 (RPE1)). Publicly available bulk-cell MNase-seq data from LCLs[6] are shown for comparison. MNase-seq data were downsampled to 70 million fragments per experiment to make them as comparable as possible to the pooled Strand-seq data.

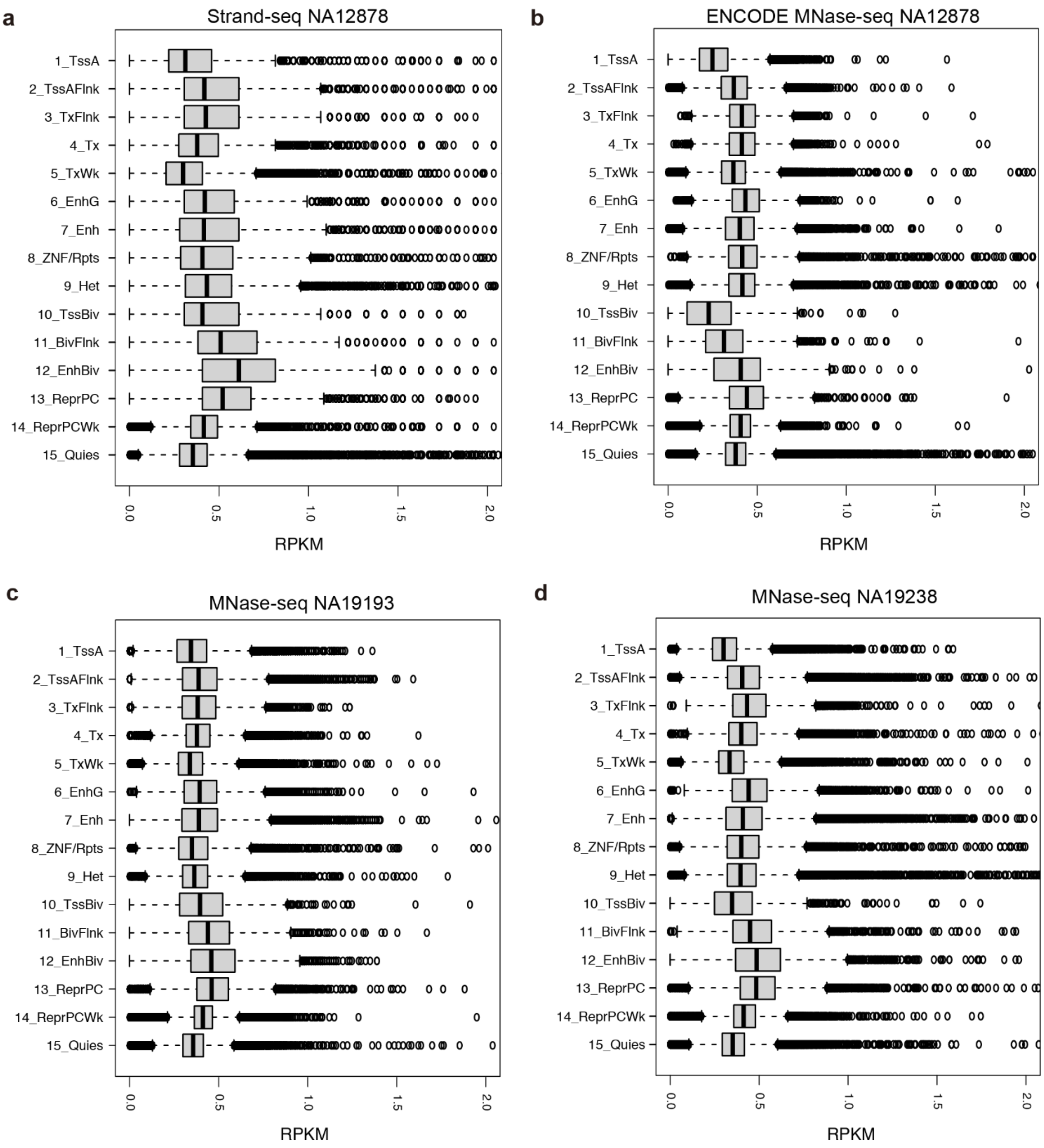


**Figure S3. Read depth measurement of Strand-seq and bulk MNase-seq data sets on genome-wide bins stratified into 15 chromatin states defined by Roadmap epigenome consortium**[7]. 15 chromatin states based on the NA12878 cell line were utilized in this analysis. Plots generated represent Strand-seq data from NA12878 **(a)**, and publicly available bulk MNase-seq from NA12878, NA19193, and NA19238 **(b-d)**. The bulk MNase-seq experiment of NA12878 was done using single-end SOLID sequencing reads, and that of NA19193 and NA19238 was done using paired-end Illumina sequencing reads. X-axis in the box plot indicates reads per kilobase per million (RPKM) measured for each genomic segment annotated by one of the 15 chromatin states. Abbreviations for chromatin states[7] are: TssA-Active TSS, TssAFlnk-Flanking Active TSS, TxFlnk - Transcription at gene 5'and 3', Tx - Strong transcription, TxWk - Weak transcription, EnhG - Genic enhancers, Enh - Enhancers, ZNV/Rpts - ZNF genes & repeats, Het - Heterochromatin, TssBiv - Bivalent/Poised TSS, BivFlnk - Flanking Bivalent TSS/Enh, EnhBiv - Bivalent Enhancer, ReprPC - Repressed PolyComb, ReprPCWk - Weak Repressed PolyComb, Quies - Quiescent/Low.

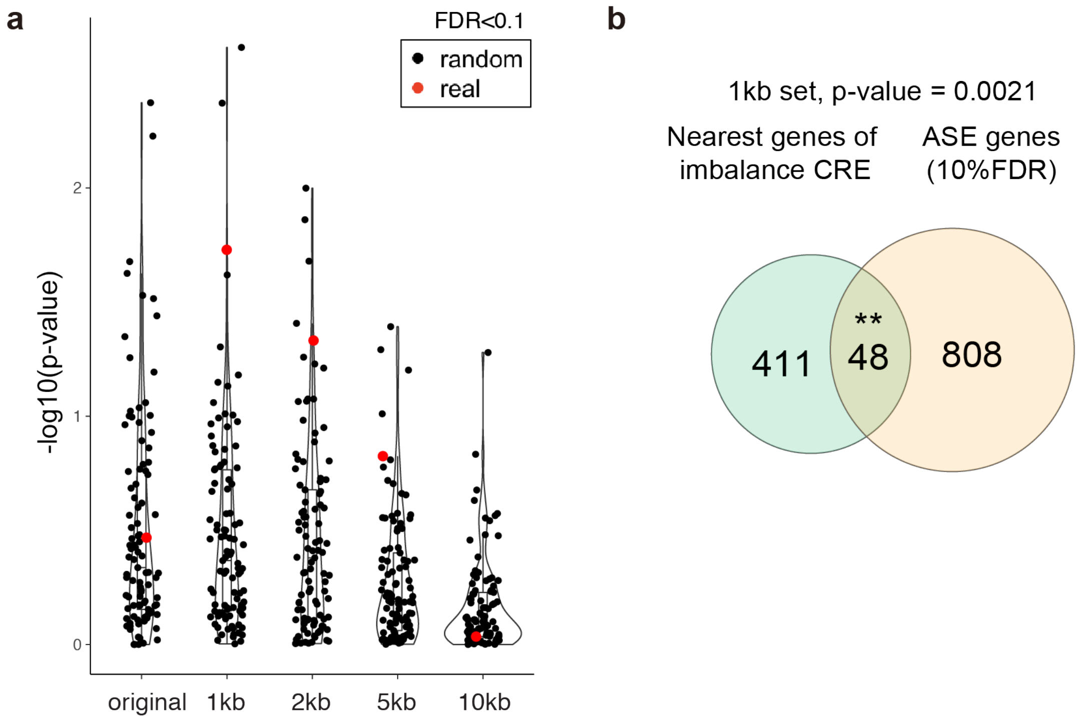


**Figure S4. NA12878 haplotype-phased NO tracks reveal haplotype-specific NO.** We extracted the genomic positions of 66,254 active CREs previously defined in NA12878 using bulk ATAC-STARR-seq[8], to parameterize the identification of haplotype-specific NO at CREs (*i.e.* scNOVA's 'Infer CRE occupancy' module). Because the average annotated CRE length was only 350bp ('original'), we expanded the search space around each CRE to examine whether this improved the ability of scNOVA to discern CREs displaying haplotype-specific NO in sparse single-cell data. We tested five sets of data, by extending CREs to 1kb, 2kb, 5kb, and 10kb intervals, each centered at the original CRE midpoint, and comparing these four sets to the originally defined[8] set of CREs. For each set, we measured haplotype-imbalances in the pseudo-bulk NO tracks generated from NA12878 Strand-seq data, using a 10% FDR cutoff. **(a)** Violin plot representing the enrichment score (-log10 p-value from hypergeometric test) of CREs with haplotype-specific NO on the inactive chromosome X. To confirm the enrichment signal is driven by the regulatory elements, we performed a randomization test by shifting the previously defined CRE locations[8] (original length preserved) +-50kb and repeating the haplotype-imbalance test for each CRE set (i.e. 'original', 1kb, 2kb, 5kb, and 10kb). Given the well-known X-inactivation process, we expect haplotype-specific NO at CREs to be enriched on the X chromosome. The chromosome X enrichment scores of 100 randomization trials are depicted as black dots in each violin plot, and the score calculated for the original (unshifted) CRE is depicted as a red dot. This randomization test shows that haplotype-specific NO at CREs is best resolved when using CREs of a 1kb length (achieved by accordingly extending previously defined[8] CRE locations). **(b)** Venn diagram showing significant overlap between target genes of CREs displaying haplotype-specific NO and genes showing allele-specific expression (ASE) in NA12878. ASE genes were defined by phasing bulk RNA-seq data generated for NA12878 using Strand-seq based haplotype information, followed by examination of haplotype-specific read counts with EdgeR[9], using a 10% FDR. The enrichment *P*-value depicted was estimated using a hypergeometric test ($P$<0.0021).

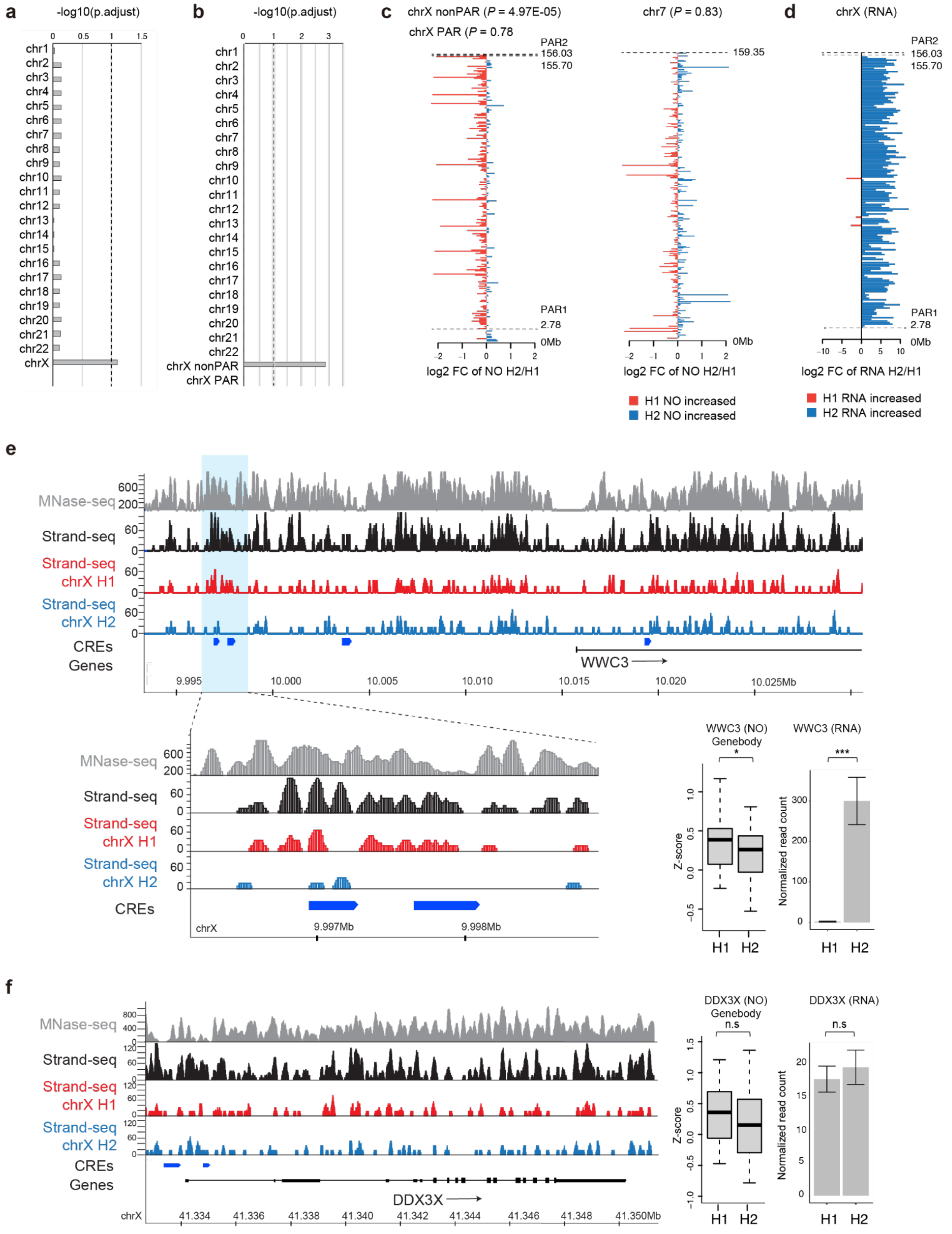

**Figure S5. NA12878 haplotype-phased NO tracks, computed for CREs and gene bodies at the level of full-length chromosomes, are consistent with patterns of X-inactivation. (a)** Single-cell level average NO signals for CREs displayed increased NO on the inactive X chromosome, indicating reduced CRE accessibility on this homologue (adjusted *P*=0.081, wilcoxon signed-rank test, followed by FDR-adjustment). **(b)** Single-cell level average NO signals at gene bodies for expressed genes (FPKM>1 in bulk-cell RNA-seq data[3]) were computed per haplotype, which revealed increased NO on the inactive X chromosome (adjusted *P*=0.0012, wilcoxon signed-rank test, followed by controlling[10] the FDR). Pseudoautosomal regions (PAR) were tested separately from the remainder of chromosome X ("nonPAR"). **(c)** Fold changes of haplotype-resolved NO in gene bodies plotted for chromosomes X and chromosome 7 (a representative autosome). For each expressed gene, fold change was calculated as the ratio between median single-cell NO signals for haplotype 1 (H1) vs. haplotype 2 (H2). Red: NO signals higher in H1, blue: higher in H2. The inactive chromosome X (represented by H1) showed significantly increased packing of nucleosomes in gene bodies suggesting decreased chromatin accessibility. **(d)** Fold changes of haplotype-resolved RNA counts on chromosome X in NA12878 cell line, derived from a female donor. For each expressed gene (FPKM>1 in bulk-cell RNA-seq data[3]), the fold-change between H2 and H1 of four biological replicates was computed using EdgeR[9]. We considered all RNA-seq reads mapping to heterozygous SNPs in non-PAR regions of the X chromosome to identify the active (H2) and inactive (H1) X chromosomal homolog. Red: RNA signals higher in H1, Blue: higher in H2. (**e**-**f**) NO tracks of NA12878 based on bulk-cell MNase-seq and pooled (pseudo-bulk) Strand-seq data. Representative loci (**e**) undergoing X-inactivation (*WWC3*) and (**f**) escaping from X-inactivation (*DDX3X*) are shown. CRE definitions are based on[8]. Adjacent boxplot represent measurements of haplotype imbalances in NO at the respective gene bodies (nominal $P<0.032$ for *WWC3*, and $P<0.52$ for *DDX3X*; likelihood ratio test). Bar graphs depict haplotype-resolved bulk RNA-seq[3] read counts (FDR-adjusted $P<2.5E-40$ for *WWC3*, and $P<1$ for *DDX3X*; likelihood ratio test).

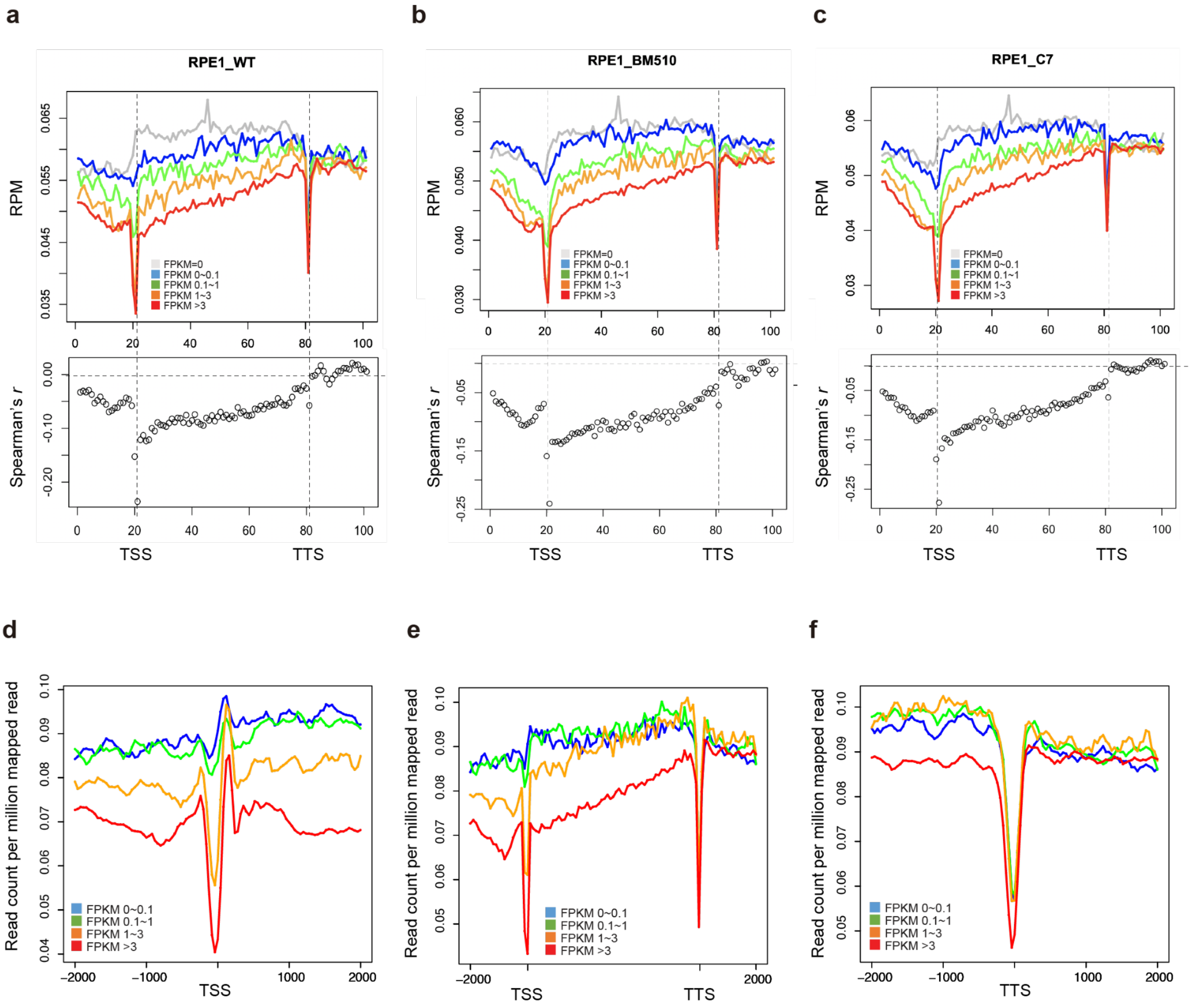


**Figure S6. Inverse correlation between NO at the body of genes and bulk RNA-seq gene expression values.** Inverse correlation shown for three RPE-1 derived cell lines[1]: the original RPE-1 cell line (denoted WT) **(a)**, BM510 **(b)**, and C7 **(c)** NO was calculated for 101 bins spanning -2kb to +2kb of gene bodies as a read count per million mapped reads using ngsplot software[11]. For each of the bins, genome-wide correlation between NO and gene expression level from bulk RNA-seq data was measured using Spearman's rho. Inverse correlation between NO and gene expression was apparent along the entire gene body (see gray dots), with the most pronounced inverse correlation measured at the TSS. **(d-f)** An equivalent inverse correlation between NO and gene expression level was also seen in published scMNase-seq data from a mouse cell line[12] (NIH3T3) - consistent with (pooled) Strand-seq and (pooled) scMNase-seq based read tracks being highly concordant along the genome (see main text and **Figure 1**). Binned NO profiles of the **(d)** TSS (±2kb), **(e)** gene bodies (±2kb), and **(f)** the transcriptional termination site (TTS; ±2kb) were extracted from pooled scMNase-seq data[12] using 45 single cells in total. We downloaded raw fastq files (GSE96688) and aligned these data to the mouse mm10 reference genome. The mono-nucleosomal fraction was extracted (140-180bp) (**Supplementary Notes**), and NO for genomic bins around TSSs (±2kb), gene bodies (±2kb), and TTSs (±2kb) computed using ngsplot[11]. Genes were divided into four groups based on expression values (FPKM) measured by bulk RNA-seq of NIH3T3 cells[13] (depicted in red, orange, green and blue).

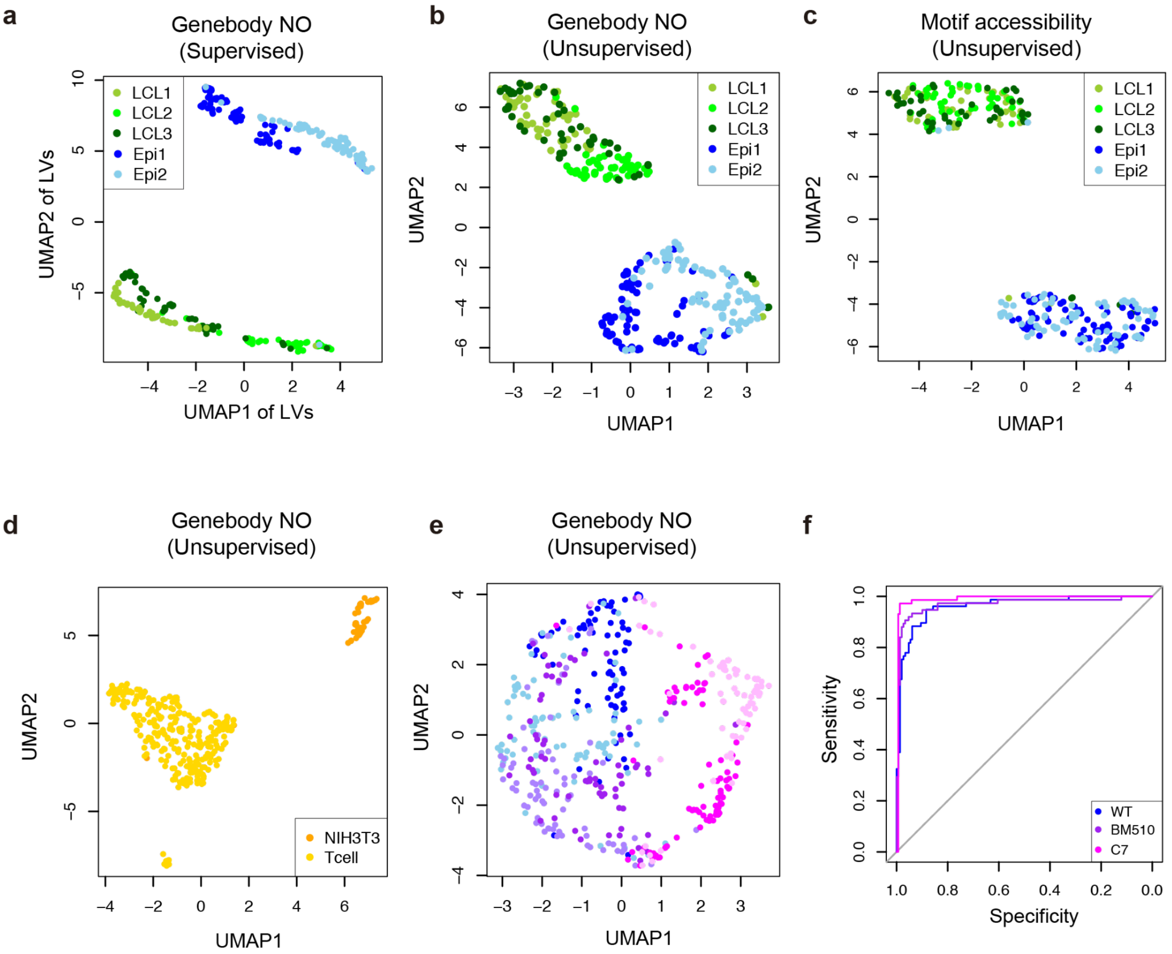


**Figure S7. Utility of NO to discern between cell types. (a)** Cell type classification based on NO at gene bodies (AUC=1). Cell line codes: Blue: RPE-1 (Epi1: RPE-1 replicate 1 (79 cells); Epi2: replicate 2 (77 cells)). Green: LCLs (LCL1: HG01573 (46 cells); LCL2: HG02018 (50 cells), LCL3: NA19036 (50 cells)). LV: latent variable. **(b-c)** Unsupervised UMAP visualization of single cell Strand-seq libraries from epithelial and lymphoblastoid (LCL) cell lines. **(b)** UMAP plot was generated based on NO at gene-bodies (normalized by ploidy status[1]) as described in the **Methods**). **(c)** We also explored the classification based on motif accessibility. To assess whether patterns of inferred chromatin accessibility at DHSs could likewise serve to classify cell types using Strand-seq data, NO was computed across DNase I hypersensitive (DHS) regions from the Roadmap epigenome database [7]. Lengths of DHS regions were adjusted to 2kb. Using the chromVAR package[14], single-cell NO profiles in these 2kb regions were transformed into a deviation Z-score, which measures how likely a certain motif accessibility would occur when randomly sampling sets of peaks with similar GC content and read depth. For each single-cell, the deviation Z-score was calculated for 870 human TF motifs from the cisBP database[15] provided by chromVAR. These dimensionality reduction plots suggest that batch effect within the same cell type (three individuals in LCL, and two batches in RPE-1 sequenced separately) is minimal, and far less than cell-type dependent variability. We concluded that when using motif accessibility, batch effects are essentially invisible. **(d)** Unsupervised UMAP using single cell MNase-seq data[12], including 45 NIH3T3 cells and 272 murine naive T cells, based on NO at the gene-bodies. **(e)** Unsupervised UMAP of RPE-1 (WT) and its transformed derived cell lines (BM510 and C7). Two Biological replicates were sequenced for each cell line. **(f)** Receiver operating characteristic (ROC) using the PLS-DA based classifier. AUC for classifying each cell line was 0.9614, 0.9694, and 0.9892 for RPE-1 (WT), BM510, and C7 respectively.

a

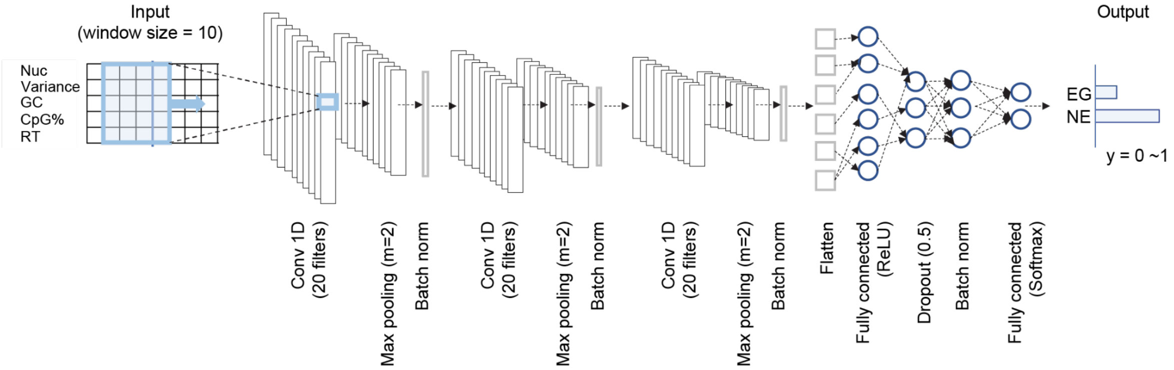


b

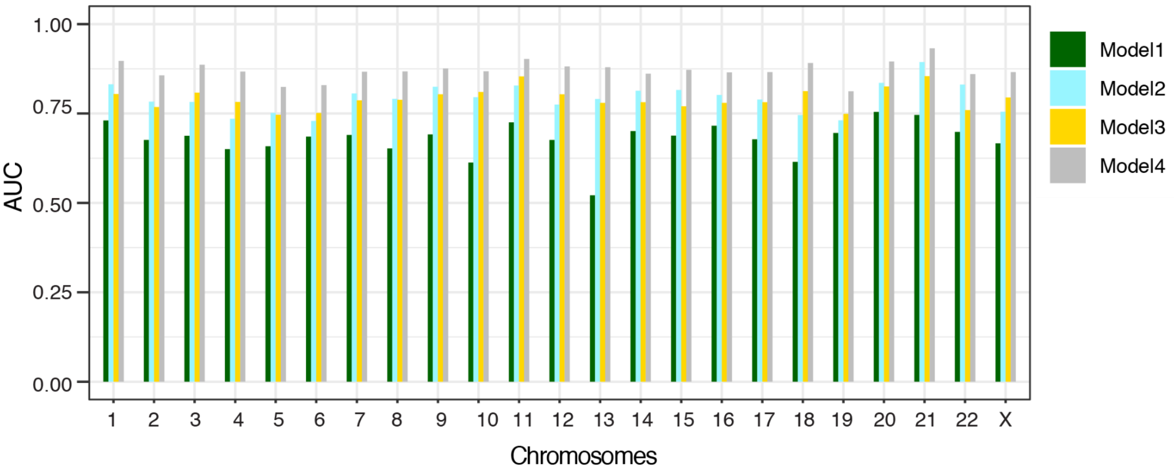


**Figure S8. Parameterization of convolutional neural network (CNN) in scNOVA. (a)** CNN architecture used by scNOVA. The 'Input' table shows a schematic representation of five layers of feature sets built into scNOVA’s CNN, which include NO, single-cell variance of NO, GC%, CpG%, and replication timing (RT); the latter three features have been reported to be associated with nucleosome positioning patterns[12,16–18], and have therefore been included in the CNNs to assist bin stratification. **(b)** Performance evaluation by cross validation, using different features and setups (“Models”), here used for choosing the optimal CNN model. Model 1 uses two features, 2K-TSS (-1kb to +1kb around the TSS) and nucleosome depleted region (-400 to +100bp around the TSS)[19]. Model 2-4 considers the region -5kb to +5kb of gene-bodies divided by 150 bins. Model 2 uses two layers of Strand-seq features (occupancy + variation). Model 3 uses three layers of genome annotation features only (CpG, GC, and RT). Model 4 uses five layers of features including Strand-seq features (occupancy + variation) and genome annotation features (CpG, GC, and RT). All models were trained by CNN except for Model 1, which was trained using a support vector machine (SVM) based setup[19]. The average AUC values of Model 1, 2, 3, and 4 were 0.679, 0.793, 0.791, and 0.871, respectively, and we thus chose Model 4 (occupancy, variation, CpG, GC, and RT) when parameterizing scNOVA. As an output, for each gene, this model provides the probability for a gene to be expressed (EG, expressed gene) or not expressed genes (NE, non-expressed gene), which when combined with scNOVA’s generalized linear models can be used to robustly infer alterations in gene activity (see **Methods**).

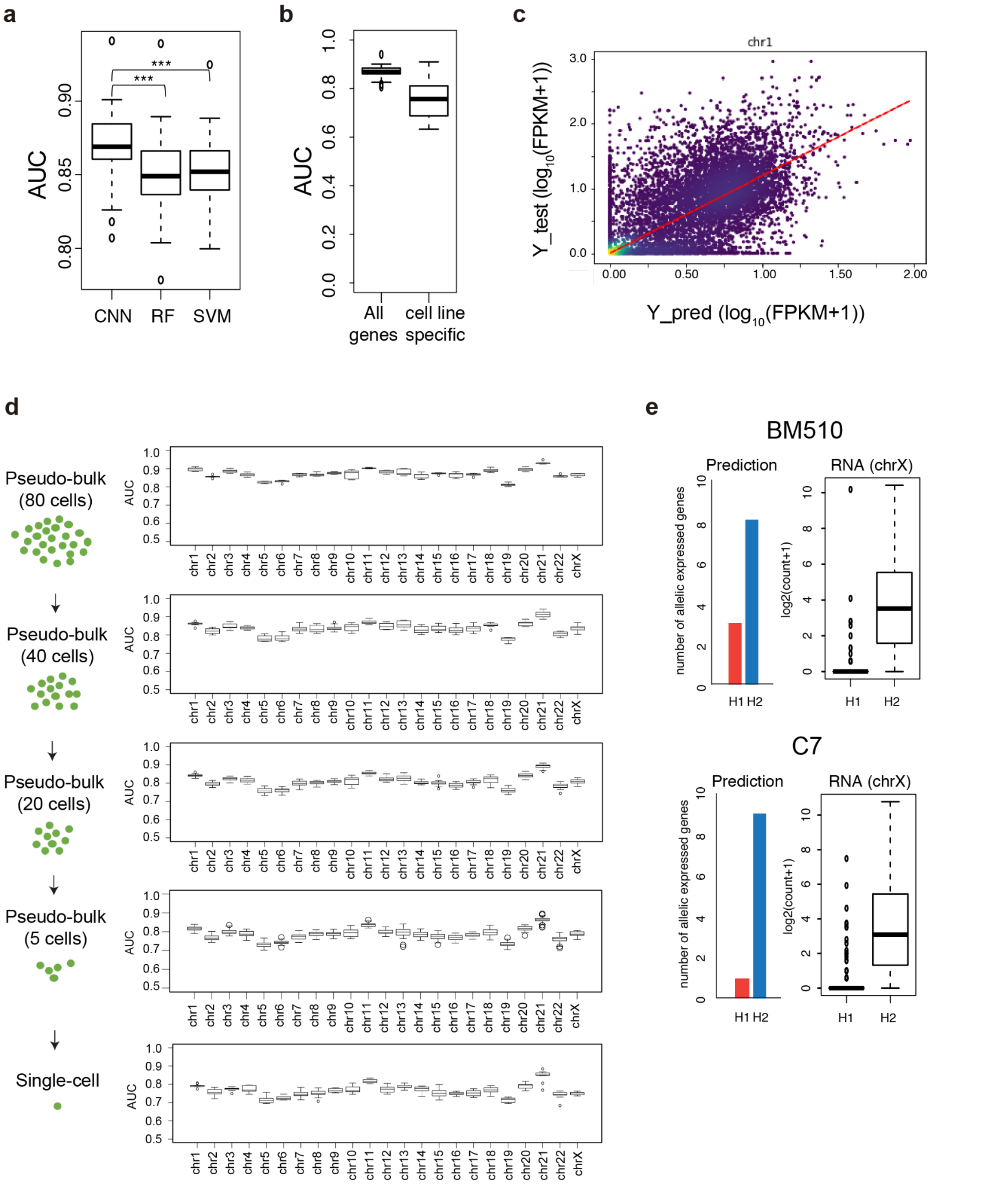


**Figure S9. Performance evaluation for scNOVA's CNN and comparison with other machine learning models. (a)** Comparison of AUC values based on leave-one-chromosome out cross validations from CNN, random forest (RF), and support vector machine (SVM). In this comparison, AUC was measured for the default (haplotype-unaware) CNN performing binary classification. All three models were trained with the same set of features, using pseudo bulk Strand-seq datasets from RPE-1 (79 cells), BM510 (plate 1: 70 cells; plate 2: 75 cells), and C7 (plate 1: 82 cells; plate 2: 72 cells). Published bulk-cell RNA-seq from these cell lines[1] was used to define ground truth labels for ~10,000 expressed genes per cell line. A boxplot depicts the AUC values from 23 cross validation experiments (one experiment per chromosome). The measured performance of the CNN surpassed random forest (RF) and support vector machine (SVM) based machine learning setups (Wilcoxon rank sum test followed by Bonferroni correction, $P = 2.4\text{e-}07$ for CNN vs. SVM, $P = 7.12\text{e-}07$ for CNN vs. RF). **(b)** AUC values for inferring gene expression ON/OFF status for all genes vs. cell type-specific genes. **(c)** Scatter plot of FPKM values measured by bulk RNA-seq (y axis), and inferred expression values predicted by the 'regression mode' of scNOVA's CNN (x axis). The scatter plot shows the result of leave-chromosome-1-out cross validation (Spearman correlation $r$=0.72; $P$<2.2e-16). The mean Spearman correlation coefficient across all 23 chromosomes was 0.68

($P$<2.2e-16). **(d)** AUC values for each chromosome, estimated in downsampled aggregated ('pseudo-bulk') Strand-seq data, as well as in single cells, using scNOVA's CNN. The overall AUC was computed as the weighted average over 23 chromosome pairs, scaled by the number of genes per chromosome. This downsampling analysis yielded AUC estimates of 0.78-0.87 in pooled, and AUC=0.76 in single cells for inferring expressed genes in RPE-1 cells. We note that coupling of the CNN with scNOVA's generalized linear models (see **Methods**) is highly recommended when using the scNOVA framework. **(e)** When exploring the utility of machine learning to infer gene activity based on NO tracks, we devised both haplotype-aware and -unaware CNNs. This panel depicts results generated by using haplotype-aware binary classification, with a haplotype-aware CNN, to infer the active X chromosome in the BM510 and C7 retinal pigment epithelial cell lines. In the case of BM510, the CNN nominated H2 as the active X chromosome. Bulk RNA-seq results[1] verified this inference. In C7, the CNN inferred the same haplotype as the active X chromosome. Bulk RNA-seq results[1] again verified this inference.

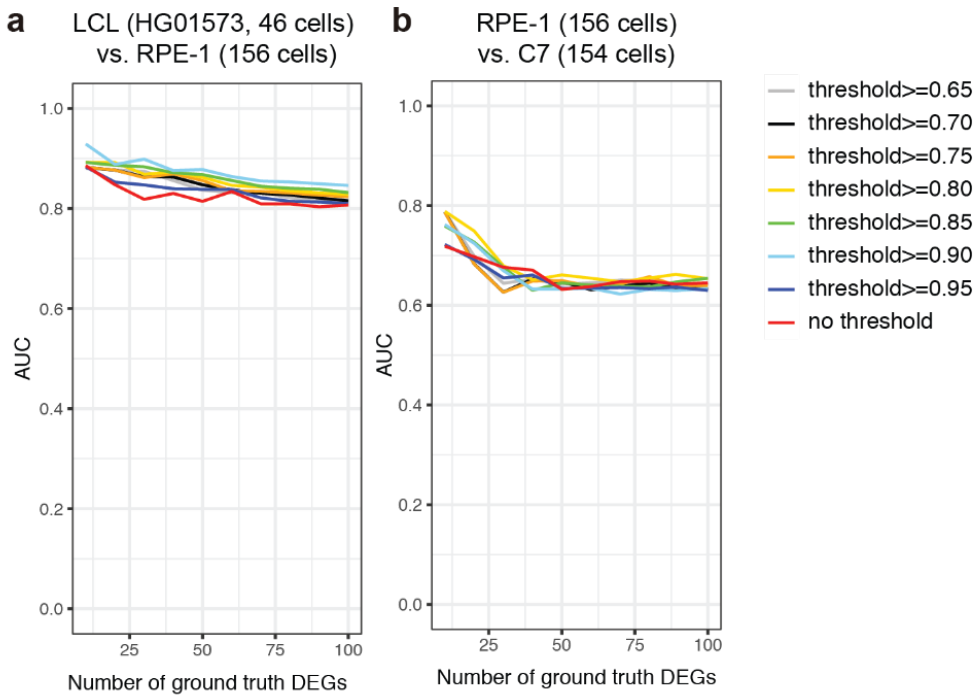


**Figure S10. Parameterization of scNOVA's differential gene activity analysis module.** As explained in the Methods, scNOVA first infers expressed genes (EGs) and non-expressed genes (NEs) using its default CNN, and then removes genes inferred to represent NEs. It then uses negative binomial generalized linear models (as available in the DESeq2 package[20]) on all remaining genes, to investigate NO changes at gene bodies, and accordingly to infer changes in gene activity. AUC, area under the curve. DEGs, differentially expressed genes (DEGs). DEGs (the "ground truth") are based on bulk-cell RNA-seq data subjected to DESeq2, comparing (**a**) RPE-1 and HG01573 (**b**) RPE-1 and C7. Coloring indicates the threshold used to filter out NEs based on the CNN: *e.g.* the threshold>=0.90 means that genes showing a probability of at least 0.9 to be not expressed (expression status='OFF') were filtered out. We chose 0.9 as the default threshold parameter, the application of which improved performance.

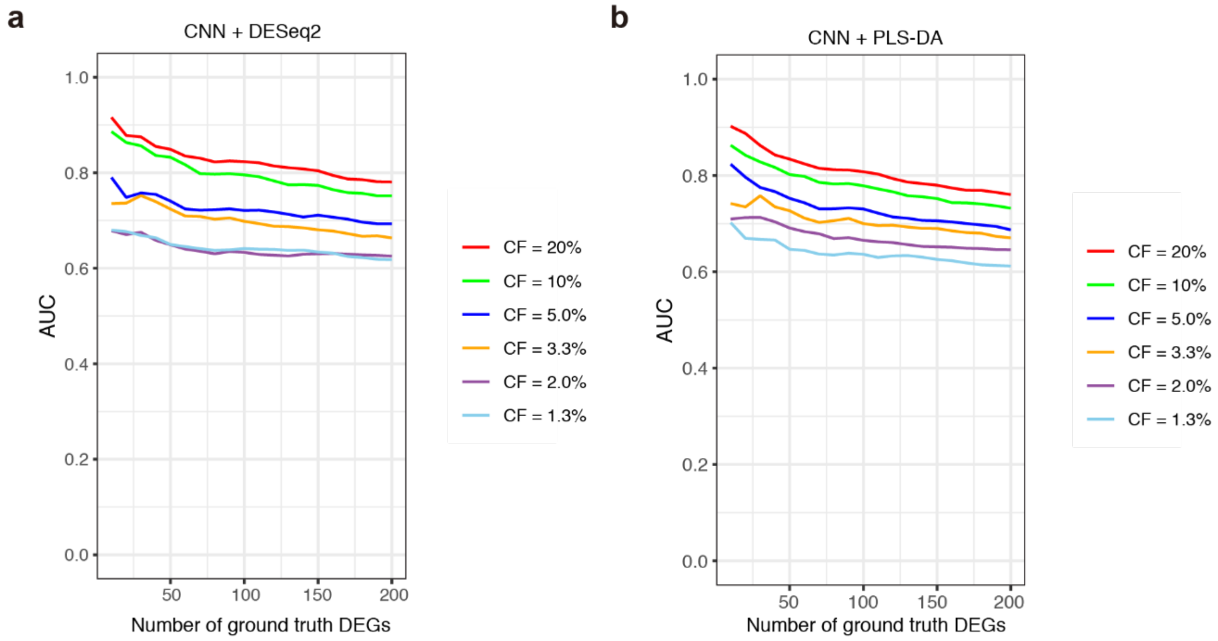


**Figure S11. *In silico* downsampling experiments using scNOVA's differential gene activity analysis module.** We performed *in silico* cell mixing of RPE-1 and HG01573 (LCL) to simulate different cell fractions (CFs). In this analysis six different CF ranges were considered (20, 10, 5, 3.3, 2, and 1.3). For each *in silico* cell mixing experiment, a total of 150 single cells were randomly subsampled from major pseudo-clone (containing RPE-1 cells) and the minor pseudo-clone (HG01573 cells), by controlling the minor pseudo-clone CF at 20, 10, 5, 3.3, 2, and 1.3%, respectively. AUC, area under the curve. DEGs, differentially expressed genes. For each CF, we performed random subsampling of single-cell libraries 10 times, and depicted the respective mean AUC in the plot. Two different analysis modes - (**a**) default (integrating convolutional neural network and negative binomial generalized linear model), and (**b**) alternative (integrating convolutional neural network and PLS-DA) were depicted here. These graphs show that when the CF is larger than 10%, default mode performs better, while for the CF smaller than 10%, alternative mode outperforms default mode.

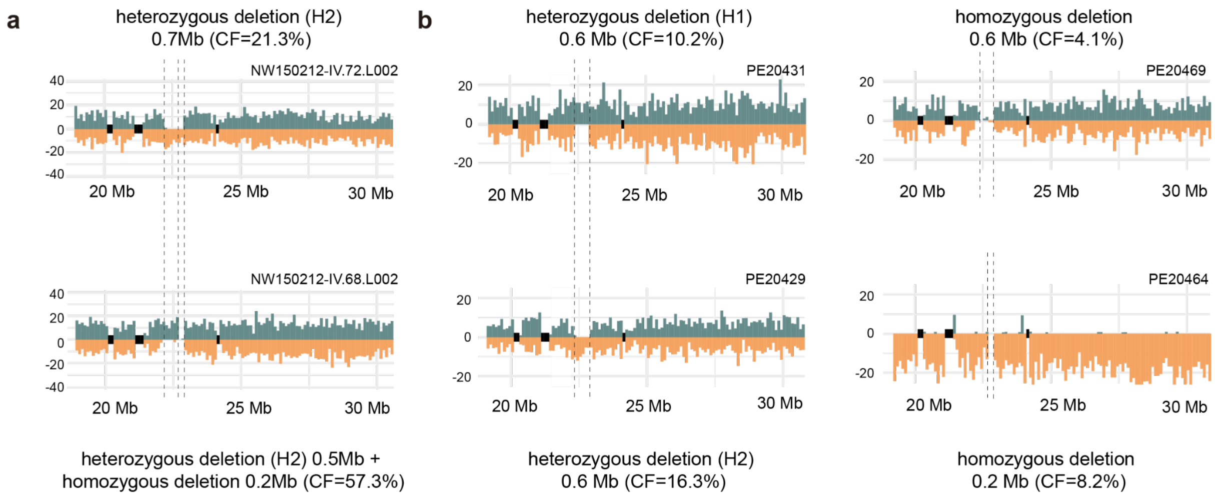


**Figure S12. Representative plots of LCLs which show evidence for at least two subclones exhibiting 22q11.2 deletions.** (**a**) Two 22q11.2 deletion bearing subclones found in NA12878, one subclone with 700kb heterozygous deletion (chr22:22.2Mb-22.9Mb), and the other subclone with 500kb heterozygous (chr22:22.2Mb-22.7Mb) and 200kb homozygous deletions (chr22:22.7Mb-22.9Mb). NA12878 additionally harbors a subclone bearing a 19q13.12 deletion (see main text). (**b**) Four 22q11.2 deletion bearing subclones detected in HG00171, which indicates that this LCL is a polyclonal cell line. Three subclones show hemizygous and/or homozygous deletion of a 0.6 Mb region (chr22:22.3Mb-22.9Mb), whereas one subclone shows homozygous loss of a 0.2 Mb region (chr22:22.7Mb-22.9Mb), at 22q11.2.

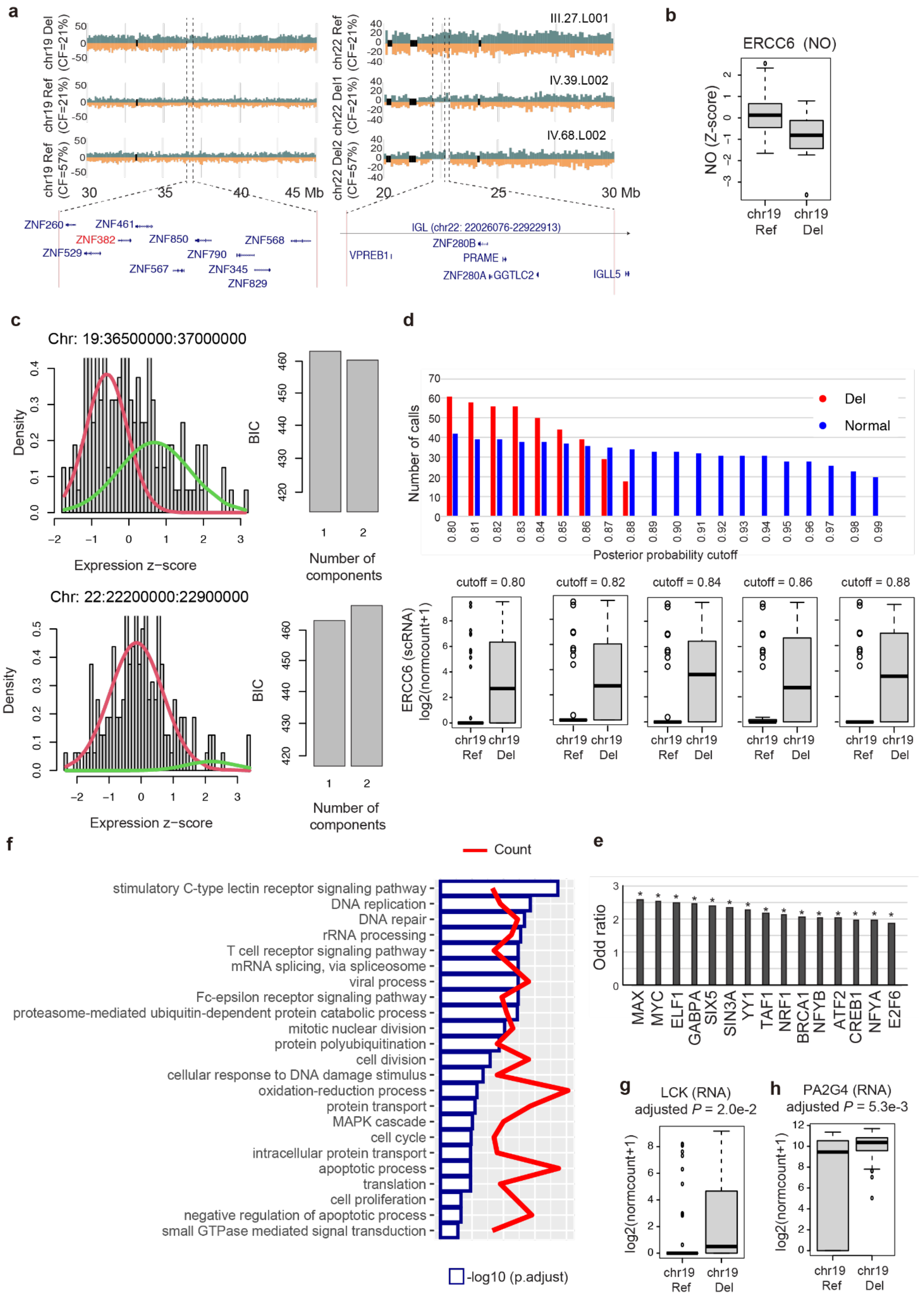


**Figure S13. scNOVA analysis of SV subclones in NA12878 and their validation using scRNA-seq.** (**a**) Mutually exclusive subclonal SVs in NA12878 single-cells. Del, Del1: hemizygous deletions; Del2: small homozygous loss region. Labels on the right side indicate single cell IDs. (**b**) Reduced NO at gene bodies predicting upregulated activity of *ERCC6* in cells of the NA12878 cell line carrying the 19q13.12 Del event (10%

FDR). **(c)** Analysis of NA12878 using CONICSmat[21] to genotype the single-cell copy-number state of the 19q13.12 deletion region using published[22,23] Fluidigm and Smart-seq scRNAseq data. Upper panel shows the histogram of mean expression Z-score of the genes located within the 19q13.12 deletion region (chr19:36.5Mb-37Mb). CONICSmat fits these distributions to 1-component (absence of subclonal copy number changes) and 2-component (presence of subclonal copy number changes) mixture models, and compares log likelihood ratio of two models to evaluate significance of difference between two models ($P$<0.00012; Chi-square likelihood ratio test). The result of 2-component model fits are shown in the plot with red and green bimodal peaks. The bar graphs show the Bayesian information criterion (BIC) value of two models. The model with the lowest BIC (2-component) is preferred, suggesting the presence of a somatic copy-number alteration (SCNA) at 19q13.12. Lower panel shows the histogram of mean expression Z-scores of the genes for 22q11.2 deletion region (chr22:22.2Mb-22.9Mb). For this region, the BIC prefers a 1-component model (absence of SCNAs). (**d**) In the case of the 19q13.12 region, for which CONICSmat inferred the 2-component model as the preferred model, the posterior probabilities of each individual cell were calculated to infer membership to the first component and the second component, and hence assign single cells as a "confident deletion" (red) or "confident normal cell" (blue). The bar graph shows the number of single-cells assigned to either confident deletion or confident normal for different posterior probability cutoffs. The box plots below show the comparison of *ERCC6* expression level from confident deletion and confident normal cells with different posterior probability cutoffs. These analyses are in strong support of the overexpression of *ERCC6* in cells exhibiting a 19q13.12 deletion ($P$=0.0067, 0.0050, 0.0051, 0.0092, and 0.052 for the posterior probability cutoffs of 0.80, 0.82, 0.84, 0.86, and 0.88 respectively; FDR-adjusted Wilcoxon rank sum test) **(e)** Odds ratio of TF targets enriched in cells bearing the 19q13.12 deletion, with c-Myc/Max target genes ranking highest (TF targets shown in this display exhibit adjusted $P$<1e-25 and combined score >100 based on EnrichR[24]), based on scRNA-seq in NA12878. **(f)** Gene ontology biological process (GOBP) terms over-represented among 1,896 up-regulated genes (FDR 1%, above 1.5 fold changes) in cells bearing the 19q13.12 deletion (61 cells) compared to WT (42 cells), based on scRNA-seq (applying a posterior probability cutoff of 0.80). GOBPs with FDR 10%, computed using Fisher's exact test with DAVID software[25], with more than 10 up-regulated genes are shown. The bar graph shows -log10 (p.adjust) values, and the red line represents the number of up-regulated genes for each term. **(g-h)** Depiction of representative up-regulated c-Myc/Max target genes implicated in cell proliferation, based on the NA12878 scRNA-seq data.

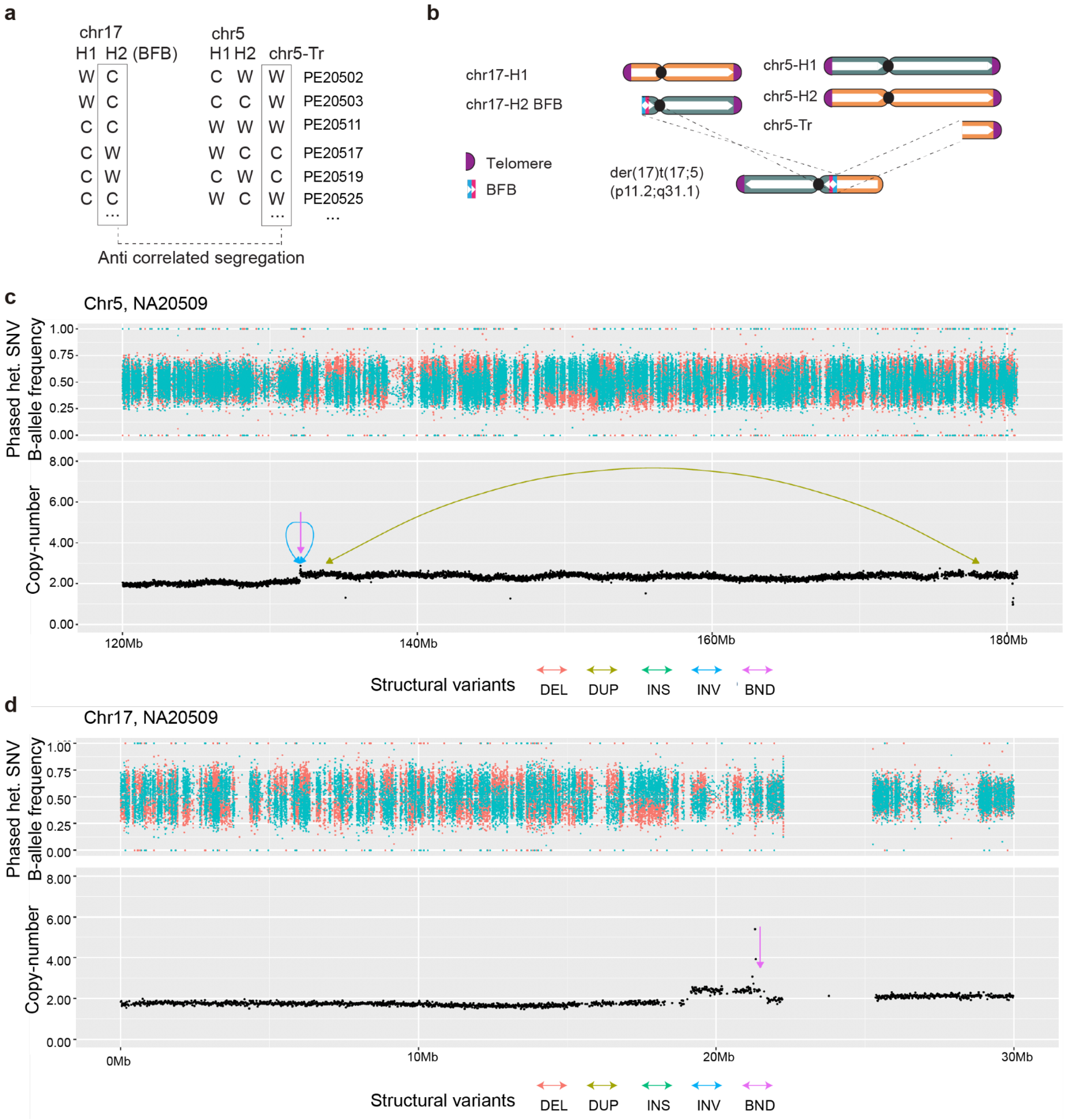


**Figure S14. Orthogonal verification of subclonal complex rearrangement found in NA20509.** (**a**) Discovered unbalanced translocation with CF=85% ($P$=1.3e-07; FDR-adjusted Fisher's exact test; inversely correlated template strand co-segregation patterns[1] used for translocation discovery shown for six representative cells). (**b**) Subclonal karyotype of NA20509, with complex derivative chromosome highlighted. **(c-d)** Read depth plot and SV calls based on NA20509 (bulk-cell) WGS data, generated at the New York Genome Center using a different cell stock of NA20509 than used for Strand-seq library preparation of NA20509 (pursued at EMBL Heidelberg). WGS data were downloaded from the data portal of the International Genome Sample Resource (IGSR)[26], and analyzed as described in the **Supplementary Notes**. Phased heterozygous sites for haplotype 1 (cyan color) and haplotype 2 (salmon color) together with read-depth based copy-numbers profiles verified **(c)** the presence of complex subclonal SVs including a large gain on chromosome 5, **(d)** subclonal terminal loss of chromosome 17 p-arm, and subclonal gain at 17p. SV analysis using Delly2[27] additionally verified the presence of a subclonal unbalanced translocation between chromosomes 5 and 17 (labeled 'BND'), showed a tail-to-tail inversion-type rearrangement at chromosome 5, and identified a tandem duplication-type rearrangement signature spanning parts of the terminal gain on chromosome 5. Further inspection of the Illumina WGS data suggested presence of the rearrangement-bearing subclone in 30-34% of cells (based on inspecting the core amplified region on chr5 and the core deleted region on chr17, respectively; see panels **(c)** and **(d)**). This is lower than the 85% CF detected in our own cell stock at the EMBL, perhaps since the CF of the respective subclone varies between distinct NA20509 cell stocks.

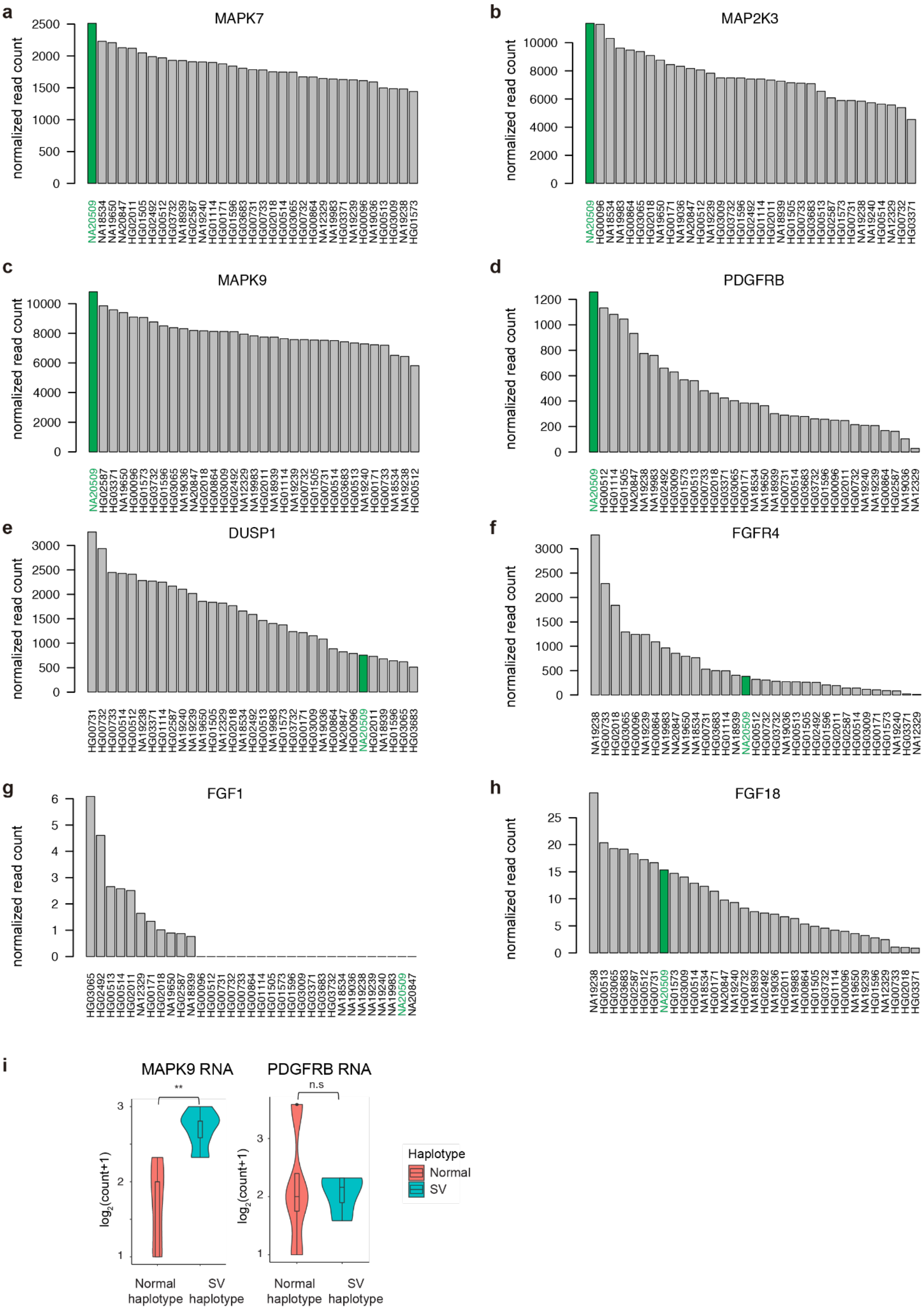


**Figure S15. Transcriptome analysis in a panel of 33 LCLs with available[28] deep bulk-cell RNA-seq data.** The figure depicts normalized read counts of MAPK pathway genes located in the duplicated region on chromosomes 17 **(a-b)** and 5 **(c-h)**. The expression level of NA20509 is highlighted with green. *MAP2K3* is an upstream regulator of c-Myc/Max in the MAPK signaling pathway. **(i)** Haplotype resolved RNA-seq read counts at heterozygous SNP sites within the *MAPK9* showed expression from both homologs, but also with increased expression in the BFB haplotype (SV) compared to the normal haplotype (Normal) (FDR-adjusted $P$=2.4e-3; likelihood ratio test) suggesting that this gene might also contribute to the molecular phenotype (activation of c-Myc/Max targets) in the genomically rearranged subclone.

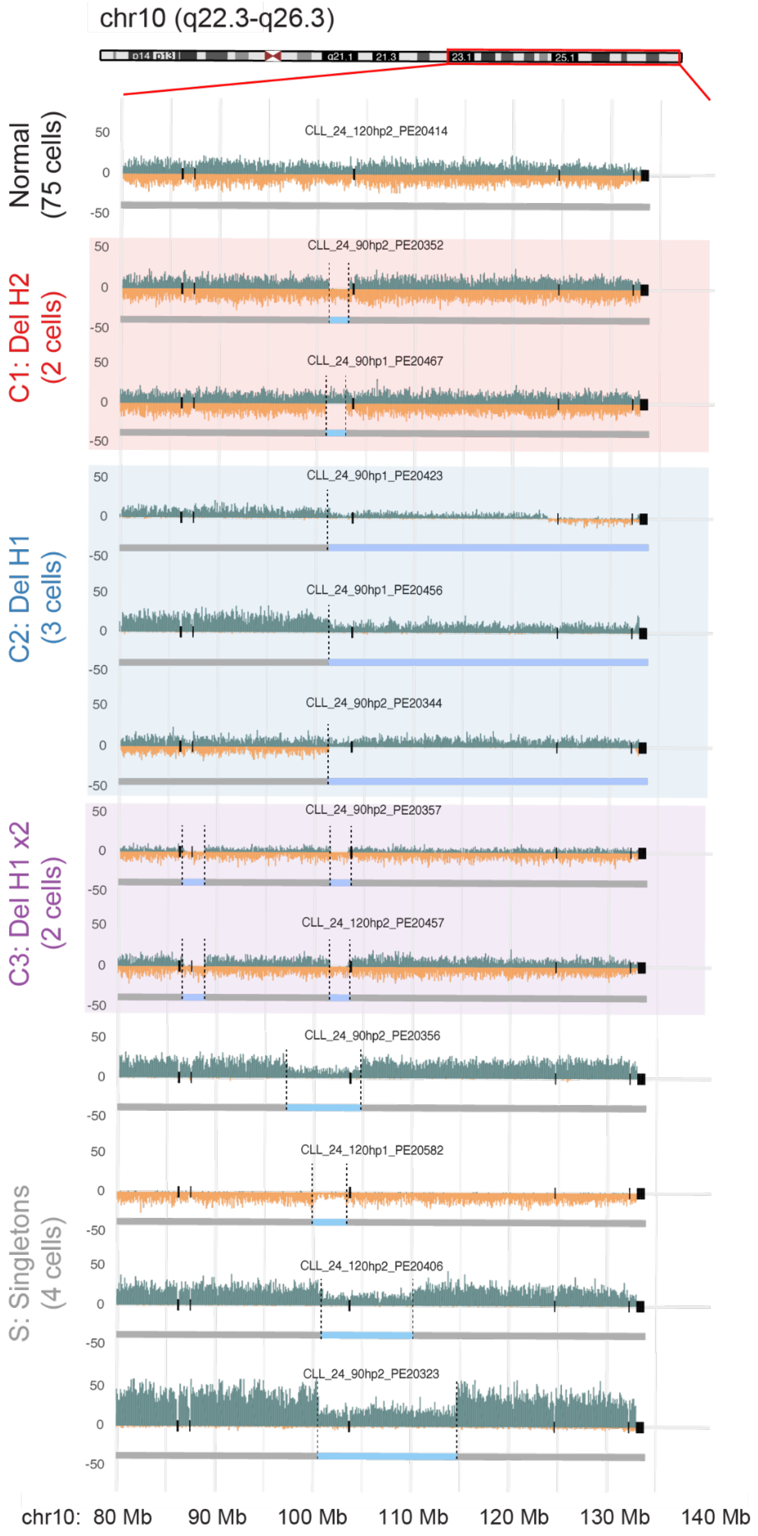


**Figure S16. Single-cell SV discovery in a CLL primary sample (CLL_24)** This panel reveals a diversity of distinct yet overlapping deletions at chromosome 10q24.32, representing subclones and single cells. Normal, normal karyotype. C1-C3 represent subclones with 10q24.32 deletions in the 'minimal region' (see main text). S (singletons), here represented as a 'group' of four single cells with distinct/individual deletions, again all affecting the 'minimal region' (see main text).

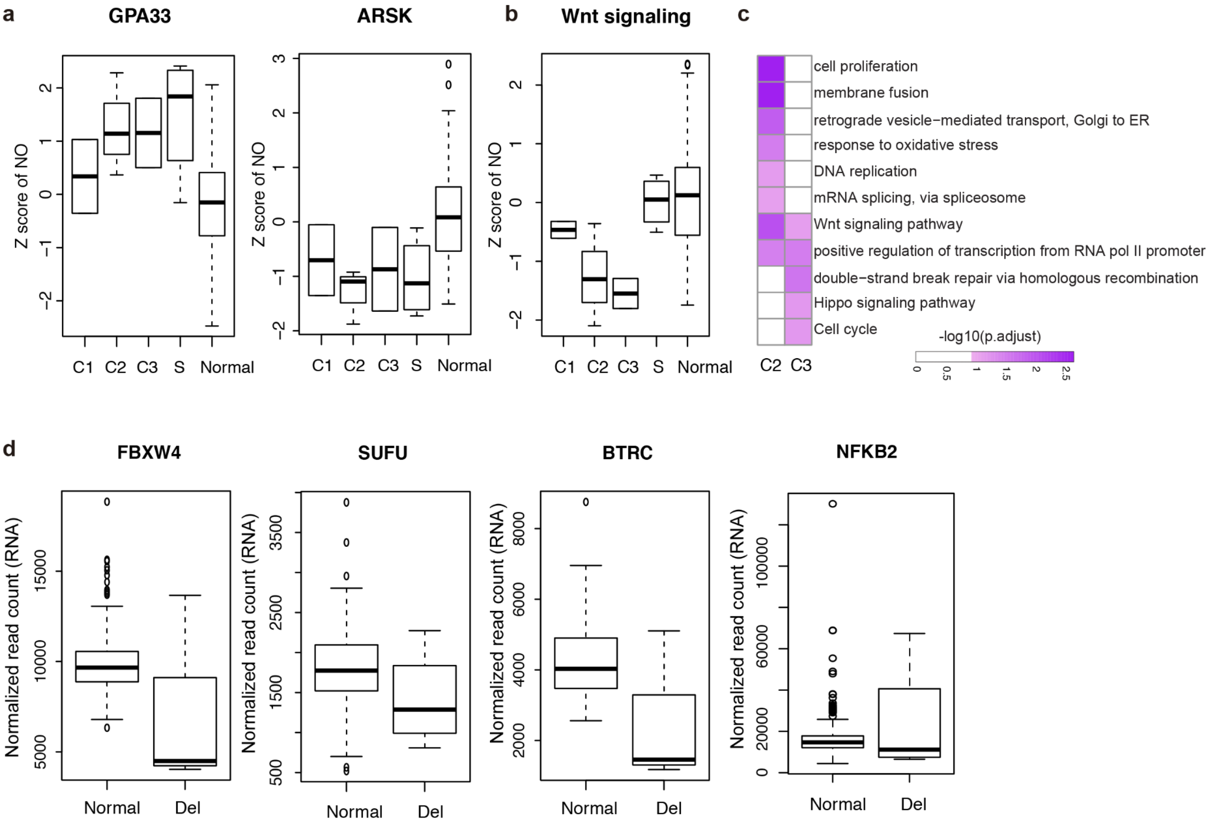


**Figure S17. Inference of altered gene activity of CLL_24 subclones using scNOVA and validation of differential expression of Wnt signaling regulators located in deleted regions via bulk RNA-seq from a large patient cohort. (a)** Inference of subclonal gene activity changes in CLL_24 based on scNOVA (10% FDR). 'Normal' indicates normal karyotype. C1, C2 and C3 denote subclones harboring deletions (Del) in the ‘minimal region’ at 10q24.32 (see main text). S combines four single cells that exhibit individual deletions in the same minimal region (seen in *N*=1 cell each). **(b)** Jointly modeled NO at the gene bodies of Wnt signaling pathway genes for each single-cells in different subclones. **(c)** Functional enrichment analysis of genes predicted from scNOVA's altered gene activity module for individual subclones with chromosome 10q deletions. As the CFs of distinct subclones were below 10%, we applied an alternative mode using [CNN+PLS-DA] for inferring altered gene activity. This analysis predicted 109, 206, 266 genes with altered activity. Gene set over-representation analysis was performed with the DAVID software[25]. **(d)** Differential expression of genes of interest in an ICGC CLL cohort[29], in samples bearing the ‘minimal region’ Del compared to donors without Del (‘normal’). Transcriptome-wide differential expression analysis revealed up-regulation of *DNM2* (FDR 10%, **Fig. 3g**), in support of the inferences initially made using scNOVA (see main text). We also checked the expression level of known or previously suspected negative regulators of Wnt signaling in the minimal deleted region in 10q. *BTRC* is a known negative regulator of Wnt signaling (**Supplementary Notes**), which is located very close to (only 58kb apart from) the minimal deleted region (deleted in 9/11 single-cells harboring the minimal region Del event). *BTRC* shows significant down-regulation in donors bearing the Del (FDR-adjusted *P*=0.000646), and hence its deletion may have caused or contributed to aberrant Wnt signaling. We also observed significant downregulation of *FBXW4* (FDR-adjusted *P*=0.00478), deleted in all eleven 10q24.32 SV bearing cells. In the case of *SUFU* (deleted in 11/11 cells with 10q24.32 SV) a slight trend of downregulation ($log_2$ fold-change = -0.34) did not reach genome-wide significance (FDR-adjusted *P*-value=0.346). For *NFKB2* (deleted in 11/11 cells with 10q24.32 SV), the expression level in the normal and deletion samples likewise did not show significant differences (FDR-adjusted P-value=0.507). Adjusted *P*-values were obtained by Wald’s test from DESeq2[20] followed by FDR adjustment.

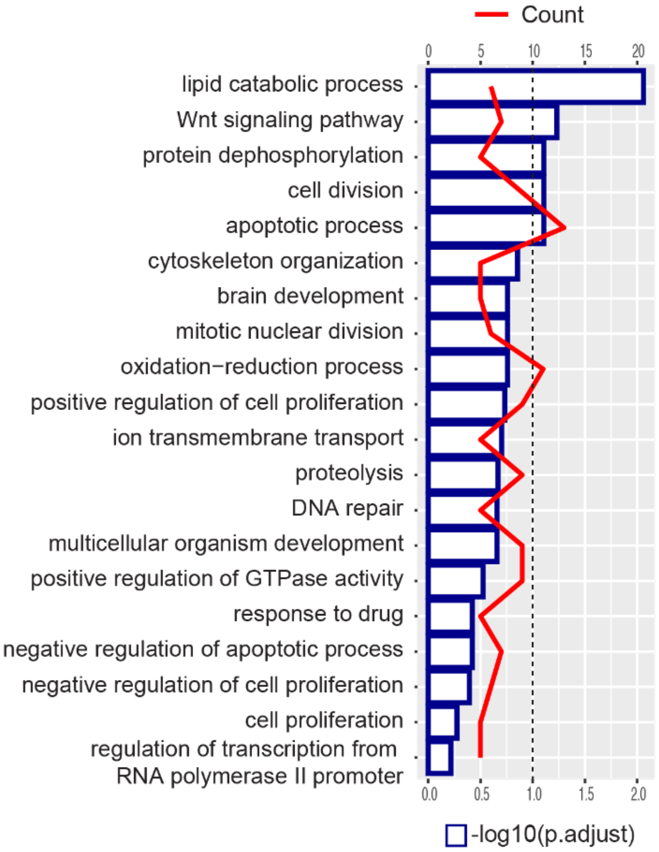


**Figure S18. Molecular phenotype analysis using gene sets with altered activity for distinct subclones in CLL_24.** Functional enrichment analysis of genes clustering genomically at 10q23.2-26.3. In total, 235 genes were affected by at least one of the Del events on 10q. These 235 genes were tested for functional enrichment with the DAVID software[25], using Fisher's exact test, followed by FDR correction. The bar graph shows the -log10(p.adjust) value, and the line graph shows the number of genes clustering in this genomic region that are involved in each biological process. Five gene ontology biological processes reach significance at 10% FDR. This includes Wnt signaling, which represents the second most significantly enriched functional category – suggesting that Wnt signaling-related genes are clustered in this chromosomal region.

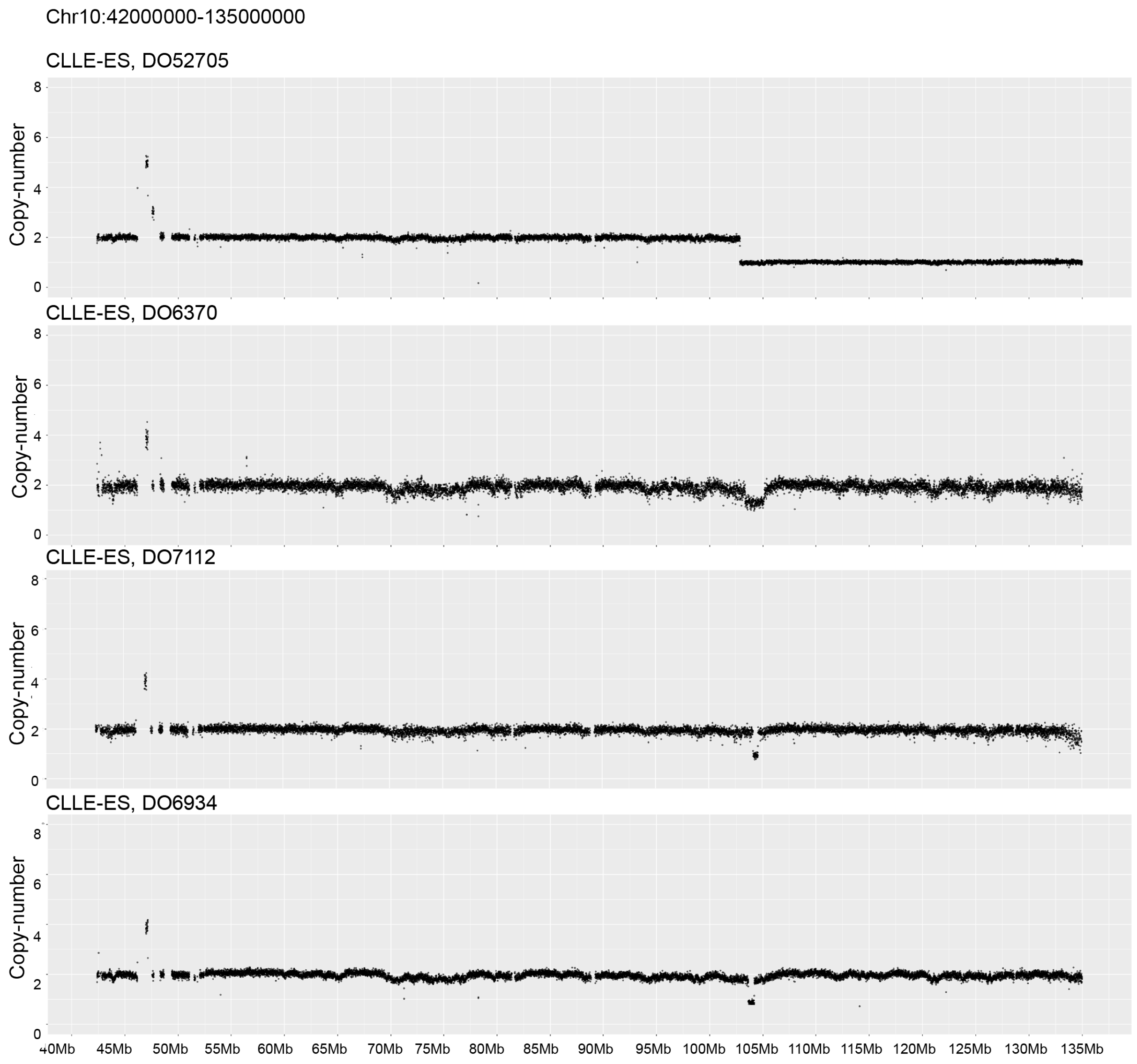


**Figure S19. Recurrence of deletions (Del) at the 10q24.32 'minimal region', in CLL donors from PCAWG[30].** Analysis of these WGS data by read depth analysis, using Delly2[27], uncovered Dels intersecting with the minimally deleted segment, initially observed in CLL_24, in 4 out of 94 (>4%) cases (all cases shown above).

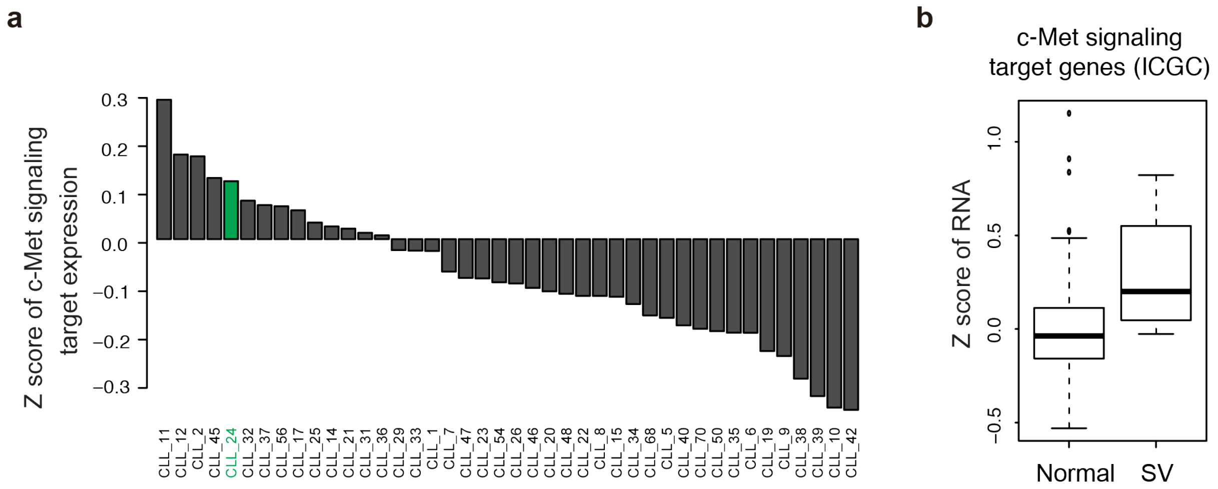


**Figure S20. Validation of increased expression of c-Met signaling target genes in the donors harboring chromosome 10q deletions. (a)** Bulk-cell RNA-seq analysis of c-Met signaling target genes. c-Met signaling target genes (**Table S7**) were collected from prior literature, where potential c-Met target genes were defined using global gene expression profiling of wildtype and c-Met-deficient primary mouse hepatocytes[31]. The expression Z-scores for those target genes are shown for each donor (CLL_24 shown in green). **(b)** CLL samples from the ICGC[32] bearing deletions at the 10q24.32 minimal segment (SV) show increased expression of c-Met pathway target genes compared to CLL samples with a normal 10q karyotype (*P*=0.0090; likelihood ratio test).

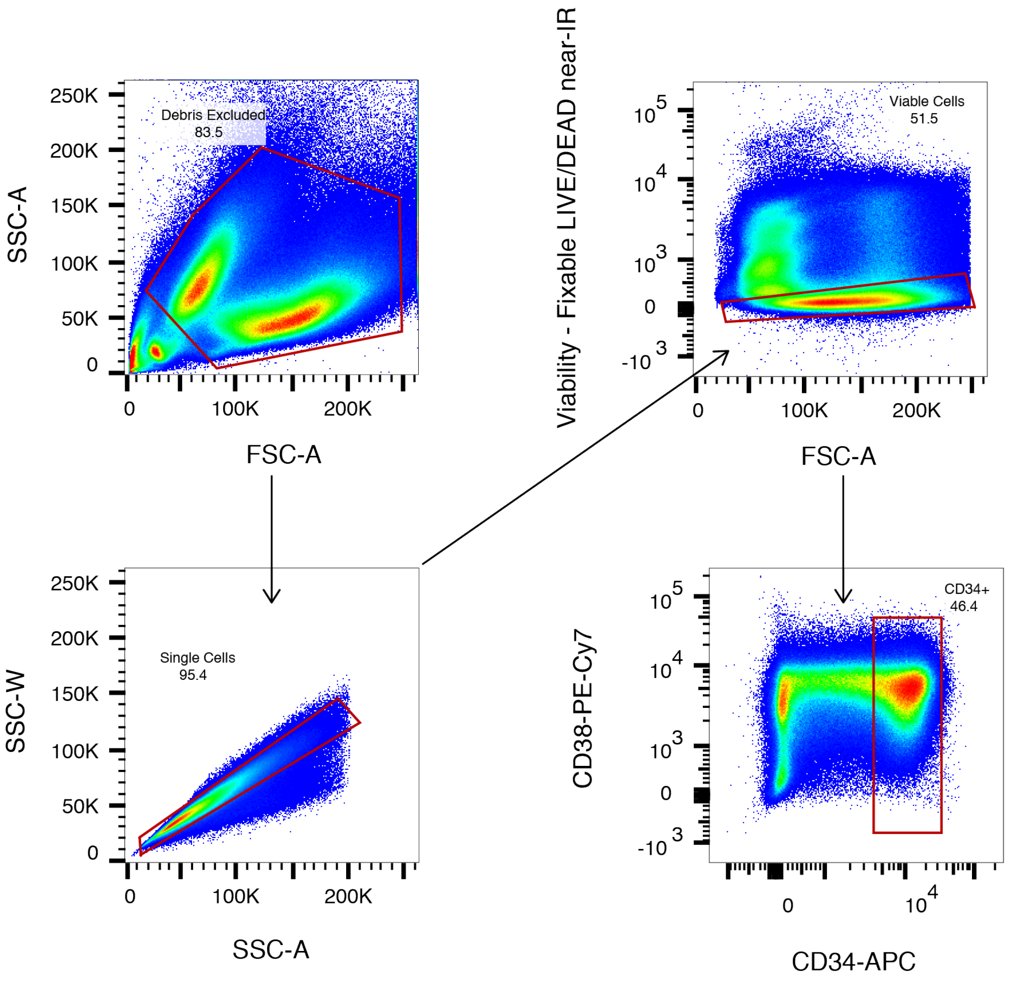


**Figure S21. Gating strategy for isolation of CD34+ cells from AML patient AML_1.**
Mononuclear cells from bone marrow aspirates were thawed and stained (**Methods**). Extracellular debris was gated out based on its low FSC-A vs SSC-A profile relative to cells. Doublets discrimination and exclusion was carried out by removing outliers in SSC-W vs SSC-A profiles; where doublets appear as outliers. Viable cells were identified by a low staining with Fixable LIVE/DEAD stain, an intracellular stain which does not strongly penetrate viable, intact cells. Finally, CD34+ cells were sorted from these single, viable cells. The red gate shows the selected population which is visualised in the consequent plot, indicated by an arrow. The final red gate indicates the population that was sorted, and used for Strand-seq library preparation.

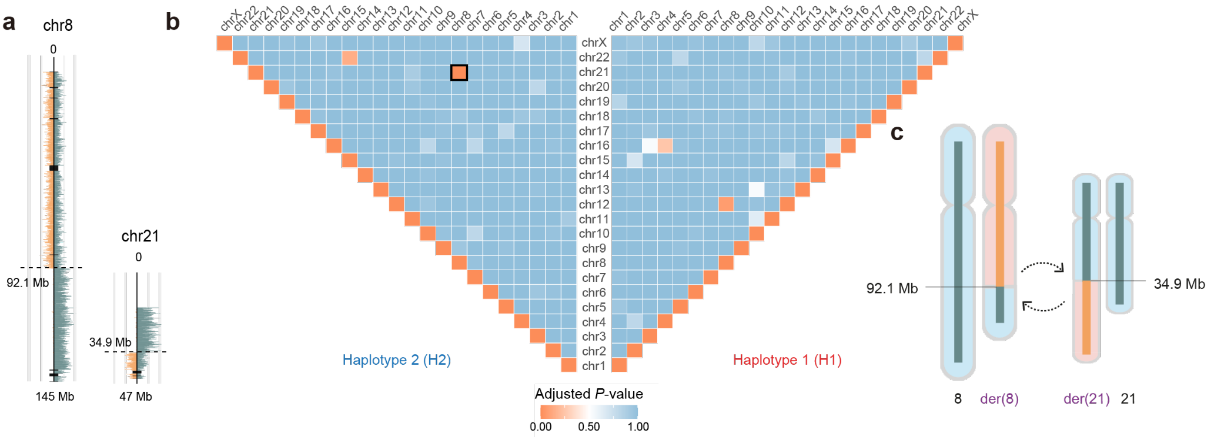


**Figure S22. Identification of balanced translocation in AML_1.** (**a**) Strand-seq based chromosome plot of representative single-cell from AML_1, which shows breakpoints at 92.1Mb of chromosome 8 and 34.9 Mb of chromosome 21. (**b**) Translocator analysis[1] of AML_1 provides the pyramid plot. Each pixel in the pyramid represents the significance of co-segregation between two chromosomal segments to detect potential translocation partners. This analysis suggests that AML_1 contains t(8;21) translocation. (*P*-value for translocation discovery using strand co-segregation: $P$=0.00003, FDR-adjusted Fisher's exact test). (**c**) The schematic diagram shows normal and derivative chromosomes resulting from t(8;21) translocation in AML_1.

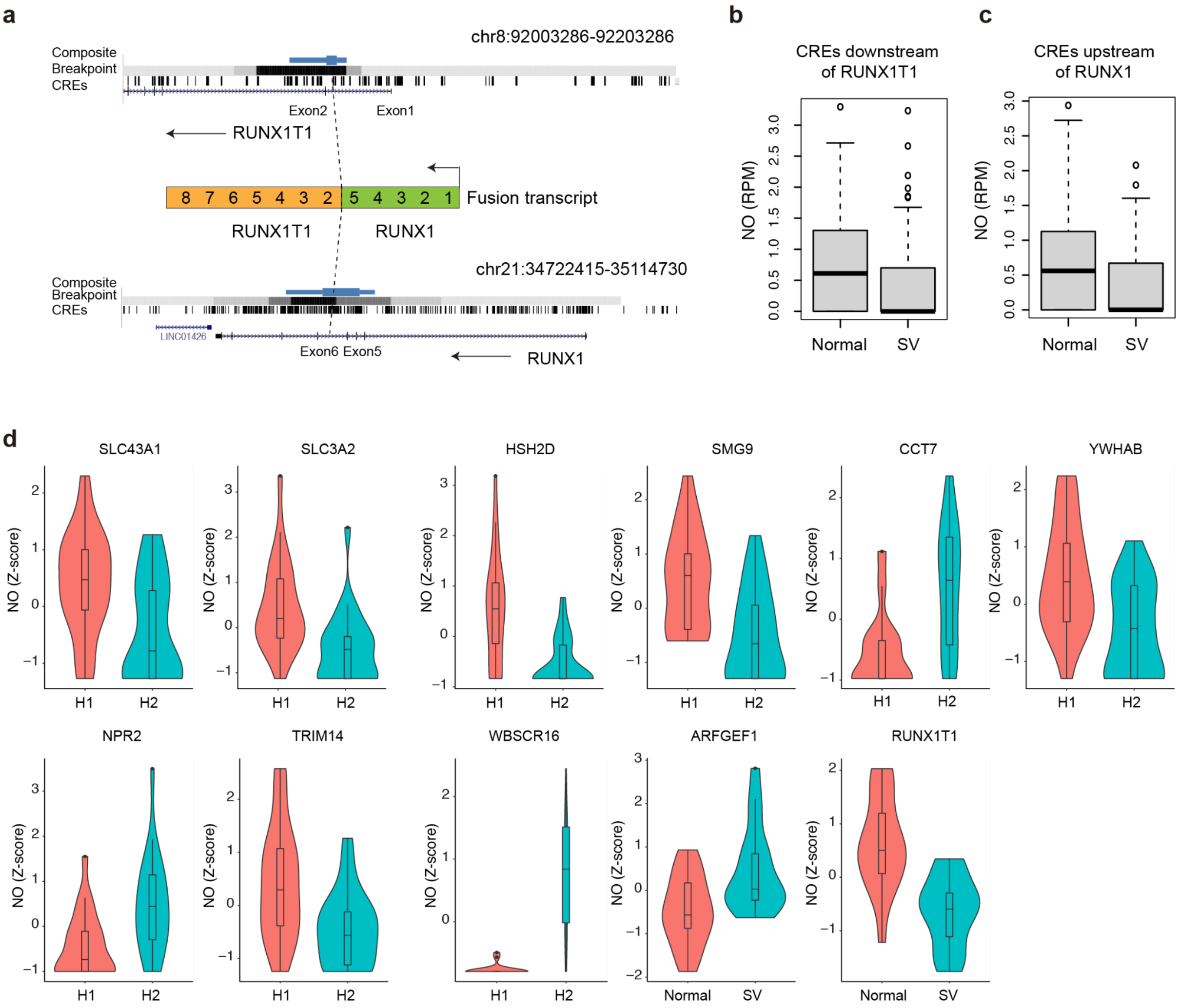


**Figure S23. Haplotype imbalance analysis of NO in AML_1.** (**a**) Strand-seq based breakpoint analysis using BreakpointR[33] located the translocation breakpoint in AML_1 within intron 1 of *RUNX1T1* on chromosome 8, and intron 5 of *RUNX1* on chromosome 21, recapitu100ing previously reported locations of t(8;21)(q22;q22.1) breakpoints[34]. CREs were defined as the union of ATAC-seq peaks from AMLs and the hematopoietic system from prior studies (**Table S5, Methods**). (**b**) Haplotype-specific NO at CREs within -278 to 22kb, adjacent to the translocation breakpoint which contains part of *RUNX1T1* ($P$<0.08; likelihood ratio test, adjusted using permutations). Normal, normal homologue of chromosome 8. SV, translocated (derivative chromosome). (**c**) Haplotype-specific NO at CREs in the upstream segment residing between 0.82Mb and 1.12Mb of *RUNX1* ($P$<0.003; likelihood ratio test, adjusted using permutations). **(d)** 11 genes demonstrate significant haplotype-specific NO in their gene bodies genome-wide (FDR<10%; likelihood ratio test).

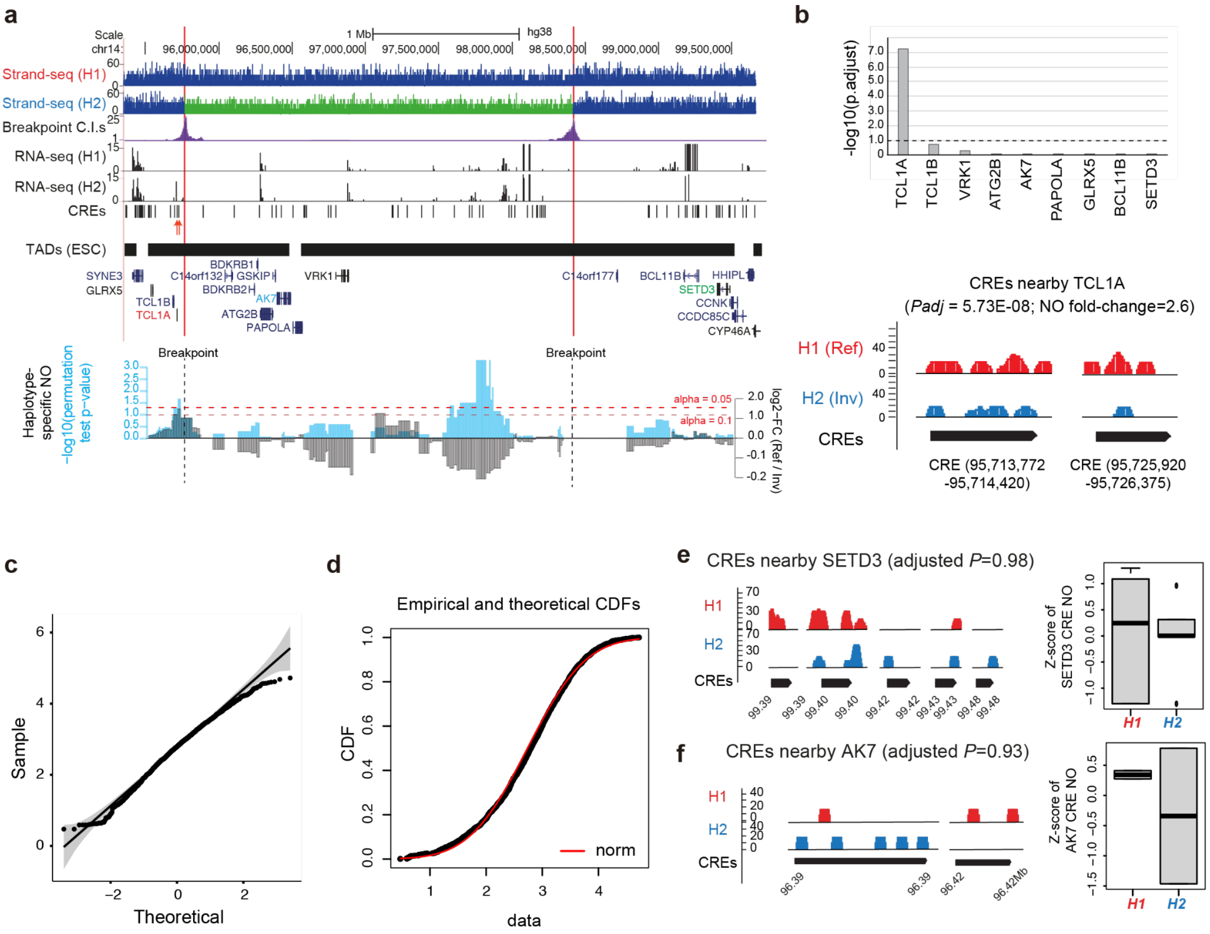


**Figure S24. Haplotype imbalance analysis of NO in T-ALL P1 to measure the cis-effect of clonal inversion in chromosome 14.** (**a**) Applying scNOVA to TALL-P1 reveals haplotype-specific NO near the breakpoints of a somatic inversion. H1: haplotype 1; H2: (rearranged) haplotype 2; RNA: allele-specific bulk RNA-seq expression values; light green: reads mapping in inverted orientation to H2, signifying the inversion[35]; dark blue: reads mapping in normal orientation. CREs were assigned to their likely target genes using the nearest gene approach (red arrows denote CREs assigned to the *TCL1A* oncogene). (**b**) haplotype-specific NO measured for CREs in the two TADs affected by the inversion. Upper panel: FDR-controlled likelihood ratio testing of haplotype-specific NO for CREs assigned to their target gene. Lower panel: haplotype-specific NO at CREs assigned to *TCL1A*. **(c)** Quantile-quantile plot of log$_2$-transformed read per million (RPM) of NO in the DHS and theoretical normal distribution suggests that log$_2$-transformed NO values exhibit normality. **(d)** Fitting of log$_2$-transformed RPM of NO (empirical, black line) to the theoretical cumulative distribution function (CDF) of normal distribution (theoretical, red line) to further evaluate the normality of log$_2$-transformed RPM. Fitting of distribution was performed using the fitdistrplus package in R[36]. Kolmogorov-Smirnov testing showed that the empirical distribution of the data is not significantly different from theoretical normal distribution (p-value > 0.05). Based on the normality checking of log$_2$-transformed NO, we chose to compare the NO of H1 and H2 using a generalized linear model fitting with gaussian family (default) and likelihood ratio tests. **(e-f)** Haplotype-aware nucleosome locations at CREs, shown for (**e**) a CRE near SETD3, and (**f**) a CRE near AK7 as example cases not displaying significant haplotype-specific NO (for comparison with panel (**b**)).

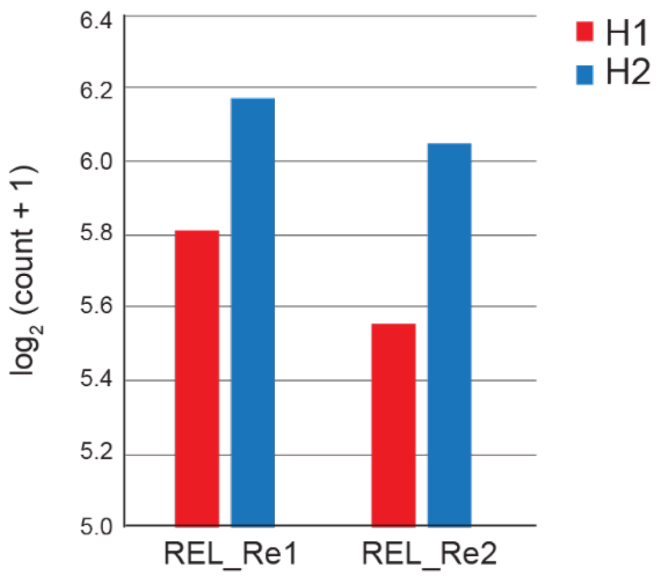


**Figure S25. Validation of allelic increase of RNA expression of *MYB* in the chromothripsis haplotype.** Increased *MYB* expression on haplotype 2 (H2) of chromosome 6, the chromosomal homolog exhibiting chromothripsis, based on haplotype-resolved bulk-cell RNA-seq pursued with two biological replicates (Re1 and Re2). Bulk RNA-seq data of TALL-P1 was analyzed to calculate allele-specific reads overlapping with heterozygous SNP sites. Allele-specific RNA-seq reads were counted using ASEReadCounter[37]. Allelic read counts were assigned to haplotype 1 (H1) or H2 using whole chromosome haplotype-phasing information from StrandPhaseR[38]. Allelic read counts along the gene were aggregated to retrieve haplotype-resolved gene-level read counts. *MYB* expression on H2 was 1.4-fold increased over H1 ($P = 0.0317$; likelihood ratio test, provided by EdgeR[9]).

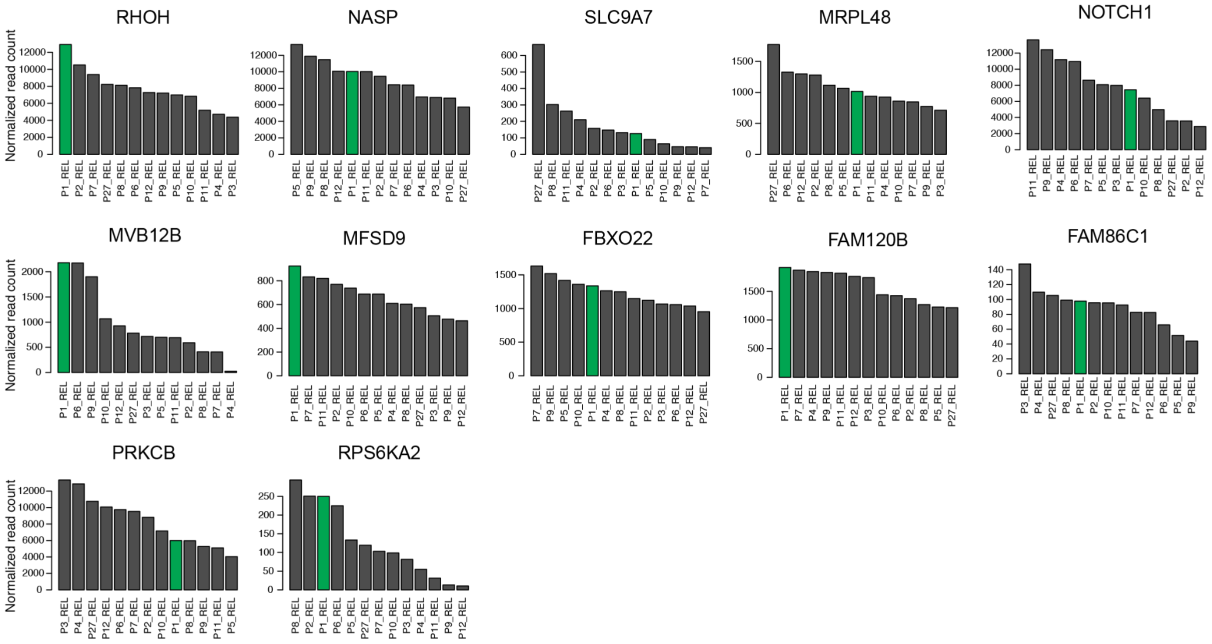


**Figure S26. Expression level of genes predicted by scNOVA for altered activity in the SV subclone in T-ALL P1.** Bulk-cell RNA-seq based gene expression measurements are shown for all genes in **Fig. 5d**, which scNOVA inferred to change in activity as a consequence of chromothripsis, in a panel of 13 T-ALL derived samples. The index sample, TALL-P1 (P1), is highlighted in green.

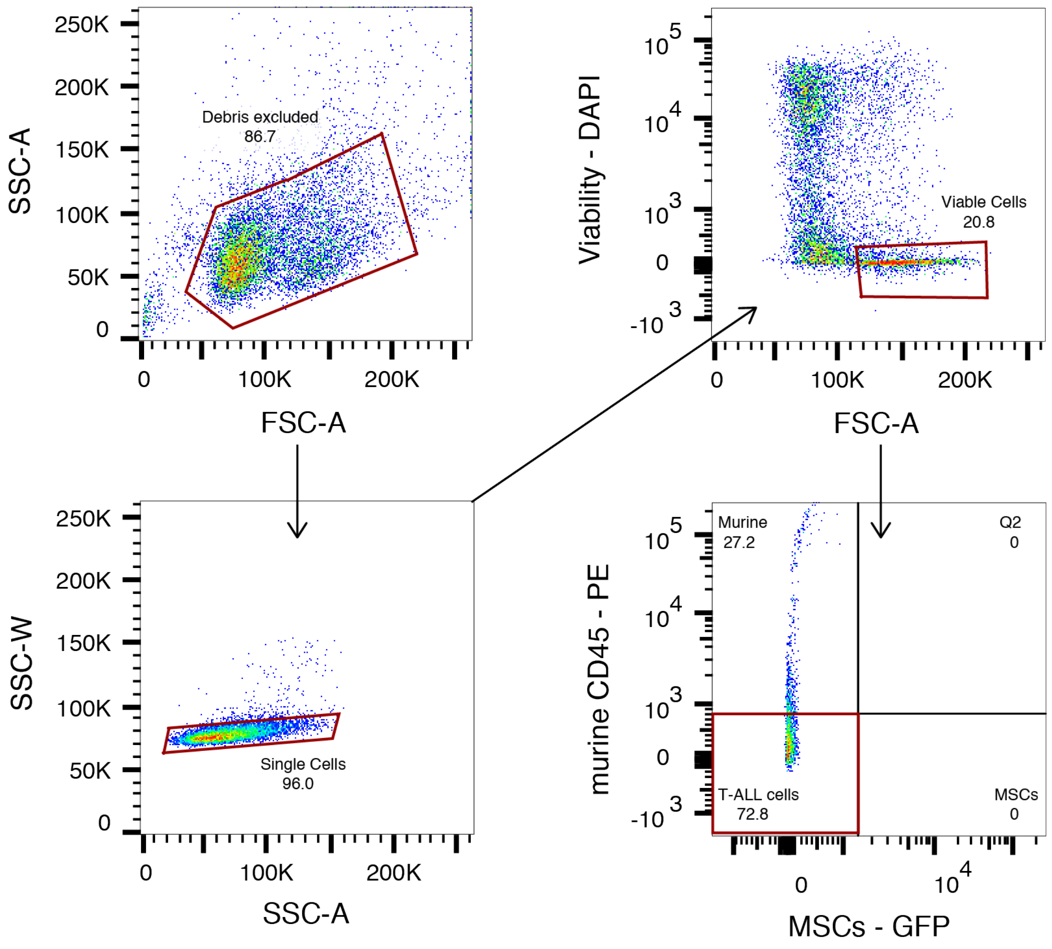


**Figure S27. Gating strategy for single, viable T-ALL cell isolation from T-ALL sample T-ALL P1 for scRNA-seq.** Viable cells are identified by low staining with DAPI, a viable cell impermeable nuclear stain. Human T-ALL cells are selectively sorted from contaminating murine and feeder layer cells by their lack of murine CD45 and GFP expression. Red gate shows selected population which is visualised in consequent plot, indicated by arrow. The final red gate indicates the population which was sorted.

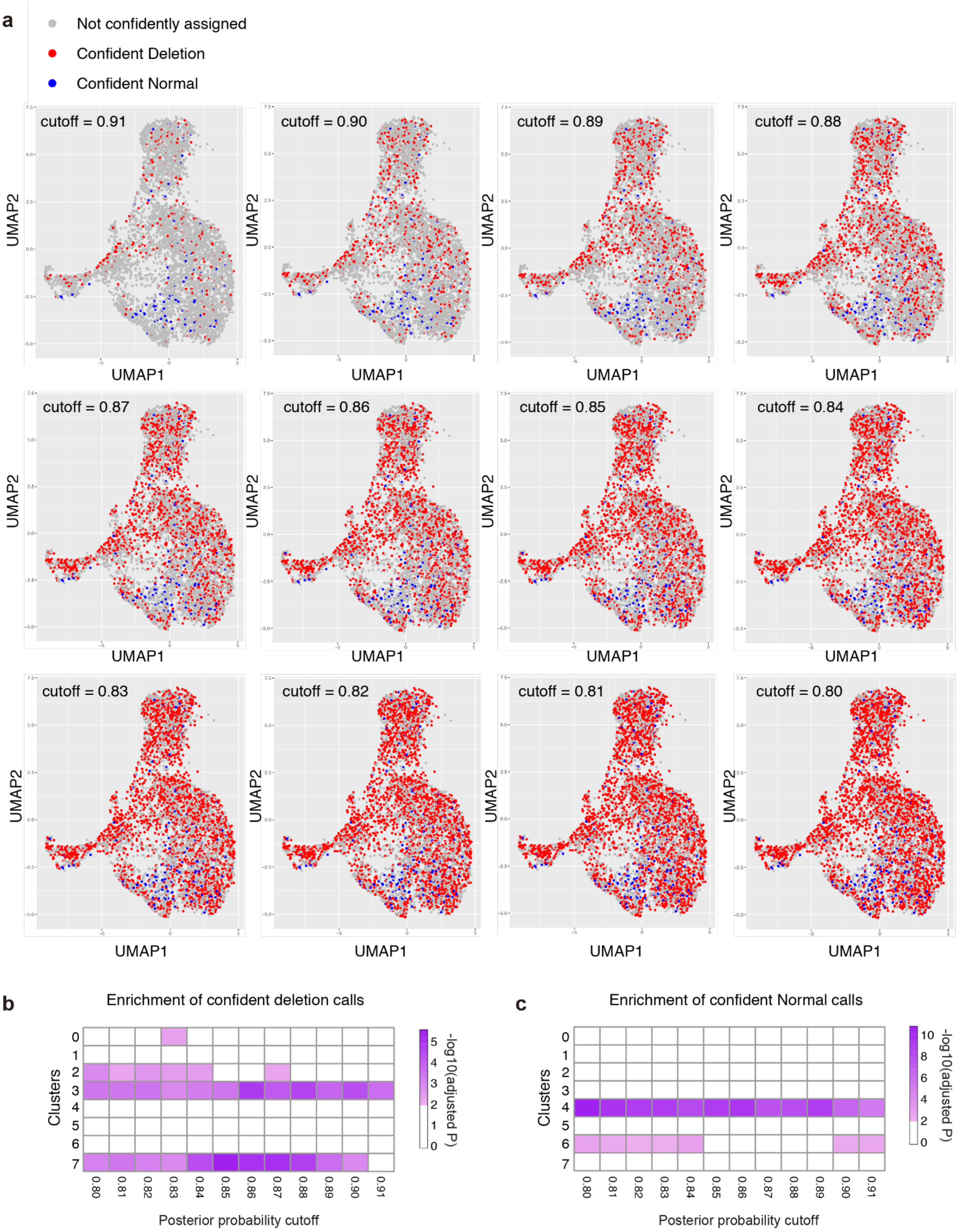


**Figure S28. Exploration of posterior probability cutoffs for the CONICSmat for the inference of SVs in T-ALL P1 using scRNA-seq.** (**a**) Single-cells harboring Deletions in chromosome 6 were inferred using CONICSmat 'genotyping mode' by applying different posterior probability cutoffs, and projected to the UMAP. As the maximum posterior probability was 0.9133, we tested the cutoff range from 0.80 to 0.91, by increasing 0.1 each time. (**b-c**) Enrichment of confident deletion calls (**b**) and confident normal calls (**c**) for each of the unsupervised clusters were tested using fisher's exact test followed by Benjamini-Hochberg multiple correction. It shows that cluster3 and cluster7 are robustly enriched by confident deletion calls, and the cluster4 and cluster6 are robustly enriched by confident normal calls.

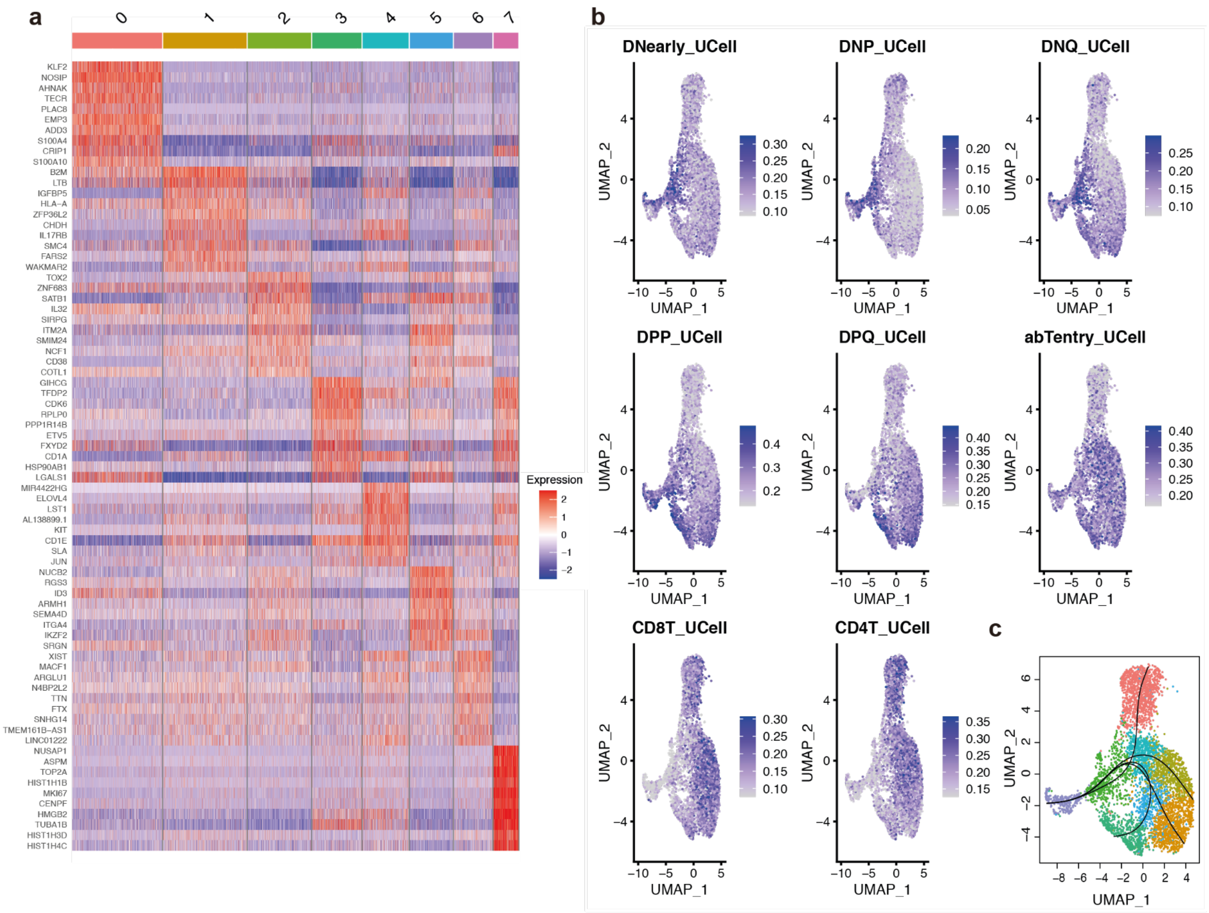


**Figure S29. Lineage trajectory analysis of scRNA-seq of T-ALL P1.** (**a**) Marker genes of eight unsupervised clusters identified from scRNA-seq of T-ALL P1 (10X Genomics). (**b**) Expression level of T-cell differentiation state marker genes were projected to the UMAP of scRNA-seq of T-ALL P1. T-cell differentiation marker genes were downloaded from the previous publication [39]. (**c**) Cell differentiation lineages inferred from scRNA-seq of T-ALL P1 using Slingshot[40].

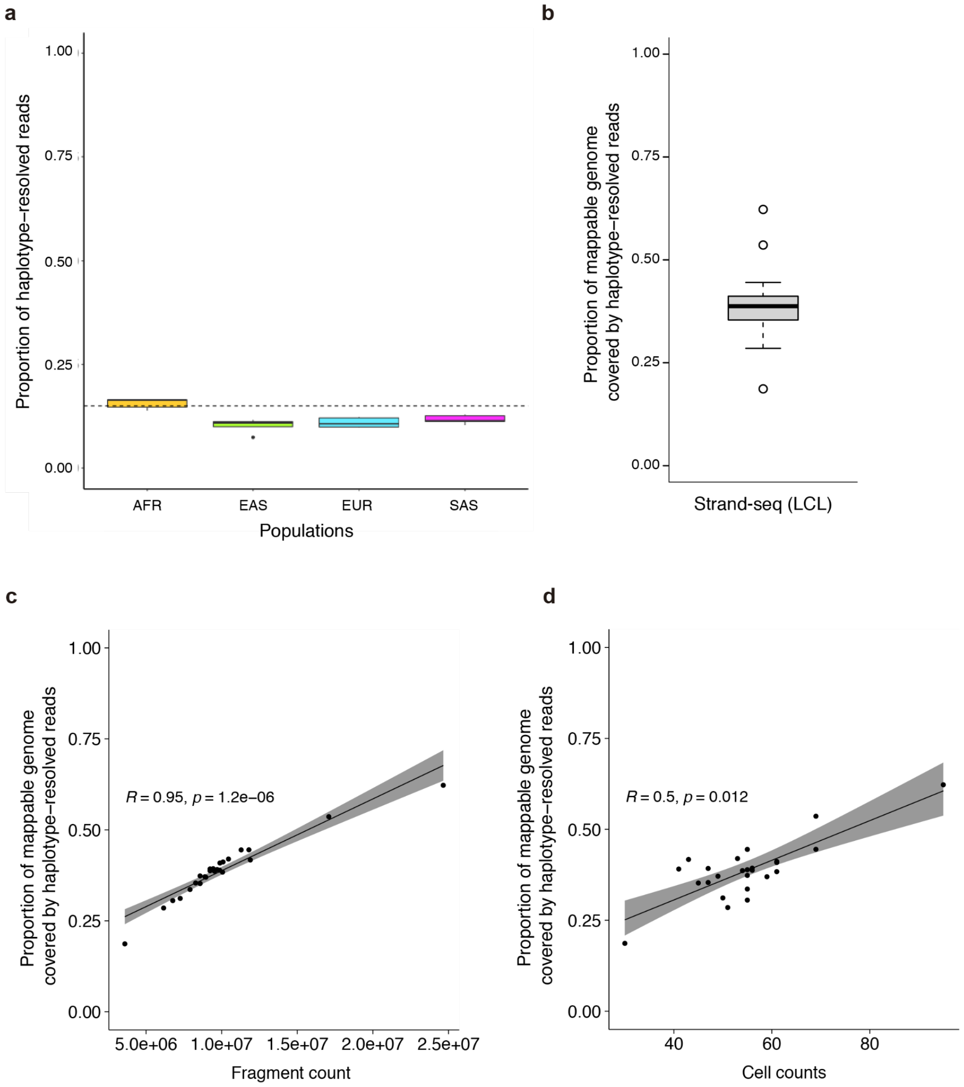


**Figure S30. Comparison of the Strand-seq to conventional WGS for the proportion of genome that can be accessible for allele specific analysis.** (**a**) Proportion of Illumina WGS reads that can be haplotype-phased, and thus are accessible for 'classical' allele-specific analyses, in the 1KG haplotype reference panel [41]. We randomly picked five 1KG samples, sequenced by Illumina short read based WGS, from each continental 'super-population' (AFR: African ancestry, EAS: East Asian ancestry, EUR: European ancestry, SAS: South Asian ancestry) and assigned reads to haplotypes using phased heterozygous sequence variants. The horizontal dashed line marks 15% of the haplotype-resolved reads, which is the approximate upper bound for the fraction of reads that can be assigned to a haplotype to pursue classical allele-specific analyses. This fraction is lower for samples from EUR, SAS and EAS populations, as expected, due to the smaller number of heterozygous SNPs compared to AFR populations. (**b**) Proportion of 1KG mappable regions that are haplotype-resolved by Strand-seq, using 25 LCLs (including 24 from a human diversity SV panel[28], as well as NA12878[2]). Single-cell libraries from chromosomes showing either a WC (Watson/Crick) or a CW configuration were haplotype-resolved using StrandPhaseR[38] and pooled into phased pseudo-bulk data sets. Evaluations were restricted to the autosomes. (**c**) Proportion of 1KG mappable regions that are haplotype-resolved using Strand-seq, plotted by the fragment count achieved for each LCL sample. (**d**) Proportion of 1KG mappable regions haplotype-resolved by Strand-seq, plotted by the number of cells sequenced for each LCL sample. The data in (**b**), (**c**), and (**d**) suggest that Strand-seq has the ability to access up to the entire mappable genome, *e.g.*, to allow haplotype-specific NO analyses from telomere to telomere. The actual fraction of genomic nucleotide bases accessed depends on the number of cells sequenced by Strand-seq, as well as on the sequencing depth (fragment count). It also depends on the specifics of nucleosome positioning, as outlined in **Fig. 1** in the main text, where the MNase digestion step used during Strand-seq library preparation directs sequencing to DNA regions protected by nucleosomes.

**Supplementary Tables**

**Table S1**. Characterization of nucleosomal fragments in Strand-seq libraries
*(Table accompanying the submission as a spreadsheet)*

**Table S2**. List of genes identified to be changing in activity when comparing major and minor clones with scNOVA. *(The Supplementary Material provides further details on some of the genes contained.)*
*(Table accompanying the submission as spreadsheet)*

**Table S3**. List of somatic SVs identified in single cells
*(Table accompanying the submission as spreadsheet)*

**Table S4**. Summary of inferring SCNAs from the scRNA-seq data sets
*(Table accompanying the submission as spreadsheet)*

**Table S5**. Reference list of ATAC-seq peaks from prior literature used to define putative CREs in AML_1
*(Table accompanying the submission as spreadsheet)*

**Table S6**. Reference list showing literature sources for sets of TF target genes, defined by TF binding (ChIP-seq) and RNA perturbation (RNA-seq or microarray) upon silencing of TFs
*(Table accompanying the submission as spreadsheet)*

**Table S7**. Gene sets used in expression analyses to verify TF target gene and pathway activities
*(Table accompanying the submission as spreadsheet)*

**Table S8**. Cell-wise genotypes and correlation matrix of NA12878 deletions on chr19 and chr22 determined using the ArbiGent tool
*(Table accompanying the submission as spreadsheet)*

***Table S9****.* List of differentially expressed genes identified for the eight unsupervised clusters of T-ALL P1 scRNA-seq
*(Table accompanying the submission as spreadsheet)*

***Table S10****.* Top20 significant TFs for Cluster3 and Cluster7 of T-ALL P1 scRNA-seq identified using EnrichR analysis
*(Table accompanying the submission as spreadsheet)*

#### Supplementary Notes for Methodological Details

##### 1. Estimating genome-wide coverage

NA12878 Strand-seq data aligned to the hg38 reference assembly was downloaded[2] and sequence reads with low quality (MAPQ<10), supplementary reads, and duplicated reads removed. Coverage in each single cell was estimated as previously described[1].

##### 2. Analysis and comparison of NO profiles derived from Strand-seq and MNase-seq

Raw reads from a previously published NA12878 MNase-seq experiment (single end, scqual and scfasta) were obtained from ENCODE (ENCSR000CXP). These MNase-seq data were generated using a SOLiD sequencer, and the data are therefore in color-space DNA sequence read format. We aligned these reads to the hg38 reference genome with bowtie (v.1.1.2)[42], using color-space read mapping enabled. After alignment, sequencing reads with poor quality (MAPQ<10), supplementary reads, and duplicated reads were removed.

```
bowtie --threads 4 -C -Q Input.csqual -f -S genome_hg38_CS Input.csfasta > output.sam
```

To obtain nucleosome positions and read depth signals (shown in **Fig. 1**, for example), the 'dpos' function provided by the DANPOS package[43] was applied to the aligned Strand-seq data, as well as the NA12878 MNase-seq data – to generate nucleosome midpoint positions genome-wide and wig files for browser track visualization with 10bp genomic bins. For the Strand-seq data, the paired-end=1 parameter was used. The Strand-seq track was haplotype-resolved into H1 and H2 tracks by pooling NO profiles generated from reads haplotype resolved using StrandPhaseR[38].

To compare nucleosomal positions obtained from pooled Strand-seq NO profiles and MNase-seq, genomic positions of human reference enhancer elements based on DNase-seq (DNase I hypersensitive sites [DHS] sequencing) and ChromHMM state analysis were downloaded [7], lifted over to hg38, and extended by 2kb centered at the midpoint. Correlation between NO in the enhancer elements based on pooled Strand-seq and MNase-seq was determined using Spearman's rho (0.68). We note that similar correlation coefficients were recently reported in a comparison of MNase-seq vs. scMNase-seq[12] – which corroborates the high quality of Strand-seq-derived NO profiles. Enhancer elements based on DHSs were defined using the Roadmap Epigenomics Consortium resource – comprising a union of 127 epigenomes in total – using data from the following URL:

https://personal.broadinstitute.org/meuleman/reg2map/HoneyBadger2_release/.

##### 3. Visualization of NO at gene bodies for genes stratified by their expression level

To visualize NO at gene bodies, each gene locus was extended by 5kb upstream of TSS and 5kb downstream of TTS. Read coverage at single base resolution was calculated on these extended loci. As each gene has different length, the coverage was normalized by fitting the coverage vector to a spline and then sampling 101 points at equal intervals. Genes were grouped into five sets based on their RNA expression level (FPKM=0, FPKM 0~0.1, FPKM 0.1~1, FPKM 1~3, FPKM>3) and their average normalized coverage was plotted as line graphs (**Fig. 1f**).

##### 4. Analysis of previously reported scMNase-seq data

Previously published single-cell MNase-seq (scMNase-seq) data from mouse cells (NIH3T3 cell line and murine naive T cells)[12] were downloaded (GSE96688) in fastq format. These raw data were aligned to the mouse reference genome (mm10) using bwa[44]. Sequencing reads with low quality (MAPQ<10), supplementary reads, and duplicated reads were removed. The mono-nucleosomal fraction was extracted (140-180bp) using samtools[45] and shell script with the following parameters.

```
samtools view -h alignment.bam | \
  awk 'substr($0,1,1)=="@" || ($9>= 140 && $9<=180) || ($9<=-140 && $9>=-180)' | \
```

```
samtools view -b > alignment_mono.bam
```

After the pre-processing, NO signals in the gene-bodies were analysed using scNOVA (in a haplotype-unaware manner) to pursue cell type classification (**Fig. S7**) and to correlate NO with gene expression (**Fig. S6**).

##### *5.* Identifying optimal parameters for inferring changes in gene activity using NO

We parameterized, and examined the performance for, inferring differentially expressed genes (DEGs) using various RPE cell lines and LCLs (see main text). The initial parameterization of scNOVA was done using RPE-1 and HG01573 (LCL): To define the ground truth set of DEGs, we compared bulk-cell RNA-seq data from RPE-1 cell line versus HG01573 using DESeq2. For scNOVA analysis, we treated the 156 single-cell libraries from RPE-1 cell line as a single "pseudo-clone", and the 46 single-cell libraries from HG01573 as a second "pseudo-clone". The CNN was trained as described in the **Methods**, to define expressed genes (EGs) and non-expressed genes (NEs) for each pseudo-clone separately. We tested different thresholds for NEs to evaluate the ideal setup. The threshold≥0.95, for example, pertains to filtering out genes whose probability to be NE (based on the CNN) is equal to or larger than 95%. After filtering out genes classified as NEs, generalized linear model analysis (as available in the DEseq2 package) was performed to compare both pseudo-clones, which yielded $\log_2$-fold changes and *P*-values. Based on this, we defined the 'differential score' as a sign of $\log_2$-fold changes multiplied by $-\log_{10}$(FDR-adjusted *P*). As a measure of prediction accuracy of the 'differential score' to infer DEGs, we calculated AUC values using different numbers of "ground truth events" (represented by the top 10 up to top 100 differentially expressed genes identified through bulk RNA-seq; see e.g. **Fig. 1**, and **Fig. S10**). These examinations revealed the best performance when using the threshold≥0.90 to filter NEs, a setting that surpassed the use of generalized linear models alone for the inference of DEGs by NO analysis. scNOVA thus uses this threshold (≥0.90) by default.

##### *6.* NO-based inference of altered gene activity by scNOVA

To identify genetic subclones, SV discovery by single-cell tri-channel processing (scTRIP) was pursued (using methods from the MosaiCatcher pipeline)[1]. Using scNOVA, genes with altered activity amongst subclones (or sets of cells in the case of CLL_24) were inferred by controlling the FDR at 10%. Genes with altered somatic copy-number were masked (removed) when investigating gene activity changes based on NO gene-bodies, since differences in copy-number status could confound differential NO measurements.

##### *7.* Molecular phenotype analysis in gene-sets in a cell lines and leukemia samples

To identify a potential upstream regulator in the NA20509 and TALL-P1 subclones, we use the molecular phenotype analysis module of scNOVA with the first mode ('gene-set overrepresentation analysis'; **Methods**). For NA20509, gene sets from the "ENCODE and ChEA consensus TFs from ChIP-X" category, provided by EnrichR[24], were used. For T-ALL P1, we used TF target gene sets curated from the literature (**Methods, Table S7**) after realising that key TFs located in the chromothriptic region are not currently annotated in EnrichR.

To infer changes of pathway activity in the single-cells harboring 10q24.32 deletions in CLL_24, we used the molecular phenotype analysis module with the second mode ('joint modeling of differential NO across predefined gene-sets'; **Methods**). Pathway level NO was compared between cells with and without 10q24.32 deletion based on linear mixed model fitting and likelihood ratio tests, using deletion status as a fixed effect and different plates (90hp1, 90hp2, 120hp1, and 120hp2) as a random effect.

#### *8.* Analysis of haplotype-specific chromatin accessibility near rearrangement breakpoints in an AML patient

To investigate haplotype-specific chromatin accessibility at the *RUNX1-RUNX1T1* locus, we defined CREs active in AML by collecting accessible chromatin regions previously profiled by subjecting AML and normal hematopoietic cells to ATAC-seq (GEO database) (**Table S5**). Consensus CREs were defined as the union of peaks detected from at least one of those ATAC-seq data sets. Haplotype-aware single-cell NO in the CREs was scaled to reads per million (RPM). NO was additionally normalized by locus copy-number, in a haplotype-aware manner. Average RPM values of single-cells at each CREs were transformed into $\log_2$ scale using a pseudocount of 1.

We also considered a sliding window (300kb in size, moving 10kb each) along the derivative chromosome, to compute chromosome-wide haplotype-specific chromatin accessibility. For each sliding window, NO at the CREs from two homologues were compared using likelihood ratio tests to obtain nominal *P*-values *[P real]*. To control the type I error from multiple testing, we performed a permutation test by shuffling haplotype labels in the single-cell RPM matrix 1000 times. For each permutation we performed likelihood ratio tests to compare NO between the two haplotypes. We then computed the number of incidences we obtained the same or lower P-value than *[P real]* from 1000 randomizations, and divided this value by the permutation trials ($N$=1000) to estimate the permutation-adjusted *P*-value.

#### *9.* Analysis of haplotype-specific chromatin accessibility near rearrangement breakpoints in T-ALL P1

To calculate haplotype-specific NO at the 2.6Mb inversion locus in T-ALL P1, ATAC-seq data of two biological replicates of TALL-P1 (EGAS00001003248)[46] were analyzed by aligning reads to the hg38 genome build using bwa[44], to define CREs in this T-ALL patient-derived sample. To do so, ATAC-seq replicates were merged using SAMtools[45]. Open chromatin regions were defined using the peak calling method provided through MACS[47] with the following parameters:

```
macs2 callpeak -t ATAC.bam -n output -g hs -q 0.05 --nomodel --shift -100 --extsize 200 -B --broad
```

As a result, genome-wide, 59,992 peaks were defined as putative CREs.

We employed scNOVA with default parameters to identify haplotype-specific NO at CREs near the breakpoints of a balanced inversion considering two rearranged TADs. We normalized NO as described above and inferred haplotype-specific NO by either considering a sliding window (300kb in size, moving 10kb each time), or CREs grouped by the nearest genes. For genes with at least two annotated CREs, differences in NO between haplotypes were evaluated using a generalized linear model likelihood ratio test, followed by FDR correction[10] (**Fig. S24**).

#### *10.* Bulk-cell RNA-seq data processing and allele-specific expression analysis

To define ground-truth differentially expressed genes between LCL and RPE-1 cell lines for parameterization of scNOVA, bulk-cell RNA-seq data was aligned to the human reference genome (GRCh38) with the STAR aligner (v2.5.3)[48], using the GTF file from ENSEMBL (GRCh38.81) with default parameters. Read counts for each gene were computed using HTSeq (v0.7.2)[49] by specifying the '-s no -t exon' option. Gene counts were normalized using the median-of-ratios method from DESeq2[20], and differentially expressed genes between conditions identified using the Wald test. To pursue allele-specific expression analysis, bulk-cell RNA-seq data were realigned to GRCh38 using GSNAP[50], using the variant-aware alignment mode to reduce allelic mapping biases. We resolved bulk-RNA-seq data by chromosome-length haplotype, using strand state and single nucleotide polymorphisms (SNPs) identified by Strand-seq (described in more detail below). ASEReadCounter[37] was used to compute allelic read counts (see also sections below).

*NA12878*. We pursued allele-specific expression analysis in NA12878 using bulk-cell RNA-seq data from ENCODE[3]. Allelic counts were assigned to either H1 (haplotype 1) or H2 (haplotype 2) based on the haplotype phasing obtained by StrandPhaseR[38]. SNP level allelic reads count were converted into gene level counts by summing up reads for each gene. Gene level counts of H1 and H2 were compared using the likelihood ratio test provided by EdgeR[9], followed by FDR adjustment. Genes with significant allele-specific expression (ASE) were identified using a FDR 10% threshold. Among ASE genes, monoallelic expressed genes were defined using definitions from a prior study, with genes showing a read-count proportion of the major allele >90% defined as monoallelic[51].

##### *11*. Bulk RNA-seq analysis in thirteen T-ALL patient-derived samples

Cells were collected from 13 pediatric T-ALL patients at the time of relapse to establish patient-derived-xenograft models as previously described[1,52]. Total RNA was extracted using TRIzol (Invitrogen Life Technologies). The RNA was then treated with TURBO DNase (Thermo Fisher Scientific) and purified using RNA Clean&Concentrator-5 (Zymo Research). We required a minimal RNA integrity number of 7, as measured using a Bioanalyzer (Agilent) with the Agilent RNA 6000 Nano kit. Cytoplasmic ribosomal RNA was depleted by Ribo-Zero rRNA Removal kit (Illumina), and RNA-seq libraries prepared from 1 µg of RNA using TruSeq RNA Library Prep (Illumina). These samples were sequenced on a Illumina HiSeq 2000 lane as 80 bp paired-end reads.

In order to confirm the subclonal perturbation of c-Myb in TALL-P1 predicted by scNOVA's infer altered gene activity module, RNA read counts were normalized using the median-of-ratios method from the DESeq2 package. Normalized read counts for each gene were standardized to obtain a Z score. After filtering out lowly variable genes (coefficient of variation (CV) of normalized read count < 25%), the average Z score of c-Myb target genes was calculated for each sample and plotted in **Fig. 5d** (we thereby considered the same c-Myb target genes as for the over-representation test in **Fig. 5c, Table S7**).

To verify allelic expression of *MYB* in the chromothripsis affected region in TALL-P1, we realigned reads to the human reference genome (GRCh38) using GSNAP as described above. Then we obtained haplotype-phased heterozygous SNP sites, based on the Strand-seq read data, using StrandPhaseR[2]. Using these phased SNP sites (input.vcf), haplotype-resolved allele-specific RNA read counts were obtained using ASEReadCounter[37], with the following parameters:

```
GenomeAnalysisTK.jar –R <reference.fasta> -T ASEReadCounter -o <output.csv> -I <input.bam> -sites <input.vcf> -U ALLOW_N_CIGAR_READS --minMappingQuality 10 --minBaseQuality 2 -drf DuplicateRead
```

SNP level allelic reads count were summarized into gene level counts by summing up reads for each gene. Gene level counts of H1 and H2 were compared using the likelihood ratio test provided by EdgeR[9], followed by FDR adjustment (**Fig. S25**).

##### *12*. Bulk RNA-seq analysis in 42 CLLs

In order to further corroborate the subclonal gene activity changes inferred in CLL_24, we performed bulk RNA-seq in a cohort of 42 CLL samples, which included CLL_24. Leukemia cells were isolated from blood using Ficoll density gradient centrifugation. Cells were viably frozen and kept on liquid nitrogen until use. Cells were thawed, allowed to recover in RPMI medium (Thermo Fisher Scientific) containing 10 % human serum (Sigma Aldrich) for 3h and filtered through a 40 µm cell strainer. Tumor cells were collected by Magnetic-activated cell sorting (MACS) using CD19 beads (Miltenyi Biotec). RNA was isolated using QIAzol Lysis Reagent (Qiagen), QIAshredder (Qiagen) and the RNeasy Mini Kit (Qiagen). Stranded mRNA sequencing, using a TruSeq Stranded Total RNA Library Preparation Kit was performed on a Illumina NextSeq 500. These RNA-seq data had originally been aligned to GRCh37.75/hg19 using STAR (v2.6.0c)[48] and counted with htseq-count[49]. For the purpose of this study, we made use of the resulting gene-level count table of protein-coding genes of interest (*i.e.*, such mapping to the relevant pathways uncovered with scNOVA). 59 CLLs were initially available to us, from which we removed *N*=8 samples exhibiting chromosome 17q13 deletions targeting *TP53*, since CLLs with *TP53* aberrations form a clinically distinct subset of CLLs[53], and since cross talk between the p53 and Wnt signaling has been reported, with p53 loss promoting Wnt signaling[54,55]. We additionally removed *N*=9 samples exhibiting trisomy 12, as this group of samples is known to express a unique set of pathways when compared to other CLL samples[56]. RNA read counts of the remaining 42 CLLs were normalized, Z-scores were derived, and lowly variable genes were filtered out as described above. To measure Wnt signaling activity from these transcriptomic data, we obtained 49 known target genes of TFs involved in Wnt canonical signaling (*CTNNB1*, *LEF1*, *TCF7*, and *TCF7L2*) from the TRRUST database[57], here called 'Wnt signaling target genes (**Table S7**)'. The mean Z score of 49 genes was computed and visualized. CLL_24 showed the most pronounced bulk RNA-seq overexpression of Wnt signalling pathway members (ranking first amongst all of the 42 considered CLL samples of this cohort; **Fig. 3f**).

We also analysed bulk RNA-seq data of CLL patients from the ICGC resource[29], by considering 395 CLL samples with available SCNA data. To be consistent with the analysis of CLL primary samples from the Heidelberg-based cohort, we removed samples exhibiting trisomy 12 or 17q13 deletions affecting *TP53*, which yielded 306 donors in total. We observed somatic 10q24.32 losses (deletion) in six out of 306 donors (**Fig. 3b**). Among those 306 donors, bulk RNA-seq data was available from 178 donors including 4 donors with 10q24.32 losses overlapping the minimal deleted regions defined in CLL_24. RNA read count of the 178 CLL samples were normalized, Z scores were derived, and lowly variable genes filtered out as described above. Mean Z score of Wnt signaling target genes of the donors with and without 10q24.32 deletions were compared using the generalized linear model (GLM) likelihood ratio test, controlling for gender and age (**Fig. 3h**).

##### *13.* Haplotype-resolved bulk RNA-seq analysis in LCLs from HGSVC consortium

To verify the subclonal activation of c-Myc/Max in NA20509, we analyzed bulk RNA-seq data in the whole panel of LCLs recently used by the Human Genome Structural Variation Consortium (HGSVC) to construct an SV germline reference resource[28]. RNA read count of the 33 LCLs were normalized, Z scores were derived, and lowly variable genes were filtered out as described in section 10. The mean Z score of c-Myc/Max target genes was calculated and visualized using a bargraph (**Fig. 2d**). c-Myc/Max heterodimer target genes were downloaded from the Molecular signatures database (Msigdb, **Table S7**)[58].

To examine the allelic expression of genes residing in regions of complex chromosomal rearrangement in NA20509, we firstly detected SNP sites from the pooled NA20509 Strand-seq libraries using freebayes[59]. Using these SNP sites (input.vcf), allele specific read counts were obtained using ASEReadCounter[37] with the following parameters:

```
GenomeAnalysisTK.jar –R <reference.fasta> -T ASEReadCounter -o <output.csv> -I <input.bam> -sites <input.vcf> -U ALLOW_N_CIGAR_READS --minMappingQuality 10 --minBaseQuality 2 -drf DuplicateRead
```

To haplotype-resolve allelic counts to either the unaffected homolog (haplotype 1) or the homolog bearing the BFB (haplotype 2, *i.e.* the derivative chromosome), we used the strand states of Strand-seq reads (Watson (W) or Crick (C)) along chromosome 17. In single cells in which the two homologs of chromosome 17 have a WW majority configuration[1] (WW strand state seen for most of chromosome 17), the BFB-mediated inverted duplication[1] will always exhibit DNA reads on the C strand belonging to haplotype 2 (the same applies to single cells with a CC majority configuration on chromosome 17, for which W reads belonging to haplotype 2 can be extracted from the inverted duplication). Among the 40 single-cells containing the BFB event, we could collect 14 cells with either a WW or CC majority configuration, allowing extraction of reads from the derivative chromosome haplotype; similarly, we collected 8 cells with either a WW or CC majority configuration for chromosome 5, which enabled extraction of reads from the terminally duplicated haplotype. These operations unambiguously phase-resolved 138 genes located on the derivative chromosome, with at least 2 phased heterozygous SNPs seen for *MAP2K3, MAPK9,* and *PDGFRB*. RNA phased-resolved allelic read counts for these SNPs were compared using the likelihood ratio test followed by FDR-adjustment (**Fig. 2e, S15i**).

##### *14.* Clinical diagnostic information for CLL_24

CLL_24 was obtained from the peripheral blood mononuclear cells of a previously untreated female CLL patient with age 61 at sampling. According to routine diagnostic methods, the patient sample showed no IGHV hypermutation, had no *TP53* mutation, and had no alterations at 6q21, 8q24, 11q22.3, 12q13, 13q14 and 17p13.

##### *15.* Clinical diagnostic information for AML_1

This sample was obtained as a diagnostic bone marrow from the first aspiration of an AML with a t(8;21) translocation (known to result in *RUNX1:RUNX1T1/ETO:AML1* gene fusion), arising after cytostatic therapy for testicular cancer in a young man. 95% of cells were identified as blasts with monocyte differentiation by microscopy. In the initial diagnostic flow cytometry characterization, 65% of cells showed monocyte differentiation markers and were positive for CD33, CD13, CD38, HLA-DR, CD11c, and CD15. A subpopulation of these blasts were CD34+/CD117+ and also partly positive for CD19 (common in t(8:21) AML). This sample also carried a *FLT3*-TKD mutation (p.Asp835Tyr, CF=44%).

##### *16.* 10q deletion discovery in CLL samples from PCAWG

To assess the frequency of deletions affecting 10q24.32 in CLL we analyzed 94 CLL samples included in the ICGC/TCGA Pan-Cancer Analysis of Whole Genomes (PCAWG) resource [30]. We first generated GC and mappability corrected fragment counts for the paired-end WGS of each sample using Delly[27]. These normalized fragment counts were then binned in 10kbp windows and screened for chr10q deletions, using Delly2[27]. As shown in **Fig. S19**, at least 4 out of the 94 CLL samples (4.3%) harbor somatic deletions intersecting with the 10q24.32 minimal region.

##### *17.* Strand-seq in a panel of lymphoblastoid cell lines (LCLs)

24 EBV-transformed LCLs (Coriell Institute) were cultured in BrdU (100uM concentration; Sigma) for 18 or 24 hours, and single isolated nuclei (0.1% NP-40 lysis buffer[60] sorted into 96-well plates using the BD FACSMelody cell sorter (NA12329, NA18534, NA18939, NA19650, NA19983, NA20509, NA20847, HG00096, HG00171, HG00864, HG01114, HG01505, HG01596, HG02011, HG02492, HG02587, HG02818, HG03009, HG03065, HG03125, HG03371, HG03486, HG03683, HG03732). In each sorted plate 94 single cells, one 100-cell positive control and one 0-cell negative control were deposited. Strand-seq libraries were prepared, sequenced and selected using the same protocol as for the primary leukemia samples. These LCLs were previously released[28] and used to construct a haplotype-resolved germline SV resource in a human population diversity panel by the Human Genome Structural Variation Consortium[28] A mean of 54 high-quality single cells (41 to 71 cells) were sequenced to a median depth of 338,271 mapped nonduplicate fragments per cell. We used a threshold of CF≥10% for discovering unwanted somatic SVs in these LCLs, using the scTRIP method[1]. Translocation discovery was pursued using the 'majority mode'[1].

#### *18.* WGS-based subclonal SV analysis in NA20509

The availability of NA20509 WGS data from the New York Genome Center[61] allowed us to attempt verification of the presence of somatic SVs. We first aligned the data to GRCh37 using bwa[44], called SNPs and InDels using FreeBayes[44] and haplotype-phased variants using eagle2[62] using the 1KG phase 3 reference panel[41]. Phased haplotype blocks were used to identify heterozygous sites deviating from the expected 1:1 ratio, in conjunction with GC and mappability corrected read-depth plots (**Fig. S14**) generated using Delly2[27]. Phased heterozygous sites and read-depth estimated copy-numbers deviated from the expected pattern for copy-number 2, and thus independently confirmed the presence of subclonal SVs in NA20509, including the terminal gain on chromosome 5, and the terminal loss of the chromosome 17 p-arm with an adjacent gain event (**Fig. S14**). SV analysis using Delly2[27] also verified the presence of a sub-clonal translocation from chromosome 17 (position 21,479,415) to chromosome 5 (position 132,093,890). Delly2 further revealed a tail-to-tail inversion-type rearrangement at chromosome 5 with breakpoint positions 132,029,510 and 132,053,373, and a tandem duplication-type rearrangement spanning the terminal gain on chromosome 5 with breakpoint positions 133,831,567 and 178,087,753 (GRCh37 coordinates).

#### *19.* Manual curation of somatic SVs in LCLs to achieve a high-quality callset

We used scTRIP (methods from the MosaiCatcher pipeline)[1] to discover somatic SVs in LCLs. The extent and diversity of deletions at 22q11.2 intersecting the immunoglobulin lambda locus (*IGL*) motivated additional curation of these somatic SV events. 17/25 LCLs harbored deletions at 22q11.2. Making use of the somatic SV calls and single-cell segmentation results from the MosaiCatcher pipeline[1], we defined putatively deleted segments, which we subjected to manual inspection in single cells followed by reanalysis using the ArbiGent SV genotyping tool[28]. ArbiGent performs analysis of Strand-seq data cell-by-cell, to allow precise haplotype-resolved genotype assignment into homozygous and heterozygous deletion events, at defined genomic intervals. These genotype assignments were accepted if the log10 likelihood ratio between SV and reference state was greater than 0.5, and as such – in a few cases – superseded variant calls created with the MosaiCatcher pipeline.

*Additional curation and analysis of NA12878 somatic SVs.* NA12878 is perhaps the single most sequenced human cell line presently existing[3,41,63]. We thus regarded the discovery of previously unknown somatic SVs in this cell line as a surprise, which motivated careful curation and manual inspection. Altogether, we analysed 75 Strand-seq libraries from NA12878, including cells prepared as a single batch[2], to allow for robust single-cell somatic SV discovery. 59 single-cell genomes harbored a somatic 22q11.2 deletion (chr22:22200000-22900000) on haplotype 2 (H2) intersecting the *IGL* locus. Application of scNOVA revealed differential gene activity patterns between these *IGL* locus deletion bearing cells (denoted clone 2), and cells unaffected by 22q11.2 SVs (clone 1) – revealing 8 significant genes (10% FDR). Out of these 8 genes, 5 genes were located in a short chromosome 19 interval and inferred to be less occupied by nucleosomes due to the decrease of read depth in clone 1. Prompted by this observation, we performed manual inspection of this region, which revealed that clone 1 harbors a ~500kb hemizygous deletion on chromosome 19, which confidently maps to haplotype 2 (H2), but which was missed in the initial single-cell population segmentation pursued using the MosaiCatcher pipeline[1]. Separate segmentation of individual cells using the MosaiCatcher pipeline fine-mapped this candidate deletion to a 500kb interval (chr19:36500000-37000000), thus corroborating its presence by single-cell SV discovery. We further subjected this candidate interval to cell-by-cell genotype analysis using ArbiGent[28], which in line with the MosaiCatcher pipeline genotyped a high-confidence somatic deletion at chr19:36500000-37000000. ArbiGent genotype calls were accepted in cells where the log10-likelihood-ratio between reference and deleted states was larger than 0.5, allowing confident genotyping in 65/75 (87%) of cells. ArbiGent revealed strict mutually exclusivity between this interstitial deletion on chromosome 19, and 22q11.2 somatic deletions (**Table S8**) – mirroring the pattern we had observed for NA20509. As described in the section below, this chromosome 19 deletion is also verified by scRNA-seq data from different NA12878 cell stocks. Finally, we further used Arbigent to analyse 20 additional NA12878 Strand-seq libraries, prepared as a separate biological replicate[2], which once again showed the presence of subclones carrying mutually exclusive interstitial deletions on chromosome 19 and at 22q11.2. These results hence show the presence of different somatic subclones in the key NA12878 human reference model cell line.

#### *20*. scRNA-seq data analysis for inferring somatic copy-number alterations (SCNAs)

*Genotyping mode*

Single-cell count matrices were normalized to obtain count per million (CPM) values. These values were converted into $\log_2$(CPM/10+1) – the standard input value for CONICSmat, a tool for inferring ('genotyping') the copy-number state of chosen candidate genomic regions using scRNA-seq data[21]. For each of the candidate SV regions, we applied CONICSmat, with parameters set to allow considering regions with at least 10 expressed genes. Firstly to verify the presence of SCNAs in regions of interest, CONICSmat generates distributions of average expression levels across single-cells, and then fits to the 1-component and 2-component mixture models[21]. It further compares the likelihood ratios of being 1-component (unimodal, absence of subconal SCNAs) and 2-component (bimodal, presence of subclonal SCNAs) to determine the most likely state in those regions based on the Bayesian information criterion (BIC). Candidate SCNAs likely to be bimodal using a 1% FDR criterion were considered further for downstream analysis. For those candidate SCNAs, the posterior probability of each single-cell to be belonging to the SV component, and the 'wildtype' (WT) (non-deletion) component was computed. Single-cells with a posterior probability above the cutoff for one of these two components were used in downstream analyses.

*Discovery mode*

Three broadly used single-cell transcriptome based SCNA analysis tools InferCNV[64], HoneyBADGER[65], and CONICSmat[21] were used for SCNA discovery. As inferCNV and HoneyBADGER require matched normal cell annotations, we first defined normal cell population if it's available within the same sample. If not, we downloaded cell-type matched normal cell profiles from the GEO database as reported in **Table S4**.

To run InferCNV, we provided single-cell count matrices with analysis_mode = 'subclusters', cutoff=0.1 for 10X, cutoff=1 for SMART-seq and Fluidigm as recommended in the manual. For HoneyBADGER, CPM normalized single-cell count matrices were converted into log(CPM+1), and put into HoneyBADGER for the CNV discovery with default parameters. CONICSmat is originally developed for 'genotyping' mode for estimating copy number of candidate SCNA regions obtained from DNA-seq data, however it also provides chromosome-arm level SCNA discovery in case no matched DNA-seq data is available. For this, above mentioned $\log_2$(CPM/10+1) – the standard input values for CONICSmat were put into CONICSmat for the chromosome-arm level discovery.

#### *21*. Pseudotime/cell-type analysis of scRNA-seq data

Pseudotime analysis was carried out using the R package Slingshot[40], implementing UMAP and cell clusters identified using Seurat (see Methods for further details.). Cell type analysis was performed using UCell (as per main methods), using the T cell differentiation stage-specific gene sets outlined in [39].

#### Supplementary Discussion

##### *I*. Scope of scNOVA compared to single-cell multiomics methods focusing on SCNAs

To evaluate the scope of scNOVA in relation to prior single-cell multiomics method focusing on large-scale (>10Mb) SCNAs, we analysed the abundance of different classes of somatic driver mutation, differentiated by variant class and size, using the PCAWG resource of 2,658 whole-cancer genomes[30]. We downloaded the PCAWG set of patient-centric drivers (https://dcc.icgc.org/releases/PCAWG/driver_mutations)[30,66] to compute the relative abundance of driver mutation classes: Across PCAWG, out of a total of 13,219 annotated drivers, 5,913 represent somatic point mutations (including single nucleotide variants, SNVs; multi-nucleotide variants, MNVs; and <50bp short insertions and deletions (indels)), 6,490 represent copy-number imbalanced SVs and another 816 are copy-number balanced SVs. Hence somatic SVs (defined as copy-number imbalanced+copy-balanced SVs) contribute 55% of all annotated somatic drivers, and significantly outnumber compared to point mutations in a typical cancer genome[30]. Second, to enable comparison of SV drivers by size and class we generated a consolidated list of somatic SV drivers from PCAWG with resolved breakpoint coordinates, and computed confident size estimates from these (achieved for *N*=3,342 SVs). Out of these, 83% (2,765 SVs) were ≥200kb in length and thus at the size range[1] accessible to scNOVA. By comparison, 37% (1,244 SVs) represented SCNAs ≥10Mb in length.

##### *II*. Known and suspected Wnt signaling regulators near 10q24.32

A closer analysis of genes located within the relevant 10q arm showed that several Wnt signaling genes reside in this genomic region (**Fig. 3b, S17**). Indeed, five genes implicated or suspected to suppress canonical Wnt signaling, *SUFU* (deleted in 11/11 cells with 10q24.32 SVs), *FBXW4* (11/11), *NFKB2* (11/11), *BTRC* (9/11) and *LZTS2* (5/11), were repeatedly deleted in CLL_24, and represent candidates for further study: *BTRC*, an F-box containing protein, has been shown to be involved in ubiquitin mediated proteolysis of β-catenin to maintain low levels of β-catenin in the cytoplasm, and as such influence Wnt signaling[67]. *BTRC* has been reported as mutated in multiple human cancer cell lines and clinical tumor samples[68], is located close to the 10q24.32 minimal region (<60kb apart), and is significantly downregulated in CLLs from the ICGC that harbor a 10q24.32 somatic deletion (FDR-adjusted *P*=0.000646; **Fig. S17**). *FBXW4*, located within the minimally deleted 10q24.32 region, has been implicated in tumor suppression [69] and as another F-box containing protein has been suspected, albeit not yet shown, to play a roles in ubiquitin mediated proteolysis of β-catenin[70]. Like *BTRC*, we observed *FBXW4* to be significantly downregulated in CLLs from the ICGC that harbor a 10q24.32 somatic deletion (FDR-adjusted *P*=0.00478; **Fig. S17**). *SUFU*, a tumor suppressor gene located in the minimally deleted region, can suppress Wnt signaling by forming a complex with β-catenin, and by enhancing β-catenin translocation to the cytoplasm[71]. *SUFU* loss of function has been shown to result in the failure to suppress Wnt signaling in medulloblastoma (with loss of function of *SUFU* mediating overactivity of both the Sonic Hedgehog signaling pathway and the Wnt signaling pathway in medulloblastoma)[72]. *LZTS2*, deleted in 5/11 cells exhibiting 10q24.32 somatic deletions, is likewise a tumor suppressor and a known negative regulator of Wnt signaling[73]. *NFKB2*, located within the minimally deleted 10q24.32 region, was previously proposed – albeit not yet shown – to be involved in the pathogenic effect of 10q24 deletion in CLL[74–76]. *NFKB2* is a gene that encodes a subunit of the transcription factor complex nuclear factor-kappa-B (NF-kB) which is essential for lymphocyte development and immune function[77]. NF-kB signaling negatively regulates the Wnt/β-catenin pathway either indirectly through the functions of NF-kB target genes (e.g., *LZTS2*) or directly by interfering with the formation of transcriptional complex β-catenin/TCF/p300[78].

#### *III*. Nucleosome repeat length measurements: considerations for future users

The main text reports very similar nucleosome repeat length[5] estimates when comparing Strand-seq (195.4 ±0.4bp) and MNase-seq (193.7 ±0.6bp). Thus, nucleosome repeat length estimates from Strand-seq are, encouragingly, highly reproducible and consistent with bulk MNase-seq; this formed the foundation of our work. Some considerations on the measurement of nucleosome repeat length are given here, to guide future users:

During Strand-seq library preparation single-cell genomic DNA is fragmented using MNase. As described extensively in [60], the protocol calls for precisely 0.5U of MNase per single cell reaction, adding the enzyme immediately after diluting into fresh Master Mix. The incubation time should then proceed for precisely 8 minutes before stopping the reaction by addition of EDTA. All steps of MNase digestion reaction were previously optimized to generate mononucleosomal fragments of 150~200 bp in length. The timing and enzymatic activity are critical for the digestion procedure, and therefore any deviations in these components can result in changes in the DNA fragmentation pattern seen in the sequenced libraries. Furthermore, after library preparation there is a final size-selection step, upon pooling the single cell libraries. During this step the 250-350bp band (including adapter sequences) is specifically excised to enrich the mononucleosome fragments. Changes to the size selection step or possible fragment contamination during size selection can change the fragment lengths present in the final sequence data, for instance if dinucleosomal fragments are also included in the excised band (e.g. as seen in the NA12878 raw data, before processing; see **Fig. S2**). Looking forward, we advise users to carefully control the size selection step and the digestion procedure, to ensure that the same reproducibility of nucleosomal patterns as observed for our automated Strand-seq library generation pipeline[1] can be achieved.

#### *IV*. Further details of functional outcomes of somatic rearrangement landscapes in lymphoblastoid cell lines

17 LCLs exhibited deletions 200–700kb in size, comprising the *IGL* locus on chromosome 22q11, present at CFs from 1.4% up to 89%. 13 LCLs (52%) showed homozygous loss events, and 8 LCLs (32%) exhibited two or more subclones harboring distinct 22q11.2 deletions (**Table S3, Fig. S12**). 22q11.2 undergoes V(D)J rearrangements during B-cell development, a physiological process resulting in up to ~1Mb-sized deletions[79], suggesting that these SVs likely emerged as a result of normal B-cell biology. However, since the expression changes conferred by 22q11.2 deletions in cultured LCLs are unknown, this alteration provided a test case for linking SV to a molecular phenotype. Using scNOVA, we inferred the lncRNA *FLJ22447* (NCBI Gene ID: 400221), present on 14q23, to be upregulated in cells with the 22q11.2 deletion compared to cells with a normal 22q status (**Table S2**). Although so far only poorly characterized, this lncRNA was shown to be upregulated in carcinoma-associated fibroblasts where it reinforced interleukin-33 signalling, promoted a proliferation phenotype, and correlated with poor disease prognosis in oral cancer[80]. How this lncRNA is dysregulated via 22q11.2 deletion remains to be seen, but it is possible that overexpression of this gene may support the expansion of cultured LCLs harbouring 22q11.2 SVs.

#### *V*. Regulatory landscape changes mediated by a 14q32 inversion in T-cell acute lymphoblastic leukemia (T-ALL)

We recently reported a 2.6 Mb sized 14q32 inversion in a T-ALL patient-derived xenograft (PDX) sample (TALL-P1), resulting in monoallelic *TCL1A* oncogene expression[35]. As changes to the nucleosome occupancy landscape emerging from this somatic inversion remain unclear, we considered it another suitable test case for scNOVA. We generated pseudo-bulk phased NO tracks by merging 77 Strand-seq libraries[35] sequenced from TALL-P1, and then used scNOVA to measure haplotype-specific NO across both of the fused TADs that emerged from this inversion (**Fig. S24**). This analysis yielded a region ~500kb upstream of *TCL1A*, falling into a gene desert near the distal inversion breakpoint, with increased NO, suggesting reduced chromatin accessibility (*P*<0.001; permutation-adjusted likelihood ratio test). A second significant region, comprising *TCL1A* and several proximal CREs (defined in bulk-cell ATAC-seq[46]) yielded reduced NO on the rearranged haplotype H2 (*P*=0.021; permutation-adjusted likelihood ratio test; **Fig. S24**), suggesting increased chromatin accessibility, which would be consistent with the outlier expression of *TCL1A* previously detected for this haplotype. We tested all CREs located within the two TADs rearranged by this inversion for haplotype-specific NO, to uncover elements that may foster aberrant oncogenic activity as a result of the SV. This analysis revealed a significant haplotype-specific NO decrease for two CREs assigned to *TCL1A*, indicative for increased CRE accessibility (*P*=0.000000045; FDR-adjusted likelihood ratio test; 2.6-fold change) – and thus uncovered elements likely to foster or support aberrant *TCL1A* expression as a result of the SV. *TCL1A*, notably, was the only gene inferred to be targeted by CREs with haplotype-specific NO at 14q32 (**Fig. S24**) indicating high specificity of predictions made with scNOVA, since *TCL1A* is the only gene exhibiting outlier monoallelic expression from the rearranged 14q32 region (**Fig. S24**).
