## Supplementary Data for "Haplotype-aware single-cell multiomics uncovers functional effects of somatic structural variation"

### Snapshots of somatic SV events in LCLs

**Single-cell SV events in LCLs (NA12329, 59 single cells profiled)**

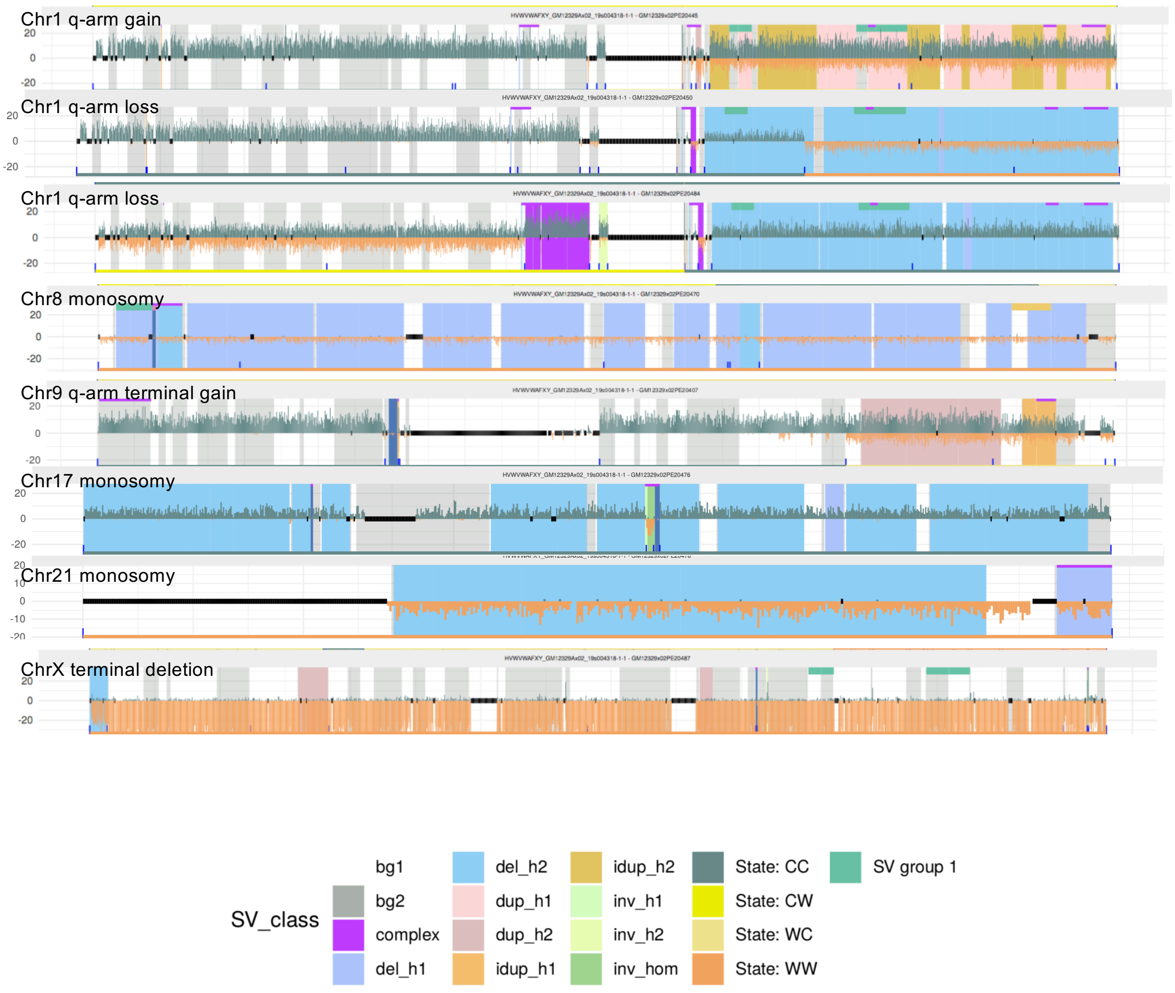

### Single-cell SV events in LCLs (NA18534, 56 single cells profiled)

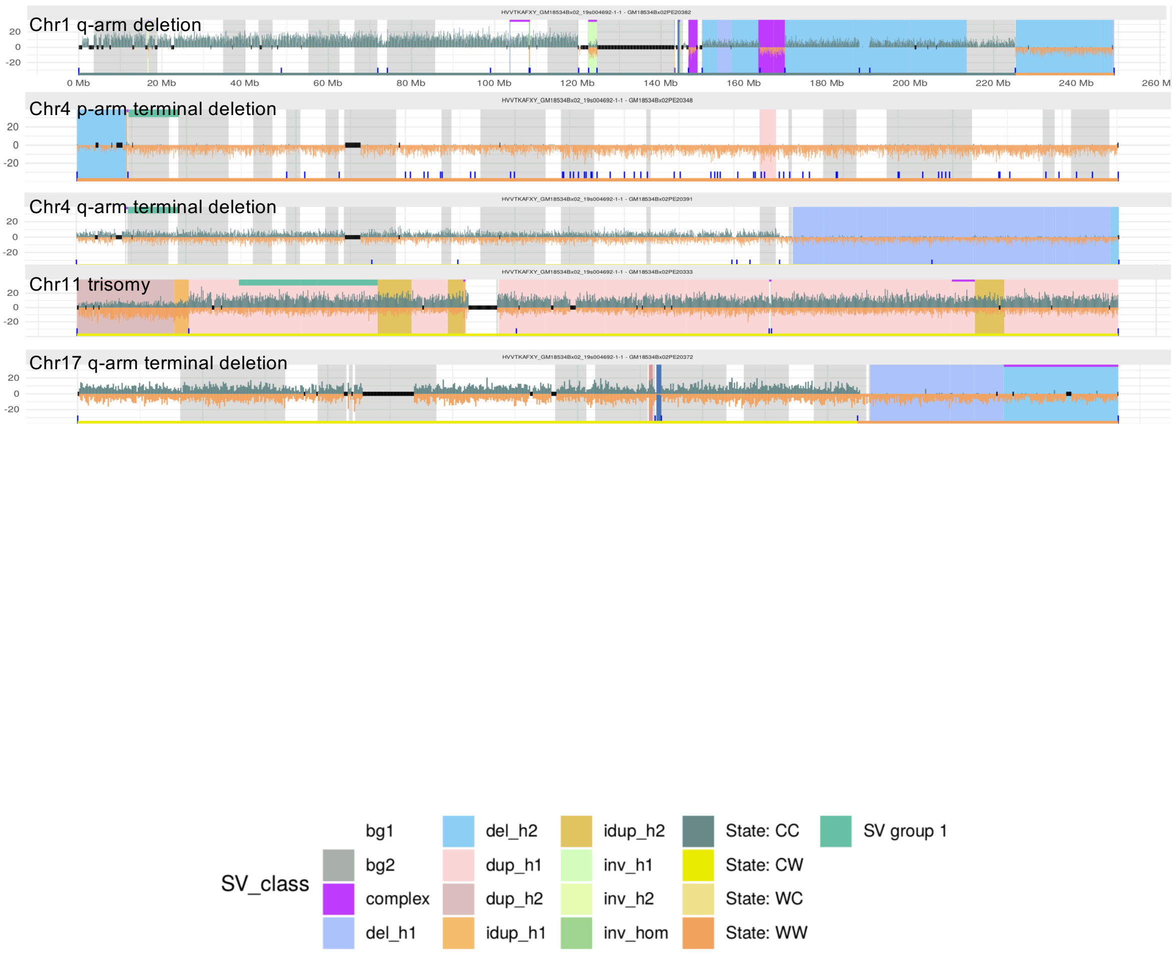

### Single-cell SV events in LCLs (NA18939, 41 single cells profiled)

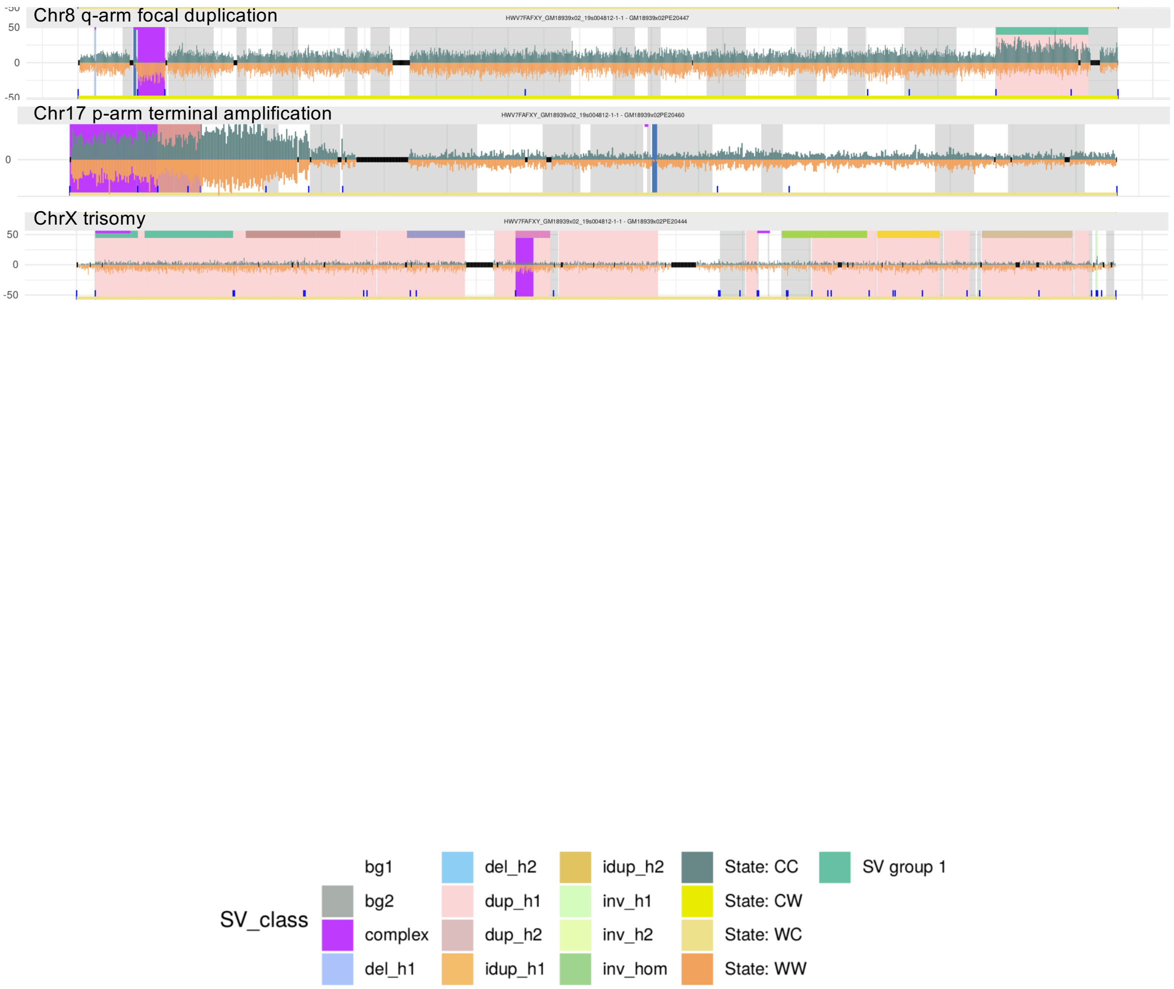

### Single-cell SV events in LCLs (NA19650, 55 single cells profiled)

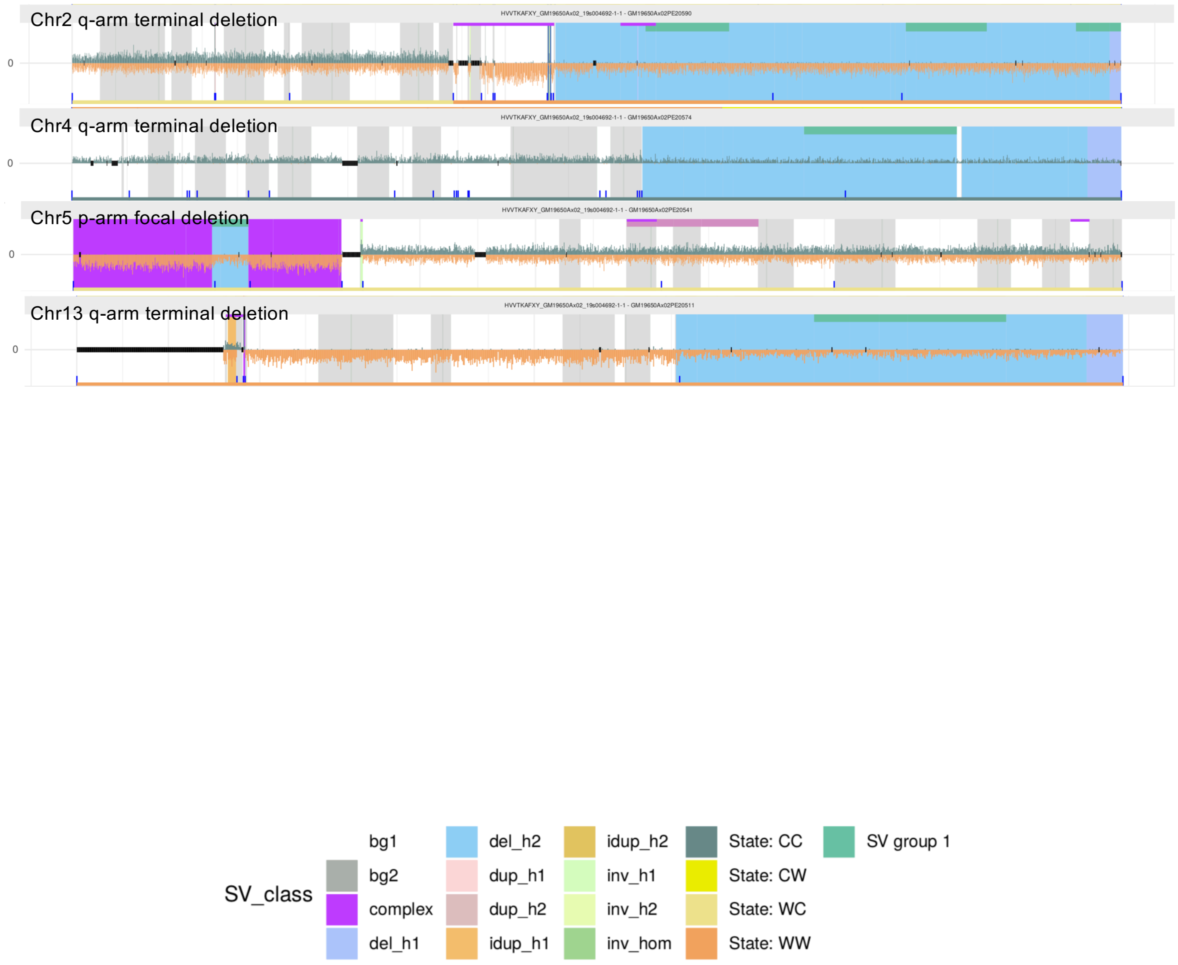

#### Single-cell SV events in LCLs (NA19983, 45 single cells profiled)

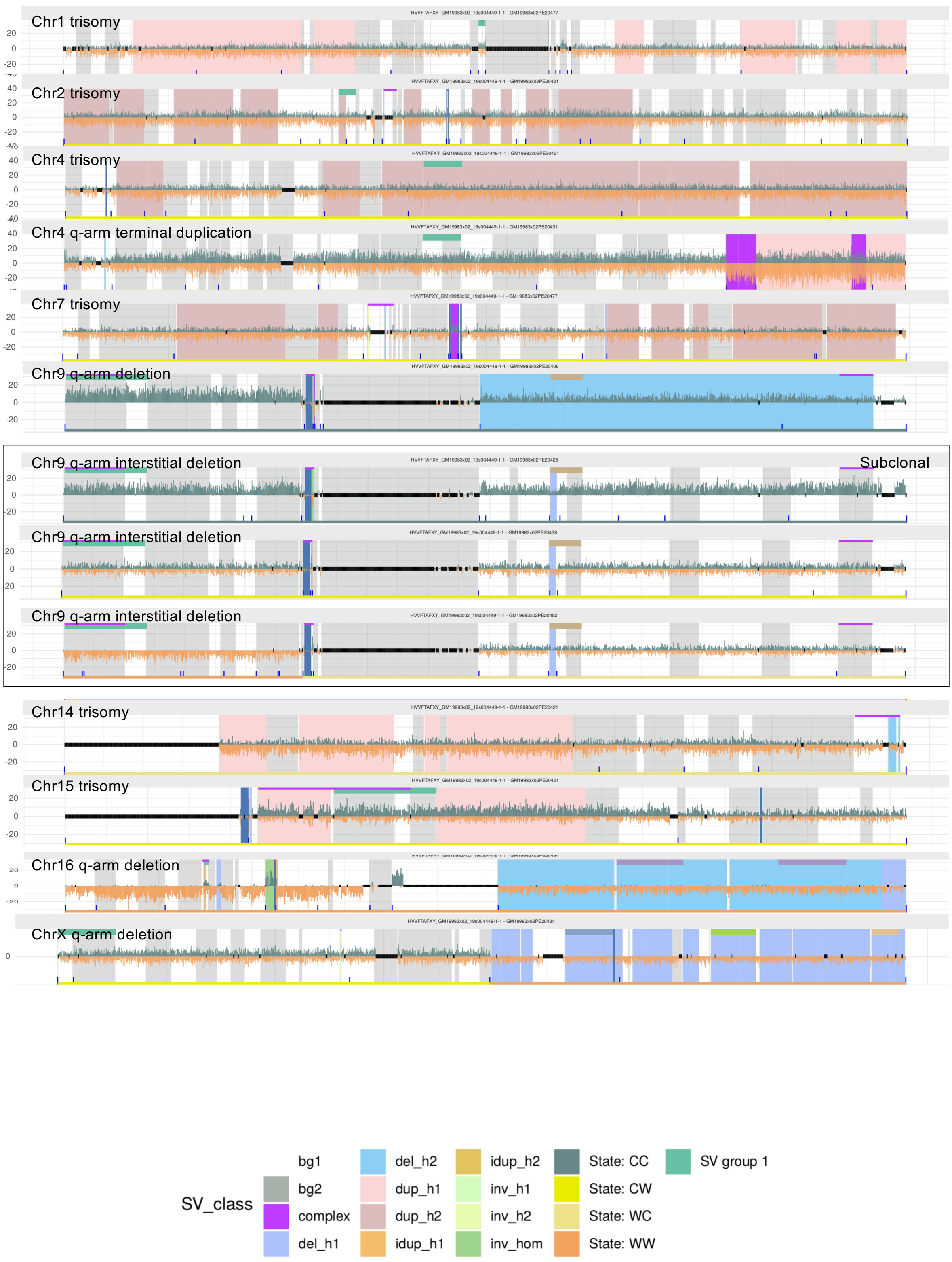

### Single-cell SV events in LCLs (NA20509, 47 single cells profiled)

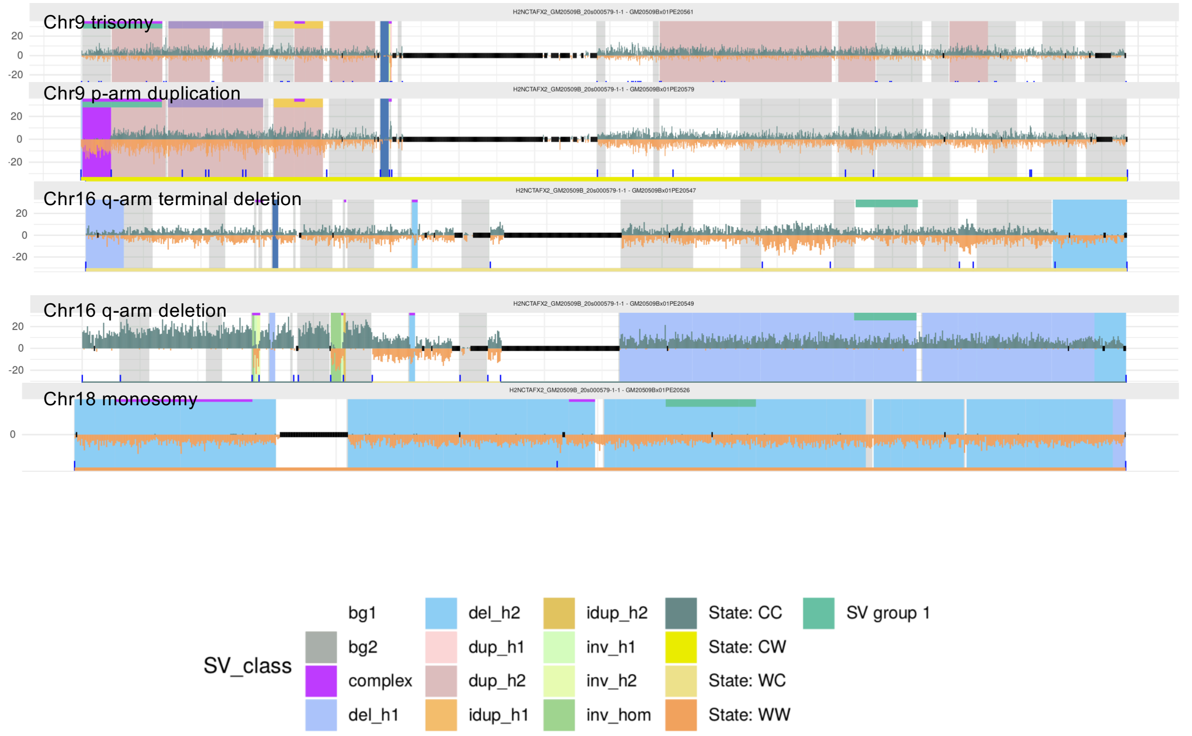

### Single-cell SV events in LCLs (NA20847, 55 single cells profiled)

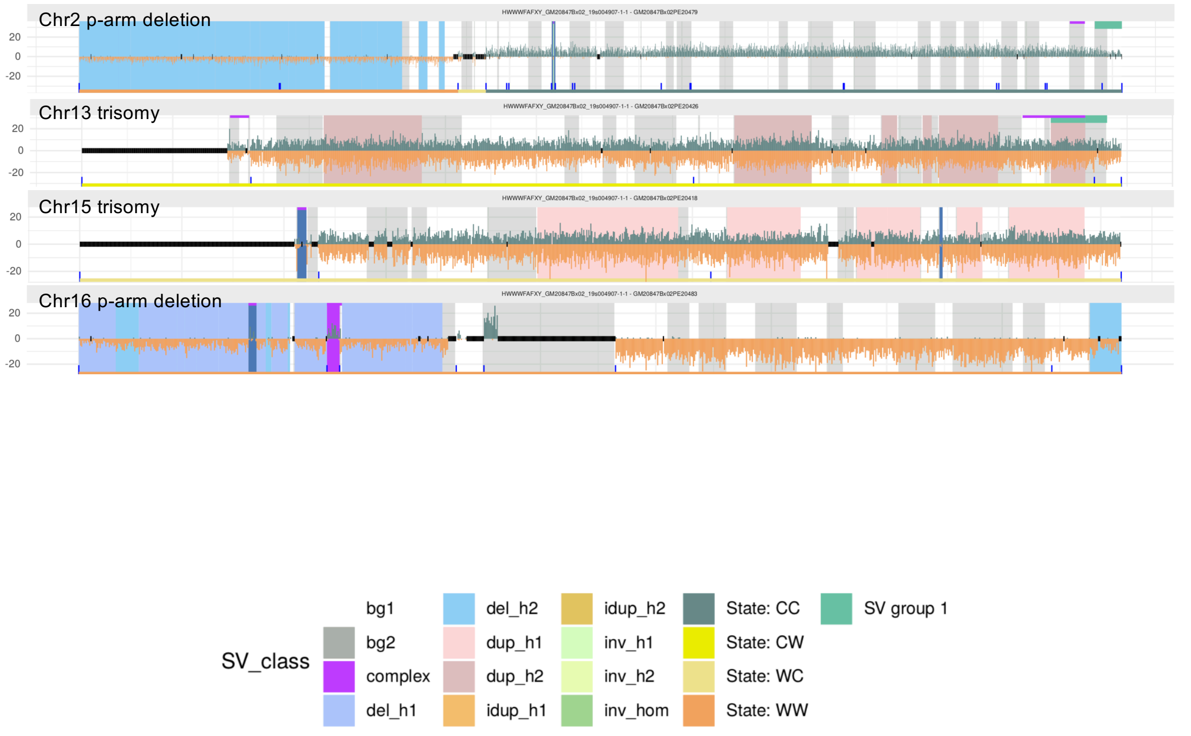

### Single-cell SV events in LCLs (HG00096, 69 single cells profiled)

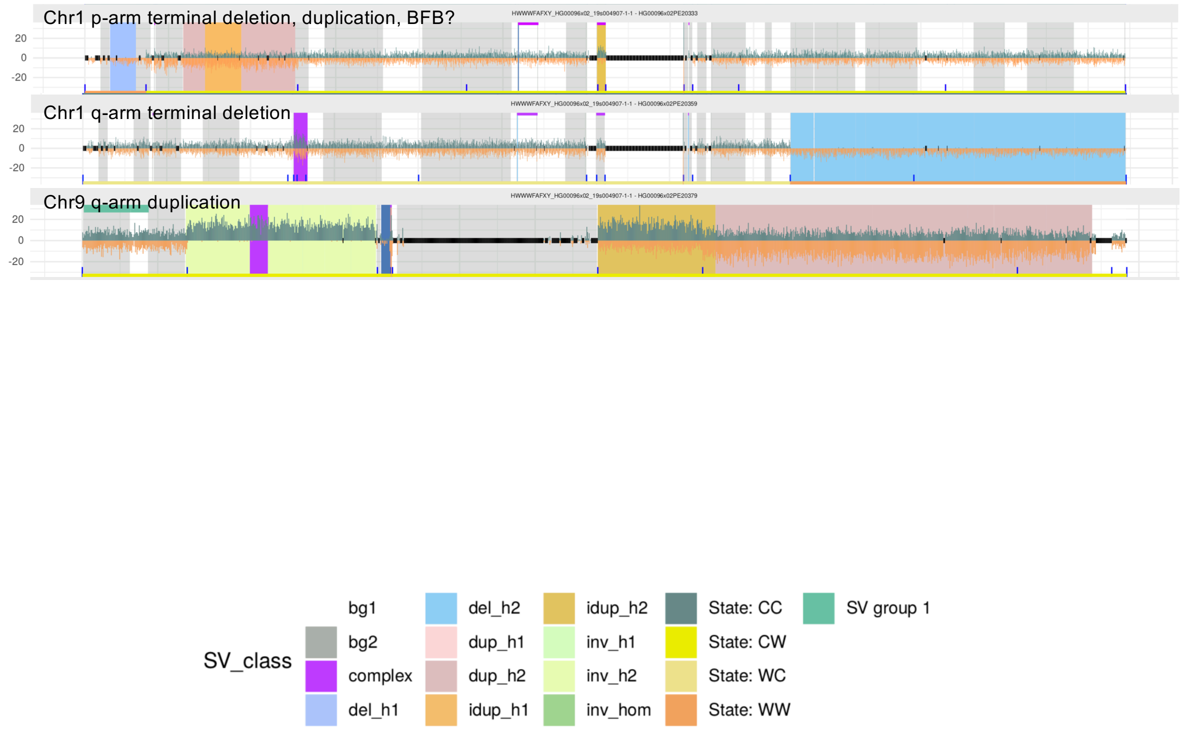

#### Single-cell SV events in LCLs (HG00171, 49 single cells profiled)

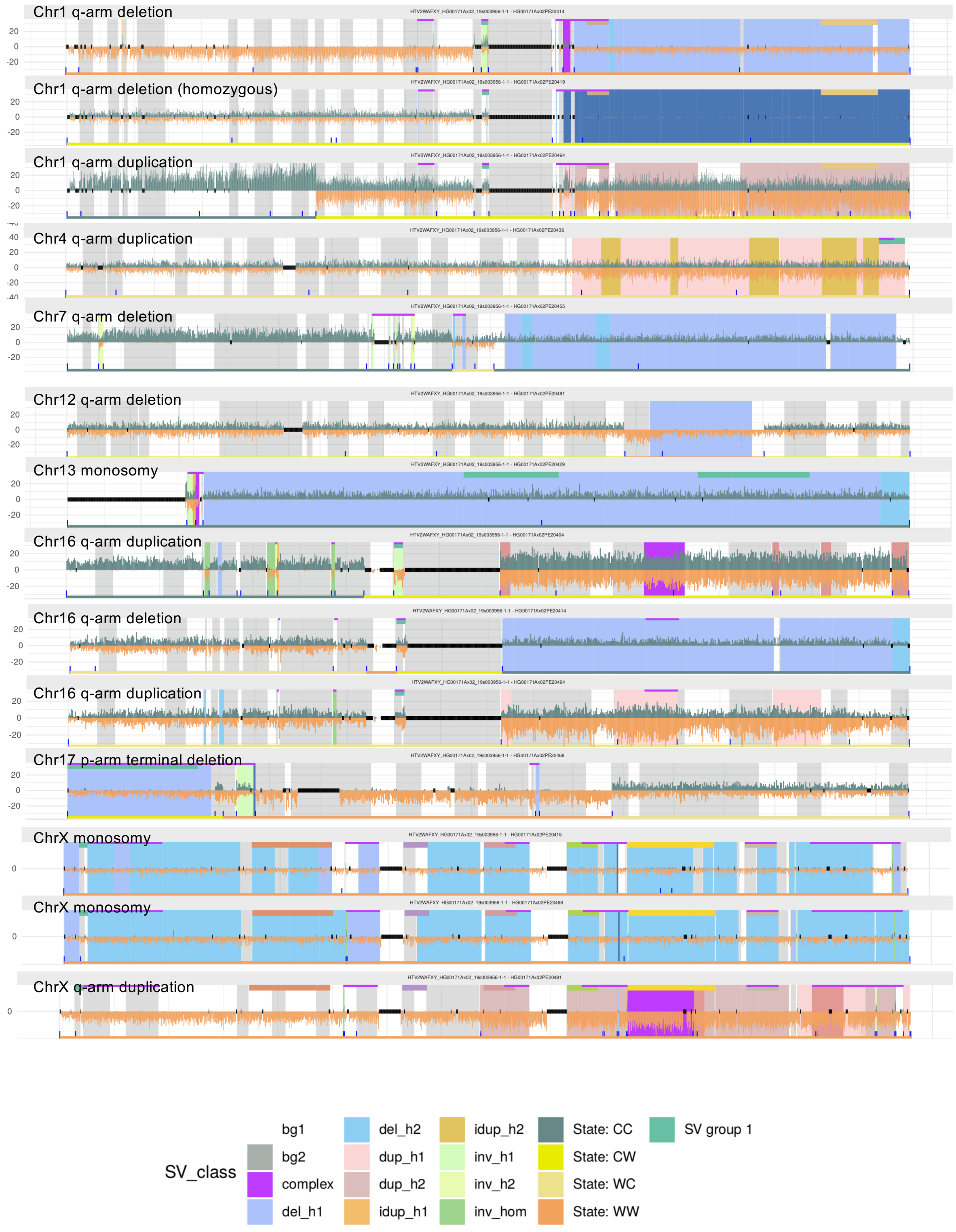

### Single-cell SV events in LCLs (HG00864, 43 single cells profiled)

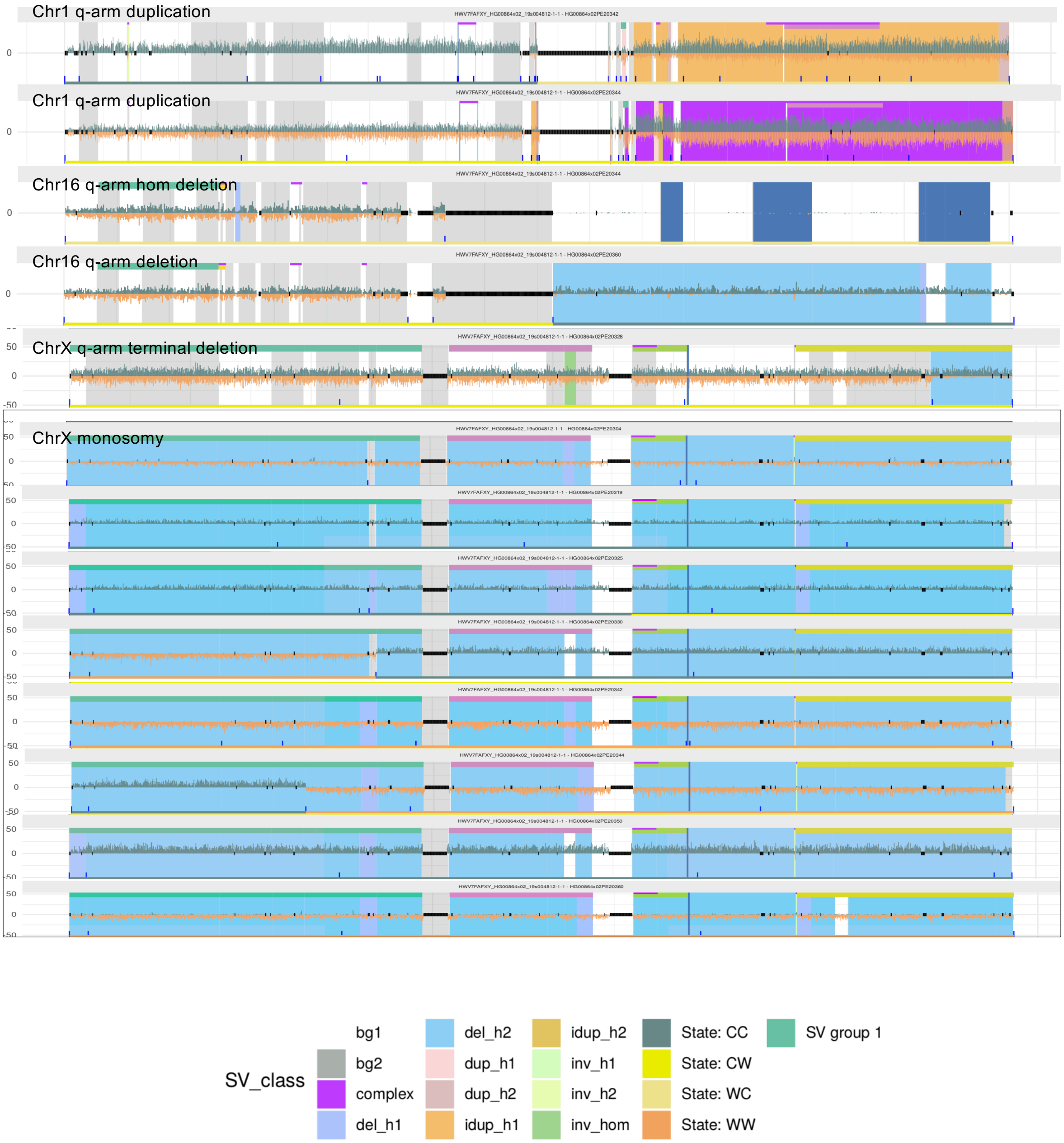

### Single-cell SV events in LCLs (HG01114, 51 single cells profiled)

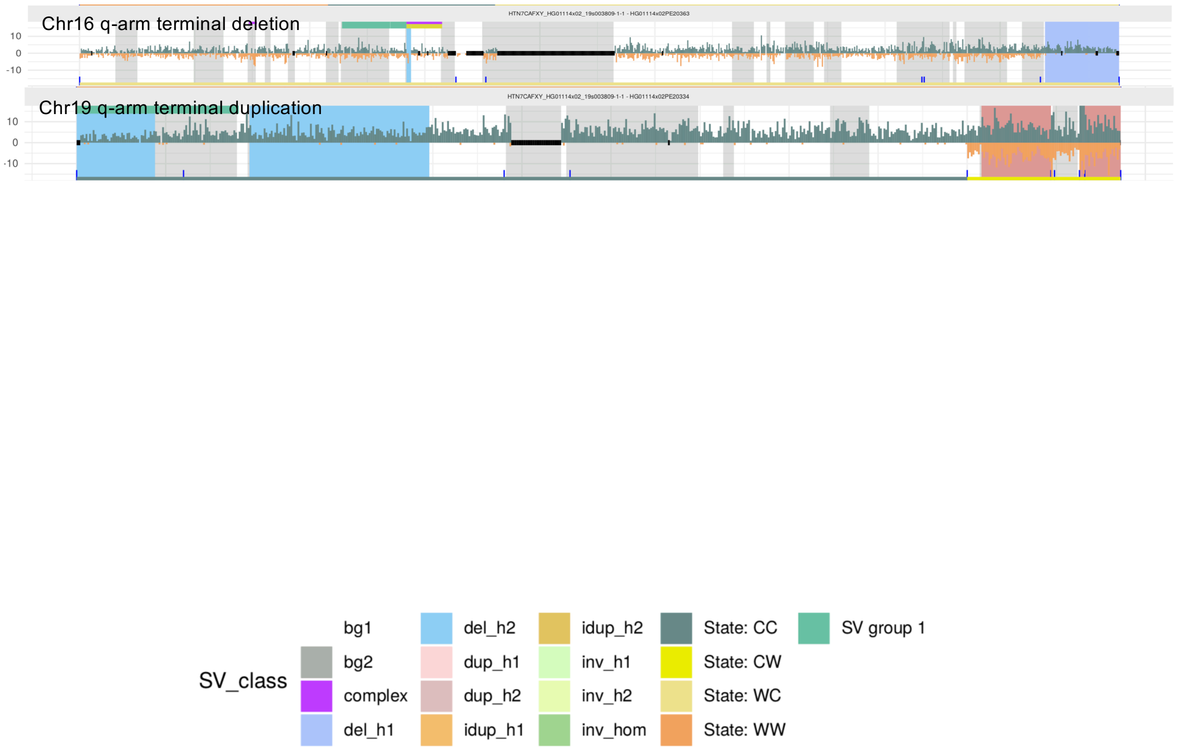

#### Single-cell SV events in LCLs (HG01596, 53 single cells profiled)

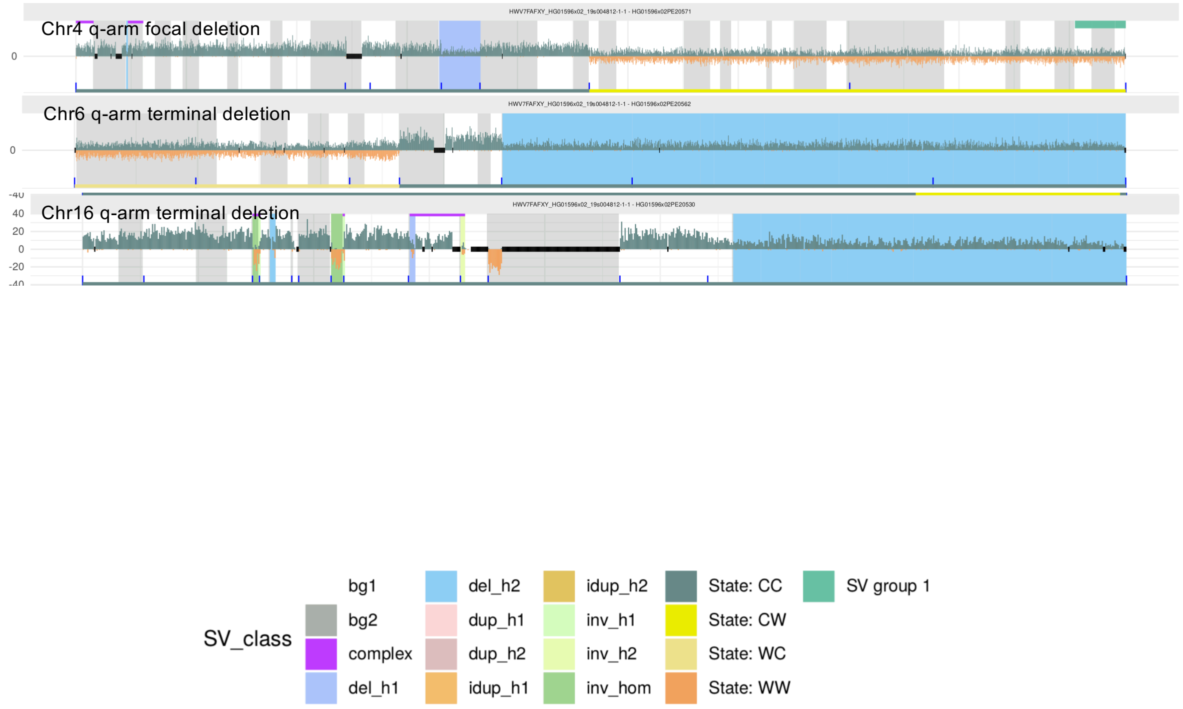

### Single-cell SV events in LCLs (HG02011, 55 single cells profiled)

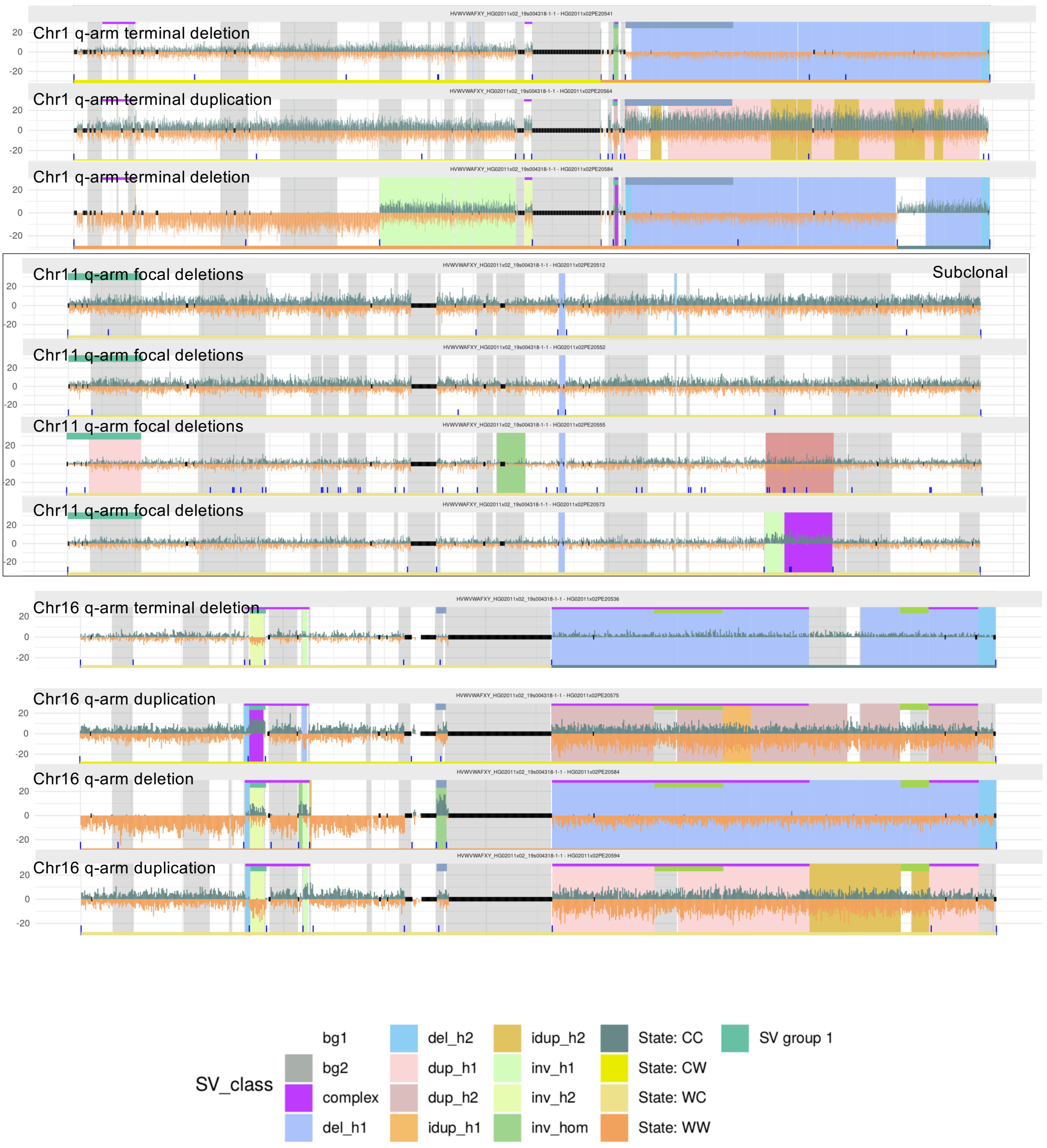

### Single-cell SV events in LCLs (HG02492, 30 single cells profiled)

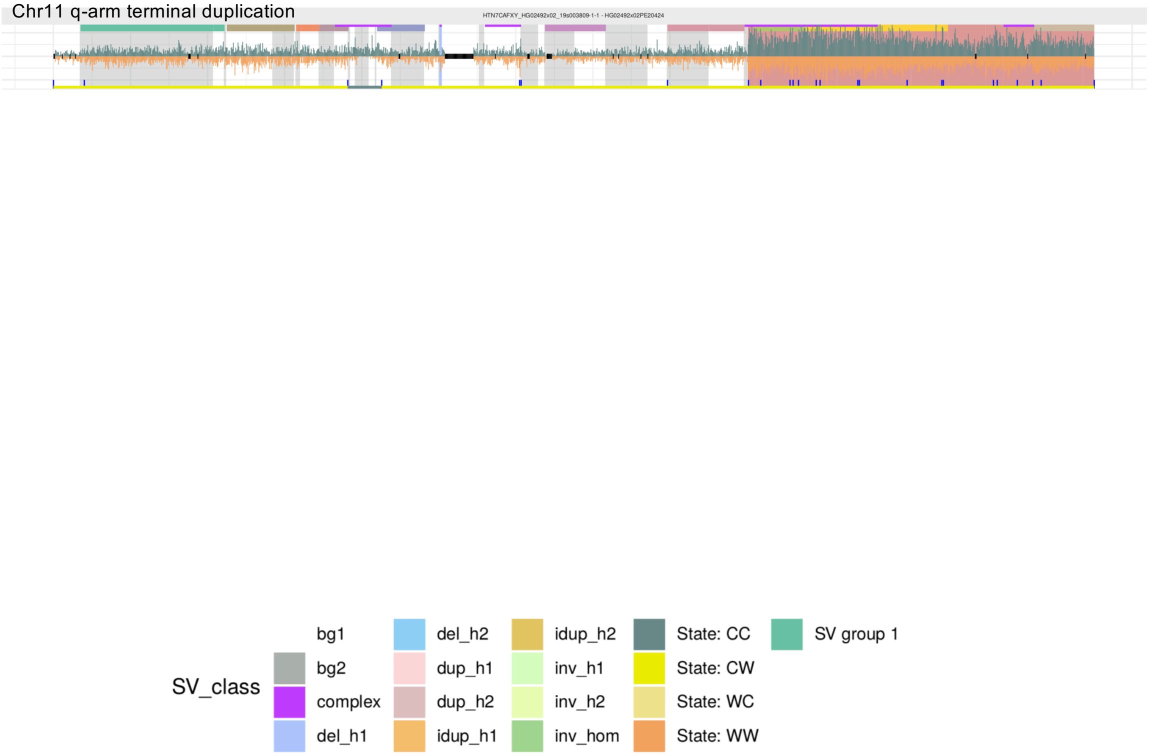

### Single-cell SV events in LCLs (HG02587, 54 single cells profiled)

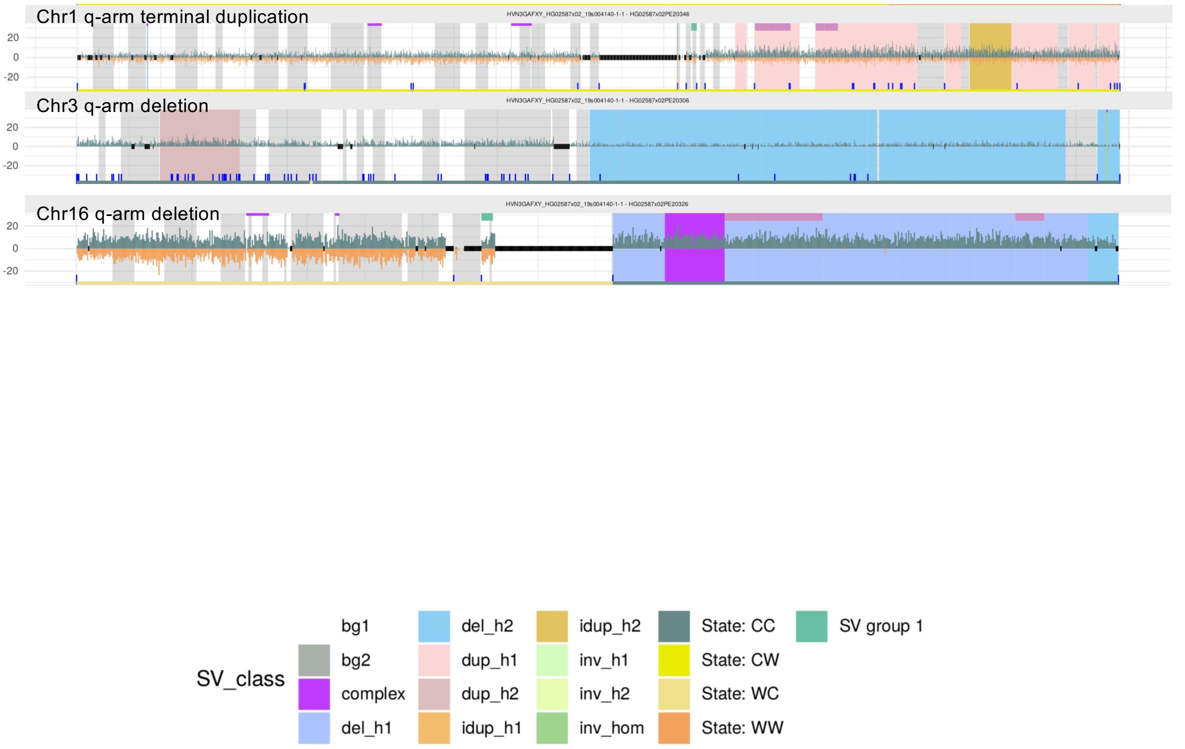

#### Single-cell SV events in LCLs (HG02818, 69 single cells profiled)

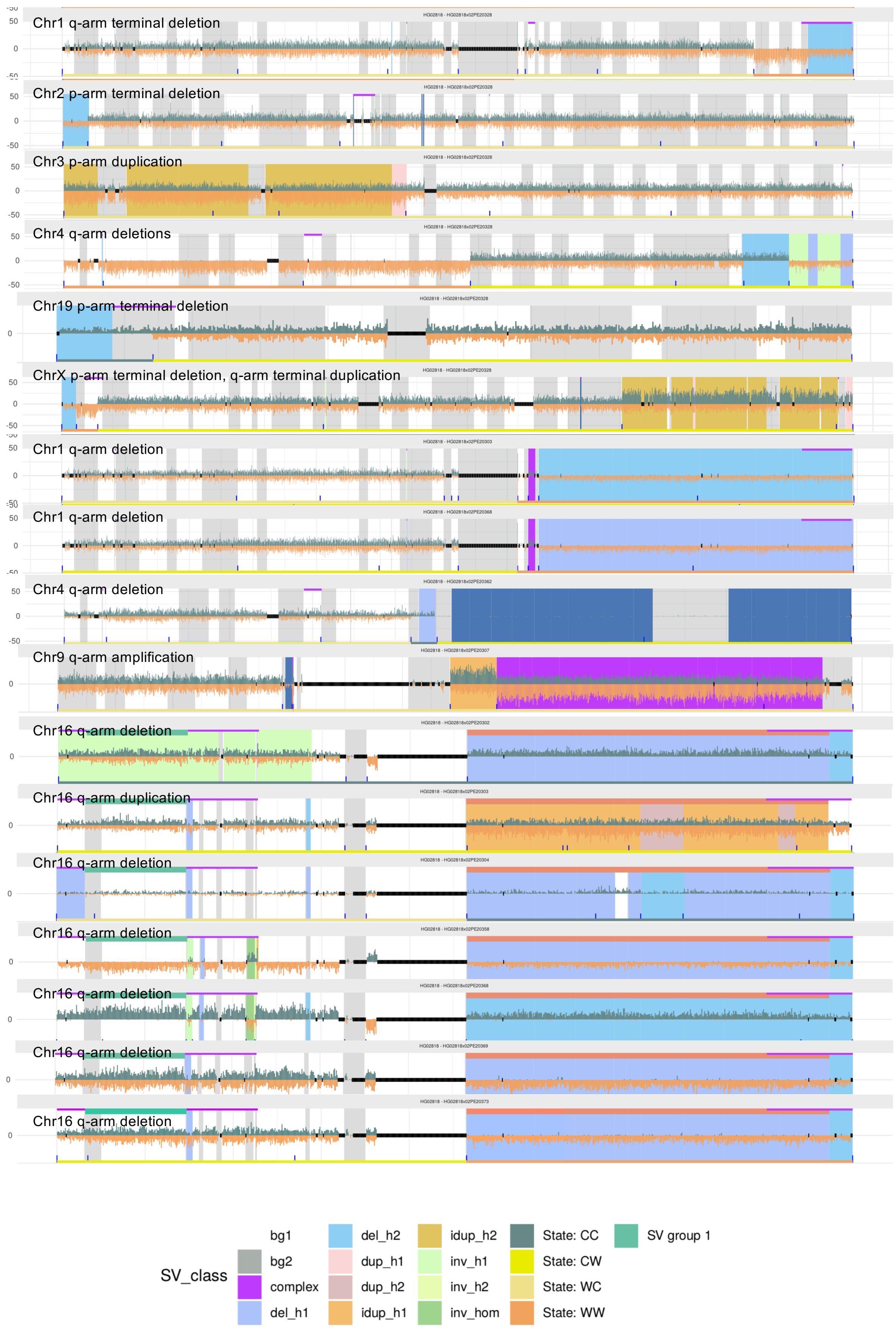

#### Single-cell SV events in LCLs (HG03009, 55 single cells profiled)

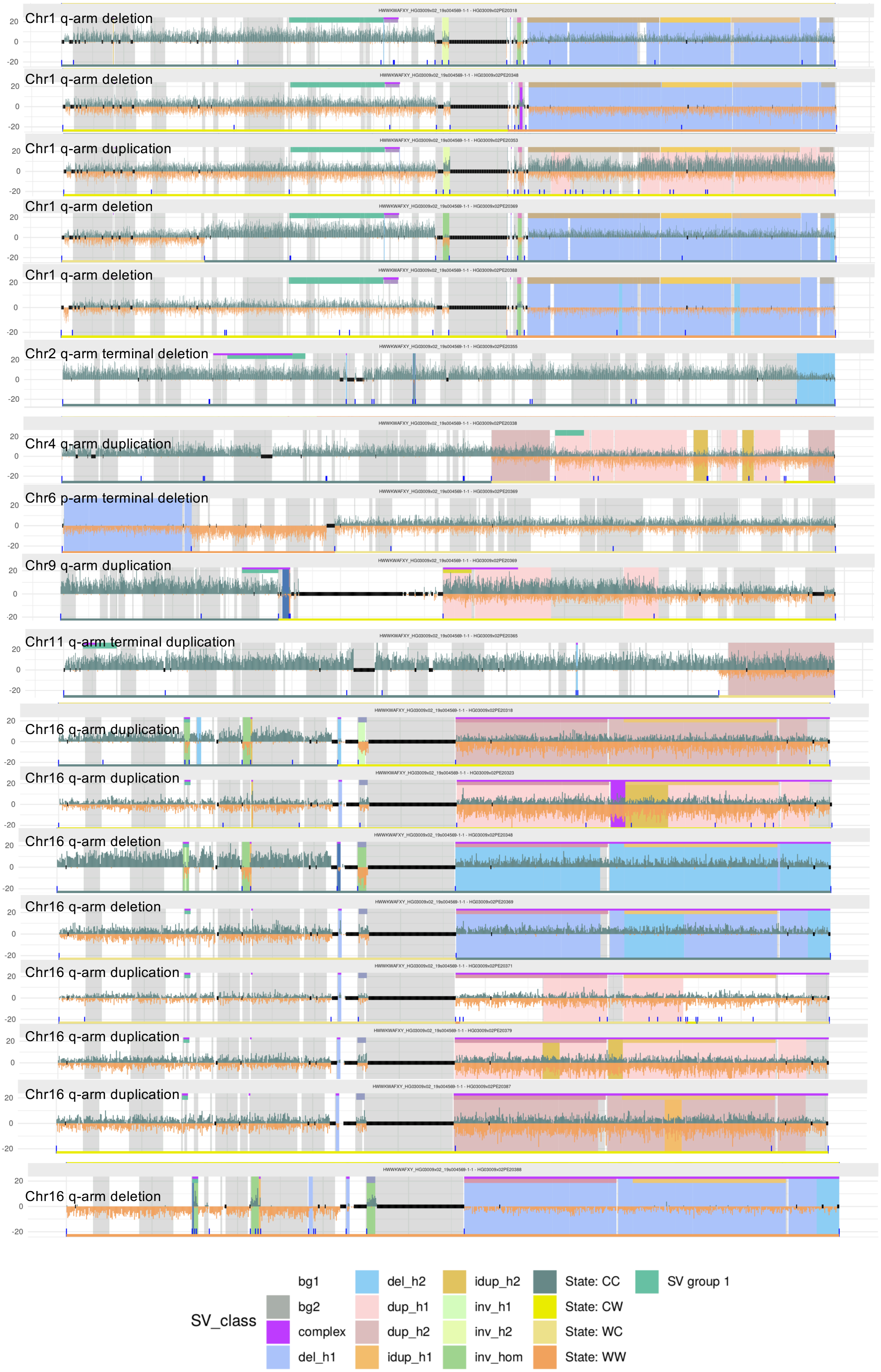

### Single-cell SV events in LCLs (HG03065, 56 single cells profiled)

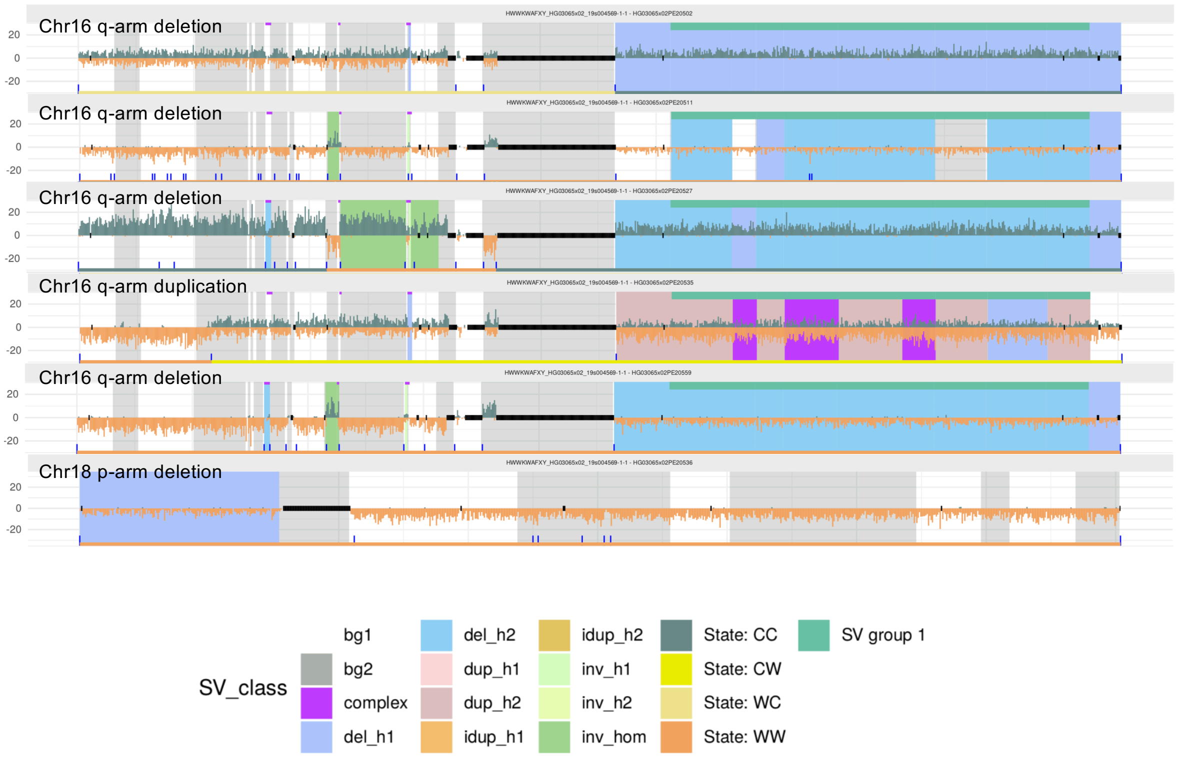

#### Single-cell SV events in LCLs (HG03125, 55 single cells profiled)

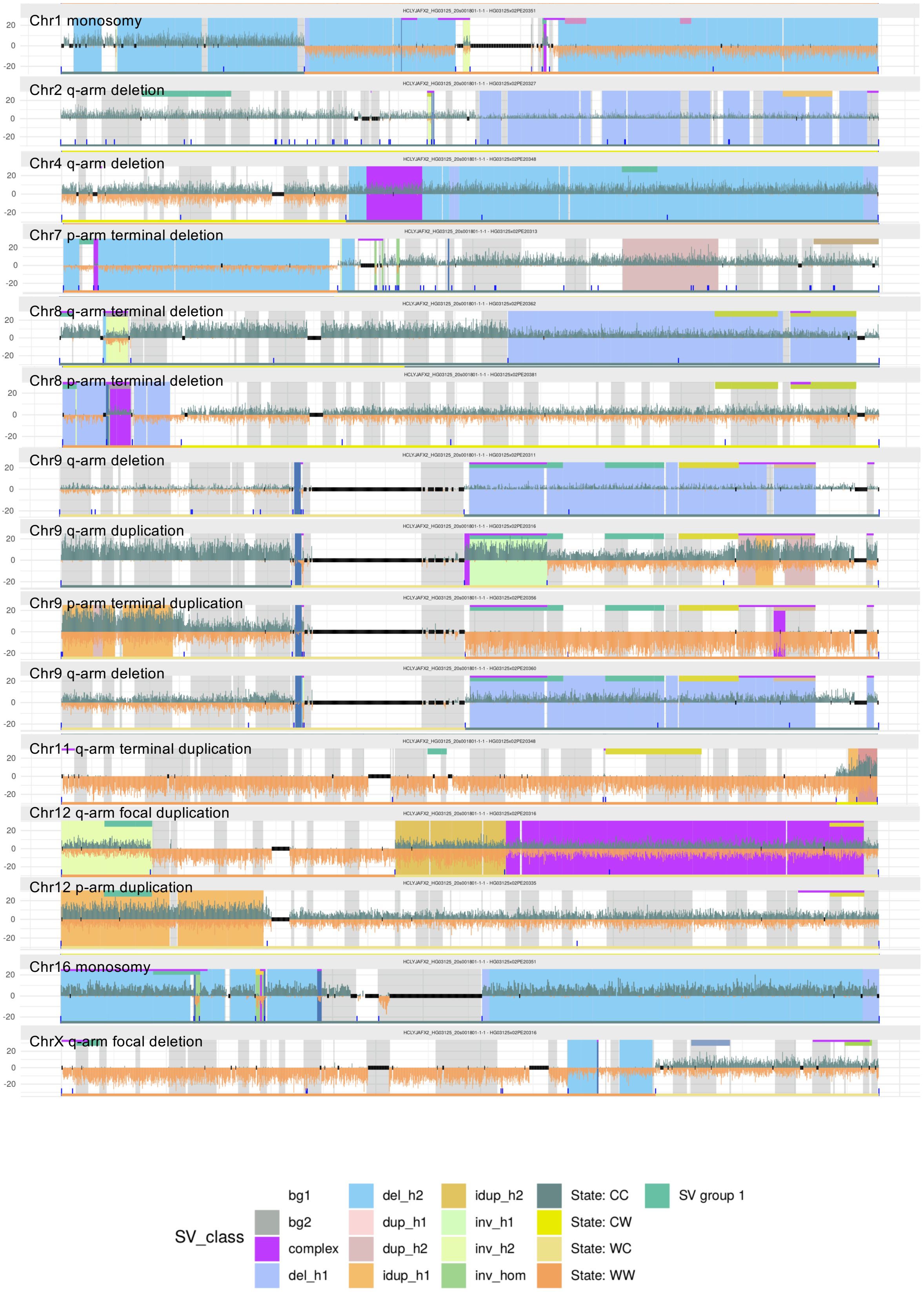

#### Single-cell SV events in LCLs (HG03371, 61 single cells profiled)

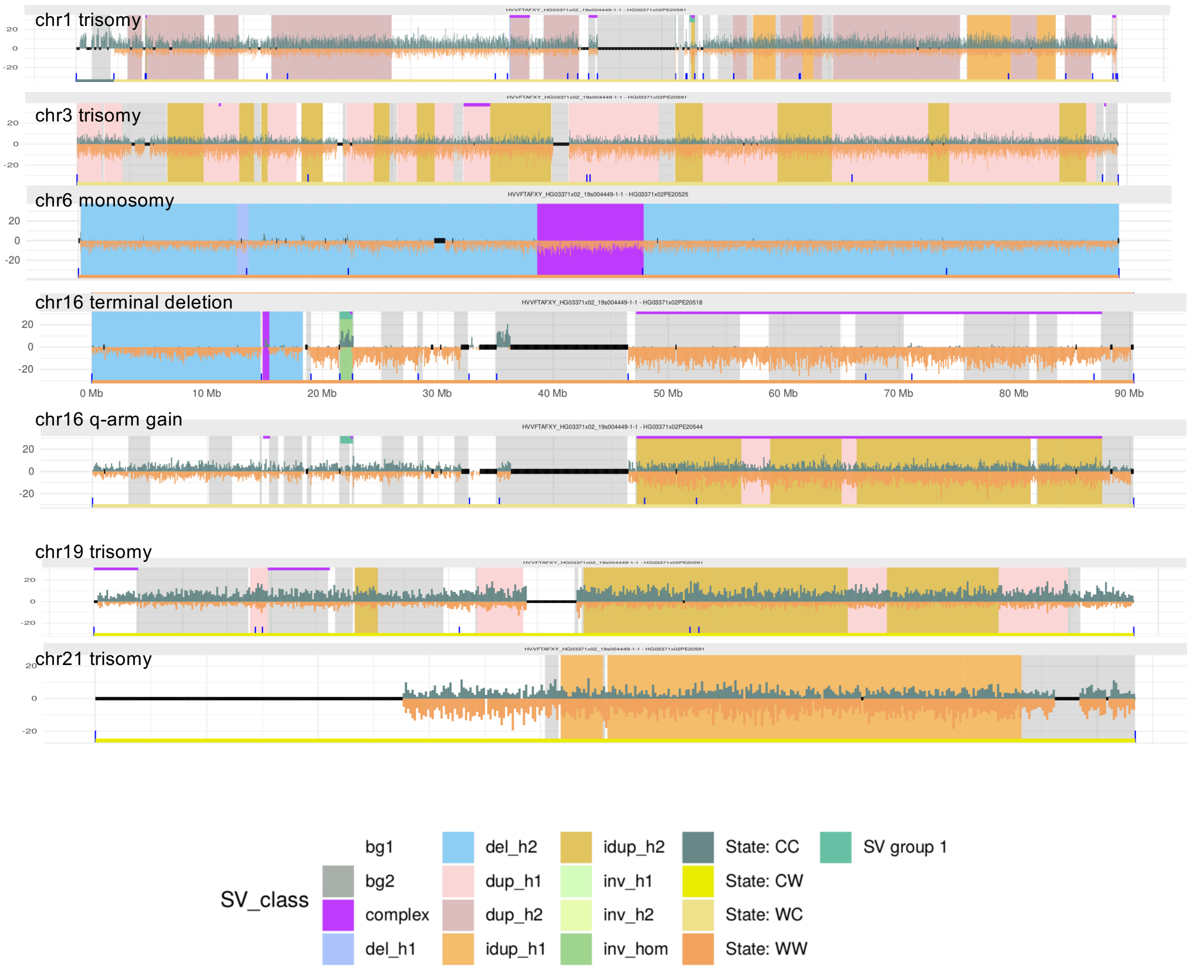

### Single-cell SV events in LCLs (HG03486, 61 single cells profiled)

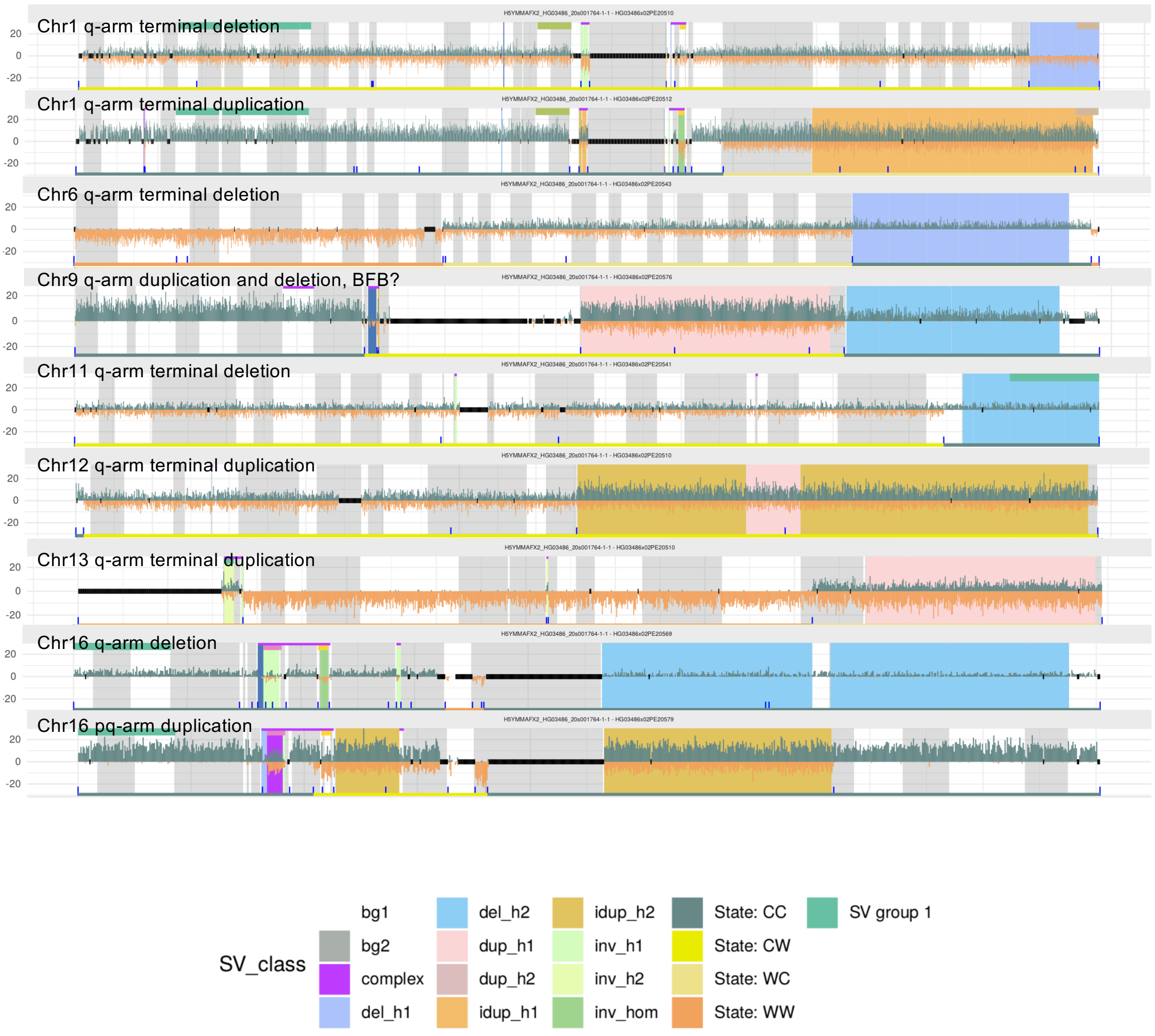

### Single-cell SV events in LCLs (HG03683, 50 single cells profiled)

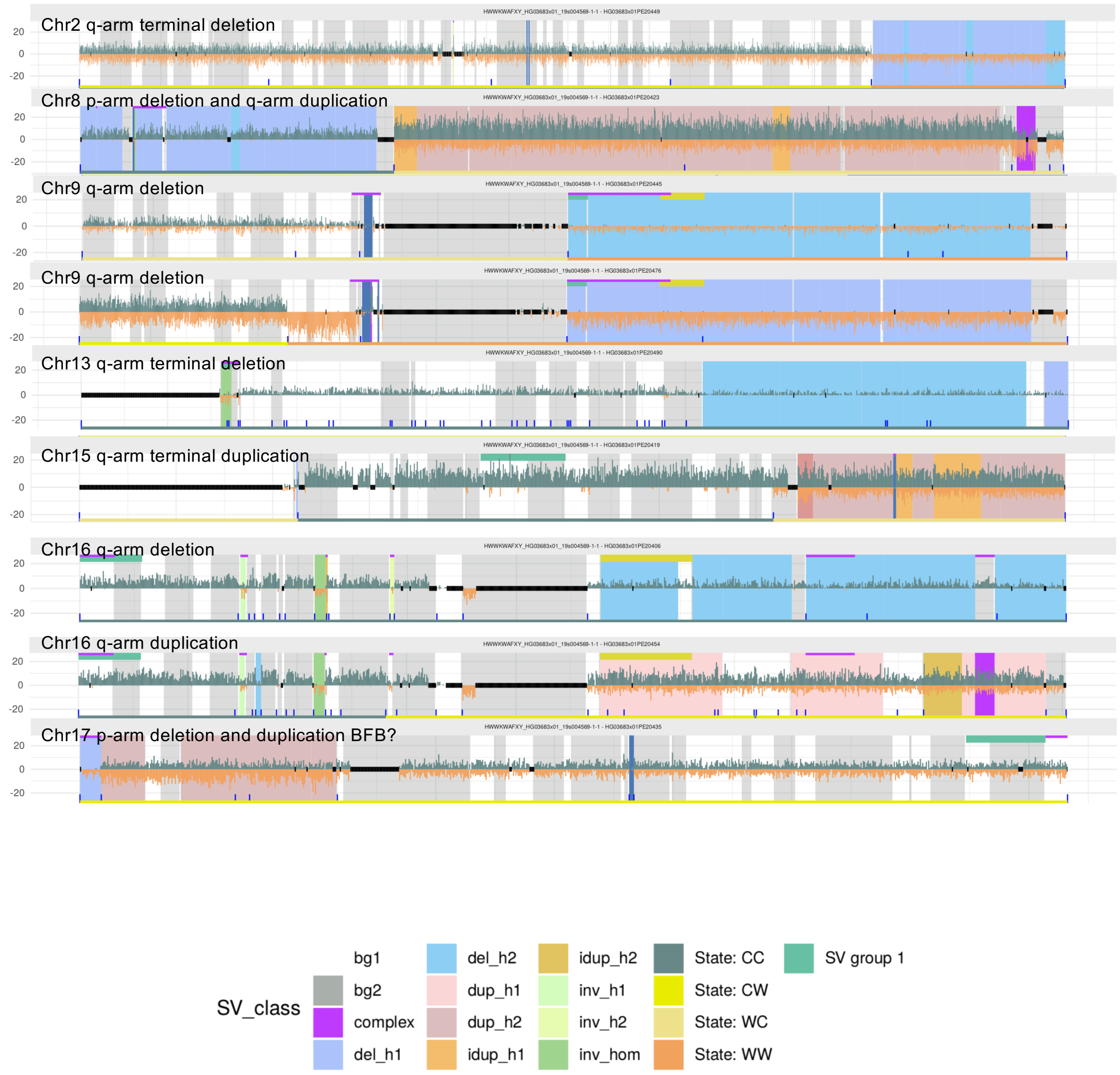

### Single-cell SV events in LCLs (HG03732, 47 single cells profiled)

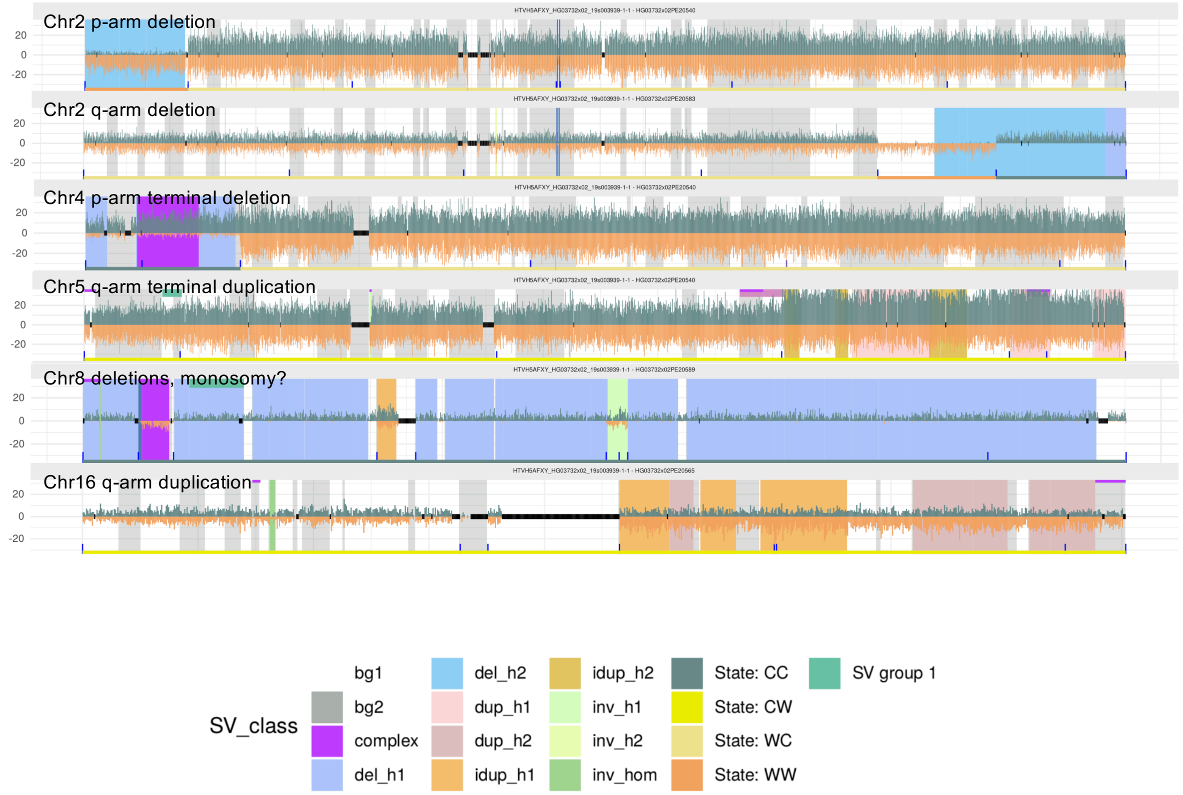
