## Supplementary Table S1 for "Haplotype-aware single-cell multiomics uncovers functional effects of somatic structural variation"

Table S1. Characterization of nucleosomal fragments of Strand-seq libraries

**a. Comparison of pooled Strand-seq and Bulk MNase-seq for well-characterzed cell line NA12878**

| Assay | Bulk Mnase-seq | Pooled Strand-seq |
|---|---|---|
| Cell line | NA12878 | NA12878 |
| Cell counts | Starting sample ~2.5×10^8 | 132 |
| Read length | 35bp, SE | 101bp, PE |
| Sequencer | SOLiD | Illumina HiSeq2500 |
| Mapped read | 1765496812 | 75330156 |
| Nucleosome peak counts | 15894540 | 13334533 |
| Reference | ENCODE | Porubsky et al. |

**b. Genomic coverage and nucleosome repeat length estimates using bulk MNase-seq (based on paired-end, Illumina sequencing)***

| Cell line and assay | Fragment count | Genomic coverage | Nucleosome Repeat Length (1st-3rd) (bp) |
|---|---|---|---|
| NA19238 MNase-seq | 636207753 (downsampled to 63627802) | 90% | 194 |
| NA18508 MNase-seq | 539077053 (downsampled to 70092387) | 91% | 194.5 |
| NA19193 MNase-seq | 197380559 (downsampled to 71050698) | 88% | 192.5 |
| | mean | | 193.6666667 |

**NA12878 MNase-seq data based on Illumina sequencing is, to our knowledge, not publicly available*

**c. Fragment size distribution of Strand-seq genomic libraries reanalyzed or generated in this study**

mean 195.4428571

| | | | | | Raw count (pooled single-cells) | | | Percentage | | | | |
|---|---|---|---|---|---|---|---|---|---|---|---|---|
| **Sample** | **Data source** | **Cell counts** | **Fragment count** | **Fragment per cell (media** | **subnucleosomal (<80bp)** | **mononucleosomal (140-180bp)** | **dinucleosomal (> 280bp)** | **subnucleosomal (<8** | **mononucleosomal (1** | **dinucleosomal (> 28** | **Nucleosome Repeat Length (1st-3rd)** | **Mappable genome covered by nucleosome (%)** |
| NA12878 (strict QC, 80-220bp) | Porubsky et al. Genome Res 2016 | 95 | 49205197 | 540379 | 0 | 29695914 | 0 | 0 | 60.35117388 | 0 | 200.5 | 81.59 |
| RPE1 WT | Sanders et al. Nat Biotechnol 2019 | 79 | 32818312 | 380889 | 34702 | 21190509 | 34249 | 0.1057397468 | 64.56916188 | 0.1043594198 | 192 | 72.93 |
| RPE1 WT second batch | This study | 77 | 36255312 | 475586 | 26510 | 23833577 | 33607 | 0.07312031958 | 65.73816549 | 0.09269538213 | 194 | 76.35 |
| RPE1 BM510 | Sanders et al. Nat Biotechnol 2019 | 145 | 70715091 | 483817 | 54479 | 47553447 | 134168 | 0.07704013278 | 67.24653299 | 0.1897303646 | 192 | 85.69 |
| RPE1 C7 | Sanders et al. Nat Biotechnol 2019 | 154 | 63600027 | 387147 | 28373 | 44505121 | 51366 | 0.04461161628 | 69.97657564 | 0.08076411666 | 193.5 | 85.21 |
| T-ALL P1 | Sanders et al. Nat Biotechnol 2019 | 77 | 31698072 | 381825 | 50169 | 17466696 | 90884 | 0.1582714558 | 55.10333878 | 0.2867177537 | 198 | 73.05 |
| CLL_24 | This study | 86 | 49895483 | 551830.5 | 38805 | 32342001 | 114541 | 0.07777257112 | 64.81949679 | 0.2295618623 | 195.5 | 84.19 |
| AML01 | This study | 42 | 16383559 | 371159 | 3449 | 10851478 | 18073 | 0.02105159203 | 66.23394831 | 0.1103118071 | 196.25 | 52.99 |
| HG01573 | HGSVC | 46 | 19602141 | 371305.5 | 5139 | 13288804 | 14393 | 0.026216524 | 67.7926151 | 0.07342565284 | 195 | 61.66 |
| HG02018 | HGSVC | 50 | 15076723 | 299160 | 7278 | 9320488 | 12018 | 0.04827308958 | 61.82038365 | 0.0797122823 | 196.25 | 51.39 |
| NA19036 | HGSVC | 50 | 20580743 | 366703 | 9564 | 12897969 | 32277 | 0.04647062548 | 62.67008436 | 0.1568310726 | 195.75 | 63.73 |
| NA12329 | HGSVC (24 LCLs in the human diversity panels | 59 | 17877573 | 289944 | 7196 | 11450374 | 12083 | 0.04025154869 | 64.04881692 | 0.06758747398 | 197 | 57.51 |
| NA18534 | HGSVC (24 LCLs in the human diversity panels | 56 | 19763483 | 357604.5 | 7203 | 13180755 | 12566 | 0.03644600499 | 66.69247015 | 0.06358191013 | 196 | 59.92 |
| NA18939 | HGSVC (24 LCLs in the human diversity panels | 41 | 19184462 | 441584 | 6366 | 12418129 | 13938 | 0.03318310412 | 64.73013942 | 0.07265254559 | 195.25 | 60.70 |
| NA19650 | HGSVC (24 LCLs in the human diversity panels | 55 | 23261627 | 406277 | 15083 | 15230069 | 12286 | 0.06484069236 | 65.47293102 | 0.05281659791 | 196.25 | 64.60 |
| NA19983 | HGSVC (24 LCLs in the human diversity panels | 45 | 17141426 | 376995 | 9910 | 10719600 | 14239 | 0.05781315977 | 62.53622073 | 0.08306776811 | 196 | 56.56 |
| NA20509 | HGSVC (24 LCLs in the human diversity panels | 47 | 19349033 | 420983 | 13157 | 12306032 | 19815 | 0.0679982302 | 63.60024297 | 0.1024082185 | 191 | 61.03 |
| NA20847 | HGSVC (24 LCLs in the human diversity panels | 55 | 19459321 | 333993 | 9942 | 11870619 | 10907 | 0.05109119686 | 61.00222613 | 0.05605025992 | 196 | 58.69 |
| HG00096 | HGSVC (24 LCLs in the human diversity panels | 69 | 24731369 | 351892 | 19930 | 16058080 | 11618 | 0.080585915 | 64.93000852 | 0.04697677674 | 200.25 | 64.00 |
| HG00171 | HGSVC (24 LCLs in the human diversity panels | 49 | 18083185 | 352412 | 3887 | 11980222 | 12905 | 0.02149510719 | 66.25061901 | 0.07136464069 | 197 | 58.10 |
| HG00864 | HGSVC (24 LCLs in the human diversity panels | 43 | 23415762 | 456657 | 21014 | 14980303 | 15289 | 0.08974296886 | 63.97529579 | 0.06529362572 | 195.75 | 69.47 |
| HG01114 | HGSVC (24 LCLs in the human diversity panels | 51 | 12623122 | 245202 | 3730 | 7865114 | 10995 | 0.02954894993 | 62.30720102 | 0.08710206556 | 198.5 | 48.06 |
| HG01505 | HGSVC (24 LCLs in the human diversity panels | 61 | 19983098 | 290889 | 11138 | 12707495 | 17840 | 0.05573710343 | 63.59121594 | 0.08927544668 | 195.5 | 60.69 |
| HG01596 | HGSVC (24 LCLs in the human diversity panels | 53 | 22252799 | 412316 | 10242 | 14787574 | 11115 | 0.04602567075 | 66.45264715 | 0.04994877274 | 196.25 | 62.26 |
| HG02011 | HGSVC (24 LCLs in the human diversity panels | 55 | 17488040 | 301683 | 5122 | 11441004 | 10958 | 0.02928858809 | 65.42187689 | 0.06265996647 | 199.5 | 57.14 |
| HG02492 | HGSVC (24 LCLs in the human diversity panels | 30 | 7558818 | 222640 | 4360 | 4695079 | 5960 | 0.05768097605 | 62.11393104 | 0.07884830671 | 192 | 33.54 |
| HG02587 | HGSVC (24 LCLs in the human diversity panels | 54 | 18935650 | 369325.5 | 3645 | 12691370 | 13851 | 0.01924940522 | 67.02368284 | 0.07314773985 | 196 | 58.88 |
| HG02818 | HGSVC (24 LCLs in the human diversity panels | 69 | 33900076 | 519497 | 27472 | 21435896 | 20219 | 0.0810381664 | 63.2325898 | 0.05964293413 | 195 | 71.81 |
| HG03009 | HGSVC (24 LCLs in the human diversity panels | 55 | 13611588 | 249392 | 8410 | 8922348 | 7694 | 0.06178559034 | 65.54964784 | 0.05652536647 | 189.5 | 48.23 |
| HG03065 | HGSVC (24 LCLs in the human diversity panels | 56 | 19731479 | 315060 | 9323 | 13244423 | 117320 | 0.04724937244 | 67.123316 | 0.5945828997 | 193 | 60.86 |
| HG03125 | HGSVC (24 LCLs in the human diversity panels | 55 | 16141668 | 294798 | 7340 | 10235031 | 11472 | 0.04547237621 | 63.40751774 | 0.07107072206 | 194.5 | 54.43 |
| HG03371 | HGSVC (24 LCLs in the human diversity panels | 61 | 21073028 | 342548 | 6123 | 14091038 | 16355 | 0.02905609958 | 66.86764712 | 0.07761105808 | 193 | 61.17 |
| HG03486 | HGSVC (24 LCLs in the human diversity panels | 61 | 19496724 | 310548 | 12069 | 12370813 | 18244 | 0.06190270735 | 63.45072639 | 0.09357469491 | 196 | 56.56 |
| HG03683 | HGSVC (24 LCLs in the human diversity panels | 50 | 14821975 | 289712 | 2245 | 9719435 | 10402 | 0.01514642954 | 65.57449328 | 0.07017958133 | 196.5 | 50.41 |
| HG03732 | HGSVC (24 LCLs in the human diversity panels | 47 | 16427583 | 332916 | 3294 | 11493108 | 10621 | 0.02005164119 | 69.96225799 | 0.06465345511 | 196 | 52.92 |
| | **Sum** | **2178** | | | | | | | | | | |
