## Supplementary Table S2 for "Haplotype-aware single-cell multiomics uncovers functional effects of somatic structural variation"

**Table S2A. List of genes inferred to be changing in activity when comparing major and**

| Index | chr | str | end | name | Symbol | EntrezID | baseMean | NO log2FoldChange (RPE1/L | lfcSE | stat | pvalue | padj | Inferred gene activit | Top10 RPE-1 (RNA-seq | Top10 HG01573 (RNA-seq) |
|---|---|---|---|---|---|---|---|---|---|---|---|---|---|---|---|
| 1 | chr1 | 25271393 | 25273393 | ENSG00000 | RHD | 6007 | 3.79945205 | -4.737291108 | 0.18866297 | -25.1098082 | 3.89E-139 | 2.60E-135 | RPE1_active | 0 | 0 |
| 2 | chr1 | 25337213 | 25339213 | ENSG00000 | RSRP1 | 57035 | 9.95575415 | -2.004299471 | 0.10838970 | -18.4916041 | 2.41E-76 | 1.07E-72 | RPE1_active | 0 | 0 |
| 3 | chr20 | 1619061 | 1621061 | ENSG00000 | SIRPB1 | 10326 | 8.19942523 | -1.008710397 | 0.11526768 | -8.75102468 | 2.11E-18 | 2.17E-15 | RPE1_active | 0 | 0 |
| 4 | chr3 | 50358610 | 50360610 | ENSG00000 | TMEM115 | 11070 | 1.60784890 | -1.00327087 | 0.25555961 | -3.92578023 | 8.64E-05 | 0.00314726 | RPE1_active | 0 | 0 |
| 5 | chr10 | 47548750 | 47550750 | ENSG00000 | AGAP10 | 642517 | 3.25325095 | -0.9270238789 | 0.18037465 | -5.13943535 | 2.76E-07 | 2.78E-05 | RPE1_active | 0 | 0 |
| 6 | chr14 | 18966434 | 18968434 | ENSG00000 | POTEG | 641455 | 2.32019691 | -0.8492667588 | 0.22369290 | -3.79657436 | 0.00014670 | 0.00481617 | RPE1_active | 0 | 0 |
| 7 | chr12 | 52191000 | 52193000 | ENSG00000 | KRT80 | 144501 | 7.19069179 | -0.8240148269 | 0.13172724 | -6.25546256 | 3.96E-10 | 8.02E-08 | RPE1_active | 0 | 0 |
| 8 | chr1 | 163067775 | 163069775 | ENSG00000 | RGS4 | 5999 | 2.19606450 | -0.7946554922 | 0.23044384 | -3.44836939 | 0.00056398 | 0.01382635 | RPE1_active | 0 | 0 |
| 9 | chr14 | 22981642 | 22983642 | ENSG00000 | AJUBA | 84962 | 3.10035728 | -0.7891001921 | 0.19553863 | -4.03552058 | 5.45E-05 | 0.00212843 | RPE1_active | 0 | 0 |
| 10 | chr5 | 93582337 | 93584337 | ENSG00000 | NR2F1 | 7025 | 1.90742988 | -0.7800484206 | 0.27999495 | -2.78593743 | 0.00533731 | 0.06654391 | RPE1_active | 0 | 0 |
| 11 | chr11 | 75561056 | 75563056 | ENSG00000 | SERPINH1 | 871 | 2.46334128 | -0.7397918432 | 0.22807204 | -3.24367610 | 0.00117997 | 0.02315079 | RPE1_active | 0 | 0 |
| 12 | chr1 | 202966280 | 202968280 | ENSG00000 | CYB5R1 | 51706 | 2.10762846 | -0.7332905764 | 0.25163596 | -2.91409286 | 0.00356723 | 0.05169399 | RPE1_active | 0 | 0 |
| 13 | chr3 | 50349695 | 50351695 | ENSG00000 | CYB561D2 | 11068 | 2.69980369 | -0.7163766579 | 0.20548314 | -3.48630368 | 0.00048974 | 0.01268116 | RPE1_active | 0 | 0 |
| 14 | chr10 | 47483158 | 47485158 | ENSG00000 | ANXA8 | 653145 | 3.05979571 | -0.7105510335 | 0.17359491 | -4.09315591 | 4.26E-05 | 0.00172816 | RPE1_active | 0 | 0 |
| 15 | chr15 | 63041632 | 63043632 | ENSG00000 | TPM1 | 7168 | 7.10865765 | -0.7026950255 | 0.11412926 | -6.15701014 | 7.41E-10 | 1.46E-07 | RPE1_active | 1 | 0 |
| 16 | chr10 | 45852875 | 45854875 | ENSG00000 | AGAP4 | 119016 | 2.51942597 | -0.6751973296 | 0.21085466 | -3.20219292 | 0.00136385 | 0.02599498 | RPE1_active | 0 | 0 |
| 17 | chr2 | 234950973 | 234952973 | ENSG00000 | SH3BP4 | 23677 | 30.0825106 | -0.6665118182 | 0.06317669 | -10.5499627 | 5.08E-26 | 6.79E-23 | RPE1_active | 0 | 0 |
| 18 | chr1 | 94540800 | 94542800 | ENSG00000 | F3 | 2152 | 2.38249104 | -0.6540858181 | 0.23377050 | -2.79798263 | 0.00514228 | 0.06510271 | RPE1_active | 0 | 0 |
| 19 | chr1 | 27011836 | 27013836 | ENSG00000 | FAM46B | 115572 | 2.27389331 | -0.652928813 | 0.23051871 | -2.83243296 | 0.00461952 | 0.06097269 | RPE1_active | 0 | 0 |
| 20 | chr3 | 134374479 | 134376479 | ENSG00000 | AMOTL2 | 51421 | 5.25545424 | -0.6368759951 | 0.15415587 | -4.13137661 | 3.61E-05 | 0.00149162 | RPE1_active | 0 | 0 |
| 21 | chr2 | 28391448 | 28393448 | ENSG00000 | FOSL2 | 2355 | 7.11837180 | -0.6157881373 | 0.12908255 | -4.77049865 | 1.84E-06 | 0.00013344 | RPE1_active | 0 | 0 |
| 22 | chr17 | 50054968 | 50056968 | ENSG00000 | ITGA3 | 3675 | 8.89074765 | -0.6154416363 | 0.11662838 | -5.27694549 | 1.31E-07 | 1.47E-05 | RPE1_active | 0 | 0 |
| 23 | chr16 | 78098413 | 78100413 | ENSG00000 | WWOX | 51741 | 261.473412 | -0.6147034072 | 0.02392453 | -25.6934320 | 1.38E-145 | 1.85E-141 | RPE1_active | 0 | 0 |
| 24 | chr1 | 150256159 | 150258159 | ENSG00000 | CA14 | 23632 | 2.32591105 | -0.6089397187 | 0.21989882 | -2.76918134 | 0.00561973 | 0.06913929 | RPE1_active | 0 | 0 |
| 25 | chr3 | 28348127 | 28350127 | ENSG00000 | AZI2 | 64343 | 7.09082018 | -0.6007897042 | 0.12726846 | -4.72064846 | 2.35E-06 | 0.00016505 | RPE1_active | 0 | 0 |
| 26 | chr1 | 94926278 | 94928278 | ENSG00000 | CNN3 | 1266 | 7.55782615 | -0.5936782753 | 0.11973309 | -4.95834755 | 7.11E-07 | 6.25E-05 | RPE1_active | 0 | 0 |
| 27 | chr2 | 130610481 | 130612481 | ENSG00000 | POTEJ | 653781 | 6.11650666 | -0.5932282331 | 0.14280226 | -4.15419358 | 3.26E-05 | 0.00137587 | RPE1_active | 0 | 0 |
| 28 | chr3 | 184297709 | 184299709 | ENSG00000 | PSMD2 | 5708 | 3.49959883 | -0.5889321903 | 0.18731966 | -3.14399553 | 0.00166657 | 0.03001524 | RPE1_active | 0 | 0 |
| 29 | chr20 | 1619009 | 1621009 | ENSG00000 | RP4-576H24 | NaN | 15.2748665 | -0.5849979351 | 0.08476935 | -6.90105464 | 5.16E-12 | 1.38E-09 | RPE1_active | 0 | 0 |
| 30 | chr15 | 39580079 | 39582079 | ENSG00000 | THBS1 | 7057 | 5.05060172 | -0.5749969477 | 0.16090241 | -3.57357557 | 0.00035213 | 0.00986359 | RPE1_active | 1 | 0 |
| 31 | chr4 | 57109385 | 57111385 | ENSG00000 | IGFBP7 | 3490 | 19.6237116 | -0.5739095096 | 0.07417566 | -7.73716716 | 1.02E-14 | 5.03E-12 | RPE1_active | 0 | 0 |
| 32 | chr11 | 114438386 | 114440386 | ENSG00000 | REXO2 | 25996 | 2.68215282 | -0.5589040885 | 0.21263260 | -2.6284966 | 0.00857632 | 0.09218682 | RPE1_active | 0 | 0 |
| 33 | chr11 | 12008769 | 12010769 | ENSG00000 | DKK3 | 27122 | 13.5258396 | -0.55336127 | 0.09065104 | -6.10430124 | 1.03E-09 | 2.00E-07 | RPE1_active | 0 | 0 |
| 34 | chr17 | 15998717 | 16000717 | ENSG00000 | ZSWIM7 | 125150 | 4.78961747 | -0.548080565 | 0.14044668 | -3.90241022 | 9.52E-05 | 0.00338429 | RPE1_active | 0 | 0 |
| 35 | chr9 | 41198261 | 41200261 | ENSG00000 | CBWD7 | 644019 | 2.65680686 | -0.5477493022 | 0.19342740 | -2.83180808 | 0.00462856 | 0.06098837 | RPE1_active | 0 | 0 |
| 36 | chr12 | 13195716 | 13197716 | ENSG00000 | EMP1 | 2012 | 6.98360799 | -0.5404980751 | 0.11877294 | -4.55068340 | 5.35E-06 | 0.00032037 | RPE1_active | 0 | 0 |
| 37 | chr7 | 128829377 | 128831377 | ENSG00000 | FLNC | 2318 | 8.69755953 | -0.535512109 | 0.11393587 | -4.70011848 | 2.60E-06 | 0.00017724 | RPE1_active | 0 | 0 |
| 38 | chr10 | 46374627 | 46376627 | ENSG00000 | ANXA8L2 | 728113 | 24.7388670 | -0.5313533558 | 0.06452522 | -8.23481596 | 1.80E-16 | 1.20E-13 | RPE1_active | 0 | 0 |
| 39 | chr12 | 85279107 | 85281107 | ENSG00000 | ALX1 | 8092 | 3.36547423 | -0.5280597836 | 0.19201492 | -2.75009763 | 0.00595775 | 0.07164852 | RPE1_active | 0 | 0 |
| 40 | chr1 | 16155087 | 16157087 | ENSG00000 | EPHA2 | 1969 | 8.67998303 | -0.5203512624 | 0.11271869 | -4.61637059 | 3.91E-06 | 0.00024381 | RPE1_active | 0 | 0 |
| 41 | chr4 | 122731702 | 122733702 | ENSG00000 | BBS12 | 166379 | 3.47414620 | -0.5184996392 | 0.16325187 | -3.17607153 | 0.00149284 | 0.02770257 | RPE1_active | 0 | 0 |
| 42 | chr1 | 230868589 | 230870589 | ENSG00000 | C1orf198 | 84886 | 9.4928065 | -0.5039194209 | 0.11516763 | -4.37552966 | 1.21E-05 | 0.00062810 | RPE1_active | 0 | 0 |
| 43 | chr3 | 184154388 | 184156388 | ENSG00000 | DVL3 | 1857 | 5.14431833 | -0.500916656 | 0.14316969 | -3.49876187 | 0.00046742 | 0.01219775 | RPE1_active | 0 | 0 |
| 44 | chr16 | 55507998 | 55509998 | ENSG00000 | LPCAT2 | 54947 | 17.4225488 | -0.4949517717 | 0.07834429 | -6.31764913 | 2.66E-10 | 5.63E-08 | RPE1_active | 0 | 0 |
| 45 | chr20 | 45415077 | 45417077 | ENSG00000 | PIGT | 51604 | 3.21060346 | -0.4893017526 | 0.17493613 | -2.79703070 | 0.00515746 | 0.06519287 | RPE1_active | 0 | 0 |
| 46 | chr8 | 123540206 | 123542206 | ENSG00000 | FBXO32 | 114907 | 11.3109833 | -0.4891719511 | 0.10139518 | -4.82440985 | 1.40E-06 | 0.00010659 | RPE1_active | 0 | 0 |
| 47 | chr1 | 1348350 | 1350350 | ENSG00000 | DVL1 | 1855 | 3.57343683 | -0.4848749941 | 0.17604239 | -2.75430812 | 0.00588163 | 0.07118165 | RPE1_active | 0 | 0 |
| 48 | chr19 | 41218203 | 41220203 | ENSG00000 | AXL | 558 | 10.8258057 | -0.4808114601 | 0.09876183 | -4.86839332 | 1.13E-06 | 9.11E-05 | RPE1_active | 1 | 0 |
| 49 | chr12 | 54418460 | 54420460 | ENSG00000 | ITGA5 | 3678 | 6.24066442 | -0.4765989296 | 0.13850747 | -3.44096187 | 0.00057965 | 0.01405572 | RPE1_active | 0 | 0 |
| 50 | chr15 | 73924989 | 73926989 | ENSG00000 | LOXL1 | 4016 | 8.15779855 | -0.4763634139 | 0.117014421 | -4.07098037 | 4.68E-05 | 0.00186162 | RPE1_active | 0 | 0 |
| 51 | chr12 | 12999273 | 13001273 | ENSG00000 | HEBP1 | 50865 | 7.70433326 | -0.4731086236 | 0.11959736 | -3.95584474 | 7.63E-05 | 0.00283836 | RPE1_active | 0 | 0 |
| 52 | chr19 | 38288151 | 38290151 | ENSG00000 | CTB-102L5. | NaN | 4.53572626 | -0.4645540947 | 0.17402096 | -2.66952941 | 0.00759576 | 0.08478443 | RPE1_active | 0 | 0 |
| 53 | chr13 | 113863067 | 113865067 | ENSG00000 | GAS6 | 2621 | 11.5661291 | -0.4589671887 | 0.10559234 | -4.34659520 | 1.38E-05 | 0.00068932 | RPE1_active | 0 | 0 |
| 54 | chr8 | 41308497 | 41310497 | ENSG00000 | SFRP1 | 6422 | 13.7222146 | -0.4586996111 | 0.09808384 | -4.67660699 | 2.92E-06 | 0.00019291 | RPE1_active | 0 | 0 |
| 55 | chr10 | 33335262 | 33337262 | ENSG00000 | NRP1 | 8829 | 38.4109799 | -0.4553740628 | 0.05303760 | -8.58587115 | 9.02E-18 | 7.53E-15 | RPE1_active | 0 | 0 |

| | | | | | | | | | | | | | | | |
|---|---|---|---|---|---|---|---|---|---|---|---|---|---|---|---|
| 56 | chr4 | 121696113 | 121698113 | ENSG00000 | ANXA5 | 308 | 6.426710877 | -0.4531773041 | 0.143966248 | -3.14780241 | 0.001645028 | 0.029782147 | RPE1_active | 0 | 0 |
| 57 | chr19 | 44642798 | 44644798 | ENSG00000 | PVR | 5817 | 4.884356873 | -0.4489733777 | 0.153803037 | -2.91914506 | 0.003509928 | 0.051252627 | RPE1_active | 0 | 0 |
| 58 | chr11 | 57323952 | 57325952 | ENSG00000 | TNKS1BP1 | 85456 | 8.006592647 | -0.4475006798 | 0.118974286 | -3.76132266 | 0.000169017 | 0.005326035 | RPE1_active | 0 | 0 |
| 59 | chr9 | 127577989 | 127579989 | ENSG00000 | FAM129B | 64855 | 20.8857201 | -0.4443606342 | 0.079132231 | -5.615419 | 1.96E-08 | 2.79E-06 | RPE1_active | 0 | 0 |
| 60 | chr8 | 23424328 | 23426328 | ENSG00000 | LOXL2 | 4017 | 36.30001614 | -0.4381749072 | 0.060130585 | -7.28705539 | 3.17E-13 | 1.21E-10 | RPE1_active | 0 | 0 |
| 61 | chr1 | 85577363 | 85579363 | ENSG00000 | DDAH1 | 23576 | 61.78021415 | -0.437756336 | 0.045219763 | -9.68064193 | 3.64E-22 | 4.43E-19 | RPE1_active | 0 | 0 |
| 62 | chr20 | 45347424 | 45349424 | ENSG00000 | SDC4 | 6385 | 6.580576608 | -0.4350009362 | 0.125864797 | -3.45609689 | 0.000548058 | 0.013636138 | RPE1_active | 0 | 0 |
| 63 | chr2 | 36354926 | 36356926 | ENSG00000 | CRIM1 | 51232 | 50.02920865 | -0.4327899235 | 0.050716685 | -8.53348204 | 1.42E-17 | 1.05E-14 | RPE1_active | 0 | 0 |
| 64 | chr3 | 120450253 | 120452253 | ENSG00000 | FSTL1 | 11167 | 17.19247907 | -0.4323926848 | 0.087816162 | -4.92383943 | 8.49E-07 | 7.16E-05 | RPE1_active | 1 | 0 |
| 65 | chr17 | 18257916 | 18259916 | ENSG00000 | FLII | 2314 | 4.246249388 | -0.4322784585 | 0.166493994 | -2.59636067 | 0.009421710 | 0.098040091 | RPE1_active | 0 | 0 |
| 66 | chr18 | 57802101 | 57804101 | ENSG00000 | ATP8B1 | 5205 | 35.6555901 | -0.4304838296 | 0.056791822 | -7.58003192 | 3.45E-14 | 1.54E-11 | RPE1_active | 0 | 0 |
| 67 | chr12 | 65823131 | 65825131 | ENSG00000 | HMGA2 | 8091 | 30.15262698 | -0.4291258135 | 0.059727690 | -7.18470462 | 6.74E-13 | 2.37E-10 | RPE1_active | 0 | 0 |
| 68 | chr14 | 74493177 | 74495177 | ENSG00000 | NPC2 | 10577 | 4.671206338 | -0.4291103453 | 0.160078542 | -2.68062377 | 0.007348508 | 0.082635696 | RPE1_active | 0 | 0 |
| 69 | chr1 | 115337236 | 115339236 | ENSG00000 | NGF | 4803 | 15.45451557 | -0.4287341118 | 0.087087457 | -4.92302935 | 8.52E-07 | 7.16E-05 | RPE1_active | 0 | 0 |
| 70 | chr6 | 138106511 | 138108511 | ENSG00000 | PERP | 64065 | 4.933018255 | -0.4243977769 | 0.148391888 | -2.85997962 | 0.004236682 | 0.057886547 | RPE1_active | 0 | 0 |
| 71 | chr7 | 108002255 | 108004255 | ENSG00000 | LAMB1 | 3912 | 20.2471929 | -0.4228675682 | 0.072386439 | -5.84180640 | 5.16E-09 | 8.31E-07 | RPE1_active | 0 | 0 |
| 72 | chr14 | 68978440 | 68980440 | ENSG00000 | ACTN1 | 87 | 30.02605814 | -0.4214284861 | 0.070162272 | -6.00648287 | 1.90E-09 | 3.42E-07 | RPE1_active | 0 | 0 |
| 73 | chr17 | 50149630 | 50151630 | ENSG00000 | PPP1R9B | 84687 | 4.790967684 | -0.4196502351 | 0.160822665 | -2.60939734 | 0.009070185 | 0.095799802 | RPE1_active | 0 | 0 |
| 74 | chr14 | 45083099 | 45085099 | ENSG00000 | PRPF39 | 55015 | 5.868938691 | -0.4194214333 | 0.141308967 | -2.96811618 | 0.002996310 | 0.045700576 | RPE1_active | 0 | 0 |
| 75 | chr19 | 38646649 | 38648649 | ENSG00000 | ACTN4 | 81 | 20.82186794 | -0.4182555664 | 0.082687826 | -5.05824841 | 4.23E-07 | 4.07E-05 | RPE1_active | 0 | 0 |
| 76 | chr1 | 209827007 | 209829007 | ENSG00000 | DIEXF | 27042 | 8.297664294 | -0.4159790486 | 0.118289429 | -3.51662063 | 0.000437077 | 0.011697232 | RPE1_active | 0 | 0 |
| 77 | chr18 | 14542585 | 14544585 | ENSG00000 | POTEC | 388468 | 6.516273972 | -0.4151680955 | 0.125453820 | -3.30933003 | 0.000935195 | 0.019791251 | RPE1_active | 0 | 0 |
| 78 | chr12 | 49058774 | 49060774 | ENSG00000 | KMT2D | 8085 | 13.52997151 | -0.4132848806 | 0.094566849 | -4.37029343 | 1.24E-05 | 0.000637611 | RPE1_active | 0 | 0 |
| 79 | chr4 | 7938926 | 7940926 | ENSG00000 | AFAP1 | 60312 | 48.51424379 | -0.4119316014 | 0.047893504 | -8.60099103 | 7.90E-18 | 7.04E-15 | RPE1_active | 0 | 0 |
| 80 | chr6 | 3751026 | 3753026 | ENSG00000 | PXDC1 | 221749 | 8.531981297 | -0.4097837974 | 0.120488171 | -3.40102927 | 0.000671326 | 0.015530189 | RPE1_active | 0 | 0 |
| 81 | chr12 | 6309436 | 6311436 | ENSG00000 | PLEKHG6 | 55200 | 5.944410811 | -0.408964594 | 0.146371833 | -2.79401155 | 0.005205861 | 0.065680378 | RPE1_active | 0 | 0 |
| 82 | chr13 | 109785568 | 109787568 | ENSG00000 | IRS2 | 8660 | 8.544511368 | -0.4076390629 | 0.116381639 | -3.50260630 | 0.000460729 | 0.012093933 | RPE1_active | 0 | 0 |
| 83 | chr2 | 230067999 | 230069999 | ENSG00000 | SLC16A14 | 151473 | 9.461540932 | -0.4055939745 | 0.106891141 | -3.79445826 | 0.000147966 | 0.004823941 | RPE1_active | 0 | 0 |
| 84 | chr22 | 46052869 | 46054869 | ENSG00000 | RP4-695O20 | 400931 | 14.68505735 | -0.4015707158 | 0.094341458 | -4.25656677 | 2.08E-05 | 0.000963059 | RPE1_active | 0 | 0 |
| 85 | chr1 | 120068626 | 120070626 | ENSG00000 | NOTCH2 | 4853 | 29.25827005 | -0.3966391755 | 0.067713618 | -5.85759829 | 4.70E-09 | 7.65E-07 | RPE1_active | 0 | 0 |
| 86 | chr12 | 6198715 | 6200715 | ENSG00000 | CD9 | 928 | 11.67274996 | -0.3933801426 | 0.097306161 | -4.04270539 | 5.28E-05 | 0.002070288 | RPE1_active | 0 | 0 |
| 87 | chr1 | 223700593 | 223702593 | ENSG00000 | CAPN2 | 824 | 21.82428982 | -0.3927621387 | 0.076437808 | -5.13832284 | 2.77E-07 | 2.78E-05 | RPE1_active | 0 | 0 |
| 88 | chr1 | 218345235 | 218347235 | ENSG00000 | TGFB2 | 7042 | 27.27334291 | -0.3915373518 | 0.064398447 | -6.07991914 | 1.20E-09 | 2.30E-07 | RPE1_active | 0 | 0 |
| 89 | chr22 | 36387018 | 36389018 | ENSG00000 | MYH9 | 4627 | 24.01738005 | -0.387529104 | 0.070324790 | -5.51056179 | 3.58E-08 | 4.77E-06 | RPE1_active | 0 | 0 |
| 90 | chr3 | 45145422 | 45147422 | ENSG00000 | CDCP1 | 64866 | 19.35696442 | -0.3851801813 | 0.080247368 | -4.79991044 | 1.59E-06 | 0.000116531 | RPE1_active | 0 | 0 |
| 91 | chr2 | 9205765 | 9207765 | ENSG00000 | ASAP2 | 8853 | 57.46286917 | -0.3844644879 | 0.049483531 | -7.76954427 | 7.88E-15 | 4.05E-12 | RPE1_active | 0 | 0 |
| 92 | chr5 | 150154872 | 150156872 | ENSG00000 | PDGFRB | 5159 | 15.51581085 | -0.38241176 | 0.090001720 | -4.24893831 | 2.15E-05 | 0.000983642 | RPE1_active | 0 | 0 |
| 93 | chr1 | 157992273 | 157994273 | ENSG00000 | KIRREL | 55243 | 35.47108201 | -0.3810527079 | 0.057683291 | -6.60594600 | 3.95E-11 | 9.42E-09 | RPE1_active | 0 | 0 |
| 94 | chr12 | 52231520 | 52233520 | ENSG00000 | KRT7 | 3855 | 5.941850651 | -0.3787890155 | 0.138031266 | -2.74422618 | 0.006065373 | 0.072486091 | RPE1_active | 0 | 0 |
| 95 | chr5 | 55021671 | 55023671 | ENSG00000 | ESM1 | 11082 | 13.55407834 | -0.3756385707 | 0.092111970 | -4.07806463 | 4.54E-05 | 0.001819591 | RPE1_active | 0 | 0 |
| 96 | chr3 | 53255052 | 53257052 | ENSG00000 | TKT | 7086 | 8.715858297 | -0.3753484567 | 0.119565331 | -3.13927501 | 0.001693664 | 0.030415385 | RPE1_active | 0 | 0 |
| 97 | chr14 | 22981258 | 22983258 | ENSG00000 | RP11-298I3. | NaN | 9.414331006 | -0.3722318933 | 0.105708103 | -3.52131844 | 0.000429406 | 0.011590508 | RPE1_active | 0 | 0 |
| 98 | chr5 | 122128513 | 122130513 | ENSG00000 | ZNF474 | 133923 | 12.84803372 | -0.3703600309 | 0.093211687 | -3.97332179 | 7.09E-05 | 0.002667577 | RPE1_active | 0 | 0 |
| 99 | chr7 | 116523785 | 116525785 | ENSG00000 | CAV1 | 857 | 8.896745492 | -0.3701239125 | 0.117487569 | -3.15032401 | 0.001630894 | 0.029687173 | RPE1_active | 0 | 0 |
| 100 | chr5 | 150484818 | 150486818 | ENSG00000 | NDST1 | 3340 | 20.632771 | -0.3679738833 | 0.084544432 | -4.35243187 | 1.35E-05 | 0.000681389 | RPE1_active | 0 | 0 |
| 101 | chr17 | 64661068 | 64663068 | ENSG00000 | SMURF2 | 64750 | 22.35887967 | -0.3678674474 | 0.074240373 | -4.95508614 | 7.23E-07 | 6.31E-05 | RPE1_active | 0 | 0 |
| 102 | chr1 | 161580311 | 161582311 | ENSG00000 | FCGR2B | 2213 | 21.39169742 | -0.3653263951 | 0.070041748 | -5.21583772 | 1.83E-07 | 1.97E-05 | RPE1_active | 0 | 0 |
| 103 | chr9 | 38068211 | 38070211 | ENSG00000 | SHB | 6461 | 46.12993924 | -0.3651291484 | 0.055489923 | -6.58009829 | 4.70E-11 | 1.10E-08 | RPE1_active | 0 | 0 |
| 104 | chr3 | 139538829 | 139540829 | ENSG00000 | RBP1 | 5947 | 8.391445972 | -0.3630206408 | 0.111525059 | -3.25505892 | 0.001133688 | 0.022574084 | RPE1_active | 0 | 0 |
| 105 | chr1 | 152736518 | 152738518 | ENSG00000 | LCE1B | 353132 | 21.97036192 | -0.3598686774 | 0.071329658 | -5.04514787 | 4.53E-07 | 4.29E-05 | RPE1_active | 0 | 0 |
| 106 | chr2 | 110115566 | 110117566 | ENSG00000 | MALL | 7851 | 6.628863216 | -0.3598524037 | 0.137432193 | -2.61839963 | 0.008834327 | 0.094104075 | RPE1_active | 0 | 0 |
| 107 | chr9 | 178147 | 180147 | ENSG00000 | CBWD1 | 55871 | 6.17685527 | -0.3571873456 | 0.130858341 | -2.72957262 | 0.006341648 | 0.074586935 | RPE1_active | 0 | 0 |
| 108 | chr11 | 130915509 | 130917509 | ENSG00000 | SNX19 | 399979 | 11.52309153 | -0.3568470997 | 0.097831555 | -3.64756644 | 0.000264735 | 0.007756876 | RPE1_active | 0 | 0 |
| 109 | chr8 | 118110853 | 118112853 | ENSG00000 | EXT1 | 2131 | 81.56064617 | -0.3567588904 | 0.043002132 | -8.29630693 | 1.07E-16 | 7.55E-14 | RPE1_active | 0 | 0 |
| 110 | chr13 | 95643703 | 95645703 | ENSG00000 | DZIP1 | 22873 | 17.42462646 | -0.3564209853 | 0.082420261 | -4.32443403 | 1.53E-05 | 0.000748431 | RPE1_active | 0 | 0 |
| 111 | chr3 | 112648530 | 112650530 | ENSG00000 | CCDC80 | 151887 | 13.37703326 | -0.3560130704 | 0.086625488 | -4.10979580 | 3.96E-05 | 0.001623030 | RPE1_active | 0 | 0 |
| 112 | chr15 | 74211267 | 74213267 | ENSG00000 | STRA6 | 64220 | 12.09008317 | -0.3526025677 | 0.102582497 | -3.43725856 | 0.000587634 | 0.014178220 | RPE1_active | 0 | 0 |
| 113 | chr3 | 50154045 | 50156045 | ENSG00000 | SEMA3F | 6405 | 10.71557858 | -0.3523370215 | 0.111053644 | -3.17267411 | 0.001510419 | 0.027951130 | RPE1_active | 0 | 0 |
| 114 | chr1 | 209651466 | 209653466 | ENSG00000 | LAMB3 | 3914 | 13.45061167 | -0.3523277287 | 0.090858322 | -3.87777057 | 0.000105418 | 0.003681380 | RPE1_active | 0 | 0 |
| 115 | chr4 | 54063690 | 54065690 | ENSG00000 | CHIC2 | 26511 | 10.56964086 | -0.3515265295 | 0.107063633 | -3.28334205 | 0.001025841 | 0.021054167 | RPE1_active | 0 | 0 |

| | | | | | | | | | | | | | | | |
|---|---|---|---|---|---|---|---|---|---|---|---|---|---|---|---|
| 116 | chr11 | 76781251 | 76783251 | ENSG00000 | TSKU | 25987 | 5.61665089 | -0.3510418068 | 0.12688644 | -2.76658233 | 0.00566472 | 0.06950085 | RPE1_active | 0 | 0 |
| 117 | chr2 | 1743852 | 1745852 | ENSG00000 | PXDN | 7837 | 31.6767596 | -0.3510139716 | 0.06030644 | -5.82050496 | 5.87E-09 | 9.33E-07 | RPE1_active | 1 | 0 |
| 118 | chr1 | 12616731 | 12618731 | ENSG00000 | DHRS3 | 9249 | 15.2594275 | -0.3479891574 | 0.09861034 | -3.52893164 | 0.00041724 | 0.01135388 | RPE1_active | 0 | 0 |
| 119 | chr20 | 45630268 | 45632268 | ENSG00000 | WFDC9 | 259240 | 6.76471344 | -0.3471559768 | 0.12188033 | -2.84833454 | 0.00439486 | 0.05903885 | RPE1_active | 0 | 0 |
| 120 | chr8 | 1972878 | 1974878 | ENSG00000 | KBTBD11 | 9920 | 9.71596726 | -0.3426367141 | 0.10747566 | -3.18804007 | 0.00143240 | 0.02687975 | RPE1_active | 0 | 0 |
| 121 | chr4 | 109847202 | 109849202 | ENSG00000 | LRIT3 | 345193 | 6.59615698 | -0.3416877218 | 0.12841316 | -2.66084650 | 0.00779444 | 0.08653044 | RPE1_active | 0 | 0 |
| 122 | chr5 | 34655237 | 34657237 | ENSG00000 | RAI14 | 26064 | 46.6750327 | -0.338785942 | 0.05020278 | -6.74834932 | 1.50E-11 | 3.70E-09 | RPE1_active | 0 | 0 |
| 123 | chr11 | 130447974 | 130449974 | ENSG00000 | ADAMTS15 | 170689 | 9.72152585 | -0.3360125636 | 0.10608044 | -3.16752587 | 0.00153742 | 0.02829403 | RPE1_active | 0 | 0 |
| 124 | chr6 | 29456071 | 29458071 | ENSG00000 | OR11A1 | 26531 | 9.06237632 | -0.3340979634 | 0.11017377 | -3.03246351 | 0.00242566 | 0.03914166 | RPE1_active | 0 | 0 |
| 125 | chr3 | 123883331 | 123885331 | ENSG00000 | MYLK | 4638 | 80.8170572 | -0.3329177291 | 0.04083192 | -8.15336753 | 3.54E-16 | 2.25E-13 | RPE1_active | 0 | 0 |
| 126 | chr18 | 9913002 | 9915002 | ENSG00000 | VAPA | 9218 | 8.97254764 | -0.3325066653 | 0.11728577 | -2.83501278 | 0.00458238 | 0.06062104 | RPE1_active | 0 | 0 |
| 127 | chr8 | 91040064 | 91042064 | ENSG00000 | TMEM55A | 55529 | 9.99473522 | -0.3316155853 | 0.09873547 | -3.35862651 | 0.00078330 | 0.01732745 | RPE1_active | 0 | 0 |
| 128 | chr5 | 72106234 | 72108234 | ENSG00000 | MAP1B | 4131 | 26.8750017 | -0.3298836546 | 0.06391092 | -5.16161609 | 2.45E-07 | 2.50E-05 | RPE1_active | 0 | 0 |
| 129 | chr12 | 57127493 | 57129493 | ENSG00000 | LRP1 | 4035 | 25.9235217 | -0.3282885878 | 0.07149811 | -4.59156989 | 4.40E-06 | 0.00027086 | RPE1_active | 0 | 0 |
| 130 | chr11 | 27471775 | 27473775 | ENSG00000 | LGR4 | 55366 | 25.3328655 | -0.3274943516 | 0.06307766 | -5.19192253 | 2.08E-07 | 2.21E-05 | RPE1_active | 0 | 0 |
| 131 | chr17 | 41396592 | 41398592 | ENSG00000 | KRT33B | 3884 | 11.3154959 | -0.3271006147 | 0.09300827 | -3.51689805 | 0.00043662 | 0.01169723 | RPE1_active | 0 | 0 |
| 132 | chr5 | 151686165 | 151688165 | ENSG00000 | SPARC | 6678 | 7.67389769 | -0.3261581543 | 0.11229835 | -2.90438949 | 0.00367970 | 0.05252920 | RPE1_active | 0 | 0 |
| 133 | chr8 | 22543986 | 22545986 | ENSG00000 | SORBS3 | 10174 | 8.61444804 | -0.3255716648 | 0.12439581 | -2.61722370 | 0.00886482 | 0.09423727 | RPE1_active | 0 | 0 |
| 134 | chr2 | 210224491 | 210226491 | ENSG00000 | ACADL | 33 | 7.70921763 | -0.3251948797 | 0.12144616 | -2.67768748 | 0.00741323 | 0.08309132 | RPE1_active | 0 | 0 |
| 135 | chr17 | 15943917 | 15945917 | ENSG00000 | ADORA2B | 136 | 8.17351200 | -0.3209149855 | 0.12088913 | -2.65462217 | 0.00793972 | 0.08738279 | RPE1_active | 0 | 0 |
| 136 | chr12 | 106139033 | 106141033 | ENSG00000 | NUAK1 | 9891 | 24.5486205 | -0.3167892063 | 0.06570065 | -4.82170551 | 1.42E-06 | 0.00010744 | RPE1_active | 0 | 0 |
| 137 | chr12 | 1528891 | 1530891 | ENSG00000 | WNT5B | 81029 | 29.1751808 | -0.3154768421 | 0.06683427 | -4.72028494 | 2.36E-06 | 0.00016505 | RPE1_active | 0 | 0 |
| 138 | chr3 | 99637475 | 99639475 | ENSG00000 | COL8A1 | 1295 | 39.5032484 | -0.3153670926 | 0.05463597 | -5.77215043 | 7.83E-09 | 1.20E-06 | RPE1_active | 1 | 0 |
| 139 | chr16 | 85010535 | 85012535 | ENSG00000 | ZDHHC7 | 55625 | 9.51394304 | -0.3147205561 | 0.11358247 | -2.77085486 | 0.00559093 | 0.06884835 | RPE1_active | 0 | 0 |
| 140 | chr2 | 119222831 | 119224831 | ENSG00000 | STEAP3 | 55240 | 13.5826161 | -0.3144206864 | 0.09345737 | -3.36432190 | 0.00076731 | 0.01708693 | RPE1_active | 0 | 0 |
| 141 | chr18 | 8704661 | 8706661 | ENSG00000 | MTCL1 | 23255 | 35.7492140 | -0.3138164853 | 0.05400337 | -5.81105281 | 6.21E-09 | 9.76E-07 | RPE1_active | 0 | 0 |
| 142 | chr2 | 105437513 | 105439513 | ENSG00000 | FHL2 | 2274 | 22.3153977 | -0.3130923416 | 0.07378302 | -4.24341964 | 2.20E-05 | 0.00100384 | RPE1_active | 0 | 0 |
| 143 | chr21 | 15064000 | 15066000 | ENSG00000 | NRIP1 | 8204 | 22.0385954 | -0.3098507109 | 0.07256295 | -4.27009478 | 1.95E-05 | 0.00091600 | RPE1_active | 0 | 0 |
| 144 | chr19 | 51794163 | 51796163 | ENSG00000 | FPR3 | 2359 | 10.4070144 | -0.3093206763 | 0.10460614 | -2.95700287 | 0.00310645 | 0.04663517 | RPE1_active | 0 | 0 |
| 145 | chr12 | 52248300 | 52250300 | ENSG00000 | KRT86 | 3892 | 18.5417442 | -0.3083768889 | 0.07756555 | -3.97569364 | 7.02E-05 | 0.00265609 | RPE1_active | 0 | 0 |
| 146 | chr5 | 83470465 | 83472465 | ENSG00000 | VCAN | 1462 | 24.5822564 | -0.3056817892 | 0.07030118 | -4.34817380 | 1.37E-05 | 0.00068694 | RPE1_active | 0 | 0 |
| 147 | chr11 | 118605440 | 118607440 | ENSG00000 | PHLDB1 | 23187 | 17.0816898 | -0.3042687824 | 0.08052118 | -3.77874208 | 0.00015762 | 0.00506248 | RPE1_active | 0 | 0 |
| 148 | chr10 | 33004792 | 33006792 | ENSG00000 | ITGB1 | 3688 | 22.8390758 | -0.3032595392 | 0.06622658 | -4.57912118 | 4.67E-06 | 0.00028617 | RPE1_active | 0 | 0 |
| 149 | chr15 | 89654451 | 89656451 | ENSG00000 | KIF7 | 374654 | 13.6273862 | -0.3006949773 | 0.10310529 | -2.91638734 | 0.00354110 | 0.05142686 | RPE1_active | 0 | 0 |
| 150 | chr1 | 205210566 | 205212566 | ENSG00000 | DSTYK | 25778 | 18.8367534 | -0.300279301 | 0.07248386 | -4.14270548 | 3.43E-05 | 0.00143310 | RPE1_active | 0 | 0 |
| 151 | chr7 | 55018021 | 55020021 | ENSG00000 | EGFR | 1956 | 68.1674153 | -0.2981974674 | 0.04425317 | -6.73844217 | 1.60E-11 | 3.89E-09 | RPE1_active | 0 | 0 |
| 152 | chr13 | 110304812 | 110306812 | ENSG00000 | COL4A2 | 1284 | 61.6030842 | -0.2973654722 | 0.04661385 | -6.37933676 | 1.78E-10 | 3.83E-08 | RPE1_active | 0 | 0 |
| 153 | chr5 | 136027895 | 136029895 | ENSG00000 | TGFBI | 7045 | 11.0347669 | -0.2971538209 | 0.10116308 | -2.93737412 | 0.00331004 | 0.04892202 | RPE1_active | 0 | 0 |
| 154 | chr11 | 12673591 | 12675591 | ENSG00000 | TEAD1 | 7003 | 75.8205892 | -0.2930472043 | 0.04007389 | -7.31267177 | 2.62E-13 | 1.03E-10 | RPE1_active | 0 | 0 |
| 155 | chr12 | 108856590 | 108858590 | ENSG00000 | SSH1 | 54434 | 20.3692339 | -0.2918086765 | 0.07931024 | -3.6793314 | 0.00023384 | 0.007021167 | RPE1_active | 0 | 0 |
| 156 | chr2 | 46697952 | 46699952 | ENSG00000 | SOCS5 | 9655 | 13.1668664 | -0.291762777 | 0.08665514 | -3.36694115 | 0.00076006 | 0.01703906 | RPE1_active | 0 | 0 |
| 157 | chr8 | 22163736 | 22165736 | ENSG00000 | BMP1 | 649 | 14.1068005 | -0.2911159648 | 0.08808033 | -3.30511878 | 0.00094936 | 0.01997546 | RPE1_active | 0 | 0 |
| 158 | chr12 | 96906223 | 96908223 | ENSG00000 | NEDD1 | 121441 | 9.22252868 | -0.2907763978 | 0.10968352 | -2.65104896 | 0.00802422 | 0.08809499 | RPE1_active | 0 | 0 |
| 159 | chr8 | 25183723 | 25185723 | ENSG00000 | DOCK5 | 80005 | 60.0140684 | -0.2905432611 | 0.04108950 | -7.07098455 | 1.54E-12 | 4.47E-10 | RPE1_active | 0 | 0 |
| 160 | chr3 | 124900418 | 124902418 | ENSG00000 | ITGB5 | 3693 | 39.8708449 | -0.2887586266 | 0.05281217 | -5.46765236 | 4.56E-08 | 5.86E-06 | RPE1_active | 0 | 0 |
| 161 | chr4 | 5710197 | 5712197 | ENSG00000 | EVC | 2121 | 30.3939520 | -0.2877647578 | 0.06634550 | -4.33736610 | 1.44E-05 | 0.00071623 | RPE1_active | 0 | 0 |
| 162 | chr8 | 96492351 | 96494351 | ENSG00000 | SDC2 | 6383 | 32.5734476 | -0.2871263715 | 0.05702139 | -5.03541459 | 4.77E-07 | 4.49E-05 | RPE1_active | 0 | 0 |
| 163 | chr10 | 58512140 | 58514140 | ENSG00000 | BICC1 | 80114 | 79.2810925 | -0.2858087227 | 0.03570921 | -8.00378161 | 1.21E-15 | 7.01E-13 | RPE1_active | 0 | 0 |
| 164 | chr2 | 69012178 | 69014178 | ENSG00000 | ANTXR1 | 84168 | 69.206734 | -0.2832360463 | 0.03948884 | -7.17255859 | 7.36E-13 | 2.46E-10 | RPE1_active | 0 | 0 |
| 165 | chr18 | 35496991 | 35498991 | ENSG00000 | INO80C | 125476 | 12.2467999 | -0.2831416894 | 0.10022630 | -2.82502379 | 0.00472771 | 0.06174676 | RPE1_active | 0 | 0 |
| 166 | chr12 | 109476544 | 109478544 | ENSG00000 | KCTD10 | 83892 | 9.56827259 | -0.2822329819 | 0.10303820 | -2.73910998 | 0.00616057 | 0.07307264 | RPE1_active | 0 | 0 |
| 167 | chr15 | 67062763 | 67064763 | ENSG00000 | SMAD3 | 4088 | 34.5348607 | -0.2819096024 | 0.06201571 | -4.54577694 | 5.47E-06 | 0.00032646 | RPE1_active | 0 | 0 |
| 168 | chr15 | 60401883 | 60403883 | ENSG00000 | ANXA2 | 302 | 12.1931191 | -0.2812341202 | 0.09883722 | -2.84542714 | 0.00443519 | 0.05943688 | RPE1_active | 0 | 0 |
| 169 | chr15 | 66701228 | 66703228 | ENSG00000 | SMAD6 | 4091 | 23.8635372 | -0.2805705744 | 0.07385113 | -3.79913696 | 0.00014520 | 0.00479019 | RPE1_active | 0 | 0 |
| 170 | chr4 | 122825708 | 122827708 | ENSG00000 | FGF2 | 2247 | 16.0680885 | -0.2789885309 | 0.08406196 | -3.31884376 | 0.00090390 | 0.01944789 | RPE1_active | 0 | 0 |
| 171 | chr2 | 46815697 | 46817697 | ENSG00000 | LINC01119 | 100134259 | 13.1562184 | -0.2764976229 | 0.09159089 | -3.01883323 | 0.00253750 | 0.04045770 | RPE1_active | 0 | 0 |
| 172 | chr6 | 149863026 | 149865026 | ENSG00000 | LRP11 | 84918 | 12.1309675 | -0.2760166969 | 0.09486529 | -2.90956460 | 0.00361932 | 0.05215503 | RPE1_active | 0 | 0 |
| 173 | chr2 | 112274594 | 112276594 | ENSG00000 | ZC3H6 | 376940 | 13.5558984 | -0.2755779681 | 0.09221793 | -2.98833385 | 0.00280502 | 0.04352845 | RPE1_active | 0 | 0 |
| 174 | chr2 | 26691690 | 26693690 | ENSG00000 | KCNK3 | 3777 | 13.7613692 | -0.2753210087 | 0.08564045 | -3.21484744 | 0.00130513 | 0.02509058 | RPE1_active | 0 | 0 |
| 175 | chr9 | 124414444 | 124416444 | ENSG00000 | PSMB7 | 5695 | 15.9330102 | -0.2752992386 | 0.07718345 | -3.56681664 | 0.00036134 | 0.01010024 | RPE1_active | 0 | 0 |

| | | | | | | | | | | | | | | | |
|---|---|---|---|---|---|---|---|---|---|---|---|---|---|---|---|
| 176 | chr11 | 86671636 | 86673636 | ENSG00000 | ME3 | 10873 | 73.34620041 | -0.2737643883 | 0.03903915 | -7.01255925 | 2.34E-12 | 6.51E-10 | RPE1_active | 0 | 0 |
| 177 | chr6 | 52575915 | 52577915 | ENSG00000 | TRAM2 | 9697 | 20.98369057 | -0.2729293059 | 0.07791260 | -3.50301883 | 0.00046001 | 0.01209393 | RPE1_active | 0 | 0 |
| 178 | chr5 | 120463278 | 120465278 | ENSG00000 | PRR16 | 51334 | 45.43180622 | -0.2725641911 | 0.04869546 | -5.59732130 | 2.18E-08 | 3.06E-06 | RPE1_active | 0 | 0 |
| 179 | chr18 | 58043367 | 58045367 | ENSG00000 | NEDD4L | 23327 | 100.1525757 | -0.2720979874 | 0.03687634 | -7.37865898 | 1.60E-13 | 6.47E-11 | RPE1_active | 0 | 0 |
| 180 | chr3 | 197298300 | 197300300 | ENSG00000 | DLG1 | 1739 | 45.6226436 | -0.2715526644 | 0.05580481 | -4.86611500 | 1.14E-06 | 9.16E-05 | RPE1_active | 0 | 0 |
| 181 | chr18 | 59696380 | 59698380 | ENSG00000 | CCBE1 | 147372 | 80.10849203 | -0.2709504441 | 0.03816009 | -7.10036012 | 1.24E-12 | 3.87E-10 | RPE1_active | 0 | 0 |
| 182 | chr14 | 64703102 | 64705102 | ENSG00000 | PLEKHG3 | 26030 | 15.19318913 | -0.2708986293 | 0.09451157 | -2.86630094 | 0.00415299 | 0.05732362 | RPE1_active | 0 | 0 |
| 183 | chr20 | 38165578 | 38167578 | ENSG00000 | TGM2 | 7052 | 11.37234989 | -0.2696422077 | 0.10208342 | -2.64139061 | 0.00825664 | 0.08983473 | RPE1_active | 0 | 0 |
| 184 | chrX | 38326564 | 38328564 | ENSG00000 | RPGR | 6103 | 13.82509652 | -0.2695417317 | 0.08572141 | -3.14439214 | 0.00166432 | 0.03001524 | RPE1_active | 0 | 0 |
| 185 | chr14 | 102304200 | 102306200 | ENSG00000 | MOK | 5891 | 20.57455187 | -0.2691082316 | 0.08139872 | -3.30604952 | 0.00094621 | 0.01994063 | RPE1_active | 0 | 0 |
| 186 | chr15 | 51828628 | 51830628 | ENSG00000 | TMOD3 | 29766 | 27.67872375 | -0.2688386378 | 0.06801545 | -3.95261084 | 7.73E-05 | 0.00286726 | RPE1_active | 0 | 0 |
| 187 | chr9 | 98254721 | 98256721 | ENSG00000 | TBC1D2 | 55357 | 17.81172301 | -0.2677545945 | 0.07930931 | -3.37607982 | 0.00073526 | 0.01659440 | RPE1_active | 0 | 0 |
| 188 | chr6 | 75205051 | 75207051 | ENSG00000 | COL12A1 | 1303 | 27.3383446 | -0.2669141245 | 0.06552137 | -4.07369547 | 4.63E-05 | 0.00184553 | RPE1_active | 0 | 0 |
| 189 | chr15 | 100876714 | 100878714 | ENSG00000 | ALDH1A3 | 220 | 11.28543357 | -0.2664714407 | 0.10015915 | -2.66048003 | 0.00780293 | 0.08653044 | RPE1_active | 0 | 0 |
| 190 | chr12 | 25958029 | 25960029 | ENSG00000 | RASSF8 | 11228 | 27.83286745 | -0.2663831016 | 0.06231556 | -4.27474449 | 1.91E-05 | 0.00090025 | RPE1_active | 0 | 0 |
| 191 | chr7 | 116671390 | 116673390 | ENSG00000 | MET | 4233 | 32.57444329 | -0.2661933676 | 0.05732690 | -4.64342851 | 3.43E-06 | 0.00022118 | RPE1_active | 0 | 0 |
| 192 | chr1 | 203625561 | 203627561 | ENSG00000 | ATP2B4 | 493 | 33.47331718 | -0.2659812513 | 0.05994575 | -4.43703235 | 9.12E-06 | 0.00050565 | RPE1_active | 0 | 0 |
| 193 | chr3 | 159838861 | 159840861 | ENSG00000 | SCHIP1 | 29970 | 14.32410515 | -0.2647266047 | 0.08898264 | -2.97503636 | 0.00292953 | 0.04493864 | RPE1_active | 0 | 0 |
| 194 | chr4 | 156970394 | 156972394 | ENSG00000 | PDGFC | 56034 | 44.35441089 | -0.2628632979 | 0.05494767 | -4.78388374 | 1.72E-06 | 0.00012553 | RPE1_active | 0 | 0 |
| 195 | chr17 | 13600927 | 13602927 | ENSG00000 | HS3ST3A1 | 9955 | 27.43637876 | -0.2627714492 | 0.06063423 | -4.33371431 | 1.47E-05 | 0.00072552 | RPE1_active | 0 | 0 |
| 196 | chr2 | 207166267 | 207168267 | ENSG00000 | KLF7 | 8609 | 25.64421983 | -0.2622853358 | 0.07071963 | -3.70880499 | 0.00020823 | 0.00639607 | RPE1_active | 0 | 0 |
| 197 | chr1 | 94274068 | 94276068 | ENSG00000 | ARHGAP29 | 9411 | 27.59351883 | -0.2621449928 | 0.06950745 | -3.77146571 | 0.00016229 | 0.00517512 | RPE1_active | 0 | 0 |
| 198 | chr5 | 33891192 | 33893192 | ENSG00000 | ADAMTS12 | 81792 | 99.61093221 | -0.2619982988 | 0.03774593 | -6.94109949 | 3.89E-12 | 1.06E-09 | RPE1_active | 0 | 0 |
| 199 | chr11 | 34512805 | 34514805 | ENSG00000 | ELF5 | 2001 | 12.18983134 | -0.2602618718 | 0.09359384 | -2.78075831 | 0.00542320 | 0.06727901 | RPE1_active | 0 | 0 |
| 200 | chr1 | 67832467 | 67834467 | ENSG00000 | GNG12 | 55970 | 35.10302709 | -0.259663434 | 0.05624572 | -4.61658935 | 3.90E-06 | 0.00024381 | RPE1_active | 0 | 0 |
| 201 | chr6 | 108558835 | 108560835 | ENSG00000 | FOXO3 | 2309 | 31.36680106 | -0.2591762709 | 0.06022001 | -4.30382295 | 1.68E-05 | 0.00079821 | RPE1_active | 0 | 0 |
| 202 | chr9 | 96874623 | 96876623 | ENSG00000 | ZNF782 | 158431 | 14.43505462 | -0.2586915518 | 0.08576945 | -3.01612671 | 0.00256026 | 0.04077195 | RPE1_active | 0 | 0 |
| 203 | chr14 | 74611378 | 74613378 | ENSG00000 | LTBP2 | 4053 | 39.9799127 | -0.2583371511 | 0.05648719 | -4.57337541 | 4.80E-06 | 0.00029146 | RPE1_active | 0 | 0 |
| 204 | chr8 | 30383479 | 30385479 | ENSG00000 | RBPMS | 11030 | 46.38557533 | -0.2583305237 | 0.05171066 | -4.99569143 | 5.86E-07 | 5.26E-05 | RPE1_active | 0 | 0 |
| 205 | chr5 | 149140483 | 149142483 | ENSG00000 | ABLIM3 | 22885 | 39.10568733 | -0.2582719915 | 0.05719309 | -4.51578961 | 6.31E-06 | 0.00036804 | RPE1_active | 0 | 0 |
| 206 | chr2 | 10301889 | 10303889 | ENSG00000 | HPCAL1 | 3241 | 37.14616131 | -0.2575260861 | 0.05981940 | -4.30505919 | 1.67E-05 | 0.00079660 | RPE1_active | 0 | 0 |
| 207 | chr11 | 12376185 | 12378185 | ENSG00000 | PARVA | 55742 | 47.40701355 | -0.2566420236 | 0.05157391 | -4.97619819 | 6.48E-07 | 5.76E-05 | RPE1_active | 0 | 0 |
| 208 | chr19 | 46917676 | 46919676 | ENSG00000 | ARHGAP35 | 2909 | 23.78653267 | -0.2563595651 | 0.07303827 | -3.50993438 | 0.00044821 | 0.01190582 | RPE1_active | 0 | 0 |
| 209 | chr1 | 31703319 | 31705319 | ENSG00000 | COL16A1 | 1307 | 19.17290722 | -0.2545952873 | 0.08568743 | -2.97120915 | 0.00296629 | 0.04538703 | RPE1_active | 0 | 0 |
| 210 | chr4 | 168496066 | 168498066 | ENSG00000 | PALLD | 23022 | 106.5169973 | -0.2545115152 | 0.03589248 | -7.09094170 | 1.33E-12 | 3.95E-10 | RPE1_active | 0 | 0 |
| 211 | chr2 | 172426354 | 172428354 | ENSG00000 | ITGA6 | 3655 | 21.93030793 | -0.2536873387 | 0.06780918 | -3.74119430 | 0.00018314 | 0.00574422 | RPE1_active | 0 | 0 |
| 212 | chr1 | 223844972 | 223846972 | ENSG00000 | TP53BP2 | 7159 | 15.38138612 | -0.2528677917 | 0.09144473 | -2.76525275 | 0.00568787 | 0.06962430 | RPE1_active | 0 | 0 |
| 213 | chr8 | 56210324 | 56212324 | ENSG00000 | PLAG1 | 5324 | 12.82188754 | -0.252648789 | 0.08814484 | -2.86629111 | 0.00415312 | 0.05732362 | RPE1_active | 0 | 0 |
| 214 | chr12 | 96193382 | 96195382 | ENSG00000 | ELK3 | 2004 | 19.80966637 | -0.252042554 | 0.07503137 | -3.35916223 | 0.00078179 | 0.01732258 | RPE1_active | 0 | 0 |
| 215 | chr9 | 39287315 | 39289315 | ENSG00000 | CNTNAP3 | 79937 | 18.11220969 | -0.2519796216 | 0.08272573 | -3.04596418 | 0.00231935 | 0.03783748 | RPE1_active | 0 | 0 |
| 216 | chr14 | 74712115 | 74714115 | ENSG00000 | AREL1 | 9870 | 15.85137597 | -0.2510974647 | 0.08460909 | -2.96773615 | 0.00300001 | 0.04570493 | RPE1_active | 0 | 0 |
| 217 | chr3 | 69738435 | 69740435 | ENSG00000 | MITF | 4286 | 62.99046575 | -0.250881089 | 0.04661593 | -5.38187388 | 7.37E-08 | 9.04E-06 | RPE1_active | 0 | 0 |
| 218 | chr1 | 63773022 | 63775022 | ENSG00000 | ROR1 | 4919 | 119.6363548 | -0.2492302368 | 0.03272303 | -7.61635427 | 2.61E-14 | 1.20E-11 | RPE1_active | 0 | 0 |
| 219 | chr6 | 52419308 | 52421308 | ENSG00000 | EFHC1 | 114327 | 19.42678189 | -0.2491453933 | 0.07597545 | -3.27928802 | 0.00104069 | 0.02116393 | RPE1_active | 0 | 0 |
| 220 | chr4 | 186725722 | 186727722 | ENSG00000 | FAT1 | 2195 | 36.70202742 | -0.2473326191 | 0.05738989 | -4.30968950 | 1.63E-05 | 0.00078841 | RPE1_active | 1 | 0 |
| 221 | chr1 | 36384896 | 36386896 | ENSG00000 | STK40 | 83931 | 14.30832702 | -0.2470761592 | 0.08472995 | -2.91604262 | 0.00354502 | 0.05142781 | RPE1_active | 0 | 0 |
| 222 | chr12 | 120264771 | 120266771 | ENSG00000 | PXN | 5829 | 15.45076804 | -0.2461436557 | 0.09070675 | -2.71362002 | 0.00665524 | 0.07737602 | RPE1_active | 0 | 0 |
| 223 | chr7 | 80921359 | 80923359 | ENSG00000 | SEMA3C | 10512 | 37.40471607 | -0.2456769131 | 0.05481037 | -4.48230647 | 7.38E-06 | 0.00041982 | RPE1_active | 0 | 0 |
| 224 | chr9 | 78030458 | 78032458 | ENSG00000 | GNAQ | 2776 | 78.73213042 | -0.2449161349 | 0.04049946 | -6.04739211 | 1.47E-09 | 2.73E-07 | RPE1_active | 0 | 0 |
| 225 | chr20 | 9067763 | 9069763 | ENSG00000 | PLCB4 | 5332 | 104.7071702 | -0.2443716204 | 0.03965261 | -6.16281177 | 7.15E-10 | 1.43E-07 | RPE1_active | 0 | 0 |
| 226 | chr14 | 102138503 | 102140503 | ENSG00000 | WDR20 | 91833 | 22.71677592 | -0.2420617975 | 0.07339074 | -3.29826038 | 0.00097285 | 0.02033010 | RPE1_active | 0 | 0 |
| 227 | chr1 | 183022460 | 183024460 | ENSG00000 | LAMC1 | 3915 | 31.26198674 | -0.2395339524 | 0.05476202 | -4.37408866 | 1.22E-05 | 0.00062905 | RPE1_active | 0 | 0 |
| 228 | chr2 | 215435172 | 215437172 | ENSG00000 | FN1 | 2335 | 17.31670052 | -0.2394748962 | 0.08241071 | -2.90587077 | 0.00366232 | 0.05244624 | RPE1_active | 1 | 0 |
| 229 | chr2 | 218001995 | 218003995 | ENSG00000 | TNS1 | 7145 | 73.67077483 | -0.2394275289 | 0.04583586 | -5.22358499 | 1.75E-07 | 1.91E-05 | RPE1_active | 1 | 0 |
| 230 | chr2 | 26969612 | 26971612 | ENSG00000 | MAPRE3 | 22924 | 18.95486263 | -0.2393922122 | 0.07647228 | -3.13044399 | 0.00174542 | 0.03113563 | RPE1_active | 0 | 0 |
| 231 | chr1 | 67053099 | 67055099 | ENSG00000 | SLC35D1 | 23169 | 13.1510855 | -0.2392562615 | 0.09125093 | -2.62195947 | 0.00874258 | 0.09337301 | RPE1_active | 0 | 0 |
| 232 | chr12 | 1687574 | 1689574 | ENSG00000 | ADIPOR2 | 79602 | 23.14886783 | -0.2389673984 | 0.07183442 | -3.32664157 | 0.00087899 | 0.01900361 | RPE1_active | 0 | 0 |
| 233 | chr22 | 20494913 | 20496913 | ENSG00000 | MED15 | 51586 | 24.33175348 | -0.2370046268 | 0.06862020 | -3.45386047 | 0.00055262 | 0.01372416 | RPE1_active | 0 | 0 |
| 234 | chr11 | 94705431 | 94707431 | ENSG00000 | AMOTL1 | 154810 | 51.51978134 | -0.2366971057 | 0.04816973 | -4.91381356 | 8.93E-07 | 7.41E-05 | RPE1_active | 0 | 0 |
| 235 | chr12 | 63843293 | 63845293 | ENSG00000 | SRGAP1 | 57522 | 78.22526679 | -0.2363052709 | 0.03979078 | -5.93869395 | 2.87E-09 | 4.80E-07 | RPE1_active | 0 | 0 |

| | | | | | | | | | | | | | | | |
|---|---|---|---|---|---|---|---|---|---|---|---|---|---|---|---|
| 236 | chrX | 38219924 | 38221924 | ENSG00000 | SRPX | 8406 | 22.12630808 | -0.2360301171 | 0.07217890 | -3.27007048 | 0.00107520 | 0.02166793 | RPE1_active | 0 | 0 |
| 237 | chr2 | 27770717 | 27772717 | ENSG00000 | MRPL33 | 9553 | 54.3112715 | -0.2352157468 | 0.04583106 | -5.13223367 | 2.86E-07 | 2.83E-05 | RPE1_active | 0 | 0 |
| 238 | chr12 | 104214779 | 104216779 | ENSG00000 | TXNRD1 | 7296 | 30.64083123 | -0.2344613133 | 0.05625454 | -4.16786413 | 3.07E-05 | 0.00132092 | RPE1_active | 0 | 0 |
| 239 | chr14 | 101963528 | 101965528 | ENSG00000 | DYNC1H1 | 1778 | 23.2775181 | -0.2341143912 | 0.06858436 | -3.41352424 | 0.00064128 | 0.01507418 | RPE1_active | 0 | 0 |
| 240 | chr16 | 68529090 | 68531090 | ENSG00000 | ZFP90 | 146198 | 13.48265216 | -0.2341047438 | 0.08431083 | -2.77668625 | 0.00549161 | 0.06781282 | RPE1_active | 0 | 0 |
| 241 | chr2 | 191244185 | 191246185 | ENSG00000 | MYO1B | 4430 | 48.4320483 | -0.233843598 | 0.04954411 | -4.71990651 | 2.36E-06 | 0.00016505 | RPE1_active | 0 | 0 |
| 242 | chr1 | 162068774 | 162070774 | ENSG00000 | NOS1AP | 9722 | 99.46274086 | -0.2333048819 | 0.03702147 | -6.30187942 | 2.94E-10 | 6.04E-08 | RPE1_active | 0 | 0 |
| 243 | chr4 | 76212893 | 76214893 | ENSG00000 | SCARB2 | 950 | 14.96107808 | -0.2325866003 | 0.08278012 | -2.80969143 | 0.00495890 | 0.06383094 | RPE1_active | 0 | 0 |
| 244 | chr13 | 110306149 | 110308149 | ENSG00000 | COL4A1 | 1282 | 46.07820328 | -0.2319716553 | 0.05236562 | -4.42984597 | 9.43E-06 | 0.00051637 | RPE1_active | 1 | 0 |
| 245 | chr8 | 19312617 | 19314617 | ENSG00000 | SH2D4A | 63898 | 23.54859438 | -0.2306124233 | 0.06897157 | -3.34358629 | 0.00082702 | 0.01808501 | RPE1_active | 0 | 0 |
| 246 | chr1 | 222871271 | 222873271 | ENSG00000 | DISP1 | 84976 | 31.84653746 | -0.2288845591 | 0.06509518 | -3.51615173 | 0.00043785 | 0.01169723 | RPE1_active | 0 | 0 |
| 247 | chr4 | 119627991 | 119629991 | ENSG00000 | PDE5A | 8654 | 30.08790283 | -0.2287030156 | 0.06583104 | -3.47409044 | 0.00051258 | 0.01309501 | RPE1_active | 0 | 0 |
| 248 | chr13 | 21060547 | 21062547 | ENSG00000 | LATS2 | 26524 | 22.00925056 | -0.2279418941 | 0.06993962 | -3.25912387 | 0.001117568 | 0.02231963 | RPE1_active | 0 | 0 |
| 249 | chr12 | 65277643 | 65279643 | ENSG00000 | MSRB3 | 253827 | 49.54923398 | -0.2271124563 | 0.04687918 | -4.84463334 | 1.27E-06 | 9.91E-05 | RPE1_active | 0 | 0 |
| 250 | chr5 | 39461300 | 39463300 | ENSG00000 | DAB2 | 1601 | 23.82143429 | -0.2267836062 | 0.06535952 | -3.46978638 | 0.00052087 | 0.01320564 | RPE1_active | 0 | 0 |
| 251 | chr6 | 37698306 | 37700306 | ENSG00000 | MDGA1 | 266727 | 21.67285397 | -0.2237404684 | 0.07950196 | -2.81427587 | 0.00488872 | 0.06329288 | RPE1_active | 0 | 0 |
| 252 | chr2 | 101473703 | 101475703 | ENSG00000 | RFX8 | 731220 | 24.60545721 | -0.2236601609 | 0.06430539 | -3.47809302 | 0.00050499 | 0.01295054 | RPE1_active | 0 | 0 |
| 253 | chr20 | 31513428 | 31515428 | ENSG00000 | HM13 | 81502 | 17.64420887 | -0.2232941148 | 0.07711950 | -2.89542990 | 0.00378639 | 0.05353442 | RPE1_active | 0 | 0 |
| 254 | chr22 | 45670798 | 45672798 | ENSG00000 | ATXN10 | 25814 | 42.44276727 | -0.2227514533 | 0.05349802 | -4.16373204 | 3.13E-05 | 0.00134075 | RPE1_active | 0 | 0 |
| 255 | chr1 | 214551449 | 214553449 | ENSG00000 | PTPN14 | 5784 | 52.53713223 | -0.2225278727 | 0.04950967 | -4.49463368 | 6.97E-06 | 0.00040246 | RPE1_active | 0 | 0 |
| 256 | chr14 | 50560126 | 50562126 | ENSG00000 | MAP4K5 | 11183 | 30.62293343 | -0.2222725189 | 0.06443824 | -3.44938808 | 0.00056185 | 0.01382503 | RPE1_active | 0 | 0 |
| 257 | chr16 | 57862053 | 57864053 | ENSG00000 | KIFC3 | 3801 | 32.3018461 | -0.2219424765 | 0.05813858 | -3.81747271 | 0.00013482 | 0.00448110 | RPE1_active | 0 | 0 |
| 258 | chr8 | 23456695 | 23458695 | ENSG00000 | ENTPD4 | 9583 | 20.38067819 | -0.2206281227 | 0.07260359 | -3.03880454 | 0.00237518 | 0.03841997 | RPE1_active | 0 | 0 |
| 259 | chr2 | 187564760 | 187566760 | ENSG00000 | TFPI | 7035 | 23.22268133 | -0.2205409288 | 0.06921671 | -3.18623802 | 0.00144135 | 0.02697199 | RPE1_active | 0 | 0 |
| 260 | chr3 | 28240584 | 28242584 | ENSG00000 | CMC1 | 152100 | 18.6037801 | -0.2204161706 | 0.08296337 | -2.65678882 | 0.00788888 | 0.08703830 | RPE1_active | 0 | 0 |
| 261 | chr11 | 47979558 | 47981558 | ENSG00000 | PTPRJ | 5795 | 49.72215884 | -0.2184981478 | 0.04919639 | -4.44134501 | 8.94E-06 | 0.00049768 | RPE1_active | 0 | 0 |
| 262 | chr8 | 141001216 | 141003216 | ENSG00000 | PTK2 | 5747 | 83.36641083 | -0.2179292841 | 0.03587482 | -6.07471319 | 1.24E-09 | 2.34E-07 | RPE1_active | 0 | 0 |
| 263 | chr15 | 48644849 | 48646849 | ENSG00000 | FBN1 | 2200 | 59.77667976 | -0.2176941147 | 0.04493000 | -4.84518337 | 1.26E-06 | 9.91E-05 | RPE1_active | 0 | 0 |
| 264 | chr6 | 35341558 | 35343558 | ENSG00000 | PPARD | 5467 | 26.41881813 | -0.2170631008 | 0.06860855 | -3.16379051 | 0.00155728 | 0.02858094 | RPE1_active | 0 | 0 |
| 265 | chr6 | 157980887 | 157982887 | ENSG00000 | SYNJ2 | 8871 | 33.53641241 | -0.2167344122 | 0.05831779 | -3.71643707 | 0.00020205 | 0.00627817 | RPE1_active | 0 | 0 |
| 266 | chr2 | 233565782 | 233567782 | ENSG00000 | USP40 | 55230 | 21.55098075 | -0.2153531476 | 0.06894864 | -3.12338475 | 0.00178783 | 0.03151360 | RPE1_active | 0 | 0 |
| 267 | chr17 | 15683311 | 15685311 | ENSG00000 | TRIM16 | 10626 | 14.81855619 | -0.2145916983 | 0.08228582 | -2.60788185 | 0.00911043 | 0.09606689 | RPE1_active | 0 | 0 |
| 268 | chr22 | 36027425 | 36029425 | ENSG00000 | RBFOX2 | 23543 | 72.16248759 | -0.2142262972 | 0.04084671 | -5.24464019 | 1.57E-07 | 1.74E-05 | RPE1_active | 0 | 0 |
| 269 | chr18 | 28176446 | 28178446 | ENSG00000 | CDH2 | 1000 | 53.48234255 | -0.2137823021 | 0.05098239 | -4.19325739 | 2.75E-05 | 0.00120458 | RPE1_active | 0 | 0 |
| 270 | chr2 | 120734623 | 120736623 | ENSG00000 | GLI2 | 2736 | 93.43902203 | -0.2136177462 | 0.04201818 | -5.08393571 | 3.70E-07 | 3.63E-05 | RPE1_active | 0 | 0 |
| 271 | chr14 | 32075114 | 32077114 | ENSG00000 | ARHGAP5 | 394 | 17.48122986 | -0.2122980277 | 0.08210490 | -2.58569253 | 0.009718357 | 0.09949960 | RPE1_active | 0 | 0 |
| 272 | chr13 | 113758240 | 113760240 | ENSG00000 | TMEM255B | 348013 | 17.15384368 | -0.2114681123 | 0.07742208 | -2.73136685 | 0.00630722 | 0.07437845 | RPE1_active | 0 | 0 |
| 273 | chr8 | 130442660 | 130444660 | ENSG00000 | ASAP1 | 50807 | 103.1445907 | -0.2106232783 | 0.03563175 | -5.91111221 | 3.40E-09 | 5.61E-07 | RPE1_active | 0 | 0 |
| 274 | chr1 | 27603431 | 27605431 | ENSG00000 | AHDC1 | 27245 | 21.47720469 | -0.2102623967 | 0.07819507 | -2.68894694 | 0.00716778 | 0.08136679 | RPE1_active | 0 | 0 |
| 275 | chr5 | 14142702 | 14144702 | ENSG00000 | TRIO | 7204 | 98.69697034 | -0.2100594085 | 0.03905417 | -5.37866736 | 7.50E-08 | 9.11E-06 | RPE1_active | 0 | 0 |
| 276 | chr3 | 111673676 | 111675676 | ENSG00000 | PLCXD2 | 257068 | 47.15854726 | -0.2099917045 | 0.04874065 | -4.30834793 | 1.64E-05 | 0.00078841 | RPE1_active | 0 | 0 |
| 277 | chr2 | 101695850 | 101697850 | ENSG00000 | MAP4K4 | 9448 | 51.45696948 | -0.2092842208 | 0.04924581 | -4.24978711 | 2.14E-05 | 0.00098364 | RPE1_active | 0 | 0 |
| 278 | chr12 | 24948459 | 24950459 | ENSG00000 | BCAT1 | 586 | 33.807246 | -0.208967236 | 0.06255494 | -3.34053915 | 0.00083615 | 0.01822499 | RPE1_active | 0 | 0 |
| 279 | chr4 | 78550519 | 78552519 | ENSG00000 | ANXA3 | 306 | 15.50424803 | -0.2086180336 | 0.07580287 | -2.75211257 | 0.00592121 | 0.07140195 | RPE1_active | 0 | 0 |
| 280 | chr8 | 26513275 | 26515275 | ENSG00000 | DPYSL2 | 1808 | 41.49909038 | -0.2080689003 | 0.05180820 | -4.01613800 | 5.92E-05 | 0.00225837 | RPE1_active | 0 | 0 |
| 281 | chr14 | 52951435 | 52953435 | ENSG00000 | FERMT2 | 10979 | 20.78980556 | -0.2079864804 | 0.07746751 | -2.68482178 | 0.00725684 | 0.08230794 | RPE1_active | 0 | 0 |
| 282 | chr11 | 96388989 | 96390989 | ENSG00000 | JRKL | 8690 | 27.86653165 | -0.2075731482 | 0.06064550 | -3.42272910 | 0.00061995 | 0.01477351 | RPE1_active | 0 | 0 |
| 283 | chr10 | 29735781 | 29737781 | ENSG00000 | SVIL | 6840 | 73.9036538 | -0.207069878 | 0.04227332 | -4.89835776 | 9.66E-07 | 7.92E-05 | RPE1_active | 0 | 0 |
| 284 | chr1 | 64744095 | 64746095 | ENSG00000 | RAVER2 | 55225 | 20.91924788 | -0.2068508003 | 0.07410005 | -2.79150657 | 0.00524632 | 0.06596133 | RPE1_active | 0 | 0 |
| 285 | chr18 | 45966274 | 45968274 | ENSG00000 | EPG5 | 57724 | 30.47515689 | -0.2060083704 | 0.06240772 | -3.30100743 | 0.00096338 | 0.02017517 | RPE1_active | 0 | 0 |
| 286 | chr1 | 178093141 | 178095141 | ENSG00000 | RASAL2 | 9462 | 87.12129803 | -0.204067049 | 0.03900325 | -5.23205193 | 1.68E-07 | 1.84E-05 | RPE1_active | 0 | 0 |
| 287 | chr14 | 64878883 | 64880883 | ENSG00000 | SPTB | 6710 | 45.70778478 | -0.2038098261 | 0.04889968 | -4.16791658 | 3.07E-05 | 0.00132092 | RPE1_active | 0 | 0 |
| 288 | chr20 | 17681295 | 17683295 | ENSG00000 | RRBP1 | 6238 | 22.22552261 | -0.2036930922 | 0.07469099 | -2.72714404 | 0.00638851 | 0.07504420 | RPE1_active | 0 | 0 |
| 289 | chr15 | 70762593 | 70764593 | ENSG00000 | UACA | 55075 | 22.4389194 | -0.2023250077 | 0.07090143 | -2.85360951 | 0.00432256 | 0.05857379 | RPE1_active | 0 | 0 |
| 290 | chr3 | 14401576 | 14403576 | ENSG00000 | SLC6A6 | 6533 | 29.9897352 | -0.2010603227 | 0.06386568 | -3.14817437 | 0.00164293 | 0.02978214 | RPE1_active | 0 | 0 |
| 291 | chr4 | 169009275 | 169011275 | ENSG00000 | CBR4 | 84869 | 34.94344493 | -0.2001379142 | 0.05885181 | -3.40070928 | 0.00067211 | 0.01553018 | RPE1_active | 0 | 0 |
| 292 | chr3 | 12286368 | 12288368 | ENSG00000 | PPARG | 5468 | 37.51101124 | -0.1998041998 | 0.05241976 | -3.81161964 | 0.00013805 | 0.00456586 | RPE1_active | 0 | 0 |
| 293 | chr2 | 189178879 | 189180879 | ENSG00000 | COL5A2 | 1290 | 30.00003006 | -0.1997886391 | 0.06682029 | -2.98993971 | 0.00279032 | 0.04340108 | RPE1_active | 0 | 0 |
| 294 | chr15 | 80778343 | 80780343 | ENSG00000 | CEMIP | 57214 | 62.77114266 | -0.1993484662 | 0.04689359 | -4.25108105 | 2.13E-05 | 0.00098354 | RPE1_active | 0 | 0 |
| 295 | chr3 | 142577826 | 142579826 | ENSG00000 | ATR | 545 | 28.50624798 | -0.1982425062 | 0.06097513 | -3.25120258 | 0.00114917 | 0.02281454 | RPE1_active | 0 | 0 |

| | | | | | | | | | | | | | | | |
|---|---|---|---|---|---|---|---|---|---|---|---|---|---|---|---|
| 296 | chr13 | 44576147 | 44578147 | ENSG00000 | TSC22D1 | 8848 | 30.76951814 | -0.1979485693 | 0.06694039 | -2.95708680 | 0.00310560 | 0.04663517 | RPE1_active | 0 | 0 |
| 297 | chr10 | 15370062 | 15372062 | ENSG00000 | FAM171A1 | 221061 | 47.8335757 | -0.1979330074 | 0.04909998 | -4.03122315 | 5.55E-05 | 0.00214280 | RPE1_active | 0 | 0 |
| 298 | chr1 | 183185238 | 183187238 | ENSG00000 | LAMC2 | 3918 | 18.64899051 | -0.1976870787 | 0.07510715 | -2.63206712 | 0.00848670 | 0.09166606 | RPE1_active | 0 | 0 |
| 299 | chr7 | 7155934 | 7157934 | ENSG00000 | C1GALT1 | 56913 | 21.24939822 | -0.1971395343 | 0.071114644 | -2.77213697 | 0.00556895 | 0.06864620 | RPE1_active | 0 | 0 |
| 300 | chr2 | 201780189 | 201782189 | ENSG00000 | ALS2 | 57679 | 17.9890669 | -0.1966727042 | 0.07114281 | -2.76447728 | 0.00570140 | 0.06969488 | RPE1_active | 0 | 0 |
| 301 | chr9 | 32383603 | 32385603 | ENSG00000 | ACO1 | 48 | 20.46047612 | -0.1961923282 | 0.07044902 | -2.78488370 | 0.00535469 | 0.06664161 | RPE1_active | 0 | 0 |
| 302 | chr3 | 157532619 | 157534619 | ENSG00000 | VEPH1 | 79674 | 68.94440095 | -0.1956854564 | 0.04048193 | -4.83389589 | 1.34E-06 | 0.00010340 | RPE1_active | 0 | 0 |
| 303 | chr9 | 94725701 | 94727701 | ENSG00000 | C9orf3 | 84909 | 101.9175541 | -0.195264532 | 0.03565068 | -5.47716022 | 4.32E-08 | 5.66E-06 | RPE1_active | 0 | 0 |
| 304 | chr4 | 94450857 | 94452857 | ENSG00000 | PDLIM5 | 10611 | 45.69311073 | -0.1946962977 | 0.05322128 | -3.65824133 | 0.00025395 | 0.00749017 | RPE1_active | 0 | 0 |
| 305 | chr9 | 88387419 | 88389419 | ENSG00000 | SPIN1 | 10927 | 20.99906985 | -0.1939754541 | 0.06842992 | -2.83465818 | 0.00458747 | 0.06062635 | RPE1_active | 0 | 0 |
| 306 | chr14 | 25049297 | 25051297 | ENSG00000 | STXBP6 | 29091 | 71.34442119 | -0.1939142477 | 0.04015072 | -4.82965802 | 1.37E-06 | 0.00010502 | RPE1_active | 0 | 0 |
| 307 | chr9 | 70257962 | 70259962 | ENSG00000 | SMC5 | 23137 | 19.09703013 | -0.1921896671 | 0.07220619 | -2.66167832 | 0.00777521 | 0.08646140 | RPE1_active | 0 | 0 |
| 308 | chr5 | 61331273 | 61333273 | ENSG00000 | ZSWIM6 | 57688 | 48.83812656 | -0.1920802945 | 0.04988242 | -3.85066070 | 0.00011779 | 0.00402537 | RPE1_active | 0 | 0 |
| 309 | chr6 | 17986623 | 17988623 | ENSG00000 | KIF13A | 63971 | 59.23704354 | -0.1920754343 | 0.04458431 | -4.30813907 | 1.65E-05 | 0.00078841 | RPE1_active | 0 | 0 |
| 310 | chr12 | 51589266 | 51591266 | ENSG00000 | SCN8A | 6334 | 63.08851828 | -0.1919203067 | 0.04376225 | -4.38552192 | 1.16E-05 | 0.00060865 | RPE1_active | 0 | 0 |
| 311 | chr2 | 147843517 | 147845517 | ENSG00000 | ACVR2A | 92 | 20.26070352 | -0.1916249375 | 0.07404665 | -2.58789452 | 0.00965654 | 0.09909493 | RPE1_active | 0 | 0 |
| 312 | chr22 | 45501238 | 45503238 | ENSG00000 | FBLN1 | 2192 | 37.32562614 | -0.1913632878 | 0.06215320 | -3.07889656 | 0.00207768 | 0.03527317 | RPE1_active | 0 | 0 |
| 313 | chr5 | 73625158 | 73627158 | ENSG00000 | ARHGEF28 | 64283 | 80.30286229 | -0.1906211495 | 0.041191177 | -4.62771786 | 3.70E-06 | 0.00023635 | RPE1_active | 0 | 0 |
| 314 | chr12 | 3076355 | 3078355 | ENSG00000 | TSPAN9 | 10867 | 73.42767989 | -0.189747469 | 0.04893532 | -3.87751531 | 0.00010552 | 0.00368138 | RPE1_active | 0 | 0 |
| 315 | chr4 | 71186286 | 71188286 | ENSG00000 | SLC4A4 | 8671 | 103.6337022 | -0.1889425028 | 0.03770603 | -5.01093475 | 5.42E-07 | 4.96E-05 | RPE1_active | 0 | 0 |
| 316 | chr1 | 93846150 | 93848150 | ENSG00000 | BCAR3 | 8412 | 83.18562321 | -0.1863492943 | 0.03877222 | -4.80625772 | 1.54E-06 | 0.00011370 | RPE1_active | 0 | 0 |
| 317 | chr1 | 204376665 | 204378665 | ENSG00000 | PLEKHA6 | 22874 | 59.81144885 | -0.1862264928 | 0.04524495 | -4.11596138 | 3.86E-05 | 0.00158510 | RPE1_active | 0 | 0 |
| 318 | chr21 | 32411006 | 32413006 | ENSG00000 | EVA1C | 59271 | 30.53895239 | -0.185512906 | 0.06221739 | -2.98168891 | 0.00286663 | 0.04422755 | RPE1_active | 0 | 0 |
| 319 | chr9 | 16869843 | 16871843 | ENSG00000 | BNC2 | 54796 | 116.3131944 | -0.1847271425 | 0.03108361 | -5.94291144 | 2.80E-09 | 4.74E-07 | RPE1_active | 0 | 0 |
| 320 | chr5 | 16935276 | 16937276 | ENSG00000 | MYO10 | 4651 | 73.23383032 | -0.1845963672 | 0.04361215 | -4.23268164 | 2.31E-05 | 0.00104943 | RPE1_active | 0 | 0 |
| 321 | chr8 | 38995869 | 38997869 | ENSG00000 | ADAM9 | 8754 | 22.67751704 | -0.1844219121 | 0.06838380 | -2.69686538 | 0.00699955 | 0.07972810 | RPE1_active | 0 | 0 |
| 322 | chrX | 38005582 | 38007582 | ENSG00000 | SYTL5 | 94122 | 33.80631886 | -0.183974873 | 0.05821860 | -3.16007029 | 0.00157731 | 0.02890870 | RPE1_active | 0 | 0 |
| 323 | chr9 | 34457570 | 34459570 | ENSG00000 | FAM219A | 203259 | 20.95514416 | -0.1838085754 | 0.06748922 | -2.72352476 | 0.00645893 | 0.07556730 | RPE1_active | 0 | 0 |
| 324 | chr11 | 122654675 | 122656675 | ENSG00000 | UBASH3B | 84959 | 47.60871826 | -0.1837490773 | 0.04927927 | -3.72872927 | 0.00019244 | 0.00600769 | RPE1_active | 0 | 0 |
| 325 | chr11 | 74747868 | 74749868 | ENSG00000 | RNF169 | 254225 | 20.26244327 | -0.1836294941 | 0.07105363 | -2.58437875 | 0.00975545 | 0.09980298 | RPE1_active | 0 | 0 |
| 326 | chr6 | 18386350 | 18388350 | ENSG00000 | RNF144B | 255488 | 21.54535997 | -0.1834037406 | 0.07042732 | -2.60415604 | 0.00921008 | 0.09659018 | RPE1_active | 0 | 0 |
| 327 | chr7 | 31513071 | 31515071 | ENSG00000 | CCDC129 | 223075 | 38.61891875 | -0.1833654537 | 0.05355420 | -3.42392242 | 0.00061724 | 0.01475308 | RPE1_active | 0 | 0 |
| 328 | chr20 | 44186578 | 44188578 | ENSG00000 | JPH2 | 57158 | 24.68720457 | -0.1821716526 | 0.06394767 | -2.84876102 | 0.00438898 | 0.05903885 | RPE1_active | 0 | 0 |
| 329 | chr6 | 45327157 | 45329157 | ENSG00000 | RUNX2 | 860 | 91.08912866 | -0.1820220847 | 0.03921123 | -4.64208994 | 3.45E-06 | 0.00022155 | RPE1_active | 0 | 0 |
| 330 | chr9 | 111037458 | 111039458 | ENSG00000 | LPAR1 | 1902 | 40.18719857 | -0.1816793438 | 0.05510630 | -3.29688870 | 0.00097762 | 0.02037755 | RPE1_active | 0 | 0 |
| 331 | chr3 | 58007400 | 58009400 | ENSG00000 | FLNB | 2317 | 45.116377 | -0.1804843231 | 0.05241333 | -3.44348076 | 0.00057427 | 0.01398394 | RPE1_active | 0 | 0 |
| 332 | chr14 | 61528128 | 61530128 | ENSG00000 | RP11-47I22. | NaN | 36.21351878 | -0.1803856332 | 0.05453910 | -3.30745513 | 0.00094147 | 0.01987217 | RPE1_active | 0 | 0 |
| 333 | chr1 | 84297366 | 84299366 | ENSG00000 | SAMD13 | 148418 | 25.94496514 | -0.1788891153 | 0.06749279 | -2.65049186 | 0.00803746 | 0.08814792 | RPE1_active | 0 | 0 |
| 334 | chr15 | 68577969 | 68579969 | ENSG00000 | CORO2B | 10391 | 51.53821541 | -0.1774583527 | 0.05038912 | -3.52175930 | 0.00042869 | 0.01159050 | RPE1_active | 0 | 0 |
| 335 | chr17 | 64262301 | 64264301 | ENSG00000 | TEX2 | 55852 | 32.59983823 | -0.1772218603 | 0.06119054 | -2.89622967 | 0.00377675 | 0.05345473 | RPE1_active | 0 | 0 |
| 336 | chr22 | 46536170 | 46538170 | ENSG00000 | CELSR1 | 9620 | 55.52710349 | -0.1771574626 | 0.05173763 | -3.42415093 | 0.00061672 | 0.01475308 | RPE1_active | 0 | 0 |
| 337 | chr2 | 157874862 | 157876862 | ENSG00000 | ACVR1 | 90 | 37.00249477 | -0.1769356024 | 0.05813071 | -3.04375401 | 0.00233646 | 0.03793129 | RPE1_active | 0 | 0 |
| 338 | chr13 | 51583455 | 51585455 | ENSG00000 | WDFY2 | 115825 | 44.62256544 | -0.1766319313 | 0.05193574 | -3.40097037 | 0.00067147 | 0.01553018 | RPE1_active | 0 | 0 |
| 339 | chr17 | 66301636 | 66303636 | ENSG00000 | PRKCA | 5578 | 146.9317398 | -0.1760744496 | 0.03177294 | -5.54164708 | 3.00E-08 | 4.17E-06 | RPE1_active | 0 | 0 |
| 340 | chr6 | 125118049 | 125120049 | ENSG00000 | TPD52L1 | 7164 | 38.23706671 | -0.1760704781 | 0.05368100 | -3.27993980 | 0.00103829 | 0.02114729 | RPE1_active | 0 | 0 |
| 341 | chr4 | 169270105 | 169272105 | ENSG00000 | SH3RF1 | 57630 | 41.66587339 | -0.1756022256 | 0.05249532 | -3.34510238 | 0.00082252 | 0.018045511 | RPE1_active | 0 | 0 |
| 342 | chr9 | 89177767 | 89179767 | ENSG00000 | SHC3 | 53358 | 51.28273309 | -0.1753262081 | 0.04737999 | -3.70042726 | 0.00021523 | 0.00658072 | RPE1_active | 0 | 0 |
| 343 | chr1 | 53737106 | 53739106 | ENSG00000 | GLIS1 | 148979 | 85.20990731 | -0.1735754802 | 0.04292438 | -4.04375034 | 5.26E-05 | 0.00206714 | RPE1_active | 0 | 0 |
| 344 | chr15 | 33193733 | 33195733 | ENSG00000 | FMN1 | 342184 | 117.2852781 | -0.1731556641 | 0.03251416 | -5.32554544 | 1.01E-07 | 1.18E-05 | RPE1_active | 0 | 0 |
| 345 | chr5 | 52988326 | 52990326 | ENSG00000 | ITGA2 | 3673 | 26.43938443 | -0.1721790595 | 0.06503898 | -2.64732089 | 0.00811323 | 0.08856282 | RPE1_active | 0 | 0 |
| 346 | chr6 | 142300854 | 142302854 | ENSG00000 | GPR126 | 57211 | 33.30903386 | -0.1705347412 | 0.06198970 | -2.75101711 | 0.00594105 | 0.07151209 | RPE1_active | 0 | 0 |
| 347 | chr15 | 62389526 | 62391526 | ENSG00000 | TLN2 | 83660 | 143.5908575 | -0.168849974 | 0.03064880 | -5.50918539 | 3.60E-08 | 4.77E-06 | RPE1_active | 0 | 0 |
| 348 | chr4 | 98657629 | 98659629 | ENSG00000 | TSPAN5 | 10098 | 53.74741269 | -0.167869951 | 0.04323244 | -3.88296249 | 0.00010319 | 0.00361874 | RPE1_active | 0 | 0 |
| 349 | chr8 | 42540926 | 42542926 | ENSG00000 | SLC20A2 | 6575 | 30.80298366 | -0.1673895999 | 0.06430577 | -2.60302606 | 0.00924049 | 0.09683311 | RPE1_active | 0 | 0 |
| 350 | chr2 | 46292667 | 46294667 | ENSG00000 | EPAS1 | 2034 | 31.72280512 | -0.1655177617 | 0.06221456 | -2.66043429 | 0.00780399 | 0.08653044 | RPE1_active | 0 | 0 |
| 351 | chr4 | 41215714 | 41217714 | ENSG00000 | APBB2 | 323 | 105.8457111 | -0.1651327472 | 0.03256540 | -5.07080215 | 3.96E-07 | 3.84E-05 | RPE1_active | 0 | 0 |
| 352 | chr9 | 81688305 | 81690305 | ENSG00000 | TLE1 | 7088 | 29.71684623 | -0.1649180164 | 0.05891519 | -2.79924435 | 0.00512223 | 0.06495456 | RPE1_active | 0 | 0 |
| 353 | chr3 | 105365909 | 105367909 | ENSG00000 | ALCAM | 214 | 41.60376795 | -0.164264757 | 0.05255804 | -3.12539716 | 0.00177565 | 0.03138158 | RPE1_active | 0 | 0 |
| 354 | chr1 | 64469792 | 64471792 | ENSG00000 | CACHD1 | 57685 | 64.78000105 | -0.1636819062 | 0.04463010 | -3.66752269 | 0.000244911 | 0.00723952 | RPE1_active | 0 | 0 |
| 355 | chr21 | 26170128 | 26172128 | ENSG00000 | APP | 351 | 68.99735378 | -0.1636087681 | 0.039452911 | -4.14693780 | 3.37E-05 | 0.00141128 | RPE1_active | 0 | 0 |

| | | | | | | | | | | | | | | | |
|---|---|---|---|---|---|---|---|---|---|---|---|---|---|---|---|
| 356 | chr3 | 8962773 | 8964773 | ENSG00000 | RAD18 | 56852 | 53.0197877 | -0.1630815524 | 0.044243893 | -3.68596747 | 0.00022783 | 0.00687157 | RPE1_active | 0 | 0 |
| 357 | chr1 | 29235516 | 29237516 | ENSG00000 | PTPRU | 10076 | 32.24092112 | -0.1627952062 | 0.060118903 | -2.70788712 | 0.00677130 | 0.07805988 | RPE1_active | 0 | 0 |
| 358 | chr12 | 95790152 | 95792152 | ENSG00000 | NTN4 | 59277 | 34.8852299 | -0.1624860806 | 0.057088179 | -2.84622986 | 0.00442402 | 0.05934678 | RPE1_active | 0 | 0 |
| 359 | chr3 | 190121437 | 190123437 | ENSG00000 | LEPREL1 | 55214 | 39.88062012 | -0.1618505689 | 0.062194946 | -2.60231061 | 0.00925979 | 0.09695933 | RPE1_active | 0 | 0 |
| 360 | chr13 | 75619434 | 75621434 | ENSG00000 | LMO7 | 4008 | 54.6476910 | -0.1613128842 | 0.047132015 | -3.42257555 | 0.00062030 | 0.01477351 | RPE1_active | 0 | 0 |
| 361 | chr1 | 113389749 | 113391749 | ENSG00000 | MAGI3 | 260425 | 65.8919332 | -0.160764461 | 0.047388007 | -3.39251364 | 0.00069254 | 0.01581725 | RPE1_active | 0 | 0 |
| 362 | chr9 | 79570773 | 79572773 | ENSG00000 | TLE4 | 7091 | 41.1739540 | -0.1604776194 | 0.054892189 | -2.92350553 | 0.00346114 | 0.05076213 | RPE1_active | 0 | 0 |
| 363 | chr12 | 19128752 | 19130752 | ENSG00000 | PLEKHA5 | 54477 | 53.5392310 | -0.1603150764 | 0.049695886 | -3.22592249 | 0.00125567 | 0.02442074 | RPE1_active | 0 | 0 |
| 364 | chr9 | 35160992 | 35162992 | ENSG00000 | UNC13B | 10497 | 73.7197871 | -0.1592155932 | 0.040911313 | -3.89172533 | 9.95E-05 | 0.00351423 | RPE1_active | 0 | 0 |
| 365 | chr15 | 82261763 | 82263763 | ENSG00000 | EFTUD1 | 79631 | 30.8729431 | -0.1583359904 | 0.059666807 | -2.65366953 | 0.00796217 | 0.08755772 | RPE1_active | 0 | 0 |
| 366 | chr1 | 17538835 | 17540835 | ENSG00000 | ARHGEF10L | 55160 | 56.2499522 | -0.1569288751 | 0.051785177 | -3.03038211 | 0.00244244 | 0.03928596 | RPE1_active | 0 | 0 |
| 367 | chr1 | 56644301 | 56646301 | ENSG00000 | PPAP2B | 8613 | 45.7340393 | -0.1568107794 | 0.054504059 | -2.87704773 | 0.00401414 | 0.05592600 | RPE1_active | 0 | 0 |
| 368 | chr8 | 19083730 | 19085730 | ENSG00000 | PSD3 | 23362 | 147.257287 | -0.1567330991 | 0.028372267 | -5.52416537 | 3.31E-08 | 4.56E-06 | RPE1_active | 0 | 0 |
| 369 | chr7 | 85185855 | 85187855 | ENSG00000 | SEMA3D | 223117 | 39.13949112 | -0.1562153421 | 0.057018148 | -2.73974771 | 0.00614863 | 0.07307264 | RPE1_active | 0 | 0 |
| 370 | chr2 | 229713558 | 229715558 | ENSG00000 | DNER | 92737 | 101.084016 | -0.1556626691 | 0.035027511 | -4.44401165 | 8.83E-06 | 0.00049361 | RPE1_active | 0 | 0 |
| 371 | chr12 | 32398529 | 32400529 | ENSG00000 | FGD4 | 121512 | 60.30814411 | -0.1553031195 | 0.045383413 | -3.42202373 | 0.00062156 | 0.01477719 | RPE1_active | 0 | 0 |
| 372 | chr15 | 39781830 | 39783830 | ENSG00000 | FSIP1 | 161835 | 47.0044803 | -0.1549177535 | 0.045399086 | -3.41235401 | 0.00064404 | 0.01509662 | RPE1_active | 0 | 0 |
| 373 | chr1 | 196608225 | 196610225 | ENSG00000 | KCNT2 | 343450 | 79.3892305 | -0.1545123005 | 0.043484189 | -3.55329838 | 0.00038043 | 0.01054556 | RPE1_active | 0 | 0 |
| 374 | chr2 | 27888941 | 27890941 | ENSG00000 | BRE | 9577 | 117.827801 | -0.1543485231 | 0.031021928 | -4.97546512 | 6.51E-07 | 5.76E-05 | RPE1_active | 0 | 0 |
| 375 | chr17 | 17971422 | 17973422 | ENSG00000 | TOM1L2 | 146691 | 39.6347256 | -0.153772139 | 0.057422652 | -2.67790031 | 0.00740852 | 0.08309132 | RPE1_active | 0 | 0 |
| 376 | chr6 | 166905451 | 166907451 | ENSG00000 | RPS6KA2 | 6196 | 157.131878 | -0.1535587245 | 0.030220463 | -5.08128295 | 3.75E-07 | 3.66E-05 | RPE1_active | 0 | 0 |
| 377 | chr1 | 185733551 | 185735551 | ENSG00000 | HMCN1 | 83872 | 109.937801 | -0.1533634867 | 0.037503326 | -4.08933020 | 4.33E-05 | 0.00175158 | RPE1_active | 0 | 0 |
| 378 | chr9 | 132078908 | 132080908 | ENSG00000 | MED27 | 9442 | 69.6776993 | -0.1530619135 | 0.045484876 | -3.36511661 | 0.00076511 | 0.01708326 | RPE1_active | 0 | 0 |
| 379 | chr2 | 56183123 | 56185123 | ENSG00000 | CCDC85A | 114800 | 54.8965133 | -0.1530109137 | 0.045971593 | -3.32837956 | 0.00087352 | 0.01891605 | RPE1_active | 0 | 0 |
| 380 | chr7 | 22631925 | 22633925 | ENSG00000 | STEAP1B | 256227 | 60.9261494 | -0.1529230264 | 0.046424294 | -3.29403013 | 0.00098761 | 0.02052189 | RPE1_active | 0 | 0 |
| 381 | chr17 | 15562561 | 15564561 | ENSG00000 | TVP23C-CD | 100533496 | 36.1549150 | -0.1528678405 | 0.055269365 | -2.76586927 | 0.00567712 | 0.06958909 | RPE1_active | 0 | 0 |
| 382 | chr3 | 71582989 | 71584989 | ENSG00000 | FOXP1 | 27086 | 166.997669 | -0.1519577348 | 0.027936063 | -5.43948269 | 5.34E-08 | 6.74E-06 | RPE1_active | 0 | 0 |
| 383 | chr2 | 70553193 | 70555193 | ENSG00000 | TGFA | 7039 | 35.7473681 | -0.1514514436 | 0.057271359 | -2.64445343 | 0.00818229 | 0.08909833 | RPE1_active | 0 | 0 |
| 384 | chr3 | 155023124 | 155025124 | ENSG00000 | MME | 4311 | 37.6743043 | -0.1512702587 | 0.052287172 | -2.89306635 | 0.00381500 | 0.05376825 | RPE1_active | 0 | 0 |
| 385 | chr2 | 9002813 | 9004813 | ENSG00000 | MBOAT2 | 129642 | 42.9708936 | -0.1510875219 | 0.053922214 | -2.80195318 | 0.00507942 | 0.06469608 | RPE1_active | 0 | 0 |
| 386 | chr3 | 185824056 | 185826056 | ENSG00000 | IGF2BP2 | 10644 | 49.8320585 | -0.1509399569 | 0.046959458 | -3.21426099 | 0.00130780 | 0.02510576 | RPE1_active | 0 | 0 |
| 387 | chr1 | 22709839 | 22711839 | ENSG00000 | EPHB2 | 2048 | 78.7108339 | -0.1503600926 | 0.041327651 | -3.6382443 | 0.00027450 | 0.00799048 | RPE1_active | 0 | 0 |
| 388 | chr15 | 44194252 | 44196252 | ENSG00000 | FRMD5 | 84978 | 88.7299547 | -0.1499433932 | 0.039845031 | -3.76316415 | 0.00016777 | 0.00529945 | RPE1_active | 0 | 0 |
| 389 | chr12 | 123149015 | 123151015 | ENSG00000 | PITPNM2 | 57605 | 56.9345609 | -0.1490042339 | 0.053324231 | -2.79430623 | 0.00520111 | 0.06568037 | RPE1_active | 0 | 0 |
| 390 | chr4 | 5892058 | 5894058 | ENSG00000 | CRMP1 | 1400 | 47.8317302 | -0.1485311213 | 0.049503766 | -3.00040042 | 0.00269624 | 0.04238186 | RPE1_active | 0 | 0 |
| 391 | chr2 | 36695690 | 36697690 | ENSG00000 | VIT | 5212 | 39.0088326 | -0.1483059434 | 0.053349956 | -2.77986997 | 0.00543806 | 0.06740075 | RPE1_active | 0 | 0 |
| 392 | chr5 | 52786896 | 52788896 | ENSG00000 | ITGA1 | 3672 | 40.4647664 | -0.1475560951 | 0.052128549 | -2.83061961 | 0.00464579 | 0.06115512 | RPE1_active | 0 | 0 |
| 393 | chr2 | 100819152 | 100821152 | ENSG00000 | NPAS2 | 4862 | 55.8409379 | -0.1474705841 | 0.044005322 | -3.35119881 | 0.00080462 | 0.01771103 | RPE1_active | 0 | 0 |
| 394 | chr3 | 111731497 | 111733497 | ENSG00000 | PHLDB2 | 90102 | 61.8109075 | -0.1469836977 | 0.043675965 | -3.36532225 | 0.00076454 | 0.01708326 | RPE1_active | 0 | 0 |
| 395 | chr4 | 183098293 | 183100293 | ENSG00000 | WWC2 | 80014 | 56.1081820 | -0.1461366913 | 0.051890318 | -2.81626121 | 0.00485861 | 0.06308644 | RPE1_active | 0 | 0 |
| 396 | chr5 | 15499196 | 15501196 | ENSG00000 | FBXL7 | 23194 | 123.579331 | -0.1456896477 | 0.02985964 | -4.8790964 | 1.07E-06 | 8.68E-05 | RPE1_active | 0 | 0 |
| 397 | chr2 | 175004369 | 175006369 | ENSG00000 | CHN1 | 1123 | 48.7873431 | -0.1451060617 | 0.050898898 | -2.85086840 | 0.00436000 | 0.05890331 | RPE1_active | 0 | 0 |
| 398 | chr2 | 168247141 | 168249141 | ENSG00000 | STK39 | 27347 | 83.6724529 | -0.1450735215 | 0.037586809 | -3.85969235 | 0.00011352 | 0.00391857 | RPE1_active | 0 | 0 |
| 399 | chr5 | 39423868 | 39425868 | ENSG00000 | C9 | 735 | 37.4587824 | -0.1444793609 | 0.053322260 | -2.70955058 | 0.00673744 | 0.07787108 | RPE1_active | 0 | 0 |
| 400 | chr2 | 46915157 | 46917157 | ENSG00000 | TTC7A | 57217 | 49.9493268 | -0.1434268591 | 0.050353343 | -2.84840787 | 0.00439385 | 0.05903885 | RPE1_active | 0 | 0 |
| 401 | chr9 | 113875282 | 113877282 | ENSG00000 | ZNF618 | 114991 | 58.8573637 | -0.1433741429 | 0.044261933 | -3.23922005 | 0.00119857 | 0.02341242 | RPE1_active | 0 | 0 |
| 402 | chr12 | 78862993 | 78864993 | ENSG00000 | SYT1 | 6857 | 133.827727 | -0.1433579534 | 0.034672887 | -4.13458367 | 3.56E-05 | 0.00147551 | RPE1_active | 0 | 0 |
| 403 | chr6 | 39791298 | 39793298 | ENSG00000 | DAAM2 | 23500 | 39.6592316 | -0.1427628339 | 0.053209485 | -2.68303353 | 0.00729576 | 0.08237839 | RPE1_active | 0 | 0 |
| 404 | chr7 | 7534853 | 7536853 | ENSG00000 | COL28A1 | 340267 | 48.2092367 | -0.1427558865 | 0.053222089 | -2.68226760 | 0.00731249 | 0.08244913 | RPE1_active | 0 | 0 |
| 405 | chr5 | 135032280 | 135034280 | ENSG00000 | C5orf66 | NaN | 116.283286 | -0.1427473363 | 0.032265053 | -4.42420884 | 9.68E-06 | 0.00052787 | RPE1_active | 0 | 0 |
| 406 | chr1 | 65419652 | 65421652 | ENSG00000 | LEPR | 3953 | 50.9885439 | -0.1426031712 | 0.045527851 | -3.13221832 | 0.00173490 | 0.03098944 | RPE1_active | 0 | 0 |
| 407 | chr15 | 29967865 | 29969865 | ENSG00000 | TJP1 | 7082 | 68.9580488 | -0.1422304007 | 0.041216942 | -3.45077518 | 0.00055897 | 0.01377955 | RPE1_active | 0 | 0 |
| 408 | chr9 | 125746345 | 125748345 | ENSG00000 | PBX3 | 5090 | 44.9423740 | -0.1419451409 | 0.051993685 | -2.73004577 | 0.00633255 | 0.07454558 | RPE1_active | 0 | 0 |
| 409 | chr4 | 84965391 | 84967391 | ENSG00000 | WDFY3 | 23001 | 67.0745834 | -0.1415593628 | 0.042923026 | -3.29798187 | 0.00097382 | 0.02033010 | RPE1_active | 0 | 0 |
| 410 | chr1 | 240013348 | 240015348 | ENSG00000 | FMN2 | 56776 | 112.2580607 | -0.1395555031 | 0.039793028 | -3.50703397 | 0.00045313 | 0.01201247 | RPE1_active | 0 | 0 |
| 411 | chr4 | 82797857 | 82799857 | ENSG00000 | SCD5 | 79966 | 51.7804999 | -0.1391689639 | 0.048669318 | -2.85948041 | 0.00424335 | 0.05791162 | RPE1_active | 0 | 0 |
| 412 | chr2 | 109128348 | 109130348 | ENSG00000 | SH3RF3 | 344558 | 133.064754 | -0.1379250092 | 0.035070104 | -3.93283712 | 8.39E-05 | 0.00307299 | RPE1_active | 0 | 0 |
| 413 | chr3 | 89106524 | 89108524 | ENSG00000 | EPHA3 | 2042 | 77.5329942 | -0.137884034 | 0.043209405 | -3.19106529 | 0.00141749 | 0.02671243 | RPE1_active | 0 | 0 |
| 414 | chr4 | 37452941 | 37454941 | ENSG00000 | C4orf19 | 55286 | 52.0263849 | -0.1376794965 | 0.045583392 | -3.02038724 | 0.00252451 | 0.04034697 | RPE1_active | 0 | 0 |
| 415 | chr14 | 54566097 | 54568097 | ENSG00000 | SAMD4A | 23034 | 64.4821963 | -0.1368185798 | 0.043054345 | -3.17781112 | 0.00148391 | 0.02757520 | RPE1_active | 0 | 0 |

| | | | | | | | | | | | | | | | |
|---|---|---|---|---|---|---|---|---|---|---|---|---|---|---|---|
| 416 | chr2 | 184597366 | 184599366 | ENSG00000 | ZNF804A | 91752 | 71.53200122 | -0.1360932458 | 0.04548775 | -2.99186417 | 0.00277279 | 0.04320322 | RPE1_active | 0 | 0 |
| 417 | chr1 | 66532383 | 66534383 | ENSG00000 | SGIP1 | 84251 | 56.88953257 | -0.1354506475 | 0.04657901 | -2.90797595 | 0.00363776 | 0.05220639 | RPE1_active | 0 | 0 |
| 418 | chrX | 44342672 | 44344672 | ENSG00000 | EFHC2 | 80258 | 55.04409761 | -0.1347338729 | 0.04872646 | -2.76510639 | 0.00569042 | 0.06962430 | RPE1_active | 0 | 0 |
| 419 | chr3 | 4466281 | 4468281 | ENSG00000 | SUMF1 | 285362 | 209.0742187 | -0.1342429602 | 0.02592592 | -5.1779427 | 2.24E-07 | 2.36E-05 | RPE1_active | 0 | 0 |
| 420 | chr6 | 56953628 | 56955628 | ENSG00000 | DST | 667 | 110.1030112 | -0.1338613579 | 0.03645525 | -3.67193606 | 0.00024071 | 0.00716315 | RPE1_active | 0 | 0 |
| 421 | chrX | 106999244 | 107001244 | ENSG00000 | MORC4 | 79710 | 53.56441333 | -0.1331756816 | 0.04644805 | -2.86719620 | 0.00414126 | 0.05732362 | RPE1_active | 0 | 0 |
| 422 | chr1 | 215004775 | 215006775 | ENSG00000 | KCNK2 | 3776 | 55.26608444 | -0.1331203638 | 0.05063126 | -2.62921251 | 0.00855828 | 0.09218682 | RPE1_active | 0 | 0 |
| 423 | chr2 | 209423058 | 209425058 | ENSG00000 | MAP2 | 4133 | 71.09992483 | -0.1324245851 | 0.04019823 | -3.29428821 | 0.00098671 | 0.02052189 | RPE1_active | 0 | 0 |
| 424 | chr4 | 150077274 | 150079274 | ENSG00000 | DCLK2 | 166614 | 46.73109816 | -0.1317701274 | 0.05069188 | -2.59943254 | 0.00933780 | 0.09739452 | RPE1_active | 0 | 0 |
| 425 | chr7 | 42236870 | 42238870 | ENSG00000 | GLI3 | 2737 | 83.17606826 | -0.1295826441 | 0.03911993 | -3.31244563 | 0.00092484 | 0.01964515 | RPE1_active | 0 | 0 |
| 426 | chr9 | 38067687 | 38069687 | ENSG00000 | RP11-613M1 | NaN | 140.326116 | -0.129525203 | 0.03102921 | -4.17429847 | 2.99E-05 | 0.00129245 | RPE1_active | 0 | 0 |
| 427 | chr4 | 94756968 | 94758968 | ENSG00000 | BMPR1B | 658 | 96.05165779 | -0.1293264185 | 0.03724350 | -3.47245603 | 0.00051571 | 0.01314986 | RPE1_active | 0 | 0 |
| 428 | chr18 | 6728718 | 6730718 | ENSG00000 | ARHGAP28 | 79822 | 49.12737943 | -0.1291572281 | 0.04899695 | -2.63602537 | 0.00838834 | 0.09085254 | RPE1_active | 0 | 0 |
| 429 | chr21 | 33641400 | 33643400 | ENSG00000 | ITSN1 | 6453 | 66.67868306 | -0.1288656561 | 0.04322565 | -2.98123112 | 0.00287092 | 0.04424263 | RPE1_active | 0 | 0 |
| 430 | chr4 | 88520048 | 88522048 | ENSG00000 | HERC3 | 8916 | 48.02826212 | -0.1284823795 | 0.04896294 | -2.62407384 | 0.00868849 | 0.09301840 | RPE1_active | 0 | 0 |
| 431 | chr5 | 168290651 | 168292651 | ENSG00000 | WWC1 | 23286 | 58.87437245 | -0.1283928304 | 0.04791164 | -2.67978348 | 0.00736697 | 0.08271446 | RPE1_active | 0 | 0 |
| 432 | chr7 | 11831198 | 11833198 | ENSG00000 | THSD7A | 221981 | 110.4642105 | -0.1280876937 | 0.03668150 | -3.49188720 | 0.00047962 | 0.01249164 | RPE1_active | 0 | 0 |
| 433 | chr1 | 227317474 | 227319474 | ENSG00000 | CDC42BPA | 8476 | 69.65104462 | -0.1278723585 | 0.04315462 | -2.96312089 | 0.00304536 | 0.04614937 | RPE1_active | 0 | 0 |
| 434 | chr2 | 199470266 | 199472266 | ENSG00000 | SATB2 | 23314 | 48.97085717 | -0.1276555072 | 0.04394296 | -2.90502734 | 0.00367221 | 0.05252920 | RPE1_active | 0 | 0 |
| 435 | chr12 | 45214987 | 45216987 | ENSG00000 | ANO6 | 196527 | 57.66385135 | -0.1275929446 | 0.04262514 | -2.99337298 | 0.00275912 | 0.04311654 | RPE1_active | 0 | 0 |
| 436 | chr7 | 34657239 | 34659239 | ENSG00000 | NPSR1 | 387129 | 65.28824293 | -0.1271909575 | 0.04371885 | -2.90929283 | 0.00362247 | 0.05215503 | RPE1_active | 0 | 0 |
| 437 | chr5 | 66595361 | 66597361 | ENSG00000 | MAST4 | 375449 | 165.4726295 | -0.126718366 | 0.02783408 | -4.55263320 | 5.30E-06 | 0.00031886 | RPE1_active | 0 | 0 |
| 438 | chr2 | 153870913 | 153872913 | ENSG00000 | GALNT13 | 114805 | 130.6923729 | -0.1264148141 | 0.03969726 | -3.18447181 | 0.00145018 | 0.02709917 | RPE1_active | 0 | 0 |
| 439 | chr15 | 37100299 | 37102299 | ENSG00000 | MEIS2 | 4212 | 51.17812921 | -0.126049884 | 0.04727251 | -2.6664518 | 0.00766565 | 0.08542191 | RPE1_active | 0 | 0 |
| 440 | chr9 | 68327308 | 68329308 | ENSG00000 | PGM5 | 5239 | 55.25676408 | -0.125332042 | 0.04717245 | -2.65689037 | 0.00788650 | 0.08703830 | RPE1_active | 0 | 0 |
| 441 | chr5 | 9545075 | 9547075 | ENSG00000 | SEMA5A | 9037 | 149.8545008 | -0.1249369767 | 0.03089404 | -4.04404751 | 5.25E-05 | 0.00206714 | RPE1_active | 0 | 0 |
| 442 | chr1 | 97920049 | 97922049 | ENSG00000 | DPYD | 1806 | 185.5574153 | -0.1247022149 | 0.02969508 | -4.19942277 | 2.68E-05 | 0.00117998 | RPE1_active | 0 | 0 |
| 443 | chr13 | 98141562 | 98143562 | ENSG00000 | FARP1 | 10160 | 94.61992548 | -0.1244249769 | 0.03701825 | -3.36117919 | 0.00077610 | 0.01723355 | RPE1_active | 0 | 0 |
| 444 | chr11 | 8909951 | 8911951 | ENSG00000 | ST5 | 6764 | 65.2865366 | -0.1243423099 | 0.04284752 | -2.90197152 | 0.00370822 | 0.05270804 | RPE1_active | 0 | 0 |
| 445 | chr5 | 84383793 | 84385793 | ENSG00000 | EDIL3 | 10085 | 97.10370287 | -0.1239573716 | 0.03904315 | -3.17488123 | 0.00149897 | 0.02777787 | RPE1_active | 0 | 0 |
| 446 | chr7 | 33903911 | 33905911 | ENSG00000 | BMPER | 168667 | 74.63207515 | -0.1228225021 | 0.04173089 | -2.94320284 | 0.00324835 | 0.04827727 | RPE1_active | 0 | 0 |
| 447 | chr22 | 28882592 | 28884592 | ENSG00000 | ZNRF3 | 84133 | 55.64217189 | -0.1226801715 | 0.04238950 | -2.89411685 | 0.00380226 | 0.05370198 | RPE1_active | 0 | 0 |
| 448 | chr2 | 32945972 | 32947972 | ENSG00000 | LTBP1 | 4052 | 128.2920804 | -0.1217685919 | 0.03211630 | -3.79148833 | 0.00014974 | 0.00484484 | RPE1_active | 0 | 0 |
| 449 | chr1 | 208243320 | 208245320 | ENSG00000 | PLXNA2 | 5362 | 82.54916239 | -0.1216875769 | 0.03855513 | -3.15619663 | 0.00159841 | 0.02921527 | RPE1_active | 0 | 0 |
| 450 | chr22 | 24951730 | 24953730 | ENSG00000 | KIAA1671 | 85379 | 77.44111912 | -0.1214682324 | 0.04046242 | -3.00200050 | 0.00268211 | 0.04220938 | RPE1_active | 0 | 0 |
| 451 | chrX | 17374420 | 17376420 | ENSG00000 | NHS | 4810 | 119.4127028 | -0.1211922815 | 0.03079295 | -3.93571487 | 8.29E-05 | 0.00304474 | RPE1_active | 0 | 0 |
| 452 | chr3 | 64687000 | 64689000 | ENSG00000 | ADAMTS9 | 56999 | 54.42247489 | -0.1179363997 | 0.04550622 | -2.59165402 | 0.00955157 | 0.09863311 | RPE1_active | 0 | 0 |
| 453 | chr7 | 103073843 | 103075843 | ENSG00000 | FBXL13 | 222235 | 68.84707271 | -0.1162148967 | 0.03895232 | -2.98351611 | 0.00284957 | 0.04406609 | RPE1_active | 0 | 0 |
| 454 | chr17 | 60676453 | 60678453 | ENSG00000 | BCAS3 | 54828 | 178.2480911 | -0.1159729943 | 0.02792629 | -4.15282420 | 3.28E-05 | 0.00137978 | RPE1_active | 0 | 0 |
| 455 | chr2 | 54455285 | 54457285 | ENSG00000 | SPTBN1 | 6711 | 57.19231065 | -0.1158774918 | 0.04393568 | -2.63743429 | 0.00835358 | 0.09064790 | RPE1_active | 0 | 0 |
| 456 | chr2 | 133567463 | 133569463 | ENSG00000 | NCKAP5 | 344148 | 258.9253828 | -0.1145749672 | 0.02477759 | -4.62413550 | 3.76E-06 | 0.00023819 | RPE1_active | 0 | 0 |
| 457 | chr5 | 107669895 | 107671895 | ENSG00000 | EFNA5 | 1946 | 76.15701281 | -0.1139762701 | 0.03576259 | -3.18702464 | 0.00143744 | 0.02693647 | RPE1_active | 0 | 0 |
| 458 | chr1 | 77281051 | 77283051 | ENSG00000 | AK5 | 26289 | 76.23545432 | -0.1135671452 | 0.04005293 | -2.83542595 | 0.00457646 | 0.06062104 | RPE1_active | 0 | 0 |
| 459 | chr17 | 66191084 | 66193084 | ENSG00000 | CEP112 | 201134 | 136.4275778 | -0.1120742985 | 0.03147538 | -3.56069698 | 0.00036987 | 0.01029553 | RPE1_active | 0 | 0 |
| 460 | chr12 | 18246614 | 18248614 | ENSG00000 | PIK3C2G | 5288 | 93.85831885 | -0.1120336929 | 0.03830967 | -2.92442261 | 0.00345095 | 0.05072415 | RPE1_active | 0 | 0 |
| 461 | chr21 | 33930607 | 33932607 | ENSG00000 | AP000304.1 | NaN | 86.13446582 | -0.1117582242 | 0.03629057 | -3.07953876 | 0.00207321 | 0.03524199 | RPE1_active | 0 | 0 |
| 462 | chr7 | 84491724 | 84493724 | ENSG00000 | SEMA3A | 10371 | 103.0812874 | -0.1112274273 | 0.03882731 | -2.86466942 | 0.00417444 | 0.05744057 | RPE1_active | 0 | 0 |
| 463 | chr2 | 124024287 | 124026287 | ENSG00000 | CNTNAP5 | 129684 | 251.6972886 | -0.1103890026 | 0.02909303 | -3.79434446 | 0.00014803 | 0.00482394 | RPE1_active | 0 | 0 |
| 464 | chr1 | 65791514 | 65793514 | ENSG00000 | PDE4B | 5142 | 157.3496873 | -0.1103547755 | 0.03212331 | -3.43534860 | 0.00059179 | 0.01422110 | RPE1_active | 0 | 0 |
| 465 | chr3 | 29279982 | 29281982 | ENSG00000 | RBMS3 | 27303 | 173.2368884 | -0.1102715529 | 0.02634878 | -4.18507127 | 2.85E-05 | 0.00124474 | RPE1_active | 0 | 0 |
| 466 | chr7 | 82442798 | 82444798 | ENSG00000 | CACNA2D1 | 781 | 103.3329543 | -0.1096026449 | 0.03615354 | -3.03158737 | 0.00243271 | 0.03920808 | RPE1_active | 0 | 0 |
| 467 | chr3 | 169662618 | 169664618 | ENSG00000 | MECOM | 2122 | 150.5710226 | -0.1092162697 | 0.03193317 | -3.42015077 | 0.00062586 | 0.01482654 | RPE1_active | 0 | 0 |
| 468 | chr13 | 67229445 | 67231445 | ENSG00000 | PCDH9 | 5101 | 195.190151 | -0.10893867 | 0.03457546 | -3.15075098 | 0.00162851 | 0.02968425 | RPE1_active | 0 | 0 |
| 469 | chr4 | 77156151 | 77158151 | ENSG00000 | CCNG2 | 901 | 109.5358463 | -0.1086297079 | 0.03713789 | -2.92503696 | 0.00344415 | 0.05067987 | RPE1_active | 0 | 0 |
| 470 | chr22 | 28678865 | 28680865 | ENSG00000 | TTC28 | 23331 | 190.2635785 | -0.1082487921 | 0.02558393 | -4.23112431 | 2.33E-05 | 0.00104958 | RPE1_active | 0 | 0 |
| 471 | chr3 | 158961235 | 158963235 | ENSG00000 | IQCJ-SCHIP | NaN | 257.9784844 | -0.1079232 | 0.02447452 | -4.40961299 | 1.04E-05 | 0.00056244 | RPE1_active | 0 | 0 |
| 472 | chr7 | 32298329 | 32300329 | ENSG00000 | PDE1C | 5137 | 158.1076828 | -0.1077712263 | 0.02992369 | -3.60153482 | 0.00031634 | 0.00903135 | RPE1_active | 0 | 0 |
| 473 | chr2 | 162837730 | 162839730 | ENSG00000 | KCNH7 | 90134 | 103.1276569 | -0.1071955368 | 0.04091197 | -2.62015093 | 0.00878908 | 0.09371985 | RPE1_active | 0 | 0 |
| 474 | chr3 | 131025850 | 131027850 | ENSG00000 | NEK11 | 79858 | 90.02578823 | -0.1071202045 | 0.03868756 | -2.76885388 | 0.00562538 | 0.06914515 | RPE1_active | 0 | 0 |
| 475 | chr1 | 211133115 | 211135115 | ENSG00000 | KCNH1 | 3756 | 144.296049 | -0.1064934092 | 0.03304044 | -3.22312226 | 0.00126801 | 0.02461269 | RPE1_active | 0 | 0 |

| | | | | | | | | | | | | | | | |
|---|---|---|---|---|---|---|---|---|---|---|---|---|---|---|---|
| 476 | chr7 | 18085949 | 18087949 | ENSG00000 | HDAC9 | 9734 | 214.2384857 | -0.1062253952 | 0.025787028 | -4.11933441 | 3.80E-05 | 0.001566900 | RPE1_active | 0 | 0 |
| 477 | chr4 | 46992723 | 46994723 | ENSG00000 | GABRB1 | 2560 | 108.2735852 | -0.104833738 | 0.036589004 | -2.86517055 | 0.004167846 | 0.057408869 | RPE1_active | 0 | 0 |
| 478 | chr7 | 79452574 | 79454574 | ENSG00000 | MAGI2 | 9863 | 363.7979985 | -0.1033381576 | 0.027389156 | -3.77295877 | 0.000161322 | 0.005168911 | RPE1_active | 0 | 0 |
| 479 | chr5 | 128658185 | 128660185 | ENSG00000 | FBN2 | 2201 | 101.7646501 | -0.1022447193 | 0.038791580 | -2.63574512 | 0.008395276 | 0.090852547 | RPE1_active | 0 | 0 |
| 480 | chr8 | 132865958 | 132867958 | ENSG00000 | TG | 7038 | 98.31848876 | -0.1012436208 | 0.038512825 | -2.62882870 | 0.008567950 | 0.092186829 | RPE1_active | 0 | 0 |
| 481 | chrX | 37348330 | 37350330 | ENSG00000 | TM4SF2 | NaN | 380.6665658 | -0.1002043938 | 0.023478013 | -4.26800987 | 1.97E-05 | 0.000921369 | RPE1_active | 0 | 0 |
| 482 | chr10 | 35246025 | 35248025 | ENSG00000 | CCNY | 219771 | 90.28363324 | -0.1001465222 | 0.038601399 | -2.59437542 | 0.009476294 | 0.098378223 | RPE1_active | 0 | 0 |
| 483 | chr5 | 113487830 | 113489830 | ENSG00000 | MCC | 4163 | 144.2667301 | -0.09892558724 | 0.031315603 | -3.15898713 | 0.001583184 | 0.028976618 | RPE1_active | 0 | 0 |
| 484 | chr8 | 103499748 | 103501748 | ENSG00000 | RIMS2 | 9699 | 164.3979564 | -0.09870613465 | 0.034655261 | -2.84822943 | 0.004396321 | 0.059038852 | RPE1_active | 0 | 0 |
| 485 | chr2 | 136764545 | 136766545 | ENSG00000 | THSD7B | 80731 | 244.7780101 | -0.09772010878 | 0.028170489 | -3.46888212 | 0.000522628 | 0.013225080 | RPE1_active | 0 | 0 |
| 486 | chr7 | 48170458 | 48172458 | ENSG00000 | ABCA13 | 154664 | 130.8675158 | -0.09632398041 | 0.034110506 | -2.82388010 | 0.004744612 | 0.061907005 | RPE1_active | 0 | 0 |
| 487 | chr6 | 128519674 | 128521674 | ENSG00000 | PTPRK | 5796 | 121.2063706 | -0.09490170399 | 0.034122751 | -2.78118558 | 0.005416076 | 0.067252973 | RPE1_active | 0 | 0 |
| 488 | chr4 | 89110398 | 89112398 | ENSG00000 | FAM13A | 10144 | 99.16042655 | -0.09478326095 | 0.035017078 | -2.70677236 | 0.006794082 | 0.078187543 | RPE1_active | 0 | 0 |
| 489 | chr2 | 17876847 | 17878847 | ENSG00000 | KCNS3 | 3790 | 141.5931506 | -0.09422868921 | 0.032878490 | -2.86596762 | 0.004157367 | 0.057323621 | RPE1_active | 0 | 0 |
| 490 | chr3 | 173395284 | 173397284 | ENSG00000 | NLGN1 | 22871 | 205.5245927 | -0.09371189201 | 0.032604225 | -2.87422536 | 0.004050198 | 0.056310824 | RPE1_active | 0 | 0 |
| 491 | chr3 | 24494282 | 24496282 | ENSG00000 | THRB | 7068 | 106.2877872 | -0.09153056768 | 0.035063468 | -2.61042540 | 0.009042969 | 0.095663587 | RPE1_active | 0 | 0 |
| 492 | chr7 | 40133977 | 40135977 | ENSG00000 | SUGCT | 79783 | 193.4661082 | -0.09134251619 | 0.029301385 | -3.11734458 | 0.001824881 | 0.031955753 | RPE1_active | 0 | 0 |
| 493 | chr6 | 39724405 | 39726405 | ENSG00000 | KIF6 | 221458 | 114.3988289 | -0.08899084834 | 0.033237562 | -2.67741797 | 0.007419201 | 0.083091325 | RPE1_active | 0 | 0 |
| 494 | chr16 | 10181754 | 10183754 | ENSG00000 | GRIN2A | 2903 | 135.675611 | -0.08766437177 | 0.027546859 | -3.18237260 | 0.001460737 | 0.027182333 | RPE1_active | 0 | 0 |
| 495 | chr21 | 36003667 | 36005667 | ENSG00000 | RUNX1 | 861 | 369.9121956 | -0.08716363355 | 0.019952890 | -4.36847151 | 1.25E-05 | 0.000638059 | RPE1_active | 0 | 0 |
| 496 | chr13 | 34941287 | 34943287 | ENSG00000 | NBEA | 26960 | 156.5892054 | -0.08544050731 | 0.029970074 | -2.85086073 | 0.004360106 | 0.058903315 | RPE1_active | 0 | 0 |
| 497 | chr2 | 72825041 | 72827041 | ENSG00000 | EXOC6B | 23233 | 157.5989245 | -0.08227654534 | 0.030042705 | -2.73865300 | 0.006169144 | 0.073072644 | RPE1_active | 0 | 0 |
| 498 | chr7 | 111561517 | 111563517 | ENSG00000 | IMMP2L | 83943 | 213.3266776 | -0.08100882929 | 0.030761130 | -2.63348020 | 0.008451476 | 0.091359362 | RPE1_active | 0 | 0 |
| 499 | chr4 | 78056570 | 78058570 | ENSG00000 | FRAS1 | 80144 | 146.3049601 | -0.07691281034 | 0.028688136 | -2.68099711 | 0.007340314 | 0.082623375 | RPE1_active | 0 | 0 |
| 500 | chr5 | 169300129 | 169302129 | ENSG00000 | SLIT3 | 6586 | 228.9064127 | -0.06788532068 | 0.024703017 | -2.74805779 | 0.005994944 | 0.071897556 | RPE1_active | 0 | 0 |
| 501 | chr18 | 48409752 | 48411752 | ENSG00000 | ZBTB7C | 201501 | 141.334479 | 0.090619469 | 0.034854040 | 2.599970249 | 0.009323184 | 0.097336704 | HG01573_active | 0 | 0 |
| 502 | chr7 | 37448249 | 37450249 | ENSG00000 | ELMO1 | 9844 | 189.8333334 | 0.09289270118 | 0.026106902 | 3.558166335 | 0.000373452 | 0.010373602 | HG01573_active | 0 | 0 |
| 503 | chr18 | 13216498 | 13218498 | ENSG00000 | LDLRAD4 | 753 | 149.4013536 | 0.09617183102 | 0.030540455 | 3.148997894 | 0.001638313 | 0.029751253 | HG01573_active | 0 | 0 |
| 504 | chr1 | 174158410 | 174160410 | ENSG00000 | RABGAP1L | 9910 | 195.4872693 | 0.09898139374 | 0.029217453 | 3.387748806 | 0.000704687 | 0.016012470 | HG01573_active | 0 | 0 |
| 505 | chr5 | 169636247 | 169638247 | ENSG00000 | DOCK2 | 1794 | 149.4353943 | 0.09923367366 | 0.032407186 | 3.06208856 | 0.002197984 | 0.036655723 | HG01573_active | 0 | 0 |
| 506 | chr16 | 11975737 | 11977737 | ENSG00000 | SNX29 | 92017 | 200.5030188 | 0.1006314388 | 0.030471637 | 3.302462486 | 0.000958399 | 0.020110590 | HG01573_active | 0 | 0 |
| 507 | chr2 | 135530236 | 135532236 | ENSG00000 | ZRANB3 | 84083 | 96.34351462 | 0.1015773905 | 0.035808254 | 2.836703187 | 0.004558196 | 0.060485947 | HG01573_active | 0 | 0 |
| 508 | chr15 | 85379571 | 85381571 | ENSG00000 | AKAP13 | 11214 | 92.26959711 | 0.1023604797 | 0.037581707 | 2.723678273 | 0.006455936 | 0.075567304 | HG01573_active | 0 | 0 |
| 509 | chr11 | 75814167 | 75816167 | ENSG00000 | UVRAG | 7405 | 80.67835884 | 0.1043667263 | 0.040023170 | 2.607657608 | 0.009116409 | 0.096066899 | HG01573_active | 0 | 0 |
| 510 | chr18 | 79068285 | 79070285 | ENSG00000 | ATP9B | 374868 | 82.81451313 | 0.1067437576 | 0.038240071 | 2.791411033 | 0.005247878 | 0.065961336 | HG01573_active | 0 | 0 |
| 511 | chr9 | 123929152 | 123931152 | ENSG00000 | DENND1A | 57706 | 167.5097174 | 0.1095857427 | 0.028714596 | 3.816377566 | 0.000135425 | 0.004489867 | HG01573_active | 0 | 0 |
| 512 | chr8 | 22999000 | 23001000 | ENSG00000 | PEBP4 | 157310 | 99.64529081 | 0.1129839463 | 0.038681149 | 2.920904569 | 0.003490167 | 0.051019828 | HG01573_active | 0 | 0 |
| 513 | chr17 | 80543819 | 80545819 | ENSG00000 | RPTOR | 57521 | 131.4611523 | 0.1148144771 | 0.035376407 | 3.245509769 | 0.001172405 | 0.023121972 | HG01573_active | 0 | 0 |
| 514 | chr2 | 42493569 | 42495569 | ENSG00000 | MTA3 | 57504 | 66.76607445 | 0.1199046469 | 0.040721304 | 2.944518807 | 0.003234572 | 0.048126072 | HG01573_active | 0 | 0 |
| 515 | chr6 | 157380133 | 157382133 | ENSG00000 | ZDHHC14 | 79683 | 93.8669911 | 0.1248827307 | 0.034812446 | 3.587301215 | 0.000334118 | 0.009478034 | HG01573_active | 0 | 0 |
| 516 | chr9 | 68704414 | 68706414 | ENSG00000 | PIP5K1B | 8395 | 90.82526068 | 0.1260985696 | 0.037040864 | 3.404309557 | 0.000663315 | 0.015466939 | HG01573_active | 0 | 0 |
| 517 | chr20 | 18566453 | 18568453 | ENSG00000 | DTD1 | 92675 | 59.82929929 | 0.1277573477 | 0.048443696 | 2.637233698 | 0.008358522 | 0.090647905 | HG01573_active | 0 | 0 |
| 518 | chr17 | 67376281 | 67378281 | ENSG00000 | PITPNC1 | 26207 | 93.50251569 | 0.1278767972 | 0.038973297 | 3.281138736 | 0.001033888 | 0.021110451 | HG01573_active | 0 | 0 |
| 519 | chr5 | 14870778 | 14872778 | ENSG00000 | ANKH | 56172 | 53.70759012 | 0.128446812 | 0.047174631 | 2.722794183 | 0.006473237 | 0.075668348 | HG01573_active | 0 | 0 |
| 520 | chr13 | 28494128 | 28496128 | ENSG00000 | FLT1 | 2321 | 59.96460536 | 0.1291871483 | 0.047804831 | 2.70238688 | 0.006884360 | 0.078886742 | HG01573_active | 0 | 0 |
| 521 | chr17 | 32876177 | 32878177 | ENSG00000 | MYO1D | 4642 | 103.9856201 | 0.1298202212 | 0.037325091 | 3.478095212 | 0.000504990 | 0.012950543 | HG01573_active | 0 | 0 |
| 522 | chr13 | 112689329 | 112691329 | ENSG00000 | ATP11A | 23250 | 66.54689348 | 0.1321644437 | 0.050880661 | 2.597537849 | 0.009389477 | 0.097780829 | HG01573_active | 0 | 0 |
| 523 | chr8 | 140457579 | 140459579 | ENSG00000 | TRAPPC9 | 83696 | 274.2689533 | 0.1331316097 | 0.028439696 | 4.681189468 | 2.85E-06 | 0.000189590 | HG01573_active | 0 | 0 |
| 524 | chr10 | 22713555 | 22715555 | ENSG00000 | PIP4K2A | 5305 | 48.05430754 | 0.1334680186 | 0.047051033 | 2.836664986 | 0.004558741 | 0.060485947 | HG01573_active | 0 | 0 |
| 525 | chr2 | 239400654 | 239402654 | ENSG00000 | HDAC4 | 9759 | 123.0074559 | 0.134590389 | 0.040625350 | 3.312965578 | 0.000923123 | 0.019645151 | HG01573_active | 0 | 0 |
| 526 | chr7 | 156892230 | 156894230 | ENSG00000 | LMBR1 | 64327 | 56.63711138 | 0.134750797 | 0.048079179 | 2.802685039 | 0.005067913 | 0.064611061 | HG01573_active | 0 | 0 |
| 527 | chr7 | 75737962 | 75739962 | ENSG00000 | HIP1 | 3092 | 59.0473064 | 0.1352027461 | 0.047048032 | 2.873717351 | 0.004056718 | 0.056342842 | HG01573_active | 0 | 0 |
| 528 | chr1 | 169661007 | 169663007 | ENSG00000 | C1orf112 | 55732 | 55.47867375 | 0.1352190868 | 0.045577680 | 2.966783007 | 0.003009332 | 0.045750238 | HG01573_active | 0 | 0 |
| 529 | chr22 | 25563849 | 25565849 | ENSG00000 | ADRBK2 | 157 | 44.80581306 | 0.1361577843 | 0.050797868 | 2.680383799 | 0.007353779 | 0.082635696 | HG01573_active | 0 | 0 |
| 530 | chr9 | 37119539 | 37121539 | ENSG00000 | ZCCHC7 | 84186 | 46.60105979 | 0.1362074902 | 0.049822632 | 2.733847713 | 0.006259898 | 0.073885609 | HG01573_active | 0 | 0 |
| 531 | chr8 | 60677778 | 60679778 | ENSG00000 | CHD7 | 55636 | 53.25209662 | 0.1362763382 | 0.046324597 | 2.941770578 | 0.003263416 | 0.048447223 | HG01573_active | 0 | 0 |
| 532 | chr9 | 2014219 | 2016219 | ENSG00000 | SMARCA2 | 6595 | 49.37619929 | 0.1368755589 | 0.052324889 | 2.615878622 | 0.008899818 | 0.094373396 | HG01573_active | 0 | 0 |
| 533 | chr14 | 92512774 | 92514774 | ENSG00000 | RIN3 | 79890 | 56.57692647 | 0.1375588804 | 0.048123999 | 2.85842576 | 0.004257487 | 0.058045190 | HG01573_active | 0 | 0 |
| 534 | chr9 | 125142506 | 125144506 | ENSG00000 | SCAI | 286205 | 47.79187078 | 0.1390514276 | 0.048703834 | 2.855040627 | 0.004303132 | 0.058429016 | HG01573_active | 0 | 0 |
| 535 | chr2 | 110731573 | 110733573 | ENSG00000 | ACOXL | 55289 | 123.5010306 | 0.1399098567 | 0.034276089 | 4.081849988 | 4.47E-05 | 0.001798045 | HG01573_active | 0 | 0 |

| | | | | | | | | | | | | | | | | |
|---|---|---|---|---|---|---|---|---|---|---|---|---|---|---|---|---|
| 536 | chr2 | 98443854 | 98445854 | ENSG00000 | INPP4A | 3631 | 49.94184445 | | 0.1406737169 | 0.04655849 | 3.02144025 | 0.00251575 | 0.04025505 | HG01573_active | 0 | 0 |
| 537 | chr13 | 111113559 | 111115559 | ENSG00000 | ARHGEF7 | 8874 | 62.1252585 | | 0.1411934787 | 0.04685832 | 3.01319969 | 0.00258508 | 0.04092340 | HG01573_active | 0 | 0 |
| 538 | chr14 | 91059333 | 91061333 | ENSG00000 | C14orf159 | 80017 | 50.6819884 | | 0.1421797272 | 0.04814805 | 2.95296934 | 0.00314733 | 0.04703747 | HG01573_active | 0 | 0 |
| 539 | chr4 | 112284509 | 112286509 | ENSG00000 | ALPK1 | 80216 | 45.43242417 | | 0.1432494222 | 0.04917822 | 2.91286291 | 0.00358131 | 0.05178570 | HG01573_active | 0 | 0 |
| 540 | chr1 | 167219829 | 167221829 | ENSG00000 | POU2F1 | 5451 | 47.843494 | | 0.1435118918 | 0.05392057 | 2.66154234 | 0.00777835 | 0.08646140 | HG01573_active | 0 | 0 |
| 541 | chr7 | 37682796 | 37684796 | ENSG00000 | GPR141 | 353345 | 40.70466609 | | 0.1439583596 | 0.05484927 | 2.62461747 | 0.00867463 | 0.09294451 | HG01573_active | 0 | 0 |
| 542 | chr1 | 10795650 | 10797650 | ENSG00000 | CASZ1 | 54897 | 58.28061997 | | 0.1444391973 | 0.05120478 | 2.82081455 | 0.00479018 | 0.06244069 | HG01573_active | 0 | 0 |
| 543 | chr4 | 86934002 | 86936002 | ENSG00000 | AFF1 | 4299 | 49.89773899 | | 0.1463895192 | 0.04733548 | 3.09259596 | 0.00198414 | 0.03403094 | HG01573_active | 0 | 0 |
| 544 | chr9 | 27528781 | 27530781 | ENSG00000 | MOB3B | 79817 | 68.0546037 | | 0.1471477406 | 0.04677894 | 3.1455977 | 0.00165747 | 0.02996693 | HG01573_active | 0 | 0 |
| 545 | chr22 | 31752951 | 31754951 | ENSG00000 | DEPDC5 | 9681 | 43.13806981 | | 0.1473972282 | 0.05347400 | 2.75642769 | 0.00584365 | 0.07078608 | HG01573_active | 0 | 0 |
| 546 | chr1 | 45490166 | 45492166 | ENSG00000 | TESK2 | 10420 | 38.18731355 | | 0.1478814095 | 0.05312939 | 2.78341995 | 0.00537891 | 0.06685363 | HG01573_active | 0 | 0 |
| 547 | chr7 | 3042945 | 3044945 | ENSG00000 | CARD11 | 84433 | 44.72449517 | | 0.1490255286 | 0.05714971 | 2.60763375 | 0.00911704 | 0.09606689 | HG01573_active | 0 | 0 |
| 548 | chr8 | 124727429 | 124729429 | ENSG00000 | MTSS1 | 9788 | 60.7682239 | | 0.1494158286 | 0.04868720 | 3.06889301 | 0.00214853 | 0.03610890 | HG01573_active | 0 | 0 |
| 549 | chr17 | 10197551 | 10199551 | ENSG00000 | GAS7 | 8522 | 94.16688176 | | 0.1495289935 | 0.04229712 | 3.53520506 | 0.00040745 | 0.01115172 | HG01573_active | 0 | 0 |
| 550 | chr1 | 247077811 | 247079811 | ENSG00000 | ZNF670 | 100533111 | 35.62616521 | | 0.1497463564 | 0.05710511 | 2.62229313 | 0.00873402 | 0.09337301 | HG01573_active | 0 | 0 |
| 551 | chr14 | 35121549 | 35123549 | ENSG00000 | KIAA0391 | NaN | 46.15739888 | | 0.1512537947 | 0.05243721 | 2.88447400 | 0.00392068 | 0.05492754 | HG01573_active | 0 | 0 |
| 552 | chr22 | 20858417 | 20860417 | ENSG00000 | PI4KA | 5297 | 46.9890793 | | 0.1513013989 | 0.05385582 | 2.80937840 | 0.00496372 | 0.06383094 | HG01573_active | 0 | 0 |
| 553 | chr11 | 3853527 | 3855527 | ENSG00000 | STIM1 | 6786 | 70.32897829 | | 0.1545806689 | 0.04088428 | 3.78093094 | 0.00015624 | 0.00503027 | HG01573_active | 0 | 0 |
| 554 | chr11 | 9137114 | 9139114 | ENSG00000 | SCUBE2 | 57758 | 38.33367233 | | 0.155925629 | 0.06029265 | 2.58614648 | 0.00970556 | 0.09944484 | HG01573_active | 0 | 0 |
| 555 | chr2 | 25672647 | 25674647 | ENSG00000 | DTNB | 1838 | 77.33159897 | | 0.1567208264 | 0.03887687 | 4.03120906 | 5.55E-05 | 0.00214280 | HG01573_active | 0 | 0 |
| 556 | chr11 | 73141261 | 73143261 | ENSG00000 | FCHSD2 | 9873 | 69.3146633 | | 0.1579402915 | 0.04699143 | 3.36104421 | 0.00077648 | 0.01723355 | HG01573_active | 0 | 0 |
| 557 | chr22 | 43410196 | 43412196 | ENSG00000 | MPPED1 | 758 | 36.04274391 | | 0.1580017099 | 0.06026901 | 2.62160779 | 0.00875160 | 0.09339475 | HG01573_active | 0 | 0 |
| 558 | chr6 | 170289703 | 170291703 | ENSG00000 | FAM120B | 84498 | 42.16118731 | | 0.1582415554 | 0.05542483 | 2.85506578 | 0.00430279 | 0.05842901 | HG01573_active | 0 | 0 |
| 559 | chr17 | 978770 | 980770 | ENSG00000 | NXN | 64359 | 57.48461615 | | 0.1590248205 | 0.06104298 | 2.60512854 | 0.00918397 | 0.09646788 | HG01573_active | 0 | 0 |
| 560 | chr14 | 89618149 | 89620149 | ENSG00000 | FOXN3 | 1112 | 152.491726 | | 0.1591727491 | 0.02993361 | 5.31752477 | 1.05E-07 | 1.22E-05 | HG01573_active | 0 | 0 |
| 561 | chr15 | 92808884 | 92810884 | ENSG00000 | FAM174B | 400451 | 57.97694102 | | 0.1622782157 | 0.04695016 | 3.45639253 | 0.00054745 | 0.01363613 | HG01573_active | 0 | 0 |
| 562 | chr2 | 213151427 | 213153427 | ENSG00000 | IKZF2 | 22807 | 32.72125593 | | 0.1630244066 | 0.06185374 | 2.63564346 | 0.00839779 | 0.09085254 | HG01573_active | 0 | 0 |
| 563 | chr13 | 99199774 | 99201774 | ENSG00000 | UBAC2 | 337867 | 50.45317653 | | 0.1635541386 | 0.05260712 | 3.10897290 | 0.00187738 | 0.03274647 | HG01573_active | 0 | 0 |
| 564 | chr3 | 185281941 | 185283941 | ENSG00000 | MAP3K13 | 9175 | 54.58105958 | | 0.1639713293 | 0.04679852 | 3.50377137 | 0.00045871 | 0.01208864 | HG01573_active | 0 | 0 |
| 565 | chr6 | 42745958 | 42747958 | ENSG00000 | GLTSCR1L | 23506 | 29.76834127 | | 0.1653579079 | 0.06251554 | 2.64506863 | 0.00816743 | 0.08900905 | HG01573_active | 0 | 0 |
| 566 | chr2 | 3376882 | 3378882 | ENSG00000 | TSSC1 | 7260 | 63.0101213 | | 0.1669256216 | 0.04497453 | 3.71155838 | 0.00020598 | 0.00635610 | HG01573_active | 0 | 0 |
| 567 | chr19 | 8444041 | 8446041 | ENSG00000 | ELAVL1 | 1994 | 130.7291903 | | 0.1679786312 | 0.03631650 | 4.62540790 | 3.74E-06 | 0.00023786 | HG01573_active | 0 | 0 |
| 568 | chr6 | 70411941 | 70413941 | ENSG00000 | FAM135A | 57579 | 32.65646086 | | 0.1688466202 | 0.06113140 | 2.76202769 | 0.00574436 | 0.07009168 | HG01573_active | 0 | 0 |
| 569 | chr11 | 121451203 | 121453203 | ENSG00000 | SORL1 | 6653 | 59.02442902 | | 0.1689480333 | 0.04582802 | 3.68656552 | 0.00022730 | 0.00687096 | HG01573_active | 0 | 1 |
| 570 | chr17 | 58691055 | 58693055 | ENSG00000 | TEX14 | 56155 | 38.90905453 | | 0.1703639373 | 0.05581730 | 3.05217068 | 0.00227192 | 0.03724569 | HG01573_active | 0 | 0 |
| 571 | chr7 | 152435005 | 152437005 | ENSG00000 | KMT2C | 58508 | 81.35451224 | | 0.1706619829 | 0.04175784 | 4.08694393 | 4.37E-05 | 0.00176434 | HG01573_active | 0 | 0 |
| 572 | chr6 | 13486662 | 13488662 | ENSG00000 | GFOD1 | 54438 | 39.19308171 | | 0.1713742314 | 0.05437948 | 3.15145024 | 0.00162461 | 0.02965372 | HG01573_active | 0 | 0 |
| 573 | chr4 | 42656105 | 42658105 | ENSG00000 | ATP8A1 | 10396 | 66.40317946 | | 0.1728273933 | 0.04386644 | 3.93985431 | 8.15E-05 | 0.00300092 | HG01573_active | 0 | 0 |
| 574 | chr9 | 90800787 | 90802787 | ENSG00000 | SYK | 6850 | 32.60758785 | | 0.1732924325 | 0.06397171 | 2.70889137 | 0.00675084 | 0.07795853 | HG01573_active | 0 | 0 |
| 575 | chr13 | 19957670 | 19959670 | ENSG00000 | ZMYM2 | 7750 | 26.78665836 | | 0.1735632737 | 0.06651277 | 2.60947274 | 0.00906818 | 0.09579980 | HG01573_active | 0 | 0 |
| 576 | chr17 | 77085716 | 77087716 | ENSG00000 | SEC14L1 | 6397 | 33.46911643 | | 0.1741066512 | 0.06347017 | 2.74312554 | 0.00608574 | 0.07259963 | HG01573_active | 0 | 0 |
| 577 | chr7 | 155643451 | 155645451 | ENSG00000 | RBM33 | 155435 | 36.23523592 | | 0.1743131972 | 0.06183611 | 2.81895472 | 0.00481803 | 0.06268132 | HG01573_active | 0 | 0 |
| 578 | chr21 | 43864186 | 43866186 | ENSG00000 | AGPAT3 | 56894 | 37.18947228 | | 0.1760755952 | 0.06691611 | 2.63128819 | 0.00850618 | 0.09180223 | HG01573_active | 0 | 0 |
| 579 | chr15 | 63832942 | 63834942 | ENSG00000 | HERC1 | 8925 | 54.51778619 | | 0.1779380791 | 0.05117136 | 3.47729757 | 0.00050649 | 0.01296414 | HG01573_active | 0 | 0 |
| 580 | chr4 | 6199591 | 6201591 | ENSG00000 | JAKMIP1 | 152789 | 64.18038606 | | 0.1780302458 | 0.04613982 | 3.85849412 | 0.00011408 | 0.00391857 | HG01573_active | 0 | 0 |
| 581 | chr13 | 28099592 | 28101592 | ENSG00000 | FLT3 | 2322 | 25.69033089 | | 0.1782487641 | 0.06814081 | 2.61588841 | 0.00889956 | 0.09437339 | HG01573_active | 0 | 0 |
| 582 | chr11 | 64722188 | 64724188 | ENSG00000 | NRXN2 | 9379 | 44.56067481 | | 0.1784884501 | 0.05799025 | 3.07790409 | 0.00208461 | 0.03534594 | HG01573_active | 0 | 0 |
| 583 | chr9 | 135701185 | 135703185 | ENSG00000 | KCNT1 | 57582 | 36.88293342 | | 0.1790791045 | 0.06603865 | 2.71173167 | 0.00669327 | 0.07756187 | HG01573_active | 0 | 0 |
| 584 | chr2 | 233353507 | 233355507 | ENSG00000 | DGKD | 8527 | 34.33088374 | | 0.179128114 | 0.06601634 | 2.71339022 | 0.00665986 | 0.07737602 | HG01573_active | 0 | 0 |
| 585 | chr15 | 52294798 | 52296798 | ENSG00000 | MYO5C | 55930 | 32.57976135 | | 0.1791509514 | 0.06141637 | 2.91699011 | 0.00353426 | 0.05141089 | HG01573_active | 0 | 0 |
| 586 | chr4 | 153465346 | 153467346 | ENSG00000 | KIAA0922 | 23240 | 44.81676207 | | 0.179460248 | 0.05410363 | 3.31697206 | 0.00090998 | 0.01945650 | HG01573_active | 0 | 0 |
| 587 | chr6 | 135102056 | 135104056 | ENSG00000 | HBS1L | 10767 | 31.94393104 | | 0.1797622743 | 0.06633745 | 2.70981577 | 0.00673205 | 0.07787108 | HG01573_active | 0 | 0 |
| 588 | chr22 | 41698499 | 41700499 | ENSG00000 | MEI1 | 150365 | 30.39415298 | | 0.1801320205 | 0.06130533 | 2.93827663 | 0.00330042 | 0.04883384 | HG01573_active | 0 | 0 |
| 589 | chr17 | 82751064 | 82753064 | ENSG00000 | TBCD | 6904 | 61.69068882 | | 0.1829574386 | 0.05379402 | 3.40107370 | 0.00067121 | 0.01553018 | HG01573_active | 0 | 0 |
| 590 | chr9 | 4665674 | 4667674 | ENSG00000 | SPATA6L | 55064 | 31.7989039 | | 0.183422159 | 0.06118362 | 2.99789603 | 0.00271850 | 0.04267054 | HG01573_active | 0 | 0 |
| 591 | chr1 | 13699198 | 13701198 | ENSG00000 | PRDM2 | 7799 | 36.88908049 | | 0.1835597737 | 0.06033196 | 3.04249636 | 0.00234624 | 0.03804393 | HG01573_active | 0 | 0 |
| 592 | chr5 | 143434512 | 143436512 | ENSG00000 | NR3C1 | 2908 | 36.08715367 | | 0.1844446988 | 0.06048242 | 3.04955846 | 0.00229178 | 0.03743334 | HG01573_active | 0 | 0 |
| 593 | chr22 | 42342616 | 42344616 | ENSG00000 | TCF20 | 6942 | 59.94686928 | | 0.1851725683 | 0.05039365 | 3.67452149 | 0.00023829 | 0.00710684 | HG01573_active | 0 | 0 |
| 594 | chr7 | 138001324 | 138003324 | ENSG00000 | AKR1D1 | 6718 | 29.85828992 | | 0.1857028194 | 0.06293214 | 2.95084184 | 0.00316909 | 0.04730975 | HG01573_active | 0 | 0 |
| 595 | chr2 | 171433802 | 171435802 | ENSG00000 | METTL8 | 79828 | 28.75940157 | | 0.1859367343 | 0.06371969 | 2.91804165 | 0.00352237 | 0.05137819 | HG01573_active | 0 | 0 |

| | | | | | | | | | | | | | | | | |
|---|---|---|---|---|---|---|---|---|---|---|---|---|---|---|---|---|
| 596 | chr3 | 46811574 | 46813574 | ENSG00000 | PRSS50 | NaN | 28.33269261 | | 0.1863771317 | 0.06271120( | 2.97199110‌9 | 0.00295875‌2 | 0.04533473‌9 | HG01573_active | 0 | 0 |
| 597 | chr2 | 47904793 | 47906793 | ENSG00000 | FBXO11 | 80204 | 25.34521202 | | 0.1873251385 | 0.06805531‌5 | 2.75254233‌7 | 0.00591344‌9 | 0.07140195‌8 | HG01573_active | 0 | 0 |
| 598 | chr10 | 5841132 | 5843132 | ENSG00000 | GDI2 | 2665 | 18.70142708 | | 0.1892654574 | 0.07313504‌9 | 2.58788993‌4 | 0.00965658‌3 | 0.09909493‌5 | HG01573_active | 0 | 0 |
| 599 | chr1 | 37946057 | 37948057 | ENSG00000 | INPP5B | 3633 | 24.27554043 | | 0.1900640326 | 0.07331248‌2 | 2.59251938‌8 | 0.00952758‌1 | 0.09863311‌5 | HG01573_active | 0 | 0 |
| 600 | chr8 | 100409015 | 100411015 | ENSG00000 | RNF19A | 25897 | 39.87317723 | | 0.1906517773 | 0.05614039‌4 | 3.39598212‌6 | 0.00068382‌8 | 0.01567174‌8 | HG01573_active | 0 | 0 |
| 601 | chr5 | 131244493 | 131246493 | ENSG00000 | CDC42SE2 | 56990 | 33.71788008 | | 0.1915827045 | 0.06448569 | 2.97093362‌1 | 0.00296895‌9 | 0.04538703‌3 | HG01573_active | 0 | 0 |
| 602 | chr2 | 60552567 | 60554567 | ENSG00000 | BCL11A | 53335 | 30.94834097 | | 0.1919801053 | 0.07022180‌6 | 2.73391009‌6 | 0.00625871‌3 | 0.07388560‌9 | HG01573_active | 0 | 0 |
| 603 | chr17 | 76452206 | 76454206 | ENSG00000 | UBE2O | 63893 | 19.19031641 | | 0.1921522757 | 0.07315089‌5 | 2.62679321 | 0.00861936‌9 | 0.09250072‌2 | HG01573_active | 0 | 0 |
| 604 | chr20 | 18466281 | 18468281 | ENSG00000 | DZANK1 | 55184 | 25.15405076 | | 0.1922176956 | 0.07120504‌9 | 2.69949530‌2 | 0.00694447‌3 | 0.07935017‌2 | HG01573_active | 0 | 0 |
| 605 | chr6 | 34391674 | 34393674 | ENSG00000 | NUDT3 | 11165 | 27.81810817 | | 0.1934715753 | 0.06674804‌8 | 2.89853532‌6 | 0.00374910( | 0.05317593‌9 | HG01573_active | 0 | 0 |
| 606 | chr19 | 16471085 | 16473085 | ENSG00000 | EPS15L1 | 58513 | 36.04080673 | | 0.1943341861 | 0.06508492‌4 | 2.98585560‌8 | 0.00282786‌1 | 0.04383185( | HG01573_active | 0 | 0 |
| 607 | chr7 | 77797792 | 77799792 | ENSG00000 | PHTF2 | 57157 | 34.26216972 | | 0.195005841 | 0.06684725‌8 | 2.91718529‌9 | 0.00353205‌8 | 0.05141089‌4 | HG01573_active | 0 | 0 |
| 608 | chr12 | 101876351 | 101878351 | ENSG00000 | DRAM1 | 55332 | 36.569456 | | 0.1954372345 | 0.05546525‌8 | 3.52359728 | 0.00042573( | 0.01154263‌4 | HG01573_active | 0 | 0 |
| 609 | chr1 | 206506530 | 206508530 | ENSG00000 | RASSF5 | 83593 | 28.19499516 | | 0.1954686466 | 0.06733422‌5 | 2.90296125‌1 | 0.00369652‌3 | 0.05259771‌3 | HG01573_active | 0 | 0 |
| 610 | chr16 | 67986746 | 67988746 | ENSG00000 | DUS2 | 54920 | 26.47088606 | | 0.19623946 | 0.06851847‌6 | 2.86403712‌4 | 0.00418278‌9 | 0.05747481‌4 | HG01573_active | 0 | 0 |
| 611 | chr22 | 29072078 | 29074078 | ENSG00000 | KREMEN1 | 83999 | 31.68595537 | | 0.1967418379 | 0.07025165( | 2.80052974‌8 | 0.00510188( | 0.06485844‌4 | HG01573_active | 0 | 0 |
| 612 | chr11 | 107927293 | 107929293 | ENSG00000 | SLC35F2 | 54733 | 36.75451237 | | 0.1972300156 | 0.05634885‌4 | 3.50016017‌7 | 0.00046497‌8 | 0.01215769( | HG01573_active | 0 | 0 |
| 613 | chr13 | 41060440 | 41062440 | ENSG00000 | ELF1 | 1997 | 30.48305685 | | 0.1982296533 | 0.06387623‌2 | 3.10333978‌2 | 0.00191349‌8 | 0.03315985‌5 | HG01573_active | 0 | 0 |
| 614 | chr21 | 46323122 | 46325122 | ENSG00000 | PCNT | 5116 | 35.33104815 | | 0.1983565709 | 0.06761870‌6 | 2.93345702‌8 | 0.00335210( | 0.04943423‌3 | HG01573_active | 0 | 0 |
| 615 | chr19 | 38432699 | 38434699 | ENSG00000 | RYR1 | 6261 | 46.78784675 | | 0.1992002991 | 0.05573307‌7 | 3.57418443 | 0.00035132‌1 | 0.00986135‌1 | HG01573_active | 0 | 0 |
| 616 | chr3 | 46811558 | 46813558 | ENSG00000 | PRSS45 | 377047 | 19.33091448 | | 0.200595751 | 0.07530086‌4 | 2.66392360‌9 | 0.00772350‌9 | 0.08599484‌7 | HG01573_active | 0 | 0 |
| 617 | chr20 | 5009293 | 5011293 | ENSG00000 | SLC23A2 | 9962 | 44.34662837 | | 0.2009262065 | 0.05606074‌1 | 3.58408045‌2 | 0.00033826‌7 | 0.00957541‌5 | HG01573_active | 0 | 0 |
| 618 | chr12 | 121580015 | 121582015 | ENSG00000 | KDM2B | 84678 | 42.43399456 | | 0.2009968809 | 0.06203549‌3 | 3.24003033‌5 | 0.00119516‌9 | 0.02338018‌1 | HG01573_active | 0 | 0 |
| 619 | chr1 | 65146549 | 65148549 | ENSG00000 | AK4 | 205 | 23.88020435 | | 0.2010264725 | 0.06658732‌4 | 3.01899007‌4 | 0.00253618‌8 | 0.04045770‌8 | HG01573_active | 0 | 0 |
| 620 | chr2 | 240819308 | 240821308 | ENSG00000 | KIF1A | 547 | 42.3552976 | | 0.2019683871 | 0.05834221‌7 | 3.46178797‌5 | 0.00053659‌9 | 0.01350189‌9 | HG01573_active | 0 | 0 |
| 621 | chr6 | 34465061 | 34467061 | ENSG00000 | PACSIN1 | 29993 | 24.61598696 | | 0.2038046277 | 0.06995117‌4 | 2.91352690‌7 | 0.00357371( | 0.05173168‌4 | HG01573_active | 0 | 0 |
| 622 | chr3 | 121748767 | 121750767 | ENSG00000 | GOLGB1 | 2804 | 21.31786267 | | 0.2052936945 | 0.07611285‌4 | 2.69722764‌6 | 0.00699194‌5 | 0.07970937( | HG01573_active | 0 | 0 |
| 623 | chr4 | 185203237 | 185205237 | ENSG00000 | SNX25 | 83891 | 40.32271281 | | 0.2070351201 | 0.05417292‌6 | 3.82174513‌7 | 0.00013251( | 0.00441514‌7 | HG01573_active | 0 | 0 |
| 624 | chr19 | 4968113 | 4970113 | ENSG00000 | KDM4B | 23030 | 59.09931955 | | 0.207823452 | 0.06000279‌7 | 3.46356271‌2 | 0.00053307‌2 | 0.01343845‌1 | HG01573_active | 0 | 0 |
| 625 | chr19 | 45078288 | 45080288 | ENSG00000 | MARK4 | 57787 | 66.95903505 | | 0.2084035897 | 0.04929916‌9 | 4.22732458‌3 | 2.36E-05 | 0.00106387( | HG01573_active | 0 | 0 |
| 626 | chr15 | 31325855 | 31327855 | ENSG00000 | KLF13 | 51621 | 33.99241069 | | 0.2090780329 | 0.05942682‌4 | 3.51824341‌4 | 0.00043441‌3 | 0.01169723‌2 | HG01573_active | 0 | 0 |
| 627 | chr6 | 37256772 | 37258772 | ENSG00000 | TBC1D22B | 55633 | 22.46475272 | | 0.2095756435 | 0.06965053‌8 | 3.00895940‌7 | 0.00262144‌1 | 0.04140080‌1 | HG01573_active | 0 | 0 |
| 628 | chr3 | 197748727 | 197750727 | ENSG00000 | KIAA0226 | 9711 | 22.94282975 | | 0.2098464785 | 0.07225194‌5 | 2.90437130‌4 | 0.00367991‌4 | 0.05252920‌7 | HG01573_active | 0 | 0 |
| 629 | chr17 | 49788180 | 49790180 | ENSG00000 | FAM117A | 81558 | 25.05298516 | | 0.2098575366 | 0.07267155( | 2.88775367‌5 | 0.00388003‌5 | 0.05448911‌2 | HG01573_active | 0 | 0 |
| 630 | chr17 | 74212514 | 74214514 | ENSG00000 | TTYH2 | 94015 | 18.50589768 | | 0.2125125168 | 0.07835972‌2 | 2.71201210‌1 | 0.00668761‌5 | 0.07756187‌9 | HG01573_active | 0 | 0 |
| 631 | chr11 | 113874570 | 113876570 | ENSG00000 | USP28 | 57646 | 20.89546722 | | 0.2125390726 | 0.08088877‌7 | 2.62754709‌1 | 0.00860029‌3 | 0.09237019‌6 | HG01573_active | 0 | 0 |
| 632 | chr5 | 303176 | 305176 | ENSG00000 | AHRR | 57491 | 35.90234641 | | 0.2125541178 | 0.06928686‌4 | 3.06774046‌4 | 0.00215683‌8 | 0.03615749 | HG01573_active | 0 | 0 |
| 633 | chr9 | 97500180 | 97502180 | ENSG00000 | TMOD1 | 7111 | 32.23018464 | | 0.2126323634 | 0.06369129‌4 | 3.33848391 | 0.00084236‌9 | 0.01830063‌9 | HG01573_active | 0 | 0 |
| 634 | chr2 | 37083342 | 37085342 | ENSG00000 | HEATR5B | 54497 | 25.36346119 | | 0.2126748701 | 0.06862502‌5 | 3.09908621‌4 | 0.00194118‌5 | 0.03342290‌7 | HG01573_active | 0 | 0 |
| 635 | chr3 | 170221365 | 170223365 | ENSG00000 | PRKCI | 5584 | 17.57280515 | | 0.2127233687 | 0.07880687‌6 | 2.69929958‌5 | 0.00694855‌9 | 0.07935017‌2 | HG01573_active | 0 | 0 |
| 636 | chr8 | 28489318 | 28491318 | ENSG00000 | FBXO16 | 157574 | 39.87833525 | | 0.2135323385 | 0.05663362‌7 | 3.77041605‌2 | 0.00016297‌5 | 0.00518456‌4 | HG01573_active | 0 | 0 |
| 637 | chr8 | 142448430 | 142450430 | ENSG00000 | BAI1 | 575 | 36.88906094 | | 0.2140055144 | 0.07320074‌2 | 2.92354294‌1 | 0.00346072‌4 | 0.05076213‌1 | HG01573_active | 0 | 0 |
| 638 | chr2 | 47694530 | 47696530 | ENSG00000 | MSH6 | 2956 | 31.25314549 | | 0.2140080549 | 0.05920366‌8 | 3.61477690‌4 | 0.00030060‌6 | 0.00861888‌7 | HG01573_active | 0 | 0 |
| 639 | chr12 | 111685056 | 111687056 | ENSG00000 | ACAD10 | 80724 | 19.82160404 | | 0.2140804293 | 0.07881718‌7 | 2.71616429‌1 | 0.00660431‌3 | 0.07688154‌8 | HG01573_active | 0 | 0 |
| 640 | chr3 | 122182683 | 122184683 | ENSG00000 | CASR | 846 | 31.45156749 | | 0.2141177002 | 0.06090347( | 3.51568963‌1 | 0.00043861‌3 | 0.01169723‌2 | HG01573_active | 0 | 0 |
| 641 | chr17 | 75404709 | 75406709 | ENSG00000 | GRB2 | 2885 | 20.87554498 | | 0.2141515982 | 0.07777649‌3 | 2.75342314‌6 | 0.00589756‌1 | 0.07130979‌1 | HG01573_active | 0 | 0 |
| 642 | chr7 | 129054173 | 129056173 | ENSG00000 | TNPO3 | 23534 | 25.05192382 | | 0.2143704252 | 0.06890064‌7 | 3.11129767 | 0.00186267( | 0.03257479‌4 | HG01573_active | 0 | 0 |
| 643 | chr3 | 13072117 | 13074117 | ENSG00000 | IQSEC1 | 9922 | 66.2561393 | | 0.2145529341 | 0.04812698‌3 | 4.45805907‌1 | 8.27E-06 | 0.00046429‌5 | HG01573_active | 0 | 0 |
| 644 | chr1 | 84996113 | 84998113 | ENSG00000 | MCOLN2 | 255231 | 19.80286029 | | 0.2148558355 | 0.07910482‌7 | 2.71609007‌4 | 0.00660579‌4 | 0.07688154‌8 | HG01573_active | 0 | 0 |
| 645 | chr9 | 5832117 | 5834117 | ENSG00000 | ERMP1 | 79956 | 17.69626837 | | 0.2149787776 | 0.08272056‌2 | 2.59885535‌5 | 0.00935351‌7 | 0.09748233‌2 | HG01573_active | 0 | 0 |
| 646 | chr9 | 37591469 | 37593469 | ENSG00000 | RP11-613M1 | 26267 | 25.25560833 | | 0.2155516829 | 0.06986208‌8 | 3.08538848‌7 | 0.00203286‌4 | 0.03464426‌1 | HG01573_active | 0 | 0 |
| 647 | chr19 | 40800297 | 40802297 | ENSG00000 | CTC-490E21 | NaN | 28.70532792 | | 0.2165825393 | 0.06581268‌1 | 3.29089370‌2 | 0.00099869‌6 | 0.02068772‌5 | HG01573_active | 0 | 0 |
| 648 | chr18 | 80246546 | 80248546 | ENSG00000 | PARD6G | 84552 | 30.47099251 | | 0.21736172 | 0.06073896‌1 | 3.57862092‌6 | 0.00034541‌2 | 0.00975697‌7 | HG01573_active | 0 | 0 |
| 649 | chr3 | 11845919 | 11847919 | ENSG00000 | TAMM41 | 132001 | 15.71267522 | | 0.2176882473 | 0.08277652‌3 | 2.62983074‌6 | 0.00854273‌8 | 0.09212230‌2 | HG01573_active | 0 | 0 |
| 650 | chr2 | 159903749 | 159905749 | ENSG00000 | LY75 | 4065 | 27.0789332 | | 0.2186594176 | 0.07720463‌5 | 2.83220581‌6 | 0.00462280‌8 | 0.06097269‌5 | HG01573_active | 0 | 0 |
| 651 | chr1 | 16436424 | 16438424 | ENSG00000 | SPATA21 | 374955 | 16.8618226 | | 0.2193671039 | 0.08045098‌7 | 2.72671736‌5 | 0.00639678( | 0.07504420‌5 | HG01573_active | 0 | 0 |
| 652 | chr4 | 1683302 | 1685302 | ENSG00000 | FAM53A | 152877 | 21.62761551 | | 0.219452088 | 0.07902731‌7 | 2.77691431‌4 | 0.00548776‌4 | 0.06781282‌9 | HG01573_active | 0 | 0 |
| 653 | chr12 | 12561383 | 12563383 | ENSG00000 | DUSP16 | 80824 | 22.68153772 | | 0.2195092776 | 0.07631481‌4 | 2.87636520‌7 | 0.00402283‌9 | 0.05598870‌9 | HG01573_active | 0 | 0 |
| 654 | chr16 | 87764992 | 87766992 | ENSG00000 | KLHDC4 | 54758 | 21.9447535 | | 0.2195468916 | 0.07499631‌7 | 2.92743562‌3 | 0.00341769‌8 | 0.05034604‌6 | HG01573_active | 0 | 0 |
| 655 | chr10 | 7410486 | 7412486 | ENSG00000 | SFMBT2 | 57713 | 78.75767339 | | 0.2217346665 | 0.04187286‌9 | 5.29542563‌8 | 1.19E-07 | 1.34E-05 | HG01573_active | 0 | 0 |

| | | | | | | | | | | | | | | | |
|---|---|---|---|---|---|---|---|---|---|---|---|---|---|---|---|
| 656 | chr5 | 139525431 | 139527431 | ENSG00000 | UBE2D2 | 7322 | 22.4590934 | 0.2218130503 | 0.07355131 | 3.01575911 | 0.00256336 | 0.04077281 | HG01573_active | 0 | 0 |
| 657 | chr17 | 78781297 | 78783297 | ENSG00000 | CYTH1 | 9267 | 30.1863572 | 0.2225684926 | 0.06011053 | 3.70265386 | 0.00021335 | 0.00653818 | HG01573_active | 0 | 0 |
| 658 | chr6 | 43368478 | 43370478 | ENSG00000 | ZNF318 | 24149 | 18.7209394 | 0.2227466703 | 0.08035268 | 2.77211239 | 0.00556938 | 0.06864620 | HG01573_active | 0 | 0 |
| 659 | chr5 | 151211150 | 151213150 | ENSG00000 | GM2A | 2760 | 19.6584741 | 0.2231907511 | 0.08412486 | 2.65308895 | 0.00797588 | 0.08763634 | HG01573_active | 0 | 0 |
| 660 | chr12 | 66187879 | 66189879 | ENSG00000 | IRAK3 | 11213 | 19.0123779 | 0.2241943672 | 0.08268468 | 2.71143758 | 0.00669921 | 0.07756344 | HG01573_active | 0 | 0 |
| 661 | chr1 | 9823526 | 9825526 | ENSG00000 | CLSTN1 | 22883 | 23.8082704 | 0.2245586179 | 0.07859114 | 2.85730191 | 0.00427259 | 0.05819174 | HG01573_active | 0 | 0 |
| 662 | chr8 | 141116278 | 141118278 | ENSG00000 | DENND3 | 22898 | 24.2278231 | 0.2246639547 | 0.07764001 | 2.89366203 | 0.00380777 | 0.05372303 | HG01573_active | 0 | 0 |
| 663 | chr4 | 498156 | 500156 | ENSG00000 | ZNF721 | 170960 | 15.2336213 | 0.2248552865 | 0.08669311 | 2.59369263 | 0.00949513 | 0.09849726 | HG01573_active | 0 | 0 |
| 664 | chr3 | 113531296 | 113533296 | ENSG00000 | SIDT1 | 54847 | 29.1384978 | 0.224971277 | 0.05935103 | 3.79052006 | 0.00015033 | 0.00485166 | HG01573_active | 0 | 0 |
| 665 | chr14 | 100586413 | 100588413 | ENSG00000 | BEGAIN | 57596 | 18.8640375 | 0.225828317 | 0.08339212 | 2.70802934 | 0.00676840 | 0.07805988 | HG01573_active | 0 | 0 |
| 666 | chr6 | 166998405 | 167000405 | ENSG00000 | CCR6 | NaN | 41.4537755 | 0.2258565478 | 0.05128982 | 4.40353535 | 1.07E-05 | 0.00057411 | HG01573_active | 0 | 0 |
| 667 | chr2 | 152716997 | 152718997 | ENSG00000 | PRPF40A | 55660 | 13.3949574 | 0.2260057883 | 0.08731342 | 2.58844253 | 0.00964110 | 0.09909493 | HG01573_active | 0 | 0 |
| 668 | chr3 | 196638775 | 196640775 | ENSG00000 | PIGX | 54965 | 26.0242318 | 0.2273912873 | 0.06669106 | 3.40962165 | 0.00065053 | 0.01521264 | HG01573_active | 0 | 0 |
| 669 | chr8 | 97643179 | 97645179 | ENSG00000 | MTDH | 92140 | 16.8054531 | 0.2277408205 | 0.08442691 | 2.69749066 | 0.00698642 | 0.07970937 | HG01573_active | 0 | 0 |
| 670 | chr12 | 109022089 | 109024089 | ENSG00000 | USP30 | 84749 | 18.7355912 | 0.2279787018 | 0.08310577 | 2.74323545 | 0.00608370 | 0.07259963 | HG01573_active | 0 | 0 |
| 671 | chr4 | 40629875 | 40631875 | ENSG00000 | RBM47 | 54502 | 52.3187766 | 0.2282303652 | 0.04737700 | 4.81732347 | 1.45E-06 | 0.00010921 | HG01573_active | 0 | 0 |
| 672 | chr1 | 111541218 | 111543218 | ENSG00000 | RAP1A | 5906 | 44.1871474 | 0.2284588382 | 0.05437475 | 4.20156116 | 2.65E-05 | 0.00117276 | HG01573_active | 0 | 0 |
| 673 | chr1 | 23580495 | 23582495 | ENSG00000 | MDS2 | NaN | 18.6549433 | 0.2286454436 | 0.08065088 | 2.83500233 | 0.00458253 | 0.06062104 | HG01573_active | 0 | 0 |
| 674 | chr3 | 47281917 | 47283917 | ENSG00000 | KLHL18 | 23276 | 18.9731986 | 0.228645452 | 0.08125658 | 2.81386969 | 0.00489490 | 0.06331156 | HG01573_active | 0 | 0 |
| 675 | chr4 | 48268864 | 48270864 | ENSG00000 | TEC | 7006 | 37.5381096 | 0.228661851 | 0.05585489 | 4.09385550 | 4.24E-05 | 0.00172816 | HG01573_active | 0 | 0 |
| 676 | chr7 | 100271218 | 100273218 | ENSG00000 | GATS | 352954 | 20.7634572 | 0.2300183205 | 0.08340514 | 2.75784339 | 0.00581840 | 0.07071690 | HG01573_active | 0 | 0 |
| 677 | chr9 | 37575349 | 37577349 | ENSG00000 | FBXO10 | 26267 | 21.7964054 | 0.2312708699 | 0.07446911 | 3.10559452 | 0.00189896 | 0.03295605 | HG01573_active | 0 | 0 |
| 678 | chr7 | 98400068 | 98402068 | ENSG00000 | BAIAP2L1 | 55971 | 29.9198681 | 0.2318219587 | 0.06913612 | 3.35312324 | 0.00079905 | 0.01761736 | HG01573_active | 0 | 0 |
| 679 | chr6 | 31356158 | 31358158 | ENSG00000 | HLA-C | 3107 | 26.7917879 | 0.2323990642 | 0.06405583 | 3.62807012 | 0.00028554 | 0.00827592 | HG01573_active | 0 | 0 |
| 680 | chr9 | 137422262 | 137424262 | ENSG00000 | EXD3 | 54932 | 35.4226549 | 0.2327966914 | 0.07083268 | 3.28657173 | 0.00101414 | 0.02091833 | HG01573_active | 0 | 0 |
| 681 | chr5 | 88903257 | 88905257 | ENSG00000 | MEF2C | 4208 | 39.1482580 | 0.2329865326 | 0.05964447 | 3.90625505 | 9.37E-05 | 0.00335771 | HG01573_active | 0 | 1 |
| 682 | chr1 | 206036231 | 206038231 | ENSG00000 | C1orf186 | 440712 | 21.8884072 | 0.2330368102 | 0.07538488 | 3.09129366 | 0.00199286 | 0.03413674 | HG01573_active | 0 | 0 |
| 683 | chr2 | 240568209 | 240570209 | ENSG00000 | ANKMY1 | 51281 | 27.6132511 | 0.2332822761 | 0.06947617 | 3.35773078 | 0.00078585 | 0.01735496 | HG01573_active | 0 | 0 |
| 684 | chr17 | 75440159 | 75442159 | ENSG00000 | KIAA0195 | 9772 | 19.8889611 | 0.2334333642 | 0.08520732 | 2.73959274 | 0.00615153 | 0.07307264 | HG01573_active | 0 | 0 |
| 685 | chr7 | 4882719 | 4884719 | ENSG00000 | RADIL | 55698 | 29.8640249 | 0.233541091 | 0.06794628 | 3.43714297 | 0.00058788 | 0.01417822 | HG01573_active | 0 | 0 |
| 686 | chr14 | 69610596 | 69612596 | ENSG00000 | KIAA0247 | 9766 | 31.6664603 | 0.2339576174 | 0.06412721 | 3.64833592 | 0.00026394 | 0.00775068 | HG01573_active | 0 | 0 |
| 687 | chr14 | 102776476 | 102778476 | ENSG00000 | TRAF3 | 7187 | 36.8321853 | 0.2341255261 | 0.06088678 | 3.84525996 | 0.00012042 | 0.00409413 | HG01573_active | 0 | 0 |
| 688 | chr12 | 121297308 | 121299308 | ENSG00000 | CAMKK2 | 10645 | 18.6390857 | 0.2352292809 | 0.07774062 | 3.02582194 | 0.00247958 | 0.03977155 | HG01573_active | 0 | 0 |
| 689 | chr19 | 10230286 | 10232286 | ENSG00000 | DNMT1 | 1786 | 26.6204641 | 0.23631939 | 0.06846085 | 3.45189059 | 0.00055667 | 0.01377354 | HG01573_active | 0 | 0 |
| 690 | chr8 | 144147016 | 144149016 | ENSG00000 | MROH1 | 727957 | 35.4336105 | 0.2370548476 | 0.06380730 | 3.71516775 | 0.00020306 | 0.00629513 | HG01573_active | 0 | 0 |
| 691 | chr9 | 133991604 | 133993604 | ENSG00000 | VAV2 | 7410 | 79.6096273 | 0.2389721347 | 0.05223644 | 4.57481634 | 4.77E-06 | 0.00029079 | HG01573_active | 0 | 0 |
| 692 | chr13 | 46210871 | 46212871 | ENSG00000 | LCP1 | 3936 | 24.4300449 | 0.2398637157 | 0.07182108 | 3.33973971 | 0.00083856 | 0.01824776 | HG01573_active | 0 | 0 |
| 693 | chr1 | 153921975 | 153923975 | ENSG00000 | GATAD2B | 57459 | 26.6419299 | 0.2403001252 | 0.07228912 | 3.32415329 | 0.00088687 | 0.01914301 | HG01573_active | 0 | 0 |
| 694 | chr14 | 105299563 | 105301563 | ENSG00000 | PACS2 | 23241 | 33.1435678 | 0.2409955123 | 0.06966898 | 3.45915066 | 0.00054188 | 0.01355004 | HG01573_active | 0 | 0 |
| 695 | chr18 | 77132683 | 77134683 | ENSG00000 | MBP | 4155 | 45.4295542 | 0.2410933341 | 0.05510375 | 4.37526136 | 1.21E-05 | 0.00062810 | HG01573_active | 0 | 0 |
| 696 | chr6 | 36870870 | 36872870 | ENSG00000 | C6orf89 | 221477 | 15.6647510 | 0.2429123855 | 0.08710851 | 2.78861824 | 0.00529334 | 0.06622129 | HG01573_active | 0 | 0 |
| 697 | chr9 | 136806922 | 136808922 | ENSG00000 | RABL6 | 55684 | 42.2817587 | 0.2430455105 | 0.06138128 | 3.95960294 | 7.51E-05 | 0.00280972 | HG01573_active | 0 | 0 |
| 698 | chr3 | 9648433 | 9650433 | ENSG00000 | MTMR14 | 64419 | 20.4120855 | 0.2432530931 | 0.07943919 | 3.06212932 | 0.00219768 | 0.03665572 | HG01573_active | 0 | 0 |
| 699 | chr20 | 48826883 | 48828883 | ENSG00000 | PREX1 | 57580 | 69.8105749 | 0.2441758154 | 0.04945734 | 4.93709876 | 7.93E-07 | 6.79E-05 | HG01573_active | 0 | 0 |
| 700 | chr19 | 19617916 | 19619916 | ENSG00000 | PBX4 | 80714 | 17.2306678 | 0.2444925139 | 0.09208599 | 2.65504561 | 0.00792976 | 0.08734512 | HG01573_active | 0 | 0 |
| 701 | chr4 | 6574175 | 6576175 | ENSG00000 | MAN2B2 | 23324 | 18.5722961 | 0.2466263971 | 0.08321703 | 2.96365264 | 0.00304011 | 0.04614937 | HG01573_active | 0 | 0 |
| 702 | chr16 | 14973591 | 14975591 | ENSG00000 | PDXDC1 | 23042 | 47.9325787 | 0.2478607739 | 0.05363341 | 4.62138674 | 3.81E-06 | 0.00024023 | HG01573_active | 0 | 0 |
| 703 | chr15 | 100918215 | 100920215 | ENSG00000 | LRRK1 | 79705 | 47.5453924 | 0.2492295889 | 0.05588603 | 4.45960428 | 8.21E-06 | 0.00046290 | HG01573_active | 0 | 0 |
| 704 | chr10 | 14953432 | 14955432 | ENSG00000 | DCLRE1C | 64421 | 14.9190336 | 0.2495259177 | 0.09277550 | 2.68956666 | 0.00715448 | 0.08136679 | HG01573_active | 0 | 0 |
| 705 | chr16 | 11248619 | 11250619 | ENSG00000 | RMI2 | 116028 | 32.2302707 | 0.2495819233 | 0.06120677 | 4.07768472 | 4.55E-05 | 0.00181959 | HG01573_active | 0 | 0 |
| 706 | chr3 | 13419309 | 13421309 | ENSG00000 | NUP210 | 23225 | 40.6004496 | 0.2501447975 | 0.05518173 | 4.53310847 | 5.81E-06 | 0.00034361 | HG01573_active | 0 | 0 |
| 707 | chr22 | 29204851 | 29206851 | ENSG00000 | EMID1 | 129080 | 18.2164313 | 0.2501551981 | 0.08360185 | 2.99222056 | 0.00276956 | 0.04320322 | HG01573_active | 0 | 0 |
| 708 | chr3 | 32684145 | 32686145 | ENSG00000 | CNOT10 | 25904 | 19.5231141 | 0.2501664871 | 0.07656709 | 3.26728457 | 0.00108584 | 0.02181649 | HG01573_active | 0 | 0 |
| 709 | chr21 | 42495008 | 42497008 | ENSG00000 | SLC37A1 | 54020 | 28.5837958 | 0.2506218712 | 0.06832478 | 3.66809604 | 0.00024436 | 0.00723933 | HG01573_active | 0 | 0 |
| 710 | chr5 | 159098761 | 159100761 | ENSG00000 | EBF1 | 1879 | 123.166313 | 0.2509627857 | 0.03341369 | 7.51077587 | 5.88E-14 | 2.53E-11 | HG01573_active | 0 | 0 |
| 711 | chr4 | 52658335 | 52660335 | ENSG00000 | USP46 | 64854 | 18.8605713 | 0.2515511875 | 0.08680965 | 2.89773269 | 0.00375870 | 0.05325566 | HG01573_active | 0 | 0 |
| 712 | chr2 | 240997838 | 240999838 | ENSG00000 | SNED1 | 25992 | 31.6193490 | 0.2519265196 | 0.06266534 | 4.02018857 | 5.82E-05 | 0.00223265 | HG01573_active | 0 | 0 |
| 713 | chr16 | 89815657 | 89817657 | ENSG00000 | FANCA | 2175 | 23.0411451 | 0.2519783589 | 0.08081407 | 3.11800088 | 0.00182082 | 0.03192652 | HG01573_active | 0 | 0 |
| 714 | chr17 | 42387568 | 42389568 | ENSG00000 | STAT3 | 6774 | 18.1789310 | 0.2520348142 | 0.08256107 | 3.05270743 | 0.00226786 | 0.03723081 | HG01573_active | 0 | 0 |
| 715 | chr9 | 37033185 | 37035185 | ENSG00000 | PAX5 | 5079 | 68.7486255 | 0.2523225075 | 0.04576459 | 5.51348665 | 3.52E-08 | 4.76E-06 | HG01573_active | 0 | 0 |

| | | | | | | | | | | | | | | | | |
|---|---|---|---|---|---|---|---|---|---|---|---|---|---|---|---|---|
| 716 | chr16 | 70438237 | 70440237 | ENSG00000 | ST3GAL2 | 6483 | 17.1873849 | | 0.2526205905 | 0.080784447 | 3.12709437 | 0.001765432 | 0.031270874 | HG01573_active | 0 | 0 |
| 717 | chr1 | 6179123 | 6181123 | ENSG00000 | CHD5 | 26038 | 30.84158485 | | 0.2548380518 | 0.066240179 | 3.847182414 | 0.000119484 | 0.004072514 | HG01573_active | 0 | 0 |
| 718 | chr14 | 35825318 | 35827318 | ENSG00000 | BRMS1L | 84312 | 24.76254873 | | 0.2550260686 | 0.070485748 | 3.618122463 | 0.000296747 | 0.008549334 | HG01573_active | 0 | 0 |
| 719 | chr22 | 43088428 | 43090428 | ENSG00000 | TTLL1 | 25809 | 15.76363672 | | 0.255363078 | 0.087846204 | 2.906933531 | 0.003649907 | 0.052324477 | HG01573_active | 0 | 0 |
| 720 | chr11 | 44563427 | 44565427 | ENSG00000 | CD82 | 3732 | 18.7315609 | | 0.2556043189 | 0.090791779 | 2.81528042 | 0.004873468 | 0.063217880 | HG01573_active | 0 | 0 |
| 721 | chr2 | 105743897 | 105745897 | ENSG00000 | NCK2 | 8440 | 46.62040674 | | 0.2563954203 | 0.052295694 | 4.902801748 | 9.45E-07 | 7.79E-05 | HG01573_active | 0 | 0 |
| 722 | chr15 | 75454842 | 75456842 | ENSG00000 | SIN3A | 25942 | 21.83966115 | | 0.2565565179 | 0.081034386 | 3.166020355 | 0.001545399 | 0.028401769 | HG01573_active | 0 | 0 |
| 723 | chr12 | 31728174 | 31730174 | ENSG00000 | AMN1 | 196394 | 13.81745478 | | 0.2570956829 | 0.093860189 | 2.739134506 | 0.006160116 | 0.073072644 | HG01573_active | 0 | 0 |
| 724 | chr9 | 123267576 | 123269576 | ENSG00000 | STRBP | 55342 | 41.25425908 | | 0.2584318854 | 0.058904761 | 4.387283473 | 1.15E-05 | 0.006073388 | HG01573_active | 0 | 0 |
| 725 | chr16 | 88452317 | 88454317 | ENSG00000 | ZFPM1 | 161882 | 28.45534329 | | 0.2588402239 | 0.082780509 | 3.126825685 | 0.001767046 | 0.031270874 | HG01573_active | 0 | 0 |
| 726 | chr5 | 179906859 | 179908859 | ENSG00000 | TBC1D9B | 23061 | 16.63807923 | | 0.259034907 | 0.091305633 | 2.837009029 | 0.004553832 | 0.060485947 | HG01573_active | 0 | 0 |
| 727 | chr22 | 43998211 | 44000211 | ENSG00000 | PARVB | 29780 | 59.73915517 | | 0.2596599349 | 0.048931017 | 5.306653053 | 1.12E-07 | 1.29E-05 | HG01573_active | 0 | 0 |
| 728 | chr4 | 931373 | 933373 | ENSG00000 | GAK | 2580 | 26.86705648 | | 0.2597015899 | 0.080989782 | 3.206596937 | 0.001343150 | 0.025698397 | HG01573_active | 0 | 0 |
| 729 | chr19 | 3223703 | 3225703 | ENSG00000 | CELF5 | 60680 | 21.7555637 | | 0.2600141804 | 0.079630515 | 3.265258018 | 0.001093644 | 0.021940207 | HG01573_active | 0 | 0 |
| 730 | chr3 | 48846846 | 48848846 | ENSG00000 | PRKAR2A | 5576 | 24.63751465 | | 0.2603855775 | 0.068642009 | 3.793385108 | 0.000148607 | 0.004823941 | HG01573_active | 0 | 0 |
| 731 | chr8 | 94718703 | 94720703 | ENSG00000 | DPY19L4 | 286148 | 16.1005539 | | 0.2610329672 | 0.087644237 | 2.978324344 | 0.002898291 | 0.044612979 | HG01573_active | 0 | 0 |
| 732 | chr5 | 32530633 | 32532633 | ENSG00000 | SUB1 | 10923 | 19.79218356 | | 0.2610680749 | 0.084182870 | 3.101201869 | 0.001927368 | 0.033313803 | HG01573_active | 0 | 0 |
| 733 | chr1 | 2002837 | 2004837 | ENSG00000 | KIAA1751 | 85452 | 27.71395594 | | 0.2610829781 | 0.085587053 | 3.050496175 | 0.002284635 | 0.037362326 | HG01573_active | 0 | 0 |
| 734 | chr19 | 19062999 | 19064999 | ENSG00000 | SLC25A42 | 284439 | 15.95155163 | | 0.2614779641 | 0.094102732 | 2.778643697 | 0.005458636 | 0.067592993 | HG01573_active | 0 | 0 |
| 735 | chr19 | 17074781 | 17076781 | ENSG00000 | MYO9B | 4650 | 38.40413772 | | 0.2615331384 | 0.062622988 | 4.176312032 | 2.96E-05 | 0.001285229 | HG01573_active | 0 | 0 |
| 736 | chr5 | 137886968 | 137888968 | ENSG00000 | PKD2L2 | 27039 | 12.58475997 | | 0.2616257516 | 0.101074035 | 2.588456578 | 0.009640710 | 0.099094935 | HG01573_active | 0 | 0 |
| 737 | chr11 | 111454630 | 111456630 | ENSG00000 | POU2AF1 | 5450 | 36.15505623 | | 0.2617424641 | 0.060633799 | 4.316774888 | 1.58E-05 | 0.000766444 | HG01573_active | 0 | 0 |
| 738 | chr5 | 134647789 | 134649789 | ENSG00000 | SEC24A | 10802 | 15.44356475 | | 0.2622580408 | 0.084409069 | 3.106988875 | 0.001890035 | 0.032881196 | HG01573_active | 0 | 0 |
| 739 | chr7 | 7967794 | 7969794 | ENSG00000 | GLCCI1 | 113263 | 28.40916375 | | 0.2625062936 | 0.069755197 | 3.763250695 | 0.000167718 | 0.005299452 | HG01573_active | 0 | 0 |
| 740 | chr5 | 172187386 | 172189386 | ENSG00000 | STK10 | 6793 | 44.31795698 | | 0.2628307487 | 0.059985820 | 4.381547921 | 1.18E-05 | 0.000617430 | HG01573_active | 0 | 0 |
| 741 | chr20 | 62641445 | 62643445 | ENSG00000 | SLCO4A1 | 28231 | 15.42408427 | | 0.2631918523 | 0.097446607 | 2.700882663 | 0.006915573 | 0.079108709 | HG01573_active | 0 | 0 |
| 742 | chr20 | 32051188 | 32053188 | ENSG00000 | HCK | 3055 | 16.6525641 | | 0.2633220362 | 0.086984035 | 3.027245573 | 0.002467933 | 0.039632286 | HG01573_active | 0 | 0 |
| 743 | chr10 | 35125791 | 35127791 | ENSG00000 | CREM | 1390 | 20.21199364 | | 0.2636564025 | 0.081903495 | 3.219110504 | 0.001285889 | 0.024827696 | HG01573_active | 0 | 0 |
| 744 | chr14 | 69725900 | 69727900 | ENSG00000 | SRSF5 | 6430 | 11.5991978 | | 0.2641287416 | 0.098225346 | 2.689007986 | 0.007166470 | 0.081366798 | HG01573_active | 0 | 0 |
| 745 | chr9 | 6644650 | 6646650 | ENSG00000 | GLDC | 2731 | 33.53112847 | | 0.2643253578 | 0.060128037 | 4.396041643 | 1.10E-05 | 0.000589181 | HG01573_active | 0 | 0 |
| 746 | chr17 | 81165077 | 81167077 | ENSG00000 | AATK | 9625 | 16.52138711 | | 0.2644327344 | 0.098624788 | 2.681199511 | 0.007335876 | 0.082623375 | HG01573_active | 0 | 0 |
| 747 | chr2 | 108447561 | 108449561 | ENSG00000 | GCC2 | 9648 | 13.26333148 | | 0.2646258898 | 0.097561456 | 2.712401993 | 0.006679753 | 0.077539688 | HG01573_active | 0 | 0 |
| 748 | chr12 | 124988122 | 124990122 | ENSG00000 | DHX37 | 57647 | 13.66250792 | | 0.2651371496 | 0.095112249 | 2.787623589 | 0.005309619 | 0.066300774 | HG01573_active | 0 | 0 |
| 749 | chr12 | 108560400 | 108562400 | ENSG00000 | SART3 | 9733 | 10.05407405 | | 0.2665909339 | 0.102739057 | 2.594835307 | 0.009463625 | 0.098320094 | HG01573_active | 0 | 0 |
| 750 | chr16 | 27401162 | 27403162 | ENSG00000 | IL21R | 50615 | 13.13549491 | | 0.2669849281 | 0.099470112 | 2.68407184 | 0.007273147 | 0.082378393 | HG01573_active | 0 | 0 |
| 751 | chr15 | 58770192 | 58772192 | ENSG00000 | FAM63B | 54629 | 19.31327991 | | 0.2672082403 | 0.082215226 | 3.250106485 | 0.001153618 | 0.022834800 | HG01573_active | 0 | 0 |
| 752 | chr20 | 63980135 | 63982135 | ENSG00000 | PRPF6 | 24148 | 16.16993766 | | 0.2680096274 | 0.094735515 | 2.82903013 | 0.004668930 | 0.061278563 | HG01573_active | 0 | 0 |
| 753 | chr5 | 100902266 | 100904266 | ENSG00000 | ST8SIA4 | 7903 | 18.46200069 | | 0.2680825282 | 0.087185601 | 3.074848651 | 0.002106093 | 0.035574616 | HG01573_active | 0 | 0 |
| 754 | chr7 | 26290895 | 26292895 | ENSG00000 | SNX10 | 29887 | 23.32096297 | | 0.2682488188 | 0.068843601 | 3.896496029 | 9.76E-05 | 0.003458776 | HG01573_active | 0 | 0 |
| 755 | chr20 | 38804705 | 38806705 | ENSG00000 | PPP1R16B | 26051 | 40.14238882 | | 0.2685484785 | 0.061345276 | 4.377655358 | 1.20E-05 | 0.000626102 | HG01573_active | 0 | 0 |
| 756 | chr20 | 43666019 | 43668019 | ENSG00000 | MYBL2 | 4605 | 14.43670213 | | 0.2686362254 | 0.095650431 | 2.808520786 | 0.004976966 | 0.063878247 | HG01573_active | 0 | 0 |
| 757 | chr1 | 233613004 | 233615004 | ENSG00000 | KCNK1 | 3775 | 18.66697129 | | 0.2688405293 | 0.078069238 | 3.443616665 | 0.000573988 | 0.013983944 | HG01573_active | 0 | 0 |
| 758 | chr8 | 11493001 | 11495001 | ENSG00000 | BLK | 640 | 20.87120405 | | 0.2692960014 | 0.081994778 | 3.284306708 | 0.001022335 | 0.021015605 | HG01573_active | 0 | 0 |
| 759 | chr1 | 40343850 | 40345850 | ENSG00000 | SMAP2 | 64744 | 20.44824669 | | 0.2697702629 | 0.077965473 | 3.460124731 | 0.000539925 | 0.013534598 | HG01573_active | 0 | 0 |
| 760 | chr1 | 26528761 | 26530761 | ENSG00000 | RPS6KA1 | 6195 | 16.27470027 | | 0.269914973 | 0.087478037 | 3.08551703 | 0.002031985 | 0.034644261 | HG01573_active | 0 | 0 |
| 761 | chr19 | 4182354 | 4184354 | ENSG00000 | ANKRD24 | 170961 | 11.66476954 | | 0.2715379396 | 0.102515083 | 2.648760855 | 0.008078747 | 0.088403068 | HG01573_active | 0 | 0 |
| 762 | chr19 | 12639098 | 12641098 | ENSG00000 | ZNF490 | 57474 | 13.2891544 | | 0.2717034449 | 0.102287737 | 2.656266056 | 0.007901125 | 0.087101429 | HG01573_active | 0 | 0 |
| 763 | chr6 | 136791518 | 136793518 | ENSG00000 | MAP3K5 | 4217 | 58.25450795 | | 0.2719444234 | 0.047891025 | 5.678400518 | 1.36E-08 | 2.00E-06 | HG01573_active | 0 | 0 |
| 764 | chr12 | 51323668 | 51325668 | ENSG00000 | BIN2 | 51411 | 11.82916157 | | 0.2725263477 | 0.097407945 | 2.797783542 | 0.005145458 | 0.065102715 | HG01573_active | 0 | 0 |
| 765 | chr1 | 151069578 | 151071578 | ENSG00000 | GABPB2 | 126626 | 12.16405855 | | 0.2733262262 | 0.096947319 | 2.819327305 | 0.004812441 | 0.062669625 | HG01573_active | 0 | 0 |
| 766 | chr19 | 3868032 | 3870032 | ENSG00000 | ZFR2 | 23217 | 20.45989649 | | 0.2738110492 | 0.083800102 | 3.267430955 | 0.001085283 | 0.021816497 | HG01573_active | 0 | 0 |
| 767 | chr18 | 69960803 | 69962803 | ENSG00000 | CD226 | 10666 | 36.61330948 | | 0.2738675554 | 0.061256599 | 4.47082534 | 7.79E-06 | 0.000441129 | HG01573_active | 0 | 0 |
| 768 | chr19 | 54965312 | 54967312 | ENSG00000 | NLRP7 | 199713 | 12.39392956 | | 0.2740135646 | 0.101268225 | 2.705819755 | 0.006813601 | 0.078344692 | HG01573_active | 0 | 0 |
| 769 | chr2 | 119365828 | 119367828 | ENSG00000 | C2orf76 | 130355 | 16.56921381 | | 0.2756411689 | 0.083112041 | 3.31650102 | 0.000911522 | 0.019456500 | HG01573_active | 0 | 0 |
| 770 | chr3 | 16513026 | 16515026 | ENSG00000 | RFTN1 | 23180 | 61.75519544 | | 0.27608725 | 0.048941703 | 5.641145119 | 1.69E-08 | 2.43E-06 | HG01573_active | 0 | 0 |
| 771 | chr3 | 23804903 | 23806903 | ENSG00000 | UBE2E1 | 7324 | 20.91202134 | | 0.27615984 | 0.081701146 | 3.380121975 | 0.000724536 | 0.016407685 | HG01573_active | 0 | 0 |
| 772 | chr9 | 92946451 | 92948451 | ENSG00000 | FGD3 | 89846 | 30.46052873 | | 0.2763219678 | 0.062350724 | 4.43173631 | 9.35E-06 | 0.000513971 | HG01573_active | 0 | 0 |
| 773 | chr6 | 166998182 | 167000182 | ENSG00000 | FGFR1OP | 11116 | 13.58552189 | | 0.2773289869 | 0.093674748 | 2.960552236 | 0.003070880 | 0.046361619 | HG01573_active | 0 | 0 |
| 774 | chr19 | 2701709 | 2703709 | ENSG00000 | GNG7 | 2788 | 49.9499494 | | 0.2776134054 | 0.059680489 | 4.651661027 | 3.29E-06 | 0.000214605 | HG01573_active | 0 | 0 |
| 775 | chr19 | 29922644 | 29924644 | ENSG00000 | URI1 | 8725 | 17.84671924 | | 0.2787289145 | 0.082199571 | 3.390880374 | 0.000696685 | 0.015884656 | HG01573_active | 0 | 0 |

| | | | | | | | | | | | | | | | |
|---|---|---|---|---|---|---|---|---|---|---|---|---|---|---|---|
| 776 | chr1 | 12166003 | 12168003 | ENSG00000 | TNFRSF1B | 7133 | 15.37729648 | 0.2789641362 | 0.093345743 | 2.98850408 | 0.002803467 | 0.043528457 | HG01573_active | 0 | 0 |
| 777 | chr2 | 230202110 | 230204110 | ENSG00000 | SP140 | 11262 | 32.872889 | 0.2807799063 | 0.063853030 | 4.397283931 | 1.10E-05 | 0.000588173 | HG01573_active | 0 | 0 |
| 778 | chr1 | 9650732 | 9652732 | ENSG00000 | PIK3CD | 5293 | 21.16930519 | 0.2810067125 | 0.080524856 | 3.489689075 | 0.000483582 | 0.012545922 | HG01573_active | 0 | 0 |
| 779 | chr5 | 126775623 | 126777623 | ENSG00000 | LMNB1 | 4001 | 13.62917006 | 0.2817179648 | 0.090714428 | 3.105547494 | 0.001899271 | 0.032956054 | HG01573_active | 0 | 0 |
| 780 | chr6 | 43242680 | 43244680 | ENSG00000 | TTBK1 | 84630 | 15.88528938 | 0.2819036924 | 0.098802123 | 2.853214902 | 0.004327934 | 0.058587162 | HG01573_active | 0 | 0 |
| 781 | chr13 | 40665597 | 40667597 | ENSG00000 | FOXO1 | 2308 | 31.43637101 | 0.2820992343 | 0.060707475 | 4.646861583 | 3.37E-06 | 0.000218590 | HG01573_active | 0 | 0 |
| 782 | chr15 | 74956219 | 74958219 | ENSG00000 | SCAMP5 | 192683 | 18.63111553 | 0.2836808579 | 0.082205402 | 3.450878532 | 0.000558764 | 0.013779554 | HG01573_active | 0 | 0 |
| 783 | chr15 | 90528925 | 90530925 | ENSG00000 | CRTC3 | 64784 | 33.49458307 | 0.2845169384 | 0.058642409 | 4.851726589 | 1.22E-06 | 9.73E-05 | HG01573_active | 0 | 0 |
| 784 | chr20 | 47500902 | 47502902 | ENSG00000 | NCOA3 | 8202 | 32.99741042 | 0.2848362313 | 0.060429186 | 4.71355394 | 2.43E-06 | 0.000168523 | HG01573_active | 0 | 0 |
| 785 | chr1 | 40160373 | 40162373 | ENSG00000 | RLF | 6018 | 17.0992204 | 0.286860233 | 0.082398360 | 3.481382765 | 0.000498832 | 0.012866593 | HG01573_active | 0 | 0 |
| 786 | chr2 | 201181881 | 201183881 | ENSG00000 | CASP10 | 843 | 13.47984545 | 0.2879957047 | 0.097518055 | 2.953255188 | 0.003144419 | 0.047037478 | HG01573_active | 0 | 0 |
| 787 | chr21 | 44930989 | 44932989 | ENSG00000 | ITGB2 | 3689 | 14.23011781 | 0.2881806333 | 0.102603329 | 2.808686969 | 0.004974398 | 0.063878247 | HG01573_active | 0 | 0 |
| 788 | chr22 | 39151674 | 39153674 | ENSG00000 | CBX7 | 23492 | 10.84118565 | 0.2895165613 | 0.105022924 | 2.756698723 | 0.005838812 | 0.070786082 | HG01573_active | 0 | 0 |
| 789 | chr19 | 45601212 | 45603212 | ENSG00000 | OPA3 | 80207 | 19.97354279 | 0.2898430751 | 0.084888307 | 3.414405154 | 0.000639214 | 0.015074187 | HG01573_active | 0 | 0 |
| 790 | chr22 | 50055935 | 50057935 | ENSG00000 | TTLL8 | 164714 | 13.12816029 | 0.2909045158 | 0.096756395 | 3.006566272 | 0.002642164 | 0.041678819 | HG01573_active | 0 | 0 |
| 791 | chr8 | 37898467 | 37900467 | ENSG00000 | RAB11FIP1 | 80223 | 10.87371547 | 0.2909751411 | 0.111603483 | 2.607222742 | 0.009127996 | 0.096106505 | HG01573_active | 0 | 0 |
| 792 | chr16 | 23835001 | 23837001 | ENSG00000 | PRKCB | 5579 | 123.3572874 | 0.2918978084 | 0.036875267 | 7.915815259 | 2.46E-15 | 1.31E-12 | HG01573_active | 0 | 0 |
| 793 | chrX | 1452762 | 1454762 | ENSG00000 | ASMTL | 8623 | 13.97051866 | 0.2919112582 | 0.104671756 | 2.788825446 | 0.005289956 | 0.066221293 | HG01573_active | 0 | 0 |
| 794 | chr1 | 160399586 | 160401586 | ENSG00000 | VANGL2 | 57216 | 11.63062934 | 0.2936060395 | 0.094727512 | 3.099480089 | 0.001938606 | 0.033422907 | HG01573_active | 0 | 0 |
| 795 | chr3 | 196081189 | 196083189 | ENSG00000 | TFRC | 7037 | 13.56196576 | 0.2936367653 | 0.095593044 | 3.071737765 | 0.002128165 | 0.035902052 | HG01573_active | 0 | 0 |
| 796 | chr19 | 11261481 | 11263481 | ENSG00000 | DOCK6 | 57572 | 18.43650169 | 0.2939007579 | 0.085969113 | 3.418678484 | 0.000629260 | 0.014880614 | HG01573_active | 0 | 0 |
| 797 | chr7 | 149125416 | 149127416 | ENSG00000 | ZNF398 | 57541 | 13.38161732 | 0.2942029196 | 0.092051919 | 3.196054151 | 0.001393209 | 0.026409085 | HG01573_active | 0 | 0 |
| 798 | chr19 | 18097944 | 18099944 | ENSG00000 | IL12RB1 | 3594 | 10.81446204 | 0.2943219764 | 0.106938341 | 2.75225865 | 0.005918575 | 0.071401958 | HG01573_active | 0 | 0 |
| 799 | chr16 | 46620626 | 46622626 | ENSG00000 | SHCBP1 | 79801 | 9.427359189 | 0.2952924639 | 0.103939811 | 2.840994791 | 0.004497304 | 0.060068975 | HG01573_active | 0 | 0 |
| 800 | chr7 | 2557972 | 2559972 | ENSG00000 | IQCE | 23288 | 18.55623547 | 0.295317894 | 0.083398561 | 3.541043015 | 0.000398548 | 0.010979397 | HG01573_active | 0 | 0 |
| 801 | chr11 | 46331905 | 46333905 | ENSG00000 | DGKZ | 8525 | 15.65314098 | 0.2961383001 | 0.097551896 | 3.035700086 | 0.002399779 | 0.038770801 | HG01573_active | 0 | 0 |
| 802 | chr1 | 24186959 | 24188959 | ENSG00000 | IFNLR1 | 163702 | 10.64457383 | 0.2966970015 | 0.107366108 | 2.763413939 | 0.005720017 | 0.069858458 | HG01573_active | 0 | 0 |
| 803 | chr3 | 197028816 | 197030816 | ENSG00000 | MFI2 | 4241 | 14.0297342 | 0.2975738721 | 0.101241456 | 2.939249234 | 0.003290083 | 0.048734823 | HG01573_active | 0 | 0 |
| 804 | chr19 | 40881210 | 40883210 | ENSG00000 | CYP2A6 | 1548 | 12.59584049 | 0.298226878 | 0.105390518 | 2.829731578 | 0.004658707 | 0.061235735 | HG01573_active | 0 | 0 |
| 805 | chr3 | 4886293 | 4888293 | ENSG00000 | AC018816.3 | NaN | 19.92547584 | 0.2983414788 | 0.077117960 | 3.868638099 | 0.000109444 | 0.003798165 | HG01573_active | 0 | 0 |
| 806 | chr1 | 236141505 | 236143505 | ENSG00000 | GPR137B | 7107 | 24.95815961 | 0.2988076174 | 0.071816182 | 4.160728214 | 3.17E-05 | 0.001349862 | HG01573_active | 0 | 0 |
| 807 | chr18 | 79394856 | 79396856 | ENSG00000 | NFATC1 | 4772 | 42.0909146 | 0.2994764129 | 0.066740089 | 4.487204265 | 7.22E-06 | 0.000412043 | HG01573_active | 0 | 0 |
| 808 | chr19 | 11504698 | 11506698 | ENSG00000 | CTC-398G3. | NaN | 11.81207344 | 0.2998672951 | 0.107681864 | 2.784752061 | 0.005356866 | 0.066641616 | HG01573_active | 0 | 0 |
| 809 | chr1 | 167629093 | 167631093 | ENSG00000 | RCSD1 | 92241 | 27.79248635 | 0.3004394572 | 0.065274034 | 4.602740748 | 4.17E-06 | 0.000259120 | HG01573_active | 0 | 0 |
| 810 | chr7 | 2353318 | 2355318 | ENSG00000 | SNX8 | 29886 | 26.06513722 | 0.3014198523 | 0.077458804 | 3.891356873 | 9.97E-05 | 0.003514231 | HG01573_active | 0 | 0 |
| 811 | chr11 | 111511354 | 111513354 | ENSG00000 | BTG4 | 54766 | 11.48992402 | 0.3019829071 | 0.105751254 | 2.855596455 | 0.004295607 | 0.058429016 | HG01573_active | 0 | 0 |
| 812 | chr1 | 52926212 | 52928212 | ENSG00000 | ECHDC2 | 55268 | 9.213469498 | 0.3036362025 | 0.115802756 | 2.622011876 | 0.008741238 | 0.093373018 | HG01573_active | 0 | 0 |
| 813 | chr9 | 94258311 | 94260311 | ENSG00000 | ZNF169 | 169841 | 10.28484624 | 0.303662482 | 0.104717081 | 2.899837131 | 0.003733566 | 0.053011876 | HG01573_active | 0 | 0 |
| 814 | chr1 | 32291086 | 32293086 | ENSG00000 | HDAC1 | 3065 | 11.11835147 | 0.3039690172 | 0.108102612 | 2.811856344 | 0.004925649 | 0.063586092 | HG01573_active | 0 | 0 |
| 815 | chr11 | 66465738 | 66467738 | ENSG00000 | MRPL11 | 65003 | 9.97761913 | 0.3039922331 | 0.117227708 | 2.593177307 | 0.009509373 | 0.098568451 | HG01573_active | 0 | 0 |
| 816 | chr20 | 5749393 | 5751393 | ENSG00000 | C20orf196 | 149840 | 32.20639712 | 0.3058544397 | 0.064842454 | 4.716885577 | 2.39E-06 | 0.000166652 | HG01573_active | 0 | 0 |
| 817 | chr5 | 133050962 | 133052962 | ENSG00000 | HSPA4 | 3308 | 11.73412963 | 0.3061951111 | 0.103370919 | 2.962101067 | 0.003055474 | 0.046181219 | HG01573_active | 0 | 0 |
| 818 | chr19 | 18606741 | 18608741 | ENSG00000 | CRLF1 | 9244 | 10.24635551 | 0.3067390227 | 0.104513254 | 2.934929389 | 0.003336235 | 0.049254639 | HG01573_active | 0 | 0 |
| 819 | chr5 | 10563330 | 10565330 | ENSG00000 | ANKRD33B | 651746 | 27.26478487 | 0.3074138052 | 0.068445148 | 4.491389274 | 7.08E-06 | 0.000405762 | HG01573_active | 0 | 0 |
| 820 | chr3 | 53155009 | 53157009 | ENSG00000 | PRKCD | 5580 | 13.13031758 | 0.3079452679 | 0.110374883 | 2.789994048 | 0.005270901 | 0.066106725 | HG01573_active | 0 | 0 |
| 821 | chr20 | 64047758 | 64049758 | ENSG00000 | ZNF512B | 57473 | 27.86215419 | 0.3080486409 | 0.078075888 | 3.945502874 | 7.96E-05 | 0.002939149 | HG01573_active | 0 | 0 |
| 822 | chr9 | 136244841 | 136246841 | ENSG00000 | QSOX2 | 169714 | 13.84090109 | 0.3082917025 | 0.104198930 | 2.958683949 | 0.003089557 | 0.046486016 | HG01573_active | 0 | 0 |
| 823 | chr2 | 201115104 | 201117104 | ENSG00000 | CFLAR | 8837 | 15.48447847 | 0.3102545968 | 0.090677187 | 3.421528682 | 0.000622701 | 0.014777825 | HG01573_active | 0 | 0 |
| 824 | chr1 | 6423788 | 6425788 | ENSG00000 | ESPN | 83715 | 13.0617649 | 0.3105289997 | 0.095727490 | 3.243885311 | 0.001179112 | 0.023150795 | HG01573_active | 0 | 0 |
| 825 | chr19 | 3699479 | 3701479 | ENSG00000 | PIP5K1C | 23396 | 21.60504038 | 0.3107072567 | 0.084647304 | 3.670610155 | 0.000241972 | 0.007184422 | HG01573_active | 0 | 0 |
| 826 | chr5 | 71586288 | 71588288 | ENSG00000 | MCCC2 | 64087 | 17.87135227 | 0.3111522216 | 0.082030138 | 3.793145161 | 0.000148751 | 0.004823941 | HG01573_active | 0 | 0 |
| 827 | chr11 | 61294424 | 61296424 | ENSG00000 | VWCE | 220001 | 12.23451576 | 0.3114533195 | 0.096681083 | 3.221450438 | 0.001275435 | 0.024661490 | HG01573_active | 0 | 0 |
| 828 | chr19 | 11558374 | 11560374 | ENSG00000 | ZNF627 | 199692 | 14.61354155 | 0.312427527 | 0.099218022 | 3.148898948 | 0.001638868 | 0.029751253 | HG01573_active | 0 | 0 |
| 829 | chr17 | 30967785 | 30969785 | ENSG00000 | RNF135 | 84282 | 7.570774621 | 0.3125482198 | 0.121000447 | 2.583033579 | 0.009793579 | 0.099963338 | HG01573_active | 0 | 0 |
| 830 | chr19 | 40591883 | 40593883 | ENSG00000 | LTBP4 | 8425 | 9.920852086 | 0.3126857717 | 0.113912666 | 2.744960516 | 0.006051817 | 0.072388843 | HG01573_active | 0 | 0 |
| 831 | chr19 | 1811276 | 1813276 | ENSG00000 | ATP8B3 | 148229 | 9.073078027 | 0.312735483 | 0.120643620 | 2.592225615 | 0.009535721 | 0.098633115 | HG01573_active | 0 | 0 |
| 832 | chr11 | 61791980 | 61793980 | ENSG00000 | FADS2 | 9415 | 23.07378011 | 0.3127906738 | 0.081059152 | 3.858795255 | 0.000113947 | 0.003918578 | HG01573_active | 0 | 0 |
| 833 | chr15 | 74872365 | 74874365 | ENSG00000 | SCAMP2 | 10066 | 9.242555897 | 0.3129965421 | 0.120387713 | 2.599043699 | 0.009324974 | 0.097336704 | HG01573_active | 0 | 0 |
| 834 | chr19 | 3046635 | 3048635 | ENSG00000 | TLE2 | 7089 | 15.08347786 | 0.3136385457 | 0.089326056 | 3.511165238 | 0.000446146 | 0.011874442 | HG01573_active | 0 | 0 |
| 835 | chr7 | 100176563 | 100178563 | ENSG00000 | STAG3 | 10734 | 12.54998095 | 0.3150111067 | 0.095999230 | 3.281391991 | 0.001032960 | 0.021110451 | HG01573_active | 0 | 0 |

| | | | | | | | | | | | | | | | |
|---|---|---|---|---|---|---|---|---|---|---|---|---|---|---|---|
| 836 | chr7 | 2402560 | 2404560 | ENSG00000 | CHST12 | 55501 | 13.41525542 | 0.3150931084 | 0.09570395€ | 3.29237286′ | 0.00099345€ | 0.02061116€ | HG01573_active | 0 | 0 |
| 837 | chr1 | 9128170 | 9130170 | ENSG00000 | GPR157 | 80045 | 8.593600162 | 0.3151207345 | 0.119119031 | 2.64542727€ | 0.00815878‹ | 0.08898733€ | HG01573_active | 0 | 0 |
| 838 | chr15 | 43825963 | 43827963 | ENSG00000 | WDR76 | 79968 | 9.12936803 | 0.3154108129 | 0.114235921 | 2.76104755‹ | 0.00576162€ | 0.07023824€ | HG01573_active | 0 | 0 |
| 839 | chr16 | 50692603 | 50694603 | ENSG00000 | NOD2 | 64127 | 14.16514264 | 0.3169816242 | 0.08978998‹ | 3.53025593€ | 0.00041515€ | 0.01132025€ | HG01573_active | 0 | 0 |
| 840 | chr17 | 63971918 | 63973918 | ENSG00000 | SCN4A | 6329 | 12.61311124 | 0.3173476806 | 0.10372261€ | 3.05958028€ | 0.00221647€ | 0.03687958‹ | HG01573_active | 0 | 0 |
| 841 | chr19 | 35868287 | 35870287 | ENSG00000 | NPHS1 | 4868 | 14.35091025 | 0.3176291748 | 0.09688757€ | 3.27832712€ | 0.00104424€ | 0.02120384€ | HG01573_active | 0 | 0 |
| 842 | chr19 | 12440082 | 12442082 | ENSG00000 | CTD-3105H′ | NaN | 12.01733162 | 0.3182124042 | 0.09873638€ | 3.22284828€ | 0.00126922€ | 0.02461269€ | HG01573_active | 0 | 0 |
| 843 | chr17 | 75720328 | 75722328 | ENSG00000 | ITGB4 | 3691 | 12.24212988 | 0.318515824 | 0.09928625€ | 3.20805562€ | 0.00133635€ | 0.02561701‹ | HG01573_active | 0 | 0 |
| 844 | chr10 | 12067972 | 12069972 | ENSG00000 | DHTKD1 | 55526 | 14.09194801 | 0.3192408473 | 0.09637604€ | 3.31245017′ | 0.00092482€ | 0.01964515′ | HG01573_active | 0 | 0 |
| 845 | chr12 | 7128698 | 7130698 | ENSG00000 | CLSTN3 | 9746 | 10.47468888 | 0.3198283738 | 0.10502326€ | 3.04530988‹ | 0.00232440€ | 0.03784986€ | HG01573_active | 0 | 0 |
| 846 | chr19 | 7394113 | 7396113 | ENSG00000 | ARHGEF18 | 23370 | 26.01152926 | 0.3199310958 | 0.07444976€ | 4.29727486€ | 1.73E-05 | 0.00081924′ | HG01573_active | 0 | 0 |
| 847 | chr19 | 19191158 | 19193158 | ENSG00000 | MEF2BNB-M | 4207 | 13.74450555 | 0.3203261481 | 0.09882581€ | 3.24132057€ | 0.00118977€ | 0.02330873€ | HG01573_active | 0 | 0 |
| 848 | chr12 | 124992700 | 124994700 | ENSG00000 | BRI3BP | 140707 | 10.92555858 | 0.3213041806 | 0.10689288‹ | 3.00585191€ | 0.00264837€ | 0.04172759€ | HG01573_active | 0 | 0 |
| 849 | chr9 | 137617963 | 137619963 | ENSG00000 | EHMT1 | 79813 | 71.45837155 | 0.3213587931 | 0.04524874€ | 7.10204831€ | 1.23E-12 | 3.87E-10 | HG01573_active | 0 | 0 |
| 850 | chr19 | 7293034 | 7295034 | ENSG00000 | INSR | 3643 | 49.32303479 | 0.3214459598 | 0.05590173€ | 5.75019643€ | 8.91E-09 | 1.35E-06 | HG01573_active | 0 | 0 |
| 851 | chr4 | 102500329 | 102502329 | ENSG00000 | NFKB1 | 4790 | 29.62314989 | 0.3218469205 | 0.06227732‹ | 5.16796335‹ | 2.37E-07 | 2.44E-05 | HG01573_active | 0 | 0 |
| 852 | chr19 | 35318261 | 35320261 | ENSG00000 | CD22 | 933 | 8.622449613 | 0.3223897871 | 0.12479018‹ | 2.58345472 | 0.00978163€ | 0.099917711 | HG01573_active | 0 | 0 |
| 853 | chr1 | 23912362 | 23914362 | ENSG00000 | CNR2 | 1269 | 12.48090139 | 0.3229717524 | 0.10024688€ | 3.22176342‹ | 0.00127404‹ | 0.02466149€ | HG01573_active | 0 | 0 |
| 854 | chr1 | 74732408 | 74734408 | ENSG00000 | CRYZ | 1429 | 7.506476751 | 0.3232284421 | 0.12481003€ | 2.58976317€ | 0.00960419€ | 0.09901364€ | HG01573_active | 0 | 0 |
| 855 | chr15 | 78076808 | 78078808 | ENSG00000 | SH2D7 | 646892 | 9.463662443 | 0.3239729149 | 0.12075757€ | 2.68283710€ | 0.00730005€ | 0.08237839€ | HG01573_active | 0 | 0 |
| 856 | chr13 | 52449628 | 52451628 | ENSG00000 | VPS36 | 51028 | 7.500272789 | 0.3244572597 | 0.12402243€ | 2.61611751€ | 0.00889359‹ | 0.09437339€ | HG01573_active | 0 | 0 |
| 857 | chr1 | 15525534 | 15527534 | ENSG00000 | CASP9 | 842 | 11.42421823 | 0.325740334 | 0.10212721€ | 3.18955461‹ | 0.00142492‹ | 0.02677691‹ | HG01573_active | 0 | 0 |
| 858 | chr5 | 176364468 | 176366468 | ENSG00000 | ARL10 | 285598 | 12.95416552 | 0.3261895426 | 0.10173334‹ | 3.20631895 | 0.00134444€ | 0.02569839€ | HG01573_active | 0 | 0 |
| 859 | chr3 | 197736179 | 197738179 | ENSG00000 | FYTTD1 | 84248 | 11.62460386 | 0.3266101364 | 0.10680570€ | 3.05798396‹ | 0.00222831‹ | 0.03699452′ | HG01573_active | 0 | 0 |
| 860 | chr2 | 112540915 | 112542915 | ENSG00000 | POLR1B | 84172 | 8.86184173 | 0.3272486112 | 0.117822201 | 2.77747832€ | 0.00547824€ | 0.06777304€ | HG01573_active | 0 | 0 |
| 861 | chr19 | 8205736 | 8207736 | ENSG00000 | CERS4 | 79603 | 16.06627675 | 0.3282908019 | 0.08977722€ | 3.65672697 | 0.00025545€ | 0.00751795‹ | HG01573_active | 0 | 0 |
| 862 | chr12 | 93441262 | 93443262 | ENSG00000 | UBE2N | 7334 | 7.64895767 | 0.3287104659 | 0.12301243€ | 2.67217262‹ | 0.00753618€ | 0.08426024€ | HG01573_active | 0 | 0 |
| 863 | chr5 | 176541511 | 176543511 | ENSG00000 | CDHR2 | 54825 | 18.34056032 | 0.3297840867 | 0.08312977‹ | 3.9670995 | 7.28E-05 | 0.00273047′ | HG01573_active | 0 | 0 |
| 864 | chr16 | 89087375 | 89089375 | ENSG00000 | ACSF3 | 197322 | 23.31163973 | 0.3301158523 | 0.08305758€ | 3.97454179‹ | 7.05E-05 | 0.00266143€ | HG01573_active | 0 | 0 |
| 865 | chr12 | 128822983 | 128824983 | ENSG00000 | SLC15A4 | 121260 | 8.12205762 | 0.3303039528 | 0.12309194€ | 2.68339209€ | 0.00728794€ | 0.08237839€ | HG01573_active | 0 | 0 |
| 866 | chr2 | 231482641 | 231484641 | ENSG00000 | NCL | 4691 | 8.358469109 | 0.3311054163 | 0.12776001€ | 2.59161998€ | 0.00955252‹ | 0.09863311€ | HG01573_active | 0 | 0 |
| 867 | chr11 | 93542508 | 93544508 | ENSG00000 | SMCO4 | 56935 | 21.84885912 | 0.331175948 | 0.07773799‹ | 4.26015552′ | 2.04E-05 | 0.00095102€ | HG01573_active | 0 | 0 |
| 868 | chr19 | 41882161 | 41884161 | ENSG00000 | ARHGEF1 | 9138 | 13.48514458 | 0.3312509062 | 0.10613416€ | 3.12105809‹ | 0.00180202‹ | 0.03168007€ | HG01573_active | 0 | 0 |
| 869 | chr15 | 66385746 | 66387746 | ENSG00000 | TIPIN | 54962 | 10.32537962 | 0.3318365338 | 0.10226501€ | 3.24486861€ | 0.00117504€ | 0.02312197‹ | HG01573_active | 0 | 0 |
| 870 | chr12 | 123711318 | 123713318 | ENSG00000 | ATP6V0A2 | 23545 | 13.37998616 | 0.3324935444 | 0.10068401€ | 3.30234694€ | 0.00095879€ | 0.02011059€ | HG01573_active | 0 | 0 |
| 871 | chr3 | 98592723 | 98594723 | ENSG00000 | CPOX | 1371 | 16.54599379 | 0.3340335808 | 0.09247020€ | 3.61233737€ | 0.00030344€ | 0.00868177‹ | HG01573_active | 0 | 0 |
| 872 | chr14 | 77376109 | 77378109 | ENSG00000 | TMED8 | 283578 | 10.54836175 | 0.3351705556 | 0.12237471€ | 2.73888723€ | 0.00616475′ | 0.07307264‹ | HG01573_active | 0 | 0 |
| 873 | chr16 | 11975643 | 11977643 | ENSG00000 | RP11-166B2 | 729978 | 10.82316285 | 0.3352001332 | 0.111946637 | 2.99428495€ | 0.00275088€ | 0.04303820€ | HG01573_active | 0 | 0 |
| 874 | chr17 | 48829988 | 48831988 | ENSG00000 | CALCOCO2 | 10241 | 9.717695185 | 0.3356168142 | 0.110425651 | 3.039301197 | 0.00237127€ | 0.03840318‹ | HG01573_active | 0 | 0 |
| 875 | chr19 | 54199742 | 54201742 | ENSG00000 | RPS9 | 6203 | 12.95132011 | 0.335685445 | 0.09575641‹ | 3.50561844‹ | 0.00045554€ | 0.01205262′ | HG01573_active | 0 | 0 |
| 876 | chr11 | 1388899 | 1390899 | ENSG00000 | BRSK2 | 9024 | 23.88808811 | 0.3370076244 | 0.08633224€ | 3.90361227 | 9.48E-05 | 0.00337650‹ | HG01573_active | 0 | 0 |
| 877 | chr2 | 96759902 | 96761902 | ENSG00000 | CNNM4 | 26504 | 15.70505777 | 0.3371910075 | 0.08592033€ | 3.924461111 | 8.69E-05 | 0.00315595€ | HG01573_active | 0 | 0 |
| 878 | chr8 | 42795392 | 42797392 | ENSG00000 | CHRNA6 | 8973 | 12.0912559 | 0.3373292994 | 0.11017436€ | 3.06177657′ | 0.00220027€ | 0.03665572€ | HG01573_active | 0 | 0 |
| 879 | chr16 | 1883294 | 1885294 | ENSG00000 | MEIOB | 254528 | 11.57627671 | 0.3397089899 | 0.10451785€ | 3.25024831‹ | 0.001153042 | 0.02283480€ | HG01573_active | 0 | 0 |
| 880 | chr7 | 149027641 | 149029641 | ENSG00000 | PDIA4 | 9601 | 8.020777521 | 0.34104806 | 0.12422855€ | 2.74532734€ | 0.00604505€ | 0.07237276‹ | HG01573_active | 0 | 0 |
| 881 | chr16 | 642262 | 644262 | ENSG00000 | MSLN | 10232 | 34.97674677 | 0.3416116617 | 0.07252273‹ | 4.71040792′ | 2.47E-06 | 0.00017026‹ | HG01573_active | 0 | 0 |
| 882 | chr19 | 45000430 | 45002430 | ENSG00000 | RELB | 5971 | 9.283525419 | 0.3418794911 | 0.111934336 | 3.05428612€ | 0.00225596€ | 0.037230817 | HG01573_active | 0 | 0 |
| 883 | chr19 | 1940149 | 1942149 | ENSG00000 | CSNK1G2 | 1455 | 10.70326989 | 0.3424325647 | 0.11715724€ | 2.92284580′ | 0.00346848‹ | 0.050814027 | HG01573_active | 0 | 0 |
| 884 | chr15 | 64774587 | 64776587 | ENSG00000 | RBPMS2 | 348093 | 11.74819815 | 0.3425457134 | 0.10763269€ | 3.18254343€ | 0.00145987€ | 0.02718233€ | HG01573_active | 0 | 0 |
| 885 | chr22 | 42720298 | 42722298 | ENSG00000 | A4GALT | 53947 | 8.366088314 | 0.3448349642 | 0.12676638‹ | 2.72023981 | 0.00652345€ | 0.076122211 | HG01573_active | 0 | 0 |
| 886 | chr19 | 49928539 | 49930539 | ENSG00000 | IL4I1 | 259307 | 11.1547748 | 0.3453866514 | 0.10643197€ | 3.24513992‹ | 0.00117392€ | 0.02312197‹ | HG01573_active | 0 | 0 |
| 887 | chr17 | 68456513 | 68458513 | ENSG00000 | WIPI1 | 55062 | 12.03256388 | 0.3458464996 | 0.10530321‹ | 3.28429201‹ | 0.00102238€ | 0.02101560€ | HG01573_active | 0 | 0 |
| 888 | chr1 | 32250239 | 32252239 | ENSG00000 | LCK | 3932 | 9.989472215 | 0.346337241 | 0.10856060€ | 3.19026622€ | 0.00142141€ | 0.02674868′ | HG01573_active | 0 | 0 |
| 889 | chr18 | 26132852 | 26134852 | ENSG00000 | PSMA8 | 143471 | 13.7941248 | 0.3466162294 | 0.09562688€ | 3.62467334€ | 0.00028932€ | 0.00834923€ | HG01573_active | 0 | 0 |
| 890 | chr7 | 100366530 | 100368530 | ENSG00000 | PILRA | 29992 | 8.928489046 | 0.3480846536 | 0.10893982€ | 3.19520126€ | 0.00139733€ | 0.02640908€ | HG01573_active | 0 | 0 |
| 891 | chr19 | 3184563 | 3186563 | ENSG00000 | NCLN | 56926 | 7.365276339 | 0.3481438041 | 0.13474878€ | 2.583650711 | 0.00977607‹ | 0.099917711 | HG01573_active | 0 | 0 |
| 892 | chr1 | 113811476 | 113813476 | ENSG00000 | RSBN1 | 54665 | 10.94925605 | 0.3489014354 | 0.10634443€ | 3.28086203€ | 0.00103490€ | 0.021110451 | HG01573_active | 0 | 0 |
| 893 | chr7 | 726650 | 728650 | ENSG00000 | PRKAR1B | 5575 | 54.88335121 | 0.3489475403 | 0.06081903′ | 5.73747284‹ | 9.61E-09 | 1.44E-06 | HG01573_active | 0 | 0 |
| 894 | chr19 | 7379964 | 7381964 | ENSG00000 | CTD-2207O‹ | NaN | 29.96533537 | 0.3492962138 | 0.07234133€ | 4.82844548€ | 1.38E-06 | 0.00010505€ | HG01573_active | 0 | 0 |
| 895 | chr8 | 55878813 | 55880813 | ENSG00000 | LYN | 4067 | 36.30784318 | 0.3495651152 | 0.05812976€ | 6.01353013‹ | 1.82E-09 | 3.32E-07 | HG01573_active | 0 | 0 |

| | | | | | | | | | | | | | | | |
|---|---|---|---|---|---|---|---|---|---|---|---|---|---|---|---|
| 896 | chr22 | 41997725 | 41999725 | ENSG00000 | WBP2NL | 164684 | 14.32491412 | 0.3498459144 | 0.094650715 | 3.696178226 | 0.000218869 | 0.006676514 | HG01573_active | 0 | 0 |
| 897 | chr19 | 41548518 | 41550518 | ENSG00000 | CEACAM21 | 90273 | 11.7197541 | 0.350498971 | 0.100707417 | 3.480368989 | 0.000500723 | 0.012890498 | HG01573_active | 0 | 0 |
| 898 | chr2 | 219308648 | 219310648 | ENSG00000 | PTPRN | 5798 | 7.311465357 | 0.3524609075 | 0.128261832 | 2.747979665 | 0.005996372 | 0.071897556 | HG01573_active | 0 | 0 |
| 899 | chr2 | 113156325 | 113158325 | ENSG00000 | PSD4 | 23550 | 18.02379564 | 0.352565438 | 0.084696306 | 4.162701431 | 3.15E-05 | 0.001342522 | HG01573_active | 0 | 0 |
| 900 | chr19 | 55070291 | 55072291 | ENSG00000 | RDH13 | 112724 | 9.845738175 | 0.3539124589 | 0.123983673 | 2.854508566 | 0.004310346 | 0.058467558 | HG01573_active | 0 | 0 |
| 901 | chr19 | 40347456 | 40349456 | ENSG00000 | PLD3 | 23646 | 9.718647739 | 0.3541211836 | 0.126411332 | 2.801340482 | 0.005089079 | 0.064757323 | HG01573_active | 0 | 0 |
| 902 | chr1 | 212949520 | 212951520 | ENSG00000 | VASH2 | 79805 | 14.83699463 | 0.3545288155 | 0.094878835 | 3.736648052 | 0.000186489 | 0.005835335 | HG01573_active | 0 | 0 |
| 903 | chr6 | 36386800 | 36388800 | ENSG00000 | ETV7 | 51513 | 10.3457705 | 0.3548307808 | 0.108469860 | 3.271238472 | 0.001070775 | 0.021660073 | HG01573_active | 0 | 0 |
| 904 | chr19 | 10415773 | 10417773 | ENSG00000 | PDE4A | 5141 | 15.14965453 | 0.3553547804 | 0.100118826 | 3.54933027 | 0.000386212 | 0.010661535 | HG01573_active | 0 | 0 |
| 905 | chr20 | 3731667 | 3733667 | ENSG00000 | HSPA12B | 116835 | 7.290622597 | 0.3557224682 | 0.129712183 | 2.742398279 | 0.006099233 | 0.072695684 | HG01573_active | 0 | 0 |
| 906 | chr11 | 72813308 | 72815308 | ENSG00000 | ATG16L2 | 89849 | 9.227966279 | 0.3561952306 | 0.127235481 | 2.799496072 | 0.005118243 | 0.064954564 | HG01573_active | 0 | 0 |
| 907 | chr17 | 42286547 | 42288547 | ENSG00000 | STAT5A | 6776 | 8.955251781 | 0.3564862816 | 0.124360587 | 2.866553528 | 0.004149679 | 0.057323621 | HG01573_active | 0 | 0 |
| 908 | chr17 | 45220444 | 45222444 | ENSG00000 | FMNL1 | 752 | 8.322038697 | 0.356890217 | 0.133812015 | 2.667101432 | 0.007650857 | 0.085328138 | HG01573_active | 0 | 0 |
| 909 | chr8 | 7384558 | 7386558 | ENSG00000 | ZNF705G | 100131980 | 8.08558057 | 0.3571083264 | 0.130790469 | 2.730384932 | 0.006326041 | 0.074534601 | HG01573_active | 0 | 0 |
| 910 | chr21 | 32611866 | 32613866 | ENSG00000 | C21orf59 | 56683 | 6.444186786 | 0.3590197217 | 0.137124043 | 2.618211317 | 0.008839204 | 0.094104075 | HG01573_active | 0 | 0 |
| 911 | chr8 | 80230232 | 80232232 | ENSG00000 | TPD52 | 7163 | 51.2987434 | 0.3592414653 | 0.050060568 | 7.176136303 | 7.17E-13 | 2.46E-10 | HG01573_active | 0 | 0 |
| 912 | chr10 | 27153824 | 27155824 | ENSG00000 | MASTL | 84930 | 7.400482585 | 0.3610274824 | 0.125666554 | 2.872900297 | 0.004067224 | 0.056430100 | HG01573_active | 0 | 0 |
| 913 | chr19 | 34358480 | 34360480 | ENSG00000 | GPI | 2821 | 21.35480667 | 0.3612927577 | 0.091290041 | 3.957636028 | 7.57E-05 | 0.002825037 | HG01573_active | 0 | 0 |
| 914 | chr19 | 40717207 | 40719207 | ENSG00000 | ADCK4 | 79934 | 8.863485428 | 0.3622962189 | 0.122284923 | 2.962721891 | 0.003049319 | 0.046149376 | HG01573_active | 0 | 0 |
| 915 | chr5 | 1522977 | 1524977 | ENSG00000 | LPCAT1 | 79888 | 20.36370203 | 0.3631380209 | 0.092393616 | 3.930336704 | 8.48E-05 | 0.003096648 | HG01573_active | 0 | 0 |
| 916 | chr5 | 1294069 | 1296069 | ENSG00000 | TERT | 7015 | 14.86695157 | 0.3635883578 | 0.105119527 | 3.458808915 | 0.000542569 | 0.013550044 | HG01573_active | 0 | 0 |
| 917 | chr1 | 220785759 | 220787759 | ENSG00000 | 36951 | 64757 | 11.30882289 | 0.3644300098 | 0.116344934 | 3.132323831 | 0.001734284 | 0.030989448 | HG01573_active | 0 | 0 |
| 918 | chr19 | 12091263 | 12093263 | ENSG00000 | ZNF788 | NaN | 12.55323634 | 0.3646018356 | 0.100429307 | 3.63043264 | 0.000282946 | 0.008218366 | HG01573_active | 0 | 0 |
| 919 | chr10 | 8052604 | 8054604 | ENSG00000 | GATA3 | 2625 | 6.422303762 | 0.364767667 | 0.129834934 | 2.809472427 | 0.004962276 | 0.063830943 | HG01573_active | 0 | 0 |
| 920 | chr12 | 31646160 | 31648160 | ENSG00000 | METTL20 | 254013 | 6.443179727 | 0.3652530392 | 0.140154036 | 2.606082915 | 0.009158428 | 0.096275191 | HG01573_active | 0 | 0 |
| 921 | chr9 | 133148334 | 133150334 | ENSG00000 | RALGDS | 5900 | 15.61876023 | 0.3655266668 | 0.089256934 | 4.095218679 | 4.22E-05 | 0.001723324 | HG01573_active | 0 | 0 |
| 922 | chr1 | 150486364 | 150488364 | ENSG00000 | TARS2 | 80222 | 6.105661749 | 0.3657616735 | 0.141218605 | 2.590038846 | 0.009596510 | 0.099010790 | HG01573_active | 0 | 0 |
| 923 | chr22 | 35399063 | 35401063 | ENSG00000 | MCM5 | 4174 | 8.463921959 | 0.3660099088 | 0.130026496 | 2.814887102 | 0.004879437 | 0.063233909 | HG01573_active | 0 | 0 |
| 924 | chr3 | 196210487 | 196212487 | ENSG00000 | SLC51A | 200931 | 11.17844568 | 0.3673171512 | 0.108030725 | 3.400117418 | 0.000673569 | 0.015530189 | HG01573_active | 0 | 0 |
| 925 | chr22 | 38109670 | 38111670 | ENSG00000 | BAIAP2L2 | 80115 | 8.263993923 | 0.3677090186 | 0.129311688 | 2.843586857 | 0.004460884 | 0.059661544 | HG01573_active | 0 | 0 |
| 926 | chr5 | 157141869 | 157143869 | ENSG00000 | HAVCR2 | 84868 | 15.53880567 | 0.3686696948 | 0.097732905 | 3.772216679 | 0.000161803 | 0.005171910 | HG01573_active | 0 | 0 |
| 927 | chr8 | 141390993 | 141392993 | ENSG00000 | PTP4A3 | 11156 | 15.75546524 | 0.3689813413 | 0.091526836 | 4.031400585 | 5.54E-05 | 0.002142805 | HG01573_active | 0 | 0 |
| 928 | chr7 | 140672153 | 140674153 | ENSG00000 | ADCK2 | 90956 | 7.810338725 | 0.3699339292 | 0.128306703 | 2.88320033 | 0.003936570 | 0.055074881 | HG01573_active | 0 | 0 |
| 929 | chr19 | 12139407 | 12141407 | ENSG00000 | ZNF20 | 7568 | 13.11584543 | 0.3710650107 | 0.096520537 | 3.844415104 | 0.000120840 | 0.004097835 | HG01573_active | 0 | 0 |
| 930 | chr1 | 151709433 | 151711433 | ENSG00000 | RIIAD1 | 284485 | 7.235760411 | 0.37179831 | 0.136355351 | 2.726686609 | 0.006397376 | 0.075044205 | HG01573_active | 0 | 0 |
| 931 | chr19 | 2268525 | 2270525 | ENSG00000 | LLfos-48D6.2 | NaN | 18.62526636 | 0.3733593266 | 0.092703932 | 4.027437866 | 5.64E-05 | 0.002171180 | HG01573_active | 0 | 0 |
| 932 | chr19 | 48512189 | 48514189 | ENSG00000 | LMTK3 | 114783 | 7.071907809 | 0.3742275494 | 0.135216572 | 2.767615991 | 0.005646794 | 0.069344507 | HG01573_active | 0 | 0 |
| 933 | chr22 | 19755703 | 19757703 | ENSG00000 | TBX1 | 6899 | 8.725820139 | 0.3753607753 | 0.131689362 | 2.850350008 | 0.004367114 | 0.058938397 | HG01573_active | 0 | 0 |
| 934 | chr2 | 69828642 | 69830642 | ENSG00000 | GMCL1 | 64395 | 11.05649645 | 0.3766020201 | 0.106883275 | 3.523488763 | 0.000425905 | 0.011542634 | HG01573_active | 0 | 0 |
| 935 | chr17 | 3963464 | 3965464 | ENSG00000 | ATP2A3 | 489 | 13.81128199 | 0.377104525 | 0.096207264 | 3.919709461 | 8.87E-05 | 0.003210108 | HG01573_active | 0 | 0 |
| 936 | chr19 | 19820751 | 19822751 | ENSG00000 | ZNF506 | 440515 | 8.743830992 | 0.3780955767 | 0.119215869 | 3.171520515 | 0.001516431 | 0.027984864 | HG01573_active | 0 | 0 |
| 937 | chr22 | 41975272 | 41977272 | ENSG00000 | 37865 | 55964 | 8.125227141 | 0.3783846085 | 0.121783836 | 3.107018311 | 0.001889846 | 0.032881196 | HG01573_active | 0 | 0 |
| 938 | chr1 | 156858018 | 156860018 | ENSG00000 | INSRR | 3645 | 7.017211815 | 0.3787283711 | 0.128202302 | 2.954146401 | 0.003135352 | 0.046963499 | HG01573_active | 0 | 0 |
| 939 | chr12 | 131827393 | 131829393 | ENSG00000 | MMP17 | 4326 | 7.3643473 | 0.3787740833 | 0.135657191 | 2.792141565 | 0.005236044 | 0.065936651 | HG01573_active | 0 | 0 |
| 940 | chr1 | 9087478 | 9089478 | ENSG00000 | SLC2A5 | 6518 | 15.14531364 | 0.3791293418 | 0.090314960 | 4.197857601 | 2.69E-05 | 0.001184259 | HG01573_active | 0 | 0 |
| 941 | chr17 | 42609623 | 42611623 | ENSG00000 | FAM134C | 162427 | 8.170674575 | 0.3792205809 | 0.128132665 | 2.959593332 | 0.003080453 | 0.046401288 | HG01573_active | 0 | 0 |
| 942 | chr19 | 5902787 | 5904787 | ENSG00000 | AC024592.1 | NaN | 11.08534265 | 0.3794554074 | 0.110230419 | 3.44238376 | 0.000576611 | 0.014007468 | HG01573_active | 0 | 0 |
| 943 | chr19 | 53158075 | 53160075 | ENSG00000 | ZNF347 | 84671 | 8.796157181 | 0.3804816326 | 0.110323460 | 3.448782619 | 0.000563119 | 0.013826358 | HG01573_active | 0 | 0 |
| 944 | chr16 | 30471658 | 30473658 | ENSG00000 | ITGAL | 3683 | 15.03249174 | 0.3807880351 | 0.087560559 | 4.348853367 | 1.37E-05 | 0.000686944 | HG01573_active | 0 | 0 |
| 945 | chr12 | 110613027 | 110615027 | ENSG00000 | TCTN1 | 79600 | 10.15631051 | 0.3825732325 | 0.113140648 | 3.381395072 | 0.000721187 | 0.016359573 | HG01573_active | 0 | 0 |
| 946 | chr5 | 179861173 | 179863173 | ENSG00000 | C5orf45 | 51149 | 7.685485521 | 0.3826634693 | 0.133789571 | 2.860189073 | 0.004233885 | 0.057886547 | HG01573_active | 0 | 0 |
| 947 | chr22 | 41832079 | 41834079 | ENSG00000 | SREBF2 | 6721 | 20.99379573 | 0.3828919553 | 0.080623025 | 4.749163835 | 2.04E-06 | 0.000146726 | HG01573_active | 0 | 0 |
| 948 | chr5 | 6447623 | 6449623 | ENSG00000 | UBE2QL1 | 134111 | 15.62219597 | 0.3832862173 | 0.082201890 | 4.662742132 | 3.12E-06 | 0.000205366 | HG01573_active | 0 | 0 |
| 949 | chr9 | 120928173 | 120930173 | ENSG00000 | TRAF1 | 7185 | 9.241814315 | 0.3833719251 | 0.122285930 | 3.135045245 | 0.001718276 | 0.030774647 | HG01573_active | 0 | 0 |
| 950 | chr17 | 8964712 | 8966712 | ENSG00000 | PIK3R5 | 23533 | 28.80789771 | 0.3843747431 | 0.067128604 | 5.725945647 | 1.03E-08 | 1.53E-06 | HG01573_active | 0 | 0 |
| 951 | chr4 | 40190053 | 40192053 | ENSG00000 | RHOH | 399 | 16.86521993 | 0.385399146 | 0.089056334 | 4.32758823 | 1.51E-05 | 0.000740508 | HG01573_active | 0 | 0 |
| 952 | chr9 | 128919895 | 128921895 | ENSG00000 | PHYHD1 | 254295 | 5.857227541 | 0.3856936689 | 0.149058466 | 2.587532785 | 0.009666599 | 0.099121594 | HG01573_active | 0 | 0 |
| 953 | chr7 | 102362872 | 102364872 | ENSG00000 | PRKRIP1 | 79706 | 15.22699662 | 0.3862666064 | 0.091274945 | 4.231901784 | 2.32E-05 | 0.001049511 | HG01573_active | 0 | 0 |
| 954 | chrX | 1335616 | 1337616 | ENSG00000 | IL3RA | 3563 | 9.688263074 | 0.3867818823 | 0.121066239 | 3.194795544 | 0.001399299 | 0.026409085 | HG01573_active | 0 | 0 |
| 955 | chr15 | 22837641 | 22839641 | ENSG00000 | NIPA2 | 81614 | 6.789781752 | 0.3867976844 | 0.135063123 | 2.863828954 | 0.004185539 | 0.057474814 | HG01573_active | 0 | 0 |

| | | | | | | | | | | | | | | | | |
|---|---|---|---|---|---|---|---|---|---|---|---|---|---|---|---|---|
| 956 | chr14 | 92747702 | 92749702 | ENSG00000 | LGMN | 5641 | 13.75349334 | | 0.3880182757 | 0.09201603 | 4.21685477 | 2.48E-05 | 0.001107012 | HG01573_active | 0 | 0 |
| 957 | chr22 | 39456344 | 39458344 | ENSG00000 | MGAT3 | 4248 | 11.34963048 | | 0.3880642571 | 0.113091492 | 3.43141864 | 0.00060043 | 0.01440285 | HG01573_active | 0 | 0 |
| 958 | chr1 | 38873494 | 38875494 | ENSG00000 | RP5-864K19 | NaN | 8.04397109 | | 0.3883089762 | 0.12416038 | 3.12747873 | 0.00176312 | 0.03127087 | HG01573_active | 0 | 0 |
| 959 | chr22 | 24416879 | 24418879 | ENSG00000 | ADORA2A | 135 | 8.343545527 | | 0.3889361443 | 0.13961220 | 2.78583206 | 0.00533905 | 0.06654391 | HG01573_active | 0 | 0 |
| 960 | chr11 | 64916248 | 64918248 | ENSG00000 | ATG2A | 23130 | 8.291449387 | | 0.389043169 | 0.11728090 | 3.31719103 | 0.00090927 | 0.01945650 | HG01573_active | 0 | 0 |
| 961 | chr12 | 14960728 | 14962728 | ENSG00000 | ARHGDIB | 397 | 5.79428515 | | 0.3891535527 | 0.14500913 | 2.68364853 | 0.00728236 | 0.08237839 | HG01573_active | 0 | 1 |
| 962 | chr2 | 85417432 | 85419432 | ENSG00000 | CAPG | 822 | 6.90052697 | | 0.3894595124 | 0.122331132 | 3.18365003 | 0.00145430 | 0.02713826 | HG01573_active | 0 | 0 |
| 963 | chr22 | 50474024 | 50476024 | ENSG00000 | SBF1 | 6305 | 9.33895984 | | 0.3897211351 | 0.12395942 | 3.14394118 | 0.00166688 | 0.03001524 | HG01573_active | 0 | 0 |
| 964 | chr7 | 74253458 | 74255458 | ENSG00000 | RFC2 | 5982 | 6.18018228 | | 0.3897575908 | 0.14982208 | 2.601469611 | 0.00928252 | 0.09704528 | HG01573_active | 0 | 0 |
| 965 | chr6 | 166956191 | 166958191 | ENSG00000 | RNASET2 | 8635 | 7.932910131 | | 0.3901043196 | 0.13202871 | 2.95469295 | 0.00312980 | 0.04693300 | HG01573_active | 0 | 0 |
| 966 | chr17 | 5077248 | 5079248 | ENSG00000 | ZFP3 | 124961 | 5.48507328 | | 0.3907212844 | 0.14170611 | 2.75726472 | 0.00582871 | 0.07073337 | HG01573_active | 0 | 0 |
| 967 | chr7 | 151340699 | 151342699 | ENSG00000 | NUB1 | 51667 | 9.70842795 | | 0.3915423979 | 0.10943909 | 3.57771976 | 0.00034660 | 0.00977001 | HG01573_active | 0 | 0 |
| 968 | chr9 | 137085927 | 137087927 | ENSG00000 | MAN1B1 | 11253 | 7.74843822 | | 0.3938170059 | 0.14000673 | 2.81284324 | 0.00491055 | 0.06345258 | HG01573_active | 0 | 0 |
| 969 | chr9 | 89497130 | 89499130 | ENSG00000 | SEMA4D | 10507 | 51.2282877 | | 0.3949925549 | 0.05456297 | 7.23920446 | 4.51E-13 | 1.68E-10 | HG01573_active | 0 | 1 |
| 970 | chr21 | 33245774 | 33247774 | ENSG00000 | AP000295.9 | 3455 | 10.6403421 | | 0.3950637142 | 0.10743359 | 3.67728269 | 0.00023573 | 0.00706190 | HG01573_active | 0 | 0 |
| 971 | chr4 | 88283769 | 88285769 | ENSG00000 | PPM1K | 152926 | 6.32325175 | | 0.3952626851 | 0.14805034 | 2.66978563 | 0.00758996 | 0.08478443 | HG01573_active | 0 | 0 |
| 972 | chr7 | 139108719 | 139110719 | ENSG00000 | ZC3HAV1 | 56829 | 17.6273649 | | 0.395268472 | 0.08299704 | 4.76244028 | 1.91E-06 | 0.00013813 | HG01573_active | 0 | 0 |
| 973 | chr12 | 110741939 | 110743939 | ENSG00000 | PPP1CC | 5501 | 5.34224138 | | 0.3960518415 | 0.14955929 | 2.64812597 | 0.00809393 | 0.08849678 | HG01573_active | 0 | 0 |
| 974 | chr2 | 74554690 | 74556690 | ENSG00000 | LOXL3 | 84695 | 7.02489565 | | 0.3970235378 | 0.12850864 | 3.08946949 | 0.00200514 | 0.03425922 | HG01573_active | 0 | 0 |
| 975 | chr1 | 161198056 | 161200056 | ENSG00000 | ADAMTS4 | 9507 | 5.22185638 | | 0.3978592755 | 0.15370456 | 2.58846755 | 0.00964040 | 0.09909493 | HG01573_active | 0 | 0 |
| 976 | chr17 | 80259866 | 80261866 | ENSG00000 | RNF213 | 57674 | 43.1889266 | | 0.3980549303 | 0.05577048 | 7.13737653 | 9.51E-13 | 3.10E-10 | HG01573_active | 0 | 0 |
| 977 | chr12 | 57027883 | 57029883 | ENSG00000 | TAC3 | 6866 | 6.77030570 | | 0.3984169083 | 0.13299683 | 2.99568708 | 0.00273827 | 0.04289104 | HG01573_active | 0 | 0 |
| 978 | chr3 | 196317299 | 196319299 | ENSG00000 | TCTEX1D2 | 255758 | 6.55342967 | | 0.3986956714 | 0.14216444 | 2.80446824 | 0.00503996 | 0.06437760 | HG01573_active | 0 | 0 |
| 979 | chr1 | 17438724 | 17440724 | ENSG00000 | RCC2 | 55920 | 9.333851121 | | 0.3987178864 | 0.11727424 | 3.39987592 | 0.00067416 | 0.01553018 | HG01573_active | 0 | 0 |
| 980 | chr16 | 88976204 | 88978204 | ENSG00000 | CBFA2T3 | 863 | 34.0548185 | | 0.3994151156 | 0.07400986 | 5.39678199 | 6.78E-08 | 8.39E-06 | HG01573_active | 0 | 0 |
| 981 | chr7 | 1137260 | 1139260 | ENSG00000 | C7orf50 | 84310 | 42.2377450 | | 0.3997562306 | 0.069172467 | 5.77912344 | 7.51E-09 | 1.17E-06 | HG01573_active | 0 | 0 |
| 982 | chr20 | 64048836 | 64050836 | ENSG00000 | TCEA2 | 6919 | 6.61956518 | | 0.4004882428 | 0.14099468 | 2.840449317 | 0.00450500 | 0.06007120 | HG01573_active | 0 | 0 |
| 983 | chr1 | 111203535 | 111205535 | ENSG00000 | DENND2D | 79961 | 6.23621851 | | 0.400735408 | 0.14566325 | 2.75110844 | 0.00593939 | 0.07151209 | HG01573_active | 0 | 1 |
| 984 | chr17 | 39636065 | 39638065 | ENSG00000 | STARD3 | 10948 | 9.15141388 | | 0.4019612521 | 0.14149680 | 2.84077961 | 0.00450034 | 0.06006897 | HG01573_active | 0 | 0 |
| 985 | chr19 | 19899913 | 19901913 | ENSG00000 | ZNF93 | 81931 | 8.21936854 | | 0.4027489435 | 0.120423827 | 3.34442903 | 0.00082452 | 0.01805971 | HG01573_active | 0 | 0 |
| 986 | chr9 | 110336884 | 110338884 | ENSG00000 | TXNDC8 | 255220 | 9.82754087 | | 0.4034297016 | 0.112410004 | 3.58891277 | 0.00033205 | 0.00943968 | HG01573_active | 0 | 0 |
| 987 | chr2 | 191019960 | 191021960 | ENSG00000 | STAT1 | 6772 | 16.6251408 | | 0.4036221937 | 0.08599377 | 4.693621115 | 2.68E-06 | 0.00018112 | HG01573_active | 0 | 0 |
| 988 | chr21 | 33542491 | 33544491 | ENSG00000 | GART | 2618 | 9.37160338 | | 0.4037675416 | 0.10930939 | 3.693804511 | 0.00022092 | 0.00672382 | HG01573_active | 0 | 0 |
| 989 | chr19 | 38425305 | 38427305 | ENSG00000 | RASGRP4 | 115727 | 6.43879042 | | 0.4038589232 | 0.14287505 | 2.82665797 | 0.00470365 | 0.06155290 | HG01573_active | 0 | 0 |
| 990 | chr12 | 53989624 | 53991624 | ENSG00000 | HOXC6 | 3223 | 11.88879441 | | 0.4039861518 | 0.10639573 | 3.79701465 | 0.00014644 | 0.00481617 | HG01573_active | 0 | 0 |
| 991 | chr17 | 16971118 | 16973118 | ENSG00000 | TNFRSF13B | 23495 | 12.6972450 | | 0.4040691581 | 0.10331718 | 3.91095774 | 9.19E-05 | 0.00331074 | HG01573_active | 0 | 0 |
| 992 | chr3 | 150601875 | 150603875 | ENSG00000 | SELT | 51714 | 5.34442420 | | 0.4041669202 | 0.15206053 | 2.65793442 | 0.00786211 | 0.08703046 | HG01573_active | 0 | 0 |
| 993 | chr11 | 9663077 | 9665077 | ENSG00000 | SWAP70 | 23075 | 18.6192390 | | 0.4071614596 | 0.08092039 | 5.03162968 | 4.86E-07 | 4.51E-05 | HG01573_active | 0 | 0 |
| 994 | chr9 | 133413358 | 133415358 | ENSG00000 | ADAMTS13 | 11093 | 14.3852531 | | 0.4093639402 | 0.09413091 | 4.34887898 | 1.37E-05 | 0.00068694 | HG01573_active | 0 | 0 |
| 995 | chr19 | 588893 | 590893 | ENSG00000 | HCN2 | 610 | 6.929004807 | | 0.4099630632 | 0.14973060 | 2.73800438 | 0.00618132 | 0.07315206 | HG01573_active | 0 | 0 |
| 996 | chr5 | 112860188 | 112862188 | ENSG00000 | CTC-487M2 | NaN | 7.88893136 | | 0.4103475489 | 0.13385027 | 3.06572066 | 0.00217146 | 0.03635700 | HG01573_active | 0 | 0 |
| 997 | chr7 | 120949749 | 120951749 | ENSG00000 | ING3 | 54556 | 5.308344417 | | 0.4104603466 | 0.15108940 | 2.71667190 | 0.00659419 | 0.07688047 | HG01573_active | 0 | 0 |
| 998 | chr20 | 2839703 | 2841703 | ENSG00000 | VPS16 | 64601 | 8.723712897 | | 0.4108921862 | 0.12732491 | 3.22711532 | 0.00125045 | 0.02439017 | HG01573_active | 0 | 0 |
| 999 | chr4 | 336814 | 338814 | ENSG00000 | ZNF141 | 7700 | 10.8340565 | | 0.4130392277 | 0.10972672 | 3.764253677 | 0.000167047 | 0.00529945 | HG01573_active | 0 | 0 |
| 1000 | chr19 | 48393875 | 48395875 | ENSG00000 | GRIN2D | 2906 | 12.4771085 | | 0.4133567075 | 0.111981486 | 3.69129505 | 0.00022311 | 0.00677509 | HG01573_active | 0 | 0 |
| 1001 | chrX | 119692370 | 119694370 | ENSG00000 | 38961 | 23157 | 23.9148681 | | 0.4133865883 | 0.08190851 | 5.04693070 | 4.49E-07 | 4.28E-05 | HG01573_active | 0 | 0 |
| 1002 | chr19 | 7594902 | 7596902 | ENSG00000 | CAMSAP3 | 57662 | 5.94931768 | | 0.4134251551 | 0.15742064 | 2.626244827 | 0.00863326 | 0.09257553 | HG01573_active | 0 | 0 |
| 1003 | chr8 | 144011772 | 144013772 | ENSG00000 | PARP10 | 84875 | 10.7660555 | | 0.4134747605 | 0.11431830 | 3.616872697 | 0.00029818 | 0.00856781 | HG01573_active | 0 | 0 |
| 1004 | chr22 | 50084902 | 50086902 | ENSG00000 | MLC1 | 23209 | 8.77298269 | | 0.4148702252 | 0.12529956 | 3.31102677 | 0.00092954 | 0.01971369 | HG01573_active | 0 | 0 |
| 1005 | chr9 | 5449503 | 5451503 | ENSG00000 | CD274 | 29126 | 5.03260881 | | 0.4149111892 | 0.15341135 | 2.70456627 | 0.00683936 | 0.07850577 | HG01573_active | 0 | 0 |
| 1006 | chr15 | 22772063 | 22774063 | ENSG00000 | NIPA1 | 123606 | 14.1970425 | | 0.4180773974 | 0.09530240 | 4.38685052 | 1.15E-05 | 0.00060733 | HG01573_active | 0 | 0 |
| 1007 | chr15 | 40564057 | 40566057 | ENSG00000 | C15orf57 | 90416 | 10.8636464 | | 0.419492562 | 0.117362487 | 3.574332567 | 0.00035112 | 0.00986135 | HG01573_active | 0 | 0 |
| 1008 | chr20 | 63707864 | 63709864 | ENSG00000 | RP4-583P15 | NaN | 7.549596967 | | 0.4214418791 | 0.13983357 | 3.01388195 | 0.00257928 | 0.04087992 | HG01573_active | 0 | 0 |
| 1009 | chr22 | 39013363 | 39015363 | ENSG00000 | APOBEC3D | 140564 | 5.53665815 | | 0.4215032702 | 0.15857603 | 2.65805150 | 0.00785938 | 0.08703046 | HG01573_active | 0 | 0 |
| 1010 | chr22 | 37485401 | 37487401 | ENSG00000 | MFNG | 4242 | 6.27841928 | | 0.421770292 | 0.15992131 | 2.63736136 | 0.00835537 | 0.09064790 | HG01573_active | 0 | 0 |
| 1011 | chr11 | 86243544 | 86245544 | ENSG00000 | EED | 8726 | 6.44726801 | | 0.4238051845 | 0.13542137 | 3.12952953 | 0.00175086 | 0.03114953 | HG01573_active | 0 | 0 |
| 1012 | chr19 | 40777216 | 40779216 | ENSG00000 | RAB4B | 53916 | 5.54642751 | | 0.4255857289 | 0.15826675 | 2.68904057 | 0.00716577 | 0.08136679 | HG01573_active | 0 | 0 |
| 1013 | chr12 | 53984845 | 53986845 | ENSG00000 | RP11-834C1 | NaN | 14.5865335 | | 0.4280130951 | 0.09510203 | 4.50056717 | 6.78E-06 | 0.00039369 | HG01573_active | 0 | 0 |
| 1014 | chr16 | 10817794 | 10819794 | ENSG00000 | TVP23A | 780776 | 17.1993028 | | 0.428536282 | 0.08701910 | 4.92462267 | 8.45E-07 | 7.16E-05 | HG01573_active | 0 | 0 |
| 1015 | chr1 | 160795074 | 160797074 | ENSG00000 | LY9 | 4063 | 11.9489810 | | 0.4306336745 | 0.10399322 | 4.14097833 | 3.46E-05 | 0.00143944 | HG01573_active | 0 | 0 |

| | | | | | | | | | | | | | | | | |
|---|---|---|---|---|---|---|---|---|---|---|---|---|---|---|---|---|
| 1016 | chr19 | 18096793 | 18098793 | ENSG00000 | MAST3 | 23031 | 15.69133483 | | 0.4317696505 | 0.09171084 | 4.70794534 | 2.50E-06 | 0.00017144 | HG01573_active | 0 | 0 |
| 1017 | chr12 | 54015931 | 54017931 | ENSG00000 | HOXC4 | 3221 | 12.51713484 | | 0.4319098243 | 0.10651612 | 4.05487736 | 5.02E-05 | 0.00198282 | HG01573_active | 0 | 0 |
| 1018 | chr19 | 57934393 | 57936393 | ENSG00000 | ZNF418 | 147686 | 4.80968964 | | 0.4323177712 | 0.16518424 | 2.61718519 | 0.00886582 | 0.09423727 | HG01573_active | 0 | 0 |
| 1019 | chr19 | 58511413 | 58513413 | ENSG00000 | SLC27A5 | 10998 | 9.99015683 | | 0.4324451331 | 0.11228533 | 3.85130538 | 0.00011748 | 0.00402508 | HG01573_active | 0 | 0 |
| 1020 | chr16 | 85897803 | 85899803 | ENSG00000 | IRF8 | 3394 | 8.12636197 | | 0.4327152373 | 0.11937497 | 3.62484029 | 0.00028914 | 0.00834923 | HG01573_active | 0 | 0 |
| 1021 | chr12 | 93376883 | 93378883 | ENSG00000 | NUDT4 | 11163 | 6.64733773 | | 0.4327631447 | 0.13048869 | 3.31647999 | 0.00091159 | 0.01945650 | HG01573_active | 0 | 0 |
| 1022 | chr14 | 24150218 | 24152218 | ENSG00000 | IRF9 | NaN | 5.00171075 | | 0.4328686837 | 0.15502115 | 2.79232009 | 0.00523315 | 0.06593665 | HG01573_active | 0 | 0 |
| 1023 | chr19 | 45405349 | 45407349 | ENSG00000 | PPP1R13L | 10848 | 6.19836013 | | 0.4330339929 | 0.14553133 | 2.97553787 | 0.00292475 | 0.04491679 | HG01573_active | 0 | 0 |
| 1024 | chr1 | 109667022 | 109669022 | ENSG00000 | GSTM2 | 2946 | 8.75688551 | | 0.433863355 | 0.13579120 | 3.19507705 | 0.00139793 | 0.02640908 | HG01573_active | 0 | 0 |
| 1025 | chr3 | 183162839 | 183164839 | ENSG00000 | LAMP3 | 27074 | 13.02583779 | | 0.4353036509 | 0.10139894 | 4.29298005 | 1.76E-05 | 0.00083230 | HG01573_active | 0 | 0 |
| 1026 | chr6 | 33079775 | 33081775 | ENSG00000 | HLA-DPA1 | 3113 | 5.72550844 | | 0.4354151086 | 0.15555364 | 2.79913153 | 0.00512402 | 0.06495456 | HG01573_active | 0 | 0 |
| 1027 | chr1 | 12062377 | 12064377 | ENSG00000 | TNFRSF8 | 943 | 24.00265367 | | 0.4361684555 | 0.07983406 | 5.46343765 | 4.67E-08 | 5.94E-06 | HG01573_active | 0 | 0 |
| 1028 | chr12 | 6450690 | 6452690 | ENSG00000 | TAPBPL | 55080 | 4.94475029 | | 0.4364095995 | 0.16692425 | 2.61441696 | 0.00893798 | 0.09462794 | HG01573_active | 0 | 0 |
| 1029 | chr3 | 196502768 | 196504768 | ENSG00000 | RNF168 | 165918 | 7.75994974 | | 0.4378689751 | 0.13615227 | 3.21602393 | 0.00129979 | 0.02502781 | HG01573_active | 0 | 0 |
| 1030 | chr17 | 81702371 | 81704371 | ENSG00000 | SLC25A10 | NaN | 4.48197929 | | 0.4387696021 | 0.16205970 | 2.70745650 | 0.00678009 | 0.07809384 | HG01573_active | 0 | 0 |
| 1031 | chr20 | 4822645 | 4824645 | ENSG00000 | RASSF2 | 9770 | 14.82354151 | | 0.4391955886 | 0.09072592 | 4.84090517 | 1.29E-06 | 0.00010040 | HG01573_active | 0 | 0 |
| 1032 | chr11 | 32828943 | 32830943 | ENSG00000 | PRRG4 | 79056 | 6.62827832 | | 0.4398459646 | 0.13188000 | 3.33519826 | 0.00085238 | 0.01848819 | HG01573_active | 0 | 0 |
| 1033 | chr14 | 91416844 | 91418844 | ENSG00000 | CCDC88C | 440193 | 50.56977002 | | 0.4414479219 | 0.05419775 | 8.14513265 | 3.79E-16 | 2.30E-13 | HG01573_active | 0 | 0 |
| 1034 | chr20 | 5610026 | 5612026 | ENSG00000 | GPCPD1 | 56261 | 14.78314901 | | 0.4420266063 | 0.09763280 | 4.52743936 | 5.97E-06 | 0.00035140 | HG01573_active | 0 | 0 |
| 1035 | chr19 | 16110629 | 16112629 | ENSG00000 | RAB8A | 4218 | 7.12034942 | | 0.4429374512 | 0.14282066 | 3.10135409 | 0.00192637 | 0.03331380 | HG01573_active | 0 | 0 |
| 1036 | chr17 | 44388505 | 44390505 | ENSG00000 | ITGA2B | 3674 | 5.06073822 | | 0.4434279697 | 0.15890456 | 2.79053007 | 0.00526218 | 0.06607894 | HG01573_active | 0 | 0 |
| 1037 | chr1 | 156146366 | 156148366 | ENSG00000 | SEMA4A | 64218 | 9.63467261 | | 0.4449120199 | 0.11070365 | 4.01894617 | 5.85E-05 | 0.00223802 | HG01573_active | 0 | 0 |
| 1038 | chr19 | 38617882 | 38619882 | ENSG00000 | MAP4K1 | 11184 | 8.27420738 | | 0.4471212892 | 0.13114709 | 3.40931135 | 0.00065127 | 0.01521264 | HG01573_active | 0 | 0 |
| 1039 | chr1 | 222588933 | 222590933 | ENSG00000 | TAF1A | 9015 | 7.15768690 | | 0.4480929477 | 0.13773902 | 3.25320258 | 0.00114112 | 0.02268827 | HG01573_active | 0 | 0 |
| 1040 | chr1 | 38858823 | 38860823 | ENSG00000 | RRAGC | 64121 | 4.98565716 | | 0.448432777 | 0.15913534 | 2.81793326 | 0.00483338 | 0.06281990 | HG01573_active | 0 | 0 |
| 1041 | chr22 | 39039961 | 39041961 | ENSG00000 | APOBEC3G | 60489 | 13.33548083 | | 0.4492471616 | 0.09867877 | 4.55262213 | 5.30E-06 | 0.00031886 | HG01573_active | 0 | 0 |
| 1042 | chr22 | 43109748 | 43111748 | ENSG00000 | BIK | 638 | 6.74453849 | | 0.449958793 | 0.14332603 | 3.13940724 | 0.00169289 | 0.03041538 | HG01573_active | 0 | 0 |
| 1043 | chr17 | 39863188 | 39865188 | ENSG00000 | IKZF3 | 22806 | 27.82563838 | | 0.450152987 | 0.07027035 | 6.40601535 | 1.49E-10 | 3.31E-08 | HG01573_active | 0 | 0 |
| 1044 | chr22 | 20964108 | 20966108 | ENSG00000 | AIFM3 | 150209 | 5.36318277 | | 0.4520070666 | 0.16604066 | 2.72226729 | 0.00648356 | 0.07572285 | HG01573_active | 0 | 0 |
| 1045 | chr9 | 136880912 | 136882912 | ENSG00000 | TRAF2 | 7186 | 12.99403544 | | 0.4541427806 | 0.10105438 | 4.49404333 | 6.99E-06 | 0.00040246 | HG01573_active | 0 | 0 |
| 1046 | chr12 | 4272772 | 4274772 | ENSG00000 | CCND2 | 894 | 10.89983773 | | 0.4544293127 | 0.11429792 | 3.97583179 | 7.01E-05 | 0.00265609 | HG01573_active | 0 | 0 |
| 1047 | chr17 | 3915500 | 3917500 | ENSG00000 | P2RX1 | 5023 | 6.63338304 | | 0.4560717754 | 0.13625043 | 3.34730498 | 0.00081601 | 0.01793217 | HG01573_active | 0 | 0 |
| 1048 | chr20 | 62815244 | 62817244 | ENSG00000 | COL9A3 | 1299 | 8.54943401 | | 0.4568873837 | 0.12820699 | 3.56366958 | 0.00036570 | 0.01020083 | HG01573_active | 0 | 0 |
| 1049 | chr19 | 12890026 | 12892026 | ENSG00000 | GCDH | 2639 | 7.05570074 | | 0.4576240593 | 0.14648762 | 3.12397765 | 0.00178424 | 0.03149171 | HG01573_active | 0 | 0 |
| 1050 | chr1 | 52841511 | 52843511 | ENSG00000 | ZYG11A | 440590 | 12.43780544 | | 0.4578871756 | 0.09664873 | 4.73764276 | 2.16E-06 | 0.00015448 | HG01573_active | 0 | 0 |
| 1051 | chr1 | 27634277 | 27636277 | ENSG00000 | FGR | 2268 | 8.01482152 | | 0.4598533115 | 0.13470265 | 3.41383994 | 0.00064054 | 0.01507418 | HG01573_active | 0 | 0 |
| 1052 | chr2 | 218879363 | 218881363 | ENSG00000 | WNT10A | 80326 | 6.54523859 | | 0.4603049747 | 0.14707931 | 3.12963781 | 0.00175021 | 0.03114953 | HG01573_active | 0 | 0 |
| 1053 | chr19 | 1445302 | 1447302 | ENSG00000 | APC2 | 10297 | 6.17265747 | | 0.4605699821 | 0.15077632 | 3.05465724 | 0.00225317 | 0.03723081 | HG01573_active | 0 | 0 |
| 1054 | chr7 | 100048774 | 100050774 | ENSG00000 | ZSCAN21 | 7589 | 4.51260121 | | 0.4607689585 | 0.16110338 | 2.86008236 | 0.00423530 | 0.05788654 | HG01573_active | 0 | 0 |
| 1055 | chr12 | 123435717 | 123437717 | ENSG00000 | RILPL2 | 196383 | 6.67101279 | | 0.4638563214 | 0.14639600 | 3.16850385 | 0.00153225 | 0.02823791 | HG01573_active | 0 | 0 |
| 1056 | chr12 | 53206307 | 53208307 | ENSG00000 | ITGB7 | 3695 | 5.24760715 | | 0.4640104373 | 0.16069310 | 2.88755667 | 0.00388246 | 0.05448911 | HG01573_active | 0 | 0 |
| 1057 | chr19 | 924781 | 926781 | ENSG00000 | ARID3A | 1820 | 12.1029223 | | 0.464804647 | 0.11760973 | 3.95209335 | 7.75E-05 | 0.00286726 | HG01573_active | 0 | 0 |
| 1058 | chr19 | 55037264 | 55039264 | ENSG00000 | GP6 | 51206 | 8.50264652 | | 0.4648978375 | 0.13282178 | 3.50016256 | 0.00046497 | 0.01215769 | HG01573_active | 0 | 0 |
| 1059 | chr19 | 11480046 | 11482046 | ENSG00000 | ELAVL3 | 1995 | 7.82790368 | | 0.4649791806 | 0.13495000 | 3.44556634 | 0.00056986 | 0.01394496 | HG01573_active | 0 | 0 |
| 1060 | chr16 | 89946925 | 89948925 | ENSG00000 | DEF8 | 54849 | 6.66047209 | | 0.4672451196 | 0.14275599 | 3.27303332 | 0.00106399 | 0.02157222 | HG01573_active | 0 | 0 |
| 1061 | chr12 | 110703950 | 110705950 | ENSG00000 | HVCN1 | 84329 | 24.90860778 | | 0.4673626857 | 0.07200187 | 6.49097892 | 8.53E-11 | 1.96E-08 | HG01573_active | 0 | 0 |
| 1062 | chr19 | 56120280 | 56122280 | ENSG00000 | ZNF787 | 126208 | 10.17992232 | | 0.4680560593 | 0.12338974 | 3.79331410 | 0.00014864 | 0.00482394 | HG01573_active | 0 | 0 |
| 1063 | chr16 | 89216703 | 89218703 | ENSG00000 | ZNF778 | 197320 | 6.08345779 | | 0.4691519302 | 0.15136994 | 3.09937314 | 0.00193930 | 0.03342290 | HG01573_active | 0 | 0 |
| 1064 | chr9 | 93057688 | 93059688 | ENSG00000 | SUSD3 | 203328 | 9.09818263 | | 0.4712249831 | 0.12817147 | 3.67651987 | 0.00023643 | 0.00706720 | HG01573_active | 0 | 0 |
| 1065 | chr22 | 18076920 | 18078920 | ENSG00000 | PEX26 | 55670 | 16.40951725 | | 0.4714201236 | 0.08896564 | 5.29890070 | 1.17E-07 | 1.33E-05 | HG01573_active | 0 | 0 |
| 1066 | chr1 | 24964121 | 24966121 | ENSG00000 | RUNX3 | 864 | 23.15232995 | | 0.4731309491 | 0.07504236 | 6.30485137 | 2.88E-10 | 6.02E-08 | HG01573_active | 0 | 0 |
| 1067 | chr2 | 85560562 | 85562562 | ENSG00000 | VAMP8 | 8673 | 5.68861779 | | 0.4738699385 | 0.15514351 | 3.05439728 | 0.00225513 | 0.03723081 | HG01573_active | 0 | 0 |
| 1068 | chr21 | 41419304 | 41421304 | ENSG00000 | MX1 | 4599 | 12.76682906 | | 0.4740328301 | 0.09590887 | 4.9425335 | 7.71E-07 | 6.69E-05 | HG01573_active | 0 | 0 |
| 1069 | chr11 | 118224094 | 118226094 | ENSG00000 | AMICA1 | 120425 | 11.25059931 | | 0.474131952 | 0.11253991 | 4.21301133 | 2.52E-05 | 0.00112227 | HG01573_active | 0 | 0 |
| 1070 | chr1 | 15584110 | 15586110 | ENSG00000 | AGMAT | 79814 | 4.22093448 | | 0.475776287 | 0.16073220 | 2.96005572 | 0.00307583 | 0.04638399 | HG01573_active | 0 | 0 |
| 1071 | chr19 | 50328653 | 50330653 | ENSG00000 | NR1H2 | 7376 | 16.2240665 | | 0.4759244186 | 0.08699014 | 5.47101539 | 4.47E-08 | 5.80E-06 | HG01573_active | 0 | 0 |
| 1072 | chr9 | 130042194 | 130044194 | ENSG00000 | FNBP1 | 23048 | 34.19483874 | | 0.4772487472 | 0.06801122 | 7.01720517 | 2.26E-12 | 6.43E-10 | HG01573_active | 0 | 0 |
| 1073 | chr1 | 1115361 | 1117361 | ENSG00000 | C1orf159 | 54991 | 11.89340451 | | 0.4784405745 | 0.11082029 | 4.31726506 | 1.58E-05 | 0.00076644 | HG01573_active | 0 | 0 |
| 1074 | chr9 | 137137390 | 137139390 | ENSG00000 | GRIN1 | 2902 | 8.66667467 | | 0.4788458648 | 0.13793183 | 3.47161248 | 0.00051734 | 0.01316612 | HG01573_active | 0 | 0 |
| 1075 | chr19 | 19137513 | 19139513 | ENSG00000 | TMEM161A | 54929 | 4.99756732 | | 0.4806814585 | 0.16679869 | 2.88180580 | 0.00395403 | 0.05526134 | HG01573_active | 0 | 0 |

| | | | | | | | | | | | | | | | |
|---|---|---|---|---|---|---|---|---|---|---|---|---|---|---|---|
| 1076 | chr19 | 1039101 | 1041101 | ENSG00000 | ABCA7 | 10347 | 6.260500858 | 0.4835613352 | 0.1434053352 | 3.371989447 | 0.00746273 | 0.01678611 | HG01573_active | 0 | 0 |
| 1077 | chr17 | 8866677 | 8868677 | ENSG00000 | PIK3R6 | 146850 | 20.05563982 | 0.4860338712 | 0.081694413 | 5.94941381 | 2.69E-09 | 4.61E-07 | HG01573_active | 0 | 0 |
| 1078 | chr17 | 38704542 | 38706542 | ENSG00000 | MLLT6 | 4302 | 7.984949936 | 0.4863143584 | 0.129412204 | 3.757870896 | 0.000171365 | 0.00538732 | HG01573_active | 0 | 0 |
| 1079 | chr6 | 167145414 | 167147414 | ENSG00000 | TCP10L2 | 401285 | 14.90684734 | 0.4878843408 | 0.096918907 | 5.033943853 | 4.80E-07 | 4.49E-05 | HG01573_active | 0 | 0 |
| 1080 | chr19 | 35247330 | 35249330 | ENSG00000 | LSR | 51599 | 6.241966781 | 0.4887625958 | 0.139459814 | 3.504684112 | 0.000457149 | 0.01207109 | HG01573_active | 0 | 0 |
| 1081 | chr11 | 57567284 | 57569284 | ENSG00000 | UBE2L6 | 9246 | 5.591401242 | 0.4942984871 | 0.159593251 | 3.097239272 | 0.00195332 | 0.03358858 | HG01573_active | 0 | 0 |
| 1082 | chr15 | 64386687 | 64388687 | ENSG00000 | KIAA0101 | 9768 | 4.604083226 | 0.4956833743 | 0.173319385 | 2.859941913 | 0.00423718 | 0.057886547 | HG01573_active | 0 | 0 |
| 1083 | chr19 | 17746718 | 17748718 | ENSG00000 | FCHO1 | 23149 | 10.81248957 | 0.4958599459 | 0.112939149 | 4.390505403 | 1.13E-05 | 0.00060197 | HG01573_active | 0 | 0 |
| 1084 | chr1 | 6259975 | 6261975 | ENSG00000 | GPR153 | 387509 | 4.778822054 | 0.4961403068 | 0.160951673 | 3.082542084 | 0.002052407 | 0.034932752 | HG01573_active | 0 | 0 |
| 1085 | chr21 | 44284838 | 44286838 | ENSG00000 | AIRE | 326 | 4.212639088 | 0.4986424194 | 0.188323809 | 2.647792764 | 0.008101917 | 0.088511625 | HG01573_active | 0 | 0 |
| 1086 | chr4 | 985895 | 987895 | ENSG00000 | DGKQ | 1609 | 8.40988212 | 0.499328997 | 0.123098317 | 4.056342975 | 4.98E-05 | 0.00197628 | HG01573_active | 0 | 0 |
| 1087 | chr1 | 3795504 | 3797504 | ENSG00000 | LRRC47 | 57470 | 6.397234578 | 0.4999303351 | 0.144480028 | 3.460203749 | 0.00053976 | 0.013534598 | HG01573_active | 0 | 0 |
| 1088 | chr1 | 1172884 | 1174884 | ENSG00000 | TTLL10 | 254173 | 7.913884407 | 0.5007502657 | 0.127928136 | 3.914309068 | 9.07E-05 | 0.00327393 | HG01573_active | 0 | 0 |
| 1089 | chr12 | 8081211 | 8083211 | ENSG00000 | NECAP1 | 25977 | 4.245631775 | 0.5007731021 | 0.192482432 | 2.601656135 | 0.009277482 | 0.09704528 | HG01573_active | 0 | 0 |
| 1090 | chr17 | 49209227 | 49211227 | ENSG00000 | ABI3 | 51225 | 4.582350381 | 0.502462454 | 0.172763807 | 2.908377977 | 0.003633089 | 0.052206393 | HG01573_active | 0 | 0 |
| 1091 | chr19 | 2268759 | 2270759 | ENSG00000 | JSRP1 | 126306 | 4.061958991 | 0.5042625046 | 0.185032584 | 2.725263254 | 0.006425023 | 0.075302403 | HG01573_active | 0 | 0 |
| 1092 | chr1 | 3651520 | 3653520 | ENSG00000 | TP73 | 7161 | 31.51752428 | 0.504743814 | 0.069809636 | 7.230288499 | 4.82E-13 | 1.74E-10 | HG01573_active | 0 | 0 |
| 1093 | chr16 | 3576400 | 3578400 | ENSG00000 | NLRC3 | 197358 | 11.99650212 | 0.5077821514 | 0.107480755 | 4.724400666 | 2.31E-06 | 0.000164024 | HG01573_active | 0 | 0 |
| 1094 | chr11 | 64339223 | 64341223 | ENSG00000 | CCDC88B | 283234 | 5.062238668 | 0.5092877217 | 0.17147741 | 2.969998899 | 0.002978008 | 0.045473333 | HG01573_active | 0 | 0 |
| 1095 | chr1 | 6623866 | 6625866 | ENSG00000 | THAP3 | 90326 | 3.597913751 | 0.5099180555 | 0.19187472 | 2.657557252 | 0.007870922 | 0.08703830 | HG01573_active | 0 | 0 |
| 1096 | chr7 | 151023499 | 151025499 | ENSG00000 | ATG9B | 285973 | 4.160369112 | 0.511182051 | 0.176800349 | 2.891295477 | 0.003836572 | 0.054015217 | HG01573_active | 0 | 0 |
| 1097 | chr19 | 4830704 | 4832704 | ENSG00000 | TICAM1 | 148022 | 4.306893649 | 0.511720698 | 0.176248036 | 2.903412193 | 0.003691204 | 0.052578022 | HG01573_active | 0 | 0 |
| 1098 | chr1 | 247856199 | 247858199 | ENSG00000 | TRIM58 | 25893 | 6.352572773 | 0.5161883778 | 0.151978551 | 3.396455437 | 0.000682646 | 0.015671715 | HG01573_active | 0 | 0 |
| 1099 | chr19 | 45177237 | 45179237 | ENSG00000 | TRAPPC6A | 79090 | 4.998222566 | 0.5164029634 | 0.171500149 | 3.011093369 | 0.002603087 | 0.041159595 | HG01573_active | 0 | 0 |
| 1100 | chr19 | 6481557 | 6483557 | ENSG00000 | DENND1C | 79958 | 4.783045488 | 0.5179251128 | 0.168391712 | 3.075716162 | 0.002099976 | 0.035516184 | HG01573_active | 0 | 0 |
| 1101 | chr2 | 218858821 | 218860821 | ENSG00000 | WNT6 | 7475 | 5.332599963 | 0.5202169816 | 0.158469193 | 3.282764124 | 0.001027946 | 0.021065017 | HG01573_active | 0 | 0 |
| 1102 | chr11 | 60951530 | 60953530 | ENSG00000 | SLC15A3 | 51296 | 5.483976157 | 0.5217630932 | 0.158761084 | 3.286467177 | 0.001014525 | 0.020918335 | HG01573_active | 0 | 0 |
| 1103 | chr19 | 7491976 | 7493976 | ENSG00000 | C19orf45 | 374877 | 4.417375514 | 0.522616525 | 0.175609134 | 2.976021298 | 0.002920145 | 0.04489766 | HG01573_active | 0 | 0 |
| 1104 | chr16 | 29678008 | 29680008 | ENSG00000 | QPRT | 23475 | 5.446211393 | 0.5227367266 | 0.171239095 | 3.052671609 | 0.00226814 | 0.037230817 | HG01573_active | 0 | 0 |
| 1105 | chr6 | 159044152 | 159046152 | ENSG00000 | TAGAP | 117289 | 3.037306608 | 0.5236098914 | 0.202035153 | 2.591677157 | 0.009550935 | 0.098633115 | HG01573_active | 0 | 0 |
| 1106 | chr3 | 169811986 | 169813986 | ENSG00000 | LRRC34 | 151827 | 4.455290224 | 0.5269796002 | 0.17309347 | 3.044479956 | 0.002330829 | 0.037885905 | HG01573_active | 0 | 0 |
| 1107 | chr1 | 206922247 | 206924247 | ENSG00000 | FAIM3 | 9214 | 6.419137301 | 0.5271226933 | 0.144839725 | 3.639351652 | 0.000273325 | 0.00797358 | HG01573_active | 0 | 0 |
| 1108 | chr7 | 151226230 | 151228230 | ENSG00000 | ABCF2 | 10061 | 7.011672849 | 0.5298856526 | 0.138039681 | 3.838647307 | 0.000123713 | 0.004163583 | HG01573_active | 0 | 0 |
| 1109 | chr19 | 49526618 | 49528618 | ENSG00000 | RCN3 | 57333 | 5.517770269 | 0.5310462621 | 0.157366672 | 3.374578965 | 0.000739287 | 0.01665702 | HG01573_active | 0 | 0 |
| 1110 | chr22 | 17083954 | 17085954 | ENSG00000 | IL17RA | 23765 | 10.5661607 | 0.5342534014 | 0.110204421 | 4.847839994 | 1.25E-06 | 9.87E-05 | HG01573_active | 0 | 0 |
| 1111 | chr19 | 18152158 | 18154158 | ENSG00000 | PIK3R2 | NaN | 6.494881401 | 0.5378799725 | 0.139944907 | 3.843512284 | 0.000121285 | 0.004102534 | HG01573_active | 0 | 0 |
| 1112 | chr19 | 17847071 | 17849071 | ENSG00000 | JAK3 | 3718 | 6.580366619 | 0.5386480281 | 0.145230622 | 3.708914971 | 0.000208149 | 0.00639607 | HG01573_active | 0 | 0 |
| 1113 | chr8 | 33566125 | 33568125 | ENSG00000 | RNF122 | 79845 | 6.323958189 | 0.53978455 | 0.156319721 | 3.453080303 | 0.000554223 | 0.013738379 | HG01573_active | 0 | 0 |
| 1114 | chr19 | 632604 | 634604 | ENSG00000 | POLRMT | 5442 | 4.370558381 | 0.5398161359 | 0.186848632 | 2.889055862 | 0.003864004 | 0.054344169 | HG01573_active | 0 | 0 |
| 1115 | chr16 | 29662279 | 29664279 | ENSG00000 | AC009133.2 | NaN | 8.4023747 | 0.5401620524 | 0.13939487 | 3.875049706 | 0.000106602 | 0.00370917 | HG01573_active | 0 | 0 |
| 1116 | chr7 | 155296776 | 155298776 | ENSG00000 | INSIG1 | 3638 | 2.94001449 | 0.5423470573 | 0.208976297 | 2.595256324 | 0.009452039 | 0.098279148 | HG01573_active | 0 | 0 |
| 1117 | chr1 | 27871543 | 27873543 | ENSG00000 | THEMIS2 | 9473 | 4.437363157 | 0.5448554813 | 0.164000221 | 3.322285025 | 0.000892834 | 0.01924058 | HG01573_active | 0 | 0 |
| 1118 | chr7 | 27179013 | 27181013 | ENSG00000 | HOXA10-HC | NaN | 4.695873122 | 0.5454695311 | 0.17867058 | 3.052934238 | 0.002266155 | 0.037230817 | HG01573_active | 0 | 0 |
| 1119 | chr22 | 18109331 | 18111331 | ENSG00000 | TUBA8 | 51807 | 11.41704788 | 0.5468727005 | 0.111218824 | 4.917087599 | 8.78E-07 | 7.34E-05 | HG01573_active | 0 | 0 |
| 1120 | chr11 | 117985348 | 117987348 | ENSG00000 | IL10RA | 3587 | 5.63344254 | 0.5472965554 | 0.16383419 | 3.3405515 | 0.00083612 | 0.01822499 | HG01573_active | 0 | 0 |
| 1121 | chr7 | 974599 | 976599 | ENSG00000 | COX19 | 90639 | 22.87928991 | 0.5497223973 | 0.077497363 | 7.093433549 | 1.31E-12 | 3.95E-10 | HG01573_active | 0 | 0 |
| 1122 | chr16 | 1332601 | 1334601 | ENSG00000 | BAIAP3 | 8938 | 4.751532473 | 0.5514878472 | 0.169778216 | 3.248283893 | 0.001161033 | 0.022947589 | HG01573_active | 0 | 0 |
| 1123 | chr22 | 45283982 | 45285982 | ENSG00000 | UPK3A | 7380 | 3.665412927 | 0.5518734503 | 0.196704311 | 2.805599163 | 0.005022312 | 0.064275016 | HG01573_active | 0 | 0 |
| 1124 | chr6 | 46128993 | 46130993 | ENSG00000 | ENPP4 | 22875 | 3.804701636 | 0.5547834307 | 0.177677543 | 3.122417265 | 0.001793725 | 0.031575706 | HG01573_active | 0 | 0 |
| 1125 | chr19 | 45487777 | 45489777 | ENSG00000 | PPM1N | 147699 | 3.283025121 | 0.5586660945 | 0.21079421 | 2.650291454 | 0.008042236 | 0.08814792 | HG01573_active | 0 | 0 |
| 1126 | chr7 | 100582982 | 100584982 | ENSG00000 | FBXO24 | 26261 | 4.949165819 | 0.5591939916 | 0.168989767 | 3.309040551 | 0.000936162 | 0.01979125 | HG01573_active | 0 | 0 |
| 1127 | chr19 | 19169289 | 19171289 | ENSG00000 | MEF2B | 100271849 | 7.669520335 | 0.5623432874 | 0.13518967 | 4.159661646 | 3.19E-05 | 0.001351876 | HG01573_active | 0 | 0 |
| 1128 | chr12 | 133078838 | 133080838 | ENSG00000 | ZNF140 | 7699 | 6.756145295 | 0.5676979742 | 0.152619523 | 3.71969433 | 0.000199464 | 0.006212211 | HG01573_active | 0 | 0 |
| 1129 | chr1 | 1723324 | 1725324 | ENSG00000 | CDK11A | 728642 | 4.405569287 | 0.5681452005 | 0.181674407 | 3.127271516 | 0.001764369 | 0.031270874 | HG01573_active | 0 | 0 |
| 1130 | chr17 | 76500790 | 76502790 | ENSG00000 | RHBDF2 | 79651 | 9.460570234 | 0.573667972 | 0.119366662 | 4.805931227 | 1.54E-06 | 0.000113703 | HG01573_active | 0 | 0 |
| 1131 | chr17 | 7328393 | 7330393 | ENSG00000 | NEURL4 | 84461 | 4.080288775 | 0.5739169722 | 0.188096298 | 3.05118695 | 0.002279386 | 0.037322152 | HG01573_active | 0 | 0 |
| 1132 | chr2 | 219542663 | 219544663 | ENSG00000 | TMEM198 | 130612 | 2.893401343 | 0.5750342141 | 0.21518258 | 2.672308388 | 0.007533138 | 0.084260245 | HG01573_active | 0 | 0 |
| 1133 | chr1 | 224174756 | 224176756 | ENSG00000 | DEGS1 | 8560 | 3.914809714 | 0.5755089963 | 0.183452592 | 3.137099286 | 0.001706283 | 0.030600875 | HG01573_active | 0 | 0 |
| 1134 | chr1 | 248858144 | 248860144 | ENSG00000 | ZNF692 | 55657 | 3.324254956 | 0.5762183832 | 0.193075578 | 2.984418788 | 0.002841176 | 0.04398720 | HG01573_active | 0 | 0 |
| 1135 | chr12 | 110500655 | 110502655 | ENSG00000 | RAD9B | 144715 | 6.739129258 | 0.5773296122 | 0.147729251 | 3.908025038 | 9.31E-05 | 0.003342177 | HG01573_active | 0 | 0 |

| | | | | | | | | | | | | | | | | |
|---|---|---|---|---|---|---|---|---|---|---|---|---|---|---|---|---|
| 1136 | chr17 | 18376662 | 18378662 | ENSG00000 | EVPLL | 645027 | 3.877379767 | | 0.5792660634 | 0.197059828 | 2.939544141 | 0.00328695 | 0.04873482 | HG01573_active | 0 | 0 |
| 1137 | chr17 | 78145353 | 78147353 | ENSG00000 | C17orf99 | 100141515 | 5.205621572 | | 0.5804784548 | 0.177489425 | 3.270495994 | 0.001073590 | 0.021667932 | HG01573_active | 0 | 0 |
| 1138 | chr11 | 67265410 | 67267410 | ENSG00000 | ADRBK1 | 156 | 6.634260206 | | 0.5828195532 | 0.139378331 | 4.181565012 | 2.90E-05 | 0.001259979 | HG01573_active | 0 | 0 |
| 1139 | chr7 | 954407 | 956407 | ENSG00000 | ADAP1 | 11033 | 17.06089407 | | 0.5836703347 | 0.091139694 | 6.404128687 | 1.51E-10 | 3.31E-08 | HG01573_active | 0 | 0 |
| 1140 | chr19 | 40412366 | 40414366 | ENSG00000 | PRX | 57716 | 6.12977736 | | 0.5848362578 | 0.152346386 | 3.838858756 | 0.000123607 | 0.004163583 | HG01573_active | 0 | 0 |
| 1141 | chr4 | 992440 | 994440 | ENSG00000 | SLC26A1 | 10861 | 4.761877652 | | 0.5848404912 | 0.157491737 | 3.713467765 | 0.000204438 | 0.006322923 | HG01573_active | 0 | 0 |
| 1142 | chr11 | 68038016 | 68040016 | ENSG00000 | TCIRG1 | 10312 | 4.181568681 | | 0.5864580008 | 0.191784388 | 3.057902714 | 0.002228919 | 0.036994521 | HG01573_active | 0 | 0 |
| 1143 | chr12 | 111368121 | 111370121 | ENSG00000 | FAM109A | 144717 | 3.204400138 | | 0.5868046956 | 0.207501190 | 2.827958209 | 0.004684592 | 0.061363564 | HG01573_active | 0 | 0 |
| 1144 | chr19 | 1651605 | 1653605 | ENSG00000 | TCF3 | 6929 | 11.12254311 | | 0.5871864665 | 0.122158184 | 4.806771394 | 1.53E-06 | 0.000113703 | HG01573_active | 0 | 0 |
| 1145 | chr19 | 48899050 | 48901050 | ENSG00000 | NUCB1 | 4924 | 5.973732712 | | 0.5903034463 | 0.141992928 | 4.15727356 | 3.22E-05 | 0.001361757 | HG01573_active | 0 | 0 |
| 1146 | chr2 | 230863639 | 230865639 | ENSG00000 | ITM2C | 81618 | 5.334549658 | | 0.59105597 | 0.164611670 | 3.590607927 | 0.000329907 | 0.009398497 | HG01573_active | 0 | 0 |
| 1147 | chr6 | 32667383 | 32669383 | ENSG00000 | HLA-DQB1 | 3119 | 2.858569732 | | 0.5913836254 | 0.205456192 | 2.878392798 | 0.003997071 | 0.055746207 | HG01573_active | 0 | 0 |
| 1148 | chr11 | 67649659 | 67651659 | ENSG00000 | ACY3 | 91703 | 2.96632284 | | 0.5917683453 | 0.210735513 | 2.808109248 | 0.004983331 | 0.063898557 | HG01573_active | 0 | 0 |
| 1149 | chr9 | 133417096 | 133419096 | ENSG00000 | REXO4 | 57109 | 3.870763426 | | 0.5923363352 | 0.185092309 | 3.20022122 | 0.001373221 | 0.026136198 | HG01573_active | 0 | 0 |
| 1150 | chr17 | 74872031 | 74874031 | ENSG00000 | FDXR | 2232 | 3.216785256 | | 0.5941920391 | 0.194647363 | 3.05265906 | 0.002268234 | 0.037230817 | HG01573_active | 0 | 0 |
| 1151 | chr19 | 16133028 | 16135028 | ENSG00000 | HSH2D | 84941 | 7.527411642 | | 0.5952431006 | 0.136652327 | 4.355894348 | 1.33E-05 | 0.000673255 | HG01573_active | 0 | 0 |
| 1152 | chr19 | 3499940 | 3501940 | ENSG00000 | DOHH | 83475 | 2.950768434 | | 0.6017779251 | 0.230143493 | 2.614794429 | 0.008928117 | 0.094598394 | HG01573_active | 0 | 0 |
| 1153 | chr19 | 35267978 | 35269978 | ENSG00000 | USF2 | 7392 | 3.487186877 | | 0.6019881692 | 0.199734136 | 3.013947334 | 0.002578725 | 0.040879927 | HG01573_active | 0 | 0 |
| 1154 | chr17 | 7335529 | 7337529 | ENSG00000 | ACAP1 | 9744 | 4.512672817 | | 0.6031494767 | 0.178490178 | 3.379174597 | 0.000727038 | 0.016436475 | HG01573_active | 0 | 0 |
| 1155 | chr17 | 78129770 | 78131770 | ENSG00000 | TMC8 | 147138 | 3.901403561 | | 0.6114550864 | 0.200797543 | 3.045132303 | 0.002325779 | 0.037849869 | HG01573_active | 0 | 0 |
| 1156 | chr2 | 130508666 | 130510666 | ENSG00000 | POTEI | 653269 | 4.257980286 | | 0.6132302231 | 0.191870333 | 3.196065867 | 0.001393153 | 0.026409085 | HG01573_active | 0 | 0 |
| 1157 | chr19 | 49674786 | 49676786 | ENSG00000 | PRMT1 | 3276 | 3.010263348 | | 0.6138208933 | 0.220024362 | 2.789786029 | 0.005274288 | 0.066106725 | HG01573_active | 0 | 0 |
| 1158 | chr4 | 109702583 | 109704583 | ENSG00000 | CASP6 | 839 | 4.041293506 | | 0.6179075051 | 0.198091063 | 3.119310358 | 0.001812749 | 0.031826728 | HG01573_active | 0 | 0 |
| 1159 | chr19 | 2388771 | 2390771 | ENSG00000 | TMPRSS9 | 360200 | 9.070612523 | | 0.6190999411 | 0.119593719 | 5.176692735 | 2.26E-07 | 2.36E-05 | HG01573_active | 0 | 0 |
| 1160 | chr19 | 15463571 | 15465571 | ENSG00000 | RASAL3 | 64926 | 4.105389213 | | 0.6191727413 | 0.182401608 | 3.394557465 | 0.000687396 | 0.015726540 | HG01573_active | 0 | 0 |
| 1161 | chr9 | 131275547 | 131277547 | ENSG00000 | FAM78A | 286336 | 6.173716214 | | 0.619332477 | 0.136490869 | 4.537537796 | 5.69E-06 | 0.000337972 | HG01573_active | 0 | 0 |
| 1162 | chr11 | 68003982 | 68005982 | ENSG00000 | UNC93B1 | 81622 | 3.979135462 | | 0.6200674129 | 0.203037181 | 3.053959919 | 0.002258422 | 0.037230817 | HG01573_active | 0 | 0 |
| 1163 | chr19 | 45691483 | 45693483 | ENSG00000 | QPCTL | 54814 | 3.203481052 | | 0.6268485775 | 0.220085439 | 2.848205581 | 0.004396651 | 0.059038852 | HG01573_active | 0 | 0 |
| 1164 | chr17 | 18475737 | 18477737 | ENSG00000 | LGALS9C | 654346 | 2.542145426 | | 0.6271224865 | 0.227308310 | 2.758906989 | 0.005799503 | 0.070571194 | HG01573_active | 0 | 0 |
| 1165 | chr16 | 648311 | 650311 | ENSG00000 | WDR90 | 197335 | 6.1177518 | | 0.6283206539 | 0.164140726 | 3.827938793 | 0.000129220 | 0.004327118 | HG01573_active | 0 | 0 |
| 1166 | chr9 | 94327644 | 94329644 | ENSG00000 | NUTM2F | 54754 | 2.764247329 | | 0.6286951868 | 0.232510002 | 2.703948996 | 0.006852080 | 0.078584247 | HG01573_active | 0 | 0 |
| 1167 | chr5 | 77638688 | 77640688 | ENSG00000 | OTP | 23440 | 2.652315306 | | 0.6295704975 | 0.229707388 | 2.740749886 | 0.006129915 | 0.072996251 | HG01573_active | 0 | 0 |
| 1168 | chr17 | 46224374 | 46226374 | ENSG00000 | KANSL1 | 284058 | 63.96865825 | | 0.6344163077 | 0.046277701 | 13.70889832 | 8.98E-43 | 1.71E-39 | HG01573_active | 0 | 0 |
| 1169 | chr14 | 105488855 | 105490855 | ENSG00000 | C14orf80 | 283643 | 2.881905878 | | 0.6348887849 | 0.224465604 | 2.828445749 | 0.004677462 | 0.061330304 | HG01573_active | 0 | 0 |
| 1170 | chr19 | 14582084 | 14584084 | ENSG00000 | CLEC17A | 388512 | 8.409681051 | | 0.6354246972 | 0.128612733 | 4.940604879 | 7.79E-07 | 6.71E-05 | HG01573_active | 0 | 0 |
| 1171 | chr18 | 72545 | 74545 | ENSG00000 | RP11-683L2 | 260334 | 6.493934916 | | 0.6367533435 | 0.149868576 | 4.248744852 | 2.15E-05 | 0.000983642 | HG01573_active | 0 | 0 |
| 1172 | chr6 | 44264788 | 44266788 | ENSG00000 | NFKBIE | 4794 | 2.596909671 | | 0.642351034 | 0.241735220 | 2.657250489 | 0.007878088 | 0.087038300 | HG01573_active | 0 | 0 |
| 1173 | chr14 | 77141734 | 77143734 | ENSG00000 | ZDHHC22 | 283576 | 4.016371422 | | 0.6450013394 | 0.182473766 | 3.534762014 | 0.000408142 | 0.011151723 | HG01573_active | 0 | 0 |
| 1174 | chr11 | 625078 | 627078 | ENSG00000 | CDHR5 | 53841 | 2.917356454 | | 0.64612854 | 0.224009977 | 2.884373932 | 0.003921927 | 0.054927540 | HG01573_active | 0 | 0 |
| 1175 | chrX | 1536107 | 1538107 | ENSG00000 | P2RY8 | 286530 | 22.39698971 | | 0.6536863079 | 0.082029565 | 7.968911061 | 1.60E-15 | 8.91E-13 | HG01573_active | 0 | 0 |
| 1176 | chr8 | 124449820 | 124451820 | ENSG00000 | TRMT12 | 55039 | 3.597412916 | | 0.657333887 | 0.188359520 | 3.489783179 | 0.000483412 | 0.012545922 | HG01573_active | 0 | 0 |
| 1177 | chr12 | 6635447 | 6637447 | ENSG00000 | LPAR5 | 57121 | 4.700070673 | | 0.6577409656 | 0.185908071 | 3.537990369 | 0.000403184 | 0.011061502 | HG01573_active | 0 | 0 |
| 1178 | chr17 | 7280667 | 7282667 | ENSG00000 | SLC2A4 | 6517 | 2.739070978 | | 0.6592962228 | 0.229974621 | 2.866821648 | 0.004146165 | 0.057323621 | HG01573_active | 0 | 0 |
| 1179 | chr19 | 45496061 | 45498061 | ENSG00000 | RTN2 | 6253 | 3.023625587 | | 0.6625693516 | 0.221469862 | 2.991690807 | 0.002774371 | 0.043203229 | HG01573_active | 0 | 0 |
| 1180 | chr17 | 29621907 | 29623907 | ENSG00000 | CORO6 | 84940 | 2.529774758 | | 0.6673855783 | 0.238004147 | 2.804092221 | 0.005045847 | 0.064391176 | HG01573_active | 0 | 0 |
| 1181 | chr1 | 109686814 | 109688814 | ENSG00000 | GSTM1 | 2944 | 3.829757687 | | 0.6707194535 | 0.190764455 | 3.515956116 | 0.000438173 | 0.011697232 | HG01573_active | 0 | 0 |
| 1182 | chr6 | 32952122 | 32954122 | ENSG00000 | XXbac-BPG | NaN | 5.579104788 | | 0.6736602302 | 0.159415204 | 4.225821712 | 2.38E-05 | 0.001067402 | HG01573_active | 0 | 0 |
| 1183 | chr6 | 43628540 | 43630540 | ENSG00000 | MAD2L1BP | 9587 | 2.757282784 | | 0.6884477598 | 0.210468344 | 3.271027575 | 0.001071574 | 0.021660073 | HG01573_active | 0 | 0 |
| 1184 | chr9 | 21801636 | 21803636 | ENSG00000 | RP11-145E5 | NaN | 86.89199646 | | 0.6890411491 | 0.051223082 | 13.4517706 | 3.01E-41 | 5.02E-38 | HG01573_active | 0 | 0 |
| 1185 | chr19 | 49579646 | 49581646 | ENSG00000 | PRRG2 | 5639 | 2.769063229 | | 0.6921050788 | 0.214509326 | 3.22645682 | 0.001253331 | 0.024410730 | HG01573_active | 0 | 0 |
| 1186 | chrX | 153971327 | 153973327 | ENSG00000 | TMEM187 | 8269 | 2.96377585 | | 0.6963910997 | 0.225099973 | 3.093696945 | 0.001976792 | 0.033948497 | HG01573_active | 0 | 0 |
| 1187 | chr7 | 65005743 | 65007743 | ENSG00000 | ERV3-1 | 2086 | 5.114134084 | | 0.698071379 | 0.161573639 | 4.320453406 | 1.56E-05 | 0.000759279 | HG01573_active | 0 | 0 |
| 1188 | chr16 | 635817 | 637817 | ENSG00000 | FAM195A | 84331 | 2.588452309 | | 0.6993676314 | 0.230794548 | 3.030260626 | 0.002443427 | 0.039285968 | HG01573_active | 0 | 0 |
| 1189 | chr6 | 31398784 | 31400784 | ENSG00000 | MICA | 100507436 | 3.731921099 | | 0.6998248561 | 0.203712915 | 3.435348488 | 0.000591792 | 0.014221109 | HG01573_active | 0 | 0 |
| 1190 | chr12 | 6443867 | 6445867 | ENSG00000 | CD27 | 939 | 2.286407292 | | 0.7019660972 | 0.24175834 | 2.903585858 | 0.003689158 | 0.052578022 | HG01573_active | 0 | 0 |
| 1191 | chr17 | 45261112 | 45263112 | ENSG00000 | SPATA32 | 124783 | 2.721265908 | | 0.7023310725 | 0.228695684 | 3.071028967 | 0.002133224 | 0.035942008 | HG01573_active | 0 | 0 |
| 1192 | chr19 | 57350307 | 57352307 | ENSG00000 | ZNF304 | 57343 | 2.852402952 | | 0.7041331444 | 0.242119159 | 2.90820911 | 0.003635051 | 0.052206393 | HG01573_active | 0 | 0 |
| 1193 | chr9 | 133478137 | 133480137 | ENSG00000 | SLC2A6 | 11182 | 2.992817008 | | 0.7056659324 | 0.209404602 | 3.369868305 | 0.000752041 | 0.016887431 | HG01573_active | 0 | 0 |
| 1194 | chr4 | 672580 | 674580 | ENSG00000 | MYL5 | 4636 | 2.401451346 | | 0.7095375927 | 0.246501695 | 2.878428854 | 0.003996614 | 0.055746207 | HG01573_active | 0 | 0 |
| 1195 | chr19 | 13951492 | 13953492 | ENSG00000 | DCAF15 | 90379 | 2.606661134 | | 0.7118562842 | 0.263132354 | 2.70531644 | 0.006823935 | 0.078396041 | HG01573_active | 0 | 0 |

| | | | | | | | | | | | | | | | | |
|---|---|---|---|---|---|---|---|---|---|---|---|---|---|---|---|---|
| 1196 | chr7 | 44978098 | 44980098 | ENSG00000 | MYO1G | 64005 | 6.339650791 | 0.714597317 | 0.158185072 | 4.517476304 | 6.26E-06 | 0.000366730 | HG01573_active | | 0 | 0 |
| 1197 | chr19 | 29810898 | 29812898 | ENSG00000 | CCNE1 | 898 | 3.036203952 | 0.715657448 | 0.207838732 | 3.443330509 | 0.000574596 | 0.013983944 | HG01573_active | | 0 | 0 |
| 1198 | chr20 | 5125626 | 5127626 | ENSG00000 | PCNA | 5111 | 2.489766933 | 0.7241050135 | 0.240243854 | 3.014041772 | 0.002577923 | 0.040879927 | HG01573_active | | 0 | 0 |
| 1199 | chr20 | 64078988 | 64080988 | ENSG00000 | RGS19 | 10287 | 2.224186554 | 0.7257842627 | 0.276105372 | 2.628649553 | 0.008572465 | 0.092186829 | HG01573_active | | 0 | 0 |
| 1200 | chr9 | 137026464 | 137028464 | ENSG00000 | C9orf139 | NaN | 3.168002111 | 0.7275093451 | 0.238256130 | 3.053475865 | 0.002262069 | 0.037230817 | HG01573_active | | 0 | 0 |
| 1201 | chr19 | 18000132 | 18002132 | ENSG00000 | ARRDC2 | 27106 | 3.267678231 | 0.7278116554 | 0.221309017 | 3.288666961 | 0.001006630 | 0.020819800 | HG01573_active | | 0 | 0 |
| 1202 | chr19 | 13861063 | 13863063 | ENSG00000 | NANOS3 | 342977 | 4.982300937 | 0.7309390998 | 0.168754848 | 4.331366502 | 1.48E-05 | 0.000730599 | HG01573_active | | 0 | 0 |
| 1203 | chr22 | 50326060 | 50328060 | ENSG00000 | DENND6B | 414918 | 5.746627748 | 0.7336359967 | 0.159682523 | 4.5943412 | 4.34E-06 | 0.000268530 | HG01573_active | | 0 | 0 |
| 1204 | chr17 | 78131407 | 78133407 | ENSG00000 | TMC6 | 11322 | 7.348459188 | 0.7395492632 | 0.144003339 | 5.13563968 | 2.81E-07 | 2.80E-05 | HG01573_active | | 0 | 0 |
| 1205 | chr9 | 132581185 | 132583185 | ENSG00000 | BARHL1 | 56751 | 2.437358874 | 0.7415119859 | 0.245173273 | 3.024440528 | 0.002490934 | 0.039905724 | HG01573_active | | 0 | 0 |
| 1206 | chr16 | 88650152 | 88652152 | ENSG00000 | CYBA | 1535 | 2.467021949 | 0.7417529003 | 0.233869751 | 3.171649581 | 0.001515757 | 0.027984864 | HG01573_active | | 0 | 0 |
| 1207 | chr17 | 7293615 | 7295615 | ENSG00000 | YBX2 | 51087 | 2.259503235 | 0.7434885616 | 0.271791376 | 2.735511961 | 0.006228332 | 0.073643139 | HG01573_active | | 0 | 0 |
| 1208 | chr19 | 50332515 | 50334515 | ENSG00000 | KCNC3 | 3748 | 6.238837961 | 0.7466250578 | 0.168311212 | 4.435979329 | 9.17E-06 | 0.000506032 | HG01573_active | | 0 | 0 |
| 1209 | chr6 | 390739 | 392739 | ENSG00000 | IRF4 | 3662 | 5.88258652 | 0.7482546342 | 0.171246462 | 4.369460381 | 1.25E-05 | 0.000637611 | HG01573_active | | 0 | 0 |
| 1210 | chr11 | 277365 | 279365 | ENSG00000 | NLRP6 | 171389 | 2.152484834 | 0.7518195293 | 0.265701135 | 2.829568372 | 0.004661083 | 0.061235735 | HG01573_active | | 0 | 0 |
| 1211 | chr17 | 74465416 | 74467416 | ENSG00000 | CD300A | 11314 | 7.007214167 | 0.7530125195 | 0.145715264 | 5.167698263 | 2.37E-07 | 2.44E-05 | HG01573_active | | 0 | 0 |
| 1212 | chr7 | 74198652 | 74200652 | ENSG00000 | LAT2 | 7462 | 6.78948828 | 0.7574327987 | 0.151273898 | 5.00702903 | 5.53E-07 | 5.02E-05 | HG01573_active | | 0 | 1 |
| 1213 | chr19 | 50822787 | 50824787 | ENSG00000 | KLK1 | 3816 | 1.848726849 | 0.7651049203 | 0.278255228 | 2.749651549 | 0.005965866 | 0.071681606 | HG01573_active | | 0 | 0 |
| 1214 | chr11 | 416455 | 418455 | ENSG00000 | SIGIRR | 59307 | 2.994190985 | 0.7679740084 | 0.238799399 | 3.215979653 | 0.001300000 | 0.025027819 | HG01573_active | | 0 | 0 |
| 1215 | chr7 | 155811273 | 155813273 | ENSG00000 | SHH | 6469 | 3.162630512 | 0.770551724 | 0.225753891 | 3.413237836 | 0.000641958 | 0.015074187 | HG01573_active | | 0 | 0 |
| 1216 | chr8 | 144516992 | 144518992 | ENSG00000 | LRRC14 | 9684 | 2.146611524 | 0.7719122875 | 0.265296449 | 2.909621625 | 0.003618665 | 0.052155034 | HG01573_active | | 0 | 0 |
| 1217 | chr16 | 3035926 | 3037926 | ENSG00000 | CCDC64B | 146439 | 2.333641549 | 0.7773356373 | 0.267160546 | 2.909619879 | 0.003618685 | 0.052155034 | HG01573_active | | 0 | 0 |
| 1218 | chr19 | 3171346 | 3173346 | ENSG00000 | S1PR4 | 8698 | 2.06433334 | 0.7781565836 | 0.282182283 | 2.757637988 | 0.005822063 | 0.070716900 | HG01573_active | | 0 | 0 |
| 1219 | chr17 | 43005725 | 43007725 | ENSG00000 | IFI35 | 3430 | 2.084108182 | 0.7792790702 | 0.273905961 | 2.845060646 | 0.004440296 | 0.059445697 | HG01573_active | | 0 | 0 |
| 1220 | chr17 | 49229766 | 49231766 | ENSG00000 | PHOSPHO1 | 162466 | 2.233362362 | 0.7827481148 | 0.277045173 | 2.825344712 | 0.004722977 | 0.061745305 | HG01573_active | | 0 | 0 |
| 1221 | chr12 | 110449422 | 110451422 | ENSG00000 | ARPC3 | 10094 | 2.890469137 | 0.7838906709 | 0.231222246 | 3.390204369 | 0.000698405 | 0.015896753 | HG01573_active | | 0 | 0 |
| 1222 | chr2 | 69892560 | 69894560 | ENSG00000 | SNRNP27 | 11017 | 2.30920106 | 0.7843971232 | 0.266177242 | 2.946897773 | 0.003209792 | 0.047863888 | HG01573_active | | 0 | 0 |
| 1223 | chr20 | 57390407 | 57392407 | ENSG00000 | RBM38 | 55544 | 6.053452098 | 0.8004892144 | 0.152682084 | 5.242849655 | 1.58E-07 | 1.75E-05 | HG01573_active | | 0 | 0 |
| 1224 | chr5 | 178112443 | 178114443 | ENSG00000 | N4BP3 | 23138 | 4.681683112 | 0.8007897451 | 0.165010152 | 4.852972589 | 1.22E-06 | 9.73E-05 | HG01573_active | | 0 | 0 |
| 1225 | chr17 | 68034519 | 68036519 | ENSG00000 | KPNA2 | 3838 | 2.187315866 | 0.8011520597 | 0.267263057 | 2.997616158 | 0.002721001 | 0.042670540 | HG01573_active | | 0 | 0 |
| 1226 | chr1 | 90716237 | 90718237 | ENSG00000 | BARHL2 | 343472 | 1.843455638 | 0.81148447 | 0.311292691 | 2.606821467 | 0.009138699 | 0.096143437 | HG01573_active | | 0 | 0 |
| 1227 | chr7 | 101320823 | 101322823 | ENSG00000 | IFT22 | 64792 | 2.10509358 | 0.814686093 | 0.295096677 | 2.760743024 | 0.005767003 | 0.070239682 | HG01573_active | | 0 | 0 |
| 1228 | chr7 | 27174180 | 27176180 | ENSG00000 | HOXA9 | 3205 | 3.358752874 | 0.8194875669 | 0.212329497 | 3.859508809 | 0.000113615 | 0.003918578 | HG01573_active | | 0 | 0 |
| 1229 | chr6 | 32438842 | 32440842 | ENSG00000 | HLA-DRA | 3122 | 1.682108889 | 0.8303818653 | 0.292415667 | 2.839731105 | 0.004515157 | 0.060146581 | HG01573_active | | 0 | 0 |
| 1230 | chr19 | 17280645 | 17282645 | ENSG00000 | ANKLE1 | 126549 | 1.548963397 | 0.8341241328 | 0.322315998 | 2.587907944 | 0.009656078 | 0.099094935 | HG01573_active | | 0 | 0 |
| 1231 | chr8 | 144526032 | 144528032 | ENSG00000 | LRRC24 | 441381 | 1.564190779 | 0.8494095092 | 0.321917567 | 2.638593211 | 0.008325081 | 0.090505620 | HG01573_active | | 0 | 0 |
| 1232 | chr5 | 150411929 | 150413929 | ENSG00000 | CD74 | 972 | 4.066061545 | 0.8610440892 | 0.185075182 | 4.65240167 | 3.28E-06 | 0.000214605 | HG01573_active | | 0 | 1 |
| 1233 | chr1 | 146020918 | 146022918 | ENSG00000 | HFE2 | 148738 | 1.733030267 | 0.8646928453 | 0.302304626 | 2.860336128 | 0.004231922 | 0.057886547 | HG01573_active | | 0 | 0 |
| 1234 | chr16 | 75218000 | 75220000 | ENSG00000 | CTRB1 | 1504 | 1.624328752 | 0.8718660164 | 0.310720881 | 2.805946009 | 0.005016909 | 0.064267431 | HG01573_active | | 0 | 0 |
| 1235 | chr11 | 825144 | 827144 | ENSG00000 | EFCAB4A | 283229 | 2.053415997 | 0.8770799244 | 0.283834556 | 3.090109734 | 0.002000825 | 0.034229232 | HG01573_active | | 0 | 0 |
| 1236 | chr2 | 176082913 | 176084913 | ENSG00000 | EVX2 | 344191 | 1.666840513 | 0.8885197047 | 0.299494502 | 2.966731269 | 0.003009839 | 0.045750238 | HG01573_active | | 0 | 0 |
| 1237 | chr1 | 161224939 | 161226939 | ENSG00000 | TOMM40L | 84134 | 1.661334793 | 0.8956635681 | 0.306617137 | 2.921113857 | 0.003487823 | 0.051019828 | HG01573_active | | 0 | 0 |
| 1238 | chr12 | 6945468 | 6947468 | ENSG00000 | PTPN6 | 5777 | 5.143413888 | 0.8989764407 | 0.178780517 | 5.028380347 | 4.95E-07 | 4.56E-05 | HG01573_active | | 0 | 0 |
| 1239 | chr19 | 1064923 | 1066923 | ENSG00000 | HMHA1 | 23526 | 5.037758416 | 0.9059431615 | 0.174490189 | 5.191943265 | 2.08E-07 | 2.21E-05 | HG01573_active | | 0 | 0 |
| 1240 | chr5 | 180987845 | 180989845 | ENSG00000 | BTNL3 | 10917 | 4.666142597 | 0.9100206447 | 0.170730567 | 5.33015652 | 9.81E-08 | 1.16E-05 | HG01573_active | | 0 | 0 |
| 1241 | chr9 | 21801543 | 21803543 | ENSG00000 | MTAP | 4507 | 62.1593951 | 0.9159221119 | 0.059819697 | 15.31137983 | 6.42E-53 | 1.72E-49 | HG01573_active | | 0 | 0 |
| 1242 | chr19 | 12680902 | 12682902 | ENSG00000 | DHPS | 1725 | 1.551191448 | 0.9584707948 | 0.311582392 | 3.076139151 | 0.002096999 | 0.035510787 | HG01573_active | | 0 | 0 |
| 1243 | chr17 | 82023107 | 82025107 | ENSG00000 | STRA13 | 201254 | 1.443592349 | 1.124550388 | 0.367166169 | 3.062783236 | 0.002192888 | 0.036655723 | HG01573_active | | 0 | 0 |
| 1244 | chr16 | 74376878 | 74378878 | ENSG00000 | NPIPB15 | 440348 | 4.397809877 | 1.222855095 | 0.204382305 | 5.983174977 | 2.19E-09 | 3.90E-07 | HG01573_active | | 0 | 0 |
| 1245 | chr16 | 70112626 | 70114626 | ENSG00000 | PDPR | 55066 | 26.80908917 | 1.278040699 | 0.078605750 | 16.25887035 | 1.93E-59 | 6.46E-56 | HG01573_active | | 0 | 0 |
| 1246 | chr17 | 36438566 | 36440566 | ENSG00000 | TBC1D3B | 729877 | 3.250540366 | 1.971419329 | 0.256574454 | 7.683615018 | 1.55E-14 | 7.38E-12 | HG01573_active | | 0 | 0 |

Table S2B. List of genes inferred to be changing in activity when comparing major and minor clones with scNOVA (C7 vs. RPE-1)

| Index | chr | str | end | name | gname | EntrezID | baseMean | NO log2FoldChange (C7/W | lfcSE | stat | pvalue | padj | Inferred gene activity |
|---|---|---|---|---|---|---|---|---|---|---|---|---|---|
| 1 | chrX | 21654690 | 21658330 | ENSG00000185 | KLHL34 | 257240 | 0.8117519 | -0.9043408 | 0.30436893 | -2.9711996 | 0.00296639 | 0.02858917 | C7_active |
| 2 | chr8 | 144082590 | 144086216 | ENSG00000197 | GPAA1 | 8733 | 0.97096439 | -0.8938194 | 0.25518801 | -3.5025917 | 0.00046076 | 0.00602282 | C7_active |
| 3 | chr19 | 14566078 | 14572062 | ENSG00000099 | NDUFB7 | 4713 | 1.58921064 | -0.8215099 | 0.23350443 | -3.5181767 | 0.00043452 | 0.00572665 | C7_active |
| 4 | chr7 | 100889994 | 100896974 | ENSG00000087 | ACHE | 43 | 1.64293498 | -0.8011397 | 0.20004994 | -4.0046983 | 6.21E-05 | 0.00097635 | C7_active |
| 5 | chr7 | 150338317 | 150341674 | ENSG00000106 | RARRES2 | 5919 | 0.79779962 | -0.7475836 | 0.29067436 | -2.571894 | 0.01011439 | 0.07406257 | C7_active |
| 6 | chr17 | 28719393 | 28724359 | ENSG00000198 | RPL23A | 6147 | 0.93388922 | -0.7446016 | 0.27595012 | -2.6983196 | 0.00696905 | 0.05594642 | C7_active |
| 7 | chr19 | 42387019 | 42390287 | ENSG00000105 | CNFN | 84518 | 0.88962376 | -0.7329238 | 0.28044835 | -2.6134004 | 0.00896462 | 0.06769907 | C7_active |
| 8 | chr22 | 20111866 | 20117392 | ENSG00000099 | TRMT2A | 27037 | 1.3690553 | -0.7315326 | 0.21623798 | -3.3829978 | 0.00071699 | 0.00874585 | C7_active |
| 9 | chr16 | 784974 | 788397 | ENSG00000007 | RPUSD1 | 113000 | 0.89505496 | -0.6854586 | 0.28202364 | -2.4305006 | 0.01507798 | 0.09908585 | C7_active |
| 10 | chr20 | 35278907 | 35284985 | ENSG00000242 | EIF6 | 3692 | 1.29135897 | -0.6720882 | 0.2394598 | -2.806685 | 0.00500542 | 0.04326624 | C7_active |
| 11 | chr19 | 32972209 | 32978222 | ENSG00000131 | C19orf40 | 91442 | 1.30809928 | -0.6673875 | 0.23370762 | -2.8556513 | 0.00429487 | 0.03865379 | C7_active |
| 12 | chr5 | 179806398 | 179838078 | ENSG00000161 | SQSTM1 | 8878 | 6.80417785 | -0.6574205 | 0.09305852 | -7.0645923 | 1.61E-12 | 3.91E-11 | C7_active |
| 13 | chr8 | 144354135 | 144361272 | ENSG00000185 | SLC52A2 | 79581 | 1.72764635 | -0.6534284 | 0.19777059 | -3.3039717 | 0.00095325 | 0.01111976 | C7_active |
| 14 | chr19 | 10289981 | 10296778 | ENSG00000105 | ICAM5 | 7087 | 1.55974904 | -0.6521334 | 0.20939202 | -3.1144141 | 0.00184311 | 0.01932648 | C7_active |
| 15 | chr4 | 87975650 | 87983426 | ENSG00000118 | SPP1 | 6696 | 1.37792506 | -0.6474608 | 0.20758897 | -3.1189558 | 0.00181493 | 0.0191005 | C7_active |
| 16 | chr17 | 36936785 | 36944612 | ENSG00000273 | LHX1 | 3975 | 1.55931631 | -0.643047 | 0.19755379 | -3.2550478 | 0.00113373 | 0.01286086 | C7_active |
| 17 | chr17 | 48621159 | 48626356 | ENSG00000170 | HOXB9 | 3219 | 1.64042216 | -0.6350807 | 0.17861205 | -3.5556429 | 0.00037706 | 0.00506182 | C7_active |
| 18 | chr22 | 50525752 | 50530085 | ENSG00000025 | TYMP | 1890 | 0.98964602 | -0.6193094 | 0.24293207 | -2.549311 | 0.0107936 | 0.07763 | C7_active |
| 19 | chr19 | 825097 | 832018 | ENSG00000172 | AZU1 | 566 | 1.92919233 | -0.6091249 | 0.21658604 | -2.8123922 | 0.00491745 | 0.04273647 | C7_active |
| 20 | chr17 | 38730235 | 38735044 | ENSG00000277 | CISD3 | 284106 | 1.33886542 | -0.580626 | 0.20840293 | -2.786074 | 0.00533507 | 0.04530096 | C7_active |
| 21 | chr16 | 31471585 | 31477960 | ENSG00000140 | TGFB1I1 | 7041 | 1.35274456 | -0.5778654 | 0.21507688 | -2.6867853 | 0.00721433 | 0.05756293 | C7_active |
| 22 | chr17 | 44070735 | 44076344 | ENSG00000141 | G6PC3 | 92579 | 1.7465189 | -0.5776958 | 0.19493022 | -2.9636031 | 0.0030406 | 0.02922627 | C7_active |
| 23 | chr15 | 39581079 | 39599466 | ENSG00000137 | THBS1 | 7057 | 3.69109552 | -0.5693253 | 0.13521689 | -4.2104598 | 2.55E-05 | 0.00042926 | C7_active |
| 24 | chr16 | 4608883 | 4615027 | ENSG00000153 | UBALD1 | 124402 | 1.39867926 | -0.5677452 | 0.22043043 | -2.5756208 | 0.01000603 | 0.07341831 | C7_active |
| 25 | chr17 | 75165586 | 75182983 | ENSG00000188 | SUMO2 | 6613 | 2.67238962 | -0.5612919 | 0.15397853 | -3.6452609 | 0.00026712 | 0.00373192 | C7_active |
| 26 | chr17 | 8339864 | 8346046 | ENSG00000184 | ODF4 | 146852 | 1.92345309 | -0.5557497 | 0.17736618 | -3.1333465 | 0.00172825 | 0.01840322 | C7_active |
| 27 | chr19 | 58432814 | 58440222 | ENSG00000131 | ZNF132 | 7691 | 2.10678165 | -0.5476928 | 0.16475901 | -3.3242053 | 0.00088671 | 0.01047057 | C7_active |
| 28 | chr11 | 747329 | 765024 | ENSG00000177 | TALDO1 | 6888 | 3.32186981 | -0.5412626 | 0.13654375 | -3.9640229 | 7.37E-05 | 0.00114254 | C7_active |
| 29 | chr8 | 143597835 | 143603224 | ENSG00000179 | TIGD5 | 84948 | 1.36172332 | -0.5353002 | 0.21767862 | -2.4591308 | 0.01392739 | 0.09405389 | C7_active |
| 30 | chr12 | 57747727 | 57756013 | ENSG00000135 | CDK4 | 1019 | 1.75163612 | -0.535247 | 0.20254851 | -2.6425617 | 0.00822815 | 0.06347427 | C7_active |
| 31 | chr22 | 41433492 | 41447023 | ENSG00000183 | TOB2 | 10766 | 2.54789064 | -0.5340132 | 0.14375714 | -3.7146901 | 0.00020345 | 0.0028986 | C7_active |
| 32 | chr19 | 10512742 | 10517931 | ENSG00000180 | S1PR5 | 53637 | 1.42733586 | -0.5321411 | 0.21790757 | -2.4420497 | 0.01460414 | 0.09732991 | C7_active |
| 33 | chr19 | 50797704 | 50804929 | ENSG00000167 | C19orf48 | 84798 | 1.51400557 | -0.5253105 | 0.21270045 | -2.4697199 | 0.01352189 | 0.09217899 | C7_active |
| 34 | chr1 | 160739057 | 160754821 | ENSG00000026 | SLAMF7 | 57823 | 5.43044356 | -0.5247053 | 0.10489494 | -5.0021985 | 5.67E-07 | 1.11E-05 | C7_active |
| 35 | chr1 | 156721891 | 156728799 | ENSG00000143 | ISG20L2 | 81875 | 1.44605736 | -0.519576 | 0.19971279 | -2.601616 | 0.00927857 | 0.06945919 | C7_active |
| 36 | chr1 | 1311585 | 1324691 | ENSG00000127 | CPSF3L | 54973 | 3.32785829 | -0.5185651 | 0.13444374 | -3.857116 | 0.00011473 | 0.00172494 | C7_active |
| 37 | chr14 | 105472962 | 105480170 | ENSG00000182 | CRIP2 | 1397 | 1.60833243 | -0.5175527 | 0.20833966 | -2.4841774 | 0.01298511 | 0.08906724 | C7_active |
| 38 | chr19 | 12610918 | 12633840 | ENSG00000173 | ZNF791 | 163049 | 3.37641852 | -0.516367 | 0.13583589 | -3.8014035 | 0.00014388 | 0.00212111 | C7_active |
| 39 | chr1 | 8861002 | 8879249 | ENSG00000074 | ENO1 | 2023 | 3.74582037 | -0.5143947 | 0.1271225 | -4.0464485 | 5.20E-05 | 0.00082936 | C7_active |
| 40 | chr1 | 112709994 | 112715477 | ENSG00000155 | PPM1J | 333926 | 2.07939858 | -0.5108852 | 0.15527484 | -3.2901995 | 0.00100116 | 0.01163324 | C7_active |
| 41 | chr16 | 1773207 | 1781708 | ENSG00000197 | EME2 | 197342 | 2.04034309 | -0.5070312 | 0.17902211 | -2.8322267 | 0.00462251 | 0.0408128 | C7_active |
| 42 | chr19 | 17511629 | 17521288 | ENSG00000130 | PGLS | 25796 | 2.26822342 | -0.5018241 | 0.14542415 | -3.4507615 | 0.00055901 | 0.00711365 | C7_active |
| 43 | chrX | 153929242 | 153935223 | ENSG00000102 | NAA10 | 8260 | 1.68842263 | -0.4982096 | 0.19908279 | -2.5025245 | 0.01233111 | 0.08547593 | C7_active |
| 44 | chr19 | 6426037 | 6433779 | ENSG00000181 | SLC25A41 | 284427 | 1.88017031 | -0.4952749 | 0.18759426 | -2.6401388 | 0.00828721 | 0.06386156 | C7_active |
| 45 | chr19 | 12965159 | 12974762 | ENSG00000179 | DAND5 | 199699 | 2.15718699 | -0.494863 | 0.17613017 | -2.8096437 | 0.00495964 | 0.04298086 | C7_active |
| 46 | chr19 | 45379634 | 45406349 | ENSG00000104 | PPP1R13L | 10848 | 5.47412414 | -0.4911223 | 0.09881679 | -4.9700294 | 6.69E-07 | 1.30E-05 | C7_active |
| 47 | chr12 | 7787794 | 7799141 | ENSG00000111 | NANOG | 79923 | 2.95777481 | -0.4881129 | 0.13774064 | -3.5437101 | 0.00039454 | 0.00526222 | C7_active |
| 48 | chr11 | 67428460 | 67435408 | ENSG00000175 | RPS6KB2 | 6199 | 2.03442918 | -0.4872817 | 0.1729411 | -2.8176163 | 0.00483816 | 0.04224745 | C7_active |
| 49 | chr11 | 62559601 | 62574086 | ENSG00000254 | EEF1G | 1937 | 3.09303413 | -0.4796853 | 0.14204126 | -3.3770844 | 0.00073259 | 0.00888345 | C7_active |
| 50 | chr12 | 6913745 | 6923698 | ENSG00000111 | ENO2 | 2026 | 2.67481984 | -0.4756862 | 0.14641532 | -3.2488825 | 0.00115859 | 0.01312223 | C7_active |
| 51 | chr1 | 150293720 | 150308979 | ENSG00000266 | MRPS21 | 54460 | 4.06243989 | -0.4747624 | 0.1182953 | -4.0133666 | 5.99E-05 | 0.00094368 | C7_active |

| | | | | | | | | | | | | | |
|---|---|---|---|---|---|---|---|---|---|---|---|---|---|
| 52 | chr3 | 51933430 | 51941941 | ENSG00000114 | RRP9 | 9136 | 2.93478192 | -0.4733314 | 0.14737849 | -3.2116719 | 0.00131965 | 0.01465848 | C7_active |
| 53 | chr6 | 12290363 | 12297194 | ENSG00000078 | EDN1 | 1906 | 1.54612524 | -0.4702305 | 0.19375749 | -2.4269025 | 0.01522835 | 0.09965423 | C7_active |
| 54 | chrX | 154541199 | 154565046 | ENSG00000269 | IKBKG | 8517 | 4.23795482 | -0.4682031 | 0.11985178 | -3.906518 | 9.36E-05 | 0.00142409 | C7_active |
| 55 | chr14 | 39175183 | 39183218 | ENSG00000100 | PNN | 5411 | 1.50518277 | -0.4673988 | 0.19114388 | -2.4452722 | 0.01447429 | 0.0968007 | C7_active |
| 56 | chr17 | 75845699 | 75856507 | ENSG00000132 | WBP2 | 23558 | 2.34695589 | -0.4661397 | 0.16935516 | -2.7524387 | 0.00591532 | 0.04937773 | C7_active |
| 57 | chr15 | 74836116 | 74843346 | ENSG00000140 | ULK3 | 25989 | 2.51400305 | -0.4644195 | 0.15582992 | -2.9802973 | 0.00287969 | 0.02788404 | C7_active |
| 58 | chr12 | 46358189 | 46372867 | ENSG00000134 | SLC38A2 | 54407 | 2.42970078 | -0.4637779 | 0.16045286 | -2.8904307 | 0.00384714 | 0.03558182 | C7_active |
| 59 | chr22 | 20115938 | 20127357 | ENSG00000099 | RANBP1 | 5902 | 2.50958772 | -0.4610957 | 0.1447856 | -3.1846794 | 0.00144914 | 0.01586467 | C7_active |
| 60 | chr1 | 160945025 | 160954799 | ENSG00000158 | ITLN2 | 142683 | 3.53869007 | -0.4591636 | 0.13025167 | -3.525203 | 0.00042316 | 0.00558709 | C7_active |
| 61 | chrX | 151912509 | 151925170 | ENSG00000147 | MAGEA4 | 4103 | 2.37669372 | -0.4551303 | 0.1591398 | -2.8599399 | 0.00423721 | 0.03823037 | C7_active |
| 62 | chr21 | 44805617 | 44818779 | ENSG00000184 | SUMO3 | 6612 | 2.969643 | -0.4543933 | 0.14875577 | -3.0546262 | 0.00225341 | 0.02284788 | C7_active |
| 63 | chr14 | 24216852 | 24232454 | ENSG00000129 | NEDD8 | 4738 | 2.8071868 | -0.4528479 | 0.1406103 | -3.2205884 | 0.00127928 | 0.01429816 | C7_active |
| 64 | chr1 | 1020123 | 1056118 | ENSG00000188 | AGRN | 375790 | 7.44799004 | -0.450252 | 0.08813261 | -5.1088013 | 3.24E-07 | 6.49E-06 | C7_active |
| 65 | chr19 | 45485289 | 45497061 | ENSG00000125 | RTN2 | 6253 | 2.77826987 | -0.4497627 | 0.14731063 | -3.0531582 | 0.00226447 | 0.02292771 | C7_active |
| 66 | chr20 | 56629302 | 56639283 | ENSG00000087 | TFAP2C | 7022 | 3.05447268 | -0.4472844 | 0.12935971 | -3.4576792 | 0.00054485 | 0.0069519 | C7_active |
| 67 | chr7 | 73568932 | 73578791 | ENSG00000106 | TBL2 | 26608 | 2.48556246 | -0.4445754 | 0.14597154 | -3.0456305 | 0.00232193 | 0.0233456 | C7_active |
| 68 | chr11 | 102835801 | 102843803 | ENSG00000149 | MMP3 | 4314 | 1.83052854 | -0.4359815 | 0.17363455 | -2.5109143 | 0.01204189 | 0.08399614 | C7_active |
| 69 | chr11 | 60040275 | 60048044 | ENSG00000149 | OOSP2 | 219990 | 2.28223932 | -0.432299 | 0.17576828 | -2.4594827 | 0.01391374 | 0.09400577 | C7_active |
| 70 | chr17 | 81843159 | 81860694 | ENSG00000185 | P4HB | 5034 | 4.18201086 | -0.4318793 | 0.1197701 | -3.6059023 | 0.00031107 | 0.00426737 | C7_active |
| 71 | chr16 | 88696495 | 88706421 | ENSG00000158 | RNF166 | 115992 | 3.04432027 | -0.4305973 | 0.14819898 | -2.905535 | 0.00366626 | 0.03423583 | C7_active |
| 72 | chr1 | 20651767 | 20661544 | ENSG00000244 | DDOST | 1650 | 2.90340533 | -0.4294271 | 0.14769833 | -2.9074604 | 0.00364376 | 0.03409202 | C7_active |
| 73 | chr19 | 50996018 | 51009581 | ENSG00000269 | KLK9 | NaN | 4.68875284 | -0.4284866 | 0.11587844 | -3.6977246 | 0.00021754 | 0.00308104 | C7_active |
| 74 | chr17 | 7243591 | 7252491 | ENSG00000175 | CTDNEP1 | 23399 | 2.23636077 | -0.426174 | 0.16010663 | -2.6618138 | 0.00777209 | 0.06086798 | C7_active |
| 75 | chr17 | 28328325 | 28335489 | ENSG00000109 | IFT20 | 90410 | 2.27009308 | -0.4205449 | 0.1573352 | -2.6729235 | 0.00751934 | 0.05956805 | C7_active |
| 76 | chr1 | 152032506 | 152047907 | ENSG00000163 | S100A11 | 6282 | 4.46737105 | -0.4191481 | 0.11940289 | -3.5103683 | 0.00044749 | 0.00588137 | C7_active |
| 77 | chr19 | 489176 | 505342 | ENSG00000099 | MADCAM1 | 8174 | 3.06942157 | -0.4099072 | 0.12941736 | -3.1673279 | 0.00153847 | 0.01670303 | C7_active |
| 78 | chr22 | 37943050 | 37953669 | ENSG00000128 | C22orf23 | 84645 | 2.52401299 | -0.4090904 | 0.1652821 | -2.4751043 | 0.01331973 | 0.09110239 | C7_active |
| 79 | chr18 | 35332212 | 35345482 | ENSG00000172 | ZNF24 | 7572 | 2.7738018 | -0.4051319 | 0.14945683 | -2.710695 | 0.00671424 | 0.05457223 | C7_active |
| 80 | chr11 | 616565 | 626078 | ENSG00000099 | CDHR5 | 53841 | 2.70719343 | -0.4032295 | 0.14208963 | -2.8378531 | 0.00454181 | 0.04029772 | C7_active |
| 81 | chr17 | 35121579 | 35130732 | ENSG00000073 | FNDC8 | 54752 | 2.76114611 | -0.4032269 | 0.15443247 | -2.6110242 | 0.00902715 | 0.06800077 | C7_active |
| 82 | chr9 | 34989641 | 34998900 | ENSG00000137 | DNAJB5 | 25822 | 2.26744809 | -0.403021 | 0.16113769 | -2.5010969 | 0.01238093 | 0.08568701 | C7_active |
| 83 | chr8 | 144466043 | 144474202 | ENSG00000167 | KIFC2 | 90990 | 2.40128382 | -0.3978191 | 0.1634863 | -2.4333482 | 0.01495991 | 0.09876775 | C7_active |
| 84 | chr16 | 30573740 | 30585771 | ENSG00000197 | ZNF785 | 146540 | 2.85237607 | -0.3962845 | 0.13980786 | -2.8344938 | 0.00458983 | 0.04059891 | C7_active |
| 85 | chr17 | 82078338 | 82098332 | ENSG00000169 | FASN | 2194 | 4.44761694 | -0.3959719 | 0.11700385 | -3.3842637 | 0.00071369 | 0.00872777 | C7_active |
| 86 | chr16 | 788046 | 800737 | ENSG00000127 | CHTF18 | 63922 | 3.15108714 | -0.394993 | 0.1440495 | -2.7420642 | 0.00610544 | 0.05050386 | C7_active |
| 87 | chr20 | 63272785 | 63289793 | ENSG00000101 | ARFGAP1 | 55738 | 5.26737238 | -0.3945211 | 0.10903098 | -3.618431 | 0.00029639 | 0.00409724 | C7_active |
| 88 | chr16 | 268301 | 275980 | ENSG00000076 | RGS11 | 8786 | 2.34013342 | -0.3896075 | 0.15794142 | -2.4667848 | 0.01363323 | 0.0927188 | C7_active |
| 89 | chr9 | 137582079 | 137590490 | ENSG00000165 | ZMYND19 | 116225 | 2.70034765 | -0.3885422 | 0.15542123 | -2.4999299 | 0.01242179 | 0.085898 | C7_active |
| 90 | chr19 | 1446302 | 1473244 | ENSG00000115 | APC2 | 10297 | 5.60857942 | -0.3863199 | 0.11243812 | -3.4358444 | 0.00059071 | 0.00745167 | C7_active |
| 91 | chr17 | 44353215 | 44363875 | ENSG00000161 | FAM171A2 | 284069 | 3.15593935 | -0.3848978 | 0.13916845 | -2.7656974 | 0.00568012 | 0.04772494 | C7_active |
| 92 | chr19 | 56176020 | 56185775 | ENSG00000197 | GALP | 85569 | 3.15420175 | -0.383219 | 0.15088656 | -2.5397821 | 0.01109215 | 0.07910767 | C7_active |
| 93 | chr7 | 100679507 | 100689448 | ENSG00000146 | GIGYF1 | 64599 | 2.58971474 | -0.3814613 | 0.15687733 | -2.4315897 | 0.01503272 | 0.09887857 | C7_active |
| 94 | chr4 | 41256413 | 41268455 | ENSG00000154 | UCHL1 | 7345 | 2.65640645 | -0.3810494 | 0.14025715 | -2.7167913 | 0.00659182 | 0.05384428 | C7_active |
| 95 | chr21 | 10521553 | 10606140 | ENSG00000274 | TPTE | 7179 | 37.7324302 | -0.3780168 | 0.04036714 | -9.3644674 | 7.64E-21 | 2.41E-19 | C7_active |
| 96 | chr9 | 88991447 | 89005010 | ENSG00000213 | S1PR3 | 1903 | 4.2902463 | -0.3770389 | 0.11942168 | -3.1572067 | 0.00159288 | 0.01720314 | C7_active |
| 97 | chr10 | 60778331 | 60794852 | ENSG00000170 | CDK1 | 983 | 3.09492596 | -0.3758188 | 0.14847176 | -2.531248 | 0.01136575 | 0.08068505 | C7_active |
| 98 | chr11 | 61873519 | 61892051 | ENSG00000221 | FADS3 | 3995 | 4.65848812 | -0.3757471 | 0.11696334 | -3.21252 | 0.00131576 | 0.01462654 | C7_active |
| 99 | chr7 | 2537877 | 2555727 | ENSG00000106 | BRAT1 | 221927 | 4.75806276 | -0.375233 | 0.11759586 | -3.1908689 | 0.00141846 | 0.01557601 | C7_active |
| 100 | chr11 | 119096503 | 119108331 | ENSG00000172 | DPAGT1 | 1798 | 3.20458645 | -0.3749995 | 0.13467124 | -2.7845554 | 0.00536012 | 0.04546009 | C7_active |
| 101 | chrX | 154348524 | 154374638 | ENSG00000196 | FLNA | 2316 | 6.36073557 | -0.3745484 | 0.09832649 | -3.8092315 | 0.0001394 | 0.00205928 | C7_active |
| 102 | chr11 | 65375192 | 65383701 | ENSG00000162 | SLC25A45 | 283130 | 2.56054329 | -0.3709575 | 0.15131514 | -2.4515554 | 0.01422403 | 0.09543139 | C7_active |
| 103 | chr17 | 49210227 | 49223225 | ENSG00000108 | ABI3 | 51225 | 4.20085867 | -0.3697098 | 0.11826127 | -3.1262117 | 0.00177074 | 0.01877245 | C7_active |
| 104 | chr16 | 58113588 | 58129450 | ENSG00000070 | C16orf80 | 29105 | 4.30647455 | -0.3696618 | 0.1077157 | -3.4318286 | 0.00059953 | 0.00753616 | C7_active |
| 105 | chr19 | 57134096 | 57145202 | ENSG00000141 | ZIM3 | 114026 | 3.07911853 | -0.3683728 | 0.13392907 | -2.7505068 | 0.00595032 | 0.04953329 | C7_active |

| | | | | | | | | | | | | | | |
|---|---|---|---|---|---|---|---|---|---|---|---|---|---|---|
| 106 | chr19 | 751126 | 764318 | ENSG00000099 | MISP | 126353 | 3.47319394 | | -0.3682863 | 0.13995749 | -2.631415 | 0.00850301 | 0.06504933 | C7_active |
| 107 | chr19 | 1177558 | 1228435 | ENSG00000118 | STK11 | 6794 | 9.4719834 | | -0.3679307 | 0.08307542 | -4.4288758 | 9.47E-06 | 0.00016737 | C7_active |
| 108 | chr20 | 63488013 | 63499315 | ENSG00000101 | EEF1A2 | 1917 | 3.14499017 | | -0.3672452 | 0.13636879 | -2.6930296 | 0.0070806 | 0.05671558 | C7_active |
| 109 | chr22 | 50185915 | 50199598 | ENSG00000170 | TRABD | 80305 | 3.50046496 | | -0.3670071 | 0.12713611 | -2.8867261 | 0.00389273 | 0.03590875 | C7_active |
| 110 | chr8 | 144428775 | 144444444 | ENSG00000160 | TONSL | 4796 | 4.52984445 | | -0.3651888 | 0.12493647 | -2.9229962 | 0.00346681 | 0.03262692 | C7_active |
| 111 | chr19 | 4044364 | 4066945 | ENSG00000178 | ZBTB7A | 51341 | 3.54328165 | | -0.3639281 | 0.13446041 | -2.7065816 | 0.00679799 | 0.05503279 | C7_active |
| 112 | chr17 | 4538897 | 4555631 | ENSG00000132 | MYBBP1A | 10514 | 4.58721496 | | -0.3626367 | 0.11355731 | -3.1934243 | 0.00140596 | 0.01548599 | C7_active |
| 113 | chr4 | 986997 | 1004506 | ENSG00000127 | IDUA | 3425 | 4.98245581 | | -0.3621925 | 0.10568826 | -3.4269894 | 0.00061031 | 0.00765173 | C7_active |
| 114 | chr19 | 2427638 | 2456996 | ENSG00000176 | LMNB2 | 84823 | 7.20708794 | | -0.3614869 | 0.09196227 | -3.9308172 | 8.47E-05 | 0.00129437 | C7_active |
| 115 | chr17 | 42968088 | 42980433 | ENSG00000267 | PTGES3L | 100885848 | 2.86567302 | | -0.3603497 | 0.14605707 | -2.4671843 | 0.01361802 | 0.09265912 | C7_active |
| 116 | chr5 | 177357837 | 177372601 | ENSG00000169 | RGS14 | 10636 | 4.95995913 | | -0.3602779 | 0.1029541 | -3.4994034 | 0.0004663 | 0.00607877 | C7_active |
| 117 | chr19 | 45692483 | 45703989 | ENSG00000011 | QPCTL | 54814 | 3.00079914 | | -0.357593 | 0.14487201 | -2.4683377 | 0.01357422 | 0.09240467 | C7_active |
| 118 | chr1 | 924880 | 944581 | ENSG00000187 | SAMD11 | 148398 | 5.43883949 | | -0.3575603 | 0.10197551 | -3.5063346 | 0.00045432 | 0.00595224 | C7_active |
| 119 | chr19 | 53333749 | 53354869 | ENSG00000213 | ZNF845 | 91664 | 5.36125144 | | -0.3572066 | 0.10290664 | -3.4711717 | 0.00051819 | 0.00667677 | C7_active |
| 120 | chr16 | 527717 | 554636 | ENSG00000103 | CAPN15 | 6650 | 5.94841826 | | -0.3566669 | 0.10449007 | -3.4134048 | 0.00064157 | 0.00799489 | C7_active |
| 121 | chr19 | 11089362 | 11133816 | ENSG00000130 | LDLR | 3949 | 8.1792899 | | -0.3510214 | 0.08551079 | -4.1049956 | 4.04E-05 | 0.00065945 | C7_active |
| 122 | chr16 | 88706463 | 88715386 | ENSG00000174 | CTU2 | 348180 | 2.96615768 | | -0.3508621 | 0.14359589 | -2.4433993 | 0.01454963 | 0.09716376 | C7_active |
| 123 | chr19 | 35550043 | 35563658 | ENSG00000105 | ATP4A | 495 | 5.02340959 | | -0.3505643 | 0.10697822 | -3.2769691 | 0.00104928 | 0.01201628 | C7_active |
| 124 | chr1 | 228082660 | 228099212 | ENSG00000143 | ARF1 | 375 | 4.06880318 | | -0.3496612 | 0.106464 | -3.284314 | 0.00102231 | 0.01179852 | C7_active |
| 125 | chr19 | 8389981 | 8404434 | ENSG00000185 | RAB11B | 9230 | 3.75578145 | | -0.3483368 | 0.12322323 | -2.8268762 | 0.00470045 | 0.04136665 | C7_active |
| 126 | chr19 | 617224 | 633604 | ENSG00000099 | POLRMT | 5442 | 4.0283601 | | -0.3477626 | 0.11877293 | -2.9279623 | 0.00341191 | 0.03221545 | C7_active |
| 127 | chr7 | 100049774 | 100065038 | ENSG00000166 | ZSCAN21 | 7589 | 4.12878789 | | -0.3470708 | 0.1017189 | -3.4120581 | 0.00064474 | 0.00802756 | C7_active |
| 128 | chr14 | 21476597 | 21499175 | ENSG00000092 | TOX4 | 9878 | 4.69262144 | | -0.347055 | 0.10918138 | -3.1787015 | 0.00147936 | 0.01612204 | C7_active |
| 129 | chr19 | 52369917 | 52385795 | ENSG00000221 | ZNF880 | 400713 | 4.15589452 | | -0.346126 | 0.11549092 | -2.9969978 | 0.00272653 | 0.02662568 | C7_active |
| 130 | chr16 | 84177847 | 84187070 | ENSG00000103 | TAF1C | 9013 | 2.8650156 | | -0.3458413 | 0.14238173 | -2.4289722 | 0.01514169 | 0.09933654 | C7_active |
| 131 | chr4 | 112636964 | 112657592 | ENSG00000174 | LARP7 | 51574 | 3.93993032 | | -0.3449647 | 0.11801373 | -2.9230894 | 0.00346577 | 0.03262692 | C7_active |
| 132 | chr17 | 81703371 | 81720539 | ENSG00000262 | SLC25A10 | NaN | 4.10003217 | | -0.3444708 | 0.11094467 | -3.1048883 | 0.00190351 | 0.01985497 | C7_active |
| 133 | chr19 | 35268978 | 35279821 | ENSG00000105 | USF2 | 7392 | 3.27715591 | | -0.3436104 | 0.13241056 | -2.5950377 | 0.00945806 | 0.07047351 | C7_active |
| 134 | chr11 | 68039016 | 68050895 | ENSG00000110 | TCIRG1 | 10312 | 3.87672782 | | -0.3391653 | 0.12151162 | -2.7912167 | 0.00525103 | 0.04479957 | C7_active |
| 135 | chr16 | 88715343 | 88785211 | ENSG00000103 | PIEZO1 | 9780 | 17.1421375 | | -0.3387336 | 0.06377259 | -5.3115868 | 1.09E-07 | 2.23E-06 | C7_active |
| 136 | chr12 | 121839527 | 121863596 | ENSG00000158 | HPD | 3242 | 5.86003384 | | -0.3371948 | 0.09988338 | -3.3758847 | 0.00073579 | 0.00889236 | C7_active |
| 137 | chr2 | 240435671 | 240468078 | ENSG00000063 | GPC1 | 2817 | 9.7737682 | | -0.3371738 | 0.07966975 | -4.2321438 | 2.31E-05 | 0.00039217 | C7_active |
| 138 | chr19 | 49085419 | 49108605 | ENSG00000104 | SNRNP70 | 6625 | 5.16800616 | | -0.3368002 | 0.10633269 | -3.1674194 | 0.00153798 | 0.01670303 | C7_active |
| 139 | chr19 | 867630 | 893218 | ENSG00000175 | MED16 | 10025 | 5.88533668 | | -0.334918 | 0.11738657 | -2.8531205 | 0.00432922 | 0.03884175 | C7_active |
| 140 | chr20 | 3208868 | 3223870 | ENSG00000125 | ITPA | 3704 | 3.79514972 | | -0.3320603 | 0.11607466 | -2.8607471 | 0.00422644 | 0.03815705 | C7_active |
| 141 | chr17 | 75087727 | 75106162 | ENSG00000170 | SLC16A5 | 9121 | 5.56102116 | | -0.3319243 | 0.09290016 | -3.572914 | 0.00035303 | 0.00478383 | C7_active |
| 142 | chr2 | 169791933 | 169812064 | ENSG00000138 | SSB | 6741 | 4.38422039 | | -0.3314051 | 0.1143074 | -2.8992442 | 0.00374063 | 0.0348176 | C7_active |
| 143 | chr19 | 54173412 | 54189882 | ENSG00000125 | MBOAT7 | 79143 | 3.92007453 | | -0.330343 | 0.12293528 | -2.687129 | 0.00720691 | 0.05753557 | C7_active |
| 144 | chr1 | 46303631 | 46316776 | ENSG00000173 | UQCRH | 7388 | 3.09576625 | | -0.3297697 | 0.13374229 | -2.4657101 | 0.0136742 | 0.09283286 | C7_active |
| 145 | chr8 | 144477969 | 144502121 | ENSG00000160 | PPP1R16A | 84988 | 7.5953678 | | -0.3279107 | 0.09083182 | -3.610086 | 0.0003061 | 0.00421517 | C7_active |
| 146 | chr19 | 10270835 | 10286615 | ENSG00000090 | ICAM1 | 3383 | 4.32872728 | | -0.325986 | 0.10883459 | -2.9952424 | 0.00274227 | 0.02673294 | C7_active |
| 147 | chr12 | 53990624 | 54030823 | ENSG00000197 | HOXC6 | 3223 | 11.0135692 | | -0.3253604 | 0.07654358 | -4.2506551 | 2.13E-05 | 0.00036455 | C7_active |
| 148 | chr19 | 647526 | 663277 | ENSG00000070 | RNF126 | 55658 | 3.81160602 | | -0.3252752 | 0.1266322 | -2.5686613 | 0.01020922 | 0.07456764 | C7_active |
| 149 | chr1 | 109466628 | 109482137 | ENSG00000143 | SYPL2 | 284612 | 5.28780734 | | -0.3251882 | 0.09743782 | -3.3373921 | 0.00084569 | 0.01005202 | C7_active |
| 150 | chr15 | 63042632 | 63071915 | ENSG00000140 | TPM1 | 7168 | 5.40801869 | | -0.324146 | 0.09334347 | -3.4726153 | 0.00051541 | 0.00665048 | C7_active |
| 151 | chr1 | 11736084 | 11754802 | ENSG00000177 | AGTRAP | 57085 | 6.229397 | | -0.3239688 | 0.09946273 | -3.2571878 | 0.00112522 | 0.01277435 | C7_active |
| 152 | chr19 | 49527618 | 49546962 | ENSG00000142 | RCN3 | 57333 | 5.14218718 | | -0.3231808 | 0.10419195 | -3.1017832 | 0.00192359 | 0.01999589 | C7_active |
| 153 | chr12 | 53985845 | 54034888 | ENSG00000273 | RP11-834C11.1 | NaN | 13.579596 | | -0.3215298 | 0.06642385 | -4.840578 | 1.29E-06 | 2.47E-05 | C7_active |
| 154 | chr21 | 45981737 | 46005050 | ENSG00000142 | COL6A1 | 1291 | 7.53897743 | | -0.3214827 | 0.09227809 | -3.4838463 | 0.00049426 | 0.00640277 | C7_active |
| 155 | chr16 | 89720400 | 89740903 | ENSG00000158 | ZNF276 | 92822 | 5.06875294 | | -0.3197923 | 0.11391248 | -2.8073508 | 0.00499508 | 0.04322874 | C7_active |
| 156 | chr9 | 134641774 | 134844843 | ENSG00000130 | COL5A1 | 1289 | 67.006659 | | -0.3172638 | 0.0380317 | -8.3420897 | 7.30E-17 | 2.01E-15 | C7_active |
| 157 | chr16 | 31214021 | 31231537 | ENSG00000177 | TRIM72 | 493829 | 5.38102401 | | -0.3164154 | 0.10390714 | -3.0451749 | 0.00232545 | 0.02334843 | C7_active |
| 158 | chr17 | 78174075 | 78187233 | ENSG00000167 | TK1 | 7083 | 3.86701577 | | -0.3161605 | 0.12263315 | -2.5781 | 0.00993452 | 0.07300508 | C7_active |
| 159 | chr3 | 134355874 | 134375479 | ENSG00000114 | AMOTL2 | 51421 | 4.0349178 | | -0.3156727 | 0.12036138 | -2.6227074 | 0.00872342 | 0.06623181 | C7_active |

| | | | | | | | | | | | | | | |
|---|---|---|---|---|---|---|---|---|---|---|---|---|---|---|
| 160 | chr17 | 39626740 | 39636626 | ENSG00000131 | PPP1R1B | 84152 | 3.37799782 | | -0.3151122 | 0.12160917 | -2.5911877 | 0.00956453 | 0.07108319 | C7_active |
| 161 | chr12 | 132725503 | 132761888 | ENSG00000176 | ANKLE2 | 23141 | 7.96893855 | | -0.3143438 | 0.08377609 | -3.7521898 | 0.0001753 | 0.00254013 | C7_active |
| 162 | chr22 | 50200979 | 50217616 | ENSG00000073 | SELO | 83642 | 4.36840909 | | -0.3127952 | 0.11509608 | -2.7176876 | 0.00657399 | 0.05376276 | C7_active |
| 163 | chr14 | 24213955 | 24232352 | ENSG00000255 | NEDD8-MDP1 | 100528064 | 3.46977584 | | -0.3125405 | 0.12541297 | -2.492091 | 0.01269935 | 0.08739821 | C7_active |
| 164 | chr19 | 57921884 | 57935393 | ENSG00000196 | ZNF418 | 147686 | 4.46474158 | | -0.3125042 | 0.10627537 | -2.9405137 | 0.00327669 | 0.03110155 | C7_active |
| 165 | chrX | 154531391 | 154547572 | ENSG00000160 | G6PD | 2539 | 3.97438445 | | -0.3110802 | 0.11913533 | -2.6111499 | 0.00902383 | 0.06800077 | C7_active |
| 166 | chr19 | 4815932 | 4831704 | ENSG00000127 | TICAM1 | 148022 | 4.0747103 | | -0.3107538 | 0.12197627 | -2.5476579 | 0.01084488 | 0.07786924 | C7_active |
| 167 | chr1 | 150268200 | 150280916 | ENSG00000118 | C1orf54 | 79630 | 3.50444531 | | -0.3104989 | 0.12381381 | -2.5077894 | 0.01214891 | 0.08447661 | C7_active |
| 168 | chr17 | 81228277 | 81239091 | ENSG00000167 | ENTHD2 | 146705 | 3.27452649 | | -0.3094981 | 0.12207818 | -2.5352447 | 0.01123688 | 0.07996709 | C7_active |
| 169 | chr2 | 107826937 | 107890841 | ENSG00000196 | RGPD4 | 285190 | 5.79795278 | | -0.3085548 | 0.09585725 | -3.2188988 | 0.00128684 | 0.01437153 | C7_active |
| 170 | chr1 | 153929501 | 153946696 | ENSG00000198 | DENND4B | 9909 | 6.15532985 | | -0.3082066 | 0.09621782 | -3.2032179 | 0.00135901 | 0.01502624 | C7_active |
| 171 | chr6 | 34757406 | 34773794 | ENSG00000124 | SNRPC | 6631 | 3.60312951 | | -0.3076896 | 0.11914368 | -2.5825085 | 0.0098085 | 0.07248533 | C7_active |
| 172 | chr1 | 228735077 | 228746669 | ENSG00000116 | RHOU | 58480 | 3.84237927 | | -0.3074021 | 0.11170789 | -2.7518382 | 0.00592618 | 0.04941796 | C7_active |
| 173 | chr3 | 53224707 | 53256052 | ENSG00000163 | TKT | 7086 | 7.15059606 | | -0.307148 | 0.09372314 | -3.2771839 | 0.00104848 | 0.01201628 | C7_active |
| 174 | chr15 | 90352245 | 90369146 | ENSG00000196 | ZNF774 | 342132 | 4.00680561 | | -0.3070606 | 0.11879865 | -2.5847142 | 0.00974597 | 0.07217127 | C7_active |
| 175 | chr12 | 6870935 | 6889358 | ENSG00000111 | SPSB2 | 84727 | 4.68322586 | | -0.3063387 | 0.10992184 | -2.7868779 | 0.00532185 | 0.04526871 | C7_active |
| 176 | chr17 | 7246829 | 7262089 | ENSG00000262 | RP1-4G17.5 | NaN | 4.09600713 | | -0.3061057 | 0.11806671 | -2.5926505 | 0.00952395 | 0.07085465 | C7_active |
| 177 | chr19 | 4472287 | 4502211 | ENSG00000167 | HDGFRP2 | 84717 | 7.1924897 | | -0.3051796 | 0.09251776 | -3.2986056 | 0.00097166 | 0.01132534 | C7_active |
| 178 | chr17 | 73283624 | 73312175 | ENSG00000179 | CDC42EP4 | 23580 | 7.04393611 | | -0.3040351 | 0.09240741 | -3.290159 | 0.00100131 | 0.01163324 | C7_active |
| 179 | chr17 | 50634777 | 50692252 | ENSG00000108 | ABCC3 | 8714 | 16.2050779 | | -0.3036685 | 0.05725868 | -5.3034487 | 1.14E-07 | 2.32E-06 | C7_active |
| 180 | chr19 | 50476400 | 50490870 | ENSG00000161 | EMC10 | 284361 | 4.172862 | | -0.3025809 | 0.11534044 | -2.6233722 | 0.00870641 | 0.06616582 | C7_active |
| 181 | chr12 | 52585589 | 52601538 | ENSG00000170 | KRT72 | 140807 | 5.83087893 | | -0.3018432 | 0.09465737 | -3.1887979 | 0.00142866 | 0.01567609 | C7_active |
| 182 | chr8 | 11842524 | 11869448 | ENSG00000164 | CTSB | 1508 | 6.40432157 | | -0.3011805 | 0.09438547 | -3.1909627 | 0.001418 | 0.01557601 | C7_active |
| 183 | chr19 | 17555594 | 17583162 | ENSG00000130 | COLGALT1 | 79709 | 6.78641652 | | -0.3006833 | 0.09767957 | -3.0782616 | 0.00208212 | 0.02129079 | C7_active |
| 184 | chr16 | 1493344 | 1510457 | ENSG00000100 | TELO2 | 9894 | 4.77703532 | | -0.3004934 | 0.10733237 | -2.7996534 | 0.00511575 | 0.04390409 | C7_active |
| 185 | chr7 | 99374249 | 99394801 | ENSG00000130 | ARPC1B | 10095 | 4.24169369 | | -0.2984036 | 0.11301618 | -2.6403619 | 0.00828175 | 0.06385366 | C7_active |
| 186 | chr19 | 19865886 | 19894674 | ENSG00000256 | ZNF253 | 56242 | 5.2508846 | | -0.2976941 | 0.0946865 | -3.1439974 | 0.00166657 | 0.01785186 | C7_active |
| 187 | chr22 | 17592136 | 17628818 | ENSG00000131 | ATP6V1E1 | 529 | 8.41786638 | | -0.2963121 | 0.08708161 | -3.4026944 | 0.00066725 | 0.00825785 | C7_active |
| 188 | chr1 | 10398592 | 10420144 | ENSG00000142 | PGD | 5226 | 5.46660857 | | -0.2956762 | 0.1026405 | -2.8806977 | 0.00396796 | 0.03639316 | C7_active |
| 189 | chr19 | 58126252 | 58155110 | ENSG00000181 | ZNF329 | 79673 | 5.97353554 | | -0.2943662 | 0.08888915 | -3.3116101 | 0.00092761 | 0.01088222 | C7_active |
| 190 | chr11 | 65961689 | 65979828 | ENSG00000175 | SART1 | 9092 | 5.17334049 | | -0.2943366 | 0.10490313 | -2.8057946 | 0.00501927 | 0.04335998 | C7_active |
| 191 | chr19 | 50311937 | 50333515 | ENSG00000131 | KCNC3 | 3748 | 6.02575244 | | -0.2941301 | 0.10255568 | -2.868004 | 0.0041307 | 0.03745691 | C7_active |
| 192 | chr1 | 1253909 | 1273885 | ENSG00000160 | UBE2J2 | 118424 | 4.47618063 | | -0.2928474 | 0.11689969 | -2.5051173 | 0.01224108 | 0.0849821 | C7_active |
| 193 | chr19 | 56087366 | 56121280 | ENSG00000142 | ZNF787 | 126208 | 9.51990521 | | -0.2922599 | 0.08083042 | -3.6157175 | 0.00029952 | 0.00413643 | C7_active |
| 194 | chr19 | 19119169 | 19138513 | ENSG00000064 | TMEM161A | 54929 | 4.69539875 | | -0.2909307 | 0.10892704 | -2.6708763 | 0.00756535 | 0.05982986 | C7_active |
| 195 | chr19 | 19900913 | 19935575 | ENSG00000184 | ZNF93 | 81931 | 7.63871173 | | -0.2907884 | 0.08089006 | -3.5948591 | 0.00032457 | 0.00443145 | C7_active |
| 196 | chr7 | 5431335 | 5513798 | ENSG00000155 | FBXL18 | 80028 | 21.44385 | | -0.2904975 | 0.05154653 | -5.6356356 | 1.74E-08 | 3.74E-07 | C7_active |
| 197 | chr19 | 48485271 | 48513189 | ENSG00000142 | LMTK3 | 114783 | 6.53034488 | | -0.2901162 | 0.0882433 | -3.2876854 | 0.00101015 | 0.01168884 | C7_active |
| 198 | chr19 | 925781 | 975934 | ENSG00000116 | ARID3A | 1820 | 11.4195475 | | -0.289355 | 0.07779361 | -3.7195209 | 0.0001996 | 0.00285784 | C7_active |
| 199 | chr17 | 39980797 | 39997960 | ENSG00000108 | PSMD3 | 5709 | 5.54358141 | | -0.2888861 | 0.1001547 | -2.8843989 | 0.00392162 | 0.03602202 | C7_active |
| 200 | chr1 | 228393673 | 228406840 | ENSG00000154 | TRIM11 | 81559 | 4.13565329 | | -0.2881506 | 0.1178409 | -2.4452514 | 0.01447512 | 0.0968007 | C7_active |
| 201 | chr11 | 46332905 | 46380554 | ENSG00000149 | DGKZ | 8525 | 14.3529581 | | -0.2862791 | 0.06820717 | -4.1971994 | 2.70E-05 | 0.00045358 | C7_active |
| 202 | chr19 | 11131520 | 11155808 | ENSG00000161 | SPC24 | 147841 | 5.02100751 | | -0.2853607 | 0.1037815 | -2.7496301 | 0.00596626 | 0.04958379 | C7_active |
| 203 | chr7 | 1838586 | 1849931 | ENSG00000176 | AC110781.3 | NaN | 4.53061495 | | -0.2853274 | 0.11128782 | -2.5638689 | 0.01035126 | 0.07549039 | C7_active |
| 204 | chr19 | 54923509 | 54966312 | ENSG00000167 | NLRP7 | 199713 | 11.3147432 | | -0.2841265 | 0.06998345 | -4.0599096 | 4.91E-05 | 0.00078645 | C7_active |
| 205 | chr1 | 224175756 | 224193441 | ENSG00000143 | DEGS1 | 8560 | 3.71499852 | | -0.2839422 | 0.11665297 | -2.434076 | 0.01492986 | 0.09869727 | C7_active |
| 206 | chr16 | 89171767 | 89195492 | ENSG00000129 | CDH15 | 1013 | 6.584242 | | -0.2839243 | 0.09578255 | -2.9642594 | 0.00303412 | 0.02918346 | C7_active |
| 207 | chr17 | 78110458 | 78132407 | ENSG00000141 | TMC6 | 11322 | 7.13402299 | | -0.2833474 | 0.08893427 | -3.1860321 | 0.00144239 | 0.01580268 | C7_active |
| 208 | chr2 | 156324432 | 156342348 | ENSG00000153 | NR4A2 | 4929 | 4.33553544 | | -0.2829967 | 0.10678902 | -2.6500546 | 0.00804788 | 0.06251847 | C7_active |
| 209 | chr1 | 2184461 | 2212720 | ENSG00000162 | C1orf86 | 199990 | 7.6859604 | | -0.2829922 | 0.09091627 | -3.1126683 | 0.00185404 | 0.01938476 | C7_active |
| 210 | chr5 | 132996985 | 133026604 | ENSG00000155 | ZCCHC10 | 54819 | 6.12942632 | | -0.2821176 | 0.09295285 | -3.0350611 | 0.00240487 | 0.02399544 | C7_active |
| 211 | chr6 | 30626842 | 30646823 | ENSG00000137 | ATAT1 | 79969 | 5.20882775 | | -0.2819199 | 0.09867872 | -2.8569475 | 0.00427737 | 0.03852034 | C7_active |
| 212 | chr18 | 57600656 | 57622213 | ENSG00000134 | NARS | 4677 | 4.48432682 | | -0.2806428 | 0.10985974 | -2.554555 | 0.01063236 | 0.07707261 | C7_active |
| 213 | chr7 | 3140234 | 3174654 | ENSG00000217 | AC091801.1 | NaN | 10.5286501 | | -0.2795636 | 0.0733063 | -3.8136374 | 0.00013694 | 0.00203122 | C7_active |

| | | | | | | | | | | | | | |
|---|---|---|---|---|---|---|---|---|---|---|---|---|---|
| 214 | chr19 | 1040101 | 1065572 | ENSG00000064 | ABCA7 | 10347 | 5.87176816 | -0.279476 | 0.1010363 | -2.7660948 | 0.0056732 | 0.04769461 | C7_active |
| 215 | chr19 | 40778216 | 40796944 | ENSG00000167 | RAB4B | 53916 | 5.1605678 | -0.2794681 | 0.10111111 | -2.7639702 | 0.00571028 | 0.04793984 | C7_active |
| 216 | chr19 | 45710629 | 45730904 | ENSG00000177 | FBXO46 | 23403 | 4.44722017 | -0.2790246 | 0.11468519 | -2.4329609 | 0.01497592 | 0.09876775 | C7_active |
| 217 | chr18 | 12407896 | 12432238 | ENSG00000141 | SLMO1 | 10650 | 6.20207427 | -0.2780666 | 0.09715007 | -2.8622381 | 0.00420661 | 0.03800179 | C7_active |
| 218 | chr19 | 7476875 | 7497449 | ENSG00000104 | PEX11G | 92960 | 5.70209997 | -0.2768996 | 0.09907334 | -2.7948954 | 0.00519165 | 0.04442328 | C7_active |
| 219 | chr22 | 49994513 | 50012659 | ENSG00000188 | IL17REL | 400935 | 5.33981803 | -0.276306 | 0.10239811 | -2.6983503 | 0.00696841 | 0.05594642 | C7_active |
| 220 | chr14 | 34515929 | 34539711 | ENSG00000129 | EAPP | 55837 | 5.07846526 | -0.2762744 | 0.10408641 | -2.6542797 | 0.00794779 | 0.06204183 | C7_active |
| 221 | chr16 | 70661204 | 70686066 | ENSG00000132 | MTSS1L | 92154 | 5.44732518 | -0.2762103 | 0.09779865 | -2.8242753 | 0.00473877 | 0.04160994 | C7_active |
| 222 | chr9 | 128787204 | 128810432 | ENSG00000107 | TBC1D13 | 54662 | 7.34056744 | -0.275853 | 0.08869956 | -3.1099703 | 0.00187106 | 0.01953437 | C7_active |
| 223 | chr16 | 69706996 | 69726951 | ENSG00000181 | NQO1 | 1728 | 4.22685461 | -0.2756426 | 0.11238577 | -2.4526466 | 0.01418096 | 0.09518671 | C7_active |
| 224 | chr19 | 16067021 | 16103005 | ENSG00000167 | TPM4 | 7171 | 6.9007341 | -0.2755617 | 0.0919609 | -2.9965092 | 0.0027309 | 0.02664013 | C7_active |
| 225 | chr19 | 19785839 | 19821751 | ENSG00000081 | ZNF506 | 440515 | 8.09478832 | -0.2752433 | 0.08151737 | -3.3764986 | 0.00073415 | 0.00889236 | C7_active |
| 226 | chr2 | 190880827 | 190965552 | ENSG00000115 | GLS | 2744 | 15.058716 | -0.2750203 | 0.06442699 | -4.2687125 | 1.97E-05 | 0.00033867 | C7_active |
| 227 | chr1 | 15572353 | 15585110 | ENSG00000116 | AGMAT | 79814 | 4.01911521 | -0.2745815 | 0.10498986 | -2.615314 | 0.00891455 | 0.06739901 | C7_active |
| 228 | chr22 | 24219654 | 24245142 | ENSG00000099 | GGT5 | 2687 | 6.88440047 | -0.2743692 | 0.0849677 | -3.2291001 | 0.0012418 | 0.01393333 | C7_active |
| 229 | chr19 | 2328615 | 2354806 | ENSG00000005 | SPPL2B | 56928 | 6.98210952 | -0.274343 | 0.10266976 | -2.6720913 | 0.00753801 | 0.05968319 | C7_active |
| 230 | chr19 | 2389771 | 2426239 | ENSG00000178 | TMPRSS9 | 360200 | 8.68053793 | -0.2730124 | 0.07648549 | -3.5694672 | 0.00035771 | 0.00483812 | C7_active |
| 231 | chr19 | 4153601 | 4173054 | ENSG00000060 | CREB3L3 | 84699 | 5.74131278 | -0.2726758 | 0.09725125 | -2.8038284 | 0.00504998 | 0.04350621 | C7_active |
| 232 | chr19 | 3185563 | 3209575 | ENSG00000125 | NCLN | 56926 | 6.87274221 | -0.27181 | 0.09351109 | -2.9067135 | 0.00365248 | 0.03415136 | C7_active |
| 233 | chr22 | 43110748 | 43129712 | ENSG00000100 | BIK | 638 | 6.29283362 | -0.2713024 | 0.09597143 | -2.8269078 | 0.00469999 | 0.04136665 | C7_active |
| 234 | chr1 | 3778558 | 3796504 | ENSG00000130 | LRRC47 | 57470 | 6.12410147 | -0.2692126 | 0.09353207 | -2.8782917 | 0.00399835 | 0.03655565 | C7_active |
| 235 | chr1 | 25800176 | 25818224 | ENSG00000162 | SEPN1 | 57190 | 6.00241413 | -0.26851 | 0.09627333 | -2.7890387 | 0.00528647 | 0.04504751 | C7_active |
| 236 | chr12 | 6946468 | 6961316 | ENSG00000111 | PTPN6 | 5777 | 5.08543912 | -0.2682308 | 0.10579413 | -2.5354034 | 0.01123179 | 0.07996709 | C7_active |
| 237 | chr7 | 73536356 | 73558002 | ENSG00000106 | BCL7B | 9275 | 4.89828418 | -0.2679789 | 0.10366242 | -2.5851117 | 0.00973475 | 0.07215821 | C7_active |
| 238 | chr22 | 20979462 | 20999038 | ENSG00000099 | LZTR1 | 8216 | 6.77934858 | -0.2679752 | 0.0890127 | -3.0105282 | 0.00260794 | 0.02564886 | C7_active |
| 239 | chr16 | 2177180 | 2196525 | ENSG00000167 | CASKIN1 | 57524 | 5.65595786 | -0.2661002 | 0.10466693 | -2.5423525 | 0.01101091 | 0.07882585 | C7_active |
| 240 | chr16 | 4882507 | 4960741 | ENSG00000118 | PPL | 5493 | 22.4104266 | -0.2649342 | 0.05800476 | -4.567456 | 4.94E-06 | 8.99E-05 | C7_active |
| 241 | chr19 | 12699194 | 12724011 | ENSG00000105 | TNPO2 | 30000 | 5.74959449 | -0.2646256 | 0.09828114 | -2.692537 | 0.00709107 | 0.0567364 | C7_active |
| 242 | chr19 | 708939 | 748329 | ENSG00000099 | PALM | 5064 | 10.6382935 | -0.2641617 | 0.07676936 | -3.4409788 | 0.00057961 | 0.00733703 | C7_active |
| 243 | chr17 | 1650629 | 1684882 | ENSG00000174 | PRPF8 | 10594 | 9.3001094 | -0.2629651 | 0.07441231 | -3.5338923 | 0.00040949 | 0.00543146 | C7_active |
| 244 | chr22 | 36281281 | 36388018 | ENSG00000100 | MYH9 | 4627 | 19.9379945 | -0.2617767 | 0.05748876 | -4.553528 | 5.28E-06 | 9.59E-05 | C7_active |
| 245 | chrX | 153907367 | 153934999 | ENSG00000089 | ARHGAP4 | 393 | 8.4852602 | -0.2610438 | 0.08249192 | -3.1644769 | 0.00155362 | 0.01684218 | C7_active |
| 246 | chr17 | 74466416 | 74484796 | ENSG00000167 | CD300A | 11314 | 6.91915935 | -0.2610152 | 0.09078262 | -2.8751667 | 0.00403814 | 0.03684933 | C7_active |
| 247 | chr22 | 41245611 | 41286251 | ENSG00000100 | RANGAP1 | 5905 | 10.0750998 | -0.2610046 | 0.07956566 | -3.2803678 | 0.00103672 | 0.01191978 | C7_active |
| 248 | chr1 | 151540305 | 151583583 | ENSG00000143 | TUFT1 | 7286 | 10.3772935 | -0.2609752 | 0.07541764 | -3.4604001 | 0.00053937 | 0.00688812 | C7_active |
| 249 | chr1 | 20588948 | 20618908 | ENSG00000158 | CDA | 978 | 8.32551534 | -0.2605844 | 0.0772954 | -3.3712791 | 0.0007482 | 0.00900463 | C7_active |
| 250 | chr8 | 143915147 | 143976734 | ENSG00000178 | PLEC | 5339 | 12.542723 | -0.2602921 | 0.07016176 | -3.7098853 | 0.00020735 | 0.00294834 | C7_active |
| 251 | chr11 | 842808 | 867116 | ENSG00000214 | TSPAN4 | 7106 | 5.9384921 | -0.2602887 | 0.10554742 | -2.4660828 | 0.01365998 | 0.09283286 | C7_active |
| 252 | chr15 | 89316305 | 89334861 | ENSG00000140 | POLG | 5428 | 5.77789586 | -0.2601063 | 0.09581889 | -2.7145618 | 0.00663635 | 0.0541193 | C7_active |
| 253 | chr19 | 3777970 | 3802129 | ENSG00000007 | MATK | 4145 | 6.81633051 | -0.2599981 | 0.08788822 | -2.958282 | 0.00309359 | 0.0296571 | C7_active |
| 254 | chr17 | 81035122 | 81117432 | ENSG00000175 | BAIAP2 | 10458 | 22.9370474 | -0.2594322 | 0.05801682 | -4.4716719 | 7.76E-06 | 0.00013849 | C7_active |
| 255 | chr1 | 9292880 | 9369532 | ENSG00000171 | SPSB1 | 80176 | 20.7557894 | -0.258856 | 0.05817396 | -4.4496892 | 8.60E-06 | 0.00015269 | C7_active |
| 256 | chr7 | 99472841 | 99503650 | ENSG00000198 | ZNF789 | 285989 | 7.73653739 | -0.2584672 | 0.08743199 | -2.9562089 | 0.00311446 | 0.02981693 | C7_active |
| 257 | chr9 | 127505339 | 127578989 | ENSG00000136 | FAM129B | 64855 | 17.0406494 | -0.2583548 | 0.06605008 | -3.911498 | 9.17E-05 | 0.00139651 | C7_active |
| 258 | chr16 | 2682475 | 2709030 | ENSG00000167 | KCTD5 | 54442 | 6.61834383 | -0.257519 | 0.0966117 | -2.6655052 | 0.00768728 | 0.06049953 | C7_active |
| 259 | chr19 | 18153158 | 18178117 | ENSG00000268 | PIK3R2 | NaN | 6.16577077 | -0.2562698 | 0.0971275 | -2.6384883 | 0.00832766 | 0.06410307 | C7_active |
| 260 | chr20 | 833715 | 857463 | ENSG00000125 | FAM110A | 83541 | 6.9229949 | -0.2557691 | 0.08844411 | -2.8918727 | 0.00382953 | 0.03550319 | C7_active |
| 261 | chr9 | 128882112 | 128918039 | ENSG00000136 | LRRC8A | 56262 | 9.34721556 | -0.2550886 | 0.07854594 | -3.2476358 | 0.00116368 | 0.01314887 | C7_active |
| 262 | chr5 | 1317744 | 1345099 | ENSG00000049 | CLPTM1L | 81037 | 8.36506873 | -0.2548385 | 0.08716198 | -2.9237348 | 0.00345859 | 0.03259215 | C7_active |
| 263 | chr12 | 104215779 | 104350305 | ENSG00000198 | TXNRD1 | 7296 | 26.1783516 | -0.2546723 | 0.04554163 | -5.5920764 | 2.24E-08 | 4.79E-07 | C7_active |
| 264 | chr19 | 45349837 | 45370918 | ENSG00000104 | ERCC2 | 2068 | 5.77140682 | -0.2531181 | 0.09887779 | -2.5599087 | 0.01046997 | 0.07608668 | C7_active |
| 265 | chr19 | 12014732 | 12035741 | ENSG00000197 | ZNF433 | 163059 | 4.6668101 | -0.251942 | 0.10285198 | -2.4495592 | 0.01430312 | 0.09591738 | C7_active |
| 266 | chr7 | 497258 | 520296 | ENSG00000197 | PDGFA | 5154 | 6.09505871 | -0.2508484 | 0.10111595 | -2.4807997 | 0.0131088 | 0.08983017 | C7_active |
| 267 | chr19 | 49591643 | 49626439 | ENSG00000126 | PRR12 | 57479 | 8.52700161 | -0.2506775 | 0.08249552 | -3.0386797 | 0.00237617 | 0.02377492 | C7_active |

| | | | | | | | | | | | | | |
|---|---|---|---|---|---|---|---|---|---|---|---|---|---|
| 268 | chr17 | 74326210 | 74355820 | ENSG00000196 | KIF19 | 124602 | 11.0294095 | -0.2504822 | 0.07520065 | -3.3308511 | 0.00086581 | 0.01025822 | C7_active |
| 269 | chr19 | 47713343 | 47743134 | ENSG00000024 | EHD2 | 30846 | 5.53763017 | -0.2493763 | 0.09143483 | -2.727367 | 0.0063842 | 0.05238894 | C7_active |
| 270 | chr16 | 85611409 | 85676204 | ENSG00000131 | GSE1 | 23199 | 19.7443559 | -0.2493436 | 0.0552983 | -4.5090655 | 6.51E-06 | 0.00011735 | C7_active |
| 271 | chr11 | 61792980 | 61867354 | ENSG00000134 | FADS2 | 9415 | 21.5071356 | -0.2489852 | 0.056877 | -4.3776076 | 1.20E-05 | 0.00020944 | C7_active |
| 272 | chr9 | 128068201 | 128109245 | ENSG00000148 | SLC25A25 | 114789 | 9.48536707 | -0.2486402 | 0.07093094 | -3.5053839 | 0.00045595 | 0.00596541 | C7_active |
| 273 | chr8 | 27596917 | 27615031 | ENSG00000120 | CLU | 1191 | 5.14743032 | -0.2485407 | 0.09953796 | -2.4969436 | 0.01252689 | 0.08650032 | C7_active |
| 274 | chr11 | 118893875 | 118925608 | ENSG00000186 | BCL9L | 283149 | 7.71063254 | -0.2481701 | 0.08703503 | -2.8513813 | 0.00435297 | 0.03903058 | C7_active |
| 275 | chr10 | 44370165 | 44386493 | ENSG00000107 | CXCL12 | 6387 | 5.46827475 | -0.2480941 | 0.09879201 | -2.5112768 | 0.01202953 | 0.08399614 | C7_active |
| 276 | chr15 | 74409289 | 74434467 | ENSG00000138 | SEMA7A | 8482 | 6.03503416 | -0.2477432 | 0.09278702 | -2.6700198 | 0.00758468 | 0.05992103 | C7_active |
| 277 | chr16 | 1363205 | 1414751 | ENSG00000059 | UNKL | 64718 | 13.4705649 | -0.2474818 | 0.06584685 | -3.7584448 | 0.00017097 | 0.0024925 | C7_active |
| 278 | chr19 | 1815246 | 1848463 | ENSG00000079 | REXO1 | 57455 | 8.49720518 | -0.2473594 | 0.08588172 | -2.8802331 | 0.00397381 | 0.03642367 | C7_active |
| 279 | chr14 | 105140981 | 105168824 | ENSG00000184 | JAG2 | 3714 | 9.24709907 | -0.246994 | 0.0789777 | -3.1273887 | 0.00176367 | 0.01871121 | C7_active |
| 280 | chr11 | 1012821 | 1036706 | ENSG00000184 | MUC6 | 4588 | 11.55092 | -0.2456979 | 0.08060731 | -3.0480844 | 0.00230305 | 0.02317195 | C7_active |
| 281 | chr16 | 89947925 | 89968060 | ENSG00000140 | DEF8 | 54849 | 6.31370845 | -0.2436167 | 0.09659101 | -2.522147 | 0.0116641 | 0.08218299 | C7_active |
| 282 | chr17 | 81976807 | 82017406 | ENSG00000169 | ASPSCR1 | 79058 | 10.5885311 | -0.2423227 | 0.07676121 | -3.1568375 | 0.0015949 | 0.01721205 | C7_active |
| 283 | chr19 | 47019820 | 47045775 | ENSG00000130 | NPAS1 | 4861 | 6.59592727 | -0.2418713 | 0.09036196 | -2.6766933 | 0.00743527 | 0.05906216 | C7_active |
| 284 | chr19 | 8413270 | 8439017 | ENSG00000099 | 37316 | 51257 | 6.12561462 | -0.2415583 | 0.09772174 | -2.471899 | 0.01343975 | 0.0917492 | C7_active |
| 285 | chr19 | 52555822 | 52587174 | ENSG00000167 | ZNF701 | 55762 | 7.39977885 | -0.2396794 | 0.08539107 | -2.8068439 | 0.00500295 | 0.04326624 | C7_active |
| 286 | chr19 | 1941149 | 1981338 | ENSG00000133 | CSNK1G2 | 1455 | 10.0174984 | -0.2396281 | 0.08459327 | -2.8327086 | 0.00461554 | 0.0407763 | C7_active |
| 287 | chr9 | 114611206 | 114646422 | ENSG00000157 | C9orf91 | 203197 | 11.0519111 | -0.2384669 | 0.06900682 | -3.455701 | 0.00054886 | 0.00699692 | C7_active |
| 288 | chr5 | 95885098 | 95962071 | ENSG00000118 | ELL2 | 22936 | 14.6334683 | -0.2376451 | 0.06326108 | -3.7565763 | 0.00017225 | 0.00250435 | C7_active |
| 289 | chrX | 24149173 | 24216255 | ENSG00000005 | ZFX | 7543 | 13.6344102 | -0.2376192 | 0.06242851 | -3.8062608 | 0.00014108 | 0.00208203 | C7_active |
| 290 | chr4 | 1211448 | 1249953 | ENSG00000159 | CTBP1 | 1487 | 10.9018357 | -0.2374702 | 0.07519722 | -3.1579646 | 0.00158875 | 0.01717134 | C7_active |
| 291 | chr1 | 15756607 | 15786594 | ENSG00000162 | FBLIM1 | 54751 | 7.67302158 | -0.2369201 | 0.08674602 | -2.7311924 | 0.00631056 | 0.05190284 | C7_active |
| 292 | chr17 | 78870910 | 78903217 | ENSG00000178 | DDC8 | 100653515 | 8.19756099 | -0.2367815 | 0.08478412 | -2.792758 | 0.00522608 | 0.04466483 | C7_active |
| 293 | chr20 | 53567065 | 53609907 | ENSG00000171 | ZNF217 | 7764 | 8.5987583 | -0.236561 | 0.08371001 | -2.8259582 | 0.00471394 | 0.04144244 | C7_active |
| 294 | chr12 | 56162983 | 56189567 | ENSG00000139 | SMARCC2 | 6601 | 7.39481578 | -0.2361649 | 0.08475245 | -2.7865265 | 0.00532763 | 0.04528146 | C7_active |
| 295 | chr7 | 6409126 | 6484190 | ENSG00000164 | DAGLB | 221955 | 16.5699142 | -0.2360626 | 0.05804529 | -4.0668685 | 4.76E-05 | 0.00076675 | C7_active |
| 296 | chr19 | 17871973 | 17895174 | ENSG00000105 | SLC5A5 | 6528 | 6.36203467 | -0.2353891 | 0.09126536 | -2.5791727 | 0.00990373 | 0.07285302 | C7_active |
| 297 | chr22 | 37848868 | 37889407 | ENSG00000100 | EIF3L | 51386 | 8.63919049 | -0.23511 | 0.08226528 | -2.8579498 | 0.00426388 | 0.03844691 | C7_active |
| 298 | chr18 | 11882622 | 11909223 | ENSG00000154 | MPPE1 | 65258 | 6.43849741 | -0.2351092 | 0.08811933 | -2.6680781 | 0.00762865 | 0.06017817 | C7_active |
| 299 | chrX | 1336616 | 1382689 | ENSG00000185 | IL3RA | 3563 | 9.20614985 | -0.2345705 | 0.08611292 | -2.7239876 | 0.00644989 | 0.05286786 | C7_active |
| 300 | chr12 | 131828393 | 131851783 | ENSG00000198 | MMP17 | 4326 | 6.96916066 | -0.2343566 | 0.08741837 | -2.6808625 | 0.00734327 | 0.05849461 | C7_active |
| 301 | chr9 | 128832942 | 128882494 | ENSG00000171 | CCBL1 | 883 | 12.8355379 | -0.2333934 | 0.07036367 | -3.3169588 | 0.00091003 | 0.01069341 | C7_active |
| 302 | chr7 | 73799542 | 73832693 | ENSG00000189 | CLDN4 | 1364 | 8.71606598 | -0.2331637 | 0.07719545 | -3.0204325 | 0.00252414 | 0.02497807 | C7_active |
| 303 | chr19 | 12577961 | 12640098 | ENSG00000188 | ZNF490 | 57474 | 12.4143093 | -0.23314 | 0.06882548 | -3.3874079 | 0.00070556 | 0.00865772 | C7_active |
| 304 | chr19 | 54953130 | 55001142 | ENSG00000022 | NLRP2 | 55655 | 12.9677053 | -0.2326236 | 0.06570146 | -3.5406152 | 0.0003992 | 0.0053145 | C7_active |
| 305 | chr1 | 6181269 | 6209389 | ENSG00000116 | RPL22 | 6146 | 5.95975562 | -0.2326176 | 0.09586359 | -2.4265477 | 0.01524325 | 0.09967218 | C7_active |
| 306 | chr7 | 73593194 | 73624543 | ENSG00000009 | MLXIPL | 51085 | 8.62660299 | -0.2325443 | 0.08855345 | -2.6260334 | 0.00863863 | 0.06576125 | C7_active |
| 307 | chr16 | 66720893 | 66751798 | ENSG00000135 | DYNC1LI2 | 1783 | 7.36605777 | -0.2324381 | 0.08247836 | -2.8181705 | 0.00482982 | 0.04220381 | C7_active |
| 308 | chr7 | 5527151 | 5563784 | ENSG00000075 | ACTB | 60 | 8.26669537 | -0.2322159 | 0.08333889 | -2.7864045 | 0.00532963 | 0.04528146 | C7_active |
| 309 | chr8 | 30678061 | 30727926 | ENSG00000104 | GSR | 2936 | 10.95224 | -0.2321973 | 0.06982313 | -3.3255069 | 0.00088258 | 0.01043035 | C7_active |
| 310 | chr19 | 45407333 | 45478828 | ENSG00000012 | ERCC1 | 2067 | 14.9210373 | -0.2316871 | 0.06202506 | -3.7353784 | 0.00018743 | 0.00269971 | C7_active |
| 311 | chr19 | 1107636 | 1174283 | ENSG00000064 | SBNO2 | 22904 | 15.8114744 | -0.2316258 | 0.06856633 | -3.3781282 | 0.00072981 | 0.00885724 | C7_active |
| 312 | chr16 | 1795620 | 1827194 | ENSG00000063 | HAGH | 3029 | 8.4579846 | -0.2312534 | 0.08887414 | -2.6020322 | 0.00926732 | 0.06944707 | C7_active |
| 313 | chr15 | 74843730 | 74873365 | ENSG00000140 | SCAMP2 | 10066 | 8.64666883 | -0.2312527 | 0.08408903 | -2.7500933 | 0.00595783 | 0.04956722 | C7_active |
| 314 | chr17 | 38869859 | 38921770 | ENSG00000002 | LASP1 | 3927 | 12.7707109 | -0.2309263 | 0.06832538 | -3.3798023 | 0.00072538 | 0.0088225 | C7_active |
| 315 | chr1 | 46394265 | 46413848 | ENSG00000117 | FAAH | 2166 | 6.70627404 | -0.2299513 | 0.09313115 | -2.4691125 | 0.01354486 | 0.09224839 | C7_active |
| 316 | chr17 | 17246820 | 17281293 | ENSG00000141 | COPS3 | 8533 | 7.8842076 | -0.2298901 | 0.08089631 | -2.8417874 | 0.00448614 | 0.03990559 | C7_active |
| 317 | chr11 | 57299638 | 57324952 | ENSG00000149 | TNKS1BP1 | 85456 | 6.62251347 | -0.2292323 | 0.09316197 | -2.4605779 | 0.01387135 | 0.09379248 | C7_active |
| 318 | chr19 | 12017888 | 12052967 | ENSG00000257 | CTD-2006C1.10 | NaN | 8.10141203 | -0.2292131 | 0.08130581 | -2.8191481 | 0.00481513 | 0.04210099 | C7_active |
| 319 | chr4 | 8846076 | 8871817 | ENSG00000215 | HMX1 | 3166 | 8.42660622 | -0.2291052 | 0.08216419 | -2.7883827 | 0.00529719 | 0.04508556 | C7_active |
| 320 | chr17 | 78852977 | 78925387 | ENSG00000035 | TIMP2 | 7077 | 17.3071208 | -0.2289657 | 0.06222734 | -3.6795028 | 0.00023369 | 0.00330003 | C7_active |
| 321 | chr18 | 48919853 | 48950711 | ENSG00000101 | SMAD7 | 4092 | 6.69060614 | -0.2276075 | 0.08598656 | -2.6470122 | 0.00812064 | 0.06287286 | C7_active |

| | | | | | | | | | | | | | |
|---|---|---|---|---|---|---|---|---|---|---|---|---|---|
| 322 | chr16 | 2275881 | 2340746 | ENSG00000167 | ABCA3 | 21 | 19.1526164 | -0.227474 | 0.05594562 | -4.065985 | 4.78E-05 | 0.0007688 | C7_active |
| 323 | chr7 | 1470277 | 1504367 | ENSG00000164 | INTS1 | 26173 | 10.2353479 | -0.2272172 | 0.08133892 | -2.7934617 | 0.00521472 | 0.0445942 | C7_active |
| 324 | chr9 | 136206333 | 136245841 | ENSG00000165 | QSOX2 | 169714 | 12.9747953 | -0.2272076 | 0.06608177 | -3.4382792 | 0.00058542 | 0.00739758 | C7_active |
| 325 | chr9 | 127940579 | 127980513 | ENSG00000167 | FAM102A | 399665 | 11.1709431 | -0.2259374 | 0.0702902 | -3.2143512 | 0.0013074 | 0.014565 | C7_active |
| 326 | chr16 | 70454421 | 70480274 | ENSG00000157 | FUK | 197258 | 8.37445544 | -0.2255236 | 0.07903053 | -2.8536267 | 0.00432233 | 0.03882825 | C7_active |
| 327 | chr3 | 126342635 | 126357442 | ENSG00000163 | KLF15 | 28999 | 5.8444961 | -0.2245461 | 0.09255062 | -2.4261975 | 0.01525796 | 0.09972318 | C7_active |
| 328 | chr19 | 17351448 | 17377350 | ENSG00000130 | PLVAP | 83483 | 6.86100324 | -0.2240842 | 0.08975455 | -2.4966333 | 0.01253785 | 0.08653459 | C7_active |
| 329 | chr22 | 47461299 | 47487111 | ENSG00000218 | LL22NC03-75H1 | 101927722 | 6.466656 | -0.2240559 | 0.08580983 | -2.6110746 | 0.00902582 | 0.06800077 | C7_active |
| 330 | chr12 | 109573255 | 109598117 | ENSG00000110 | MVK | 4598 | 8.77127075 | -0.2240489 | 0.07800842 | -2.872112 | 0.00407738 | 0.03706666 | C7_active |
| 331 | chr19 | 3708109 | 3750813 | ENSG00000105 | TJP3 | 27134 | 11.7633062 | -0.224017 | 0.0735634 | -3.0452243 | 0.00232507 | 0.02334843 | C7_active |
| 332 | chr11 | 118606440 | 118658038 | ENSG00000019 | PHLDB1 | 23187 | 14.4846388 | -0.2232849 | 0.06387888 | -3.4954419 | 0.00047328 | 0.00615303 | C7_active |
| 333 | chr11 | 62559603 | 62591531 | ENSG00000255 | MIR3654 | NaN | 7.55340651 | -0.2228984 | 0.08851118 | -2.5183082 | 0.01179201 | 0.08285438 | C7_active |
| 334 | chr19 | 3359563 | 3469217 | ENSG00000141 | NFIC | 4782 | 23.5850858 | -0.222794 | 0.05692408 | -3.9138807 | 9.08E-05 | 0.00138426 | C7_active |
| 335 | chrX | 41447434 | 41475710 | ENSG00000188 | NYX | 60506 | 7.31611264 | -0.2223843 | 0.08564895 | -2.5964629 | 0.00941891 | 0.07030112 | C7_active |
| 336 | chr19 | 40778242 | 40808198 | ENSG00000171 | RAB4B-EGLN2 | NaN | 8.28666624 | -0.2221158 | 0.07939631 | -2.797558 | 0.00514905 | 0.04408493 | C7_active |
| 337 | chr9 | 128702523 | 128720918 | ENSG00000160 | PKN3 | 29941 | 5.74927487 | -0.2220364 | 0.0912282 | -2.4338572 | 0.01493889 | 0.09871167 | C7_active |
| 338 | chr1 | 16124337 | 16156087 | ENSG00000142 | EPHA2 | 1969 | 7.12348239 | -0.2215117 | 0.08554427 | -2.5894391 | 0.00961324 | 0.07140841 | C7_active |
| 339 | chr17 | 48607227 | 48633572 | ENSG00000260 | HOXB7 | 3217 | 7.61303515 | -0.2197956 | 0.08159262 | -2.6938166 | 0.0070639 | 0.05667631 | C7_active |
| 340 | chr19 | 40691529 | 40718207 | ENSG00000123 | ADCK4 | 79934 | 8.36671542 | -0.2193393 | 0.07942791 | -2.7614895 | 0.00575384 | 0.04822477 | C7_active |
| 341 | chr9 | 128411751 | 128437351 | ENSG00000167 | CERCAM | 51148 | 7.63471965 | -0.2191194 | 0.08137137 | -2.6928312 | 0.00708481 | 0.05671784 | C7_active |
| 342 | chr9 | 136006537 | 136095285 | ENSG00000148 | NACC2 | 138151 | 22.8482253 | -0.2188748 | 0.0543088 | -4.0301892 | 5.57E-05 | 0.00088594 | C7_active |
| 343 | chr19 | 53124072 | 53159075 | ENSG00000197 | ZNF347 | 84671 | 8.40995853 | -0.218807 | 0.07535704 | -2.9036036 | 0.00368895 | 0.03440314 | C7_active |
| 344 | chr1 | 6521347 | 6554535 | ENSG00000162 | NOL9 | 79707 | 8.09000099 | -0.2186717 | 0.08670461 | -2.5220306 | 0.01166796 | 0.08218299 | C7_active |
| 345 | chr11 | 1389899 | 1462689 | ENSG00000174 | BRSK2 | 9024 | 22.5297557 | -0.2184601 | 0.05487642 | -3.9809462 | 6.86E-05 | 0.00106876 | C7_active |
| 346 | chr19 | 1985438 | 2034881 | ENSG00000133 | BTBD2 | 55643 | 10.7224145 | -0.2178268 | 0.07500497 | -2.9041648 | 0.00368234 | 0.03436376 | C7_active |
| 347 | chr12 | 51186936 | 51217693 | ENSG00000184 | POU6F1 | 5463 | 9.55259312 | -0.2156691 | 0.07663482 | -2.8142443 | 0.00488921 | 0.04254229 | C7_active |
| 348 | chr19 | 38647649 | 38731583 | ENSG00000130 | ACTN4 | 81 | 17.3645142 | -0.2149571 | 0.06444371 | -3.3355787 | 0.00085122 | 0.01010947 | C7_active |
| 349 | chr19 | 45039040 | 45070956 | ENSG00000104 | CLASRP | 11129 | 8.84792415 | -0.2145214 | 0.08315525 | -2.5797694 | 0.00988663 | 0.07276439 | C7_active |
| 350 | chr7 | 4905989 | 4959213 | ENSG00000136 | MMD2 | 221938 | 15.1336158 | -0.2126071 | 0.06036648 | -3.5219403 | 0.0004284 | 0.00565113 | C7_active |
| 351 | chr19 | 3630183 | 3700479 | ENSG00000186 | PIP5K1C | 23396 | 20.3303593 | -0.2123947 | 0.0565952 | -3.7528756 | 0.00017482 | 0.00253573 | C7_active |
| 352 | chr2 | 240586716 | 240617705 | ENSG00000142 | CAPN10 | 11132 | 11.0748004 | -0.2122665 | 0.06961891 | -3.0489782 | 0.00229621 | 0.02311925 | C7_active |
| 353 | chr5 | 150113837 | 150155872 | ENSG00000113 | PDGFRB | 5159 | 13.0355244 | -0.2119592 | 0.0672224 | -3.1531044 | 0.00161544 | 0.01735575 | C7_active |
| 354 | chr16 | 425619 | 523011 | ENSG00000090 | RAB11FIP3 | 9727 | 27.4571336 | -0.2115844 | 0.05586672 | -3.7873073 | 0.00015229 | 0.0022314 | C7_active |
| 355 | chr4 | 138164097 | 138242349 | ENSG00000151 | SLC7A11 | 23657 | 15.981584 | -0.2108746 | 0.05488058 | -3.8424261 | 0.00012182 | 0.00182017 | C7_active |
| 356 | chr19 | 49976467 | 50025946 | ENSG00000105 | VRK3 | 51231 | 14.7811242 | -0.2104233 | 0.06201632 | -3.3930307 | 0.00069124 | 0.00851819 | C7_active |
| 357 | chr19 | 3804024 | 3869032 | ENSG00000105 | ZFR2 | 23217 | 19.2414244 | -0.2103528 | 0.05885203 | -3.5742658 | 0.00035121 | 0.00476366 | C7_active |
| 358 | chr20 | 32358344 | 32439319 | ENSG00000171 | ASXL1 | 171023 | 16.1885421 | -0.2103333 | 0.05795656 | -3.6291548 | 0.00028435 | 0.00395349 | C7_active |
| 359 | chr11 | 62856102 | 62888875 | ENSG00000168 | SLC3A2 | 6520 | 6.92017978 | -0.2102942 | 0.08594679 | -2.4467954 | 0.01441326 | 0.09647652 | C7_active |
| 360 | chr19 | 39083913 | 39111493 | ENSG00000183 | PAPL | 390928 | 7.60831141 | -0.2100553 | 0.08458999 | -2.4832172 | 0.01302017 | 0.08926522 | C7_active |
| 361 | chr1 | 2228695 | 2310119 | ENSG00000157 | SKI | 6497 | 20.6848047 | -0.2093859 | 0.05734428 | -3.6513833 | 0.00026083 | 0.00365824 | C7_active |
| 362 | chr5 | 1456480 | 1523977 | ENSG00000153 | LPCAT1 | 79888 | 19.3295661 | -0.2091925 | 0.06091878 | -3.4339574 | 0.00059484 | 0.00749028 | C7_active |
| 363 | chr16 | 1153241 | 1221771 | ENSG00000196 | CACNA1H | 8912 | 22.0568946 | -0.2089965 | 0.05587768 | -3.7402506 | 0.00018384 | 0.00265056 | C7_active |
| 364 | chr19 | 4675224 | 4724673 | ENSG00000142 | DPP9 | 91039 | 11.4097053 | -0.2089302 | 0.07566761 | -2.7611581 | 0.00575968 | 0.04822477 | C7_active |
| 365 | chr9 | 111549722 | 111599855 | ENSG00000106 | PTGR1 | 22949 | 12.9649058 | -0.2081281 | 0.06707746 | -3.1028028 | 0.00191697 | 0.0199559 | C7_active |
| 366 | chr18 | 11981025 | 12030883 | ENSG00000141 | IMPA2 | 3613 | 14.6796739 | -0.2080463 | 0.06167079 | -3.3734981 | 0.0007422 | 0.00895479 | C7_active |
| 367 | chr22 | 30576625 | 30607083 | ENSG00000100 | PES1 | 23481 | 8.90617258 | -0.2049527 | 0.07942017 | -2.5806131 | 0.0098625 | 0.07273534 | C7_active |
| 368 | chr22 | 41301522 | 41360147 | ENSG00000100 | ZC3H7B | 23264 | 16.3279061 | -0.2047743 | 0.0577661 | -3.5448869 | 0.00039278 | 0.00524363 | C7_active |
| 369 | chr19 | 18636965 | 18671714 | ENSG00000167 | KLHL26 | 55295 | 10.5750929 | -0.2034382 | 0.08029591 | -2.5336059 | 0.01128956 | 0.08026277 | C7_active |
| 370 | chr22 | 19783224 | 19854939 | ENSG00000185 | GNB1L | 54584 | 22.8714461 | -0.2030213 | 0.04976381 | -4.079697 | 4.51E-05 | 0.00072971 | C7_active |
| 371 | chr11 | 46277661 | 46321422 | ENSG00000157 | CREB3L1 | 90993 | 10.8941568 | -0.2029257 | 0.07564904 | -2.682462 | 0.00730825 | 0.0582478 | C7_active |
| 372 | chr19 | 2100988 | 2164465 | ENSG00000065 | AP3D1 | 8943 | 16.0186425 | -0.202621 | 0.06503854 | -3.1153997 | 0.00183696 | 0.01927603 | C7_active |
| 373 | chr22 | 37367960 | 37427470 | ENSG00000166 | ELFN2 | 114794 | 20.4611848 | -0.2017098 | 0.05369621 | -3.7564998 | 0.00017231 | 0.00250435 | C7_active |
| 374 | chr16 | 75228187 | 75268053 | ENSG00000050 | BCAR1 | 9564 | 8.98480027 | -0.20169 | 0.08269636 | -2.4389229 | 0.01473111 | 0.09769694 | C7_active |
| 375 | chr7 | 99325898 | 99388164 | ENSG00000241 | ARPC1A | 10552 | 13.8309048 | -0.2004282 | 0.06102893 | -3.2841517 | 0.0010229 | 0.01179852 | C7_active |

| | | | | | | | | | | | | | |
|---|---|---|---|---|---|---|---|---|---|---|---|---|---|
| 376 | chr22 | 31618491 | 31662432 | ENSG00000241 | PISD | 23761 | 11.2404959 | -0.1998457 | 0.07463153 | -2.6777644 | 0.00741153 | 0.05890821 | C7_active |
| 377 | chr19 | 58479512 | 58512413 | ENSG00000083 | SLC27A5 | 10998 | 9.55898487 | -0.1991591 | 0.0761093 | -2.6167515 | 0.00887709 | 0.06723758 | C7_active |
| 378 | chr16 | 67562407 | 67639183 | ENSG00000102 | CTCF | 10664 | 16.9879011 | -0.1990248 | 0.0589546 | -3.375899 | 0.00073575 | 0.00889236 | C7_active |
| 379 | chr19 | 2997638 | 3047635 | ENSG00000065 | TLE2 | 7089 | 14.2392333 | -0.1986702 | 0.0603924 | -3.2896553 | 0.0010031 | 0.01164464 | C7_active |
| 380 | chr1 | 45694324 | 45750650 | ENSG00000197 | IPP | 3652 | 10.8448161 | -0.1983567 | 0.07771949 | -2.5522127 | 0.01070412 | 0.07731031 | C7_active |
| 381 | chr20 | 57495966 | 57525652 | ENSG00000124 | CTCFL | 140690 | 8.72762163 | -0.1981787 | 0.07835825 | -2.5291361 | 0.01143437 | 0.08093311 | C7_active |
| 382 | chr5 | 139526431 | 139628433 | ENSG00000131 | UBE2D2 | 7322 | 21.1609697 | -0.1976005 | 0.0499636 | -3.95489 | 7.66E-05 | 0.001182 | C7_active |
| 383 | chr1 | 11934205 | 11975538 | ENSG00000083 | PLOD1 | 5351 | 12.2430469 | -0.1966482 | 0.07002106 | -2.8084149 | 0.0049786 | 0.04311202 | C7_active |
| 384 | chr22 | 43039644 | 43089428 | ENSG00000100 | TTLL1 | 25809 | 14.7659621 | -0.1966228 | 0.05669166 | -3.4682844 | 0.00052379 | 0.00672488 | C7_active |
| 385 | chr22 | 48489460 | 48850912 | ENSG00000219 | FAM19A5 | 25817 | 112.686801 | -0.1956497 | 0.03374635 | -5.7976562 | 6.72E-09 | 1.46E-07 | C7_active |
| 386 | chr4 | 3244369 | 3271738 | ENSG00000188 | MSANTD1 | 345222 | 9.31451261 | -0.1955813 | 0.07990166 | -2.447775 | 0.01437414 | 0.09625933 | C7_active |
| 387 | chr11 | 125591712 | 125625215 | ENSG00000134 | STT3A | 3703 | 8.3106799 | -0.1953256 | 0.07899349 | -2.4726803 | 0.0134104 | 0.09159223 | C7_active |
| 388 | chr19 | 41966582 | 41997497 | ENSG00000105 | ATP1A3 | 478 | 9.9259129 | -0.1948677 | 0.08010406 | -2.4326825 | 0.01498744 | 0.09876775 | C7_active |
| 389 | chr16 | 70289663 | 70335283 | ENSG00000157 | DDX19B | 11269 | 9.87497575 | -0.1947073 | 0.07662916 | -2.5409034 | 0.01105665 | 0.07903556 | C7_active |
| 390 | chr15 | 41332694 | 41381050 | ENSG00000137 | NUSAP1 | 51203 | 10.5628375 | -0.1946968 | 0.06801678 | -2.8624811 | 0.00420338 | 0.03799647 | C7_active |
| 391 | chr3 | 15070073 | 15099163 | ENSG00000131 | ZFYVE20 | 64145 | 9.04925285 | -0.1946842 | 0.07550097 | -2.5785659 | 0.00992114 | 0.07294388 | C7_active |
| 392 | chr16 | 643262 | 768865 | ENSG00000102 | MSLN | 10232 | 33.4490688 | -0.194575 | 0.0515361 | -3.7755084 | 0.00015968 | 0.00233261 | C7_active |
| 393 | chr9 | 35490127 | 35561898 | ENSG00000198 | RUSC2 | 9853 | 15.8729004 | -0.1945164 | 0.05745167 | -3.3857407 | 0.00070986 | 0.00870308 | C7_active |
| 394 | chr15 | 85759323 | 85795030 | ENSG00000183 | KLHL25 | 64410 | 11.4807013 | -0.1940126 | 0.06957573 | -2.7885092 | 0.00529512 | 0.04508556 | C7_active |
| 395 | chr3 | 45082278 | 45146422 | ENSG00000163 | CDCP1 | 64866 | 16.4283965 | -0.1939714 | 0.06061507 | -3.2000523 | 0.00137403 | 0.015169 | C7_active |
| 396 | chr14 | 104937244 | 104978357 | ENSG00000185 | AHNAK2 | 113146 | 11.0100932 | -0.1934787 | 0.07421591 | -2.6069706 | 0.00913472 | 0.06862482 | C7_active |
| 397 | chr7 | 75842602 | 75888926 | ENSG00000005 | RHBDD2 | 57414 | 12.3596133 | -0.1928567 | 0.06673343 | -2.8899558 | 0.00385296 | 0.03561024 | C7_active |
| 398 | chr16 | 1632259 | 1686715 | ENSG00000261 | LA16c-431H6.6 | NaN | 17.1484879 | -0.1927088 | 0.06137845 | -3.1396827 | 0.00169131 | 0.01806318 | C7_active |
| 399 | chr5 | 176542511 | 176595974 | ENSG00000074 | CDHR2 | 54825 | 17.3866622 | -0.1923445 | 0.0575047 | -3.3448475 | 0.00082328 | 0.00981805 | C7_active |
| 400 | chr7 | 74289475 | 74405943 | ENSG00000106 | CLIP2 | 7461 | 29.1866567 | -0.1920808 | 0.04684481 | -4.1003654 | 4.12E-05 | 0.00067126 | C7_active |
| 401 | chr2 | 96760902 | 96811891 | ENSG00000158 | CNNM4 | 26504 | 14.8377488 | -0.1913923 | 0.05892606 | -3.2480083 | 0.00116216 | 0.01314887 | C7_active |
| 402 | chr16 | 87881546 | 87936529 | ENSG00000174 | CA5A | 763 | 17.0402868 | -0.1913209 | 0.05520586 | -3.4655906 | 0.00052907 | 0.00677452 | C7_active |
| 403 | chr9 | 98198999 | 98255721 | ENSG00000095 | TBC1D2 | 55357 | 15.4405507 | -0.1904641 | 0.06334982 | -3.0065456 | 0.00264234 | 0.0259341 | C7_active |
| 404 | chr12 | 52897187 | 52949954 | ENSG00000170 | KRT8 | 3856 | 14.1034066 | -0.1900682 | 0.06024898 | -3.154712 | 0.00160657 | 0.01728617 | C7_active |
| 405 | chr1 | 46247688 | 46278473 | ENSG00000085 | RAD54L | 8438 | 10.5921693 | -0.1889176 | 0.06954131 | -2.7166235 | 0.00659516 | 0.05384428 | C7_active |
| 406 | chr16 | 84145287 | 84178767 | ENSG00000154 | DNAAF1 | 123872 | 9.55281204 | -0.188442 | 0.07747509 | -2.4322914 | 0.01500363 | 0.09877733 | C7_active |
| 407 | chr17 | 4498838 | 4539035 | ENSG00000183 | SPNS2 | 124976 | 13.3384041 | -0.1882518 | 0.07112238 | -2.6468715 | 0.00812402 | 0.06287286 | C7_active |
| 408 | chr2 | 30231531 | 30323730 | ENSG00000213 | LBH | 81606 | 25.9420605 | -0.1880897 | 0.04439349 | -4.2368758 | 2.27E-05 | 0.00038445 | C7_active |
| 409 | chr15 | 64387748 | 64455303 | ENSG00000103 | TRIP4 | 9325 | 14.3975208 | -0.1869755 | 0.06026502 | -3.1025537 | 0.00191859 | 0.01995829 | C7_active |
| 410 | chr1 | 10032832 | 10181239 | ENSG00000130 | UBE4B | 10277 | 35.0617758 | -0.1867491 | 0.03910582 | -4.7754816 | 1.79E-06 | 3.36E-05 | C7_active |
| 411 | chr11 | 68312609 | 68449275 | ENSG00000162 | LRP5 | 4041 | 41.900698 | -0.186233 | 0.04325879 | -4.3050905 | 1.67E-05 | 0.00028856 | C7_active |
| 412 | chr19 | 41397460 | 41431345 | ENSG00000248 | BCKDHA | 593 | 10.8653875 | -0.1860439 | 0.07443592 | -2.4993831 | 0.01244097 | 0.08598943 | C7_active |
| 413 | chr7 | 2558972 | 2614734 | ENSG00000106 | IQCE | 23288 | 17.5952702 | -0.185165 | 0.05463375 | -3.3892059 | 0.00070095 | 0.00861231 | C7_active |
| 414 | chr7 | 75416787 | 75486271 | ENSG00000272 | POM121C | 100101267 | 13.1593674 | -0.1848517 | 0.06663745 | -2.7739908 | 0.00553732 | 0.04676642 | C7_active |
| 415 | chr1 | 32108038 | 32176568 | ENSG00000025 | KPNA6 | 23633 | 15.812277 | -0.1846165 | 0.05649015 | -3.2681177 | 0.00108265 | 0.01237882 | C7_active |
| 416 | chr16 | 89818179 | 89871319 | ENSG00000204 | SPIRE2 | 84501 | 14.7723708 | -0.1835665 | 0.06676102 | -2.7496058 | 0.0059667 | 0.04958379 | C7_active |
| 417 | chrX | 1268800 | 1310381 | ENSG00000198 | CSF2RA | 1438 | 11.6457546 | -0.1834163 | 0.07440193 | -2.4652083 | 0.01369336 | 0.09290865 | C7_active |
| 418 | chr16 | 70687439 | 70801161 | ENSG00000103 | VAC14 | 55697 | 36.8128805 | -0.1827563 | 0.04456731 | -4.1006799 | 4.12E-05 | 0.00067111 | C7_active |
| 419 | chr7 | 74453790 | 74602604 | ENSG00000006 | GTF2IRD1 | 9569 | 36.3877097 | -0.1824813 | 0.04362054 | -4.1833791 | 2.87E-05 | 0.00048039 | C7_active |
| 420 | chr20 | 63981135 | 64033100 | ENSG00000101 | PRPF6 | 24148 | 15.3172662 | -0.1818325 | 0.05493398 | -3.3100194 | 0.0009329 | 0.01092647 | C7_active |
| 421 | chr4 | 150078274 | 150257457 | ENSG00000170 | DCLK2 | 166614 | 41.7631043 | -0.1806903 | 0.03450867 | -5.2360836 | 1.64E-07 | 3.33E-06 | C7_active |
| 422 | chr7 | 4682309 | 4771443 | ENSG00000164 | FOXK1 | 221937 | 24.0129372 | -0.1805517 | 0.05039046 | -3.5830536 | 0.0003396 | 0.00461921 | C7_active |
| 423 | chr11 | 133908564 | 133956985 | ENSG00000080 | IGSF9B | 22997 | 17.8051994 | -0.1799288 | 0.05510964 | -3.2649246 | 0.00109493 | 0.01247963 | C7_active |
| 424 | chr12 | 132768909 | 132828858 | ENSG00000090 | GOLGA3 | 2802 | 15.017081 | -0.1798236 | 0.06395772 | -2.8116009 | 0.00492956 | 0.04281593 | C7_active |
| 425 | chr3 | 14402576 | 14489349 | ENSG00000131 | SLC6A6 | 6533 | 26.3736622 | -0.1796788 | 0.04655877 | -3.8591826 | 0.00011377 | 0.0017122 | C7_active |
| 426 | chr17 | 15229777 | 15265326 | ENSG00000109 | PMP22 | 5376 | 9.81828239 | -0.1794252 | 0.07312015 | -2.4538402 | 0.01413398 | 0.09495978 | C7_active |
| 427 | chr20 | 62307955 | 62367312 | ENSG00000130 | LAMA5 | 3911 | 17.564126 | -0.1788797 | 0.06307245 | -2.8360995 | 0.00456682 | 0.0404865 | C7_active |
| 428 | chr12 | 54016931 | 54056030 | ENSG00000198 | HOXC4 | 3221 | 12.1684569 | -0.1784794 | 0.06881699 | -2.5935365 | 0.00949945 | 0.07070883 | C7_active |
| 429 | chr1 | 54849633 | 54887218 | ENSG00000116 | DHCR24 | 1718 | 10.1132566 | -0.1781135 | 0.06826016 | -2.6093325 | 0.00907191 | 0.06826656 | C7_active |

| | | | | | | | | | | | | | | |
|---|---|---|---|---|---|---|---|---|---|---|---|---|---|---|
| 430 | chr16 | 67438014 | 67481237 | ENSG00000159 | ATP6V0D1 | 9114 | 15.2599835 | | -0.1780257 | 0.06349568 | -2.8037451 | 0.00505128 | 0.04350621 | C7_active |
| 431 | chr2 | 160272151 | 160493794 | ENSG00000153 | RBMS1 | 5937 | 51.9131253 | | -0.1776976 | 0.03345658 | -5.3112909 | 1.09E-07 | 2.23E-06 | C7_active |
| 432 | chr17 | 75441159 | 75500090 | ENSG00000177 | KIAA0195 | 9772 | 18.807746 | | -0.1770663 | 0.0571157 | -3.1001331 | 0.00193434 | 0.0200498 | C7_active |
| 433 | chr4 | 6575175 | 6623362 | ENSG00000013 | MAN2B2 | 23324 | 17.6188489 | | -0.1768963 | 0.05817737 | -3.0406375 | 0.00236078 | 0.02365372 | C7_active |
| 434 | chr12 | 122774327 | 122827528 | ENSG00000130 | CCDC62 | 84660 | 11.8173193 | | -0.1762807 | 0.06656564 | -2.6482236 | 0.0080916 | 0.06275669 | C7_active |
| 435 | chr22 | 37686414 | 37726503 | ENSG00000100 | RP1-37E16.12 | NaN | 11.4164096 | | -0.176188 | 0.07033212 | -2.5050854 | 0.01224218 | 0.0849821 | C7_active |
| 436 | chr19 | 50516892 | 50568045 | ENSG00000131 | LRRC4B | 94030 | 14.819579 | | -0.1758486 | 0.06639474 | -2.6485314 | 0.00808423 | 0.06275669 | C7_active |
| 437 | chr3 | 13816258 | 13880121 | ENSG00000154 | WNT7A | 7476 | 23.7068438 | | -0.1752996 | 0.04769133 | -3.6757113 | 0.00023719 | 0.00334288 | C7_active |
| 438 | chr11 | 27654893 | 27722058 | ENSG00000176 | BDNF | 627 | 15.2669069 | | -0.1747238 | 0.05759864 | -3.0334709 | 0.00241758 | 0.02408892 | C7_active |
| 439 | chr16 | 88570381 | 88631966 | ENSG00000158 | ZC3H18 | 124245 | 19.1416095 | | -0.1743359 | 0.05898769 | -2.9554625 | 0.00312201 | 0.02986934 | C7_active |
| 440 | chr16 | 3443611 | 3486963 | ENSG00000122 | NAA60 | 79903 | 11.5719429 | | -0.1742299 | 0.07130519 | -2.4434398 | 0.014548 | 0.09716376 | C7_active |
| 441 | chr19 | 8444767 | 8489114 | ENSG00000099 | HNRNPM | 4670 | 10.8760658 | | -0.1730331 | 0.06788526 | -2.548905 | 0.01080617 | 0.07763 | C7_active |
| 442 | chr14 | 74303069 | 74360008 | ENSG00000133 | VRTN | 55237 | 15.1685415 | | -0.1727811 | 0.06030977 | -2.8648947 | 0.00417148 | 0.03777914 | C7_active |
| 443 | chr17 | 81117295 | 81166077 | ENSG00000181 | AATK | 9625 | 15.7275279 | | -0.1717341 | 0.06538752 | -2.6264054 | 0.0086292 | 0.06572413 | C7_active |
| 444 | chr16 | 4556490 | 4600714 | ENSG00000205 | C16orf96 | 342346 | 13.355349 | | -0.1693726 | 0.06387519 | -2.651618 | 0.00801071 | 0.06236148 | C7_active |
| 445 | chr12 | 56521929 | 56596196 | ENSG00000076 | RBMS2 | 5939 | 16.8931697 | | -0.1693618 | 0.05535828 | -3.0593768 | 0.00221798 | 0.02254031 | C7_active |
| 446 | chr11 | 61752642 | 61788518 | ENSG00000124 | MYRF | 745 | 12.9209101 | | -0.1682056 | 0.06821049 | -2.4659791 | 0.01366393 | 0.09283286 | C7_active |
| 447 | chr21 | 42974510 | 43033931 | ENSG00000160 | PKNOX1 | 5316 | 15.8288367 | | -0.1681762 | 0.05604145 | -3.0009257 | 0.0026916 | 0.0263638 | C7_active |
| 448 | chr7 | 151085831 | 151144436 | ENSG00000133 | AGAP3 | 116988 | 16.6026102 | | -0.1676046 | 0.06493277 | -2.5812024 | 0.00984569 | 0.07272289 | C7_active |
| 449 | chr20 | 63956702 | 64048758 | ENSG00000196 | ZNF512B | 57473 | 26.676008 | | -0.1675959 | 0.04964981 | -3.3755607 | 0.00073665 | 0.00889538 | C7_active |
| 450 | chr5 | 154818491 | 154859252 | ENSG00000170 | FAXDC2 | 10826 | 10.5935312 | | -0.1674951 | 0.06887739 | -2.4317868 | 0.01502455 | 0.0988699 | C7_active |
| 451 | chr18 | 2847030 | 2915993 | ENSG00000132 | EMILIN2 | 84034 | 21.9910668 | | -0.1672218 | 0.05125275 | -3.2626884 | 0.00110361 | 0.0125498 | C7_active |
| 452 | chr22 | 19479459 | 19520612 | ENSG00000093 | CDC45 | 8318 | 12.421074 | | -0.1664927 | 0.06416364 | -2.5948131 | 0.00946424 | 0.07048314 | C7_active |
| 453 | chr16 | 1706183 | 1770317 | ENSG00000138 | MAPK8IP3 | 23162 | 17.3683848 | | -0.1664797 | 0.06582227 | -2.5292308 | 0.01143128 | 0.08093311 | C7_active |
| 454 | chr19 | 12763003 | 12872740 | ENSG00000095 | HOOK2 | 29911 | 26.8873511 | | -0.166354 | 0.04704637 | -3.5359582 | 0.0004063 | 0.00539412 | C7_active |
| 455 | chr7 | 816615 | 896435 | ENSG00000164 | SUN1 | 23353 | 20.9486744 | | -0.1662726 | 0.05170502 | -3.2157911 | 0.00130085 | 0.01451681 | C7_active |
| 456 | chr2 | 42442017 | 42494097 | ENSG00000171 | KCNG3 | 170850 | 12.7612453 | | -0.1656728 | 0.06594717 | -2.5122056 | 0.01199792 | 0.08393302 | C7_active |
| 457 | chr1 | 156525405 | 156572604 | ENSG00000183 | IQGAP3 | 128239 | 15.6010435 | | -0.1654863 | 0.05859379 | -2.824297 | 0.00473845 | 0.04160994 | C7_active |
| 458 | chr12 | 111452214 | 111599676 | ENSG00000204 | ATXN2 | 6311 | 28.5114701 | | -0.1654498 | 0.04430643 | -3.7342173 | 0.0001883 | 0.00270949 | C7_active |
| 459 | chr5 | 157085832 | 157142869 | ENSG00000135 | HAVCR2 | 84868 | 15.0599291 | | -0.165405 | 0.06210736 | -2.6632113 | 0.00773988 | 0.06074774 | C7_active |
| 460 | chr12 | 50392383 | 50480004 | ENSG00000161 | LARP4 | 113251 | 17.1636933 | | -0.1652094 | 0.05896392 | -2.801873 | 0.00508069 | 0.04369225 | C7_active |
| 461 | chr19 | 36054881 | 36105106 | ENSG00000075 | WDR62 | 284403 | 14.6837651 | | -0.164339 | 0.06173538 | -2.6619901 | 0.00776802 | 0.06086798 | C7_active |
| 462 | chr8 | 139730343 | 140458579 | ENSG00000167 | TRAPPC9 | 83696 | 256.620676 | | -0.1641391 | 0.01921438 | -8.5425137 | 1.31E-17 | 3.70E-16 | C7_active |
| 463 | chr12 | 79574979 | 79691097 | ENSG00000177 | PAWR | 5074 | 19.4912218 | | -0.1636178 | 0.04945001 | -3.3087518 | 0.00093713 | 0.01096715 | C7_active |
| 464 | chr19 | 52764195 | 52818004 | ENSG00000189 | ZNF600 | 162966 | 14.1006138 | | -0.1634777 | 0.06343808 | -2.5769643 | 0.00996723 | 0.0731708 | C7_active |
| 465 | chr19 | 23304991 | 23395560 | ENSG00000167 | ZNF91 | 7644 | 21.7921776 | | -0.1633073 | 0.04975741 | -3.2820708 | 0.00103048 | 0.01186695 | C7_active |
| 466 | chr22 | 20017014 | 20065926 | ENSG00000183 | TANGO2 | 128989 | 16.230904 | | -0.1625799 | 0.06261713 | -2.5964124 | 0.00942029 | 0.07030112 | C7_active |
| 467 | chr16 | 46884378 | 46931297 | ENSG00000166 | GPT2 | 84706 | 14.5481345 | | -0.1618069 | 0.06429544 | -2.5166154 | 0.01184881 | 0.08313193 | C7_active |
| 468 | chr9 | 33817567 | 33920404 | ENSG00000107 | UBE2R2 | 54926 | 19.2890571 | | -0.1612863 | 0.05110111 | -3.1562201 | 0.00159828 | 0.017228 | C7_active |
| 469 | chr11 | 72754729 | 72794168 | ENSG00000214 | STARD10 | 10809 | 12.6555808 | | -0.1612369 | 0.06372722 | -2.530111 | 0.01140265 | 0.08086736 | C7_active |
| 470 | chr12 | 108645109 | 108731596 | ENSG00000110 | CORO1C | 23603 | 17.0764929 | | -0.1609261 | 0.05644955 | -2.8507949 | 0.00436101 | 0.03907832 | C7_active |
| 471 | chr7 | 996986 | 1138260 | ENSG00000146 | C7orf50 | 84310 | 40.9854343 | | -0.1607084 | 0.04390551 | -3.6603251 | 0.0002519 | 0.00353979 | C7_active |
| 472 | chr1 | 2050470 | 2185395 | ENSG00000067 | PRKCZ | 5590 | 44.515232 | | -0.1602702 | 0.03947109 | -4.0604456 | 4.90E-05 | 0.00078639 | C7_active |
| 473 | chr22 | 17178790 | 17221989 | ENSG00000093 | CECR1 | 51816 | 12.9303596 | | -0.1600758 | 0.06204742 | -2.579895 | 0.00988304 | 0.07276439 | C7_active |
| 474 | chr7 | 5306790 | 5425414 | ENSG00000182 | TNRC18 | 84629 | 28.7709665 | | -0.1597407 | 0.05020669 | -3.1816609 | 0.00146433 | 0.01600381 | C7_active |
| 475 | chr1 | 154582062 | 154627999 | ENSG00000160 | ADAR | 103 | 13.7173007 | | -0.159677 | 0.06161942 | -2.5913422 | 0.00956024 | 0.07108319 | C7_active |
| 476 | chr6 | 41546426 | 41602384 | ENSG00000137 | FOXP4 | 116113 | 18.0801414 | | -0.1589479 | 0.05856337 | -2.7141179 | 0.00664525 | 0.05416123 | C7_active |
| 477 | chr7 | 73440398 | 73522278 | ENSG00000009 | BAZ1B | 9031 | 18.5976635 | | -0.1570088 | 0.05132143 | -3.0593218 | 0.00221839 | 0.02254031 | C7_active |
| 478 | chr19 | 47113274 | 47210636 | ENSG00000142 | SAE1 | 10055 | 20.7026344 | | -0.1561437 | 0.05411697 | -2.8853003 | 0.0039104 | 0.03600269 | C7_active |
| 479 | chr9 | 115019578 | 115118257 | ENSG00000041 | TNC | 3371 | 28.0337706 | | -0.1561102 | 0.0455045 | -3.4306546 | 0.00060213 | 0.00756225 | C7_active |
| 480 | chr4 | 83535914 | 83605875 | ENSG00000138 | AGPAT9 | 84803 | 15.9788838 | | -0.1542339 | 0.0612933 | -2.5163249 | 0.01185858 | 0.08316002 | C7_active |
| 481 | chr7 | 74657667 | 74760692 | ENSG00000263 | GTF2I | 2969 | 14.2534019 | | -0.1541632 | 0.06094512 | -2.5295406 | 0.0114212 | 0.08091932 | C7_active |
| 482 | chr19 | 12995608 | 13098796 | ENSG00000008 | NFIX | 4784 | 29.1228183 | | -0.15318 | 0.04661604 | -3.2859948 | 0.00101623 | 0.01174037 | C7_active |
| 483 | chr19 | 8520790 | 8577577 | ENSG00000142 | MYO1F | 4542 | 15.1212402 | | -0.1525782 | 0.06277029 | -2.4307383 | 0.01506809 | 0.09906601 | C7_active |

| | | | | | | | | | | | | | | |
|---|---|---|---|---|---|---|---|---|---|---|---|---|---|---|
| 484 | chr19 | 18097793 | 18151692 | ENSG00000099 | MAST3 | 23031 | 15.316719 | | -0.1525266 | 0.06093602 | -2.5030615 | 0.01231242 | 0.0853874 | C7_active |
| 485 | chr16 | 68310974 | 68358563 | ENSG00000132 | PRMT7 | 54496 | 14.3389288 | | -0.150546 | 0.0604546 | -2.4902322 | 0.01276596 | 0.08777286 | C7_active |
| 486 | chr17 | 81556887 | 81648465 | ENSG00000182 | NPLOC4 | 55666 | 20.6466867 | | -0.1497178 | 0.05622443 | -2.6628604 | 0.00774796 | 0.06077804 | C7_active |
| 487 | chr3 | 125446644 | 125520197 | ENSG00000114 | SNX4 | 8723 | 14.8532003 | | -0.1490235 | 0.05716663 | -2.606827 | 0.00913855 | 0.06862482 | C7_active |
| 488 | chr22 | 44668547 | 44737681 | ENSG00000186 | PRR5 | 55615 | 22.611051 | | -0.1484551 | 0.04994025 | -2.972654 | 0.00295237 | 0.02853035 | C7_active |
| 489 | chr7 | 898778 | 975599 | ENSG00000240 | COX19 | 90639 | 22.6953172 | | -0.1469286 | 0.05132822 | -2.8625303 | 0.00420273 | 0.03799647 | C7_active |
| 490 | chr12 | 64610513 | 64697567 | ENSG00000153 | RASSF3 | 283349 | 20.2311762 | | -0.1457069 | 0.05332634 | -2.7323628 | 0.00628819 | 0.05177788 | C7_active |
| 491 | chr2 | 201116104 | 201176687 | ENSG00000003 | CFLAR | 8837 | 15.0520801 | | -0.1454998 | 0.05942549 | -2.4484411 | 0.01434759 | 0.09612618 | C7_active |
| 492 | chrX | 9725346 | 9786297 | ENSG00000101 | GPR143 | 4935 | 18.1030121 | | -0.1454012 | 0.05492575 | -2.6472323 | 0.00811536 | 0.06287286 | C7_active |
| 493 | chr8 | 144148016 | 144261940 | ENSG00000179 | MROH1 | 727957 | 33.8563105 | | -0.1452165 | 0.04446383 | -3.2659464 | 0.00109099 | 0.01245438 | C7_active |
| 494 | chr1 | 21822245 | 21937297 | ENSG00000142 | HSPG2 | 3339 | 33.7525687 | | -0.1451684 | 0.04701708 | -3.0875662 | 0.00201803 | 0.02076797 | C7_active |
| 495 | chr20 | 9068763 | 9481242 | ENSG00000101 | PLCB4 | 5332 | 92.6727567 | | -0.1449732 | 0.03220125 | -4.5020977 | 6.73E-06 | 0.00012096 | C7_active |
| 496 | chr7 | 102363872 | 102426676 | ENSG00000128 | PRKRIP1 | 79706 | 14.7867935 | | -0.1448697 | 0.05853842 | -2.4747802 | 0.01333182 | 0.09114186 | C7_active |
| 497 | chr21 | 46324122 | 46445769 | ENSG00000160 | PCNT | 5116 | 33.654327 | | -0.1444543 | 0.04504351 | -3.2069953 | 0.00134129 | 0.01486453 | C7_active |
| 498 | chr12 | 118063593 | 118136026 | ENSG00000176 | VSIG10 | 54621 | 18.9482685 | | -0.1444284 | 0.05581471 | -2.5876411 | 0.00966356 | 0.07174523 | C7_active |
| 499 | chr3 | 123490820 | 123585185 | ENSG00000206 | PTPLB | 201562 | 23.1258729 | | -0.1441989 | 0.04888162 | -2.9499619 | 0.00317813 | 0.03028572 | C7_active |
| 500 | chr17 | 3565444 | 3636249 | ENSG00000262 | TRPV1 | NaN | 20.6325156 | | -0.1439859 | 0.05733696 | -2.511224 | 0.01203133 | 0.08399614 | C7_active |
| 501 | chr16 | 87696485 | 87765992 | ENSG00000104 | KLHDC4 | 54758 | 20.8572155 | | -0.1436986 | 0.0543046 | -2.6461581 | 0.00814117 | 0.06288789 | C7_active |
| 502 | chr20 | 63743666 | 63831244 | ENSG00000130 | ZBTB46 | 140685 | 23.4792176 | | -0.1436979 | 0.05188903 | -2.7693308 | 0.00561716 | 0.04730617 | C7_active |
| 503 | chrX | 54445454 | 54496166 | ENSG00000102 | FGD1 | 2245 | 16.5037182 | | -0.1430999 | 0.05655159 | -2.5304312 | 0.01139224 | 0.08083335 | C7_active |
| 504 | chr5 | 710360 | 766952 | ENSG00000206 | ZDHHC11B | 653082 | 15.4110787 | | -0.1429842 | 0.05860686 | -2.4397173 | 0.01469876 | 0.09757217 | C7_active |
| 505 | chr19 | 54724459 | 54767884 | ENSG00000243 | KIR2DL3 | 3804 | 15.8374803 | | -0.1426865 | 0.05865427 | -2.4326709 | 0.01498792 | 0.09876775 | C7_active |
| 506 | chr8 | 143437655 | 143541453 | ENSG00000014 | ZC3H3 | 23144 | 33.3832392 | | -0.1422977 | 0.04525557 | -3.144314 | 0.00166477 | 0.01785186 | C7_active |
| 507 | chr1 | 2321253 | 2391707 | ENSG00000116 | MORN1 | 79906 | 23.6092843 | | -0.1418372 | 0.05103275 | -2.7793361 | 0.00544701 | 0.04606161 | C7_active |
| 508 | chr19 | 7395113 | 7472477 | ENSG00000104 | ARHGEF18 | 23370 | 24.9671474 | | -0.1410069 | 0.05525542 | -2.5519101 | 0.01071341 | 0.07731031 | C7_active |
| 509 | chr19 | 41350853 | 41425004 | ENSG00000255 | CTC-435M10.3 | NaN | 19.8812571 | | -0.140906 | 0.05301331 | -2.657936 | 0.00786208 | 0.06146786 | C7_active |
| 510 | chr2 | 98346155 | 98398601 | ENSG00000144 | CNGA3 | 1261 | 18.3182641 | | -0.1408665 | 0.05485689 | -2.5678903 | 0.01023195 | 0.07469585 | C7_active |
| 511 | chr12 | 122271432 | 122422632 | ENSG00000130 | CLIP1 | 6249 | 32.1459897 | | -0.1406892 | 0.04290643 | -3.2789772 | 0.00104184 | 0.01196913 | C7_active |
| 512 | chr9 | 136807922 | 136957733 | ENSG00000196 | RABL6 | 55684 | 40.4807387 | | -0.1406736 | 0.04375593 | -3.2149615 | 0.00130462 | 0.01454758 | C7_active |
| 513 | chr14 | 96204844 | 96263929 | ENSG00000258 | RP11-404P21.8 | NaN | 21.5294159 | | -0.1404347 | 0.05094179 | -2.7567681 | 0.00583757 | 0.04882027 | C7_active |
| 514 | chr12 | 112418351 | 112509913 | ENSG00000179 | PTPN11 | 5781 | 22.0743987 | | -0.1399719 | 0.05106701 | -2.7409452 | 0.00612627 | 0.0506181 | C7_active |
| 515 | chr19 | 7380964 | 7472477 | ENSG00000268 | CTD-2207O23.3 | NaN | 28.9422018 | | -0.1388731 | 0.05237129 | -2.6517033 | 0.00800869 | 0.06236148 | C7_active |
| 516 | chr22 | 20495913 | 20587632 | ENSG00000099 | MED15 | 51586 | 21.614331 | | -0.1382069 | 0.0515616 | -2.6804233 | 0.00735291 | 0.05853908 | C7_active |
| 517 | chr4 | 2462220 | 2625320 | ENSG00000063 | RNF4 | 6047 | 38.3024444 | | -0.1374342 | 0.038085 | -3.6086164 | 0.00030783 | 0.00423507 | C7_active |
| 518 | chr12 | 113298759 | 113359493 | ENSG00000089 | SLC8B1 | 80024 | 17.3050427 | | -0.1372322 | 0.05390304 | -2.5459095 | 0.01089934 | 0.07814359 | C7_active |
| 519 | chr1 | 4654732 | 4792534 | ENSG00000196 | AJAP1 | 55966 | 50.821379 | | -0.1372002 | 0.03676261 | -3.7320567 | 0.00018992 | 0.00273012 | C7_active |
| 520 | chr3 | 124965710 | 125055958 | ENSG00000173 | HEG1 | 57493 | 22.2279366 | | -0.1366092 | 0.04981824 | -2.7421516 | 0.00610382 | 0.05050386 | C7_active |
| 521 | chr11 | 72685069 | 72793599 | ENSG00000186 | ARAP1 | 116985 | 36.124845 | | -0.1364654 | 0.04673389 | -2.9200516 | 0.00349973 | 0.03287242 | C7_active |
| 522 | chr6 | 33620365 | 33696574 | ENSG00000096 | ITPR3 | 3710 | 21.7808707 | | -0.1364373 | 0.05411304 | -2.5213388 | 0.01169092 | 0.08230456 | C7_active |
| 523 | chr4 | 5748084 | 5893058 | ENSG00000072 | CRMP1 | 1400 | 43.1537432 | | -0.1363225 | 0.03960584 | -3.4419801 | 0.00057747 | 0.00731635 | C7_active |
| 524 | chr11 | 126355640 | 126440344 | ENSG00000110 | ST3GAL4 | 6484 | 25.6827881 | | -0.1362096 | 0.04818612 | -2.8267399 | 0.00470245 | 0.04136665 | C7_active |
| 525 | chr16 | 24845841 | 24911628 | ENSG00000158 | SLC5A11 | 115584 | 18.0816924 | | -0.1356709 | 0.05531525 | -2.452686 | 0.0141794 | 0.09518671 | C7_active |
| 526 | chr4 | 5711197 | 5814305 | ENSG00000072 | EVC | 2121 | 26.7637378 | | -0.1356122 | 0.05023525 | -2.6995426 | 0.00694349 | 0.05580333 | C7_active |
| 527 | chr1 | 1921951 | 2003837 | ENSG00000142 | KIAA1751 | 85452 | 26.7400484 | | -0.1354932 | 0.05311672 | -2.5508582 | 0.0107458 | 0.07746649 | C7_active |
| 528 | chr16 | 89267627 | 89490561 | ENSG00000167 | ANKRD11 | 29123 | 51.6733784 | | -0.1352946 | 0.03896206 | -3.4724709 | 0.00051569 | 0.00665048 | C7_active |
| 529 | chr14 | 68874143 | 68979440 | ENSG00000072 | ACTN1 | 87 | 25.7433694 | | -0.1351234 | 0.05294297 | -2.5522443 | 0.01070314 | 0.07731031 | C7_active |
| 530 | chr3 | 9649433 | 9702394 | ENSG00000163 | MTMR14 | 64419 | 19.7093132 | | -0.1351187 | 0.05251485 | -2.5729619 | 0.01008323 | 0.07387195 | C7_active |
| 531 | chr19 | 23525631 | 23687220 | ENSG00000197 | ZNF675 | 171392 | 39.5486315 | | -0.1349456 | 0.03778093 | -3.5717929 | 0.00035455 | 0.00479985 | C7_active |
| 532 | chr19 | 2511219 | 2702709 | ENSG00000176 | GNG7 | 2788 | 48.1008553 | | -0.1349132 | 0.03980642 | -3.3892324 | 0.00070089 | 0.00861231 | C7_active |
| 533 | chr15 | 75369379 | 75455842 | ENSG00000169 | SIN3A | 25942 | 20.9167596 | | -0.1339947 | 0.05370808 | -2.4948699 | 0.01260033 | 0.08692421 | C7_active |
| 534 | chr15 | 78810487 | 78898133 | ENSG00000185 | MORF4L1 | 10933 | 20.3409443 | | -0.1331552 | 0.0509216 | -2.614906 | 0.0089252 | 0.0674442 | C7_active |
| 535 | chr19 | 9853718 | 9936552 | ENSG00000105 | OLFM2 | 93145 | 23.839135 | | -0.1331265 | 0.04958413 | -2.6848606 | 0.00725601 | 0.05786345 | C7_active |
| 536 | chr11 | 3671083 | 3797792 | ENSG00000110 | NUP98 | 4928 | 27.3950633 | | -0.1329623 | 0.04730645 | -2.8106583 | 0.00494403 | 0.04289092 | C7_active |
| 537 | chr7 | 44606572 | 44709066 | ENSG00000105 | OGDH | 4967 | 26.2424482 | | -0.1325 | 0.04668074 | -2.8384292 | 0.00453362 | 0.04027461 | C7_active |

| | | | | | | | | | | | | | |
|---|---|---|---|---|---|---|---|---|---|---|---|---|---|
| 538 | chr9 | 132161676 | 132244534 | ENSG00000196 | NTNG2 | 84628 | 28.7549024 | -0.1320856 | 0.04763176 | -2.7730561 | 0.00555325 | 0.0468501 | C7_active |
| 539 | chr11 | 117314557 | 117413268 | ENSG00000110 | CEP164 | 22897 | 29.7794946 | -0.1318398 | 0.04470299 | -2.9492398 | 0.00318557 | 0.03033653 | C7_active |
| 540 | chr17 | 74776483 | 74839779 | ENSG00000109 | TMEM104 | 54868 | 24.421396 | -0.1313549 | 0.04963807 | -2.6462535 | 0.00813888 | 0.06288789 | C7_active |
| 541 | chr1 | 3652520 | 3736201 | ENSG00000078 | TP73 | 7161 | 31.3091049 | -0.1311787 | 0.04440995 | -2.9538124 | 0.00313875 | 0.0299897 | C7_active |
| 542 | chr11 | 62433542 | 62556235 | ENSG00000124 | AHNAK | 79026 | 31.6773131 | -0.1296755 | 0.04493416 | -2.8859001 | 0.00390296 | 0.03595711 | C7_active |
| 543 | chr9 | 132410043 | 132573317 | ENSG00000188 | C9orf171 | 389799 | 53.6394236 | -0.1295329 | 0.03591976 | -3.6061739 | 0.00031074 | 0.00426697 | C7_active |
| 544 | chr16 | 24729695 | 24827632 | ENSG00000090 | TNRC6A | 27327 | 23.6130546 | -0.1293542 | 0.04612879 | -2.8041965 | 0.00504422 | 0.04350621 | C7_active |
| 545 | chr1 | 53242364 | 53328070 | ENSG00000157 | LRP8 | 7804 | 31.519249 | -0.1270144 | 0.04130803 | -3.0748122 | 0.00210635 | 0.0215233 | C7_active |
| 546 | chr4 | 7192538 | 7742836 | ENSG00000184 | SORCS2 | 57537 | 204.728812 | -0.1265361 | 0.02616651 | -4.8358049 | 1.33E-06 | 2.52E-05 | C7_active |
| 547 | chr3 | 185643739 | 185825056 | ENSG00000073 | IGF2BP2 | 10644 | 44.9573864 | -0.1256558 | 0.03467975 | -3.6233198 | 0.00029085 | 0.00403213 | C7_active |
| 548 | chr22 | 43197283 | 43343388 | ENSG00000159 | SCUBE1 | 80274 | 55.2390242 | -0.1238063 | 0.03391656 | -3.6503208 | 0.00026191 | 0.00366984 | C7_active |
| 549 | chr19 | 12461559 | 12551474 | ENSG00000196 | ZNF709 | NaN | 20.8946215 | -0.1213943 | 0.04668967 | -2.6000246 | 0.00932171 | 0.06974591 | C7_active |
| 550 | chr17 | 2405562 | 2511891 | ENSG00000127 | METTL16 | 79066 | 24.5044442 | -0.1213552 | 0.04940132 | -2.4565181 | 0.01402907 | 0.09456344 | C7_active |
| 551 | chr1 | 218346235 | 218444619 | ENSG00000092 | TGFB2 | 7042 | 23.6368224 | -0.1208156 | 0.04877869 | -2.4768111 | 0.0132562 | 0.09075401 | C7_active |
| 552 | chr7 | 6161776 | 6272644 | ENSG00000008 | CYTH3 | 9265 | 27.3572521 | -0.1201273 | 0.04503856 | -2.66721 | 0.00764839 | 0.0602922 | C7_active |
| 553 | chr16 | 15949577 | 16143074 | ENSG00000103 | ABCC1 | 4363 | 49.5073069 | -0.119942 | 0.04002228 | -2.996881 | 0.00272757 | 0.02662568 | C7_active |
| 554 | chr5 | 172042073 | 172188386 | ENSG00000072 | STK10 | 6793 | 42.8724823 | -0.1198416 | 0.03840874 | -3.120166 | 0.00180749 | 0.01903609 | C7_active |
| 555 | chr16 | 50548330 | 50640739 | ENSG00000140 | NKD1 | 85407 | 34.8795348 | -0.1198409 | 0.04486733 | -2.6710063 | 0.00756242 | 0.05982986 | C7_active |
| 556 | chr9 | 137306896 | 137423262 | ENSG00000187 | EXD3 | 54932 | 33.959854 | -0.1197112 | 0.04611467 | -2.5959471 | 0.00943306 | 0.07032359 | C7_active |
| 557 | chr7 | 2906141 | 3043945 | ENSG00000198 | CARD11 | 84433 | 42.6335555 | -0.1191757 | 0.03905735 | -3.0513004 | 0.00227852 | 0.02300544 | C7_active |
| 558 | chr17 | 59707192 | 59842255 | ENSG00000062 | VMP1 | 81671 | 23.0172497 | -0.1188466 | 0.04705695 | -2.52559 | 0.01155042 | 0.08163429 | C7_active |
| 559 | chr4 | 40423267 | 40630875 | ENSG00000163 | RBM47 | 54502 | 50.351953 | -0.1187894 | 0.0331234 | -3.5862682 | 0.00033544 | 0.00456698 | C7_active |
| 560 | chr4 | 39822863 | 39977956 | ENSG00000121 | PDS5A | 23244 | 29.9594977 | -0.1185393 | 0.04530888 | -2.6162495 | 0.00889016 | 0.06728519 | C7_active |
| 561 | chr4 | 7965310 | 8158832 | ENSG00000163 | ABLIM2 | 84448 | 67.7129913 | -0.1176423 | 0.03258072 | -3.610796 | 0.00030526 | 0.00420767 | C7_active |
| 562 | chr14 | 103562962 | 103685924 | ENSG00000256 | KLC1 | NaN | 29.8559373 | -0.1176231 | 0.04411001 | -2.666584 | 0.00766265 | 0.06037159 | C7_active |
| 563 | chr5 | 14143702 | 14532128 | ENSG00000038 | TRIO | 7204 | 88.5554776 | -0.117224 | 0.02961778 | -3.9578939 | 7.56E-05 | 0.00116848 | C7_active |
| 564 | chr7 | 4797055 | 4883719 | ENSG00000157 | RADIL | 55698 | 28.8340011 | -0.1167919 | 0.04756371 | -2.455484 | 0.0140695 | 0.09471434 | C7_active |
| 565 | chr17 | 65137431 | 65227703 | ENSG00000108 | RGS9 | 8787 | 29.4874732 | -0.1162215 | 0.04523791 | -2.5691178 | 0.01019578 | 0.07450723 | C7_active |
| 566 | chr18 | 57646426 | 57803101 | ENSG00000081 | ATP8B1 | 5205 | 30.8117568 | -0.1161655 | 0.04560413 | -2.5472573 | 0.01085734 | 0.07791989 | C7_active |
| 567 | chr14 | 103733752 | 103847590 | ENSG00000088 | PPP1R13B | 23368 | 30.2615876 | -0.1161595 | 0.04401545 | -2.6390616 | 0.00831359 | 0.06403061 | C7_active |
| 568 | chr2 | 46916157 | 47076137 | ENSG00000068 | TTC7A | 57217 | 45.2959596 | -0.1158085 | 0.03737961 | -3.0981721 | 0.00194718 | 0.02016845 | C7_active |
| 569 | chr15 | 22867052 | 22981063 | ENSG00000273 | CYFIP1 | 23191 | 29.79806 | -0.1146564 | 0.04416378 | -2.5961644 | 0.00942709 | 0.07031549 | C7_active |
| 570 | chr19 | 17075781 | 17214537 | ENSG00000099 | MYO9B | 4650 | 37.1006931 | -0.1137722 | 0.04174696 | -2.725282 | 0.00642466 | 0.05269098 | C7_active |
| 571 | chr17 | 77086716 | 77217101 | ENSG00000129 | SEC14L1 | 6397 | 32.0144019 | -0.1109253 | 0.04484895 | -2.473309 | 0.01338683 | 0.09147459 | C7_active |
| 572 | chr3 | 58008400 | 58172251 | ENSG00000136 | FLNB | 2317 | 40.8366648 | -0.1108883 | 0.04041725 | -2.7435875 | 0.00607719 | 0.05032789 | C7_active |
| 573 | chr19 | 16355239 | 16472085 | ENSG00000127 | EPS15L1 | 58513 | 34.5106645 | -0.1104121 | 0.04529481 | -2.4376328 | 0.01478378 | 0.09795616 | C7_active |
| 574 | chr6 | 150143076 | 150250357 | ENSG00000198 | PPP1R14C | 81706 | 33.3818407 | -0.1103308 | 0.03879562 | -2.8438978 | 0.00445653 | 0.03971218 | C7_active |
| 575 | chr7 | 47275154 | 47582558 | ENSG00000136 | TNS3 | 64759 | 77.5773314 | -0.1098875 | 0.03246672 | -3.3846206 | 0.00071277 | 0.00872383 | C7_active |
| 576 | chr7 | 157539056 | 158587788 | ENSG00000155 | PTPRN2 | 5799 | 345.603365 | -0.1084383 | 0.02404259 | -4.5102583 | 6.47E-06 | 0.00011684 | C7_active |
| 577 | chr9 | 6532464 | 6645650 | ENSG00000178 | GLDC | 2731 | 32.6294776 | -0.106874 | 0.0374441 | -2.8542272 | 0.00431417 | 0.03880327 | C7_active |
| 578 | chr7 | 5620047 | 5781739 | ENSG00000011 | RNF216 | 54476 | 40.0085331 | -0.1063071 | 0.03934002 | -2.7022642 | 0.0068869 | 0.05544129 | C7_active |
| 579 | chr5 | 172834275 | 172952685 | ENSG00000113 | ERGIC1 | 57222 | 34.4569849 | -0.1052727 | 0.04080124 | -2.5801347 | 0.00987618 | 0.07276174 | C7_active |
| 580 | chr18 | 11688956 | 11885685 | ENSG00000141 | GNAL | 2774 | 54.4975739 | -0.1040925 | 0.03092125 | -3.3663751 | 0.00076163 | 0.00914643 | C7_active |
| 581 | chr4 | 163524298 | 164384050 | ENSG00000145 | 36951 | 55016 | 204.903816 | -0.103533 | 0.02679003 | -3.8646111 | 0.00011127 | 0.00167632 | C7_active |
| 582 | chr20 | 51596514 | 51768634 | ENSG00000054 | ATP9A | 10079 | 46.5485081 | -0.102197 | 0.03525357 | -2.8989111 | 0.00374461 | 0.03483213 | C7_active |
| 583 | chr19 | 7958579 | 8445041 | ENSG00000066 | ELAVL1 | 1994 | 125.215977 | -0.1019598 | 0.02663631 | -3.8278493 | 0.00012927 | 0.0019254 | C7_active |
| 584 | chr22 | 40410281 | 40636702 | ENSG00000196 | MKL1 | 57591 | 49.3680191 | -0.0987995 | 0.03391726 | -2.9129554 | 0.00358026 | 0.03356317 | C7_active |
| 585 | chr12 | 64264762 | 64391192 | ENSG00000185 | C12orf56 | 115749 | 33.161818 | -0.098136 | 0.03834609 | -2.5592177 | 0.0104908 | 0.0761997 | C7_active |
| 586 | chr22 | 20707691 | 20859417 | ENSG00000241 | PI4KA | 5297 | 44.9580744 | -0.0980957 | 0.03760474 | -2.6085994 | 0.00909136 | 0.06837728 | C7_active |
| 587 | chr1 | 10636604 | 10796650 | ENSG00000130 | CASZ1 | 54897 | 55.8164571 | -0.0979368 | 0.03463262 | -2.8278782 | 0.00468576 | 0.0413207 | C7_active |
| 588 | chr3 | 123610049 | 123884331 | ENSG00000065 | MYLK | 4638 | 71.3041377 | -0.0974266 | 0.03357023 | -2.9021723 | 0.00370585 | 0.03453839 | C7_active |
| 589 | chr4 | 6320578 | 6563600 | ENSG00000074 | PPP2R2C | 5522 | 86.5258448 | -0.0974228 | 0.03138917 | -3.1037069 | 0.00191113 | 0.0199094 | C7_active |
| 590 | chr7 | 1815793 | 2233243 | ENSG00000002 | MAD1L1 | 8379 | 142.14741 | -0.097168 | 0.03180436 | -3.0551771 | 0.00224928 | 0.02282201 | C7_active |
| 591 | chr17 | 6876635 | 7012349 | ENSG00000215 | AC027763.2 | 100506713 | 37.6564249 | -0.0948426 | 0.03558952 | -2.6649013 | 0.00770109 | 0.06054217 | C7_active |

| | | | | | | | | | | | | |
|---|---|---|---|---|---|---|---|---|---|---|---|---|
| 592 | chr22 | 44702233 | 44862784 | ENSG00000248 | PRR5-ARHGAP8 | 553158 | 50.7724012 | -0.0945697 | 0.03488907 | -2.7105827 | 0.00671651 | 0.05457223 C7_active |
| 593 | chr18 | 12446512 | 12658134 | ENSG00000134 | SPIRE1 | 56907 | 45.7188349 | -0.0939606 | 0.03746876 | -2.5077055 | 0.01215179 | 0.08447661 C7_active |
| 594 | chr17 | 799313 | 979770 | ENSG00000167 | NXN | 64359 | 55.3780402 | -0.0935616 | 0.03840363 | -2.4362692 | 0.01483963 | 0.09828105 C7_active |
| 595 | chr19 | 45079288 | 45305283 | ENSG00000007 | MARK4 | 57787 | 64.8439563 | -0.092676 | 0.03587519 | -2.5832899 | 0.00978631 | 0.07239557 C7_active |
| 596 | chr6 | 3268962 | 3457022 | ENSG00000137 | SLC22A23 | 63027 | 59.2489692 | -0.089641 | 0.0322141 | -2.7826634 | 0.00539147 | 0.04567229 C7_active |
| 597 | chr18 | 59430940 | 59697380 | ENSG00000183 | CCBE1 | 147372 | 72.1938373 | -0.0875775 | 0.031041 | -2.8213491 | 0.00478221 | 0.04186396 C7_active |
| 598 | chr11 | 8693351 | 8910951 | ENSG00000166 | ST5 | 6764 | 60.0352003 | -0.0860962 | 0.03049546 | -2.8232465 | 0.004754 | 0.04171266 C7_active |
| 599 | chr21 | 45643694 | 45942454 | ENSG00000183 | PCBP3 | 54039 | 93.8414339 | -0.0852605 | 0.02719865 | -3.1347336 | 0.0017201 | 0.01832996 C7_active |
| 600 | chr4 | 86935002 | 87141054 | ENSG00000172 | AFF1 | 4299 | 48.0130416 | -0.0839307 | 0.03333215 | -2.5180103 | 0.01180199 | 0.08288409 C7_active |
| 601 | chr3 | 11556070 | 11771350 | ENSG00000144 | VGLL4 | 9686 | 56.6429967 | -0.083604 | 0.03262563 | -2.5625259 | 0.01039138 | 0.07574467 C7_active |
| 602 | chr1 | 39081316 | 39487177 | ENSG00000127 | MACF1 | 23499 | 96.6054667 | -0.081212 | 0.02584252 | -3.1425731 | 0.0016747 | 0.01791232 C7_active |
| 603 | chr7 | 75533300 | 75738962 | ENSG00000127 | HIP1 | 3092 | 56.868556 | -0.0805318 | 0.03280401 | -2.4549383 | 0.01409088 | 0.09471434 C7_active |
| 604 | chr17 | 66302636 | 66810743 | ENSG00000154 | PRKCA | 5578 | 134.321311 | -0.079148 | 0.02527294 | -3.1317284 | 0.00173781 | 0.01846402 C7_active |
| 605 | chr1 | 22710839 | 22921500 | ENSG00000133 | EPHB2 | 2048 | 72.4272208 | -0.0779104 | 0.02958024 | -2.6338659 | 0.00844188 | 0.06464843 C7_active |
| 606 | chr16 | 81445170 | 81711762 | ENSG00000153 | CMIP | 80790 | 83.9083976 | -0.0769303 | 0.02992872 | -2.5704509 | 0.01015662 | 0.07429638 C7_active |
| 607 | chr18 | 37243047 | 37566037 | ENSG00000101 | CELF4 | 56853 | 123.038193 | -0.0767189 | 0.02717553 | -2.8230859 | 0.00475638 | 0.04171266 C7_active |
| 608 | chr7 | 101815904 | 102283957 | ENSG00000257 | CUX1 | 1523 | 131.369109 | -0.0757973 | 0.02382918 | -3.1808607 | 0.00146838 | 0.0160266 C7_active |
| 609 | chr5 | 78485215 | 78770021 | ENSG00000145 | LHFPL2 | 10184 | 77.1233911 | -0.0744535 | 0.02892674 | -2.5738624 | 0.01005703 | 0.07375494 C7_active |
| 610 | chr2 | 109129348 | 109504632 | ENSG00000172 | SH3RF3 | 344558 | 122.709311 | -0.0729136 | 0.02695442 | -2.7050695 | 0.00682901 | 0.05516004 C7_active |
| 611 | chr9 | 94726701 | 95087218 | ENSG00000148 | C9orf3 | 84909 | 93.3402997 | -0.0722775 | 0.02505823 | -2.8843822 | 0.00392182 | 0.03602202 C7_active |
| 612 | chr20 | 61252426 | 61940617 | ENSG00000179 | CDH4 | 1002 | 238.854469 | -0.0714678 | 0.02548884 | -2.803884 | 0.00504911 | 0.04350621 C7_active |
| 613 | chr1 | 36795527 | 37034129 | ENSG00000163 | GRIK3 | 2899 | 94.6778427 | -0.0703658 | 0.02801162 | -2.5120224 | 0.01200415 | 0.08393588 C7_active |
| 614 | chr7 | 71132169 | 71713600 | ENSG00000185 | WBSCR17 | 64409 | 177.336899 | -0.0576927 | 0.02186723 | -2.6383161 | 0.00833189 | 0.06410307 C7_active |
| 615 | chr3 | 134597801 | 135260467 | ENSG00000154 | EPHB1 | 2047 | 234.07993 | -0.0484717 | 0.01924193 | -2.519065 | 0.01176669 | 0.0827168 C7_active |
| 616 | chr5 | 167284799 | 168264157 | ENSG00000145 | TENM2 | 57451 | 299.811195 | 0.04425804 | 0.01762025 | 2.51177086 | 0.01201271 | 0.08395501 WT_active |
| 617 | chr14 | 67819779 | 68730218 | ENSG00000182 | RAD51B | 5890 | 246.383392 | 0.04705632 | 0.01741362 | 2.70227089 | 0.00688676 | 0.05544129 WT_active |
| 618 | chr3 | 55508308 | 56468363 | ENSG00000187 | ERC2 | 26059 | 294.419006 | 0.04745484 | 0.01666552 | 2.84748695 | 0.00440659 | 0.03938885 WT_active |
| 619 | chr3 | 29280982 | 30010391 | ENSG00000144 | RBMS3 | 27303 | 167.906136 | 0.04900825 | 0.02014599 | 2.43265477 | 0.01498858 | 0.09876775 WT_active |
| 620 | chr5 | 168661733 | 169301129 | ENSG00000184 | SLIT3 | 6586 | 223.42391 | 0.05069986 | 0.01964358 | 2.58098916 | 0.00985177 | 0.07273056 WT_active |
| 621 | chr2 | 45651345 | 46187990 | ENSG00000171 | PRKCE | 5581 | 173.034226 | 0.05199286 | 0.0194134 | 2.67819395 | 0.00740203 | 0.05886516 WT_active |
| 622 | chr18 | 36297714 | 36780055 | ENSG00000134 | FHOD3 | 80206 | 148.683552 | 0.05367328 | 0.02129075 | 2.52096699 | 0.01170328 | 0.08235136 WT_active |
| 623 | chrX | 31097677 | 33339441 | ENSG00000198 | DMD | 1756 | 535.690708 | 0.05374288 | 0.01825288 | 2.94435082 | 0.00323633 | 0.03077928 WT_active |
| 624 | chr15 | 71096952 | 71783383 | ENSG00000187 | THSD4 | 79875 | 206.427325 | 0.05697113 | 0.01707001 | 3.33749858 | 0.00084536 | 0.01005202 WT_active |
| 625 | chr3 | 65353525 | 66038834 | ENSG00000151 | MAGI1 | 9223 | 186.585742 | 0.05711581 | 0.01912034 | 2.98717543 | 0.00281568 | 0.02736251 WT_active |
| 626 | chr9 | 75890644 | 76362339 | ENSG00000099 | PCSK5 | 5125 | 141.30094 | 0.05717206 | 0.02238844 | 2.55364168 | 0.01066029 | 0.07715553 WT_active |
| 627 | chr3 | 54122547 | 55074557 | ENSG00000157 | CACNA2D3 | 55799 | 305.487736 | 0.05782471 | 0.01629041 | 3.54961699 | 0.00038579 | 0.00515988 WT_active |
| 628 | chr10 | 50991358 | 52298350 | ENSG00000185 | PRKG1 | 5592 | 322.58098 | 0.05795188 | 0.01812391 | 3.19753681 | 0.00138607 | 0.01529022 WT_active |
| 629 | chr10 | 60026298 | 60733490 | ENSG00000151 | ANK3 | 288 | 176.781802 | 0.05844693 | 0.01991288 | 2.93513168 | 0.00333406 | 0.03156303 WT_active |
| 630 | chr6 | 44809317 | 45377953 | ENSG00000196 | SUPT3H | 8464 | 126.910488 | 0.05860183 | 0.02386774 | 2.45527334 | 0.01407775 | 0.09471434 WT_active |
| 631 | chr7 | 69598919 | 70793068 | ENSG00000158 | AUTS2 | 26053 | 352.590783 | 0.05891737 | 0.01583085 | 3.72168087 | 0.0001979 | 0.00283632 WT_active |
| 632 | chrX | 41514934 | 41923463 | ENSG00000147 | CASK | 8573 | 104.698753 | 0.05969291 | 0.02443819 | 2.44260709 | 0.0145816 | 0.09728716 WT_active |
| 633 | chr8 | 35235457 | 35796550 | ENSG00000156 | UNC5D | 137970 | 159.577216 | 0.06009812 | 0.02384477 | 2.52039036 | 0.01172248 | 0.08244617 WT_active |
| 634 | chr2 | 72175984 | 72826041 | ENSG00000144 | EXOC6B | 23233 | 154.195171 | 0.06198202 | 0.02109582 | 2.93811867 | 0.00330211 | 0.03132221 WT_active |
| 635 | chr7 | 48171458 | 48647496 | ENSG00000179 | ABCA13 | 154664 | 127.575948 | 0.06215277 | 0.0243544 | 2.55201434 | 0.01071021 | 0.07731031 WT_active |
| 636 | chr8 | 47260575 | 47736306 | ENSG00000164 | SPIDR | 23514 | 126.910073 | 0.06220652 | 0.02183727 | 2.84864031 | 0.00439065 | 0.0392707 WT_active |
| 637 | chr2 | 27889941 | 28338901 | ENSG00000158 | BRE | 9577 | 113.878258 | 0.06270645 | 0.02188668 | 2.86505115 | 0.00416942 | 0.03777914 WT_active |
| 638 | chr14 | 37595847 | 38041442 | ENSG00000139 | TTC6 | 319089 | 111.798614 | 0.06361412 | 0.02253502 | 2.82289988 | 0.00475914 | 0.04171266 WT_active |
| 639 | chr3 | 2098813 | 3057956 | ENSG00000144 | CNTN4 | 152330 | 260.804096 | 0.06416445 | 0.0211806 | 3.02939702 | 0.00245042 | 0.02439932 WT_active |
| 640 | chr1 | 185734551 | 186190949 | ENSG00000143 | HMCN1 | 83872 | 106.47519 | 0.06416704 | 0.02634635 | 2.43551912 | 0.01487044 | 0.09843985 WT_active |
| 641 | chr18 | 7566782 | 8406861 | ENSG00000173 | PTPRM | 5797 | 231.305181 | 0.06427839 | 0.01891524 | 3.39823263 | 0.00067823 | 0.00837609 WT_active |
| 642 | chr2 | 15166909 | 15561330 | ENSG00000151 | NBAS | 51594 | 104.510616 | 0.06433773 | 0.02461972 | 2.61325943 | 0.00896832 | 0.06769907 WT_active |
| 643 | chr7 | 141074232 | 141480380 | ENSG00000261 | TMEM178B | 100507421 | 139.581128 | 0.06492176 | 0.0210893 | 3.07842147 | 0.002081 | 0.02129079 WT_active |
| 644 | chr5 | 170861870 | 171300015 | ENSG00000204 | RANBP17 | 64901 | 110.158455 | 0.06500233 | 0.02592377 | 2.50744138 | 0.01216088 | 0.08449903 WT_active |
| 645 | chr21 | 20998315 | 21543329 | ENSG00000154 | NCAM2 | 4685 | 108.685364 | 0.065034 | 0.02643158 | 2.46046614 | 0.01387567 | 0.09379248 WT_active |

| | | | | | | | | | | | | | | |
|---|---|---|---|---|---|---|---|---|---|---|---|---|---|---|
| 646 | chr1 | 231626815 | 232041272 | ENSG00000162 | DISC1 | 27185 | 130.326696 | | 0.06525212 | 0.02376121 | 2.74616167 | 0.0060297 | 0.05002087 | WT_active |
| 647 | chr1 | 171841498 | 172418466 | ENSG00000197 | DNM3 | 26052 | 144.640356 | | 0.06546258 | 0.02171966 | 3.01397826 | 0.00257846 | 0.02541099 | WT_active |
| 648 | chrX | 11137543 | 11665701 | ENSG00000047 | ARHGAP6 | 395 | 148.050681 | | 0.06607045 | 0.0218806 | 3.01959026 | 0.00253117 | 0.02503044 | WT_active |
| 649 | chr22 | 27978014 | 28679865 | ENSG00000100 | TTC28 | 23331 | 185.572218 | | 0.06651494 | 0.01904591 | 3.49234813 | 0.00047879 | 0.00621355 | WT_active |
| 650 | chr12 | 78863993 | 79452008 | ENSG00000067 | SYT1 | 6857 | 129.645176 | | 0.06652501 | 0.02587355 | 2.57115896 | 0.01013588 | 0.07418229 | WT_active |
| 651 | chr1 | 85729233 | 86156943 | ENSG00000171 | COL24A1 | 255631 | 100.770048 | | 0.06652983 | 0.02694392 | 2.4691959 | 0.01354171 | 0.09224839 | WT_active |
| 652 | chr5 | 58969038 | 60522120 | ENSG00000113 | PDE4D | 5144 | 407.076145 | | 0.06665382 | 0.01774326 | 3.75657022 | 0.00017226 | 0.00250435 | WT_active |
| 653 | chr12 | 10845849 | 11171598 | ENSG00000275 | PRH1 | 100533464 | 74.4794332 | | 0.06672023 | 0.0273313 | 2.44116584 | 0.01463993 | 0.09745084 | WT_active |
| 654 | chr12 | 10845849 | 11171613 | ENSG00000111 | PRR4 | 11272 | 74.4794332 | | 0.06672023 | 0.0273313 | 2.44116584 | 0.01463993 | 0.09745084 | WT_active |
| 655 | chr6 | 161347420 | 162727771 | ENSG00000185 | PARK2 | 5071 | 393.133093 | | 0.06699641 | 0.01650766 | 4.05850424 | 4.94E-05 | 0.00079032 | WT_active |
| 656 | chr8 | 140657900 | 141002216 | ENSG00000169 | PTK2 | 5747 | 79.627196 | | 0.06746954 | 0.02637593 | 2.55799627 | 0.01052772 | 0.07642935 | WT_active |
| 657 | chr12 | 61708259 | 62279150 | ENSG00000198 | FAM19A2 | 338811 | 145.541684 | | 0.06763198 | 0.02473926 | 2.73379168 | 0.00626096 | 0.05161268 | WT_active |
| 658 | chr7 | 116953238 | 117230103 | ENSG00000004 | ST7 | 7982 | 70.1336621 | | 0.06772324 | 0.02758553 | 2.45502802 | 0.01408736 | 0.09471434 | WT_active |
| 659 | chr7 | 126438598 | 127253294 | ENSG00000179 | GRM8 | 2918 | 220.089706 | | 0.06780876 | 0.0202194 | 3.35364812 | 0.00079754 | 0.00953474 | WT_active |
| 660 | chr14 | 71932439 | 72566529 | ENSG00000182 | RGS6 | 9628 | 205.112108 | | 0.06843478 | 0.02025734 | 3.37827027 | 0.00072943 | 0.00885724 | WT_active |
| 661 | chr2 | 43230836 | 43596046 | ENSG00000115 | THADA | 63892 | 96.3353558 | | 0.06855687 | 0.02340449 | 2.92921927 | 0.00339815 | 0.03210647 | WT_active |
| 662 | chr1 | 50437028 | 50960263 | ENSG00000185 | FAF1 | 11124 | 130.041465 | | 0.06870352 | 0.02199482 | 3.12362348 | 0.00178639 | 0.01886898 | WT_active |
| 663 | chr5 | 94152966 | 94618604 | ENSG00000185 | KIAA0825 | 285600 | 113.127312 | | 0.06895901 | 0.02490997 | 2.76832947 | 0.00563445 | 0.04739641 | WT_active |
| 664 | chr7 | 38977998 | 39493095 | ENSG00000106 | POU6F2 | 11281 | 148.682199 | | 0.06900938 | 0.02401371 | 2.87374928 | 0.00405631 | 0.03694495 | WT_active |
| 665 | chr4 | 61201258 | 62078335 | ENSG00000150 | LPHN3 | 23284 | 193.548707 | | 0.06902274 | 0.02542041 | 2.71524869 | 0.0066226 | 0.05403773 | WT_active |
| 666 | chr6 | 25279078 | 25620530 | ENSG00000079 | LRRC16A | 55604 | 93.2227503 | | 0.06906094 | 0.02721123 | 2.53795706 | 0.01115017 | 0.07946767 | WT_active |
| 667 | chr3 | 24117160 | 24495282 | ENSG00000151 | THRB | 7068 | 104.399538 | | 0.06978837 | 0.02619947 | 2.66373196 | 0.00772791 | 0.0607199 | WT_active |
| 668 | chr1 | 239386565 | 239915452 | ENSG00000133 | CHRM3 | 1131 | 144.349068 | | 0.070213 | 0.0259647 | 2.7041711 | 0.0068475 | 0.0552166 | WT_active |
| 669 | chr5 | 128257909 | 128659185 | ENSG00000138 | FBN2 | 2201 | 99.6532553 | | 0.07047764 | 0.02751424 | 2.5614963 | 0.01042224 | 0.07581624 | WT_active |
| 670 | chr9 | 77716087 | 78031458 | ENSG00000156 | GNAQ | 2776 | 75.0902717 | | 0.07058019 | 0.02755406 | 2.56151657 | 0.01042163 | 0.07581624 | WT_active |
| 671 | chr4 | 37891087 | 38139175 | ENSG00000065 | TBC1D1 | 23216 | 69.5401234 | | 0.07083683 | 0.02737838 | 2.58732759 | 0.00967236 | 0.07177358 | WT_active |
| 672 | chr5 | 15500196 | 15939795 | ENSG00000183 | FBXL7 | 23194 | 120.016178 | | 0.07103648 | 0.02215328 | 3.20659 | 0.00134318 | 0.01487405 | WT_active |
| 673 | chr7 | 41960950 | 42237870 | ENSG00000106 | GLI3 | 2737 | 80.9697528 | | 0.07113855 | 0.02797247 | 2.54316274 | 0.01098541 | 0.07870279 | WT_active |
| 674 | chr1 | 107571160 | 107965144 | ENSG00000134 | VAV3 | 10451 | 108.147354 | | 0.07121029 | 0.02439769 | 2.91873097 | 0.00351459 | 0.03299051 | WT_active |
| 675 | chr11 | 17788048 | 18013162 | ENSG00000129 | SERGEF | 26297 | 78.9007891 | | 0.07129577 | 0.0271308 | 2.62785333 | 0.00859256 | 0.06552567 | WT_active |
| 676 | chr7 | 137381037 | 137847092 | ENSG00000157 | DGKI | 9162 | 128.966718 | | 0.07179026 | 0.0201202 | 3.56806943 | 0.00035962 | 0.00485944 | WT_active |
| 677 | chr1 | 56994778 | 58546734 | ENSG00000173 | DAB1 | 1600 | 512.181228 | | 0.07197077 | 0.01897097 | 3.793732 | 0.0001484 | 0.00218107 | WT_active |
| 678 | chr12 | 18247614 | 18648416 | ENSG00000139 | PIK3C2G | 5288 | 91.5285433 | | 0.0723155 | 0.02732447 | 2.64654701 | 0.00813182 | 0.06288789 | WT_active |
| 679 | chr9 | 84668551 | 85027070 | ENSG00000148 | NTRK2 | 4915 | 116.072697 | | 0.07240928 | 0.02443538 | 2.96329709 | 0.00304363 | 0.02923306 | WT_active |
| 680 | chr6 | 39329990 | 39725405 | ENSG00000164 | KIF6 | 221458 | 112.221549 | | 0.07245364 | 0.02305427 | 3.14274268 | 0.00167373 | 0.01791232 | WT_active |
| 681 | chr1 | 59296638 | 59767675 | ENSG00000172 | FGGY | 55277 | 143.486397 | | 0.07249455 | 0.02261921 | 3.20499958 | 0.00135063 | 0.014945 | WT_active |
| 682 | chr9 | 121821928 | 122093606 | ENSG00000175 | TTLL11 | 158135 | 85.6220845 | | 0.07289631 | 0.02571515 | 2.83476177 | 0.00458599 | 0.04058978 | WT_active |
| 683 | chr1 | 210678315 | 211134115 | ENSG00000143 | KCNH1 | 3756 | 141.320916 | | 0.07327209 | 0.02365308 | 3.09778168 | 0.00194975 | 0.02018054 | WT_active |
| 684 | chr10 | 53802771 | 55627942 | ENSG00000150 | PCDH15 | 65217 | 378.464927 | | 0.07386205 | 0.026652 | 2.77135081 | 0.00558242 | 0.04706866 | WT_active |
| 685 | chr1 | 240014348 | 240475189 | ENSG00000155 | FMN2 | 56776 | 109.395672 | | 0.07459894 | 0.02728209 | 2.73435624 | 0.00625024 | 0.05155374 | WT_active |
| 686 | chr5 | 139846779 | 140043299 | ENSG00000158 | NRG2 | 9542 | 71.1012917 | | 0.07465585 | 0.02832333 | 2.63584308 | 0.00839285 | 0.06436604 | WT_active |
| 687 | chr5 | 142770384 | 143229011 | ENSG00000145 | ARHGAP26 | 23092 | 136.286137 | | 0.07498937 | 0.02085688 | 3.59542619 | 0.00032386 | 0.004426 | WT_active |
| 688 | chr20 | 17226107 | 17484578 | ENSG00000125 | PCSK2 | 5126 | 85.4506685 | | 0.07507723 | 0.02834841 | 2.64837585 | 0.00808796 | 0.06275669 | WT_active |
| 689 | chr8 | 132866958 | 133134903 | ENSG00000042 | TG | 7038 | 96.0355335 | | 0.07515059 | 0.02481747 | 3.02813253 | 0.0024607 | 0.02448474 | WT_active |
| 690 | chr4 | 71187286 | 71572087 | ENSG00000080 | SLC4A4 | 8671 | 99.9461052 | | 0.07522626 | 0.02667973 | 2.81960351 | 0.0048083 | 0.04206681 | WT_active |
| 691 | chr18 | 79069285 | 79378283 | ENSG00000166 | ATP9B | 374868 | 83.8004087 | | 0.07537955 | 0.02692846 | 2.7992525 | 0.00512211 | 0.0439325 | WT_active |
| 692 | chr2 | 54723499 | 54972025 | ENSG00000214 | EML6 | 400954 | 70.8394883 | | 0.07549177 | 0.02809077 | 2.68742233 | 0.00720058 | 0.05751691 | WT_active |
| 693 | chr1 | 233904933 | 234324516 | ENSG00000183 | SLC35F3 | 148641 | 132.435871 | | 0.07573894 | 0.02524684 | 2.99993732 | 0.00270035 | 0.02643155 | WT_active |
| 694 | chr2 | 177623252 | 178108339 | ENSG00000128 | PDE11A | 50940 | 134.756691 | | 0.07574129 | 0.02292292 | 3.30417363 | 0.00095257 | 0.01111976 | WT_active |
| 695 | chr9 | 16409503 | 16870843 | ENSG00000173 | BNC2 | 54796 | 112.446847 | | 0.07584585 | 0.02285599 | 3.31842343 | 0.00090527 | 0.01064617 | WT_active |
| 696 | chr15 | 57953424 | 58497866 | ENSG00000128 | ALDH1A2 | 8854 | 163.706656 | | 0.07585922 | 0.01925748 | 3.93920774 | 8.18E-05 | 0.00125126 | WT_active |
| 697 | chr2 | 63121383 | 63827843 | ENSG00000143 | WDPCP | 51057 | 182.226444 | | 0.07594051 | 0.01972977 | 3.84903168 | 0.00011859 | 0.00177916 | WT_active |
| 698 | chr11 | 4704927 | 4992429 | ENSG00000167 | MMP26 | 56547 | 79.80305 | | 0.07602067 | 0.02981049 | 2.55013108 | 0.01076824 | 0.07755071 | WT_active |
| 699 | chr1 | 76074719 | 76634601 | ENSG00000184 | ST6GALNAC3 | 256435 | 166.591448 | | 0.0763252 | 0.02368739 | 3.22218726 | 0.00127216 | 0.01422963 | WT_active |

| | | | | | | | | | | | | | |
|---|---|---|---|---|---|---|---|---|---|---|---|---|---|
| 700 | chr6 | 45328157 | 45664349 | ENSG00000124 | RUNX2 | 860 | 87.9326159 | 0.07646275 | 0.02507517 | 3.04934082 | 0.00229344 | 0.0231075 | WT_active |
| 701 | chr6 | 146027646 | 146437598 | ENSG00000152 | GRM1 | 2911 | 109.11839 | 0.07672769 | 0.02538025 | 3.02312582 | 0.00250178 | 0.02482498 | WT_active |
| 702 | chr15 | 49326962 | 49620931 | ENSG00000166 | FAM227B | 196951 | 68.3801783 | 0.07718194 | 0.03014527 | 2.56033352 | 0.01045718 | 0.07603205 | WT_active |
| 703 | chr5 | 107376889 | 107670895 | ENSG00000184 | EFNA5 | 1946 | 74.4744451 | 0.07737802 | 0.02903153 | 2.66530966 | 0.00769175 | 0.06050169 | WT_active |
| 704 | chr4 | 113451032 | 113761927 | ENSG00000145 | CAMK2D | 817 | 64.4547144 | 0.07756633 | 0.0319843 | 2.42513796 | 0.01530257 | 0.0999694 | WT_active |
| 705 | chr1 | 180632004 | 180890251 | ENSG00000143 | XPR1 | 9213 | 56.0186192 | 0.07756841 | 0.03120504 | 2.48576566 | 0.01292731 | 0.08875519 | WT_active |
| 706 | chr6 | 118460772 | 118710075 | ENSG00000111 | CEP85L | 387119 | 61.6377588 | 0.07763059 | 0.03056661 | 2.53971865 | 0.01109417 | 0.07910767 | WT_active |
| 707 | chr4 | 46993723 | 47426444 | ENSG00000163 | GABRB1 | 2560 | 106.327406 | 0.07786665 | 0.02519976 | 3.08997613 | 0.00200173 | 0.02061492 | WT_active |
| 708 | chr2 | 179441982 | 179861505 | ENSG00000144 | ZNF385B | 151126 | 107.398682 | 0.07802395 | 0.02962392 | 2.63381587 | 0.00844313 | 0.06464843 | WT_active |
| 709 | chr17 | 60677453 | 61392838 | ENSG00000141 | BCAS3 | 54828 | 174.320163 | 0.07832873 | 0.02152138 | 3.63957795 | 0.00027309 | 0.00380788 | WT_active |
| 710 | chr1 | 162069774 | 162370475 | ENSG00000198 | NOS1AP | 9722 | 94.9539132 | 0.07848404 | 0.0257134 | 3.05226269 | 0.00227123 | 0.02296398 | WT_active |
| 711 | chr7 | 27830573 | 28180743 | ENSG00000153 | JAZF1 | 221895 | 101.042566 | 0.07852614 | 0.02445829 | 3.21061467 | 0.00132451 | 0.01470119 | WT_active |
| 712 | chr4 | 52872982 | 53366075 | ENSG00000184 | SCFD2 | 152579 | 136.37677 | 0.07858753 | 0.02306132 | 3.40776433 | 0.00065497 | 0.00813387 | WT_active |
| 713 | chr3 | 3700814 | 4467281 | ENSG00000144 | SUMF1 | 285362 | 204.047549 | 0.0792988 | 0.02108628 | 3.76068255 | 0.00016945 | 0.00247281 | WT_active |
| 714 | chr4 | 20728616 | 21948799 | ENSG00000185 | KCNIP4 | 80333 | 329.476755 | 0.07980655 | 0.01886276 | 4.2309062 | 2.33E-05 | 0.00039387 | WT_active |
| 715 | chr1 | 215622894 | 216423396 | ENSG00000042 | USH2A | 7399 | 197.203278 | 0.07982618 | 0.02301073 | 3.46908556 | 0.00052223 | 0.00671272 | WT_active |
| 716 | chr14 | 60734742 | 60969953 | ENSG00000020 | MNAT1 | 4331 | 49.856592 | 0.07989691 | 0.03090845 | 2.58495374 | 0.00973921 | 0.07215821 | WT_active |
| 717 | chr5 | 76403249 | 76708132 | ENSG00000145 | IQGAP2 | 10788 | 86.5563628 | 0.0799024 | 0.02862646 | 2.7912085 | 0.00525116 | 0.04479957 | WT_active |
| 718 | chr4 | 42408373 | 42657105 | ENSG00000124 | ATP8A1 | 10396 | 68.1924736 | 0.08005075 | 0.03209651 | 2.49406364 | 0.01262899 | 0.08699703 | WT_active |
| 719 | chrX | 35919734 | 36385319 | ENSG00000165 | CXorf22 | 286464 | 101.791052 | 0.08014139 | 0.03026471 | 2.64801433 | 0.00809661 | 0.06276178 | WT_active |
| 720 | chr2 | 100820152 | 100996829 | ENSG00000170 | NPAS2 | 4862 | 54.3300297 | 0.08042971 | 0.03311461 | 2.42882876 | 0.01514768 | 0.09933654 | WT_active |
| 721 | chr3 | 170418865 | 170860380 | ENSG00000013 | CLDN11 | 5010 | 132.758867 | 0.08046234 | 0.02159078 | 3.72669867 | 0.000194 | 0.00278324 | WT_active |
| 722 | chr8 | 6406592 | 6648504 | ENSG00000147 | MCPH1 | 79648 | 67.7221584 | 0.08050985 | 0.02830435 | 2.84443384 | 0.00444904 | 0.03966994 | WT_active |
| 723 | chr1 | 224896262 | 225399292 | ENSG00000185 | DNAH14 | 127602 | 129.546672 | 0.0805959 | 0.02484049 | 3.24453698 | 0.00117642 | 0.01327198 | WT_active |
| 724 | chr11 | 15966449 | 16739591 | ENSG00000110 | SOX6 | 55553 | 186.591644 | 0.0807639 | 0.02042547 | 3.95407784 | 7.68E-05 | 0.00118475 | WT_active |
| 725 | chr4 | 37453941 | 37623495 | ENSG00000154 | C4orf19 | 55286 | 50.5194686 | 0.08078895 | 0.03318417 | 2.43456328 | 0.01490977 | 0.09860968 | WT_active |
| 726 | chr3 | 42809446 | 43055871 | ENSG00000273 | RP11-136C24.3 | NaN | 81.6093995 | 0.08081461 | 0.02719059 | 2.97215342 | 0.00295719 | 0.02855777 | WT_active |
| 727 | chr8 | 99013266 | 99877580 | ENSG00000132 | VPS13B | 157680 | 194.013533 | 0.08103339 | 0.02056441 | 3.94046799 | 8.13E-05 | 0.00124804 | WT_active |
| 728 | chr17 | 65635538 | 66192084 | ENSG00000154 | CEP112 | 201134 | 133.783869 | 0.08104531 | 0.02395164 | 3.38370655 | 0.00071514 | 0.00873198 | WT_active |
| 729 | chr6 | 9596110 | 10211608 | ENSG00000181 | OFCC1 | NaN | 162.013438 | 0.08131857 | 0.02328391 | 3.49247864 | 0.00047856 | 0.00621355 | WT_active |
| 730 | chr4 | 86015123 | 86594131 | ENSG00000109 | MAPK10 | 5602 | 144.155863 | 0.08143912 | 0.02513907 | 3.23954351 | 0.00119721 | 0.01349601 | WT_active |
| 731 | chr8 | 101492432 | 101669726 | ENSG00000083 | GRHL2 | 79977 | 50.5918504 | 0.08149139 | 0.03257644 | 2.50154356 | 0.01236532 | 0.08563075 | WT_active |
| 732 | chr7 | 86643914 | 86864884 | ENSG00000198 | GRM3 | 2913 | 59.6415493 | 0.08149462 | 0.03299184 | 2.47014511 | 0.01350583 | 0.09211305 | WT_active |
| 733 | chr2 | 76747719 | 77593319 | ENSG00000176 | LRRTM4 | 80059 | 181.841222 | 0.08161322 | 0.02542676 | 3.20973785 | 0.00132856 | 0.01473476 | WT_active |
| 734 | chr9 | 17134982 | 17503923 | ENSG00000044 | CNTLN | 54875 | 83.1775633 | 0.08199115 | 0.02785984 | 2.94298727 | 0.00325062 | 0.03088865 | WT_active |
| 735 | chr1 | 75202131 | 75611116 | ENSG00000137 | SLC44A5 | 204962 | 103.161028 | 0.08207044 | 0.02823941 | 2.90623765 | 0.00365804 | 0.03418117 | WT_active |
| 736 | chr3 | 99817834 | 100178603 | ENSG00000184 | CMSS1 | 84319 | 91.9486747 | 0.08230835 | 0.0261544 | 3.1470178 | 0.00164945 | 0.01770794 | WT_active |
| 737 | chr18 | 32124877 | 32470484 | ENSG00000141 | GAREM | 64762 | 94.2136055 | 0.08238601 | 0.02683183 | 3.07045777 | 0.00213731 | 0.0217933 | WT_active |
| 738 | chr5 | 111662621 | 111997464 | ENSG00000134 | NREP | 9315 | 88.8778073 | 0.08242759 | 0.02781783 | 2.9631214 | 0.00304536 | 0.02923306 | WT_active |
| 739 | chr4 | 118033618 | 118258648 | ENSG00000164 | NDST3 | 9348 | 55.6483806 | 0.08262735 | 0.03263204 | 2.5320928 | 0.0113384 | 0.08053055 | WT_active |
| 740 | chr10 | 48446034 | 48656265 | ENSG00000128 | ARHGAP22 | 58504 | 63.834558 | 0.08293309 | 0.03183681 | 2.60494374 | 0.00918893 | 0.06893135 | WT_active |
| 741 | chr11 | 46936689 | 47164385 | ENSG00000149 | C11orf49 | 79096 | 58.5047751 | 0.08296465 | 0.02859525 | 2.90134376 | 0.00371566 | 0.03460749 | WT_active |
| 742 | chr9 | 113876282 | 114056591 | ENSG00000157 | ZNF618 | 114991 | 57.2613353 | 0.08308972 | 0.03215299 | 2.58419852 | 0.00976056 | 0.07224217 | WT_active |
| 743 | chr8 | 98401403 | 98942827 | ENSG00000104 | STK3 | 6788 | 140.732414 | 0.0831464 | 0.02160399 | 3.84865954 | 0.00011877 | 0.00178001 | WT_active |
| 744 | chr3 | 157259742 | 157533619 | ENSG00000197 | VEPH1 | 79674 | 66.7621241 | 0.08327592 | 0.02804028 | 2.96986732 | 0.00297928 | 0.02869427 | WT_active |
| 745 | chr14 | 66507407 | 67181803 | ENSG00000171 | GPHN | 10243 | 156.846258 | 0.08355151 | 0.02175612 | 3.84036751 | 0.00012285 | 0.0018336 | WT_active |
| 746 | chr6 | 154154496 | 154356792 | ENSG00000074 | IPCEF1 | 26034 | 54.6670577 | 0.08356536 | 0.03251879 | 2.56975637 | 0.01017701 | 0.07440774 | WT_active |
| 747 | chr7 | 88759368 | 89337057 | ENSG00000182 | ZNF804B | 219578 | 127.036342 | 0.08361065 | 0.02892693 | 2.89040837 | 0.00384742 | 0.03558182 | WT_active |
| 748 | chr4 | 24806117 | 24980204 | ENSG00000181 | CCDC149 | 91050 | 55.8975917 | 0.0837855 | 0.03243637 | 2.5830725 | 0.00979247 | 0.07240405 | WT_active |
| 749 | chr6 | 122471917 | 122726373 | ENSG00000135 | PKIB | 5570 | 63.8181952 | 0.08391684 | 0.02985673 | 2.81065043 | 0.00494415 | 0.04289092 | WT_active |
| 750 | chr7 | 90466424 | 91210590 | ENSG00000058 | CDK14 | 5218 | 189.669413 | 0.08426451 | 0.01890613 | 4.45699382 | 8.31E-06 | 0.00014777 | WT_active |
| 751 | chr4 | 92303622 | 93774556 | ENSG00000152 | GRID2 | 2895 | 321.85564 | 0.08434069 | 0.02737259 | 3.0812099 | 0.00206161 | 0.02114106 | WT_active |
| 752 | chr2 | 25377198 | 25673647 | ENSG00000138 | DTNB | 1838 | 78.9460067 | 0.08461485 | 0.02632683 | 3.21401611 | 0.00130892 | 0.014565 | WT_active |
| 753 | chr9 | 15552897 | 16061663 | ENSG00000164 | CCDC171 | 203238 | 127.475462 | 0.08472537 | 0.02330071 | 3.63617162 | 0.00027672 | 0.00385483 | WT_active |

| | | | | | | | | | | | | | |
|---|---|---|---|---|---|---|---|---|---|---|---|---|---|
| 754 | chr2 | 213284379 | 214410501 | ENSG00000144 | SPAG16 | 79582 | 260.819392 | 0.08477585 | 0.02424854 | 3.49612225 | 0.00047207 | 0.00614845 | WT_active |
| 755 | chr5 | 77425970 | 77620611 | ENSG00000164 | WDR41 | 55255 | 47.8520035 | 0.08478834 | 0.03485248 | 2.43277819 | 0.01498348 | 0.09876775 | WT_active |
| 756 | chr17 | 32492522 | 32877177 | ENSG00000176 | MYO1D | 4642 | 106.071693 | 0.0851431 | 0.02526633 | 3.36982474 | 0.00075216 | 0.00904474 | WT_active |
| 757 | chr5 | 35617844 | 35814611 | ENSG00000152 | SPEF2 | 79925 | 50.2824358 | 0.08515402 | 0.03455916 | 2.46400684 | 0.01373935 | 0.09317683 | WT_active |
| 758 | chr2 | 99545419 | 100142739 | ENSG00000144 | AFF3 | 3899 | 164.642 | 0.08546039 | 0.02208075 | 3.87035617 | 0.00010868 | 0.00163901 | WT_active |
| 759 | chr3 | 158962235 | 159897366 | ENSG00000250 | IQCJ-SCHIP1 | NaN | 253.512635 | 0.08581094 | 0.0178549 | 4.80601534 | 1.54E-06 | 2.91E-05 | WT_active |
| 760 | chr3 | 99830141 | 100114513 | ENSG00000168 | FILIP1L | 11259 | 72.3595736 | 0.086065 | 0.02831322 | 3.03974624 | 0.00236778 | 0.02370735 | WT_active |
| 761 | chrX | 123960212 | 124422664 | ENSG00000101 | STAG2 | 10735 | 119.626642 | 0.08608828 | 0.02441809 | 3.52559377 | 0.00042253 | 0.00558396 | WT_active |
| 762 | chr12 | 26336515 | 26833198 | ENSG00000123 | ITPR2 | 3709 | 128.830716 | 0.08618116 | 0.02214779 | 3.891186 | 9.98E-05 | 0.00151397 | WT_active |
| 763 | chr7 | 146116002 | 148420998 | ENSG00000174 | CNTNAP2 | 26047 | 582.200599 | 0.08670549 | 0.01613767 | 5.37286303 | 7.75E-08 | 1.60E-06 | WT_active |
| 764 | chr12 | 96489571 | 96875555 | ENSG00000188 | C12orf55 | 144535 | 99.9818025 | 0.08702135 | 0.02621289 | 3.31979169 | 0.00090085 | 0.01061144 | WT_active |
| 765 | chr2 | 151485336 | 151734487 | ENSG00000183 | NEB | 4703 | 60.8678359 | 0.08721131 | 0.0341138 | 2.55648189 | 0.01057366 | 0.0767242 | WT_active |
| 766 | chr6 | 114055586 | 114343045 | ENSG00000249 | HS3ST5 | 222537 | 79.383527 | 0.08729748 | 0.02915761 | 2.99398574 | 0.00275359 | 0.02682516 | WT_active |
| 767 | chr2 | 169827458 | 170084131 | ENSG00000144 | UBR3 | 130507 | 53.7051482 | 0.08731091 | 0.03578543 | 2.43984552 | 0.01469354 | 0.09757217 | WT_active |
| 768 | chr5 | 78000525 | 78294755 | ENSG00000132 | AP3B1 | 8546 | 65.2629236 | 0.08768918 | 0.03187918 | 2.75067261 | 0.0059473 | 0.04953329 | WT_active |
| 769 | chr2 | 196194370 | 196593692 | ENSG00000138 | HECW2 | 57520 | 107.401785 | 0.08773816 | 0.02620131 | 3.34861728 | 0.00081216 | 0.00969347 | WT_active |
| 770 | chr12 | 11649854 | 11895402 | ENSG00000139 | ETV6 | 2120 | 73.6677271 | 0.08785697 | 0.02914611 | 3.01436329 | 0.00257519 | 0.02539612 | WT_active |
| 771 | chr2 | 84516455 | 84819589 | ENSG00000115 | DNAH6 | 1768 | 83.9277913 | 0.08790857 | 0.02778786 | 3.16356026 | 0.00155852 | 0.01688261 | WT_active |
| 772 | chr3 | 133038391 | 133397792 | ENSG00000144 | TMEM108 | 66000 | 113.49358 | 0.08796519 | 0.02756186 | 3.19155534 | 0.00141509 | 0.01556275 | WT_active |
| 773 | chr1 | 213051233 | 213274773 | ENSG00000136 | RPS6KC1 | 26750 | 51.4704763 | 0.08798719 | 0.03300325 | 2.66601556 | 0.00767561 | 0.06044074 | WT_active |
| 774 | chr7 | 133253073 | 134066589 | ENSG00000131 | EXOC4 | 60412 | 219.308424 | 0.08802805 | 0.01760827 | 4.99924596 | 5.76E-07 | 1.13E-05 | WT_active |
| 775 | chr3 | 151085697 | 151437072 | ENSG00000144 | MED12L | 116931 | 94.136701 | 0.08810576 | 0.02606598 | 3.3801051 | 0.00072458 | 0.0088225 | WT_active |
| 776 | chr3 | 62398346 | 62875389 | ENSG00000163 | CADPS | 8618 | 143.837494 | 0.08848674 | 0.02256806 | 3.92088358 | 8.82E-05 | 0.00134606 | WT_active |
| 777 | chr22 | 35738736 | 36028425 | ENSG00000100 | RBFOX2 | 23543 | 69.771963 | 0.08862627 | 0.02694459 | 3.28920396 | 0.00100471 | 0.01164464 | WT_active |
| 778 | chr9 | 35161992 | 35405338 | ENSG00000198 | UNC13B | 10497 | 71.6317295 | 0.0888668 | 0.02837203 | 3.13219749 | 0.00173503 | 0.01844814 | WT_active |
| 779 | chr2 | 136765545 | 137677717 | ENSG00000144 | THSD7B | 80731 | 241.502936 | 0.08896036 | 0.01978246 | 4.4969302 | 6.89E-06 | 0.00012363 | WT_active |
| 780 | chr2 | 132671799 | 133568463 | ENSG00000176 | NCKAP5 | 344148 | 254.428555 | 0.08903758 | 0.01819079 | 4.89465224 | 9.85E-07 | 1.90E-05 | WT_active |
| 781 | chr3 | 23203020 | 23591793 | ENSG00000182 | UBE2E2 | 7325 | 97.0003007 | 0.08912432 | 0.02616137 | 3.40671488 | 0.0006575 | 0.00815116 | WT_active |
| 782 | chr18 | 36829106 | 37232172 | ENSG00000150 | KIAA1328 | 57536 | 99.8700003 | 0.08936125 | 0.02526518 | 3.53693355 | 0.0004048 | 0.0053792 | WT_active |
| 783 | chr2 | 169127109 | 169362685 | ENSG00000081 | LRP2 | 4036 | 67.4360028 | 0.08944868 | 0.03177461 | 2.81509871 | 0.00487623 | 0.04245496 | WT_active |
| 784 | chr7 | 93591573 | 93911265 | ENSG00000127 | GNGT1 | 2792 | 74.8567446 | 0.08950091 | 0.0314538 | 2.84547216 | 0.00443456 | 0.03958981 | WT_active |
| 785 | chr8 | 97868840 | 98036716 | ENSG00000132 | MATN2 | 4147 | 49.5343325 | 0.08958051 | 0.03277633 | 2.73308568 | 0.0062744 | 0.05169389 | WT_active |
| 786 | chr14 | 32934933 | 33804176 | ENSG00000151 | NPAS3 | 64067 | 239.176826 | 0.09009657 | 0.01692249 | 5.32407416 | 1.01E-07 | 2.09E-06 | WT_active |
| 787 | chr14 | 31489956 | 31861224 | ENSG00000151 | NUBPL | 80224 | 95.723974 | 0.09014527 | 0.02566743 | 3.51204933 | 0.00044467 | 0.00584962 | WT_active |
| 788 | chr8 | 73420369 | 73747708 | ENSG00000040 | STAU2 | 27067 | 77.294249 | 0.09016158 | 0.02645713 | 3.40783706 | 0.0006548 | 0.00813387 | WT_active |
| 789 | chr21 | 38380027 | 38661780 | ENSG00000157 | ERG | 2078 | 84.9327997 | 0.09018995 | 0.02538179 | 3.55333263 | 0.00038038 | 0.00509699 | WT_active |
| 790 | chr20 | 37693955 | 37872129 | ENSG00000132 | CTNNBL1 | 56259 | 56.7755318 | 0.09062795 | 0.03416861 | 2.65237459 | 0.00799278 | 0.06230291 | WT_active |
| 791 | chr4 | 94757968 | 95158448 | ENSG00000138 | BMPR1B | 658 | 94.1334023 | 0.09063876 | 0.02660577 | 3.40673391 | 0.00065745 | 0.00815116 | WT_active |
| 792 | chr8 | 49909789 | 50794118 | ENSG00000147 | SNTG1 | 54212 | 205.418938 | 0.09102047 | 0.02119383 | 4.29466819 | 1.75E-05 | 0.0003021 | WT_active |
| 793 | chr3 | 37452115 | 37823514 | ENSG00000144 | ITGA9 | 3680 | 135.059065 | 0.09128957 | 0.02283506 | 3.99778087 | 6.39E-05 | 0.00100313 | WT_active |
| 794 | chr1 | 81306160 | 81992436 | ENSG00000117 | LPHN2 | 23266 | 167.421926 | 0.0913375 | 0.02434696 | 3.75149481 | 0.00017578 | 0.00254462 | WT_active |
| 795 | chr4 | 84669610 | 84966391 | ENSG00000163 | WDFY3 | 23001 | 65.6243523 | 0.09143288 | 0.0312349 | 2.9272662 | 0.00341956 | 0.03226651 | WT_active |
| 796 | chr12 | 91984976 | 92142914 | ENSG00000257 | C12orf79 | 256021 | 39.2990024 | 0.09159079 | 0.03750052 | 2.44238748 | 0.01459048 | 0.09730134 | WT_active |
| 797 | chr3 | 108380369 | 108529322 | ENSG00000144 | MYH15 | 22989 | 42.2722858 | 0.09164953 | 0.03631567 | 2.52369108 | 0.01161299 | 0.08195601 | WT_active |
| 798 | chr18 | 56651343 | 57036606 | ENSG00000091 | WDR7 | 23335 | 89.8258085 | 0.09183698 | 0.02867727 | 3.20243076 | 0.00136273 | 0.01505582 | WT_active |
| 799 | chr5 | 42423777 | 42721878 | ENSG00000112 | GHR | 2690 | 88.0078001 | 0.09196607 | 0.02683973 | 3.42649034 | 0.00061144 | 0.00765915 | WT_active |
| 800 | chr15 | 80779343 | 80951776 | ENSG00000103 | CEMIP | 57214 | 60.6596976 | 0.09214481 | 0.03412321 | 2.70035599 | 0.00692653 | 0.05569812 | WT_active |
| 801 | chr6 | 13357830 | 13487662 | ENSG00000145 | GFOD1 | 54438 | 40.2049391 | 0.09232981 | 0.03728985 | 2.47600354 | 0.01328623 | 0.09091638 | WT_active |
| 802 | chr2 | 197804702 | 198572581 | ENSG00000115 | PLCL1 | 5334 | 205.360333 | 0.09237285 | 0.02375617 | 3.88837285 | 0.00010092 | 0.00153001 | WT_active |
| 803 | chr1 | 63774022 | 64181498 | ENSG00000185 | ROR1 | 4919 | 115.085351 | 0.09239419 | 0.02277065 | 4.05760003 | 4.96E-05 | 0.0007925 | WT_active |
| 804 | chr6 | 111087503 | 111231194 | ENSG00000112 | SLC16A10 | 117247 | 39.1258912 | 0.0924665 | 0.03707207 | 2.49423632 | 0.01262285 | 0.08699628 | WT_active |
| 805 | chr18 | 74597870 | 75065671 | ENSG00000215 | ZNF407 | 55628 | 118.395947 | 0.09250464 | 0.02245995 | 4.11864796 | 3.81E-05 | 0.0006244 | WT_active |
| 806 | chr6 | 77690658 | 77926974 | ENSG00000269 | MEI4 | 101928601 | 59.6974137 | 0.0927148 | 0.0317399 | 2.92108023 | 0.0034882 | 0.03280683 | WT_active |
| 807 | chr5 | 94703741 | 95284575 | ENSG00000175 | MCTP1 | 79772 | 148.145362 | 0.09306819 | 0.02449982 | 3.7987302 | 0.00014544 | 0.00213974 | WT_active |

| | | | | | | | | | | | | | |
|---|---|---|---|---|---|---|---|---|---|---|---|---|---|
| 808 | chr6 | 109492856 | 109691217 | ENSG00000155 | AK9 | 221264 | 51.5754905 | 0.09317765 | 0.03437778 | 2.71040339 | 0.00672014 | 0.05457223 | WT_active |
| 809 | chr3 | 108958239 | 109118142 | ENSG00000114 | MORC1 | 27136 | 44.3949315 | 0.09318973 | 0.03750271 | 2.48487959 | 0.01295953 | 0.08893405 | WT_active |
| 810 | chr4 | 165873258 | 166103895 | ENSG00000038 | TLL1 | 7092 | 52.5574525 | 0.09347053 | 0.03648362 | 2.5619861 | 0.01040755 | 0.07578588 | WT_active |
| 811 | chr5 | 145936579 | 146081791 | ENSG00000156 | SH3RF2 | 153769 | 43.6582137 | 0.09355011 | 0.0363521 | 2.57344429 | 0.01006919 | 0.07380657 | WT_active |
| 812 | chr2 | 236324147 | 236507542 | ENSG00000132 | IQCA1 | 79781 | 54.7311544 | 0.09355917 | 0.03457052 | 2.70632794 | 0.00680318 | 0.05504393 | WT_active |
| 813 | chr4 | 88725955 | 89111398 | ENSG00000138 | FAM13A | 10144 | 97.9789164 | 0.09362315 | 0.02473598 | 3.78489675 | 0.00015377 | 0.00224943 | WT_active |
| 814 | chr7 | 26667062 | 26995239 | ENSG00000005 | SKAP2 | 8935 | 82.8090018 | 0.0938667 | 0.03143086 | 2.98644998 | 0.00282237 | 0.02740265 | WT_active |
| 815 | chr11 | 28108248 | 28333507 | ENSG00000169 | METTL15 | 196074 | 49.6502166 | 0.09395677 | 0.03473874 | 2.70466852 | 0.00683726 | 0.05519574 | WT_active |
| 816 | chr15 | 84980440 | 85139145 | ENSG00000073 | PDE8A | 5151 | 45.3991477 | 0.09409042 | 0.03262126 | 2.88432815 | 0.0039225 | 0.03602202 | WT_active |
| 817 | chr6 | 151494030 | 151621193 | ENSG00000120 | CCDC170 | 80129 | 35.3477025 | 0.09424061 | 0.03827381 | 2.46227398 | 0.01380592 | 0.09354027 | WT_active |
| 818 | chr5 | 26880600 | 27121150 | ENSG00000113 | CDH9 | 1007 | 49.3988995 | 0.09427836 | 0.03587738 | 2.6277941 | 0.00859405 | 0.06552567 | WT_active |
| 819 | chr3 | 36380344 | 36548007 | ENSG00000144 | STAC | 6769 | 48.3466646 | 0.09447795 | 0.03327089 | 2.83965789 | 0.00451619 | 0.04014457 | WT_active |
| 820 | chr1 | 202193901 | 202319781 | ENSG00000133 | LGR6 | 59352 | 46.0451838 | 0.09481887 | 0.03812427 | 2.48710023 | 0.01287891 | 0.08846505 | WT_active |
| 821 | chr4 | 110045846 | 110199199 | ENSG00000170 | ELOVL6 | 79071 | 37.5008462 | 0.09486814 | 0.03750172 | 2.52970059 | 0.01141599 | 0.08091932 | WT_active |
| 822 | chr12 | 43835967 | 44389762 | ENSG00000139 | TMEM117 | 84216 | 142.495824 | 0.09488081 | 0.0233886 | 4.05671207 | 4.98E-05 | 0.00079464 | WT_active |
| 823 | chr18 | 69401055 | 69849087 | ENSG00000206 | DOK6 | 220164 | 104.362304 | 0.09489343 | 0.02621644 | 3.61961475 | 0.00029504 | 0.00408245 | WT_active |
| 824 | chr6 | 135851696 | 136195574 | ENSG00000171 | PDE7B | 27115 | 91.4547936 | 0.09497969 | 0.02682496 | 3.54072039 | 0.00039904 | 0.0053145 | WT_active |
| 825 | chr11 | 30830369 | 31369810 | ENSG00000170 | DCDC1 | 341019 | 134.958589 | 0.09529083 | 0.02543085 | 3.74705663 | 0.00017892 | 0.00258486 | WT_active |
| 826 | chr2 | 152335237 | 152649834 | ENSG00000157 | FMNL2 | 114793 | 82.9791905 | 0.09550435 | 0.02717177 | 3.5148371 | 0.00044002 | 0.00579385 | WT_active |
| 827 | chr15 | 70853239 | 71053657 | ENSG00000137 | LRRC49 | 54839 | 50.1518734 | 0.09591189 | 0.03498775 | 2.7412994 | 0.00611967 | 0.05059256 | WT_active |
| 828 | chr2 | 49918505 | 51032561 | ENSG00000179 | NRXN1 | 9378 | 266.870616 | 0.09604201 | 0.02188605 | 4.38827524 | 1.14E-05 | 0.00019992 | WT_active |
| 829 | chr8 | 39106990 | 39284911 | ENSG00000197 | ADAM32 | 203102 | 45.0177606 | 0.09618307 | 0.03448013 | 2.78952179 | 0.00527859 | 0.04500696 | WT_active |
| 830 | chr1 | 66533383 | 66748299 | ENSG00000118 | SGIP1 | 84251 | 56.0802135 | 0.09646786 | 0.03309896 | 2.91452841 | 0.00356226 | 0.03341621 | WT_active |
| 831 | chr3 | 185282941 | 185489097 | ENSG00000073 | MAP3K13 | 9175 | 56.1606207 | 0.09651901 | 0.0321867 | 2.99872352 | 0.00271113 | 0.02650875 | WT_active |
| 832 | chr18 | 23689443 | 23955066 | ENSG00000053 | LAMA3 | 3909 | 78.1580427 | 0.09655092 | 0.02700247 | 3.57563224 | 0.00034938 | 0.00474331 | WT_active |
| 833 | chr1 | 174159410 | 174995308 | ENSG00000152 | RABGAP1L | 9910 | 199.517212 | 0.0965895 | 0.02272986 | 4.24945514 | 2.14E-05 | 0.00036607 | WT_active |
| 834 | chr7 | 18086949 | 19002416 | ENSG00000048 | HDAC9 | 9734 | 211.390161 | 0.09683491 | 0.01868854 | 5.18151365 | 2.20E-07 | 4.43E-06 | WT_active |
| 835 | chr1 | 169132531 | 169367967 | ENSG00000143 | NME7 | 29922 | 55.0438717 | 0.09690599 | 0.03391365 | 2.85743291 | 0.00427083 | 0.03848551 | WT_active |
| 836 | chr12 | 72087266 | 72670757 | ENSG00000072 | TRHDE | 29953 | 141.64167 | 0.09694217 | 0.02272776 | 4.2653643 | 2.00E-05 | 0.00034297 | WT_active |
| 837 | chr7 | 114086327 | 114693772 | ENSG00000128 | FOXP2 | 93986 | 130.912227 | 0.09705554 | 0.02728387 | 3.55724929 | 0.00037476 | 0.00503566 | WT_active |
| 838 | chr2 | 151832776 | 152099079 | ENSG00000182 | CACNB4 | 785 | 78.4770455 | 0.09712639 | 0.02699535 | 3.59789287 | 0.00032081 | 0.00439675 | WT_active |
| 839 | chr4 | 152321259 | 152536101 | ENSG00000109 | FBXW7 | 55294 | 45.2716597 | 0.09744779 | 0.0386307 | 2.52254751 | 0.01165082 | 0.08214256 | WT_active |
| 840 | chr3 | 43354859 | 43691594 | ENSG00000160 | ANO10 | 55129 | 93.3839941 | 0.09753119 | 0.02515346 | 3.87744676 | 0.00010556 | 0.00159603 | WT_active |
| 841 | chr12 | 26121991 | 26299290 | ENSG00000123 | SSPN | 8082 | 50.8451737 | 0.09785183 | 0.03416994 | 2.86368165 | 0.00418749 | 0.0379003 | WT_active |
| 842 | chr14 | 93333219 | 93707876 | ENSG00000133 | UNC79 | 57578 | 102.811383 | 0.09789471 | 0.02456043 | 3.98587043 | 6.72E-05 | 0.00104797 | WT_active |
| 843 | chr1 | 231528653 | 231819244 | ENSG00000270 | TSNAX-DISC1 | 100303453 | 88.1958466 | 0.09798045 | 0.0281905 | 3.47565465 | 0.00050961 | 0.00658971 | WT_active |
| 844 | chr6 | 100508194 | 100881372 | ENSG00000112 | ASCC3 | 10973 | 75.6845015 | 0.09801743 | 0.02865382 | 3.42074533 | 0.0006245 | 0.0078092 | WT_active |
| 845 | chr5 | 120464278 | 120687332 | ENSG00000184 | PRR16 | 51334 | 43.6694321 | 0.09802641 | 0.03520947 | 2.784092 | 0.00536778 | 0.04549833 | WT_active |
| 846 | chr7 | 80369575 | 80679277 | ENSG00000135 | CD36 | 948 | 70.7787594 | 0.09810456 | 0.02773552 | 3.53714579 | 0.00040448 | 0.0053792 | WT_active |
| 847 | chr3 | 64515654 | 64688000 | ENSG00000163 | ADAMTS9 | 56999 | 53.6099318 | 0.09813469 | 0.03416762 | 2.87215432 | 0.00407684 | 0.03706666 | WT_active |
| 848 | chr2 | 209424058 | 209734118 | ENSG00000078 | MAP2 | 4133 | 70.1793005 | 0.0983492 | 0.03031176 | 3.24458935 | 0.0011762 | 0.01327198 | WT_active |
| 849 | chr7 | 144451941 | 144836395 | ENSG00000196 | TPK1 | 27010 | 101.93377 | 0.09835848 | 0.02559579 | 3.84276057 | 0.00012166 | 0.00182017 | WT_active |
| 850 | chr6 | 125781161 | 125932030 | ENSG00000111 | NCOA7 | 135112 | 38.8791519 | 0.09847111 | 0.037001 | 2.66130961 | 0.00778373 | 0.06092611 | WT_active |
| 851 | chrX | 106813871 | 107000244 | ENSG00000133 | MORC4 | 79710 | 52.4672922 | 0.09883662 | 0.03310241 | 2.98578354 | 0.00282853 | 0.02743173 | WT_active |
| 852 | chrX | 16946691 | 17153280 | ENSG00000169 | REPS2 | 9185 | 60.4392998 | 0.09890171 | 0.03241858 | 3.05077221 | 0.00228254 | 0.02302982 | WT_active |
| 853 | chr6 | 47477789 | 47627263 | ENSG00000198 | CD2AP | 23607 | 31.0371028 | 0.09891197 | 0.04046733 | 2.44424219 | 0.01451568 | 0.09702691 | WT_active |
| 854 | chr1 | 219685427 | 219958647 | ENSG00000196 | SLC30A10 | 55532 | 77.9303818 | 0.0990167 | 0.03007047 | 3.29282185 | 0.00099187 | 0.01154223 | WT_active |
| 855 | chr5 | 118837322 | 118988545 | ENSG00000169 | DTWD2 | 285605 | 35.9711229 | 0.09915756 | 0.03781431 | 2.62222356 | 0.00873581 | 0.06629101 | WT_active |
| 856 | chr1 | 65248219 | 65415869 | ENSG00000116 | DNAJC6 | 9829 | 46.4580639 | 0.09917052 | 0.03682438 | 2.69306736 | 0.0070798 | 0.05671558 | WT_active |
| 857 | chr6 | 123216339 | 123637093 | ENSG00000186 | TRDN | 10345 | 92.2654426 | 0.09918915 | 0.02993344 | 3.31365748 | 0.00092084 | 0.01081165 | WT_active |
| 858 | chrX | 37349330 | 38687674 | ENSG00000250 | TM4SF2 | NaN | 375.964893 | 0.09928233 | 0.01656408 | 5.99383415 | 2.05E-09 | 4.55E-08 | WT_active |
| 859 | chr7 | 86876906 | 87059699 | ENSG00000164 | KIAA1324L | 222223 | 48.6525108 | 0.09940967 | 0.03980763 | 2.49725134 | 0.01251602 | 0.08646669 | WT_active |
| 860 | chr8 | 118189459 | 118621995 | ENSG00000177 | SAMD12 | 401474 | 130.17315 | 0.09951723 | 0.0211994 | 4.69434138 | 2.67E-06 | 4.96E-05 | WT_active |
| 861 | chr6 | 135283532 | 135497776 | ENSG00000135 | AHI1 | 54806 | 46.4970065 | 0.09962579 | 0.03395271 | 2.9342516 | 0.00334353 | 0.03163191 | WT_active |

| | | | | | | | | | | | | | |
|---|---|---|---|---|---|---|---|---|---|---|---|---|---|
| 862 | chr7 | 78017057 | 79453574 | ENSG00000187 | MAGI2 | 9863 | 359.25209 | 0.09965513 | 0.01657156 | 6.01362258 | 1.81E-09 | 4.03E-08 | WT_active |
| 863 | chr1 | 107140007 | 107483458 | ENSG00000162 | NTNG1 | 22854 | 93.2682287 | 0.09979399 | 0.02734416 | 3.6495534 | 0.0002627 | 0.00367724 | WT_active |
| 864 | chr5 | 81972025 | 82276857 | ENSG00000152 | ATG10 | 83734 | 76.2407758 | 0.09987223 | 0.02817267 | 3.54500434 | 0.00039261 | 0.00524363 | WT_active |
| 865 | chr11 | 63137867 | 63369718 | ENSG00000184 | SLC22A10 | 387775 | 70.7796864 | 0.1000092 | 0.02915541 | 3.43021037 | 0.00060311 | 0.00756805 | WT_active |
| 866 | chr3 | 41246599 | 41962430 | ENSG00000168 | ULK4 | 54986 | 189.236092 | 0.10005421 | 0.01971559 | 5.07487654 | 3.88E-07 | 7.70E-06 | WT_active |
| 867 | chr15 | 76347904 | 76905444 | ENSG00000140 | SCAPER | 49855 | 140.41583 | 0.1001749 | 0.02273761 | 4.40569221 | 1.05E-05 | 0.00018518 | WT_active |
| 868 | chr11 | 86791059 | 86952910 | ENSG00000150 | PRSS23 | 11098 | 46.9019489 | 0.10022803 | 0.03526972 | 2.84175879 | 0.00448654 | 0.03990559 | WT_active |
| 869 | chr5 | 128538013 | 129033642 | ENSG00000113 | SLC27A6 | 28965 | 128.415982 | 0.10028636 | 0.02676893 | 3.7463714 | 0.00017941 | 0.00258934 | WT_active |
| 870 | chr12 | 80936414 | 81261205 | ENSG00000111 | ACSS3 | 79611 | 73.1893207 | 0.10112894 | 0.03098496 | 3.26380794 | 0.00109926 | 0.01251902 | WT_active |
| 871 | chr15 | 67254800 | 67502260 | ENSG00000103 | IQCH | 64799 | 66.8491607 | 0.10122652 | 0.03039079 | 3.33082832 | 0.00086588 | 0.01025822 | WT_active |
| 872 | chr1 | 197268204 | 197478455 | ENSG00000134 | CRB1 | 23418 | 51.1918395 | 0.10141763 | 0.03507426 | 2.89151132 | 0.00383394 | 0.03550319 | WT_active |
| 873 | chr11 | 128965060 | 129279324 | ENSG00000134 | ARHGAP32 | 9743 | 75.4172606 | 0.1015067 | 0.02916834 | 3.48003012 | 0.00050136 | 0.00648884 | WT_active |
| 874 | chr4 | 166733384 | 167234796 | ENSG00000196 | SPOCK3 | 50859 | 108.90144 | 0.10153125 | 0.02852634 | 3.55921019 | 0.00037197 | 0.00500288 | WT_active |
| 875 | chr15 | 48191664 | 48304078 | ENSG00000074 | SLC12A1 | 6557 | 31.4582853 | 0.10163126 | 0.0398463 | 2.55058214 | 0.01075432 | 0.07748913 | WT_active |
| 876 | chr15 | 57375967 | 57550727 | ENSG00000128 | CGNL1 | 84952 | 53.862027 | 0.1021001 | 0.03578355 | 2.85326921 | 0.00432719 | 0.03884175 | WT_active |
| 877 | chr2 | 45388680 | 45612165 | ENSG00000068 | SRBD1 | 55133 | 51.5793323 | 0.10211866 | 0.03458332 | 2.95283015 | 0.00314875 | 0.0300256 | WT_active |
| 878 | chr2 | 191949043 | 192195709 | ENSG00000144 | TMEFF2 | 23671 | 58.9942781 | 0.10217557 | 0.0353982 | 2.8864623 | 0.00389599 | 0.03591589 | WT_active |
| 879 | chr6 | 38168451 | 38640148 | ENSG00000183 | BTBD9 | 114781 | 134.41815 | 0.10225165 | 0.02115768 | 4.83283828 | 1.35E-06 | 2.56E-05 | WT_active |
| 880 | chr6 | 108294894 | 108526796 | ENSG00000135 | LACE1 | 246269 | 58.7770081 | 0.10235593 | 0.02872127 | 3.56376716 | 0.00036557 | 0.00493058 | WT_active |
| 881 | chr7 | 23710167 | 23832513 | ENSG00000196 | STK31 | 56164 | 29.8249533 | 0.10241656 | 0.04044635 | 2.53215852 | 0.01133627 | 0.08053055 | WT_active |
| 882 | chr8 | 9555914 | 9782346 | ENSG00000173 | TNKS | 8658 | 51.2719262 | 0.10262475 | 0.03184929 | 3.22219907 | 0.00127211 | 0.01422963 | WT_active |
| 883 | chr3 | 17157162 | 18444817 | ENSG00000131 | TBC1D5 | 9779 | 328.416023 | 0.1026308 | 0.02032344 | 5.04987253 | 4.42E-07 | 8.73E-06 | WT_active |
| 884 | chr7 | 87503633 | 87713323 | ENSG00000085 | ABCB1 | 5243 | 51.816397 | 0.10267966 | 0.03365066 | 3.05134117 | 0.00227822 | 0.02300544 | WT_active |
| 885 | chr2 | 166195185 | 166375993 | ENSG00000169 | SCN9A | 6335 | 37.7577433 | 0.10270565 | 0.04083167 | 2.51534289 | 0.01189167 | 0.08335151 | WT_active |
| 886 | chr6 | 116254173 | 116444860 | ENSG00000111 | DSE | 29940 | 48.3182968 | 0.10281846 | 0.03362811 | 3.05751543 | 0.0022318 | 0.02266064 | WT_active |
| 887 | chr1 | 203626561 | 203744081 | ENSG00000058 | ATP2B4 | 493 | 32.1415569 | 0.10282828 | 0.04216842 | 2.43851391 | 0.01474779 | 0.0977626 | WT_active |
| 888 | chr5 | 119268692 | 119399688 | ENSG00000145 | TNFAIP8 | 25816 | 34.9803574 | 0.10296058 | 0.04102213 | 2.5098789 | 0.01207726 | 0.08416138 | WT_active |
| 889 | chr7 | 104328656 | 104907232 | ENSG00000187 | LHFPL3 | 375612 | 173.584323 | 0.10302103 | 0.01979344 | 5.20480749 | 1.94E-07 | 3.93E-06 | WT_active |
| 890 | chr1 | 42463330 | 42654664 | ENSG00000186 | CCDC30 | 728621 | 52.7804146 | 0.10324951 | 0.03366608 | 3.06687029 | 0.00216313 | 0.02204097 | WT_active |
| 891 | chr4 | 72280969 | 72568799 | ENSG00000156 | ADAMTS3 | 9508 | 67.5297123 | 0.10329246 | 0.03091441 | 3.34124035 | 0.00083405 | 0.00993509 | WT_active |
| 892 | chr15 | 71822289 | 72118577 | ENSG00000066 | MYO9A | 4649 | 68.6933814 | 0.10335442 | 0.02886618 | 3.58046743 | 0.00034298 | 0.00466078 | WT_active |
| 893 | chr1 | 69568398 | 70151945 | ENSG00000033 | LRRC7 | 57554 | 137.869794 | 0.10336517 | 0.02365395 | 4.36989063 | 1.24E-05 | 0.0002162 | WT_active |
| 894 | chr15 | 59372693 | 59523549 | ENSG00000157 | FAM81A | 145773 | 41.873598 | 0.10341704 | 0.0369533 | 2.79858761 | 0.00513266 | 0.04397074 | WT_active |
| 895 | chr15 | 36579611 | 36810248 | ENSG00000186 | C15orf41 | 84529 | 56.2699419 | 0.10368717 | 0.03178424 | 3.26221937 | 0.00110544 | 0.01255963 | WT_active |
| 896 | chr17 | 28041737 | 28196381 | ENSG00000087 | NLK | 51701 | 34.0589424 | 0.10378905 | 0.04105141 | 2.52827026 | 0.01146261 | 0.08109317 | WT_active |
| 897 | chr2 | 148021011 | 148516971 | ENSG00000204 | MBD5 | 55777 | 110.44934 | 0.10432958 | 0.02548404 | 4.09391884 | 4.24E-05 | 0.00068944 | WT_active |
| 898 | chr3 | 19148454 | 19535646 | ENSG00000183 | KCNH8 | 131096 | 97.4110264 | 0.10440012 | 0.02938815 | 3.55245679 | 0.00038165 | 0.00510924 | WT_active |
| 899 | chr7 | 87627548 | 87832296 | ENSG00000105 | RUNDC3B | 154661 | 45.5846465 | 0.1044858 | 0.03617504 | 2.88833964 | 0.00387281 | 0.03574791 | WT_active |
| 900 | chr4 | 105682627 | 105895986 | ENSG00000138 | INTS12 | 57117 | 50.1962779 | 0.10460554 | 0.03378875 | 3.09586874 | 0.00196237 | 0.02028206 | WT_active |
| 901 | chr14 | 70907405 | 71115382 | ENSG00000100 | PCNX | 22990 | 43.5930292 | 0.10465526 | 0.03643755 | 2.87218156 | 0.00407649 | 0.03706666 | WT_active |
| 902 | chr4 | 78057570 | 78544269 | ENSG00000138 | FRAS1 | 80144 | 145.385974 | 0.10489142 | 0.0221446 | 4.73665883 | 2.17E-06 | 4.06E-05 | WT_active |
| 903 | chr12 | 104802553 | 104958744 | ENSG00000136 | SLC41A2 | 84102 | 35.8352841 | 0.1049534 | 0.0381168 | 2.75346849 | 0.00589674 | 0.04925798 | WT_active |
| 904 | chr5 | 38474963 | 38608354 | ENSG00000113 | LIFR | 3977 | 37.3465224 | 0.10498294 | 0.03702313 | 2.83560377 | 0.00457392 | 0.04050781 | WT_active |
| 905 | chr7 | 17790761 | 17940501 | ENSG00000071 | SNX13 | 23161 | 32.3234524 | 0.10503129 | 0.04163083 | 2.52292102 | 0.01163845 | 0.08209551 | WT_active |
| 906 | chr2 | 157876702 | 158136154 | ENSG00000007 | UPP2 | 151531 | 76.5211095 | 0.10505787 | 0.02999774 | 3.50219257 | 0.00046145 | 0.00602638 | WT_active |
| 907 | chr4 | 169094256 | 169271105 | ENSG00000154 | SH3RF1 | 57630 | 40.8170496 | 0.10520087 | 0.03883164 | 2.70915352 | 0.00674551 | 0.05470011 | WT_active |
| 908 | chrX | 17375420 | 17735994 | ENSG00000188 | NHS | 4810 | 117.821105 | 0.10522145 | 0.02257055 | 4.66189123 | 3.13E-06 | 5.78E-05 | WT_active |
| 909 | chr11 | 111352252 | 111455630 | ENSG00000110 | POU2AF1 | 5450 | 37.8345766 | 0.10526107 | 0.03868609 | 2.72090198 | 0.00651041 | 0.05330326 | WT_active |
| 910 | chr6 | 87408012 | 87511634 | ENSG00000213 | C6orf165 | NaN | 25.2898003 | 0.10529907 | 0.04312094 | 2.44194771 | 0.01460826 | 0.09732991 | WT_active |
| 911 | chr1 | 67092165 | 67231853 | ENSG00000203 | C1orf141 | 400757 | 38.3907294 | 0.10544166 | 0.04059799 | 2.59721403 | 0.00939833 | 0.07023276 | WT_active |
| 912 | chr7 | 132784868 | 133082088 | ENSG00000106 | CHCHD3 | 54927 | 80.7278437 | 0.10559188 | 0.02872489 | 3.67597182 | 0.00023695 | 0.00334274 | WT_active |
| 913 | chrX | 80670854 | 80809688 | ENSG00000165 | BRWD3 | 254065 | 30.2304042 | 0.10563096 | 0.04329463 | 2.43981671 | 0.01469472 | 0.09757217 | WT_active |
| 914 | chr4 | 97184093 | 98143240 | ENSG00000163 | STPG2 | 285555 | 228.695758 | 0.10585863 | 0.0227844 | 4.64610195 | 3.38E-06 | 6.22E-05 | WT_active |
| 915 | chr14 | 59919713 | 60063559 | ENSG00000131 | LRRC9 | 341883 | 35.1896411 | 0.10591129 | 0.04133497 | 2.56226869 | 0.01039908 | 0.0757625 | WT_active |

| | | | | | | | | | | | | | |
|---|---|---|---|---|---|---|---|---|---|---|---|---|---|
| 916 | chr12 | 106357658 | 106510198 | ENSG00000013 | POLR3B | 55703 | 42.5599992 | 0.10595563 | 0.03506902 | 3.02134583 | 0.00251654 | 0.02491994 | WT_active |
| 917 | chr3 | 120908072 | 121424761 | ENSG00000145 | STXBP5L | 9515 | 126.301512 | 0.10598737 | 0.02675571 | 3.96129934 | 7.45E-05 | 0.00115318 | WT_active |
| 918 | chr6 | 88610272 | 88963721 | ENSG00000111 | RNGTT | 8732 | 79.0082897 | 0.10602714 | 0.0265281 | 3.99678545 | 6.42E-05 | 0.00100626 | WT_active |
| 919 | chr11 | 88504576 | 89065945 | ENSG00000168 | GRM5 | 2915 | 137.788964 | 0.10685331 | 0.02505567 | 4.2646355 | 2.00E-05 | 0.00034368 | WT_active |
| 920 | chr3 | 114338094 | 115147271 | ENSG00000181 | ZBTB20 | 26137 | 191.618721 | 0.10698419 | 0.02160449 | 4.95194216 | 7.35E-07 | 1.42E-05 | WT_active |
| 921 | chr4 | 120684919 | 120922870 | ENSG00000138 | PRDM5 | 11107 | 57.8542411 | 0.10707711 | 0.03071952 | 3.4856372 | 0.00049097 | 0.00636579 | WT_active |
| 922 | chr7 | 117710651 | 117874139 | ENSG00000077 | CTTNBP2 | 83992 | 43.7748399 | 0.10727383 | 0.03481105 | 3.08160296 | 0.00205889 | 0.02112819 | WT_active |
| 923 | chr7 | 29806486 | 29917066 | ENSG00000122 | WIPF3 | 644150 | 33.4536497 | 0.10729238 | 0.04009793 | 2.67575858 | 0.00745603 | 0.05913151 | WT_active |
| 924 | chr3 | 43286512 | 43424764 | ENSG00000163 | SNRK | 54861 | 39.9666169 | 0.1074483 | 0.03836792 | 2.80047252 | 0.00510279 | 0.04381891 | WT_active |
| 925 | chr6 | 119177209 | 119349761 | ENSG00000111 | MAN1A1 | 4121 | 43.2452133 | 0.10763875 | 0.03482867 | 3.09052163 | 0.00199805 | 0.0205918 | WT_active |
| 926 | chr7 | 122318425 | 122886759 | ENSG00000081 | CADPS2 | 93664 | 139.796992 | 0.10773795 | 0.02698387 | 3.99267912 | 6.53E-05 | 0.00102163 | WT_active |
| 927 | chr6 | 83552880 | 83709691 | ENSG00000065 | SNAP91 | 9892 | 37.8291603 | 0.10793275 | 0.0404541 | 2.66802991 | 0.00762975 | 0.06017817 | WT_active |
| 928 | chr4 | 26576437 | 26755351 | ENSG00000109 | TBC1D19 | 55296 | 42.0767823 | 0.10826352 | 0.0364325 | 2.97161901 | 0.00296234 | 0.02858242 | WT_active |
| 929 | chr8 | 67173511 | 67343677 | ENSG00000066 | ARFGEF1 | 10565 | 38.5760799 | 0.10831464 | 0.03851012 | 2.81262804 | 0.00491385 | 0.0427309 | WT_active |
| 930 | chr15 | 77100656 | 77420144 | ENSG00000173 | PEAK1 | 79834 | 79.409316 | 0.10873906 | 0.02707241 | 4.01660016 | 5.90E-05 | 0.00093549 | WT_active |
| 931 | chr2 | 227979950 | 228181645 | ENSG00000153 | SPHKAP | 80309 | 53.3075053 | 0.10873994 | 0.03430547 | 3.16975493 | 0.00152568 | 0.01658913 | WT_active |
| 932 | chr5 | 145471799 | 145835369 | ENSG00000186 | PRELID2 | 153768 | 101.437823 | 0.10891529 | 0.0267104 | 4.07763645 | 4.55E-05 | 0.00073538 | WT_active |
| 933 | chr2 | 172735274 | 173052893 | ENSG00000091 | RAPGEF4 | 11069 | 95.2371534 | 0.10894884 | 0.02674626 | 4.07342295 | 4.63E-05 | 0.00074714 | WT_active |
| 934 | chr1 | 161766294 | 161964070 | ENSG00000118 | ATF6 | 22926 | 48.4636058 | 0.10898478 | 0.03231648 | 3.37242109 | 0.0007451 | 0.00897487 | WT_active |
| 935 | chr8 | 79918717 | 80030289 | ENSG00000147 | MRPS28 | 28957 | 24.8593152 | 0.10909793 | 0.04489311 | 2.43017108 | 0.0150917 | 0.09913079 | WT_active |
| 936 | chr4 | 171812254 | 173041559 | ENSG00000174 | GALNTL6 | 442117 | 294.978897 | 0.10913179 | 0.02432536 | 4.48633894 | 7.25E-06 | 0.00012961 | WT_active |
| 937 | chr7 | 22889371 | 23014130 | ENSG00000122 | FAM126A | 84668 | 27.4630384 | 0.10918789 | 0.04118327 | 2.65126805 | 0.00801902 | 0.06236148 | WT_active |
| 938 | chr7 | 24573268 | 24694193 | ENSG00000105 | MPP6 | 51678 | 26.5600464 | 0.10918974 | 0.04469155 | 2.44318547 | 0.01455825 | 0.09717634 | WT_active |
| 939 | chr16 | 9753404 | 10182754 | ENSG00000183 | GRIN2A | 2903 | 134.701488 | 0.10947607 | 0.0228777 | 4.78527437 | 1.71E-06 | 3.21E-05 | WT_active |
| 940 | chr16 | 61647242 | 62037035 | ENSG00000150 | CDH8 | 1006 | 93.6014945 | 0.10951886 | 0.02778011 | 3.94234785 | 8.07E-05 | 0.00124025 | WT_active |
| 941 | chr2 | 227164565 | 227314792 | ENSG00000169 | COL4A3 | 1285 | 39.342811 | 0.10952803 | 0.04014752 | 2.72813922 | 0.00636927 | 0.05229622 | WT_active |
| 942 | chr7 | 91692008 | 91880720 | ENSG00000127 | MTERF | 7978 | 50.7282638 | 0.10954615 | 0.03533569 | 3.10015587 | 0.00193419 | 0.0200498 | WT_active |
| 943 | chr3 | 179794958 | 180037053 | ENSG00000114 | PEX5L | 51555 | 65.3358468 | 0.10961397 | 0.03349085 | 3.27295267 | 0.0010643 | 0.01217867 | WT_active |
| 944 | chr3 | 188947214 | 189325304 | ENSG00000188 | TPRG1 | 285386 | 111.487444 | 0.10970243 | 0.02481154 | 4.42142698 | 9.81E-06 | 0.00017304 | WT_active |
| 945 | chr6 | 97142161 | 97283217 | ENSG00000146 | MMS22L | 253714 | 30.6577203 | 0.10970869 | 0.04182204 | 2.62322651 | 0.00871013 | 0.06616582 | WT_active |
| 946 | chr14 | 80476983 | 80959517 | ENSG00000100 | CEP128 | 145508 | 115.338143 | 0.10997491 | 0.02504162 | 4.39168544 | 1.12E-05 | 0.00019704 | WT_active |
| 947 | chr4 | 175632934 | 176002664 | ENSG00000150 | GPM6A | 2823 | 81.6254407 | 0.11008857 | 0.03526223 | 3.12199688 | 0.00179629 | 0.01894578 | WT_active |
| 948 | chr7 | 14145049 | 14974777 | ENSG00000136 | DGKB | 1607 | 181.319665 | 0.11027985 | 0.02495925 | 4.41839519 | 9.94E-06 | 0.00017527 | WT_active |
| 949 | chr7 | 81946444 | 82443798 | ENSG00000153 | CACNA2D1 | 781 | 102.418036 | 0.11069837 | 0.02648502 | 4.17965901 | 2.92E-05 | 0.00048719 | WT_active |
| 950 | chr22 | 45671798 | 45845307 | ENSG00000130 | ATXN10 | 25814 | 41.0914749 | 0.11080778 | 0.03639927 | 3.04423124 | 0.00233276 | 0.02338922 | WT_active |
| 951 | chr15 | 56244009 | 56445997 | ENSG00000151 | TEX9 | 374618 | 48.5245083 | 0.1108233 | 0.03480158 | 3.18443278 | 0.00145038 | 0.01586614 | WT_active |
| 952 | chr6 | 72621792 | 73198851 | ENSG00000185 | KCNQ5 | 56479 | 136.962197 | 0.11105392 | 0.0220979 | 5.02554163 | 5.02E-07 | 9.87E-06 | WT_active |
| 953 | chr18 | 34976928 | 35143470 | ENSG00000166 | MAPRE2 | 10982 | 46.0387126 | 0.11105748 | 0.03567062 | 3.11341638 | 0.00184935 | 0.01937138 | WT_active |
| 954 | chr6 | 133240598 | 133532120 | ENSG00000112 | EYA4 | 2070 | 74.1647135 | 0.11116796 | 0.02965492 | 3.74871849 | 0.00017774 | 0.00257037 | WT_active |
| 955 | chr7 | 83955777 | 84492724 | ENSG00000075 | SEMA3A | 10371 | 102.176789 | 0.11118145 | 0.02684532 | 4.14155861 | 3.45E-05 | 0.0005684 | WT_active |
| 956 | chr6 | 142751467 | 142945201 | ENSG00000010 | HIVEP2 | 3097 | 47.2809728 | 0.11121131 | 0.03281456 | 3.38908378 | 0.00070127 | 0.00861231 | WT_active |
| 957 | chr2 | 161424332 | 161985282 | ENSG00000144 | SLC4A10 | 57282 | 131.034512 | 0.11122482 | 0.02726294 | 4.07970815 | 4.51E-05 | 0.00072971 | WT_active |
| 958 | chr2 | 184598366 | 184939492 | ENSG00000170 | ZNF804A | 91752 | 70.6630618 | 0.11151467 | 0.03429882 | 3.25126815 | 0.00114891 | 0.01302284 | WT_active |
| 959 | chr2 | 225400648 | 225654018 | ENSG00000144 | NYAP2 | 57624 | 71.4691509 | 0.11166232 | 0.03281992 | 3.40227253 | 0.00066828 | 0.00826351 | WT_active |
| 960 | chr4 | 35948221 | 36244509 | ENSG00000047 | ARAP2 | 116984 | 69.19696 | 0.11167125 | 0.03142161 | 3.55396278 | 0.00037947 | 0.00508953 | WT_active |
| 961 | chr2 | 204545793 | 205620162 | ENSG00000116 | PARD3B | 117583 | 282.100141 | 0.11171276 | 0.02057553 | 5.42939863 | 5.65E-08 | 1.18E-06 | WT_active |
| 962 | chrX | 84058346 | 84187907 | ENSG00000072 | RPS6KA6 | 27330 | 29.1611988 | 0.11173992 | 0.04579451 | 2.44002885 | 0.01468609 | 0.09757217 | WT_active |
| 963 | chr2 | 199269500 | 199471266 | ENSG00000119 | SATB2 | 23314 | 48.6370232 | 0.11202023 | 0.03390412 | 3.30402973 | 0.00095306 | 0.01111976 | WT_active |
| 964 | chr11 | 31509750 | 31783998 | ENSG00000109 | ELP4 | 26610 | 62.3732194 | 0.11223372 | 0.03180891 | 3.52837318 | 0.00041812 | 0.0055358 | WT_active |
| 965 | chr7 | 93424487 | 93574730 | ENSG00000004 | CALCR | 799 | 36.2316684 | 0.1122459 | 0.04408322 | 2.54622718 | 0.01088943 | 0.07811134 | WT_active |
| 966 | chr14 | 96797304 | 96931722 | ENSG00000100 | VRK1 | 7443 | 36.2008933 | 0.11244476 | 0.03808016 | 2.95284404 | 0.00314861 | 0.0300256 | WT_active |
| 967 | chr1 | 180269653 | 180502954 | ENSG00000230 | LHX4-AS1 | 84320 | 55.106771 | 0.11245089 | 0.03093092 | 3.63554977 | 0.00027739 | 0.00386041 | WT_active |
| 968 | chr9 | 131860107 | 132079908 | ENSG00000160 | MED27 | 9442 | 68.5943781 | 0.11254891 | 0.03135695 | 3.58928101 | 0.00033159 | 0.00452307 | WT_active |
| 969 | chr8 | 29067279 | 29263124 | ENSG00000197 | KIF13B | 23303 | 51.0945539 | 0.11257981 | 0.03266419 | 3.44658264 | 0.00056772 | 0.00720551 | WT_active |

| | | | | | | | | | | | | | |
|---|---|---|---|---|---|---|---|---|---|---|---|---|---|
| 970 | chr3 | 33496245 | 33718356 | ENSG00000163 | CLASP2 | 23122 | 49.7636423 | 0.11261231 | 0.03636214 | 3.09696607 | 0.00195512 | 0.02022163 | WT_active |
| 971 | chr6 | 123804141 | 124825657 | ENSG00000188 | NKAIN2 | 154215 | 253.867837 | 0.11285771 | 0.02351249 | 4.79990456 | 1.59E-06 | 2.99E-05 | WT_active |
| 972 | chr5 | 113022099 | 113488830 | ENSG00000171 | MCC | 4163 | 143.379384 | 0.11288246 | 0.02188311 | 5.15842726 | 2.49E-07 | 5.01E-06 | WT_active |
| 973 | chr5 | 126867714 | 127030808 | ENSG00000173 | 37681 | 115123 | 45.3524405 | 0.11303034 | 0.03315772 | 3.40886912 | 0.00065233 | 0.00811498 | WT_active |
| 974 | chr2 | 48962157 | 49154537 | ENSG00000170 | FSHR | 2492 | 57.9418919 | 0.11314021 | 0.03097065 | 3.65314275 | 0.00025905 | 0.00363679 | WT_active |
| 975 | chr5 | 19472951 | 20575873 | ENSG00000145 | CDH18 | 1016 | 240.921338 | 0.11370546 | 0.01942487 | 5.85360133 | 4.81E-09 | 1.05E-07 | WT_active |
| 976 | chr3 | 113286930 | 113441610 | ENSG00000206 | WDR52 | 55779 | 42.2226531 | 0.11386506 | 0.03671968 | 3.10092753 | 0.00192915 | 0.02002488 | WT_active |
| 977 | chr11 | 103109431 | 103479863 | ENSG00000187 | DYNC2H1 | 79659 | 79.2629132 | 0.11411986 | 0.02843091 | 4.01393615 | 5.97E-05 | 0.00094368 | WT_active |
| 978 | chr8 | 103140710 | 103230305 | ENSG00000164 | BAALC | 79870 | 28.8187669 | 0.11413115 | 0.04370134 | 2.61161661 | 0.00901152 | 0.06798962 | WT_active |
| 979 | chr11 | 120336914 | 120489936 | ENSG00000196 | ARHGEF12 | 23365 | 33.8447246 | 0.11422684 | 0.04161743 | 2.74468756 | 0.00605685 | 0.0502172 | WT_active |
| 980 | chr2 | 167954020 | 168248141 | ENSG00000198 | STK39 | 27347 | 82.6525282 | 0.11422812 | 0.02641213 | 4.3248359 | 1.53E-05 | 0.00026421 | WT_active |
| 981 | chr9 | 110873263 | 111038458 | ENSG00000198 | LPAR1 | 1902 | 39.3677815 | 0.11432487 | 0.03790096 | 3.01641116 | 0.00255786 | 0.02525976 | WT_active |
| 982 | chr2 | 97113496 | 97264521 | ENSG00000135 | ANKRD36 | 375248 | 31.3924602 | 0.1145017 | 0.04065458 | 2.81645236 | 0.00485572 | 0.0423532 | WT_active |
| 983 | chr1 | 65792514 | 66374579 | ENSG00000184 | PDE4B | 5142 | 156.615275 | 0.1151038 | 0.02303123 | 4.99772678 | 5.80E-07 | 1.13E-05 | WT_active |
| 984 | chr6 | 128883141 | 129516569 | ENSG00000196 | LAMA2 | 3908 | 147.60614 | 0.11514136 | 0.02452755 | 4.69436909 | 2.67E-06 | 4.96E-05 | WT_active |
| 985 | chr3 | 169083499 | 169663618 | ENSG00000085 | MECOM | 2122 | 149.433882 | 0.11521597 | 0.02401234 | 4.79819843 | 1.60E-06 | 3.01E-05 | WT_active |
| 986 | chr4 | 37244220 | 37449465 | ENSG00000174 | NWD2 | 57495 | 60.6421324 | 0.11534077 | 0.03207709 | 3.59573631 | 0.00032348 | 0.00442492 | WT_active |
| 987 | chr3 | 180602858 | 180871005 | ENSG00000145 | CCDC39 | NaN | 64.0299278 | 0.11534977 | 0.03101844 | 3.7187478 | 0.00020021 | 0.00286376 | WT_active |
| 988 | chr3 | 102099244 | 102479841 | ENSG00000170 | ZPLD1 | 131368 | 94.9391845 | 0.11538274 | 0.02648482 | 4.35656071 | 1.32E-05 | 0.00022923 | WT_active |
| 989 | chr2 | 211375717 | 212538841 | ENSG00000178 | ERBB4 | 2066 | 254.268628 | 0.11547718 | 0.02562912 | 4.50570294 | 6.62E-06 | 0.00011908 | WT_active |
| 990 | chr4 | 65319563 | 65670495 | ENSG00000145 | EPHA5 | 2044 | 66.4313563 | 0.11549431 | 0.03349308 | 3.44830343 | 0.00056412 | 0.00716606 | WT_active |
| 991 | chr7 | 96481626 | 96709891 | ENSG00000127 | SHFM1 | 7979 | 61.4039296 | 0.11561923 | 0.03110971 | 3.71649948 | 0.000202 | 0.00288078 | WT_active |
| 992 | chr12 | 18080869 | 18320107 | ENSG00000111 | RERGL | 79785 | 51.3392777 | 0.11575857 | 0.0356431 | 3.24771358 | 0.00116336 | 0.01314887 | WT_active |
| 993 | chr1 | 60865259 | 61462793 | ENSG00000162 | NFIA | 4774 | 164.751841 | 0.11597315 | 0.02185122 | 5.30739928 | 1.11E-07 | 2.28E-06 | WT_active |
| 994 | chr3 | 131026850 | 131350465 | ENSG00000114 | NEK11 | 79858 | 89.5287284 | 0.11600411 | 0.02742239 | 4.23026971 | 2.33E-05 | 0.00039453 | WT_active |
| 995 | chr9 | 5881596 | 6007901 | ENSG00000183 | KIAA2026 | 158358 | 30.8481671 | 0.11610533 | 0.04254477 | 2.72901492 | 0.00635238 | 0.05221702 | WT_active |
| 996 | chr9 | 3218297 | 3526004 | ENSG00000080 | RFX3 | 5991 | 67.6593317 | 0.11615925 | 0.0331162 | 3.50762616 | 0.00045212 | 0.0059315 | WT_active |
| 997 | chr2 | 36696690 | 36814792 | ENSG00000205 | VIT | 5212 | 38.3551382 | 0.11622137 | 0.03774398 | 3.07920235 | 0.00207556 | 0.02126263 | WT_active |
| 998 | chr8 | 106270178 | 106752694 | ENSG00000164 | OXR1 | 55074 | 118.129882 | 0.11629559 | 0.02727313 | 4.26410813 | 2.01E-05 | 0.00034408 | WT_active |
| 999 | chr2 | 181954241 | 182131398 | ENSG00000150 | PPP1R1C | 151242 | 42.7064677 | 0.11642753 | 0.0372729 | 3.12365074 | 0.00178622 | 0.01886898 | WT_active |
| 1000 | chr6 | 71886703 | 72403143 | ENSG00000079 | RIMS1 | 22999 | 116.88967 | 0.11661302 | 0.02547136 | 4.57820117 | 4.69E-06 | 8.55E-05 | WT_active |
| 1001 | chr3 | 156825481 | 157046129 | ENSG00000197 | LEKR1 | 389170 | 51.9264245 | 0.11670498 | 0.03477837 | 3.35567721 | 0.00079171 | 0.00947292 | WT_active |
| 1002 | chr1 | 51354263 | 51519328 | ENSG00000085 | EPS15 | 2060 | 35.7458778 | 0.11693964 | 0.03875202 | 3.01763965 | 0.00254752 | 0.02517484 | WT_active |
| 1003 | chr4 | 86594315 | 86815171 | ENSG00000163 | PTPN13 | 5783 | 50.571566 | 0.11698537 | 0.03191012 | 3.66609054 | 0.00024629 | 0.00346774 | WT_active |
| 1004 | chr1 | 147541412 | 147626216 | ENSG00000116 | BCL9 | 607 | 26.6836709 | 0.11746503 | 0.04708952 | 2.49450467 | 0.01261331 | 0.0869721 | WT_active |
| 1005 | chr11 | 22625642 | 22813055 | ENSG00000148 | GAS2 | 2620 | 45.413497 | 0.11764476 | 0.03605788 | 3.26266467 | 0.0011037 | 0.0125498 | WT_active |
| 1006 | chr5 | 129904472 | 130186634 | ENSG00000198 | CHSY3 | 337876 | 68.3008259 | 0.1176846 | 0.03463243 | 3.39810401 | 0.00067855 | 0.00837609 | WT_active |
| 1007 | chr2 | 208359714 | 208854503 | ENSG00000144 | PTH2R | 5746 | 126.01534 | 0.11769354 | 0.02634101 | 4.4680731 | 7.89E-06 | 0.00014066 | WT_active |
| 1008 | chr1 | 33323623 | 33431052 | ENSG00000134 | PHC2 | 1912 | 33.1354891 | 0.11771653 | 0.0391449 | 3.00719939 | 0.00263667 | 0.0258961 | WT_active |
| 1009 | chrX | 72578814 | 72714319 | ENSG00000067 | PHKA1 | 5255 | 35.3123905 | 0.11776878 | 0.03963307 | 2.97147746 | 0.00296371 | 0.02858242 | WT_active |
| 1010 | chr3 | 105366909 | 105576900 | ENSG00000170 | ALCAM | 214 | 41.0558796 | 0.1177887 | 0.03893412 | 3.02533369 | 0.00248359 | 0.02466143 | WT_active |
| 1011 | chr2 | 10302889 | 10427617 | ENSG00000115 | HPCAL1 | 3241 | 36.0024155 | 0.1178047 | 0.04346551 | 2.7103032 | 0.00672217 | 0.05457223 | WT_active |
| 1012 | chr1 | 22451851 | 22531157 | ENSG00000184 | ZBTB40 | 9923 | 25.3053675 | 0.11780595 | 0.04697397 | 2.50789855 | 0.01214515 | 0.08447661 | WT_active |
| 1013 | chr3 | 158962928 | 159266307 | ENSG00000214 | IQCJ | 654502 | 86.4827198 | 0.11820501 | 0.0278194 | 4.24901284 | 2.15E-05 | 0.00036636 | WT_active |
| 1014 | chr2 | 161992241 | 162074542 | ENSG00000197 | DPP4 | 1803 | 22.586435 | 0.11821293 | 0.04843131 | 2.4408367 | 0.01465328 | 0.09749468 | WT_active |
| 1015 | chr2 | 132416574 | 132646559 | ENSG00000183 | GPR39 | 2863 | 74.9473898 | 0.11826801 | 0.02802379 | 4.22027242 | 2.44E-05 | 0.00041147 | WT_active |
| 1016 | chr6 | 154010496 | 154246867 | ENSG00000112 | OPRM1 | 4988 | 64.9464466 | 0.11851093 | 0.02895165 | 4.09340847 | 4.25E-05 | 0.00069018 | WT_active |
| 1017 | chr1 | 114854803 | 114995370 | ENSG00000198 | SYCP1 | 6847 | 32.3283898 | 0.11853292 | 0.042071 | 2.81744963 | 0.00484067 | 0.04224745 | WT_active |
| 1018 | chr2 | 169694488 | 169776989 | ENSG00000213 | KLHL23 | 151230 | 21.4342834 | 0.11885935 | 0.04839457 | 2.45604732 | 0.01404747 | 0.09464316 | WT_active |
| 1019 | chr3 | 160755602 | 161078907 | ENSG00000163 | PPM1L | 151742 | 90.9735339 | 0.11887401 | 0.02676009 | 4.44221346 | 8.90E-06 | 0.00015752 | WT_active |
| 1020 | chr4 | 161383897 | 162164035 | ENSG00000168 | FSTL5 | 56884 | 152.57842 | 0.11905803 | 0.0300474 | 3.96234018 | 7.42E-05 | 0.00114939 | WT_active |
| 1021 | chr5 | 66596361 | 67169595 | ENSG00000069 | MAST4 | 375449 | 163.711371 | 0.11916178 | 0.02073891 | 5.74580809 | 9.15E-09 | 1.97E-07 | WT_active |
| 1022 | chr6 | 117907526 | 118317676 | ENSG00000196 | SLC35F1 | 222553 | 118.848649 | 0.1191907 | 0.02595235 | 4.59267484 | 4.38E-06 | 8.00E-05 | WT_active |
| 1023 | chr14 | 70722526 | 70809534 | ENSG00000006 | MAP3K9 | 4293 | 27.5851009 | 0.11938576 | 0.04394069 | 2.71697477 | 0.00658816 | 0.05384428 | WT_active |

| | | | | | | | | | | | | | |
|---|---|---|---|---|---|---|---|---|---|---|---|---|---|
| 1024 | chr2 | 159318979 | 159616692 | ENSG00000123 | BAZ2B | 29994 | 65.5110001 | 0.11958971 | 0.02979688 | 4.01349843 | 5.98E-05 | 0.00094368 | WT_active |
| 1025 | chrX | 69616067 | 70039469 | ENSG00000158 | EDA | 1896 | 120.19061 | 0.11964862 | 0.02414823 | 4.95475631 | 7.24E-07 | 1.41E-05 | WT_active |
| 1026 | chr8 | 116950273 | 117176714 | ENSG00000164 | SLC30A8 | 169026 | 66.9799205 | 0.11969099 | 0.03051889 | 3.9218652 | 8.79E-05 | 0.00134201 | WT_active |
| 1027 | chr3 | 167478684 | 167653983 | ENSG00000174 | WDR49 | NaN | 44.5391514 | 0.11980861 | 0.03909104 | 3.06486151 | 0.00217771 | 0.02217387 | WT_active |
| 1028 | chr1 | 113390749 | 113685923 | ENSG00000081 | MAGI3 | 260425 | 65.2085802 | 0.11989392 | 0.03155022 | 3.80009779 | 0.00014464 | 0.00213014 | WT_active |
| 1029 | chr1 | 159062484 | 159147096 | ENSG00000163 | AIM2 | 9447 | 25.8149426 | 0.11998638 | 0.04522801 | 2.65292173 | 0.00797984 | 0.06225828 | WT_active |
| 1030 | chr1 | 100351734 | 100520277 | ENSG00000079 | CDC14A | 8556 | 44.2547232 | 0.12010009 | 0.03658863 | 3.28244308 | 0.00102912 | 0.01186077 | WT_active |
| 1031 | chr5 | 68215720 | 68301821 | ENSG00000145 | PIK3R1 | 5295 | 23.7977886 | 0.12034667 | 0.04889374 | 2.46139189 | 0.01383991 | 0.09368255 | WT_active |
| 1032 | chrX | 107825755 | 107927193 | ENSG00000080 | MID2 | 11043 | 31.6561312 | 0.12039501 | 0.03982972 | 3.02274342 | 0.00250495 | 0.02483927 | WT_active |
| 1033 | chr1 | 117367449 | 117528872 | ENSG00000198 | MAN1A2 | 10905 | 34.3940857 | 0.12044632 | 0.04188866 | 2.87539235 | 0.00403526 | 0.03684632 | WT_active |
| 1034 | chr3 | 139935185 | 140577397 | ENSG00000158 | CLSTN2 | 64084 | 224.854096 | 0.12053864 | 0.01848844 | 6.51967768 | 7.05E-11 | 1.65E-09 | WT_active |
| 1035 | chr3 | 190514051 | 190659750 | ENSG00000196 | IL1RAP | 3556 | 39.1199095 | 0.12058748 | 0.04021412 | 2.99863545 | 0.00271192 | 0.02650875 | WT_active |
| 1036 | chr1 | 35268967 | 35422058 | ENSG00000146 | ZMYM4 | 9202 | 32.8814399 | 0.1207642 | 0.04179291 | 2.88958614 | 0.00385749 | 0.0356293 | WT_active |
| 1037 | chr2 | 233475520 | 233566782 | ENSG00000085 | USP40 | 55230 | 21.0391938 | 0.12082301 | 0.04629103 | 2.6100739 | 0.00905227 | 0.06815435 | WT_active |
| 1038 | chr6 | 83859643 | 83967424 | ENSG00000065 | CYB5R4 | 51167 | 27.0962382 | 0.12083373 | 0.04682913 | 2.58031142 | 0.00987113 | 0.0727617 | WT_active |
| 1039 | chr4 | 169393275 | 169612629 | ENSG00000137 | NEK1 | 4750 | 48.2921463 | 0.121037 | 0.03394516 | 3.56566291 | 0.00036294 | 0.00489966 | WT_active |
| 1040 | chr5 | 119452443 | 119637199 | ENSG00000133 | HSD17B4 | 3295 | 47.0376708 | 0.12107399 | 0.03911942 | 3.0949839 | 0.00196824 | 0.02031355 | WT_active |
| 1041 | chr6 | 116100849 | 116249497 | ENSG00000178 | NT5DC1 | 221294 | 33.9020405 | 0.12117128 | 0.04003128 | 3.02691527 | 0.00247063 | 0.02455466 | WT_active |
| 1042 | chr2 | 178525989 | 178830802 | ENSG00000155 | TTN | 7273 | 85.7006605 | 0.12142066 | 0.02729322 | 4.44874864 | 8.64E-06 | 0.00015299 | WT_active |
| 1043 | chr6 | 56056590 | 56394094 | ENSG00000124 | COL21A1 | 81578 | 82.0274503 | 0.1215627 | 0.02530896 | 4.80314962 | 1.56E-06 | 2.95E-05 | WT_active |
| 1044 | chr1 | 241590102 | 241677376 | ENSG00000054 | OPN3 | 23596 | 22.608488 | 0.12160488 | 0.04982941 | 2.44042393 | 0.01467004 | 0.09756115 | WT_active |
| 1045 | chr15 | 38252326 | 38357249 | ENSG00000166 | SPRED1 | 161742 | 20.4503565 | 0.12165783 | 0.04841217 | 2.51295948 | 0.01197231 | 0.08383523 | WT_active |
| 1046 | chr4 | 80335720 | 80963756 | ENSG00000197 | C4orf22 | 255119 | 149.812157 | 0.12172976 | 0.02493247 | 4.88237907 | 1.05E-06 | 2.02E-05 | WT_active |
| 1047 | chrX | 38006582 | 38128819 | ENSG00000147 | SYTL5 | 94122 | 33.1948905 | 0.1218128 | 0.04230439 | 2.87943664 | 0.00398386 | 0.03649259 | WT_active |
| 1048 | chr2 | 214411065 | 214578959 | ENSG00000174 | VWC2L | 402117 | 43.3638955 | 0.12200059 | 0.03909099 | 3.12093862 | 0.00180276 | 0.0190001 | WT_active |
| 1049 | chr1 | 67166400 | 67259979 | ENSG00000162 | IL23R | 149233 | 26.5702026 | 0.12218343 | 0.04515986 | 2.70557605 | 0.0068186 | 0.05513463 | WT_active |
| 1050 | chr5 | 116084991 | 116412762 | ENSG00000145 | COMMD10 | 51397 | 86.0118442 | 0.12223964 | 0.02977234 | 4.10581287 | 4.03E-05 | 0.00065787 | WT_active |
| 1051 | chr2 | 44361950 | 44772592 | ENSG00000143 | CAMKMT | 79823 | 101.980764 | 0.12224155 | 0.02438115 | 5.01377321 | 5.34E-07 | 1.05E-05 | WT_active |
| 1052 | chr3 | 142449235 | 142578826 | ENSG00000175 | ATR | 545 | 27.8652149 | 0.12226907 | 0.04351849 | 2.80958926 | 0.00496048 | 0.04298086 | WT_active |
| 1053 | chr7 | 117465784 | 117715971 | ENSG00000001 | CFTR | 1080 | 63.9324057 | 0.12263894 | 0.03212192 | 3.81792047 | 0.00013458 | 0.00199835 | WT_active |
| 1054 | chr2 | 66433452 | 66573869 | ENSG00000143 | MEIS1 | 4211 | 36.4873219 | 0.12283006 | 0.04045767 | 3.03601411 | 0.00239728 | 0.02395233 | WT_active |
| 1055 | chr4 | 148078762 | 148444698 | ENSG00000151 | NR3C2 | 4306 | 94.9770601 | 0.12304519 | 0.02703704 | 4.55098684 | 5.34E-06 | 9.70E-05 | WT_active |
| 1056 | chr7 | 87401697 | 87480435 | ENSG00000005 | ABCB4 | 5244 | 21.5833599 | 0.12325583 | 0.05078992 | 2.42677759 | 0.01523359 | 0.09965423 | WT_active |
| 1057 | chr2 | 97756333 | 97995891 | ENSG00000075 | TMEM131 | 23505 | 62.1587625 | 0.12342619 | 0.03132377 | 3.94033628 | 8.14E-05 | 0.00124804 | WT_active |
| 1058 | chr6 | 131135666 | 131283535 | ENSG00000118 | AKAP7 | 9465 | 38.6598358 | 0.12360618 | 0.03821311 | 3.2346534 | 0.0012179 | 0.01370785 | WT_active |
| 1059 | chr5 | 33987174 | 34124528 | ENSG00000273 | C1QTNF3-AMA | 100534612 | 32.2765037 | 0.12372229 | 0.03786052 | 3.26784413 | 0.0010837 | 0.01238098 | WT_active |
| 1060 | chr5 | 122311354 | 122464219 | ENSG00000064 | SNCAIP | 9627 | 44.8619928 | 0.12373017 | 0.03935426 | 3.14400979 | 0.0016665 | 0.01785186 | WT_active |
| 1061 | chr1 | 74198235 | 74543982 | ENSG00000259 | FPGT-TNNI3K | 100526835 | 89.8674734 | 0.124164 | 0.0298704 | 4.15675701 | 3.23E-05 | 0.00053557 | WT_active |
| 1062 | chr8 | 120445400 | 120542133 | ENSG00000172 | MTBP | 27085 | 21.1167949 | 0.12425094 | 0.04874125 | 2.54919451 | 0.01079721 | 0.07763 | WT_active |
| 1063 | chr5 | 83940554 | 84384793 | ENSG00000164 | EDIL3 | 10085 | 96.8092113 | 0.12430761 | 0.02989242 | 4.15849926 | 3.20E-05 | 0.00053273 | WT_active |
| 1064 | chr6 | 83067666 | 83171350 | ENSG00000083 | DOPEY1 | 23033 | 23.8102146 | 0.12436363 | 0.04956993 | 2.50885237 | 0.01211241 | 0.08436557 | WT_active |
| 1065 | chr7 | 110663051 | 111562517 | ENSG00000184 | IMMP2L | 83943 | 213.542033 | 0.12439145 | 0.02059809 | 6.03897965 | 1.55E-09 | 3.46E-08 | WT_active |
| 1066 | chr12 | 28133249 | 28581511 | ENSG00000123 | CCDC91 | 55297 | 101.004131 | 0.12461527 | 0.02561752 | 4.86445391 | 1.15E-06 | 2.20E-05 | WT_active |
| 1067 | chr8 | 119730775 | 119832863 | ENSG00000064 | TAF2 | 6873 | 22.8067968 | 0.12466473 | 0.04937032 | 2.52509477 | 0.01156671 | 0.08170937 | WT_active |
| 1068 | chr7 | 134986508 | 135147963 | ENSG00000146 | AGBL3 | 340351 | 42.4255315 | 0.12478098 | 0.03948886 | 3.15990309 | 0.00157822 | 0.01707031 | WT_active |
| 1069 | chr2 | 174559572 | 174682916 | ENSG00000115 | WIPF1 | 7456 | 36.6267667 | 0.12480329 | 0.04010697 | 3.11176071 | 0.00185975 | 0.01943036 | WT_active |
| 1070 | chr12 | 43353866 | 43551921 | ENSG00000173 | ADAMTS20 | 80070 | 44.8990555 | 0.12483226 | 0.03957907 | 3.15399709 | 0.00161051 | 0.01731565 | WT_active |
| 1071 | chr5 | 38937919 | 39074408 | ENSG00000164 | RICTOR | 253260 | 27.8104167 | 0.12488672 | 0.0449878 | 2.77601277 | 0.005503 | 0.04650781 | WT_active |
| 1072 | chr15 | 68054179 | 68198603 | ENSG00000033 | PIAS1 | 8554 | 29.727978 | 0.12518015 | 0.04263515 | 2.93607842 | 0.0033239 | 0.03149801 | WT_active |
| 1073 | chr18 | 57435685 | 57491297 | ENSG00000119 | ONECUT2 | 9480 | 17.5874127 | 0.12542768 | 0.05168128 | 2.42694587 | 0.01522653 | 0.09965423 | WT_active |
| 1074 | chr5 | 94618347 | 94739436 | ENSG00000133 | ANKRD32 | 84250 | 26.9285202 | 0.12547733 | 0.04921332 | 2.54966196 | 0.01078274 | 0.07761636 | WT_active |
| 1075 | chr1 | 198638671 | 198757283 | ENSG00000081 | PTPRC | 5788 | 24.2996078 | 0.12552371 | 0.04633102 | 2.7092803 | 0.00674293 | 0.05470011 | WT_active |
| 1076 | chr2 | 201288271 | 201357398 | ENSG00000155 | ALS2CR12 | 130540 | 20.4432289 | 0.12558102 | 0.04736483 | 2.65135589 | 0.00801693 | 0.06236148 | WT_active |
| 1077 | chr1 | 197504748 | 197775696 | ENSG00000213 | DENND1B | 163486 | 53.7797164 | 0.12568661 | 0.03297601 | 3.81145543 | 0.00013815 | 0.00204293 | WT_active |

| | | | | | | | | | | | | | |
|---|---|---|---|---|---|---|---|---|---|---|---|---|---|
| 1078 | chr2 | 40097270 | 40611053 | ENSG00000183 | SLC8A1 | 6546 | 130.153569 | 0.12583409 | 0.02274605 | 5.53212839 | 3.16E-08 | 6.73E-07 | WT_active |
| 1079 | chr4 | 105708778 | 105847728 | ENSG00000138 | GSTCD | 79807 | 30.7439264 | 0.12598018 | 0.04372773 | 2.88101338 | 0.00396399 | 0.03637987 | WT_active |
| 1080 | chr1 | 74235401 | 74544428 | ENSG00000116 | TNNI3K | 51086 | 81.6523393 | 0.12620189 | 0.03008396 | 4.19498894 | 2.73E-05 | 0.00045749 | WT_active |
| 1081 | chr7 | 11370357 | 11832198 | ENSG00000005 | THSD7A | 221981 | 109.845166 | 0.12640043 | 0.03042352 | 4.15469402 | 3.26E-05 | 0.0005398 | WT_active |
| 1082 | chrX | 38149336 | 38220924 | ENSG00000101 | SRPX | 8406 | 21.5486839 | 0.12641266 | 0.05047428 | 2.50449662 | 0.01226258 | 0.0850827 | WT_active |
| 1083 | chr3 | 190322541 | 190412143 | ENSG00000113 | CLDN16 | 10686 | 22.912669 | 0.12658236 | 0.05155869 | 2.45511214 | 0.01408406 | 0.09471434 | WT_active |
| 1084 | chr4 | 153466346 | 153636711 | ENSG00000121 | KIAA0922 | 23240 | 46.8138235 | 0.12660827 | 0.03669965 | 3.4498496 | 0.0005609 | 0.00713143 | WT_active |
| 1085 | chr6 | 106184476 | 106325820 | ENSG00000057 | ATG5 | 9474 | 31.5615099 | 0.12664505 | 0.04053289 | 3.12450081 | 0.00178107 | 0.01885424 | WT_active |
| 1086 | chr17 | 37686432 | 37745247 | ENSG00000275 | HNF1B | 6928 | 19.5068439 | 0.12679455 | 0.05069813 | 2.50097097 | 0.01238533 | 0.08568701 | WT_active |
| 1087 | chr7 | 116672390 | 116798386 | ENSG00000105 | MET | 4233 | 31.5146044 | 0.12736785 | 0.03854192 | 3.30465761 | 0.00095092 | 0.01111957 | WT_active |
| 1088 | chr5 | 64768930 | 65018763 | ENSG00000153 | CWC27 | 10283 | 57.1421668 | 0.12738835 | 0.03287696 | 3.8747 | 0.00010676 | 0.00161174 | WT_active |
| 1089 | chr4 | 157204182 | 157366075 | ENSG00000120 | GRIA2 | 2891 | 34.7446377 | 0.12746582 | 0.04433716 | 2.8749205 | 0.00404129 | 0.03685475 | WT_active |
| 1090 | chr3 | 66378797 | 66501263 | ENSG00000144 | LRIG1 | 26018 | 37.4904237 | 0.12773883 | 0.03810196 | 3.35255298 | 0.0008007 | 0.00956461 | WT_active |
| 1091 | chr9 | 95875701 | 96014571 | ENSG00000182 | ERCC6L2 | 375748 | 33.077378 | 0.12814484 | 0.03962344 | 3.23406682 | 0.00122041 | 0.01372532 | WT_active |
| 1092 | chr8 | 17922840 | 18029944 | ENSG00000078 | PCM1 | 5108 | 23.4103452 | 0.12817129 | 0.04832432 | 2.65231463 | 0.0079942 | 0.06230291 | WT_active |
| 1093 | chr7 | 12570577 | 12660179 | ENSG00000006 | SCIN | 85477 | 23.5379372 | 0.12824862 | 0.04976063 | 2.57731118 | 0.00995723 | 0.07313464 | WT_active |
| 1094 | chr10 | 49455368 | 49539538 | ENSG00000225 | ERCC6 | 2074 | 23.7716563 | 0.12829966 | 0.04733556 | 2.71042869 | 0.00671963 | 0.05457223 | WT_active |
| 1095 | chr9 | 18473894 | 18910950 | ENSG00000178 | ADAMTSL1 | 92949 | 116.820303 | 0.12830484 | 0.02338748 | 5.48604842 | 4.11E-08 | 8.66E-07 | WT_active |
| 1096 | chr4 | 100395341 | 100880126 | ENSG00000164 | EMCN | 51705 | 118.511204 | 0.12841324 | 0.02519872 | 5.09602192 | 3.47E-07 | 6.93E-06 | WT_active |
| 1097 | chr2 | 189879609 | 190203484 | ENSG00000187 | C2orf88 | 84281 | 79.3486085 | 0.12842012 | 0.02877491 | 4.46291959 | 8.09E-06 | 0.00014391 | WT_active |
| 1098 | chr8 | 103500748 | 104256094 | ENSG00000176 | RIMS2 | 9699 | 164.170262 | 0.12847303 | 0.02394506 | 5.36532401 | 8.08E-08 | 1.67E-06 | WT_active |
| 1099 | chr2 | 69013178 | 69249327 | ENSG00000169 | ANTXR1 | 84168 | 66.8927947 | 0.12865757 | 0.02858446 | 4.50096251 | 6.76E-06 | 0.00012146 | WT_active |
| 1100 | chr6 | 57086844 | 57170307 | ENSG00000112 | ZNF451 | 26036 | 17.9406862 | 0.12867693 | 0.05039275 | 2.55348105 | 0.01066521 | 0.07715553 | WT_active |
| 1101 | chr1 | 179954738 | 180114880 | ENSG00000135 | CEP350 | 9857 | 32.0528855 | 0.12918296 | 0.04319981 | 2.99035953 | 0.00278649 | 0.02712738 | WT_active |
| 1102 | chr7 | 7356203 | 7535853 | ENSG00000215 | COL28A1 | 340267 | 47.8960802 | 0.12944187 | 0.03635126 | 3.56086351 | 0.00036964 | 0.00497612 | WT_active |
| 1103 | chr1 | 93179919 | 93278730 | ENSG00000122 | CCDC18 | 343099 | 21.1560719 | 0.129939 | 0.05000309 | 2.59861954 | 0.00935994 | 0.06999569 | WT_active |
| 1104 | chr15 | 48823735 | 48963444 | ENSG00000185 | SHC4 | 399694 | 39.1433877 | 0.13030688 | 0.03716456 | 3.50621348 | 0.00045453 | 0.00595224 | WT_active |
| 1105 | chr3 | 111292719 | 111665750 | ENSG00000153 | CD96 | 10225 | 102.716577 | 0.13060448 | 0.02402359 | 5.43651082 | 5.43E-08 | 1.14E-06 | WT_active |
| 1106 | chr1 | 71395940 | 72282734 | ENSG00000172 | NEGR1 | 257194 | 198.461006 | 0.13074914 | 0.02597699 | 5.03326827 | 4.82E-07 | 9.50E-06 | WT_active |
| 1107 | chr1 | 190097662 | 190478404 | ENSG00000162 | BRINP3 | 339479 | 72.1272433 | 0.13075861 | 0.03620993 | 3.61112545 | 0.00030487 | 0.00420635 | WT_active |
| 1108 | chrX | 55089045 | 55288785 | ENSG00000158 | PAGE5 | 90737 | 56.4637855 | 0.13078648 | 0.0316191 | 4.13631276 | 3.53E-05 | 0.00057956 | WT_active |
| 1109 | chr4 | 168863770 | 169010275 | ENSG00000145 | CBR4 | 84869 | 34.3571178 | 0.13081464 | 0.04056043 | 3.22517848 | 0.00125894 | 0.01411463 | WT_active |
| 1110 | chr7 | 16087527 | 16421322 | ENSG00000214 | ISPD | 729920 | 78.8060766 | 0.13085589 | 0.03010196 | 4.34708907 | 1.38E-05 | 0.00023907 | WT_active |
| 1111 | chr2 | 188291669 | 188595931 | ENSG00000144 | GULP1 | 51454 | 67.3134401 | 0.13109358 | 0.02969135 | 4.41521147 | 1.01E-05 | 0.00017765 | WT_active |
| 1112 | chrX | 24693919 | 24996986 | ENSG00000101 | POLA1 | 5422 | 80.4620465 | 0.13118519 | 0.02597017 | 5.05138004 | 4.39E-07 | 8.68E-06 | WT_active |
| 1113 | chr5 | 58582221 | 58859394 | ENSG00000152 | RAB3C | 115827 | 79.7158208 | 0.13119638 | 0.02675328 | 4.90393546 | 9.39E-07 | 1.81E-05 | WT_active |
| 1114 | chr3 | 18345387 | 18445588 | ENSG00000182 | SATB1 | 6304 | 22.4104417 | 0.13131391 | 0.05235115 | 2.50832912 | 0.01213036 | 0.08444981 | WT_active |
| 1115 | chr9 | 32455705 | 32526324 | ENSG00000107 | DDX58 | 23586 | 18.415066 | 0.13181451 | 0.05299825 | 2.48714863 | 0.01287716 | 0.08846505 | WT_active |
| 1116 | chr2 | 237485428 | 237555318 | ENSG00000115 | MLPH | 79083 | 22.9985969 | 0.13208385 | 0.05174218 | 2.55273063 | 0.01068821 | 0.07728317 | WT_active |
| 1117 | chr15 | 79311062 | 79427432 | ENSG00000166 | TMED3 | 23423 | 39.1254385 | 0.13211883 | 0.03954351 | 3.34110047 | 0.00083447 | 0.00993509 | WT_active |
| 1118 | chr3 | 21415071 | 22373321 | ENSG00000151 | ZNF385D | 79750 | 235.416857 | 0.13213844 | 0.02438837 | 5.41809199 | 6.02E-08 | 1.25E-06 | WT_active |
| 1119 | chr4 | 168216293 | 168318807 | ENSG00000137 | DDX60 | 55601 | 24.9829473 | 0.13214401 | 0.05033501 | 2.62529013 | 0.00865752 | 0.06587022 | WT_active |
| 1120 | chr11 | 110577042 | 110713189 | ENSG00000137 | ARHGAP20 | 57569 | 34.3346223 | 0.13220324 | 0.04246483 | 3.11324122 | 0.00185045 | 0.01937138 | WT_active |
| 1121 | chr6 | 96924620 | 97140754 | ENSG00000186 | KLHL32 | 114792 | 62.1751646 | 0.13224057 | 0.03162374 | 4.18168603 | 2.89E-05 | 0.00048342 | WT_active |
| 1122 | chr1 | 97077743 | 97921049 | ENSG00000188 | DPYD | 1806 | 184.842081 | 0.13231256 | 0.01959851 | 6.75115307 | 1.47E-11 | 3.46E-10 | WT_active |
| 1123 | chr7 | 152025674 | 152122347 | ENSG00000178 | GALNT11 | 63917 | 28.0280055 | 0.13263642 | 0.04388441 | 3.02240385 | 0.00250776 | 0.02485007 | WT_active |
| 1124 | chr1 | 62454728 | 62688368 | ENSG00000116 | DOCK7 | 85440 | 47.6134772 | 0.13271684 | 0.03481522 | 3.8120351 | 0.00013783 | 0.00204024 | WT_active |
| 1125 | chr3 | 89107524 | 89482134 | ENSG00000044 | EPHA3 | 2042 | 77.1388264 | 0.13293041 | 0.02924753 | 4.54501298 | 5.49E-06 | 9.96E-05 | WT_active |
| 1126 | chrX | 110194186 | 110440233 | ENSG00000101 | AMMECR1 | 9949 | 69.2652429 | 0.13294979 | 0.02768783 | 4.80174177 | 1.57E-06 | 2.97E-05 | WT_active |
| 1127 | chr9 | 121074863 | 121177610 | ENSG00000119 | CNTRL | 11064 | 26.9193202 | 0.13311476 | 0.04433195 | 3.00268232 | 0.00267612 | 0.02622994 | WT_active |
| 1128 | chr1 | 167809388 | 167914215 | ENSG00000143 | ADCY10 | 55811 | 30.256642 | 0.13329255 | 0.04067484 | 3.27702728 | 0.00104906 | 0.01201628 | WT_active |
| 1129 | chr8 | 97775057 | 97853013 | ENSG00000104 | LAPTM4B | 55353 | 21.6092561 | 0.13330198 | 0.04814507 | 2.76875685 | 0.00562706 | 0.04736192 | WT_active |
| 1130 | chr2 | 190408355 | 190509205 | ENSG00000151 | MFSD6 | 54842 | 28.7555083 | 0.13335568 | 0.043271 | 3.08187208 | 0.00205703 | 0.02112639 | WT_active |
| 1131 | chr4 | 174636914 | 174829314 | ENSG00000145 | GLRA3 | 8001 | 40.588314 | 0.13357224 | 0.04233195 | 3.15535316 | 0.00160304 | 0.01726111 | WT_active |

| | | | | | | | | | | | | |
|---|---|---|---|---|---|---|---|---|---|---|---|---|
| 1132 | chr18 | 35581117 | 35711834 | ENSG00000141 | GALNT1 | 2589 | 31.9123931 | 0.13366263 | 0.0402474 | 3.32102548 | 0.00089687 | 0.01057328 | WT_active |
| 1133 | chr15 | 49155656 | 49367869 | ENSG00000156 | GALK2 | 2585 | 54.350258 | 0.13366553 | 0.03312323 | 4.035401 | 5.45E-05 | 0.00086745 | WT_active |
| 1134 | chr9 | 105447796 | 105552433 | ENSG00000106 | FSD1L | 83856 | 23.5170044 | 0.13374892 | 0.04865873 | 2.74871373 | 0.00598296 | 0.04969029 | WT_active |
| 1135 | chr3 | 174438573 | 175810552 | ENSG00000177 | NAALADL2 | 254827 | 307.343937 | 0.13375892 | 0.02232939 | 5.99026221 | 2.10E-09 | 4.64E-08 | WT_active |
| 1136 | chr14 | 51826195 | 51979342 | ENSG00000186 | GNG2 | 54331 | 43.5845207 | 0.13393186 | 0.03566667 | 3.7550983 | 0.00017327 | 0.00251587 | WT_active |
| 1137 | chr11 | 100687653 | 100991937 | ENSG00000165 | ARHGAP42 | 143872 | 78.8066813 | 0.1340095 | 0.02747145 | 4.87813742 | 1.07E-06 | 2.06E-05 | WT_active |
| 1138 | chr3 | 98732236 | 98821201 | ENSG00000064 | ST3GAL6 | 10402 | 22.4876256 | 0.13438572 | 0.04753894 | 2.82685549 | 0.00470075 | 0.04136665 | WT_active |
| 1139 | chr2 | 32946972 | 33399509 | ENSG00000049 | LTBP1 | 4052 | 128.128094 | 0.13469451 | 0.02194329 | 6.13830117 | 8.34E-10 | 1.87E-08 | WT_active |
| 1140 | chr20 | 53936777 | 54070594 | ENSG00000064 | BCAS1 | 8537 | 40.9753972 | 0.13477193 | 0.0347544 | 3.87783821 | 0.00010539 | 0.00159603 | WT_active |
| 1141 | chr14 | 50532509 | 50633068 | ENSG00000198 | ATL1 | 51062 | 24.1135752 | 0.13486179 | 0.04689064 | 2.87609157 | 0.00402633 | 0.03678809 | WT_active |
| 1142 | chr20 | 9537389 | 9839041 | ENSG00000101 | PAK7 | 57144 | 92.8916706 | 0.13493764 | 0.02497912 | 5.40201649 | 6.59E-08 | 1.37E-06 | WT_active |
| 1143 | chr1 | 55066359 | 55215113 | ENSG00000162 | USP24 | 23358 | 35.9363847 | 0.1349708 | 0.03993607 | 3.37967193 | 0.00072572 | 0.0088225 | WT_active |
| 1144 | chr11 | 107591091 | 107666779 | ENSG00000110 | ELMOD1 | 55531 | 18.8642352 | 0.13502212 | 0.05363817 | 2.5172765 | 0.0118266 | 0.0830165 | WT_active |
| 1145 | chr5 | 24487100 | 24644978 | ENSG00000040 | CDH10 | 1008 | 31.2956884 | 0.13523284 | 0.04609266 | 2.93393455 | 0.00334695 | 0.03164347 | WT_active |
| 1146 | chr7 | 112206588 | 112343096 | ENSG00000198 | ZNF277 | 11179 | 34.6411398 | 0.1352709 | 0.04064266 | 3.32829876 | 0.00087378 | 0.01033484 | WT_active |
| 1147 | chr12 | 65824131 | 65966295 | ENSG00000149 | HMGA2 | 8091 | 28.3869158 | 0.13543821 | 0.04464369 | 3.03375957 | 0.00241527 | 0.02408253 | WT_active |
| 1148 | chr20 | 33993646 | 34108308 | ENSG00000125 | RALY | 22913 | 34.3432067 | 0.1354456 | 0.04127731 | 3.28135737 | 0.00103309 | 0.01188752 | WT_active |
| 1149 | chr8 | 80967810 | 81112068 | ENSG00000076 | PAG1 | 55824 | 40.5684904 | 0.13559344 | 0.03582577 | 3.78480154 | 0.00015383 | 0.00224943 | WT_active |
| 1150 | chr2 | 227871054 | 227924344 | ENSG00000123 | DAW1 | 164781 | 15.6020008 | 0.13560252 | 0.05499623 | 2.46566932 | 0.01367575 | 0.09283286 | WT_active |
| 1151 | chr11 | 82973133 | 83071923 | ENSG00000137 | RAB30 | 27314 | 24.9667134 | 0.13653046 | 0.04497329 | 3.0358121 | 0.00239889 | 0.02395233 | WT_active |
| 1152 | chr6 | 127708072 | 127918631 | ENSG00000172 | THEMIS | 387357 | 48.3854134 | 0.13713134 | 0.03782236 | 3.62566858 | 0.00028821 | 0.00400246 | WT_active |
| 1153 | chr1 | 95117923 | 95247225 | ENSG00000271 | TMEM56-RWDI | 100527978 | 34.3051845 | 0.13745326 | 0.04047598 | 3.3959217 | 0.00068398 | 0.00843594 | WT_active |
| 1154 | chr5 | 53883945 | 54310582 | ENSG00000185 | ARL15 | 54622 | 111.013714 | 0.13754296 | 0.02514543 | 5.46989965 | 4.50E-08 | 9.46E-07 | WT_active |
| 1155 | chr3 | 119703022 | 119767102 | ENSG00000183 | MAATS1 | 89876 | 16.6870626 | 0.13760106 | 0.0541082 | 2.543072 | 0.01098826 | 0.07870279 | WT_active |
| 1156 | chr8 | 104489231 | 104589024 | ENSG00000147 | LRP12 | 29967 | 21.3521663 | 0.13772635 | 0.05086709 | 2.70757277 | 0.00677772 | 0.05489954 | WT_active |
| 1157 | chr2 | 58159243 | 58241372 | ENSG00000115 | FANCL | 55120 | 16.3809464 | 0.13799609 | 0.05433095 | 2.53991658 | 0.01108789 | 0.07910767 | WT_active |
| 1158 | chr4 | 146706638 | 146945882 | ENSG00000137 | TTC29 | 83894 | 60.6954514 | 0.138279 | 0.03270254 | 4.22838674 | 2.35E-05 | 0.00039738 | WT_active |
| 1159 | chr9 | 97411950 | 97496125 | ENSG00000196 | TDRD7 | 23424 | 23.343266 | 0.13830809 | 0.04817988 | 2.87066071 | 0.00409615 | 0.03721379 | WT_active |
| 1160 | chr18 | 62187258 | 62307829 | ENSG00000134 | KIAA1468 | 57614 | 25.9001704 | 0.13850854 | 0.04599281 | 3.01152583 | 0.00259938 | 0.02558219 | WT_active |
| 1161 | chr12 | 70242994 | 70354993 | ENSG00000111 | CNOT2 | 4848 | 21.5249289 | 0.13855573 | 0.0495078 | 2.79866438 | 0.00513144 | 0.04397074 | WT_active |
| 1162 | chr5 | 102753981 | 103031105 | ENSG00000145 | PAM | 5066 | 64.6840508 | 0.1386571 | 0.03340346 | 4.15098048 | 3.31E-05 | 0.00054729 | WT_active |
| 1163 | chr4 | 177728757 | 177990750 | ENSG00000231 | LINC01098 | 285501 | 61.3814308 | 0.13919723 | 0.03628437 | 3.83628605 | 0.00012491 | 0.0018624 | WT_active |
| 1164 | chr4 | 77157151 | 77433388 | ENSG00000138 | CCNG2 | 901 | 109.707249 | 0.1394794 | 0.026602 | 5.24319278 | 1.58E-07 | 3.20E-06 | WT_active |
| 1165 | chr7 | 107923799 | 108003255 | ENSG00000091 | LAMB1 | 3912 | 19.0272758 | 0.13959103 | 0.05161525 | 2.70445348 | 0.00684168 | 0.05520057 | WT_active |
| 1166 | chr6 | 127118604 | 127197765 | ENSG00000146 | RSPO3 | 84870 | 18.5094899 | 0.13961276 | 0.05256397 | 2.6560544 | 0.00790609 | 0.0617497 | WT_active |
| 1167 | chrX | 74732849 | 74925485 | ENSG00000050 | KIAA2022 | 340533 | 58.3872913 | 0.13969559 | 0.03360316 | 4.1572153 | 3.22E-05 | 0.00053511 | WT_active |
| 1168 | chr6 | 47878028 | 48068689 | ENSG00000244 | PTCHD4 | 442213 | 50.8738877 | 0.13971924 | 0.03398853 | 4.11077639 | 3.94E-05 | 0.00064534 | WT_active |
| 1169 | chr3 | 158105855 | 158545730 | ENSG00000174 | RSRC1 | 51319 | 96.7503623 | 0.13995958 | 0.02637601 | 5.30632172 | 1.12E-07 | 2.29E-06 | WT_active |
| 1170 | chr15 | 69160584 | 69272217 | ENSG00000138 | GLCE | 26035 | 26.3289064 | 0.14008892 | 0.04491937 | 3.11867476 | 0.00181666 | 0.01910478 | WT_active |
| 1171 | chr2 | 143938068 | 144332568 | ENSG00000121 | GTDC1 | 79712 | 97.9149811 | 0.14022205 | 0.02704895 | 5.1840106 | 2.17E-07 | 4.38E-06 | WT_active |
| 1172 | chr1 | 212035553 | 212107400 | ENSG00000143 | DTL | 51514 | 18.478853 | 0.14078861 | 0.05358464 | 2.62740632 | 0.00860385 | 0.06556572 | WT_active |
| 1173 | chr5 | 34838833 | 34899456 | ENSG00000205 | TTC23L | 153657 | 17.7275396 | 0.14081962 | 0.05261294 | 2.67652075 | 0.0074391 | 0.05906216 | WT_active |
| 1174 | chr5 | 144170832 | 144485686 | ENSG00000183 | KCTD16 | 57528 | 89.7690893 | 0.14096187 | 0.02962337 | 4.75846834 | 1.95E-06 | 3.65E-05 | WT_active |
| 1175 | chr11 | 89177452 | 89295759 | ENSG00000077 | TYR | 7299 | 28.56664 | 0.14097844 | 0.0449078 | 3.1392867 | 0.0016936 | 0.01807422 | WT_active |
| 1176 | chr3 | 101222546 | 101320560 | ENSG00000081 | IMPG2 | 50939 | 27.2771077 | 0.14110555 | 0.0446522 | 3.16010282 | 0.00157713 | 0.01707031 | WT_active |
| 1177 | chr2 | 153871913 | 154453849 | ENSG00000144 | GALNT13 | 114805 | 130.811048 | 0.14125783 | 0.03017477 | 4.68132274 | 2.85E-06 | 5.28E-05 | WT_active |
| 1178 | chr6 | 53001279 | 53061802 | ENSG00000112 | ICK | 22858 | 17.4791471 | 0.14171621 | 0.05834922 | 2.42875947 | 0.01515058 | 0.09933654 | WT_active |
| 1179 | chr8 | 131904088 | 132013642 | ENSG00000132 | EFR3A | 23167 | 24.1954398 | 0.14176969 | 0.05176414 | 2.73876241 | 0.00616709 | 0.05092619 | WT_active |
| 1180 | chr2 | 216412414 | 216483053 | ENSG00000138 | SMARCAL1 | 50485 | 20.3757072 | 0.14205075 | 0.05561981 | 2.55395959 | 0.01065056 | 0.07715553 | WT_active |
| 1181 | chr7 | 123452400 | 123535077 | ENSG00000164 | IQUB | 154865 | 19.5514034 | 0.14211166 | 0.0544843 | 2.60830461 | 0.00909919 | 0.06840051 | WT_active |
| 1182 | chr14 | 26443093 | 26597754 | ENSG00000139 | NOVA1 | 4857 | 28.1383736 | 0.14271098 | 0.05069032 | 2.81534959 | 0.00487242 | 0.04245496 | WT_active |
| 1183 | chr2 | 112055223 | 112119318 | ENSG00000153 | TMEM87B | 84910 | 13.9029339 | 0.1427423 | 0.05589341 | 2.55383037 | 0.01065451 | 0.07715553 | WT_active |
| 1184 | chr15 | 61852389 | 62060473 | ENSG00000129 | VPS13C | 54832 | 40.4863877 | 0.14300121 | 0.03847558 | 3.71667449 | 0.00020186 | 0.00288078 | WT_active |
| 1185 | chr1 | 227563543 | 227677443 | ENSG00000181 | ZNF678 | 339500 | 32.9697412 | 0.1431755 | 0.04060146 | 3.52636313 | 0.00042131 | 0.00557287 | WT_active |

| | | | | | | | | | | | | |
|---|---|---|---|---|---|---|---|---|---|---|---|---|
| 1186 | chr3 | 142724074 | 142807888 | ENSG00000144 | TRPC1 | 7220 | 17.4106748 | 0.1433293 | 0.05901143 | 2.42883978 | 0.01514722 | 0.09933654 | WT_active |
| 1187 | chr6 | 117288300 | 117425855 | ENSG00000047 | ROS1 | 6098 | 37.4824239 | 0.14355682 | 0.04178226 | 3.43583178 | 0.00059074 | 0.00745167 | WT_active |
| 1188 | chr4 | 186208979 | 186258471 | ENSG00000164 | KLKB1 | 3818 | 14.3788244 | 0.14361028 | 0.0562819 | 2.55162436 | 0.01072221 | 0.07733505 | WT_active |
| 1189 | chr1 | 29236516 | 29326813 | ENSG00000060 | PTPRU | 10076 | 32.2581642 | 0.14364639 | 0.04267793 | 3.36582387 | 0.00076315 | 0.00914643 | WT_active |
| 1190 | chr5 | 123512099 | 123617045 | ENSG00000151 | CSNK1G3 | 1456 | 21.2781432 | 0.14364677 | 0.04959186 | 2.89657971 | 0.00377255 | 0.03506937 | WT_active |
| 1191 | chr7 | 13891228 | 13991425 | ENSG00000006 | ETV1 | 2115 | 21.7829695 | 0.14430366 | 0.05472544 | 2.63686631 | 0.00836758 | 0.06424056 | WT_active |
| 1192 | chr5 | 110289233 | 110738956 | ENSG00000186 | TMEM232 | 642987 | 109.589254 | 0.14541873 | 0.02914118 | 4.99014543 | 6.03E-07 | 1.18E-05 | WT_active |
| 1193 | chr5 | 107859035 | 108382098 | ENSG00000145 | FBXL17 | 64839 | 120.579896 | 0.14585043 | 0.02645755 | 5.51262001 | 3.54E-08 | 7.50E-07 | WT_active |
| 1194 | chr2 | 113215997 | 113278950 | ENSG00000125 | PAX8 | 7849 | 20.5441701 | 0.14589867 | 0.0537784 | 2.71296068 | 0.0066685 | 0.05432003 | WT_active |
| 1195 | chr5 | 81413021 | 81751797 | ENSG00000145 | SSBP2 | 23635 | 72.5349934 | 0.14590022 | 0.03166117 | 4.60817578 | 4.06E-06 | 7.44E-05 | WT_active |
| 1196 | chr14 | 52004803 | 52069228 | ENSG00000087 | NID2 | 22795 | 17.9059083 | 0.14623841 | 0.05480075 | 2.66854762 | 0.007618 | 0.06015131 | WT_active |
| 1197 | chr5 | 121961961 | 122075570 | ENSG00000151 | SRFBP1 | 153443 | 25.0527632 | 0.14652347 | 0.04798467 | 3.05354788 | 0.00226153 | 0.02291405 | WT_active |
| 1198 | chr3 | 171061339 | 171460408 | ENSG00000154 | TNIK | 23043 | 111.808435 | 0.14659873 | 0.02350351 | 6.23731183 | 4.45E-10 | 1.01E-08 | WT_active |
| 1199 | chr1 | 110150486 | 110202202 | ENSG00000197 | SLC6A17 | 388662 | 19.8221478 | 0.14668553 | 0.05543534 | 2.64606524 | 0.00814341 | 0.06288789 | WT_active |
| 1200 | chr6 | 52420308 | 52495785 | ENSG00000096 | EFHC1 | 114327 | 19.0712595 | 0.14684276 | 0.05167005 | 2.84193204 | 0.0044841 | 0.03990559 | WT_active |
| 1201 | chr5 | 115841481 | 115914081 | ENSG00000177 | AP3S1 | 1176 | 17.5134613 | 0.14711586 | 0.05593762 | 2.62999871 | 0.00853852 | 0.06517119 | WT_active |
| 1202 | chr9 | 108867517 | 108934116 | ENSG00000070 | IKBKAP | 8518 | 19.2122688 | 0.14741256 | 0.05472939 | 2.69348064 | 0.00707103 | 0.05670191 | WT_active |
| 1203 | chr3 | 138187248 | 138298389 | ENSG00000114 | ARMC8 | 25852 | 28.6216489 | 0.14746538 | 0.04286912 | 3.43989765 | 0.00058193 | 0.00735994 | WT_active |
| 1204 | chr2 | 168834132 | 168913371 | ENSG00000152 | SPC25 | 57405 | 21.9893926 | 0.14762748 | 0.04723894 | 3.12512261 | 0.00177731 | 0.01882825 | WT_active |
| 1205 | chr3 | 193593144 | 193697823 | ENSG00000198 | OPA1 | 4976 | 24.8477885 | 0.14811251 | 0.04922123 | 3.00911839 | 0.00262007 | 0.02575063 | WT_active |
| 1206 | chr6 | 127438406 | 127519355 | ENSG00000255 | C6ORF174 | 387104 | 19.9210539 | 0.14827066 | 0.05202699 | 2.84987948 | 0.00437358 | 0.03916663 | WT_active |
| 1207 | chr6 | 78935867 | 79078236 | ENSG00000146 | PHIP | 55023 | 27.4089294 | 0.14842629 | 0.0459623 | 3.22930509 | 0.00124091 | 0.01393333 | WT_active |
| 1208 | chr3 | 171600405 | 171810950 | ENSG00000075 | PLD1 | 5337 | 55.9392629 | 0.14874074 | 0.03028053 | 4.91209204 | 9.01E-07 | 1.74E-05 | WT_active |
| 1209 | chr7 | 90245174 | 90311063 | ENSG00000105 | C7orf63 | 79846 | 14.4640791 | 0.14886222 | 0.05716655 | 2.60400897 | 0.00921403 | 0.0690837 | WT_active |
| 1210 | chr6 | 111660332 | 111873452 | ENSG00000010 | FYN | 2534 | 63.3491539 | 0.14886345 | 0.02919862 | 5.09830417 | 3.43E-07 | 6.85E-06 | WT_active |
| 1211 | chr2 | 187464230 | 187565760 | ENSG00000003 | TFPI | 7035 | 23.005061 | 0.14901464 | 0.05184398 | 2.87429001 | 0.00404937 | 0.03690506 | WT_active |
| 1212 | chr2 | 120013005 | 120179119 | ENSG00000115 | EPB41L5 | 57669 | 40.6467064 | 0.14975473 | 0.03630055 | 4.12541234 | 3.70E-05 | 0.00060702 | WT_active |
| 1213 | chr7 | 95478444 | 95540232 | ENSG00000005 | ASB4 | 51666 | 17.2199023 | 0.14980699 | 0.05531726 | 2.70814188 | 0.00676611 | 0.05483629 | WT_active |
| 1214 | chr6 | 55434369 | 55579214 | ENSG00000146 | HMGCLL1 | 54511 | 33.9097446 | 0.14981009 | 0.04360814 | 3.43537008 | 0.00059175 | 0.00745785 | WT_active |
| 1215 | chr9 | 4985033 | 5128183 | ENSG00000096 | JAK2 | 3717 | 34.701798 | 0.15057629 | 0.04111132 | 3.66264827 | 0.00024962 | 0.00351126 | WT_active |
| 1216 | chr15 | 77420412 | 77485607 | ENSG00000140 | HMG20A | 10363 | 16.2284702 | 0.15103217 | 0.05499644 | 2.74621701 | 0.00602869 | 0.05002087 | WT_active |
| 1217 | chr4 | 94207611 | 94291292 | ENSG00000163 | SMARCAD1 | 56916 | 17.3177874 | 0.15161316 | 0.05575918 | 2.71907052 | 0.00654656 | 0.05356888 | WT_active |
| 1218 | chr9 | 124353466 | 124415444 | ENSG00000136 | PSMB7 | 5695 | 15.541523 | 0.15169327 | 0.05505338 | 2.75538536 | 0.0058623 | 0.04899867 | WT_active |
| 1219 | chr1 | 75796882 | 75913238 | ENSG00000057 | MSH4 | 4438 | 28.7536812 | 0.15211547 | 0.04803119 | 3.16701428 | 0.00154013 | 0.01670847 | WT_active |
| 1220 | chr4 | 122152333 | 122362758 | ENSG00000138 | KIAA1109 | 84162 | 45.7001606 | 0.15229889 | 0.03676864 | 4.14208681 | 3.44E-05 | 0.00056775 | WT_active |
| 1221 | chr6 | 46549580 | 46652830 | ENSG00000146 | CYP39A1 | 51302 | 25.9083084 | 0.1523045 | 0.04825671 | 3.15613102 | 0.00159877 | 0.017228 | WT_active |
| 1222 | chr2 | 195737703 | 196068812 | ENSG00000118 | DNAH7 | 56171 | 80.6141448 | 0.15246255 | 0.03077917 | 4.95343263 | 7.29E-07 | 1.41E-05 | WT_active |
| 1223 | chr7 | 37848597 | 37900401 | ENSG00000086 | NME8 | 51314 | 14.3560669 | 0.15329873 | 0.06260264 | 2.44875809 | 0.01433497 | 0.09608628 | WT_active |
| 1224 | chr8 | 109240919 | 109334385 | ENSG00000120 | NUDCD1 | 84955 | 21.7544663 | 0.15340936 | 0.0497024 | 3.08655845 | 0.00202488 | 0.02082364 | WT_active |
| 1225 | chr9 | 105558130 | 105641118 | ENSG00000106 | FKTN | 2218 | 19.7505982 | 0.15357888 | 0.05622804 | 2.73135781 | 0.0063074 | 0.05190284 | WT_active |
| 1226 | chr14 | 30622112 | 30735812 | ENSG00000092 | SCFD1 | 23256 | 23.7392931 | 0.15389216 | 0.04935291 | 3.11819866 | 0.0018196 | 0.0191078 | WT_active |
| 1227 | chr5 | 100806935 | 100903266 | ENSG00000113 | ST8SIA4 | 7903 | 19.7664755 | 0.15397859 | 0.05681219 | 2.71030873 | 0.00672206 | 0.05457223 | WT_active |
| 1228 | chr5 | 52989326 | 53094779 | ENSG00000164 | ITGA2 | 3673 | 26.3716469 | 0.1540564 | 0.04435904 | 3.47294226 | 0.00051479 | 0.00665048 | WT_active |
| 1229 | chr6 | 143940300 | 144064599 | ENSG00000118 | PLAGL1 | 5325 | 36.2283292 | 0.15420969 | 0.04163285 | 3.704039 | 0.00021219 | 0.00300827 | WT_active |
| 1230 | chr18 | 73151241 | 73264480 | ENSG00000263 | RP11-169F17.1 | 400655 | 25.3258405 | 0.15442169 | 0.04804728 | 3.21395254 | 0.00130921 | 0.014565 | WT_active |
| 1231 | chr6 | 106571971 | 106629487 | ENSG00000130 | RTN4IP1 | 84816 | 15.6284657 | 0.15443021 | 0.06348642 | 2.43249193 | 0.01499533 | 0.09876775 | WT_active |
| 1232 | chr2 | 64524305 | 64593005 | ENSG00000119 | AFTPH | 54812 | 13.9477411 | 0.1544402 | 0.06080835 | 2.53978601 | 0.01109203 | 0.07910767 | WT_active |
| 1233 | chr5 | 75068275 | 75236878 | ENSG00000145 | ANKRD31 | 256006 | 44.0948067 | 0.15457032 | 0.03642506 | 4.24351641 | 2.20E-05 | 0.00037413 | WT_active |
| 1234 | chr5 | 129460265 | 129738683 | ENSG00000145 | ADAMTS19 | 171019 | 60.8737911 | 0.15470146 | 0.03536925 | 4.37389656 | 1.22E-05 | 0.00021252 | WT_active |
| 1235 | chr11 | 74807739 | 74949200 | ENSG00000166 | XRRA1 | 143570 | 40.3668273 | 0.15507875 | 0.03627791 | 4.27474341 | 1.91E-05 | 0.00033002 | WT_active |
| 1236 | chr6 | 127968779 | 128520674 | ENSG00000152 | PTPRK | 5796 | 122.398037 | 0.1553786 | 0.0246478 | 6.30395485 | 2.90E-10 | 6.62E-09 | WT_active |
| 1237 | chr12 | 95867048 | 95942974 | ENSG00000165 | CCDC38 | 120935 | 20.1045348 | 0.15560549 | 0.04881463 | 3.18768145 | 0.00143418 | 0.01572477 | WT_active |
| 1238 | chr6 | 55106460 | 55282620 | ENSG00000137 | HCRTR2 | 3062 | 39.8521642 | 0.1557327 | 0.03952563 | 3.94004344 | 8.15E-05 | 0.00124823 | WT_active |
| 1239 | chr4 | 144646021 | 144745271 | ENSG00000164 | HHIP | 64399 | 21.1708767 | 0.15591537 | 0.05134142 | 3.03683405 | 0.00239077 | 0.02390439 | WT_active |

| | | | | | | | | | | | | | | |
|---|---|---|---|---|---|---|---|---|---|---|---|---|---|---|
| 1240 | chr14 | 63204114 | 63293219 | ENSG00000126 | RHOJ | 57381 | 25.0084172 | | 0.15689678 | 0.04441342 | 3.5326433 | 0.00041143 | 0.00545217 | WT_active |
| 1241 | chr1 | 64745095 | 64833232 | ENSG00000162 | RAVER2 | 55225 | 20.7187916 | | 0.15696966 | 0.05452871 | 2.87866097 | 0.00399367 | 0.03653604 | WT_active |
| 1242 | chr4 | 48986247 | 49062081 | ENSG00000109 | CWH43 | 80157 | 22.5514584 | | 0.15750801 | 0.05173901 | 3.0442796 | 0.00233238 | 0.02338922 | WT_active |
| 1243 | chr7 | 120273668 | 120750331 | ENSG00000184 | KCND2 | 3751 | 109.315748 | | 0.15793459 | 0.02617589 | 6.03359099 | 1.60E-09 | 3.57E-08 | WT_active |
| 1244 | chr2 | 68467561 | 68655862 | ENSG00000169 | APLF | 200558 | 49.2427453 | | 0.15794139 | 0.0338635 | 4.66406017 | 3.10E-06 | 5.72E-05 | WT_active |
| 1245 | chr2 | 101397361 | 101474703 | ENSG00000196 | RFX8 | 731220 | 24.2770597 | | 0.15796909 | 0.0468352 | 3.37287102 | 0.00074389 | 0.00896771 | WT_active |
| 1246 | chr6 | 115931149 | 116060758 | ENSG00000111 | FRK | 2444 | 32.2836007 | | 0.15812307 | 0.04147502 | 3.81248921 | 0.00013757 | 0.00203859 | WT_active |
| 1247 | chr1 | 215005775 | 215237093 | ENSG00000082 | KCNK2 | 3776 | 55.5052241 | | 0.15860244 | 0.03424067 | 4.6319897 | 3.62E-06 | 6.65E-05 | WT_active |
| 1248 | chr1 | 84298366 | 84389957 | ENSG00000203 | SAMD13 | 148418 | 25.863529 | | 0.15891062 | 0.04652967 | 3.41525362 | 0.00063723 | 0.00796147 | WT_active |
| 1249 | chr12 | 109477402 | 109536705 | ENSG00000151 | UBE3B | 89910 | 21.1486899 | | 0.15909654 | 0.05085976 | 3.12814206 | 0.00175915 | 0.01867706 | WT_active |
| 1250 | chr7 | 117276631 | 117323289 | ENSG00000105 | WNT2 | 7472 | 14.7016923 | | 0.1591752 | 0.05942504 | 2.67858777 | 0.00739334 | 0.05882843 | WT_active |
| 1251 | chr5 | 50665899 | 50846522 | ENSG00000151 | PARP8 | 79668 | 38.7597635 | | 0.16127824 | 0.03779164 | 4.26756397 | 1.98E-05 | 0.00034001 | WT_active |
| 1252 | chr12 | 65278643 | 65488244 | ENSG00000174 | MSRB3 | 253827 | 49.0379759 | | 0.16302985 | 0.03384415 | 4.81707692 | 1.46E-06 | 2.76E-05 | WT_active |
| 1253 | chr11 | 27365961 | 27472775 | ENSG00000205 | LGR4 | 55366 | 24.6275775 | | 0.16392469 | 0.0475886 | 3.44462072 | 0.00057186 | 0.00725162 | WT_active |
| 1254 | chr3 | 39107704 | 39138903 | ENSG00000168 | TTC21A | 199223 | 12.4726605 | | 0.16404092 | 0.06675356 | 2.45741101 | 0.01399425 | 0.09437281 | WT_active |
| 1255 | chrX | 106802680 | 106876145 | ENSG00000133 | TBC1D8B | 54885 | 16.2696894 | | 0.16434038 | 0.05615178 | 2.92671705 | 0.0034256 | 0.03230239 | WT_active |
| 1256 | chr12 | 101155493 | 101210407 | ENSG00000256 | SLC5A8 | 160728 | 15.4528565 | | 0.16436959 | 0.05709875 | 2.87868976 | 0.00399331 | 0.03653604 | WT_active |
| 1257 | chr4 | 78776342 | 78916372 | ENSG00000138 | BMP2K | 55589 | 29.5267773 | | 0.16454776 | 0.04052818 | 4.06008266 | 4.91E-05 | 0.00078645 | WT_active |
| 1258 | chr17 | 63484817 | 63521848 | ENSG00000264 | ACE | NaN | 13.2774151 | | 0.16482543 | 0.06190772 | 2.66243753 | 0.0077577 | 0.06082136 | WT_active |
| 1259 | chr6 | 110424687 | 110476641 | ENSG00000004 | SLC22A16 | 85413 | 16.5000219 | | 0.16501097 | 0.05750377 | 2.86956774 | 0.00411033 | 0.03729564 | WT_active |
| 1260 | chr11 | 94415578 | 94493908 | ENSG00000020 | MRE11A | 4361 | 19.2205213 | | 0.16552363 | 0.05390502 | 3.07065338 | 0.00213591 | 0.0217933 | WT_active |
| 1261 | chr9 | 99104038 | 99154192 | ENSG00000106 | TGFBR1 | 7046 | 13.7803799 | | 0.16553877 | 0.06264182 | 2.64262405 | 0.00822663 | 0.06347427 | WT_active |
| 1262 | chr12 | 75275979 | 75390928 | ENSG00000180 | CAPS2 | 84698 | 23.886156 | | 0.16575169 | 0.04887842 | 3.39110148 | 0.00069612 | 0.00857105 | WT_active |
| 1263 | chr2 | 165469647 | 165689407 | ENSG00000178 | CSRNP3 | 80034 | 52.2249922 | | 0.16658764 | 0.03613514 | 4.61012844 | 4.02E-06 | 7.38E-05 | WT_active |
| 1264 | chr2 | 101991844 | 102028544 | ENSG00000115 | IL1R2 | 7850 | 12.4466464 | | 0.16671442 | 0.06162264 | 2.70540868 | 0.00682204 | 0.05513463 | WT_active |
| 1265 | chr10 | 58513140 | 58831437 | ENSG00000122 | BICC1 | 80114 | 77.7405625 | | 0.16679398 | 0.02679398 | 6.22505421 | 4.81E-10 | 1.09E-08 | WT_active |
| 1266 | chr8 | 39743735 | 39838289 | ENSG00000104 | ADAM2 | 2515 | 21.1186329 | | 0.16705629 | 0.05162033 | 3.23624976 | 0.00121111 | 0.01364206 | WT_active |
| 1267 | chr5 | 64165844 | 64372869 | ENSG00000164 | RNF180 | 285671 | 51.876664 | | 0.16715146 | 0.03452816 | 4.84101861 | 1.29E-06 | 2.47E-05 | WT_active |
| 1268 | chr8 | 94378377 | 94436952 | ENSG00000265 | FSBP | 25788 | 12.2975723 | | 0.16724506 | 0.06439679 | 2.5971024 | 0.00940139 | 0.07023276 | WT_active |
| 1269 | chr8 | 78516139 | 78605267 | ENSG00000171 | PKIA | 5569 | 19.2501791 | | 0.16744003 | 0.05569565 | 3.00633967 | 0.00264413 | 0.0259341 | WT_active |
| 1270 | chr20 | 45363200 | 45410610 | ENSG00000254 | SYS1-DBNDD2 | 767557 | 13.565401 | | 0.16781234 | 0.06234273 | 2.69177072 | 0.00710738 | 0.05683538 | WT_active |
| 1271 | chr3 | 93980139 | 94055668 | ENSG00000169 | ARL13B | 200894 | 14.6533104 | | 0.16843279 | 0.05912408 | 2.84880191 | 0.00438842 | 0.0392707 | WT_active |
| 1272 | chr9 | 96816472 | 96875623 | ENSG00000196 | ZNF782 | 158431 | 14.2723774 | | 0.16850678 | 0.0603695 | 2.79125685 | 0.00525038 | 0.04479957 | WT_active |
| 1273 | chr6 | 108986437 | 109094819 | ENSG00000080 | SESN1 | 27244 | 24.5119655 | | 0.16874071 | 0.04866143 | 3.46764765 | 0.00052504 | 0.00673484 | WT_active |
| 1274 | chr7 | 121873089 | 122062036 | ENSG00000106 | PTPRZ1 | 5803 | 44.4973501 | | 0.16896593 | 0.04206167 | 4.01709999 | 5.89E-05 | 0.00093453 | WT_active |
| 1275 | chr16 | 46580554 | 46621626 | ENSG00000171 | SHCBP1 | 79801 | 10.0488157 | | 0.16914739 | 0.06787072 | 2.49219953 | 0.01269547 | 0.08739821 | WT_active |
| 1276 | chr11 | 22193176 | 22283357 | ENSG00000171 | ANO5 | 203859 | 19.3433063 | | 0.16980766 | 0.05162434 | 3.28929471 | 0.00100439 | 0.01164464 | WT_active |
| 1277 | chr21 | 26917912 | 26966513 | ENSG00000154 | ADAMTS5 | 11096 | 12.3695726 | | 0.16999543 | 0.06004224 | 2.83126408 | 0.00463644 | 0.04091079 | WT_active |
| 1278 | chr1 | 196225779 | 196609225 | ENSG00000162 | KCNT2 | 343450 | 80.1285223 | | 0.1702788 | 0.0344748 | 4.93922576 | 7.84E-07 | 1.52E-05 | WT_active |
| 1279 | chr18 | 35452230 | 35497991 | ENSG00000153 | INO80C | 125476 | 12.0854461 | | 0.17082442 | 0.0668473 | 2.55544223 | 0.0106053 | 0.07691507 | WT_active |
| 1280 | chr2 | 210431249 | 210477652 | ENSG00000115 | LANCL1 | 10314 | 12.6491268 | | 0.17084321 | 0.07017172 | 2.43464462 | 0.01490642 | 0.09860968 | WT_active |
| 1281 | chr5 | 115578642 | 115626161 | ENSG00000251 | TMED7-TICAM2 | 100302736 | 11.7058823 | | 0.1710683 | 0.06503671 | 2.63033428 | 0.00853009 | 0.06517119 | WT_active |
| 1282 | chr8 | 115408496 | 115809673 | ENSG00000104 | TRPS1 | 7227 | 94.5385839 | | 0.17113719 | 0.02569104 | 6.66135591 | 2.71E-11 | 6.36E-10 | WT_active |
| 1283 | chr3 | 108043298 | 108091025 | ENSG00000196 | CD47 | 961 | 11.410441 | | 0.17124685 | 0.07021172 | 2.43900655 | 0.0147277 | 0.09769694 | WT_active |
| 1284 | chr7 | 15611212 | 15686812 | ENSG00000106 | MEOX2 | 4223 | 15.5639503 | | 0.17227487 | 0.06184535 | 2.78557505 | 0.00534329 | 0.04534403 | WT_active |
| 1285 | chr3 | 132417526 | 132539032 | ENSG00000138 | DNAJC13 | 23317 | 26.1955138 | | 0.17271735 | 0.04815732 | 3.58652351 | 0.00033512 | 0.00456682 | WT_active |
| 1286 | chr5 | 136132845 | 136188747 | ENSG00000113 | SMAD5 | 4090 | 13.4024586 | | 0.17280892 | 0.06211671 | 2.78200357 | 0.00540244 | 0.04573837 | WT_active |
| 1287 | chr6 | 29355230 | 29431967 | ENSG00000243 | OR5V1 | 81696 | 19.4913295 | | 0.17374236 | 0.0523098 | 3.32141107 | 0.00089564 | 0.01056732 | WT_active |
| 1288 | chr5 | 10353703 | 10440388 | ENSG00000145 | 38777 | 10299 | 20.2759464 | | 0.17374238 | 0.0487897 | 3.56104612 | 0.00036938 | 0.00497612 | WT_active |
| 1289 | chr12 | 96000828 | 96043520 | ENSG00000111 | LTA4H | 4048 | 10.6016867 | | 0.1739296 | 0.06858672 | 2.53590784 | 0.01121562 | 0.0798947 | WT_active |
| 1290 | chr1 | 109548611 | 109618321 | ENSG00000065 | GNAI3 | 2773 | 18.0779283 | | 0.17454051 | 0.0562176 | 3.10473067 | 0.00190452 | 0.01985497 | WT_active |
| 1291 | chr6 | 131573966 | 131628229 | ENSG00000112 | MED23 | 9439 | 12.2570543 | | 0.17460505 | 0.06378221 | 2.73751976 | 0.00619044 | 0.05108973 | WT_active |
| 1292 | chr10 | 49818279 | 49970203 | ENSG00000227 | PARG | 8505 | 27.468512 | | 0.17486772 | 0.04330275 | 4.03825912 | 5.38E-05 | 0.0008579 | WT_active |
| 1293 | chr8 | 98189833 | 98294393 | ENSG00000104 | NIPAL2 | 79815 | 27.3219025 | | 0.17516386 | 0.04281093 | 4.09156909 | 4.28E-05 | 0.00069489 | WT_active |

| | | | | | | | | | | | | |
|---|---|---|---|---|---|---|---|---|---|---|---|---|
| 1294 | chrX | 38269163 | 38327564 | ENSG00000156 | RPGR | 6103 | 13.6349714 | 0.17524431 | 0.05706041 | 3.07120686 | 0.00213195 | 0.02176948 | WT_active |
| 1295 | chr2 | 162267079 | 162318703 | ENSG00000115 | IFIH1 | 64135 | 12.1505239 | 0.1754553 | 0.06950484 | 2.52436088 | 0.01159089 | 0.08184007 | WT_active |
| 1296 | chr2 | 190189735 | 190344193 | ENSG00000198 | HIBCH | 26275 | 40.415389 | 0.17578023 | 0.03885626 | 4.52385842 | 6.07E-06 | 0.00010971 | WT_active |
| 1297 | chr1 | 65420652 | 65641559 | ENSG00000116 | LEPR | 3953 | 51.8764739 | 0.17591116 | 0.03322636 | 5.29432594 | 1.19E-07 | 2.43E-06 | WT_active |
| 1298 | chr7 | 121348851 | 121396364 | ENSG00000196 | FAM3C | 10447 | 10.1465647 | 0.17594483 | 0.06998377 | 2.51408047 | 0.01193432 | 0.08360983 | WT_active |
| 1299 | chr17 | 48997412 | 49055650 | ENSG00000159 | IGF2BP1 | 10642 | 15.2364558 | 0.17694659 | 0.06165931 | 2.86974666 | 0.00410801 | 0.03729564 | WT_active |
| 1300 | chrX | 106611930 | 106679442 | ENSG00000147 | CXorf57 | 55086 | 14.988768 | 0.17698279 | 0.06113285 | 2.89505234 | 0.00379095 | 0.03519508 | WT_active |
| 1301 | chr19 | 55891699 | 55932336 | ENSG00000173 | NLRP13 | 126204 | 12.9970915 | 0.17714555 | 0.06440333 | 2.7505652 | 0.00594926 | 0.04953329 | WT_active |
| 1302 | chr12 | 88049014 | 88142216 | ENSG00000198 | CEP290 | 80184 | 17.8711419 | 0.17737784 | 0.05582496 | 3.17739293 | 0.00148606 | 0.01618274 | WT_active |
| 1303 | chr3 | 108160812 | 108222570 | ENSG00000114 | IFT57 | 55081 | 14.6078354 | 0.1775075 | 0.06423619 | 2.76335647 | 0.00572102 | 0.04798472 | WT_active |
| 1304 | chr4 | 119494395 | 119628991 | ENSG00000138 | PDE5A | 8654 | 29.9248234 | 0.17753517 | 0.04437587 | 4.00071376 | 6.32E-05 | 0.00099185 | WT_active |
| 1305 | chr8 | 90001405 | 90052092 | ENSG00000104 | DECR1 | 1666 | 13.6701735 | 0.17810133 | 0.06125561 | 2.90751032 | 0.00364318 | 0.03409202 | WT_active |
| 1306 | chr1 | 117863485 | 117929630 | ENSG00000196 | GDAP2 | 54834 | 16.7140688 | 0.17987754 | 0.06080486 | 2.95827575 | 0.00309365 | 0.0296571 | WT_active |
| 1307 | chrX | 135520659 | 135582510 | ENSG00000165 | DDX26B | 203522 | 17.0217087 | 0.18050369 | 0.05286696 | 3.41430045 | 0.00063946 | 0.00797555 | WT_active |
| 1308 | chr3 | 150541994 | 150603228 | ENSG00000120 | SERP1 | 27230 | 13.1420966 | 0.18050389 | 0.06434124 | 2.80541499 | 0.00502518 | 0.04338509 | WT_active |
| 1309 | chr1 | 185157080 | 185291765 | ENSG00000116 | SWT1 | 54823 | 32.9434953 | 0.18074205 | 0.04062466 | 4.44907284 | 8.62E-06 | 0.00015294 | WT_active |
| 1310 | chr1 | 46675159 | 46719064 | ENSG00000159 | EFCAB14 | 9813 | 12.5510279 | 0.18170859 | 0.06907409 | 2.63063308 | 0.0085226 | 0.06515315 | WT_active |
| 1311 | chr6 | 130365734 | 130443063 | ENSG00000164 | TMEM200A | 114801 | 19.6892621 | 0.18205849 | 0.05317499 | 3.42376163 | 0.00061761 | 0.00772975 | WT_active |
| 1312 | chr3 | 43079232 | 43106076 | ENSG00000144 | POMGNT2 | 84892 | 9.99782573 | 0.18286343 | 0.07284962 | 2.51014928 | 0.01206801 | 0.08413763 | WT_active |
| 1313 | chr7 | 81699006 | 81770438 | ENSG00000019 | HGF | 3082 | 14.2461091 | 0.18426991 | 0.06254585 | 2.9461572 | 0.00321749 | 0.0306203 | WT_active |
| 1314 | chr2 | 162371407 | 162838730 | ENSG00000184 | KCNH7 | 90134 | 105.400785 | 0.18508242 | 0.02981035 | 6.20866407 | 5.34E-10 | 1.20E-08 | WT_active |
| 1315 | chr3 | 108589682 | 108694846 | ENSG00000198 | DZIP3 | 9666 | 24.568474 | 0.18516089 | 0.04805425 | 3.85316378 | 0.0001166 | 0.00175121 | WT_active |
| 1316 | chr2 | 216498189 | 216579180 | ENSG00000197 | RPL37A | 6168 | 25.7916654 | 0.18588107 | 0.04653422 | 3.99450299 | 6.48E-05 | 0.0010149 | WT_active |
| 1317 | chr2 | 161136908 | 161236221 | ENSG00000136 | TANK | 10010 | 22.2908294 | 0.18777192 | 0.04607351 | 4.07548586 | 4.59E-05 | 0.00074138 | WT_active |
| 1318 | chr16 | 90056566 | 90092072 | ENSG00000126 | PRDM7 | 11105 | 9.78885277 | 0.18924495 | 0.07140501 | 2.65030361 | 0.00804195 | 0.06250608 | WT_active |
| 1319 | chr11 | 74748868 | 74842413 | ENSG00000166 | RNF169 | 254225 | 20.4101742 | 0.189386 | 0.05141974 | 3.68313773 | 0.00023038 | 0.0032565 | WT_active |
| 1320 | chr1 | 186311652 | 186375693 | ENSG00000047 | TPR | 7175 | 12.3496215 | 0.18943731 | 0.06882887 | 2.75229432 | 0.00591793 | 0.04937773 | WT_active |
| 1321 | chr8 | 123768456 | 123815452 | ENSG00000176 | FAM91A1 | 157769 | 10.5139022 | 0.19035379 | 0.07488036 | 2.54210583 | 0.01101868 | 0.07884235 | WT_active |
| 1322 | chr11 | 95789965 | 95832693 | ENSG00000166 | CEP57 | 9702 | 10.2672446 | 0.19055556 | 0.06723811 | 2.83404097 | 0.00459634 | 0.04063157 | WT_active |
| 1323 | chr7 | 136868669 | 137020255 | ENSG00000181 | CHRM2 | 1129 | 37.9047954 | 0.19057703 | 0.03976585 | 4.79247921 | 1.65E-06 | 3.10E-05 | WT_active |
| 1324 | chr5 | 74721204 | 74767371 | ENSG00000164 | GFM2 | 84340 | 10.6550125 | 0.19058235 | 0.06879215 | 2.77040859 | 0.0055986 | 0.04717746 | WT_active |
| 1325 | chr7 | 141656728 | 141702153 | ENSG00000257 | KIAA1147 | 57189 | 14.3574872 | 0.19075921 | 0.06482136 | 2.9428449 | 0.00325211 | 0.03088865 | WT_active |
| 1326 | chr1 | 59893308 | 59926790 | ENSG00000134 | CYP2J2 | 1573 | 10.9634933 | 0.19236937 | 0.07547102 | 2.54891721 | 0.01080579 | 0.07763 | WT_active |
| 1327 | chr2 | 165747591 | 165794682 | ENSG00000115 | GALNT3 | 2591 | 10.6955091 | 0.19370034 | 0.07288052 | 2.65777935 | 0.00786574 | 0.06146786 | WT_active |
| 1328 | chr2 | 74654228 | 74683853 | ENSG00000135 | SEMA4F | 10505 | 10.3656308 | 0.19383523 | 0.07353319 | 2.63602356 | 0.00838839 | 0.06436604 | WT_active |
| 1329 | chr7 | 16753535 | 16784536 | ENSG00000106 | TSPAN13 | 27075 | 7.63752341 | 0.19433566 | 0.07898217 | 2.46050046 | 0.01387434 | 0.09379248 | WT_active |
| 1330 | chr5 | 96390415 | 96434143 | ENSG00000175 | PCSK1 | 5122 | 12.6097195 | 0.19440342 | 0.06723256 | 2.89150708 | 0.00383399 | 0.03550319 | WT_active |
| 1331 | chr4 | 121879027 | 121951754 | ENSG00000138 | TRPC3 | 7222 | 19.5672198 | 0.19454597 | 0.05394769 | 3.60619661 | 0.00031072 | 0.00426697 | WT_active |
| 1332 | chr2 | 165989160 | 166128047 | ENSG00000144 | SCN1A | 6323 | 30.3032642 | 0.19487346 | 0.04592556 | 4.24324588 | 2.20E-05 | 0.00037414 | WT_active |
| 1333 | chrX | 136848004 | 136880764 | ENSG00000147 | RBMX | 27316 | 7.92057911 | 0.19515509 | 0.0794066 | 2.45766833 | 0.01398423 | 0.09434935 | WT_active |
| 1334 | chr5 | 80487969 | 80542563 | ENSG00000152 | FAM151B | 167555 | 13.3002256 | 0.19576314 | 0.06582732 | 2.97388893 | 0.00294051 | 0.02843484 | WT_active |
| 1335 | chr12 | 56416373 | 56449403 | ENSG00000111 | TIMELESS | 8914 | 9.65096887 | 0.19599239 | 0.07098048 | 2.76121521 | 0.00575867 | 0.04822477 | WT_active |
| 1336 | chr4 | 127781821 | 127840733 | ENSG00000164 | HSPA4L | 22824 | 11.4501718 | 0.19610596 | 0.07434737 | 2.63769851 | 0.00834708 | 0.06415146 | WT_active |
| 1337 | chr5 | 83471465 | 83582303 | ENSG00000038 | VCAN | 1462 | 24.2824044 | 0.19749327 | 0.04930948 | 4.00517849 | 6.20E-05 | 0.00097543 | WT_active |
| 1338 | chr1 | 78889764 | 79006718 | ENSG00000162 | ELTD1 | 64123 | 25.2520742 | 0.19760728 | 0.04918625 | 4.01753075 | 5.88E-05 | 0.00093385 | WT_active |
| 1339 | chr8 | 11284416 | 11328146 | ENSG00000104 | MTMR9 | 66036 | 10.9795052 | 0.19809521 | 0.0703664 | 2.81519588 | 0.00487475 | 0.04245496 | WT_active |
| 1340 | chr3 | 194640788 | 194672477 | ENSG00000041 | LSG1 | 55341 | 8.99039913 | 0.19851648 | 0.0807472 | 2.45849372 | 0.01395212 | 0.09417682 | WT_active |
| 1341 | chr4 | 47847233 | 47914667 | ENSG00000170 | NFXL1 | 152518 | 14.4902756 | 0.19864332 | 0.06222319 | 3.19243234 | 0.0014108 | 0.01552741 | WT_active |
| 1342 | chr14 | 52553148 | 52637713 | ENSG00000180 | GPR137C | 283554 | 19.8437514 | 0.19887316 | 0.05366191 | 3.70603975 | 0.00021053 | 0.00298755 | WT_active |
| 1343 | chr7 | 127346790 | 127431924 | ENSG00000048 | ZNF800 | 168850 | 21.5585301 | 0.1993133 | 0.04780331 | 4.16944585 | 3.05E-05 | 0.00050895 | WT_active |
| 1344 | chr8 | 23020133 | 23069179 | ENSG00000120 | TNFRSF10B | 8795 | 11.8867547 | 0.19997839 | 0.06699827 | 2.98482921 | 0.00283737 | 0.02749271 | WT_active |
| 1345 | chr20 | 38127387 | 38166578 | ENSG00000198 | TGM2 | 7052 | 11.3042337 | 0.2017026 | 0.07068098 | 2.85370412 | 0.00432128 | 0.03882825 | WT_active |
| 1346 | chr4 | 78887127 | 78939438 | ENSG00000163 | PAQR3 | 152559 | 12.9550205 | 0.20343143 | 0.06808909 | 2.98772447 | 0.00281063 | 0.02734388 | WT_active |
| 1347 | chr1 | 207880972 | 207911402 | ENSG00000174 | CD34 | 947 | 9.70415367 | 0.20375047 | 0.07875576 | 2.58711837 | 0.00967823 | 0.07178023 | WT_active |

| | | | | | | | | | | | | |
|---|---|---|---|---|---|---|---|---|---|---|---|---|
| 1348 | chr6 | 117675502 | 117710640 | ENSG00000153 | NUS1 | 116150 | 8.38549288 | 0.20389916 | 0.08070419 | 2.5265002 | 0.01152053 | 0.08146299 | WT_active |
| 1349 | chr1 | 15465909 | 15491400 | ENSG00000215 | CELA2B | 51032 | 9.11614364 | 0.20477831 | 0.07651984 | 2.67614654 | 0.00744741 | 0.05909561 | WT_active |
| 1350 | chr3 | 100334701 | 100361635 | ENSG00000114 | NIT2 | 56954 | 7.72694709 | 0.20650012 | 0.07894917 | 2.6156084 | 0.00890686 | 0.06737627 | WT_active |
| 1351 | chr5 | 72320367 | 72368395 | ENSG00000049 | PTCD2 | 79810 | 12.9549814 | 0.20747056 | 0.06436991 | 3.22309832 | 0.00126812 | 0.01420649 | WT_active |
| 1352 | chr12 | 26973195 | 27014434 | ENSG00000064 | TM7SF3 | 51768 | 11.3525297 | 0.20803507 | 0.0690412 | 3.01320186 | 0.00258507 | 0.02545869 | WT_active |
| 1353 | chr4 | 39045039 | 39126857 | ENSG00000109 | KLHL5 | 51088 | 17.2091884 | 0.20817165 | 0.05646913 | 3.68646803 | 0.00022739 | 0.00321735 | WT_active |
| 1354 | chr1 | 192316992 | 192367285 | ENSG00000253 | RGS21 | 431704 | 10.3659678 | 0.20837957 | 0.07437323 | 2.80180888 | 0.0050817 | 0.04369225 | WT_active |
| 1355 | chr2 | 176269395 | 176338025 | ENSG00000128 | MTX2 | 10651 | 14.5548225 | 0.20847667 | 0.0645243 | 3.23097899 | 0.00123367 | 0.01386364 | WT_active |
| 1356 | chr1 | 171781664 | 171814023 | ENSG00000010 | METTL13 | 51603 | 10.5304676 | 0.20898406 | 0.07561334 | 2.76385172 | 0.00571235 | 0.04793984 | WT_active |
| 1357 | chr12 | 96907223 | 96953777 | ENSG00000139 | NEDD1 | 121441 | 9.15822336 | 0.21001795 | 0.07984971 | 2.63016563 | 0.00853433 | 0.06517119 | WT_active |
| 1358 | chr12 | 43758944 | 43789543 | ENSG00000198 | IRAK4 | 51135 | 7.60511686 | 0.21101703 | 0.08099431 | 2.60533157 | 0.00917854 | 0.06888921 | WT_active |
| 1359 | chr3 | 33798352 | 33869707 | ENSG00000170 | PDCD6IP | 10015 | 14.4402606 | 0.21169023 | 0.06270004 | 3.37623736 | 0.00073484 | 0.00889236 | WT_active |
| 1360 | chr1 | 86346824 | 86396342 | ENSG00000122 | ODF2L | 57489 | 9.54622697 | 0.21254443 | 0.08282067 | 2.56632097 | 0.01027837 | 0.07499671 | WT_active |
| 1361 | chr7 | 124822386 | 124929983 | ENSG00000128 | POT1 | 25913 | 23.2129218 | 0.21276107 | 0.05142825 | 4.13704656 | 3.52E-05 | 0.00057837 | WT_active |
| 1362 | chr20 | 18288283 | 18316996 | ENSG00000125 | ZNF133 | 7692 | 7.58119415 | 0.21306084 | 0.08655803 | 2.46147983 | 0.01383652 | 0.09368255 | WT_active |
| 1363 | chr6 | 43509702 | 43529585 | ENSG00000171 | POLR1C | 9533 | 6.20157546 | 0.21317903 | 0.08780812 | 2.42778267 | 0.01519144 | 0.09951396 | WT_active |
| 1364 | chr21 | 25734570 | 25772460 | ENSG00000154 | GABPA | 2551 | 7.80168011 | 0.21362006 | 0.0815762 | 2.61865671 | 0.00882767 | 0.06695286 | WT_active |
| 1365 | chr16 | 66552563 | 66579135 | ENSG00000254 | CKLF-CMTM1 | 100529251 | 7.20780374 | 0.21410841 | 0.08526994 | 2.51094822 | 0.01204074 | 0.08399614 | WT_active |
| 1366 | chr2 | 112275594 | 112340063 | ENSG00000188 | ZC3H6 | 376940 | 13.5602774 | 0.21423696 | 0.06365007 | 3.36585564 | 0.00076307 | 0.00914643 | WT_active |
| 1367 | chr3 | 131462212 | 131502983 | ENSG00000114 | MRPL3 | 11222 | 9.94971439 | 0.21450731 | 0.07656603 | 2.80159896 | 0.005085 | 0.04369225 | WT_active |
| 1368 | chr17 | 30248195 | 30292056 | ENSG00000108 | BLMH | 642 | 11.9896178 | 0.21562272 | 0.06557166 | 3.28835209 | 0.00100776 | 0.01167056 | WT_active |
| 1369 | chr6 | 127472794 | 127519191 | ENSG00000214 | SOGA3 | NaN | 11.3843318 | 0.21609473 | 0.0696861 | 3.10097313 | 0.00192886 | 0.02002488 | WT_active |
| 1370 | chr2 | 46490750 | 46542557 | ENSG00000250 | ATP6V1E2 | 90423 | 16.601323 | 0.21614191 | 0.05434843 | 3.97696697 | 6.98E-05 | 0.00108446 | WT_active |
| 1371 | chr4 | 133149315 | 133208606 | ENSG00000138 | PCDH10 | 57575 | 10.2540065 | 0.21642465 | 0.07089949 | 3.05255568 | 0.00226902 | 0.02295767 | WT_active |
| 1372 | chr4 | 147480917 | 147544954 | ENSG00000151 | EDNRA | 1909 | 16.9361679 | 0.21650168 | 0.05269063 | 4.10892179 | 3.98E-05 | 0.00064981 | WT_active |
| 1373 | chr20 | 59863563 | 59933655 | ENSG00000196 | SYCP2 | 10388 | 12.8487154 | 0.21779871 | 0.072913 | 2.98710408 | 0.00281634 | 0.02736251 | WT_active |
| 1374 | chr11 | 31369840 | 31431849 | ENSG00000170 | DNAJC24 | 120526 | 14.5679901 | 0.21897545 | 0.06344719 | 3.45130277 | 0.00055789 | 0.00710567 | WT_active |
| 1375 | chr1 | 207104262 | 207144972 | ENSG00000123 | C4BPA | 722 | 11.8152081 | 0.2212458 | 0.06713114 | 3.29572551 | 0.00098168 | 0.01143283 | WT_active |
| 1376 | chr7 | 130293134 | 130324180 | ENSG00000128 | CPA4 | 51200 | 9.81714467 | 0.22251008 | 0.07470069 | 2.97868834 | 0.00289485 | 0.02801205 | WT_active |
| 1377 | chr15 | 49620980 | 49656232 | ENSG00000104 | DTWD1 | 56986 | 7.98914653 | 0.22290647 | 0.08517283 | 2.61710769 | 0.00886784 | 0.06722211 | WT_active |
| 1378 | chr4 | 155666711 | 155732349 | ENSG00000164 | GUCY1A3 | 2982 | 15.9381029 | 0.23124701 | 0.06339746 | 3.64757541 | 0.00026473 | 0.00370206 | WT_active |
| 1379 | chr2 | 19910260 | 19990131 | ENSG00000118 | WDR35 | 57539 | 18.2739927 | 0.23253013 | 0.05314799 | 4.3751445 | 1.21E-05 | 0.00021157 | WT_active |
| 1380 | chr2 | 171522247 | 171558133 | ENSG00000071 | CYBRD1 | 79901 | 8.01611946 | 0.23258452 | 0.08201239 | 2.83596808 | 0.0045687 | 0.0404865 | WT_active |
| 1381 | chr8 | 81657961 | 81686693 | ENSG00000133 | IMPA1 | 3612 | 6.75332796 | 0.23309132 | 0.08883842 | 2.62376699 | 0.00869633 | 0.0661306 | WT_active |
| 1382 | chr15 | 51723011 | 51751585 | ENSG00000140 | LYSMD2 | 256586 | 7.80237842 | 0.23425241 | 0.08234362 | 2.84481534 | 0.00444372 | 0.03964698 | WT_active |
| 1383 | chr7 | 106865278 | 106907145 | ENSG00000105 | PIK3CG | 5294 | 11.9475513 | 0.23445081 | 0.06681024 | 3.50920453 | 0.00044945 | 0.00590178 | WT_active |
| 1384 | chr5 | 33523535 | 33892192 | ENSG00000151 | ADAMTS12 | 81792 | 100.828897 | 0.23469054 | 0.02538541 | 9.24509534 | 2.35E-20 | 7.32E-19 | WT_active |
| 1385 | chr6 | 88139864 | 88166359 | ENSG00000118 | CNR1 | 1268 | 8.11882098 | 0.23538197 | 0.08468115 | 2.77962664 | 0.00544214 | 0.04604743 | WT_active |
| 1386 | chr3 | 28315003 | 28349127 | ENSG00000163 | AZI2 | 64343 | 6.66768429 | 0.23573803 | 0.08826776 | 2.67071505 | 0.00756899 | 0.05982986 | WT_active |
| 1387 | chr11 | 130875436 | 130916509 | ENSG00000120 | SNX19 | 399979 | 11.449589 | 0.2384307 | 0.07150785 | 3.33432904 | 0.00085505 | 0.01014665 | WT_active |
| 1388 | chr4 | 140524158 | 140553770 | ENSG00000179 | ELMOD2 | 255520 | 6.67959146 | 0.23885147 | 0.08928015 | 2.67530327 | 0.00746617 | 0.05917933 | WT_active |
| 1389 | chr6 | 28571630 | 28616212 | ENSG00000232 | ZBED9 | 114821 | 10.3956669 | 0.23910071 | 0.07069111 | 3.38233077 | 0.00071874 | 0.0087597 | WT_active |
| 1390 | chr12 | 75334639 | 75370560 | ENSG00000173 | GLIPR1L1 | 256710 | 6.48602 | 0.24100023 | 0.08914671 | 2.70341146 | 0.00686317 | 0.05531203 | WT_active |
| 1391 | chr6 | 122443354 | 122471822 | ENSG00000111 | SERINC1 | 57515 | 5.89498477 | 0.24261953 | 0.09325602 | 2.60165007 | 0.00927765 | 0.06945919 | WT_active |
| 1392 | chr9 | 113255835 | 113275589 | ENSG00000176 | CDC26 | 246184 | 5.9039627 | 0.24663058 | 0.09425359 | 2.61667041 | 0.0088792 | 0.06723758 | WT_active |
| 1393 | chr8 | 27733311 | 27772653 | ENSG00000147 | CCDC25 | 55246 | 11.2630887 | 0.24746587 | 0.07310898 | 3.38489021 | 0.00071207 | 0.00872267 | WT_active |
| 1394 | chr6 | 143450807 | 143490010 | ENSG00000034 | PEX3 | 8504 | 8.7022406 | 0.24749998 | 0.08381477 | 2.95293984 | 0.00314763 | 0.0300256 | WT_active |
| 1395 | chr2 | 190504342 | 190534722 | ENSG00000189 | TMEM194B | 100131211 | 8.92473601 | 0.24797183 | 0.07937963 | 3.12387228 | 0.00178488 | 0.01886898 | WT_active |
| 1396 | chr3 | 69084944 | 69106066 | ENSG00000144 | ARL6IP5 | 10550 | 4.58280858 | 0.24828241 | 0.10225524 | 2.4280654 | 0.01517961 | 0.09948162 | WT_active |
| 1397 | chr6 | 155257509 | 155314493 | ENSG00000029 | TFB1M | 51106 | 14.2012556 | 0.24829178 | 0.0599554 | 4.14127507 | 3.45E-05 | 0.00056846 | WT_active |
| 1398 | chr1 | 109603267 | 109619929 | ENSG00000134 | GNAT2 | 2780 | 5.74125073 | 0.25217915 | 0.10204587 | 2.47123333 | 0.01346479 | 0.09187666 | WT_active |
| 1399 | chr22 | 36189124 | 36204840 | ENSG00000100 | APOL4 | 80832 | 5.90551968 | 0.25362726 | 0.09315852 | 2.72253423 | 0.00647833 | 0.05307079 | WT_active |
| 1400 | chr6 | 49834257 | 49877096 | ENSG00000124 | CRISP1 | 167 | 11.2640745 | 0.25383866 | 0.07501883 | 3.38366612 | 0.00071525 | 0.00873198 | WT_active |
| 1401 | chr1 | 98661701 | 98760500 | ENSG00000162 | SNX7 | 51375 | 25.2008872 | 0.25455874 | 0.04692057 | 5.42531247 | 5.79E-08 | 1.21E-06 | WT_active |

| | | | | | | | | | | | | |
|---|---|---|---|---|---|---|---|---|---|---|---|---|
| 1402 | chr3 | 190852877 | 190892429 | ENSG00000205 | GMNC | 647309 | 8.4327176 | 0.25475614 | 0.08035918 | 3.17021819 | 0.00152325 | 0.01657521 | WT_active |
| 1403 | chr4 | 81030965 | 81057531 | ENSG00000152 | BMP3 | 651 | 7.40326605 | 0.25564895 | 0.08831851 | 2.89462473 | 0.00379612 | 0.03522038 | WT_active |
| 1404 | chr7 | 94394561 | 94431232 | ENSG00000164 | COL1A2 | 1278 | 8.71738281 | 0.2556853 | 0.07487823 | 3.41468138 | 0.00063857 | 0.0079713 | WT_active |
| 1405 | chr3 | 169966635 | 169998373 | ENSG00000008 | SEC62 | 7095 | 6.71894418 | 0.25716131 | 0.09032461 | 2.84707909 | 0.00441224 | 0.03941492 | WT_active |
| 1406 | chr22 | 37180167 | 37199385 | ENSG00000133 | C1QTNF6 | 114904 | 5.693327 | 0.25914881 | 0.09633119 | 2.69018588 | 0.00714122 | 0.05707436 | WT_active |
| 1407 | chr8 | 52534037 | 52565507 | ENSG00000196 | FAM150A | 389658 | 7.80152652 | 0.25937227 | 0.08278798 | 3.13297026 | 0.00173047 | 0.01841322 | WT_active |
| 1408 | chr3 | 123968521 | 123992178 | ENSG00000065 | ROPN1 | 54763 | 6.84898138 | 0.26037652 | 0.08783078 | 2.9645246 | 0.00303151 | 0.02917778 | WT_active |
| 1409 | chr6 | 46652915 | 46678193 | ENSG00000153 | SLC25A27 | 9481 | 6.45080856 | 0.26343906 | 0.09402706 | 2.80173669 | 0.00508283 | 0.04369225 | WT_active |
| 1410 | chr5 | 163460203 | 163491945 | ENSG00000072 | HMMR | 3161 | 6.1284133 | 0.2642492 | 0.09579608 | 2.75845523 | 0.00580753 | 0.04859716 | WT_active |
| 1411 | chr20 | 45607939 | 45631268 | ENSG00000180 | WFDC9 | 259240 | 6.87914703 | 0.26431875 | 0.08315256 | 3.17872063 | 0.00147927 | 0.01612204 | WT_active |
| 1412 | chrX | 49303669 | 49319844 | ENSG00000215 | GAGE10 | 102724473 | 5.08240456 | 0.26755709 | 0.10864654 | 2.46263802 | 0.01379191 | 0.09348929 | WT_active |
| 1413 | chr9 | 120815496 | 120842984 | ENSG00000095 | PSMD5 | 5711 | 6.84966986 | 0.26788324 | 0.09171422 | 2.92084755 | 0.00349081 | 0.03280994 | WT_active |
| 1414 | chr16 | 16232495 | 16294814 | ENSG00000103 | NOMO3 | 408050 | 5.65012806 | 0.26856116 | 0.09273863 | 2.89589318 | 0.00378081 | 0.03512354 | WT_active |
| 1415 | chr8 | 95244919 | 95269201 | ENSG00000156 | C8orf37 | 157657 | 5.50937617 | 0.27181671 | 0.09799335 | 2.77382823 | 0.00554009 | 0.04676642 | WT_active |
| 1416 | chr7 | 12211241 | 12243367 | ENSG00000106 | TMEM106B | 54664 | 5.89500313 | 0.27206219 | 0.10068737 | 2.70204884 | 0.00689136 | 0.05544625 | WT_active |
| 1417 | chr8 | 41261958 | 41309497 | ENSG00000104 | SFRP1 | 6422 | 13.3948982 | 0.27271413 | 0.06914807 | 3.94391543 | 8.02E-05 | 0.00123348 | WT_active |
| 1418 | chr12 | 75391070 | 75432688 | ENSG00000180 | GLIPR1L2 | 144321 | 9.07135192 | 0.2765837 | 0.07415899 | 3.72960461 | 0.00019178 | 0.00275407 | WT_active |
| 1419 | chr15 | 43330674 | 43354555 | ENSG00000168 | ADAL | 161823 | 6.46651522 | 0.27774481 | 0.0889341 | 3.1230407 | 0.00178993 | 0.01889254 | WT_active |
| 1420 | chr12 | 65851340 | 65882167 | ENSG00000197 | RP11-366L20.2 | 100129940 | 7.08197649 | 0.28126607 | 0.08840696 | 3.18149233 | 0.00146518 | 0.01600381 | WT_active |
| 1421 | chr17 | 56834099 | 56869567 | ENSG00000153 | DGKE | 8526 | 7.84562262 | 0.28490935 | 0.08229342 | 3.46211594 | 0.00053595 | 0.0068565 | WT_active |
| 1422 | chr6 | 7590199 | 7611967 | ENSG00000168 | SNRNP48 | 154007 | 4.8012042 | 0.28547928 | 0.11237052 | 2.54051755 | 0.01106885 | 0.07908361 | WT_active |
| 1423 | chr5 | 100535305 | 100586741 | ENSG00000174 | FAM174A | 345757 | 11.8365147 | 0.28976569 | 0.06808766 | 4.25577402 | 2.08E-05 | 0.00035673 | WT_active |
| 1424 | chr4 | 182890060 | 182917936 | ENSG00000129 | DCTD | 1635 | 7.9512079 | 0.29090203 | 0.08735505 | 3.33011114 | 0.00086811 | 0.01027624 | WT_active |
| 1425 | chr4 | 169729465 | 169757953 | ENSG00000056 | C4orf27 | 54969 | 7.10432457 | 0.29120747 | 0.087957 | 3.31079362 | 0.00093032 | 0.01090514 | WT_active |
| 1426 | chr3 | 42784298 | 42804531 | ENSG00000181 | HIGD1A | 25994 | 3.89843093 | 0.29217013 | 0.117942 | 2.47723559 | 0.01324045 | 0.0906892 | WT_active |
| 1427 | chr1 | 172532349 | 172611833 | ENSG00000094 | SUCO | 51430 | 16.2232452 | 0.29639572 | 0.05760106 | 5.14566433 | 2.67E-07 | 5.35E-06 | WT_active |
| 1428 | chr4 | 75479037 | 75514764 | ENSG00000163 | RCHY1 | 25898 | 7.21188539 | 0.29719548 | 0.08829596 | 3.36590117 | 0.00076294 | 0.00914643 | WT_active |
| 1429 | chr4 | 51993702 | 52017620 | ENSG00000188 | LRRC66 | 339977 | 6.50315431 | 0.29987213 | 0.08644314 | 3.46901025 | 0.00052238 | 0.00671272 | WT_active |
| 1430 | chr14 | 30874514 | 30895065 | ENSG00000100 | COCH | 1690 | 4.84823037 | 0.30682939 | 0.10635194 | 2.88503796 | 0.00391366 | 0.03600971 | WT_active |
| 1431 | chr12 | 76858715 | 76879060 | ENSG00000175 | CSRP2 | 1466 | 6.0741631 | 0.30731359 | 0.0987167 | 3.11308619 | 0.00185142 | 0.01937138 | WT_active |
| 1432 | chr12 | 9827481 | 9845007 | ENSG00000150 | KLRF1 | 51348 | 4.24022311 | 0.30840893 | 0.11580208 | 2.66324165 | 0.00773918 | 0.06074774 | WT_active |
| 1433 | chr15 | 72398273 | 72408029 | ENSG00000187 | TMEM202 | 338949 | 3.4843083 | 0.32171205 | 0.12691379 | 2.53488644 | 0.01124838 | 0.08000943 | WT_active |
| 1434 | chr19 | 57697367 | 57709194 | ENSG00000179 | ZNF154 | 7710 | 3.92342118 | 0.32206998 | 0.11413754 | 2.82177088 | 0.00477593 | 0.04183435 | WT_active |
| 1435 | chr6 | 112087599 | 112102790 | ENSG00000203 | FAM229B | 619208 | 3.9744009 | 0.3253543 | 0.12352612 | 2.63389067 | 0.00844127 | 0.06464843 | WT_active |
| 1436 | chr17 | 47200446 | 47223679 | ENSG00000198 | MYL4 | 4635 | 7.43968789 | 0.32784307 | 0.09473074 | 3.46078871 | 0.0005386 | 0.00688428 | WT_active |
| 1437 | chr11 | 167784 | 207428 | ENSG00000177 | BET1L | 51272 | 9.66181803 | 0.33151044 | 0.0798548 | 4.15141547 | 3.30E-05 | 0.00054697 | WT_active |
| 1438 | chrX | 55143102 | 55161310 | ENSG00000182 | FAM104B | 90736 | 4.65955866 | 0.34023942 | 0.11502625 | 2.95792855 | 0.00309714 | 0.0296708 | WT_active |
| 1439 | chr20 | 31465506 | 31476757 | ENSG00000180 | DEFB124 | 245937 | 3.28370533 | 0.34433854 | 0.13826897 | 2.490353 | 0.01276163 | 0.08777286 | WT_active |
| 1440 | chr1 | 65420587 | 65436007 | ENSG00000213 | LEPROT | 54741 | 3.32436736 | 0.34810644 | 0.13856132 | 2.51229164 | 0.01199499 | 0.08393302 | WT_active |
| 1441 | chrX | 17737449 | 17754988 | ENSG00000047 | SCML1 | 6322 | 4.29802861 | 0.3586126 | 0.11219551 | 3.19631859 | 0.00139193 | 0.01534319 | WT_active |
| 1442 | chr1 | 95234155 | 95247225 | ENSG00000122 | RWDD3 | 25950 | 3.28097919 | 0.37053643 | 0.13056243 | 2.83800199 | 0.00453969 | 0.04029772 | WT_active |
| 1443 | chr21 | 30480048 | 30491985 | ENSG00000244 | KRTAP19-3 | 337970 | 2.73167943 | 0.37757462 | 0.1463105 | 2.58063926 | 0.00986176 | 0.07273534 | WT_active |
| 1444 | chr1 | 100719742 | 100739045 | ENSG00000162 | VCAM1 | 7412 | 5.63588537 | 0.38134433 | 0.10067036 | 3.78804956 | 0.00015183 | 0.00222701 | WT_active |
| 1445 | chr1 | 113878168 | 113887547 | ENSG00000188 | BCL2L15 | 440603 | 2.56153285 | 0.38469813 | 0.15139982 | 2.54094179 | 0.01105543 | 0.07903556 | WT_active |
| 1446 | chr16 | 21639537 | 21652660 | ENSG00000140 | IGSF6 | 10261 | 3.2596911 | 0.38891668 | 0.13025919 | 2.98571399 | 0.00282917 | 0.02743173 | WT_active |
| 1447 | chr8 | 63015079 | 63039171 | ENSG00000137 | GGH | 8836 | 5.81756621 | 0.41076925 | 0.10290565 | 3.99170728 | 6.56E-05 | 0.00102471 | WT_active |
| 1448 | chr17 | 6643315 | 6651634 | ENSG00000108 | MED31 | 51003 | 2.43963994 | 0.43655463 | 0.16555467 | 2.63692133 | 0.00836622 | 0.06424056 | WT_active |
| 1449 | chr6 | 8413068 | 8435483 | ENSG00000124 | SLC35B3 | 51000 | 4.15214255 | 0.44821927 | 0.1070238 | 4.18803368 | 2.81E-05 | 0.00047119 | WT_active |
| 1450 | chr3 | 48293264 | 48301685 | ENSG00000172 | NME6 | 10201 | 2.62138766 | 0.45361746 | 0.14732245 | 3.07907889 | 0.00207642 | 0.02126263 | WT_active |
| 1451 | chrX | 49687450 | 49695993 | ENSG00000068 | PAGE1 | 8712 | 2.22244991 | 0.45939412 | 0.16835616 | 2.72870388 | 0.00635838 | 0.05223651 | WT_active |
| 1452 | chr6 | 31718594 | 31726714 | ENSG00000204 | C6orf25 | 80739 | 3.27717307 | 0.47492773 | 0.13837744 | 3.43211816 | 0.00059889 | 0.00753468 | WT_active |
| 1453 | chr4 | 41981696 | 41986467 | ENSG00000182 | DCAF4L1 | 285429 | 1.63136018 | 0.51350767 | 0.19304415 | 2.66005295 | 0.00781284 | 0.06112072 | WT_active |
| 1454 | chr17 | 41420753 | 41424523 | ENSG00000108 | KRT37 | 8688 | 1.72293748 | 0.6155535 | 0.19894739 | 3.09405162 | 0.00197443 | 0.02036291 | WT_active |

**Table S2C. List of Gene Sets over-represented in C7 (615 genes inferred, using scNOVA, to be upregulated compared to RPE-1).**

| Term | Overlap | P-value | Adjusted P-valu | Odds Ratio | Combined Scor | Genes | GeneSets |
|---|---|---|---|---|---|---|---|
| Carcinoma | 408/11318 | 3.34E-07 | 2.20E-04 | 1.53110429 | 22.8327114 | CYFIP1;ANKLE2 | Jensen_DISEASES |
| Williams-Beuren syndrome | 8/45 | 6.03E-05 | 0.01982378 | 6.89184737 | 66.9676037 | CLIP2;MLXIPL;V | Jensen_DISEASES |
| Epithelial Mesenchymal Transition | 17/200 | 1.57E-04 | 0.00770401 | 2.98293035 | 26.1240111 | PDGFRB;TPM4; | MSigDB_Hallmark_2020 |
| Mitotic Spindle | 16/199 | 4.53E-04 | 0.00782543 | 2.80277694 | 21.5781036 | TRIO;MYO9B;A | MSigDB_Hallmark_2020 |
| Apical Junction | 16/200 | 4.79E-04 | 0.00782543 | 2.78739929 | 21.3057222 | ACTN1;ATP1A3; | MSigDB_Hallmark_2020 |
| Reactive Oxygen Species Pathway | 7/49 | 7.03E-04 | 0.0086091 | 5.30235746 | 38.4975631 | NQO1;G6PD;AE | MSigDB_Hallmark_2020 |
| Cholesterol Homeostasis | 8/74 | 0.00193287 | 0.01894212 | 3.85782038 | 24.1065542 | FADS2;ATXN2;N | MSigDB_Hallmark_2020 |
| Glycolysis | 14/200 | 0.00366525 | 0.02993284 | 2.40446925 | 13.4863315 | G6PD;IDUA;BIK | MSigDB_Hallmark_2020 |

**Table S2D. List of genes predicted from scNOVA's altered gene activity inference module comparing cells with chr22q 11.2 deletion and without deletion in the LCL sample panel.**

| Rank | chr | str | end | name | Symbol | EntrezID | baseMean | log2FoldChang | lfcSE | stat | pvalue | padj | NO | Gene activity in SV subclone |
|---|---|---|---|---|---|---|---|---|---|---|---|---|---|---|
| 1 | chr14 | 61570540 | 61658696 | ENSG00000232 | FLJ22447 | 400221 | 21.4458239 | -0.2015601 | 0.0384518 | -5.2418893 | 1.59E-07 | 0.00082546 | Down | Up |

**Table S2E. List of genes inferred to be changing in activity when comparing major and minor clone in NA20509**

| Rank | chr | start | end | name | Symnbol | EntrezID | baseMean | log2FoldChang | lfcSE | stat | pvalue | padj | NO | Gene activity in SV subclone |
|---|---|---|---|---|---|---|---|---|---|---|---|---|---|---|
| 1 | chr13 | 44988529 | 44990529 | ENSG00000133 | GPALPP1 | 55425 | 10.9209219 | -0.8623882 | 0.22839307 | -3.7758948 | 0.00015943 | 0.02009799 | Down | Up |
| 2 | chr22 | 43811337 | 43813337 | ENSG00000186 | EFCAB6 | 64800 | 67.9191792 | 0.43916485 | 0.11949352 | 3.67521891 | 0.00023765 | 0.02740761 | Up | Down |
| 3 | chr20 | 13783950 | 13785950 | ENSG00000101 | NDUFAF5 | 79133 | 8.66487265 | -0.9634579 | 0.26676484 | -3.6116375 | 0.00030427 | 0.03332559 | Down | Up |
| 4 | chr20 | 35225654 | 35227654 | ENSG00000125 | MMP24 | 10893 | 14.9532227 | -0.696152 | 0.19501293 | -3.5697738 | 0.00035729 | 0.03654129 | Down | Up |
| 5 | chr18 | 157383 | 159383 | ENSG00000101 | USP14 | 9097 | 10.6027307 | -0.81962 | 0.23556093 | -3.4794395 | 0.00050246 | 0.04567979 | Down | Up |
| 6 | chr7 | 725701 | 727701 | ENSG00000164 | HEATR2 | 54919 | 13.0383102 | 0.95813905 | 0.27585678 | 3.47332061 | 0.00051406 | 0.04567979 | Up | Down |
| 7 | chr14 | 50531509 | 50533509 | ENSG00000198 | ATL1 | 51062 | 22.6920837 | 0.64497177 | 0.1939958 | 3.32466863 | 0.00088524 | 0.0678975 | Up | Down |
| 8 | chr11 | 57323952 | 57325952 | ENSG00000149 | TNKS1BP1 | 85456 | 6.88363494 | -1.0164372 | 0.31298533 | -3.2475555 | 0.00116401 | 0.08194173 | Down | Up |
| 9 | chr14 | 64504213 | 64506213 | ENSG00000089 | ZBTB25 | 7597 | 12.5955426 | 0.88624292 | 0.2721481 | 3.25647297 | 0.00112806 | 0.08194173 | Up | Down |
| 10 | chr3 | 3125916 | 3127916 | ENSG00000072 | TRNT1 | 51095 | 5.68244872 | 1.36372788 | 0.41918843 | 3.25325744 | 0.0011409 | 0.08194173 | Up | Down |
| 11 | chr4 | 78550519 | 78552519 | ENSG00000138 | ANXA3 | 306 | 16.0139543 | 0.80519337 | 0.24787407 | 3.24839691 | 0.00116057 | 0.08194173 | Up | Down |
| 12 | chr7 | 139339359 | 139341359 | ENSG00000146 | C7orf55-LUC7L2 | 100996928 | 15.1518153 | -0.6210448 | 0.19229491 | -3.2296475 | 0.00123943 | 0.08604871 | Down | Up |
| 13 | chr10 | 96358365 | 96360365 | ENSG00000197 | OPALIN | 93377 | 4.62081183 | 1.55397757 | 0.48493387 | 3.20451443 | 0.00135291 | 0.08800514 | Up | Down |
| 14 | chr17 | 69326244 | 69328244 | ENSG00000154 | ABCA5 | 23461 | 14.9728287 | 0.70466263 | 0.2194556 | 3.21095761 | 0.00132293 | 0.08800514 | Up | Down |
| 15 | chr7 | 139340360 | 139342360 | ENSG00000269 | LUC7L2 | 51631 | 14.74325 | -0.6187392 | 0.19351367 | -3.1973928 | 0.00138676 | 0.08895775 | Down | Up |
| 16 | chr3 | 51941345 | 51943345 | ENSG00000041 | PARP3 | 10039 | 1.46699042 | 3.23372852 | 1.01687834 | 3.18005447 | 0.00147247 | 0.09227218 | Up | Down |
| 17 | chr11 | 14642723 | 14644723 | ENSG00000152 | PDE3B | 5140 | 52.8484989 | 0.37224055 | 0.1174921 | 3.16821769 | 0.00153377 | 0.09447477 | Up | Down |
| 18 | chr6 | 42878618 | 42880618 | ENSG00000146 | RPL7L1 | 285855 | 1.61670707 | 2.68655646 | 0.85338455 | 3.14811939 | 0.00164325 | 0.09896903 | Up | Down |

**Table S2F. List of genes inferred to be changing in activity when comparing the chr19 deletion clone with the remainder of cells in NA12878, using scNOVA**

| Rank | chr | str | end | name | Symbol | EntrezID | baseMean | log2FoldChai | lfcSE | stat | pvalue | padj | NO | Gene activity in SV subclone |
|---|---|---|---|---|---|---|---|---|---|---|---|---|---|---|
| 1 | chr8 | 139730343 | 140458579 | ENSG000001 | TRAPPC9 | 83696 | 344.3894 | 0.383692 | 0.063958 | -5.999158 | 1.98E-09 | 1.39E-05 | Up | Down |
| 2 | chr18 | 10666483 | 11148762 | ENSG000001 | PIEZO2 | 63895 | 151.1817 | 0.248563 | 0.051708 | -4.807066 | 1.53E-06 | 0.004294 | Up | Down |
| 3 | chr10 | 49455368 | 49539538 | ENSG000002 | ERCC6 | 2074 | 30.98761 | -0.417368 | 0.100995 | 4.132574 | 0.0000359 | 0.062862 | Down | Up |

**Table S2G. List of genes inferred to be changing in activity in a haplotype-specific manner in AML_1**

| | | | | | | | Haplotype comparison SV/WT (before Tr breakpoint), LRT, GLMfit | | | | | | | |
|---|---|---|---|---|---|---|---|---|---|---|---|---|---|---|
| **Rank** | **chr** | **str** | **end** | **name** | **Symbol** | **EntrezID** | **median_wt** | **median_sv** | **log2FC** | **pvalue** | **median_all** | **var** | **p.adjust** | **Active_haplotype** |
| 1 | chr7 | 75027122 | 75074228 | ENSG00000274523 | WBSCR16 | 81554 | 0 | 4.8877396 | 4.8877396 | 2.66E-29 | 2.49002288 | 17.4622034 | 1.84E-26 | H1 |
| 2 | chr19 | 16134028 | 16158575 | ENSG00000196684 | HSH2D | 84941 | 3.54041664 | 0 | -3.5404166 | 8.06E-08 | 2.64833969 | 7.67723212 | 4.59E-05 | H2 |
| 3 | chr11 | 62856102 | 62888875 | ENSG00000168003 | SLC3A2 | 6520 | 3.43575537 | 2.52769262 | -0.9080627 | 6.03E-06 | 2.94285204 | 7.39055498 | 0.00324481 | H2 |
| 4 | chr19 | 43727992 | 43754990 | ENSG00000105771 | SMG9 | 56006 | 4.01867707 | 2.61819401 | -1.4004831 | 3.82E-05 | 3.30363666 | 7.99127751 | 0.01947396 | H2 |
| 5 | chr9 | 98069275 | 98119212 | ENSG00000106785 | TRIM14 | 9830 | 4.17028923 | 3.08403091 | -1.0862583 | 7.71E-05 | 3.66776554 | 11.1088067 | 0.03556146 | H2 |
| 6 | chr9 | 35792154 | 35809732 | ENSG00000159899 | NPR2 | 4882 | 1.20352277 | 3.67622743 | 2.47270466 | 9.37E-05 | 2.85246772 | 8.16583713 | 0.04125355 | H1 |
| 7 | chr20 | 44885676 | 44908532 | ENSG00000166913 | YWHAB | 7529 | 3.63134811 | 2.78402348 | -0.8473246 | 0.00012905 | 3.17642964 | 6.73708085 | 0.05208324 | H2 |
| 8 | chr2 | 73233420 | 73253021 | ENSG00000135624 | CCT7 | 10574 | 0 | 4.14632713 | 4.14632713 | 0.00015163 | 2.80499074 | 10.3257373 | 0.05874675 | H1 |
| 9 | chr8 | 67173511 | 67343677 | ENSG00000066777 | ARFGEF1 | 10565 | 5.31852336 | 5.70301223 | 0.38448887 | 0.00022656 | 5.61964492 | 20.4857205 | 0.0812781 | WT |
| 10 | chr11 | 57484534 | 57515786 | ENSG00000149150 | SLC43A1 | 8501 | 4.25630487 | 2.59720738 | -1.6590975 | 0.00026887 | 3.89305326 | 10.3784091 | 0.08980124 | H2 |
| 11 | chr8 | 91954967 | 92103286 | ENSG00000079102 | RUNX1T1 | 862 | 5.64619896 | 4.83459987 | -0.8115991 | 0.00026557 | 5.25926698 | 19.6676662 | 0.08980124 | Tr |

**Table S2H. List of genes inferred to be changing in activity when analyzing intra-sample heterogeneity in CLL_24 cells, using scNOVA (i.e. performing an analysis of cells with vs. cells without a Del event intersecting th**

| Rank | chr | start | end | name | gname | EntrezID | baseMean | log2FoldChang | lfcSE | stat | pvalue | padj | NO | Gene activity in SV subclone |
|---|---|---|---|---|---|---|---|---|---|---|---|---|---|---|
| 1 | chr19 | 10717079 | 10719079 | ENSG00000079805 | DNM2 | 1785 | 38.5923675 | -0.5218082 | 0.13321497 | -3.9170389 | 8.96E-05 | 0.05020899 | Down | Up |
| 2 | chr1 | 167165479 | 167167479 | ENSG00000143167 | GPA33 | 10223 | 53.6997046 | 0.29358657 | 0.07660393 | 3.83252629 | 0.00012683 | 0.06282095 | Up | Down |
| 3 | chr5 | 95554074 | 95556074 | ENSG00000164291 | ARSK | 153642 | 15.649635 | -0.65391 | 0.17064993 | -3.8318797 | 0.00012717 | 0.06282095 | Down | Up |

**Table S2I. List of genes inferred to be changing in activity when comparing major and minor clone in TALL-P1, using scNOVA**

| Rank | chr | str | end | name | Symbol | EntreÌD | baseMean | log2FoldChai | lfcSE | stat | pvalue | padj | NO | Gene activity in SV subclo | MYB_targets |
|---|---|---|---|---|---|---|---|---|---|---|---|---|---|---|---|
| 1 | chr6 | 166409364 | 166906451 | ENSG000000 | RPS6KA2 | 6196 | 148.4822072 | 0.304876051 | 0.0573489852 | 5.316154236 | 0.0000001059 | 0.0000264751 | Up | Down | 0 |
| 2 | chr6 | 170290703 | 170407065 | ENSG000001 | FAM120B | 84498 | 43.01382083 | 0.4480533412 | 0.0859640856 | 5.212099192 | 0.0000001867 | 0.000045531 | Up | Down | 0 |
| 3 | chrX | 46599252 | 46759172 | ENSG000000 | SLC9A7 | 84679 | 41.21641742 | -0.318699391 | 0.0784601065 | -4.06192912 | 0.0000486688 | 0.008594248 | Down | Up | 1 |
| 4 | chr4 | 40191053 | 40246967 | ENSG000001 | RHOH | 399 | 10.54336321 | -0.678797521 | 0.1762366283 | -3.851625671 | 0.0001173362 | 0.019383197 | Down | Up | 1 |
| 5 | chr2 | 102715840 | 102736888 | ENSG000001 | MFSD9 | 84804 | 5.073381786 | -0.912512582 | 0.2520561395 | -3.620275168 | 0.0002942898 | 0.044986819 | Down | Up | 0 |
| 6 | chr16 | 23836001 | 24220611 | ENSG000001 | PRKCB | 5579 | 97.89333621 | 0.2155832445 | 0.0584637265 | 3.687470113 | 0.0002264946 | 0.036246227 | Up | Down | 1 |
| 7 | chr15 | 75903859 | 75942510 | ENSG000001 | FBXO22 | 26263 | 8.597075467 | -0.681077474 | 0.1861445412 | -3.658863538 | 0.0002533361 | 0.0393131697 | Down | Up | 0 |
| 8 | chr9 | 136494444 | 136545862 | ENSG000001 | NOTCH1 | 4851 | 8.960143279 | -0.605297250 | 0.1758667021 | -3.441795649 | 0.0005778666 | 0.079979862 | Down | Up | 1 |
| 9 | chr1 | 45583846 | 45618904 | ENSG000001 | NASP | 4678 | 8.076289178 | -0.653576188 | 0.1845902887 | -3.540685659 | 0.0003990887 | 0.0592386504 | Down | Up | 1 |
| 10 | chr11 | 71787510 | 71801236 | ENSG000001 | FAM86C1 | 55199 | 4.242881353 | -0.952500341 | 0.2745965771 | -3.468726202 | 0.000522932 | 0.0743870907 | Down | Up | 0 |
| 11 | chr11 | 73787316 | 73865133 | ENSG000001 | MRPL48 | 51642 | 16.72959558 | -0.426647793 | 0.1232613206 | -3.461327459 | 0.0005375185 | 0.075414589 | Down | Up | 1 |
| 12 | chr9 | 126326849 | 126507041 | ENSG000001 | MVB12B | 89853 | 45.76202572 | -0.262798956 | 0.0766105581 | -3.430322965 | 0.0006028632 | 0.082327001 | Down | Up | 0 |
