## Supplementary Table S3 for "Haplotype-aware single-cell multiomics uncovers functional effects of somatic structural variation"

**Table S3A. List of SVs and linked molecular phenotypes analyzed in this study.**

| scNOVA locus ID | Sample | Chromosome | Start | End | Size (bp) | Size (Mb) | SV Class (scTRIP) | Singleton/Subclonal | CF (%) | Grouping | Is SCNA >10Mb (Y=1/N=0) | Associated to molecular phenotype by scNOVA (Y/N) | Reference |
|---|---|---|---|---|---|---|---|---|---|---|---|---|---|
| 1 | NA12329 | chr22 | 22700000 | 22900000 | 200000 | 0.2 | del_het | Subclonal | 66.1 | IGL_associated | 0 | Y (association with FLJ2247 expression, analysis across samples) | This study |
| 1 | NA12329 | chr22 | 22700000 | 22900000 | 200000 | 0.2 | del_hom | Singleton | 1.7 | IGL_associated | 0 | Y (association with FLJ2247 expression, analysis across samples) | This study |
| 1 | NA18939 | chr22 | 22300000 | 22900000 | 600000 | 0.6 | del_het | Subclonal | 9.8 | IGL_associated | 0 | Y (association with FLJ2247 expression, analysis across samples) | This study |
| 1 | NA18939 | chr22 | 22300000 | 22900000 | 600000 | 0.6 | del_hom | Subclonal | 14.6 | IGL_associated | 0 | Y (association with FLJ2247 expression, analysis across samples) | This study |
| 1 | NA19650 | chr22 | 2.20E+07 | 22900000 | 900000 | 0.9 | del_het | Singleton | 1.8 | IGL_associated | 0 | Y (association with FLJ2247 expression, analysis across samples) | This study |
| 1 | NA19650 | chr22 | 22300000 | 22900000 | 600000 | 0.6 | del_hom | Singleton | 1.8 | IGL_associated | 0 | Y (association with FLJ2247 expression, analysis across samples) | This study |
| 1 | NA19650 | chr22 | 22300000 | 22900000 | 600000 | 0.6 | del_het_h1 | Subclonal | 23.6 | IGL_associated | 0 | Y (association with FLJ2247 expression, analysis across samples) | This study |
| 1 | NA19650 | chr22 | 22300000 | 22900000 | 600000 | 0.6 | del_het_h2 | Subclonal | 3.6 | IGL_associated | 0 | Y (association with FLJ2247 expression, analysis across samples) | This study |
| 1 | NA19983 | chr22 | 22300000 | 22900000 | 600000 | 0.6 | del_het+del_hom | Subclonal | 4.4 | IGL_associated | 0 | Y (association with FLJ2247 expression, analysis across samples) | This study |
| 1 | NA19983 | chr22 | 22300000 | 22900000 | 600000 | 0.6 | del_hom | Subclonal | 4.4 | IGL_associated | 0 | Y (association with FLJ2247 expression, analysis across samples) | This study |
| 1 | NA19983 | chr22 | 22300000 | 22900000 | 600000 | 0.6 | del_het | Subclonal | 11.1 | IGL_associated | 0 | Y (association with FLJ2247 expression, analysis across samples) | This study |
| 1 | NA20847 | chr22 | 22700000 | 22900000 | 200000 | 0.2 | del_het_h1 | Singleton | 1.8 | IGL_associated | 0 | Y (association with FLJ2247 expression, analysis across samples) | This study |
| 1 | NA20847 | chr22 | 22700000 | 22900000 | 200000 | 0.2 | del_het_h2 | Singleton | 1.8 | IGL_associated | 0 | Y (association with FLJ2247 expression, analysis across samples) | This study |
| 1 | NA20847 | chr22 | 22700000 | 22900000 | 200000 | 0.2 | del_hom | Subclonal | 18.2 | IGL_associated | 0 | Y (association with FLJ2247 expression, analysis across samples) | This study |
| 1 | HG00096 | chr22 | 22700000 | 22900000 | 200000 | 0.2 | del_hom | Subclonal | 2.9 | IGL_associated | 0 | Y (association with FLJ2247 expression, analysis across samples) | This study |
| 1 | HG00096 | chr22 | 22300000 | 22900000 | 600000 | 0.6 | del_hom | Singleton | 1.4 | IGL_associated | 0 | Y (association with FLJ2247 expression, analysis across samples) | This study |
| 1 | HG00096 | chr22 | 22300000 | 22900000 | 600000 | 0.6 | del_het_h1 | Subclonal | 1.4 | IGL_associated | 0 | Y (association with FLJ2247 expression, analysis across samples) | This study |
| 1 | HG00096 | chr22 | 22300000 | 22900000 | 600000 | 0.6 | del_het_h2 | Subclonal | 7.2 | IGL_associated | 0 | Y (association with FLJ2247 expression, analysis across samples) | This study |
| 1 | HG00171 | chr22 | 22700000 | 22900000 | 200000 | 0.2 | del_hom | Subclonal | 8.2 | IGL_associated | 0 | Y (association with FLJ2247 expression, analysis across samples) | This study |
| 1 | HG00171 | chr22 | 22300000 | 22900000 | 600000 | 0.6 | del_hom | Subclonal | 4.1 | IGL_associated | 0 | Y (association with FLJ2247 expression, analysis across samples) | This study |
| 1 | HG00171 | chr22 | 22300000 | 22900000 | 600000 | 0.6 | del_het_h1 | Subclonal | 10.2 | IGL_associated | 0 | Y (association with FLJ2247 expression, analysis across samples) | This study |
| 1 | HG00171 | chr22 | 22300000 | 22900000 | 600000 | 0.6 | del_het_h2 | Subclonal | 16.3 | IGL_associated | 0 | Y (association with FLJ2247 expression, analysis across samples) | This study |
| 1 | HG00864 | chr22 | 22400000 | 22900000 | 500000 | 0.5 | del_het | Subclonal | 39.5 | IGL_associated | 0 | Y (association with FLJ2247 expression, analysis across samples) | This study |
| 1 | HG01114 | chr22 | 22400000 | 22900000 | 500000 | 0.5 | del_het_h1 | Subclonal | 3.9 | IGL_associated | 0 | Y (association with FLJ2247 expression, analysis across samples) | This study |
| 1 | HG01114 | chr22 | 22400000 | 22900000 | 500000 | 0.5 | del_het_h2 | Subclonal | 5.9 | IGL_associated | 0 | Y (association with FLJ2247 expression, analysis across samples) | This study |
| 1 | HG01114 | chr22 | 22400000 | 22900000 | 500000 | 0.5 | del_hom | Subclonal | 2 | IGL_associated | 0 | Y (association with FLJ2247 expression, analysis across samples) | This study |
| 1 | HG01596 | chr22 | 22300000 | 22900000 | 600000 | 0.6 | del_het_h1 | Subclonal | 7.5 | IGL_associated | 0 | Y (association with FLJ2247 expression, analysis across samples) | This study |
| 1 | HG01596 | chr22 | 22300000 | 22900000 | 600000 | 0.6 | del_het_h2 | Subclonal | 5.7 | IGL_associated | 0 | Y (association with FLJ2247 expression, analysis across samples) | This study |
| 1 | HG01596 | chr22 | 22300000 | 22900000 | 600000 | 0.6 | del_hom | Singleton | 1.9 | IGL_associated | 0 | Y (association with FLJ2247 expression, analysis across samples) | This study |
| 1 | HG02011 | chr22 | 22500000 | 22900000 | 400000 | 0.4 | del_het | Subclonal | 14.5 | IGL_associated | 0 | Y (association with FLJ2247 expression, analysis across samples) | This study |
| 1 | HG02587 | chr22 | 22400000 | 22900000 | 500000 | 0.5 | del_het | Subclonal | 88.9 | IGL_associated | 0 | Y (association with FLJ2247 expression, analysis across samples) | This study |
| 1 | HG02818 | chr22 | 22500000 | 22900000 | 400000 | 0.4 | del_het+del_hom | Subclonal | 21.7 | IGL_associated | 0 | Y (association with FLJ2247 expression, analysis across samples) | This study |
| 1 | HG03486 | chr22 | 22700000 | 22900000 | 200000 | 0.2 | del_het | Subclonal | 44.3 | IGL_associated | 0 | Y (association with FLJ2247 expression, analysis across samples) | This study |
| 1 | HG03486 | chr22 | 22500000 | 22900000 | 400000 | 0.4 | del_het | Singleton | 1.6 | IGL_associated | 0 | Y (association with FLJ2247 expression, analysis across samples) | This study |
| 1 | HG03683 | chr22 | 22300000 | 22900000 | 600000 | 0.6 | del_het | Subclonal | 8 | IGL_associated | 0 | Y (association with FLJ2247 expression, analysis across samples) | This study |
| 1 | HG03683 | chr22 | 22600000 | 22900000 | 300000 | 0.3 | del_het | Subclonal | 10 | IGL_associated | 0 | Y (association with FLJ2247 expression, analysis across samples) | This study |
| 1 | HG03683 | chr22 | 22600000 | 22900000 | 300000 | 0.3 | del_hom | Singleton | 2 | IGL_associated | 0 | Y (association with FLJ2247 expression, analysis across samples) | This study |
| 1 | HG03732 | chr22 | 22000000 | 22900000 | 900000 | 0.9 | del_het | Singleton | 2.1 | IGL_associated | 0 | Y (association with FLJ2247 expression, analysis across samples) | This study |
| 1 | HG03732 | chr22 | 22500000 | 22900000 | 400000 | 0.4 | del_het | Subclonal | 4.3 | IGL_associated | 0 | Y (association with FLJ2247 expression, analysis across samples) | This study |
| 1 | HG03732 | chr22 | 22300000 | 22900000 | 600000 | 0.6 | del_hom | Subclonal | 4.3 | IGL_associated | 0 | Y (association with FLJ2247 expression, analysis across samples) | This study |
| 1 | HG03732 | chr22 | 22300000 | 22900000 | 600000 | 0.6 | del_het_h1 | Singleton | 2.1 | IGL_associated | 0 | Y (association with FLJ2247 expression, analysis across samples) | This study |
| 1 | HG03732 | chr22 | 22300000 | 22900000 | 600000 | 0.6 | del_het_h2 | Subclonal | 8.5 | IGL_associated | 0 | Y (association with FLJ2247 expression, analysis across samples) | This study |
| 1 | NA12878 | chr22 | 22200000 | 22900000 | 700000 | 0.7 | del_het | Subclonal | 21.3 | IGL_associated | 0 | Y (association with FLJ2247 expression, analysis across samples) | This study |
| 1 | NA12878 | chr22 | 22200000 | 22900000 | 700000 | 0.7 | del_het+del_hom | Subclonal | 57.3 | IGL_associated | 0 | Y (association with FLJ2247 expression, analysis across samples) | This study |
| 2 | NA12878 | chr19 | 36500000 | 37000000 | 500000 | 0.5 | del_h2 | Subclonal | 21.3 | Mutually exclusive with chr22 Del | 0 | Y (association with ERCC6 expression) | This study |
| 3 | NA20509 | chr17 | 0 | 19200000 | 19200000 | 19.2 | del_h2 | Subclonal | 85.1 | BFB | 1 | Y (association with c-Myc/Max target deregulation) | This study |
| 3 | NA20509 | chr17 | 19200000 | 21700000 | 2500000 | 2.5 | idup_h2 | Subclonal | 85.1 | BFB | 1 | Y (association with c-Myc/Max target deregulation) | This study |
| 3 | NA20509 | chr5 | 132700000 | 181538259 | 48838259 | 48.8 | dup_h2 | Subclonal | 85.1 | - | 1 | Y (association with c-Myc/Max target deregulation) | This study |
| 3 | NA20509 | chr17 | 19200000 | 83257441 | 64057441 | 64.1 | translocation | Subclonal | 85.1 | t(17;5) | 0 | Y (association with c-Myc/Max target deregulation) | This study |
| 3 | NA20509 | chr5 | 132700000 | 181538259 | 48838259 | 48.8 | translocation | Subclonal | 85.1 | t(17;5) | - | - (refers to same rearrangement as row above) | This study |
| 4 | CLL_24 | chr10 | 101021000 | 103195000 | 2174000 | 2.2 | del_h2 | Subclonal | 2.3 | C1 | 0 | Y (WNT & MET signaling, from combined analysis of cells with minimal region deletion) | This study |
| 4 | CLL_24 | chr10 | 101343000 | 132793000 | 31450000 | 31.5 | del_h1 | Subclonal | 3.5 | C2 | 1 | Y (WNT & MET signaling, from combined analysis of cells with minimal region deletion) | This study |
| 4 | CLL_24 | chr10 | 86651000 | 88715000 | 2064000 | 2.1 | del_h1 | Subclonal | 2.3 | C3 | 0 | Y (WNT & MET signaling, from combined analysis of cells with minimal region deletion) | This study |
| 4 | CLL_24 | chr10 | 101589000 | 103479000 | 1890000 | 1.9 | del_h1 | Subclonal | 2.3 | C3 | 0 | Y (WNT & MET signaling, from combined analysis of cells with minimal region deletion) | This study |
| 4 | CLL_24 | chr10 | 100397000 | 103261000 | 2864000 | 2.9 | del_h2 | Singleton | 1.2 | S | 0 | Y (WNT & MET signaling, from combined analysis of cells with minimal region deletion) | This study |
| 4 | CLL_24 | chr10 | 100998000 | 110278000 | 9280000 | 9.3 | del_h2 | Singleton | 1.2 | S | 0 | Y (WNT & MET signaling, from combined analysis of cells with minimal region deletion) | This study |
| 4 | CLL_24 | chr10 | 100654000 | 114619000 | 13965000 | 14 | del_h2 | Singleton | 1.2 | S | 1 | Y (WNT & MET signaling, from combined analysis of cells with minimal region deletion) | This study |
| 4 | CLL_24 | chr10 | 97253000 | 104888000 | 7635000 | 7.6 | del_h2 | Singleton | 1.2 | S | 0 | Y (WNT & MET signaling, from combined analysis of cells with minimal region deletion) | This study |
| - | CLL_24 | chr11 | 81300000 | 135086622 | 53786622 | 53.8 | inv_dup_h2 | Singleton | 1.2 | - | 1 | N | This study |
| - | CLL_24 | chr19 | 18000000 | 19500000 | 1500000 | 1.5 | del_h2 | Singleton | 1.2 | - | 0 | N | This study |
| - | CLL_24 | chr22 | 36800000 | 42600000 | 5800000 | 5.8 | del_h1 | Singleton | 1.2 | - | 0 | N | This study |
| - | CLL_24 | chr6 | 98100000 | 170805979 | 72705979 | 72.7 | del_h1 | Singleton | 1.2 | - | 1 | N | This study |
| - | CLL_24 | chr7 | 60800000 | 159345973 | 98545973 | 98.5 | del | Singleton | 1.2 | - | 1 | N | This study |
| 5 | AML01 | chr8 | 92000000 | 145138636 | 53138636 | 53.1 | translocation | Clonal | 100 | t(8;21) | 0 | Y (haplotype-specific deregulated activity of several genes including RUNX1T1) | This study |
| 5 | AML01 | chr21 | 34900000 | 50818468 | 15918468 | 15.9 | translocation | Clonal | 100 | t(8;21) | - | - (refers to same rearrangement as row above) | This study |
| 6* | TALL_P1 | chr14 | 95800000 | 98400000 | 2600000 | 2.6 | inv_h2 | Clonal | 100 | - | 0 | Y (increased chromatin accessibility in cis of TCL1A) | PMID: 31873213 |
| 7* | TALL_P1 | chr6 | 83200000 | 170600000 | 87400000 | 87.4 | complex_h2 | Subclonal | 29.9 | - | 1 | Y (deregulation of c-Myb targets) | PMID: 31873213 |
| 8 | NA12329 | chr1 | 124932724 | 248956422 | 124023698 | 124 | dup_h1 | Singleton | 1.7 | - | 1 | - | This study |
| 9 | NA12329 | chr1 | 124932724 | 248956422 | 124023698 | 124 | del_h2 | Subclonal | 3.4 | - | 1 | - | This study |
| 10 | NA12329 | chr8 | 0 | 145138636 | 145138636 | 145.1 | del_h1 | Singleton | 1.7 | - | 1 | - | This study |
| 11 | NA12329 | chr9 | 101800000 | 138394717 | 36594717 | 36.6 | dup_h2 | Singleton | 1.7 | - | 1 | - | This study |
| 12 | NA12329 | chr17 | 0 | 83257441 | 83257441 | 83.3 | del_h2 | Singleton | 1.7 | - | 1 | - | This study |
| 13 | NA12329 | chr21 | 0 | 46709983 | 46709983 | 46.7 | del_h2 | Singleton | 1.7 | - | 1 | - | This study |
| 14 | NA12329 | chrX | 0 | 2800000 | 2800000 | 2.8 | del_h2 | Singleton | 1.7 | - | 0 | - | This study |
| 15 | NA18534 | chr1 | 124932724 | 248956422 | 124023698 | 124 | del_h2 | Singleton | 1.8 | - | 1 | - | This study |
| 16 | NA18534 | chr4 | 0 | 9000000 | 9000000 | 9 | del_h2 | Singleton | 1.8 | - | 0 | - | This study |

Table S3A. List of SVs and linked molecular phenotypes analyzed in this study.

| scNOVA locus ID | Sample | Chromosome | Start | End | Size (bp) | Size (Mb) | SV Class (scTRIP) | Singleton/Subclonal | CF (%) | Grouping | Is SCNA >10Mb (Y=1/N=0) | Associated to molecular phenotype by scNOVA (Y/N) | Reference |
|---|---|---|---|---|---|---|---|---|---|---|---|---|---|
| 17 | NA18534 | chr4 | 128300000 | 190214555 | 61914555 | 61.9 | del_h1 | Singleton | 1.8 | - | 1 | - | This study |
| 18 | NA18534 | chr11 | 0 | 135086622 | 135086622 | 135.1 | dup_h1 | Singleton | 1.8 | - | 1 | - | This study |
| 19 | NA18534 | chr17 | 62400000 | 83257441 | 20857441 | 20.9 | del_h1 | Singleton | 1.8 | - | 1 | - | This study |
| 20 | NA18939 | chr8 | 128100000 | 145138636 | 17038636 | 17 | dup_h1 | Singleton | 2.4 | - | 1 | - | This study |
| 21 | NA18939 | chr17 | 0 | 19000000 | 19000000 | 19 | dup_hom | Singleton | 2.4 | - | 1 | - | This study |
| 22 | NA18939 | chrX | 0 | 156040895 | 156040895 | 156 | dup_h1 | Singleton | 2.4 | - | 1 | - | This study |
| 23 | NA19650 | chr2 | 111600000 | 242193529 | 130593529 | 130.6 | del_h2 | Singleton | 1.8 | - | 1 | - | This study |
| 24 | NA19650 | chr4 | 103400000 | 190214555 | 86814555 | 86.8 | del_h2 | Singleton | 1.8 | - | 1 | - | This study |
| 25 | NA19650 | chr5 | 24000000 | 30300000 | 6300000 | 6.3 | del_h2 | Singleton | 1.8 | - | 0 | - | This study |
| 26 | NA19650 | chr13 | 65500000 | 114364328 | 48864328 | 48.9 | del_h2 | Singleton | 1.8 | - | 1 | - | This study |
| 27 | NA19983 | chr1 | 0 | 248956422 | 248956422 | 249 | dup_h1 | Singleton | 2.2 | - | 1 | - | This study |
| 28 | NA19983 | chr2 | 0 | 242193529 | 242193529 | 242.2 | dup_h2 | Singleton | 2.2 | - | 1 | - | This study |
| 29 | NA19983 | chr4 | 0 | 190214555 | 190214555 | 190.2 | dup_h2 | Singleton | 2.2 | - | 1 | - | This study |
| 30 | NA19983 | chr4 | 156400000 | 190214555 | 33814555 | 33.8 | dup_h1 | Singleton | 2.2 | - | 1 | - | This study |
| 31 | NA19983 | chr7 | 0 | 159345973 | 159345973 | 159.3 | dup_h2 | Singleton | 2.2 | - | 1 | - | This study |
| 32 | NA19983 | chr9 | 45518558 | 138394717 | 92876159 | 92.9 | del_h2 | Singleton | 2.2 | - | 1 | - | This study |
| 33 | NA19983 | chr9 | 79900000 | 81000000 | 1100000 | 1.1 | del_h1 | Subclonal | 6.7 | - | 0 | - | This study |
| 34 | NA19983 | chr14 | 0 | 107043718 | 107043718 | 107 | dup_h1 | Singleton | 2.2 | - | 1 | - | This study |
| 35 | NA19983 | chr15 | 0 | 101991189 | 101991189 | 102 | dup_h1 | Singleton | 2.2 | - | 1 | - | This study |
| 36 | NA19983 | chr16 | 36334460 | 90338345 | 54003885 | 54 | del_h2 | Singleton | 2.2 | - | 1 | - | This study |
| 37 | NA19983 | chrX | 79800000 | 156040895 | 76240895 | 76.2 | del_h1 | Singleton | 2.2 | - | 1 | - | This study |
| 38 | NA20509 | chr9 | 0 | 138394717 | 138394717 | 138.4 | dup_h2 | Singleton | 2.1 | - | 1 | - | This study |
| 39 | NA20509 | chr9 | 0 | 32000000 | 32000000 | 32 | dup_h2 | Singleton | 2.1 | - | 1 | - | This study |
| 40 | NA20509 | chr16 | 83900000 | 90338345 | 6438345 | 6.4 | del_h2 | Singleton | 2.1 | - | 0 | - | This study |
| 41 | NA20509 | chr16 | 36334460 | 90338345 | 54003885 | 54 | del_h1 | Singleton | 2.1 | - | 1 | - | This study |
| 42 | NA20509 | chr18 | 0 | 80373285 | 80373285 | 80.4 | del_h2 | Singleton | 2.1 | - | 1 | - | This study |
| 43 | NA20847 | chr2 | 0 | 92188145 | 92188145 | 92.2 | del_h2 | Singleton | 1.8 | - | 1 | - | This study |
| 44 | NA20847 | chr13 | 0 | 114364328 | 114364328 | 114.4 | dup_h2 | Singleton | 1.8 | - | 1 | - | This study |
| 45 | NA20847 | chr15 | 0 | 101991189 | 101991189 | 102 | dup_h1 | Singleton | 1.8 | - | 1 | - | This study |
| 46 | NA20847 | chr16 | 0 | 36311158 | 36311158 | 36.3 | del_h1 | Singleton | 1.8 | - | 1 | - | This study |
| 47 | HG00096 | chr1 | 0 | 14600000 | 14600000 | 14.6 | del_h1 | Singleton | 1.4 | - | 1 | - | This study |
| 48 | HG00096 | chr1 | 14600000 | 50300000 | 35700000 | 35.7 | dup_h2 | Singleton | 1.4 | - | 1 | - | This study |
| 49 | HG00096 | chr1 | 168900000 | 248956422 | 80056422 | 80.1 | del_h2 | Singleton | 1.4 | - | 1 | - | This study |
| 50 | HG00096 | chr9 | 45518558 | 138394717 | 92876159 | 92.9 | dup_h2 | Singleton | 1.4 | - | 1 | - | This study |
| 51 | HG00171 | chr1 | 124932724 | 248956422 | 124023698 | 124 | del_h1 | Singleton | 2 | - | 1 | - | This study |
| 52 | HG00171 | chr1 | 124932724 | 248956422 | 124023698 | 124 | del_hom | Singleton | 2 | - | 1 | - | This study |
| 53 | HG00171 | chr1 | 124932724 | 248956422 | 124023698 | 124 | dup_h2 | Singleton | 2 | - | 1 | - | This study |
| 54 | HG00171 | chr4 | 114200000 | 190214555 | 76014555 | 76 | dup_h1 | Singleton | 2 | - | 1 | - | This study |
| 55 | HG00171 | chr7 | 82800000 | 159345973 | 76545973 | 76.5 | del_h1 | Singleton | 2 | - | 1 | - | This study |
| 56 | HG00171 | chr12 | 94100000 | 110200000 | 16100000 | 16.1 | del_h1 | Singleton | 2 | - | 1 | - | This study |
| 57 | HG00171 | chr13 | 0 | 114364328 | 114364328 | 114.4 | del_h1 | Singleton | 2 | - | 1 | - | This study |
| 58 | HG00171 | chr16 | 36334460 | 90338345 | 54003885 | 54 | dup_hom | Singleton | 2 | - | 1 | - | This study |
| 59 | HG00171 | chr16 | 36334460 | 90338345 | 54003885 | 54 | del_h1 | Singleton | 2 | - | 1 | - | This study |
| 60 | HG00171 | chr16 | 36334460 | 90338345 | 54003885 | 54 | dup_h1 | Singleton | 2 | - | 1 | - | This study |
| 61 | HG00171 | chr17 | 0 | 14200000 | 14200000 | 14.2 | del_h1 | Singleton | 2 | - | 1 | - | This study |
| 62 | HG00171 | chrX | 0 | 156040895 | 156040895 | 156 | del_h2 | Subclonal | 4.1 | - | 1 | - | This study |
| 63 | HG00171 | chrX | 77000000 | 156040895 | 79040895 | 79 | dup_h2 | Singleton | 2 | - | 1 | - | This study |
| 64 | HG00864 | chr1 | 124932724 | 248956422 | 124023698 | 124 | idup_h1 | Singleton | 2.3 | - | 1 | - | This study |
| 65 | HG00864 | chr1 | 124932724 | 248956422 | 124023698 | 124 | dup_hom | Singleton | 2.3 | - | 1 | - | This study |
| 66 | HG00864 | chr16 | 36334460 | 90338345 | 54003885 | 54 | del_hom | Singleton | 2.3 | - | 1 | - | This study |
| 67 | HG00864 | chr16 | 36334460 | 90338345 | 54003885 | 54 | del_h2 | Singleton | 2.3 | - | 1 | - | This study |
| 68 | HG00864 | chrX | 0 | 156040895 | 156040895 | 156 | del_h2 | Subclonal | 18.6 | - | 1 | - | This study |
| 69 | HG00864 | chrX | 142500000 | 156040895 | 13540895 | 13.5 | del_h2 | Singleton | 2.3 | - | 1 | - | This study |
| 70 | HG01114 | chr16 | 83900000 | 90338345 | 6438345 | 6.4 | del_h1 | Singleton | 2 | - | 0 | - | This study |
| 71 | HG01114 | chr16 | 50800000 | 90338345 | 39538345 | 39.5 | dup_hom | Singleton | 2 | - | 1 | - | This study |
| 72 | HG01596 | chr4 | 65800000 | 73300000 | 7500000 | 7.5 | del_h1 | Singleton | 1.9 | - | 0 | - | This study |
| 73 | HG01596 | chr6 | 69500000 | 170805979 | 101305979 | 101.3 | del_h2 | Singleton | 1.9 | - | 1 | - | This study |
| 74 | HG01596 | chr16 | 56300000 | 90338345 | 34038345 | 34 | del_h2 | Singleton | 1.9 | - | 1 | - | This study |
| 75 | HG02011 | chr1 | 124932724 | 248956422 | 124023698 | 124 | del_h1 | Subclonal | 3.6 | - | 1 | - | This study |
| 76 | HG02011 | chr1 | 124932724 | 248956422 | 124023698 | 124 | dup_h1 | Singleton | 1.8 | - | 1 | - | This study |
| 77 | HG02011 | chr11 | 72700000 | 73600000 | 900000 | 0.9 | del_h1 | Subclonal | 7.3 | - | 0 | - | This study |
| 78 | HG02011 | chr16 | 36334460 | 90338345 | 54003885 | 54 | del_h1 | Subclonal | 3.6 | - | 1 | - | This study |
| 79 | HG02011 | chr16 | 36334460 | 90338345 | 54003885 | 54 | dup_h2 | Singleton | 1.8 | - | 1 | - | This study |
| 80 | HG02011 | chr16 | 36334460 | 90338345 | 54003885 | 54 | dup_h1 | Singleton | 1.8 | - | 1 | - | This study |
| 81 | HG02492 | chr11 | 90100000 | 135086622 | 44986622 | 45 | dup_hom | Singleton | 3.3 | - | 1 | - | This study |
| 82 | HG02587 | chr1 | 124932724 | 248956422 | 124023698 | 124 | dup_h1 | Singleton | 1.9 | - | 1 | - | This study |
| 83 | HG02587 | chr3 | 93655574 | 198295559 | 104639985 | 104.6 | del_h2 | Singleton | 1.9 | - | 1 | - | This study |
| 84 | HG02587 | chr16 | 36334460 | 90338345 | 54003885 | 54 | del_h1 | Singleton | 1.9 | - | 1 | - | This study |
| 85 | HG02818 | chr1 | 124932724 | 248956422 | 124023698 | 124 | del_h2 | Singleton | 1.4 | - | 1 | - | This study |
| 86 | HG02818 | chr1 | 234600000 | 248956422 | 14356422 | 14.4 | del_h2 | Singleton | 1.4 | - | 1 | - | This study |
| 87 | HG02818 | chr1 | 124932724 | 248956422 | 124023698 | 124 | del_h1 | Singleton | 1.4 | - | 1 | - | This study |
| 88 | HG02818 | chr2 | 0 | 8000000 | 8000000 | 8 | del_h2 | Singleton | 1.4 | - | 0 | - | This study |
| 89 | HG02818 | chr3 | 0 | 86200000 | 86200000 | 86.2 | idup_h2 | Singleton | 1.4 | - | 1 | - | This study |
| 90 | HG02818 | chr4 | 163500000 | 190214555 | 26714555 | 26.7 | del_h2 | Singleton | 1.4 | - | 1 | - | This study |
| 91 | HG02818 | chr4 | 83800000 | 90100000 | 6300000 | 6.3 | del_h1 | Singleton | 1.4 | - | 0 | - | This study |
| 92 | HG02818 | chr4 | 90100000 | 190214555 | 100114555 | 100.1 | del_hom | Singleton | 1.4 | - | 1 | - | This study |

**Table S3A. List of SVs and linked molecular phenotypes analyzed in this study.**

| scNOVA locus ID | Sample | Chromosome | Start | End | Size (bp) | Size (Mb) | SV Class (scTRIP) | Singleton/Subclonal | CF (%) | Grouping | Is SCNA >10Mb (Y=1/N=0) | Associated to molecular phenotype by scNOVA (Y/N) | Reference |
|---|---|---|---|---|---|---|---|---|---|---|---|---|---|
| 93 | HG02818 | chr9 | 45518558 | 138394717 | 92876159 | 92.9 | idup_h1 | Singleton | 1.4 | - | 1 | - | This study |
| 94 | HG02818 | chr16 | 36334460 | 90338345 | 54003885 | 54 | del_h1 | Subclonal | 7.2 | - | 1 | - | This study |
| 95 | HG02818 | chr16 | 36334460 | 90338345 | 54003885 | 54 | dup_h2 | Singleton | 1.4 | - | 1 | - | This study |
| 96 | HG02818 | chr16 | 36334460 | 90338345 | 54003885 | 54 | del_h2 | Singleton | 1.4 | - | 1 | - | This study |
| 97 | HG02818 | chr19 | 0 | 4100000 | 4100000 | 4.1 | del_h2 | Singleton | 1.4 | - | 0 | - | This study |
| 98 | HG02818 | chrX | 0 | 2800000 | 2800000 | 2.8 | del_h2 | Singleton | 1.4 | - | 0 | - | This study |
| 99 | HG02818 | chrX | 110500000 | 156040895 | 45540895 | 45.5 | idup_h2 | Singleton | 1.4 | - | 1 | - | This study |
| 100 | HG03009 | chr1 | 124932724 | 248956422 | 124023698 | 124 | del_h1 | Subclonal | 7.3 | - | 1 | - | This study |
| 101 | HG03009 | chr1 | 124932724 | 248956422 | 124023698 | 124 | dup_h1 | Singleton | 1.8 | - | 1 | - | This study |
| 102 | HG03009 | chr2 | 230200000 | 242193529 | 11993529 | 12 | del_h2 | Singleton | 1.8 | - | 1 | - | This study |
| 103 | HG03009 | chr4 | 105800000 | 190214555 | 84414555 | 84.4 | dup_h1 | Singleton | 1.8 | - | 1 | - | This study |
| 104 | HG03009 | chr6 | 0 | 28600000 | 28600000 | 28.6 | del_h1 | Singleton | 1.8 | - | 1 | - | This study |
| 105 | HG03009 | chr9 | 68300000 | 106800000 | 38500000 | 38.5 | dup_h1 | Singleton | 1.8 | - | 1 | - | This study |
| 106 | HG03009 | chr11 | 114800000 | 135086622 | 20286622 | 20.3 | dup_h2 | Singleton | 1.8 | - | 1 | - | This study |
| 107 | HG03009 | chr16 | 36334460 | 90338345 | 54003885 | 54 | dup_h2 | Subclonal | 3.6 | - | 1 | - | This study |
| 108 | HG03009 | chr16 | 36334460 | 90338345 | 54003885 | 54 | dup_h1 | Subclonal | 5.5 | - | 1 | - | This study |
| 109 | HG03009 | chr16 | 36334460 | 90338345 | 54003885 | 54 | del_h1 | Subclonal | 3.6 | - | 1 | - | This study |
| 110 | HG03009 | chr16 | 36334460 | 90338345 | 54003885 | 54 | del_h2 | Singleton | 1.8 | - | 1 | - | This study |
| 111 | HG03065 | chr16 | 36334460 | 90338345 | 54003885 | 54 | del_h1 | Singleton | 1.8 | - | 1 | - | This study |
| 112 | HG03065 | chr16 | 36334460 | 90338345 | 54003885 | 54 | del_h2 | Subclonal | 5.4 | - | 1 | - | This study |
| 113 | HG03065 | chr16 | 36334460 | 90338345 | 54003885 | 54 | dup_h2 | Singleton | 1.8 | - | 1 | - | This study |
| 114 | HG03065 | chr18 | 0 | 15460899 | 15460899 | 15.5 | del_h1 | Singleton | 1.8 | - | 1 | - | This study |
| 115 | HG03125 | chr1 | 0 | 248956422 | 248956422 | 249 | del_h2 | Singleton | 1.8 | - | 1 | - | This study |
| 116 | HG03125 | chr2 | 124000000 | 242193529 | 118193529 | 118.2 | del_h1 | Singleton | 1.8 | - | 1 | - | This study |
| 117 | HG03125 | chr4 | 66900000 | 190214555 | 123314555 | 123.3 | del_h2 | Singleton | 1.8 | - | 1 | - | This study |
| 118 | HG03125 | chr7 | 0 | 58169653 | 58169653 | 58.2 | del_h2 | Singleton | 1.8 | - | 1 | - | This study |
| 119 | HG03125 | chr8 | 79400000 | 145138636 | 65738636 | 65.7 | del_h1 | Singleton | 1.8 | - | 1 | - | This study |
| 120 | HG03125 | chr8 | 0 | 21100000 | 21100000 | 21.1 | del_h1 | Singleton | 1.8 | - | 1 | - | This study |
| 121 | HG03125 | chr9 | 45518558 | 138394717 | 92876159 | 92.9 | del_h1 | Subclonal | 3.6 | - | 1 | - | This study |
| 122 | HG03125 | chr9 | 114700000 | 138394717 | 23694717 | 23.7 | dup_h2 | Singleton | 1.8 | - | 1 | - | This study |
| 123 | HG03125 | chr9 | 0 | 20800000 | 20800000 | 20.8 | idup_h1 | Singleton | 1.8 | - | 1 | - | This study |
| 124 | HG03125 | chr11 | 128300000 | 135086622 | 6786622 | 6.8 | idup_h1 | Singleton | 1.8 | - | 0 | - | This study |
| 125 | HG03125 | chr12 | 54500000 | 133275309 | 78775309 | 78.8 | idup_h2 | Singleton | 1.8 | - | 1 | - | This study |
| 126 | HG03125 | chr12 | 0 | 34769407 | 34769407 | 34.8 | idup_h1 | Singleton | 1.8 | - | 1 | - | This study |
| 127 | HG03125 | chr16 | 0 | 90338345 | 90338345 | 90.3 | del_h2 | Singleton | 1.8 | - | 1 | - | This study |
| 128 | HG03125 | chrX | 96600000 | 112800000 | 16200000 | 16.2 | del_h2 | Singleton | 1.8 | - | 1 | - | This study |
| 129 | HG03371 | chr1 | 0 | 248956422 | 248956422 | 249 | dup_h2 | Singleton | 1.6 | - | 1 | - | This study |
| 130 | HG03371 | chr3 | 0 | 198295559 | 198295559 | 198.3 | dup_h1 | Singleton | 1.6 | - | 1 | - | This study |
| 131 | HG03371 | chr6 | 0 | 170805979 | 170805979 | 170.8 | del_h2 | Singleton | 1.6 | - | 1 | - | This study |
| 132 | HG03371 | chr16 | 0 | 18300000 | 18300000 | 18.3 | del_h2 | Singleton | 1.6 | - | 1 | - | This study |
| 133 | HG03371 | chr16 | 36334460 | 90338345 | 54003885 | 54 | idup_h2 | Singleton | 1.6 | - | 1 | - | This study |
| 134 | HG03371 | chr19 | 0 | 58617616 | 58617616 | 58.6 | idup_h2 | Singleton | 1.6 | - | 1 | - | This study |
| 135 | HG03371 | chr21 | 0 | 46709983 | 46709983 | 46.7 | idup_h1 | Singleton | 1.6 | - | 1 | - | This study |
| 136 | HG03486 | chr1 | 232000000 | 248956422 | 16956422 | 17 | del_h1 | Singleton | 1.6 | - | 1 | - | This study |
| 137 | HG03486 | chr1 | 157600000 | 248956422 | 91356422 | 91.4 | idup_h1 | Singleton | 1.6 | - | 1 | - | This study |
| 138 | HG03486 | chr6 | 129700000 | 170805979 | 41105979 | 41.1 | del_h1 | Singleton | 1.6 | - | 1 | - | This study |
| 139 | HG03486 | chr9 | 68300000 | 104200000 | 35900000 | 35.9 | dup_h1 | Singleton | 1.6 | - | 1 | - | This study |
| 140 | HG03486 | chr9 | 104200000 | 138394717 | 34194717 | 34.2 | del_h2 | Singleton | 1.6 | - | 1 | - | This study |
| 141 | HG03486 | chr11 | 114600000 | 135086622 | 20486622 | 20.5 | del_h2 | Singleton | 1.6 | - | 1 | - | This study |
| 142 | HG03486 | chr12 | 65400000 | 133275309 | 67875309 | 67.9 | idup_h2 | Singleton | 1.6 | - | 1 | - | This study |
| 143 | HG03486 | chr13 | 82000000 | 114364328 | 32364328 | 32.4 | dup_h1 | Singleton | 1.6 | - | 1 | - | This study |
| 144 | HG03486 | chr16 | 36334460 | 90338345 | 54003885 | 54 | del_h2 | Singleton | 1.6 | - | 1 | - | This study |
| 145 | HG03486 | chr16 | 22800000 | 66600000 | 43800000 | 43.8 | idup_h2 | Singleton | 1.6 | - | 1 | - | This study |
| 146 | HG03683 | chr2 | 195000000 | 242193529 | 47193529 | 47.2 | del_h1 | Singleton | 2 | - | 1 | - | This study |
| 147 | HG03683 | chr8 | 0 | 44033744 | 44033744 | 44 | del_h1 | Singleton | 2 | - | 1 | - | This study |
| 148 | HG03683 | chr8 | 45877265 | 145138636 | 99261371 | 99.3 | dup_h2 | Singleton | 2 | - | 1 | - | This study |
| 149 | HG03683 | chr9 | 45518558 | 138394717 | 92876159 | 92.9 | del_h2 | Singleton | 2 | - | 1 | - | This study |
| 150 | HG03683 | chr9 | 45518558 | 138394717 | 92876159 | 92.9 | del_h1 | Singleton | 2 | - | 1 | - | This study |
| 151 | HG03683 | chr13 | 72000000 | 114364328 | 42364328 | 42.4 | del_h2 | Singleton | 2 | - | 1 | - | This study |
| 152 | HG03683 | chr15 | 74300000 | 101991189 | 27691189 | 27.7 | dup_h2 | Singleton | 2 | - | 1 | - | This study |
| 153 | HG03683 | chr16 | 36334460 | 90338345 | 54003885 | 54 | del_h2 | Singleton | 2 | - | 1 | - | This study |
| 154 | HG03683 | chr16 | 36334460 | 90338345 | 54003885 | 54 | dup_h1 | Singleton | 2 | - | 1 | - | This study |
| 155 | HG03683 | chr17 | 0 | 1800000 | 1800000 | 1.8 | del_h1 | Singleton | 2 | - | 0 | - | This study |
| 156 | HG03683 | chr17 | 1800000 | 21600000 | 19800000 | 19.8 | dup_h2 | Singleton | 2 | - | 1 | - | This study |
| 157 | HG03732 | chr2 | 0 | 23300000 | 23300000 | 23.3 | del_h2 | Singleton | 2.1 | - | 1 | - | This study |
| 158 | HG03732 | chr2 | 184500000 | 242193529 | 57693529 | 57.7 | del_h2 | Singleton | 2.1 | - | 1 | - | This study |
| 159 | HG03732 | chr4 | 0 | 27400000 | 27400000 | 27.4 | del_h1 | Singleton | 2.1 | - | 1 | - | This study |
| 160 | HG03732 | chr5 | 122100000 | 181538259 | 59438259 | 59.4 | dup_h1 | Singleton | 2.1 | - | 1 | - | This study |
| 161 | HG03732 | chr8 | 0 | 145138636 | 145138636 | 145.1 | del_h1 | Singleton | 2.1 | - | 1 | - | This study |
| 162 | HG03732 | chr16 | 36334460 | 90338345 | 54003885 | 54 | dup_h2 | Singleton | 2.1 | - | 1 | - | This study |

*For TALL P1, among several SVs reported in the original publication (PMID: 31873213), a somatic inversion and a chromothripsis event were associated with molecular phenotypes in this study.

Table S3B. List of SVs found in the 24 LCLs from HGSVC

| Cell | Chromosome | Start | End | Size (bp) | SV Class | Singleton/Sub | Sample | Copy Number estimation |
|---|---|---|---|---|---|---|---|---|
| GM12329x02PE20406 | chr22 | 22700000 | 22900000 | 200000 | del_hom | Singleton | NA12329 | 0 |
| GM12329x02PE20401 | chr22 | 22700000 | 22900000 | 200000 | del_h1 | Subclonal | NA12329 | 1 |
| GM12329x02PE20404 | chr22 | 22700000 | 22900000 | 200000 | del_h1 | Subclonal | NA12329 | 1 |
| GM12329x02PE20410 | chr22 | 22700000 | 22900000 | 200000 | del_h1 | Subclonal | NA12329 | 1 |
| GM12329x02PE20413 | chr22 | 22700000 | 22900000 | 200000 | del_h1 | Subclonal | NA12329 | 1 |
| GM12329x02PE20418 | chr22 | 22700000 | 22900000 | 200000 | del_h1 | Subclonal | NA12329 | 1 |
| GM12329x02PE20420 | chr22 | 22700000 | 22900000 | 200000 | del_h1 | Subclonal | NA12329 | 1 |
| GM12329x02PE20423 | chr22 | 22700000 | 22900000 | 200000 | del_h1 | Subclonal | NA12329 | 1 |
| GM12329x02PE20425 | chr22 | 22700000 | 22900000 | 200000 | del_h1 | Subclonal | NA12329 | 1 |
| GM12329x02PE20427 | chr22 | 22700000 | 22900000 | 200000 | del_h1 | Subclonal | NA12329 | 1 |
| GM12329x02PE20430 | chr22 | 22700000 | 22900000 | 200000 | del_h1 | Subclonal | NA12329 | 1 |
| GM12329x02PE20431 | chr22 | 22700000 | 22900000 | 200000 | del_h1 | Subclonal | NA12329 | 1 |
| GM12329x02PE20435 | chr22 | 22700000 | 22900000 | 200000 | del_h1 | Subclonal | NA12329 | 1 |
| GM12329x02PE20436 | chr22 | 22700000 | 22900000 | 200000 | del_h1 | Subclonal | NA12329 | 1 |
| GM12329x02PE20445 | chr22 | 22700000 | 22900000 | 200000 | del_h1 | Subclonal | NA12329 | 1 |
| GM12329x02PE20446 | chr22 | 22700000 | 22900000 | 200000 | del_h1 | Subclonal | NA12329 | 1 |
| GM12329x02PE20450 | chr22 | 22700000 | 22900000 | 200000 | del_h1 | Subclonal | NA12329 | 1 |
| GM12329x02PE20451 | chr22 | 22700000 | 22900000 | 200000 | del_h1 | Subclonal | NA12329 | 1 |
| GM12329x02PE20453 | chr22 | 22700000 | 22900000 | 200000 | del_h1 | Subclonal | NA12329 | 1 |
| GM12329x02PE20457 | chr22 | 22700000 | 22900000 | 200000 | del_h1 | Subclonal | NA12329 | 1 |
| GM12329x02PE20458 | chr22 | 22700000 | 22900000 | 200000 | del_h1 | Subclonal | NA12329 | 1 |
| GM12329x02PE20459 | chr22 | 22700000 | 22900000 | 200000 | del_h1 | Subclonal | NA12329 | 1 |
| GM12329x02PE20460 | chr22 | 22700000 | 22900000 | 200000 | del_h1 | Subclonal | NA12329 | 1 |
| GM12329x02PE20461 | chr22 | 22700000 | 22900000 | 200000 | del_h1 | Subclonal | NA12329 | 1 |
| GM12329x02PE20462 | chr22 | 22700000 | 22900000 | 200000 | del_h1 | Subclonal | NA12329 | 1 |
| GM12329x02PE20463 | chr22 | 22700000 | 22900000 | 200000 | del_h1 | Subclonal | NA12329 | 1 |
| GM12329x02PE20465 | chr22 | 22700000 | 22900000 | 200000 | del_h1 | Subclonal | NA12329 | 1 |
| GM12329x02PE20467 | chr22 | 22700000 | 22900000 | 200000 | del_h1 | Subclonal | NA12329 | 1 |
| GM12329x02PE20469 | chr22 | 22700000 | 22900000 | 200000 | del_h1 | Subclonal | NA12329 | 1 |
| GM12329x02PE20473 | chr22 | 22700000 | 22900000 | 200000 | del_h1 | Subclonal | NA12329 | 1 |
| GM12329x02PE20474 | chr22 | 22700000 | 22900000 | 200000 | del_h1 | Subclonal | NA12329 | 1 |
| GM12329x02PE20477 | chr22 | 22700000 | 22900000 | 200000 | del_h1 | Subclonal | NA12329 | 1 |
| GM12329x02PE20478 | chr22 | 22700000 | 22900000 | 200000 | del_h1 | Subclonal | NA12329 | 1 |
| GM12329x02PE20481 | chr22 | 22700000 | 22900000 | 200000 | del_h1 | Subclonal | NA12329 | 1 |

| | | | | | | | | |
|---|---|---|---|---|---|---|---|---|
| GM12329x02PE20484 | chr22 | 22700000 | 22900000 | 200000 | del_h1 | Subclonal | NA12329 | 1 |
| GM12329x02PE20486 | chr22 | 22700000 | 22900000 | 200000 | del_h1 | Subclonal | NA12329 | 1 |
| GM12329x02PE20487 | chr22 | 22700000 | 22900000 | 200000 | del_h1 | Subclonal | NA12329 | 1 |
| GM12329x02PE20489 | chr22 | 22700000 | 22900000 | 200000 | del_h1 | Subclonal | NA12329 | 1 |
| GM12329x02PE20491 | chr22 | 22700000 | 22900000 | 200000 | del_h1 | Subclonal | NA12329 | 1 |
| GM12329x02PE20493 | chr22 | 22700000 | 22900000 | 200000 | del_h1 | Subclonal | NA12329 | 1 |
| GM18939x02PE20413 | chr22 | 22300000 | 22900000 | 600000 | del_h2 | Subclonal | NA18939 | 1 |
| GM18939x02PE20442 | chr22 | 22300000 | 22900000 | 600000 | del_h2 | Subclonal | NA18939 | 1 |
| GM18939x02PE20458 | chr22 | 22300000 | 22900000 | 600000 | del_h2 | Subclonal | NA18939 | 1 |
| GM18939x02PE20478 | chr22 | 22300000 | 22900000 | 600000 | del_h2 | Subclonal | NA18939 | 1 |
| GM18939x02PE20401 | chr22 | 22300000 | 22900000 | 600000 | del_hom | Subclonal | NA18939 | 0 |
| GM18939x02PE20412 | chr22 | 22300000 | 22900000 | 600000 | del_hom | Subclonal | NA18939 | 0 |
| GM18939x02PE20416 | chr22 | 22300000 | 22900000 | 600000 | del_hom | Subclonal | NA18939 | 0 |
| GM18939x02PE20422 | chr22 | 22300000 | 22900000 | 600000 | del_hom | Subclonal | NA18939 | 0 |
| GM18939x02PE20449 | chr22 | 22300000 | 22900000 | 600000 | del_hom | Subclonal | NA18939 | 0 |
| GM18939x02PE20462 | chr22 | 22300000 | 22900000 | 600000 | del_hom | Subclonal | NA18939 | 0 |
| GM19650Ax02PE20524 | chr22 | 2.20E+07 | 22900000 | 900000 | del_h1 | Singleton | NA19650 | 1 |
| GM19650Ax02PE20526 | chr22 | 22300000 | 22900000 | 600000 | del_hom | Singleton | NA19650 | 0 |
| GM19650Ax02PE20501 | chr22 | 22300000 | 22900000 | 600000 | del_h1 | Subclonal | NA19650 | 1 |
| GM19650Ax02PE20506 | chr22 | 22300000 | 22900000 | 600000 | del_h1 | Subclonal | NA19650 | 1 |
| GM19650Ax02PE20511 | chr22 | 22300000 | 22900000 | 600000 | del_h1 | Subclonal | NA19650 | 1 |
| GM19650Ax02PE20541 | chr22 | 22300000 | 22900000 | 600000 | del_h1 | Subclonal | NA19650 | 1 |
| GM19650Ax02PE20544 | chr22 | 22300000 | 22900000 | 600000 | del_h1 | Subclonal | NA19650 | 1 |
| GM19650Ax02PE20547 | chr22 | 22300000 | 22900000 | 600000 | del_h1 | Subclonal | NA19650 | 1 |
| GM19650Ax02PE20548 | chr22 | 22300000 | 22900000 | 600000 | del_h1 | Subclonal | NA19650 | 1 |
| GM19650Ax02PE20550 | chr22 | 22300000 | 22900000 | 600000 | del_h1 | Subclonal | NA19650 | 1 |
| GM19650Ax02PE20553 | chr22 | 22300000 | 22900000 | 600000 | del_h1 | Subclonal | NA19650 | 1 |
| GM19650Ax02PE20563 | chr22 | 22300000 | 22900000 | 600000 | del_h1 | Subclonal | NA19650 | 1 |
| GM19650Ax02PE20568 | chr22 | 22300000 | 22900000 | 600000 | del_h1 | Subclonal | NA19650 | 1 |
| GM19650Ax02PE20573 | chr22 | 22300000 | 22900000 | 600000 | del_h1 | Subclonal | NA19650 | 1 |
| GM19650Ax02PE20582 | chr22 | 22300000 | 22900000 | 600000 | del_h1 | Subclonal | NA19650 | 1 |
| GM19650Ax02PE20536 | chr22 | 22300000 | 22900000 | 600000 | del_h2 | Subclonal | NA19650 | 1 |
| GM19650Ax02PE20580 | chr22 | 22300000 | 22900000 | 600000 | del_h2 | Subclonal | NA19650 | 1 |
| GM19983x02PE20453 | chr22 | 22300000 | 22900000 | 600000 | del_het+del_hom | Subclonal | NA19983 | 1,0 |
| GM19983x02PE20414 | chr22 | 22300000 | 22900000 | 600000 | del_het+del_hom | Subclonal | NA19983 | 1,0 |
| GM19983x02PE20412 | chr22 | 22300000 | 22900000 | 600000 | del_hom | Subclonal | NA19983 | 0 |
| GM19983x02PE20458 | chr22 | 22300000 | 22900000 | 600000 | del_hom | Subclonal | NA19983 | 0 |
| GM19983x02PE20434 | chr22 | 22300000 | 22900000 | 600000 | del_het | Subclonal | NA19983 | 1 |

| | | | | | | | | |
|---|---|---|---|---|---|---|---|---|
| GM19983x02PE20454 | chr22 | 22300000 | 22900000 | 600000 | del_het | Subclonal | NA19983 | 1 |
| GM19983x02PE20493 | chr22 | 22300000 | 22900000 | 600000 | del_het | Subclonal | NA19983 | 1 |
| GM19983x02PE20470 | chr22 | 22300000 | 22900000 | 600000 | del_het | Subclonal | NA19983 | 1 |
| GM19983x02PE20475 | chr22 | 22300000 | 22900000 | 600000 | del_het | Subclonal | NA19983 | 1 |
| GM20847Bx02PE20422 | chr22 | 22700000 | 22900000 | 200000 | del_het_h1 | Singleton | NA20847 | 1 |
| GM20847Bx02PE20437 | chr22 | 22700000 | 22900000 | 200000 | del_het_h2 | Singleton | NA20847 | 1 |
| GM20847Bx02PE20406 | chr22 | 22700000 | 22900000 | 200000 | del_hom | Subclonal | NA20847 | 0 |
| GM20847Bx02PE20419 | chr22 | 22700000 | 22900000 | 200000 | del_hom | Singleton | NA20847 | 0 |
| GM20847Bx02PE20423 | chr22 | 22700000 | 22900000 | 200000 | del_hom | Singleton | NA20847 | 0 |
| GM20847Bx02PE20426 | chr22 | 22700000 | 22900000 | 200000 | del_hom | Subclonal | NA20847 | 0 |
| GM20847Bx02PE20438 | chr22 | 22700000 | 22900000 | 200000 | del_hom | Singleton | NA20847 | 0 |
| GM20847Bx02PE20443 | chr22 | 22700000 | 22900000 | 200000 | del_hom | Singleton | NA20847 | 0 |
| GM20847Bx02PE20446 | chr22 | 22700000 | 22900000 | 200000 | del_hom | Subclonal | NA20847 | 0 |
| GM20847Bx02PE20447 | chr22 | 22700000 | 22900000 | 200000 | del_hom | Singleton | NA20847 | 0 |
| GM20847Bx02PE20457 | chr22 | 22700000 | 22900000 | 200000 | del_hom | Singleton | NA20847 | 0 |
| GM20847Bx02PE20470 | chr22 | 22700000 | 22900000 | 200000 | del_hom | Subclonal | NA20847 | 0 |
| HG00096x02PE20367 | chr22 | 22700000 | 22900000 | 200000 | del_hom | Subclonal | HG00096 | 0 |
| HG00096x02PE20377 | chr22 | 22700000 | 22900000 | 200000 | del_hom | Subclonal | HG00096 | 0 |
| HG00096x02PE20368 | chr22 | 22300000 | 22900000 | 600000 | del_hom | Singleton | HG00096 | 0 |
| HG00096x02PE20370 | chr22 | 22300000 | 22900000 | 600000 | del_h1 | Singleton | HG00096 | 1 |
| HG00096x02PE20307 | chr22 | 22300000 | 22900000 | 600000 | del_h2 | Subclonal | HG00096 | 1 |
| HG00096x02PE20330 | chr22 | 22300000 | 22900000 | 600000 | del_h2 | Subclonal | HG00096 | 1 |
| HG00096x02PE20342 | chr22 | 22300000 | 22900000 | 600000 | del_h2 | Subclonal | HG00096 | 1 |
| HG00096x02PE20344 | chr22 | 22300000 | 22900000 | 600000 | del_h2 | Subclonal | HG00096 | 1 |
| HG00096x02PE20373 | chr22 | 22300000 | 22900000 | 600000 | del_h2 | Subclonal | HG00096 | 1 |
| HG00171Ax02PE20420 | chr22 | 22700000 | 22900000 | 200000 | del_hom_small | Subclonal | HG00171 | 0 |
| HG00171Ax02PE20445 | chr22 | 22700000 | 22900000 | 200000 | del_hom_small | Subclonal | HG00171 | 0 |
| HG00171Ax02PE20457 | chr22 | 22700000 | 22900000 | 200000 | del_hom_small | Subclonal | HG00171 | 0 |
| HG00171Ax02PE20464 | chr22 | 22700000 | 22900000 | 200000 | del_hom_small | Subclonal | HG00171 | 0 |
| HG00171Ax02PE20436 | chr22 | 22300000 | 22900000 | 600000 | del_hom | Subclonal | HG00171 | 0 |
| HG00171Ax02PE20469 | chr22 | 22300000 | 22900000 | 600000 | del_hom | Subclonal | HG00171 | 0 |
| HG00171Ax02PE20405 | chr22 | 22300000 | 22900000 | 600000 | del_h1 | Subclonal | HG00171 | 1 |
| HG00171Ax02PE20415 | chr22 | 22300000 | 22900000 | 600000 | del_h1 | Subclonal | HG00171 | 1 |
| HG00171Ax02PE20431 | chr22 | 22300000 | 22900000 | 600000 | del_h1 | Subclonal | HG00171 | 1 |
| HG00171Ax02PE20458 | chr22 | 22300000 | 22900000 | 600000 | del_h1 | Subclonal | HG00171 | 1 |
| HG00171Ax02PE20468 | chr22 | 22300000 | 22900000 | 600000 | del_h1 | Subclonal | HG00171 | 1 |
| HG00171Ax02PE20416 | chr22 | 22300000 | 22900000 | 600000 | del_h2 | Subclonal | HG00171 | 1 |
| HG00171Ax02PE20417 | chr22 | 22300000 | 22900000 | 600000 | del_h2 | Subclonal | HG00171 | 1 |

| | | | | | | | | |
|---|---|---|---|---|---|---|---|---|
| HG00171Ax02PE20419 | chr22 | 22300000 | 22900000 | 600000 | del_h2 | Subclonal | HG00171 | 1 |
| HG00171Ax02PE20429 | chr22 | 22300000 | 22900000 | 600000 | del_h2 | Subclonal | HG00171 | 1 |
| HG00171Ax02PE20447 | chr22 | 22300000 | 22900000 | 600000 | del_h2 | Subclonal | HG00171 | 1 |
| HG00171Ax02PE20452 | chr22 | 22300000 | 22900000 | 600000 | del_h2 | Subclonal | HG00171 | 1 |
| HG00171Ax02PE20455 | chr22 | 22300000 | 22900000 | 600000 | del_h2 | Subclonal | HG00171 | 1 |
| HG00171Ax02PE20491 | chr22 | 22300000 | 22900000 | 600000 | del_h2 | Subclonal | HG00171 | 1 |
| HG00864x02PE20304 | chr22 | 22400000 | 22900000 | 500000 | del_h2 | Subclonal | HG00864 | 1 |
| HG00864x02PE20305 | chr22 | 22400000 | 22900000 | 500000 | del_h2 | Subclonal | HG00864 | 1 |
| HG00864x02PE20315 | chr22 | 22400000 | 22900000 | 500000 | del_h2 | Subclonal | HG00864 | 1 |
| HG00864x02PE20319 | chr22 | 22400000 | 22900000 | 500000 | del_h2 | Subclonal | HG00864 | 1 |
| HG00864x02PE20322 | chr22 | 22400000 | 22900000 | 500000 | del_h2 | Subclonal | HG00864 | 1 |
| HG00864x02PE20323 | chr22 | 22400000 | 22900000 | 500000 | del_h2 | Subclonal | HG00864 | 1 |
| HG00864x02PE20325 | chr22 | 22400000 | 22900000 | 500000 | del_h2 | Subclonal | HG00864 | 1 |
| HG00864x02PE20330 | chr22 | 22400000 | 22900000 | 500000 | del_h2 | Subclonal | HG00864 | 1 |
| HG00864x02PE20336 | chr22 | 22400000 | 22900000 | 500000 | del_h2 | Subclonal | HG00864 | 1 |
| HG00864x02PE20339 | chr22 | 22400000 | 22900000 | 500000 | del_h2 | Subclonal | HG00864 | 1 |
| HG00864x02PE20344 | chr22 | 22400000 | 22900000 | 500000 | del_h2 | Subclonal | HG00864 | 1 |
| HG00864x02PE20347 | chr22 | 22400000 | 22900000 | 500000 | del_h2 | Subclonal | HG00864 | 1 |
| HG00864x02PE20350 | chr22 | 22400000 | 22900000 | 500000 | del_h2 | Subclonal | HG00864 | 1 |
| HG00864x02PE20353 | chr22 | 22400000 | 22900000 | 500000 | del_h2 | Subclonal | HG00864 | 1 |
| HG00864x02PE20359 | chr22 | 22400000 | 22900000 | 500000 | del_h2 | Subclonal | HG00864 | 1 |
| HG00864x02PE20360 | chr22 | 22400000 | 22900000 | 500000 | del_h2 | Subclonal | HG00864 | 1 |
| HG00864x02PE20370 | chr22 | 22400000 | 22900000 | 500000 | del_h2 | Subclonal | HG00864 | 1 |
| HG01114x02PE20319 | chr22 | 22400000 | 22900000 | 500000 | del_h1 | Subclonal | HG01114 | 1 |
| HG01114x02PE20366 | chr22 | 22400000 | 22900000 | 500000 | del_h1 | Subclonal | HG01114 | 1 |
| HG01114x02PE20320 | chr22 | 22400000 | 22900000 | 500000 | del_h2 | Subclonal | HG01114 | 1 |
| HG01114x02PE20334 | chr22 | 22400000 | 22900000 | 500000 | del_h2 | Subclonal | HG01114 | 1 |
| HG01114x02PE20382 | chr22 | 22400000 | 22900000 | 500000 | del_h2 | Subclonal | HG01114 | 1 |
| HG01114x02PE20348 | chr22 | 22400000 | 22900000 | 500000 | del_hom | Singleton | HG01114 | 0 |
| HG01596x02PE20508 | chr22 | 22300000 | 22900000 | 600000 | del_h1 | Subclonal | HG01596 | 1 |
| HG01596x02PE20575 | chr22 | 22300000 | 22900000 | 600000 | del_h1 | Subclonal | HG01596 | 1 |
| HG01596x02PE20578 | chr22 | 22300000 | 22900000 | 600000 | del_h1 | Subclonal | HG01596 | 1 |
| HG01596x02PE20587 | chr22 | 22300000 | 22900000 | 600000 | del_h1 | Subclonal | HG01596 | 1 |
| HG01596x02PE20521 | chr22 | 22300000 | 22900000 | 600000 | del_h2 | Subclonal | HG01596 | 1 |
| HG01596x02PE20544 | chr22 | 22300000 | 22900000 | 600000 | del_h2 | Subclonal | HG01596 | 1 |
| HG01596x02PE20571 | chr22 | 22300000 | 22900000 | 600000 | del_h2 | Subclonal | HG01596 | 1 |
| HG01596x02PE20573 | chr22 | 22300000 | 22900000 | 600000 | del_hom | Singleton | HG01596 | 0 |
| HG02011x02PE20533 | chr22 | 22500000 | 22900000 | 400000 | del_h1 | Subclonal | HG02011 | 1 |

| | | | | | | | | |
|---|---|---|---|---|---|---|---|---|
| HG02011x02PE20534 | chr22 | 22500000 | 22900000 | 400000 | del_h1 | Subclonal | HG02011 | 1 |
| HG02011x02PE20539 | chr22 | 22500000 | 22900000 | 400000 | del_h1 | Subclonal | HG02011 | 1 |
| HG02011x02PE20544 | chr22 | 22500000 | 22900000 | 400000 | del_h1 | Subclonal | HG02011 | 1 |
| HG02011x02PE20557 | chr22 | 22500000 | 22900000 | 400000 | del_h1 | Subclonal | HG02011 | 1 |
| HG02011x02PE20569 | chr22 | 22500000 | 22900000 | 400000 | del_h1 | Subclonal | HG02011 | 1 |
| HG02011x02PE20575 | chr22 | 22500000 | 22900000 | 400000 | del_h1 | Subclonal | HG02011 | 1 |
| HG02011x02PE20584 | chr22 | 22500000 | 22900000 | 400000 | del_h1 | Subclonal | HG02011 | 1 |
| HG02587x02PE20301 | chr22 | 22400000 | 22900000 | 500000 | del_h2 | Subclonal | HG02587 | 1 |
| HG02587x02PE20305 | chr22 | 22400000 | 22900000 | 500000 | del_h2 | Subclonal | HG02587 | 1 |
| HG02587x02PE20306 | chr22 | 22400000 | 22900000 | 500000 | del_h2 | Subclonal | HG02587 | 1 |
| HG02587x02PE20309 | chr22 | 22400000 | 22900000 | 500000 | del_h2 | Subclonal | HG02587 | 1 |
| HG02587x02PE20310 | chr22 | 22400000 | 22900000 | 500000 | del_h2 | Subclonal | HG02587 | 1 |
| HG02587x02PE20312 | chr22 | 22400000 | 22900000 | 500000 | del_h2 | Subclonal | HG02587 | 1 |
| HG02587x02PE20313 | chr22 | 22400000 | 22900000 | 500000 | del_h2 | Subclonal | HG02587 | 1 |
| HG02587x02PE20314 | chr22 | 22400000 | 22900000 | 500000 | del_h2 | Subclonal | HG02587 | 1 |
| HG02587x02PE20315 | chr22 | 22400000 | 22900000 | 500000 | del_h2 | Subclonal | HG02587 | 1 |
| HG02587x02PE20318 | chr22 | 22400000 | 22900000 | 500000 | del_h2 | Subclonal | HG02587 | 1 |
| HG02587x02PE20320 | chr22 | 22400000 | 22900000 | 500000 | del_h2 | Subclonal | HG02587 | 1 |
| HG02587x02PE20322 | chr22 | 22400000 | 22900000 | 500000 | del_h2 | Subclonal | HG02587 | 1 |
| HG02587x02PE20325 | chr22 | 22400000 | 22900000 | 500000 | del_h2 | Subclonal | HG02587 | 1 |
| HG02587x02PE20326 | chr22 | 22400000 | 22900000 | 500000 | del_h2 | Subclonal | HG02587 | 1 |
| HG02587x02PE20328 | chr22 | 22400000 | 22900000 | 500000 | del_h2 | Subclonal | HG02587 | 1 |
| HG02587x02PE20330 | chr22 | 22400000 | 22900000 | 500000 | del_h2 | Subclonal | HG02587 | 1 |
| HG02587x02PE20331 | chr22 | 22400000 | 22900000 | 500000 | del_h2 | Subclonal | HG02587 | 1 |
| HG02587x02PE20333 | chr22 | 22400000 | 22900000 | 500000 | del_h2 | Subclonal | HG02587 | 1 |
| HG02587x02PE20334 | chr22 | 22400000 | 22900000 | 500000 | del_h2 | Subclonal | HG02587 | 1 |
| HG02587x02PE20336 | chr22 | 22400000 | 22900000 | 500000 | del_h2 | Subclonal | HG02587 | 1 |
| HG02587x02PE20337 | chr22 | 22400000 | 22900000 | 500000 | del_h2 | Subclonal | HG02587 | 1 |
| HG02587x02PE20338 | chr22 | 22400000 | 22900000 | 500000 | del_h2 | Subclonal | HG02587 | 1 |
| HG02587x02PE20341 | chr22 | 22400000 | 22900000 | 500000 | del_h2 | Subclonal | HG02587 | 1 |
| HG02587x02PE20344 | chr22 | 22400000 | 22900000 | 500000 | del_h2 | Subclonal | HG02587 | 1 |
| HG02587x02PE20346 | chr22 | 22400000 | 22900000 | 500000 | del_h2 | Subclonal | HG02587 | 1 |
| HG02587x02PE20347 | chr22 | 22400000 | 22900000 | 500000 | del_h2 | Subclonal | HG02587 | 1 |
| HG02587x02PE20349 | chr22 | 22400000 | 22900000 | 500000 | del_h2 | Subclonal | HG02587 | 1 |
| HG02587x02PE20351 | chr22 | 22400000 | 22900000 | 500000 | del_h2 | Subclonal | HG02587 | 1 |
| HG02587x02PE20353 | chr22 | 22400000 | 22900000 | 500000 | del_h2 | Subclonal | HG02587 | 1 |
| HG02587x02PE20358 | chr22 | 22400000 | 22900000 | 500000 | del_h2 | Subclonal | HG02587 | 1 |
| HG02587x02PE20360 | chr22 | 22400000 | 22900000 | 500000 | del_h2 | Subclonal | HG02587 | 1 |

| | | | | | | | | |
|---|---|---|---|---|---|---|---|---|
| HG02587x02PE20361 | chr22 | 22400000 | 22900000 | 500000 | del_h2 | Subclonal | HG02587 | 1 |
| HG02587x02PE20364 | chr22 | 22400000 | 22900000 | 500000 | del_h2 | Subclonal | HG02587 | 1 |
| HG02587x02PE20365 | chr22 | 22400000 | 22900000 | 500000 | del_h2 | Subclonal | HG02587 | 1 |
| HG02587x02PE20367 | chr22 | 22400000 | 22900000 | 500000 | del_h2 | Subclonal | HG02587 | 1 |
| HG02587x02PE20368 | chr22 | 22400000 | 22900000 | 500000 | del_h2 | Subclonal | HG02587 | 1 |
| HG02587x02PE20373 | chr22 | 22400000 | 22900000 | 500000 | del_h2 | Subclonal | HG02587 | 1 |
| HG02587x02PE20374 | chr22 | 22400000 | 22900000 | 500000 | del_h2 | Subclonal | HG02587 | 1 |
| HG02587x02PE20375 | chr22 | 22400000 | 22900000 | 500000 | del_h2 | Subclonal | HG02587 | 1 |
| HG02587x02PE20376 | chr22 | 22400000 | 22900000 | 500000 | del_h2 | Subclonal | HG02587 | 1 |
| HG02587x02PE20377 | chr22 | 22400000 | 22900000 | 500000 | del_h2 | Subclonal | HG02587 | 1 |
| HG02587x02PE20381 | chr22 | 22400000 | 22900000 | 500000 | del_h2 | Subclonal | HG02587 | 1 |
| HG02587x02PE20383 | chr22 | 22400000 | 22900000 | 500000 | del_h2 | Subclonal | HG02587 | 1 |
| HG02587x02PE20384 | chr22 | 22400000 | 22900000 | 500000 | del_h2 | Subclonal | HG02587 | 1 |
| HG02587x02PE20385 | chr22 | 22400000 | 22900000 | 500000 | del_h2 | Subclonal | HG02587 | 1 |
| HG02587x02PE20389 | chr22 | 22400000 | 22900000 | 500000 | del_h2 | Subclonal | HG02587 | 1 |
| HG02587x02PE20391 | chr22 | 22400000 | 22900000 | 500000 | del_h2 | Subclonal | HG02587 | 1 |
| HG02587x02PE20392 | chr22 | 22400000 | 22900000 | 500000 | del_h2 | Subclonal | HG02587 | 1 |
| HG02818x02PE20307 | chr22 | 22500000 | 22900000 | 400000 | del_het+del_hom | Subclonal | HG02818 | 1,0 |
| HG02818x02PE20310 | chr22 | 22500000 | 22900000 | 400000 | del_het+del_hom | Subclonal | HG02818 | 1,0 |
| HG02818x02PE20324 | chr22 | 22500000 | 22900000 | 400000 | del_het+del_hom | Subclonal | HG02818 | 1,0 |
| HG02818x02PE20339 | chr22 | 22500000 | 22900000 | 400000 | del_het+del_hom | Subclonal | HG02818 | 1,0 |
| HG02818x02PE20340 | chr22 | 22500000 | 22900000 | 400000 | del_het+del_hom | Subclonal | HG02818 | 1,0 |
| HG02818x02PE20342 | chr22 | 22500000 | 22900000 | 400000 | del_het+del_hom | Subclonal | HG02818 | 1,0 |
| HG02818x02PE20350 | chr22 | 22500000 | 22900000 | 400000 | del_het+del_hom | Subclonal | HG02818 | 1,0 |
| HG02818x02PE20352 | chr22 | 22500000 | 22900000 | 400000 | del_het+del_hom | Subclonal | HG02818 | 1,0 |
| HG02818x02PE20354 | chr22 | 22500000 | 22900000 | 400000 | del_het+del_hom | Subclonal | HG02818 | 1,0 |
| HG02818x02PE20355 | chr22 | 22500000 | 22900000 | 400000 | del_het+del_hom | Subclonal | HG02818 | 1,0 |
| HG02818x02PE20362 | chr22 | 22500000 | 22900000 | 400000 | del_het+del_hom | Subclonal | HG02818 | 1,0 |
| HG02818x02PE20371 | chr22 | 22500000 | 22900000 | 400000 | del_het+del_hom | Subclonal | HG02818 | 1,0 |
| HG02818x02PE20378 | chr22 | 22500000 | 22900000 | 400000 | del_het+del_hom | Subclonal | HG02818 | 1,0 |
| HG02818x02PE20381 | chr22 | 22500000 | 22900000 | 400000 | del_het+del_hom | Subclonal | HG02818 | 1,0 |
| HG02818x02PE20390 | chr22 | 22500000 | 22900000 | 400000 | del_het+del_hom | Subclonal | HG02818 | 1,0 |
| HG03486x02PE20503 | chr22 | 22700000 | 22900000 | 200000 | del_h1 | Subclonal | HG03486 | 1 |
| HG03486x02PE20510 | chr22 | 22700000 | 22900000 | 200000 | del_h1 | Subclonal | HG03486 | 1 |
| HG03486x02PE20511 | chr22 | 22700000 | 22900000 | 200000 | del_h1 | Subclonal | HG03486 | 1 |
| HG03486x02PE20519 | chr22 | 22700000 | 22900000 | 200000 | del_h1 | Subclonal | HG03486 | 1 |
| HG03486x02PE20543 | chr22 | 22700000 | 22900000 | 200000 | del_h1 | Subclonal | HG03486 | 1 |
| HG03486x02PE20545 | chr22 | 22700000 | 22900000 | 200000 | del_h1 | Subclonal | HG03486 | 1 |

| | | | | | | | | |
|---|---|---|---|---|---|---|---|---|
| HG03486x02PE20546 | chr22 | 22700000 | 22900000 | 200000 | del_h1 | Subclonal | HG03486 | 1 |
| HG03486x02PE20547 | chr22 | 22700000 | 22900000 | 200000 | del_h1 | Subclonal | HG03486 | 1 |
| HG03486x02PE20552 | chr22 | 22700000 | 22900000 | 200000 | del_h1 | Subclonal | HG03486 | 1 |
| HG03486x02PE20556 | chr22 | 22700000 | 22900000 | 200000 | del_h1 | Subclonal | HG03486 | 1 |
| HG03486x02PE20557 | chr22 | 22700000 | 22900000 | 200000 | del_h1 | Subclonal | HG03486 | 1 |
| HG03486x02PE20560 | chr22 | 22700000 | 22900000 | 200000 | del_h1 | Subclonal | HG03486 | 1 |
| HG03486x02PE20562 | chr22 | 22700000 | 22900000 | 200000 | del_h1 | Subclonal | HG03486 | 1 |
| HG03486x02PE20567 | chr22 | 22700000 | 22900000 | 200000 | del_h1 | Subclonal | HG03486 | 1 |
| HG03486x02PE20568 | chr22 | 22700000 | 22900000 | 200000 | del_h1 | Subclonal | HG03486 | 1 |
| HG03486x02PE20572 | chr22 | 22700000 | 22900000 | 200000 | del_h1 | Subclonal | HG03486 | 1 |
| HG03486x02PE20578 | chr22 | 22700000 | 22900000 | 200000 | del_h1 | Subclonal | HG03486 | 1 |
| HG03486x02PE20579 | chr22 | 22700000 | 22900000 | 200000 | del_h1 | Subclonal | HG03486 | 1 |
| HG03486x02PE20580 | chr22 | 22700000 | 22900000 | 200000 | del_h1 | Subclonal | HG03486 | 1 |
| HG03486x02PE20584 | chr22 | 22700000 | 22900000 | 200000 | del_h1 | Subclonal | HG03486 | 1 |
| HG03486x02PE20585 | chr22 | 22700000 | 22900000 | 200000 | del_h1 | Subclonal | HG03486 | 1 |
| HG03486x02PE20587 | chr22 | 22700000 | 22900000 | 200000 | del_h1 | Subclonal | HG03486 | 1 |
| HG03486x02PE20590 | chr22 | 22700000 | 22900000 | 200000 | del_h1 | Subclonal | HG03486 | 1 |
| HG03486x02PE20591 | chr22 | 22700000 | 22900000 | 200000 | del_h1 | Subclonal | HG03486 | 1 |
| HG03486x02PE20592 | chr22 | 22700000 | 22900000 | 200000 | del_h1 | Subclonal | HG03486 | 1 |
| HG03486x02PE20594 | chr22 | 22700000 | 22900000 | 200000 | del_h1 | Subclonal | HG03486 | 1 |
| HG03486x02PE20596 | chr22 | 22700000 | 22900000 | 200000 | del_h1 | Subclonal | HG03486 | 1 |
| HG03486x02PE20563 | chr22 | 22500000 | 22900000 | 400000 | del_h1 | Singleton | HG03486 | 1 |
| HG03683x01PE20401 | chr22 | 22300000 | 22900000 | 600000 | del_h1 | Subclonal | HG03683 | 1 |
| HG03683x01PE20441 | chr22 | 22300000 | 22900000 | 600000 | del_h1 | Subclonal | HG03683 | 1 |
| HG03683x01PE20448 | chr22 | 22300000 | 22900000 | 600000 | del_h1 | Subclonal | HG03683 | 1 |
| HG03683x01PE20495 | chr22 | 22300000 | 22900000 | 600000 | del_h1 | Subclonal | HG03683 | 1 |
| HG03683x01PE20428 | chr22 | 22600000 | 22900000 | 300000 | del_h1 | Subclonal | HG03683 | 1 |
| HG03683x01PE20442 | chr22 | 22600000 | 22900000 | 300000 | del_h1 | Subclonal | HG03683 | 1 |
| HG03683x01PE20452 | chr22 | 22600000 | 22900000 | 300000 | del_h1 | Subclonal | HG03683 | 1 |
| HG03683x01PE20462 | chr22 | 22600000 | 22900000 | 300000 | del_h1 | Subclonal | HG03683 | 1 |
| HG03683x01PE20471 | chr22 | 22600000 | 22900000 | 300000 | del_h1 | Subclonal | HG03683 | 1 |
| HG03683x01PE20483 | chr22 | 22600000 | 22900000 | 300000 | del_hom | Singleton | HG03683 | 0 |
| HG03732x02PE20562 | chr22 | 22000000 | 22900000 | 900000 | del_h1 | Singleton | HG03732 | 1 |
| HG03732x02PE20514 | chr22 | 22500000 | 22900000 | 400000 | del_h2 | Subclonal | HG03732 | 1 |
| HG03732x02PE20582 | chr22 | 22500000 | 22900000 | 400000 | del_h2 | Subclonal | HG03732 | 1 |
| HG03732x02PE20510 | chr22 | 22300000 | 22900000 | 600000 | del_hom | Subclonal | HG03732 | 0 |
| HG03732x02PE20522 | chr22 | 22300000 | 22900000 | 600000 | del_hom | Subclonal | HG03732 | 0 |
| HG03732x02PE20546 | chr22 | 22300000 | 22900000 | 600000 | del_h1 | Singleton | HG03732 | 1 |

| | | | | | | | | |
|---|---|---|---|---|---|---|---|---|
| HG03732x02PE20555 | chr22 | 22300000 | 22900000 | 600000 | del_h2 | Subclonal | HG03732 | 1 |
| HG03732x02PE20557 | chr22 | 22300000 | 22900000 | 600000 | del_h2 | Subclonal | HG03732 | 1 |
| HG03732x02PE20577 | chr22 | 22300000 | 22900000 | 600000 | del_h2 | Subclonal | HG03732 | 1 |
| HG03732x02PE20584 | chr22 | 22300000 | 22900000 | 600000 | del_h2 | Subclonal | HG03732 | 1 |
| NW130711.291.L008 | chr22 | 22200000 | 22900000 | 700000 | del_h2 | Subclonal | NA12878 | 1 |
| NW130711.296.L008 | chr22 | 22200000 | 22900000 | 700000 | del_h2 | Subclonal | NA12878 | 1 |
| NW130711.300.L008 | chr22 | 22200000 | 22900000 | 700000 | del_h2 | Subclonal | NA12878 | 1 |
| NW130711.307.L008 | chr22 | 22200000 | 22900000 | 700000 | del_h2 | Subclonal | NA12878 | 1 |
| NW130711.324.L008 | chr22 | 22200000 | 22900000 | 700000 | del_h2 | Subclonal | NA12878 | 1 |
| NW130711.327.L008 | chr22 | 22200000 | 22900000 | 700000 | del_h2 | Subclonal | NA12878 | 1 |
| NW130711.329.L008 | chr22 | 22200000 | 22900000 | 700000 | del_h2 | Subclonal | NA12878 | 1 |
| NW130711.350.L008 | chr22 | 22200000 | 22900000 | 700000 | del_h2 | Subclonal | NA12878 | 1 |
| NW130711.355.L008 | chr22 | 22200000 | 22900000 | 700000 | del_h2 | Subclonal | NA12878 | 1 |
| NW130711.362.L008 | chr22 | 22200000 | 22900000 | 700000 | del_h2 | Subclonal | NA12878 | 1 |
| NW130711.365.L008 | chr22 | 22200000 | 22900000 | 700000 | del_h2 | Subclonal | NA12878 | 1 |
| NW130711.374.L008 | chr22 | 22200000 | 22900000 | 700000 | del_h2 | Subclonal | NA12878 | 1 |
| NW130711.376.L008 | chr22 | 22200000 | 22900000 | 700000 | del_h2 | Subclonal | NA12878 | 1 |
| NW130711.377.L008 | chr22 | 22200000 | 22900000 | 700000 | del_h2 | Subclonal | NA12878 | 1 |
| NW150212-III.05.L001 | chr22 | 22200000 | 22900000 | 700000 | del_h2 | Subclonal | NA12878 | 1 |
| NW150212-III.06.L001 | chr22 | 22200000 | 22900000 | 700000 | del_h2 | Subclonal | NA12878 | 1 |
| NW150212-III.09.L001 | chr22 | 22200000 | 22900000 | 700000 | del_h2 | Subclonal | NA12878 | 1 |
| NW150212-III.12.L001 | chr22 | 22200000 | 22900000 | 700000 | del_h2 | Subclonal | NA12878 | 1 |
| NW150212-III.16.L001 | chr22 | 22200000 | 22900000 | 700000 | del_h2 | Subclonal | NA12878 | 1 |
| NW150212-III.55.L001 | chr22 | 22200000 | 22900000 | 700000 | del_h2 | Subclonal | NA12878 | 1 |
| NW150212-III.57.L001 | chr22 | 22200000 | 22900000 | 700000 | del_h2 | Subclonal | NA12878 | 1 |
| NW150212-III.63.L001 | chr22 | 22200000 | 22900000 | 700000 | del_h2 | Subclonal | NA12878 | 1 |
| NW150212-III.68.L001 | chr22 | 22200000 | 22900000 | 700000 | del_h2 | Subclonal | NA12878 | 1 |
| NW150212-III.77.L001 | chr22 | 22200000 | 22900000 | 700000 | del_h2 | Subclonal | NA12878 | 1 |
| NW150212-III.84.L001 | chr22 | 22200000 | 22900000 | 700000 | del_h2 | Subclonal | NA12878 | 1 |
| NW150212-III.94.L001 | chr22 | 22200000 | 22900000 | 700000 | del_h2 | Subclonal | NA12878 | 1 |
| NW150212-IV.39.L002 | chr22 | 22200000 | 22900000 | 700000 | del_h2 | Subclonal | NA12878 | 1 |
| NW150212-IV.60.L002 | chr22 | 22200000 | 22900000 | 700000 | del_h2 | Subclonal | NA12878 | 1 |
| NW150212-IV.72.L002 | chr22 | 22200000 | 22900000 | 700000 | del_h2 | Subclonal | NA12878 | 1 |
| NW150212-IV.77.L002 | chr22 | 22200000 | 22900000 | 700000 | del_h2 | Subclonal | NA12878 | 1 |
| NW150212-III.03.L001 | chr22 | 22200000 | 22900000 | 700000 | del_h2+del_hom | Subclonal | NA12878 | 1,0 |
| NW150212-III.11.L001 | chr22 | 22200000 | 22900000 | 700000 | del_h2+del_hom | Subclonal | NA12878 | 1,0 |
| NW150212-III.21.L001 | chr22 | 22200000 | 22900000 | 700000 | del_h2+del_hom | Subclonal | NA12878 | 1,0 |
| NW150212-III.26.L001 | chr22 | 22200000 | 22900000 | 700000 | del_h2+del_hom | Subclonal | NA12878 | 1,0 |

| | | | | | | | | |
|---|---|---|---|---|---|---|---|---|
| NW150212-III.30.L001 | chr22 | 22200000 | 22900000 | 700000 | del_h2+del_hom | Subclonal | NA12878 | 1,0 |
| NW150212-III.33.L001 | chr22 | 22200000 | 22900000 | 700000 | del_h2+del_hom | Subclonal | NA12878 | 1,0 |
| NW150212-III.35.L001 | chr22 | 22200000 | 22900000 | 700000 | del_h2+del_hom | Subclonal | NA12878 | 1,0 |
| NW150212-III.36.L001 | chr22 | 22200000 | 22900000 | 700000 | del_h2+del_hom | Subclonal | NA12878 | 1,0 |
| NW150212-III.48.L001 | chr22 | 22200000 | 22900000 | 700000 | del_h2+del_hom | Subclonal | NA12878 | 1,0 |
| NW150212-III.52.L001 | chr22 | 22200000 | 22900000 | 700000 | del_h2+del_hom | Subclonal | NA12878 | 1,0 |
| NW150212-III.59.L001 | chr22 | 22200000 | 22900000 | 700000 | del_h2+del_hom | Subclonal | NA12878 | 1,0 |
| NW150212-III.69.L001 | chr22 | 22200000 | 22900000 | 700000 | del_h2+del_hom | Subclonal | NA12878 | 1,0 |
| NW150212-III.92.L001 | chr22 | 22200000 | 22900000 | 700000 | del_h2+del_hom | Subclonal | NA12878 | 1,0 |
| NW150212-III.93.L001 | chr22 | 22200000 | 22900000 | 700000 | del_h2+del_hom | Subclonal | NA12878 | 1,0 |
| NW150212-IV.09.L002 | chr22 | 22200000 | 22900000 | 700000 | del_h2+del_hom | Subclonal | NA12878 | 1,0 |
| NW150212-IV.10.L002 | chr22 | 22200000 | 22900000 | 700000 | del_h2+del_hom | Subclonal | NA12878 | 1,0 |
| NW150212-IV.11.L002 | chr22 | 22200000 | 22900000 | 700000 | del_h2+del_hom | Subclonal | NA12878 | 1,0 |
| NW150212-IV.12.L002 | chr22 | 22200000 | 22900000 | 700000 | del_h2+del_hom | Subclonal | NA12878 | 1,0 |
| NW150212-IV.15.L002 | chr22 | 22200000 | 22900000 | 700000 | del_h2+del_hom | Subclonal | NA12878 | 1,0 |
| NW150212-IV.17.L002 | chr22 | 22200000 | 22900000 | 700000 | del_h2+del_hom | Subclonal | NA12878 | 1,0 |
| NW150212-IV.24.L002 | chr22 | 22200000 | 22900000 | 700000 | del_h2+del_hom | Subclonal | NA12878 | 1,0 |
| NW150212-IV.25.L002 | chr22 | 22200000 | 22900000 | 700000 | del_h2+del_hom | Subclonal | NA12878 | 1,0 |
| NW150212-IV.31.L002 | chr22 | 22200000 | 22900000 | 700000 | del_h2+del_hom | Subclonal | NA12878 | 1,0 |
| NW150212-IV.34.L002 | chr22 | 22200000 | 22900000 | 700000 | del_h2+del_hom | Subclonal | NA12878 | 1,0 |
| NW150212-IV.41.L002 | chr22 | 22200000 | 22900000 | 700000 | del_h2+del_hom | Subclonal | NA12878 | 1,0 |
| NW150212-IV.43.L002 | chr22 | 22200000 | 22900000 | 700000 | del_h2+del_hom | Subclonal | NA12878 | 1,0 |
| NW150212-IV.44.L002 | chr22 | 22200000 | 22900000 | 700000 | del_h2+del_hom | Subclonal | NA12878 | 1,0 |
| NW150212-IV.45.L002 | chr22 | 22200000 | 22900000 | 700000 | del_h2+del_hom | Subclonal | NA12878 | 1,0 |
| NW150212-IV.46.L002 | chr22 | 22200000 | 22900000 | 700000 | del_h2+del_hom | Subclonal | NA12878 | 1,0 |
| NW150212-IV.50.L002 | chr22 | 22200000 | 22900000 | 700000 | del_h2+del_hom | Subclonal | NA12878 | 1,0 |
| NW150212-IV.58.L002 | chr22 | 22200000 | 22900000 | 700000 | del_h2+del_hom | Subclonal | NA12878 | 1,0 |
| NW150212-IV.62.L002 | chr22 | 22200000 | 22900000 | 700000 | del_h2+del_hom | Subclonal | NA12878 | 1,0 |
| NW150212-IV.65.L002 | chr22 | 22200000 | 22900000 | 700000 | del_h2+del_hom | Subclonal | NA12878 | 1,0 |
| NW150212-IV.68.L002 | chr22 | 22200000 | 22900000 | 700000 | del_h2+del_hom | Subclonal | NA12878 | 1,0 |
| NW150212-IV.69.L002 | chr22 | 22200000 | 22900000 | 700000 | del_h2+del_hom | Subclonal | NA12878 | 1,0 |
| NW150212-IV.74.L002 | chr22 | 22200000 | 22900000 | 700000 | del_h2+del_hom | Subclonal | NA12878 | 1,0 |
| NW150212-IV.81.L002 | chr22 | 22200000 | 22900000 | 700000 | del_h2+del_hom | Subclonal | NA12878 | 1,0 |
| NW150212-IV.83.L002 | chr22 | 22200000 | 22900000 | 700000 | del_h2+del_hom | Subclonal | NA12878 | 1,0 |
| NW150212-IV.84.L002 | chr22 | 22200000 | 22900000 | 700000 | del_h2+del_hom | Subclonal | NA12878 | 1,0 |
| NW150212-IV.85.L002 | chr22 | 22200000 | 22900000 | 700000 | del_h2+del_hom | Subclonal | NA12878 | 1,0 |
| NW150212-IV.87.L002 | chr22 | 22200000 | 22900000 | 700000 | del_h2+del_hom | Subclonal | NA12878 | 1,0 |
| NW150212-IV.89.L002 | chr22 | 22200000 | 22900000 | 700000 | del_h2+del_hom | Subclonal | NA12878 | 1,0 |

| | | | | | | | | |
|---|---|---|---|---|---|---|---|---|
| NW150212-IV.91.L002 | chr22 | 22200000 | 22900000 | 700000 | del_h2+del_hom | Subclonal | NA12878 | 1,0 |
| NW130711.320.L008 | chr19 | 36500000 | 37000000 | 500000 | del_h2 | Subclonal | NA12878 | 1 |
| NW150212-III.07.L001 | chr19 | 36500000 | 37000000 | 500000 | del_h2 | Subclonal | NA12878 | 1 |
| NW150212-III.13.L001 | chr19 | 36500000 | 37000000 | 500000 | del_h2 | Subclonal | NA12878 | 1 |
| NW150212-III.27.L001 | chr19 | 36500000 | 37000000 | 500000 | del_h2 | Subclonal | NA12878 | 1 |
| NW150212-III.39.L001 | chr19 | 36500000 | 37000000 | 500000 | del_h2 | Subclonal | NA12878 | 1 |
| NW150212-III.42.L001 | chr19 | 36500000 | 37000000 | 500000 | del_h2 | Subclonal | NA12878 | 1 |
| NW150212-III.66.L001 | chr19 | 36500000 | 37000000 | 500000 | del_h2 | Subclonal | NA12878 | 1 |
| NW150212-III.75.L001 | chr19 | 36500000 | 37000000 | 500000 | del_h2 | Subclonal | NA12878 | 1 |
| NW150212-III.76.L001 | chr19 | 36500000 | 37000000 | 500000 | del_h2 | Subclonal | NA12878 | 1 |
| NW150212-III.80.L001 | chr19 | 36500000 | 37000000 | 500000 | del_h2 | Subclonal | NA12878 | 1 |
| NW150212-III.81.L001 | chr19 | 36500000 | 37000000 | 500000 | del_h2 | Subclonal | NA12878 | 1 |
| NW150212-III.90.L001 | chr19 | 36500000 | 37000000 | 500000 | del_h2 | Subclonal | NA12878 | 1 |
| NW150212-IV.08.L002 | chr19 | 36500000 | 37000000 | 500000 | del_h2 | Subclonal | NA12878 | 1 |
| NW150212-IV.26.L002 | chr19 | 36500000 | 37000000 | 500000 | del_h2 | Subclonal | NA12878 | 1 |
| NW150212-IV.63.L002 | chr19 | 36500000 | 37000000 | 500000 | del_h2 | Subclonal | NA12878 | 1 |
| NW150212-IV.64.L002 | chr19 | 36500000 | 37000000 | 500000 | del_h2 | Subclonal | NA12878 | 1 |
| NW150212-IV.93.L002 | chr19 | 36500000 | 37000000 | 500000 | del_h2 | Subclonal | NA12878 | 1 |
| GM20509Bx01PE20561 | chr17 | 19200000 | 21700000 | 2500000 | idup_h2 | Subclonal | NA20509 | 5 |
| GM20509Bx01PE20514 | chr17 | 19200000 | 21700000 | 2500000 | idup_h2 | Subclonal | NA20509 | 4 |
| GM20509Bx01PE20502 | chr17 | 19200000 | 21700000 | 2500000 | idup_h2 | Subclonal | NA20509 | 3 |
| GM20509Bx01PE20503 | chr17 | 19200000 | 21700000 | 2500000 | idup_h2 | Subclonal | NA20509 | 3 |
| GM20509Bx01PE20506 | chr17 | 19200000 | 21700000 | 2500000 | idup_h2 | Subclonal | NA20509 | 3 |
| GM20509Bx01PE20508 | chr17 | 19200000 | 21700000 | 2500000 | idup_h2 | Subclonal | NA20509 | 3 |
| GM20509Bx01PE20509 | chr17 | 19200000 | 21700000 | 2500000 | idup_h2 | Subclonal | NA20509 | 3 |
| GM20509Bx01PE20510 | chr17 | 19200000 | 21700000 | 2500000 | idup_h2 | Subclonal | NA20509 | 3 |
| GM20509Bx01PE20511 | chr17 | 19200000 | 21700000 | 2500000 | idup_h2 | Subclonal | NA20509 | 3 |
| GM20509Bx01PE20517 | chr17 | 19200000 | 21700000 | 2500000 | idup_h2 | Subclonal | NA20509 | 3 |
| GM20509Bx01PE20518 | chr17 | 19200000 | 21700000 | 2500000 | idup_h2 | Subclonal | NA20509 | 3 |
| GM20509Bx01PE20519 | chr17 | 19200000 | 21700000 | 2500000 | idup_h2 | Subclonal | NA20509 | 3 |
| GM20509Bx01PE20525 | chr17 | 19200000 | 21700000 | 2500000 | idup_h2 | Subclonal | NA20509 | 3 |
| GM20509Bx01PE20526 | chr17 | 19200000 | 21700000 | 2500000 | idup_h2 | Subclonal | NA20509 | 3 |
| GM20509Bx01PE20529 | chr17 | 19200000 | 21700000 | 2500000 | idup_h2 | Subclonal | NA20509 | 3 |
| GM20509Bx01PE20530 | chr17 | 19200000 | 21700000 | 2500000 | idup_h2 | Subclonal | NA20509 | 3 |
| GM20509Bx01PE20531 | chr17 | 19200000 | 21700000 | 2500000 | idup_h2 | Subclonal | NA20509 | 3 |
| GM20509Bx01PE20533 | chr17 | 19200000 | 21700000 | 2500000 | idup_h2 | Subclonal | NA20509 | 3 |
| GM20509Bx01PE20535 | chr17 | 19200000 | 21700000 | 2500000 | idup_h2 | Subclonal | NA20509 | 3 |
| GM20509Bx01PE20536 | chr17 | 19200000 | 21700000 | 2500000 | idup_h2 | Subclonal | NA20509 | 3 |

| | | | | | | | | |
|---|---|---|---|---|---|---|---|---|
| GM20509Bx01PE20539 | chr17 | 19200000 | 21700000 | 2500000 | idup_h2 | Subclonal | NA20509 | 3 |
| GM20509Bx01PE20541 | chr17 | 19200000 | 21700000 | 2500000 | idup_h2 | Subclonal | NA20509 | 3 |
| GM20509Bx01PE20545 | chr17 | 19200000 | 21700000 | 2500000 | idup_h2 | Subclonal | NA20509 | 3 |
| GM20509Bx01PE20546 | chr17 | 19200000 | 21700000 | 2500000 | idup_h2 | Subclonal | NA20509 | 3 |
| GM20509Bx01PE20547 | chr17 | 19200000 | 21700000 | 2500000 | idup_h2 | Subclonal | NA20509 | 3 |
| GM20509Bx01PE20552 | chr17 | 19200000 | 21700000 | 2500000 | idup_h2 | Subclonal | NA20509 | 3 |
| GM20509Bx01PE20555 | chr17 | 19200000 | 21700000 | 2500000 | idup_h2 | Subclonal | NA20509 | 3 |
| GM20509Bx01PE20556 | chr17 | 19200000 | 21700000 | 2500000 | idup_h2 | Subclonal | NA20509 | 3 |
| GM20509Bx01PE20557 | chr17 | 19200000 | 21700000 | 2500000 | idup_h2 | Subclonal | NA20509 | 3 |
| GM20509Bx01PE20560 | chr17 | 19200000 | 21700000 | 2500000 | idup_h2 | Subclonal | NA20509 | 3 |
| GM20509Bx01PE20562 | chr17 | 19200000 | 21700000 | 2500000 | idup_h2 | Subclonal | NA20509 | 3 |
| GM20509Bx01PE20565 | chr17 | 19200000 | 21700000 | 2500000 | idup_h2 | Subclonal | NA20509 | 3 |
| GM20509Bx01PE20566 | chr17 | 19200000 | 21700000 | 2500000 | idup_h2 | Subclonal | NA20509 | 3 |
| GM20509Bx01PE20568 | chr17 | 19200000 | 21700000 | 2500000 | idup_h2 | Subclonal | NA20509 | 3 |
| GM20509Bx01PE20569 | chr17 | 19200000 | 21700000 | 2500000 | idup_h2 | Subclonal | NA20509 | 3 |
| GM20509Bx01PE20576 | chr17 | 19200000 | 21700000 | 2500000 | idup_h2 | Subclonal | NA20509 | 3 |
| GM20509Bx01PE20581 | chr17 | 19200000 | 21700000 | 2500000 | idup_h2 | Subclonal | NA20509 | 3 |
| GM20509Bx01PE20587 | chr17 | 19200000 | 21700000 | 2500000 | idup_h2 | Subclonal | NA20509 | 3 |
| GM20509Bx01PE20588 | chr17 | 19200000 | 21700000 | 2500000 | idup_h2 | Subclonal | NA20509 | 3 |
| GM20509Bx01PE20520 | chr17 | 19200000 | 21700000 | 2500000 | idup_h2 | Subclonal | NA20509 | 2 |
| GM20509Bx01PE20502 | chr17 | 0 | 19200000 | 19200000 | del_h2 | Subclonal | NA20509 | 1 |
| GM20509Bx01PE20503 | chr17 | 0 | 19200000 | 19200000 | del_h2 | Subclonal | NA20509 | 1 |
| GM20509Bx01PE20506 | chr17 | 0 | 19200000 | 19200000 | del_h2 | Subclonal | NA20509 | 1 |
| GM20509Bx01PE20508 | chr17 | 0 | 19200000 | 19200000 | del_h2 | Subclonal | NA20509 | 1 |
| GM20509Bx01PE20509 | chr17 | 0 | 19200000 | 19200000 | del_h2 | Subclonal | NA20509 | 1 |
| GM20509Bx01PE20510 | chr17 | 0 | 19200000 | 19200000 | del_h2 | Subclonal | NA20509 | 1 |
| GM20509Bx01PE20511 | chr17 | 0 | 19200000 | 19200000 | del_h2 | Subclonal | NA20509 | 1 |
| GM20509Bx01PE20514 | chr17 | 0 | 19200000 | 19200000 | del_h2 | Subclonal | NA20509 | 1 |
| GM20509Bx01PE20517 | chr17 | 0 | 19200000 | 19200000 | del_h2 | Subclonal | NA20509 | 1 |
| GM20509Bx01PE20518 | chr17 | 0 | 19200000 | 19200000 | del_h2 | Subclonal | NA20509 | 1 |
| GM20509Bx01PE20519 | chr17 | 0 | 19200000 | 19200000 | del_h2 | Subclonal | NA20509 | 1 |
| GM20509Bx01PE20520 | chr17 | 0 | 19200000 | 19200000 | del_h2 | Subclonal | NA20509 | 1 |
| GM20509Bx01PE20525 | chr17 | 0 | 19200000 | 19200000 | del_h2 | Subclonal | NA20509 | 1 |
| GM20509Bx01PE20526 | chr17 | 0 | 19200000 | 19200000 | del_h2 | Subclonal | NA20509 | 1 |
| GM20509Bx01PE20529 | chr17 | 0 | 19200000 | 19200000 | del_h2 | Subclonal | NA20509 | 1 |
| GM20509Bx01PE20530 | chr17 | 0 | 19200000 | 19200000 | del_h2 | Subclonal | NA20509 | 1 |
| GM20509Bx01PE20531 | chr17 | 0 | 19200000 | 19200000 | del_h2 | Subclonal | NA20509 | 1 |
| GM20509Bx01PE20533 | chr17 | 0 | 19200000 | 19200000 | del_h2 | Subclonal | NA20509 | 1 |

| | | | | | | | | |
|---|---|---|---|---|---|---|---|---|
| GM20509Bx01PE20535 | chr17 | 0 | 19200000 | 19200000 | del_h2 | Subclonal | NA20509 | 1 |
| GM20509Bx01PE20536 | chr17 | 0 | 19200000 | 19200000 | del_h2 | Subclonal | NA20509 | 1 |
| GM20509Bx01PE20539 | chr17 | 0 | 19200000 | 19200000 | del_h2 | Subclonal | NA20509 | 1 |
| GM20509Bx01PE20541 | chr17 | 0 | 19200000 | 19200000 | del_h2 | Subclonal | NA20509 | 1 |
| GM20509Bx01PE20545 | chr17 | 0 | 19200000 | 19200000 | del_h2 | Subclonal | NA20509 | 1 |
| GM20509Bx01PE20546 | chr17 | 0 | 19200000 | 19200000 | del_h2 | Subclonal | NA20509 | 1 |
| GM20509Bx01PE20547 | chr17 | 0 | 19200000 | 19200000 | del_h2 | Subclonal | NA20509 | 1 |
| GM20509Bx01PE20552 | chr17 | 0 | 19200000 | 19200000 | del_h2 | Subclonal | NA20509 | 1 |
| GM20509Bx01PE20555 | chr17 | 0 | 19200000 | 19200000 | del_h2 | Subclonal | NA20509 | 1 |
| GM20509Bx01PE20556 | chr17 | 0 | 19200000 | 19200000 | del_h2 | Subclonal | NA20509 | 1 |
| GM20509Bx01PE20557 | chr17 | 0 | 19200000 | 19200000 | del_h2 | Subclonal | NA20509 | 1 |
| GM20509Bx01PE20560 | chr17 | 0 | 19200000 | 19200000 | del_h2 | Subclonal | NA20509 | 1 |
| GM20509Bx01PE20561 | chr17 | 0 | 19200000 | 19200000 | del_h2 | Subclonal | NA20509 | 1 |
| GM20509Bx01PE20562 | chr17 | 0 | 19200000 | 19200000 | del_h2 | Subclonal | NA20509 | 1 |
| GM20509Bx01PE20565 | chr17 | 0 | 19200000 | 19200000 | del_h2 | Subclonal | NA20509 | 1 |
| GM20509Bx01PE20566 | chr17 | 0 | 19200000 | 19200000 | del_h2 | Subclonal | NA20509 | 1 |
| GM20509Bx01PE20568 | chr17 | 0 | 19200000 | 19200000 | del_h2 | Subclonal | NA20509 | 1 |
| GM20509Bx01PE20569 | chr17 | 0 | 19200000 | 19200000 | del_h2 | Subclonal | NA20509 | 1 |
| GM20509Bx01PE20576 | chr17 | 0 | 19200000 | 19200000 | del_h2 | Subclonal | NA20509 | 1 |
| GM20509Bx01PE20581 | chr17 | 0 | 19200000 | 19200000 | del_h2 | Subclonal | NA20509 | 1 |
| GM20509Bx01PE20587 | chr17 | 0 | 19200000 | 19200000 | del_h2 | Subclonal | NA20509 | 1 |
| GM20509Bx01PE20588 | chr17 | 0 | 19200000 | 19200000 | del_h2 | Subclonal | NA20509 | 1 |
| GM20509Bx01PE20561 | chr5 | 132700000 | 181538259 | 48838259 | dup_h2 | Subclonal | NA20509 | 3 |
| GM20509Bx01PE20514 | chr5 | 132700000 | 181538259 | 48838259 | dup_h2 | Subclonal | NA20509 | 3 |
| GM20509Bx01PE20502 | chr5 | 132700000 | 181538259 | 48838259 | dup_h2 | Subclonal | NA20509 | 3 |
| GM20509Bx01PE20503 | chr5 | 132700000 | 181538259 | 48838259 | dup_h2 | Subclonal | NA20509 | 3 |
| GM20509Bx01PE20506 | chr5 | 132700000 | 181538259 | 48838259 | dup_h2 | Subclonal | NA20509 | 3 |
| GM20509Bx01PE20508 | chr5 | 132700000 | 181538259 | 48838259 | dup_h2 | Subclonal | NA20509 | 3 |
| GM20509Bx01PE20509 | chr5 | 132700000 | 181538259 | 48838259 | dup_h2 | Subclonal | NA20509 | 3 |
| GM20509Bx01PE20510 | chr5 | 132700000 | 181538259 | 48838259 | dup_h2 | Subclonal | NA20509 | 3 |
| GM20509Bx01PE20511 | chr5 | 132700000 | 181538259 | 48838259 | dup_h2 | Subclonal | NA20509 | 3 |
| GM20509Bx01PE20517 | chr5 | 132700000 | 181538259 | 48838259 | dup_h2 | Subclonal | NA20509 | 3 |
| GM20509Bx01PE20518 | chr5 | 132700000 | 181538259 | 48838259 | dup_h2 | Subclonal | NA20509 | 3 |
| GM20509Bx01PE20519 | chr5 | 132700000 | 181538259 | 48838259 | dup_h2 | Subclonal | NA20509 | 3 |
| GM20509Bx01PE20525 | chr5 | 132700000 | 181538259 | 48838259 | dup_h2 | Subclonal | NA20509 | 3 |
| GM20509Bx01PE20526 | chr5 | 132700000 | 181538259 | 48838259 | dup_h2 | Subclonal | NA20509 | 3 |
| GM20509Bx01PE20529 | chr5 | 132700000 | 181538259 | 48838259 | dup_h2 | Subclonal | NA20509 | 3 |
| GM20509Bx01PE20530 | chr5 | 132700000 | 181538259 | 48838259 | dup_h2 | Subclonal | NA20509 | 3 |

| | | | | | | | | |
|---|---|---|---|---|---|---|---|---|
| GM20509Bx01PE20531 | chr5 | 132700000 | 181538259 | 48838259 | dup_h2 | Subclonal | NA20509 | 3 |
| GM20509Bx01PE20533 | chr5 | 132700000 | 181538259 | 48838259 | dup_h2 | Subclonal | NA20509 | 3 |
| GM20509Bx01PE20535 | chr5 | 132700000 | 181538259 | 48838259 | dup_h2 | Subclonal | NA20509 | 3 |
| GM20509Bx01PE20536 | chr5 | 132700000 | 181538259 | 48838259 | dup_h2 | Subclonal | NA20509 | 3 |
| GM20509Bx01PE20539 | chr5 | 132700000 | 181538259 | 48838259 | dup_h2 | Subclonal | NA20509 | 3 |
| GM20509Bx01PE20541 | chr5 | 132700000 | 181538259 | 48838259 | dup_h2 | Subclonal | NA20509 | 3 |
| GM20509Bx01PE20545 | chr5 | 132700000 | 181538259 | 48838259 | dup_h2 | Subclonal | NA20509 | 3 |
| GM20509Bx01PE20546 | chr5 | 132700000 | 181538259 | 48838259 | dup_h2 | Subclonal | NA20509 | 3 |
| GM20509Bx01PE20547 | chr5 | 132700000 | 181538259 | 48838259 | dup_h2 | Subclonal | NA20509 | 3 |
| GM20509Bx01PE20552 | chr5 | 132700000 | 181538259 | 48838259 | dup_h2 | Subclonal | NA20509 | 3 |
| GM20509Bx01PE20555 | chr5 | 132700000 | 181538259 | 48838259 | dup_h2 | Subclonal | NA20509 | 3 |
| GM20509Bx01PE20556 | chr5 | 132700000 | 181538259 | 48838259 | dup_h2 | Subclonal | NA20509 | 3 |
| GM20509Bx01PE20557 | chr5 | 132700000 | 181538259 | 48838259 | dup_h2 | Subclonal | NA20509 | 3 |
| GM20509Bx01PE20560 | chr5 | 132700000 | 181538259 | 48838259 | dup_h2 | Subclonal | NA20509 | 3 |
| GM20509Bx01PE20562 | chr5 | 132700000 | 181538259 | 48838259 | dup_h2 | Subclonal | NA20509 | 3 |
| GM20509Bx01PE20565 | chr5 | 132700000 | 181538259 | 48838259 | dup_h2 | Subclonal | NA20509 | 3 |
| GM20509Bx01PE20566 | chr5 | 132700000 | 181538259 | 48838259 | dup_h2 | Subclonal | NA20509 | 3 |
| GM20509Bx01PE20568 | chr5 | 132700000 | 181538259 | 48838259 | dup_h2 | Subclonal | NA20509 | 3 |
| GM20509Bx01PE20569 | chr5 | 132700000 | 181538259 | 48838259 | dup_h2 | Subclonal | NA20509 | 3 |
| GM20509Bx01PE20576 | chr5 | 132700000 | 181538259 | 48838259 | dup_h2 | Subclonal | NA20509 | 3 |
| GM20509Bx01PE20581 | chr5 | 132700000 | 181538259 | 48838259 | dup_h2 | Subclonal | NA20509 | 3 |
| GM20509Bx01PE20587 | chr5 | 132700000 | 181538259 | 48838259 | dup_h2 | Subclonal | NA20509 | 3 |
| GM20509Bx01PE20588 | chr5 | 132700000 | 181538259 | 48838259 | dup_h2 | Subclonal | NA20509 | 3 |
| GM20509Bx01PE20520 | chr5 | 132700000 | 181538259 | 48838259 | dup_h2 | Subclonal | NA20509 | 3 |
| GM12329x02PE20445 | chr1 | 124932724 | 248956422 | 124023698 | dup_h1 | Singleton | NA12329 | 3 |
| GM12329x02PE20450 | chr1 | 124932724 | 248956422 | 124023698 | del_h2 | Subclonal | NA12329 | 1 |
| GM12329x02PE20484 | chr1 | 124932724 | 248956422 | 124023698 | del_h2 | Subclonal | NA12329 | 1 |
| GM12329x02PE20470 | chr8 | 0 | 145138636 | 145138636 | del_h1 | Singleton | NA12329 | 1 |
| GM12329x02PE20407 | chr9 | 101800000 | 138394717 | 36594717 | dup_h2 | Singleton | NA12329 | 3 |
| GM12329x02PE20476 | chr17 | 0 | 83257441 | 83257441 | del_h2 | Singleton | NA12329 | 1 |
| GM12329x02PE20476 | chr21 | 0 | 46709983 | 46709983 | del_h2 | Singleton | NA12329 | 1 |
| GM12329x02PE20487 | chrX | 0 | 2800000 | 2800000 | del_h2 | Singleton | NA12329 | 1 |
| GM18534Bx02PE20382 | chr1 | 124932724 | 248956422 | 124023698 | del_h2 | Singleton | NA18534 | 1 |
| GM18534Bx02PE20348 | chr4 | 0 | 9000000 | 9000000 | del_h2 | Singleton | NA18534 | 1 |
| GM18534Bx02PE20391 | chr4 | 128300000 | 190214555 | 61914555 | del_h1 | Singleton | NA18534 | 1 |
| GM18534Bx02PE20333 | chr11 | 0 | 135086622 | 135086622 | dup_h1 | Singleton | NA18534 | 3 |
| GM18534Bx02PE20372 | chr17 | 62400000 | 83257441 | 20857441 | del_h1 | Singleton | NA18534 | 1 |
| GM18939x02PE20447 | chr8 | 128100000 | 145138636 | 17038636 | dup_h1 | Singleton | NA18939 | 3 |

| | | | | | | | | |
|---|---|---|---|---|---|---|---|---|
| GM18939x02PE20460 | chr17 | 0 | 19000000 | 19000000 | dup_hom | Singleton | NA18939 | 4 |
| GM18939x02PE20444 | chrX | 0 | 156040895 | 156040895 | dup_h1 | Singleton | NA18939 | 3 |
| GM19650Ax02PE20590 | chr2 | 111600000 | 242193529 | 130593529 | del_h2 | Singleton | NA19650 | 1 |
| GM19650Ax02PE20574 | chr4 | 103400000 | 190214555 | 86814555 | del_h2 | Singleton | NA19650 | 1 |
| GM19650Ax02PE20541 | chr5 | 24000000 | 30300000 | 6300000 | del_h2 | Singleton | NA19650 | 1 |
| GM19650Ax02PE20511 | chr13 | 65500000 | 114364328 | 48864328 | del_h2 | Singleton | NA19650 | 1 |
| GM19983x02PE20477 | chr1 | 0 | 248956422 | 248956422 | dup_h1 | Singleton | NA19983 | 3 |
| GM19983x02PE20421 | chr2 | 0 | 242193529 | 242193529 | dup_h2 | Singleton | NA19983 | 3 |
| GM19983x02PE20421 | chr4 | 0 | 190214555 | 190214555 | dup_h2 | Singleton | NA19983 | 3 |
| GM19983x02PE20431 | chr4 | 156400000 | 190214555 | 33814555 | dup_h1 | Singleton | NA19983 | 3 |
| GM19983x02PE20477 | chr7 | 0 | 159345973 | 159345973 | dup_h2 | Singleton | NA19983 | 3 |
| GM19983x02PE20406 | chr9 | 45518558 | 138394717 | 92876159 | del_h2 | Singleton | NA19983 | 1 |
| GM19983x02PE20425 | chr9 | 79900000 | 81000000 | 1100000 | del_h1 | Subclonal | NA19983 | 1 |
| GM19983x02PE20438 | chr9 | 79900000 | 81000000 | 1100000 | del_h1 | Subclonal | NA19983 | 1 |
| GM19983x02PE20482 | chr9 | 79900000 | 81000000 | 1100000 | del_h1 | Subclonal | NA19983 | 1 |
| GM19983x02PE20421 | chr14 | 0 | 107043718 | 107043718 | dup_h1 | Singleton | NA19983 | 3 |
| GM19983x02PE20421 | chr15 | 0 | 101991189 | 101991189 | dup_h1 | Singleton | NA19983 | 3 |
| GM19983x02PE20494 | chr16 | 36334460 | 90338345 | 54003885 | del_h2 | Singleton | NA19983 | 1 |
| GM19983x02PE20434 | chrX | 79800000 | 156040895 | 76240895 | del_h1 | Singleton | NA19983 | 1 |
| GM20509Bx01PE20561 | chr9 | 0 | 138394717 | 138394717 | dup_h2 | Singleton | NA20509 | 3 |
| GM20509Bx01PE20579 | chr9 | 0 | 32000000 | 32000000 | dup_h2 | Singleton | NA20509 | 3 |
| GM20509Bx01PE20547 | chr16 | 83900000 | 90338345 | 6438345 | del_h2 | Singleton | NA20509 | 1 |
| GM20509Bx01PE20549 | chr16 | 36334460 | 90338345 | 54003885 | del_h1 | Singleton | NA20509 | 1 |
| GM20509Bx01PE20526 | chr18 | 0 | 80373285 | 80373285 | del_h2 | Singleton | NA20509 | 1 |
| GM20847Bx02PE20479 | chr2 | 0 | 92188145 | 92188145 | del_h2 | Singleton | NA20847 | 1 |
| GM20847Bx02PE20423 | chr13 | 0 | 114364328 | 114364328 | dup_h2 | Singleton | NA20847 | 3 |
| GM20847Bx02PE20418 | chr15 | 0 | 101991189 | 101991189 | dup_h1 | Singleton | NA20847 | 3 |
| GM20847Bx02PE20483 | chr16 | 0 | 36311158 | 36311158 | del_h1 | Singleton | NA20847 | 1 |
| HG00096x02PE20333 | chr1 | 0 | 14600000 | 14600000 | del_h1 | Singleton | HG00096 | 1 |
| HG00096x02PE20333 | chr1 | 14600000 | 50300000 | 35700000 | dup_h2 | Singleton | HG00096 | 3 |
| HG00096x02PE20359 | chr1 | 168900000 | 248956422 | 80056422 | del_h2 | Singleton | HG00096 | 1 |
| HG00096x02PE20379 | chr9 | 45518558 | 138394717 | 92876159 | dup_h2 | Singleton | HG00096 | 3 |
| HG00171Ax02PE20414 | chr1 | 124932724 | 248956422 | 124023698 | del_h1 | Singleton | HG00171 | 1 |
| HG00171Ax02PE20419 | chr1 | 124932724 | 248956422 | 124023698 | del_hom | Singleton | HG00171 | 0 |
| HG00171Ax02PE20464 | chr1 | 124932724 | 248956422 | 124023698 | dup_h2 | Singleton | HG00171 | 3 |
| HG00171Ax02PE20436 | chr4 | 114200000 | 190214555 | 76014555 | dup_h1 | Singleton | HG00171 | 3 |
| HG00171Ax02PE20455 | chr7 | 82800000 | 159345973 | 76545973 | del_h1 | Singleton | HG00171 | 1 |
| HG00171Ax02PE20481 | chr12 | 94100000 | 110200000 | 16100000 | del_h1 | Singleton | HG00171 | 1 |

| | | | | | | | | |
|---|---|---|---|---|---|---|---|---|
| HG00171Ax02PE20429 | chr13 | 0 | 114364328 | 114364328 | del_h1 | Singleton | HG00171 | 1 |
| HG00171Ax02PE20404 | chr16 | 36334460 | 90338345 | 54003885 | dup_hom | Singleton | HG00171 | 3 |
| HG00171Ax02PE20414 | chr16 | 36334460 | 90338345 | 54003885 | del_h1 | Singleton | HG00171 | 1 |
| HG00171Ax02PE20464 | chr16 | 36334460 | 90338345 | 54003885 | dup_h1 | Singleton | HG00171 | 3 |
| HG00171Ax02PE20468 | chr17 | 0 | 14200000 | 14200000 | del_h1 | Singleton | HG00171 | 1 |
| HG00171Ax02PE20415 | chrX | 0 | 156040895 | 156040895 | del_h2 | Subclonal | HG00171 | 1 |
| HG00171Ax02PE20468 | chrX | 0 | 156040895 | 156040895 | del_h2 | Subclonal | HG00171 | 1 |
| HG00171Ax02PE20481 | chrX | 77000000 | 156040895 | 79040895 | dup_h2 | Singleton | HG00171 | 3 |
| HG00864x02PE20342 | chr1 | 124932724 | 248956422 | 124023698 | idup_h1 | Singleton | HG00864 | 3 |
| HG00864x02PE20344 | chr1 | 124932724 | 248956422 | 124023698 | dup_hom | Singleton | HG00864 | 4 |
| HG00864x02PE20344 | chr16 | 36334460 | 90338345 | 54003885 | del_hom | Singleton | HG00864 | 0 |
| HG00864x02PE20360 | chr16 | 36334460 | 90338345 | 54003885 | del_h2 | Singleton | HG00864 | 1 |
| HG00864x02PE20304 | chrX | 0 | 156040895 | 156040895 | del_h2 | Subclonal | HG00864 | 1 |
| HG00864x02PE20319 | chrX | 0 | 156040895 | 156040895 | del_h2 | Subclonal | HG00864 | 1 |
| HG00864x02PE20325 | chrX | 0 | 156040895 | 156040895 | del_h2 | Subclonal | HG00864 | 1 |
| HG00864x02PE20330 | chrX | 0 | 156040895 | 156040895 | del_h2 | Subclonal | HG00864 | 1 |
| HG00864x02PE20342 | chrX | 0 | 156040895 | 156040895 | del_h2 | Subclonal | HG00864 | 1 |
| HG00864x02PE20344 | chrX | 0 | 156040895 | 156040895 | del_h2 | Subclonal | HG00864 | 1 |
| HG00864x02PE20350 | chrX | 0 | 156040895 | 156040895 | del_h2 | Subclonal | HG00864 | 1 |
| HG00864x02PE20360 | chrX | 0 | 156040895 | 156040895 | del_h2 | Subclonal | HG00864 | 1 |
| HG00864x02PE20328 | chrX | 142500000 | 156040895 | 13540895 | del_h2 | Singleton | HG00864 | 1 |
| HG01114x02PE20363 | chr16 | 83900000 | 90338345 | 6438345 | del_h1 | Singleton | HG01114 | 1 |
| HG01114x02PE20334 | chr16 | 50800000 | 90338345 | 39538345 | dup_hom | Singleton | HG01114 | 4 |
| HG01596x02PE20571 | chr4 | 65800000 | 73300000 | 7500000 | del_h1 | Singleton | HG01596 | 1 |
| HG01596x02PE20562 | chr6 | 69500000 | 170805979 | 101305979 | del_h2 | Singleton | HG01596 | 1 |
| HG01596x02PE20530 | chr16 | 56300000 | 90338345 | 34038345 | del_h2 | Singleton | HG01596 | 1 |
| HG02011x02PE20541 | chr1 | 124932724 | 248956422 | 124023698 | del_h1 | Subclonal | HG02011 | 1 |
| HG02011x02PE20564 | chr1 | 124932724 | 248956422 | 124023698 | dup_h1 | Singleton | HG02011 | 3 |
| HG02011x02PE20584 | chr1 | 124932724 | 248956422 | 124023698 | del_h1 | Subclonal | HG02011 | 1 |
| HG02011x02PE20512 | chr11 | 72700000 | 73600000 | 900000 | del_h1 | Subclonal | HG02011 | 1 |
| HG02011x02PE20552 | chr11 | 72700000 | 73600000 | 900000 | del_h1 | Subclonal | HG02011 | 1 |
| HG02011x02PE20555 | chr11 | 72700000 | 73600000 | 900000 | del_h1 | Subclonal | HG02011 | 1 |
| HG02011x02PE20573 | chr11 | 72700000 | 73600000 | 900000 | del_h1 | Subclonal | HG02011 | 1 |
| HG02011x02PE20536 | chr16 | 36334460 | 90338345 | 54003885 | del_h1 | Subclonal | HG02011 | 1 |
| HG02011x02PE20575 | chr16 | 36334460 | 90338345 | 54003885 | dup_h2 | Singleton | HG02011 | 3 |
| HG02011x02PE20584 | chr16 | 36334460 | 90338345 | 54003885 | del_h1 | Subclonal | HG02011 | 1 |
| HG02011x02PE20594 | chr16 | 36334460 | 90338345 | 54003885 | dup_h1 | Singleton | HG02011 | 3 |
| HG02492x02PE20424 | chr11 | 90100000 | 135086622 | 44986622 | dup_hom | Singleton | HG02492 | 4 |

| | | | | | | | | |
|---|---|---|---|---|---|---|---|---|
| HG02587x02PE20346 | chr1 | 124932724 | 248956422 | 124023698 | dup_h1 | Singleton | HG02587 | 3 |
| HG02587x02PE20306 | chr3 | 93655574 | 198295559 | 104639985 | del_h2 | Singleton | HG02587 | 1 |
| HG02587x02PE20326 | chr16 | 36334460 | 90338345 | 54003885 | del_h1 | Singleton | HG02587 | 1 |
| HG02818x02PE20303 | chr1 | 124932724 | 248956422 | 124023698 | del_h2 | Singleton | HG02818 | 1 |
| HG02818x02PE20328 | chr1 | 234600000 | 248956422 | 14356422 | del_h2 | Singleton | HG02818 | 1 |
| HG02818x02PE20368 | chr1 | 124932724 | 248956422 | 124023698 | del_h1 | Singleton | HG02818 | 1 |
| HG02818x02PE20328 | chr2 | 0 | 8000000 | 8000000 | del_h2 | Singleton | HG02818 | 1 |
| HG02818x02PE20328 | chr3 | 0 | 86200000 | 86200000 | idup_h2 | Singleton | HG02818 | 3 |
| HG02818x02PE20328 | chr4 | 163500000 | 190214555 | 26714555 | del_h2 | Singleton | HG02818 | 1 |
| HG02818x02PE20362 | chr4 | 83800000 | 90100000 | 6300000 | del_h1 | Singleton | HG02818 | 1 |
| HG02818x02PE20362 | chr4 | 90100000 | 190214555 | 100114555 | del_hom | Singleton | HG02818 | 0 |
| HG02818x02PE20307 | chr9 | 45518558 | 138394717 | 92876159 | idup_h1 | Singleton | HG02818 | 3 |
| HG02818x02PE20302 | chr16 | 36334460 | 90338345 | 54003885 | del_h1 | Subclonal | HG02818 | 1 |
| HG02818x02PE20303 | chr16 | 36334460 | 90338345 | 54003885 | dup_h2 | Singleton | HG02818 | 3 |
| HG02818x02PE20304 | chr16 | 36334460 | 90338345 | 54003885 | del_h1 | Subclonal | HG02818 | 1 |
| HG02818x02PE20358 | chr16 | 36334460 | 90338345 | 54003885 | del_h1 | Subclonal | HG02818 | 1 |
| HG02818x02PE20368 | chr16 | 36334460 | 90338345 | 54003885 | del_h2 | Singleton | HG02818 | 1 |
| HG02818x02PE20369 | chr16 | 36334460 | 90338345 | 54003885 | del_h1 | Subclonal | HG02818 | 1 |
| HG02818x02PE20373 | chr16 | 36334460 | 90338345 | 54003885 | del_h1 | Subclonal | HG02818 | 1 |
| HG02818x02PE20328 | chr19 | 0 | 4100000 | 4100000 | del_h2 | Singleton | HG02818 | 1 |
| HG02818x02PE20328 | chrX | 0 | 2800000 | 2800000 | del_h2 | Singleton | HG02818 | 1 |
| HG02818x02PE20328 | chrX | 110500000 | 156040895 | 45540895 | idup_h2 | Singleton | HG02818 | 3 |
| HG03009x02PE20318 | chr1 | 124932724 | 248956422 | 124023698 | del_h1 | Subclonal | HG03009 | 1 |
| HG03009x02PE20348 | chr1 | 124932724 | 248956422 | 124023698 | del_h1 | Subclonal | HG03009 | 1 |
| HG03009x02PE20353 | chr1 | 124932724 | 248956422 | 124023698 | dup_h1 | Singleton | HG03009 | 3 |
| HG03009x02PE20369 | chr1 | 124932724 | 248956422 | 124023698 | del_h1 | Subclonal | HG03009 | 1 |
| HG03009x02PE20388 | chr1 | 124932724 | 248956422 | 124023698 | del_h1 | Subclonal | HG03009 | 1 |
| HG03009x02PE20355 | chr2 | 230200000 | 242193529 | 11993529 | del_h2 | Singleton | HG03009 | 1 |
| HG03009x02PE20338 | chr4 | 105800000 | 190214555 | 84414555 | dup_h1 | Singleton | HG03009 | 3 |
| HG03009x02PE20369 | chr6 | 0 | 28600000 | 28600000 | del_h1 | Singleton | HG03009 | 1 |
| HG03009x02PE20369 | chr9 | 68300000 | 106800000 | 38500000 | dup_h1 | Singleton | HG03009 | 3 |
| HG03009x02PE20365 | chr11 | 114800000 | 135086622 | 20286622 | dup_h2 | Singleton | HG03009 | 3 |
| HG03009x02PE20318 | chr16 | 36334460 | 90338345 | 54003885 | dup_h2 | Subclonal | HG03009 | 3 |
| HG03009x02PE20323 | chr16 | 36334460 | 90338345 | 54003885 | dup_h1 | Subclonal | HG03009 | 3 |
| HG03009x02PE20348 | chr16 | 36334460 | 90338345 | 54003885 | del_h2 | Singleton | HG03009 | 1 |
| HG03009x02PE20369 | chr16 | 36334460 | 90338345 | 54003885 | del_h1 | Subclonal | HG03009 | 1 |
| HG03009x02PE20371 | chr16 | 36334460 | 90338345 | 54003885 | dup_h1 | Subclonal | HG03009 | 3 |
| HG03009x02PE20379 | chr16 | 36334460 | 90338345 | 54003885 | dup_h1 | Subclonal | HG03009 | 3 |

| | | | | | | | | |
|---|---|---|---|---|---|---|---|---|
| HG03009x02PE20387 | chr16 | 36334460 | 90338345 | 54003885 | dup_h2 | Subclonal | HG03009 | 3 |
| HG03009x02PE20388 | chr16 | 36334460 | 90338345 | 54003885 | del_h1 | Subclonal | HG03009 | 1 |
| HG03065x02PE20502 | chr16 | 36334460 | 90338345 | 54003885 | del_h1 | Singleton | HG03065 | 1 |
| HG03065x02PE20511 | chr16 | 36334460 | 90338345 | 54003885 | del_h2 | Subclonal | HG03065 | 1 |
| HG03065x02PE20527 | chr16 | 36334460 | 90338345 | 54003885 | del_h2 | Subclonal | HG03065 | 1 |
| HG03065x02PE20535 | chr16 | 36334460 | 90338345 | 54003885 | dup_h2 | Singleton | HG03065 | 3 |
| HG03065x02PE20559 | chr16 | 36334460 | 90338345 | 54003885 | del_h2 | Subclonal | HG03065 | 1 |
| HG03065x02PE20536 | chr18 | 0 | 15460899 | 15460899 | del_h1 | Singleton | HG03065 | 1 |
| HG03125x02PE20351 | chr1 | 0 | 248956422 | 248956422 | del_h2 | Singleton | HG03125 | 1 |
| HG03125x02PE20327 | chr2 | 124000000 | 242193529 | 118193529 | del_h1 | Singleton | HG03125 | 1 |
| HG03125x02PE20348 | chr4 | 66900000 | 190214555 | 123314555 | del_h2 | Singleton | HG03125 | 1 |
| HG03125x02PE20313 | chr7 | 0 | 58169653 | 58169653 | del_h2 | Singleton | HG03125 | 1 |
| HG03125x02PE20362 | chr8 | 79400000 | 145138636 | 65738636 | del_h1 | Singleton | HG03125 | 1 |
| HG03125x02PE20381 | chr8 | 0 | 21100000 | 21100000 | del_h1 | Singleton | HG03125 | 1 |
| HG03125x02PE20311 | chr9 | 45518558 | 138394717 | 92876159 | del_h1 | Subclonal | HG03125 | 1 |
| HG03125x02PE20316 | chr9 | 114700000 | 138394717 | 23694717 | dup_h2 | Singleton | HG03125 | 3 |
| HG03125x02PE20356 | chr9 | 0 | 20800000 | 20800000 | idup_h1 | Singleton | HG03125 | 3 |
| HG03125x02PE20360 | chr9 | 45518558 | 138394717 | 92876159 | del_h1 | Subclonal | HG03125 | 1 |
| HG03125x02PE20348 | chr11 | 128300000 | 135086622 | 6786622 | idup_h1 | Singleton | HG03125 | 3 |
| HG03125x02PE20316 | chr12 | 54500000 | 133275309 | 78775309 | idup_h2 | Singleton | HG03125 | 3 |
| HG03125x02PE20335 | chr12 | 0 | 34769407 | 34769407 | idup_h1 | Singleton | HG03125 | 3 |
| HG03125x02PE20351 | chr16 | 0 | 90338345 | 90338345 | del_h2 | Singleton | HG03125 | 1 |
| HG03125x02PE20316 | chrX | 96600000 | 112800000 | 16200000 | del_h2 | Singleton | HG03125 | 1 |
| HG03371x02PE20591 | chr1 | 0 | 248956422 | 248956422 | dup_h2 | Singleton | HG03371 | 3 |
| HG03371x02PE20591 | chr3 | 0 | 198295559 | 198295559 | dup_h1 | Singleton | HG03371 | 3 |
| HG03371x02PE20525 | chr6 | 0 | 170805979 | 170805979 | del_h2 | Singleton | HG03371 | 1 |
| HG03371x02PE20518 | chr16 | 0 | 18300000 | 18300000 | del_h2 | Singleton | HG03371 | 1 |
| HG03371x02PE20544 | chr16 | 36334460 | 90338345 | 54003885 | idup_h2 | Singleton | HG03371 | 3 |
| HG03371x02PE20591 | chr19 | 0 | 58617616 | 58617616 | idup_h2 | Singleton | HG03371 | 3 |
| HG03371x02PE20591 | chr21 | 0 | 46709983 | 46709983 | idup_h1 | Singleton | HG03371 | 3 |
| HG03486x02PE20510 | chr1 | 232000000 | 248956422 | 16956422 | del_h1 | Singleton | HG03486 | 1 |
| HG03486x02PE20512 | chr1 | 157600000 | 248956422 | 91356422 | idup_h1 | Singleton | HG03486 | 3 |
| HG03486x02PE20543 | chr6 | 129700000 | 170805979 | 41105979 | del_h1 | Singleton | HG03486 | 1 |
| HG03486x02PE20576 | chr9 | 68300000 | 104200000 | 35900000 | dup_h1 | Singleton | HG03486 | 3 |
| HG03486x02PE20576 | chr9 | 104200000 | 138394717 | 34194717 | del_h2 | Singleton | HG03486 | 1 |
| HG03486x02PE20541 | chr11 | 114600000 | 135086622 | 20486622 | del_h2 | Singleton | HG03486 | 1 |
| HG03486x02PE20510 | chr12 | 65400000 | 133275309 | 67875309 | idup_h2 | Singleton | HG03486 | 3 |
| HG03486x02PE20510 | chr13 | 82000000 | 114364328 | 32364328 | dup_h1 | Singleton | HG03486 | 3 |

| | | | | | | | | |
|---|---|---|---|---|---|---|---|---|
| HG03486x02PE20569 | chr16 | 36334460 | 90338345 | 54003885 | del_h2 | Singleton | HG03486 | 1 |
| HG03486x02PE20579 | chr16 | 22800000 | 66600000 | 43800000 | idup_h2 | Singleton | HG03486 | 3 |
| HG03683x01PE20449 | chr2 | 195000000 | 242193529 | 47193529 | del_h1 | Singleton | HG03683 | 1 |
| HG03683x01PE20423 | chr8 | 0 | 44033744 | 44033744 | del_h1 | Singleton | HG03683 | 1 |
| HG03683x01PE20423 | chr8 | 45877265 | 145138636 | 99261371 | dup_h2 | Singleton | HG03683 | 3 |
| HG03683x01PE20445 | chr9 | 45518558 | 138394717 | 92876159 | del_h2 | Singleton | HG03683 | 1 |
| HG03683x01PE20476 | chr9 | 45518558 | 138394717 | 92876159 | del_h1 | Singleton | HG03683 | 1 |
| HG03683x01PE20490 | chr13 | 72000000 | 114364328 | 42364328 | del_h2 | Singleton | HG03683 | 1 |
| HG03683x01PE20419 | chr15 | 74300000 | 101991189 | 27691189 | dup_h2 | Singleton | HG03683 | 3 |
| HG03683x01PE20406 | chr16 | 36334460 | 90338345 | 54003885 | del_h2 | Singleton | HG03683 | 1 |
| HG03683x01PE20454 | chr16 | 36334460 | 90338345 | 54003885 | dup_h1 | Singleton | HG03683 | 3 |
| HG03683x01PE20435 | chr17 | 0 | 1800000 | 1800000 | del_h1 | Singleton | HG03683 | 1 |
| HG03683x01PE20435 | chr17 | 1800000 | 21600000 | 19800000 | dup_h2 | Singleton | HG03683 | 3 |
| HG03732x02PE20540 | chr2 | 0 | 23300000 | 23300000 | del_h2 | Singleton | HG03732 | 1 |
| HG03732x02PE20583 | chr2 | 184500000 | 242193529 | 57693529 | del_h2 | Singleton | HG03732 | 1 |
| HG03732x02PE20540 | chr4 | 0 | 27400000 | 27400000 | del_h1 | Singleton | HG03732 | 1 |
| HG03732x02PE20540 | chr5 | 122100000 | 181538259 | 59438259 | dup_h1 | Singleton | HG03732 | 3 |
| HG03732x02PE20589 | chr8 | 0 | 145138636 | 145138636 | del_h1 | Singleton | HG03732 | 1 |
| HG03732x02PE20565 | chr16 | 36334460 | 90338345 | 54003885 | dup_h2 | Singleton | HG03732 | 3 |

**Table S3C. List of SVs found in the NA20509**

| Cell | Chromosome | Start | End | Size (bp) | SV Class | Singleton/Sub | Sample | Copy Number estimation |
|---|---|---|---|---|---|---|---|---|
| GM20509Bx01PE20561 | chr17 | 19200000 | 21700000 | 2500000 | dup_h2 | Subclonal | NA20509 | 5 |
| GM20509Bx01PE20514 | chr17 | 19200000 | 21700000 | 2500000 | dup_h2 | Subclonal | NA20509 | 4 |
| GM20509Bx01PE20502 | chr17 | 19200000 | 21700000 | 2500000 | dup_h2 | Subclonal | NA20509 | 3 |
| GM20509Bx01PE20503 | chr17 | 19200000 | 21700000 | 2500000 | dup_h2 | Subclonal | NA20509 | 3 |
| GM20509Bx01PE20506 | chr17 | 19200000 | 21700000 | 2500000 | dup_h2 | Subclonal | NA20509 | 3 |
| GM20509Bx01PE20508 | chr17 | 19200000 | 21700000 | 2500000 | dup_h2 | Subclonal | NA20509 | 3 |
| GM20509Bx01PE20509 | chr17 | 19200000 | 21700000 | 2500000 | dup_h2 | Subclonal | NA20509 | 3 |
| GM20509Bx01PE20510 | chr17 | 19200000 | 21700000 | 2500000 | dup_h2 | Subclonal | NA20509 | 3 |
| GM20509Bx01PE20511 | chr17 | 19200000 | 21700000 | 2500000 | dup_h2 | Subclonal | NA20509 | 3 |
| GM20509Bx01PE20517 | chr17 | 19200000 | 21700000 | 2500000 | dup_h2 | Subclonal | NA20509 | 3 |
| GM20509Bx01PE20518 | chr17 | 19200000 | 21700000 | 2500000 | dup_h2 | Subclonal | NA20509 | 3 |
| GM20509Bx01PE20519 | chr17 | 19200000 | 21700000 | 2500000 | dup_h2 | Subclonal | NA20509 | 3 |
| GM20509Bx01PE20525 | chr17 | 19200000 | 21700000 | 2500000 | dup_h2 | Subclonal | NA20509 | 3 |
| GM20509Bx01PE20526 | chr17 | 19200000 | 21700000 | 2500000 | dup_h2 | Subclonal | NA20509 | 3 |
| GM20509Bx01PE20529 | chr17 | 19200000 | 21700000 | 2500000 | dup_h2 | Subclonal | NA20509 | 3 |
| GM20509Bx01PE20530 | chr17 | 19200000 | 21700000 | 2500000 | dup_h2 | Subclonal | NA20509 | 3 |
| GM20509Bx01PE20531 | chr17 | 19200000 | 21700000 | 2500000 | dup_h2 | Subclonal | NA20509 | 3 |
| GM20509Bx01PE20533 | chr17 | 19200000 | 21700000 | 2500000 | dup_h2 | Subclonal | NA20509 | 3 |
| GM20509Bx01PE20535 | chr17 | 19200000 | 21700000 | 2500000 | dup_h2 | Subclonal | NA20509 | 3 |
| GM20509Bx01PE20536 | chr17 | 19200000 | 21700000 | 2500000 | dup_h2 | Subclonal | NA20509 | 3 |
| GM20509Bx01PE20539 | chr17 | 19200000 | 21700000 | 2500000 | dup_h2 | Subclonal | NA20509 | 3 |
| GM20509Bx01PE20541 | chr17 | 19200000 | 21700000 | 2500000 | dup_h2 | Subclonal | NA20509 | 3 |
| GM20509Bx01PE20545 | chr17 | 19200000 | 21700000 | 2500000 | dup_h2 | Subclonal | NA20509 | 3 |
| GM20509Bx01PE20546 | chr17 | 19200000 | 21700000 | 2500000 | dup_h2 | Subclonal | NA20509 | 3 |
| GM20509Bx01PE20547 | chr17 | 19200000 | 21700000 | 2500000 | dup_h2 | Subclonal | NA20509 | 3 |
| GM20509Bx01PE20552 | chr17 | 19200000 | 21700000 | 2500000 | dup_h2 | Subclonal | NA20509 | 3 |
| GM20509Bx01PE20555 | chr17 | 19200000 | 21700000 | 2500000 | dup_h2 | Subclonal | NA20509 | 3 |
| GM20509Bx01PE20556 | chr17 | 19200000 | 21700000 | 2500000 | dup_h2 | Subclonal | NA20509 | 3 |
| GM20509Bx01PE20557 | chr17 | 19200000 | 21700000 | 2500000 | dup_h2 | Subclonal | NA20509 | 3 |
| GM20509Bx01PE20560 | chr17 | 19200000 | 21700000 | 2500000 | dup_h2 | Subclonal | NA20509 | 3 |
| GM20509Bx01PE20562 | chr17 | 19200000 | 21700000 | 2500000 | dup_h2 | Subclonal | NA20509 | 3 |
| GM20509Bx01PE20565 | chr17 | 19200000 | 21700000 | 2500000 | dup_h2 | Subclonal | NA20509 | 3 |
| GM20509Bx01PE20566 | chr17 | 19200000 | 21700000 | 2500000 | dup_h2 | Subclonal | NA20509 | 3 |

| | | | | | | | | |
|---|---|---|---|---|---|---|---|---|
| GM20509Bx01PE20568 | chr17 | 19200000 | 21700000 | 2500000 | dup_h2 | Subclonal | NA20509 | 3 |
| GM20509Bx01PE20569 | chr17 | 19200000 | 21700000 | 2500000 | dup_h2 | Subclonal | NA20509 | 3 |
| GM20509Bx01PE20576 | chr17 | 19200000 | 21700000 | 2500000 | dup_h2 | Subclonal | NA20509 | 3 |
| GM20509Bx01PE20581 | chr17 | 19200000 | 21700000 | 2500000 | dup_h2 | Subclonal | NA20509 | 3 |
| GM20509Bx01PE20587 | chr17 | 19200000 | 21700000 | 2500000 | dup_h2 | Subclonal | NA20509 | 3 |
| GM20509Bx01PE20588 | chr17 | 19200000 | 21700000 | 2500000 | dup_h2 | Subclonal | NA20509 | 3 |
| GM20509Bx01PE20520 | chr17 | 19200000 | 21700000 | 2500000 | dup_h2 | Subclonal | NA20509 | 2 |
| GM20509Bx01PE20502 | chr17 | 0 | 19200000 | 19200000 | del_h2 | Subclonal | NA20509 | - |
| GM20509Bx01PE20503 | chr17 | 0 | 19200000 | 19200000 | del_h2 | Subclonal | NA20509 | - |
| GM20509Bx01PE20506 | chr17 | 0 | 19200000 | 19200000 | del_h2 | Subclonal | NA20509 | - |
| GM20509Bx01PE20508 | chr17 | 0 | 19200000 | 19200000 | del_h2 | Subclonal | NA20509 | - |
| GM20509Bx01PE20509 | chr17 | 0 | 19200000 | 19200000 | del_h2 | Subclonal | NA20509 | - |
| GM20509Bx01PE20510 | chr17 | 0 | 19200000 | 19200000 | del_h2 | Subclonal | NA20509 | - |
| GM20509Bx01PE20511 | chr17 | 0 | 19200000 | 19200000 | del_h2 | Subclonal | NA20509 | - |
| GM20509Bx01PE20514 | chr17 | 0 | 19200000 | 19200000 | del_h2 | Subclonal | NA20509 | - |
| GM20509Bx01PE20517 | chr17 | 0 | 19200000 | 19200000 | del_h2 | Subclonal | NA20509 | - |
| GM20509Bx01PE20518 | chr17 | 0 | 19200000 | 19200000 | del_h2 | Subclonal | NA20509 | - |
| GM20509Bx01PE20519 | chr17 | 0 | 19200000 | 19200000 | del_h2 | Subclonal | NA20509 | - |
| GM20509Bx01PE20520 | chr17 | 0 | 19200000 | 19200000 | del_h2 | Subclonal | NA20509 | - |
| GM20509Bx01PE20525 | chr17 | 0 | 19200000 | 19200000 | del_h2 | Subclonal | NA20509 | - |
| GM20509Bx01PE20526 | chr17 | 0 | 19200000 | 19200000 | del_h2 | Subclonal | NA20509 | - |
| GM20509Bx01PE20529 | chr17 | 0 | 19200000 | 19200000 | del_h2 | Subclonal | NA20509 | - |
| GM20509Bx01PE20530 | chr17 | 0 | 19200000 | 19200000 | del_h2 | Subclonal | NA20509 | - |
| GM20509Bx01PE20531 | chr17 | 0 | 19200000 | 19200000 | del_h2 | Subclonal | NA20509 | - |
| GM20509Bx01PE20533 | chr17 | 0 | 19200000 | 19200000 | del_h2 | Subclonal | NA20509 | - |
| GM20509Bx01PE20535 | chr17 | 0 | 19200000 | 19200000 | del_h2 | Subclonal | NA20509 | - |
| GM20509Bx01PE20536 | chr17 | 0 | 19200000 | 19200000 | del_h2 | Subclonal | NA20509 | - |
| GM20509Bx01PE20539 | chr17 | 0 | 19200000 | 19200000 | del_h2 | Subclonal | NA20509 | - |
| GM20509Bx01PE20541 | chr17 | 0 | 19200000 | 19200000 | del_h2 | Subclonal | NA20509 | - |
| GM20509Bx01PE20545 | chr17 | 0 | 19200000 | 19200000 | del_h2 | Subclonal | NA20509 | - |
| GM20509Bx01PE20546 | chr17 | 0 | 19200000 | 19200000 | del_h2 | Subclonal | NA20509 | - |
| GM20509Bx01PE20547 | chr17 | 0 | 19200000 | 19200000 | del_h2 | Subclonal | NA20509 | - |
| GM20509Bx01PE20552 | chr17 | 0 | 19200000 | 19200000 | del_h2 | Subclonal | NA20509 | - |
| GM20509Bx01PE20555 | chr17 | 0 | 19200000 | 19200000 | del_h2 | Subclonal | NA20509 | - |
| GM20509Bx01PE20556 | chr17 | 0 | 19200000 | 19200000 | del_h2 | Subclonal | NA20509 | - |
| GM20509Bx01PE20557 | chr17 | 0 | 19200000 | 19200000 | del_h2 | Subclonal | NA20509 | - |
| GM20509Bx01PE20560 | chr17 | 0 | 19200000 | 19200000 | del_h2 | Subclonal | NA20509 | - |

| | | | | | | | | |
|---|---|---|---|---|---|---|---|---|
| GM20509Bx01PE20561 | chr17 | 0 | 19200000 | 19200000 | del_h2 | Subclonal | NA20509 | - |
| GM20509Bx01PE20562 | chr17 | 0 | 19200000 | 19200000 | del_h2 | Subclonal | NA20509 | - |
| GM20509Bx01PE20565 | chr17 | 0 | 19200000 | 19200000 | del_h2 | Subclonal | NA20509 | - |
| GM20509Bx01PE20566 | chr17 | 0 | 19200000 | 19200000 | del_h2 | Subclonal | NA20509 | - |
| GM20509Bx01PE20568 | chr17 | 0 | 19200000 | 19200000 | del_h2 | Subclonal | NA20509 | - |
| GM20509Bx01PE20569 | chr17 | 0 | 19200000 | 19200000 | del_h2 | Subclonal | NA20509 | - |
| GM20509Bx01PE20576 | chr17 | 0 | 19200000 | 19200000 | del_h2 | Subclonal | NA20509 | - |
| GM20509Bx01PE20581 | chr17 | 0 | 19200000 | 19200000 | del_h2 | Subclonal | NA20509 | - |
| GM20509Bx01PE20587 | chr17 | 0 | 19200000 | 19200000 | del_h2 | Subclonal | NA20509 | - |
| GM20509Bx01PE20588 | chr17 | 0 | 19200000 | 19200000 | del_h2 | Subclonal | NA20509 | - |
| GM20509Bx01PE20502 | chr17 | 19200000 | 21700000 | 2500000 | idup_h2 | Subclonal | NA20509 | - |
| GM20509Bx01PE20503 | chr17 | 19200000 | 21700000 | 2500000 | idup_h2 | Subclonal | NA20509 | - |
| GM20509Bx01PE20506 | chr17 | 19200000 | 21700000 | 2500000 | idup_h2 | Subclonal | NA20509 | - |
| GM20509Bx01PE20508 | chr17 | 19200000 | 21700000 | 2500000 | idup_h2 | Subclonal | NA20509 | - |
| GM20509Bx01PE20509 | chr17 | 19200000 | 21700000 | 2500000 | idup_h2 | Subclonal | NA20509 | - |
| GM20509Bx01PE20511 | chr17 | 19200000 | 21700000 | 2500000 | idup_h2 | Subclonal | NA20509 | - |
| GM20509Bx01PE20517 | chr17 | 19200000 | 21700000 | 2500000 | idup_h2 | Subclonal | NA20509 | - |
| GM20509Bx01PE20518 | chr17 | 19200000 | 21700000 | 2500000 | idup_h2 | Subclonal | NA20509 | - |
| GM20509Bx01PE20519 | chr17 | 19200000 | 21700000 | 2500000 | idup_h2 | Subclonal | NA20509 | - |
| GM20509Bx01PE20525 | chr17 | 19200000 | 21700000 | 2500000 | idup_h2 | Subclonal | NA20509 | - |
| GM20509Bx01PE20526 | chr17 | 19200000 | 21700000 | 2500000 | idup_h2 | Subclonal | NA20509 | - |
| GM20509Bx01PE20529 | chr17 | 19200000 | 21700000 | 2500000 | idup_h2 | Subclonal | NA20509 | - |
| GM20509Bx01PE20530 | chr17 | 19200000 | 21700000 | 2500000 | idup_h2 | Subclonal | NA20509 | - |
| GM20509Bx01PE20531 | chr17 | 19200000 | 21700000 | 2500000 | idup_h2 | Subclonal | NA20509 | - |
| GM20509Bx01PE20533 | chr17 | 19200000 | 21700000 | 2500000 | idup_h2 | Subclonal | NA20509 | - |
| GM20509Bx01PE20535 | chr17 | 19200000 | 21700000 | 2500000 | idup_h2 | Subclonal | NA20509 | - |
| GM20509Bx01PE20536 | chr17 | 19200000 | 21700000 | 2500000 | idup_h2 | Subclonal | NA20509 | - |
| GM20509Bx01PE20539 | chr17 | 19200000 | 21700000 | 2500000 | idup_h2 | Subclonal | NA20509 | - |
| GM20509Bx01PE20541 | chr17 | 19200000 | 21700000 | 2500000 | idup_h2 | Subclonal | NA20509 | - |
| GM20509Bx01PE20545 | chr17 | 19200000 | 21700000 | 2500000 | idup_h2 | Subclonal | NA20509 | - |
| GM20509Bx01PE20546 | chr17 | 19200000 | 21700000 | 2500000 | idup_h2 | Subclonal | NA20509 | - |
| GM20509Bx01PE20547 | chr17 | 19200000 | 21700000 | 2500000 | idup_h2 | Subclonal | NA20509 | - |
| GM20509Bx01PE20552 | chr17 | 19200000 | 21700000 | 2500000 | idup_h2 | Subclonal | NA20509 | - |
| GM20509Bx01PE20557 | chr17 | 19200000 | 21700000 | 2500000 | idup_h2 | Subclonal | NA20509 | - |
| GM20509Bx01PE20560 | chr17 | 19200000 | 21700000 | 2500000 | idup_h2 | Subclonal | NA20509 | - |
| GM20509Bx01PE20561 | chr17 | 19200000 | 21700000 | 2500000 | idup_h2 | Subclonal | NA20509 | - |
| GM20509Bx01PE20562 | chr17 | 19200000 | 21700000 | 2500000 | idup_h2 | Subclonal | NA20509 | - |

| | | | | | | | | |
|---|---|---|---|---|---|---|---|---|
| GM20509Bx01PE20565 | chr17 | 19200000 | 21700000 | 2500000 | idup_h2 | Subclonal | NA20509 | - |
| GM20509Bx01PE20566 | chr17 | 19200000 | 21700000 | 2500000 | idup_h2 | Subclonal | NA20509 | - |
| GM20509Bx01PE20569 | chr17 | 19200000 | 21700000 | 2500000 | idup_h2 | Subclonal | NA20509 | - |
| GM20509Bx01PE20576 | chr17 | 19200000 | 21700000 | 2500000 | idup_h2 | Subclonal | NA20509 | - |
| GM20509Bx01PE20581 | chr17 | 19200000 | 21700000 | 2500000 | idup_h2 | Subclonal | NA20509 | - |
| GM20509Bx01PE20587 | chr17 | 19200000 | 21700000 | 2500000 | idup_h2 | Subclonal | NA20509 | - |
| GM20509Bx01PE20588 | chr17 | 19200000 | 21700000 | 2500000 | idup_h2 | Subclonal | NA20509 | - |
| GM20509Bx01PE20561 | chr5 | 132700000 | 181538259 | 48838259 | dup_h2 | Subclonal | NA20509 | - |
| GM20509Bx01PE20514 | chr5 | 132700000 | 181538259 | 48838259 | dup_h2 | Subclonal | NA20509 | - |
| GM20509Bx01PE20502 | chr5 | 132700000 | 181538259 | 48838259 | dup_h2 | Subclonal | NA20509 | - |
| GM20509Bx01PE20503 | chr5 | 132700000 | 181538259 | 48838259 | dup_h2 | Subclonal | NA20509 | - |
| GM20509Bx01PE20506 | chr5 | 132700000 | 181538259 | 48838259 | dup_h2 | Subclonal | NA20509 | - |
| GM20509Bx01PE20508 | chr5 | 132700000 | 181538259 | 48838259 | dup_h2 | Subclonal | NA20509 | - |
| GM20509Bx01PE20509 | chr5 | 132700000 | 181538259 | 48838259 | dup_h2 | Subclonal | NA20509 | - |
| GM20509Bx01PE20510 | chr5 | 132700000 | 181538259 | 48838259 | dup_h2 | Subclonal | NA20509 | - |
| GM20509Bx01PE20511 | chr5 | 132700000 | 181538259 | 48838259 | dup_h2 | Subclonal | NA20509 | - |
| GM20509Bx01PE20517 | chr5 | 132700000 | 181538259 | 48838259 | dup_h2 | Subclonal | NA20509 | - |
| GM20509Bx01PE20518 | chr5 | 132700000 | 181538259 | 48838259 | dup_h2 | Subclonal | NA20509 | - |
| GM20509Bx01PE20519 | chr5 | 132700000 | 181538259 | 48838259 | dup_h2 | Subclonal | NA20509 | - |
| GM20509Bx01PE20525 | chr5 | 132700000 | 181538259 | 48838259 | dup_h2 | Subclonal | NA20509 | - |
| GM20509Bx01PE20526 | chr5 | 132700000 | 181538259 | 48838259 | dup_h2 | Subclonal | NA20509 | - |
| GM20509Bx01PE20529 | chr5 | 132700000 | 181538259 | 48838259 | dup_h2 | Subclonal | NA20509 | - |
| GM20509Bx01PE20530 | chr5 | 132700000 | 181538259 | 48838259 | dup_h2 | Subclonal | NA20509 | - |
| GM20509Bx01PE20531 | chr5 | 132700000 | 181538259 | 48838259 | dup_h2 | Subclonal | NA20509 | - |
| GM20509Bx01PE20533 | chr5 | 132700000 | 181538259 | 48838259 | dup_h2 | Subclonal | NA20509 | - |
| GM20509Bx01PE20535 | chr5 | 132700000 | 181538259 | 48838259 | dup_h2 | Subclonal | NA20509 | - |
| GM20509Bx01PE20536 | chr5 | 132700000 | 181538259 | 48838259 | dup_h2 | Subclonal | NA20509 | - |
| GM20509Bx01PE20539 | chr5 | 132700000 | 181538259 | 48838259 | dup_h2 | Subclonal | NA20509 | - |
| GM20509Bx01PE20541 | chr5 | 132700000 | 181538259 | 48838259 | dup_h2 | Subclonal | NA20509 | - |
| GM20509Bx01PE20545 | chr5 | 132700000 | 181538259 | 48838259 | dup_h2 | Subclonal | NA20509 | - |
| GM20509Bx01PE20546 | chr5 | 132700000 | 181538259 | 48838259 | dup_h2 | Subclonal | NA20509 | - |
| GM20509Bx01PE20547 | chr5 | 132700000 | 181538259 | 48838259 | dup_h2 | Subclonal | NA20509 | - |
| GM20509Bx01PE20552 | chr5 | 132700000 | 181538259 | 48838259 | dup_h2 | Subclonal | NA20509 | - |
| GM20509Bx01PE20555 | chr5 | 132700000 | 181538259 | 48838259 | dup_h2 | Subclonal | NA20509 | - |
| GM20509Bx01PE20556 | chr5 | 132700000 | 181538259 | 48838259 | dup_h2 | Subclonal | NA20509 | - |
| GM20509Bx01PE20557 | chr5 | 132700000 | 181538259 | 48838259 | dup_h2 | Subclonal | NA20509 | - |
| GM20509Bx01PE20560 | chr5 | 132700000 | 181538259 | 48838259 | dup_h2 | Subclonal | NA20509 | - |

| | | | | | | | | |
|---|---|---|---|---|---|---|---|---|
| GM20509Bx01PE20562 | chr5 | 132700000 | 181538259 | 48838259 | dup_h2 | Subclonal | NA20509 | - |
| GM20509Bx01PE20565 | chr5 | 132700000 | 181538259 | 48838259 | dup_h2 | Subclonal | NA20509 | - |
| GM20509Bx01PE20566 | chr5 | 132700000 | 181538259 | 48838259 | dup_h2 | Subclonal | NA20509 | - |
| GM20509Bx01PE20568 | chr5 | 132700000 | 181538259 | 48838259 | dup_h2 | Subclonal | NA20509 | - |
| GM20509Bx01PE20569 | chr5 | 132700000 | 181538259 | 48838259 | dup_h2 | Subclonal | NA20509 | - |
| GM20509Bx01PE20576 | chr5 | 132700000 | 181538259 | 48838259 | dup_h2 | Subclonal | NA20509 | - |
| GM20509Bx01PE20581 | chr5 | 132700000 | 181538259 | 48838259 | dup_h2 | Subclonal | NA20509 | - |
| GM20509Bx01PE20587 | chr5 | 132700000 | 181538259 | 48838259 | dup_h2 | Subclonal | NA20509 | - |
| GM20509Bx01PE20588 | chr5 | 132700000 | 181538259 | 48838259 | dup_h2 | Subclonal | NA20509 | - |
| GM20509Bx01PE20520 | chr5 | 132700000 | 181538259 | 48838259 | dup_h2 | Subclonal | NA20509 | - |

**Table S3D. List of SVs found in the CLL_24 sample**

| Cell | Chromosome | Start | End | Size (bp) | SV Class | Singleton/Subclonal |
|---|---|---|---|---|---|---|
| BCLL01_90hp2_PE20356 | chr10 | 97300000 | 104700000 | 7400000 | Del_h2 | singleton |
| BCLL01_90hp2_PE20357 | chr10 | 86600000 | 88700000 | 2100000 | Del_h1 | subclonal |
| BCLL01_90hp2_PE20357 | chr10 | 101600000 | 103400000 | 1800000 | Del_h1 | subclonal |
| BCLL01_90hp2_PE20305 | chr6 | 98100000 | 170805979 | 72705979 | Del_h1 | singleton |
| BCLL01_90hp2_PE20323 | chr10 | 100600000 | 114600000 | 14000000 | Del_h2 | singleton |
| BCLL01_90hp1_PE20456 | chr10 | 101400000 | 133797422 | 32397422 | Del_h1 | Subclonal |
| BCLL01_90hp2_PE20344 | chr10 | 101400000 | 133797422 | 32397422 | Del_h1 | Subclonal |
| BCLL01_90hp1_PE20467 | chr10 | 100900000 | 103000000 | 2100000 | Del_h2 | subclonal |
| BCLL01_90hp1_PE20478 | chr19 | 18000000 | 19500000 | 1500000 | del_h2 | singleton |
| BCLL01_90hp1_PE20478 | chr22 | 36800000 | 42600000 | 5800000 | del_h1 | singleton |
| BCLL01_90hp2_PE20352 | chr10 | 101200000 | 103300000 | 2100000 | Del_h2 | subclonal |
| BCLL01_90hp2_PE20354 | chr7 | 60800000 | 159345973 | 98545973 | Del_h2 | singleton |
| BCLL01_90hp1_PE20423 | chr10 | 101400000 | 133797422 | 32397422 | Del_h1 | subclonal |
| BCLL01_120hp2_PE20406 | chr10 | 100900000 | 110100000 | 9200000 | Del_h2 | singleton |
| BCLL01_120hp2_PE20457 | chr10 | 86600000 | 88700000 | 2100000 | Del_h1 | subclonal |
| BCLL01_120hp2_PE20457 | chr10 | 101600000 | 103500000 | 1900000 | Del_h1 | subclonal |
| BCLL01_120hp2_PE20457 | chr11 | 81300000 | 135086622 | 53786622 | inv_dup_h2 | singleton |
| BCLL01_120hp1_PE20582 | chr10 | 100000000 | 133797422 | 33797422 | Del_h2 | singleton |

**Table S3E. List of SVs found in the AML_1 sample**

| Cell | Chromosome | Start | End | Size (bp) | SV Class | Singleton/Subclonal |
|---|---|---|---|---|---|---|
| AML01_PE20311 | chr21 | 34900000 | 50818468 | 15918468 | translocation | Clonal |
| AML01_PE20311 | chr8 | 92000000 | 145138636 | 53138636 | translocation | Clonal |
| AML01_PE20312 | chr21 | 34900000 | 50818468 | 15918468 | translocation | Clonal |
| AML01_PE20312 | chr8 | 92000000 | 145138636 | 53138636 | translocation | Clonal |
| AML01_PE20314 | chr21 | 34900000 | 50818468 | 15918468 | translocation | Clonal |
| AML01_PE20314 | chr8 | 92000000 | 145138636 | 53138636 | translocation | Clonal |
| AML01_PE20316 | chr21 | 34900000 | 50818468 | 15918468 | translocation | Clonal |
| AML01_PE20316 | chr8 | 92000000 | 145138636 | 53138636 | translocation | Clonal |
| AML01_PE20317 | chr21 | 34900000 | 50818468 | 15918468 | translocation | Clonal |
| AML01_PE20317 | chr8 | 92000000 | 145138636 | 53138636 | translocation | Clonal |
| AML01_PE20319 | chr21 | 34900000 | 50818468 | 15918468 | translocation | Clonal |
| AML01_PE20319 | chr8 | 92000000 | 145138636 | 53138636 | translocation | Clonal |
| AML01_PE20321 | chr21 | 34900000 | 50818468 | 15918468 | translocation | Clonal |
| AML01_PE20321 | chr8 | 92000000 | 145138636 | 53138636 | translocation | Clonal |
| AML01_PE20323 | chr21 | 34900000 | 50818468 | 15918468 | translocation | Clonal |
| AML01_PE20323 | chr8 | 92000000 | 145138636 | 53138636 | translocation | Clonal |
| AML01_PE20327 | chr21 | 34900000 | 50818468 | 15918468 | translocation | Clonal |
| AML01_PE20327 | chr8 | 92000000 | 145138636 | 53138636 | translocation | Clonal |
| AML01_PE20328 | chr21 | 34900000 | 50818468 | 15918468 | translocation | Clonal |
| AML01_PE20328 | chr8 | 92000000 | 145138636 | 53138636 | translocation | Clonal |
| AML01_PE20331 | chr21 | 34900000 | 50818468 | 15918468 | translocation | Clonal |
| AML01_PE20331 | chr8 | 92000000 | 145138636 | 53138636 | translocation | Clonal |
| AML01_PE20332 | chr21 | 34900000 | 50818468 | 15918468 | translocation | Clonal |
| AML01_PE20332 | chr8 | 92000000 | 145138636 | 53138636 | translocation | Clonal |
| AML01_PE20334 | chr21 | 34900000 | 50818468 | 15918468 | translocation | Clonal |
| AML01_PE20334 | chr8 | 92000000 | 145138636 | 53138636 | translocation | Clonal |
| AML01_PE20336 | chr21 | 34900000 | 50818468 | 15918468 | translocation | Clonal |
| AML01_PE20336 | chr8 | 92000000 | 145138636 | 53138636 | translocation | Clonal |

| | | | | | | |
|---|---|---|---|---|---|---|
| AML01_PE20337 | chr21 | 34900000 | 50818468 | 15918468 | translocation | Clonal |
| AML01_PE20337 | chr8 | 92000000 | 145138636 | 53138636 | translocation | Clonal |
| AML01_PE20338 | chr21 | 34900000 | 50818468 | 15918468 | translocation | Clonal |
| AML01_PE20338 | chr8 | 92000000 | 145138636 | 53138636 | translocation | Clonal |
| AML01_PE20339 | chr21 | 34900000 | 50818468 | 15918468 | translocation | Clonal |
| AML01_PE20339 | chr8 | 92000000 | 145138636 | 53138636 | translocation | Clonal |
| AML01_PE20340 | chr21 | 34900000 | 50818468 | 15918468 | translocation | Clonal |
| AML01_PE20340 | chr8 | 92000000 | 145138636 | 53138636 | translocation | Clonal |
| AML01_PE20342 | chr21 | 34900000 | 50818468 | 15918468 | translocation | Clonal |
| AML01_PE20342 | chr8 | 92000000 | 145138636 | 53138636 | translocation | Clonal |
| AML01_PE20343 | chr21 | 34900000 | 50818468 | 15918468 | translocation | Clonal |
| AML01_PE20343 | chr8 | 92000000 | 145138636 | 53138636 | translocation | Clonal |
| AML01_PE20346 | chr21 | 34900000 | 50818468 | 15918468 | translocation | Clonal |
| AML01_PE20346 | chr8 | 92000000 | 145138636 | 53138636 | translocation | Clonal |
| AML01_PE20350 | chr21 | 34900000 | 50818468 | 15918468 | translocation | Clonal |
| AML01_PE20350 | chr8 | 92000000 | 145138636 | 53138636 | translocation | Clonal |
| AML01_PE20351 | chr21 | 34900000 | 50818468 | 15918468 | translocation | Clonal |
| AML01_PE20351 | chr8 | 92000000 | 145138636 | 53138636 | translocation | Clonal |
| AML01_PE20353 | chr21 | 34900000 | 50818468 | 15918468 | translocation | Clonal |
| AML01_PE20353 | chr8 | 92000000 | 145138636 | 53138636 | translocation | Clonal |
| AML01_PE20356 | chr21 | 34900000 | 50818468 | 15918468 | translocation | Clonal |
| AML01_PE20356 | chr8 | 92000000 | 145138636 | 53138636 | translocation | Clonal |
| AML01_PE20357 | chr21 | 34900000 | 50818468 | 15918468 | translocation | Clonal |
| AML01_PE20357 | chr8 | 92000000 | 145138636 | 53138636 | translocation | Clonal |
| AML01_PE20359 | chr21 | 34900000 | 50818468 | 15918468 | translocation | Clonal |
| AML01_PE20359 | chr8 | 92000000 | 145138636 | 53138636 | translocation | Clonal |
| AML01_PE20361 | chr21 | 34900000 | 50818468 | 15918468 | translocation | Clonal |
| AML01_PE20361 | chr8 | 92000000 | 145138636 | 53138636 | translocation | Clonal |
| AML01_PE20371 | chr21 | 34900000 | 50818468 | 15918468 | translocation | Clonal |
| AML01_PE20371 | chr8 | 92000000 | 145138636 | 53138636 | translocation | Clonal |
| AML01_PE20372 | chr21 | 34900000 | 50818468 | 15918468 | translocation | Clonal |

| | | | | | | |
|---|---|---|---|---|---|---|
| AML01_PE20372 | chr8 | 92000000 | 145138636 | 53138636 | translocation | Clonal |
| AML01_PE20374 | chr21 | 34900000 | 50818468 | 15918468 | translocation | Clonal |
| AML01_PE20374 | chr8 | 92000000 | 145138636 | 53138636 | translocation | Clonal |
| AML01_PE20375 | chr21 | 34900000 | 50818468 | 15918468 | translocation | Clonal |
| AML01_PE20375 | chr8 | 92000000 | 145138636 | 53138636 | translocation | Clonal |
| AML01_PE20376 | chr21 | 34900000 | 50818468 | 15918468 | translocation | Clonal |
| AML01_PE20376 | chr8 | 92000000 | 145138636 | 53138636 | translocation | Clonal |
| AML01_PE20377 | chr21 | 34900000 | 50818468 | 15918468 | translocation | Clonal |
| AML01_PE20377 | chr8 | 92000000 | 145138636 | 53138636 | translocation | Clonal |
| AML01_PE20382 | chr21 | 34900000 | 50818468 | 15918468 | translocation | Clonal |
| AML01_PE20382 | chr8 | 92000000 | 145138636 | 53138636 | translocation | Clonal |
| AML01_PE20383 | chr21 | 34900000 | 50818468 | 15918468 | translocation | Clonal |
| AML01_PE20383 | chr8 | 92000000 | 145138636 | 53138636 | translocation | Clonal |
| AML01_PE20384 | chr21 | 34900000 | 50818468 | 15918468 | translocation | Clonal |
| AML01_PE20384 | chr8 | 92000000 | 145138636 | 53138636 | translocation | Clonal |
| AML01_PE20385 | chr21 | 34900000 | 50818468 | 15918468 | translocation | Clonal |
| AML01_PE20385 | chr8 | 92000000 | 145138636 | 53138636 | translocation | Clonal |
| AML01_PE20387 | chr21 | 34900000 | 50818468 | 15918468 | translocation | Clonal |
| AML01_PE20387 | chr8 | 92000000 | 145138636 | 53138636 | translocation | Clonal |
| AML01_PE20388 | chr21 | 34900000 | 50818468 | 15918468 | translocation | Clonal |
| AML01_PE20388 | chr8 | 92000000 | 145138636 | 53138636 | translocation | Clonal |
| AML01_PE20395 | chr21 | 34900000 | 50818468 | 15918468 | translocation | Clonal |
| AML01_PE20395 | chr8 | 92000000 | 145138636 | 53138636 | translocation | Clonal |
| AML01_PE20396 | chr21 | 34900000 | 50818468 | 15918468 | translocation | Clonal |
| AML01_PE20396 | chr8 | 92000000 | 145138636 | 53138636 | translocation | Clonal |

**Table S3F. List of SVs found in the TALL_P1 sample**

| Cell | Chromosome | Start | End | Size (bp) | SV Class | Singleton/Subclonal |
|---|---|---|---|---|---|---|
| TALL3x1_DEA5_PE20406 | chr14 | 95800000 | 98400000 | 2600000 | inv_h2 | Clonal |
| TALL3x1_DEA5_PE20406 | chr6 | 83200000 | 170600000 | 87400000 | complex_h2 | Subclonal |
| TALL3x1_DEA5_PE20414 | chr14 | 95800000 | 98400000 | 2600000 | inv_h2 | Clonal |
| TALL3x1_DEA5_PE20414 | chr6 | 83200000 | 170600000 | 87400000 | complex_h2 | Subclonal |
| TALL3x1_DEA5_PE20415 | chr14 | 95800000 | 98400000 | 2600000 | inv_h2 | Clonal |
| TALL3x1_DEA5_PE20416 | chr14 | 95800000 | 98400000 | 2600000 | inv_h2 | Clonal |
| TALL3x1_DEA5_PE20417 | chr14 | 95800000 | 98400000 | 2600000 | inv_h2 | Clonal |
| TALL3x1_DEA5_PE20418 | chr14 | 95800000 | 98400000 | 2600000 | inv_h2 | Clonal |
| TALL3x1_DEA5_PE20419 | chr14 | 95800000 | 98400000 | 2600000 | inv_h2 | Clonal |
| TALL3x1_DEA5_PE20421 | chr14 | 95800000 | 98400000 | 2600000 | inv_h2 | Clonal |
| TALL3x1_DEA5_PE20422 | chr14 | 95800000 | 98400000 | 2600000 | inv_h2 | Clonal |
| TALL3x1_DEA5_PE20424 | chr14 | 95800000 | 98400000 | 2600000 | inv_h2 | Clonal |
| TALL3x1_DEA5_PE20424 | chr6 | 83200000 | 170600000 | 87400000 | complex_h2 | Subclonal |
| TALL3x1_DEA5_PE20427 | chr14 | 95800000 | 98400000 | 2600000 | inv_h2 | Clonal |
| TALL3x1_DEA5_PE20430 | chr14 | 95800000 | 98400000 | 2600000 | inv_h2 | Clonal |
| TALL3x1_DEA5_PE20433 | chr14 | 95800000 | 98400000 | 2600000 | inv_h2 | Clonal |
| TALL3x1_DEA5_PE20435 | chr14 | 95800000 | 98400000 | 2600000 | inv_h2 | Clonal |
| TALL3x1_DEA5_PE20435 | chr6 | 83200000 | 170600000 | 87400000 | complex_h2 | Subclonal |
| TALL3x1_DEA5_PE20438 | chr14 | 95800000 | 98400000 | 2600000 | inv_h2 | Clonal |
| TALL3x1_DEA5_PE20439 | chr14 | 95800000 | 98400000 | 2600000 | inv_h2 | Clonal |
| TALL3x1_DEA5_PE20442 | chr14 | 95800000 | 98400000 | 2600000 | inv_h2 | Clonal |
| TALL3x1_DEA5_PE20442 | chr6 | 83200000 | 170600000 | 87400000 | complex_h2 | Subclonal |
| TALL3x1_DEA5_PE20444 | chr14 | 95800000 | 98400000 | 2600000 | inv_h2 | Clonal |
| TALL3x1_DEA5_PE20447 | chr14 | 95800000 | 98400000 | 2600000 | inv_h2 | Clonal |
| TALL3x1_DEA5_PE20447 | chr6 | 83200000 | 170600000 | 87400000 | complex_h2 | Subclonal |
| TALL3x1_DEA5_PE20448 | chr14 | 95800000 | 98400000 | 2600000 | inv_h2 | Clonal |
| TALL3x1_DEA5_PE20448 | chr6 | 83200000 | 170600000 | 87400000 | complex_h2 | Subclonal |
| TALL3x1_DEA5_PE20449 | chr14 | 95800000 | 98400000 | 2600000 | inv_h2 | Clonal |

| | | | | | | |
|---|---|---|---|---|---|---|
| TALL3x1_DEA5_PE20451 | chr14 | 95800000 | 98400000 | 2600000 | inv_h2 | Clonal |
| TALL3x1_DEA5_PE20452 | chr14 | 95800000 | 98400000 | 2600000 | inv_h2 | Clonal |
| TALL3x1_DEA5_PE20453 | chr14 | 95800000 | 98400000 | 2600000 | inv_h2 | Clonal |
| TALL3x1_DEA5_PE20453 | chr6 | 83200000 | 170600000 | 87400000 | complex_h2 | Subclonal |
| TALL3x1_DEA5_PE20455 | chr14 | 95800000 | 98400000 | 2600000 | inv_h2 | Clonal |
| TALL3x1_DEA5_PE20457 | chr14 | 95800000 | 98400000 | 2600000 | inv_h2 | Clonal |
| TALL3x1_DEA5_PE20458 | chr14 | 95800000 | 98400000 | 2600000 | inv_h2 | Clonal |
| TALL3x1_DEA5_PE20458 | chr6 | 83200000 | 170600000 | 87400000 | complex_h2 | Subclonal |
| TALL3x1_DEA5_PE20461 | chr14 | 95800000 | 98400000 | 2600000 | inv_h2 | Clonal |
| TALL3x1_DEA5_PE20463 | chr14 | 95800000 | 98400000 | 2600000 | inv_h2 | Clonal |
| TALL3x1_DEA5_PE20466 | chr14 | 95800000 | 98400000 | 2600000 | inv_h2 | Clonal |
| TALL3x1_DEA5_PE20469 | chr14 | 95800000 | 98400000 | 2600000 | inv_h2 | Clonal |
| TALL3x1_DEA5_PE20471 | chr14 | 95800000 | 98400000 | 2600000 | inv_h2 | Clonal |
| TALL3x1_DEA5_PE20471 | chr6 | 83200000 | 170600000 | 87400000 | complex_h2 | Subclonal |
| TALL3x1_DEA5_PE20474 | chr14 | 95800000 | 98400000 | 2600000 | inv_h2 | Clonal |
| TALL3x1_DEA5_PE20474 | chr6 | 83200000 | 170600000 | 87400000 | complex_h2 | Subclonal |
| TALL3x1_DEA5_PE20475 | chr14 | 95800000 | 98400000 | 2600000 | inv_h2 | Clonal |
| TALL3x1_DEA5_PE20481 | chr14 | 95800000 | 98400000 | 2600000 | inv_h2 | Clonal |
| TALL3x1_DEA5_PE20485 | chr14 | 95800000 | 98400000 | 2600000 | inv_h2 | Clonal |
| TALL3x1_DEA5_PE20487 | chr14 | 95800000 | 98400000 | 2600000 | inv_h2 | Clonal |
| TALL3x1_DEA5_PE20488 | chr14 | 95800000 | 98400000 | 2600000 | inv_h2 | Clonal |
| TALL3x1_DEA5_PE20494 | chr14 | 95800000 | 98400000 | 2600000 | inv_h2 | Clonal |
| TALL3x1_DEA5_PE20494 | chr6 | 83200000 | 170600000 | 87400000 | complex_h2 | Subclonal |
| TALL3x1_DEA5_PE20495 | chr14 | 95800000 | 98400000 | 2600000 | inv_h2 | Clonal |
| TALL3x1_DEA5_PE20495 | chr6 | 83200000 | 170600000 | 87400000 | complex_h2 | Subclonal |
| TALL3x2_DEA5_PE20402 | chr14 | 95800000 | 98400000 | 2600000 | inv_h2 | Clonal |
| TALL3x2_DEA5_PE20408 | chr14 | 95800000 | 98400000 | 2600000 | inv_h2 | Clonal |
| TALL3x2_DEA5_PE20410 | chr14 | 95800000 | 98400000 | 2600000 | inv_h2 | Clonal |
| TALL3x2_DEA5_PE20410 | chr6 | 83200000 | 170600000 | 87400000 | complex_h2 | Subclonal |
| TALL3x2_DEA5_PE20411 | chr14 | 95800000 | 98400000 | 2600000 | inv_h2 | Clonal |
| TALL3x2_DEA5_PE20412 | chr14 | 95800000 | 98400000 | 2600000 | inv_h2 | Clonal |

| | | | | | | |
|---|---|---|---|---|---|---|
| TALL3x2_DEA5_PE20414 | chr14 | 95800000 | 98400000 | 2600000 | inv_h2 | Clonal |
| TALL3x2_DEA5_PE20416 | chr14 | 95800000 | 98400000 | 2600000 | inv_h2 | Clonal |
| TALL3x2_DEA5_PE20418 | chr14 | 95800000 | 98400000 | 2600000 | inv_h2 | Clonal |
| TALL3x2_DEA5_PE20420 | chr14 | 95800000 | 98400000 | 2600000 | inv_h2 | Clonal |
| TALL3x2_DEA5_PE20420 | chr6 | 83200000 | 170600000 | 87400000 | complex_h2 | Subclonal |
| TALL3x2_DEA5_PE20423 | chr14 | 95800000 | 98400000 | 2600000 | inv_h2 | Clonal |
| TALL3x2_DEA5_PE20428 | chr14 | 95800000 | 98400000 | 2600000 | inv_h2 | Clonal |
| TALL3x2_DEA5_PE20431 | chr14 | 95800000 | 98400000 | 2600000 | inv_h2 | Clonal |
| TALL3x2_DEA5_PE20435 | chr14 | 95800000 | 98400000 | 2600000 | inv_h2 | Clonal |
| TALL3x2_DEA5_PE20435 | chr6 | 83200000 | 170600000 | 87400000 | complex_h2 | Subclonal |
| TALL3x2_DEA5_PE20436 | chr14 | 95800000 | 98400000 | 2600000 | inv_h2 | Clonal |
| TALL3x2_DEA5_PE20436 | chr6 | 83200000 | 170600000 | 87400000 | complex_h2 | Subclonal |
| TALL3x2_DEA5_PE20440 | chr14 | 95800000 | 98400000 | 2600000 | inv_h2 | Clonal |
| TALL3x2_DEA5_PE20441 | chr14 | 95800000 | 98400000 | 2600000 | inv_h2 | Clonal |
| TALL3x2_DEA5_PE20442 | chr14 | 95800000 | 98400000 | 2600000 | inv_h2 | Clonal |
| TALL3x2_DEA5_PE20444 | chr14 | 95800000 | 98400000 | 2600000 | inv_h2 | Clonal |
| TALL3x2_DEA5_PE20445 | chr14 | 95800000 | 98400000 | 2600000 | inv_h2 | Clonal |
| TALL3x2_DEA5_PE20446 | chr14 | 95800000 | 98400000 | 2600000 | inv_h2 | Clonal |
| TALL3x2_DEA5_PE20446 | chr6 | 83200000 | 170600000 | 87400000 | complex_h2 | Subclonal |
| TALL3x2_DEA5_PE20448 | chr14 | 95800000 | 98400000 | 2600000 | inv_h2 | Clonal |
| TALL3x2_DEA5_PE20448 | chr6 | 83200000 | 170600000 | 87400000 | complex_h2 | Subclonal |
| TALL3x2_DEA5_PE20450 | chr14 | 95800000 | 98400000 | 2600000 | inv_h2 | Clonal |
| TALL3x2_DEA5_PE20453 | chr14 | 95800000 | 98400000 | 2600000 | inv_h2 | Clonal |
| TALL3x2_DEA5_PE20454 | chr14 | 95800000 | 98400000 | 2600000 | inv_h2 | Clonal |
| TALL3x2_DEA5_PE20454 | chr6 | 83200000 | 170600000 | 87400000 | complex_h2 | Subclonal |
| TALL3x2_DEA5_PE20456 | chr14 | 95800000 | 98400000 | 2600000 | inv_h2 | Clonal |
| TALL3x2_DEA5_PE20456 | chr6 | 83200000 | 170600000 | 87400000 | complex_h2 | Subclonal |
| TALL3x2_DEA5_PE20457 | chr14 | 95800000 | 98400000 | 2600000 | inv_h2 | Clonal |
| TALL3x2_DEA5_PE20461 | chr14 | 95800000 | 98400000 | 2600000 | inv_h2 | Clonal |
| TALL3x2_DEA5_PE20462 | chr14 | 95800000 | 98400000 | 2600000 | inv_h2 | Clonal |
| TALL3x2_DEA5_PE20463 | chr14 | 95800000 | 98400000 | 2600000 | inv_h2 | Clonal |

| | | | | | | |
|---|---|---|---|---|---|---|
| TALL3x2_DEA5_PE20466 | chr14 | 95800000 | 98400000 | 2600000 | inv_h2 | Clonal |
| TALL3x2_DEA5_PE20471 | chr14 | 95800000 | 98400000 | 2600000 | inv_h2 | Clonal |
| TALL3x2_DEA5_PE20474 | chr14 | 95800000 | 98400000 | 2600000 | inv_h2 | Clonal |
| TALL3x2_DEA5_PE20474 | chr6 | 83200000 | 170600000 | 87400000 | complex_h2 | Subclonal |
| TALL3x2_DEA5_PE20477 | chr14 | 95800000 | 98400000 | 2600000 | inv_h2 | Clonal |
| TALL3x2_DEA5_PE20479 | chr14 | 95800000 | 98400000 | 2600000 | inv_h2 | Clonal |
| TALL3x2_DEA5_PE20482 | chr14 | 95800000 | 98400000 | 2600000 | inv_h2 | Clonal |
| TALL3x2_DEA5_PE20482 | chr6 | 83200000 | 170600000 | 87400000 | complex_h2 | Subclonal |
| TALL3x2_DEA5_PE20485 | chr14 | 95800000 | 98400000 | 2600000 | inv_h2 | Clonal |
| TALL3x2_DEA5_PE20486 | chr14 | 95800000 | 98400000 | 2600000 | inv_h2 | Clonal |
| TALL3x2_DEA5_PE20495 | chr14 | 95800000 | 98400000 | 2600000 | inv_h2 | Clonal |
