## Supplementary Table S4 for "Haplotype-aware single-cell multiomics uncovers functional effects of somatic structural variation"

**Table S4. Summary of inferring SCNAs from the scRNA-seq data sets.**

| Sample | SV | CF (%) | Size (Mb) | Discovery mode<br>InferCNV | <br>HoneyBADGER | <br>CONICSmat | Genotyping mode<br>CONICSmat |
|---|---|---|---|---|---|---|---|
| NA12878 | chr19 small deletion | 21.3 | 0.5 | N | N | N | Y |
| NA12878 | chr22 small deletion | 78.6 | 0.7 | N | N | N | N |
| Sample to test | - | - | - | NA12878 from SMART-seq and Fluidigm data sets (GSE44618, GSE81861) | | - | |
| Normal control | - | - | - | *SMART-seq of LCL (GSE123028) | | - | |
| Predicted SCNAs | - | - | - | 137 | 0 | 0 | - |
| NA12878 | chr19 small deletion | 21.3 | 0.5 | N | N | N | Y |
| NA12878 | chr22 small deletion | 78.6 | 0.7 | N | N | N | N |
| Tumor | - | - | - | NA12878 10X transcriptome (GSM3596321) | | - | |
| Sample to test | - | - | - | *NA18502 10X transcriptome (GSM3596320) | | - | |
| Predicted SCNAs | - | - | - | 8 | 1 | 13 | - |
| TALL_P1 | chr7p loss | 100 | 58.2 | Y | N | Not available (clonal) | Not available (clonal) |
| TALL_P1 | chr7q gain | 100 | 97.8 | Y | N | Not available (clonal) | Not available (clonal) |
| TALL_P1 | chr6 Del1 | 29.9 | 2.4 | N | N | N | Y (Del group) |
| TALL_P1 | chr6 Dup | 29.9 | 1 | N | N | N | Y (Dup group) |
| TALL_P1 | chr6 Del2 | 29.9 | 7 | N | N | N | Y (Del group) |
| TALL_P1 | chr6 Inv1 | 29.9 | 1.6 | N | N | N | Not available (copy neutral) |
| TALL_P1 | chr6 InvDup | 29.9 | 39.9 | N | N | N | Y (Dup group) |
| TALL_P1 | chr6 Del3 | 29.9 | 1.2 | N | N | N | Y (Del group) |
| TALL_P1 | chr6 Inv2 | 29.9 | 0.6 | N | N | N | Not available (copy neutral) |
| Tumor | - | - | - | T-ALL P1 10X transcriptome (This study) | | - | |
| Sample to test | - | - | - | *CD4 T cells, and CD8 T cells from PBMC data sets (10X genomics), https://www.10xgenomics.com/reso | | - | |
| Predicted SCNAs | - | - | - | 194 | 22 | 2 | - |

Discovery mode

| | |
|---|---|
| Y | two SV breakpoints (start and end) found in +-5Mb of ground truth |
| N | two SV breakpoints (start and end) was not found in +-5Mb of ground truth |

Genotyping mode

| | |
|---|---|
| Y | BIC for 2-component model is lower than 1-component model and LRT of difference between two models show P.adj<0.01 |
| N | BIC for 1-component model is lower than 2-component model |

* Two packages (InferCNV, HoneyBADGER) require karyotypically normal cell control population for the discovery of SCNAs
