## Supplementary Table S5 for "Haplotype-aware single-cell multiomics uncovers functional effects of somatic structural variation"

**Table S5. Reference list of ATAC-seq peaks from previous publications collected to define putative CREs in the AML system.**

| AccessionID | Sample | Available peak information | Reference |
|---|---|---|---|
| GSE135025 | ATAC-seq in THP1 cells | GSM3983900_THP1_ATAC_peaks.txt.gz | PMID: 32242051 |
| GSE124967 | Comparison of chromatin accessibility (ATAC-seq) in genetically clonal iLSCs vs iBlasts | GSE124967_ATAC_atlas.bed.gz | - |
| GSE122577 | Single-cell ATAC-seq of pre-leukemia hematopoietic stem cells from AML patient | GSE122577_SU353_pHSC_CountByCell.bed.gz | PMID: 30958261 |
| GSE122576 | Single-cell ATAC-seq of leukemia stem cells and leukemic blasts from AML patients | GSE122576_AMLSU070_blast_LSC_CountByCell.bed.gz | PMID: 30958261 |
| GSE122576 | Single-cell ATAC-seq of leukemia stem cells and leukemic blasts from AML patients | GSE122576_AMLSU353_blast_LSC_CountByCell.bed.gz | PMID: 30958261 |
| GSM3099673 | WT_1_ATAC-seq (MOLM13, Human-derived acute myeloid leukemia cells) | GSM3099673_WT_1_ATAC-seq_peaks.bed.gz | PMID: 29760161 |
| GSM3099674 | WT_2_ATAC-seq (MOLM13, Human-derived acute myeloid leukemia cells) | GSM3099674_WT_2_ATAC-seq_peaks.bed.gz | PMID: 29760161 |
| GSE74912 | ATAC-seq profiles of hematopoietic and leukemic cell types, across 13 normal hematopoietic cell types and 3 acute myeloid leukemia cell types. | GSE74912_ATACseq_All_Counts.txt.gz | PMID: 27526324 |
| GSE137667 | Expression of RUNX1-ETO Rapidly Alters the Chromatin Landscape and Growth of Early Human Myeloid Precursor Cells (ATAC-Seq) | GSM4083935_Runx1cNeg_0Dox_treat_pileup.bedgraph.gz, GSM4083936_R | PMID: 32460028 |
