## Supplementary Table S6 for "Haplotype-aware single-cell multiomics uncovers functional effects of somatic structural variation"

**Table S6. Reference list showing literature sources for sets of TF target genes.** Target genes were defined by TF binding (ChIP-seq) and RNA perturbation (RNA-seq or microarray) upon silencing of TFs. TFs located in the chromothripsis affe

| Index | chr | str | end | gname | EntrezID | TF family | CNA (copy num | Reference | Method | Targets | Overlap | Pvalue | P.adjust |
|---|---|---|---|---|---|---|---|---|---|---|---|---|---|
| 1 | chr6 | 135181315 | 135219173 | MYB | 4602 | MYB | inv_dup | PMID:21317192 | Expression MYB KD | 915 | 6 | 7.03E-06 | 0.00015466 |
| 2 | chr6 | 98834592 | 98839490 | POU3F2 | 5454 | Pou | del | PMID:25991548 | RNA-seq following | 205 | 2 | 0.0066 | 0.1452 |
| 3 | chr6 | 106086320 | 106109939 | PRDM1 | 639 | zf-C2H2 | none | PMID:26779602 | RNA-seq upon Prd | 267 | 1 | 0.1506 | 1 |
| 4 | chr6 | 108166058 | 108188809 | NR2E1 | 7101 | RXR-like | none | PMID:26204903 | LongSAGE libraries | 754 | 2 | 0.0744 | 1 |
| 5 | chr6 | 87152833 | 87264196 | ZNF292 | 23036 | zf-C2H2 | none | PMID:22955616 | ChIP-seq of ZNF29 | 1049 | 1 | 0.4803 | 1 |
| 6 | chr6 | 99606730 | 99615578 | PRDM13 | 59336 | zf-C2H2 | del | PMID:28850031 | RNA-seq Prdm13 v | 52 | 0 | 1 | 1 |
| 7 | chr6 | 116877212 | 116932163 | RFX6 | 222546 | RFX | inv_dup | PMID:30509498 | Microarray of siRfx | 31 | 0 | 1 | 1 |
| 8 | chr6 | 122399546 | 122433119 | HSF2 | 3298 | HSF | inv_dup | PMID:23959860 (C | RNA-seq HSF2 KO i | 40 | 0 | 1 | 1 |
| 9 | chr6 | 125747664 | 125761269 | HEY2 | 23493 | bHLH | inv_dup | PMID:22615585 | Microarray express | 1225 | 0 | 1 | 1 |
| 10 | chr6 | 130013699 | 130141451 | L3MBTL3 | 84456 | Others | inv_dup | PMID:29030483 | Expression using q | 4 | 0 | 1 | 1 |
| 11 | chr6 | 133889138 | 133895553 | TCF21 | 6943 | bHLH | inv_dup | PMID:28481916 fo | RNA-seq, ChIP-seq | 358 | 0 | 1 | 1 |
| 12 | chr6 | 136256863 | 136289851 | BCLAF1 | 9774 | Others | inv_dup | PMID:31644907 | RNA-seq shBclaf1 i | 169 | 0 | 1 | 1 |
| 13 | chr6 | 142751467 | 142945201 | HIVEP2 | 3097 | zf-C2H2 | inv_dup | PMID:1409593 | EMSA assay | 6 | 0 | 1 | 1 |
| 14 | chr6 | 143940300 | 144064599 | PLAGL1 | 5325 | zf-C2H2 | inv_dup | PMID:28985358 | Digital Gene Expre | 313 | 0 | 1 | 1 |
| 15 | chr6 | 151364117 | 151391548 | ZBTB2 | 57621 | ZBTB | inv_dup | PMID:29437775 | RNA-seq upon Zbtl | 271 | 0 | 1 | 1 |
| 16 | chr6 | 151656691 | 152129619 | ESR1 | 2099 | ESR-like | inv_dup | PMID:31408468 | RNA-seq DEG regu | 157 | 0 | 1 | 1 |
| 17 | chr6 | 156777374 | 157210779 | ARID1B | 57492 | ARID | del | PMID:26716708 | RNA-seq RNAi agai | 349 | 0 | 1 | 1 |
| 18 | chr6 | 89926529 | 90296908 | BACH2 | 60468 | TF_bZIP | none | PMID:27158840 | RNA-seq upon KO | 276 | 0 | 1 | 1 |
| 19 | chr6 | 104957048 | 105083332 | LIN28B | 389421 | CSD | none | PMID:32601179 | RNA-seq upon LIN | 488 | 0 | 1 | 1 |
| 20 | chr6 | 108559835 | 108684774 | FOXO3 | 2309 | Fork_head | none | PMID:23340844 fo | RNAPII ChIP-seq u | 805 | 0 | 1 | 1 |
| 21 | chr6 | 109462594 | 109483237 | ZBTB24 | 9841 | ZBTB | none | PMID:30085123 | RNA-seq of HCT11 | 106 | 0 | 1 | 1 |
| 22 | chr6 | 166157656 | 166168700 | T | 6862 | T-box | none | PMID:24616493 | RNA-seq in the E | 165 | 0 | 1 | 1 |
