## Supplementary Table S7 for "Haplotype-aware single-cell multiomics uncovers functional effects of somatic structural variation"

**Table S7. Gene sets used in expression analyses to verify TF target gene and pathway activities**

| MYB targets | WNT canonical pathway targets | WNT_SIGNALING | MET pathway ta | MYC/MAX targets (1) | MYC/MAX targets (2) | MAPK signaling pathway |
|---|---|---|---|---|---|---|
| Zhao et al. 2011 | Trrust (CTNNB1, LEF1, TCF7, TCF7L2) | Msigdb | Kaposi-Novak et | Msigdb (MYCMAX_B) | Msigdb (MYCMAX_01) | KEGG pathway |
| C4orf19 | AR | DVL1 | CAP1 | MMP23B | ATAD3B | CACNA1A |
| ACAD9 | BIRC5 | CTNNBIP1 | CYP4A11 | DNAJC11 | ATAD3A | CACNA1B |
| ACBD6 | CCND1 | WNT4 | SCP2 | SLC25A33 | UBIAD1 | CACNA1C |
| ACOT7 | CD44 | JUN | ANGPTL3 | EPHA2 | UBR4 | CACNA1D |
| ACSL1 | CHGA | WNT2B | VCAM1 | SDHB | UBXN10 | CACNA1E |
| ADORA2B | CLDN2 | BCL9 | NGF | UBXN10 | STMN1 | CACNA1F |
| ALOX5AP | FST | WNT9A | PKLR | HNRNPR | LIN28A | CACNA1G |
| ANKRD28 | GLCE | WNT3A | PEA15 | STMN1 | NUDC | CACNA1H |
| ANKS1A | GLI2 | RHOU | DNM3 | YTHDF2 | OPRD1 | CACNA1I |
| ARHGAP24 | JUN | FZD8 | G0S2 | HPCA | EPB41 | CACNA1S |
| ATP6V1C1 | MMP14 | DKK1 | CAPN2 | ZNF362 | MYCL | CACNA2D1 |
| B4GALT1 | MMP7 | FRAT1 | VIM | HPCAL4 | FOXJ3 | CACNA2D2 |
| BCL2 | MYC | BTRC | ITGB1 | RIMKLA | IPO13 | CACNA2D3 |
| BCL6 | NCOA2 | FBXW4 | FAS | LRP8 | TESK2 | CACNA2D4 |
| BLM | NRCAM | CTBP2 | LIPA | SERBP1 | SLC1A7 | CACNB1 |
| BLVRB | PLD1 | FSHB | ANKRD1 | GFI1 | LRP8 | CACNB2 |
| CDCA4 | PPARD | FOSL1 | GPAM | HENMT1 | FOXD3 | CACNB3 |
| CDCA7L | PTGS2 | LRP5 | BNIP3 | RORC | SERBP1 | CACNB4 |
| CITED2 | RB1 | CCND1 | GLYAT | CRABP2 | CDC14A | CACNG1 |
| COQ9 | STARD7 | FGF4 | DGAT2 | C1orf21 | AMPD2 | CACNG2 |
| CR1L | TCF7 | WNT11 | MMP7 | IVNS1ABP | NRAS | CACNG3 |
| CTSG | VIM | FZD4 | MGST1 | AHCTF1 | CSDE1 | CACNG4 |
| CTSK | CCL7 | DIXDC1 | TUBA1A | OR2L13 | CA14 | CACNG5 |
| DAAM1 | CD1D | WNT5B | KRT8 | PTF1A | RFX5 | CACNG6 |
| DGKG | CEBPA | CCND2 | NFYB | NEUROG3 | RORC | CACNG7 |
| DNAJC21 | CTLA4 | GAPDH | GPC6 | PAX2 | KRTCAP2 | CACNG8 |
| DTYMK | CTNNB1 | LRP6 | MOAP1 | PITX3 | TRIM46 | PRKACA |
| DUSP1 | CYLD | WNT1 | LIPC | TRIM8 | POGK | PRKACB |
| DUSP6 | DSG4 | WIF1 | ANXA2 | ARL3 | IVNS1ABP | PRKACG |
| DUT | EDA | DAAM1 | NEO1 | VAX1 | TGFB2 | PRKCA |
| EIF2S2 | ELANE | B2M | ACSM1 | ASCL2 | H3F3A | PRKCB |
| EMILIN2 | ESR1 | PYGO1 | RRAD | IPO7 | HNRNPH3 | PRKCG |
| EMP3 | HERC2 | CSNK1G1 | CTCF | WEE1 | CNNM1 | GNA12 |
| EXTL2 | MITF | AXIN1 | INPP5K | MICAL2 | LZTS2 | GNG12 |
| FAM20C | MYCBP | NKD1 | KPNB1 | CSRP3 | PPRC1 | PPP3CA |
| FGD4 | NT5E | DVL2 | ITGA3 | ABTB2 | PITX3 | PPP3CB |
| FGR | OCA2 | NLK | SMURF2 | ZDHHC5 | ARL3 | PPP3CC |
| FKBP5 | PRL | FOXN1 | MRPL38 | SYT7 | SFXN2 | PPP3R1 |
| FLOT1 | RAG2 | FZD2 | ACOX1 | SYT12 | SMC3 | PPP3R2 |
| FOXO3 | SKP2 | WNT3 | FASN | CCND1 | NRIP3 | RASGRF1 |
| FXYD5 | SNAI2 | SLC9A3R1 | RAB12 | FCHSD2 | IPO7 | RASGRF2 |
| GBE1 | TCF3 | CSNK1D | TUBB6 | KLHL35 | IGSF22 | RASGRP1 |
| GCLM | TCF4 | TLE2 | TJP3 | CREBZF | RTN4RL2 | RASGRP2 |
| GCNT2 | ABCB1 | AES | ANGPTL4 | PICALM | SLC43A1 | RASGRP3 |
| GFI1 | ACVRL1 | GSK3A | GCDH | C11orf57 | TIMM10 | RASGRP4 |
| GLIPR2 | GATA3 | RPL13A | CEBPA | PIH1D2 | RAB3IL1 | RAPGEF2 |
| GLRX | HECA | PPP2R1A | ECH1 | GRIN2B | CHRM1 | NF1 |
| GNAI2 | WWC1 | TCF7L1 | GCKR | EPS8 | RCOR2 | RASA1 |
| GPR160 | STAT3 | FRZB | RTN4 | BHLHE41 | ESRRA | RASA2 |
| GSTM5 | | FZD7 | FABP1 | WNT1 | REXO2 | RAP1A |
| HEBP1 | | FZD5 | ACVR1 | HOXC4 | APOA5 | RAP1B |
| HK1 | | WNT6 | NFE2L2 | C12orf66 | KMT2A | EGF |
| HSPA2 | | WNT10A | ITGAV | CCER1 | FGF6 | TGFA |
| IFI30 | | CSNK2A1 | FN1 | PLXNC1 | CHD4 | EREG |
| IFNGR2 | | KREMEN1 | NCL | ASCL1 | ATF7IP | AREG |
| IL13RA1 | | EP300 | MAFF | STARD13 | COL2A1 | FGF1 |
| IL17RA | | WNT7B | FOXP1 | FGF14 | FKBP11 | FGF2 |
| IQSEC1 | | WNT7A | ROBO1 | AJUBA | RHEBL1 | FGF3 |
| KIF17 | | CTNNB1 | EIF4A2 | NFATC4 | HOXC11 | FGF4 |
| KLF13 | | WNT5A | GSX2 | NKX2-8 | CBX5 | FGF17 |
| KLF6 | | GSK3B | CXCL3 | LRFN5 | HNRNPA1 | FGF6 |
| LBR | | SENP2 | CXCL10 | ARF6 | PA2G4 | FGF7 |
| LCP1 | | CTBP1 | SPP1 | SIX1 | ZC3H10 | FGF8 |
| LMNA | | CXXC4 | ADH1C | ZFP36L1 | USP15 | FGF9 |
| LYPD6B | | LEF1 | PIK3R1 | IRF2BPL | C12orf66 | FGF10 |
| MAP1LC3B | | PITX2 | CHD1 | GTF2A1 | SLC6A15 | FGF16 |

| | | | | | |
|---|---|---|---|---|---|
| MAP2K3 | APC | EGR1 | BCL11B | RFX4 | FGF5 |
| MAPK3 | TCF7 | HBEGF | KLF13 | DIABLO | FGF18 |
| MBP | PPP2CA | PPP1R10 | DLL4 | PUS1 | FGF20 |
| MCM6 | WNT8A | CLIC1 | CATSPER2 | FLT3 | FGF22 |
| MFSD6 | CSNK1A1 | BAK1 | ALDH1A2 | FGF14 | FGF19 |
| MGAT4A | FBXW11 | HMGA1 | SPG21 | DCAF11 | FGF21 |
| MIDN | CCND3 | EPHA7 | PARP6 | CTAGE5 | FGF23 |
| MITF | T | CTGF | SEMA7A | FAM179B | NGF |
| MPO | ACTB | CROT | MEX3B | KLHL28 | BDNF |
| MS4A3 | SFRP4 | ARPC1B | PDIA2 | BMP4 | NTF3 |
| MSH6 | FZD1 | RAB19 | NTN3 | KIAA0586 | NTF4 |
| MXI1 | WNT2 | PDIA4 | TFAP4 | TIMM9 | INS |
| MYC | WNT16 | SLC25A37 | SEZ6L2 | MTHFD1 | IGF1 |
| MYH9 | FZD3 | CDH17 | ZNF771 | RPS6KA5 | IGF2 |
| MYO9B | SFRP1 | GLDC | SALL1 | C2CD4A | PDGFA |
| NAP1L4 | SOX17 | TPM2 | PDP2 | CSK | PDGFB |
| NASP | FZD6 | HSPA5 | HSF4 | ADAMTS17 | PDGFC |
| NAV1 | MYC | ASS1 | NUTF2 | PDIA2 | PDGFD |
| NCF2 | WISP1 | MID1 | ST3GAL2 | NTN3 | CSF1 |
| NDEL1 | TLE1 | OTC | ZFP1 | TFAP4 | KITLG |
| NDUFAF3 | FBXW2 | MSN | CRK | CLN3 | FLT3LG |
| NFE2 | PORCN | CLDN2 | PITPNA | ZNF771 | VEGFA |
| NFIL3 | HPRT1 | | SERPINF1 | DCTPP1 | VEGFB |
| NFKBIZ | | | PFN1 | BCL7C | PGF |
| NOTCH1 | | | EIF5A | ESRP2 | VEGFC |
| NUP205 | | | FXR2 | GCSH | VEGFD |
| OLA1 | | | ALOXE3 | MNT | HGF |
| OSTF1 | | | PER1 | CLUH | ANGPT1 |
| PBX4 | | | HS3ST3B1 | ENO3 | ANGPT2 |
| PDCD4 | | | GIT1 | PFN1 | ANGPT4 |
| PEX10 | | | TMEM132E | PER1 | EFNA1 |
| PFKL | | | PIGW | GIT1 | EFNA2 |
| PHLDA1 | | | MYO19 | LEPREL4 | EFNA3 |
| PLEKHA2 | | | PIP4K2B | FKBP10 | EFNA4 |
| PLEKHO2 | | | NR1D1 | KCNH4 | EFNA5 |
| PLP2 | | | KCNH4 | RAMP2 | EGFR |
| POLR1E | | | G6PC3 | PSME3 | ERBB2 |
| POLR2I | | | NSF | ETV4 | ERBB3 |
| PPP1R12A | | | KPNB1 | MPP3 | ERBB4 |
| PRAM1 | | | HOXB9 | G6PC3 | FGFR1 |
| PRPF19 | | | SPOP | HOXB5 | FGFR2 |
| PRPS1 | | | PPP1R9B | HOXB7 | FGFR3 |
| PRPS1L1 | | | XYLT2 | IGF2BP1 | FGFR4 |
| PSEN2 | | | SPAG9 | PPP1R9B | NGFR |
| PTGS1 | | | CA10 | BCAS3 | NTRK1 |
| PTK2B | | | TBX2 | WIPI1 | NTRK2 |
| PYGL | | | CEP95 | GPRC5C | INSR |
| RAB8A | | | HN1 | GPS1 | IGF1R |
| RAD54B | | | SOCS3 | SMAD2 | PDGFRA |
| RAPGEF1 | | | PCYT2 | CTIF | PDGFRB |
| RASGRP2 | | | TGIF1 | DYM | CSF1R |
| RBMS1 | | | VAPA | ZADH2 | KIT |
| RGL1 | | | RNF125 | DAZAP1 | FLT3 |
| RHBDF2 | | | CELF4 | STXBP2 | FLT1 |
| RIT1 | | | CTIF | ILF3 | FLT4 |
| RSU1 | | | SMAD7 | QTRT1 | KDR |
| RUNX1 | | | DYM | ELAVL3 | MET |
| S100A10 | | | DAZAP1 | ARMC6 | TEK |
| SAMSN1 | | | ILF3 | SUGP2 | EPHA2 |
| SCARB1 | | | ELAVL3 | CEBPA | GRB2 |
| SDCBP | | | TNPO2 | KCTD15 | SOS1 |
| SERTAD1 | | | RTBDN | ZNF565 | SOS2 |
| SH3KBP1 | | | SUGP2 | RPS19 | HRAS |
| SKP1 | | | CEBPA | KCNN4 | KRAS |
| SLC25A26 | | | SHKBP1 | CD3EAP | NRAS |
| SLC39A8 | | | BCKDHA | PRMT1 | RRAS |
| SLTM | | | EXOSC5 | SYT3 | RRAS2 |
| SMC3 | | | RPS19 | U2AF2 | MRAS |
| SNRPF | | | ZNF574 | ODC1 | ARAF |
| SNX11 | | | SIX5 | OSR1 | BRAF |

| | | | |
|---|---|---|---|
| SORT1 | CALM3 | SDC1 | RAF1 |
| SPRED1 | DBP | CGREF1 | MAP2K1 |
| SQSTM1 | SNRNP70 | CAD | MAP2K2 |
| ST7 | PTOV1 | USP34 | LAMTOR3 |
| STK17B | UBE2S | KCMF1 | MAPK1 |
| STRBP | CALM2 | MAT2A | MAPK3 |
| SWAP70 | KDM3A | ATOH8 | MKNK1 |
| TBL1X | KANSL3 | KDM3A | MKNK2 |
| TBXAS1 | AMMECR1L | METAP1D | RPS6KA3 |
| TCF7L2 | ACVR2A | NEUROD1 | RPS6KA1 |
| TEC | RND3 | SATB2 | RPS6KA2 |
| THBD | NEUROD1 | NOP58 | RPS6KA6 |
| THBS1 | FAM117B | EEF1B2 | ATF4 |
| THOC5 | EEF1B2 | NDUFS1 | ELK1 |
| TK1 | NDUFS1 | KANSL1L | ELK4 |
| TMEM107 | ANKZF1 | TUBA4A | MYC |
| TMX4 | NCL | NCL | SRF |
| TOB1 | PTMA | PTMA | FOS |
| TRIM27 | SOX12 | BOK | MAPT |
| TRPV2 | SCRT2 | SCRT2 | STMN1 |
| TULP4 | SNAP25 | NOP56 | PLA2G4E |
| UCK2 | SNX5 | SNX5 | PLA2G4A |
| VAMP3 | INSM1 | ASXL1 | JMJD7-PLA2G4B |
| VCAN | XRN2 | NOL4L | PLA2G4B |
| VGLL4 | FOXA2 | TOP1 | PLA2G4C |
| VOPP1 | ID1 | SLC12A5 | PLA2G4D |
| WIPI1 | TRPC4AP | SLC17A9 | PLA2G4F |
| ZBTB44 | DLGAP4 | MRPL40 | TNF |
| ZNF281 | TGIF2 | HIRA | IL1A |
| ZFP36 | KLRD1 | RANBP1 | IL1B |
| ZFP36L1 | TFAP2C | TRMT2A | TGFB1 |
| ZYX | RANBP1 | EWSR1 | TGFB2 |
| C7orf50 | TRMT2A | CYP2D6 | TGFB3 |
| C4orf46 | MMP11 | DAZL | TNFRSF1A |
| CXorf21 | EWSR1 | CAMKV | IL1R1 |
| ABCA13 | PES1 | MON1A | IL1RAP |
| ABCG1 | SUN2 | IFRD2 | TGFBR1 |
| ABHD2 | PDGFB | MANF | TGFBR2 |
| ABI2 | CYP2D6 | ACY1 | FASLG |
| ACOT9 | TSEN2 | SUCLG2 | FAS |
| ACOX1 | TMEM43 | FAM19A4 | CD14 |
| ACOX2 | DAZL | QTRTD1 | RAC1 |
| ACP6 | OSBPL10 | KIAA1407 | RAC2 |
| ACSL3 | PLCD1 | HSPBAP1 | RAC3 |
| ACVR1B | CAMKV | DIRC2 | CDC42 |
| ACYP1 | PCBP4 | TMEM108 | TRADD |
| ADAMTS9 | ACY1 | YEATS2 | CASP3 |
| ADAMTSL4 | BAP1 | TFRC | TRAF2 |
| ADCY7 | PHF7 | MXD4 | DAXX |
| ADCY9 | SEC61A1 | LYAR | MYD88 |
| ADK | CNBP | ZBTB49 | IRAK1 |
| ADORA2A | H1FX | CRMP1 | IRAK4 |
| ADORA3 | NAT8L | LAP3 | TRAF6 |
| AGTRAP | CRMP1 | BEND4 | GADD45A |
| AHCY | KIT | PPAT | GADD45B |
| AHI1 | HNRNPD | PAICS | GADD45G |
| AK1 | HNRNPDL | SRP72 | TAB1 |
| AK2 | ENOPH1 | HNRNPDL | TAB2 |
| AK3 | UBE2D3 | ENOPH1 | ECSIT |
| AKAP13 | ELOVL6 | EIF4E | MAP4K3 |
| AKAP7 | PGRMC2 | TET2 | MAP4K4 |
| AKIRIN1 | ABCE1 | HHIP | MAP4K1 |
| AKIRIN2 | ANAPC10 | RAI14 | PAK1 |
| ALAS1 | ENPP6 | IQGAP2 | PAK2 |
| ALDH1A2 | IRX2 | SNCAIP | STK4 |
| ALKBH3 | C5orf38 | ADAMTS19 | STK3 |
| ANKRD26 | NSUN2 | UBE2B | MAP4K2 |
| ANKRD33 | FAM173B | HSPA9 | MAP3K8 |
| ANKRD46 | CCT5 | PCDHA10 | MAP3K1 |
| ANKRD55 | RAI14 | DIAPH1 | MAP3K11 |

| | | | |
|---|---|---|---|
| ANTXR1 | NADK2 | TCERG1 | MAP3K2 |
| ANTXR2 | ISL1 | PPARGC1B | MAP3K3 |
| ANXA2 | RAPGEF6 | TCOF1 | MAP3K13 |
| AP3M2 | CXCL14 | NPM1 | MAP3K12 |
| APAF1 | ETF1 | PRR7 | MAP3K20 |
| APBB1IP | PAIP2 | HMGA1 | MAP3K6 |
| ARHGAP15 | DIAPH1 | RUNX2 | MAP3K5 |
| ARHGAP20 | PPARGC1B | KCNQ5 | MAP3K7 |
| ARHGAP26 | TCOF1 | SYNCRIP | MAP3K4 |
| ARID2 | WWC1 | POU3F2 | TAOK2 |
| ARL8B | NPM1 | ATG5 | TAOK3 |
| ARPC2 | RBM24 | FOXO3 | TAOK1 |
| ASAP1 | SOX4 | GJA1 | MAP2K4 |
| ASH2L | HMGA1 | MAP7 | MAP2K7 |
| ASRGL1 | TAF11 | ETV1 | MAP2K3 |
| ATP2B3 | ANKS1A | MEOX2 | MAP2K6 |
| ATP6V0D1 | CPNE5 | HOXA3 | MAPK8IP1 |
| ATP6V0E1 | PTK7 | HOXA11 | MAPK8IP2 |
| ATP8B1 | SRF | BHLHA15 | MAPK8IP3 |
| ATP8B4 | VEGFA | PDIA4 | FLNA |
| C7orf25 | PTCHD4 | ZBTB10 | FLNC |
| B4GALT5 | NR2E1 | PABPC1 | FLNB |
| BAMBI | RNF146 | UBR5 | CRK |
| BANK1 | ZDHHC14 | DCAF13 | CRKL |
| BASP1 | EIF3B | SLC25A32 | ARRB1 |
| BCAP31 | SDK1 | RSPO2 | ARRB2 |
| BCAR3 | ANKMY2 | ZHX2 | MAPK8 |
| BCL2L12 | BZW2 | TRIB1 | MAPK10 |
| BCLAF1 | HOXA9 | AGO2 | MAPK9 |
| BICD1 | ZNRF2 | RCL1 | MAPK11 |
| BID | TMEM248 | SIGMAR1 | MAPK12 |
| BMP2K | TSC22D4 | DNAJB5 | MAPK13 |
| BMP8A | LAMB1 | GADD45G | MAPK14 |
| BMX | ZNF800 | ABCA1 | MAPKAPK5 |
| BNIP3 | KCNH2 | NR6A1 | MAPKAPK2 |
| BRI3BP | PRKAG2 | PTGES2 | MAPKAPK3 |
| BTG3 | EGR3 | GPM6B | RPS6KA5 |
| BUB1B | PABPC1 | PRDX4 | RPS6KA4 |
| C5AR1 | KLF10 | UBA1 | CDC25B |
| CAB39 | EBAG9 | EFNB1 | NFATC1 |
| CABLES1 | TRIB1 | TIMM8A | NFATC3 |
| CACNA2D3 | SLC1A1 | PRPS1 | JUN |
| CAMP | AK3 | GPC3 | JUND |
| CAPN3 | RCL1 | | ATF2 |
| CAPRIN1 | FAM214B | | TP53 |
| CAPZA2 | ZNF367 | | DDIT3 |
| CAPZB | NSMF | | MAX |
| CA2 | TXLNG | | MEF2C |
| CASK | RBBP7 | | HSPB1 |
| CBLB | SCML2 | | AKT1 |
| CBR4 | RPS6KA3 | | AKT2 |
| CCDC14 | CDK16 | | AKT3 |
| CCDC34 | SLC25A5 | | PPM1A |
| CCL24 | STAG2 | | PTPRR |
| CCL4 | APLN | | PTPN5 |
| CCL5 | | | PTPN7 |
| CCM2 | | | DUSP1 |
| CCNB2 | | | DUSP4 |
| CCND1 | | | DUSP2 |
| CCND3 | | | DUSP7 |
| CD109 | | | DUSP8 |
| CD163 | | | DUSP5 |
| CD300A | | | DUSP16 |
| CD300C | | | DUSP6 |
| CD36 | | | DUSP9 |
| CD44 | | | DUSP10 |
| CD53 | | | DUSP3 |
| CD84 | | | PPP5C |
| CD86 | | | PPM1B |
| CD9 | | | HSPA8 |

CDC14A
CDC25C
CDC42BPB
CDC42EP3
CDC42EP4
CDCA7
CDH23
CDK5R1
CDK5RAP2
CDKN1A
CDKN3
CENPC
CENPF
CENPJ
CEP76
CHD7
CHMP2B
CHST12
CHST13
CKS2
CLCC1
CLINT1
CLIP2
CLIP4
COL6A1
COMMD9
COQ10A
COQ2
CREG1
CRTAP
CSNK1D
CSTB
CTNS
CTSC
CX3CR1
CXCL16
CXCR4
CYCS
CYP1B1
CYTH1
CYTH4
CYTIP
DAB2
DACH1
DAG1
DAPK2
DCK
DCLRE1A
DCLRE1B
DDAH1
DDX54
DENND4C
DEPDC1
DEPDC1B
DHDDS
DHRS11
DHRS3
DHRS9
DIP2B
DLGAP5
DMRT2
DMXL2
DNAJB12
DNAJC15
DNAJC19
DOCK10
DOCK11
DOCK5
DPYD
DPYSL2

HSPA1A
HSPA2
HSPA1L
HSPA1B
HSPA6
MECOM
MAP2K5
MAPK7
NR4A1
MAP3K14
CHUK
IKBKB
IKBKG
NLK
NFKB1
NFKB2
RELA
RELB

DSN1
DSTYK
DTL
DUSP4
DUSP5
DYNC2LI1
E2F3
E2F7
EBF3
EBNA1BP2
ECM1
ECT2
EFR3A
EGR2
EHD4
EID1
EIF1AX
EIF2B2
EIF3L
ELMO1
EMP1
ENTPD1
EPDR1
EPRS
ERCC6L
ERLIN2
ERMP1
ESD
ETNK1
ETV6
EXOSC6
EZH2
EZR
FABP4
FAM102B
FAM107B
FAM110B
FAM117A
FAM117B
FAM122A
FAM129B
FAM174A
FAM49B
FAM76B
FARS2
FBP1
FBXO3
FBXO4
FBXO43
FBXW11
FECH
FGFRL1
FIGNL1
FILIP1L
FKBPL
FLNA
FLRT2
FLT3
FMNL2
FNBP1
FNDC3B
FOS
FOSB
FOXN2
FOXO1
FRAT2
FST
FTH1
FYB1
GAB3

GALC
GALNT2
GALNT6
GARS
GAS2L3
GAS7
GATA2
GFOD1
GLDC
GLIPR1
GM2A
C17orf67
GMDS
GPD1L
GPR18
GPR180
GPR65
GPR88
GPX1
GSR
GTSF1
H2AFX
H2AFY
HACL1
HBP1
HCK
HECW2
HES6
HEXB
HIC2
HIST1H3I
HIST1H4A
HIST2H2AB
HIST1H4B
HIVEP1
HIVEP2
HK3
HLTF
HLX
HNF4G
HNRNPU
HP
HPCAL1
HPS5
HSPB1
HYLS1
IARS
IDH1
IDH2
IER3
IFI16
IFNAR1
IFNAR2
IFNGR1
IFT122
IFT52
IFT57
IFT88
IGFBP3
IGFBP7
IL10RA
IL12RB1
IL12RB2
IL1A
IL1R1
IL21R
IL23A
IL4R
IMPA2
IMPAD1

IMPDH1
INPP5D
INTS12
IP6K2
IPCEF1
IPO11
IQGAP1
IQGAP2
IRAK1BP1
IRF8
IRX3
ISM1
ITGA4
ITGA5
ITGAL
ITGAV
ITGB5
ITM2B
ITPR1
IVNS1ABP
JAZF1
JUN
JUND
KCMF1
KCND2
KCNJ2
KCNK13
KHDRBS3
KIF20B
KIF24
KLHL2
KLHL5
KPNA4
KRAS
KYNU
LACTB
LAMP2
LDHD
LFNG
LGALS1
LGALS3
LGI2
LGMN
LHFPL2
LIMK1
LIPT1
LMNB1
LPAR1
LPAR6
LPGAT1
LPL
LPP
LRIG1
LRP1
LRRC39
LRRC3B
LTA4H
LTB4R
LY86
LYL1
LYN
LYST
MAD1L1
MAD2L1
MAF
MALT1
MAN2A2
MANBA
MANF
MAP3K1

MAPK9
MAPKAPK2
MB
MCCC1
MCCC2
MDC1
MDK
MEF2D
MEFV
MEIS2
MERTK
METTL9
MFSD5
MID1
MINPP1
MMACHC
MMP14
MND1
MNX1
MORF4L2
MPP1
MPZL1
MRPL44
MRPL45
MRPL48
MRPS27
MRPS33
MRPS6
MSI2
MSRA
MSRB3
MTSS1
MYCBP
MYCBP2
MYL6B
MYO1E
MYO1F
MYO5C
MYOF
NAB2
NACC2
NAMPT
NAP1L3
NAP1L5
NAPEPLD
NAV3
NCALD
NCAM2
NCF1
NCF4
NDRG1
NECAP1
NEIL3
NEK2
NEK6
NEU1
NGLY1
NINJ1
NIPAL3
NIT2
NLN
NME7
NPAS1
NPC1
NT5C3A
NT5DC2
NT5E
NUB1
NUCB2
NUF2

NUMB
NUP214
NUP35
NXT1
ODC1
OLIG2
OPLAH
OSBP2
OSBPL8
OSM
OSTM1
OVOL2
OXCT1
OXSM
P2RX4
P2RY2
PACSIN2
PALLD
PAPSS1
PAPSS2
PARVG
PATL1
PCDH7
PCSK6
PDE7B
PDGFC
PDSS2
PDXK
PECAM1
PELI2
PENK
PEX19
PHACTR4
PHF21A
PHF23
PHF7
PHLPP1
PIGK
PIGV
PIK3CB
PIK3R1
PIM1
PIN4
PIP4K2A
PKDCC
PLA2G15
PLAU
PLAUR
PLCL2
PLD1
PLEK
PLEKHB2
PLEKHO1
PLIN2
PLOD3
PMP22
POR
POU2F2
PPARGC1A
PPM1E
PRDM1
PREB
PRKCA
PRKCB
PRKCD
PRKCQ
PRMT6
PRNP
PRSS12
PSAP

PSMG1
PSTPIP2
PTGDS
PTPRC
PTPRE
PTPRF
PTPRG
PTPRN2
PTTG1
PWWP2B
PXK
PYROXD2
RAB11FIP1
RAB27A
RAB31
RABAC1
RAC2
RAP1B
RASA3
RASSF2
RASSF4
RBM17
RBM38
RBM47
RCAN1
RCBTB2
RCN3
RDH11
REPS2
RFC3
RGS2
RHOH
RIMS3
RIN3
RND3
RNF149
RNF24
RNLS
ROPN1L
RPLP0
RPS17
RRAGC
RTKN2
RTN4R
RUSC1
RUSC2
RYBP
S100A4
S1PR3
SAE1
SASH1
SCARB2
SCG2
SCLT1
SDC2
SEMA3A
SEMA4D
SERPINB2
SERPINI2
SESN3
SETBP1
SETD4
SGK1
SH2D2A
SH3BGRL2
SH3RF1
SH3TC1
SIGLEC9
SIGLEC10
SIL1

SIPA1L1
SIRPA
SIX1
SIX4
SLA
SLC16A3
SLC20A1
SLC20A2
SLC22A4
SLC25A10
SLC25A28
SLC26A1
SLC26A11
SLC27A3
SLC29A3
SLC37A1
SLC38A2
SLC44A1
SLC9A1
SLC9A7
SLPI
SMAD3
SMAD7
SMAP1
SMARCA2
SMARCD3
SMYD3
SNAI3
SNAPC3
SNTB2
SNX10
SNX16
SNX19
SNX29
SORCS2
SORL1
SPIRE1
SPN
SPOCD1
SPOCK1
SPOPL
SPP1
SPRED2
SPRY2
SPSB1
SRGAP1
SSH1
SSH2
SSX2IP
ST3GAL1
ST3GAL2
ST3GAL3
ST3GAL4
ST3GAL6
STAT3
STBD1
STEAP1
STEAP3
STK39
STK4
STXBP5
SULF2
SYN2
SYPL1
SYS1
TAF9B
TAP2
TARSL2
TASP1
TAX1BP1

TBC1D2
TBC1D23
TBC1D4
TBC1D9B
TBRG4
TCIRG1
TCTEX1D2
TDRD7
TEX2
TGM2
TIMELESS
TLE3
TLE6
TLR4
TM2D3
TM4SF19
TMBIM4
TMEM120A
TMEM30A
TMEM38B
TMEM44
TMEM87B
TMEM97
TMPO
TNFAIP3
TNFRSF12A
TNFRSF14
TNFRSF1B
TNFRSF21
TNS3
TOR1A
TRAP1
TRIB1
TRIB2
TRIM33
TRIM45
TRMT112
TROAP
TP53BP2
TRPM2
TSC22D4
TSHZ3
TSPAN14
TSPYL5
TTC26
TTLL5
TUBGCP3
UBE2F
UGT3A1
ULK1
UPB1
USP1
USP38
USP46
USP47
USP54
USP9X
UVRAG
VAC14
VAMP8
VAV1
VAV3
VIM
VPS11
VWF
WDR26
WDR48
WEE1
XPO6
YIF1B

ZC3H12A
ZDHHC14
ZNF213
ZNF239
ZNF395
ZNF22
ZNF438
ZNF777
ZFYVE26
ZHX3
ZMIZ1
ZSCAN2
ZSWIM6
