## Supplementary Table S8 for "Haplotype-aware single-cell multiomics uncovers functional effects of somatic structural variation"

**Table S8. Cell-wise genotypes and correlation matrix of NA12878 deletions on chr19 and chr22 determined using the ArbiGent tool**

| | | chr22:22200000-22900000 | | |
|---|---|---|---|---|
| | | Reference | Deleted | Unclear |
| chr19:36500000-3700000 | Reference | 0 | 62 | 7 |
| | Deleted | 15 | 0 | 0 |
| | Unclear | 2 | 8 | 1 |

| | | chr22:22694696-22899260 | | | |
|---|---|---|---|---|---|
| | | Reference | Deleted (hom) | Deleted (het) | Unclear |
| chr19:36500000 | Reference | 7 | 36 | 21 | 5 |
| | Deleted | 15 | 0 | 0 | 0 |
| | Unclear | 3 | 8 | 0 | 0 |

| cellname | chrom | start | end | class | expected | W observed | C observed | del_h1 | del_h2 | del_hom | top.score | TOP_SV | Batch |
|---|---|---|---|---|---|---|---|---|---|---|---|---|---|
| NW130711.291.L008 | chr19 | 36500000 | 37000000 | WC | 109.699 | 28 | 68 | -1.68 | -5.238 | -6.798 | -1.68 | ref_hom | Batch2 |
| NW130711.320.L008 | chr19 | 36500000 | 37000000 | CC | 40.192 | 0 | 10 | 1.196 | 1.196 | 0.194 | 1.196 | del | Batch2 |
| NW150212-IV.43.L002 | chr19 | 36500000 | 37000000 | CC | 76.674 | 0 | 47 | 0.35 | 0.35 | -3.415 | 0.35 | unclear | Batch1 |
| NW150212-IV.44.L002 | chr19 | 36500000 | 37000000 | WW | 85.522 | 60 | 1 | -0.285 | -0.285 | -4.84 | -0.285 | unclear | Batch1 |
| NW150212-IV.45.L002 | chr19 | 36500000 | 37000000 | WW | 53.826 | 32 | 1 | -0.088 | -0.088 | -2.897 | -0.088 | unclear | Batch1 |
| NW150212-IV.46.L002 | chr19 | 36500000 | 37000000 | WC | 101.307 | 37 | 38 | -2.877 | -2.975 | -5.76 | -2.877 | ref_hom | Batch1 |
| NW150212-IV.50.L002 | chr19 | 36500000 | 37000000 | CC | 105.116 | 1 | 92 | -1.06 | -1.06 | -7.388 | -1.06 | ref_hom | Batch1 |
| NW150212-IV.58.L002 | chr19 | 36500000 | 37000000 | CC | 120.565 | 1 | 93 | -0.566 | -0.566 | -7.131 | -0.566 | ref_hom | Batch1 |
| NW150212-IV.60.L002 | chr19 | 36500000 | 37000000 | CC | 123.439 | 2 | 131 | -2.175 | -2.175 | -10.408 | -2.175 | ref_hom | Batch1 |
| NW150212-IV.62.L002 | chr19 | 36500000 | 37000000 | WW | 176.057 | 133 | 3 | -0.704 | -0.704 | -9.826 | -0.704 | ref_hom | Batch1 |
| NW150212-IV.63.L002 | chr19 | 36500000 | 37000000 | CC | 66.55 | 3 | 36 | 0.092 | 0.092 | -2.96 | 0.092 | unclear | Batch1 |
| NW150212-IV.64.L002 | chr19 | 36500000 | 37000000 | CC | 127.539 | 3 | 50 | 2.139 | 2.139 | -1.516 | 2.139 | del | Batch1 |
| NW150212-IV.65.L002 | chr19 | 36500000 | 37000000 | CC | 137.659 | 2 | 123 | -1.456 | -1.456 | -9.633 | -1.456 | ref_hom | Batch1 |
| NW150212-IV.68.L002 | chr19 | 36500000 | 37000000 | CW | 138.21 | 49 | 70 | -5.337 | -3.523 | -8.736 | -3.523 | ref_hom | Batch1 |
| NW150212-IV.69.L002 | chr19 | 36500000 | 37000000 | CC | 101.868 | 0 | 75 | -0.099 | -0.099 | -5.574 | -0.099 | unclear | Batch1 |
| NW150212-IV.72.L002 | chr19 | 36500000 | 37000000 | CW | 94.801 | 35 | 28 | -1.988 | -2.749 | -4.676 | -1.988 | ref_hom | Batch1 |
| NW150212-IV.74.L002 | chr19 | 36500000 | 37000000 | CC | 125.037 | 1 | 80 | 0.273 | 0.273 | -5.613 | 0.273 | unclear | Batch1 |
| NW150212-IV.77.L002 | chr19 | 36500000 | 37000000 | WC | 145.788 | 69 | 60 | -5.233 | -4.481 | -9.574 | -4.481 | ref_hom | Batch1 |
| NW150212-IV.81.L002 | chr19 | 36500000 | 37000000 | CW | 125.162 | 50 | 53 | -4.07 | -3.802 | -7.756 | -3.802 | ref_hom | Batch1 |
| NW150212-IV.83.L002 | chr19 | 36500000 | 37000000 | WW | 159.371 | 139 | 2 | -1.434 | -1.434 | -10.641 | -1.434 | ref_hom | Batch1 |
| NW150212-IV.84.L002 | chr19 | 36500000 | 37000000 | WW | 148.021 | 117 | 2 | -0.846 | -0.846 | -8.884 | -0.846 | ref_hom | Batch1 |
| NW150212-IV.85.L002 | chr19 | 36500000 | 37000000 | WW | 128.61 | 86 | 4 | -0.221 | -0.221 | -6.471 | -0.221 | unclear | Batch1 |
| NW150212-IV.87.L002 | chr19 | 36500000 | 37000000 | CC | 143.369 | 2 | 105 | -0.444 | -0.444 | -7.831 | -0.444 | ref_hom | Batch1 |
| NW150212-IV.89.L002 | chr19 | 36500000 | 37000000 | WW | 136.763 | 128 | 1 | -1.536 | -1.536 | -9.911 | -1.536 | ref_hom | Batch1 |
| NW150212-IV.91.L002 | chr19 | 36500000 | 37000000 | WW | 96.617 | 85 | 2 | -1.167 | -1.167 | -7.063 | -1.167 | ref_hom | Batch1 |
| NW130711.323.L008 | chr19 | 36500000 | 37000000 | CW | 53.724 | 28 | 24 | -2.418 | -2.73 | -4.954 | -2.418 | ref_hom | Batch2 |
| NW150212-IV.93.L002 | chr19 | 36500000 | 37000000 | CC | 159.948 | 0 | 65 | 3.256 | 3.256 | -1.366 | 3.256 | del | Batch1 |
| NW130711.324.L008 | chr19 | 36500000 | 37000000 | CC | 52.949 | 0 | 47 | -0.564 | -0.564 | -4.161 | -0.564 | ref_hom | Batch2 |
| NW130711.325.L008 | chr19 | 36500000 | 37000000 | WC | 52.68 | 28 | 23 | -2.735 | -2.344 | -4.886 | -2.344 | ref_hom | Batch2 |
| NW130711.327.L008 | chr19 | 36500000 | 37000000 | CC | 75.184 | 1 | 80 | -1.484 | -1.484 | -6.797 | -1.484 | ref_hom | Batch2 |
| NW130711.329.L008 | chr19 | 36500000 | 37000000 | CW | 78.252 | 34 | 15 | -0.597 | -2.866 | -3.411 | -0.597 | ref_hom | Batch2 |
| NW130711.345.L008 | chr19 | 36500000 | 37000000 | CC | 70.528 | 0 | 57 | -0.403 | -0.403 | -4.677 | -0.403 | ref_hom | Batch2 |
| NW130711.350.L008 | chr19 | 36500000 | 37000000 | CC | 85.546 | 0 | 76 | -0.722 | -0.722 | -6.06 | -0.722 | ref_hom | Batch2 |
| NW130711.354.L008 | chr19 | 36500000 | 37000000 | CW | 54.525 | 30 | 32 | -3.036 | -2.902 | -5.711 | -2.902 | ref_hom | Batch2 |
| NW130711.355.L008 | chr19 | 36500000 | 37000000 | CW | 46.209 | 25 | 20 | -2.163 | -2.552 | -4.521 | -2.163 | ref_hom | Batch2 |
| NW130711.362.L008 | chr19 | 36500000 | 37000000 | CC | 82.627 | 0 | 76 | -0.815 | -0.815 | -6.108 | -0.815 | ref_hom | Batch2 |
| NW130711.365.L008 | chr19 | 36500000 | 37000000 | CW | 62.058 | 33 | 35 | -3.218 | -3.08 | -6.075 | -3.08 | ref_hom | Batch2 |
| NW130711.296.L008 | chr19 | 36500000 | 37000000 | CC | 95.336 | 0 | 92 | -1.064 | -1.064 | -7.235 | -1.064 | ref_hom | Batch2 |
| NW130711.374.L008 | chr19 | 36500000 | 37000000 | CC | 65.62 | 0 | 86 | -1.6 | -1.6 | -6.873 | -1.6 | ref_hom | Batch2 |

| | | | | | | | | | | | | | |
|---|---|---|---|---|---|---|---|---|---|---|---|---|---|
| NW130711.376.L008 | chr19 | 36500000 | 37000000 | CC | 65.189 | 0 | 77 | -1.329 | -1.329 | -6.305 | -1.329 | ref_hom | Batch2 |
| NW130711.377.L008 | chr19 | 36500000 | 37000000 | WW | 35.749 | 33 | 0 | -0.535 | -0.535 | -3.287 | -0.535 | ref_hom | Batch2 |
| NW150212-III.03.L001 | chr19 | 36500000 | 37000000 | WC | 129.33 | 47 | 50 | -3.41 | -3.701 | -7.041 | -3.41 | ref_hom | Batch1 |
| NW150212-III.05.L001 | chr19 | 36500000 | 37000000 | CW | 150.226 | 84 | 67 | -5.105 | -6.373 | -11.27 | -5.105 | ref_hom | Batch1 |
| NW150212-III.06.L001 | chr19 | 36500000 | 37000000 | WW | 153.692 | 133 | 1 | -1.203 | -1.203 | -10.06 | -1.203 | ref_hom | Batch1 |
| NW150212-III.07.L001 | chr19 | 36500000 | 37000000 | WC | 117.827 | 57 | 15 | -4.282 | 1.116 | -3.228 | 1.116 | del | Batch1 |
| NW150212-III.09.L001 | chr19 | 36500000 | 37000000 | CW | 145.457 | 47 | 50 | -3.373 | -3.051 | -6.431 | -3.051 | ref_hom | Batch1 |
| NW130711.300.L008 | chr19 | 36500000 | 37000000 | WW | 39.24 | 38 | 0 | -0.628 | -0.628 | -3.639 | -0.628 | ref_hom | Batch2 |
| NW150212-III.11.L001 | chr19 | 36500000 | 37000000 | CW | 131.484 | 37 | 73 | -5.544 | -2.282 | -7.729 | -2.282 | ref_hom | Batch1 |
| NW150212-III.12.L001 | chr19 | 36500000 | 37000000 | WC | 128.212 | 44 | 63 | -3.166 | -4.855 | -7.907 | -3.166 | ref_hom | Batch1 |
| NW150212-III.13.L001 | chr19 | 36500000 | 37000000 | CW | 84.279 | 10 | 28 | -2.091 | 0.809 | -1.338 | 0.809 | del | Batch1 |
| NW150212-III.16.L001 | chr19 | 36500000 | 37000000 | WW | 47.548 | 42 | 2 | -0.91 | -0.91 | -4.213 | -0.91 | ref_hom | Batch1 |
| NW150212-III.21.L001 | chr19 | 36500000 | 37000000 | CW | 151.944 | 52 | 64 | -4.679 | -3.532 | -8.157 | -3.532 | ref_hom | Batch1 |
| NW150212-III.26.L001 | chr19 | 36500000 | 37000000 | CW | 128.913 | 44 | 42 | -2.841 | -3.056 | -5.88 | -2.841 | ref_hom | Batch1 |
| NW130711.302.L008 | chr19 | 36500000 | 37000000 | WW | 70.81 | 56 | 0 | -0.347 | -0.347 | -4.575 | -0.347 | unclear | Batch2 |
| NW150212-III.27.L001 | chr19 | 36500000 | 37000000 | CW | 203.072 | 13 | 86 | -5.548 | 6.519 | 0.523 | 6.519 | del | Batch1 |
| NW150212-III.30.L001 | chr19 | 36500000 | 37000000 | WW | 164.829 | 127 | 1 | -0.545 | -0.545 | -9.255 | -0.545 | ref_hom | Batch1 |
| NW150212-III.33.L001 | chr19 | 36500000 | 37000000 | CW | 109.056 | 55 | 51 | -4.129 | -4.437 | -8.373 | -4.129 | ref_hom | Batch1 |
| NW150212-III.35.L001 | chr19 | 36500000 | 37000000 | WC | 87.346 | 32 | 22 | -2.555 | -1.388 | -3.897 | -1.388 | ref_hom | Batch1 |
| NW150212-III.36.L001 | chr19 | 36500000 | 37000000 | WW | 155.432 | 108 | 1 | 0.031 | 0.031 | -7.606 | 0.031 | unclear | Batch1 |
| NW150212-III.39.L001 | chr19 | 36500000 | 37000000 | WC | 58.106 | 30 | 3 | -2.678 | 1.627 | -1.032 | 1.627 | del | Batch1 |
| NW150212-III.42.L001 | chr19 | 36500000 | 37000000 | WW | 118.871 | 41 | 0 | 2.954 | 2.954 | 0.053 | 2.954 | del | Batch1 |
| NW150212-III.48.L001 | chr19 | 36500000 | 37000000 | CW | 155.347 | 82 | 66 | -4.94 | -6.193 | -10.955 | -4.94 | ref_hom | Batch1 |
| NW150212-III.52.L001 | chr19 | 36500000 | 37000000 | CW | 130.697 | 66 | 48 | -3.554 | -5.088 | -8.506 | -3.554 | ref_hom | Batch1 |
| NW150212-III.55.L001 | chr19 | 36500000 | 37000000 | WW | 119.11 | 123 | 0 | -1.557 | -1.557 | -9.4 | -1.557 | ref_hom | Batch1 |
| NW150212-III.57.L001 | chr19 | 36500000 | 37000000 | CC | 105.033 | 0 | 86 | -0.5 | -0.5 | -6.553 | -0.5 | ref_hom | Batch1 |
| NW150212-III.59.L001 | chr19 | 36500000 | 37000000 | CW | 80.19 | 28 | 39 | -3.348 | -2.367 | -5.57 | -2.367 | ref_hom | Batch1 |
| NW150212-III.63.L001 | chr19 | 36500000 | 37000000 | CW | 84.757 | 29 | 34 | -2.846 | -2.354 | -5.091 | -2.354 | ref_hom | Batch1 |
| NW150212-III.66.L001 | chr19 | 36500000 | 37000000 | WC | 107.377 | 40 | 12 | -2.878 | 1.434 | -1.553 | 1.434 | del | Batch1 |
| NW150212-III.68.L001 | chr19 | 36500000 | 37000000 | CW | 112.314 | 47 | 49 | -3.883 | -3.71 | -7.453 | -3.71 | ref_hom | Batch1 |
| NW130711.307.L008 | chr19 | 36500000 | 37000000 | WC | 110.536 | 59 | 57 | -4.752 | -4.609 | -9.136 | -4.609 | ref_hom | Batch2 |
| NW150212-III.69.L001 | chr19 | 36500000 | 37000000 | WC | 116.039 | 45 | 39 | -3.409 | -2.807 | -6.149 | -2.807 | ref_hom | Batch1 |
| NW150212-III.75.L001 | chr19 | 36500000 | 37000000 | CW | 107.701 | 11 | 40 | -2.86 | 1.736 | -1.246 | 1.736 | del | Batch1 |
| NW150212-III.76.L001 | chr19 | 36500000 | 37000000 | WW | 135.072 | 75 | 1 | 1.008 | 1.008 | -4.568 | 1.008 | del | Batch1 |
| NW150212-III.77.L001 | chr19 | 36500000 | 37000000 | CW | 44.762 | 22 | 21 | -2.26 | -2.339 | -4.407 | -2.26 | ref_hom | Batch1 |
| NW150212-III.80.L001 | chr19 | 36500000 | 37000000 | CW | 116.386 | 9 | 45 | -3.163 | 2.862 | -0.475 | 2.862 | del | Batch1 |
| NW150212-III.81.L001 | chr19 | 36500000 | 37000000 | WW | 84.339 | 37 | 0 | 1.371 | 1.371 | -1.635 | 1.371 | del | Batch1 |
| NW150212-III.84.L001 | chr19 | 36500000 | 37000000 | CC | 160.949 | 1 | 137 | -1.133 | -1.133 | -10.273 | -1.133 | ref_hom | Batch1 |
| NW150212-III.90.L001 | chr19 | 36500000 | 37000000 | CC | 67.994 | 2 | 26 | 0.946 | 0.946 | -1.275 | 0.946 | del | Batch1 |
| NW150212-III.92.L001 | chr19 | 36500000 | 37000000 | WC | 112.811 | 40 | 45 | -2.988 | -3.471 | -6.372 | -2.988 | ref_hom | Batch1 |
| NW150212-III.93.L001 | chr19 | 36500000 | 37000000 | CW | 148.199 | 58 | 60 | -4.395 | -4.211 | -8.523 | -4.211 | ref_hom | Batch1 |
| NW150212-III.94.L001 | chr19 | 36500000 | 37000000 | CW | 128.65 | 44 | 38 | -2.367 | -3.04 | -5.411 | -2.367 | ref_hom | Batch1 |
| NW150212-IV.08.L002 | chr19 | 36500000 | 37000000 | WC | 76.18 | 26 | 11 | -2.075 | 0.163 | -1.921 | 0.163 | unclear | Batch1 |
| NW150212-IV.09.L002 | chr19 | 36500000 | 37000000 | CW | 77.265 | 24 | 21 | -1.525 | -1.892 | -3.366 | -1.525 | ref_hom | Batch1 |
| NW150212-IV.10.L002 | chr19 | 36500000 | 37000000 | WC | 87.157 | 32 | 43 | -2.663 | -3.615 | -6.127 | -2.663 | ref_hom | Batch1 |
| NW150212-IV.11.L002 | chr19 | 36500000 | 37000000 | CW | 76.818 | 30 | 35 | -3.065 | -2.627 | -5.538 | -2.627 | ref_hom | Batch1 |
| NW150212-IV.12.L002 | chr19 | 36500000 | 37000000 | WW | 80.107 | 49 | 0 | 0.385 | 0.385 | -3.508 | 0.385 | unclear | Batch1 |
| NW150212-IV.15.L002 | chr19 | 36500000 | 37000000 | CW | 132.986 | 46 | 41 | -2.631 | -3.179 | -5.81 | -2.631 | ref_hom | Batch1 |
| NW150212-IV.17.L002 | chr19 | 36500000 | 37000000 | CC | 92.732 | 1 | 75 | -0.736 | -0.736 | -6.122 | -0.736 | ref_hom | Batch1 |
| NW150212-IV.24.L002 | chr19 | 36500000 | 37000000 | CW | 80.817 | 28 | 25 | -1.958 | -2.288 | -4.169 | -1.958 | ref_hom | Batch1 |
| NW150212-IV.25.L002 | chr19 | 36500000 | 37000000 | CC | 100.964 | 1 | 118 | -2.145 | -2.145 | -9.394 | -2.145 | ref_hom | Batch1 |

| | | | | | | | | | | | | | |
|---|---|---|---|---|---|---|---|---|---|---|---|---|---|
| NW150212-IV.26.L002 | chr19 | 36500000 | 37000000 | WW | 132.822 | 58 | 1 | 2.047 | 2.047 | -2.258 | 2.047 | del | Batch1 |
| NW150212-IV.31.L002 | chr19 | 36500000 | 37000000 | CW | 58.191 | 26 | 11 | -0.65 | -2.436 | -2.995 | -0.65 | ref_hom | Batch1 |
| NW150212-IV.34.L002 | chr19 | 36500000 | 37000000 | CW | 123.021 | 49 | 50 | -3.82 | -3.729 | -7.441 | -3.729 | ref_hom | Batch1 |
| NW150212-IV.39.L002 | chr19 | 36500000 | 37000000 | WC | 121.222 | 53 | 59 | -4.164 | -4.647 | -8.641 | -4.164 | ref_hom | Batch1 |
| NW150212-IV.41.L002 | chr19 | 36500000 | 37000000 | CW | 106.232 | 36 | 34 | -2.423 | -2.641 | -5.021 | -2.423 | ref_hom | Batch1 |
| NW130711.291.L008 | chr22 | 22200000 | 22900000 | WC | 153.578 | 94 | 0 | -6.452 | 11.377 | 4.565 | 11.377 | del | Batch2 |
| NW130711.320.L008 | chr22 | 22200000 | 22900000 | WC | 56.269 | 39 | 29 | -3.472 | -2.836 | -6.068 | -2.836 | ref_hom | Batch2 |
| NW150212-IV.43.L002 | chr22 | 22200000 | 22900000 | WW | 107.344 | 20 | 0 | 4.464 | 4.464 | 4.069 | 4.464 | del | Batch1 |
| NW150212-IV.44.L002 | chr22 | 22200000 | 22900000 | WW | 119.731 | 37 | 4 | 2.768 | 2.768 | 0.322 | 2.768 | del | Batch1 |
| NW150212-IV.45.L002 | chr22 | 22200000 | 22900000 | CW | 75.356 | 0 | 26 | -1.936 | 5.569 | 3.47 | 5.569 | del | Batch1 |
| NW150212-IV.46.L002 | chr22 | 22200000 | 22900000 | WC | 141.829 | 47 | 0 | -2.647 | 10.654 | 7.528 | 10.654 | del | Batch1 |
| NW150212-IV.50.L002 | chr22 | 22200000 | 22900000 | CW | 147.162 | 0 | 42 | -1.896 | 11.1 | 8.661 | 11.1 | del | Batch1 |
| NW150212-IV.58.L002 | chr22 | 22200000 | 22900000 | CW | 168.791 | 2 | 58 | -3.262 | 10.407 | 6.593 | 10.407 | del | Batch1 |
| NW150212-IV.60.L002 | chr22 | 22200000 | 22900000 | WW | 172.814 | 80 | 2 | 2.343 | 2.343 | -3.426 | 2.343 | del | Batch1 |
| NW150212-IV.62.L002 | chr22 | 22200000 | 22900000 | CW | 246.479 | 4 | 70 | -2.656 | 14.594 | 10.988 | 14.594 | del | Batch1 |
| NW150212-IV.63.L002 | chr22 | 22200000 | 22900000 | CC | 93.17 | 5 | 75 | -1.023 | -1.023 | -6.414 | -1.023 | ref_hom | Batch1 |
| NW150212-IV.64.L002 | chr22 | 22200000 | 22900000 | WW | 178.554 | 196 | 0 | -2.593 | -2.593 | -14.424 | -2.593 | ref_hom | Batch1 |
| NW150212-IV.65.L002 | chr22 | 22200000 | 22900000 | WC | 192.722 | 61 | 0 | -2.962 | 14.559 | 10.863 | 14.559 | del | Batch1 |
| NW150212-IV.68.L002 | chr22 | 22200000 | 22900000 | CW | 193.494 | 0 | 56 | -2.329 | 14.648 | 11.552 | 14.648 | del | Batch1 |
| NW150212-IV.69.L002 | chr22 | 22200000 | 22900000 | WC | 142.616 | 0 | 29 | 10.829 | -0.128 | 10.103 | 10.829 | del | Batch1 |
| NW150212-IV.72.L002 | chr22 | 22200000 | 22900000 | WC | 132.721 | 103 | 0 | -6.963 | 9.744 | 2.558 | 9.744 | del | Batch1 |
| NW150212-IV.74.L002 | chr22 | 22200000 | 22900000 | WC | 175.052 | 43 | 2 | -1.185 | 10.977 | 9.112 | 10.977 | del | Batch1 |
| NW150212-IV.77.L002 | chr22 | 22200000 | 22900000 | CW | 204.103 | 0 | 93 | -5.92 | 15.296 | 8.722 | 15.296 | del | Batch1 |
| NW150212-IV.81.L002 | chr22 | 22200000 | 22900000 | WW | 175.226 | 62 | 2 | 3.882 | 3.882 | -0.236 | 3.882 | del | Batch1 |
| NW150212-IV.83.L002 | chr22 | 22200000 | 22900000 | WW | 223.12 | 58 | 2 | 7.141 | 7.141 | 4.503 | 7.141 | del | Batch1 |
| NW150212-IV.84.L002 | chr22 | 22200000 | 22900000 | WW | 207.23 | 49 | 0 | 7.602 | 7.602 | 5.78 | 7.602 | del | Batch1 |
| NW150212-IV.85.L002 | chr22 | 22200000 | 22900000 | WW | 180.054 | 37 | 0 | 7.229 | 7.229 | 6.416 | 7.229 | del | Batch1 |
| NW150212-IV.87.L002 | chr22 | 22200000 | 22900000 | CC | 200.717 | 2 | 77 | 4.045 | 4.045 | -1.165 | 4.045 | del | Batch1 |
| NW150212-IV.89.L002 | chr22 | 22200000 | 22900000 | CC | 191.468 | 0 | 31 | 8.805 | 8.805 | 9.467 | 9.467 | del | Batch1 |
| NW150212-IV.91.L002 | chr22 | 22200000 | 22900000 | WC | 135.264 | 43 | 0 | -2.353 | 10.162 | 7.351 | 10.162 | del | Batch1 |
| NW130711.323.L008 | chr22 | 22200000 | 22900000 | CC | 75.213 | 0 | 51 | 0.069 | 0.069 | -3.929 | 0.069 | unclear | Batch2 |
| NW150212-IV.93.L002 | chr22 | 22200000 | 22900000 | WW | 223.927 | 265 | 2 | -4.305 | -4.305 | -19.629 | -4.305 | ref_hom | Batch1 |
| NW130711.324.L008 | chr22 | 22200000 | 22900000 | CC | 74.129 | 0 | 45 | 0.354 | 0.354 | -3.284 | 0.354 | unclear | Batch2 |
| NW130711.325.L008 | chr22 | 22200000 | 22900000 | CC | 73.752 | 0 | 42 | 0.514 | 0.514 | -2.929 | 0.514 | del | Batch2 |
| NW130711.327.L008 | chr22 | 22200000 | 22900000 | CC | 105.257 | 0 | 67 | 0.454 | 0.454 | -4.592 | 0.454 | unclear | Batch2 |
| NW130711.329.L008 | chr22 | 22200000 | 22900000 | CC | 109.552 | 2 | 62 | 0.498 | 0.498 | -4.237 | 0.498 | unclear | Batch2 |
| NW130711.345.L008 | chr22 | 22200000 | 22900000 | WW | 98.74 | 65 | 2 | -0.139 | -0.139 | -5.044 | -0.139 | unclear | Batch2 |
| NW130711.350.L008 | chr22 | 22200000 | 22900000 | CW | 119.764 | 2 | 64 | -4.623 | 6.641 | 1.791 | 6.641 | del | Batch2 |
| NW130711.354.L008 | chr22 | 22200000 | 22900000 | WW | 76.336 | 38 | 2 | 0.483 | 0.483 | -2.667 | 0.483 | unclear | Batch2 |
| NW130711.355.L008 | chr22 | 22200000 | 22900000 | CW | 64.693 | 0 | 41 | -3.338 | 4.68 | 1.303 | 4.68 | del | Batch2 |
| NW130711.362.L008 | chr22 | 22200000 | 22900000 | CW | 115.678 | 0 | 94 | -6.4 | 8.458 | 1.898 | 8.458 | del | Batch2 |
| NW130711.365.L008 | chr22 | 22200000 | 22900000 | WC | 86.882 | 49 | 0 | -3.756 | 6.369 | 2.476 | 6.369 | del | Batch2 |
| NW130711.296.L008 | chr22 | 22200000 | 22900000 | WW | 133.47 | 91 | 0 | 0.332 | 0.332 | -6.227 | 0.332 | unclear | Batch2 |
| NW130711.374.L008 | chr22 | 22200000 | 22900000 | WW | 91.868 | 49 | 0 | 0.921 | 0.921 | -2.96 | 0.921 | del | Batch2 |
| NW130711.376.L008 | chr22 | 22200000 | 22900000 | WC | 91.265 | 54 | 0 | -4.074 | 6.691 | 2.473 | 6.691 | del | Batch2 |
| NW130711.377.L008 | chr22 | 22200000 | 22900000 | CW | 50.049 | 0 | 47 | -3.606 | 3.539 | -0.015 | 3.539 | del | Batch2 |
| NW150212-III.03.L001 | chr22 | 22200000 | 22900000 | WC | 181.063 | 57 | 0 | -2.797 | 13.669 | 10.192 | 13.669 | del | Batch1 |
| NW150212-III.05.L001 | chr22 | 22200000 | 22900000 | WC | 210.316 | 162 | 2 | -10.502 | 13.212 | 2.289 | 13.212 | del | Batch1 |
| NW150212-III.06.L001 | chr22 | 22200000 | 22900000 | WC | 215.169 | 124 | 0 | -8.178 | 16.039 | 7.257 | 16.039 | del | Batch1 |
| NW150212-III.07.L001 | chr22 | 22200000 | 22900000 | CW | 164.958 | 102 | 122 | -8.861 | -7.733 | -16.19 | -7.733 | ref_hom | Batch1 |
| NW150212-III.09.L001 | chr22 | 22200000 | 22900000 | CC | 203.64 | 0 | 118 | 1.827 | 1.827 | -6.562 | 1.827 | del | Batch1 |

| | | | | | | | | | | | | | |
|---|---|---|---|---|---|---|---|---|---|---|---|---|---|
| NW130711.300.L008 | chr22 | 22200000 | 22900000 | WC | 54.936 | 34 | 2 | -2.969 | 1.928 | -1.007 | 1.928 | del | Batch2 |
| NW150212-III.11.L001 | chr22 | 22200000 | 22900000 | CC | 184.078 | 5 | 62 | 4.095 | 4.095 | 0.128 | 4.095 | del | Batch1 |
| NW150212-III.12.L001 | chr22 | 22200000 | 22900000 | WC | 179.497 | 107 | 0 | -7.206 | 13.338 | 5.671 | 13.338 | del | Batch1 |
| NW150212-III.13.L001 | chr22 | 22200000 | 22900000 | CW | 117.99 | 63 | 91 | -6.76 | -5.107 | -11.546 | -5.107 | ref_hom | Batch1 |
| NW150212-III.16.L001 | chr22 | 22200000 | 22900000 | WW | 66.567 | 16 | 0 | 2.226 | 2.226 | 1.143 | 2.226 | del | Batch1 |
| NW150212-III.21.L001 | chr22 | 22200000 | 22900000 | WW | 212.722 | 66 | 0 | 6.185 | 6.185 | 2.323 | 6.185 | del | Batch1 |
| NW150212-III.26.L001 | chr22 | 22200000 | 22900000 | CW | 180.478 | 0 | 51 | -2.07 | 13.657 | 10.877 | 13.657 | del | Batch1 |
| NW130711.302.L008 | chr22 | 22200000 | 22900000 | WW | 99.134 | 73 | 0 | -0.106 | -0.106 | -5.454 | -0.106 | unclear | Batch2 |
| NW150212-III.27.L001 | chr22 | 22200000 | 22900000 | CW | 284.3 | 183 | 204 | -14.319 | -13.141 | -26.9 | -13.141 | ref_hom | Batch1 |
| NW150212-III.30.L001 | chr22 | 22200000 | 22900000 | CW | 230.761 | 0 | 53 | -0.692 | 17.592 | 15.863 | 17.592 | del | Batch1 |
| NW150212-III.33.L001 | chr22 | 22200000 | 22900000 | CC | 152.679 | 2 | 46 | 4.031 | 4.031 | 1.234 | 4.031 | del | Batch1 |
| NW150212-III.35.L001 | chr22 | 22200000 | 22900000 | CW | 122.285 | 0 | 44 | -2.781 | 9.143 | 5.991 | 9.143 | del | Batch1 |
| NW150212-III.36.L001 | chr22 | 22200000 | 22900000 | CC | 217.604 | 0 | 45 | 8.757 | 8.757 | 7.858 | 8.757 | del | Batch1 |
| NW150212-III.39.L001 | chr22 | 22200000 | 22900000 | WW | 81.348 | 71 | 0 | -0.649 | -0.649 | -5.705 | -0.649 | ref_hom | Batch1 |
| NW150212-III.42.L001 | chr22 | 22200000 | 22900000 | WW | 166.419 | 220 | 0 | -3.629 | -3.629 | -15.92 | -3.629 | ref_hom | Batch1 |
| NW150212-III.48.L001 | chr22 | 22200000 | 22900000 | WC | 217.486 | 51 | 0 | -0.859 | 16.562 | 14.743 | 16.562 | del | Batch1 |
| NW150212-III.52.L001 | chr22 | 22200000 | 22900000 | CC | 182.975 | 2 | 51 | 5.374 | 5.374 | 2.653 | 5.374 | del | Batch1 |
| NW150212-III.55.L001 | chr22 | 22200000 | 22900000 | WC | 166.755 | 84 | 2 | -5.689 | 10.137 | 4.022 | 10.137 | del | Batch1 |
| NW150212-III.57.L001 | chr22 | 22200000 | 22900000 | CW | 147.047 | 0 | 84 | -5.811 | 10.905 | 4.738 | 10.905 | del | Batch1 |
| NW150212-III.59.L001 | chr22 | 22200000 | 22900000 | WW | 112.266 | 36 | 0 | 3.021 | 3.021 | 0.537 | 3.021 | del | Batch1 |
| NW150212-III.63.L001 | chr22 | 22200000 | 22900000 | CW | 118.66 | 3 | 60 | -4.349 | 5.832 | 1.266 | 5.832 | del | Batch1 |
| NW150212-III.66.L001 | chr22 | 22200000 | 22900000 | WW | 150.327 | 193 | 0 | -3.138 | -3.138 | -14.135 | -3.138 | ref_hom | Batch1 |
| NW150212-III.68.L001 | chr22 | 22200000 | 22900000 | WW | 157.239 | 90 | 0 | 1.409 | 1.409 | -5.15 | 1.409 | del | Batch1 |
| NW130711.307.L008 | chr22 | 22200000 | 22900000 | CW | 154.751 | 0 | 90 | -6.18 | 11.481 | 4.922 | 11.481 | del | Batch2 |
| NW150212-III.69.L001 | chr22 | 22200000 | 22900000 | CW | 162.454 | 0 | 55 | -3.039 | 12.224 | 8.616 | 12.224 | del | Batch1 |
| NW150212-III.75.L001 | chr22 | 22200000 | 22900000 | WC | 150.782 | 122 | 88 | -8.707 | -6.825 | -15.136 | -6.825 | ref_hom | Batch1 |
| NW150212-III.76.L001 | chr22 | 22200000 | 22900000 | WW | 189.101 | 284 | 9 | -6.02 | -6.02 | -20.844 | -6.02 | ref_hom | Batch1 |
| NW150212-III.77.L001 | chr22 | 22200000 | 22900000 | CW | 62.667 | 2 | 26 | -2.253 | 2.523 | 0.228 | 2.523 | del | Batch1 |
| NW150212-III.80.L001 | chr22 | 22200000 | 22900000 | CW | 162.94 | 77 | 142 | -9.764 | -5.974 | -15.367 | -5.974 | ref_hom | Batch1 |
| NW150212-III.81.L001 | chr22 | 22200000 | 22900000 | CC | 118.074 | 0 | 144 | -2.297 | -2.297 | -10.849 | -2.297 | ref_hom | Batch1 |
| NW150212-III.84.L001 | chr22 | 22200000 | 22900000 | WC | 225.328 | 153 | 4 | -10.082 | 12.671 | 2.104 | 12.671 | del | Batch1 |
| NW150212-III.90.L001 | chr22 | 22200000 | 22900000 | WC | 95.192 | 68 | 43 | -5.305 | -3.655 | -8.709 | -3.655 | ref_hom | Batch1 |
| NW150212-III.92.L001 | chr22 | 22200000 | 22900000 | CW | 157.936 | 0 | 69 | -4.516 | 11.806 | 6.814 | 11.806 | del | Batch1 |
| NW150212-III.93.L001 | chr22 | 22200000 | 22900000 | CC | 207.478 | 0 | 63 | 6.157 | 6.157 | 2.532 | 6.157 | del | Batch1 |
| NW150212-III.94.L001 | chr22 | 22200000 | 22900000 | WW | 180.109 | 98 | 9 | 0.996 | 0.996 | -6.074 | 0.996 | del | Batch1 |
| NW150212-IV.08.L002 | chr22 | 22200000 | 22900000 | CW | 106.651 | 64 | 47 | -3.847 | -5.082 | -8.712 | -3.847 | ref_hom | Batch1 |
| NW150212-IV.09.L002 | chr22 | 22200000 | 22900000 | CC | 108.171 | 2 | 28 | 3.134 | 3.134 | 1.534 | 3.134 | del | Batch1 |
| NW150212-IV.10.L002 | chr22 | 22200000 | 22900000 | CW | 122.02 | 0 | 20 | 0.672 | 9.277 | 9.426 | 9.426 | del | Batch1 |
| NW150212-IV.11.L002 | chr22 | 22200000 | 22900000 | CW | 107.546 | 2 | 28 | -1.296 | 5.883 | 4.266 | 5.883 | del | Batch1 |
| NW150212-IV.12.L002 | chr22 | 22200000 | 22900000 | WW | 112.149 | 30 | 0 | 3.608 | 3.608 | 1.845 | 3.608 | del | Batch1 |
| NW150212-IV.15.L002 | chr22 | 22200000 | 22900000 | CC | 186.18 | 0 | 41 | 7.152 | 7.152 | 5.901 | 7.152 | del | Batch1 |
| NW150212-IV.17.L002 | chr22 | 22200000 | 22900000 | WW | 129.825 | 39 | 0 | 3.788 | 3.788 | 1.324 | 3.788 | del | Batch1 |
| NW150212-IV.24.L002 | chr22 | 22200000 | 22900000 | CW | 113.144 | 0 | 31 | -1.485 | 8.499 | 6.631 | 8.499 | del | Batch1 |
| NW150212-IV.25.L002 | chr22 | 22200000 | 22900000 | WW | 141.35 | 51 | 0 | 3.371 | 3.371 | -0.179 | 3.371 | del | Batch1 |
| NW150212-IV.26.L002 | chr22 | 22200000 | 22900000 | WC | 185.951 | 119 | 108 | -8.773 | -8.09 | -16.508 | -8.09 | ref_hom | Batch1 |
| NW150212-IV.31.L002 | chr22 | 22200000 | 22900000 | WC | 81.467 | 20 | 2 | -1.03 | 3.957 | 2.731 | 3.957 | del | Batch1 |
| NW150212-IV.34.L002 | chr22 | 22200000 | 22900000 | CW | 172.23 | 0 | 51 | -2.31 | 13.011 | 10.045 | 13.011 | del | Batch1 |
| NW150212-IV.39.L002 | chr22 | 22200000 | 22900000 | CW | 169.71 | 0 | 98 | -6.645 | 12.611 | 5.529 | 12.611 | del | Batch1 |
| NW150212-IV.41.L002 | chr22 | 22200000 | 22900000 | CC | 148.725 | 0 | 29 | 6.121 | 6.121 | 5.598 | 6.121 | del | Batch1 |
| NW130711.291.L008 | chr22 | 22694696 | 22899260 | WC | 44.881 | 20 | 0 | -1.983 | 3.221 | 1.236 | 3.221 | del | Batch2 |
| NW130711.320.L008 | chr22 | 22694696 | 22899260 | WC | 16.444 | 15 | 6 | -1.959 | -1.365 | -3.115 | -1.365 | ref_hom | Batch2 |

| | | | | | | | | | | | | | |
|---|---|---|---|---|---|---|---|---|---|---|---|---|---|
| NW150212-IV.43.L002 | chr22 | 22694696 | 22899260 | WW | 31.370 | 1 | 0 | 2.023 | 2.023 | 2.995 | 2.995 | del_hom | Batch1 |
| NW150212-IV.44.L002 | chr22 | 22694696 | 22899260 | WW | 34.990 | 1 | 2 | 1.948 | 1.948 | 3.180 | 3.180 | del_hom | Batch1 |
| NW150212-IV.45.L002 | chr22 | 22694696 | 22899260 | CW | 22.022 | 0 | 0 | 1.579 | 1.579 | 3.159 | 3.159 | del_hom | Batch1 |
| NW150212-IV.46.L002 | chr22 | 22694696 | 22899260 | WC | 41.447 | 1 | 0 | 1.827 | 3.105 | 4.800 | 4.800 | del_hom | Batch1 |
| NW150212-IV.50.L002 | chr22 | 22694696 | 22899260 | CW | 43.006 | 0 | 0 | 3.252 | 3.252 | 6.337 | 6.337 | del_hom | Batch1 |
| NW150212-IV.58.L002 | chr22 | 22694696 | 22899260 | CW | 49.326 | 0 | 0 | 3.756 | 3.756 | 7.294 | 7.294 | del_hom | Batch1 |
| NW150212-IV.60.L002 | chr22 | 22694696 | 22899260 | WW | 50.502 | 22 | 1 | 0.413 | 0.413 | -1.677 | 0.413 | unclear | Batch1 |
| NW150212-IV.62.L002 | chr22 | 22694696 | 22899260 | CW | 72.030 | 0 | 0 | 5.565 | 5.565 | 10.733 | 10.733 | del_hom | Batch1 |
| NW150212-IV.63.L002 | chr22 | 22694696 | 22899260 | CC | 27.227 | 1 | 15 | -0.272 | -0.272 | -2.026 | -0.272 | ref_hom | Batch1 |
| NW150212-IV.64.L002 | chr22 | 22694696 | 22899260 | WW | 52.180 | 51 | 0 | -0.749 | -0.749 | -4.502 | -0.749 | ref_hom | Batch1 |
| NW150212-IV.65.L002 | chr22 | 22694696 | 22899260 | WC | 56.320 | 0 | 0 | 4.313 | 4.313 | 8.353 | 8.353 | del_hom | Batch1 |
| NW150212-IV.68.L002 | chr22 | 22694696 | 22899260 | CW | 56.546 | 0 | 3 | 1.765 | 4.273 | 5.822 | 5.822 | del_hom | Batch1 |
| NW150212-IV.69.L002 | chr22 | 22694696 | 22899260 | WC | 41.677 | 0 | 0 | 3.146 | 3.146 | 6.136 | 6.136 | del_hom | Batch1 |
| NW150212-IV.72.L002 | chr22 | 22694696 | 22899260 | WC | 38.786 | 23 | 0 | -2.292 | 2.736 | 0.490 | 2.736 | del | Batch1 |
| NW150212-IV.74.L002 | chr22 | 22694696 | 22899260 | WC | 51.156 | 0 | 1 | 3.879 | 2.601 | 6.270 | 6.270 | del_hom | Batch1 |
| NW150212-IV.77.L002 | chr22 | 22694696 | 22899260 | CW | 59.646 | 0 | 17 | -1.295 | 4.389 | 2.984 | 4.389 | del | Batch1 |
| NW150212-IV.81.L002 | chr22 | 22694696 | 22899260 | WW | 51.207 | 1 | 0 | 3.605 | 3.605 | 5.999 | 5.999 | del_hom | Batch1 |
| NW150212-IV.83.L002 | chr22 | 22694696 | 22899260 | WW | 65.203 | 0 | 1 | 4.720 | 4.720 | 9.398 | 9.398 | del_hom | Batch1 |
| NW150212-IV.84.L002 | chr22 | 22694696 | 22899260 | WW | 60.560 | 1 | 0 | 4.350 | 4.350 | 7.416 | 7.416 | del_hom | Batch1 |
| NW150212-IV.85.L002 | chr22 | 22694696 | 22899260 | WW | 52.618 | 1 | 0 | 3.717 | 3.717 | 6.213 | 6.213 | del_hom | Batch1 |
| NW150212-IV.87.L002 | chr22 | 22694696 | 22899260 | CC | 58.656 | 0 | 0 | 4.499 | 4.499 | 8.707 | 8.707 | del_hom | Batch1 |
| NW150212-IV.89.L002 | chr22 | 22694696 | 22899260 | CC | 55.953 | 0 | 2 | 3.716 | 3.716 | 5.746 | 5.746 | del_hom | Batch1 |
| NW150212-IV.91.L002 | chr22 | 22694696 | 22899260 | WC | 39.529 | 0 | 0 | 2.975 | 2.975 | 5.810 | 5.810 | del_hom | Batch1 |
| NW130711.323.L008 | chr22 | 22694696 | 22899260 | CC | 21.980 | 0 | 13 | -0.092 | -0.092 | -1.740 | -0.092 | ref_hom | Batch2 |
| NW150212-IV.93.L002 | chr22 | 22694696 | 22899260 | WW | 65.439 | 72 | 0 | -1.154 | -1.154 | -5.964 | -1.154 | ref_hom | Batch1 |
| NW130711.324.L008 | chr22 | 22694696 | 22899260 | CC | 21.663 | 0 | 12 | -0.044 | -0.044 | -1.627 | -0.044 | ref_hom | Batch2 |
| NW130711.325.L008 | chr22 | 22694696 | 22899260 | CC | 21.553 | 0 | 12 | -0.050 | -0.050 | -1.633 | -0.050 | ref_hom | Batch2 |
| NW130711.327.L008 | chr22 | 22694696 | 22899260 | CC | 30.760 | 0 | 18 | 0.008 | 0.008 | -1.933 | 0.008 | ref_hom | Batch2 |
| NW130711.329.L008 | chr22 | 22694696 | 22899260 | CC | 32.015 | 1 | 16 | -0.105 | -0.105 | -1.896 | -0.105 | ref_hom | Batch2 |
| NW130711.345.L008 | chr22 | 22694696 | 22899260 | WW | 28.855 | 15 | 1 | -0.194 | -0.194 | -1.936 | -0.194 | ref_hom | Batch2 |
| NW130711.350.L008 | chr22 | 22694696 | 22899260 | CW | 34.999 | 0 | 19 | -2.044 | 2.457 | 0.467 | 2.457 | del | Batch2 |
| NW130711.354.L008 | chr22 | 22694696 | 22899260 | WW | 22.308 | 14 | 0 | -0.136 | -0.136 | -1.847 | -0.136 | ref_hom | Batch2 |
| NW130711.355.L008 | chr22 | 22694696 | 22899260 | CW | 18.905 | 0 | 14 | -1.841 | 1.229 | -0.484 | 1.229 | del | Batch2 |
| NW130711.362.L008 | chr22 | 22694696 | 22899260 | CW | 33.805 | 0 | 23 | -2.319 | 2.352 | 0.106 | 2.352 | del | Batch2 |
| NW130711.365.L008 | chr22 | 22694696 | 22899260 | WC | 25.390 | 16 | 0 | -1.925 | 1.724 | -0.104 | 1.724 | del | Batch2 |
| NW130711.296.L008 | chr22 | 22694696 | 22899260 | WW | 39.005 | 23 | 0 | 0.082 | 0.082 | -2.164 | 0.082 | unclear | Batch2 |
| NW130711.374.L008 | chr22 | 22694696 | 22899260 | WW | 26.847 | 14 | 0 | 0.076 | 0.076 | -1.609 | 0.076 | unclear | Batch2 |
| NW130711.376.L008 | chr22 | 22694696 | 22899260 | WC | 26.671 | 12 | 0 | -1.604 | 1.838 | 0.309 | 1.838 | del | Batch2 |
| NW130711.377.L008 | chr22 | 22694696 | 22899260 | CW | 14.626 | 0 | 10 | -1.634 | 0.911 | -0.584 | 0.911 | del | Batch2 |
| NW150212-III.03.L001 | chr22 | 22694696 | 22899260 | WC | 52.913 | 0 | 0 | 4.042 | 4.042 | 7.837 | 7.837 | del_hom | Batch1 |
| NW150212-III.05.L001 | chr22 | 22694696 | 22899260 | WC | 61.462 | 40 | 1 | -3.307 | 3.133 | -0.177 | 3.133 | del | Batch1 |
| NW150212-III.06.L001 | chr22 | 22694696 | 22899260 | WC | 62.880 | 34 | 0 | -2.872 | 4.563 | 1.639 | 4.563 | del | Batch1 |
| NW150212-III.07.L001 | chr22 | 22694696 | 22899260 | CW | 48.206 | 21 | 38 | -3.354 | -2.262 | -5.386 | -2.262 | ref_hom | Batch1 |
| NW150212-III.09.L001 | chr22 | 22694696 | 22899260 | CC | 59.511 | 0 | 25 | 0.965 | 0.965 | -1.280 | 0.965 | del | Batch1 |
| NW130711.300.L008 | chr22 | 22694696 | 22899260 | WC | 16.054 | 12 | 0 | -1.740 | 1.015 | -0.589 | 1.015 | del | Batch2 |
| NW150212-III.11.L001 | chr22 | 22694696 | 22899260 | CC | 53.794 | 1 | 0 | 3.811 | 3.811 | 7.670 | 7.670 | del_hom | Batch1 |
| NW150212-III.12.L001 | chr22 | 22694696 | 22899260 | WC | 52.455 | 25 | 0 | -2.292 | 3.789 | 1.471 | 3.789 | del | Batch1 |
| NW150212-III.13.L001 | chr22 | 22694696 | 22899260 | CW | 34.481 | 21 | 22 | -2.416 | -2.354 | -4.542 | -2.354 | ref_hom | Batch1 |
| NW150212-III.16.L001 | chr22 | 22694696 | 22899260 | WW | 19.453 | 1 | 0 | 1.073 | 1.073 | 1.190 | 1.190 | del_hom | Batch1 |
| NW150212-III.21.L001 | chr22 | 22694696 | 22899260 | WW | 62.165 | 0 | 0 | 4.779 | 4.779 | 9.239 | 9.239 | del_hom | Batch1 |
| NW150212-III.26.L001 | chr22 | 22694696 | 22899260 | CW | 52.742 | 0 | 1 | 2.727 | 4.006 | 6.511 | 6.511 | del_hom | Batch1 |

| | | | | | | | | | | | | | |
|---|---|---|---|---|---|---|---|---|---|---|---|---|---|
| NW130711.302.L008 | chr22 | 22694696 | 22899260 | WW | 28.970 | 21 | 0 | -0.234 | -0.234 | -2.353 | -0.234 | ref_hom | Batch2 |
| NW150212-III.27.L001 | chr22 | 22694696 | 22899260 | CW | 83.082 | 48 | 65 | -5.095 | -4.130 | -8.925 | -4.130 | ref_hom | Batch1 |
| NW150212-III.30.L001 | chr22 | 22694696 | 22899260 | CW | 67.436 | 0 | 0 | 5.199 | 5.199 | 10.037 | 10.037 | del_hom | Batch1 |
| NW150212-III.33.L001 | chr22 | 22694696 | 22899260 | CC | 44.618 | 0 | 2 | 2.820 | 2.820 | 4.098 | 4.098 | del_hom | Batch1 |
| NW150212-III.35.L001 | chr22 | 22694696 | 22899260 | CW | 35.736 | 0 | 0 | 2.672 | 2.672 | 5.236 | 5.236 | del_hom | Batch1 |
| NW150212-III.36.L001 | chr22 | 22694696 | 22899260 | CC | 63.591 | 0 | 0 | 4.893 | 4.893 | 9.455 | 9.455 | del_hom | Batch1 |
| NW150212-III.39.L001 | chr22 | 22694696 | 22899260 | WW | 23.773 | 25 | 0 | -0.581 | -0.581 | -2.829 | -0.581 | ref_hom | Batch1 |
| NW150212-III.42.L001 | chr22 | 22694696 | 22899260 | WW | 48.633 | 55 | 0 | -0.990 | -0.990 | -4.822 | -0.990 | ref_hom | Batch1 |
| NW150212-III.48.L001 | chr22 | 22694696 | 22899260 | WC | 63.557 | 1 | 0 | 3.589 | 4.868 | 8.148 | 8.148 | del_hom | Batch1 |
| NW150212-III.52.L001 | chr22 | 22694696 | 22899260 | CC | 53.472 | 1 | 1 | 3.484 | 3.484 | 6.041 | 6.041 | del_hom | Batch1 |
| NW150212-III.55.L001 | chr22 | 22694696 | 22899260 | WC | 48.731 | 18 | 1 | -1.735 | 2.229 | 0.482 | 2.229 | del | Batch1 |
| NW150212-III.57.L001 | chr22 | 22694696 | 22899260 | CW | 42.972 | 0 | 22 | -2.178 | 3.064 | 0.905 | 3.064 | del | Batch1 |
| NW150212-III.59.L001 | chr22 | 22694696 | 22899260 | WW | 32.808 | 0 | 0 | 2.439 | 2.439 | 4.793 | 4.793 | del_hom | Batch1 |
| NW150212-III.63.L001 | chr22 | 22694696 | 22899260 | CW | 34.677 | 0 | 17 | -1.892 | 2.440 | 0.595 | 2.440 | del | Batch1 |
| NW150212-III.66.L001 | chr22 | 22694696 | 22899260 | WW | 43.931 | 45 | 0 | -0.759 | -0.759 | -4.128 | -0.759 | ref_hom | Batch1 |
| NW150212-III.68.L001 | chr22 | 22694696 | 22899260 | WW | 45.951 | 22 | 0 | 0.479 | 0.479 | -1.658 | 0.479 | unclear | Batch1 |
| NW130711.307.L008 | chr22 | 22694696 | 22899260 | CW | 45.223 | 0 | 26 | -2.450 | 3.224 | 0.794 | 3.224 | del | Batch2 |
| NW150212-III.69.L001 | chr22 | 22694696 | 22899260 | CW | 47.475 | 0 | 1 | 2.307 | 3.586 | 5.713 | 5.713 | del_hom | Batch1 |
| NW150212-III.75.L001 | chr22 | 22694696 | 22899260 | WC | 44.064 | 33 | 23 | -3.076 | -2.464 | -5.302 | -2.464 | ref_hom | Batch1 |
| NW150212-III.76.L001 | chr22 | 22694696 | 22899260 | WW | 55.262 | 71 | 0 | -1.355 | -1.355 | -5.877 | -1.355 | ref_hom | Batch1 |
| NW150212-III.77.L001 | chr22 | 22694696 | 22899260 | CW | 18.313 | 0 | 6 | -1.207 | 1.214 | 0.107 | 1.214 | del | Batch1 |
| NW150212-III.80.L001 | chr22 | 22694696 | 22899260 | CW | 47.617 | 17 | 40 | -3.432 | -1.884 | -5.097 | -1.884 | ref_hom | Batch1 |
| NW150212-III.81.L001 | chr22 | 22694696 | 22899260 | CC | 34.505 | 0 | 45 | -0.986 | -0.986 | -4.143 | -0.986 | ref_hom | Batch1 |
| NW150212-III.84.L001 | chr22 | 22694696 | 22899260 | WC | 65.849 | 38 | 0 | -3.133 | 4.779 | 1.590 | 4.779 | del | Batch1 |
| NW150212-III.90.L001 | chr22 | 22694696 | 22899260 | WC | 27.818 | 10 | 11 | -1.490 | -1.590 | -2.911 | -1.490 | ref_hom | Batch1 |
| NW150212-III.92.L001 | chr22 | 22694696 | 22899260 | CW | 46.154 | 0 | 0 | 3.503 | 3.503 | 6.814 | 6.814 | del_hom | Batch1 |
| NW150212-III.93.L001 | chr22 | 22694696 | 22899260 | CC | 60.632 | 0 | 1 | 4.356 | 4.356 | 7.427 | 7.427 | del_hom | Batch1 |
| NW150212-III.94.L001 | chr22 | 22694696 | 22899260 | WW | 52.634 | 26 | 5 | 0.039 | 0.039 | -2.355 | 0.039 | unclear | Batch1 |
| NW150212-IV.08.L002 | chr22 | 22694696 | 22899260 | CW | 31.167 | 18 | 19 | -2.224 | -2.159 | -4.163 | -2.159 | ref_hom | Batch1 |
| NW150212-IV.09.L002 | chr22 | 22694696 | 22899260 | CC | 31.611 | 1 | 0 | 2.043 | 2.043 | 4.310 | 4.310 | del_hom | Batch1 |
| NW150212-IV.10.L002 | chr22 | 22694696 | 22899260 | CW | 35.658 | 0 | 0 | 2.666 | 2.666 | 5.224 | 5.224 | del_hom | Batch1 |
| NW150212-IV.11.L002 | chr22 | 22694696 | 22899260 | CW | 31.429 | 0 | 2 | 0.462 | 2.290 | 2.716 | 2.716 | del_hom | Batch1 |
| NW150212-IV.12.L002 | chr22 | 22694696 | 22899260 | WW | 32.774 | 0 | 0 | 2.436 | 2.436 | 4.787 | 4.787 | del_hom | Batch1 |
| NW150212-IV.15.L002 | chr22 | 22694696 | 22899260 | CC | 54.408 | 0 | 0 | 4.161 | 4.161 | 8.064 | 8.064 | del_hom | Batch1 |
| NW150212-IV.17.L002 | chr22 | 22694696 | 22899260 | WW | 37.939 | 0 | 0 | 2.848 | 2.848 | 5.570 | 5.570 | del_hom | Batch1 |
| NW150212-IV.24.L002 | chr22 | 22694696 | 22899260 | CW | 33.065 | 0 | 1 | 1.158 | 2.437 | 3.530 | 3.530 | del_hom | Batch1 |
| NW150212-IV.25.L002 | chr22 | 22694696 | 22899260 | WW | 41.307 | 1 | 0 | 2.815 | 2.815 | 4.500 | 4.500 | del_hom | Batch1 |
| NW150212-IV.26.L002 | chr22 | 22694696 | 22899260 | WC | 54.341 | 36 | 27 | -3.284 | -2.688 | -5.742 | -2.688 | ref_hom | Batch1 |
| NW150212-IV.31.L002 | chr22 | 22694696 | 22899260 | WC | 23.807 | 0 | 0 | 1.722 | 1.722 | 3.430 | 3.430 | del_hom | Batch1 |
| NW150212-IV.34.L002 | chr22 | 22694696 | 22899260 | CW | 50.331 | 0 | 0 | 3.836 | 3.836 | 7.446 | 7.446 | del_hom | Batch1 |
| NW150212-IV.39.L002 | chr22 | 22694696 | 22899260 | CW | 49.595 | 0 | 24 | -2.250 | 3.571 | 1.308 | 3.571 | del | Batch1 |
| NW150212-IV.41.L002 | chr22 | 22694696 | 22899260 | CC | 43.462 | 0 | 1 | 2.987 | 2.987 | 4.826 | 4.826 | del_hom | Batch1 |
