## Supplementary Table S9 for "Haplotype-aware single-cell multiomics uncovers functional effects of somatic structural variation"

**Table S9. List of differentially expressed genes identified for the eight unsupervised clusters of T-**

| gene | avg_log2FC | p_val_adj | cluster |
|---|---|---|---|
| KLF2 | 1.8780238 | 0 | 0 |
| NOSIP | 1.79113642 | 0 | 0 |
| AHNAK | 1.77032796 | 0 | 0 |
| TECR | 1.64827804 | 0 | 0 |
| PLAC8 | 1.55070948 | 0 | 0 |
| EMP3 | 1.42718218 | 0 | 0 |
| ADD3 | 1.35544249 | 0 | 0 |
| TMSB10 | 1.23979655 | 0 | 0 |
| CLIC3 | 1.18421309 | 0 | 0 |
| FUT7 | 0.80402825 | 0 | 0 |
| GIMAP4 | 0.81292725 | 2.25E-284 | 0 |
| CD52 | 0.95075707 | 6.31E-278 | 0 |
| GIMAP7 | 1.01139547 | 2.62E-263 | 0 |
| KLRK1 | 0.94694303 | 4.87E-261 | 0 |
| CAPN2 | 0.76331376 | 4.49E-260 | 0 |
| RIPOR2 | 1.10018762 | 1.87E-254 | 0 |
| S100A4 | 1.47870303 | 1.48E-251 | 0 |
| LITAF | 0.8530669 | 1.02E-249 | 0 |
| S100A6 | 1.2649281 | 8.25E-235 | 0 |
| SAMHD1 | 0.96599059 | 6.91E-231 | 0 |
| TSC22D3 | 0.98547902 | 1.55E-223 | 0 |
| CNN2 | 0.97647324 | 2.34E-217 | 0 |
| CRIP1 | 1.73608293 | 9.39E-216 | 0 |
| MYOM2 | 0.92219648 | 4.50E-210 | 0 |
| CTSW | 0.96926139 | 8.83E-204 | 0 |
| S100A10 | 1.2689284 | 8.64E-200 | 0 |
| FLNA | 0.90374779 | 4.00E-197 | 0 |
| NKG7 | 0.84316951 | 3.40E-195 | 0 |
| VIM | 0.94167288 | 3.78E-195 | 0 |
| S100A11 | 0.99524813 | 1.71E-190 | 0 |
| IL10RA | 0.51209635 | 1.60E-187 | 0 |
| ZNRF1 | -0.9225729 | 1.28E-180 | 0 |
| CALM1 | 0.80456868 | 2.91E-178 | 0 |
| COL6A2 | 0.88496673 | 1.57E-176 | 0 |
| MYO1F | 0.77859881 | 1.40E-175 | 0 |
| ISG20 | 0.77456696 | 5.77E-172 | 0 |
| FCER1G | 0.7467453 | 3.35E-169 | 0 |
| LGALS1 | 0.90390935 | 4.08E-157 | 0 |
| CD1E | -1.3164143 | 9.69E-156 | 0 |

| | | | |
|---|---|---|---|
| SAMD3 | 0.75578562 | 1.24E-154 | 0 |
| CD1A | -1.3023236 | 5.45E-148 | 0 |
| TSPAN2 | 0.59329368 | 5.19E-146 | 0 |
| TPST2 | 0.69785685 | 7.13E-145 | 0 |
| ANXA2 | 0.67806661 | 2.92E-142 | 0 |
| TXK | 0.53077468 | 5.08E-142 | 0 |
| MSN | 0.6998764 | 6.45E-141 | 0 |
| SH3BGRL3 | 0.63757457 | 1.39E-139 | 0 |
| CD1B | -1.0428747 | 5.73E-139 | 0 |
| MBP | 0.73586771 | 2.30E-131 | 0 |
| CDC42 | 0.58245842 | 3.66E-129 | 0 |
| CD2 | -0.7594091 | 8.61E-129 | 0 |
| BIN2 | 0.65573515 | 1.03E-127 | 0 |
| S1PR1 | 0.48957547 | 1.98E-127 | 0 |
| RASA3 | 0.41489191 | 6.18E-127 | 0 |
| TAGLN2 | 0.88952296 | 8.96E-127 | 0 |
| CD38 | -0.9493893 | 7.87E-126 | 0 |
| CYTH1 | 0.6197007 | 1.29E-124 | 0 |
| ANXA1 | 1.00124658 | 2.58E-123 | 0 |
| HCST | 0.64173106 | 3.10E-123 | 0 |
| SELL | 0.70879985 | 1.92E-122 | 0 |
| EZR | 0.79669193 | 1.27E-121 | 0 |
| CDC25B | 0.58492456 | 1.50E-121 | 0 |
| SATB1 | -1.2190072 | 1.16E-119 | 0 |
| DOK2 | 0.61108219 | 1.73E-119 | 0 |
| STK38 | 0.53364178 | 2.69E-119 | 0 |
| SULT1B1 | 0.29526946 | 1.35E-117 | 0 |
| SYTL1 | 0.60365232 | 2.11E-115 | 0 |
| FXYD5 | 0.65201162 | 1.10E-114 | 0 |
| PTPN6 | 0.67673719 | 2.87E-113 | 0 |
| RASGRP2 | 0.61545775 | 5.98E-110 | 0 |
| RAB29 | 0.47145172 | 1.11E-109 | 0 |
| VCL | 0.32935034 | 5.23E-108 | 0 |
| TCEA3 | 0.42697041 | 1.25E-106 | 0 |
| PI16 | 0.2680081 | 1.38E-106 | 0 |
| NUCB2 | -0.6619755 | 2.71E-106 | 0 |
| TMBIM1 | 0.29579448 | 7.66E-105 | 0 |
| CDC42SE1 | 0.58677959 | 2.26E-104 | 0 |
| MYL12A | 0.50873328 | 3.23E-104 | 0 |
| LSP1 | 0.61881718 | 5.39E-104 | 0 |
| SBK1 | 0.51405537 | 5.60E-104 | 0 |
| GIMAP1 | 0.49495926 | 8.68E-104 | 0 |
| WDR86-AS1 | 0.27267996 | 1.30E-101 | 0 |

| | | | |
|---|---|---|---|
| CD55 | 0.35791641 | 2.55E-100 | 0 |
| IGFBP5 | -1.1086071 | 6.14E-100 | 0 |
| KLRB1 | 0.66455211 | 7.33E-99 | 0 |
| TSPO | 0.58687459 | 7.73E-99 | 0 |
| C19orf33 | 0.38488688 | 4.93E-97 | 0 |
| SEPTIN6 | -0.5159746 | 3.17E-96 | 0 |
| S1PR4 | 0.46481289 | 2.48E-93 | 0 |
| TSPAN32 | 0.4356208 | 3.39E-92 | 0 |
| KLRC3 | 0.54475565 | 5.10E-92 | 0 |
| WAKMAR2 | -0.9427942 | 7.13E-92 | 0 |
| KLF6 | 0.80754726 | 2.29E-91 | 0 |
| APOBEC3G | 0.49538882 | 5.06E-90 | 0 |
| RASSF1 | 0.51587049 | 9.40E-90 | 0 |
| ITGA6 | 0.55221899 | 3.88E-88 | 0 |
| ITGB1 | 0.70396039 | 6.38E-88 | 0 |
| RAP1B | 0.5274337 | 5.91E-86 | 0 |
| TES | 0.55995265 | 1.14E-85 | 0 |
| SMIM24 | -0.7157827 | 1.95E-85 | 0 |
| ATP2B1 | 0.6555417 | 3.13E-85 | 0 |
| OPTN | 0.50438045 | 3.49E-85 | 0 |
| SLCO3A1 | 0.34462059 | 2.01E-84 | 0 |
| ARPC1B | 0.45083522 | 1.36E-81 | 0 |
| ESYT2 | 0.66711385 | 2.93E-81 | 0 |
| SMPD3 | -0.8139251 | 3.95E-81 | 0 |
| GLIPR2 | 0.53654475 | 7.12E-81 | 0 |
| ITGB7 | 0.3128065 | 2.03E-79 | 0 |
| KRAS | 0.54924872 | 1.86E-78 | 0 |
| MZB1 | -0.8058579 | 2.53E-78 | 0 |
| CST7 | 0.44659487 | 3.14E-78 | 0 |
| TGFB1 | 0.49801044 | 7.36E-78 | 0 |
| TRIB2 | 0.36827026 | 7.80E-78 | 0 |
| TLE5 | 0.42670992 | 2.36E-77 | 0 |
| CAST | 0.59182084 | 5.46E-77 | 0 |
| NME2 | 0.49184065 | 1.69E-76 | 0 |
| LZTFL1 | -0.6780416 | 9.70E-76 | 0 |
| CLIC1 | 0.46758149 | 2.16E-75 | 0 |
| NEAT1 | 0.66827029 | 2.69E-75 | 0 |
| LINC00861 | 0.38625947 | 5.70E-75 | 0 |
| CYTIP | 0.39839143 | 1.14E-74 | 0 |
| ICAM2 | 0.46113926 | 2.68E-74 | 0 |
| GRK6 | 0.47806831 | 6.56E-73 | 0 |
| PLEC | 0.46483472 | 1.27E-72 | 0 |
| ID3 | 0.62709056 | 3.53E-72 | 0 |

| | | | |
|---|---|---|---|
| CD300A | 0.39180791 | 4.22E-72 | 0 |
| IQGAP1 | 0.48931034 | 5.05E-72 | 0 |
| NCAPG2 | 0.65620178 | 2.16E-71 | 0 |
| MAP3K1 | 0.45368792 | 2.69E-70 | 0 |
| RNF125 | 0.49035031 | 5.40E-70 | 0 |
| IL32 | 0.40508523 | 6.68E-70 | 0 |
| GNG2 | 0.45588165 | 2.85E-69 | 0 |
| CR1 | 0.53548278 | 2.25E-68 | 0 |
| CYBA | 0.44477556 | 5.34E-68 | 0 |
| PRMT2 | 0.49505509 | 7.10E-68 | 0 |
| SVIL | 0.3756635 | 1.42E-67 | 0 |
| TPM3 | 0.43419887 | 2.30E-67 | 0 |
| UBASH3B | 0.48779977 | 3.64E-67 | 0 |
| EOMES | 0.30706043 | 1.69E-65 | 0 |
| HMGB1 | -0.6298878 | 2.51E-65 | 0 |
| GPR183 | 0.25299491 | 4.81E-65 | 0 |
| HLA-B | 0.63318641 | 6.60E-65 | 0 |
| LY6E | 0.45231835 | 1.37E-63 | 0 |
| ESYT1 | 0.42856898 | 1.84E-63 | 0 |
| LRRFIP1 | 0.45423898 | 3.79E-63 | 0 |
| HMOX2 | 0.38006509 | 1.54E-62 | 0 |
| ATP8B2 | 0.41641735 | 1.58E-62 | 0 |
| TMEM63A | 0.31664897 | 2.29E-62 | 0 |
| TFDP2 | -0.8215646 | 1.13E-61 | 0 |
| CDC14A | 0.3778364 | 9.86E-61 | 0 |
| MACROD2 | 0.62748363 | 1.05E-60 | 0 |
| B2M | 0.37756271 | 1.63E-60 | 0 |
| BIN1 | 0.4025114 | 1.93E-60 | 0 |
| HMGN2 | -0.6980098 | 1.42E-59 | 0 |
| CCND3 | 0.4025092 | 1.49E-59 | 0 |
| HLA-E | 0.42895804 | 2.27E-59 | 0 |
| SIVA1 | -0.680962 | 6.98E-59 | 0 |
| ABHD17A | 0.40618626 | 1.32E-57 | 0 |
| CASC15 | -0.5558963 | 4.03E-57 | 0 |
| CORO1A | 0.37799954 | 8.57E-57 | 0 |
| HLA-C | 0.47200254 | 1.34E-56 | 0 |
| SH2D1A | -0.5051747 | 1.90E-56 | 0 |
| CTSA | 0.39551184 | 2.54E-56 | 0 |
| GSTK1 | 0.43028736 | 2.36E-55 | 0 |
| SELPLG | 0.36410598 | 3.46E-55 | 0 |
| PAXX | 0.443156 | 3.83E-55 | 0 |
| C1orf56 | 0.41766307 | 3.91E-54 | 0 |
| CD9 | 0.39092759 | 4.79E-54 | 0 |

| | | | |
|---|---|---|---|
| GLG1 | 0.38565992 | 6.21E-54 | 0 |
| RTKN2 | 0.46154673 | 9.68E-54 | 0 |
| PPP2R5A | 0.38484435 | 1.66E-53 | 0 |
| CCR9 | -0.6169124 | 3.63E-53 | 0 |
| AL138899.1 | -0.8005529 | 1.11E-52 | 0 |
| DGKD | 0.41149452 | 1.19E-52 | 0 |
| PDE7A | -0.5812719 | 1.43E-52 | 0 |
| SOX4 | -0.5784629 | 3.38E-52 | 0 |
| MYH9 | 0.43392704 | 8.41E-52 | 0 |
| KLRG1 | 0.42180359 | 9.51E-52 | 0 |
| RPL6 | 0.30718038 | 1.46E-51 | 0 |
| INF2 | 0.295396 | 1.49E-51 | 0 |
| GIHCG | -0.7165443 | 2.33E-51 | 0 |
| UCP2 | 0.49909316 | 4.94E-51 | 0 |
| STK10 | 0.38223088 | 6.25E-51 | 0 |
| CFLAR | 0.45411241 | 1.40E-50 | 0 |
| S1PR3 | -0.4594292 | 1.44E-50 | 0 |
| AL365440.2 | -0.5595618 | 1.76E-50 | 0 |
| TXNIP | 0.44741804 | 7.17E-50 | 0 |
| TMEM173 | 0.34773102 | 1.01E-49 | 0 |
| PPP3CC | 0.25034348 | 2.72E-49 | 0 |
| VIPR2 | -0.502813 | 3.31E-49 | 0 |
| CAP1 | 0.38752837 | 4.99E-49 | 0 |
| MYO1G | 0.39080398 | 5.28E-49 | 0 |
| TOX2 | -0.5915016 | 9.18E-49 | 0 |
| SNHG18 | -0.4122927 | 1.32E-48 | 0 |
| ECE1 | 0.25161364 | 1.70E-48 | 0 |
| PTMA | -0.3899975 | 2.04E-48 | 0 |
| CAMK1D | -0.492834 | 2.35E-48 | 0 |
| ITGB2 | 0.40944623 | 3.59E-48 | 0 |
| RASSF3 | 0.28713865 | 5.04E-48 | 0 |
| ANXA6 | 0.40930118 | 8.67E-48 | 0 |
| POU2F2 | 0.35109036 | 8.28E-47 | 0 |
| RPS14 | 0.2832344 | 1.56E-46 | 0 |
| RNPEPL1 | 0.32808988 | 1.67E-46 | 0 |
| AXIN1 | 0.32309271 | 1.84E-46 | 0 |
| TIMP1 | 0.36615689 | 1.87E-46 | 0 |
| RGS19 | 0.34374373 | 2.47E-45 | 0 |
| ADGRE5 | 0.38521386 | 3.83E-45 | 0 |
| PLSCR3 | 0.25118867 | 8.02E-45 | 0 |
| PRKCB | 0.42009621 | 9.35E-45 | 0 |
| CA5B | 0.26754719 | 1.15E-44 | 0 |
| MYADM | 0.41779948 | 1.29E-44 | 0 |

| | | | |
|---|---|---|---|
| CAPNS1 | 0.36534136 | 1.84E-44 | 0 |
| SPTBN1 | 0.39508034 | 7.89E-44 | 0 |
| CD53 | 0.38636935 | 8.35E-44 | 0 |
| NCR3 | 0.3985589 | 1.10E-43 | 0 |
| SLA | -0.5491272 | 5.83E-43 | 0 |
| RTN4 | 0.36978172 | 8.91E-43 | 0 |
| TRBC2 | -0.3390276 | 8.94E-43 | 0 |
| IL17RB | -0.8216068 | 1.01E-42 | 0 |
| CLEC2D | 0.41385785 | 1.12E-42 | 0 |
| CXCR4 | -0.5228188 | 7.53E-42 | 0 |
| LINC-PINT | 0.35753695 | 1.36E-41 | 0 |
| RPS7 | 0.27350807 | 1.37E-41 | 0 |
| FOXO1 | 0.26511512 | 1.44E-41 | 0 |
| SP100 | 0.40071669 | 3.15E-41 | 0 |
| NPNT | -0.4311517 | 3.62E-41 | 0 |
| RAP2B | 0.31636981 | 4.70E-41 | 0 |
| ITM2C | -0.5049677 | 5.54E-41 | 0 |
| NSMCE1 | 0.33805602 | 1.56E-40 | 0 |
| GIMAP6 | 0.29744074 | 3.36E-40 | 0 |
| ARL4C | 0.33581063 | 7.88E-40 | 0 |
| IL27RA | 0.36220534 | 9.72E-40 | 0 |
| PRKCQ-AS1 | 0.35556286 | 1.76E-38 | 0 |
| HELB | 0.29955444 | 2.68E-38 | 0 |
| MYL12B | 0.29405198 | 2.73E-38 | 0 |
| ZNF683 | 0.35495161 | 1.78E-37 | 0 |
| YWHAZ | 0.28224317 | 2.37E-37 | 0 |
| ARHGAP45 | 0.32499618 | 3.89E-37 | 0 |
| KDM7A | 0.28157061 | 5.76E-37 | 0 |
| SRGN | 0.30317744 | 6.02E-37 | 0 |
| RAC2 | 0.27616092 | 8.83E-37 | 0 |
| CASP10 | 0.30348275 | 1.04E-36 | 0 |
| CD48 | 0.35472315 | 1.49E-36 | 0 |
| YWHAH | 0.38138565 | 1.97E-36 | 0 |
| ARPC3 | 0.26976945 | 4.30E-36 | 0 |
| MLLT11 | 0.27888038 | 5.13E-36 | 0 |
| HK1 | 0.27752649 | 5.94E-36 | 0 |
| ITGA4 | -0.6861827 | 6.44E-36 | 0 |
| PFN1 | 0.26253539 | 9.21E-36 | 0 |
| PRF1 | 0.29910747 | 1.09E-35 | 0 |
| TPO | 0.27449532 | 1.54E-35 | 0 |
| RORB | -0.5259411 | 1.79E-35 | 0 |
| NDUFA12 | 0.31770484 | 2.19E-35 | 0 |
| ENOSF1 | -0.388143 | 2.84E-35 | 0 |

| | | | |
|---|---|---|---|
| H1FX | -0.4941001 | 5.10E-35 | 0 |
| PTGDR | 0.323514 | 1.47E-34 | 0 |
| TMOD3 | 0.33036023 | 1.72E-34 | 0 |
| FCMR | 0.31631358 | 3.49E-34 | 0 |
| STMN1 | -0.5745089 | 3.77E-34 | 0 |
| FGR | 0.31604641 | 1.37E-33 | 0 |
| CD27 | 0.30955821 | 1.86E-33 | 0 |
| ERCC6 | 0.2797884 | 2.26E-32 | 0 |
| RBM38 | 0.28377301 | 3.38E-32 | 0 |
| MCUB | 0.27535509 | 3.56E-32 | 0 |
| LIME1 | 0.2909943 | 3.91E-32 | 0 |
| SUN2 | 0.29326389 | 4.55E-32 | 0 |
| ZMAT3 | 0.30562789 | 7.39E-32 | 0 |
| CHDH | -0.569417 | 8.04E-32 | 0 |
| SP110 | 0.30177558 | 8.53E-32 | 0 |
| CSTB | 0.31413133 | 8.55E-32 | 0 |
| DSTN | 0.35235599 | 9.42E-32 | 0 |
| RHOA | 0.30327475 | 1.51E-31 | 0 |
| IL7R | 0.35266007 | 2.14E-31 | 0 |
| ARHGEF18 | 0.25232584 | 4.48E-31 | 0 |
| NLRC5 | 0.26237682 | 5.34E-31 | 0 |
| SRSF10 | -0.4049883 | 8.93E-31 | 0 |
| PDCD10 | 0.29315613 | 9.07E-31 | 0 |
| LIMS1 | 0.36902708 | 1.03E-30 | 0 |
| STK4 | 0.28640771 | 2.12E-30 | 0 |
| SKAP1 | 0.30440868 | 2.76E-30 | 0 |
| CRBN | 0.27505083 | 6.51E-30 | 0 |
| ITGAE | -0.4167662 | 1.57E-29 | 0 |
| DOCK11 | 0.27146155 | 1.75E-29 | 0 |
| A2M-AS1 | 0.25177169 | 2.13E-29 | 0 |
| ARF6 | 0.28651987 | 3.11E-29 | 0 |
| GNAI2 | 0.29251147 | 3.19E-29 | 0 |
| RASGRP1 | 0.29570754 | 3.67E-29 | 0 |
| RPL14 | 0.25454757 | 4.26E-29 | 0 |
| GBP1 | 0.26266989 | 4.34E-29 | 0 |
| FBXW5 | 0.25270265 | 4.56E-29 | 0 |
| CBLB | 0.26257162 | 4.60E-29 | 0 |
| NTM | -0.2998458 | 6.54E-29 | 0 |
| TRG-AS1 | 0.26545181 | 1.04E-28 | 0 |
| HSP90AA1 | -0.3915765 | 1.51E-28 | 0 |
| SERPINB1 | 0.30568323 | 2.01E-28 | 0 |
| B3GNT2 | 0.27700585 | 3.47E-28 | 0 |
| SPRY1 | -0.5535198 | 7.46E-28 | 0 |

| | | | |
|---|---|---|---|
| VAMP8 | 0.26581652 | 2.03E-27 | 0 |
| REEP5 | 0.25776553 | 2.31E-27 | 0 |
| RBMX | -0.3774835 | 2.60E-27 | 0 |
| HMGN1 | -0.2892498 | 2.80E-27 | 0 |
| IFITM2 | 0.42424763 | 3.56E-27 | 0 |
| PLP2 | 0.31782138 | 3.57E-27 | 0 |
| OXNAD1 | -0.3695824 | 4.17E-27 | 0 |
| WASF2 | 0.28460626 | 5.86E-27 | 0 |
| GPSM3 | 0.28596727 | 7.44E-27 | 0 |
| LDLRAD4 | -0.3862589 | 8.41E-27 | 0 |
| IDI1 | 0.26999964 | 1.00E-26 | 0 |
| XIST | -0.4779365 | 1.22E-26 | 0 |
| CUX1 | -0.3490825 | 1.73E-26 | 0 |
| FDFT1 | 0.28840128 | 2.13E-26 | 0 |
| IL2RG | 0.28438748 | 3.56E-26 | 0 |
| CCDC88A | -0.4204049 | 3.90E-26 | 0 |
| FTL | 0.26266906 | 4.96E-26 | 0 |
| CXXC5 | 0.3300861 | 6.31E-26 | 0 |
| DBI | 0.25837225 | 1.09E-25 | 0 |
| HIST1H1D | -0.5814998 | 1.16E-25 | 0 |
| SEMA5A | -0.2730422 | 1.53E-25 | 0 |
| PPARA | -0.4188942 | 1.61E-25 | 0 |
| DIP2A | 0.3011156 | 2.02E-25 | 0 |
| ARPP21 | -0.4180681 | 2.26E-25 | 0 |
| TWF2 | 0.25933892 | 2.54E-25 | 0 |
| COTL1 | 0.27124672 | 2.78E-25 | 0 |
| TFF3 | -0.43986 | 3.27E-25 | 0 |
| LTB | 0.30513265 | 3.64E-25 | 0 |
| JUND | 0.33284778 | 4.03E-25 | 0 |
| MAP4 | 0.25663335 | 7.76E-25 | 0 |
| STMN3 | 0.26091605 | 8.82E-25 | 0 |
| RHOF | 0.25799945 | 9.20E-25 | 0 |
| PRMT7 | -0.4610066 | 9.83E-25 | 0 |
| AC144521.1 | -0.3284135 | 1.77E-24 | 0 |
| ARL6IP5 | 0.30076952 | 1.97E-24 | 0 |
| MVB12B | -0.2713792 | 2.66E-24 | 0 |
| CENPV | -0.3085825 | 3.36E-24 | 0 |
| PPP2R5C | 0.3003998 | 3.37E-24 | 0 |
| QSOX1 | -0.2818831 | 3.74E-24 | 0 |
| ATP1A1 | 0.27057113 | 3.82E-24 | 0 |
| TRA2B | -0.3421313 | 4.85E-24 | 0 |
| NDUFB7 | 0.26043497 | 9.46E-24 | 0 |
| ZNF107 | -0.3792172 | 1.26E-23 | 0 |

| | | | |
|---|---|---|---|
| KLF13 | 0.28591747 | 1.40E-23 | 0 |
| TLE4 | 0.26160972 | 1.59E-23 | 0 |
| C12orf75 | 0.30272511 | 1.93E-23 | 0 |
| AP005482.1 | -0.4469244 | 2.14E-23 | 0 |
| PCLAF | -0.7175479 | 2.28E-23 | 0 |
| IKZF2 | -0.4947273 | 2.34E-23 | 0 |
| H2AFZ | -0.6389416 | 2.38E-23 | 0 |
| TCL1A | -0.4295599 | 2.41E-23 | 0 |
| PCAT18 | -0.2942715 | 2.66E-23 | 0 |
| ARID5B | 0.27973683 | 2.72E-23 | 0 |
| RHOC | 0.25037314 | 2.86E-23 | 0 |
| GTF3A | 0.26170532 | 4.94E-23 | 0 |
| GTF2I | -0.280295 | 6.95E-23 | 0 |
| DGKZ | 0.25098381 | 1.41E-22 | 0 |
| STK17A | -0.3704607 | 1.62E-22 | 0 |
| TMC6 | 0.26197182 | 2.25E-22 | 0 |
| SARAF | 0.26016996 | 2.81E-22 | 0 |
| ARHGAP30 | 0.27573473 | 5.01E-22 | 0 |
| GNB2 | 0.25243567 | 8.21E-22 | 0 |
| EIF3A | 0.27547828 | 1.78E-21 | 0 |
| IRF1 | 0.25669346 | 1.89E-21 | 0 |
| CDPF1 | -0.2588866 | 3.02E-21 | 0 |
| REPIN1 | -0.2928764 | 4.05E-21 | 0 |
| CD44 | 0.25692117 | 5.10E-21 | 0 |
| MTPN | 0.25848324 | 1.53E-20 | 0 |
| FUS | -0.2800737 | 1.85E-20 | 0 |
| CDK6 | -0.5700405 | 2.52E-20 | 0 |
| COX6C | -0.3105984 | 3.71E-20 | 0 |
| TAOK3 | 0.25078187 | 3.93E-20 | 0 |
| RGCC | 0.36668679 | 5.10E-20 | 0 |
| AIF1 | -0.3456104 | 5.35E-20 | 0 |
| FMNL1 | 0.26387008 | 5.96E-20 | 0 |
| TNRC6C | -0.3670236 | 1.21E-19 | 0 |
| LAPTM5 | 0.26672894 | 1.32E-19 | 0 |
| AFF3 | -0.3446047 | 2.07E-19 | 0 |
| SLC5A3 | -0.3845246 | 2.71E-19 | 0 |
| HIPK1 | -0.3095983 | 7.28E-19 | 0 |
| NEGR1 | -0.39048 | 8.26E-19 | 0 |
| RGS10 | -0.2935559 | 9.90E-19 | 0 |
| NIBAN3 | -0.2607322 | 1.03E-18 | 0 |
| LINC01222 | -0.4642108 | 1.08E-18 | 0 |
| LEF1 | -0.2546686 | 1.61E-18 | 0 |
| BCL11A | -0.2643503 | 3.50E-18 | 0 |

| | | | |
|---|---|---|---|
| CRNDE | -0.3598395 | 4.01E-18 | 0 |
| OSBPL8 | 0.26120803 | 5.37E-18 | 0 |
| NAP1L1 | -0.2893985 | 7.25E-18 | 0 |
| HLA-A | 0.25881001 | 8.16E-18 | 0 |
| MKI67 | -0.8186985 | 8.86E-18 | 0 |
| HIST1H2AC | -0.39165 | 1.17E-17 | 0 |
| S100B | 0.39322735 | 2.36E-17 | 0 |
| CKS2 | -0.369148 | 6.90E-17 | 0 |
| HMGA1 | -0.3778358 | 7.85E-17 | 0 |
| RNASEH2B | -0.3172172 | 9.25E-17 | 0 |
| ARAP2 | 0.26802292 | 1.02E-16 | 0 |
| PALM2-AKAP2 | -0.3521685 | 1.34E-16 | 0 |
| ARHGEF7 | -0.2600654 | 2.88E-16 | 0 |
| ANXA5 | 0.25462665 | 3.67E-16 | 0 |
| KAT6B | -0.2891157 | 3.73E-16 | 0 |
| SESN3 | -0.3212283 | 6.20E-16 | 0 |
| ELOVL4 | -0.3945717 | 7.94E-16 | 0 |
| CD226 | 0.2501574 | 1.19E-15 | 0 |
| MXD4 | -0.3125843 | 1.73E-15 | 0 |
| GALNT6 | -0.3447049 | 2.79E-15 | 0 |
| PLEKHG1 | -0.3735477 | 5.77E-15 | 0 |
| LINC00891 | -0.3158359 | 7.32E-15 | 0 |
| NUCKS1 | -0.3924878 | 7.32E-15 | 0 |
| DNTT | -0.3248786 | 8.85E-15 | 0 |
| LDHB | -0.2596966 | 1.14E-14 | 0 |
| APBB1IP | -0.2618158 | 1.23E-14 | 0 |
| C21orf58 | -0.3121633 | 1.27E-14 | 0 |
| RFLNB | -0.2715295 | 1.53E-14 | 0 |
| TNFAIP3 | -0.3486335 | 2.17E-14 | 0 |
| ZWINT | -0.262218 | 1.20E-13 | 0 |
| FARS2 | -0.3404496 | 1.60E-13 | 0 |
| GZMM | 0.26779741 | 1.61E-13 | 0 |
| CBFA2T3 | -0.2596065 | 1.76E-13 | 0 |
| ARHGEF6 | -0.284324 | 2.10E-13 | 0 |
| LRP12 | -0.2957775 | 3.07E-13 | 0 |
| HSPA1B | -0.3897938 | 4.21E-13 | 0 |
| HIST1H2AG | -0.2684328 | 4.66E-13 | 0 |
| UHRF1 | -0.2954334 | 9.07E-13 | 0 |
| RRM2 | -0.3979312 | 1.03E-12 | 0 |
| CR2 | -0.2593561 | 1.08E-12 | 0 |
| HMGB2 | -0.8880016 | 1.22E-12 | 0 |
| MYB | -0.349635 | 3.28E-12 | 0 |
| MLXIP | -0.284089 | 3.63E-12 | 0 |

| | | | |
|---|---|---|---|
| INSIG1 | 0.25856802 | 3.79E-12 | 0 |
| GSTP1 | -0.3650509 | 4.28E-12 | 0 |
| SLC12A6 | -0.2907665 | 6.86E-12 | 0 |
| NASP | -0.330461 | 8.71E-12 | 0 |
| GLUL | -0.3410073 | 9.21E-12 | 0 |
| TYMS | -0.6724109 | 1.23E-11 | 0 |
| AC068587.4 | -0.3308955 | 1.63E-11 | 0 |
| ANP32B | -0.2921477 | 1.89E-11 | 0 |
| H3F3B | -0.2555138 | 2.11E-11 | 0 |
| AQP3 | -0.3589038 | 3.02E-11 | 0 |
| PTPN22 | -0.2760384 | 6.53E-11 | 0 |
| CDK5RAP3 | -0.3201791 | 1.44E-10 | 0 |
| AHI1 | -0.2614573 | 1.51E-10 | 0 |
| HIST1H1B | -0.5791263 | 1.51E-10 | 0 |
| GAPDH | -0.3596894 | 8.18E-10 | 0 |
| GTSE1 | -0.3715224 | 1.10E-09 | 0 |
| CD8A | -0.3044737 | 3.22E-09 | 0 |
| ZFP36L2 | -0.301428 | 3.89E-09 | 0 |
| NCF1 | -0.2920211 | 6.91E-09 | 0 |
| HIST1H3B | -0.3770089 | 9.03E-09 | 0 |
| KNL1 | -0.3043782 | 9.83E-09 | 0 |
| CDK1 | -0.2834491 | 1.99E-08 | 0 |
| AC002454.1 | -0.2667346 | 2.86E-08 | 0 |
| TUBB | -0.5689186 | 3.54E-08 | 0 |
| PIK3R3 | -0.3196566 | 4.17E-08 | 0 |
| TPX2 | -0.293224 | 4.27E-08 | 0 |
| NUSAP1 | -0.5894038 | 6.23E-08 | 0 |
| HIST1H1C | -0.4794332 | 9.18E-08 | 0 |
| AAK1 | -0.2829989 | 1.07E-07 | 0 |
| MDM4 | -0.2570713 | 1.17E-07 | 0 |
| ZNF280D | -0.2601477 | 1.67E-07 | 0 |
| TMPO | -0.3316435 | 2.93E-07 | 0 |
| HIST1H3D | -0.4986241 | 3.26E-07 | 0 |
| SSBP2 | -0.2545995 | 5.85E-07 | 0 |
| CBX5 | -0.2807608 | 8.57E-07 | 0 |
| CALR | -0.2996391 | 9.07E-07 | 0 |
| BTG2 | -0.3284002 | 1.13E-06 | 0 |
| H2AFY | -0.2531161 | 2.07E-06 | 0 |
| TRIM14 | -0.2898961 | 2.68E-06 | 0 |
| PLIN2 | -0.2512948 | 3.95E-06 | 0 |
| CLSPN | -0.3124197 | 5.07E-06 | 0 |
| HIST1H4C | -1.1550562 | 8.10E-06 | 0 |
| LMNB1 | -0.2833874 | 9.81E-06 | 0 |

| | | | |
|---|---|---|---|
| FABP5 | -0.3057656 | 2.14E-05 | 0 |
| PPP1R14B | -0.2758672 | 5.94E-05 | 0 |
| ASPM | -0.5491266 | 6.39E-05 | 0 |
| UBE2C | -0.2850742 | 0.00013561 | 0 |
| IDH2 | -0.2557987 | 0.00014647 | 0 |
| H2AFX | -0.3108445 | 0.00023896 | 0 |
| HSP90B1 | -0.2983406 | 0.00062189 | 0 |
| GTF3C5 | -0.2716209 | 0.00066157 | 0 |
| DLEU2 | -0.2606595 | 0.00088374 | 0 |
| SMC2 | -0.2995867 | 0.00262355 | 0 |
| RAN | -0.2801464 | 0.01374398 | 0 |
| EZH2 | -0.3114108 | 0.04173454 | 0 |
| DUT | -0.395622 | 0.09838596 | 0 |
| TOP2A | -0.7243716 | 0.13596851 | 0 |
| MCM7 | -0.3175886 | 0.15342042 | 0 |
| PTTG1 | -0.2663224 | 0.37648727 | 0 |
| ATAD2 | -0.304997 | 0.4033895 | 0 |
| ITM2A | -0.3825342 | 0.42471027 | 0 |
| PCNA | -0.3628194 | 1 | 0 |
| LST1 | -0.2734408 | 1 | 0 |
| UBE2S | -0.2959849 | 1 | 0 |
| B2M | 0.93622372 | 9.20E-262 | 1 |
| TRBC2 | 0.6288928 | 3.50E-216 | 1 |
| LTB | 1.25126637 | 8.82E-210 | 1 |
| VIM | -1.3091002 | 1.03E-178 | 1 |
| IGFBP5 | 1.0493591 | 7.42E-153 | 1 |
| ZFP36L2 | 0.80665332 | 1.66E-147 | 1 |
| HLA-A | 0.8396172 | 3.28E-145 | 1 |
| CHDH | 0.93210718 | 8.15E-144 | 1 |
| BTG1 | 0.78297458 | 1.70E-143 | 1 |
| S100A4 | -1.5393287 | 8.34E-134 | 1 |
| IL17RB | 1.11524578 | 1.14E-133 | 1 |
| TXNIP | 0.7534189 | 5.51E-133 | 1 |
| SMC4 | 0.93292989 | 2.19E-132 | 1 |
| PDE7A | 0.73859989 | 3.53E-132 | 1 |
| FARS2 | 0.84261711 | 1.11E-129 | 1 |
| LGALS1 | -1.7814344 | 1.91E-129 | 1 |
| SMPD3 | 0.78508008 | 5.35E-128 | 1 |
| GAPDH | -0.9126519 | 8.70E-126 | 1 |
| PNRC1 | 0.6515734 | 1.45E-116 | 1 |
| CD2 | 0.58659238 | 2.18E-116 | 1 |
| WAKMAR2 | 0.86063563 | 3.60E-115 | 1 |
| BTG2 | 0.75353231 | 9.29E-107 | 1 |

| | | | |
|---|---|---|---|
| HLA-C | 0.65018279 | 4.20E-106 | 1 |
| CHMP7 | 0.71809545 | 4.24E-101 | 1 |
| RPLP0 | -0.6368176 | 1.98E-99 | 1 |
| TNRC6C | 0.67655782 | 1.93E-92 | 1 |
| CAPG | -0.7610955 | 2.51E-92 | 1 |
| CD8B | 0.54986129 | 5.41E-87 | 1 |
| HLA-B | 0.70187759 | 3.53E-84 | 1 |
| AHNAK | -1.0894223 | 4.99E-83 | 1 |
| TMSB10 | -0.7413064 | 9.24E-82 | 1 |
| MACROD2 | 0.63865826 | 1.38E-80 | 1 |
| NEGR1 | 0.67476063 | 2.50E-78 | 1 |
| RHOH | 0.50399301 | 4.54E-77 | 1 |
| PTPN6 | -0.7415549 | 5.44E-77 | 1 |
| RPS27A | 0.37411946 | 3.59E-76 | 1 |
| STK17B | 0.48667511 | 4.86E-73 | 1 |
| TNFAIP3 | 0.67856112 | 9.37E-73 | 1 |
| HSP90B1 | 0.74028188 | 8.48E-71 | 1 |
| MZB1 | 0.58705935 | 2.82E-70 | 1 |
| RFLNB | 0.54670928 | 6.22E-70 | 1 |
| CRIP1 | -1.3525054 | 7.40E-67 | 1 |
| NME2 | -0.6365288 | 6.77E-66 | 1 |
| CD7 | 0.50526869 | 8.88E-66 | 1 |
| PLXDC1 | 0.36856654 | 1.41E-64 | 1 |
| ATP5F1E | 0.38378806 | 8.23E-64 | 1 |
| HNRNPA1 | -0.5307791 | 1.91E-63 | 1 |
| KLF2 | -0.8996171 | 3.74E-63 | 1 |
| RPS29 | 0.31631314 | 7.60E-63 | 1 |
| RPL14 | -0.4547275 | 7.56E-62 | 1 |
| CLNK | 0.42818701 | 1.71E-61 | 1 |
| S100A10 | -0.7515827 | 3.30E-60 | 1 |
| CD79A | 0.71559837 | 5.70E-60 | 1 |
| SCRIB | 0.52252913 | 8.74E-59 | 1 |
| RPS3A | 0.30782486 | 4.16E-58 | 1 |
| MAL | 0.52557808 | 7.68E-57 | 1 |
| ARMH1 | -0.6212566 | 2.46E-55 | 1 |
| FTL | 0.46109503 | 2.60E-55 | 1 |
| GIHCG | -0.7409771 | 2.64E-55 | 1 |
| EMP3 | -0.7552615 | 1.33E-54 | 1 |
| YWHAH | -0.4878968 | 2.16E-53 | 1 |
| FXYD2 | -1.207626 | 2.19E-53 | 1 |
| SLC25A3 | -0.4869498 | 1.55E-52 | 1 |
| TECR | -0.8508003 | 2.05E-52 | 1 |
| CCND3 | -0.4749753 | 2.52E-51 | 1 |

| | | | |
|---|---|---|---|
| ID3 | -0.8546076 | 1.58E-50 | 1 |
| XBP1 | 0.63779705 | 5.59E-50 | 1 |
| STMN1 | -0.6471348 | 1.13E-49 | 1 |
| TAGLN2 | -0.640503 | 1.96E-49 | 1 |
| GSTP1 | -0.5670672 | 1.26E-48 | 1 |
| PKM | -0.5161848 | 9.29E-48 | 1 |
| RPL30 | 0.29534913 | 3.54E-46 | 1 |
| SELL | -0.4967611 | 6.51E-46 | 1 |
| COX5A | -0.4304694 | 1.35E-45 | 1 |
| YWHAE | -0.4317757 | 5.47E-45 | 1 |
| TPT1 | 0.27371693 | 1.23E-44 | 1 |
| TPM3 | -0.4373219 | 1.55E-44 | 1 |
| CD8A | 0.46986638 | 1.68E-43 | 1 |
| STK4 | 0.38203789 | 6.00E-43 | 1 |
| S100A11 | -0.5979236 | 9.14E-43 | 1 |
| MDM4 | 0.41220644 | 9.60E-43 | 1 |
| MT-ND3 | 0.26564056 | 9.78E-43 | 1 |
| SPTBN1 | -0.4641871 | 1.65E-42 | 1 |
| TIMP1 | -0.4987785 | 1.75E-42 | 1 |
| VDAC1 | -0.4173729 | 2.00E-42 | 1 |
| FYB1 | 0.35443445 | 2.86E-42 | 1 |
| RGS3 | -0.5464269 | 3.66E-42 | 1 |
| KRAS | -0.4339681 | 6.47E-42 | 1 |
| ANXA2 | -0.4286223 | 6.75E-42 | 1 |
| SELENOH | -0.3846 | 9.86E-42 | 1 |
| ADORA2A | 0.33213112 | 1.15E-41 | 1 |
| ARL4C | 0.34930326 | 1.66E-41 | 1 |
| SLC25A6 | -0.4232034 | 1.68E-41 | 1 |
| SNHG29 | -0.496307 | 2.75E-41 | 1 |
| RPL21 | 0.27485791 | 3.57E-41 | 1 |
| PTPRC | 0.26791986 | 4.18E-41 | 1 |
| CD28 | -0.4580014 | 6.65E-41 | 1 |
| EZR | -0.5147553 | 1.49E-40 | 1 |
| IDH2 | -0.471316 | 2.63E-40 | 1 |
| APEX1 | -0.4082808 | 4.26E-40 | 1 |
| CR2 | 0.40942096 | 6.45E-40 | 1 |
| CD1E | 0.41813412 | 1.60E-39 | 1 |
| PLAC8 | -0.63273 | 3.51E-39 | 1 |
| LIMS1 | -0.420633 | 9.64E-39 | 1 |
| MCUB | -0.3455636 | 1.09E-38 | 1 |
| ESYT1 | -0.3712623 | 2.63E-38 | 1 |
| LIME1 | 0.36174868 | 4.28E-38 | 1 |
| YBX1 | -0.416064 | 5.64E-38 | 1 |

| | | | |
|---|---|---|---|
| RANBP1 | -0.4598711 | 3.62E-37 | 1 |
| CRNDE | -0.4329911 | 4.00E-37 | 1 |
| RPL22 | 0.27499809 | 5.77E-37 | 1 |
| HNRNPAB | -0.4529157 | 7.51E-37 | 1 |
| RPL34 | 0.26672156 | 1.20E-36 | 1 |
| SEPTIN11 | -0.3722031 | 1.25E-36 | 1 |
| AL365361.1 | 0.45923027 | 2.52E-36 | 1 |
| CAPN2 | -0.3439377 | 7.80E-36 | 1 |
| PPIA | -0.4118481 | 1.58E-35 | 1 |
| RPSA | -0.2972178 | 1.94E-35 | 1 |
| ANXA1 | -0.7037689 | 3.03E-35 | 1 |
| CD3G | 0.30816032 | 5.22E-35 | 1 |
| AKNA | 0.37597426 | 9.46E-35 | 1 |
| ACTG1 | -0.3204344 | 1.30E-34 | 1 |
| MLLT11 | -0.3148163 | 1.36E-34 | 1 |
| MYL6B | -0.3380174 | 1.02E-33 | 1 |
| TUBB2A | -0.3049541 | 1.86E-33 | 1 |
| NKG7 | -0.431729 | 2.77E-33 | 1 |
| MT-ND2 | 0.28292274 | 3.43E-33 | 1 |
| RAN | -0.461651 | 3.62E-33 | 1 |
| RHOC | -0.3113851 | 6.07E-33 | 1 |
| FGFR1 | -0.4466093 | 7.78E-33 | 1 |
| SNRPE | -0.3680702 | 1.62E-32 | 1 |
| LEF1 | 0.28931166 | 6.29E-32 | 1 |
| ESYT2 | -0.4315039 | 7.48E-32 | 1 |
| CARHSP1 | -0.3873206 | 2.49E-31 | 1 |
| CLEC2D | -0.5701746 | 3.30E-31 | 1 |
| MCM7 | -0.5026627 | 3.66E-31 | 1 |
| BIRC2 | 0.42084074 | 8.21E-31 | 1 |
| SNRPF | -0.3866791 | 8.84E-31 | 1 |
| EZH2 | -0.4874465 | 9.55E-31 | 1 |
| MYO1G | -0.3562596 | 1.62E-30 | 1 |
| ANXA5 | -0.3540431 | 2.09E-30 | 1 |
| CTSW | -0.5605119 | 2.42E-30 | 1 |
| RHOA | -0.3529525 | 3.39E-30 | 1 |
| PCLAF | -0.7281544 | 4.28E-30 | 1 |
| C4orf48 | -0.3045851 | 4.56E-30 | 1 |
| SLC8A1-AS1 | 0.263962 | 4.98E-30 | 1 |
| GBP2 | -0.3451884 | 7.57E-30 | 1 |
| ENO1 | -0.4076697 | 9.91E-30 | 1 |
| SPRY1 | 0.47103376 | 1.17E-29 | 1 |
| MSN | -0.3827845 | 1.65E-29 | 1 |
| ATP5F1B | -0.3323873 | 1.94E-29 | 1 |

| | | | |
|---|---|---|---|
| CD3E | 0.30223011 | 2.93E-29 | 1 |
| MYH10 | -0.3310771 | 4.73E-29 | 1 |
| DDX5 | 0.26268161 | 8.57E-29 | 1 |
| CENPK | -0.2973665 | 1.09E-28 | 1 |
| PLP2 | -0.3517712 | 1.30E-28 | 1 |
| RBM3 | -0.3366679 | 1.43E-28 | 1 |
| CST7 | -0.3555946 | 1.66E-28 | 1 |
| SYNRG | 0.36795923 | 2.31E-28 | 1 |
| ILF2 | -0.3413117 | 4.34E-28 | 1 |
| CXCR4 | 0.41003315 | 5.64E-28 | 1 |
| TUBB | -0.7332415 | 6.52E-28 | 1 |
| CALM1 | -0.4026058 | 7.14E-28 | 1 |
| SNRPC | -0.3039042 | 8.12E-28 | 1 |
| RPS2 | -0.2813389 | 8.86E-28 | 1 |
| EIF3A | -0.3615299 | 8.87E-28 | 1 |
| SMS | -0.3203203 | 1.09E-27 | 1 |
| SNRPA | -0.2871681 | 1.18E-27 | 1 |
| SARAF | 0.33443379 | 1.19E-27 | 1 |
| MLXIP | 0.35838562 | 1.27E-27 | 1 |
| ACTB | -0.2555605 | 1.28E-27 | 1 |
| CIRBP | 0.32388491 | 1.93E-27 | 1 |
| ATP5PD | -0.3216643 | 2.57E-27 | 1 |
| UBASH3B | -0.3798338 | 2.88E-27 | 1 |
| TXNDC17 | -0.2957687 | 3.82E-27 | 1 |
| NME1 | -0.3009202 | 4.57E-27 | 1 |
| ATP5MC3 | -0.3243652 | 4.57E-27 | 1 |
| YPEL3 | 0.37784345 | 6.87E-27 | 1 |
| RPLP1 | -0.2780918 | 7.88E-27 | 1 |
| HIST1H2AC | 0.44341437 | 8.47E-27 | 1 |
| NSD2 | -0.3617862 | 8.55E-27 | 1 |
| NCF1 | 0.32490284 | 1.25E-26 | 1 |
| CBX5 | -0.4055841 | 1.25E-26 | 1 |
| MAN2A1 | 0.35571592 | 1.26E-26 | 1 |
| NAB2 | -0.2876995 | 1.28E-26 | 1 |
| MCRIP1 | -0.274414 | 2.67E-26 | 1 |
| ATP2B1 | -0.394148 | 2.78E-26 | 1 |
| S100A6 | -0.594173 | 2.98E-26 | 1 |
| NPM1 | -0.3590822 | 3.68E-26 | 1 |
| PSMA4 | -0.3116817 | 5.02E-26 | 1 |
| TYMS | -0.7128209 | 7.38E-26 | 1 |
| BCL11B | 0.32014249 | 7.75E-26 | 1 |
| MCM5 | -0.349519 | 8.44E-26 | 1 |
| RTRAF | -0.2951677 | 1.04E-25 | 1 |

| | | | |
|---|---|---|---|
| HIVEP3 | -0.287325 | 1.11E-25 | 1 |
| MCM6 | -0.3427083 | 1.38E-25 | 1 |
| HELLS | -0.5097041 | 1.83E-25 | 1 |
| SVIL | -0.2559715 | 2.06E-25 | 1 |
| SEC31B | -0.3255736 | 2.30E-25 | 1 |
| C12orf57 | -0.3783579 | 3.10E-25 | 1 |
| ACAP1 | 0.27013879 | 3.40E-25 | 1 |
| MRPS15 | -0.2608183 | 5.44E-25 | 1 |
| HLA-E | 0.30697934 | 5.47E-25 | 1 |
| PPA1 | -0.3212224 | 5.55E-25 | 1 |
| MCM4 | -0.3607906 | 7.40E-25 | 1 |
| EIF4A1 | -0.3416154 | 9.22E-25 | 1 |
| HSPD1 | -0.3814175 | 9.88E-25 | 1 |
| TMEM173 | -0.2663158 | 1.32E-24 | 1 |
| HNRNPD | -0.3594574 | 1.45E-24 | 1 |
| SH2D2A | -0.2519903 | 1.71E-24 | 1 |
| FUT7 | -0.281499 | 2.40E-24 | 1 |
| H3F3B | 0.30946747 | 2.90E-24 | 1 |
| UNG | -0.3026873 | 3.69E-24 | 1 |
| EIF5A | -0.3101811 | 3.72E-24 | 1 |
| SIRPG | 0.33855312 | 4.46E-24 | 1 |
| PHB2 | -0.287971 | 1.02E-23 | 1 |
| HSP90AA1 | -0.3771777 | 1.10E-23 | 1 |
| P2RX5 | -0.354978 | 1.72E-23 | 1 |
| HMGB3 | -0.2937772 | 1.94E-23 | 1 |
| NOSIP | -0.6915499 | 3.07E-23 | 1 |
| APBB1IP | 0.29236043 | 3.18E-23 | 1 |
| LDHA | -0.3668129 | 3.21E-23 | 1 |
| ATP5MC1 | -0.2657725 | 4.71E-23 | 1 |
| FKBP3 | -0.282185 | 5.24E-23 | 1 |
| RORB | 0.44907526 | 7.13E-23 | 1 |
| PTTG1 | -0.3976255 | 7.93E-23 | 1 |
| CDT1 | -0.2537477 | 8.18E-23 | 1 |
| PIK3IP1 | 0.29077439 | 8.61E-23 | 1 |
| IKZF3 | -0.4092845 | 8.66E-23 | 1 |
| PSMB7 | -0.2543473 | 1.06E-22 | 1 |
| SLC18A2 | -0.2880602 | 1.27E-22 | 1 |
| COPE | -0.2709361 | 1.40E-22 | 1 |
| ZNF683 | 0.33384376 | 1.54E-22 | 1 |
| RBX1 | -0.2900649 | 1.55E-22 | 1 |
| TUBA1B | -0.9109264 | 1.68E-22 | 1 |
| SSRP1 | -0.2997905 | 1.84E-22 | 1 |
| PNISR | 0.2661606 | 1.98E-22 | 1 |

| | | | |
|---|---|---|---|
| HNRNPDL | -0.3119232 | 2.21E-22 | 1 |
| AAK1 | 0.32173181 | 2.39E-22 | 1 |
| HSPA1B | -0.44022 | 3.43E-22 | 1 |
| SNRPD1 | -0.3322649 | 3.95E-22 | 1 |
| COX7A2 | -0.2817311 | 4.42E-22 | 1 |
| TPST2 | -0.3262875 | 5.31E-22 | 1 |
| OXNAD1 | 0.34362005 | 5.55E-22 | 1 |
| TRAC | 0.28757356 | 6.46E-22 | 1 |
| GAS5 | -0.3412579 | 6.98E-22 | 1 |
| TRIM14 | 0.34297421 | 8.83E-22 | 1 |
| PDCD1 | -0.2536442 | 9.87E-22 | 1 |
| ADD3 | -0.4977965 | 1.19E-21 | 1 |
| IQGAP1 | -0.2975918 | 1.39E-21 | 1 |
| YBX3 | -0.2579489 | 1.40E-21 | 1 |
| H2AFY | -0.328385 | 2.29E-21 | 1 |
| TSPAN2 | -0.2615826 | 2.53E-21 | 1 |
| EIF4A2 | 0.27021845 | 2.83E-21 | 1 |
| ILF3 | -0.2935261 | 3.05E-21 | 1 |
| YWHAQ | -0.2959197 | 3.60E-21 | 1 |
| ANP32E | -0.4384029 | 3.89E-21 | 1 |
| TKT | -0.2956734 | 5.03E-21 | 1 |
| EDEM1 | 0.35928509 | 5.19E-21 | 1 |
| ZWINT | -0.2955174 | 5.21E-21 | 1 |
| H1FX | 0.31384896 | 5.59E-21 | 1 |
| SLC25A5 | -0.3373383 | 5.66E-21 | 1 |
| CDK6 | -0.5720295 | 5.93E-21 | 1 |
| SEPTIN9 | 0.27891187 | 8.95E-21 | 1 |
| NEAT1 | -0.470502 | 9.17E-21 | 1 |
| NIBAN3 | 0.2729014 | 9.73E-21 | 1 |
| IGFLR1 | -0.2828796 | 1.12E-20 | 1 |
| SNHG18 | 0.25708975 | 1.17E-20 | 1 |
| CDK2AP1 | -0.2936944 | 1.39E-20 | 1 |
| FAM102A | 0.30403705 | 1.45E-20 | 1 |
| FBLN2 | 0.27763284 | 1.76E-20 | 1 |
| RBBP8 | -0.3219918 | 2.05E-20 | 1 |
| PAICS | -0.2811901 | 2.87E-20 | 1 |
| NDUFAB1 | -0.2558132 | 2.87E-20 | 1 |
| CTSD | 0.33091799 | 2.93E-20 | 1 |
| RNASEH2B | 0.2992374 | 3.10E-20 | 1 |
| SNRPG | -0.3044117 | 3.35E-20 | 1 |
| FNBP1 | 0.25982402 | 3.47E-20 | 1 |
| DIAPH1 | -0.2809487 | 4.04E-20 | 1 |
| FXYD5 | -0.325805 | 4.34E-20 | 1 |

| | | | |
|---|---|---|---|
| SRSF2 | -0.3278942 | 4.53E-20 | 1 |
| HSP90AB1 | -0.4517236 | 5.08E-20 | 1 |
| RRM1 | -0.317231 | 5.17E-20 | 1 |
| SYNE2 | -0.4311793 | 5.26E-20 | 1 |
| CLSPN | -0.3823932 | 5.59E-20 | 1 |
| ISCU | 0.29432321 | 6.43E-20 | 1 |
| WDR34 | -0.2798556 | 6.97E-20 | 1 |
| EEF1G | -0.2555498 | 7.54E-20 | 1 |
| PSMB3 | -0.2587529 | 7.80E-20 | 1 |
| SMC2 | -0.4020869 | 1.07E-19 | 1 |
| UCP2 | -0.3497032 | 1.26E-19 | 1 |
| TSC22D3 | -0.3985827 | 1.46E-19 | 1 |
| NONO | -0.2664936 | 1.48E-19 | 1 |
| TRIM28 | -0.2926916 | 2.12E-19 | 1 |
| EIF3L | -0.2717514 | 2.26E-19 | 1 |
| PPM1G | -0.2550157 | 2.33E-19 | 1 |
| FBL | -0.2724572 | 2.36E-19 | 1 |
| RUNX1 | -0.3496252 | 2.73E-19 | 1 |
| P2RY11 | -0.271735 | 2.80E-19 | 1 |
| KLHL23 | -0.2667372 | 3.35E-19 | 1 |
| PPARA | 0.29999299 | 4.41E-19 | 1 |
| FAM111B | -0.2647915 | 4.45E-19 | 1 |
| MXD4 | 0.31167878 | 5.36E-19 | 1 |
| LMNB1 | -0.3634288 | 5.78E-19 | 1 |
| SUSD3 | 0.25898757 | 5.92E-19 | 1 |
| MT-ND6 | -0.3308917 | 6.21E-19 | 1 |
| POLR2L | -0.2774796 | 6.73E-19 | 1 |
| FABP5 | -0.364869 | 8.53E-19 | 1 |
| NDUFB7 | -0.2511813 | 1.11E-18 | 1 |
| SNRPB | -0.2815579 | 1.18E-18 | 1 |
| SCD | -0.2535471 | 1.31E-18 | 1 |
| CAPNS1 | -0.2555389 | 1.43E-18 | 1 |
| MAZ | -0.2597886 | 1.47E-18 | 1 |
| SPN | -0.2858119 | 1.87E-18 | 1 |
| NOP53 | -0.3020659 | 4.04E-18 | 1 |
| TOMM7 | 0.27635634 | 4.37E-18 | 1 |
| GMNN | -0.2538344 | 4.63E-18 | 1 |
| HNRNPM | -0.2948854 | 5.50E-18 | 1 |
| C12orf75 | -0.294169 | 6.38E-18 | 1 |
| SLFN5 | 0.25975719 | 7.33E-18 | 1 |
| DDX6 | 0.30351109 | 7.83E-18 | 1 |
| C17orf49 | -0.2508231 | 1.02E-17 | 1 |
| PCNA | -0.4938795 | 1.34E-17 | 1 |

| | | | |
|---|---|---|---|
| CCR9 | 0.31190551 | 1.39E-17 | 1 |
| EIF5B | -0.260709 | 1.43E-17 | 1 |
| CENPU | -0.2827878 | 1.67E-17 | 1 |
| DAZAP1 | -0.2687524 | 2.35E-17 | 1 |
| RGS10 | 0.25796012 | 2.48E-17 | 1 |
| CLIC1 | -0.3013855 | 2.49E-17 | 1 |
| CCT2 | -0.2644516 | 2.73E-17 | 1 |
| NUSAP1 | -0.6306592 | 3.00E-17 | 1 |
| ELOVL5 | -0.2619922 | 3.13E-17 | 1 |
| PA2G4 | -0.2887646 | 3.63E-17 | 1 |
| ANP32B | -0.3325053 | 3.72E-17 | 1 |
| MSH6 | -0.2918111 | 3.95E-17 | 1 |
| UHRF1 | -0.3060912 | 4.00E-17 | 1 |
| CD84 | -0.3698151 | 4.20E-17 | 1 |
| MYOM2 | -0.3256259 | 4.33E-17 | 1 |
| SH3BGRL3 | -0.372418 | 5.38E-17 | 1 |
| NDUFA12 | -0.2510237 | 8.20E-17 | 1 |
| CDC25B | -0.2784092 | 9.68E-17 | 1 |
| RNASEH2C | -0.2563083 | 9.76E-17 | 1 |
| REC8 | -0.2534717 | 9.96E-17 | 1 |
| ESD | -0.2536467 | 1.17E-16 | 1 |
| TIMM13 | -0.2566318 | 1.21E-16 | 1 |
| SBK1 | -0.2516297 | 1.37E-16 | 1 |
| CD82 | -0.2553987 | 1.41E-16 | 1 |
| PSMA6 | -0.2889161 | 1.57E-16 | 1 |
| HNRNPA2B1 | -0.2725962 | 2.76E-16 | 1 |
| HIPK1 | 0.28541434 | 3.30E-16 | 1 |
| RTKN2 | -0.3105377 | 4.00E-16 | 1 |
| C1QBP | -0.2977987 | 4.81E-16 | 1 |
| TPI1 | -0.2690647 | 5.20E-16 | 1 |
| GIMAP4 | -0.2887923 | 5.44E-16 | 1 |
| H2AFV | -0.3033546 | 5.67E-16 | 1 |
| HSPB1 | 0.3204722 | 6.30E-16 | 1 |
| ALYREF | -0.2621866 | 6.94E-16 | 1 |
| PHACTR2 | -0.2699679 | 1.12E-15 | 1 |
| FDFT1 | -0.2639469 | 1.32E-15 | 1 |
| TUBB4B | -0.4109022 | 1.48E-15 | 1 |
| CD247 | -0.3476369 | 1.72E-15 | 1 |
| RALY | -0.2577904 | 1.72E-15 | 1 |
| HMGA1 | -0.3307959 | 2.12E-15 | 1 |
| TUBA1A | -0.3039457 | 3.36E-15 | 1 |
| UQCRQ | -0.2579602 | 3.61E-15 | 1 |
| NASP | -0.318563 | 3.93E-15 | 1 |

| | | | |
|---|---|---|---|
| COX8A | -0.2691958 | 3.96E-15 | 1 |
| BACH2 | -0.2831531 | 4.65E-15 | 1 |
| HSPA1A | -0.4028983 | 4.76E-15 | 1 |
| ACTN1 | 0.2591731 | 7.35E-15 | 1 |
| ITGA6 | -0.3047744 | 8.72E-15 | 1 |
| RASGRP1 | -0.2900517 | 1.02E-14 | 1 |
| PPP1R14B | -0.3163372 | 1.48E-14 | 1 |
| AC002454.1 | -0.2902907 | 1.84E-14 | 1 |
| RASGRP2 | -0.2593869 | 2.06E-14 | 1 |
| HSPE1 | -0.2903639 | 2.10E-14 | 1 |
| GTF3C5 | -0.2994282 | 2.78E-14 | 1 |
| PRDX2 | -0.299469 | 3.14E-14 | 1 |
| PTMA | -0.2859088 | 3.70E-14 | 1 |
| ITGA4 | -0.545286 | 4.26E-14 | 1 |
| USP1 | -0.3185862 | 6.77E-14 | 1 |
| DOCK10 | 0.25496159 | 2.50E-13 | 1 |
| ARHGAP26 | 0.27055527 | 2.93E-13 | 1 |
| DUSP2 | -0.3174871 | 3.67E-13 | 1 |
| ATP2A1 | 0.30110771 | 4.19E-13 | 1 |
| PRDX1 | -0.2598239 | 5.18E-13 | 1 |
| NCAPG2 | -0.3234614 | 6.45E-13 | 1 |
| RORA | -0.2600795 | 8.18E-13 | 1 |
| NR4A1 | -0.2632433 | 1.06E-12 | 1 |
| LITAF | -0.2687592 | 1.79E-12 | 1 |
| DNTT | -0.3038407 | 1.80E-12 | 1 |
| FDPS | -0.2532353 | 1.83E-12 | 1 |
| GIMAP7 | -0.3524132 | 1.92E-12 | 1 |
| PTPN22 | 0.26917468 | 1.97E-12 | 1 |
| LRRFIP1 | -0.2505072 | 2.28E-12 | 1 |
| LSM4 | -0.2714907 | 2.32E-12 | 1 |
| CENPF | -0.7232044 | 2.56E-12 | 1 |
| JTB | 0.27151886 | 3.03E-12 | 1 |
| IL7R | -0.2638396 | 3.17E-12 | 1 |
| CDCA7 | -0.2577955 | 3.35E-12 | 1 |
| H2AFZ | -0.5234883 | 8.05E-12 | 1 |
| C21orf58 | -0.2834798 | 8.32E-12 | 1 |
| HDAC7 | 0.26883461 | 8.95E-12 | 1 |
| PLEKHG1 | 0.25594377 | 1.46E-11 | 1 |
| SELENOW | -0.2948466 | 1.69E-11 | 1 |
| ANKRD36C | -0.2577304 | 2.42E-11 | 1 |
| H2AFX | -0.34911 | 2.67E-11 | 1 |
| TSPAN7 | -0.2988916 | 2.71E-11 | 1 |
| MIF | -0.3010177 | 5.17E-11 | 1 |

| | | | |
|---|---|---|---|
| SAMHD1 | -0.2947822 | 7.27E-11 | 1 |
| ZNF92 | 0.31063594 | 9.45E-11 | 1 |
| TCF12 | 0.28382699 | 1.27E-10 | 1 |
| PTPN7 | -0.2961839 | 1.29E-10 | 1 |
| KLRB1 | -0.3885746 | 1.29E-10 | 1 |
| ITM2A | -0.5589226 | 1.81E-10 | 1 |
| ATAD2 | -0.3594572 | 3.40E-10 | 1 |
| DEK | -0.2601026 | 4.32E-10 | 1 |
| MYO1F | -0.2665387 | 7.40E-10 | 1 |
| TXN | -0.3015104 | 9.32E-10 | 1 |
| PRKCH | -0.2617805 | 1.28E-09 | 1 |
| RIPOR2 | -0.313775 | 1.38E-09 | 1 |
| STAT5A | -0.3112528 | 2.00E-09 | 1 |
| SEMA4D | -0.2795451 | 2.43E-09 | 1 |
| DUT | -0.383749 | 3.33E-09 | 1 |
| UBE2S | -0.3483068 | 3.55E-09 | 1 |
| TCF7 | -0.2602685 | 3.55E-09 | 1 |
| CCDC88A | -0.2526222 | 4.45E-09 | 1 |
| ZNF280D | 0.2695836 | 4.59E-09 | 1 |
| KNL1 | -0.3071852 | 6.02E-09 | 1 |
| CDK1 | -0.2739268 | 1.10E-08 | 1 |
| GTSE1 | -0.3490531 | 1.17E-08 | 1 |
| MCM3 | -0.2549131 | 1.26E-08 | 1 |
| CDK5RAP3 | -0.269421 | 1.62E-08 | 1 |
| KLRK1 | -0.291951 | 1.81E-08 | 1 |
| SRGN | -0.3777012 | 6.49E-08 | 1 |
| DHFR | -0.2667726 | 1.53E-07 | 1 |
| NUCKS1 | -0.3282821 | 1.54E-07 | 1 |
| CCDC18-AS1 | 0.2511848 | 1.96E-07 | 1 |
| LAT2 | 0.25109954 | 2.44E-07 | 1 |
| CKS2 | -0.2680518 | 4.23E-07 | 1 |
| SLC5A3 | -0.2564636 | 9.68E-07 | 1 |
| RRM2 | -0.3547427 | 1.16E-06 | 1 |
| TPX2 | -0.2663101 | 1.93E-06 | 1 |
| TOP2A | -0.7049901 | 2.37E-06 | 1 |
| FLNA | -0.2696848 | 3.70E-06 | 1 |
| LST1 | -0.3177958 | 0.00013524 | 1 |
| ASPM | -0.5011328 | 0.00099859 | 1 |
| HIST1H4C | -0.7991649 | 0.01467164 | 1 |
| HIST1H3B | -0.3197544 | 0.01592604 | 1 |
| MACF1 | -0.2521747 | 0.02146744 | 1 |
| UBE2C | -0.2629065 | 0.02201753 | 1 |
| TFDP2 | -0.332 | 0.08672518 | 1 |

| | | | |
|---|---|---|---|
| HIST1H1B | -0.4774564 | 0.41331823 | 1 |
| HMGB2 | -0.7521484 | 0.75201347 | 1 |
| MKI67 | -0.6378827 | 0.80367024 | 1 |
| TOX2 | 0.8527964 | 1.28E-128 | 2 |
| RPL37 | 0.47073757 | 3.50E-116 | 2 |
| RPL34 | 0.49258916 | 1.49E-111 | 2 |
| RPS27 | 0.49805468 | 8.45E-111 | 2 |
| TMSB4X | 0.4388937 | 5.96E-110 | 2 |
| ZNF683 | 1.13985243 | 3.23E-108 | 2 |
| SATB1 | 0.90849495 | 6.89E-105 | 2 |
| RPL39 | 0.42952273 | 1.00E-94 | 2 |
| RPL13 | 0.40884459 | 3.49E-89 | 2 |
| IL32 | 0.57857775 | 1.08E-83 | 2 |
| SIRPG | 0.63496223 | 7.89E-83 | 2 |
| RPL11 | 0.38333952 | 3.04E-82 | 2 |
| FXYD2 | -1.4082373 | 5.21E-81 | 2 |
| RPS4X | 0.39608171 | 6.27E-80 | 2 |
| RPLP2 | 0.38588962 | 6.58E-80 | 2 |
| RPS29 | 0.38810168 | 9.06E-80 | 2 |
| SMIM24 | 0.67991678 | 1.43E-78 | 2 |
| ITM2A | 0.93423345 | 6.00E-78 | 2 |
| GZMM | 0.52234513 | 7.26E-77 | 2 |
| RPS23 | 0.36747123 | 1.74E-76 | 2 |
| ANXA1 | -1.0798259 | 5.33E-76 | 2 |
| RPS15A | 0.38679561 | 9.12E-76 | 2 |
| NCF1 | 0.59657936 | 3.37E-75 | 2 |
| CD38 | 0.6412089 | 5.98E-73 | 2 |
| RPL30 | 0.38408514 | 1.03E-72 | 2 |
| RPS24 | 0.36957931 | 1.25E-72 | 2 |
| CD1E | -0.9504977 | 3.08E-71 | 2 |
| RPL26 | 0.38890837 | 5.13E-71 | 2 |
| RPL32 | 0.37140373 | 4.88E-70 | 2 |
| RPS18 | 0.37901466 | 7.81E-70 | 2 |
| RPS28 | 0.34418912 | 2.26E-69 | 2 |
| RPL10 | 0.39067686 | 2.33E-68 | 2 |
| RPL12 | 0.4203139 | 3.15E-68 | 2 |
| COTL1 | 0.55685816 | 4.07E-68 | 2 |
| RPS3 | 0.35318776 | 9.72E-68 | 2 |
| RPL35A | 0.37215779 | 2.88E-67 | 2 |
| RPS8 | 0.37278685 | 4.07E-67 | 2 |
| RPS12 | 0.40417364 | 2.70E-65 | 2 |
| RPS27A | 0.34803428 | 1.80E-64 | 2 |
| RPL13A | 0.34591453 | 2.43E-64 | 2 |

| | | | |
|---|---|---|---|
| RPL19 | 0.35152401 | 1.24E-61 | 2 |
| RPL18 | 0.3458424 | 7.95E-61 | 2 |
| EEF1A1 | 0.3139265 | 1.32E-60 | 2 |
| RPL41 | 0.32261282 | 2.52E-59 | 2 |
| RPL9 | 0.33649332 | 8.08E-58 | 2 |
| RPS19 | 0.32762827 | 2.58E-57 | 2 |
| CD99 | -0.6757296 | 3.89E-57 | 2 |
| RPL7A | 0.32454835 | 4.53E-57 | 2 |
| RPS25 | 0.34301434 | 1.11E-56 | 2 |
| TPT1 | 0.3164675 | 1.20E-54 | 2 |
| AL138899.1 | -0.8893005 | 4.52E-52 | 2 |
| RPL28 | 0.30258908 | 3.26E-51 | 2 |
| RPS13 | 0.34269952 | 9.89E-51 | 2 |
| RPS3A | 0.29650866 | 6.00E-50 | 2 |
| TUBA1B | -1.1264609 | 7.98E-50 | 2 |
| EVL | 0.38152512 | 8.40E-50 | 2 |
| CRIP1 | -1.333558 | 3.13E-48 | 2 |
| ETS1 | 0.397833 | 3.33E-48 | 2 |
| MACROD2 | -0.7509296 | 5.54E-48 | 2 |
| FAU | 0.30179227 | 2.62E-47 | 2 |
| CD247 | 0.47886058 | 3.85E-47 | 2 |
| RPL36 | 0.32381765 | 6.93E-47 | 2 |
| RPL3 | 0.29115778 | 8.12E-47 | 2 |
| ABLIM1 | 0.48388789 | 5.93E-46 | 2 |
| P2RX5 | 0.4342328 | 2.14E-45 | 2 |
| VIM | -0.787608 | 2.83E-45 | 2 |
| RPL27A | 0.34458533 | 7.74E-45 | 2 |
| RPL17 | 0.31178068 | 1.69E-44 | 2 |
| PFDN5 | 0.36058521 | 7.84E-43 | 2 |
| RPL15 | 0.28695423 | 8.15E-43 | 2 |
| CD1A | -0.858293 | 3.67E-42 | 2 |
| UBA52 | 0.29403888 | 1.18E-41 | 2 |
| GYPC | 0.39700197 | 1.31E-41 | 2 |
| RPL29 | 0.30400549 | 3.17E-41 | 2 |
| ACVR1B | 0.37227809 | 5.28E-41 | 2 |
| IKZF2 | 0.47981556 | 8.35E-41 | 2 |
| ACTN1 | 0.39463142 | 2.31E-40 | 2 |
| HCST | 0.36745989 | 2.55E-40 | 2 |
| ABCB1 | 0.33580189 | 6.22E-39 | 2 |
| RPS14 | 0.29059172 | 9.03E-39 | 2 |
| RPS11 | 0.29344129 | 1.40E-38 | 2 |
| TFDP2 | -0.7749857 | 1.55E-38 | 2 |
| VASP | 0.40564203 | 2.12E-38 | 2 |

| | | | |
|---|---|---|---|
| CD1B | -0.6554339 | 3.36E-38 | 2 |
| LCP2 | 0.40025595 | 5.93E-38 | 2 |
| NEGR1 | -0.6586998 | 1.18E-37 | 2 |
| HMGB2 | -1.1261628 | 8.08E-37 | 2 |
| IL17RB | -0.8680681 | 1.26E-36 | 2 |
| PRKCH | 0.40982867 | 3.65E-36 | 2 |
| RPL38 | 0.28197513 | 4.48E-36 | 2 |
| CD8A | 0.46066432 | 6.40E-36 | 2 |
| CD63 | 0.40736601 | 1.72E-35 | 2 |
| AP005482.1 | -0.5876095 | 4.11E-35 | 2 |
| NACA | 0.28540331 | 4.48E-35 | 2 |
| CHDH | -0.6156514 | 5.70E-35 | 2 |
| ETV5 | -0.6053817 | 5.89E-35 | 2 |
| RPL31 | 0.31221702 | 6.63E-35 | 2 |
| PFN1 | 0.28919753 | 7.54E-35 | 2 |
| CTSW | 0.48375035 | 7.56E-35 | 2 |
| AL365440.2 | -0.5314732 | 8.57E-35 | 2 |
| PABPC1 | 0.32868182 | 9.75E-35 | 2 |
| RPL27 | 0.28676825 | 3.32E-34 | 2 |
| CDK6 | -0.722416 | 4.33E-34 | 2 |
| JAK1 | 0.37488588 | 5.90E-34 | 2 |
| C9orf16 | 0.35302096 | 1.03E-33 | 2 |
| CD27 | 0.36972164 | 1.47E-33 | 2 |
| ZYX | 0.38208218 | 3.91E-33 | 2 |
| QSOX1 | 0.34349941 | 4.08E-33 | 2 |
| RPL24 | 0.27491503 | 4.52E-33 | 2 |
| RPS10 | 0.31205128 | 4.95E-33 | 2 |
| STK17A | 0.43133969 | 7.76E-33 | 2 |
| FTL | 0.40740495 | 7.91E-33 | 2 |
| LIME1 | 0.35139697 | 1.03E-32 | 2 |
| CXCR3 | 0.29240679 | 4.18E-32 | 2 |
| LGALS1 | -1.2011885 | 1.29E-31 | 2 |
| CD7 | 0.38717653 | 1.39E-31 | 2 |
| ATP5F1E | 0.26826122 | 1.89E-31 | 2 |
| XBP1 | -0.5958771 | 9.63E-31 | 2 |
| KLF2 | -0.7712529 | 1.75E-30 | 2 |
| RPL22 | 0.26237033 | 2.59E-30 | 2 |
| ACTB | 0.25270573 | 8.14E-30 | 2 |
| RACK1 | 0.26345541 | 9.62E-30 | 2 |
| IL16 | 0.3684432 | 1.98E-29 | 2 |
| SCRIB | 0.40348439 | 2.14E-29 | 2 |
| RUNX3 | 0.32338405 | 2.19E-29 | 2 |
| RAC2 | 0.30093795 | 2.94E-29 | 2 |

| | | | |
|---|---|---|---|
| HELLS | -0.6283777 | 3.09E-29 | 2 |
| AC144521.1 | 0.30522925 | 4.60E-29 | 2 |
| RGS3 | 0.31855974 | 6.25E-29 | 2 |
| PNRC1 | 0.31390493 | 1.30E-28 | 2 |
| LST1 | -0.6328104 | 1.54E-28 | 2 |
| RAB33A | 0.27448496 | 2.71E-28 | 2 |
| ELF1 | 0.33805622 | 4.96E-28 | 2 |
| RUNX1 | -0.4932863 | 6.60E-28 | 2 |
| SELL | -0.4543477 | 9.39E-28 | 2 |
| LY9 | 0.27061276 | 2.90E-27 | 2 |
| HSP90AA1 | -0.419574 | 6.37E-27 | 2 |
| ANXA2 | -0.3969384 | 7.04E-27 | 2 |
| TYMS | -0.7801469 | 7.17E-27 | 2 |
| JAML | 0.28718711 | 1.45E-26 | 2 |
| MACF1 | 0.30383569 | 1.47E-26 | 2 |
| TRAC | 0.34931297 | 2.59E-26 | 2 |
| SYNE2 | -0.6047165 | 5.21E-26 | 2 |
| CD79B | 0.29421178 | 5.67E-26 | 2 |
| EMP3 | -0.6319222 | 1.17E-25 | 2 |
| EEF1G | 0.26731353 | 7.45E-25 | 2 |
| SHISA2 | 0.36428527 | 9.20E-25 | 2 |
| EZR | -0.4888272 | 1.58E-24 | 2 |
| E2F2 | -0.3222567 | 3.90E-24 | 2 |
| DOCK10 | 0.31877973 | 2.08E-23 | 2 |
| DEK | -0.3899666 | 6.42E-23 | 2 |
| WAKMAR2 | -0.6028048 | 7.71E-23 | 2 |
| C12orf57 | 0.34515169 | 1.23E-22 | 2 |
| SESN3 | 0.2966063 | 2.94E-22 | 2 |
| KLF6 | -0.5228859 | 6.69E-22 | 2 |
| TUBB | -0.7211035 | 7.03E-22 | 2 |
| ARPP21 | -0.4254332 | 7.06E-22 | 2 |
| ITGB2 | 0.29646516 | 7.32E-22 | 2 |
| SYNE1 | 0.29814746 | 1.05E-21 | 2 |
| RIPOR2 | -0.498674 | 1.78E-21 | 2 |
| TMSB10 | -0.5327676 | 2.14E-21 | 2 |
| ARL4C | -0.3647273 | 2.87E-21 | 2 |
| CNN2 | 0.35367256 | 8.29E-21 | 2 |
| IRF2BP2 | 0.3769447 | 1.62E-20 | 2 |
| PPP1R18 | 0.2922618 | 2.36E-20 | 2 |
| TAGLN2 | -0.5379021 | 2.38E-20 | 2 |
| CD44 | 0.35349025 | 3.84E-20 | 2 |
| MZB1 | -0.5328003 | 7.50E-20 | 2 |
| AQP3 | -0.446216 | 1.31E-19 | 2 |

| | | | |
|---|---|---|---|
| HNRNPU | -0.3238643 | 1.67E-19 | 2 |
| IGFLR1 | 0.26065252 | 2.01E-19 | 2 |
| COMMD6 | 0.27625368 | 3.80E-19 | 2 |
| WDR34 | -0.3098755 | 1.06E-18 | 2 |
| PDCD4 | 0.31555679 | 1.34E-18 | 2 |
| SIVA1 | -0.5161117 | 1.43E-18 | 2 |
| SRGN | 0.36912852 | 1.87E-18 | 2 |
| FYN | 0.25945019 | 2.24E-18 | 2 |
| PCLAF | -0.7186481 | 2.26E-18 | 2 |
| HNRNPA2B1 | -0.3111508 | 2.32E-18 | 2 |
| FAM102A | 0.26686103 | 2.53E-18 | 2 |
| SRSF2 | -0.358628 | 3.10E-18 | 2 |
| RNASEH2B | -0.3671316 | 3.12E-18 | 2 |
| DBN1 | 0.27982855 | 3.98E-18 | 2 |
| TXN | -0.4517749 | 4.90E-18 | 2 |
| CAPN2 | -0.2904317 | 5.06E-18 | 2 |
| COX6C | 0.28199774 | 9.77E-18 | 2 |
| PLAC8 | -0.5550505 | 1.00E-17 | 2 |
| AC093673.1 | 0.25786761 | 1.52E-17 | 2 |
| MXRA7 | 0.25555729 | 2.65E-17 | 2 |
| TMEM173 | -0.2685575 | 3.13E-17 | 2 |
| STMN1 | -0.5320586 | 3.82E-17 | 2 |
| MCM4 | -0.3555574 | 4.13E-17 | 2 |
| SRSF10 | -0.3689378 | 7.04E-17 | 2 |
| ST6GAL1 | 0.25960026 | 7.38E-17 | 2 |
| NCAPG2 | -0.417037 | 1.03E-16 | 2 |
| ARGLU1 | -0.3612703 | 1.04E-16 | 2 |
| LINC00342 | -0.4521846 | 1.24E-16 | 2 |
| HNRNPM | -0.3452531 | 1.27E-16 | 2 |
| ICAM3 | 0.28016443 | 1.54E-16 | 2 |
| CLSPN | -0.4064699 | 2.05E-16 | 2 |
| SPINT2 | 0.26543351 | 2.36E-16 | 2 |
| AHNAK | -0.725983 | 2.49E-16 | 2 |
| SEMA4D | 0.29737785 | 5.25E-16 | 2 |
| UHRF1 | -0.3315604 | 5.49E-16 | 2 |
| FAM111B | -0.2696543 | 9.87E-16 | 2 |
| PLEKHG1 | -0.4260334 | 1.75E-15 | 2 |
| OXNAD1 | 0.26451155 | 1.84E-15 | 2 |
| RPS6KA3 | 0.30655877 | 2.52E-15 | 2 |
| ATAD5 | -0.3095338 | 2.57E-15 | 2 |
| MCM2 | -0.2501593 | 2.92E-15 | 2 |
| ZFP36 | 0.28446644 | 2.97E-15 | 2 |
| JUN | -0.4648729 | 3.75E-15 | 2 |

| | | | |
|---|---|---|---|
| ARHGEF6 | 0.25448287 | 1.11E-14 | 2 |
| WDR76 | -0.3118477 | 3.59E-14 | 2 |
| SSBP2 | -0.3557191 | 4.10E-14 | 2 |
| CD37 | 0.25829999 | 5.04E-14 | 2 |
| BCL7A | -0.2928907 | 6.24E-14 | 2 |
| RRM1 | -0.319187 | 6.35E-14 | 2 |
| VAMP5 | 0.25373719 | 6.68E-14 | 2 |
| IRF2BPL | -0.2744671 | 8.47E-14 | 2 |
| S100A4 | -0.8126814 | 1.10E-13 | 2 |
| CDK5RAP3 | -0.3683139 | 1.12E-13 | 2 |
| BRCA1 | -0.2870785 | 1.54E-13 | 2 |
| SRSF7 | -0.3040937 | 2.32E-13 | 2 |
| GAS5 | 0.28175748 | 2.66E-13 | 2 |
| CBX5 | -0.3838489 | 2.98E-13 | 2 |
| LITAF | -0.2946674 | 4.82E-13 | 2 |
| LDHA | -0.3640859 | 7.30E-13 | 2 |
| NASP | -0.3676423 | 7.77E-13 | 2 |
| PLP2 | -0.292831 | 1.94E-12 | 2 |
| NCL | -0.2928495 | 3.17E-12 | 2 |
| TNNT3 | -0.3044994 | 3.29E-12 | 2 |
| RTN4 | -0.2908917 | 3.45E-12 | 2 |
| HNRNPA3 | -0.263999 | 3.81E-12 | 2 |
| C21orf58 | -0.313557 | 3.97E-12 | 2 |
| DNMT1 | -0.3312931 | 4.41E-12 | 2 |
| ZWINT | -0.2681229 | 4.63E-12 | 2 |
| TCF12 | -0.3073242 | 5.79E-12 | 2 |
| EZH2 | -0.4180879 | 5.85E-12 | 2 |
| EIF3A | -0.2905463 | 6.21E-12 | 2 |
| FKBP5 | -0.3421643 | 7.91E-12 | 2 |
| IL7R | -0.2940486 | 8.32E-12 | 2 |
| GLUL | -0.3368086 | 1.80E-11 | 2 |
| SRSF3 | -0.2889847 | 2.02E-11 | 2 |
| HNRNPAB | -0.3362877 | 2.49E-11 | 2 |
| CAPG | -0.4084293 | 2.64E-11 | 2 |
| TMPO | -0.3928209 | 3.36E-11 | 2 |
| NUSAP1 | -0.6243772 | 3.67E-11 | 2 |
| ITGA6 | -0.3390596 | 6.91E-11 | 2 |
| DUT | -0.4986792 | 7.62E-11 | 2 |
| BAZ1B | -0.2906416 | 7.72E-11 | 2 |
| ARMH1 | 0.2756838 | 8.38E-11 | 2 |
| GMNN | -0.2557281 | 8.49E-11 | 2 |
| RTKN2 | -0.3106804 | 9.48E-11 | 2 |
| CCR9 | 0.30611425 | 1.51E-10 | 2 |

| | | | |
|---|---|---|---|
| GIHCG | -0.5162871 | 1.56E-10 | 2 |
| YWHAQ | -0.2796973 | 1.91E-10 | 2 |
| DNTT | -0.3127571 | 3.01E-10 | 2 |
| HMGB1 | -0.425295 | 3.27E-10 | 2 |
| MYOM2 | -0.3116662 | 3.28E-10 | 2 |
| ADA | -0.3045338 | 3.35E-10 | 2 |
| NPNT | -0.3016846 | 3.36E-10 | 2 |
| H2AFY | -0.3117057 | 3.72E-10 | 2 |
| GTSE1 | -0.3767251 | 5.18E-10 | 2 |
| YBX1 | -0.2942296 | 6.16E-10 | 2 |
| SH2D1A | 0.32849423 | 6.49E-10 | 2 |
| PRMT7 | -0.4092844 | 7.64E-10 | 2 |
| CENPF | -0.7383779 | 8.49E-10 | 2 |
| USP1 | -0.3473409 | 8.50E-10 | 2 |
| HNRNPR | -0.257009 | 9.12E-10 | 2 |
| CLIC3 | -0.3995072 | 9.50E-10 | 2 |
| LMNB1 | -0.3381146 | 1.20E-09 | 2 |
| MCM7 | -0.4214093 | 1.29E-09 | 2 |
| PRKDC | -0.2971267 | 1.35E-09 | 2 |
| S100A6 | -0.5412284 | 1.40E-09 | 2 |
| PCNA | -0.496317 | 1.64E-09 | 2 |
| HSPA1B | -0.3862003 | 2.82E-09 | 2 |
| YWHAE | -0.2703957 | 4.27E-09 | 2 |
| SFPQ | -0.2685293 | 7.21E-09 | 2 |
| H2AFZ | -0.5289753 | 7.30E-09 | 2 |
| TSC22D3 | -0.3560769 | 8.33E-09 | 2 |
| TMEM106C | -0.2538793 | 8.56E-09 | 2 |
| RAN | -0.3613218 | 9.54E-09 | 2 |
| RBBP8 | -0.281311 | 1.10E-08 | 2 |
| NME2 | -0.3590708 | 1.52E-08 | 2 |
| ELOVL4 | -0.3649924 | 1.69E-08 | 2 |
| NAP1L1 | -0.2538219 | 1.96E-08 | 2 |
| ANP32B | -0.2932273 | 2.01E-08 | 2 |
| ITGB1 | -0.3182818 | 2.05E-08 | 2 |
| S100A11 | -0.3911988 | 2.31E-08 | 2 |
| KLRK1 | -0.2922295 | 2.40E-08 | 2 |
| SMC1A | -0.3119585 | 2.46E-08 | 2 |
| NOSIP | -0.5990966 | 2.47E-08 | 2 |
| HMGN2 | -0.4522798 | 4.20E-08 | 2 |
| SLBP | -0.296295 | 4.24E-08 | 2 |
| MCM3 | -0.3241373 | 4.61E-08 | 2 |
| SNRPB | -0.2614693 | 6.44E-08 | 2 |
| MSH6 | -0.2552037 | 1.29E-07 | 2 |

| | | | |
|---|---|---|---|
| RBMX | -0.2670393 | 1.38E-07 | 2 |
| GALNT2 | -0.3164312 | 1.53E-07 | 2 |
| DLEU2 | -0.3113601 | 2.10E-07 | 2 |
| ADD3 | -0.432213 | 2.85E-07 | 2 |
| THEMIS | -0.2894115 | 3.92E-07 | 2 |
| RRM2 | -0.3679008 | 8.51E-07 | 2 |
| SNRPF | -0.2933676 | 9.69E-07 | 2 |
| MT-ND1 | -0.2541608 | 1.26E-06 | 2 |
| SNRPG | -0.2564632 | 1.32E-06 | 2 |
| AUTS2 | -0.2530087 | 1.39E-06 | 2 |
| ENO1 | -0.3175691 | 1.41E-06 | 2 |
| HSPD1 | -0.3378704 | 1.42E-06 | 2 |
| MCM5 | -0.2586549 | 1.53E-06 | 2 |
| HSPA1A | -0.3117997 | 2.64E-06 | 2 |
| FOXP1 | -0.2948543 | 3.58E-06 | 2 |
| GALNT6 | -0.2965275 | 4.22E-06 | 2 |
| NSD2 | -0.2658345 | 5.22E-06 | 2 |
| ZNF280D | -0.2687928 | 1.12E-05 | 2 |
| SCN3A | -0.2752001 | 1.43E-05 | 2 |
| HSP90AB1 | -0.3652366 | 1.56E-05 | 2 |
| HIST1H4C | -1.1272148 | 1.89E-05 | 2 |
| SMC3 | -0.2565726 | 2.40E-05 | 2 |
| CDKN2D | -0.2599696 | 2.71E-05 | 2 |
| ATAD2 | -0.3605608 | 5.83E-05 | 2 |
| TSPAN7 | -0.2570267 | 7.70E-05 | 2 |
| ANKRD36C | -0.2597437 | 9.92E-05 | 2 |
| GSTP1 | -0.294272 | 0.00011851 | 2 |
| MAL | -0.3354896 | 0.00013265 | 2 |
| SCAI | -0.2639658 | 0.00016244 | 2 |
| UBE2S | -0.3495331 | 0.00025637 | 2 |
| AC068587.4 | -0.3022916 | 0.00026394 | 2 |
| IDH2 | -0.2752954 | 0.00029768 | 2 |
| TOP2A | -0.7163161 | 0.00031055 | 2 |
| CDK1 | -0.2599904 | 0.00033077 | 2 |
| NUCKS1 | -0.3161697 | 0.00044122 | 2 |
| GTF3C5 | -0.2686584 | 0.00047904 | 2 |
| UBE2C | -0.2789805 | 0.00053611 | 2 |
| CALR | -0.2867588 | 0.00171921 | 2 |
| ANP32E | -0.2908072 | 0.00227532 | 2 |
| HIST1H3B | -0.3295369 | 0.00454206 | 2 |
| NEAT1 | -0.2874587 | 0.00478137 | 2 |
| KNL1 | -0.2616601 | 0.00950456 | 2 |
| ASPM | -0.5113926 | 0.01059588 | 2 |

| | | | |
|---|---|---|---|
| CENPE | -0.2574715 | 0.01198847 | 2 |
| RANBP1 | -0.2501094 | 0.01377234 | 2 |
| MKI67 | -0.6840599 | 0.01401616 | 2 |
| HSP90B1 | -0.3129286 | 0.02500399 | 2 |
| SMC2 | -0.311398 | 0.02720931 | 2 |
| TPX2 | -0.2659847 | 0.05817787 | 2 |
| PTTG1 | -0.2839591 | 0.08208012 | 2 |
| SELENOW | -0.2528384 | 0.16429003 | 2 |
| TUBB4B | -0.3324768 | 0.77278087 | 2 |
| BTG2 | -0.2753513 | 0.83166268 | 2 |
| XIST | -0.3003857 | 0.99336254 | 2 |
| H2AFX | -0.2682636 | 1 | 2 |
| LINC01222 | -0.2570462 | 1 | 2 |
| TECR | -0.3787922 | 1 | 2 |
| GIHCG | 1.25380787 | 2.37E-178 | 3 |
| TFDP2 | 1.18108585 | 7.74E-155 | 3 |
| CDK6 | 1.382242 | 7.48E-152 | 3 |
| SMC4 | -1.5912113 | 8.80E-151 | 3 |
| B2M | -1.164922 | 4.15E-139 | 3 |
| RPLP0 | 0.88711052 | 1.16E-136 | 3 |
| GNA15 | 0.56544665 | 8.65E-136 | 3 |
| HNRNPA1 | 0.79070414 | 4.41E-133 | 3 |
| STMN1 | 0.83946539 | 8.12E-133 | 3 |
| LINC02694 | 0.39011368 | 5.64E-127 | 3 |
| CAPG | 0.88517481 | 1.40E-118 | 3 |
| CCDC26 | 0.50833438 | 1.70E-117 | 3 |
| GINS2 | 0.5659394 | 8.67E-117 | 3 |
| PAICS | 0.73427457 | 1.26E-116 | 3 |
| TYMS | 0.85779604 | 6.96E-115 | 3 |
| GAPDH | 0.882596 | 2.85E-112 | 3 |
| NKAIN4 | 0.63427303 | 1.19E-111 | 3 |
| DNTT | 0.83920228 | 6.99E-110 | 3 |
| NOTCH1 | 0.68568855 | 4.96E-108 | 3 |
| PTPRC | -0.8022596 | 7.63E-108 | 3 |
| ETS1 | -0.8756204 | 2.38E-106 | 3 |
| GSTP1 | 0.87062011 | 5.06E-104 | 3 |
| ETV5 | 0.89871034 | 1.67E-103 | 3 |
| MCM4 | 0.69288618 | 2.09E-103 | 3 |
| NME1 | 0.67365516 | 7.62E-103 | 3 |
| PPP1R14B | 0.93952603 | 7.87E-103 | 3 |
| SIRPG | -1.036554 | 3.93E-102 | 3 |
| ZNF683 | -1.4512374 | 1.73E-101 | 3 |
| RPLP1 | 0.56743107 | 3.47E-100 | 3 |

| | | | |
|---|---|---|---|
| AC002454.1 | 0.77128299 | 4.61E-100 | 3 |
| NPM1 | 0.72557902 | 6.83E-100 | 3 |
| CTHRC1 | 0.50854184 | 2.13E-99 | 3 |
| PTMA | 0.52211987 | 9.74E-98 | 3 |
| FXYD2 | 1.4664723 | 2.61E-97 | 3 |
| CD1B | 0.80580437 | 1.83E-95 | 3 |
| TXN | 0.82967117 | 3.80E-93 | 3 |
| CD1A | 0.9737693 | 1.04E-92 | 3 |
| CD27 | -0.8636012 | 1.37E-91 | 3 |
| CD1E | 0.7870784 | 3.01E-91 | 3 |
| BCL7A | 0.58780251 | 6.16E-91 | 3 |
| SNHG29 | 0.7608492 | 3.32E-90 | 3 |
| GALNT2 | 0.74735057 | 4.00E-90 | 3 |
| TXNIP | -1.0741288 | 7.89E-90 | 3 |
| MCM2 | 0.53527394 | 4.16E-89 | 3 |
| AC027601.6 | 0.41131633 | 1.69E-88 | 3 |
| HSP90AB1 | 1.0547814 | 4.07E-88 | 3 |
| LTB | -1.2512916 | 4.83E-88 | 3 |
| TUBB | 0.69747138 | 6.55E-88 | 3 |
| UHRF1 | 0.63715008 | 7.97E-88 | 3 |
| NAP1L1 | 0.66084491 | 4.87E-87 | 3 |
| QPRT | 0.45115191 | 2.42E-86 | 3 |
| TUBA1B | 0.84220477 | 3.60E-86 | 3 |
| PNRC1 | -0.8357874 | 3.82E-86 | 3 |
| SATB1 | -1.3370349 | 4.68E-86 | 3 |
| HLA-E | -0.8373653 | 5.86E-86 | 3 |
| BCL11A | 0.54421653 | 3.59E-84 | 3 |
| UNG | 0.606707 | 4.07E-84 | 3 |
| ARPP21 | 0.63743336 | 5.05E-84 | 3 |
| CD52 | -0.8604528 | 5.27E-84 | 3 |
| MCM5 | 0.69833291 | 3.41E-83 | 3 |
| GLUL | 0.67544881 | 8.23E-82 | 3 |
| HSPA1B | 0.65348644 | 5.74E-81 | 3 |
| KLHL23 | 0.48922824 | 5.02E-78 | 3 |
| RUNX1 | 0.75098354 | 5.05E-78 | 3 |
| SCN3A | 0.73657762 | 6.04E-78 | 3 |
| UST | 0.40275561 | 1.27E-77 | 3 |
| LDHA | 0.74262539 | 2.16E-77 | 3 |
| IDH2 | 0.63854441 | 2.23E-77 | 3 |
| MALAT1 | -0.6329447 | 3.65E-77 | 3 |
| SNHG25 | 0.43440143 | 5.51E-77 | 3 |
| MYC | 0.57734199 | 2.31E-76 | 3 |
| RAN | 0.76352374 | 4.90E-76 | 3 |

| | | | |
|---|---|---|---|
| HELLS | 0.73294596 | 7.55E-76 | 3 |
| CDK5RAP3 | 0.68100833 | 7.87E-76 | 3 |
| AEBP1 | 0.53664045 | 9.12E-76 | 3 |
| VAT1 | 0.50079751 | 1.74E-75 | 3 |
| MCM7 | 0.85893471 | 1.83E-75 | 3 |
| AL365440.2 | 0.58811685 | 4.12E-75 | 3 |
| DIP2A | -0.8323709 | 5.02E-75 | 3 |
| HSPD1 | 0.80685734 | 5.85E-75 | 3 |
| NME2 | 0.78331108 | 1.07E-74 | 3 |
| PCNA | 0.81327781 | 1.50E-74 | 3 |
| PCLAF | 0.75921652 | 1.58E-74 | 3 |
| ANXA1 | 0.71705561 | 1.67E-74 | 3 |
| ITGB2 | -0.7640182 | 7.24E-74 | 3 |
| SIVA1 | 0.82073612 | 8.02E-74 | 3 |
| BTG1 | -0.874734 | 2.02E-73 | 3 |
| PPA1 | 0.66826754 | 6.04E-73 | 3 |
| NOP53 | 0.61894407 | 1.49E-72 | 3 |
| TDRD9 | 0.3893352 | 3.52E-72 | 3 |
| VPREB1 | 0.34813141 | 3.97E-72 | 3 |
| PPIA | 0.58908445 | 5.00E-72 | 3 |
| HLA-B | -1.0416564 | 1.04E-70 | 3 |
| RPS5 | 0.51679471 | 1.92E-70 | 3 |
| FABP5 | 0.74571374 | 4.58E-70 | 3 |
| HCST | -0.7639742 | 7.51E-69 | 3 |
| PLCH1 | 0.33235861 | 2.45E-68 | 3 |
| SNRPF | 0.71767746 | 4.05E-68 | 3 |
| HLA-A | -0.8952951 | 5.00E-68 | 3 |
| MSH6 | 0.58407333 | 1.01E-67 | 3 |
| MCM3 | 0.66562621 | 1.18E-67 | 3 |
| FADS2 | 0.34350131 | 1.23E-67 | 3 |
| PPIF | 0.32176525 | 1.91E-67 | 3 |
| YBX3 | 0.52294831 | 3.43E-67 | 3 |
| HLA-C | -0.8320562 | 5.82E-67 | 3 |
| FIRRE | 0.29738314 | 6.90E-66 | 3 |
| RPSA | 0.42393013 | 8.73E-66 | 3 |
| HS6ST1 | 0.34042325 | 1.32E-64 | 3 |
| CD8B | -0.6869709 | 1.35E-64 | 3 |
| C1QBP | 0.68965288 | 4.57E-64 | 3 |
| CDK2AP1 | 0.53815297 | 5.99E-64 | 3 |
| WDR34 | 0.42101505 | 9.66E-64 | 3 |
| MIF | 0.73859382 | 1.55E-63 | 3 |
| CDT1 | 0.41155434 | 2.84E-63 | 3 |
| CLEC2D | -0.9315296 | 7.41E-63 | 3 |

| | | | |
|---|---|---|---|
| SNRPD1 | 0.61375395 | 8.87E-63 | 3 |
| E2F1 | 0.38715976 | 1.24E-62 | 3 |
| ENO1 | 0.7876823 | 2.57E-62 | 3 |
| RPL35 | 0.42578157 | 4.43E-62 | 3 |
| SELENOW | 0.78722502 | 2.06E-61 | 3 |
| DOCK10 | -0.7252499 | 2.42E-61 | 3 |
| CD79A | -0.8496895 | 2.95E-61 | 3 |
| RPL14 | 0.47027658 | 1.25E-60 | 3 |
| C19orf48 | 0.45649973 | 1.31E-60 | 3 |
| ANP32B | 0.51921063 | 1.70E-60 | 3 |
| TSPAN7 | 0.70941655 | 2.34E-60 | 3 |
| CD99 | 0.71167906 | 5.20E-60 | 3 |
| PGD | 0.35593759 | 7.17E-60 | 3 |
| IGLL1 | 0.67734732 | 9.62E-60 | 3 |
| DTL | 0.31517533 | 4.20E-59 | 3 |
| ENOSF1 | 0.5270174 | 4.40E-59 | 3 |
| NASP | 0.63588866 | 6.79E-59 | 3 |
| DMC1 | 0.28634721 | 9.71E-59 | 3 |
| APEX1 | 0.55861813 | 1.32E-58 | 3 |
| ZFP36L1 | -0.6710574 | 1.51E-58 | 3 |
| SLFN5 | -0.7195994 | 1.71E-58 | 3 |
| AHCY | 0.31142267 | 1.99E-58 | 3 |
| LIME1 | -0.6459088 | 2.99E-58 | 3 |
| UCK2 | 0.2995922 | 5.90E-58 | 3 |
| SNRPE | 0.592357 | 6.93E-58 | 3 |
| CRNDE | 0.48120425 | 1.73E-57 | 3 |
| TRIM28 | 0.53478306 | 1.91E-57 | 3 |
| CHAF1A | 0.36746103 | 7.58E-57 | 3 |
| H2AFZ | 0.57360862 | 9.26E-57 | 3 |
| ZFP36L2 | -0.7480432 | 1.87E-56 | 3 |
| VIM | 0.65744538 | 2.91E-56 | 3 |
| SSRP1 | 0.49870205 | 3.50E-55 | 3 |
| CLSPN | 0.46743526 | 1.17E-54 | 3 |
| VANGL1 | 0.32706991 | 1.36E-54 | 3 |
| SLC25A3 | 0.54737219 | 4.18E-54 | 3 |
| RPL36A | 0.43569718 | 1.55E-53 | 3 |
| EXOSC5 | 0.321621 | 1.90E-53 | 3 |
| DCTPP1 | 0.44701589 | 2.05E-53 | 3 |
| SLIT1 | 0.40160773 | 2.05E-53 | 3 |
| RPS17 | 0.51009018 | 4.16E-53 | 3 |
| LINC02384 | 0.29862898 | 5.33E-53 | 3 |
| MCM6 | 0.55296786 | 1.02E-52 | 3 |
| GMNN | 0.44993858 | 1.22E-52 | 3 |

| | | | |
|---|---|---|---|
| TPO | 0.32702396 | 1.88E-52 | 3 |
| VDAC1 | 0.53517675 | 3.06E-52 | 3 |
| CDCA7 | 0.53560394 | 5.35E-52 | 3 |
| SAMD3 | -0.6893064 | 6.29E-52 | 3 |
| RANBP1 | 0.60225398 | 1.50E-51 | 3 |
| FAM111B | 0.35289836 | 3.65E-51 | 3 |
| TMSB15A | 0.31656793 | 5.21E-51 | 3 |
| MACF1 | -0.684518 | 7.07E-51 | 3 |
| AL138899.1 | 0.51978521 | 8.78E-51 | 3 |
| CDK4 | 0.46372743 | 1.10E-50 | 3 |
| TKT | 0.56168597 | 1.69E-50 | 3 |
| SARAF | -0.5808539 | 1.79E-50 | 3 |
| FTH1 | 0.42656742 | 3.23E-50 | 3 |
| TIPIN | 0.30620402 | 4.28E-50 | 3 |
| CHCHD2 | 0.48692868 | 4.85E-50 | 3 |
| HMGA1 | 0.59508861 | 1.07E-49 | 3 |
| TIMM13 | 0.5242867 | 3.26E-49 | 3 |
| SRM | 0.50742388 | 3.83E-49 | 3 |
| JAK1 | -0.5930106 | 3.90E-49 | 3 |
| CARHSP1 | 0.4972634 | 4.50E-49 | 3 |
| STK4 | -0.6059589 | 5.45E-49 | 3 |
| MRTO4 | 0.31854373 | 1.03E-48 | 3 |
| SLC25A5 | 0.50124407 | 1.24E-48 | 3 |
| YBX1 | 0.54641768 | 1.42E-48 | 3 |
| LYL1 | 0.45113495 | 1.73E-48 | 3 |
| TALDO1 | 0.49428147 | 1.89E-48 | 3 |
| PDCD5 | 0.47048064 | 2.75E-48 | 3 |
| RPS2 | 0.39432156 | 2.76E-48 | 3 |
| ODC1 | 0.34171957 | 3.09E-48 | 3 |
| SNRPA | 0.41416791 | 3.69E-48 | 3 |
| IMPDH2 | 0.42571514 | 1.09E-47 | 3 |
| DUT | 0.72352295 | 1.14E-47 | 3 |
| EEF1G | 0.40673263 | 1.17E-47 | 3 |
| MRPL57 | 0.48098821 | 1.42E-47 | 3 |
| FEN1 | 0.36069058 | 2.35E-47 | 3 |
| HINT1 | 0.43221878 | 2.80E-47 | 3 |
| RPA3 | 0.33464223 | 2.87E-47 | 3 |
| TAGLN2 | 0.52571052 | 3.70E-47 | 3 |
| CBX5 | 0.54750694 | 3.72E-47 | 3 |
| SNHG19 | 0.35196356 | 4.21E-47 | 3 |
| COPRS | 0.2546613 | 4.43E-47 | 3 |
| CALR | 0.7538376 | 6.08E-47 | 3 |
| SUB1 | 0.47094544 | 8.82E-47 | 3 |

| | | | |
|---|---|---|---|
| LDHB | 0.5054543 | 1.01E-46 | 3 |
| CD59 | 0.34027925 | 1.03E-46 | 3 |
| HSPE1 | 0.62750089 | 2.46E-46 | 3 |
| FADS1 | 0.36090683 | 3.42E-46 | 3 |
| MZB1 | 0.61824409 | 3.56E-46 | 3 |
| EIF4A1 | 0.5646297 | 3.70E-46 | 3 |
| HSP90AA1 | 0.4479063 | 3.85E-46 | 3 |
| EVL | -0.4942364 | 6.97E-46 | 3 |
| PSMA6 | 0.53813286 | 1.26E-45 | 3 |
| CHML | 0.41571026 | 1.31E-45 | 3 |
| CHEK1 | 0.35339405 | 1.53E-45 | 3 |
| SNRPC | 0.43405914 | 1.97E-45 | 3 |
| RPS6 | 0.34899558 | 2.00E-45 | 3 |
| SMS | 0.42596151 | 3.02E-45 | 3 |
| CD8A | -0.6756191 | 5.92E-45 | 3 |
| ATP5MC1 | 0.45165775 | 6.30E-45 | 3 |
| FKBP3 | 0.44358476 | 7.02E-45 | 3 |
| PXMP2 | 0.3451315 | 8.95E-45 | 3 |
| ACAP1 | -0.5335766 | 9.21E-45 | 3 |
| ST18 | 0.27974402 | 1.80E-44 | 3 |
| CCT2 | 0.4868463 | 4.34E-44 | 3 |
| ILF2 | 0.47299103 | 4.36E-44 | 3 |
| DLEU7 | 0.37037783 | 4.40E-44 | 3 |
| GADD45GIP1 | 0.43828455 | 5.21E-44 | 3 |
| MTHFD1 | 0.34908141 | 7.90E-44 | 3 |
| NT5DC2 | 0.35488521 | 9.56E-44 | 3 |
| CBFA2T3 | 0.46594183 | 1.06E-43 | 3 |
| SEM1 | 0.43680166 | 1.08E-43 | 3 |
| ADA | 0.536061 | 1.18E-43 | 3 |
| POLR2L | 0.50320245 | 1.24E-43 | 3 |
| HNRNPM | 0.51158214 | 1.58E-43 | 3 |
| HMGB3 | 0.29957261 | 2.13E-43 | 3 |
| TEX30 | 0.26735814 | 4.34E-43 | 3 |
| POLD2 | 0.26646593 | 4.79E-43 | 3 |
| UQCRH | 0.42890961 | 5.93E-43 | 3 |
| RPS12 | 0.38646046 | 7.08E-43 | 3 |
| IPO5 | 0.38482389 | 9.35E-43 | 3 |
| MYL6B | 0.40745274 | 1.05E-42 | 3 |
| MYO7B | 0.40872834 | 1.57E-42 | 3 |
| FAM102A | -0.5934336 | 1.68E-42 | 3 |
| PHGDH | 0.46331137 | 1.91E-42 | 3 |
| AKR1B1 | 0.42418465 | 3.57E-42 | 3 |
| HNRNPA1P48 | 0.32501903 | 5.22E-42 | 3 |

| | | | |
|---|---|---|---|
| RPL4 | 0.3792398 | 6.01E-42 | 3 |
| HILPDA | 0.27604307 | 9.16E-42 | 3 |
| FBL | 0.46641573 | 1.75E-41 | 3 |
| RUVBL2 | 0.28469375 | 3.00E-41 | 3 |
| HNRNPD | 0.47268812 | 3.52E-41 | 3 |
| COX5A | 0.48493302 | 3.58E-41 | 3 |
| HSPA1A | 0.50628994 | 3.65E-41 | 3 |
| PALM2-AKAP2 | 0.4619793 | 3.87E-41 | 3 |
| SRSF2 | 0.48141072 | 4.14E-41 | 3 |
| HNRNPAB | 0.55456799 | 4.52E-41 | 3 |
| STK17B | -0.5746593 | 5.17E-41 | 3 |
| RSRP1 | -0.5026223 | 6.01E-41 | 3 |
| SNRPB | 0.47299236 | 1.05E-40 | 3 |
| SRSF7 | 0.47137627 | 1.16E-40 | 3 |
| MRPL12 | 0.303042 | 1.18E-40 | 3 |
| ATP5F1B | 0.47634332 | 1.55E-40 | 3 |
| ZWINT | 0.33572862 | 1.58E-40 | 3 |
| CENPH | 0.2870913 | 2.37E-40 | 3 |
| NETO2 | 0.366259 | 4.13E-40 | 3 |
| LGALS1 | 1.1492189 | 4.63E-40 | 3 |
| MRPL11 | 0.38223353 | 4.63E-40 | 3 |
| SNHG8 | 0.46923749 | 5.77E-40 | 3 |
| EEF1A1 | 0.29181806 | 5.77E-40 | 3 |
| SLIRP | 0.38824661 | 8.28E-40 | 3 |
| POP7 | 0.3189112 | 1.12E-39 | 3 |
| CENPX | 0.39848182 | 1.17E-39 | 3 |
| PSMG1 | 0.28270872 | 1.65E-39 | 3 |
| ABLIM1 | -0.6443836 | 1.69E-39 | 3 |
| H2AFV | 0.4435841 | 3.13E-39 | 3 |
| CD2 | -0.5234851 | 3.94E-39 | 3 |
| PCAT18 | 0.34236065 | 4.91E-39 | 3 |
| YWHAE | 0.47459643 | 5.72E-39 | 3 |
| TCEAL9 | 0.30917279 | 6.64E-39 | 3 |
| IL16 | -0.5753749 | 8.17E-39 | 3 |
| SNHG3 | 0.42602164 | 8.63E-39 | 3 |
| SPOCK2 | -0.5691288 | 9.25E-39 | 3 |
| N4BP2L2 | -0.5281606 | 1.23E-38 | 3 |
| SLC1A5 | 0.28340229 | 1.40E-38 | 3 |
| PDE7A | -0.6493962 | 1.60E-38 | 3 |
| PRMT2 | -0.5647339 | 1.74E-38 | 3 |
| ATP5F1A | 0.477446 | 2.04E-38 | 3 |
| SREBF2 | 0.27170032 | 2.10E-38 | 3 |
| RPL27A | 0.32344235 | 5.28E-38 | 3 |

| | | | |
|---|---|---|---|
| COX7C | 0.34111898 | 5.42E-38 | 3 |
| MZT2B | 0.38757509 | 5.91E-38 | 3 |
| COTL1 | -0.5562989 | 7.44E-38 | 3 |
| MTHFD2 | 0.33243595 | 7.97E-38 | 3 |
| BOP1 | 0.30725781 | 8.63E-38 | 3 |
| GCSH | 0.3069595 | 8.87E-38 | 3 |
| MRPL14 | 0.35426537 | 9.79E-38 | 3 |
| GLO1 | 0.33305533 | 1.07E-37 | 3 |
| EIF5A | 0.47515767 | 1.40E-37 | 3 |
| NSG1 | 0.28486068 | 1.96E-37 | 3 |
| E2F2 | 0.32848724 | 1.97E-37 | 3 |
| EIF4EBP1 | 0.25802827 | 2.04E-37 | 3 |
| BMP2K | 0.30242995 | 2.18E-37 | 3 |
| TCTEX1D2 | 0.30436094 | 3.47E-37 | 3 |
| TCL1A | 0.68111491 | 3.58E-37 | 3 |
| BTF3 | 0.37292666 | 4.02E-37 | 3 |
| EBNA1BP2 | 0.32126815 | 4.22E-37 | 3 |
| EEF1B2 | 0.38113611 | 4.37E-37 | 3 |
| PRXL2A | 0.27173327 | 4.41E-37 | 3 |
| COX7A2 | 0.40643317 | 4.54E-37 | 3 |
| PRPF19 | 0.32194637 | 5.97E-37 | 3 |
| SRSF3 | 0.45200638 | 6.72E-37 | 3 |
| NTM | 0.31804555 | 9.27E-37 | 3 |
| ZNF423 | 0.29090627 | 9.79E-37 | 3 |
| DANCR | 0.40235458 | 1.84E-36 | 3 |
| BAHCC1 | 0.44794884 | 2.81E-36 | 3 |
| NHP2 | 0.45445536 | 3.04E-36 | 3 |
| PA2G4 | 0.50508551 | 3.14E-36 | 3 |
| TFRC | 0.38616681 | 3.45E-36 | 3 |
| PMAIP1 | 0.26501301 | 4.39E-36 | 3 |
| NDUFS5 | 0.43313357 | 4.42E-36 | 3 |
| HEMGN | 0.35545174 | 4.56E-36 | 3 |
| WDR76 | 0.36947406 | 4.80E-36 | 3 |
| LSM7 | 0.44328694 | 5.00E-36 | 3 |
| CD44 | -0.5874174 | 5.45E-36 | 3 |
| SRGN | -0.6083526 | 8.02E-36 | 3 |
| RRM1 | 0.34843902 | 1.10E-35 | 3 |
| ITGB2-AS1 | -0.4924435 | 1.32E-35 | 3 |
| LMNB1 | 0.39243864 | 1.38E-35 | 3 |
| DDX5 | -0.4143201 | 2.01E-35 | 3 |
| SLC25A6 | 0.45799279 | 2.18E-35 | 3 |
| CTSW | -0.7152538 | 2.21E-35 | 3 |
| CENPU | 0.35281535 | 2.23E-35 | 3 |

| | | | |
|---|---|---|---|
| PRDX4 | 0.26624081 | 2.60E-35 | 3 |
| CNN2 | -0.6855867 | 3.82E-35 | 3 |
| ATM | -0.5454563 | 4.22E-35 | 3 |
| RAP1GAP | 0.30302444 | 4.35E-35 | 3 |
| RAC2 | -0.4311074 | 4.96E-35 | 3 |
| YWHAG | 0.36102196 | 6.36E-35 | 3 |
| SNRPD2 | 0.41199021 | 1.02E-34 | 3 |
| MT2A | 0.28940509 | 1.03E-34 | 3 |
| MRPL15 | 0.27457441 | 1.39E-34 | 3 |
| TNNT3 | 0.39390899 | 1.57E-34 | 3 |
| SLBP | 0.42543153 | 1.68E-34 | 3 |
| CDPF1 | 0.33703821 | 2.22E-34 | 3 |
| RPS16 | 0.29432114 | 2.39E-34 | 3 |
| NDUFAB1 | 0.39581319 | 2.69E-34 | 3 |
| ARPC1B | -0.4473018 | 2.76E-34 | 3 |
| SNHG7 | 0.36045526 | 3.02E-34 | 3 |
| CCT7 | 0.42402072 | 5.33E-34 | 3 |
| ERH | 0.41778004 | 5.71E-34 | 3 |
| ACOT7 | 0.25082359 | 7.06E-34 | 3 |
| CD320 | 0.28059935 | 8.22E-34 | 3 |
| PRMT1 | 0.4387663 | 9.61E-34 | 3 |
| GIMAP7 | -0.6329426 | 1.05E-33 | 3 |
| PRMT7 | 0.47681681 | 1.22E-33 | 3 |
| IGFBP2 | 0.41738962 | 1.35E-33 | 3 |
| ATP5PF | 0.40746793 | 1.57E-33 | 3 |
| RBBP8 | 0.37312434 | 1.75E-33 | 3 |
| MBP | -0.5745037 | 2.45E-33 | 3 |
| SNHG15 | 0.28874259 | 2.76E-33 | 3 |
| HNRNPDL | 0.41015696 | 3.34E-33 | 3 |
| SERBP1 | 0.44317629 | 3.54E-33 | 3 |
| IKZF2 | -0.6600864 | 3.71E-33 | 3 |
| FH | 0.27793158 | 7.35E-33 | 3 |
| MARCKSL1 | 0.48828796 | 1.11E-32 | 3 |
| HHIP | 0.34849097 | 3.10E-32 | 3 |
| ALYREF | 0.36444588 | 3.23E-32 | 3 |
| SLFN11 | 0.25923215 | 3.62E-32 | 3 |
| CCT8 | 0.3800616 | 4.01E-32 | 3 |
| PDXP | 0.25019726 | 4.11E-32 | 3 |
| WEE1 | 0.33857797 | 4.59E-32 | 3 |
| CHST11 | 0.30687304 | 5.05E-32 | 3 |
| FGFR1 | 0.51037883 | 6.82E-32 | 3 |
| POLR2I | 0.36693695 | 7.38E-32 | 3 |
| PDCD4 | -0.5071061 | 8.03E-32 | 3 |

| | | | |
|---|---|---|---|
| HMGN2 | 0.38146942 | 8.32E-32 | 3 |
| GNL3 | 0.32490925 | 9.91E-32 | 3 |
| ADGRG1 | 0.26249035 | 1.07E-31 | 3 |
| BLK | 0.34128766 | 1.25E-31 | 3 |
| TOMM40 | 0.32363409 | 1.46E-31 | 3 |
| UQCRQ | 0.43058194 | 2.34E-31 | 3 |
| RCAN1 | 0.27860737 | 2.85E-31 | 3 |
| UQCC2 | 0.3213964 | 2.92E-31 | 3 |
| DDX17 | -0.3826573 | 3.13E-31 | 3 |
| CYBA | -0.4471513 | 3.50E-31 | 3 |
| NDUFA6 | 0.35485486 | 3.78E-31 | 3 |
| PSMB6 | 0.35388376 | 3.86E-31 | 3 |
| CTPS1 | 0.3247292 | 4.42E-31 | 3 |
| COL6A2 | -0.5406019 | 4.69E-31 | 3 |
| PSMB2 | 0.34535506 | 5.54E-31 | 3 |
| MYB | 0.42776702 | 5.85E-31 | 3 |
| MRPL37 | 0.27944165 | 6.08E-31 | 3 |
| RPL5 | 0.31731017 | 6.88E-31 | 3 |
| RPL18A | 0.29429349 | 7.31E-31 | 3 |
| PNISR | -0.4766651 | 9.15E-31 | 3 |
| CCT5 | 0.43103442 | 1.08E-30 | 3 |
| S1PR3 | 0.29769678 | 1.31E-30 | 3 |
| NXT1 | 0.27275893 | 1.33E-30 | 3 |
| TBCA | 0.39813532 | 1.34E-30 | 3 |
| ATP5PD | 0.39810755 | 3.21E-30 | 3 |
| NUCKS1 | 0.37213409 | 3.25E-30 | 3 |
| MRPL16 | 0.32028565 | 3.26E-30 | 3 |
| ELOVL5 | 0.37665157 | 3.91E-30 | 3 |
| CUEDC2 | 0.28548452 | 3.95E-30 | 3 |
| MAL | 0.65567173 | 4.27E-30 | 3 |
| SNHG6 | 0.33025814 | 4.46E-30 | 3 |
| EZH2 | 0.31509827 | 4.61E-30 | 3 |
| ARHGEF1 | -0.4667222 | 5.53E-30 | 3 |
| CLEC11A | 0.39058831 | 5.72E-30 | 3 |
| PSMA4 | 0.39222914 | 8.20E-30 | 3 |
| TRAF3IP3 | -0.4675821 | 9.44E-30 | 3 |
| CHMP7 | -0.548755 | 9.87E-30 | 3 |
| ESD | 0.34617843 | 1.28E-29 | 3 |
| ATP5MC2 | 0.33021584 | 2.11E-29 | 3 |
| PEBP1 | 0.41900132 | 2.21E-29 | 3 |
| ANXA2 | 0.35458249 | 3.00E-29 | 3 |
| TOX2 | -0.577602 | 3.46E-29 | 3 |
| NCF1 | -0.5373949 | 4.23E-29 | 3 |

| | | | |
|---|---|---|---|
| PRKCH | -0.4883502 | 5.52E-29 | 3 |
| RBX1 | 0.36495633 | 5.82E-29 | 3 |
| TOP1MT | 0.34275562 | 1.28E-28 | 3 |
| PSMB7 | 0.34215925 | 2.07E-28 | 3 |
| LBH | -0.4954264 | 2.22E-28 | 3 |
| MRPS15 | 0.30601015 | 2.30E-28 | 3 |
| AC245060.5 | 0.45697429 | 2.32E-28 | 3 |
| CIRBP | -0.4238847 | 2.78E-28 | 3 |
| DNMT1 | 0.41186684 | 3.20E-28 | 3 |
| SNRPG | 0.40902645 | 3.66E-28 | 3 |
| BTG2 | -0.5740997 | 4.43E-28 | 3 |
| TRBC2 | -0.3428129 | 6.05E-28 | 3 |
| NDUFS8 | 0.33720924 | 7.25E-28 | 3 |
| RPS6KA3 | -0.5129341 | 7.30E-28 | 3 |
| GLRX3 | 0.30829792 | 7.95E-28 | 3 |
| FAT1 | 0.31938818 | 9.48E-28 | 3 |
| PABPC4 | 0.32502168 | 1.14E-27 | 3 |
| MCMBP | 0.31685845 | 1.19E-27 | 3 |
| PGAM1 | 0.32903656 | 1.24E-27 | 3 |
| ILF3 | 0.38016979 | 1.28E-27 | 3 |
| HPF1 | 0.31714037 | 1.38E-27 | 3 |
| YWHAQ | 0.37141167 | 1.57E-27 | 3 |
| TBCD | 0.31331956 | 2.13E-27 | 3 |
| NDUFS3 | 0.31179545 | 2.29E-27 | 3 |
| SET | 0.30176279 | 2.46E-27 | 3 |
| ATAD5 | 0.30018543 | 2.84E-27 | 3 |
| SCD | 0.3236627 | 2.90E-27 | 3 |
| NCL | 0.44094091 | 2.94E-27 | 3 |
| SCRIB | -0.522807 | 2.94E-27 | 3 |
| ATP5MC3 | 0.3449789 | 3.03E-27 | 3 |
| COX7B | 0.39647048 | 3.26E-27 | 3 |
| PRKDC | 0.42181536 | 3.31E-27 | 3 |
| IL2RG | -0.4209625 | 4.37E-27 | 3 |
| CZIB | 0.2738405 | 4.51E-27 | 3 |
| SNHG1 | 0.30165659 | 5.78E-27 | 3 |
| NAA50 | 0.30046885 | 5.90E-27 | 3 |
| ATIC | 0.3104334 | 6.57E-27 | 3 |
| TXNRD1 | 0.30601779 | 7.22E-27 | 3 |
| RFC2 | 0.33052303 | 7.37E-27 | 3 |
| VDAC3 | 0.29036905 | 8.42E-27 | 3 |
| GPATCH4 | 0.28146828 | 9.78E-27 | 3 |
| XRCC2 | 0.28164113 | 1.22E-26 | 3 |
| SUPT16H | 0.39271368 | 1.29E-26 | 3 |

| | | | |
|---|---|---|---|
| TMA16 | 0.31236966 | 1.48E-26 | 3 |
| COPS3 | 0.30234807 | 1.67E-26 | 3 |
| MLLT11 | 0.28653456 | 2.04E-26 | 3 |
| LYAR | 0.25890106 | 2.86E-26 | 3 |
| GNAS | 0.27743231 | 2.94E-26 | 3 |
| HMGB1 | 0.27424077 | 2.94E-26 | 3 |
| GOT2 | 0.27297256 | 3.14E-26 | 3 |
| SSBP1 | 0.37493221 | 3.21E-26 | 3 |
| POMP | 0.35617164 | 3.57E-26 | 3 |
| ATP1B3 | 0.33479842 | 4.64E-26 | 3 |
| NDUFAF8 | 0.34146704 | 4.78E-26 | 3 |
| ACAT2 | 0.29473524 | 5.12E-26 | 3 |
| PMVK | 0.25159571 | 9.38E-26 | 3 |
| PIK3IP1 | -0.3947117 | 9.98E-26 | 3 |
| RPS11 | 0.2577397 | 1.11E-25 | 3 |
| NOLC1 | 0.37299673 | 1.13E-25 | 3 |
| HDAC2 | 0.34745883 | 1.40E-25 | 3 |
| NOP58 | 0.39371142 | 1.56E-25 | 3 |
| SRI | 0.41344517 | 1.60E-25 | 3 |
| NOP56 | 0.40403039 | 1.79E-25 | 3 |
| MDH2 | 0.3794667 | 2.30E-25 | 3 |
| LSM4 | 0.39194672 | 2.34E-25 | 3 |
| PSMD7 | 0.31225065 | 2.58E-25 | 3 |
| ELF1 | -0.4115964 | 2.80E-25 | 3 |
| TMEM106C | 0.29544179 | 2.98E-25 | 3 |
| SMC2 | 0.3463356 | 3.00E-25 | 3 |
| AAK1 | -0.5222365 | 3.47E-25 | 3 |
| CNOT6 | 0.27693026 | 3.78E-25 | 3 |
| PRDX1 | 0.4429975 | 3.88E-25 | 3 |
| MRPS24 | 0.25051047 | 4.33E-25 | 3 |
| MAGOHB | 0.25362734 | 4.45E-25 | 3 |
| MAT2A | 0.40955787 | 6.26E-25 | 3 |
| FMC1 | 0.28509916 | 7.80E-25 | 3 |
| PLP2 | 0.29645292 | 9.57E-25 | 3 |
| CELF2 | -0.455861 | 1.00E-24 | 3 |
| SNX5 | 0.34666741 | 1.00E-24 | 3 |
| FARSA | 0.26379384 | 1.12E-24 | 3 |
| MT-ND3 | -0.3505174 | 1.15E-24 | 3 |
| RPL23 | 0.33601939 | 1.25E-24 | 3 |
| PSMA2 | 0.30998373 | 1.80E-24 | 3 |
| GZMM | -0.4398353 | 1.90E-24 | 3 |
| NCR3 | -0.4545239 | 2.11E-24 | 3 |
| USP20 | 0.27916415 | 2.15E-24 | 3 |

| | | | |
|---|---|---|---|
| EEF1E1 | 0.2667306 | 2.30E-24 | 3 |
| DAZAP1 | 0.32656968 | 2.50E-24 | 3 |
| H2AFY | 0.33773353 | 2.96E-24 | 3 |
| SELL | 0.33787708 | 3.51E-24 | 3 |
| RBM3 | 0.39489847 | 3.93E-24 | 3 |
| TRAC | -0.4150584 | 4.11E-24 | 3 |
| ID3 | -0.6933939 | 4.70E-24 | 3 |
| DBI | 0.35571027 | 5.20E-24 | 3 |
| DEK | 0.31640358 | 5.51E-24 | 3 |
| NDUFA11 | 0.34216927 | 5.80E-24 | 3 |
| CENPV | 0.37330994 | 6.50E-24 | 3 |
| IQGAP2 | 0.44773529 | 6.61E-24 | 3 |
| HSPA9 | 0.3103803 | 6.72E-24 | 3 |
| SNU13 | 0.32609559 | 7.03E-24 | 3 |
| RNF213 | -0.4594509 | 8.45E-24 | 3 |
| SNRPA1 | 0.32635826 | 8.72E-24 | 3 |
| STOML2 | 0.29075711 | 1.05E-23 | 3 |
| MRPL23 | 0.25800833 | 1.30E-23 | 3 |
| LCP2 | -0.4187404 | 1.79E-23 | 3 |
| NDUFA4 | 0.32705038 | 2.16E-23 | 3 |
| GMPS | 0.28449533 | 2.19E-23 | 3 |
| GPX4 | 0.33214174 | 2.27E-23 | 3 |
| YEATS4 | 0.28558416 | 2.30E-23 | 3 |
| SNHG32 | 0.35230249 | 2.86E-23 | 3 |
| PSMB1 | 0.3398272 | 3.03E-23 | 3 |
| AP005482.1 | 0.35583631 | 3.05E-23 | 3 |
| PELI2 | 0.25279637 | 3.07E-23 | 3 |
| CYCS | 0.27314313 | 3.13E-23 | 3 |
| NDUFB6 | 0.33087941 | 3.62E-23 | 3 |
| LAGE3 | 0.29909182 | 3.87E-23 | 3 |
| NDUFS6 | 0.37285965 | 4.39E-23 | 3 |
| MRPL20 | 0.33899146 | 5.36E-23 | 3 |
| DDX39A | 0.3341761 | 5.54E-23 | 3 |
| FDPS | 0.3833064 | 6.48E-23 | 3 |
| PSMA3 | 0.28893463 | 7.82E-23 | 3 |
| ACTG1 | 0.2767295 | 8.20E-23 | 3 |
| RRP15 | 0.27291173 | 8.48E-23 | 3 |
| HHIP-AS1 | 0.38516767 | 1.10E-22 | 3 |
| PHF5A | 0.27947509 | 1.30E-22 | 3 |
| GTPBP4 | 0.27220177 | 1.46E-22 | 3 |
| JAML | -0.3408444 | 2.01E-22 | 3 |
| NUDT1 | 0.25118168 | 2.22E-22 | 3 |
| ATP5MF | 0.31918961 | 2.25E-22 | 3 |

| | | | |
|---|---|---|---|
| TIMM10 | 0.28131641 | 2.38E-22 | 3 |
| SYNRG | -0.4282415 | 2.61E-22 | 3 |
| SYNE1 | -0.4580546 | 2.65E-22 | 3 |
| FAM49B | -0.38217 | 3.12E-22 | 3 |
| HCFC1 | 0.28141069 | 3.45E-22 | 3 |
| PRDX2 | 0.42311286 | 3.91E-22 | 3 |
| POLE3 | 0.31380545 | 4.12E-22 | 3 |
| BAZ1B | 0.3537848 | 4.22E-22 | 3 |
| HNRNPR | 0.33197162 | 4.33E-22 | 3 |
| SEPTIN6 | 0.29602066 | 4.41E-22 | 3 |
| HACD3 | 0.28134823 | 4.46E-22 | 3 |
| LRRC75A | 0.30961608 | 4.86E-22 | 3 |
| SOD1 | 0.38569433 | 4.87E-22 | 3 |
| LRPPRC | 0.28338949 | 5.26E-22 | 3 |
| UQCRC1 | 0.26915942 | 5.55E-22 | 3 |
| ACTN1 | -0.4147386 | 5.59E-22 | 3 |
| MDH1 | 0.31317516 | 6.11E-22 | 3 |
| KLRB1 | -0.4214285 | 6.36E-22 | 3 |
| SUMO2 | 0.26865147 | 6.65E-22 | 3 |
| CXCR3 | -0.3027665 | 8.20E-22 | 3 |
| POLR2E | 0.29061728 | 8.28E-22 | 3 |
| FCMR | -0.4232087 | 8.49E-22 | 3 |
| SMC1A | 0.37927581 | 9.87E-22 | 3 |
| AKNA | -0.414559 | 1.02E-21 | 3 |
| EIF3B | 0.26206412 | 1.12E-21 | 3 |
| HDGFL3 | 0.27036319 | 1.15E-21 | 3 |
| ANAPC15 | 0.26641049 | 1.18E-21 | 3 |
| RPL26L1 | 0.25382036 | 1.37E-21 | 3 |
| ATP5MPL | 0.32062351 | 1.39E-21 | 3 |
| UMODL1 | 0.31775225 | 1.51E-21 | 3 |
| KNOP1 | 0.25179531 | 1.52E-21 | 3 |
| MRPS26 | 0.27735174 | 1.77E-21 | 3 |
| LDLRAD4 | 0.34427427 | 2.01E-21 | 3 |
| RABL6 | 0.26826659 | 2.05E-21 | 3 |
| SINHCAF | 0.27536513 | 2.16E-21 | 3 |
| GRHPR | 0.2792248 | 2.44E-21 | 3 |
| UBE2H | -0.4002691 | 3.03E-21 | 3 |
| ZNRF1 | 0.31356262 | 3.18E-21 | 3 |
| KLF13 | -0.4219032 | 3.41E-21 | 3 |
| HSBP1 | 0.27395572 | 4.52E-21 | 3 |
| RTRAF | 0.32527059 | 4.92E-21 | 3 |
| SSBP2 | 0.33776422 | 5.25E-21 | 3 |
| SMARCB1 | 0.28524077 | 6.79E-21 | 3 |

| | | | |
|---|---|---|---|
| IPO7 | 0.27576108 | 7.43E-21 | 3 |
| SH3BGRL3 | -0.4112617 | 1.00E-20 | 3 |
| FUNDC2 | 0.27372795 | 1.03E-20 | 3 |
| CHCHD3 | 0.27884852 | 1.19E-20 | 3 |
| ST6GAL1 | -0.4054725 | 1.29E-20 | 3 |
| SF3B5 | 0.31484231 | 1.35E-20 | 3 |
| SMARCA4 | 0.30026752 | 1.35E-20 | 3 |
| HINT2 | 0.28807414 | 1.41E-20 | 3 |
| PIK3R1 | -0.3842744 | 1.59E-20 | 3 |
| JPT1 | 0.29408159 | 1.65E-20 | 3 |
| PLIN2 | 0.3958244 | 1.79E-20 | 3 |
| MRPL52 | 0.28153009 | 1.84E-20 | 3 |
| GALNT6 | 0.28814601 | 2.11E-20 | 3 |
| DKC1 | 0.30696101 | 2.28E-20 | 3 |
| HADH | 0.26378718 | 2.30E-20 | 3 |
| NDUFB2 | 0.3210983 | 2.36E-20 | 3 |
| LSM5 | 0.25596666 | 2.61E-20 | 3 |
| HNRNPC | 0.31968482 | 2.67E-20 | 3 |
| MAZ | 0.3156122 | 3.23E-20 | 3 |
| MCUR1 | 0.25384066 | 3.60E-20 | 3 |
| MAGOH | 0.32345256 | 4.00E-20 | 3 |
| AP2S1 | 0.27937927 | 4.23E-20 | 3 |
| PAFAH1B3 | 0.26581909 | 4.26E-20 | 3 |
| ATOX1 | 0.27728735 | 4.43E-20 | 3 |
| DPYSL2 | 0.27493878 | 4.89E-20 | 3 |
| NUDC | 0.32047582 | 5.46E-20 | 3 |
| TTN | -0.5017827 | 6.64E-20 | 3 |
| GDI2 | 0.32222965 | 7.14E-20 | 3 |
| NREP | 0.3503856 | 7.15E-20 | 3 |
| RIF1 | 0.33063264 | 7.34E-20 | 3 |
| HDGF | 0.29592274 | 8.05E-20 | 3 |
| MYO1F | -0.433961 | 9.08E-20 | 3 |
| AKR1A1 | 0.26104428 | 9.14E-20 | 3 |
| CD37 | -0.3867603 | 1.06E-19 | 3 |
| RSL1D1 | 0.29566695 | 1.07E-19 | 3 |
| ELOB | 0.30387005 | 1.23E-19 | 3 |
| ISG20 | -0.4304682 | 1.41E-19 | 3 |
| RASGRP1 | 0.41866034 | 1.47E-19 | 3 |
| ADD3 | -0.5508951 | 1.47E-19 | 3 |
| XIST | -0.4192825 | 1.50E-19 | 3 |
| CACYBP | 0.3273504 | 1.61E-19 | 3 |
| USP1 | 0.32937197 | 1.68E-19 | 3 |
| GIMAP1 | -0.3419724 | 1.70E-19 | 3 |

| | | | |
|---|---|---|---|
| UQCRFS1 | 0.28928364 | 2.27E-19 | 3 |
| NDUFA13 | 0.2768004 | 2.36E-19 | 3 |
| BCL9L | -0.3301446 | 2.46E-19 | 3 |
| SUN2 | -0.387724 | 2.51E-19 | 3 |
| ANAPC11 | 0.30933436 | 2.78E-19 | 3 |
| CDKN2A | 0.272673 | 2.96E-19 | 3 |
| MSI2 | 0.35695448 | 3.27E-19 | 3 |
| ARL6IP5 | -0.4096824 | 3.34E-19 | 3 |
| SEC61G | 0.31562696 | 3.48E-19 | 3 |
| HNRNPA2B1 | 0.27915788 | 4.92E-19 | 3 |
| CYC1 | 0.26569256 | 5.00E-19 | 3 |
| ENY2 | 0.29615388 | 5.73E-19 | 3 |
| DHX9 | 0.33009617 | 5.96E-19 | 3 |
| UBE2J1 | 0.29101776 | 6.25E-19 | 3 |
| TPI1 | 0.38034377 | 6.49E-19 | 3 |
| FOXO1 | -0.2909873 | 8.01E-19 | 3 |
| MACROD2 | -0.5525613 | 8.17E-19 | 3 |
| MBNL1 | -0.3721569 | 8.83E-19 | 3 |
| SF3B6 | 0.27021407 | 8.95E-19 | 3 |
| TNRC6C-AS1 | -0.3785487 | 9.33E-19 | 3 |
| NDUFAF3 | 0.25868425 | 1.00E-18 | 3 |
| PSMA7 | 0.31173243 | 1.02E-18 | 3 |
| CCDC85B | 0.28805292 | 1.03E-18 | 3 |
| C4orf48 | 0.28018763 | 1.16E-18 | 3 |
| KMT2E | -0.3304556 | 1.27E-18 | 3 |
| ZNF22 | 0.27810199 | 1.73E-18 | 3 |
| ATP5PO | 0.29743004 | 1.75E-18 | 3 |
| ACTR2 | -0.3377707 | 2.49E-18 | 3 |
| SND1 | 0.2888917 | 2.82E-18 | 3 |
| SEPTIN11 | 0.2853519 | 2.86E-18 | 3 |
| DCUN1D1 | 0.31477604 | 3.02E-18 | 3 |
| POLR2J3 | -0.3931997 | 3.19E-18 | 3 |
| CYB5A | 0.31485366 | 3.23E-18 | 3 |
| XRCC6 | 0.30965789 | 3.38E-18 | 3 |
| RTL6 | 0.31664305 | 3.42E-18 | 3 |
| CAP1 | -0.3653801 | 3.49E-18 | 3 |
| DYRK2 | -0.3792738 | 3.85E-18 | 3 |
| TOMM5 | 0.30990806 | 3.90E-18 | 3 |
| RNASEH2C | 0.27423882 | 5.29E-18 | 3 |
| EIF1AX | 0.30719708 | 5.97E-18 | 3 |
| PRDX3 | 0.26489584 | 6.04E-18 | 3 |
| WDR43 | 0.27122942 | 6.57E-18 | 3 |
| PDAP1 | 0.29557382 | 6.68E-18 | 3 |

| | | | |
|---|---|---|---|
| COX6B1 | 0.2695366 | 7.39E-18 | 3 |
| RBM39 | -0.3086519 | 7.56E-18 | 3 |
| TPGS2 | 0.26156978 | 7.98E-18 | 3 |
| ANP32A | 0.29638805 | 1.04E-17 | 3 |
| ABCE1 | 0.2806747 | 1.04E-17 | 3 |
| CD3G | -0.3190488 | 1.06E-17 | 3 |
| ATP5MD | 0.29339647 | 1.30E-17 | 3 |
| MLEC | 0.29673971 | 1.31E-17 | 3 |
| TMC6 | -0.3949521 | 1.43E-17 | 3 |
| HNRNPH3 | 0.29476376 | 1.47E-17 | 3 |
| PMF1 | 0.25767325 | 1.62E-17 | 3 |
| DDX21 | 0.34784646 | 1.87E-17 | 3 |
| COX8A | 0.25636479 | 1.96E-17 | 3 |
| PSMC3 | 0.28561714 | 2.20E-17 | 3 |
| MXD4 | -0.3724094 | 2.28E-17 | 3 |
| ELOC | 0.27656731 | 2.51E-17 | 3 |
| SLC38A2 | 0.25762921 | 2.56E-17 | 3 |
| ITGA6 | 0.40741755 | 2.82E-17 | 3 |
| TNRC6B | -0.4151307 | 3.08E-17 | 3 |
| KAT6B | -0.3579549 | 3.21E-17 | 3 |
| BCL11B | -0.3915718 | 4.02E-17 | 3 |
| CD226 | -0.4322225 | 4.23E-17 | 3 |
| RHOA | 0.27880861 | 4.45E-17 | 3 |
| HIST1H2AC | -0.4303468 | 4.50E-17 | 3 |
| CD247 | -0.3334991 | 4.64E-17 | 3 |
| TOPBP1 | 0.27112426 | 4.89E-17 | 3 |
| RGS3 | -0.4481254 | 5.79E-17 | 3 |
| NAA38 | 0.26837889 | 6.29E-17 | 3 |
| OXNAD1 | -0.3915486 | 6.39E-17 | 3 |
| DYNC1H1 | 0.376991 | 6.55E-17 | 3 |
| NDUFB9 | 0.27708745 | 6.71E-17 | 3 |
| MT-ATP6 | -0.2769847 | 6.80E-17 | 3 |
| TPM4 | 0.34288724 | 9.04E-17 | 3 |
| TXNDC17 | 0.27315823 | 1.06E-16 | 3 |
| SUMO1 | 0.25878741 | 1.09E-16 | 3 |
| RBBP4 | 0.28051048 | 1.12E-16 | 3 |
| BRCA1 | 0.27102181 | 1.16E-16 | 3 |
| YWHAH | 0.2907083 | 1.19E-16 | 3 |
| COPE | 0.28718852 | 1.24E-16 | 3 |
| CCDC69 | -0.3927431 | 1.24E-16 | 3 |
| PSMA1 | 0.27961771 | 1.30E-16 | 3 |
| POLR2F | 0.25875691 | 1.34E-16 | 3 |
| PTBP1 | 0.27388184 | 1.45E-16 | 3 |

| | | | |
|---|---|---|---|
| HNRNPF | 0.27879683 | 1.57E-16 | 3 |
| FTL | -0.4174269 | 1.60E-16 | 3 |
| PGK1 | 0.29090942 | 1.84E-16 | 3 |
| NRROS | 0.25847501 | 1.90E-16 | 3 |
| AMD1 | 0.27530029 | 1.93E-16 | 3 |
| RPL34 | -0.2525742 | 1.99E-16 | 3 |
| CBX3 | 0.27887626 | 2.22E-16 | 3 |
| SQLE | 0.34026383 | 2.28E-16 | 3 |
| LY9 | -0.283527 | 2.43E-16 | 3 |
| NDUFV2 | 0.27510146 | 2.45E-16 | 3 |
| PKM | 0.29166444 | 3.50E-16 | 3 |
| BAG1 | 0.25815183 | 3.51E-16 | 3 |
| CLIC3 | -0.4620886 | 3.82E-16 | 3 |
| CCND3 | 0.32839035 | 4.66E-16 | 3 |
| CLNS1A | 0.25738477 | 4.76E-16 | 3 |
| NEGR1 | 0.31332566 | 5.05E-16 | 3 |
| GSTO1 | 0.26618291 | 5.16E-16 | 3 |
| NEAT1 | -0.4600761 | 5.51E-16 | 3 |
| RCN1 | 0.25110048 | 5.53E-16 | 3 |
| SRSF9 | 0.27035575 | 5.68E-16 | 3 |
| TGOLN2 | -0.3537394 | 5.75E-16 | 3 |
| FKBP5 | 0.29168886 | 5.90E-16 | 3 |
| GLRX | 0.25802086 | 5.92E-16 | 3 |
| NOSIP | -0.6693655 | 6.09E-16 | 3 |
| NDUFB3 | 0.27002451 | 6.39E-16 | 3 |
| OLA1 | 0.26837122 | 6.71E-16 | 3 |
| CLTA | 0.25618108 | 7.20E-16 | 3 |
| RBL2 | -0.3844169 | 7.38E-16 | 3 |
| CHD2 | -0.36994 | 7.72E-16 | 3 |
| ZSCAN16-AS1 | 0.27613667 | 7.87E-16 | 3 |
| COX6A1 | 0.26538008 | 9.22E-16 | 3 |
| TMEM131L | 0.34430256 | 9.45E-16 | 3 |
| SAMHD1 | -0.4117424 | 9.53E-16 | 3 |
| DDX6 | -0.3576968 | 9.87E-16 | 3 |
| RBM8A | 0.25393605 | 9.97E-16 | 3 |
| ATP5ME | 0.29097167 | 1.11E-15 | 3 |
| TIMM8B | 0.26633372 | 1.14E-15 | 3 |
| ALOX5AP | -0.2711986 | 1.15E-15 | 3 |
| TSPAN32 | -0.275367 | 1.27E-15 | 3 |
| IL27RA | -0.3632743 | 1.35E-15 | 3 |
| CCT6A | 0.26463983 | 1.40E-15 | 3 |
| GTF3C5 | 0.25372701 | 1.46E-15 | 3 |
| METTL26 | 0.28881361 | 1.58E-15 | 3 |

| | | | |
|---|---|---|---|
| EIF3L | 0.29406814 | 1.79E-15 | 3 |
| PNRC2 | -0.358539 | 1.80E-15 | 3 |
| VDAC2 | 0.25206301 | 1.89E-15 | 3 |
| DHFR | 0.28720104 | 1.96E-15 | 3 |
| DGKZ | -0.3434341 | 1.99E-15 | 3 |
| CTSA | -0.3575435 | 2.86E-15 | 3 |
| SESN3 | -0.3715745 | 3.01E-15 | 3 |
| PHB2 | 0.26847716 | 3.13E-15 | 3 |
| PTGES3 | 0.28489266 | 3.82E-15 | 3 |
| UBE2N | 0.25622347 | 4.20E-15 | 3 |
| DCP2 | -0.3568192 | 4.24E-15 | 3 |
| AKAP13 | -0.3654187 | 4.80E-15 | 3 |
| EWSR1 | 0.25892619 | 6.47E-15 | 3 |
| RUVBL1 | 0.25052449 | 7.19E-15 | 3 |
| CYTH4 | -0.2847732 | 7.21E-15 | 3 |
| ATF4 | 0.35093555 | 7.66E-15 | 3 |
| BIN2 | -0.3748836 | 8.66E-15 | 3 |
| RORA | 0.28495114 | 1.20E-14 | 3 |
| TGFB1 | -0.3518875 | 1.23E-14 | 3 |
| AIF1 | 0.32157207 | 1.32E-14 | 3 |
| MT-ND6 | 0.28407697 | 1.38E-14 | 3 |
| MICOS10 | 0.27584883 | 1.46E-14 | 3 |
| CCT4 | 0.26265389 | 1.56E-14 | 3 |
| AC144521.1 | -0.3295839 | 1.83E-14 | 3 |
| THOC7 | 0.2535148 | 2.69E-14 | 3 |
| CAMK4 | -0.3436048 | 3.25E-14 | 3 |
| CCNG2 | -0.3315797 | 3.48E-14 | 3 |
| RSL24D1 | 0.25313787 | 4.24E-14 | 3 |
| SMC3 | 0.27917724 | 4.52E-14 | 3 |
| RBMX | 0.28187801 | 6.36E-14 | 3 |
| CFLAR | -0.389664 | 7.18E-14 | 3 |
| TUBA1A | 0.37711669 | 9.41E-14 | 3 |
| TRA2B | 0.25812009 | 9.67E-14 | 3 |
| CANX | 0.32498263 | 1.11E-13 | 3 |
| MRPS34 | 0.25879587 | 1.15E-13 | 3 |
| SMARCC1 | 0.25097644 | 1.18E-13 | 3 |
| ST3GAL1 | -0.3007885 | 1.19E-13 | 3 |
| ATAD2 | 0.28563332 | 1.41E-13 | 3 |
| CPT1A | 0.27034586 | 1.52E-13 | 3 |
| IL7R | 0.2913444 | 1.61E-13 | 3 |
| CSNK2B | 0.2572038 | 1.79E-13 | 3 |
| SPRY1 | 0.50006639 | 1.84E-13 | 3 |
| NDUFB11 | 0.26224469 | 1.92E-13 | 3 |

| | | | |
|---|---|---|---|
| CEP85L | -0.341819 | 2.70E-13 | 3 |
| PTPRA | -0.3419468 | 2.83E-13 | 3 |
| NDUFB7 | 0.25797548 | 2.87E-13 | 3 |
| SDF2L1 | 0.25093603 | 4.11E-13 | 3 |
| ACSF3 | 0.29474711 | 4.57E-13 | 3 |
| FYB1 | -0.2782517 | 5.14E-13 | 3 |
| PSMB3 | 0.25890358 | 8.89E-13 | 3 |
| CDC42SE1 | -0.3532819 | 9.56E-13 | 3 |
| JUN | 0.29568616 | 9.75E-13 | 3 |
| TCP1 | 0.250911 | 1.01E-12 | 3 |
| MT-ND2 | -0.3098711 | 1.11E-12 | 3 |
| KLRK1 | -0.3411751 | 1.27E-12 | 3 |
| STAT5A | 0.55632003 | 1.47E-12 | 3 |
| TMBIM4 | -0.2998008 | 1.69E-12 | 3 |
| CCT3 | 0.26798495 | 1.73E-12 | 3 |
| ATRX | -0.3241155 | 2.08E-12 | 3 |
| PLEC | -0.3083611 | 2.19E-12 | 3 |
| IARS | 0.259206 | 2.26E-12 | 3 |
| SEC61B | 0.28521682 | 2.58E-12 | 3 |
| ZEB1 | 0.25257219 | 2.73E-12 | 3 |
| ARL4C | -0.3077806 | 2.74E-12 | 3 |
| SAMD9 | -0.335593 | 2.90E-12 | 3 |
| SEMA4D | -0.3715266 | 2.94E-12 | 3 |
| PARK7 | 0.25317309 | 3.25E-12 | 3 |
| ZYX | -0.2963984 | 5.44E-12 | 3 |
| CD53 | -0.3099209 | 5.57E-12 | 3 |
| LSP1 | -0.3562736 | 5.64E-12 | 3 |
| GAS5 | 0.27020425 | 5.74E-12 | 3 |
| NDUFA12 | 0.25092898 | 6.07E-12 | 3 |
| HERC1 | -0.324943 | 6.33E-12 | 3 |
| TRMT112 | 0.25661066 | 6.50E-12 | 3 |
| C12orf75 | 0.28254941 | 8.88E-12 | 3 |
| SYNCRIP | 0.30158213 | 9.36E-12 | 3 |
| GIMAP4 | -0.299289 | 1.00E-11 | 3 |
| FDFT1 | 0.30417129 | 1.08E-11 | 3 |
| DGKA | -0.3497154 | 1.37E-11 | 3 |
| RAB8B | -0.3302403 | 1.40E-11 | 3 |
| TMBIM6 | -0.2714336 | 1.59E-11 | 3 |
| LINC01578 | -0.2840882 | 1.64E-11 | 3 |
| ITM2A | -0.4104144 | 1.87E-11 | 3 |
| DOK2 | -0.3217253 | 1.98E-11 | 3 |
| AL365361.1 | -0.3896455 | 2.24E-11 | 3 |
| G3BP1 | 0.26427877 | 2.61E-11 | 3 |

| | | | |
|---|---|---|---|
| TCERG1 | 0.25221714 | 3.13E-11 | 3 |
| SATB1-AS1 | -0.2782106 | 4.00E-11 | 3 |
| PLCL2 | -0.2747347 | 4.11E-11 | 3 |
| CDC42EP3 | -0.3543207 | 4.94E-11 | 3 |
| BCL6 | -0.3219442 | 5.27E-11 | 3 |
| PTPN22 | -0.3294155 | 7.56E-11 | 3 |
| TECR | -0.6042439 | 8.98E-11 | 3 |
| NFATC3 | -0.3039149 | 1.03E-10 | 3 |
| LST1 | 0.258422 | 1.10E-10 | 3 |
| LRRFIP1 | -0.2720985 | 1.10E-10 | 3 |
| ITK | -0.3151983 | 1.15E-10 | 3 |
| CD48 | -0.2797286 | 1.32E-10 | 3 |
| FYN | -0.3152503 | 1.47E-10 | 3 |
| DDAH2 | 0.29676181 | 1.72E-10 | 3 |
| TSC22D4 | -0.2987054 | 2.51E-10 | 3 |
| ACAP2 | -0.3095436 | 2.58E-10 | 3 |
| PPP1R2 | -0.3282202 | 2.73E-10 | 3 |
| YPEL3 | -0.319852 | 3.32E-10 | 3 |
| ANKRD44 | -0.336396 | 3.60E-10 | 3 |
| CREBRF | -0.3255865 | 4.91E-10 | 3 |
| LEPROTL1 | -0.322968 | 6.16E-10 | 3 |
| TNRC6C | -0.3694762 | 7.04E-10 | 3 |
| CEMIP2 | -0.2865654 | 7.44E-10 | 3 |
| PSME2 | 0.28944572 | 8.29E-10 | 3 |
| ARAP2 | -0.3314225 | 1.33E-09 | 3 |
| GBP2 | -0.2982276 | 1.61E-09 | 3 |
| MDM4 | -0.3419198 | 1.74E-09 | 3 |
| CD3E | -0.2500921 | 1.97E-09 | 3 |
| LNPEP | -0.2753099 | 2.40E-09 | 3 |
| STK10 | -0.3013763 | 2.44E-09 | 3 |
| PCSK7 | -0.3278374 | 2.56E-09 | 3 |
| PPP1R18 | -0.2805753 | 2.58E-09 | 3 |
| FARS2 | -0.3871528 | 3.40E-09 | 3 |
| C9orf16 | -0.2895165 | 3.71E-09 | 3 |
| STK17A | -0.3130889 | 5.62E-09 | 3 |
| S100A4 | 0.63503951 | 6.06E-09 | 3 |
| GPSM3 | -0.2595756 | 6.47E-09 | 3 |
| KDM5B | -0.2778794 | 7.59E-09 | 3 |
| ICAM3 | -0.2759558 | 7.83E-09 | 3 |
| PPP2R5C | -0.3246461 | 7.88E-09 | 3 |
| ATF7IP | -0.3003811 | 8.29E-09 | 3 |
| FLNA | -0.3162667 | 8.63E-09 | 3 |
| SENP7 | -0.3075892 | 9.55E-09 | 3 |

| | | | |
|---|---|---|---|
| LIMD2 | -0.2712274 | 1.06E-08 | 3 |
| HDAC7 | -0.3000318 | 1.21E-08 | 3 |
| LAPTM5 | -0.2552707 | 1.41E-08 | 3 |
| FOXP1 | -0.3444796 | 1.45E-08 | 3 |
| TMC8 | -0.2821342 | 1.59E-08 | 3 |
| HCLS1 | -0.2838782 | 1.65E-08 | 3 |
| RAPGEF6 | -0.300838 | 1.83E-08 | 3 |
| TIAM1 | -0.3215557 | 2.45E-08 | 3 |
| ZNF655 | -0.2842081 | 2.87E-08 | 3 |
| IGFLR1 | -0.268371 | 3.38E-08 | 3 |
| IGFBP5 | -0.5837887 | 3.53E-08 | 3 |
| UBE2F | 0.29160329 | 3.78E-08 | 3 |
| TRAF5 | -0.2826419 | 4.69E-08 | 3 |
| DDX3X | -0.2752016 | 6.05E-08 | 3 |
| IKZF1 | -0.2683868 | 6.99E-08 | 3 |
| TMEM161B-AS1 | -0.3671719 | 7.58E-08 | 3 |
| ITGA4 | 0.31397727 | 7.90E-08 | 3 |
| GCC2 | -0.2984612 | 8.38E-08 | 3 |
| FAM107B | -0.2547878 | 8.77E-08 | 3 |
| SLC16A7 | -0.3184257 | 9.13E-08 | 3 |
| POLD4 | -0.2638059 | 1.03E-07 | 3 |
| OGA | -0.2720867 | 1.05E-07 | 3 |
| FGD3 | -0.2596837 | 1.16E-07 | 3 |
| KIT | -0.2994409 | 1.57E-07 | 3 |
| ITGAL | -0.294413 | 1.83E-07 | 3 |
| VASP | -0.2986834 | 2.42E-07 | 3 |
| UCP2 | -0.2822915 | 3.45E-07 | 3 |
| SP100 | -0.3162931 | 3.45E-07 | 3 |
| ERICH1 | -0.2667704 | 4.99E-07 | 3 |
| MKNK2 | -0.2628082 | 5.32E-07 | 3 |
| SPINT2 | -0.2537136 | 5.77E-07 | 3 |
| ADGRE5 | -0.2726066 | 6.39E-07 | 3 |
| ZFP36 | -0.2514604 | 7.11E-07 | 3 |
| KMT2C | -0.258813 | 7.75E-07 | 3 |
| CR1 | -0.3507527 | 9.11E-07 | 3 |
| ZNF217 | -0.261697 | 9.30E-07 | 3 |
| PSMA3-AS1 | -0.2756636 | 1.11E-06 | 3 |
| CLDND1 | -0.2979085 | 1.14E-06 | 3 |
| TTC14 | -0.2766268 | 1.22E-06 | 3 |
| OGT | -0.2678954 | 1.63E-06 | 3 |
| SIPA1 | -0.2533099 | 1.82E-06 | 3 |
| LAT2 | -0.2815801 | 2.04E-06 | 3 |
| MPHOSPH8 | -0.2746523 | 2.09E-06 | 3 |

| | | | |
|---|---|---|---|
| SAMD12 | -0.2966057 | 2.91E-06 | 3 |
| ABHD17A | -0.2696979 | 3.05E-06 | 3 |
| CASK | -0.2666548 | 3.73E-06 | 3 |
| CNOT6L | -0.2878168 | 6.33E-06 | 3 |
| HIST1H1E | -0.3533567 | 7.12E-06 | 3 |
| ZNF92 | -0.2704306 | 9.02E-06 | 3 |
| ANKRD12 | -0.2826657 | 9.09E-06 | 3 |
| FTX | -0.2579782 | 9.68E-06 | 3 |
| PRKACB | -0.2638705 | 1.24E-05 | 3 |
| DNAJB14 | -0.259969 | 1.36E-05 | 3 |
| PLAC8 | -0.4785176 | 1.75E-05 | 3 |
| TACC1 | -0.2634612 | 2.81E-05 | 3 |
| CAST | -0.2615823 | 3.54E-05 | 3 |
| CARD8 | -0.2620608 | 3.61E-05 | 3 |
| SP110 | -0.2503424 | 5.37E-05 | 3 |
| ZBTB20 | -0.2641499 | 5.81E-05 | 3 |
| PILRB | -0.2810036 | 6.28E-05 | 3 |
| EPB41 | -0.2992622 | 8.76E-05 | 3 |
| DGKH | -0.2501743 | 0.00011029 | 3 |
| JPX | -0.2617013 | 0.00021612 | 3 |
| ESYT2 | -0.2644225 | 0.00025988 | 3 |
| ITSN2 | -0.2500505 | 0.0003603 | 3 |
| ZC3HAV1 | -0.2526072 | 0.00037211 | 3 |
| KLF2 | -0.5522834 | 0.00044209 | 3 |
| RIPOR2 | -0.3387288 | 0.000508 | 3 |
| HIST1H3D | -0.4828011 | 0.00191077 | 3 |
| HSPA5 | 0.30527359 | 0.00226229 | 3 |
| PMEPA1 | 0.26135838 | 0.00351588 | 3 |
| HMGCS1 | 0.30172139 | 0.00388796 | 3 |
| CCDC186 | -0.2585916 | 0.00459687 | 3 |
| RESF1 | -0.2594343 | 0.04083778 | 3 |
| ARL6IP1 | -0.2852867 | 0.0688696 | 3 |
| HIST1H1C | -0.4151106 | 1 | 3 |
| HIST1H1D | -0.3560406 | 1 | 3 |
| IFITM2 | 0.31901022 | 1 | 3 |
| MIR4422HG | 0.89378362 | 2.01E-252 | 4 |
| ELOVL4 | 1.26131828 | 3.72E-161 | 4 |
| LST1 | 1.29627325 | 1.01E-141 | 4 |
| CCR6 | 0.37980325 | 2.00E-129 | 4 |
| AL138899.1 | 1.28979937 | 7.01E-125 | 4 |
| KIT | 0.96520884 | 8.07E-117 | 4 |
| CD1E | 0.97412083 | 4.68E-115 | 4 |
| RORC | 0.52333321 | 1.05E-112 | 4 |

| | | | |
|---|---|---|---|
| RPL13 | -0.6390598 | 4.89E-102 | 4 |
| RPS18 | -0.6430211 | 4.60E-100 | 4 |
| CD1A | 0.9699536 | 4.71E-97 | 4 |
| IL32 | -0.9429914 | 9.69E-94 | 4 |
| RPS27 | -0.6701644 | 3.53E-90 | 4 |
| RPL10 | -0.6580017 | 6.31E-89 | 4 |
| SLA | 0.90264975 | 1.15E-88 | 4 |
| RPL41 | -0.5713309 | 7.92E-88 | 4 |
| IL17RB | 0.95773356 | 1.32E-87 | 4 |
| RPL13A | -0.5598234 | 6.03E-85 | 4 |
| RPL19 | -0.5851051 | 4.92E-84 | 4 |
| FLNB | 0.57069978 | 2.90E-82 | 4 |
| NPNT | 0.74783811 | 9.00E-82 | 4 |
| RPS14 | -0.5881059 | 3.09E-80 | 4 |
| RPL12 | -0.6608217 | 3.85E-80 | 4 |
| RPS8 | -0.5571872 | 1.10E-79 | 4 |
| PLPP3 | 0.31048092 | 2.53E-78 | 4 |
| CD1B | 0.76198002 | 1.17E-75 | 4 |
| PLXNA4 | 0.39573308 | 1.19E-74 | 4 |
| FTL | -0.8433928 | 1.34E-71 | 4 |
| AQP3 | 0.89776904 | 1.88E-71 | 4 |
| PLCB1 | 0.53874796 | 1.37E-70 | 4 |
| RPLP2 | -0.4951816 | 1.43E-70 | 4 |
| EML4 | 0.76386946 | 5.48E-70 | 4 |
| AL365440.2 | 0.73912739 | 1.13E-69 | 4 |
| RPS2 | -0.5589181 | 1.19E-69 | 4 |
| HCST | -0.8053673 | 1.26E-69 | 4 |
| RPL7A | -0.5092852 | 5.01E-69 | 4 |
| EEF1A1 | -0.4930331 | 8.52E-69 | 4 |
| RPL26 | -0.5420931 | 1.36E-67 | 4 |
| RPS3 | -0.4994825 | 6.48E-67 | 4 |
| RPS4X | -0.5169628 | 3.92E-66 | 4 |
| CNN2 | -0.93151 | 1.18E-64 | 4 |
| RPL37 | -0.5015551 | 8.42E-64 | 4 |
| RPL11 | -0.4784389 | 1.05E-63 | 4 |
| RPL39 | -0.5068749 | 5.23E-63 | 4 |
| RPL32 | -0.4991328 | 9.57E-63 | 4 |
| RPS27A | -0.4972174 | 1.13E-61 | 4 |
| RPL18 | -0.4777159 | 1.89E-61 | 4 |
| PLEKHG1 | 0.76802489 | 2.44E-61 | 4 |
| MBP | -0.7811154 | 2.82E-60 | 4 |
| RPS25 | -0.4704044 | 3.70E-60 | 4 |
| XIST | 0.75036706 | 2.83E-59 | 4 |

| | | | |
|---|---|---|---|
| ARSG | 0.58973211 | 3.34E-59 | 4 |
| BCAR3 | 0.39014806 | 3.67E-59 | 4 |
| RPS29 | -0.4569113 | 4.95E-59 | 4 |
| RPS12 | -0.5416509 | 9.97E-59 | 4 |
| AC097518.2 | 0.34837208 | 2.12E-58 | 4 |
| RPS24 | -0.4545601 | 3.43E-58 | 4 |
| RPS23 | -0.4491758 | 3.86E-58 | 4 |
| RAG1 | 0.31555397 | 7.24E-57 | 4 |
| RPS28 | -0.4384032 | 2.40E-56 | 4 |
| AP005482.1 | 0.71976283 | 2.85E-56 | 4 |
| RPL10A | -0.4840869 | 3.92E-56 | 4 |
| RPL34 | -0.4902081 | 2.84E-55 | 4 |
| SH3BGRL3 | -0.6848657 | 1.26E-54 | 4 |
| RPL8 | -0.457298 | 2.16E-54 | 4 |
| ID3 | -1.1568016 | 5.96E-54 | 4 |
| RPL27A | -0.4883188 | 7.52E-54 | 4 |
| RPL36 | -0.4662767 | 2.41E-53 | 4 |
| TMSB4X | -0.443657 | 1.46E-51 | 4 |
| RPS15A | -0.4502656 | 3.57E-51 | 4 |
| SIRPG | -0.7438421 | 5.16E-51 | 4 |
| WAKMAR2 | 0.66419231 | 1.25E-50 | 4 |
| SSBP2 | 0.65552092 | 2.61E-49 | 4 |
| RPL3 | -0.4134331 | 3.24E-49 | 4 |
| CD27 | -0.7277193 | 2.56E-48 | 4 |
| RPL27 | -0.4324494 | 4.00E-48 | 4 |
| RPS15 | -0.3996063 | 3.27E-47 | 4 |
| CTSW | -0.8116902 | 4.51E-47 | 4 |
| DPP4 | 0.38397489 | 5.65E-47 | 4 |
| RPL28 | -0.409267 | 8.68E-47 | 4 |
| JUN | 0.93905368 | 3.19E-46 | 4 |
| CHDH | 0.59275956 | 6.28E-45 | 4 |
| RPS10 | -0.4635441 | 6.96E-45 | 4 |
| ABCC4 | 0.45509154 | 1.07E-44 | 4 |
| RPL9 | -0.3959887 | 3.10E-44 | 4 |
| ARGLU1 | 0.5762726 | 3.67E-44 | 4 |
| RPL29 | -0.4154279 | 5.44E-44 | 4 |
| RPL38 | -0.4021301 | 1.29E-43 | 4 |
| RACK1 | -0.4257815 | 1.64E-43 | 4 |
| RPL15 | -0.3868161 | 2.04E-43 | 4 |
| ZNF683 | -1.1055678 | 7.55E-43 | 4 |
| FAU | -0.3978042 | 1.27E-42 | 4 |
| RPL23A | -0.3730643 | 2.00E-42 | 4 |
| RAC2 | -0.4724471 | 2.03E-42 | 4 |

| | | | |
|---|---|---|---|
| RPL37A | -0.3348004 | 2.10E-42 | 4 |
| PDE7A | 0.51731262 | 3.28E-42 | 4 |
| PRMT7 | 0.63799373 | 7.68E-42 | 4 |
| BCL11B | 0.57350365 | 8.48E-42 | 4 |
| RPL14 | -0.4777464 | 1.60E-41 | 4 |
| CD2 | 0.50140123 | 1.83E-41 | 4 |
| CLIC1 | -0.5144761 | 7.36E-41 | 4 |
| RPS21 | -0.3579293 | 7.70E-41 | 4 |
| SEMA5A | 0.47535381 | 1.71E-40 | 4 |
| THEMIS | 0.5954185 | 4.31E-40 | 4 |
| RPL24 | -0.4156054 | 5.61E-40 | 4 |
| ACTG1 | -0.4083877 | 8.92E-40 | 4 |
| AAK1 | 0.56489578 | 1.82E-39 | 4 |
| BTG2 | 0.60185571 | 2.00E-39 | 4 |
| RPL18A | -0.3795649 | 1.25E-38 | 4 |
| TFDP2 | 0.60574428 | 1.38E-38 | 4 |
| GIMAP7 | -0.6388533 | 2.54E-38 | 4 |
| SYNE2 | 0.75790357 | 3.17E-38 | 4 |
| RPL17 | -0.3748744 | 9.97E-38 | 4 |
| RPS9 | -0.3663055 | 1.17E-37 | 4 |
| VASP | -0.517474 | 1.82E-37 | 4 |
| RPL30 | -0.3911711 | 2.38E-37 | 4 |
| ZP1 | 0.45900925 | 3.09E-37 | 4 |
| TPT1 | -0.3597619 | 3.52E-37 | 4 |
| VPS13C | 0.53776891 | 1.09E-36 | 4 |
| SMIM24 | -0.698026 | 4.17E-36 | 4 |
| AUTS2 | 0.59252482 | 5.44E-36 | 4 |
| AHNAK | -0.9398374 | 7.45E-36 | 4 |
| MT-ND1 | 0.44830923 | 1.27E-35 | 4 |
| ARPP21 | 0.59017473 | 2.31E-35 | 4 |
| LINC01222 | 0.63280626 | 3.29E-35 | 4 |
| NOP53 | -0.4826205 | 1.04E-34 | 4 |
| COTL1 | -0.5611198 | 1.18E-34 | 4 |
| SCART1 | 0.38694861 | 2.08E-34 | 4 |
| CTSA | -0.4831109 | 2.50E-34 | 4 |
| DNMBP | 0.31243112 | 4.00E-34 | 4 |
| AFF3 | 0.61623513 | 5.27E-34 | 4 |
| PIK3R3 | 0.76325004 | 6.53E-34 | 4 |
| COL6A2 | -0.5629429 | 7.94E-34 | 4 |
| TIAM1 | 0.57013846 | 1.08E-33 | 4 |
| SERPINB1 | -0.485555 | 1.34E-33 | 4 |
| PTPN6 | -0.6099457 | 1.60E-33 | 4 |
| RPS19 | -0.3420156 | 1.95E-33 | 4 |

| | | | |
|---|---|---|---|
| RPL4 | -0.3766908 | 2.42E-33 | 4 |
| SRSF10 | 0.52621699 | 2.50E-33 | 4 |
| UBA52 | -0.3429524 | 2.85E-33 | 4 |
| P2RY11 | -0.4106383 | 3.58E-33 | 4 |
| SAMHD1 | -0.563897 | 4.68E-33 | 4 |
| S100A11 | -0.6914242 | 5.95E-33 | 4 |
| POR | 0.51789284 | 8.74E-33 | 4 |
| PTPRC | 0.41796869 | 2.07E-32 | 4 |
| RPS6 | -0.3461846 | 2.91E-32 | 4 |
| EEF1G | -0.4017882 | 3.77E-32 | 4 |
| LGALS1 | -1.3524483 | 7.94E-32 | 4 |
| RPS26 | -0.3490943 | 8.78E-32 | 4 |
| RASGRP2 | -0.443537 | 9.41E-32 | 4 |
| TSPO | -0.4831589 | 2.33E-31 | 4 |
| SAMD3 | -0.5034026 | 6.04E-31 | 4 |
| NACA | -0.3683142 | 6.54E-31 | 4 |
| RPL31 | -0.3745223 | 8.48E-31 | 4 |
| SHISA2 | -0.5045923 | 9.63E-31 | 4 |
| FCER1G | -0.4618828 | 2.48E-30 | 4 |
| TPM3 | -0.451875 | 3.54E-30 | 4 |
| HELLS | 0.77116545 | 3.81E-30 | 4 |
| C9orf16 | -0.4427144 | 5.34E-30 | 4 |
| RPS11 | -0.3367631 | 6.31E-30 | 4 |
| HMGB2 | 0.28243379 | 8.63E-30 | 4 |
| IL27RA | -0.4567889 | 1.48E-29 | 4 |
| SLC18A2 | 0.49061988 | 1.52E-29 | 4 |
| AC068587.4 | 0.60805908 | 2.76E-29 | 4 |
| KCNMA1 | 0.26741381 | 2.81E-29 | 4 |
| RPL22 | -0.3381583 | 2.82E-29 | 4 |
| RASSF1 | -0.4505766 | 2.82E-29 | 4 |
| SRGN | -0.698844 | 5.43E-29 | 4 |
| IGFLR1 | -0.4162577 | 5.67E-29 | 4 |
| HSP90AA1 | 0.40113151 | 5.79E-29 | 4 |
| CD63 | -0.5217716 | 5.95E-29 | 4 |
| SAMD12 | 0.50419554 | 7.71E-29 | 4 |
| RPSA | -0.3421788 | 8.02E-29 | 4 |
| IGFBP5 | 0.62591295 | 1.57E-28 | 4 |
| SMC4 | -0.7619699 | 1.91E-28 | 4 |
| ARID5B | -0.4729214 | 2.07E-28 | 4 |
| RPL7 | -0.3207013 | 3.73E-28 | 4 |
| CYBA | -0.4363517 | 3.84E-28 | 4 |
| AC007952.4 | 0.4198955 | 4.43E-28 | 4 |
| TPST2 | -0.4696782 | 1.05E-27 | 4 |

| | | | |
|---|---|---|---|
| LINC-PINT | -0.3961469 | 1.09E-27 | 4 |
| CD247 | -0.4988537 | 2.08E-27 | 4 |
| ISG20 | -0.4760935 | 2.35E-27 | 4 |
| MYO1F | -0.446358 | 2.54E-27 | 4 |
| PFDN5 | -0.3574146 | 3.02E-27 | 4 |
| TSC22D1 | 0.55883473 | 4.15E-27 | 4 |
| ACTN1 | -0.4411611 | 4.26E-27 | 4 |
| WDR76 | 0.52491681 | 5.07E-27 | 4 |
| DBN1 | -0.3952506 | 5.93E-27 | 4 |
| ARPC2 | -0.3661129 | 6.43E-27 | 4 |
| MKNK1 | 0.38881327 | 8.18E-27 | 4 |
| CCNE2 | 0.39836666 | 8.47E-27 | 4 |
| FCMR | -0.4321706 | 1.06E-26 | 4 |
| GZMM | -0.4648923 | 1.39E-26 | 4 |
| RPL21 | -0.3087727 | 1.54E-26 | 4 |
| RPS16 | -0.2940587 | 4.34E-26 | 4 |
| S100A10 | -0.7173074 | 5.58E-26 | 4 |
| PTGDR2 | 0.35658339 | 6.05E-26 | 4 |
| REC8 | 0.52504197 | 6.86E-26 | 4 |
| ITM2A | -0.8487897 | 6.99E-26 | 4 |
| KLF2 | -0.7823147 | 8.57E-26 | 4 |
| PFN1 | -0.3515017 | 1.10E-25 | 4 |
| PAXIP1 | 0.43723386 | 1.60E-25 | 4 |
| ITM2C | 0.52252726 | 2.21E-25 | 4 |
| RPS5 | -0.3607354 | 2.48E-25 | 4 |
| STAT5A | -0.4874923 | 2.48E-25 | 4 |
| EEF1B2 | -0.3651788 | 2.84E-25 | 4 |
| RPL35A | -0.3062458 | 5.67E-25 | 4 |
| GBP2 | -0.4144693 | 5.68E-25 | 4 |
| BTF3 | -0.3630675 | 1.78E-24 | 4 |
| DOK2 | -0.4210919 | 1.79E-24 | 4 |
| RPS3A | -0.2943511 | 1.85E-24 | 4 |
| FOXO1 | -0.3226538 | 1.95E-24 | 4 |
| SCRIB | -0.5929974 | 2.52E-24 | 4 |
| LIMD2 | -0.4395908 | 2.86E-24 | 4 |
| PRDX1 | -0.400712 | 9.40E-24 | 4 |
| CDC42EP3 | 0.52918024 | 1.09E-23 | 4 |
| NME2 | -0.5385386 | 1.40E-23 | 4 |
| SUPT3H | -0.3349349 | 1.46E-23 | 4 |
| COMMD6 | -0.3615751 | 1.61E-23 | 4 |
| MTRNR2L12 | 0.45811067 | 1.79E-23 | 4 |
| INPP4A | 0.46536164 | 2.31E-23 | 4 |
| RAB37 | -0.3890025 | 2.88E-23 | 4 |

| | | | |
|---|---|---|---|
| PSAP | 0.41470459 | 3.45E-23 | 4 |
| POLR2J3 | 0.42460513 | 3.59E-23 | 4 |
| MZB1 | 0.49124592 | 3.92E-23 | 4 |
| JAK1 | -0.4429607 | 4.19E-23 | 4 |
| CXCR3 | -0.3129588 | 5.04E-23 | 4 |
| CD52 | -0.4653417 | 6.83E-23 | 4 |
| SNHG5 | -0.3856366 | 1.74E-22 | 4 |
| PABPC1 | -0.3310843 | 2.04E-22 | 4 |
| LINC00342 | 0.56198132 | 2.57E-22 | 4 |
| EEF2 | -0.3285238 | 3.20E-22 | 4 |
| TRA2B | 0.40051492 | 3.61E-22 | 4 |
| PNISR | 0.38828105 | 4.58E-22 | 4 |
| XBP1 | 0.52611094 | 7.12E-22 | 4 |
| OST4 | -0.3106279 | 8.75E-22 | 4 |
| MT-ND4 | 0.33603064 | 1.73E-21 | 4 |
| ARPC3 | -0.3222542 | 1.82E-21 | 4 |
| EVI2B | -0.3592926 | 2.00E-21 | 4 |
| NEGR1 | 0.48657276 | 2.18E-21 | 4 |
| FXYD5 | -0.4476661 | 2.21E-21 | 4 |
| RHOC | -0.3236035 | 3.37E-21 | 4 |
| PNN | 0.38386084 | 3.71E-21 | 4 |
| RUNX3 | -0.3550057 | 3.83E-21 | 4 |
| CXXC5 | -0.4312458 | 4.74E-21 | 4 |
| LEPROTL1 | -0.3609877 | 8.47E-21 | 4 |
| KMT2C | 0.40996773 | 9.46E-21 | 4 |
| GALNT6 | 0.47324656 | 9.78E-21 | 4 |
| HMGN2 | 0.39460792 | 9.95E-21 | 4 |
| RPS13 | -0.3035514 | 1.37E-20 | 4 |
| LIME1 | -0.4117516 | 1.68E-20 | 4 |
| HNRNPU | 0.35380384 | 2.21E-20 | 4 |
| SUN2 | -0.3576057 | 2.24E-20 | 4 |
| CPNE1 | -0.336162 | 2.72E-20 | 4 |
| RUFY2 | 0.50450184 | 2.81E-20 | 4 |
| RALGDS | -0.3170512 | 2.95E-20 | 4 |
| RPL5 | -0.3019033 | 3.80E-20 | 4 |
| PTPRD | 0.32657705 | 3.91E-20 | 4 |
| ARHGEF7 | 0.41837406 | 6.22E-20 | 4 |
| RORB | 0.57963834 | 6.45E-20 | 4 |
| BIN1 | -0.3366072 | 6.46E-20 | 4 |
| ETS1 | -0.3818833 | 7.28E-20 | 4 |
| MCM4 | 0.44288409 | 8.47E-20 | 4 |
| PRKCH | -0.463528 | 8.59E-20 | 4 |
| ZYX | -0.4149387 | 1.37E-19 | 4 |

| | | | |
|---|---|---|---|
| ZNF107 | 0.43317915 | 1.37E-19 | 4 |
| RPL6 | -0.282462 | 1.42E-19 | 4 |
| ARPC1B | -0.3447996 | 1.45E-19 | 4 |
| SH3TC1 | 0.38240244 | 1.93E-19 | 4 |
| CCNL1 | 0.44083339 | 2.43E-19 | 4 |
| FGFR1 | -0.4094349 | 2.50E-19 | 4 |
| DLEU2 | 0.43896056 | 2.54E-19 | 4 |
| TFF3 | 0.57328202 | 2.61E-19 | 4 |
| ATP5F1E | -0.2920301 | 2.62E-19 | 4 |
| CST7 | -0.3761051 | 2.87E-19 | 4 |
| UXT | -0.361938 | 2.95E-19 | 4 |
| COX7C | -0.2770517 | 3.32E-19 | 4 |
| TSPAN32 | -0.297981 | 3.33E-19 | 4 |
| GPR15 | -0.3221634 | 3.51E-19 | 4 |
| SATB1 | 0.36497978 | 3.81E-19 | 4 |
| FOXP1 | 0.39347273 | 4.41E-19 | 4 |
| HDAC7 | 0.39615974 | 5.46E-19 | 4 |
| ZNRF1 | 0.3593471 | 6.38E-19 | 4 |
| TRBC2 | 0.3017907 | 6.95E-19 | 4 |
| TMEM160 | -0.2954502 | 6.98E-19 | 4 |
| LUC7L3 | 0.36704076 | 7.10E-19 | 4 |
| RAB33A | -0.2652699 | 9.83E-19 | 4 |
| SDCBP | -0.3528263 | 1.11E-18 | 4 |
| GRAP2 | 0.37859263 | 1.15E-18 | 4 |
| GMFG | -0.3511746 | 1.17E-18 | 4 |
| DGKZ | -0.3358022 | 1.27E-18 | 4 |
| FLT3LG | 0.32518855 | 1.41E-18 | 4 |
| GCNT4 | -0.3226082 | 1.44E-18 | 4 |
| TWF2 | -0.3226993 | 1.52E-18 | 4 |
| RTKN2 | -0.3985281 | 1.71E-18 | 4 |
| TRIM14 | 0.4644112 | 1.80E-18 | 4 |
| STAG3 | 0.43879063 | 1.84E-18 | 4 |
| CERK | 0.33570532 | 1.84E-18 | 4 |
| GLUL | 0.47655892 | 2.08E-18 | 4 |
| FAM111B | 0.36805453 | 2.15E-18 | 4 |
| LINC01226 | 0.34426285 | 3.76E-18 | 4 |
| N4BP2 | 0.46279011 | 3.76E-18 | 4 |
| DDAH2 | -0.3668749 | 3.84E-18 | 4 |
| SNHG18 | 0.39656203 | 5.09E-18 | 4 |
| VAMP5 | -0.347603 | 6.13E-18 | 4 |
| ANXA2 | -0.366555 | 1.11E-17 | 4 |
| S1PR1 | -0.2689494 | 1.14E-17 | 4 |
| E2F2 | 0.347288 | 1.25E-17 | 4 |

| | | | |
|---|---|---|---|
| TXK | -0.2771631 | 1.30E-17 | 4 |
| ABLIM1 | -0.3721411 | 1.35E-17 | 4 |
| PAXIP1-AS1 | 0.347246 | 1.43E-17 | 4 |
| PLAC8 | -0.5697823 | 1.50E-17 | 4 |
| DEK | 0.40205002 | 1.57E-17 | 4 |
| CD44 | -0.4443373 | 1.84E-17 | 4 |
| TAF10 | -0.3223457 | 1.89E-17 | 4 |
| PTPRF | 0.28013947 | 2.02E-17 | 4 |
| UBASH3B | -0.3800952 | 2.15E-17 | 4 |
| ARHGAP5 | 0.31544111 | 2.31E-17 | 4 |
| TLE1 | 0.40887466 | 3.21E-17 | 4 |
| SLC5A3 | 0.39534068 | 3.37E-17 | 4 |
| CALM2 | -0.2885109 | 3.78E-17 | 4 |
| CD84 | -0.4705443 | 4.67E-17 | 4 |
| HSPB1 | -0.3496581 | 4.72E-17 | 4 |
| SMPD3 | 0.42744437 | 5.13E-17 | 4 |
| RPL36AL | -0.2881935 | 6.73E-17 | 4 |
| RGS3 | -0.4609855 | 7.04E-17 | 4 |
| EIF3K | -0.3041289 | 7.39E-17 | 4 |
| RPS7 | -0.2616688 | 1.08E-16 | 4 |
| RBBP4 | 0.38136714 | 1.22E-16 | 4 |
| FAM102A | -0.331205 | 1.26E-16 | 4 |
| LAT | 0.32158994 | 1.55E-16 | 4 |
| GAS5 | -0.4052029 | 1.69E-16 | 4 |
| S1PR3 | 0.39892542 | 2.01E-16 | 4 |
| EIF3I | -0.3127202 | 2.33E-16 | 4 |
| IL16 | -0.3686722 | 2.66E-16 | 4 |
| PDCD1 | -0.2838632 | 2.77E-16 | 4 |
| MME | -0.3461548 | 3.01E-16 | 4 |
| PPAN | -0.2527397 | 3.02E-16 | 4 |
| NSMCE1 | -0.2903963 | 3.84E-16 | 4 |
| SSR2 | -0.3119049 | 4.24E-16 | 4 |
| CHMP7 | -0.3666686 | 4.30E-16 | 4 |
| COX6C | -0.3141468 | 4.38E-16 | 4 |
| CCNE1 | 0.36165705 | 4.66E-16 | 4 |
| JPH1 | 0.31896473 | 5.50E-16 | 4 |
| OPTN | -0.3357677 | 5.53E-16 | 4 |
| NCOA7 | -0.3305726 | 6.67E-16 | 4 |
| CASP8 | 0.52049611 | 8.07E-16 | 4 |
| SLC14A1 | 0.37707331 | 1.01E-15 | 4 |
| ATP5MC2 | -0.2836891 | 1.15E-15 | 4 |
| FUT7 | -0.2838034 | 1.83E-15 | 4 |
| LY9 | -0.2561344 | 1.85E-15 | 4 |

| | | | |
|---|---|---|---|
| CDC25B | -0.3067601 | 2.15E-15 | 4 |
| ALKBH7 | -0.2901435 | 2.71E-15 | 4 |
| NASP | 0.39824771 | 2.88E-15 | 4 |
| VIM | -0.6318659 | 3.12E-15 | 4 |
| RBM38 | -0.293599 | 3.40E-15 | 4 |
| SMC3 | 0.40373392 | 3.49E-15 | 4 |
| LSM2 | -0.2790337 | 3.50E-15 | 4 |
| MAPK13 | -0.2512731 | 3.63E-15 | 4 |
| GADD45GIP1 | -0.2923737 | 4.27E-15 | 4 |
| AL355075.4 | 0.38259702 | 5.06E-15 | 4 |
| SPINT2 | -0.3342877 | 5.08E-15 | 4 |
| CRTAM | -0.2761056 | 5.32E-15 | 4 |
| GYPC | -0.3057731 | 5.51E-15 | 4 |
| ATP2B4 | 0.40101762 | 6.16E-15 | 4 |
| GLCCI1 | 0.39763693 | 6.71E-15 | 4 |
| ARL4C | 0.31643048 | 7.93E-15 | 4 |
| CD6 | -0.2831275 | 8.67E-15 | 4 |
| PARVG | -0.2940835 | 8.94E-15 | 4 |
| LINC00891 | 0.42519151 | 9.44E-15 | 4 |
| AC093673.1 | -0.2646933 | 1.06E-14 | 4 |
| NHP2 | -0.3033522 | 1.36E-14 | 4 |
| C4orf48 | -0.2822604 | 1.50E-14 | 4 |
| AIP | -0.2906128 | 1.55E-14 | 4 |
| ZMIZ1 | -0.2859118 | 1.61E-14 | 4 |
| FYN | -0.3053125 | 1.93E-14 | 4 |
| VAMP8 | -0.2764727 | 2.03E-14 | 4 |
| FUS | 0.30249629 | 2.23E-14 | 4 |
| MDM4 | 0.37248079 | 2.24E-14 | 4 |
| ABHD17A | -0.3318556 | 2.58E-14 | 4 |
| ST8SIA4 | -0.2817891 | 2.70E-14 | 4 |
| RPL36A | -0.2770124 | 2.81E-14 | 4 |
| RNF7 | -0.277052 | 2.86E-14 | 4 |
| P2RX5 | -0.3393701 | 2.90E-14 | 4 |
| KDM7A | -0.2551991 | 3.93E-14 | 4 |
| CAPNS1 | -0.2936826 | 4.17E-14 | 4 |
| PGLS | -0.302116 | 4.51E-14 | 4 |
| NCF1 | -0.3656175 | 4.86E-14 | 4 |
| LRP12 | 0.40969154 | 5.63E-14 | 4 |
| ADGRE5 | -0.315436 | 5.68E-14 | 4 |
| FIS1 | -0.2842936 | 6.27E-14 | 4 |
| TECR | -0.743156 | 6.53E-14 | 4 |
| RSL24D1 | -0.2772937 | 8.01E-14 | 4 |
| CALM1 | -0.3372807 | 9.89E-14 | 4 |

| | | | |
|---|---|---|---|
| NIFK | -0.2719492 | 1.14E-13 | 4 |
| MMS22L | 0.42420988 | 1.19E-13 | 4 |
| PLCL2 | -0.2738198 | 1.30E-13 | 4 |
| APBB1IP | 0.31682332 | 1.32E-13 | 4 |
| KLRB1 | -0.452828 | 1.34E-13 | 4 |
| CBX5 | 0.3409789 | 1.43E-13 | 4 |
| ISG15 | -0.2681222 | 1.49E-13 | 4 |
| PYHIN1 | -0.3131999 | 1.60E-13 | 4 |
| NDUFB9 | -0.278646 | 1.79E-13 | 4 |
| RGCC | -0.3670886 | 1.86E-13 | 4 |
| FLNA | -0.4004069 | 2.14E-13 | 4 |
| NOSIP | -0.6649176 | 3.03E-13 | 4 |
| SOX4 | 0.3466269 | 3.07E-13 | 4 |
| PPP1R18 | -0.3014553 | 3.38E-13 | 4 |
| HNRNPH1 | 0.34720758 | 3.62E-13 | 4 |
| DUSP2 | -0.3597584 | 3.80E-13 | 4 |
| MIF | -0.3634914 | 4.58E-13 | 4 |
| ZFP36 | -0.2938487 | 5.10E-13 | 4 |
| GIMAP4 | -0.3171554 | 5.30E-13 | 4 |
| ID2 | 0.29545509 | 8.32E-13 | 4 |
| PRPF4B | 0.36669132 | 8.38E-13 | 4 |
| NSA2 | -0.3211618 | 9.01E-13 | 4 |
| CLEC2D | -0.5466054 | 9.51E-13 | 4 |
| CDK5RAP3 | 0.35709156 | 1.07E-12 | 4 |
| FDFT1 | -0.2910161 | 1.13E-12 | 4 |
| EIF3F | -0.3051353 | 1.21E-12 | 4 |
| ICAM2 | -0.2884268 | 1.23E-12 | 4 |
| REPIN1 | 0.35303334 | 1.27E-12 | 4 |
| NKG7 | -0.3525077 | 1.50E-12 | 4 |
| MZT2B | -0.2573485 | 1.59E-12 | 4 |
| INPP5D | -0.2615673 | 1.60E-12 | 4 |
| MVB12B | 0.29040912 | 1.79E-12 | 4 |
| GRK6 | -0.2659456 | 1.82E-12 | 4 |
| BAZ1B | 0.3203332 | 1.86E-12 | 4 |
| SRGAP3 | -0.3183132 | 1.91E-12 | 4 |
| HLA-E | -0.3158914 | 1.93E-12 | 4 |
| RHOG | -0.2748695 | 1.99E-12 | 4 |
| CYTH4 | -0.2523845 | 2.09E-12 | 4 |
| ARHGAP25 | -0.2628411 | 2.38E-12 | 4 |
| C1QBP | -0.31812 | 2.42E-12 | 4 |
| LIMA1 | -0.262034 | 3.14E-12 | 4 |
| IGBP1 | -0.281039 | 3.96E-12 | 4 |
| LCP1 | 0.27798274 | 4.70E-12 | 4 |

| | | | |
|---|---|---|---|
| PHB2 | -0.2618223 | 5.33E-12 | 4 |
| TRIM22 | -0.3016404 | 5.40E-12 | 4 |
| IKZF2 | -0.5141001 | 6.17E-12 | 4 |
| GTF3A | -0.3004873 | 7.20E-12 | 4 |
| ROMO1 | -0.2770444 | 9.34E-12 | 4 |
| ATP2A1 | -0.2659298 | 9.79E-12 | 4 |
| IRF2BPL | 0.36032272 | 9.99E-12 | 4 |
| MRPL51 | -0.2571439 | 1.21E-11 | 4 |
| CHST2 | -0.2601755 | 1.25E-11 | 4 |
| STMN1 | 0.38202859 | 1.26E-11 | 4 |
| UFC1 | -0.27967 | 1.33E-11 | 4 |
| PSMB6 | -0.2594911 | 1.41E-11 | 4 |
| EIF3G | -0.2736505 | 1.50E-11 | 4 |
| ITGB2 | -0.3522596 | 1.69E-11 | 4 |
| PPARA | 0.39320789 | 2.29E-11 | 4 |
| ARHGEF6 | -0.2554178 | 2.39E-11 | 4 |
| ELF1 | -0.3290497 | 2.84E-11 | 4 |
| APOBEC3G | -0.2599455 | 2.94E-11 | 4 |
| IRF2BP2 | -0.3205942 | 2.96E-11 | 4 |
| MED13 | 0.38936534 | 3.11E-11 | 4 |
| UQCR11 | -0.2503077 | 3.21E-11 | 4 |
| WDR83OS | -0.273992 | 3.37E-11 | 4 |
| AKAP9 | 0.30486512 | 3.59E-11 | 4 |
| RNASEH2B | 0.34764524 | 3.71E-11 | 4 |
| MYL12A | -0.2512322 | 4.51E-11 | 4 |
| B3GNT2 | -0.2513223 | 5.30E-11 | 4 |
| RBM33 | 0.34197347 | 5.32E-11 | 4 |
| HNRNPH3 | 0.31699707 | 6.78E-11 | 4 |
| C12orf57 | -0.3560642 | 7.42E-11 | 4 |
| CAST | -0.3086355 | 8.25E-11 | 4 |
| SIVA1 | 0.28690231 | 8.34E-11 | 4 |
| TRIM8 | 0.36539013 | 8.61E-11 | 4 |
| UQCRB | -0.2526262 | 1.00E-10 | 4 |
| ICAM3 | -0.2952357 | 1.03E-10 | 4 |
| HSP90AB1 | -0.4246811 | 1.10E-10 | 4 |
| MCM7 | 0.35748651 | 1.12E-10 | 4 |
| HNRNPA3 | 0.2650814 | 1.15E-10 | 4 |
| ITM2B | -0.2610862 | 1.16E-10 | 4 |
| CDC42SE1 | -0.275181 | 1.52E-10 | 4 |
| KLF13 | 0.30226617 | 1.91E-10 | 4 |
| YPEL3 | -0.2659609 | 2.00E-10 | 4 |
| ANKRD36C | 0.42694029 | 2.23E-10 | 4 |
| BACH2 | -0.2826606 | 2.54E-10 | 4 |

| | | | |
|---|---|---|---|
| SRSF7 | 0.28234709 | 2.98E-10 | 4 |
| KRAS | -0.3011512 | 5.38E-10 | 4 |
| MAP3K1 | -0.2548461 | 5.57E-10 | 4 |
| ADD3 | -0.4186146 | 6.36E-10 | 4 |
| NKTR | 0.33986661 | 6.88E-10 | 4 |
| DUT | 0.27221313 | 7.16E-10 | 4 |
| SH3KBP1 | -0.2964807 | 7.18E-10 | 4 |
| AHI1 | 0.39468973 | 7.60E-10 | 4 |
| CAMK1D | 0.33442717 | 7.61E-10 | 4 |
| APRT | -0.2760503 | 7.87E-10 | 4 |
| TIMP1 | -0.3283147 | 9.72E-10 | 4 |
| ANKRD44 | 0.3569167 | 1.07E-09 | 4 |
| TGFB1 | -0.2847633 | 1.75E-09 | 4 |
| TOMM7 | -0.2684484 | 1.95E-09 | 4 |
| WDR34 | 0.3209246 | 2.14E-09 | 4 |
| ZNF704 | 0.26159461 | 2.51E-09 | 4 |
| LSP1 | -0.2598125 | 2.58E-09 | 4 |
| EIF3E | -0.2620648 | 2.65E-09 | 4 |
| IL2RG | -0.2652396 | 2.85E-09 | 4 |
| TYMS | 0.32233321 | 3.12E-09 | 4 |
| ZNF280D | 0.35426978 | 3.26E-09 | 4 |
| H2AFY | 0.33249372 | 3.66E-09 | 4 |
| SELENOW | -0.3569907 | 4.82E-09 | 4 |
| PTGDR | -0.2761559 | 5.13E-09 | 4 |
| CHD1 | 0.37724873 | 5.43E-09 | 4 |
| MT-ATP6 | 0.26982104 | 5.70E-09 | 4 |
| ST3GAL5 | 0.34224601 | 5.84E-09 | 4 |
| N4BP2L2 | 0.32297522 | 6.98E-09 | 4 |
| PCNA | 0.52132348 | 8.63E-09 | 4 |
| EPG5 | 0.26603152 | 8.77E-09 | 4 |
| OXR1 | 0.34600268 | 1.34E-08 | 4 |
| TMSB10 | -0.3332205 | 1.35E-08 | 4 |
| REV3L | 0.40846358 | 1.46E-08 | 4 |
| ESYT2 | -0.3165995 | 1.57E-08 | 4 |
| FKBP5 | 0.38116628 | 2.38E-08 | 4 |
| TAGLN2 | -0.4343314 | 2.49E-08 | 4 |
| SLC25A6 | -0.2706506 | 2.90E-08 | 4 |
| ERAP1 | 0.30758693 | 2.98E-08 | 4 |
| JPX | 0.31671391 | 3.82E-08 | 4 |
| POLR2L | -0.2547125 | 4.43E-08 | 4 |
| STK17A | -0.3017796 | 4.59E-08 | 4 |
| CLIC3 | -0.3474066 | 4.62E-08 | 4 |
| SARAF | -0.2609466 | 4.65E-08 | 4 |

| | | | |
|---|---|---|---|
| RASGRP1 | -0.2544582 | 4.96E-08 | 4 |
| TRIM56 | 0.25793528 | 6.03E-08 | 4 |
| SMC1A | 0.29202025 | 6.14E-08 | 4 |
| SLC12A6 | -0.2779523 | 6.14E-08 | 4 |
| RAP1B | -0.260591 | 6.29E-08 | 4 |
| CUX1 | 0.27374862 | 6.41E-08 | 4 |
| STK17B | 0.25922652 | 6.95E-08 | 4 |
| ASH1L | 0.3378084 | 7.72E-08 | 4 |
| H1FX | 0.26341891 | 8.43E-08 | 4 |
| PCLAF | 0.35554269 | 9.44E-08 | 4 |
| ARFGEF1 | 0.30194754 | 1.10E-07 | 4 |
| KMT2E | 0.25684009 | 1.15E-07 | 4 |
| MSH6 | 0.35849242 | 1.15E-07 | 4 |
| HIST1H3D | -0.4498541 | 1.21E-07 | 4 |
| TBL1XR1 | 0.26377703 | 1.22E-07 | 4 |
| PBRM1 | 0.2628326 | 1.49E-07 | 4 |
| DNMT1 | 0.27738264 | 1.66E-07 | 4 |
| C21orf58 | 0.30328729 | 1.83E-07 | 4 |
| BCL7A | 0.25803179 | 1.88E-07 | 4 |
| ADA | 0.2598641 | 1.99E-07 | 4 |
| GATA3 | -0.3112828 | 2.08E-07 | 4 |
| HLA-C | -0.3350085 | 2.18E-07 | 4 |
| HLA-A | -0.3708089 | 2.22E-07 | 4 |
| AC092683.1 | 0.39721012 | 2.58E-07 | 4 |
| HLA-B | -0.5114383 | 2.74E-07 | 4 |
| DDX39B | 0.25019794 | 2.98E-07 | 4 |
| RUFY3 | 0.28417954 | 3.29E-07 | 4 |
| TMEM123 | 0.2804581 | 3.95E-07 | 4 |
| BTG1 | 0.2933501 | 4.10E-07 | 4 |
| BOD1L1 | 0.26855901 | 4.37E-07 | 4 |
| NUP210 | 0.32133675 | 4.50E-07 | 4 |
| PSMA3-AS1 | 0.28486322 | 5.11E-07 | 4 |
| SRSF5 | 0.25211166 | 5.19E-07 | 4 |
| VIPR2 | 0.2525313 | 6.18E-07 | 4 |
| GPR174 | -0.2824138 | 8.61E-07 | 4 |
| KLHL23 | 0.29159426 | 1.00E-06 | 4 |
| TOX2 | -0.3282947 | 1.67E-06 | 4 |
| CLSPN | 0.29528963 | 1.92E-06 | 4 |
| CD2AP | 0.29007605 | 2.40E-06 | 4 |
| PTTG1 | -0.3109539 | 3.17E-06 | 4 |
| RPLP0 | -0.2664002 | 3.53E-06 | 4 |
| VPS13A | 0.27993282 | 4.03E-06 | 4 |
| LCP2 | -0.2726866 | 5.53E-06 | 4 |

| | | | |
|---|---|---|---|
| SEMA4D | -0.288641 | 6.08E-06 | 4 |
| KAT6B | 0.25680858 | 6.18E-06 | 4 |
| PRKDC | 0.27037951 | 6.30E-06 | 4 |
| LAPTM5 | -0.2670175 | 7.14E-06 | 4 |
| UHRF1 | 0.27349648 | 7.44E-06 | 4 |
| IER2 | -0.2624914 | 8.61E-06 | 4 |
| NSD3 | 0.25055505 | 9.37E-06 | 4 |
| TNNT3 | 0.26872189 | 1.03E-05 | 4 |
| KLRK1 | -0.2871589 | 1.69E-05 | 4 |
| MDM1 | 0.27059066 | 1.95E-05 | 4 |
| PSIP1 | 0.26893128 | 2.67E-05 | 4 |
| ATP2A3 | 0.29467859 | 2.86E-05 | 4 |
| EZH2 | 0.34860287 | 3.53E-05 | 4 |
| HIPK1 | 0.2769022 | 3.58E-05 | 4 |
| TMEM106C | 0.27124454 | 5.36E-05 | 4 |
| ITGAE | 0.29776263 | 0.00012172 | 4 |
| ATRX | 0.26112509 | 0.0001545 | 4 |
| ARID1B | 0.28069048 | 0.00017694 | 4 |
| CCDC107 | 0.27050421 | 0.00023206 | 4 |
| NUCB2 | -0.2569062 | 0.00023562 | 4 |
| MLXIP | 0.2603995 | 0.0003736 | 4 |
| ARMH1 | -0.3235504 | 0.00041593 | 4 |
| GAPDH | -0.2878676 | 0.00046677 | 4 |
| ARL6IP1 | -0.2711269 | 0.00050834 | 4 |
| SPTBN1 | -0.2700759 | 0.00052547 | 4 |
| PCBP1-AS1 | 0.25328609 | 0.00062218 | 4 |
| TSC22D3 | -0.2839942 | 0.00073734 | 4 |
| CCDC14 | 0.31151706 | 0.00076511 | 4 |
| CD7 | -0.2552929 | 0.00079247 | 4 |
| FXYD2 | -0.768352 | 0.00087071 | 4 |
| ATAD2 | 0.26851996 | 0.00157793 | 4 |
| TMEM161B-AS: | 0.25379091 | 0.00170658 | 4 |
| NMT2 | 0.25332583 | 0.00174918 | 4 |
| PDS5B | 0.2878231 | 0.00229888 | 4 |
| KCNAB2 | 0.28083047 | 0.00286843 | 4 |
| MYB | -0.2645246 | 0.00333348 | 4 |
| EPM2AIP1 | 0.26691917 | 0.00403316 | 4 |
| MYOM2 | -0.2520503 | 0.00662245 | 4 |
| USP15 | 0.25006923 | 0.00760686 | 4 |
| RBBP8 | 0.30627369 | 0.00784917 | 4 |
| RIPOR2 | -0.3058703 | 0.01068118 | 4 |
| SLBP | 0.36710376 | 0.01251635 | 4 |
| TSPAN7 | -0.2938098 | 0.01329253 | 4 |

| | | | |
|---|---|---|---|
| SLFN13 | 0.30642979 | 0.02793315 | 4 |
| SAMD1 | 0.34824996 | 0.0327711 | 4 |
| RIF1 | 0.35026879 | 0.05764136 | 4 |
| EMP3 | -0.4334473 | 0.10472322 | 4 |
| SCAI | 0.28034072 | 0.10719988 | 4 |
| MCM3 | 0.31894379 | 0.10941597 | 4 |
| NSD2 | 0.2896132 | 0.11699954 | 4 |
| AC114760.2 | 0.25949779 | 0.18088619 | 4 |
| ITGA4 | -0.3678951 | 0.27356568 | 4 |
| SLC16A7 | 0.28694129 | 0.28849 | 4 |
| HIST1H3B | -0.2946574 | 0.49555805 | 4 |
| USP1 | 0.32219848 | 0.76080658 | 4 |
| HIST1H4C | -0.6843022 | 1 | 4 |
| HIST2H2AC | -0.285465 | 1 | 4 |
| NUCB2 | 1.03352838 | 4.18E-145 | 5 |
| SCG2 | 0.82271117 | 6.45E-144 | 5 |
| RGS3 | 1.08902544 | 2.78E-142 | 5 |
| GNG4 | 0.72178297 | 7.45E-137 | 5 |
| ID3 | 1.34744392 | 8.70E-126 | 5 |
| P2RX5 | 0.85563354 | 7.59E-124 | 5 |
| ATP9A | 0.53976695 | 3.64E-118 | 5 |
| B2M | -1.1499276 | 1.76E-115 | 5 |
| COL6A3 | 0.6262163 | 4.28E-115 | 5 |
| ARMH1 | 1.00888034 | 1.36E-107 | 5 |
| GBP2 | 0.78486749 | 6.69E-103 | 5 |
| SEMA4D | 0.9294009 | 3.07E-101 | 5 |
| LTB | -1.6044444 | 3.85E-101 | 5 |
| SMIM24 | 0.89086567 | 3.73E-99 | 5 |
| FGFR1 | 0.79598589 | 2.23E-98 | 5 |
| ITM2A | 1.15891893 | 2.24E-97 | 5 |
| ITGA4 | 1.34620952 | 4.71E-97 | 5 |
| SNTA1 | 0.4994511 | 3.06E-96 | 5 |
| EGR3 | 0.66787795 | 1.34E-93 | 5 |
| LCP2 | 0.73111041 | 2.71E-90 | 5 |
| AC124798.1 | 0.3264663 | 5.16E-90 | 5 |
| HIVEP3 | 0.73366261 | 1.08E-89 | 5 |
| IKZF2 | 0.92510139 | 2.29E-84 | 5 |
| ARL4C | -0.8702951 | 6.81E-84 | 5 |
| TXNIP | -1.1503226 | 1.12E-83 | 5 |
| PDCD1 | 0.55625836 | 1.72E-80 | 5 |
| IGFLR1 | 0.58572747 | 1.71E-78 | 5 |
| SATB1 | 1.03599191 | 1.18E-77 | 5 |
| CLDN1 | 0.28477857 | 1.36E-76 | 5 |

| | | | |
|---|---|---|---|
| TUBB2A | 0.52685413 | 1.36E-74 | 5 |
| GAPDH | 0.62062833 | 2.57E-73 | 5 |
| IKZF3 | 0.8708411 | 2.84E-72 | 5 |
| DMD | 0.54783469 | 1.18E-71 | 5 |
| CD247 | 0.68309312 | 6.66E-71 | 5 |
| BACH2 | 0.69083854 | 9.29E-71 | 5 |
| ACTG1 | 0.54280902 | 2.69E-70 | 5 |
| NCR3LG1 | 0.25951255 | 4.54E-70 | 5 |
| SRGN | 0.89370241 | 8.00E-69 | 5 |
| SRGAP3 | 0.57053749 | 4.15E-68 | 5 |
| GPC3 | 0.27101275 | 9.91E-68 | 5 |
| LAYN | 0.30268892 | 2.82E-67 | 5 |
| PTPN7 | 0.74344871 | 6.16E-66 | 5 |
| AAK1 | -0.8537896 | 6.36E-66 | 5 |
| NAB2 | 0.49940005 | 7.39E-64 | 5 |
| TIMP1 | 0.78308097 | 1.03E-63 | 5 |
| CD28 | 0.65945611 | 1.04E-63 | 5 |
| SLC39A14 | 0.30251659 | 2.83E-63 | 5 |
| PDE7A | -0.8630872 | 7.92E-62 | 5 |
| SPTBN2 | 0.29347227 | 8.59E-61 | 5 |
| LGALS1 | 0.916488 | 2.67E-60 | 5 |
| SLC7A5 | 0.44966168 | 4.34E-59 | 5 |
| EEF1A1 | 0.37340407 | 5.97E-59 | 5 |
| SH3BGRL3 | 0.63716504 | 1.69E-58 | 5 |
| AC144521.1 | 0.485284 | 1.70E-58 | 5 |
| PCAT18 | 0.47149712 | 1.94E-58 | 5 |
| C9orf16 | 0.56362208 | 2.43E-58 | 5 |
| P2RY11 | 0.49793587 | 1.14E-57 | 5 |
| RHOC | 0.46160363 | 1.85E-57 | 5 |
| WAKMAR2 | -1.0452784 | 5.38E-57 | 5 |
| NFATC1 | 0.54690385 | 6.79E-57 | 5 |
| PTPRC | -0.571391 | 2.96E-56 | 5 |
| BTG1 | -0.854169 | 1.72E-55 | 5 |
| NR4A1 | 0.71882042 | 2.38E-55 | 5 |
| MACROD2 | -0.946774 | 7.41E-55 | 5 |
| LZTFL1 | 0.67120254 | 8.44E-55 | 5 |
| CD84 | 0.74238323 | 8.66E-53 | 5 |
| XBP1 | -0.8806807 | 3.39E-52 | 5 |
| MCCC2 | 0.45788604 | 6.81E-51 | 5 |
| SMIM10 | 0.27032653 | 9.35E-51 | 5 |
| RPL24 | 0.39704379 | 1.94E-50 | 5 |
| MZB1 | -0.9276484 | 2.51E-50 | 5 |
| CLEC2D | 0.84386068 | 5.08E-50 | 5 |

| | | | |
|---|---|---|---|
| SERTAD2 | 0.52234089 | 6.92E-49 | 5 |
| PRKCH | 0.6247154 | 1.06E-48 | 5 |
| AC011893.1 | 0.36896101 | 2.17E-48 | 5 |
| TSPAN7 | 0.67588401 | 7.74E-48 | 5 |
| MT-ND3 | -0.5337732 | 9.03E-48 | 5 |
| GIHCG | 0.59835371 | 1.03E-47 | 5 |
| DBN1 | 0.43243684 | 1.04E-46 | 5 |
| GCNT4 | 0.43231385 | 2.87E-46 | 5 |
| NR4A3 | 0.2958932 | 4.10E-46 | 5 |
| AGMAT | 0.32943867 | 1.30E-45 | 5 |
| LIMD2 | 0.54002643 | 1.32E-45 | 5 |
| PKM | 0.76569148 | 3.12E-45 | 5 |
| SNHG29 | 0.540923 | 5.78E-45 | 5 |
| HLA-C | -0.7459072 | 7.49E-45 | 5 |
| TCF7 | 0.57347473 | 7.76E-45 | 5 |
| HLA-B | -0.8995462 | 7.96E-45 | 5 |
| BTF3 | 0.46232306 | 1.25E-44 | 5 |
| HNRNPA1 | 0.45306586 | 3.34E-44 | 5 |
| TOX | 0.54935141 | 4.86E-44 | 5 |
| SMPD3 | -0.8341949 | 5.62E-44 | 5 |
| CHD4 | 0.53619092 | 5.83E-44 | 5 |
| BCL6 | 0.54212053 | 1.49E-43 | 5 |
| IRF2BP2 | 0.54765807 | 4.81E-43 | 5 |
| NISCH | 0.39661914 | 5.20E-43 | 5 |
| RPL14 | 0.4293636 | 7.75E-43 | 5 |
| TNRC6C | -0.7196743 | 1.25E-42 | 5 |
| TOX2 | 0.57225748 | 1.48E-42 | 5 |
| STAT5A | 0.63567612 | 4.79E-42 | 5 |
| SELENOH | 0.47600708 | 1.42E-41 | 5 |
| RPLP1 | 0.36462199 | 3.47E-41 | 5 |
| PDE4D | 0.5114616 | 1.23E-40 | 5 |
| ZYX | 0.52155808 | 4.03E-40 | 5 |
| TPM3 | 0.48551968 | 4.45E-40 | 5 |
| RHOH | -0.5869024 | 6.84E-40 | 5 |
| CD7 | -0.644104 | 9.75E-40 | 5 |
| STK17B | -0.5662423 | 4.58E-39 | 5 |
| GAS5 | 0.49987408 | 4.79E-39 | 5 |
| STRBP | 0.44664813 | 6.78E-39 | 5 |
| MT-ND2 | -0.5359204 | 3.32E-38 | 5 |
| C12orf57 | 0.55902549 | 2.58E-37 | 5 |
| HMGB2 | -1.1883977 | 2.76E-37 | 5 |
| LIMS1 | 0.49750452 | 9.29E-37 | 5 |
| ATP5MC3 | 0.45264341 | 3.24E-36 | 5 |

| | | | |
|---|---|---|---|
| MAL | -0.7139674 | 3.39E-36 | 5 |
| MYB | 0.74131456 | 3.55E-36 | 5 |
| CHDH | -0.7530051 | 4.40E-36 | 5 |
| MT-ND4 | -0.4057935 | 6.31E-36 | 5 |
| MSL2 | 0.40076943 | 8.21E-36 | 5 |
| RPL28 | 0.30070861 | 1.64E-35 | 5 |
| PLEKHO1 | 0.41589106 | 3.58E-35 | 5 |
| RPS8 | 0.30014966 | 5.82E-35 | 5 |
| BCAT1 | 0.2661007 | 7.17E-35 | 5 |
| EEF1G | 0.36895331 | 7.51E-35 | 5 |
| ABRACL | 0.42563415 | 8.78E-35 | 5 |
| EIF3E | 0.45184141 | 1.01E-34 | 5 |
| MLLT3 | 0.29714249 | 1.08E-34 | 5 |
| MT-ND1 | -0.4725039 | 1.86E-34 | 5 |
| CNIH1 | 0.34000852 | 3.76E-34 | 5 |
| ZFP36L2 | -0.6538655 | 6.50E-34 | 5 |
| PTPN6 | 0.48480333 | 1.06E-33 | 5 |
| AK6 | 0.36511776 | 1.52E-33 | 5 |
| HLA-A | -0.7028143 | 3.18E-33 | 5 |
| ATP6V0A2 | 0.43133315 | 3.65E-33 | 5 |
| TAF9 | 0.36592319 | 4.83E-33 | 5 |
| QSOX1 | 0.30612447 | 1.06E-32 | 5 |
| TEC | 0.29408222 | 1.59E-32 | 5 |
| SDCBP | 0.4783379 | 4.39E-32 | 5 |
| GMFG | 0.41868792 | 4.56E-32 | 5 |
| CD8B | -0.5178749 | 5.80E-32 | 5 |
| UBXN11 | 0.2920365 | 7.74E-32 | 5 |
| IL17RB | -0.9269179 | 8.00E-32 | 5 |
| IGBP1 | 0.38071269 | 1.31E-31 | 5 |
| RPS2 | 0.33667635 | 1.69E-31 | 5 |
| ECI2 | 0.38214253 | 3.77E-31 | 5 |
| C4orf48 | 0.37298155 | 5.47E-31 | 5 |
| GPR174 | 0.4548144 | 5.84E-31 | 5 |
| HBS1L | 0.49389745 | 6.71E-31 | 5 |
| CD6 | 0.37108823 | 7.88E-31 | 5 |
| HIST1H4C | -1.3972753 | 1.04E-30 | 5 |
| MT-CYB | -0.412241 | 1.29E-30 | 5 |
| RACK1 | 0.3111349 | 2.78E-30 | 5 |
| CD38 | 0.53178752 | 3.18E-30 | 5 |
| DUSP2 | 0.64858124 | 3.21E-30 | 5 |
| FAM110A | 0.29734974 | 3.41E-30 | 5 |
| CD5 | 0.32956531 | 3.13E-29 | 5 |
| LST1 | -0.793065 | 5.14E-29 | 5 |

| | | | |
|---|---|---|---|
| STK4 | -0.5062891 | 7.37E-29 | 5 |
| RPS14 | 0.29176007 | 1.18E-28 | 5 |
| DDIT4L | 0.28836744 | 1.33E-28 | 5 |
| RPSA | 0.31536967 | 2.08E-28 | 5 |
| ECHS1 | 0.35658751 | 3.14E-28 | 5 |
| ARHGEF6 | 0.39216184 | 3.30E-28 | 5 |
| SH2D2A | 0.32397798 | 7.26E-28 | 5 |
| RAC2 | 0.33870208 | 7.45E-28 | 5 |
| MT-ATP6 | -0.3379122 | 7.64E-28 | 5 |
| SPINT2 | 0.36958194 | 9.84E-28 | 5 |
| KLRB1 | 0.48934066 | 1.28E-27 | 5 |
| SLAMF1 | 0.42787682 | 1.57E-27 | 5 |
| VDAC1 | 0.35799281 | 1.63E-27 | 5 |
| ZMIZ1 | 0.40938235 | 1.76E-27 | 5 |
| PAK2 | 0.42733624 | 2.33E-27 | 5 |
| CRTAM | 0.4019565 | 5.03E-27 | 5 |
| FAM117B | 0.26513022 | 6.37E-27 | 5 |
| SIVA1 | -0.654831 | 7.16E-27 | 5 |
| TCL1A | -0.5818793 | 7.74E-27 | 5 |
| ENC1 | 0.30786718 | 8.98E-27 | 5 |
| PPAN | 0.31830618 | 1.54E-26 | 5 |
| PITPNM2 | 0.26314263 | 2.39E-26 | 5 |
| TMCO3 | 0.30530905 | 2.39E-26 | 5 |
| RBMX2 | 0.33500011 | 2.52E-26 | 5 |
| CDKN2D | -0.5286645 | 2.85E-26 | 5 |
| ST8SIA4 | 0.37228652 | 6.18E-26 | 5 |
| BICDL1 | 0.27701247 | 7.00E-26 | 5 |
| RPS11 | 0.29490952 | 8.91E-26 | 5 |
| EBPL | 0.28955193 | 1.68E-25 | 5 |
| CD1E | -0.6027096 | 2.64E-25 | 5 |
| IL6ST | 0.37589966 | 2.89E-25 | 5 |
| RPL36A | 0.30837809 | 3.07E-25 | 5 |
| WIPF1 | -0.4737187 | 3.19E-25 | 5 |
| APEX1 | 0.35492052 | 4.14E-25 | 5 |
| ELF1 | 0.36269534 | 5.53E-25 | 5 |
| ARPC2 | 0.34467285 | 5.92E-25 | 5 |
| FDX1 | 0.37555422 | 6.52E-25 | 5 |
| FHL2 | 0.27699331 | 7.73E-25 | 5 |
| CRIP1 | -1.1021556 | 1.32E-24 | 5 |
| HSP90B1 | -0.6372622 | 1.43E-24 | 5 |
| MYL6B | 0.34795573 | 1.78E-24 | 5 |
| VASP | 0.38969706 | 2.04E-24 | 5 |
| RPLP0 | 0.30023862 | 3.50E-24 | 5 |

| | | | |
|---|---|---|---|
| CCNB1IP1 | 0.27139626 | 4.55E-24 | 5 |
| CD200 | 0.27279223 | 5.71E-24 | 5 |
| CCDC88A | 0.44072029 | 8.60E-24 | 5 |
| EIF3H | 0.36424275 | 9.14E-24 | 5 |
| ALOX5AP | 0.39619653 | 9.51E-24 | 5 |
| COTL1 | 0.36325558 | 1.07E-23 | 5 |
| GCNT1 | 0.25405589 | 1.50E-23 | 5 |
| LRMP | 0.30341476 | 1.50E-23 | 5 |
| AL138899.1 | -0.7323287 | 3.32E-23 | 5 |
| ALDH5A1 | 0.26784709 | 3.38E-23 | 5 |
| LUZP1 | 0.30471224 | 3.51E-23 | 5 |
| DGKH | 0.3517048 | 4.12E-23 | 5 |
| VIM | 0.43106832 | 9.25E-23 | 5 |
| BCL11B | -0.4852888 | 1.09E-22 | 5 |
| RNASET2 | 0.30934044 | 1.14E-22 | 5 |
| SLC25A3 | 0.35697738 | 2.29E-22 | 5 |
| ARGLU1 | -0.4353593 | 2.78E-22 | 5 |
| KDM5B | 0.38474391 | 2.83E-22 | 5 |
| GOLIM4 | 0.28319885 | 2.88E-22 | 5 |
| MPG | 0.29046659 | 3.45E-22 | 5 |
| ARMCX6 | 0.2969891 | 3.58E-22 | 5 |
| SHISA2 | 0.39758008 | 5.24E-22 | 5 |
| BZW1 | 0.36635903 | 5.33E-22 | 5 |
| PHACTR2 | 0.36522834 | 6.80E-22 | 5 |
| MIB1 | 0.31552188 | 7.26E-22 | 5 |
| SHMT2 | 0.25421226 | 7.67E-22 | 5 |
| JARID2 | 0.38091409 | 7.69E-22 | 5 |
| SNHG6 | 0.32141962 | 1.18E-21 | 5 |
| BTG2 | -0.5488472 | 1.26E-21 | 5 |
| MYH10 | 0.28666603 | 1.43E-21 | 5 |
| PLEKHG1 | -0.5197872 | 3.21E-21 | 5 |
| MAGED2 | 0.2503901 | 3.75E-21 | 5 |
| TTC39C | 0.3613709 | 4.08E-21 | 5 |
| CD27 | 0.45683043 | 5.17E-21 | 5 |
| NEGR1 | -0.6108697 | 6.01E-21 | 5 |
| EEF2 | 0.28353704 | 7.94E-21 | 5 |
| CD63 | 0.33381923 | 8.44E-21 | 5 |
| GCLC | 0.26015247 | 9.97E-21 | 5 |
| RFLNB | -0.4665155 | 1.12E-20 | 5 |
| ETHE1 | 0.26454777 | 1.56E-20 | 5 |
| MRPS6 | 0.46431293 | 1.86E-20 | 5 |
| RNASEH2B | -0.4586973 | 2.43E-20 | 5 |
| VAMP5 | 0.29057381 | 2.46E-20 | 5 |

| | | | |
|---|---|---|---|
| SCRIB | -0.4955393 | 2.59E-20 | 5 |
| EGR1 | 0.5055938 | 3.16E-20 | 5 |
| ANXA1 | -0.5890168 | 3.77E-20 | 5 |
| TUBA1B | -0.8970078 | 3.81E-20 | 5 |
| INPP4A | -0.4589396 | 3.83E-20 | 5 |
| NPM1 | 0.29351639 | 4.67E-20 | 5 |
| RPL4 | 0.2672974 | 4.67E-20 | 5 |
| CCDC50 | 0.2705215 | 5.21E-20 | 5 |
| TRBC2 | -0.3261051 | 5.35E-20 | 5 |
| TCEAL4 | 0.29489904 | 7.07E-20 | 5 |
| FOXP1 | -0.5054743 | 1.86E-19 | 5 |
| TMEM256 | 0.30887036 | 2.18E-19 | 5 |
| N4BP2L2 | -0.4244282 | 2.50E-19 | 5 |
| SMC4 | -0.5818367 | 2.87E-19 | 5 |
| H1FX | -0.4933689 | 2.93E-19 | 5 |
| EIF3F | 0.35008763 | 3.57E-19 | 5 |
| RPL31 | 0.2669764 | 3.83E-19 | 5 |
| GATA3 | 0.48731444 | 7.11E-19 | 5 |
| TOP2B | 0.36140338 | 7.41E-19 | 5 |
| EIF3M | 0.31954348 | 7.60E-19 | 5 |
| AP005482.1 | -0.5320063 | 8.09E-19 | 5 |
| RPL27A | 0.25455112 | 9.16E-19 | 5 |
| HSP90AB1 | 0.31625296 | 1.16E-18 | 5 |
| EIF4A1 | 0.30337572 | 1.21E-18 | 5 |
| AC090152.1 | 0.34467589 | 1.24E-18 | 5 |
| RAP1A | 0.34300405 | 1.38E-18 | 5 |
| TRIM14 | -0.4900876 | 1.64E-18 | 5 |
| TMEM126B | 0.28152162 | 1.76E-18 | 5 |
| SIPA1L1 | 0.27258698 | 2.10E-18 | 5 |
| PHB2 | 0.30146879 | 4.45E-18 | 5 |
| ERO1B | 0.26451571 | 8.06E-18 | 5 |
| BRK1 | 0.30067014 | 8.49E-18 | 5 |
| SLC25A6 | 0.32394154 | 8.89E-18 | 5 |
| AL365440.2 | -0.4729119 | 9.20E-18 | 5 |
| KLF13 | -0.4079033 | 9.62E-18 | 5 |
| ERGIC3 | 0.2574838 | 9.78E-18 | 5 |
| BCOR | 0.35066171 | 1.59E-17 | 5 |
| TRAF3IP3 | -0.3952508 | 1.60E-17 | 5 |
| RTRAF | 0.31337898 | 1.61E-17 | 5 |
| MEF2D | 0.2553102 | 2.29E-17 | 5 |
| EEF1B2 | 0.27756944 | 3.67E-17 | 5 |
| S100A11 | 0.31218277 | 3.82E-17 | 5 |
| EIF3K | 0.29245319 | 4.02E-17 | 5 |

| | | | |
|---|---|---|---|
| CXCR4 | -0.4485054 | 7.16E-17 | 5 |
| ACADVL | 0.28503019 | 7.39E-17 | 5 |
| ATP5MC2 | 0.28738345 | 7.96E-17 | 5 |
| HELB | -0.399644 | 9.67E-17 | 5 |
| COX5A | 0.27545873 | 1.00E-16 | 5 |
| C19orf53 | 0.31643437 | 1.33E-16 | 5 |
| THEMIS | -0.4396702 | 1.92E-16 | 5 |
| CCNI | 0.28225017 | 2.00E-16 | 5 |
| GPRASP1 | 0.25869735 | 3.23E-16 | 5 |
| PBDC1 | 0.25851038 | 3.31E-16 | 5 |
| PRKAR1A | 0.30678693 | 3.81E-16 | 5 |
| TXN2 | 0.28150469 | 5.18E-16 | 5 |
| MDM4 | -0.42796 | 5.26E-16 | 5 |
| PARVG | 0.30207741 | 5.87E-16 | 5 |
| ROMO1 | 0.28674201 | 5.92E-16 | 5 |
| EVI2B | 0.29652737 | 8.17E-16 | 5 |
| BEX2 | 0.26017851 | 8.49E-16 | 5 |
| ARID5B | 0.31281949 | 8.90E-16 | 5 |
| IGFBP5 | -0.6601471 | 9.43E-16 | 5 |
| CCDC167 | -0.3805664 | 1.03E-15 | 5 |
| SLC9A3R1 | -0.3539749 | 1.12E-15 | 5 |
| CD37 | 0.28350301 | 1.38E-15 | 5 |
| RPS5 | 0.2583626 | 1.54E-15 | 5 |
| ARFGAP3 | 0.30289087 | 1.99E-15 | 5 |
| SYNRG | -0.4114643 | 3.65E-15 | 5 |
| CD1B | -0.5125546 | 3.90E-15 | 5 |
| PRMT7 | -0.4814194 | 3.99E-15 | 5 |
| CD47 | -0.3626486 | 4.24E-15 | 5 |
| NOSIP | -0.7048455 | 4.47E-15 | 5 |
| PRDX2 | 0.30497703 | 4.59E-15 | 5 |
| MAPRE2 | 0.30831896 | 6.49E-15 | 5 |
| CMTM6 | 0.2545539 | 7.10E-15 | 5 |
| RPL22L1 | 0.27631047 | 9.30E-15 | 5 |
| FAM89B | 0.2632538 | 9.83E-15 | 5 |
| NFATC3 | 0.32293142 | 1.77E-14 | 5 |
| SSR2 | 0.26813664 | 1.82E-14 | 5 |
| CLIC3 | -0.4808886 | 2.34E-14 | 5 |
| AKNA | -0.3696954 | 2.59E-14 | 5 |
| ZNF683 | -0.6660113 | 2.81E-14 | 5 |
| SESN3 | 0.29241741 | 3.85E-14 | 5 |
| PLXDC1 | -0.275315 | 4.93E-14 | 5 |
| SLFN5 | -0.3920103 | 5.85E-14 | 5 |
| MLXIP | -0.4108522 | 5.85E-14 | 5 |

| | | | |
|---|---|---|---|
| PLIN2 | 0.33916906 | 6.70E-14 | 5 |
| HIGD2A | 0.26356463 | 6.90E-14 | 5 |
| GALNT6 | -0.4090957 | 7.75E-14 | 5 |
| E2F2 | -0.300265 | 9.95E-14 | 5 |
| SEC31B | 0.25215585 | 1.04E-13 | 5 |
| HNRNPLL | 0.25760444 | 1.25E-13 | 5 |
| PYHIN1 | 0.28446741 | 1.51E-13 | 5 |
| LMO4 | 0.28493015 | 2.16E-13 | 5 |
| SLC5A3 | 0.41495932 | 2.68E-13 | 5 |
| CAPG | 0.28172745 | 2.82E-13 | 5 |
| SNHG5 | 0.27046922 | 2.89E-13 | 5 |
| REST | 0.26719494 | 3.80E-13 | 5 |
| MACF1 | 0.34735575 | 4.09E-13 | 5 |
| CR1 | -0.4563861 | 4.35E-13 | 5 |
| NFKBID | 0.26459374 | 4.40E-13 | 5 |
| SLA | -0.3927588 | 6.19E-13 | 5 |
| ZC3H8 | 0.25343449 | 9.98E-13 | 5 |
| SMCHD1 | -0.3477114 | 1.08E-12 | 5 |
| NSA2 | 0.28881025 | 1.70E-12 | 5 |
| NOP53 | 0.27310657 | 2.28E-12 | 5 |
| SLC12A6 | 0.29795361 | 2.97E-12 | 5 |
| RSL1D1 | 0.2543906 | 3.00E-12 | 5 |
| LDHB | 0.27518593 | 3.27E-12 | 5 |
| ADD3 | -0.5097417 | 5.09E-12 | 5 |
| TESPA1 | 0.25127407 | 5.37E-12 | 5 |
| BIN2 | -0.3728804 | 5.67E-12 | 5 |
| ETS1 | 0.2625951 | 5.90E-12 | 5 |
| ITGA6 | -0.3765767 | 6.70E-12 | 5 |
| FTH1 | -0.3107229 | 7.45E-12 | 5 |
| LSP1 | -0.4000671 | 1.04E-11 | 5 |
| MAP4K4 | 0.25451968 | 1.06E-11 | 5 |
| CD3G | -0.2930368 | 1.46E-11 | 5 |
| CASC15 | 0.31344835 | 1.46E-11 | 5 |
| CD1A | -0.5561633 | 1.69E-11 | 5 |
| IRF2BPL | -0.3075413 | 1.69E-11 | 5 |
| ST13 | 0.27687265 | 2.08E-11 | 5 |
| FYB1 | -0.2662314 | 2.08E-11 | 5 |
| EVI2A | 0.2667739 | 2.14E-11 | 5 |
| SMDT1 | 0.28628598 | 3.61E-11 | 5 |
| GSE1 | 0.28348425 | 6.35E-11 | 5 |
| AL365361.1 | -0.3876617 | 6.55E-11 | 5 |
| VPS13C | -0.3604438 | 6.88E-11 | 5 |
| GPX4 | 0.27697546 | 7.30E-11 | 5 |

| | | | |
|---|---|---|---|
| FYN | 0.27800992 | 7.96E-11 | 5 |
| RNF213 | -0.3462247 | 9.37E-11 | 5 |
| ITGAE | 0.27562061 | 9.91E-11 | 5 |
| UBASH3B | 0.31885312 | 1.14E-10 | 5 |
| DCP2 | 0.26980744 | 1.27E-10 | 5 |
| AUTS2 | -0.3318958 | 1.43E-10 | 5 |
| NCF1 | 0.26224015 | 1.44E-10 | 5 |
| SPRY1 | -0.423384 | 1.71E-10 | 5 |
| SLC3A2 | 0.27621811 | 2.21E-10 | 5 |
| HIST1H1D | -0.5610012 | 2.36E-10 | 5 |
| NCAPG2 | -0.4005743 | 2.60E-10 | 5 |
| STK17A | 0.25983333 | 2.81E-10 | 5 |
| TLE5 | -0.2725509 | 3.58E-10 | 5 |
| CR2 | -0.3190954 | 4.00E-10 | 5 |
| CLIC1 | 0.27091109 | 4.61E-10 | 5 |
| GIMAP7 | -0.4544873 | 5.61E-10 | 5 |
| KLF6 | -0.4770462 | 8.12E-10 | 5 |
| AQP3 | -0.4035869 | 1.06E-09 | 5 |
| SEPTIN9 | -0.2936241 | 1.30E-09 | 5 |
| MTRNR2L12 | -0.2930325 | 1.60E-09 | 5 |
| HIST1H1E | -0.4039281 | 2.45E-09 | 5 |
| PRKCQ-AS1 | -0.3277396 | 2.62E-09 | 5 |
| SAMD12 | -0.3306819 | 2.79E-09 | 5 |
| PTP4A2 | -0.302679 | 4.14E-09 | 5 |
| IRF1 | -0.3337897 | 4.40E-09 | 5 |
| AC068587.4 | -0.3889736 | 4.77E-09 | 5 |
| GIMAP4 | -0.2844283 | 5.29E-09 | 5 |
| SPN | 0.25715199 | 5.45E-09 | 5 |
| IPCEF1 | 0.34229824 | 6.94E-09 | 5 |
| KLRK1 | -0.33086 | 8.06E-09 | 5 |
| ACAP1 | -0.2805008 | 1.03E-08 | 5 |
| MTF2 | 0.27338234 | 1.28E-08 | 5 |
| RGCC | 0.28097115 | 1.49E-08 | 5 |
| ZNF280D | -0.3592592 | 1.67E-08 | 5 |
| PRDX1 | 0.28504977 | 1.70E-08 | 5 |
| FBLN2 | -0.2696944 | 2.67E-08 | 5 |
| EIF4A2 | -0.2703892 | 2.82E-08 | 5 |
| RUNX1 | -0.2998315 | 3.49E-08 | 5 |
| ADA | -0.3243188 | 4.42E-08 | 5 |
| HELLS | -0.4380496 | 4.72E-08 | 5 |
| HMGB1 | -0.4236763 | 5.00E-08 | 5 |
| CD79A | -0.4145959 | 6.60E-08 | 5 |
| TNNT3 | -0.3178173 | 7.29E-08 | 5 |

| | | | |
|---|---|---|---|
| XIST | -0.3764745 | 7.60E-08 | 5 |
| TFDP2 | -0.4700866 | 8.22E-08 | 5 |
| SLC38A1 | 0.27937479 | 8.85E-08 | 5 |
| PLAC8 | -0.4979026 | 1.01E-07 | 5 |
| ADGRE5 | 0.2784298 | 1.08E-07 | 5 |
| CHMP7 | -0.3737935 | 1.17E-07 | 5 |
| TIAM1 | -0.3370721 | 1.30E-07 | 5 |
| RAD21 | -0.3457386 | 1.51E-07 | 5 |
| NMT2 | -0.3130368 | 1.90E-07 | 5 |
| CIRBP | -0.2756567 | 1.94E-07 | 5 |
| TYMS | -0.6507593 | 1.97E-07 | 5 |
| TRIM56 | -0.3323658 | 2.81E-07 | 5 |
| ITGB1 | -0.3586138 | 3.04E-07 | 5 |
| ANKRD44 | -0.3193781 | 4.67E-07 | 5 |
| CDK5RAP3 | -0.3500851 | 4.98E-07 | 5 |
| ERAP2 | -0.2934675 | 5.55E-07 | 5 |
| APBB1IP | -0.272611 | 6.91E-07 | 5 |
| PSAP | -0.2874358 | 7.34E-07 | 5 |
| DNMT1 | -0.317748 | 7.93E-07 | 5 |
| DDX6 | -0.2685923 | 9.67E-07 | 5 |
| ATP2A3 | -0.2823167 | 1.09E-06 | 5 |
| DUT | -0.502016 | 1.22E-06 | 5 |
| TNFAIP3 | -0.3773183 | 1.32E-06 | 5 |
| MKI67 | -0.6970143 | 1.72E-06 | 5 |
| TCF12 | -0.3084226 | 1.76E-06 | 5 |
| ERAP1 | -0.2710281 | 2.34E-06 | 5 |
| MPHOSPH8 | -0.2766056 | 2.49E-06 | 5 |
| POLR2J3 | -0.2799008 | 2.66E-06 | 5 |
| C12orf75 | -0.3129858 | 3.40E-06 | 5 |
| COL6A2 | -0.3340012 | 3.96E-06 | 5 |
| CASP8 | -0.3146485 | 4.13E-06 | 5 |
| MAN2A1 | -0.3037616 | 4.44E-06 | 5 |
| LITAF | -0.2819552 | 5.97E-06 | 5 |
| TLE1 | -0.2581145 | 1.40E-05 | 5 |
| IL7R | -0.2659097 | 1.72E-05 | 5 |
| GLIPR2 | -0.2797956 | 1.93E-05 | 5 |
| RIPOR2 | -0.3807494 | 2.19E-05 | 5 |
| CDC42EP3 | -0.2891883 | 3.09E-05 | 5 |
| PNISR | -0.2536911 | 3.23E-05 | 5 |
| EPB41 | -0.2901305 | 5.89E-05 | 5 |
| PTGDR | -0.3092802 | 6.30E-05 | 5 |
| DNTT | -0.3046081 | 7.36E-05 | 5 |
| CD99 | -0.2783935 | 0.00011883 | 5 |

| | | | |
|---|---|---|---|
| WDR76 | -0.2729027 | 0.00013246 | 5 |
| RESF1 | -0.321868 | 0.00014103 | 5 |
| EDEM1 | -0.2963888 | 0.00018264 | 5 |
| NPNT | -0.2688596 | 0.00018367 | 5 |
| PAXX | -0.290354 | 0.00020648 | 5 |
| YPEL3 | -0.2816411 | 0.00027636 | 5 |
| PAG1 | -0.2624304 | 0.00035622 | 5 |
| ETV5 | -0.3709113 | 0.00039021 | 5 |
| SCAI | -0.2683426 | 0.00042402 | 5 |
| UHRF1 | -0.2743833 | 0.0004885 | 5 |
| CLSPN | -0.3311854 | 0.00055967 | 5 |
| SNRK | -0.2576375 | 0.00071017 | 5 |
| IQGAP2 | -0.2672074 | 0.00078825 | 5 |
| SLFN13 | -0.267275 | 0.00094441 | 5 |
| ARHGAP26 | -0.2634331 | 0.00103131 | 5 |
| ANKRD12 | -0.2623118 | 0.00127796 | 5 |
| DYRK2 | -0.250834 | 0.00147834 | 5 |
| KLF2 | -0.4781854 | 0.00205578 | 5 |
| CDK1 | -0.2672445 | 0.00262827 | 5 |
| CENPF | -0.6199432 | 0.00314901 | 5 |
| DLEU2 | -0.2929096 | 0.00347872 | 5 |
| HIST1H1C | -0.4825285 | 0.00467605 | 5 |
| C21orf58 | -0.2685036 | 0.00502208 | 5 |
| ARPP21 | -0.302856 | 0.00563181 | 5 |
| EML4 | -0.2874683 | 0.01526443 | 5 |
| ZFP36L1 | 0.342597 | 0.01879151 | 5 |
| MCM4 | -0.2643458 | 0.01917086 | 5 |
| LINC00342 | -0.3002587 | 0.0197399 | 5 |
| GTSE1 | -0.3125593 | 0.02113057 | 5 |
| ELOVL4 | -0.3128532 | 0.03659853 | 5 |
| CNN2 | -0.3092772 | 0.06129563 | 5 |
| ATAD2 | -0.3610174 | 0.06196018 | 5 |
| PPARA | -0.263442 | 0.06637709 | 5 |
| PCNA | -0.3883722 | 0.07870464 | 5 |
| TTN | -0.3047817 | 0.09562307 | 5 |
| NUSAP1 | -0.5261886 | 0.18643483 | 5 |
| TMPO | -0.2986892 | 0.19192445 | 5 |
| ASPM | -0.4734887 | 0.75347756 | 5 |
| CFLAR | -0.2570192 | 0.77276066 | 5 |
| TECR | -0.3767266 | 0.78447094 | 5 |
| USP1 | -0.2820136 | 0.85216322 | 5 |
| HMGN2 | -0.3290987 | 0.98330794 | 5 |
| RRM2 | -0.3335968 | 1 | 5 |

| | | | |
|---|---|---|---|
| EMP3 | -0.2542822 | 1 | 5 |
| SYNE2 | -0.310411 | 1 | 5 |
| TOP2A | -0.6727937 | 1 | 5 |
| HIST1H1B | -0.478833 | 1 | 5 |
| FARS2 | -0.2827264 | 1 | 5 |
| RORB | -0.2757411 | 1 | 5 |
| MCM7 | -0.2808606 | 1 | 5 |
| HIST1H3B | -0.3041379 | 1 | 5 |
| TMSB10 | -0.2706791 | 1 | 5 |
| XIST | 1.03164942 | 1.39E-119 | 6 |
| MALAT1 | 0.64804332 | 2.88E-113 | 6 |
| RPL41 | -0.7309056 | 6.39E-103 | 6 |
| RPL18A | -0.7050859 | 4.55E-97 | 6 |
| RPS15 | -0.6614962 | 7.24E-97 | 6 |
| RPS12 | -0.801225 | 1.66E-95 | 6 |
| RPL13 | -0.7143504 | 3.58E-95 | 6 |
| RPS18 | -0.7082217 | 2.08E-94 | 6 |
| RPS3A | -0.661651 | 8.23E-94 | 6 |
| RPLP1 | -0.7334024 | 1.95E-91 | 6 |
| EEF1A1 | -0.6914129 | 3.94E-91 | 6 |
| RPS3 | -0.6637035 | 4.03E-91 | 6 |
| RPL28 | -0.6478672 | 4.37E-88 | 6 |
| TPT1 | -0.7179661 | 8.98E-88 | 6 |
| RPS23 | -0.6428001 | 1.81E-87 | 6 |
| RPL32 | -0.6853503 | 5.26E-87 | 6 |
| RPS27A | -0.6707025 | 4.57E-85 | 6 |
| RPS28 | -0.6235245 | 1.32E-84 | 6 |
| RPL37A | -0.5539618 | 5.67E-84 | 6 |
| RPS4X | -0.6661261 | 1.04E-83 | 6 |
| RPL23A | -0.6056719 | 1.86E-82 | 6 |
| RPL10 | -0.7126357 | 1.95E-82 | 6 |
| RPS21 | -0.604108 | 3.68E-82 | 6 |
| RPS15A | -0.6728628 | 7.67E-82 | 6 |
| RPS27 | -0.7194378 | 4.64E-81 | 6 |
| RPS14 | -0.666078 | 1.45E-80 | 6 |
| RPS6 | -0.6216337 | 6.62E-80 | 6 |
| RPS9 | -0.5970408 | 4.61E-79 | 6 |
| RPS16 | -0.5805154 | 1.00E-78 | 6 |
| RPL19 | -0.6132916 | 2.80E-78 | 6 |
| RPS7 | -0.6159976 | 6.09E-78 | 6 |
| RPL26 | -0.66484 | 7.56E-77 | 6 |
| RPS24 | -0.6087511 | 5.49E-76 | 6 |
| RPS8 | -0.6254461 | 6.34E-76 | 6 |

| | | | |
|---|---|---|---|
| RPL35 | -0.6024423 | 4.42E-75 | 6 |
| RPL37 | -0.6310477 | 1.22E-74 | 6 |
| RPS5 | -0.6933338 | 2.40E-74 | 6 |
| TMSB10 | -0.9214205 | 6.86E-74 | 6 |
| RPS2 | -0.6310563 | 9.34E-74 | 6 |
| MACF1 | 0.84617174 | 1.37E-72 | 6 |
| RPL8 | -0.5813828 | 5.25E-71 | 6 |
| RPL6 | -0.6098524 | 8.53E-71 | 6 |
| RPL7A | -0.5672006 | 1.03E-69 | 6 |
| RPL15 | -0.5631697 | 1.34E-69 | 6 |
| RPL10A | -0.5987519 | 1.57E-69 | 6 |
| RPL11 | -0.5681892 | 2.55E-69 | 6 |
| RPL35A | -0.6031511 | 2.83E-69 | 6 |
| RPL18 | -0.5631989 | 9.25E-69 | 6 |
| RPL39 | -0.5939549 | 2.71E-68 | 6 |
| RPL30 | -0.5967734 | 2.50E-67 | 6 |
| RPS19 | -0.5534653 | 1.04E-66 | 6 |
| TMSB4X | -0.6325303 | 1.63E-66 | 6 |
| RPLP2 | -0.5246422 | 7.47E-65 | 6 |
| FTH1 | -0.659168 | 2.13E-64 | 6 |
| RPL27 | -0.5550509 | 2.06E-63 | 6 |
| ARGLU1 | 0.65895177 | 2.13E-63 | 6 |
| RPL21 | -0.5365214 | 7.91E-63 | 6 |
| PTMA | -0.6198387 | 8.97E-63 | 6 |
| RPL29 | -0.5635831 | 9.76E-63 | 6 |
| RPL24 | -0.5788849 | 9.50E-62 | 6 |
| RPL3 | -0.5171802 | 2.45E-61 | 6 |
| RPL5 | -0.5800782 | 2.72E-61 | 6 |
| RPS13 | -0.5737307 | 4.20E-61 | 6 |
| FAU | -0.531761 | 1.92E-60 | 6 |
| RPSA | -0.5550855 | 3.66E-60 | 6 |
| RPL36A | -0.5851484 | 2.54E-59 | 6 |
| SERF2 | -0.5152854 | 2.62E-59 | 6 |
| UBA52 | -0.5151092 | 5.59E-59 | 6 |
| RPS25 | -0.5289505 | 9.73E-59 | 6 |
| RACK1 | -0.5438764 | 6.93E-58 | 6 |
| N4BP2L2 | 0.66472859 | 1.23E-57 | 6 |
| ATP5F1E | -0.5488812 | 1.28E-57 | 6 |
| ACTG1 | -0.5629709 | 2.38E-57 | 6 |
| RPL17 | -0.5186199 | 1.04E-56 | 6 |
| RPL36 | -0.5240506 | 6.17E-56 | 6 |
| RPL9 | -0.5066558 | 6.30E-56 | 6 |
| RPL34 | -0.558713 | 9.17E-56 | 6 |

| | | | |
|---|---|---|---|
| RPL7 | -0.510435 | 1.96E-55 | 6 |
| TTN | 0.96293909 | 8.10E-55 | 6 |
| NACA | -0.5353868 | 1.15E-53 | 6 |
| RPS29 | -0.4932884 | 1.44E-53 | 6 |
| MYL6 | -0.5109032 | 4.99E-53 | 6 |
| RPL12 | -0.5711123 | 5.28E-53 | 6 |
| COX7C | -0.5102628 | 8.62E-53 | 6 |
| RPL13A | -0.475163 | 9.88E-53 | 6 |
| IKZF2 | 0.80598386 | 1.12E-52 | 6 |
| SRP14 | -0.5189794 | 5.03E-51 | 6 |
| CFL1 | -0.4848398 | 2.41E-50 | 6 |
| EEF1B2 | -0.5695716 | 3.73E-50 | 6 |
| RPL14 | -0.5809254 | 6.42E-50 | 6 |
| RPS10 | -0.5339086 | 2.59E-49 | 6 |
| RPL22 | -0.492434 | 2.64E-49 | 6 |
| RPLP0 | -0.6392201 | 3.30E-49 | 6 |
| YBX1 | -0.5957099 | 8.51E-49 | 6 |
| PFN1 | -0.5024947 | 8.91E-48 | 6 |
| RPS20 | -0.4705721 | 1.21E-47 | 6 |
| RPL38 | -0.4675952 | 7.77E-47 | 6 |
| COX4I1 | -0.4880749 | 1.22E-45 | 6 |
| ATP5MG | -0.4711597 | 1.65E-45 | 6 |
| RPL4 | -0.4964626 | 3.14E-45 | 6 |
| CLIC1 | -0.5777137 | 1.32E-44 | 6 |
| H3F3A | -0.4112573 | 6.85E-44 | 6 |
| FTX | 0.65954562 | 1.02E-43 | 6 |
| BTF3 | -0.5324378 | 1.46E-43 | 6 |
| DDX17 | 0.48382503 | 1.88E-43 | 6 |
| MT-ND3 | 0.61036642 | 1.71E-42 | 6 |
| PPIA | -0.5870515 | 8.13E-42 | 6 |
| MT-ATP6 | 0.57504087 | 9.22E-42 | 6 |
| EEF1G | -0.4998186 | 2.27E-41 | 6 |
| SUMO2 | -0.4851608 | 4.35E-41 | 6 |
| UQCRH | -0.4851445 | 5.01E-41 | 6 |
| UBL5 | -0.4559569 | 7.58E-41 | 6 |
| MTRNR2L12 | 0.64846732 | 1.21E-40 | 6 |
| RPS26 | -0.4438777 | 3.25E-40 | 6 |
| HINT1 | -0.4995307 | 7.58E-40 | 6 |
| MZT2B | -0.5040799 | 2.57E-39 | 6 |
| TMA7 | -0.4189411 | 6.62E-39 | 6 |
| UBB | -0.4840132 | 1.50E-38 | 6 |
| SNHG14 | 0.67442992 | 2.43E-38 | 6 |
| ATM | 0.58561412 | 3.19E-38 | 6 |

| | | | |
|---|---|---|---|
| ARPC3 | -0.4445254 | 8.50E-38 | 6 |
| CD52 | -0.6341314 | 7.91E-37 | 6 |
| RPS17 | -0.4626974 | 6.74E-36 | 6 |
| TMEM161B-AS: | 0.6842021 | 7.15E-36 | 6 |
| GNAS | -0.4108534 | 9.52E-35 | 6 |
| RPL36AL | -0.4416521 | 9.90E-35 | 6 |
| AAK1 | 0.52979157 | 1.71E-33 | 6 |
| KAT6B | 0.51882179 | 2.34E-33 | 6 |
| RPS11 | -0.3685987 | 2.82E-32 | 6 |
| LINC01222 | 0.72645508 | 3.25E-32 | 6 |
| MYL12B | -0.4093658 | 3.59E-32 | 6 |
| ACTB | -0.3855119 | 3.67E-32 | 6 |
| EEF1D | -0.3998821 | 1.53E-31 | 6 |
| PSMA7 | -0.4785987 | 1.82E-31 | 6 |
| ATP5MC2 | -0.4311949 | 2.05E-31 | 6 |
| RPL27A | -0.4044422 | 2.26E-31 | 6 |
| JPX | 0.51541117 | 3.61E-31 | 6 |
| VIM | -0.8543641 | 4.41E-31 | 6 |
| ATP5MF | -0.4337192 | 5.20E-31 | 6 |
| SH3BGRL3 | -0.5777332 | 1.06E-30 | 6 |
| PNISR | 0.50868921 | 1.26E-30 | 6 |
| RPL23 | -0.4216957 | 1.59E-30 | 6 |
| GAPDH | -0.6664622 | 2.87E-30 | 6 |
| MT-CO2 | 0.46100031 | 3.06E-30 | 6 |
| MIF | -0.5278911 | 8.08E-30 | 6 |
| UQCRB | -0.4282084 | 1.08E-29 | 6 |
| RPL31 | -0.4002482 | 1.55E-29 | 6 |
| ANAPC11 | -0.4920151 | 2.92E-29 | 6 |
| NDUFB2 | -0.452931 | 4.42E-29 | 6 |
| POLR2J3 | 0.5016252 | 7.12E-29 | 6 |
| SMG1 | 0.47942147 | 7.28E-29 | 6 |
| EIF1 | -0.3870316 | 8.03E-29 | 6 |
| SATB1 | 0.56890673 | 8.68E-29 | 6 |
| ELOB | -0.4138796 | 1.31E-28 | 6 |
| PSMB6 | -0.4003008 | 1.76E-28 | 6 |
| ARHGDIB | -0.339764 | 1.99E-28 | 6 |
| PSMB4 | -0.4159963 | 2.76E-28 | 6 |
| MT-ND4 | 0.47467931 | 3.84E-28 | 6 |
| CASC15 | 0.51416792 | 5.30E-28 | 6 |
| NPM1 | -0.4694002 | 7.03E-28 | 6 |
| TTC14 | 0.47863434 | 7.38E-28 | 6 |
| DBI | -0.4725753 | 1.17E-27 | 6 |
| COX6C | -0.4232514 | 2.13E-27 | 6 |

| | | | |
|---|---|---|---|
| SF3B6 | -0.4582921 | 3.93E-27 | 6 |
| ANP32B | -0.4851022 | 4.66E-27 | 6 |
| MYL12A | -0.428315 | 5.01E-27 | 6 |
| ATP5PD | -0.4788904 | 8.60E-27 | 6 |
| SEC61G | -0.4118628 | 1.44E-26 | 6 |
| PFDN5 | -0.3831042 | 1.63E-26 | 6 |
| CD3D | -0.3654182 | 2.81E-26 | 6 |
| POLR2L | -0.4447389 | 5.00E-26 | 6 |
| MT-CO3 | 0.4202708 | 7.56E-26 | 6 |
| DIP2A | 0.49344408 | 9.40E-26 | 6 |
| TOMM6 | -0.4397942 | 1.06E-25 | 6 |
| SNHG6 | -0.4190966 | 1.21E-25 | 6 |
| SCAND1 | -0.3971873 | 1.34E-25 | 6 |
| HCST | -0.5224845 | 2.68E-25 | 6 |
| NKTR | 0.51268165 | 4.39E-25 | 6 |
| PSMB3 | -0.4340262 | 5.17E-25 | 6 |
| SNHG29 | -0.5286679 | 5.19E-25 | 6 |
| RNF213 | 0.46787399 | 5.26E-25 | 6 |
| COX7A2L | -0.4166854 | 6.50E-25 | 6 |
| SLC25A5 | -0.4461335 | 6.91E-25 | 6 |
| PFDN2 | -0.390774 | 7.20E-25 | 6 |
| NDUFA6 | -0.4124937 | 8.08E-25 | 6 |
| SMIM26 | -0.411756 | 1.16E-24 | 6 |
| TALDO1 | -0.4370722 | 1.23E-24 | 6 |
| MICOS10 | -0.4106728 | 1.57E-24 | 6 |
| PTPRC | 0.38011966 | 1.85E-24 | 6 |
| NDUFAB1 | -0.3967079 | 1.89E-24 | 6 |
| LSM4 | -0.4655858 | 2.11E-24 | 6 |
| AP2S1 | -0.3698618 | 2.14E-24 | 6 |
| C17orf49 | -0.3734706 | 2.66E-24 | 6 |
| VDAC1 | -0.4452903 | 2.78E-24 | 6 |
| TRBC2 | -0.3483317 | 3.27E-24 | 6 |
| PSMB2 | -0.3966722 | 3.93E-24 | 6 |
| RHOA | -0.4050899 | 6.48E-24 | 6 |
| CD99 | -0.5727535 | 7.46E-24 | 6 |
| OST4 | -0.356431 | 9.48E-24 | 6 |
| LSM3 | -0.4179406 | 1.32E-23 | 6 |
| PSMB10 | -0.3961375 | 1.61E-23 | 6 |
| POLR2G | -0.3796427 | 1.63E-23 | 6 |
| FIS1 | -0.3993734 | 1.90E-23 | 6 |
| SMC4 | 0.51521999 | 2.49E-23 | 6 |
| MAGOH | -0.3859853 | 3.50E-23 | 6 |
| EIF5A | -0.445192 | 3.79E-23 | 6 |

| | | | |
|---|---|---|---|
| LDHB | -0.4382131 | 3.86E-23 | 6 |
| NDUFS5 | -0.4265892 | 4.14E-23 | 6 |
| TUT4 | 0.45172586 | 1.00E-22 | 6 |
| MT-CYB | 0.48136206 | 1.17E-22 | 6 |
| SNRPD2 | -0.4209805 | 1.45E-22 | 6 |
| KANSL1 | 0.44025062 | 1.47E-22 | 6 |
| MT-ND2 | 0.47350564 | 1.47E-22 | 6 |
| EIF3I | -0.3825346 | 1.71E-22 | 6 |
| GADD45GIP1 | -0.3690174 | 1.88E-22 | 6 |
| POMP | -0.4273687 | 2.42E-22 | 6 |
| NDUFA12 | -0.3982687 | 3.34E-22 | 6 |
| RALY | -0.3969744 | 3.64E-22 | 6 |
| STMN1 | -0.6141877 | 3.90E-22 | 6 |
| PGLS | -0.411474 | 4.33E-22 | 6 |
| ATP5ME | -0.3943555 | 4.80E-22 | 6 |
| CSNK2B | -0.412309 | 5.05E-22 | 6 |
| CALM1 | -0.4628085 | 5.28E-22 | 6 |
| NDUFA7 | -0.3481372 | 5.53E-22 | 6 |
| HNRNPA1 | -0.4430994 | 5.71E-22 | 6 |
| SYNE1 | 0.5010619 | 5.92E-22 | 6 |
| NDUFB3 | -0.3886323 | 8.11E-22 | 6 |
| TIMM8B | -0.3727694 | 1.20E-21 | 6 |
| PGAM1 | -0.3665225 | 1.24E-21 | 6 |
| GSTO1 | -0.3659602 | 1.51E-21 | 6 |
| TOMM5 | -0.3761065 | 3.45E-21 | 6 |
| MT-ND1 | 0.51308099 | 4.79E-21 | 6 |
| NDUFA13 | -0.3534399 | 5.39E-21 | 6 |
| MT-CO1 | 0.34722421 | 5.93E-21 | 6 |
| MDM4 | 0.45513385 | 7.13E-21 | 6 |
| TIMM13 | -0.3609143 | 7.72E-21 | 6 |
| NDUFS6 | -0.4005963 | 9.03E-21 | 6 |
| C1QBP | -0.4130268 | 9.63E-21 | 6 |
| PPA1 | -0.4089203 | 9.92E-21 | 6 |
| DOCK10 | 0.43210975 | 1.15E-20 | 6 |
| FTL | -0.475108 | 1.20E-20 | 6 |
| BANF1 | -0.3864581 | 1.27E-20 | 6 |
| SF3B5 | -0.3835657 | 1.42E-20 | 6 |
| POLR2J | -0.3706528 | 1.47E-20 | 6 |
| NDUFB10 | -0.3433553 | 1.75E-20 | 6 |
| HSPA8 | -0.3806005 | 2.02E-20 | 6 |
| RAB5IF | -0.3398969 | 2.17E-20 | 6 |
| PSMB8 | -0.376142 | 2.23E-20 | 6 |
| PPDPF | -0.4251873 | 2.27E-20 | 6 |

| | | | |
|---|---|---|---|
| OGT | 0.41851255 | 2.41E-20 | 6 |
| LAMTOR1 | -0.3302735 | 2.67E-20 | 6 |
| PSMA3-AS1 | 0.43224396 | 2.75E-20 | 6 |
| COX7A2 | -0.3890798 | 3.43E-20 | 6 |
| SMS | -0.3815911 | 3.49E-20 | 6 |
| TGOLN2 | 0.41560887 | 3.74E-20 | 6 |
| ILF2 | -0.4199391 | 3.80E-20 | 6 |
| COX6A1 | -0.3634594 | 4.20E-20 | 6 |
| TSPO | -0.4161597 | 4.63E-20 | 6 |
| NOP53 | -0.4049666 | 6.42E-20 | 6 |
| PRDX1 | -0.3940621 | 7.15E-20 | 6 |
| DCTN3 | -0.3545007 | 7.88E-20 | 6 |
| NME2 | -0.5793522 | 8.01E-20 | 6 |
| TNRC6B | 0.42191094 | 8.98E-20 | 6 |
| ATP5MC3 | -0.3931839 | 9.64E-20 | 6 |
| SLFN5 | 0.436835 | 1.15E-19 | 6 |
| ZNF207 | 0.41176951 | 1.54E-19 | 6 |
| ELOC | -0.3261182 | 1.70E-19 | 6 |
| SNRPC | -0.3183702 | 1.76E-19 | 6 |
| NDUFA2 | -0.3691876 | 1.77E-19 | 6 |
| AURKAIP1 | -0.3294835 | 2.08E-19 | 6 |
| NDUFA4 | -0.3750462 | 2.13E-19 | 6 |
| PSMB7 | -0.3282075 | 2.41E-19 | 6 |
| PDE7A | 0.38555587 | 2.93E-19 | 6 |
| PPP4C | -0.3217554 | 3.09E-19 | 6 |
| DYNLRB1 | -0.3544122 | 3.67E-19 | 6 |
| SNU13 | -0.3771527 | 3.85E-19 | 6 |
| CAMK4 | 0.45295515 | 3.86E-19 | 6 |
| LSM2 | -0.3551352 | 4.00E-19 | 6 |
| ATRX | 0.41217871 | 4.24E-19 | 6 |
| MIEN1 | -0.3151543 | 5.36E-19 | 6 |
| ZMAT2 | -0.3474462 | 9.21E-19 | 6 |
| NHP2 | -0.3585266 | 9.79E-19 | 6 |
| NDUFC1 | -0.338212 | 1.02E-18 | 6 |
| ROMO1 | -0.3695989 | 1.03E-18 | 6 |
| ATP5PB | -0.3362384 | 1.06E-18 | 6 |
| ZNF428 | -0.3661802 | 1.09E-18 | 6 |
| KRT10 | -0.3651274 | 1.10E-18 | 6 |
| DYNLL1 | -0.4219803 | 1.13E-18 | 6 |
| NUDC | -0.3569821 | 1.13E-18 | 6 |
| TAF10 | -0.3400291 | 1.19E-18 | 6 |
| MRPL54 | -0.3344457 | 1.29E-18 | 6 |
| COMMD6 | -0.3590097 | 1.33E-18 | 6 |

| | | | |
|---|---|---|---|
| GABARAPL2 | -0.3130903 | 1.54E-18 | 6 |
| SUB1 | -0.3573728 | 1.56E-18 | 6 |
| FKBP1A | -0.3618848 | 1.85E-18 | 6 |
| DDT | -0.3745996 | 1.97E-18 | 6 |
| PSMC3 | -0.3436054 | 2.43E-18 | 6 |
| CLEC2D | 0.38798124 | 2.63E-18 | 6 |
| EVL | 0.34911423 | 3.08E-18 | 6 |
| SNRPD1 | -0.3878566 | 3.15E-18 | 6 |
| RGCC | -0.4597471 | 3.46E-18 | 6 |
| OAZ1 | -0.3229362 | 4.37E-18 | 6 |
| ARID1B | 0.41891853 | 5.80E-18 | 6 |
| MRPL33 | -0.3242269 | 6.24E-18 | 6 |
| PHF5A | -0.3102232 | 6.85E-18 | 6 |
| NAA38 | -0.3507155 | 6.86E-18 | 6 |
| MRPL51 | -0.344687 | 7.07E-18 | 6 |
| TXNDC17 | -0.3311898 | 7.52E-18 | 6 |
| BCL11B | 0.40721256 | 7.57E-18 | 6 |
| COX6B1 | -0.3267427 | 8.06E-18 | 6 |
| SNRPB2 | -0.372401 | 9.19E-18 | 6 |
| CHCHD2 | -0.3793421 | 9.89E-18 | 6 |
| S100A11 | -0.6377935 | 1.06E-17 | 6 |
| LAGE3 | -0.3049591 | 1.26E-17 | 6 |
| OGA | 0.41762243 | 1.38E-17 | 6 |
| SNRPE | -0.414253 | 1.49E-17 | 6 |
| FBL | -0.3544064 | 1.49E-17 | 6 |
| ITGB1BP1 | -0.3228902 | 1.49E-17 | 6 |
| TMEM258 | -0.3702817 | 1.81E-17 | 6 |
| VAMP8 | -0.2977292 | 2.07E-17 | 6 |
| ORAI1 | -0.3345055 | 2.07E-17 | 6 |
| VDAC2 | -0.3251138 | 2.10E-17 | 6 |
| TXNL4A | -0.3399207 | 2.30E-17 | 6 |
| APEX1 | -0.3744717 | 2.31E-17 | 6 |
| NDUFB9 | -0.3345626 | 2.47E-17 | 6 |
| LY6E | -0.3537488 | 2.49E-17 | 6 |
| MCRIP1 | -0.338241 | 2.59E-17 | 6 |
| TMEM219 | -0.330191 | 2.99E-17 | 6 |
| SEM1 | -0.3356602 | 3.25E-17 | 6 |
| FAAP20 | -0.3180681 | 3.28E-17 | 6 |
| UQCR11 | -0.3192692 | 3.43E-17 | 6 |
| SNRPB | -0.4174141 | 3.63E-17 | 6 |
| CIAO2B | -0.3175669 | 3.76E-17 | 6 |
| AP2M1 | -0.3258942 | 3.80E-17 | 6 |
| ARHGDIA | -0.3336322 | 5.21E-17 | 6 |

| | | | |
|---|---|---|---|
| COX14 | -0.3105291 | 5.71E-17 | 6 |
| NEAT1 | 0.43961267 | 5.87E-17 | 6 |
| RSL24D1 | -0.3242674 | 5.98E-17 | 6 |
| GTF3A | -0.377188 | 6.30E-17 | 6 |
| GTF3C6 | -0.3031737 | 7.99E-17 | 6 |
| PCNA | -0.5476662 | 8.15E-17 | 6 |
| PLIN2 | -0.3946822 | 8.82E-17 | 6 |
| MRPS36 | -0.3055692 | 9.48E-17 | 6 |
| NDUFB11 | -0.3988313 | 1.09E-16 | 6 |
| UQCRFS1 | -0.3121651 | 1.12E-16 | 6 |
| NDUFS8 | -0.3235908 | 1.12E-16 | 6 |
| CORO1A | -0.3405436 | 1.24E-16 | 6 |
| PRDX6 | -0.3198361 | 1.38E-16 | 6 |
| ESD | -0.3188153 | 1.42E-16 | 6 |
| TOMM20 | -0.3160347 | 1.47E-16 | 6 |
| RSL1D1 | -0.3205655 | 1.55E-16 | 6 |
| CCT7 | -0.3135448 | 1.83E-16 | 6 |
| TBCB | -0.3230307 | 1.93E-16 | 6 |
| NDUFB6 | -0.3105723 | 2.05E-16 | 6 |
| MRPL18 | -0.2882138 | 2.30E-16 | 6 |
| CCT8 | -0.3385787 | 2.40E-16 | 6 |
| HIGD2A | -0.3070625 | 2.56E-16 | 6 |
| SUCLG1 | -0.2961249 | 2.58E-16 | 6 |
| TXN | -0.4819521 | 2.66E-16 | 6 |
| BLOC1S1 | -0.3601152 | 3.00E-16 | 6 |
| ARPP19 | -0.3327819 | 3.16E-16 | 6 |
| FXYD2 | -1.0837055 | 3.43E-16 | 6 |
| RPS19BP1 | -0.3037908 | 4.84E-16 | 6 |
| DEK | -0.3935236 | 5.06E-16 | 6 |
| MDH2 | -0.3441375 | 6.29E-16 | 6 |
| IGFBP2 | -0.3259137 | 6.83E-16 | 6 |
| MBNL1 | 0.38824006 | 7.11E-16 | 6 |
| IDH2 | -0.4412786 | 8.41E-16 | 6 |
| PLP2 | -0.3382182 | 8.95E-16 | 6 |
| TIMP1 | -0.4059555 | 9.19E-16 | 6 |
| UQCR10 | -0.3496294 | 9.50E-16 | 6 |
| ETFB | -0.325722 | 9.65E-16 | 6 |
| ZRANB2 | 0.3797772 | 1.00E-15 | 6 |
| ALYREF | -0.3310974 | 1.08E-15 | 6 |
| UQCRQ | -0.396212 | 1.14E-15 | 6 |
| CCT2 | -0.353279 | 1.17E-15 | 6 |
| AIF1 | -0.3895767 | 1.18E-15 | 6 |
| MRPL57 | -0.3355687 | 1.46E-15 | 6 |

| | | | |
|---|---|---|---|
| CCT4 | -0.3095559 | 1.49E-15 | 6 |
| SUMO1 | -0.3069817 | 1.79E-15 | 6 |
| LAMTOR5 | -0.3135415 | 2.02E-15 | 6 |
| PET100 | -0.3550031 | 2.08E-15 | 6 |
| PTPN6 | -0.4923965 | 2.15E-15 | 6 |
| LDHA | -0.4214644 | 2.29E-15 | 6 |
| TRAPPC5 | -0.2854635 | 2.32E-15 | 6 |
| PSMD7 | -0.3139794 | 2.59E-15 | 6 |
| GTF2A2 | -0.2884912 | 2.61E-15 | 6 |
| EEF2 | -0.2961109 | 2.65E-15 | 6 |
| PPP1CA | -0.3154202 | 2.70E-15 | 6 |
| ANXA1 | -0.7198823 | 2.85E-15 | 6 |
| PAXX | -0.3561488 | 2.93E-15 | 6 |
| FAM89B | -0.3255138 | 2.94E-15 | 6 |
| C4orf48 | -0.3177463 | 3.14E-15 | 6 |
| TRIR | -0.3222878 | 3.27E-15 | 6 |
| GHITM | -0.2868787 | 3.30E-15 | 6 |
| TCEAL8 | -0.2947209 | 3.47E-15 | 6 |
| UFC1 | -0.3570121 | 3.61E-15 | 6 |
| RBM8A | -0.3397924 | 3.66E-15 | 6 |
| FIBP | -0.3145072 | 3.68E-15 | 6 |
| BUD31 | -0.3270966 | 3.74E-15 | 6 |
| IGBP1 | -0.328549 | 3.85E-15 | 6 |
| PDCD5 | -0.3360483 | 4.29E-15 | 6 |
| PSMD9 | -0.2825467 | 4.34E-15 | 6 |
| PSMA5 | -0.307132 | 5.25E-15 | 6 |
| TSPAN7 | -0.4513769 | 5.34E-15 | 6 |
| CRIP1 | -1.1777531 | 5.51E-15 | 6 |
| TRIM28 | -0.3086014 | 5.67E-15 | 6 |
| COX8A | -0.3705915 | 5.76E-15 | 6 |
| MPC2 | -0.2884367 | 5.95E-15 | 6 |
| SUMO3 | -0.2928254 | 6.86E-15 | 6 |
| MRPL11 | -0.2883974 | 6.90E-15 | 6 |
| PHB2 | -0.3145876 | 7.16E-15 | 6 |
| NDUFS3 | -0.2844136 | 7.57E-15 | 6 |
| ECHS1 | -0.2706337 | 7.72E-15 | 6 |
| NDUFB7 | -0.3375377 | 8.05E-15 | 6 |
| C9orf16 | -0.3271685 | 8.16E-15 | 6 |
| SYNRG | 0.36903185 | 8.27E-15 | 6 |
| RSRP1 | 0.35257051 | 8.82E-15 | 6 |
| CYBA | -0.3302182 | 9.01E-15 | 6 |
| CCNI | -0.2802626 | 9.43E-15 | 6 |
| CIB1 | -0.3249608 | 9.61E-15 | 6 |

| | | | |
|---|---|---|---|
| RHOG | -0.2971259 | 9.71E-15 | 6 |
| MDH1 | -0.304481 | 9.85E-15 | 6 |
| IMP3 | -0.2810847 | 1.02E-14 | 6 |
| GLIPR2 | -0.3530663 | 1.11E-14 | 6 |
| H2AFV | -0.3720562 | 1.17E-14 | 6 |
| ISG15 | -0.3003394 | 1.32E-14 | 6 |
| AL138899.1 | -0.5431743 | 1.33E-14 | 6 |
| RANBP1 | -0.4213871 | 1.35E-14 | 6 |
| REX1BD | -0.2985003 | 1.40E-14 | 6 |
| TLE5 | -0.2810041 | 1.46E-14 | 6 |
| ARF5 | -0.2844761 | 1.49E-14 | 6 |
| GLRX | -0.2982242 | 1.54E-14 | 6 |
| PSMA1 | -0.3205549 | 1.58E-14 | 6 |
| MRPL41 | -0.2952604 | 1.67E-14 | 6 |
| KDM5B | 0.37725463 | 1.69E-14 | 6 |
| LAMTOR2 | -0.2612554 | 1.78E-14 | 6 |
| ATP6V1F | -0.3198859 | 1.84E-14 | 6 |
| CKLF | -0.3729309 | 2.12E-14 | 6 |
| NSMCE1 | -0.3086316 | 2.33E-14 | 6 |
| FKBP8 | -0.2815126 | 2.79E-14 | 6 |
| PNN | 0.35408474 | 2.93E-14 | 6 |
| RAC1 | -0.286751 | 3.24E-14 | 6 |
| COPS9 | -0.2950915 | 3.32E-14 | 6 |
| HSPB1 | -0.3833153 | 3.72E-14 | 6 |
| NDUFA1 | -0.3102994 | 3.82E-14 | 6 |
| SNX3 | -0.2858236 | 3.82E-14 | 6 |
| SNRPG | -0.3916043 | 4.06E-14 | 6 |
| GNG5 | -0.3252529 | 4.17E-14 | 6 |
| VPS13C | 0.35305015 | 4.21E-14 | 6 |
| MRPL34 | -0.2754701 | 4.22E-14 | 6 |
| CCDC167 | -0.3197434 | 6.03E-14 | 6 |
| NDUFV2 | -0.285968 | 6.60E-14 | 6 |
| CD38 | 0.49798302 | 7.73E-14 | 6 |
| NDUFA11 | -0.3362901 | 8.08E-14 | 6 |
| BEX3 | -0.3173584 | 1.03E-13 | 6 |
| MAP2K2 | -0.3050367 | 1.14E-13 | 6 |
| MT-ND5 | 0.3651848 | 1.18E-13 | 6 |
| STK4 | 0.3466481 | 1.24E-13 | 6 |
| TMEM14C | -0.299933 | 1.25E-13 | 6 |
| IKZF1 | 0.39687933 | 1.27E-13 | 6 |
| COX16 | -0.2673689 | 1.37E-13 | 6 |
| PSMA4 | -0.3609682 | 1.52E-13 | 6 |
| STK17B | 0.33144563 | 1.60E-13 | 6 |

| | | | |
|---|---|---|---|
| AKAP9 | 0.38717829 | 1.70E-13 | 6 |
| SELL | -0.3624557 | 1.71E-13 | 6 |
| UBE2L3 | -0.289826 | 1.71E-13 | 6 |
| SNHG8 | -0.3197943 | 1.84E-13 | 6 |
| TRAC | -0.3615576 | 2.00E-13 | 6 |
| RAC2 | -0.2968162 | 2.24E-13 | 6 |
| JPT1 | -0.3634581 | 2.25E-13 | 6 |
| RPL22L1 | -0.3219758 | 2.48E-13 | 6 |
| NDUFA3 | -0.3356799 | 2.52E-13 | 6 |
| RTF2 | -0.2752967 | 2.60E-13 | 6 |
| SLBP | -0.3353703 | 2.80E-13 | 6 |
| CCT5 | -0.304714 | 2.89E-13 | 6 |
| NOP10 | -0.3138958 | 3.20E-13 | 6 |
| CCDC141 | 0.49649428 | 3.23E-13 | 6 |
| SPCS1 | -0.31212 | 3.77E-13 | 6 |
| WDR83OS | -0.3153894 | 3.91E-13 | 6 |
| SNRPD3 | -0.2830624 | 3.94E-13 | 6 |
| BIN1 | -0.2994756 | 4.23E-13 | 6 |
| ATP5PF | -0.3250541 | 4.23E-13 | 6 |
| PCSK7 | 0.38651386 | 4.28E-13 | 6 |
| CNPY3 | -0.2874414 | 4.44E-13 | 6 |
| MRPS15 | -0.2613094 | 4.45E-13 | 6 |
| CLTA | -0.2892429 | 4.51E-13 | 6 |
| AC004687.1 | 0.36456108 | 5.14E-13 | 6 |
| ZNF107 | 0.38813188 | 5.38E-13 | 6 |
| TMEM256 | -0.2902257 | 5.54E-13 | 6 |
| PSMB1 | -0.3368238 | 5.66E-13 | 6 |
| HINT2 | -0.2809614 | 5.70E-13 | 6 |
| ATP5MC1 | -0.2990423 | 5.78E-13 | 6 |
| DAD1 | -0.3052454 | 5.98E-13 | 6 |
| CD82 | -0.3032768 | 6.43E-13 | 6 |
| EIF3L | -0.3260392 | 6.64E-13 | 6 |
| NDUFAF3 | -0.2690386 | 7.84E-13 | 6 |
| AIP | -0.3072273 | 7.85E-13 | 6 |
| IFITM1 | -0.3917552 | 8.28E-13 | 6 |
| SLC25A6 | -0.3718701 | 8.78E-13 | 6 |
| ADRM1 | -0.2561746 | 1.02E-12 | 6 |
| GYPC | -0.2964824 | 1.05E-12 | 6 |
| TEX264 | -0.2561253 | 1.05E-12 | 6 |
| TCEAL9 | -0.2515475 | 1.06E-12 | 6 |
| SSB | -0.3049944 | 1.07E-12 | 6 |
| CNN2 | -0.4390609 | 1.12E-12 | 6 |
| DGUOK | -0.2575208 | 1.15E-12 | 6 |

| | | | |
|---|---|---|---|
| AKR1B1 | -0.285875 | 1.22E-12 | 6 |
| OSTC | -0.269159 | 1.25E-12 | 6 |
| COPZ1 | -0.2568986 | 1.39E-12 | 6 |
| JTB | -0.31785 | 1.42E-12 | 6 |
| NDUFC2 | -0.2998878 | 1.49E-12 | 6 |
| LNPEP | 0.33030522 | 1.51E-12 | 6 |
| ACAP1 | 0.3472406 | 1.69E-12 | 6 |
| CHD2 | 0.31454451 | 1.87E-12 | 6 |
| E2F2 | -0.2841462 | 1.92E-12 | 6 |
| ABRACL | -0.3120136 | 1.95E-12 | 6 |
| PSMA2 | -0.2533826 | 2.17E-12 | 6 |
| TUFM | -0.2768593 | 2.29E-12 | 6 |
| RBX1 | -0.345194 | 2.46E-12 | 6 |
| CLNS1A | -0.2610951 | 2.82E-12 | 6 |
| LGALS1 | -1.2073684 | 2.89E-12 | 6 |
| CLIC3 | -0.437635 | 3.08E-12 | 6 |
| CAPNS1 | -0.2910318 | 3.15E-12 | 6 |
| DDAH2 | -0.3116116 | 3.27E-12 | 6 |
| SSR4 | -0.3229453 | 3.40E-12 | 6 |
| HEBP2 | -0.2777108 | 3.88E-12 | 6 |
| NUTF2 | -0.2537298 | 4.03E-12 | 6 |
| SLC25A3 | -0.3722506 | 4.05E-12 | 6 |
| PLAAT4 | -0.2922913 | 4.22E-12 | 6 |
| PIN1 | -0.2631778 | 4.60E-12 | 6 |
| EIF3K | -0.2909983 | 4.69E-12 | 6 |
| CALM3 | -0.3369573 | 4.70E-12 | 6 |
| FABP5 | -0.3626376 | 4.70E-12 | 6 |
| ERH | -0.3166808 | 5.01E-12 | 6 |
| THOC7 | -0.2657453 | 5.21E-12 | 6 |
| ATP5PO | -0.3262191 | 5.25E-12 | 6 |
| CTDNEP1 | -0.2579816 | 5.53E-12 | 6 |
| RTRAF | -0.311076 | 5.75E-12 | 6 |
| UBE2S | -0.4215209 | 5.80E-12 | 6 |
| DCTPP1 | -0.2598654 | 5.85E-12 | 6 |
| MRPS34 | -0.2636041 | 6.29E-12 | 6 |
| PRELID1 | -0.2963192 | 6.33E-12 | 6 |
| SIT1 | -0.3242215 | 6.39E-12 | 6 |
| MRPS21 | -0.2700779 | 7.27E-12 | 6 |
| COPS3 | -0.2633634 | 7.41E-12 | 6 |
| NAP1L1 | -0.3180388 | 7.50E-12 | 6 |
| TIMM10 | -0.2676048 | 7.58E-12 | 6 |
| TMEM50A | -0.2796453 | 7.92E-12 | 6 |
| NMI | -0.2722071 | 8.04E-12 | 6 |

| | | | |
|---|---|---|---|
| EAPP | -0.2665194 | 8.25E-12 | 6 |
| SRI | -0.3061116 | 8.38E-12 | 6 |
| HNRNPH1 | 0.3327183 | 8.78E-12 | 6 |
| IL32 | -0.349927 | 8.98E-12 | 6 |
| SAP18 | -0.300628 | 9.28E-12 | 6 |
| EDF1 | -0.2734226 | 9.29E-12 | 6 |
| COX5A | -0.3513981 | 9.72E-12 | 6 |
| SOD1 | -0.3162647 | 9.93E-12 | 6 |
| SELENOF | -0.2695495 | 9.94E-12 | 6 |
| PARK7 | -0.3035359 | 1.00E-11 | 6 |
| NKG7 | -0.3628983 | 1.01E-11 | 6 |
| CDK2AP1 | -0.2861838 | 1.08E-11 | 6 |
| TUBB4B | -0.4221146 | 1.13E-11 | 6 |
| FKBP3 | -0.2761264 | 1.16E-11 | 6 |
| HMGA1 | -0.3713025 | 1.30E-11 | 6 |
| ATP5MD | -0.3286033 | 1.46E-11 | 6 |
| NDUFA5 | -0.2539347 | 1.48E-11 | 6 |
| RNF7 | -0.2674759 | 1.51E-11 | 6 |
| POLR3GL | -0.273264 | 1.62E-11 | 6 |
| GTF2H5 | -0.2635232 | 1.62E-11 | 6 |
| BCL7C | -0.2541661 | 1.66E-11 | 6 |
| ATOX1 | -0.259031 | 1.80E-11 | 6 |
| PAFAH1B3 | -0.2703348 | 1.81E-11 | 6 |
| ISCU | -0.2591 | 1.82E-11 | 6 |
| CD3E | -0.2733342 | 1.84E-11 | 6 |
| ZNF292 | 0.33133475 | 1.94E-11 | 6 |
| SET | -0.2595022 | 2.05E-11 | 6 |
| RBBP8 | -0.3291891 | 2.06E-11 | 6 |
| COX17 | -0.2852192 | 2.06E-11 | 6 |
| SNRPF | -0.3829949 | 2.09E-11 | 6 |
| TYMS | -0.6761511 | 2.11E-11 | 6 |
| HNRNPU | 0.27758439 | 2.32E-11 | 6 |
| PSMD8 | -0.2749668 | 2.45E-11 | 6 |
| TNRC6C | 0.32372317 | 2.57E-11 | 6 |
| GLRX3 | -0.2548577 | 2.69E-11 | 6 |
| VPS51 | -0.2602111 | 2.78E-11 | 6 |
| DGKA | 0.40036601 | 2.81E-11 | 6 |
| GSTP1 | -0.4855006 | 2.91E-11 | 6 |
| RAD23A | -0.2595745 | 2.97E-11 | 6 |
| TMEM14B | -0.2596854 | 3.14E-11 | 6 |
| STK17A | 0.34200186 | 3.19E-11 | 6 |
| RAN | -0.4327424 | 3.60E-11 | 6 |
| TRMT112 | -0.3147381 | 3.61E-11 | 6 |

| | | | |
|---|---|---|---|
| ARPC4 | -0.2775958 | 4.55E-11 | 6 |
| GCHFR | -0.2609425 | 4.67E-11 | 6 |
| APOBEC3G | -0.2705445 | 4.86E-11 | 6 |
| CCND3 | -0.3002972 | 5.46E-11 | 6 |
| PRDX5 | -0.3074768 | 5.46E-11 | 6 |
| CST7 | -0.3050461 | 5.64E-11 | 6 |
| CCT3 | -0.2611154 | 5.79E-11 | 6 |
| TAGLN2 | -0.5155285 | 5.89E-11 | 6 |
| RBM39 | 0.28005415 | 5.91E-11 | 6 |
| GLRX5 | -0.2660637 | 6.13E-11 | 6 |
| NDUFB4 | -0.316065 | 6.69E-11 | 6 |
| DMAC1 | -0.273564 | 6.70E-11 | 6 |
| PSMC5 | -0.2874959 | 7.00E-11 | 6 |
| KLF2 | -0.5954102 | 7.00E-11 | 6 |
| TMEM59 | -0.2507685 | 7.97E-11 | 6 |
| HNRNPA0 | -0.29661 | 8.63E-11 | 6 |
| HLA-C | -0.4457777 | 9.43E-11 | 6 |
| DDX3X | 0.31335124 | 9.51E-11 | 6 |
| HSP90AB1 | -0.4639719 | 9.84E-11 | 6 |
| ISG20 | -0.3139681 | 1.03E-10 | 6 |
| KRTCAP2 | -0.2619405 | 1.08E-10 | 6 |
| PSME1 | -0.2872843 | 1.08E-10 | 6 |
| HSPE1 | -0.3350841 | 1.09E-10 | 6 |
| DSTN | -0.3412978 | 1.17E-10 | 6 |
| ANXA2 | -0.3293984 | 1.25E-10 | 6 |
| PPP1R14B | -0.3602394 | 1.47E-10 | 6 |
| ABHD17A | -0.2633804 | 1.51E-10 | 6 |
| CCDC57 | -0.2581044 | 1.66E-10 | 6 |
| RUVBL1 | -0.2500368 | 1.87E-10 | 6 |
| AKNA | 0.29227208 | 2.49E-10 | 6 |
| EIF3G | -0.2994198 | 2.56E-10 | 6 |
| DDX5 | 0.27987212 | 2.66E-10 | 6 |
| CSTB | -0.307135 | 2.83E-10 | 6 |
| SEC61B | -0.2815933 | 2.87E-10 | 6 |
| C19orf53 | -0.3183614 | 3.09E-10 | 6 |
| HDAC2 | -0.2722286 | 3.20E-10 | 6 |
| DDX39A | -0.2595894 | 3.36E-10 | 6 |
| MYL6B | -0.2531829 | 3.71E-10 | 6 |
| WDR34 | -0.2724284 | 3.76E-10 | 6 |
| USP1 | -0.3452442 | 3.89E-10 | 6 |
| CALM2 | -0.2542197 | 3.92E-10 | 6 |
| GCC2 | 0.36272454 | 4.79E-10 | 6 |
| GIHCG | -0.5940438 | 5.54E-10 | 6 |

| | | | |
|---|---|---|---|
| B2M | -0.3936918 | 5.85E-10 | 6 |
| ANKRD11 | 0.34746335 | 6.75E-10 | 6 |
| ZFP36L2 | 0.31020984 | 7.59E-10 | 6 |
| HHIP-AS1 | -0.2575461 | 7.65E-10 | 6 |
| GUK1 | -0.2904906 | 7.99E-10 | 6 |
| LITAF | -0.2969191 | 8.88E-10 | 6 |
| FDFT1 | -0.2544253 | 9.54E-10 | 6 |
| RBM17 | -0.25811 | 1.12E-09 | 6 |
| UXT | -0.2964335 | 1.13E-09 | 6 |
| ZNF22 | -0.2578316 | 1.23E-09 | 6 |
| CELF2 | 0.37218414 | 1.33E-09 | 6 |
| EIF3D | -0.2709996 | 1.35E-09 | 6 |
| ETS1 | 0.29382876 | 1.46E-09 | 6 |
| LAMTOR4 | -0.2892972 | 1.50E-09 | 6 |
| ATP5F1D | -0.286833 | 1.56E-09 | 6 |
| KLRB1 | -0.4969694 | 1.76E-09 | 6 |
| HMGN2 | -0.4970833 | 1.85E-09 | 6 |
| XBP1 | -0.3633039 | 1.94E-09 | 6 |
| DDX6 | 0.26448059 | 2.14E-09 | 6 |
| GMFG | -0.2846294 | 2.18E-09 | 6 |
| CEP85L | 0.3720116 | 2.38E-09 | 6 |
| NOP58 | -0.2521044 | 2.48E-09 | 6 |
| LARP7 | -0.2521413 | 2.49E-09 | 6 |
| TUBB | -0.6418049 | 2.67E-09 | 6 |
| ZFAS1 | -0.2891259 | 2.90E-09 | 6 |
| RPS27L | -0.2676985 | 3.55E-09 | 6 |
| C12orf57 | -0.3700483 | 3.72E-09 | 6 |
| NOP56 | -0.283652 | 3.80E-09 | 6 |
| KMT2C | 0.27543627 | 3.95E-09 | 6 |
| CAPG | -0.4379127 | 4.21E-09 | 6 |
| RPS6KA3 | 0.33241447 | 4.22E-09 | 6 |
| ZNF638 | 0.32893373 | 4.56E-09 | 6 |
| HIST2H2AC | -0.4610578 | 4.63E-09 | 6 |
| PLAC8 | -0.5087858 | 4.92E-09 | 6 |
| EIF3H | -0.26089 | 5.36E-09 | 6 |
| UNG | -0.2508743 | 5.87E-09 | 6 |
| DDX21 | -0.2501428 | 6.28E-09 | 6 |
| PCM1 | 0.3030404 | 6.34E-09 | 6 |
| NEDD8 | -0.290188 | 6.36E-09 | 6 |
| ZC3HAV1 | 0.31564149 | 6.45E-09 | 6 |
| COX7B | -0.3082859 | 6.47E-09 | 6 |
| TFF3 | -0.3670331 | 6.86E-09 | 6 |
| COPE | -0.2712293 | 7.03E-09 | 6 |

| | | | |
|---|---|---|---|
| SMDT1 | -0.3016805 | 7.05E-09 | 6 |
| PDIA6 | -0.25484 | 7.09E-09 | 6 |
| SELENOH | -0.2940895 | 9.18E-09 | 6 |
| GABARAP | -0.2681979 | 9.45E-09 | 6 |
| ADA | -0.300001 | 9.49E-09 | 6 |
| SSRP1 | -0.2555944 | 1.02E-08 | 6 |
| H1FX | -0.4368247 | 1.43E-08 | 6 |
| NPIPB5 | 0.35453546 | 1.44E-08 | 6 |
| CKS2 | -0.3425749 | 1.46E-08 | 6 |
| YWHAH | -0.3208331 | 1.47E-08 | 6 |
| CD81 | -0.252815 | 1.51E-08 | 6 |
| ANP32E | -0.3533245 | 1.53E-08 | 6 |
| SELENOW | -0.353789 | 1.68E-08 | 6 |
| TKT | -0.286835 | 2.28E-08 | 6 |
| PKM | -0.4054698 | 2.45E-08 | 6 |
| SNHG32 | -0.30075 | 2.49E-08 | 6 |
| H2AFZ | -0.5528354 | 2.52E-08 | 6 |
| APRT | -0.291633 | 2.67E-08 | 6 |
| GIMAP4 | -0.2899078 | 2.83E-08 | 6 |
| LMNB1 | -0.2975764 | 2.88E-08 | 6 |
| PCLAF | -0.5671586 | 3.15E-08 | 6 |
| PEBP1 | -0.2856146 | 3.50E-08 | 6 |
| NDUFB1 | -0.2973386 | 3.55E-08 | 6 |
| EIF4A1 | -0.296429 | 3.76E-08 | 6 |
| MCM6 | -0.2907792 | 4.31E-08 | 6 |
| FOXP1 | 0.31015413 | 4.62E-08 | 6 |
| MYOM2 | -0.3121766 | 4.98E-08 | 6 |
| PRDX2 | -0.370689 | 5.02E-08 | 6 |
| RBM6 | 0.31972249 | 5.15E-08 | 6 |
| ERICH1 | 0.31869849 | 5.45E-08 | 6 |
| SDCBP | -0.2670999 | 5.55E-08 | 6 |
| NOSIP | -0.6385151 | 5.90E-08 | 6 |
| ITGA4 | 0.27420359 | 5.91E-08 | 6 |
| SCP2 | -0.2529765 | 6.04E-08 | 6 |
| ASH1L | 0.32278679 | 9.57E-08 | 6 |
| FXYD5 | -0.3520759 | 9.84E-08 | 6 |
| PDAP1 | -0.258566 | 1.11E-07 | 6 |
| GPX4 | -0.3055249 | 1.29E-07 | 6 |
| HNRNPF | -0.2716161 | 1.36E-07 | 6 |
| SIVA1 | -0.4226225 | 1.50E-07 | 6 |
| C12orf75 | -0.3279551 | 1.54E-07 | 6 |
| AC144521.1 | 0.33866033 | 1.70E-07 | 6 |
| CD226 | 0.37936588 | 1.83E-07 | 6 |

| | | | |
|---|---|---|---|
| BRK1 | -0.2551336 | 1.85E-07 | 6 |
| NUCKS1 | -0.3974632 | 2.11E-07 | 6 |
| TPI1 | -0.3142521 | 2.12E-07 | 6 |
| RBM25 | 0.29973731 | 2.18E-07 | 6 |
| XRCC6 | -0.2641208 | 2.29E-07 | 6 |
| PCBP1 | -0.2540257 | 2.34E-07 | 6 |
| AKAP13 | 0.28185034 | 2.46E-07 | 6 |
| FCER1G | -0.2786783 | 2.90E-07 | 6 |
| MCM3 | -0.3106375 | 3.07E-07 | 6 |
| CD96 | 0.25264275 | 3.58E-07 | 6 |
| TUBA1B | -0.797625 | 3.65E-07 | 6 |
| ZNF655 | 0.34356312 | 3.71E-07 | 6 |
| ANKRD12 | 0.31670383 | 3.74E-07 | 6 |
| TRAPPC1 | -0.2527006 | 3.80E-07 | 6 |
| CCNL1 | 0.27919633 | 3.90E-07 | 6 |
| SLC16A7 | 0.36911759 | 4.24E-07 | 6 |
| POLR1D | -0.2553277 | 4.39E-07 | 6 |
| ETV5 | -0.3913267 | 4.51E-07 | 6 |
| GNB2 | -0.2603671 | 4.60E-07 | 6 |
| TBCA | -0.2546842 | 4.99E-07 | 6 |
| AQP3 | -0.4090532 | 5.58E-07 | 6 |
| EIF2S2 | -0.2762749 | 6.90E-07 | 6 |
| PCDHGA6 | 0.30175325 | 7.94E-07 | 6 |
| GAS5 | -0.3252913 | 1.08E-06 | 6 |
| S100A4 | -0.9465549 | 1.27E-06 | 6 |
| ZNF117 | 0.40617829 | 1.39E-06 | 6 |
| ZBTB37 | 0.34476344 | 1.43E-06 | 6 |
| EMP3 | -0.570855 | 1.43E-06 | 6 |
| MCM4 | -0.2546731 | 1.59E-06 | 6 |
| DIRC3 | 0.28936315 | 1.69E-06 | 6 |
| RBM33 | 0.3011496 | 1.79E-06 | 6 |
| RNASEK | -0.2542819 | 1.82E-06 | 6 |
| TENM1 | 0.38673122 | 2.09E-06 | 6 |
| ATP5F1C | -0.2540886 | 2.11E-06 | 6 |
| PRMT7 | -0.3300743 | 2.12E-06 | 6 |
| KMT2A | 0.29958355 | 2.38E-06 | 6 |
| KMT2E | 0.25035486 | 2.69E-06 | 6 |
| AC108066.2 | 0.29728319 | 3.00E-06 | 6 |
| MZT2A | -0.2655404 | 3.90E-06 | 6 |
| AHNAK | -0.5109913 | 3.92E-06 | 6 |
| DCP2 | 0.26318399 | 3.98E-06 | 6 |
| CDKN2D | -0.3509219 | 4.16E-06 | 6 |
| AL365440.2 | -0.2665734 | 4.62E-06 | 6 |

| | | | |
|---|---|---|---|
| DGKH | 0.3674662 | 4.64E-06 | 6 |
| FUS | 0.26169038 | 4.66E-06 | 6 |
| RRM1 | -0.2589957 | 4.66E-06 | 6 |
| IFITM2 | -0.4721581 | 4.85E-06 | 6 |
| GOLGB1 | 0.33835934 | 5.11E-06 | 6 |
| ADD3 | -0.3910151 | 5.32E-06 | 6 |
| ERP29 | -0.2596409 | 5.35E-06 | 6 |
| LINC00342 | 0.29272617 | 5.47E-06 | 6 |
| FAM49B | 0.25242975 | 6.18E-06 | 6 |
| ARPP21 | -0.2754691 | 7.04E-06 | 6 |
| TBL1XR1 | 0.32289752 | 7.45E-06 | 6 |
| MARCKSL1 | -0.3055412 | 8.94E-06 | 6 |
| SCAF11 | 0.26990269 | 1.03E-05 | 6 |
| LIMS1 | -0.2555364 | 1.18E-05 | 6 |
| DUT | -0.4730122 | 1.43E-05 | 6 |
| ENO1 | -0.3381368 | 1.93E-05 | 6 |
| PCDHGA5 | 0.2508518 | 1.99E-05 | 6 |
| PPIB | -0.2722815 | 2.17E-05 | 6 |
| ACAP2 | 0.25235803 | 2.61E-05 | 6 |
| GABPB1-AS1 | 0.32636439 | 2.70E-05 | 6 |
| GIMAP7 | -0.3104469 | 2.80E-05 | 6 |
| ARL6IP1 | -0.2833406 | 2.88E-05 | 6 |
| CTSW | -0.3572686 | 3.25E-05 | 6 |
| PRMT2 | 0.25796225 | 3.28E-05 | 6 |
| S100A6 | -0.5609816 | 3.50E-05 | 6 |
| LST1 | -0.5890601 | 3.77E-05 | 6 |
| PSME2 | -0.2601043 | 4.28E-05 | 6 |
| HSP90AA1 | -0.2505692 | 4.58E-05 | 6 |
| BOD1L1 | 0.27754413 | 4.60E-05 | 6 |
| AC068587.4 | 0.30798238 | 5.62E-05 | 6 |
| CLSPN | -0.2974702 | 7.15E-05 | 6 |
| HSPA1A | -0.3012445 | 0.00010036 | 6 |
| MBD5 | 0.36175416 | 0.0001339 | 6 |
| HLA-A | -0.3825462 | 0.00015299 | 6 |
| KLRK1 | -0.2534907 | 0.00015833 | 6 |
| LAPTM5 | -0.2637662 | 0.00015916 | 6 |
| TRIM56 | 0.2563801 | 0.00016573 | 6 |
| MZB1 | -0.3373526 | 0.00018638 | 6 |
| H2AFX | -0.313019 | 0.00019508 | 6 |
| ZNF66 | 0.26395945 | 0.00020998 | 6 |
| STX16 | 0.29987357 | 0.00023276 | 6 |
| ZNF708 | 0.33280523 | 0.00027769 | 6 |
| SMC2 | -0.2975875 | 0.00034571 | 6 |

| | | | |
|---|---|---|---|
| MSN | -0.2644635 | 0.00048514 | 6 |
| MAL | -0.3889971 | 0.00052878 | 6 |
| SERINC5 | 0.32559064 | 0.00055952 | 6 |
| GTF2IRD2 | 0.30923657 | 0.00066229 | 6 |
| CD7 | -0.2895211 | 0.00066392 | 6 |
| DNAJB14 | 0.32967519 | 0.00077296 | 6 |
| ANXA5 | -0.2540934 | 0.00079414 | 6 |
| MPHOSPH8 | 0.25176805 | 0.00083201 | 6 |
| NEGR1 | -0.2966357 | 0.00086773 | 6 |
| FAM133B | 0.26458754 | 0.00100958 | 6 |
| IL17RB | -0.4103236 | 0.00102237 | 6 |
| JAML | 0.26246418 | 0.00107805 | 6 |
| ZBTB20 | 0.32672573 | 0.00136602 | 6 |
| NUSAP1 | -0.4979844 | 0.00140487 | 6 |
| CALR | -0.339869 | 0.0014411 | 6 |
| ATAD2 | -0.3062782 | 0.00172273 | 6 |
| FARS2 | -0.2700465 | 0.00179581 | 6 |
| DMTF1 | 0.25183399 | 0.00193109 | 6 |
| ZNF720 | 0.33470417 | 0.0020089 | 6 |
| PCSK5 | 0.28456538 | 0.00264588 | 6 |
| ELOVL4 | -0.2889653 | 0.00272503 | 6 |
| SRGN | -0.3512815 | 0.00278357 | 6 |
| WSB1 | 0.26203725 | 0.00333968 | 6 |
| EZR | -0.3316788 | 0.00379699 | 6 |
| S100A10 | -0.4983995 | 0.00545458 | 6 |
| PTTG1 | -0.2776317 | 0.00601006 | 6 |
| NCR3 | -0.2697668 | 0.00687967 | 6 |
| CHST2 | 0.27081044 | 0.00771557 | 6 |
| MCM7 | -0.2800593 | 0.00803389 | 6 |
| HERC1 | 0.27588747 | 0.00834958 | 6 |
| RNPC3 | 0.27437331 | 0.00967544 | 6 |
| RAPGEF6 | 0.28285667 | 0.01051362 | 6 |
| HELLS | -0.3062429 | 0.01454747 | 6 |
| RBM12B | 0.31216034 | 0.01719301 | 6 |
| GTSE1 | -0.3134623 | 0.01784672 | 6 |
| ZNF138 | 0.2745889 | 0.0181457 | 6 |
| ABLIM1 | 0.33783279 | 0.01941959 | 6 |
| NIBAN3 | 0.25978774 | 0.0216768 | 6 |
| MIR181A1HG | 0.32288988 | 0.1105908 | 6 |
| MLXIP | 0.27611562 | 0.12959614 | 6 |
| CARD8 | 0.27410745 | 0.17731337 | 6 |
| ATXN7 | 0.28857112 | 0.18110702 | 6 |
| ANKRD44 | 0.28288394 | 0.2232324 | 6 |

| | | | |
|---|---|---|---|
| SETD2 | 0.29825681 | 0.25258371 | 6 |
| ODF2L | 0.28430825 | 0.33442113 | 6 |
| NUTM2A-AS1 | 0.294115 | 0.3608771 | 6 |
| SHPRH | 0.34074885 | 0.49634046 | 6 |
| RBM5 | 0.28049096 | 0.5326034 | 6 |
| ITM2A | -0.4368199 | 0.58613845 | 6 |
| AC025171.2 | 0.27789125 | 0.62860541 | 6 |
| ZBED5 | 0.25483763 | 0.67388355 | 6 |
| CD1E | -0.3979482 | 0.72117129 | 6 |
| AC114760.2 | 0.283639 | 0.84019727 | 6 |
| RRM2 | -0.2771433 | 0.946094 | 6 |
| CHD9 | 0.26119685 | 1 | 6 |
| TECR | -0.5205474 | 1 | 6 |
| TRIM73 | 0.27593345 | 1 | 6 |
| DNASE1 | 0.25785082 | 1 | 6 |
| TCL1A | -0.3075206 | 1 | 6 |
| USP15 | 0.25375211 | 1 | 6 |
| ZNF431 | 0.25166178 | 1 | 6 |
| SENP7 | 0.25272787 | 1 | 6 |
| ASPM | -0.4405145 | 1 | 6 |
| RPAP2 | 0.31683215 | 1 | 6 |
| CD79A | -0.2551019 | 1 | 6 |
| CD1A | -0.5064021 | 1 | 6 |
| CCDC14 | 0.26288923 | 1 | 6 |
| CEMIP2 | 0.28332672 | 1 | 6 |
| HLA-B | -0.4624203 | 1 | 6 |
| CD1B | -0.362 | 1 | 6 |
| TOP2A | -0.5380738 | 1 | 6 |
| ZNF737 | 0.26770375 | 1 | 6 |
| CENPF | -0.458972 | 1 | 6 |
| CREBRF | 0.25759386 | 1 | 6 |
| LTB | -0.4904378 | 1 | 6 |
| HCG18 | 0.25177055 | 1 | 6 |
| MKI67 | -0.5100716 | 1 | 6 |
| CR1 | 0.25383348 | 1 | 6 |
| NUSAP1 | 2.80530065 | 0 | 7 |
| ASPM | 2.80505535 | 0 | 7 |
| HIST1H3B | 2.23058408 | 0 | 7 |
| GTSE1 | 2.20364679 | 0 | 7 |
| RRM2 | 2.08330778 | 0 | 7 |
| UBE2C | 1.98618134 | 0 | 7 |
| KNL1 | 1.9761637 | 0 | 7 |
| TPX2 | 1.88741501 | 0 | 7 |

| | | | |
|---|---|---|---|
| CDK1 | 1.82856944 | 0 | 7 |
| CENPE | 1.77008286 | 0 | 7 |
| KIF11 | 1.67129893 | 0 | 7 |
| BIRC5 | 1.63108405 | 0 | 7 |
| DLGAP5 | 1.61710376 | 0 | 7 |
| CKS1B | 1.61390297 | 0 | 7 |
| AURKB | 1.54021054 | 0 | 7 |
| PRC1 | 1.49466668 | 0 | 7 |
| KIF15 | 1.43789242 | 0 | 7 |
| HMMR | 1.42787547 | 0 | 7 |
| NCAPG | 1.34412853 | 0 | 7 |
| CCNB2 | 1.30396381 | 0 | 7 |
| HIST1H2AH | 1.2955256 | 0 | 7 |
| ARHGAP11A | 1.29369333 | 0 | 7 |
| CEP55 | 1.2663825 | 0 | 7 |
| CKAP2L | 1.23919874 | 0 | 7 |
| CENPW | 1.22020139 | 0 | 7 |
| NUF2 | 1.18177207 | 0 | 7 |
| CCNA2 | 1.14834672 | 0 | 7 |
| HIST1H2AL | 1.13167585 | 0 | 7 |
| HIST1H3G | 1.13154482 | 0 | 7 |
| KIFC1 | 1.13036573 | 0 | 7 |
| RAD51AP1 | 1.12530594 | 0 | 7 |
| CDC20 | 1.12255621 | 0 | 7 |
| KIF14 | 1.1174608 | 0 | 7 |
| FOXM1 | 1.11183265 | 0 | 7 |
| HJURP | 1.10068139 | 0 | 7 |
| CDKN3 | 1.06578759 | 0 | 7 |
| CDCA8 | 1.0539291 | 0 | 7 |
| KIF23 | 1.02662732 | 0 | 7 |
| TTK | 1.01984695 | 0 | 7 |
| BUB1 | 1.01290157 | 0 | 7 |
| ESCO2 | 0.98784394 | 0 | 7 |
| KIF2C | 0.98770905 | 0 | 7 |
| KIF4A | 0.97546138 | 0 | 7 |
| NCAPH | 0.95040898 | 0 | 7 |
| CDCA5 | 0.94388528 | 0 | 7 |
| E2F8 | 0.94199676 | 0 | 7 |
| SKA3 | 0.93091769 | 0 | 7 |
| DEPDC1 | 0.93051076 | 0 | 7 |
| CENPA | 0.92318122 | 0 | 7 |
| CDCA3 | 0.92263824 | 0 | 7 |
| RACGAP1 | 0.91769735 | 0 | 7 |

| | | | |
|---|---|---|---|
| SPC25 | 0.87630767 | 0 | 7 |
| DIAPH3 | 0.84991995 | 0 | 7 |
| CIT | 0.81830267 | 0 | 7 |
| CDCA2 | 0.80510848 | 0 | 7 |
| PBK | 0.79098422 | 0 | 7 |
| MELK | 0.77498277 | 0 | 7 |
| SPC24 | 0.76928733 | 0 | 7 |
| KIF20A | 0.72649471 | 0 | 7 |
| TROAP | 0.70718327 | 0 | 7 |
| DEPDC1B | 0.69589552 | 0 | 7 |
| SHCBP1 | 0.67732087 | 0 | 7 |
| PKMYT1 | 0.65727777 | 0 | 7 |
| KIF18B | 0.64510608 | 0 | 7 |
| HIST1H3C | 0.54081632 | 0 | 7 |
| SKA1 | 0.57879273 | 1.73E-297 | 7 |
| FBXO43 | 0.71459644 | 4.17E-297 | 7 |
| SGO1 | 1.12785401 | 5.44E-297 | 7 |
| ORC6 | 1.08694813 | 2.35E-294 | 7 |
| AC091057.6 | 0.78802936 | 7.17E-294 | 7 |
| TOP2A | 3.26978599 | 2.05E-292 | 7 |
| POLQ | 0.72623644 | 2.07E-290 | 7 |
| E2F7 | 0.83227977 | 5.26E-287 | 7 |
| NDC80 | 1.23537551 | 3.35E-285 | 7 |
| HIST1H2BB | 0.50921168 | 1.47E-284 | 7 |
| ESPL1 | 0.45704273 | 2.81E-281 | 7 |
| PIMREG | 0.52265236 | 2.92E-281 | 7 |
| MXD3 | 1.14412881 | 1.14E-280 | 7 |
| PLK1 | 1.01079868 | 1.72E-280 | 7 |
| SPAG5 | 0.60098759 | 1.12E-279 | 7 |
| HIST1H1B | 2.77336983 | 1.11E-278 | 7 |
| BUB1B | 1.01763931 | 2.81E-278 | 7 |
| HIST1H2AM | 0.82530561 | 4.77E-277 | 7 |
| ASF1B | 1.17860666 | 5.34E-277 | 7 |
| HIST1H4D | 0.66605074 | 7.61E-277 | 7 |
| UBE2T | 1.12581727 | 6.33E-274 | 7 |
| KIF18A | 0.60356064 | 1.97E-270 | 7 |
| CENPM | 1.31025017 | 1.39E-268 | 7 |
| MKI67 | 3.12206724 | 2.56E-266 | 7 |
| CENPF | 3.02856754 | 3.56E-265 | 7 |
| STIL | 0.73053295 | 1.01E-264 | 7 |
| AURKA | 0.69944215 | 1.78E-264 | 7 |
| ECT2 | 0.90559369 | 5.34E-262 | 7 |
| HIST1H2AI | 0.53816077 | 6.41E-262 | 7 |

| | | | |
|---|---|---|---|
| HIST1H2BH | 0.91681585 | 9.41E-260 | 7 |
| HIST1H2AG | 1.49705552 | 6.29E-251 | 7 |
| CDC25C | 0.43018954 | 6.80E-250 | 7 |
| ANLN | 0.50370564 | 1.20E-249 | 7 |
| HIST1H2AB | 0.49187068 | 1.04E-243 | 7 |
| MYBL2 | 0.88901007 | 4.26E-242 | 7 |
| NEK2 | 0.50784199 | 9.68E-241 | 7 |
| ARHGEF39 | 0.68106096 | 2.76E-239 | 7 |
| PLK4 | 0.62616029 | 1.37E-238 | 7 |
| MAD2L1 | 1.20263135 | 4.25E-234 | 7 |
| SGO2 | 1.35388359 | 3.46E-231 | 7 |
| CIP2A | 0.76948303 | 7.08E-230 | 7 |
| TK1 | 0.69442438 | 7.29E-229 | 7 |
| HIST1H3F | 0.52314962 | 4.53E-228 | 7 |
| BRCA2 | 1.23796536 | 2.58E-224 | 7 |
| OIP5 | 0.43486434 | 2.02E-222 | 7 |
| MND1 | 0.92873885 | 8.81E-222 | 7 |
| MCM10 | 0.72432214 | 3.17E-221 | 7 |
| H2AFX | 1.89724402 | 4.46E-218 | 7 |
| INCENP | 0.6073962 | 9.60E-216 | 7 |
| HIST1H2AJ | 0.72695663 | 2.09E-212 | 7 |
| KNSTRN | 0.69488413 | 1.13E-208 | 7 |
| KIF20B | 1.47902691 | 4.33E-207 | 7 |
| TRAIP | 0.36251858 | 1.03E-205 | 7 |
| POC1A | 0.41144021 | 4.46E-202 | 7 |
| CENPU | 1.25436244 | 5.98E-202 | 7 |
| TMSB15A | 1.02143624 | 8.10E-201 | 7 |
| ZWINT | 1.16568945 | 2.67E-199 | 7 |
| CCDC34 | 0.96733522 | 2.68E-197 | 7 |
| PTTG1 | 1.80711856 | 2.46E-195 | 7 |
| PRR11 | 0.76594759 | 2.10E-191 | 7 |
| HMGB3 | 1.25375335 | 1.84E-190 | 7 |
| CCNF | 0.78282809 | 1.24E-189 | 7 |
| PCLAF | 2.1015924 | 9.73E-189 | 7 |
| RPL39L | 0.8123462 | 5.02E-188 | 7 |
| PSRC1 | 0.40169452 | 1.51E-187 | 7 |
| C21orf58 | 1.3594028 | 1.30E-182 | 7 |
| CCDC150 | 0.42065111 | 2.15E-182 | 7 |
| HIST1H2BL | 0.35027616 | 3.46E-182 | 7 |
| TYMS | 2.19001522 | 3.50E-182 | 7 |
| WDR62 | 0.51706511 | 7.41E-182 | 7 |
| LMNB2 | 0.79534208 | 7.73E-182 | 7 |
| RAD54L | 0.43322217 | 2.95E-181 | 7 |

| | | | |
|---|---|---|---|
| CCNB1 | 1.19967911 | 7.10E-181 | 7 |
| HIST2H2AB | 0.62288532 | 3.93E-180 | 7 |
| RAD51 | 0.55453743 | 2.93E-179 | 7 |
| CENPN | 0.53055699 | 3.45E-179 | 7 |
| TRIP13 | 0.39155107 | 4.17E-177 | 7 |
| HMGB2 | 3.08477309 | 9.69E-177 | 7 |
| MTFR2 | 0.37466956 | 1.84E-174 | 7 |
| GGH | 0.4876674 | 8.05E-174 | 7 |
| FANCD2 | 0.63326913 | 1.07E-172 | 7 |
| FBXO5 | 1.37573817 | 1.12E-171 | 7 |
| HMGB1 | 1.90242602 | 1.60E-171 | 7 |
| KIF22 | 1.17276686 | 1.91E-169 | 7 |
| FANCI | 1.04474633 | 2.94E-168 | 7 |
| DSCC1 | 0.58588411 | 3.74E-167 | 7 |
| SPDL1 | 0.83619693 | 7.24E-167 | 7 |
| HIST1H3I | 0.39538344 | 8.73E-162 | 7 |
| TUBA1B | 2.59010586 | 1.13E-159 | 7 |
| FAM111B | 0.94686611 | 1.23E-158 | 7 |
| EXO1 | 0.41322933 | 2.65E-158 | 7 |
| HIST1H3D | 2.41388519 | 1.98E-157 | 7 |
| TUBB4B | 2.01258786 | 5.17E-157 | 7 |
| SMC2 | 1.54897682 | 6.86E-157 | 7 |
| XRCC2 | 0.84641747 | 2.08E-156 | 7 |
| UBE2S | 1.9428331 | 6.02E-156 | 7 |
| CLSPN | 1.40870572 | 2.00E-155 | 7 |
| GEN1 | 0.59220672 | 1.08E-154 | 7 |
| TUBB | 2.2414015 | 1.06E-153 | 7 |
| ANP32E | 1.68709899 | 4.64E-153 | 7 |
| DTYMK | 0.93582071 | 3.17E-151 | 7 |
| CDC45 | 0.55134193 | 3.69E-146 | 7 |
| HIST1H2AE | 0.59719648 | 1.23E-144 | 7 |
| CENPS | 0.45237832 | 3.88E-144 | 7 |
| H2AFZ | 2.12179821 | 6.61E-143 | 7 |
| TACC3 | 1.15573288 | 2.99E-142 | 7 |
| FEN1 | 0.93734423 | 6.34E-141 | 7 |
| HMGN2 | 1.80850762 | 1.60E-138 | 7 |
| MIS18A | 0.57485609 | 2.26E-138 | 7 |
| SAC3D1 | 0.95765991 | 2.42E-138 | 7 |
| HIST1H2BC | 0.65175474 | 6.64E-137 | 7 |
| HIST1H4C | 3.99298368 | 6.83E-137 | 7 |
| RFC3 | 0.6540909 | 1.82E-136 | 7 |
| GPSM2 | 0.69283876 | 2.96E-136 | 7 |
| PARPBP | 0.53687128 | 7.03E-136 | 7 |

| | | | |
|---|---|---|---|
| STMN1 | 1.4863067 | 9.58E-135 | 7 |
| ATAD5 | 1.0598257 | 1.16E-134 | 7 |
| TCF19 | 0.66567916 | 5.80E-134 | 7 |
| HIST2H2BF | 0.37330732 | 1.91E-133 | 7 |
| C18orf54 | 0.42787547 | 5.03E-132 | 7 |
| GINS1 | 0.41990754 | 5.30E-132 | 7 |
| FAM83D | 0.31301349 | 5.77E-132 | 7 |
| HIST1H2BI | 0.26650781 | 6.96E-132 | 7 |
| NUCKS1 | 1.55622809 | 6.44E-131 | 7 |
| CDC25A | 0.36987417 | 6.31E-130 | 7 |
| Z94721.3 | 0.47081692 | 9.39E-130 | 7 |
| LMNB1 | 1.30800749 | 1.16E-129 | 7 |
| C1orf112 | 0.49414381 | 3.12E-128 | 7 |
| CENPH | 0.72771748 | 9.38E-128 | 7 |
| ATAD2 | 1.58322305 | 1.28E-127 | 7 |
| RCCD1 | 0.48303626 | 2.75E-127 | 7 |
| KPNA2 | 1.29755718 | 1.55E-126 | 7 |
| TMPO | 1.49535862 | 5.09E-126 | 7 |
| ERCC6L | 0.31911308 | 4.25E-125 | 7 |
| NCAPD2 | 1.10904537 | 1.37E-123 | 7 |
| NEIL3 | 0.93665359 | 2.10E-122 | 7 |
| GINS4 | 0.46319937 | 2.37E-122 | 7 |
| HIST1H1C | 2.17775803 | 1.01E-120 | 7 |
| IQGAP3 | 0.29368017 | 8.58E-120 | 7 |
| WDR34 | 0.93452574 | 1.66E-119 | 7 |
| UHRF1 | 0.94151547 | 6.61E-119 | 7 |
| CKS2 | 1.56007005 | 4.79E-118 | 7 |
| TEX30 | 0.62302447 | 5.51E-118 | 7 |
| NEURL1B | 0.32588254 | 2.12E-116 | 7 |
| BRIP1 | 0.55384116 | 1.49E-113 | 7 |
| CENPO | 0.36201352 | 2.87E-113 | 7 |
| HIRIP3 | 0.83961127 | 8.73E-113 | 7 |
| HIST1H1D | 1.95738307 | 3.14E-111 | 7 |
| GMNN | 0.90111854 | 5.87E-111 | 7 |
| HYI | 0.49236961 | 1.33E-110 | 7 |
| TUBA1C | 1.16108999 | 1.74E-110 | 7 |
| H2AFV | 1.17129604 | 3.15E-110 | 7 |
| PTMA | 0.99496537 | 9.90E-110 | 7 |
| ASRGL1 | 0.65883909 | 5.45E-109 | 7 |
| FANCA | 0.58974041 | 8.25E-109 | 7 |
| HSPA1B | 1.36949838 | 1.39E-107 | 7 |
| DDIAS | 0.30938894 | 1.83E-107 | 7 |
| TTF2 | 0.80654805 | 1.98E-107 | 7 |

| | | | |
|---|---|---|---|
| DHFR | 1.23534481 | 9.82E-107 | 7 |
| SUV39H2 | 0.37245654 | 2.83E-106 | 7 |
| HIST1H4E | 0.68947563 | 3.06E-106 | 7 |
| PKP4 | 0.44266078 | 5.90E-106 | 7 |
| CDKN2C | 0.98947016 | 3.28E-105 | 7 |
| EZH2 | 1.43853145 | 5.29E-105 | 7 |
| GTF3C5 | 1.23018754 | 7.44E-105 | 7 |
| LRR1 | 0.59230344 | 1.05E-103 | 7 |
| ANP32B | 1.13527913 | 1.20E-103 | 7 |
| CDC6 | 0.47764622 | 8.97E-103 | 7 |
| HIST1H2BF | 0.81588289 | 1.54E-102 | 7 |
| RRM1 | 1.08746415 | 1.75E-102 | 7 |
| CENPJ | 0.60112471 | 7.24E-102 | 7 |
| TEDC1 | 0.43662145 | 9.24E-102 | 7 |
| RAN | 1.2272583 | 1.93E-101 | 7 |
| HIST2H2AC | 2.16264631 | 3.89E-101 | 7 |
| ERI2 | 0.33251243 | 3.99E-101 | 7 |
| BRCA1 | 0.8430195 | 4.35E-100 | 7 |
| MGME1 | 0.5810543 | 1.50E-99 | 7 |
| DBF4 | 1.02655373 | 5.58E-99 | 7 |
| CEP152 | 0.75713676 | 1.31E-98 | 7 |
| MAD2L2 | 0.92491338 | 1.59E-98 | 7 |
| HIST1H3H | 0.40043835 | 4.64E-98 | 7 |
| CDT1 | 0.7774636 | 1.21E-97 | 7 |
| NRM | 0.49458957 | 1.80E-97 | 7 |
| TUBG1 | 0.46598949 | 1.85E-97 | 7 |
| CENPK | 0.85793659 | 5.09E-97 | 7 |
| RPL34 | -1.0387119 | 5.10E-96 | 7 |
| RECQL4 | 0.29010148 | 1.88E-95 | 7 |
| CTNNAL1 | 0.46556836 | 2.18E-94 | 7 |
| ETS1 | -1.3029034 | 1.18E-93 | 7 |
| HNRNPA2B1 | 0.87702769 | 1.99E-93 | 7 |
| USP1 | 1.10978891 | 2.49E-93 | 7 |
| CRNDE | 1.03412782 | 1.63E-92 | 7 |
| RAD21 | 1.31243069 | 2.48E-92 | 7 |
| CBX5 | 1.07720376 | 1.16E-91 | 7 |
| DNTT | 0.95530728 | 3.26E-91 | 7 |
| HIST1H4I | 0.27533854 | 3.32E-91 | 7 |
| PIF1 | 0.4568741 | 4.55E-91 | 7 |
| CEP128 | 0.51842144 | 3.80E-90 | 7 |
| FAM111A | 1.1780938 | 5.90E-90 | 7 |
| MIS18BP1 | 1.10546722 | 2.01E-89 | 7 |
| EPB41L2 | 0.66890589 | 2.77E-89 | 7 |

| | | | |
|---|---|---|---|
| TEDC2 | 0.26422902 | 3.11E-89 | 7 |
| CKAP5 | 0.89572481 | 5.30E-88 | 7 |
| NCAPD3 | 0.80577262 | 9.94E-88 | 7 |
| PHF19 | 0.73491194 | 1.18E-87 | 7 |
| HIST1H2BK | 0.35446437 | 1.34E-87 | 7 |
| RPS27 | -0.9854301 | 2.73E-87 | 7 |
| HAUS8 | 0.43400342 | 2.20E-85 | 7 |
| RPL30 | -0.9057921 | 5.82E-85 | 7 |
| HSP90AA1 | 1.00201813 | 4.50E-84 | 7 |
| DEK | 0.93473063 | 1.27E-83 | 7 |
| TMEM237 | 0.36422567 | 3.28E-83 | 7 |
| HCST | -1.3064017 | 4.85E-83 | 7 |
| DTL | 0.41740969 | 7.44E-83 | 7 |
| GNA15 | 0.67093967 | 1.11E-82 | 7 |
| HIST2H4B | 0.25889002 | 1.16E-82 | 7 |
| DDX39A | 0.89106504 | 1.19E-82 | 7 |
| PNRC1 | -1.2449784 | 1.36E-82 | 7 |
| TIMELESS | 0.53212853 | 5.47E-82 | 7 |
| CHTF18 | 0.30404168 | 1.42E-81 | 7 |
| CCDC18 | 0.66231146 | 2.10E-81 | 7 |
| ZGRF1 | 0.4352124 | 1.33E-80 | 7 |
| SAE1 | 0.69763185 | 2.18E-80 | 7 |
| COX8A | 1.0220416 | 2.18E-80 | 7 |
| APOLD1 | 0.79872826 | 2.20E-80 | 7 |
| RPL37 | -0.8406445 | 2.72E-80 | 7 |
| C9orf40 | 0.50026534 | 3.50E-80 | 7 |
| CENPP | 0.37421768 | 1.02E-79 | 7 |
| MASTL | 0.42097521 | 1.59E-79 | 7 |
| TMEM106C | 0.8766336 | 1.64E-79 | 7 |
| NSD2 | 0.97030592 | 1.99E-79 | 7 |
| VRK1 | 0.70180028 | 2.08E-79 | 7 |
| TXNIP | -1.5472557 | 2.68E-79 | 7 |
| TAF5 | 0.34492376 | 5.39E-79 | 7 |
| TPT1 | -0.8045638 | 8.10E-78 | 7 |
| ALYREF | 0.91027509 | 8.67E-78 | 7 |
| ZNF730 | 0.30870019 | 1.05E-77 | 7 |
| RPS27A | -0.840693 | 1.42E-77 | 7 |
| CD52 | -1.2723697 | 1.51E-77 | 7 |
| E2F2 | 0.68578588 | 2.29E-77 | 7 |
| B2M | -1.2296338 | 5.17E-77 | 7 |
| SLFN5 | -1.25165 | 7.78E-77 | 7 |
| SIVA1 | 1.23578051 | 1.40E-76 | 7 |
| RPL26 | -0.8560373 | 1.45E-76 | 7 |

| | | | |
|---|---|---|---|
| HELLPAR | 0.56241113 | 2.04E-76 | 7 |
| RANBP1 | 1.05278344 | 2.66E-76 | 7 |
| NUDT1 | 0.66573404 | 2.75E-76 | 7 |
| SAP30 | 0.5013382 | 6.12E-76 | 7 |
| HIST1H1E | 1.30344535 | 1.05E-75 | 7 |
| ZNF704 | 0.60323833 | 1.87E-75 | 7 |
| LSM4 | 0.98818674 | 2.55E-75 | 7 |
| SASS6 | 0.50600713 | 4.82E-75 | 7 |
| LIG1 | 0.69828842 | 8.64E-75 | 7 |
| HIST1H2AK | 0.43012825 | 2.25E-74 | 7 |
| C4orf46 | 0.3594028 | 7.16E-74 | 7 |
| FANCG | 0.4085422 | 8.66E-74 | 7 |
| RPL10 | -0.8802172 | 1.24E-73 | 7 |
| H2AFY | 0.95505162 | 2.09E-73 | 7 |
| ZWILCH | 0.43933448 | 3.29E-73 | 7 |
| HIST1H2BN | 0.4191618 | 3.66E-73 | 7 |
| C5orf34 | 0.27049618 | 1.09E-72 | 7 |
| DUT | 1.52087367 | 4.54E-72 | 7 |
| NET1 | 0.49651727 | 4.62E-72 | 7 |
| SLC16A1 | 0.41172146 | 1.22E-71 | 7 |
| CENPL | 0.5001083 | 2.11E-71 | 7 |
| RPS15A | -0.7839428 | 5.29E-71 | 7 |
| CDCA4 | 0.68086618 | 9.37E-70 | 7 |
| SRSF2 | 0.90531557 | 1.53E-69 | 7 |
| SRSF3 | 0.86197515 | 1.77E-69 | 7 |
| HLA-E | -1.1203605 | 2.20E-69 | 7 |
| TMPO-AS1 | 0.32422213 | 3.00E-69 | 7 |
| TDP1 | 0.35975048 | 3.35E-69 | 7 |
| SNRPG | 0.92300021 | 5.75E-69 | 7 |
| RPS13 | -0.8000433 | 7.11E-69 | 7 |
| ANAPC11 | 0.9379043 | 7.82E-69 | 7 |
| RPL13A | -0.7187715 | 2.33E-68 | 7 |
| GAPDH | 1.04660517 | 2.57E-68 | 7 |
| BRI3BP | 0.49071269 | 6.39E-68 | 7 |
| CHAF1A | 0.63434898 | 1.42E-67 | 7 |
| WEE1 | 0.70339475 | 2.73E-67 | 7 |
| CEP78 | 0.85122036 | 3.61E-67 | 7 |
| RPS28 | -0.6923755 | 4.00E-67 | 7 |
| CARHSP1 | 0.92571549 | 6.60E-67 | 7 |
| NDST3 | 0.34161482 | 2.32E-66 | 7 |
| IDH2 | 0.9985542 | 2.77E-66 | 7 |
| HDGF | 0.88532931 | 4.08E-66 | 7 |
| RPL35A | -0.7645967 | 1.60E-65 | 7 |

| | | | |
|---|---|---|---|
| GIHCG | 0.97278965 | 3.55E-65 | 7 |
| RPS29 | -0.6742059 | 2.26E-64 | 7 |
| HNRNPD | 0.8704102 | 2.56E-64 | 7 |
| RPL11 | -0.6663916 | 5.21E-64 | 7 |
| HMGN1 | 0.66953046 | 5.24E-64 | 7 |
| SYNE2 | 1.14788255 | 7.02E-64 | 7 |
| YBX1 | 0.82781895 | 7.58E-64 | 7 |
| GSTP1 | 1.01923983 | 1.19E-63 | 7 |
| RPL9 | -0.7044574 | 1.20E-63 | 7 |
| LTB | -1.6967789 | 1.39E-63 | 7 |
| STK17B | -1.0628655 | 1.47E-63 | 7 |
| KLRB1 | -1.2677101 | 1.55E-63 | 7 |
| BCL2 | 0.34539594 | 2.26E-63 | 7 |
| PRELID2 | 0.32731968 | 2.94E-63 | 7 |
| RBMX | 0.82598266 | 6.73E-63 | 7 |
| MZT1 | 0.65085644 | 8.20E-63 | 7 |
| DNA2 | 0.53312274 | 1.51E-62 | 7 |
| TNF | 0.39202633 | 2.92E-62 | 7 |
| PCNA | 1.22151552 | 7.15E-62 | 7 |
| IL32 | -1.0508657 | 8.74E-62 | 7 |
| CD27 | -0.9825435 | 9.00E-62 | 7 |
| ORC1 | 0.27000536 | 1.02E-61 | 7 |
| TFDP2 | 1.03401155 | 3.33E-61 | 7 |
| BOLA3 | 0.44259674 | 7.70E-61 | 7 |
| HNRNPAB | 0.96849035 | 1.57E-60 | 7 |
| DBF4B | 0.27037339 | 1.72E-60 | 7 |
| EVL | -0.8711378 | 1.88E-60 | 7 |
| LDHA | 0.84479629 | 1.93E-60 | 7 |
| AC011447.3 | 0.42171151 | 2.27E-60 | 7 |
| ARPP21 | 0.81289308 | 4.01E-60 | 7 |
| CCDC15 | 0.39440578 | 8.02E-60 | 7 |
| RPL17 | -0.7081771 | 1.03E-59 | 7 |
| CSE1L | 0.50370123 | 1.33E-59 | 7 |
| RPL39 | -0.6866492 | 2.05E-59 | 7 |
| BTG1 | -1.1702653 | 2.45E-59 | 7 |
| HIST1H4H | 0.31408628 | 7.19E-59 | 7 |
| CLGN | 0.35905852 | 1.09E-58 | 7 |
| ZNF683 | -1.5326814 | 1.37E-58 | 7 |
| CHEK1 | 0.56958414 | 3.63E-58 | 7 |
| RPS24 | -0.635215 | 4.68E-58 | 7 |
| LSM3 | 0.80488857 | 6.36E-58 | 7 |
| TRIM28 | 0.74901394 | 6.50E-58 | 7 |
| CD99 | 0.90086002 | 6.62E-58 | 7 |

| | | | |
|---|---|---|---|
| VPREB1 | 0.41598633 | 7.46E-58 | 7 |
| FIGNL1 | 0.35116995 | 1.19E-57 | 7 |
| STK4 | -0.9498528 | 2.11E-57 | 7 |
| LIME1 | -0.9434095 | 2.16E-57 | 7 |
| RPL13 | -0.6679592 | 2.43E-57 | 7 |
| AC002454.1 | 0.84438918 | 2.92E-57 | 7 |
| ZNF724 | 0.39256183 | 5.99E-57 | 7 |
| SNRPD1 | 0.78616959 | 7.66E-57 | 7 |
| RPS4X | -0.6661523 | 1.29E-56 | 7 |
| LSM5 | 0.66846664 | 3.35E-56 | 7 |
| ZFP36L1 | -1.105673 | 6.39E-56 | 7 |
| PTGES3 | 0.76372429 | 1.86E-55 | 7 |
| RPL18 | -0.6235012 | 2.78E-55 | 7 |
| RCC1 | 0.45888389 | 3.56E-55 | 7 |
| C19orf48 | 0.53415435 | 1.67E-54 | 7 |
| NAA38 | 0.74434181 | 2.76E-54 | 7 |
| RPL32 | -0.6630212 | 3.27E-54 | 7 |
| GSTCD | 0.3127696 | 3.32E-54 | 7 |
| FTL | -1.0083541 | 3.40E-54 | 7 |
| ACAP1 | -0.8405794 | 7.37E-54 | 7 |
| ZNF519 | 0.42473073 | 8.64E-54 | 7 |
| CCNE2 | 0.47590545 | 8.87E-54 | 7 |
| POLD1 | 0.38107125 | 8.94E-54 | 7 |
| LIN9 | 0.29377747 | 1.16E-53 | 7 |
| MT2A | 0.51432338 | 1.52E-53 | 7 |
| ZNF367 | 0.39845883 | 1.80E-53 | 7 |
| SLC29A1 | 0.35675322 | 2.09E-53 | 7 |
| HADH | 0.61776447 | 2.51E-53 | 7 |
| SNRPF | 0.78497971 | 2.55E-53 | 7 |
| CDK6 | 1.0347195 | 3.78E-53 | 7 |
| TXN | 0.92736268 | 1.31E-52 | 7 |
| SMC1A | 0.86964142 | 1.45E-52 | 7 |
| HNRNPR | 0.7391507 | 1.90E-52 | 7 |
| POLE2 | 0.30934647 | 1.97E-52 | 7 |
| HMGN5 | 0.44858581 | 2.22E-52 | 7 |
| SKA2 | 0.74130462 | 2.22E-52 | 7 |
| MACF1 | -0.9989833 | 2.39E-52 | 7 |
| RPS3A | -0.5911619 | 5.06E-52 | 7 |
| CAPG | 0.86193472 | 1.05E-51 | 7 |
| RPA3 | 0.60680711 | 1.26E-51 | 7 |
| CCP110 | 0.58118675 | 1.71E-51 | 7 |
| JPT1 | 0.95860118 | 4.17E-51 | 7 |
| ILF2 | 0.72663372 | 6.76E-51 | 7 |

| | | | |
|---|---|---|---|
| DNMT3B | 0.29139963 | 6.92E-51 | 7 |
| JAK1 | -0.8528899 | 1.07E-50 | 7 |
| RPS18 | -0.6549836 | 1.16E-50 | 7 |
| FYB1 | -0.868606 | 1.32E-50 | 7 |
| ERH | 0.70693348 | 1.35E-50 | 7 |
| MAP3K20 | 0.45177882 | 1.42E-50 | 7 |
| CALM2 | 0.7088229 | 1.54E-50 | 7 |
| CDK5RAP3 | 0.75615407 | 2.27E-50 | 7 |
| CEP57L1 | 0.41759379 | 2.45E-50 | 7 |
| ACOT7 | 0.49479023 | 2.58E-50 | 7 |
| FANCB | 0.29057963 | 3.60E-50 | 7 |
| NDC1 | 0.31597101 | 4.22E-50 | 7 |
| PSIP1 | 0.70254027 | 4.59E-50 | 7 |
| SET | 0.66484782 | 4.92E-50 | 7 |
| MIR4435-2HG | 0.25488769 | 5.09E-50 | 7 |
| PSMC3IP | 0.27361183 | 8.40E-50 | 7 |
| NEMP1 | 0.42687238 | 9.13E-50 | 7 |
| H1FX | 0.94050812 | 1.17E-49 | 7 |
| N4BP2L2 | -0.8493091 | 1.53E-49 | 7 |
| PSMA6 | 0.75684539 | 2.08E-49 | 7 |
| RFC5 | 0.38353844 | 2.22E-49 | 7 |
| CD1B | 0.88826752 | 3.62E-49 | 7 |
| CD247 | -0.9076122 | 3.69E-49 | 7 |
| CALM3 | 0.78650166 | 4.35E-49 | 7 |
| RPL29 | -0.6388037 | 4.66E-49 | 7 |
| ADA | 0.7854525 | 4.82E-49 | 7 |
| PPIA | 0.78588511 | 8.02E-49 | 7 |
| ZNF726 | 0.37237813 | 8.99E-49 | 7 |
| SMC3 | 0.76416365 | 9.85E-49 | 7 |
| IFI27L1 | 0.35509132 | 1.03E-48 | 7 |
| HMGA1 | 0.90398272 | 1.52E-48 | 7 |
| CDKN2D | 0.98105144 | 1.56E-48 | 7 |
| HNRNPA0 | 0.69666514 | 2.74E-48 | 7 |
| SSRP1 | 0.75665232 | 3.44E-48 | 7 |
| ARL6IP1 | 1.47123664 | 3.95E-48 | 7 |
| LIMD2 | -0.8454788 | 6.18E-48 | 7 |
| RPL6 | -0.6352841 | 6.19E-48 | 7 |
| FXYD2 | 1.46137704 | 6.95E-48 | 7 |
| XPO1 | 0.71782962 | 7.59E-48 | 7 |
| POLA2 | 0.33119681 | 9.49E-48 | 7 |
| PPP2R3B | 0.37746638 | 9.75E-48 | 7 |
| MRPL51 | 0.68315979 | 1.03E-47 | 7 |
| PSMC3 | 0.72454114 | 1.11E-47 | 7 |

| | | | |
|---|---|---|---|
| USP13 | 0.36813509 | 1.75E-47 | 7 |
| CKAP2 | 0.85332374 | 2.05E-47 | 7 |
| NOTCH1 | 0.63146338 | 2.77E-47 | 7 |
| YWHAE | 0.75421527 | 3.65E-47 | 7 |
| PIM1 | 0.30856425 | 1.27E-46 | 7 |
| CEP76 | 0.31481343 | 1.38E-46 | 7 |
| ST18 | 0.42097881 | 1.69E-46 | 7 |
| COPRS | 0.35337582 | 1.77E-46 | 7 |
| CD1A | 0.98607984 | 3.20E-46 | 7 |
| TAF9B | 0.25312502 | 4.13E-46 | 7 |
| RSRP1 | -0.7809978 | 7.00E-46 | 7 |
| CEP295 | 0.52082212 | 1.24E-45 | 7 |
| MRPL57 | 0.7370196 | 1.30E-45 | 7 |
| SCRIB | -0.9911982 | 1.37E-45 | 7 |
| CD7 | -0.9589983 | 1.39E-45 | 7 |
| AAMDC | 0.33547099 | 1.67E-45 | 7 |
| RPL36 | -0.6007107 | 2.65E-45 | 7 |
| ARHGAP33 | 0.37034831 | 4.28E-45 | 7 |
| RPS8 | -0.5873827 | 6.69E-45 | 7 |
| RPL19 | -0.5902431 | 6.85E-45 | 7 |
| CNTROB | 0.30318165 | 7.77E-45 | 7 |
| NACA | -0.6257273 | 1.09E-44 | 7 |
| CD8B | -0.825291 | 1.28E-44 | 7 |
| PA2G4 | 0.75675215 | 1.62E-44 | 7 |
| RPL3 | -0.5317407 | 1.71E-44 | 7 |
| FABP5 | 0.93649706 | 1.90E-44 | 7 |
| RPS3 | -0.5362079 | 4.65E-44 | 7 |
| MCM7 | 0.93674041 | 6.11E-44 | 7 |
| SMS | 0.6522728 | 6.12E-44 | 7 |
| IKZF2 | -1.0832617 | 6.13E-44 | 7 |
| EFCAB2 | 0.3513083 | 1.52E-43 | 7 |
| MCM8 | 0.39641981 | 1.77E-43 | 7 |
| MXD4 | -0.8250891 | 2.37E-43 | 7 |
| RPS7 | -0.5788367 | 3.03E-43 | 7 |
| SNRNP25 | 0.43700875 | 3.84E-43 | 7 |
| TRAC | -0.8627533 | 4.10E-43 | 7 |
| DIP2A | -0.8801078 | 4.24E-43 | 7 |
| NASP | 0.75375261 | 4.99E-43 | 7 |
| HSPA1A | 1.17102441 | 1.06E-42 | 7 |
| DSN1 | 0.34689722 | 1.25E-42 | 7 |
| SLC43A3 | 0.39715682 | 1.53E-42 | 7 |
| EIF4A3 | 0.67319511 | 2.62E-42 | 7 |
| DONSON | 0.28062569 | 3.20E-42 | 7 |

| | | | |
|---|---|---|---|
| BCL7A | 0.53137213 | 3.34E-42 | 7 |
| TRAF3IP3 | -0.7677762 | 3.79E-42 | 7 |
| RPL41 | -0.5468191 | 6.60E-42 | 7 |
| RNASEH2C | 0.67175155 | 7.29E-42 | 7 |
| SIRPG | -0.9446024 | 9.97E-42 | 7 |
| SPATS2L | 0.40018405 | 1.06E-41 | 7 |
| MAZ | 0.64473374 | 1.19E-41 | 7 |
| PCDH10 | 0.28742896 | 1.27E-41 | 7 |
| DARS2 | 0.31385754 | 1.33E-41 | 7 |
| COX17 | 0.63926684 | 1.59E-41 | 7 |
| PHGDH | 0.54961705 | 1.73E-41 | 7 |
| BTG3 | 0.52702477 | 1.89E-41 | 7 |
| IFI16 | 0.69327248 | 3.58E-41 | 7 |
| RPL12 | -0.6312766 | 4.63E-41 | 7 |
| PTPRC | -0.6879986 | 5.92E-41 | 7 |
| NKAIN4 | 0.49896235 | 6.59E-41 | 7 |
| RPL22 | -0.5852716 | 7.50E-41 | 7 |
| POP7 | 0.47463252 | 2.82E-40 | 7 |
| FANCM | 0.30982315 | 3.59E-40 | 7 |
| CMSS1 | 0.41215271 | 3.87E-40 | 7 |
| ATP5F1B | 0.61824068 | 4.10E-40 | 7 |
| CLEC2D | -1.0204649 | 4.12E-40 | 7 |
| ITGB2 | -0.8147623 | 4.37E-40 | 7 |
| SFPQ | 0.70452681 | 4.42E-40 | 7 |
| SNRPC | 0.61655455 | 5.51E-40 | 7 |
| HNRNPA1 | 0.56271421 | 5.56E-40 | 7 |
| CDK5RAP2 | 0.64806015 | 7.65E-40 | 7 |
| UQCRQ | 0.68911797 | 9.56E-40 | 7 |
| CD1E | 0.72854214 | 1.27E-39 | 7 |
| PHIP | 0.70391706 | 1.41E-39 | 7 |
| BANF1 | 0.65362558 | 1.62E-39 | 7 |
| LIN54 | 0.37399909 | 1.75E-39 | 7 |
| NCAPH2 | 0.43953219 | 1.98E-39 | 7 |
| RPS25 | -0.5174297 | 2.04E-39 | 7 |
| YEATS4 | 0.5210737 | 2.26E-39 | 7 |
| PRMT2 | -0.8228908 | 2.43E-39 | 7 |
| RPL38 | -0.5398147 | 2.75E-39 | 7 |
| CBX1 | 0.60602533 | 4.35E-39 | 7 |
| ATP5F1E | -0.5705232 | 5.30E-39 | 7 |
| LCP2 | -0.7495311 | 6.28E-39 | 7 |
| SLC25A5 | 0.63886491 | 6.50E-39 | 7 |
| RPS15 | -0.4825052 | 8.09E-39 | 7 |
| PXMP2 | 0.51276245 | 8.45E-39 | 7 |

| | | | |
|---|---|---|---|
| SEPHS1 | 0.52166936 | 9.60E-39 | 7 |
| TTN | -1.040769 | 1.20E-38 | 7 |
| CIB2 | 0.3616127 | 1.27E-38 | 7 |
| PMAIP1 | 0.31104751 | 1.61E-38 | 7 |
| PFDN5 | -0.6210404 | 1.82E-38 | 7 |
| RACK1 | -0.5483143 | 1.90E-38 | 7 |
| AC026401.3 | 0.27215814 | 2.13E-38 | 7 |
| YWHAQ | 0.61291774 | 3.06E-38 | 7 |
| ATP6AP1L | 0.35470972 | 3.64E-38 | 7 |
| DAZAP1 | 0.61065415 | 3.80E-38 | 7 |
| RFWD3 | 0.4621814 | 4.28E-38 | 7 |
| IL16 | -0.795332 | 6.23E-38 | 7 |
| TALDO1 | 0.63277408 | 8.24E-38 | 7 |
| POLA1 | 0.40506578 | 9.47E-38 | 7 |
| DCK | 0.5847014 | 1.05E-37 | 7 |
| RPL21 | -0.5074993 | 1.16E-37 | 7 |
| PIDD1 | 0.32134552 | 2.21E-37 | 7 |
| RPL15 | -0.4934334 | 2.58E-37 | 7 |
| RPL7A | -0.4778834 | 3.48E-37 | 7 |
| EXOSC8 | 0.47933043 | 3.90E-37 | 7 |
| ANP32A | 0.63033014 | 4.38E-37 | 7 |
| MED30 | 0.50195683 | 4.64E-37 | 7 |
| RPLP2 | -0.4686701 | 8.38E-37 | 7 |
| SRGN | -0.871084 | 1.18E-36 | 7 |
| COX5A | 0.62098201 | 1.54E-36 | 7 |
| CALR | 0.80802596 | 1.74E-36 | 7 |
| RPS23 | -0.4644339 | 2.08E-36 | 7 |
| POLH | 0.34934914 | 2.08E-36 | 7 |
| IGLL1 | 0.74591901 | 2.24E-36 | 7 |
| LST1 | 0.83149705 | 2.63E-36 | 7 |
| HPF1 | 0.55471631 | 3.10E-36 | 7 |
| CDK2 | 0.38092992 | 3.17E-36 | 7 |
| SNRPB | 0.6423455 | 3.40E-36 | 7 |
| CYTOR | 0.33997652 | 4.71E-36 | 7 |
| POLD2 | 0.33683655 | 5.39E-36 | 7 |
| RPS10 | -0.5843919 | 7.18E-36 | 7 |
| PPIH | 0.40254114 | 8.64E-36 | 7 |
| ATM | -0.7890866 | 9.84E-36 | 7 |
| BLM | 0.45309056 | 1.04E-35 | 7 |
| RBM8A | 0.60343336 | 1.09E-35 | 7 |
| PSMA4 | 0.60240718 | 1.09E-35 | 7 |
| GMPPB | 0.26994962 | 1.55E-35 | 7 |
| SHMT1 | 0.413359 | 1.79E-35 | 7 |

| | | | |
|---|---|---|---|
| SPIN4 | 0.35255305 | 3.13E-35 | 7 |
| ATAD3A | 0.28671964 | 3.57E-35 | 7 |
| RPL27 | -0.5229934 | 4.53E-35 | 7 |
| RMI2 | 0.38633147 | 4.91E-35 | 7 |
| CACYBP | 0.57415239 | 1.13E-34 | 7 |
| RALY | 0.63796293 | 1.20E-34 | 7 |
| NOP56 | 0.61544777 | 1.33E-34 | 7 |
| RPL31 | -0.5566978 | 1.46E-34 | 7 |
| MAGOH | 0.56566521 | 1.50E-34 | 7 |
| ELF1 | -0.7367348 | 1.94E-34 | 7 |
| CRIP1 | 0.8377729 | 2.19E-34 | 7 |
| RUSC1 | 0.32912531 | 2.43E-34 | 7 |
| CCSAP | 0.36593192 | 2.82E-34 | 7 |
| GINS2 | 0.42256234 | 3.77E-34 | 7 |
| ZFP36L2 | -0.8543062 | 3.94E-34 | 7 |
| LEF1 | -0.5852291 | 4.94E-34 | 7 |
| VDAC3 | 0.50142181 | 5.10E-34 | 7 |
| KLHL23 | 0.55019566 | 5.72E-34 | 7 |
| CTHRC1 | 0.35716587 | 7.86E-34 | 7 |
| BCL11A | 0.50514086 | 1.00E-33 | 7 |
| BARD1 | 0.59775926 | 1.35E-33 | 7 |
| GNB4 | 0.26321747 | 1.35E-33 | 7 |
| COMMD6 | -0.6500102 | 1.37E-33 | 7 |
| HSPD1 | 0.82081689 | 1.76E-33 | 7 |
| NDUFAF3 | 0.54455439 | 1.86E-33 | 7 |
| NCF1 | -0.7410177 | 1.87E-33 | 7 |
| BCL2L12 | 0.37033696 | 1.89E-33 | 7 |
| RUVBL2 | 0.34437207 | 1.93E-33 | 7 |
| COTL1 | -0.6968257 | 2.09E-33 | 7 |
| CD44 | -0.8508134 | 2.86E-33 | 7 |
| CHCHD3 | 0.53581095 | 3.72E-33 | 7 |
| RBBP8 | 0.65247218 | 4.29E-33 | 7 |
| MAGOHB | 0.43025582 | 5.51E-33 | 7 |
| NDUFB11 | 0.58426724 | 5.89E-33 | 7 |
| PSMG1 | 0.34160261 | 1.19E-32 | 7 |
| RPL28 | -0.4235816 | 1.54E-32 | 7 |
| CMC2 | 0.56975916 | 1.63E-32 | 7 |
| MPP1 | 0.28021495 | 1.90E-32 | 7 |
| SLC20A1 | 0.49123774 | 2.09E-32 | 7 |
| SNRPD3 | 0.51047181 | 3.16E-32 | 7 |
| C12orf75 | 0.6427541 | 3.44E-32 | 7 |
| ABLIM1 | -0.8010619 | 3.87E-32 | 7 |
| VIM | 0.58984276 | 4.53E-32 | 7 |

| | | | |
|---|---|---|---|
| ECI1 | 0.44412339 | 4.74E-32 | 7 |
| CTSH | 0.27635024 | 4.91E-32 | 7 |
| HLA-A | -0.9183334 | 5.18E-32 | 7 |
| ARHGAP19 | 0.42741709 | 6.06E-32 | 7 |
| SH3BGRL3 | -0.688631 | 6.61E-32 | 7 |
| SORT1 | 0.25515261 | 6.78E-32 | 7 |
| POGLUT3 | 0.33775781 | 7.39E-32 | 7 |
| HMGXB4 | 0.49418138 | 7.65E-32 | 7 |
| CEP85 | 0.25377629 | 8.95E-32 | 7 |
| HEMGN | 0.47286983 | 1.20E-31 | 7 |
| GPN3 | 0.30696015 | 1.34E-31 | 7 |
| MPHOSPH9 | 0.58009347 | 1.77E-31 | 7 |
| XRCC6 | 0.59669356 | 1.91E-31 | 7 |
| ZNF714 | 0.34087794 | 2.34E-31 | 7 |
| NAP1L1 | 0.48315642 | 2.46E-31 | 7 |
| ACYP1 | 0.47036198 | 2.88E-31 | 7 |
| SLC4A4 | 0.27382707 | 3.41E-31 | 7 |
| EIF5A | 0.60374253 | 4.17E-31 | 7 |
| HNRNPM | 0.58589064 | 4.35E-31 | 7 |
| PTBP1 | 0.55978706 | 5.27E-31 | 7 |
| HNRNPDL | 0.58391839 | 5.73E-31 | 7 |
| AMOTL1 | 0.32790512 | 5.99E-31 | 7 |
| HELLS | 0.7677207 | 6.09E-31 | 7 |
| MZT2B | 0.52044746 | 6.20E-31 | 7 |
| KHDRBS1 | 0.55392118 | 6.94E-31 | 7 |
| ABHD12 | 0.27338338 | 8.80E-31 | 7 |
| SCML2 | 0.26404656 | 1.10E-30 | 7 |
| RPS14 | -0.5037739 | 1.11E-30 | 7 |
| CTCF | 0.60356979 | 1.15E-30 | 7 |
| UBE2H | -0.7081918 | 1.27E-30 | 7 |
| SATB1 | -1.0567527 | 1.49E-30 | 7 |
| RNF213 | -0.6995349 | 1.57E-30 | 7 |
| SKP2 | 0.30145272 | 1.69E-30 | 7 |
| NME1 | 0.5188301 | 1.99E-30 | 7 |
| HLA-B | -0.9603692 | 2.14E-30 | 7 |
| DDX17 | -0.5312648 | 2.21E-30 | 7 |
| ATP23 | 0.3284292 | 2.22E-30 | 7 |
| CKLF | 0.63104346 | 2.42E-30 | 7 |
| WDR76 | 0.53610255 | 3.10E-30 | 7 |
| POLE | 0.43143146 | 3.36E-30 | 7 |
| HILPDA | 0.32402498 | 3.42E-30 | 7 |
| SNRPA1 | 0.50243202 | 4.24E-30 | 7 |
| FAU | -0.4376996 | 4.44E-30 | 7 |

| | | | |
|---|---|---|---|
| NONO | 0.53052604 | 4.47E-30 | 7 |
| RPL24 | -0.4914647 | 4.54E-30 | 7 |
| ACTN1 | -0.6751313 | 6.07E-30 | 7 |
| DLEU2 | 0.71467704 | 6.25E-30 | 7 |
| HP1BP3 | 0.72615462 | 7.50E-30 | 7 |
| PRKCH | -0.7131774 | 1.28E-29 | 7 |
| SAMD3 | -0.7281665 | 1.31E-29 | 7 |
| MT-ND3 | -0.5497202 | 1.38E-29 | 7 |
| PPIF | 0.32244221 | 1.45E-29 | 7 |
| DNAJB1 | 0.71583232 | 1.64E-29 | 7 |
| RBX1 | 0.58975828 | 1.80E-29 | 7 |
| CSNK2B | 0.52816481 | 2.29E-29 | 7 |
| SCD5 | 0.29662934 | 3.08E-29 | 7 |
| MYO1F | -0.676064 | 3.74E-29 | 7 |
| THORLNC | 0.34707793 | 5.92E-29 | 7 |
| PRPSAP1 | 0.37371674 | 6.23E-29 | 7 |
| RTTN | 0.29347926 | 9.33E-29 | 7 |
| ANAPC15 | 0.44656206 | 1.00E-28 | 7 |
| HNRNPH3 | 0.52797497 | 1.04E-28 | 7 |
| PNISR | -0.6428988 | 1.15E-28 | 7 |
| SRGAP2 | 0.26774486 | 1.32E-28 | 7 |
| IMPDH2 | 0.43325421 | 1.35E-28 | 7 |
| MICB | 0.27647601 | 1.37E-28 | 7 |
| GSS | 0.29494236 | 1.59E-28 | 7 |
| PBX3 | 0.25437453 | 1.85E-28 | 7 |
| XIST | -0.6772298 | 2.36E-28 | 7 |
| PALM2-AKAP2 | 0.57409331 | 3.28E-28 | 7 |
| SNHG14 | -0.7905006 | 3.58E-28 | 7 |
| CELF2 | -0.6627333 | 5.76E-28 | 7 |
| GAS5 | -0.6911209 | 6.64E-28 | 7 |
| FLNB | 0.40898702 | 7.73E-28 | 7 |
| HSPB11 | 0.61880524 | 9.19E-28 | 7 |
| FCMR | -0.6264962 | 1.02E-27 | 7 |
| CCT5 | 0.50467273 | 1.07E-27 | 7 |
| PAXIP1 | 0.39504467 | 1.21E-27 | 7 |
| PSMB2 | 0.515809 | 1.38E-27 | 7 |
| HCFC1 | 0.49543788 | 1.43E-27 | 7 |
| RHNO1 | 0.32329695 | 1.65E-27 | 7 |
| RPS9 | -0.4264547 | 2.75E-27 | 7 |
| TOMM5 | 0.54858594 | 2.79E-27 | 7 |
| PRDX3 | 0.49832188 | 2.86E-27 | 7 |
| CFAP20 | 0.39636913 | 2.94E-27 | 7 |
| ILF3 | 0.52229854 | 3.44E-27 | 7 |

| | | | |
|---|---|---|---|
| CD79A | -0.8256291 | 3.65E-27 | 7 |
| CD2 | -0.6585191 | 3.80E-27 | 7 |
| FADS2 | 0.27951709 | 3.82E-27 | 7 |
| CDC7 | 0.27733368 | 5.26E-27 | 7 |
| HLA-C | -0.7649596 | 5.35E-27 | 7 |
| ICAM3 | -0.5990285 | 5.65E-27 | 7 |
| TRA2B | 0.53979608 | 5.76E-27 | 7 |
| LBH | -0.6654669 | 6.05E-27 | 7 |
| HSPA8 | 0.57883307 | 6.59E-27 | 7 |
| ETV5 | 0.64376611 | 1.02E-26 | 7 |
| PSMB3 | 0.50526463 | 1.05E-26 | 7 |
| PDCD4 | -0.6358682 | 1.22E-26 | 7 |
| GNAS | 0.4406879 | 1.37E-26 | 7 |
| TCTEX1D2 | 0.38814115 | 1.51E-26 | 7 |
| CTSA | -0.62797 | 1.62E-26 | 7 |
| CENPQ | 0.27307193 | 2.09E-26 | 7 |
| ARPC1B | -0.5526083 | 2.86E-26 | 7 |
| LRRC59 | 0.33784498 | 2.90E-26 | 7 |
| KDM5B | -0.6654004 | 2.96E-26 | 7 |
| PI4K2B | 0.27433071 | 3.18E-26 | 7 |
| CYBA | -0.5908673 | 3.28E-26 | 7 |
| TMSB4X | -0.4055408 | 3.34E-26 | 7 |
| RBBP7 | 0.48115435 | 3.42E-26 | 7 |
| IER5 | 0.37495505 | 3.48E-26 | 7 |
| UBALD2 | 0.5209602 | 3.61E-26 | 7 |
| SUPT16H | 0.63447792 | 4.57E-26 | 7 |
| MCM2 | 0.38039199 | 5.07E-26 | 7 |
| TFDP1 | 0.48967581 | 5.36E-26 | 7 |
| ACAT2 | 0.38948062 | 6.96E-26 | 7 |
| MAT2A | 0.49645555 | 7.16E-26 | 7 |
| SAMHD1 | -0.6988302 | 7.34E-26 | 7 |
| COX7B | 0.46738055 | 8.33E-26 | 7 |
| SRSF10 | 0.52452103 | 8.82E-26 | 7 |
| BTG2 | -0.7277497 | 1.00E-25 | 7 |
| NDUFS8 | 0.47813652 | 1.01E-25 | 7 |
| GLRX5 | 0.52201598 | 1.19E-25 | 7 |
| DOCK10 | -0.6571037 | 1.25E-25 | 7 |
| H3F3A | 0.39705131 | 1.41E-25 | 7 |
| CCDC26 | 0.29330308 | 1.44E-25 | 7 |
| BLOC1S1 | 0.48188218 | 1.68E-25 | 7 |
| ESD | 0.4835391 | 1.79E-25 | 7 |
| NUP205 | 0.38090668 | 1.81E-25 | 7 |
| PCBP1 | 0.51576363 | 1.85E-25 | 7 |

| | | | |
|---|---|---|---|
| FTX | -0.7258619 | 1.96E-25 | 7 |
| NEAT1 | -0.7505347 | 2.10E-25 | 7 |
| SUMO2 | 0.46142975 | 3.15E-25 | 7 |
| CXCR4 | 0.54869398 | 3.31E-25 | 7 |
| RPL10A | -0.4528303 | 3.36E-25 | 7 |
| SRP9 | 0.50104127 | 3.83E-25 | 7 |
| CTSW | -0.798044 | 4.27E-25 | 7 |
| LRRCC1 | 0.37270385 | 4.76E-25 | 7 |
| CLEC11A | 0.51483272 | 4.99E-25 | 7 |
| SRSF7 | 0.54917076 | 5.04E-25 | 7 |
| SRRM1 | 0.50410912 | 5.33E-25 | 7 |
| PGP | 0.43963863 | 6.69E-25 | 7 |
| CD3G | -0.4963459 | 8.80E-25 | 7 |
| SARAF | -0.5908262 | 9.31E-25 | 7 |
| RHOH | -0.6324861 | 9.50E-25 | 7 |
| EEF1A1 | -0.3768033 | 9.53E-25 | 7 |
| PLCH1 | 0.27012107 | 1.15E-24 | 7 |
| PAICS | 0.51055949 | 1.20E-24 | 7 |
| CCDC88A | 0.56681642 | 1.38E-24 | 7 |
| MTHFD2 | 0.39143982 | 1.53E-24 | 7 |
| ICMT | 0.29801286 | 1.56E-24 | 7 |
| PKM | 0.56820252 | 2.40E-24 | 7 |
| WDHD1 | 0.33351864 | 2.75E-24 | 7 |
| TTF1 | 0.53003585 | 2.95E-24 | 7 |
| NENF | 0.31435157 | 3.03E-24 | 7 |
| CCT7 | 0.44643769 | 4.17E-24 | 7 |
| PRXL2A | 0.33815287 | 4.43E-24 | 7 |
| RPL7 | -0.4251124 | 5.21E-24 | 7 |
| CD8A | -0.681598 | 5.37E-24 | 7 |
| PARP2 | 0.2711041 | 5.69E-24 | 7 |
| NETO2 | 0.33961544 | 5.99E-24 | 7 |
| COX6A1 | 0.47494312 | 8.66E-24 | 7 |
| FAM102A | -0.6405508 | 9.98E-24 | 7 |
| COPS6 | 0.47811744 | 1.02E-23 | 7 |
| THOC6 | 0.25034124 | 1.10E-23 | 7 |
| VDAC1 | 0.5037115 | 1.11E-23 | 7 |
| ENO1 | 0.72046693 | 1.13E-23 | 7 |
| RABL6 | 0.43320358 | 1.40E-23 | 7 |
| NDUFB3 | 0.47540922 | 1.46E-23 | 7 |
| ATP5MC3 | 0.49613347 | 1.53E-23 | 7 |
| TENT4A | 0.27673932 | 1.91E-23 | 7 |
| MDH1 | 0.45365701 | 2.15E-23 | 7 |
| GLO1 | 0.4425894 | 2.16E-23 | 7 |

| | | | |
|---|---|---|---|
| PSMG2 | 0.46188576 | 2.83E-23 | 7 |
| CDK4 | 0.44543524 | 3.08E-23 | 7 |
| POLR2L | 0.52318306 | 3.09E-23 | 7 |
| CD37 | -0.5734507 | 3.28E-23 | 7 |
| ITGB2-AS1 | -0.536368 | 3.44E-23 | 7 |
| NELFE | 0.39134834 | 3.48E-23 | 7 |
| TSPAN5 | 0.28455076 | 3.96E-23 | 7 |
| NDUFS6 | 0.50743231 | 4.12E-23 | 7 |
| CNN2 | -0.7617504 | 4.29E-23 | 7 |
| RRP7A | 0.44237637 | 4.49E-23 | 7 |
| DDX5 | -0.4473112 | 4.50E-23 | 7 |
| HNRNPA3 | 0.47949751 | 6.00E-23 | 7 |
| LYAR | 0.35373593 | 9.32E-23 | 7 |
| LBR | 0.53317212 | 9.63E-23 | 7 |
| S100A6 | -0.7670774 | 1.11E-22 | 7 |
| FNBP1 | -0.5402833 | 1.30E-22 | 7 |
| NUP37 | 0.29490732 | 1.31E-22 | 7 |
| RAD51C | 0.32687302 | 1.32E-22 | 7 |
| SPINT2 | -0.4577636 | 1.39E-22 | 7 |
| ELOC | 0.42617354 | 1.42E-22 | 7 |
| SYNRG | -0.5911378 | 1.49E-22 | 7 |
| SNRPE | 0.52033392 | 1.57E-22 | 7 |
| ISG20 | -0.5633654 | 1.61E-22 | 7 |
| C9orf16 | -0.5061622 | 1.69E-22 | 7 |
| TCP1 | 0.49087908 | 1.72E-22 | 7 |
| COX7A2 | 0.47757797 | 2.23E-22 | 7 |
| AHCY | 0.31463578 | 2.24E-22 | 7 |
| PSMA7 | 0.4536224 | 2.42E-22 | 7 |
| RPS11 | -0.4188173 | 2.44E-22 | 7 |
| YPEL3 | -0.6073873 | 2.97E-22 | 7 |
| PBRM1 | 0.47021573 | 3.15E-22 | 7 |
| RPLP0 | 0.47736571 | 3.25E-22 | 7 |
| CTPS1 | 0.37912468 | 3.80E-22 | 7 |
| THOC3 | 0.31108234 | 3.92E-22 | 7 |
| CEP70 | 0.33718807 | 4.09E-22 | 7 |
| SPINDOC | 0.25654709 | 4.33E-22 | 7 |
| TCF7 | -0.5649123 | 4.35E-22 | 7 |
| RCAN1 | 0.33203961 | 4.37E-22 | 7 |
| RPS19 | -0.3622487 | 4.38E-22 | 7 |
| UCK2 | 0.26058506 | 5.03E-22 | 7 |
| CR1 | -0.7369342 | 5.17E-22 | 7 |
| GPANK1 | 0.3656034 | 5.40E-22 | 7 |
| CDK2AP1 | 0.49930141 | 5.54E-22 | 7 |

| | | | |
|---|---|---|---|
| IMMP1L | 0.30972618 | 5.71E-22 | 7 |
| MBP | -0.6816543 | 6.65E-22 | 7 |
| PMF1 | 0.42955106 | 7.40E-22 | 7 |
| SLBP | 0.55138995 | 8.09E-22 | 7 |
| POLR2K | 0.43998683 | 8.36E-22 | 7 |
| MYL12A | -0.4735561 | 8.57E-22 | 7 |
| PSMB6 | 0.43802823 | 1.09E-21 | 7 |
| CHMP7 | -0.6056976 | 1.17E-21 | 7 |
| RPL37A | -0.3282235 | 1.27E-21 | 7 |
| FAM107B | -0.6088112 | 1.35E-21 | 7 |
| FKBP3 | 0.45520296 | 1.36E-21 | 7 |
| CDKN2A | 0.34748363 | 1.48E-21 | 7 |
| PAFAH1B3 | 0.41825735 | 1.56E-21 | 7 |
| FLNA | -0.6132899 | 1.74E-21 | 7 |
| NDUFA6 | 0.46162456 | 1.98E-21 | 7 |
| PSME2 | 0.46264179 | 2.08E-21 | 7 |
| POLR2E | 0.40038269 | 2.10E-21 | 7 |
| CDC25B | 0.51910461 | 2.11E-21 | 7 |
| NDUFA2 | 0.45916788 | 2.18E-21 | 7 |
| TMC6 | -0.6004036 | 2.24E-21 | 7 |
| RAD18 | 0.28681219 | 3.24E-21 | 7 |
| ATP5MF | 0.41079101 | 3.94E-21 | 7 |
| ACTL6A | 0.39278532 | 3.94E-21 | 7 |
| FTH1 | 0.47008942 | 4.01E-21 | 7 |
| CCDC18-AS1 | -0.6053193 | 4.53E-21 | 7 |
| LSM7 | 0.48680853 | 4.69E-21 | 7 |
| HSPE1 | 0.62914075 | 4.91E-21 | 7 |
| CEP135 | 0.40455562 | 5.03E-21 | 7 |
| RFC4 | 0.38480832 | 5.78E-21 | 7 |
| ST6GAL1 | -0.5709667 | 5.92E-21 | 7 |
| PON1 | 0.26029466 | 5.98E-21 | 7 |
| ADK | 0.27133821 | 6.61E-21 | 7 |
| ODC1 | 0.36003709 | 8.16E-21 | 7 |
| ENY2 | 0.42367799 | 9.43E-21 | 7 |
| CHCHD2 | 0.5120007 | 1.00E-20 | 7 |
| RBM15 | 0.25343837 | 1.10E-20 | 7 |
| AP005482.1 | 0.51472128 | 1.11E-20 | 7 |
| VASP | -0.5629838 | 1.18E-20 | 7 |
| IKZF1 | -0.5489213 | 1.53E-20 | 7 |
| ZFAS1 | -0.5034101 | 1.55E-20 | 7 |
| CEP57 | 0.41656844 | 1.55E-20 | 7 |
| PCBP2 | 0.39687577 | 1.86E-20 | 7 |
| SNHG1 | 0.40276463 | 2.09E-20 | 7 |

| | | | |
|---|---|---|---|
| SLFN11 | 0.26413672 | 2.18E-20 | 7 |
| NUP107 | 0.33516998 | 2.21E-20 | 7 |
| TRMU | 0.28568545 | 2.24E-20 | 7 |
| IL27RA | -0.5493333 | 2.32E-20 | 7 |
| NT5DC2 | 0.33074259 | 2.47E-20 | 7 |
| DOK2 | -0.5330315 | 2.55E-20 | 7 |
| RAB5IF | 0.4068313 | 2.62E-20 | 7 |
| NDUFB2 | 0.45375377 | 2.77E-20 | 7 |
| GIMAP7 | -0.6603547 | 2.97E-20 | 7 |
| POLD3 | 0.35932703 | 3.00E-20 | 7 |
| PABPC1 | -0.4267168 | 3.39E-20 | 7 |
| ATP5IF1 | 0.46018057 | 3.67E-20 | 7 |
| AC004687.1 | -0.5773721 | 3.76E-20 | 7 |
| OGT | -0.5625257 | 3.87E-20 | 7 |
| MPDU1 | 0.26938805 | 4.65E-20 | 7 |
| UBA52 | -0.3567503 | 4.73E-20 | 7 |
| SUZ12 | 0.51730305 | 5.14E-20 | 7 |
| NUDT21 | 0.44477848 | 5.15E-20 | 7 |
| CD48 | -0.5479466 | 5.16E-20 | 7 |
| UQCRC1 | 0.39917251 | 6.13E-20 | 7 |
| KAT6B | -0.5396779 | 6.54E-20 | 7 |
| TOX2 | -0.6912305 | 6.73E-20 | 7 |
| RPS26 | -0.3948364 | 7.56E-20 | 7 |
| MT-CO1 | -0.3338614 | 8.34E-20 | 7 |
| NTAN1 | 0.34124448 | 8.74E-20 | 7 |
| HAUS1 | 0.42117441 | 8.93E-20 | 7 |
| RCSD1 | -0.5080279 | 9.01E-20 | 7 |
| DNMT1 | 0.53658289 | 1.04E-19 | 7 |
| NPM1 | 0.53879153 | 1.09E-19 | 7 |
| ATP1B3 | 0.42544224 | 1.16E-19 | 7 |
| CCNE1 | 0.41820757 | 1.39E-19 | 7 |
| TPRKB | 0.38327707 | 1.39E-19 | 7 |
| SFMBT1 | 0.31598052 | 1.54E-19 | 7 |
| TOMM7 | -0.485944 | 1.90E-19 | 7 |
| COL6A2 | -0.5807183 | 1.94E-19 | 7 |
| PSMD8 | 0.43704453 | 2.54E-19 | 7 |
| RBM17 | 0.42565175 | 2.95E-19 | 7 |
| RNASEH2B | 0.47837189 | 3.16E-19 | 7 |
| ARGLU1 | -0.4874533 | 3.35E-19 | 7 |
| PIK3IP1 | -0.4668128 | 3.64E-19 | 7 |
| CDC27 | 0.31040714 | 3.75E-19 | 7 |
| STK17A | -0.608358 | 4.12E-19 | 7 |
| EMC9 | 0.304179 | 5.85E-19 | 7 |

| | | | |
|---|---|---|---|
| PARD3 | 0.27867325 | 5.90E-19 | 7 |
| ELOB | 0.4362343 | 5.96E-19 | 7 |
| ATRX | -0.5634293 | 7.50E-19 | 7 |
| SLIRP | 0.39791531 | 7.78E-19 | 7 |
| RMI1 | 0.330508 | 7.95E-19 | 7 |
| BRD8 | 0.37729902 | 8.19E-19 | 7 |
| NUP85 | 0.30056169 | 8.37E-19 | 7 |
| CNTRL | 0.51231109 | 9.22E-19 | 7 |
| NUP93 | 0.28639796 | 9.53E-19 | 7 |
| RPL18A | -0.3730858 | 1.28E-18 | 7 |
| PDS5B | 0.47982199 | 1.39E-18 | 7 |
| CENPX | 0.41015269 | 1.41E-18 | 7 |
| BOP1 | 0.28125703 | 1.94E-18 | 7 |
| CYC1 | 0.41313478 | 2.04E-18 | 7 |
| ANXA1 | 0.49078162 | 2.17E-18 | 7 |
| NDUFV1 | 0.38495598 | 2.58E-18 | 7 |
| BUD13 | 0.26407828 | 2.65E-18 | 7 |
| TMEM14C | 0.4580295 | 2.83E-18 | 7 |
| GALNT2 | 0.48280613 | 2.99E-18 | 7 |
| FARSA | 0.30734075 | 3.97E-18 | 7 |
| DDB2 | 0.32026441 | 4.11E-18 | 7 |
| HCLS1 | -0.4929051 | 4.62E-18 | 7 |
| NUDC | 0.45211207 | 4.67E-18 | 7 |
| PSMD14 | 0.38995904 | 5.20E-18 | 7 |
| PAK1IP1 | 0.26014866 | 5.43E-18 | 7 |
| MZB1 | 0.47035006 | 5.67E-18 | 7 |
| TAF15 | 0.44273118 | 6.10E-18 | 7 |
| MCM4 | 0.47711272 | 6.26E-18 | 7 |
| CBFB | 0.38243905 | 7.80E-18 | 7 |
| AL138899.1 | 0.4642872 | 8.00E-18 | 7 |
| ZNF22 | 0.4303519 | 8.92E-18 | 7 |
| TOPBP1 | 0.40264113 | 1.01E-17 | 7 |
| ELAVL1 | 0.35624352 | 1.03E-17 | 7 |
| RAC2 | -0.4336872 | 1.18E-17 | 7 |
| PFDN2 | 0.40441367 | 1.20E-17 | 7 |
| HTATSF1 | 0.41700201 | 1.31E-17 | 7 |
| SAMD1 | 0.67459188 | 1.48E-17 | 7 |
| MPHOSPH6 | 0.28005992 | 1.49E-17 | 7 |
| MDFIC | 0.33343578 | 1.51E-17 | 7 |
| NEDD1 | 0.33625184 | 1.64E-17 | 7 |
| VPS29 | 0.41900959 | 2.22E-17 | 7 |
| CD63 | -0.5416725 | 2.52E-17 | 7 |
| SSBP1 | 0.40979648 | 2.58E-17 | 7 |

| | | | |
|---|---|---|---|
| PPP2R3C | 0.37282841 | 2.99E-17 | 7 |
| PRPF19 | 0.32347881 | 3.01E-17 | 7 |
| AC246817.2 | 0.32500067 | 3.35E-17 | 7 |
| CCT2 | 0.43309475 | 3.47E-17 | 7 |
| ATP5PF | 0.43297925 | 4.40E-17 | 7 |
| MRPL27 | 0.30295266 | 5.24E-17 | 7 |
| SPOCK2 | -0.5492686 | 5.43E-17 | 7 |
| SMARCB1 | 0.39109487 | 6.02E-17 | 7 |
| SNRPD2 | 0.40415588 | 6.12E-17 | 7 |
| PLP2 | 0.40994019 | 6.45E-17 | 7 |
| PGD | 0.28642187 | 7.23E-17 | 7 |
| PRKDC | 0.5255743 | 7.68E-17 | 7 |
| NDUFAB1 | 0.39337779 | 8.15E-17 | 7 |
| DPYSL2 | 0.40540005 | 8.17E-17 | 7 |
| RHEB | 0.43436486 | 1.01E-16 | 7 |
| HNRNPF | 0.43727854 | 1.18E-16 | 7 |
| DCTN3 | 0.48405224 | 1.20E-16 | 7 |
| MRPL37 | 0.35122887 | 1.35E-16 | 7 |
| CBX3 | 0.39537335 | 1.66E-16 | 7 |
| MMS22L | 0.43075404 | 1.68E-16 | 7 |
| ATF4 | 0.43983951 | 1.72E-16 | 7 |
| TRIM59 | 0.42116597 | 1.73E-16 | 7 |
| FCER1G | -0.485417 | 1.79E-16 | 7 |
| HINT1 | 0.38942541 | 1.81E-16 | 7 |
| NUTF2 | 0.34515352 | 1.83E-16 | 7 |
| SLC25A11 | 0.3623245 | 1.91E-16 | 7 |
| CD59 | 0.28735615 | 2.03E-16 | 7 |
| NUP50 | 0.38995087 | 2.35E-16 | 7 |
| PPP4C | 0.38760341 | 2.47E-16 | 7 |
| ID3 | -0.7186435 | 2.55E-16 | 7 |
| AL365440.2 | 0.35916076 | 2.80E-16 | 7 |
| KPNB1 | 0.43106272 | 3.24E-16 | 7 |
| YIF1B | 0.25704574 | 3.27E-16 | 7 |
| TKT | 0.41911798 | 3.38E-16 | 7 |
| NUP188 | 0.29208933 | 3.66E-16 | 7 |
| SLC25A3 | 0.44305571 | 3.70E-16 | 7 |
| LSP1 | -0.5373743 | 4.17E-16 | 7 |
| TCEAL3 | 0.29596177 | 4.44E-16 | 7 |
| PIN1 | 0.39504818 | 4.73E-16 | 7 |
| SEM1 | 0.40735136 | 4.75E-16 | 7 |
| SNX5 | 0.34063598 | 4.80E-16 | 7 |
| RALGDS | -0.4057353 | 5.17E-16 | 7 |
| DNAJC9 | 0.41309284 | 5.32E-16 | 7 |

| | | | |
|---|---|---|---|
| EBNA1BP2 | 0.32429973 | 5.41E-16 | 7 |
| EWSR1 | 0.37406021 | 5.61E-16 | 7 |
| EIF3E | -0.4575227 | 5.71E-16 | 7 |
| ZNRD2 | 0.25069296 | 6.57E-16 | 7 |
| LINC-PINT | -0.4494757 | 6.79E-16 | 7 |
| DYNLL1 | 0.44634244 | 7.31E-16 | 7 |
| TRIM24 | 0.38094691 | 8.50E-16 | 7 |
| CORO1C | 0.27362647 | 8.99E-16 | 7 |
| LAPTM5 | -0.4882551 | 9.61E-16 | 7 |
| FAM49B | -0.4400619 | 9.82E-16 | 7 |
| FADS1 | 0.28001031 | 1.07E-15 | 7 |
| MALAT1 | -0.3688468 | 1.53E-15 | 7 |
| CCR9 | -0.6370065 | 1.54E-15 | 7 |
| SF3B2 | 0.39487214 | 1.63E-15 | 7 |
| RTN3 | 0.33635431 | 1.80E-15 | 7 |
| DHX9 | 0.39264538 | 1.86E-15 | 7 |
| SLC7A1 | 0.26390129 | 1.97E-15 | 7 |
| DBI | 0.40690531 | 2.04E-15 | 7 |
| MRPL12 | 0.2774255 | 2.28E-15 | 7 |
| ALOX5AP | -0.4648567 | 2.31E-15 | 7 |
| TRIM22 | -0.4658319 | 2.35E-15 | 7 |
| FARS2 | -0.5691508 | 2.55E-15 | 7 |
| CHML | 0.36667487 | 2.60E-15 | 7 |
| TNRC6C-AS1 | -0.4551935 | 2.62E-15 | 7 |
| RAB37 | -0.4915378 | 2.95E-15 | 7 |
| CYTH4 | -0.394781 | 2.97E-15 | 7 |
| POLR3K | 0.3359941 | 3.01E-15 | 7 |
| ING2 | 0.30200879 | 3.07E-15 | 7 |
| CD226 | -0.5635985 | 3.45E-15 | 7 |
| CLN6 | 0.32362916 | 3.82E-15 | 7 |
| ANKRD36C | 0.55775668 | 3.86E-15 | 7 |
| AEBP1 | 0.30190033 | 4.45E-15 | 7 |
| HNRNPUL2 | 0.4110526 | 4.66E-15 | 7 |
| SESN3 | -0.4931435 | 5.17E-15 | 7 |
| HNRNPC | 0.33564481 | 6.21E-15 | 7 |
| MACROD2 | -0.7111205 | 6.23E-15 | 7 |
| SSBP2 | 0.41618725 | 6.37E-15 | 7 |
| TXNDC17 | 0.36308661 | 6.54E-15 | 7 |
| UBL7-AS1 | 0.25208342 | 6.88E-15 | 7 |
| SNRPB2 | 0.36749954 | 7.66E-15 | 7 |
| SRPK1 | 0.35205432 | 8.62E-15 | 7 |
| TRBC2 | -0.3532563 | 9.23E-15 | 7 |
| COPE | 0.39306848 | 9.62E-15 | 7 |

| | | | |
|---|---|---|---|
| PHTF2 | 0.32160844 | 9.80E-15 | 7 |
| FGFR1OP | 0.29168678 | 1.05E-14 | 7 |
| HNRNPUL1 | 0.37521675 | 1.50E-14 | 7 |
| JPX | -0.4742215 | 1.53E-14 | 7 |
| TMEM161B-AS1 | -0.606095 | 1.70E-14 | 7 |
| VDAC2 | 0.36967998 | 1.78E-14 | 7 |
| BAZ1B | 0.44947291 | 2.01E-14 | 7 |
| PSMD1 | 0.34730057 | 2.01E-14 | 7 |
| IPO9 | 0.29602769 | 2.06E-14 | 7 |
| EIF2AK2 | 0.33776247 | 2.06E-14 | 7 |
| AKNA | -0.4657551 | 2.09E-14 | 7 |
| KMT2E | -0.422064 | 2.11E-14 | 7 |
| DGKA | -0.4905487 | 2.15E-14 | 7 |
| PSMD2 | 0.35057234 | 2.31E-14 | 7 |
| CCDC69 | -0.5421758 | 2.71E-14 | 7 |
| NCL | 0.44886598 | 2.94E-14 | 7 |
| PCSK7 | -0.5001892 | 3.50E-14 | 7 |
| RPL5 | -0.3282818 | 3.63E-14 | 7 |
| SYNE1 | -0.5278205 | 3.90E-14 | 7 |
| LY9 | -0.3622249 | 3.91E-14 | 7 |
| CD320 | 0.27277104 | 3.97E-14 | 7 |
| NDUFV2 | 0.37661093 | 4.09E-14 | 7 |
| KDM7A | -0.3798281 | 4.88E-14 | 7 |
| ZCCHC8 | 0.2747469 | 5.21E-14 | 7 |
| CD84 | -0.5134778 | 6.05E-14 | 7 |
| POLR2I | 0.35125484 | 6.21E-14 | 7 |
| UQCR10 | 0.35078154 | 6.33E-14 | 7 |
| NDUFS5 | 0.41421403 | 6.44E-14 | 7 |
| DLEU7 | 0.27621642 | 6.85E-14 | 7 |
| HAUS6 | 0.30545709 | 7.58E-14 | 7 |
| CAP1 | -0.4311261 | 7.65E-14 | 7 |
| BIN1 | -0.4232103 | 7.69E-14 | 7 |
| PRRC2A | 0.36750309 | 1.01E-13 | 7 |
| RHOA | 0.31415778 | 1.03E-13 | 7 |
| NUDCD2 | 0.35117473 | 1.04E-13 | 7 |
| LUC7L2 | 0.41207641 | 1.07E-13 | 7 |
| ZBTB20 | -0.5096925 | 1.07E-13 | 7 |
| PAK1 | 0.25474816 | 1.12E-13 | 7 |
| SUMO3 | 0.38214702 | 1.29E-13 | 7 |
| TRIOBP | 0.26571998 | 1.34E-13 | 7 |
| SLC16A1-AS1 | 0.30571602 | 1.42E-13 | 7 |
| C1orf35 | 0.33243188 | 1.45E-13 | 7 |
| MKNK2 | -0.4002131 | 1.51E-13 | 7 |

| | | | |
|---|---|---|---|
| PPP1R14B | 0.47338006 | 1.77E-13 | 7 |
| CCT4 | 0.3525194 | 1.99E-13 | 7 |
| NDUFC2 | 0.36197168 | 2.01E-13 | 7 |
| CD3D | -0.3569139 | 2.07E-13 | 7 |
| HLTF | 0.34961153 | 2.16E-13 | 7 |
| G2E3 | 0.40084562 | 2.24E-13 | 7 |
| SLC35D2 | -0.3247507 | 2.37E-13 | 7 |
| FXYD5 | -0.4651319 | 2.78E-13 | 7 |
| COX5B | 0.36349675 | 3.37E-13 | 7 |
| KLHL24 | -0.4311734 | 3.40E-13 | 7 |
| PMVK | 0.26901656 | 3.62E-13 | 7 |
| FDPS | 0.40660744 | 3.68E-13 | 7 |
| CCAR1 | 0.3750059 | 3.87E-13 | 7 |
| PRIM1 | 0.27104693 | 4.08E-13 | 7 |
| TTC14 | -0.4452123 | 4.16E-13 | 7 |
| AKAP13 | -0.4456445 | 4.17E-13 | 7 |
| ATP5PD | 0.37837387 | 4.27E-13 | 7 |
| CHRAC1 | 0.34493889 | 4.39E-13 | 7 |
| SNRPA | 0.32774723 | 4.53E-13 | 7 |
| MANF | 0.3279072 | 4.62E-13 | 7 |
| FAT1 | 0.32826561 | 4.63E-13 | 7 |
| UBE2I | 0.3475506 | 4.63E-13 | 7 |
| CCDC141 | -0.4424494 | 4.86E-13 | 7 |
| ATP5MPL | 0.3707202 | 6.09E-13 | 7 |
| KIF5B | 0.44727556 | 6.43E-13 | 7 |
| RANGAP1 | 0.26237835 | 6.56E-13 | 7 |
| STOML2 | 0.29409375 | 7.01E-13 | 7 |
| HINT2 | 0.37580441 | 7.31E-13 | 7 |
| RAPGEF6 | -0.4470954 | 7.32E-13 | 7 |
| ITGB3BP | 0.29168101 | 7.39E-13 | 7 |
| GZMM | -0.4585184 | 7.58E-13 | 7 |
| TNRC6B | -0.4758733 | 7.85E-13 | 7 |
| GIMAP1 | -0.3813089 | 8.35E-13 | 7 |
| PRMT7 | 0.49974076 | 8.37E-13 | 7 |
| NDUFA4 | 0.36946426 | 8.37E-13 | 7 |
| DDX11 | 0.34158345 | 8.45E-13 | 7 |
| MOB3A | -0.3944796 | 8.48E-13 | 7 |
| THRAP3 | 0.3488801 | 9.83E-13 | 7 |
| ELOVL4 | 0.40526896 | 1.10E-12 | 7 |
| CUTA | -0.4225578 | 1.11E-12 | 7 |
| TBC1D10C | -0.4186199 | 1.16E-12 | 7 |
| HAUS3 | 0.25214818 | 1.21E-12 | 7 |
| SENP7 | -0.4591186 | 1.28E-12 | 7 |

| | | | |
|---|---|---|---|
| MRPL11 | 0.36013498 | 1.34E-12 | 7 |
| YPEL5 | -0.4030296 | 1.54E-12 | 7 |
| POMP | 0.3802954 | 1.66E-12 | 7 |
| SF3B5 | 0.36212473 | 1.72E-12 | 7 |
| TSPO | -0.4880146 | 1.73E-12 | 7 |
| TIMM10 | 0.35356401 | 1.91E-12 | 7 |
| NME3 | -0.3224919 | 1.94E-12 | 7 |
| HSBP1 | 0.34617564 | 1.99E-12 | 7 |
| HDAC2 | 0.37196951 | 2.10E-12 | 7 |
| DYRK2 | -0.4361018 | 2.12E-12 | 7 |
| FH | 0.25584661 | 2.13E-12 | 7 |
| PPP1CA | 0.37388436 | 2.26E-12 | 7 |
| ABHD17A | -0.4491047 | 2.34E-12 | 7 |
| ARL6IP6 | 0.2631978 | 2.38E-12 | 7 |
| EIF4A1 | 0.41247014 | 2.70E-12 | 7 |
| SCCPDH | 0.29977544 | 2.73E-12 | 7 |
| NFATC3 | -0.4682263 | 2.87E-12 | 7 |
| TPGS2 | 0.31678479 | 2.87E-12 | 7 |
| PIK3R1 | -0.4292691 | 3.00E-12 | 7 |
| HNRNPL | 0.35240669 | 3.23E-12 | 7 |
| QPRT | 0.26034811 | 3.26E-12 | 7 |
| MYH10 | 0.36739559 | 3.45E-12 | 7 |
| TTC3 | -0.4294844 | 3.59E-12 | 7 |
| WBP11 | 0.34325961 | 4.76E-12 | 7 |
| CCT8 | 0.34571562 | 4.97E-12 | 7 |
| TCERG1 | 0.36313478 | 5.16E-12 | 7 |
| PSMA3 | 0.32277208 | 5.41E-12 | 7 |
| S1PR3 | 0.33227004 | 6.22E-12 | 7 |
| NDUFS7 | 0.33614745 | 6.57E-12 | 7 |
| RCOR1 | 0.25831429 | 6.63E-12 | 7 |
| PDE7A | -0.5285941 | 8.15E-12 | 7 |
| RHOF | -0.3695141 | 8.57E-12 | 7 |
| GATAD1 | -0.5100851 | 8.91E-12 | 7 |
| MLH1 | 0.25431407 | 9.50E-12 | 7 |
| MLLT6 | -0.4394984 | 9.51E-12 | 7 |
| GLUL | 0.34495001 | 9.61E-12 | 7 |
| UQCRFS1 | 0.28423252 | 1.01E-11 | 7 |
| CHD2 | -0.4617557 | 1.07E-11 | 7 |
| BIN2 | -0.469821 | 1.09E-11 | 7 |
| TGFB1 | -0.4340428 | 1.11E-11 | 7 |
| NDUFA12 | 0.33977586 | 1.12E-11 | 7 |
| CCDC167 | 0.35754844 | 1.14E-11 | 7 |
| NDE1 | 0.33325782 | 1.15E-11 | 7 |

| | | | |
|---|---|---|---|
| PGAM1 | 0.38011197 | 1.16E-11 | 7 |
| DCXR | 0.3382556 | 1.18E-11 | 7 |
| SUB1 | 0.38960593 | 1.21E-11 | 7 |
| TBCA | 0.32293522 | 1.26E-11 | 7 |
| SIPA1 | -0.4133909 | 1.26E-11 | 7 |
| KMT5A | 0.34016839 | 1.28E-11 | 7 |
| TTC7A | 0.3028649 | 1.29E-11 | 7 |
| PABPC4 | 0.26999837 | 1.38E-11 | 7 |
| EEF2 | -0.3526503 | 1.57E-11 | 7 |
| DPM2 | 0.26241926 | 1.70E-11 | 7 |
| BRD7 | 0.38302986 | 1.71E-11 | 7 |
| MT-ND6 | 0.43670853 | 1.75E-11 | 7 |
| NDUFB6 | 0.36372553 | 1.87E-11 | 7 |
| GTF2A1 | 0.29218446 | 2.04E-11 | 7 |
| TRMT112 | 0.3446494 | 2.11E-11 | 7 |
| METAP2 | 0.32294996 | 2.24E-11 | 7 |
| NCAPG2 | 0.31071441 | 2.27E-11 | 7 |
| PRDX1 | 0.39659536 | 2.34E-11 | 7 |
| EIF3A | 0.34722423 | 2.34E-11 | 7 |
| MDH2 | 0.32532181 | 2.38E-11 | 7 |
| HPS4 | 0.31814251 | 2.46E-11 | 7 |
| TCEAL9 | 0.28500413 | 2.71E-11 | 7 |
| NDUFS3 | 0.29350938 | 3.24E-11 | 7 |
| BCL9L | -0.3597316 | 3.28E-11 | 7 |
| KDM6B | -0.393718 | 3.60E-11 | 7 |
| PSMA2 | 0.32477394 | 3.84E-11 | 7 |
| NDUFA13 | 0.32637936 | 4.21E-11 | 7 |
| NDUFA11 | 0.32999218 | 4.27E-11 | 7 |
| SRRT | 0.32661212 | 4.27E-11 | 7 |
| TPST2 | -0.4180408 | 4.37E-11 | 7 |
| GSTK1 | -0.4342489 | 4.57E-11 | 7 |
| CDC42SE1 | -0.4373069 | 4.92E-11 | 7 |
| UQCC2 | 0.32953114 | 5.00E-11 | 7 |
| KLF13 | -0.4290772 | 5.15E-11 | 7 |
| RPS21 | -0.2617429 | 5.29E-11 | 7 |
| GCNT4 | -0.3622657 | 5.34E-11 | 7 |
| POLR2D | 0.25737416 | 5.35E-11 | 7 |
| NDUFB8 | 0.34432194 | 5.49E-11 | 7 |
| RCC2 | 0.33400661 | 6.21E-11 | 7 |
| SLIT1 | 0.30657184 | 6.61E-11 | 7 |
| POLE3 | 0.36301405 | 1.04E-10 | 7 |
| CREBRF | -0.4147902 | 1.06E-10 | 7 |
| YPEL2 | -0.3242759 | 1.09E-10 | 7 |

| | | | |
|---|---|---|---|
| MEA1 | 0.32700003 | 1.12E-10 | 7 |
| EBP | 0.25025966 | 1.14E-10 | 7 |
| PLEC | -0.3898389 | 1.15E-10 | 7 |
| UBE2M | 0.31486891 | 1.16E-10 | 7 |
| MYL6 | -0.3167832 | 1.17E-10 | 7 |
| RGS3 | -0.4896952 | 1.17E-10 | 7 |
| CLTA | 0.33758002 | 1.23E-10 | 7 |
| CYCS | 0.28650225 | 1.25E-10 | 7 |
| LAT2 | -0.4685207 | 1.32E-10 | 7 |
| ZFP36 | -0.3936086 | 1.35E-10 | 7 |
| BCL6 | -0.4084674 | 1.36E-10 | 7 |
| DCAF15 | 0.25545815 | 1.43E-10 | 7 |
| GNG5 | 0.36231276 | 1.44E-10 | 7 |
| GDI2 | 0.32781045 | 1.54E-10 | 7 |
| SYNCRIP | 0.35429262 | 1.54E-10 | 7 |
| TSPAN32 | -0.3030607 | 1.58E-10 | 7 |
| UBB | 0.40881547 | 1.59E-10 | 7 |
| RPL27A | -0.3095142 | 1.95E-10 | 7 |
| SVIP | 0.31214506 | 2.00E-10 | 7 |
| GYPC | -0.3878842 | 2.04E-10 | 7 |
| FAF1 | 0.30516595 | 2.11E-10 | 7 |
| OAZ1 | 0.31223246 | 2.21E-10 | 7 |
| JAML | -0.3421603 | 2.24E-10 | 7 |
| CIRBP | -0.397229 | 2.36E-10 | 7 |
| CXCR3 | -0.3015509 | 2.39E-10 | 7 |
| MYL6B | 0.30619852 | 2.49E-10 | 7 |
| CLIC3 | -0.5096334 | 2.69E-10 | 7 |
| PGK1 | 0.35914428 | 2.70E-10 | 7 |
| KNTC1 | 0.26509961 | 2.75E-10 | 7 |
| USP49 | 0.26944959 | 3.15E-10 | 7 |
| ARPP19 | 0.32005413 | 3.42E-10 | 7 |
| ATOX1 | 0.27210475 | 3.53E-10 | 7 |
| IL2RG | -0.385235 | 3.55E-10 | 7 |
| STK10 | -0.4066663 | 3.56E-10 | 7 |
| PDIA6 | 0.32865759 | 3.69E-10 | 7 |
| STAT3 | -0.3385489 | 3.76E-10 | 7 |
| NAA50 | 0.30172091 | 3.89E-10 | 7 |
| UBE2N | 0.30737824 | 4.15E-10 | 7 |
| STAG2 | 0.36250831 | 4.36E-10 | 7 |
| PHF6 | 0.31677191 | 4.36E-10 | 7 |
| SLFN13 | 0.43078082 | 4.41E-10 | 7 |
| ZYX | -0.4267254 | 4.45E-10 | 7 |
| AP2S1 | 0.3242029 | 4.59E-10 | 7 |

| | | | |
|---|---|---|---|
| SNHG5 | -0.3731422 | 4.64E-10 | 7 |
| MRPL23 | 0.28769293 | 4.64E-10 | 7 |
| FUBP1 | 0.30452115 | 4.97E-10 | 7 |
| CCND3 | 0.36892047 | 5.19E-10 | 7 |
| SUGP2 | 0.30529856 | 5.33E-10 | 7 |
| GPAA1 | 0.2692194 | 5.48E-10 | 7 |
| P2RX5 | -0.3986364 | 5.58E-10 | 7 |
| SATB1-AS1 | -0.3286758 | 5.69E-10 | 7 |
| NOSIP | -0.7149931 | 5.82E-10 | 7 |
| NDUFB7 | 0.33533144 | 6.11E-10 | 7 |
| PPM1M | -0.3329606 | 6.59E-10 | 7 |
| TMC8 | -0.4351044 | 6.73E-10 | 7 |
| PSMA1 | 0.29408284 | 6.75E-10 | 7 |
| EEF1B2 | -0.3305623 | 7.06E-10 | 7 |
| FUS | 0.328771 | 7.23E-10 | 7 |
| AHNAK | -0.7908522 | 7.91E-10 | 7 |
| TIMM8B | 0.31886219 | 8.35E-10 | 7 |
| PTPRE | -0.3675056 | 8.50E-10 | 7 |
| RPL8 | -0.26068 | 8.62E-10 | 7 |
| SLC38A1 | -0.4122187 | 9.91E-10 | 7 |
| KLF2 | -0.6929989 | 1.06E-09 | 7 |
| FKBP5 | 0.34694665 | 1.07E-09 | 7 |
| RNF125 | -0.4351475 | 1.09E-09 | 7 |
| SOX4 | -0.4496279 | 1.12E-09 | 7 |
| TMX1 | 0.30256 | 1.22E-09 | 7 |
| MRPL52 | 0.30481918 | 1.22E-09 | 7 |
| SEPTIN9 | -0.3560644 | 1.22E-09 | 7 |
| BCL11B | -0.4583975 | 1.23E-09 | 7 |
| RUNX1 | 0.3396809 | 1.25E-09 | 7 |
| CAMK1D | 0.3522073 | 1.26E-09 | 7 |
| P2RY11 | -0.3211006 | 1.27E-09 | 7 |
| LTBP3 | -0.3506499 | 1.30E-09 | 7 |
| PDAP1 | 0.29750827 | 1.39E-09 | 7 |
| LINC00954 | -0.379228 | 1.42E-09 | 7 |
| SEMA4D | -0.4615055 | 1.46E-09 | 7 |
| ARRDC5 | -0.2638004 | 1.49E-09 | 7 |
| HAT1 | 0.30876716 | 1.66E-09 | 7 |
| TRIM73 | -0.3748908 | 1.67E-09 | 7 |
| LSM2 | 0.28901347 | 1.74E-09 | 7 |
| MCUB | 0.28492999 | 1.83E-09 | 7 |
| POLR2J | 0.30997589 | 1.84E-09 | 7 |
| PLCL2 | -0.3459314 | 1.96E-09 | 7 |
| MBD5 | -0.3943142 | 2.10E-09 | 7 |

| | | | |
|---|---|---|---|
| SYTL1 | -0.4093965 | 2.28E-09 | 7 |
| RIF1 | 0.32698132 | 2.91E-09 | 7 |
| ITM2A | -0.6271382 | 3.06E-09 | 7 |
| OXNAD1 | -0.4423203 | 3.13E-09 | 7 |
| JADE1 | 0.33769769 | 3.30E-09 | 7 |
| MZT2A | 0.29897574 | 3.34E-09 | 7 |
| NHP2 | 0.32807359 | 3.44E-09 | 7 |
| CAMK4 | -0.4116924 | 3.44E-09 | 7 |
| AKAP9 | -0.4045873 | 3.99E-09 | 7 |
| ZMAT3 | -0.4124045 | 4.05E-09 | 7 |
| IPCEF1 | -0.367664 | 4.14E-09 | 7 |
| ELP5 | 0.27158343 | 4.17E-09 | 7 |
| ATP5PB | 0.30821865 | 4.36E-09 | 7 |
| ACAP2 | -0.4019067 | 4.44E-09 | 7 |
| MCM3 | 0.43088574 | 4.46E-09 | 7 |
| CDC42 | -0.3484521 | 4.60E-09 | 7 |
| CCSER2 | -0.3885405 | 4.72E-09 | 7 |
| EIF5 | 0.33528108 | 4.87E-09 | 7 |
| CEP192 | 0.3022074 | 4.99E-09 | 7 |
| SF3B6 | 0.29887512 | 5.13E-09 | 7 |
| HDGFL3 | 0.28454 | 5.52E-09 | 7 |
| GPSM3 | -0.3783121 | 5.64E-09 | 7 |
| PYHIN1 | -0.396275 | 5.96E-09 | 7 |
| AC114760.2 | -0.4010456 | 6.05E-09 | 7 |
| OLA1 | 0.28299753 | 6.25E-09 | 7 |
| ZNF708 | -0.3766098 | 6.30E-09 | 7 |
| CYLD | -0.3953203 | 6.58E-09 | 7 |
| NREP | 0.31566853 | 6.72E-09 | 7 |
| ITK | -0.3966486 | 7.49E-09 | 7 |
| YWHAH | 0.26660573 | 8.04E-09 | 7 |
| RBM39 | -0.2847793 | 8.38E-09 | 7 |
| MCRIP1 | 0.33549664 | 8.81E-09 | 7 |
| RNF138 | 0.2935574 | 8.83E-09 | 7 |
| FOXO1 | -0.2900296 | 8.95E-09 | 7 |
| PHYKPL | -0.364502 | 9.01E-09 | 7 |
| G3BP1 | 0.31379854 | 9.04E-09 | 7 |
| DCAF7 | 0.33861271 | 9.46E-09 | 7 |
| PSMD7 | 0.31791651 | 9.52E-09 | 7 |
| HMGN3 | 0.35948639 | 9.68E-09 | 7 |
| SSNA1 | 0.30581695 | 1.08E-08 | 7 |
| EEF1D | -0.2930129 | 1.09E-08 | 7 |
| GBP2 | -0.3628589 | 1.10E-08 | 7 |
| AC022706.1 | -0.2778958 | 1.25E-08 | 7 |

| | | | |
|---|---|---|---|
| ANKRD12 | -0.4566875 | 1.25E-08 | 7 |
| RTRAF | 0.31413134 | 1.30E-08 | 7 |
| PSMA5 | 0.31119371 | 1.39E-08 | 7 |
| TSPAN14 | -0.3410989 | 1.42E-08 | 7 |
| ZNRF1 | 0.28521357 | 1.55E-08 | 7 |
| COPS3 | 0.30389482 | 1.59E-08 | 7 |
| TSC22D3 | -0.44664 | 1.62E-08 | 7 |
| IQGAP2 | 0.36812159 | 1.62E-08 | 7 |
| ARL6IP5 | -0.427561 | 1.69E-08 | 7 |
| PSMB9 | 0.2679975 | 1.70E-08 | 7 |
| EIF2S2 | 0.29744538 | 1.71E-08 | 7 |
| SERINC1 | -0.3926977 | 1.81E-08 | 7 |
| TRIR | 0.28793225 | 1.82E-08 | 7 |
| FYN | -0.4090728 | 1.87E-08 | 7 |
| RASSF1 | -0.3742753 | 2.03E-08 | 7 |
| MT-ATP6 | -0.261866 | 2.18E-08 | 7 |
| CD6 | -0.3094702 | 2.22E-08 | 7 |
| PRPF8 | 0.31278876 | 2.28E-08 | 7 |
| NCKAP1L | -0.3828909 | 2.31E-08 | 7 |
| DCAF5 | -0.3410964 | 2.32E-08 | 7 |
| PPA1 | 0.34577623 | 2.55E-08 | 7 |
| SRP14 | 0.26001595 | 2.57E-08 | 7 |
| CD53 | -0.3575978 | 2.61E-08 | 7 |
| DHPS | 0.27937839 | 2.62E-08 | 7 |
| C17orf49 | 0.29055235 | 2.65E-08 | 7 |
| GSTO1 | 0.31138646 | 2.68E-08 | 7 |
| GTF3C6 | 0.28461947 | 2.81E-08 | 7 |
| SIGIRR | -0.368898 | 2.81E-08 | 7 |
| HNRNPU | 0.28403988 | 3.49E-08 | 7 |
| TSPOAP1-AS1 | -0.3448524 | 3.70E-08 | 7 |
| PRDX6 | 0.27061835 | 3.73E-08 | 7 |
| SERF2 | -0.2671411 | 3.75E-08 | 7 |
| ZFP91 | 0.29222732 | 3.83E-08 | 7 |
| ZNF655 | -0.3928969 | 3.97E-08 | 7 |
| SAP18 | 0.28668654 | 3.98E-08 | 7 |
| ABCB1 | -0.2953108 | 4.05E-08 | 7 |
| MRPL16 | 0.2587689 | 4.07E-08 | 7 |
| MRPS6 | 0.29253654 | 4.09E-08 | 7 |
| ATP5PO | 0.31293343 | 4.17E-08 | 7 |
| ERICH1 | -0.3966668 | 4.59E-08 | 7 |
| DNAJC21 | 0.31997982 | 4.95E-08 | 7 |
| DNAJC8 | 0.29477121 | 5.09E-08 | 7 |
| LINC00342 | 0.52475741 | 5.18E-08 | 7 |

| | | | |
|---|---|---|---|
| HNRNPH1 | 0.33668703 | 5.27E-08 | 7 |
| NFKBIA | 0.26846435 | 5.57E-08 | 7 |
| SNRNP200 | 0.29675156 | 5.78E-08 | 7 |
| PSMC5 | 0.27307839 | 5.98E-08 | 7 |
| RBM38 | -0.3166454 | 6.22E-08 | 7 |
| IGFBP5 | -0.7397264 | 6.26E-08 | 7 |
| RBM3 | 0.29442021 | 6.70E-08 | 7 |
| ARID4B | -0.3518597 | 6.72E-08 | 7 |
| LY6H | 0.32179177 | 6.87E-08 | 7 |
| YBX3 | 0.29751725 | 6.94E-08 | 7 |
| RPS12 | -0.3047893 | 6.95E-08 | 7 |
| FUNDC2 | 0.26598158 | 7.16E-08 | 7 |
| CHAMP1 | 0.26495512 | 7.33E-08 | 7 |
| TFRC | 0.26288989 | 7.39E-08 | 7 |
| PPP1CC | 0.30919135 | 7.86E-08 | 7 |
| IL6ST | -0.3352045 | 7.93E-08 | 7 |
| SARNP | 0.28392274 | 8.07E-08 | 7 |
| IKZF3 | -0.4269723 | 8.55E-08 | 7 |
| UBE2E3 | 0.29735945 | 8.80E-08 | 7 |
| AURKAIP1 | 0.29007078 | 9.00E-08 | 7 |
| CD96 | -0.3755459 | 9.10E-08 | 7 |
| MAPRE1 | 0.29317032 | 9.17E-08 | 7 |
| ERV3-1 | -0.361587 | 9.44E-08 | 7 |
| DBN1 | -0.2951614 | 9.95E-08 | 7 |
| MCUR1 | 0.26211237 | 1.00E-07 | 7 |
| ELP6 | 0.25167815 | 1.06E-07 | 7 |
| C1QBP | 0.46113269 | 1.11E-07 | 7 |
| PILRB | -0.3861224 | 1.24E-07 | 7 |
| SP100 | -0.4508396 | 1.30E-07 | 7 |
| USP14 | 0.27659366 | 1.32E-07 | 7 |
| MICOS10 | 0.27808279 | 1.37E-07 | 7 |
| UCP2 | -0.4044861 | 1.38E-07 | 7 |
| NDUFB10 | 0.25583511 | 1.39E-07 | 7 |
| SSB | 0.27547742 | 1.44E-07 | 7 |
| RNF44 | -0.329711 | 1.45E-07 | 7 |
| DDX39B | 0.32154982 | 1.51E-07 | 7 |
| DNAJA1 | 0.35393974 | 1.59E-07 | 7 |
| PBXIP1 | -0.3408579 | 1.62E-07 | 7 |
| PPIG | 0.28511113 | 1.62E-07 | 7 |
| TMEM219 | -0.3303873 | 1.63E-07 | 7 |
| RPS6KA3 | -0.4518335 | 1.78E-07 | 7 |
| STAT6 | -0.2891453 | 1.80E-07 | 7 |
| TMBIM4 | -0.3599725 | 1.85E-07 | 7 |

| | | | |
|---|---|---|---|
| UBE2A | 0.2996804 | 1.88E-07 | 7 |
| FBXL20 | -0.2737548 | 1.94E-07 | 7 |
| RSRC1 | 0.29280838 | 2.06E-07 | 7 |
| AHSA1 | 0.27506335 | 2.19E-07 | 7 |
| POLR2G | 0.28455382 | 2.36E-07 | 7 |
| PHB2 | 0.27796534 | 2.39E-07 | 7 |
| TRIM56 | -0.38153 | 2.44E-07 | 7 |
| PSMA3-AS1 | -0.359464 | 2.61E-07 | 7 |
| MTRNR2L12 | -0.2646285 | 2.70E-07 | 7 |
| SNHG7 | 0.28680981 | 2.74E-07 | 7 |
| GIMAP6 | -0.3021628 | 2.85E-07 | 7 |
| AL365361.1 | -0.4384489 | 2.99E-07 | 7 |
| PSMD11 | 0.27634204 | 2.99E-07 | 7 |
| ATP5MD | 0.31067864 | 3.15E-07 | 7 |
| TSEN54 | -0.2957499 | 3.27E-07 | 7 |
| CNOT6L | -0.3887784 | 3.28E-07 | 7 |
| ZNF276 | -0.3198346 | 3.35E-07 | 7 |
| ARL4C | -0.386155 | 3.42E-07 | 7 |
| PTPN22 | -0.4199687 | 3.51E-07 | 7 |
| EIF3F | -0.2884465 | 3.66E-07 | 7 |
| TUFM | 0.3110964 | 3.88E-07 | 7 |
| IGFBP2 | 0.32119164 | 4.10E-07 | 7 |
| ATP5F1A | 0.28200441 | 4.18E-07 | 7 |
| IGFLR1 | -0.2929671 | 4.25E-07 | 7 |
| PPP1R18 | -0.3242633 | 4.40E-07 | 7 |
| EIF4E | 0.26592354 | 4.62E-07 | 7 |
| CCT3 | 0.27269788 | 4.63E-07 | 7 |
| DGKZ | -0.3185567 | 4.82E-07 | 7 |
| RFC2 | 0.30511246 | 5.16E-07 | 7 |
| MYO7B | 0.28201536 | 5.28E-07 | 7 |
| TUBA1A | 0.37595172 | 5.30E-07 | 7 |
| GMPS | 0.27988676 | 5.54E-07 | 7 |
| TPI1 | 0.41733005 | 5.58E-07 | 7 |
| AHCTF1 | 0.31741431 | 5.76E-07 | 7 |
| SP110 | -0.3694965 | 5.92E-07 | 7 |
| AZIN1 | 0.26205833 | 5.98E-07 | 7 |
| INPP5D | -0.3447314 | 6.20E-07 | 7 |
| FGR | -0.3204493 | 6.22E-07 | 7 |
| NLRC3 | -0.3335585 | 6.29E-07 | 7 |
| C16orf54 | -0.3144477 | 6.37E-07 | 7 |
| AKR1B1 | 0.30065111 | 6.50E-07 | 7 |
| SYPL1 | 0.25787453 | 6.69E-07 | 7 |
| FBRS | -0.3030122 | 6.76E-07 | 7 |

| | | | |
|---|---|---|---|
| NCR3 | -0.3765948 | 7.53E-07 | 7 |
| PSMB7 | 0.26139957 | 7.68E-07 | 7 |
| ZNF292 | -0.3334407 | 8.59E-07 | 7 |
| ESYT2 | -0.3889167 | 9.76E-07 | 7 |
| RGS14 | -0.289279 | 9.95E-07 | 7 |
| HECA | -0.3248723 | 1.03E-06 | 7 |
| ARHGDIA | 0.25807621 | 1.04E-06 | 7 |
| ATXN2L | 0.25560739 | 1.07E-06 | 7 |
| HSPH1 | 0.32327389 | 1.09E-06 | 7 |
| CEMIP2 | -0.3458311 | 1.12E-06 | 7 |
| NUP62 | 0.25981055 | 1.16E-06 | 7 |
| NUDT5 | 0.27454887 | 1.16E-06 | 7 |
| BEX2 | -0.3032636 | 1.17E-06 | 7 |
| PDCD5 | 0.30523382 | 1.19E-06 | 7 |
| CCDC107 | -0.312492 | 1.20E-06 | 7 |
| OFD1 | -0.3045577 | 1.23E-06 | 7 |
| RIPOR2 | -0.464665 | 1.26E-06 | 7 |
| AKR7A2 | 0.26248399 | 1.30E-06 | 7 |
| RSRC2 | 0.26353014 | 1.34E-06 | 7 |
| SNRNP70 | 0.25796294 | 1.37E-06 | 7 |
| PPM1G | 0.27106927 | 1.40E-06 | 7 |
| RNASET2 | -0.2692489 | 1.41E-06 | 7 |
| CD82 | 0.26735044 | 1.53E-06 | 7 |
| SLC25A24 | 0.26704047 | 1.53E-06 | 7 |
| SLC38A2 | 0.3151292 | 1.73E-06 | 7 |
| PRDX2 | 0.43818293 | 1.76E-06 | 7 |
| SSR4 | -0.3165272 | 1.94E-06 | 7 |
| PLEKHO1 | 0.27059457 | 2.07E-06 | 7 |
| TECPR1 | -0.2873101 | 2.10E-06 | 7 |
| SRSF9 | 0.27909872 | 2.30E-06 | 7 |
| RAD23A | 0.26971292 | 2.31E-06 | 7 |
| ZNF720 | -0.3625521 | 2.34E-06 | 7 |
| SUMO1 | 0.29031118 | 2.35E-06 | 7 |
| DIAPH1 | 0.28993021 | 2.36E-06 | 7 |
| TXK | -0.2621886 | 2.39E-06 | 7 |
| ANKRD44 | -0.3853654 | 2.74E-06 | 7 |
| CD3E | -0.2747167 | 2.75E-06 | 7 |
| SH3TC1 | 0.34159885 | 2.80E-06 | 7 |
| ATP6V0E1 | -0.3238124 | 3.06E-06 | 7 |
| NUP210 | 0.2988538 | 3.09E-06 | 7 |
| AIF1 | 0.30432073 | 3.11E-06 | 7 |
| AL359220.1 | -0.3949221 | 3.12E-06 | 7 |
| CFLAR | -0.3721102 | 3.13E-06 | 7 |

| | | | |
|---|---|---|---|
| EIF1AX | 0.28194615 | 3.15E-06 | 7 |
| DDT | 0.2518038 | 3.28E-06 | 7 |
| CTSS | -0.2802314 | 3.50E-06 | 7 |
| MDM4 | -0.3604723 | 3.62E-06 | 7 |
| MRPS15 | 0.25607707 | 3.72E-06 | 7 |
| GLIPR1 | -0.3373329 | 3.80E-06 | 7 |
| GTF2I | -0.2976573 | 3.93E-06 | 7 |
| ARHGAP30 | -0.3093204 | 3.99E-06 | 7 |
| POLD4 | -0.3265826 | 4.53E-06 | 7 |
| DDX6 | -0.3392008 | 4.69E-06 | 7 |
| KMT5B | -0.3134255 | 4.82E-06 | 7 |
| SELENOH | 0.27286099 | 5.02E-06 | 7 |
| ITPKB | -0.3755351 | 5.03E-06 | 7 |
| HERC1 | -0.3707255 | 5.50E-06 | 7 |
| ST3GAL1 | -0.3412159 | 5.56E-06 | 7 |
| TRAF5 | -0.3742884 | 5.61E-06 | 7 |
| TENM1 | -0.330087 | 6.13E-06 | 7 |
| ITGB1BP1 | 0.258023 | 6.14E-06 | 7 |
| DDIT4 | -0.3956057 | 6.38E-06 | 7 |
| HEBP2 | 0.26079192 | 6.54E-06 | 7 |
| SMARCA4 | 0.26198051 | 6.69E-06 | 7 |
| NDUFB1 | 0.28434731 | 7.01E-06 | 7 |
| STAG1 | 0.26623113 | 7.22E-06 | 7 |
| RAB8B | -0.3187183 | 7.37E-06 | 7 |
| TECR | -0.7010997 | 8.10E-06 | 7 |
| RAB29 | -0.2577601 | 8.78E-06 | 7 |
| DDAH2 | 0.38183705 | 9.03E-06 | 7 |
| LYL1 | 0.30247718 | 9.09E-06 | 7 |
| AQP3 | 0.33044707 | 1.02E-05 | 7 |
| TTC39B | -0.2851395 | 1.08E-05 | 7 |
| CALU | 0.25357575 | 1.17E-05 | 7 |
| S100A4 | -0.5035595 | 1.22E-05 | 7 |
| ZBTB1 | -0.3431593 | 1.27E-05 | 7 |
| SHISA2 | -0.3770796 | 1.28E-05 | 7 |
| SLC3A2 | 0.31393748 | 1.39E-05 | 7 |
| TPM4 | 0.31022348 | 1.40E-05 | 7 |
| MPHOSPH8 | -0.3295758 | 1.45E-05 | 7 |
| AC108066.2 | -0.25081 | 1.46E-05 | 7 |
| PABPN1 | 0.27066349 | 1.49E-05 | 7 |
| CDCA7 | 0.32449302 | 1.57E-05 | 7 |
| NCOA3 | -0.2625245 | 1.61E-05 | 7 |
| NLRC5 | -0.2587657 | 1.63E-05 | 7 |
| GATA3 | -0.3557922 | 1.79E-05 | 7 |

| | | | |
|---|---|---|---|
| MT-ND2 | -0.3121217 | 1.97E-05 | 7 |
| ATP13A3 | 0.28984308 | 1.98E-05 | 7 |
| BIRC6 | -0.3444891 | 2.06E-05 | 7 |
| CALCOCO1 | -0.2540434 | 2.11E-05 | 7 |
| SCN3A | 0.41633937 | 2.16E-05 | 7 |
| SPEN | 0.30901077 | 2.19E-05 | 7 |
| CHURC1 | -0.2845664 | 2.29E-05 | 7 |
| ZBTB7A | -0.3452466 | 2.30E-05 | 7 |
| GIMAP4 | -0.2859419 | 2.44E-05 | 7 |
| ANXA5 | 0.27533091 | 2.50E-05 | 7 |
| SNU13 | 0.27539386 | 2.71E-05 | 7 |
| BUB3 | 0.28965434 | 3.17E-05 | 7 |
| ZMYND8 | -0.306682 | 3.19E-05 | 7 |
| CHCHD1 | 0.26324007 | 3.30E-05 | 7 |
| CHD7 | -0.268204 | 3.32E-05 | 7 |
| LIMA1 | -0.2696917 | 3.34E-05 | 7 |
| PRDM2 | -0.3116436 | 3.43E-05 | 7 |
| PARK7 | 0.26577146 | 3.78E-05 | 7 |
| FAM172A | -0.2743905 | 4.02E-05 | 7 |
| MAN1A2 | -0.3145884 | 4.04E-05 | 7 |
| S100A11 | -0.3706922 | 4.06E-05 | 7 |
| GABPB1-AS1 | -0.4501078 | 4.19E-05 | 7 |
| TIMM13 | 0.25936189 | 4.27E-05 | 7 |
| RASGRP2 | -0.2923477 | 4.32E-05 | 7 |
| UBE2J1 | 0.27270299 | 4.34E-05 | 7 |
| TMEM243 | -0.3036153 | 4.61E-05 | 7 |
| PRRC2B | -0.2782111 | 4.81E-05 | 7 |
| RNF168 | 0.25835205 | 4.92E-05 | 7 |
| GSE1 | -0.3423131 | 5.32E-05 | 7 |
| ZNF101 | -0.3013971 | 5.52E-05 | 7 |
| GLG1 | -0.3204652 | 5.95E-05 | 7 |
| BTF3 | -0.2834601 | 6.82E-05 | 7 |
| MIA2 | -0.2586747 | 7.05E-05 | 7 |
| OGA | -0.330489 | 7.11E-05 | 7 |
| PRF1 | -0.2864288 | 8.08E-05 | 7 |
| FGD3 | -0.3004188 | 8.14E-05 | 7 |
| SLC7A6 | -0.2806207 | 8.57E-05 | 7 |
| AMD1 | 0.2581022 | 8.90E-05 | 7 |
| ATP2A3 | 0.27223857 | 9.44E-05 | 7 |
| ATP6AP2 | -0.281607 | 9.72E-05 | 7 |
| SSR2 | -0.2808392 | 0.00010624 | 7 |
| PCMTD1 | -0.3026956 | 0.00011601 | 7 |
| ORMDL1 | -0.3186691 | 0.00011917 | 7 |

| | | | |
|---|---|---|---|
| MIF | 0.52379834 | 0.00012184 | 7 |
| CAST | -0.4051481 | 0.00012299 | 7 |
| BEX3 | 0.2862231 | 0.00012339 | 7 |
| CASC15 | -0.3626548 | 0.0001236 | 7 |
| HSP90AB1 | 0.47634131 | 0.00012799 | 7 |
| ARHGAP25 | -0.2628852 | 0.00012911 | 7 |
| UBA2 | 0.27671363 | 0.0001374 | 7 |
| LEPROTL1 | -0.331755 | 0.00014363 | 7 |
| ARHGAP45 | -0.3194626 | 0.00015004 | 7 |
| CCDC57 | -0.2972386 | 0.00015942 | 7 |
| MAP1LC3B | -0.2785055 | 0.00016301 | 7 |
| TBC1D2B | -0.2867844 | 0.00016445 | 7 |
| TGOLN2 | -0.3143074 | 0.00017059 | 7 |
| EIF3H | -0.2921531 | 0.00018188 | 7 |
| TNRC6C | -0.3802806 | 0.00018355 | 7 |
| TSC22D4 | -0.2950661 | 0.00018827 | 7 |
| SH3KBP1 | -0.2991838 | 0.00019242 | 7 |
| TES | -0.3492341 | 0.00020451 | 7 |
| FAM214A | -0.2767279 | 0.00021644 | 7 |
| RUNX3 | -0.2729333 | 0.00021901 | 7 |
| AC144521.1 | -0.2852776 | 0.00021998 | 7 |
| SAMD9 | -0.3067558 | 0.00022255 | 7 |
| PTGDR | -0.3646279 | 0.00022305 | 7 |
| STMN3 | -0.3028995 | 0.00024806 | 7 |
| C1orf56 | -0.303664 | 0.00025358 | 7 |
| SAMD12 | -0.2892969 | 0.00025497 | 7 |
| PLAC8 | -0.5006406 | 0.00025969 | 7 |
| SKAP1 | -0.3271498 | 0.00027309 | 7 |
| ETFB | 0.25027313 | 0.00027864 | 7 |
| ARHGEF1 | -0.2944523 | 0.00028816 | 7 |
| AC245060.5 | 0.27934714 | 0.00030325 | 7 |
| CD244 | -0.2539681 | 0.00030559 | 7 |
| KIAA2026 | -0.3095896 | 0.00030645 | 7 |
| ZNF91 | -0.3576827 | 0.00030892 | 7 |
| ZNF217 | -0.3364017 | 0.00032417 | 7 |
| NOL4L | -0.3020318 | 0.00036773 | 7 |
| BIRC2 | -0.3450937 | 0.0003919 | 7 |
| DHRS7 | -0.3135161 | 0.00040109 | 7 |
| PHF3 | -0.3051282 | 0.00043006 | 7 |
| PRKCQ-AS1 | -0.3330848 | 0.00044107 | 7 |
| CLDND1 | -0.366586 | 0.00044281 | 7 |
| HCG18 | -0.3116915 | 0.00045142 | 7 |
| AC108134.3 | -0.3062688 | 0.00047948 | 7 |

| | | | |
|---|---|---|---|
| CDK2AP2 | 0.2596868 | 0.00049812 | 7 |
| LPGAT1 | -0.3448939 | 0.00052114 | 7 |
| PCSK5 | -0.2921589 | 0.00058123 | 7 |
| RAP1B | -0.2859822 | 0.00058282 | 7 |
| ZSCAN18 | -0.2522862 | 0.00062697 | 7 |
| JAKMIP2 | -0.2585468 | 0.00065431 | 7 |
| CCDC88C | -0.2522961 | 0.00071429 | 7 |
| IGBP1 | -0.2846481 | 0.0007489 | 7 |
| HSPA9 | 0.26660478 | 0.00074924 | 7 |
| TLE4 | -0.2818977 | 0.00075314 | 7 |
| SERBP1 | 0.25395158 | 0.00076339 | 7 |
| PRKACB | -0.3229821 | 0.00081274 | 7 |
| DENND2D | -0.3132254 | 0.00088643 | 7 |
| ATP2A1 | -0.2893281 | 0.000893 | 7 |
| SERINC5 | -0.3668598 | 0.00102745 | 7 |
| DDX21 | 0.26656408 | 0.0010429 | 7 |
| CYB5A | 0.27634066 | 0.00106659 | 7 |
| ITGAL | -0.2953776 | 0.00112826 | 7 |
| ERBIN | -0.296745 | 0.00132527 | 7 |
| DST | -0.2633617 | 0.00134257 | 7 |
| SUPT3H | -0.2507009 | 0.00141641 | 7 |
| MYEF2 | 0.36032318 | 0.00145336 | 7 |
| KMT2C | -0.2803134 | 0.0014624 | 7 |
| RPS27L | 0.30993668 | 0.00150582 | 7 |
| PCM1 | 0.26121427 | 0.00157242 | 7 |
| SMIM24 | -0.4088456 | 0.00180007 | 7 |
| CR2 | -0.293749 | 0.00213505 | 7 |
| ZEB1 | 0.27505966 | 0.00225332 | 7 |
| FOXN3 | -0.2568904 | 0.0022849 | 7 |
| PPP1R2 | -0.3431376 | 0.00233745 | 7 |
| AC008105.3 | -0.2795138 | 0.00273593 | 7 |
| RBBP4 | 0.26243287 | 0.00277756 | 7 |
| Mar 06 | -0.3204379 | 0.0028742 | 7 |
| AC025171.2 | -0.2574891 | 0.00334533 | 7 |
| TOP1 | 0.28913302 | 0.00349002 | 7 |
| KLRK1 | -0.2678966 | 0.00354906 | 7 |
| CTSD | -0.3015742 | 0.00400964 | 7 |
| BAZ2B | -0.289494 | 0.00411361 | 7 |
| CASK | -0.277247 | 0.00452379 | 7 |
| CANX | 0.27015068 | 0.00455367 | 7 |
| YIPF4 | -0.3426622 | 0.0045588 | 7 |
| RBL2 | -0.3128857 | 0.00543947 | 7 |
| RPL14 | -0.2512608 | 0.00573557 | 7 |

| | | | |
|---|---|---|---|
| MXRA7 | -0.2752744 | 0.00627396 | 7 |
| TGFBR2 | -0.2741994 | 0.00655114 | 7 |
| ADGRE5 | -0.2527898 | 0.00700648 | 7 |
| DDX24 | -0.2525339 | 0.00705748 | 7 |
| GCC2 | -0.3070408 | 0.0070692 | 7 |
| ABI1 | -0.2560795 | 0.00708228 | 7 |
| HELB | -0.2957715 | 0.00725775 | 7 |
| PSAP | 0.25539673 | 0.00770074 | 7 |
| ST8SIA4 | -0.2607772 | 0.00788355 | 7 |
| KLRG1 | -0.2716477 | 0.00815835 | 7 |
| MIDN | -0.2528125 | 0.00905431 | 7 |
| PLCG2 | -0.2678801 | 0.01021955 | 7 |
| TAOK3 | -0.2963364 | 0.01062284 | 7 |
| CASD1 | -0.260582 | 0.01207221 | 7 |
| HDAC7 | -0.2950549 | 0.01556288 | 7 |
| CEP170 | -0.2683859 | 0.01654461 | 7 |
| MSL3 | -0.2588798 | 0.0167402 | 7 |
| NCOA7 | -0.2943338 | 0.01937909 | 7 |
| ZMYM2 | -0.287001 | 0.02028292 | 7 |
| ESCO1 | -0.2571896 | 0.02042864 | 7 |
| TCL1A | 0.29515211 | 0.02318829 | 7 |
| HERC4 | -0.2684508 | 0.02440052 | 7 |
| ARID5B | -0.2569038 | 0.02561097 | 7 |
| SORL1 | -0.2832152 | 0.02753651 | 7 |
| CASP4 | -0.2693046 | 0.02866696 | 7 |
| CHST2 | -0.2647515 | 0.03202679 | 7 |
| ODF2L | -0.2655844 | 0.03347084 | 7 |
| ZC3HAV1 | -0.2550761 | 0.03715137 | 7 |
| VAMP5 | -0.2630956 | 0.04158741 | 7 |
| XIAP | -0.2781316 | 0.04399669 | 7 |
| RPAP2 | -0.3853538 | 0.04475039 | 7 |
| ZNF721 | -0.2733468 | 0.05572394 | 7 |
| RASSF5 | -0.2569349 | 0.05668375 | 7 |
| EMB | -0.2656972 | 0.05800594 | 7 |
| GLS | -0.2904819 | 0.07658512 | 7 |
| LINC01222 | -0.4188712 | 0.07797569 | 7 |
| ZKSCAN1 | -0.2897477 | 0.08712604 | 7 |
| DNMT3A | -0.2816555 | 0.09027406 | 7 |
| HSP90B1 | 0.31533123 | 0.09118184 | 7 |
| ARHGAP26 | -0.2675821 | 0.12031909 | 7 |
| MCM5 | 0.25441653 | 0.12476523 | 7 |
| ARAP2 | -0.3150893 | 0.12551399 | 7 |
| MCM6 | 0.29357175 | 0.18383434 | 7 |

| | | | |
|---|---|---|---|
| CDKN1B | 0.26887981 | 0.19224597 | 7 |
| NMT2 | -0.2761858 | 0.19760695 | 7 |
| DNAJB14 | -0.2663003 | 0.20720486 | 7 |
| IRF2BP2 | -0.2688973 | 0.251299 | 7 |
| SETX | -0.2572213 | 0.30579552 | 7 |
| KDM5A | -0.254258 | 0.31027711 | 7 |
| TRIM38 | -0.259947 | 0.44022535 | 7 |
| CCDC186 | -0.2602694 | 0.48361784 | 7 |
| ZNF431 | -0.2794201 | 0.51767851 | 7 |
| MME | -0.2550465 | 0.58938936 | 7 |
| CEP85L | -0.2785754 | 0.60072704 | 7 |
| AAK1 | -0.2742929 | 0.73925948 | 7 |
| EPB41 | -0.2702776 | 1 | 7 |
| AC068587.4 | 0.25328261 | 1 | 7 |
| S100A10 | -0.3547956 | 1 | 7 |
| ORAI2 | -0.2517886 | 1 | 7 |
| LPP | -0.2700075 | 1 | 7 |
| TNFAIP3 | -0.261711 | 1 | 7 |
| SELENOW | 0.2565536 | 1 | 7 |
| ITGA4 | -0.3123508 | 1 | 7 |
| TMSB10 | -0.2864069 | 1 | 7 |
| GZMK | 2.04373702 | 5.68E-220 | 8 |
| SLAMF7 | 0.75804556 | 6.06E-190 | 8 |
| FASLG | 0.42649943 | 1.94E-101 | 8 |
| IFNG-AS1 | 1.09540859 | 1.59E-91 | 8 |
| PLEK | 0.56358078 | 4.14E-86 | 8 |
| ZNF80 | 1.8570533 | 1.57E-70 | 8 |
| LINC01871 | 0.251398 | 2.03E-62 | 8 |
| EOMES | 1.25666692 | 7.85E-56 | 8 |
| GIMAP4 | 1.39943141 | 1.33E-38 | 8 |
| CST7 | 1.39330284 | 8.08E-31 | 8 |
| FCRL3 | 0.27561931 | 7.05E-28 | 8 |
| EFNB1 | 0.31209014 | 2.90E-27 | 8 |
| GZMM | 1.23479213 | 4.81E-22 | 8 |
| SH2D1A | 1.34558322 | 2.94E-21 | 8 |
| IL4 | 0.43727082 | 7.48E-19 | 8 |
| HSBP1L1 | 0.77896859 | 1.00E-18 | 8 |
| GIMAP7 | 1.1310467 | 8.97E-18 | 8 |
| MEF2C | 0.42475601 | 4.82E-15 | 8 |
| KLRC3 | 0.98978238 | 1.54E-13 | 8 |
| NKG7 | 1.34934316 | 6.68E-13 | 8 |
| FCRL6 | 0.48044795 | 8.94E-13 | 8 |
| VIM | -1.5573396 | 9.87E-13 | 8 |

| | | | |
|---|---|---|---|
| IL10RA | 0.60687064 | 8.93E-12 | 8 |
| KLRD1 | 0.34486601 | 1.46E-11 | 8 |
| CD81 | 1.20415188 | 1.74E-11 | 8 |
| STX11 | 0.27123552 | 4.16E-11 | 8 |
| KLRC2 | 0.54050943 | 5.99E-11 | 8 |
| YPEL1 | 0.77081318 | 7.08E-11 | 8 |
| SRGN | 1.06954243 | 9.78E-11 | 8 |
| XCL1 | 0.32173304 | 2.63E-10 | 8 |
| VIPR2 | 0.89402323 | 1.52E-08 | 8 |
| PYHIN1 | 1.06608802 | 2.10E-08 | 8 |
| PILRB | 0.76641865 | 3.14E-08 | 8 |
| HLA-B | 0.9236694 | 3.60E-08 | 8 |
| S100A4 | -1.6822295 | 2.14E-07 | 8 |
| PTGDR | 0.94143478 | 3.63E-07 | 8 |
| SAMD3 | 0.68659203 | 3.76E-07 | 8 |
| HLA-A | 0.77456489 | 6.18E-07 | 8 |
| SFMBT2 | 0.34211866 | 2.05E-06 | 8 |
| GIMAP6 | 0.56275928 | 4.63E-06 | 8 |
| BTG1 | 0.73335101 | 5.04E-06 | 8 |
| PTPRJ | 0.38830444 | 6.39E-06 | 8 |
| ARAP2 | 0.69584045 | 7.12E-06 | 8 |
| DNAJC1 | 0.52952596 | 7.95E-06 | 8 |
| CNN2 | 0.71576112 | 8.55E-06 | 8 |
| KLRB1 | 0.77155895 | 1.36E-05 | 8 |
| HLA-C | 0.63429332 | 3.31E-05 | 8 |
| RASGEF1A | 0.29930214 | 0.0001317 | 8 |
| PTMA | -0.565173 | 0.00013794 | 8 |
| EPB41 | 0.57179048 | 0.00014872 | 8 |
| CCSER2 | 0.62636266 | 0.00015351 | 8 |
| STMN1 | -0.866072 | 0.00019827 | 8 |
| CD1B | -0.8830029 | 0.00061569 | 8 |
| ITM2C | 0.79808475 | 0.00089217 | 8 |
| LST1 | -1.0033956 | 0.00141218 | 8 |
| B2M | 0.46261108 | 0.00162057 | 8 |
| CXCR4 | 0.64317757 | 0.00165159 | 8 |
| TIGIT | 0.38187376 | 0.00180765 | 8 |
| YPEL3 | 0.69293005 | 0.00213629 | 8 |
| PDE4DIP | 0.28199795 | 0.00256158 | 8 |
| IKZF3 | 0.54062079 | 0.0031914 | 8 |
| ERN1 | 0.41407064 | 0.0049074 | 8 |
| ITM2A | 0.68702459 | 0.00720344 | 8 |
| KLRK1 | 0.40268715 | 0.0073586 | 8 |
| YBX1 | -0.582993 | 0.00871621 | 8 |

| | | | |
|---|---|---|---|
| STK17A | 0.68741054 | 0.01096273 | 8 |
| CD99 | -0.7330027 | 0.01275216 | 8 |
| CD52 | -0.6769915 | 0.01326504 | 8 |
| AQP3 | -0.7737722 | 0.01349946 | 8 |
| PTP4A2 | 0.67621907 | 0.016497 | 8 |
| CD3G | 0.45414926 | 0.01894094 | 8 |
| RGS10 | -0.5840352 | 0.02047433 | 8 |
| APMAP | 0.4385871 | 0.02124597 | 8 |
| IFI16 | 0.55069915 | 0.02273037 | 8 |
| FAM118A | 0.32011133 | 0.02478478 | 8 |
| A2M-AS1 | 0.30087027 | 0.0292283 | 8 |
| EMP3 | -0.7757656 | 0.03016447 | 8 |
| ITGB2 | 0.46118059 | 0.03227948 | 8 |
| RPL41 | 0.34769129 | 0.03342918 | 8 |
| CAPG | -0.6829451 | 0.03389848 | 8 |
| CLDND1 | 0.77074742 | 0.03428862 | 8 |
| CDC42SE2 | 0.46504806 | 0.03481817 | 8 |
| GIMAP1 | 0.46867056 | 0.03727497 | 8 |
| SH3BGRL3 | -0.6263703 | 0.04257951 | 8 |
| FCMR | 0.63307489 | 0.04417206 | 8 |
| CD247 | -0.6546615 | 0.05203181 | 8 |
| SEPTIN6 | -0.4942349 | 0.0537409 | 8 |
| STAT4 | 0.34377818 | 0.05922529 | 8 |
| TMEM229B | 0.38371938 | 0.05951471 | 8 |
| ATP5F1E | 0.32527357 | 0.11159067 | 8 |
| UTRN | 0.52448927 | 0.11293571 | 8 |
| CXCR3 | 0.35105118 | 0.13030884 | 8 |
| CNR2 | 0.32234886 | 0.15245864 | 8 |
| PDE7A | 0.48644171 | 0.19828899 | 8 |
| ZFP36L2 | 0.4464592 | 0.19904425 | 8 |
| PKM | -0.5070069 | 0.21446644 | 8 |
| LGALS1 | -1.2671772 | 0.21706959 | 8 |
| TAGAP | 0.52474349 | 0.21884006 | 8 |
| VDAC1 | -0.5573346 | 0.22917664 | 8 |
| SPN | -0.5232541 | 0.23631381 | 8 |
| COX6C | 0.5081691 | 0.26076399 | 8 |
| ACTG1 | -0.4089372 | 0.2647277 | 8 |
| YWHAQ | -0.4963829 | 0.48443675 | 8 |
| IL2RB | 0.28639786 | 0.48770935 | 8 |
| DIAPH1 | -0.4926273 | 0.52896965 | 8 |
| GNAS | -0.3511538 | 0.53105313 | 8 |
| FLNA | -0.5224001 | 0.5472908 | 8 |
| ITGAL | 0.40939878 | 0.55391174 | 8 |

| | | | |
|---|---|---|---|
| AP005482.1 | 0.71562564 | 0.56464414 | 8 |
| CRYBG1 | -0.3799167 | 0.61296052 | 8 |
| RPS15A | 0.30229005 | 0.62864845 | 8 |
| FYB1 | 0.36449712 | 0.63027901 | 8 |
| SLC25A5 | -0.5143416 | 0.63887026 | 8 |
| TPM3 | -0.4996941 | 0.65061635 | 8 |
| GAPDH | -0.6649092 | 0.68364112 | 8 |
| REEP5 | 0.26771893 | 0.72256072 | 8 |
| H3F3A | -0.3191658 | 0.79801343 | 8 |
| CCR9 | -0.6134633 | 0.80532334 | 8 |
| GIHCG | -0.7982626 | 0.81649259 | 8 |
| SASH3 | 0.40516342 | 0.92175396 | 8 |
| RPS29 | 0.28493859 | 1 | 8 |
| PLXDC1 | 0.25903768 | 1 | 8 |
| PXK | 0.27980255 | 1 | 8 |
| CYLD | 0.51482124 | 1 | 8 |
| ATM | 0.47155358 | 1 | 8 |
| ID3 | -0.8839072 | 1 | 8 |
| WIPF1 | 0.47812703 | 1 | 8 |
| RPLP2 | 0.25088303 | 1 | 8 |
| RUNX3 | 0.43959207 | 1 | 8 |
| PPP1R12B | 0.30997104 | 1 | 8 |
| MYO1G | -0.4522088 | 1 | 8 |
| MEX3C | 0.28884952 | 1 | 8 |
| CRIP1 | -1.1050284 | 1 | 8 |
| HSPB1 | 0.30959455 | 1 | 8 |
| SHISA2 | 0.44309258 | 1 | 8 |
| MAPK1 | 0.43440393 | 1 | 8 |
| SIK3 | 0.39705291 | 1 | 8 |
| HLA-E | 0.33210062 | 1 | 8 |
| CD1E | -0.7487782 | 1 | 8 |
| TMSB10 | -0.5739431 | 1 | 8 |
| TXNIP | 0.38267635 | 1 | 8 |
| RPL27 | 0.27996646 | 1 | 8 |
| BCR | 0.35228806 | 1 | 8 |
| ANXA1 | -0.7916933 | 1 | 8 |
| CD27 | 0.32527839 | 1 | 8 |
| LINC-PINT | 0.30391697 | 1 | 8 |
| SYNRG | 0.33223765 | 1 | 8 |
| SNRPF | -0.4708874 | 1 | 8 |
| SSRP1 | -0.4389436 | 1 | 8 |
| CD1A | -0.8585121 | 1 | 8 |
| TRAC | -0.421712 | 1 | 8 |

| | | | |
|---|---|---|---|
| INPP1 | 0.25759954 | 1 | 8 |
| TUBA1B | -1.0267258 | 1 | 8 |
| CD79A | -0.5549005 | 1 | 8 |
| FTL | 0.38392425 | 1 | 8 |
| PPIA | -0.4597297 | 1 | 8 |
| RASSF5 | 0.30818166 | 1 | 8 |
| HNRNPM | -0.4149294 | 1 | 8 |
| PLEKHG1 | 0.30442345 | 1 | 8 |
| RPS27 | 0.28604612 | 1 | 8 |
| NAP1L1 | -0.447213 | 1 | 8 |
| ERGIC3 | -0.3468161 | 1 | 8 |
| TIMP1 | -0.5138942 | 1 | 8 |
| PAFAH2 | 0.35034761 | 1 | 8 |
| GLUL | -0.4376807 | 1 | 8 |
| PSIP1 | -0.4417725 | 1 | 8 |
| MIB2 | 0.30078459 | 1 | 8 |
| DDX5 | 0.28622984 | 1 | 8 |
| S1PR3 | -0.425396 | 1 | 8 |
| SMAP2 | 0.36872871 | 1 | 8 |
| COTL1 | 0.45427756 | 1 | 8 |
| DTNBP1 | 0.40519639 | 1 | 8 |
| ILF2 | -0.4331472 | 1 | 8 |
| NREP | -0.3682991 | 1 | 8 |
| SUCLG2 | 0.28596185 | 1 | 8 |
| TFDP2 | -0.6447976 | 1 | 8 |
| RALGPS2 | 0.39793427 | 1 | 8 |
| BTN3A2 | 0.41995301 | 1 | 8 |
| MCMBP | -0.3395452 | 1 | 8 |
| GALNT2 | -0.4407356 | 1 | 8 |
| TXN | -0.560124 | 1 | 8 |
| FNBP4 | 0.43149412 | 1 | 8 |
| CHST2 | 0.32680917 | 1 | 8 |
| KIAA1324L | 0.34269949 | 1 | 8 |
| BAHCC1 | -0.4207766 | 1 | 8 |
| ARMH1 | -0.5095781 | 1 | 8 |
| NR3C1 | 0.29356549 | 1 | 8 |
| ETV5 | -0.5564651 | 1 | 8 |
| CFLAR | 0.41869647 | 1 | 8 |
| DDIT4 | -0.4452961 | 1 | 8 |
| ANXA5 | -0.4166976 | 1 | 8 |
| JUND | -0.4179382 | 1 | 8 |
| CD2 | 0.31988391 | 1 | 8 |
| S100A6 | -0.6107363 | 1 | 8 |

| | | | |
|---|---|---|---|
| RPL12 | 0.29463636 | 1 | 8 |
| LINC00861 | 0.28006441 | 1 | 8 |
| CTSZ | 0.33249345 | 1 | 8 |
| ATF7IP2 | -0.3056666 | 1 | 8 |
| EVL | 0.35134194 | 1 | 8 |
| C12orf75 | -0.4394924 | 1 | 8 |
| IDS | 0.35797781 | 1 | 8 |
| TFCP2 | 0.34334561 | 1 | 8 |
| S100A10 | -0.5972048 | 1 | 8 |
| TPM4 | -0.419297 | 1 | 8 |
| TUBB | -0.7170696 | 1 | 8 |
| PRKCB | -0.37835 | 1 | 8 |
| NME2 | -0.4888093 | 1 | 8 |
| PLCG2 | 0.33735693 | 1 | 8 |
| CDPF1 | -0.3096647 | 1 | 8 |
| INSIG1 | 0.30896116 | 1 | 8 |
| IPCEF1 | 0.31908365 | 1 | 8 |
| HK1 | -0.2980634 | 1 | 8 |
| RBL2 | 0.35759685 | 1 | 8 |
| UBASH3B | -0.3942014 | 1 | 8 |
| MSH6 | -0.3951021 | 1 | 8 |
| ZNF706 | -0.3862198 | 1 | 8 |
| HCST | 0.2833934 | 1 | 8 |
| TXNL4A | 0.39281499 | 1 | 8 |
| RERE | 0.26558521 | 1 | 8 |
| KLRG1 | 0.27540572 | 1 | 8 |
| AUTS2 | 0.36329326 | 1 | 8 |
| SBK1 | -0.3166481 | 1 | 8 |
| NEK7 | 0.33942653 | 1 | 8 |
| YWHAZ | -0.2851471 | 1 | 8 |
| RPS18 | 0.26175913 | 1 | 8 |
| MAZ | -0.3559967 | 1 | 8 |
| TNRC6B | 0.35878236 | 1 | 8 |
| ST3GAL1 | 0.32347921 | 1 | 8 |
| ARPP21 | -0.4159045 | 1 | 8 |
| PIK3R1 | 0.31076396 | 1 | 8 |
| SLC25A3 | -0.3335351 | 1 | 8 |
| GLIPR2 | -0.3006012 | 1 | 8 |
| HELLS | -0.602167 | 1 | 8 |
| OSBPL8 | -0.4194158 | 1 | 8 |
| MAN2A1 | 0.33522356 | 1 | 8 |
| SRPK2 | 0.30877931 | 1 | 8 |
| TOMM7 | 0.32545183 | 1 | 8 |

| | | | |
|---|---|---|---|
| RGL4 | -0.3194076 | 1 | 8 |
| CASC4 | 0.25952285 | 1 | 8 |
| PLP2 | -0.3421321 | 1 | 8 |
| SUB1 | -0.2862694 | 1 | 8 |
| HLA-F | 0.25642804 | 1 | 8 |
| SEPTIN7 | 0.40666966 | 1 | 8 |
| SNHG29 | -0.4363286 | 1 | 8 |
| SMPD3 | 0.26120958 | 1 | 8 |
| RPLP0 | -0.348232 | 1 | 8 |
| SEC24C | 0.2502076 | 1 | 8 |
| IGFBP5 | -0.6751167 | 1 | 8 |
| TYMS | -0.7063345 | 1 | 8 |
| IRF1 | 0.32096829 | 1 | 8 |
| MSH2 | -0.289952 | 1 | 8 |
| PTPN2 | 0.42819719 | 1 | 8 |
| FOXO1 | 0.25727694 | 1 | 8 |
| CDK6 | -0.5817628 | 1 | 8 |
| USP20 | -0.2845911 | 1 | 8 |
| ANP32B | -0.4051506 | 1 | 8 |
| SEPHS2 | -0.2827308 | 1 | 8 |
| PSMA6 | -0.315451 | 1 | 8 |
| ZWINT | -0.3218861 | 1 | 8 |
| STAU2 | -0.3013496 | 1 | 8 |
| ITGB2-AS1 | 0.27176859 | 1 | 8 |
| JADE2 | 0.29108474 | 1 | 8 |
| INPP5D | 0.41268261 | 1 | 8 |
| CNOT6L | 0.32239358 | 1 | 8 |
| PRDX2 | -0.2780694 | 1 | 8 |
| GRAMD1A | 0.42480411 | 1 | 8 |
| ATP5F1C | -0.3485538 | 1 | 8 |
| LDLRAD4 | -0.3653456 | 1 | 8 |
| NAA10 | -0.309235 | 1 | 8 |
| FAM200B | -0.2918217 | 1 | 8 |
| CBX5 | -0.4380888 | 1 | 8 |
| ARF6 | -0.2873153 | 1 | 8 |
| HTATSF1 | -0.2741497 | 1 | 8 |
| SPATA13 | 0.34971557 | 1 | 8 |
| MLLT10 | -0.2642037 | 1 | 8 |
| TKT | -0.3566268 | 1 | 8 |
| DYNC1I2 | -0.2924387 | 1 | 8 |
| DDX6 | 0.29899566 | 1 | 8 |
| ASPM | -0.6078862 | 1 | 8 |
| CASP2 | -0.3660872 | 1 | 8 |

| | | | |
|---|---|---|---|
| JAK1 | 0.36972562 | 1 | 8 |
| NCK1 | 0.27592321 | 1 | 8 |
| CCNK | 0.2800254 | 1 | 8 |
| KMT2E | 0.25954065 | 1 | 8 |
| ELOVL4 | -0.4512159 | 1 | 8 |
| MCM3 | -0.3807059 | 1 | 8 |
| SHPRH | 0.2834437 | 1 | 8 |
| CCDC50 | -0.2790131 | 1 | 8 |
| POLD3 | -0.2626618 | 1 | 8 |
| RPL14 | -0.325964 | 1 | 8 |
| FAM89B | -0.3134276 | 1 | 8 |
| PTPN6 | -0.4301737 | 1 | 8 |
| LCP2 | -0.3890967 | 1 | 8 |
| TLE4 | -0.4097383 | 1 | 8 |
| CAPNS1 | -0.3071877 | 1 | 8 |
| RALY | -0.3401006 | 1 | 8 |
| ARID4B | 0.37949437 | 1 | 8 |
| RAPGEF6 | 0.32323 | 1 | 8 |
| AL138899.1 | -0.6558237 | 1 | 8 |
| CIRBP | 0.34221954 | 1 | 8 |
| HMGN2 | -0.5611034 | 1 | 8 |
| SYNE2 | -0.4461679 | 1 | 8 |
| CENPX | -0.260363 | 1 | 8 |
| COX5A | -0.3476188 | 1 | 8 |
| LRMP | 0.30853929 | 1 | 8 |
| LINC01222 | -0.5678425 | 1 | 8 |
| SMC2 | -0.4883481 | 1 | 8 |
| SH3GLB2 | -0.2796249 | 1 | 8 |
| KLHL6 | -0.3063232 | 1 | 8 |
| BARD1 | -0.3250455 | 1 | 8 |
| CAPN2 | -0.2991836 | 1 | 8 |
| ATAD2 | -0.4683781 | 1 | 8 |
| MARCKSL1 | -0.2970393 | 1 | 8 |
| TAOK3 | 0.3009205 | 1 | 8 |
| TIAM1 | -0.3693795 | 1 | 8 |
| CDK2AP1 | -0.339981 | 1 | 8 |
| BRK1 | -0.2968406 | 1 | 8 |
| TMEM218 | 0.25391921 | 1 | 8 |
| RANBP1 | -0.3591047 | 1 | 8 |
| MCM6 | -0.3353005 | 1 | 8 |
| SPRY1 | 0.4152203 | 1 | 8 |
| CBLB | 0.29815883 | 1 | 8 |
| NFATC3 | -0.3697907 | 1 | 8 |

| | | | |
|---|---|---|---|
| JPT1 | -0.3716032 | 1 | 8 |
| GALNT6 | -0.3666072 | 1 | 8 |
| HUWE1 | -0.271913 | 1 | 8 |
| COA3 | -0.256067 | 1 | 8 |
| SRSF2 | -0.3546142 | 1 | 8 |
| MAD2L2 | -0.2728971 | 1 | 8 |
| CRNDE | -0.3880971 | 1 | 8 |
| WDR34 | -0.3010229 | 1 | 8 |
| PCAT18 | -0.3042869 | 1 | 8 |
| HMGB1 | -0.4048633 | 1 | 8 |
| NDUFB1 | -0.2702401 | 1 | 8 |
| FYN | 0.25200033 | 1 | 8 |
| CDT1 | -0.2615949 | 1 | 8 |
| ACBD3 | -0.2705513 | 1 | 8 |
| WAKMAR2 | 0.39094808 | 1 | 8 |
| PRDX3 | -0.2941515 | 1 | 8 |
| IFITM2 | -0.5085408 | 1 | 8 |
| CORO1A | -0.3001745 | 1 | 8 |
| HCG18 | 0.29949676 | 1 | 8 |
| CD79B | 0.25475146 | 1 | 8 |
| H2AFX | -0.3964169 | 1 | 8 |
| GTF3A | 0.25821466 | 1 | 8 |
| ARPC5L | 0.2936587 | 1 | 8 |
| WDR1 | -0.2935115 | 1 | 8 |
| PDAP1 | -0.3124593 | 1 | 8 |
| NDUFS3 | -0.2687933 | 1 | 8 |
| DGKZ | 0.26228869 | 1 | 8 |
| NASP | -0.3717174 | 1 | 8 |
| LSM2 | -0.3164616 | 1 | 8 |
| KIF20B | -0.3366623 | 1 | 8 |
| MLLT11 | -0.2507122 | 1 | 8 |
| NDUFA4 | -0.3197523 | 1 | 8 |
| TNFAIP3 | 0.29135278 | 1 | 8 |
| NDUFS6 | -0.3146156 | 1 | 8 |
| CAMK4 | 0.30009107 | 1 | 8 |
| LAX1 | 0.39233716 | 1 | 8 |
| CD44 | 0.39389171 | 1 | 8 |
| SEC31B | 0.28004551 | 1 | 8 |
| HNRNPAB | -0.3967317 | 1 | 8 |
| HMGB3 | -0.2768819 | 1 | 8 |
| CD7 | -0.3377981 | 1 | 8 |
| RBM28 | 0.30309381 | 1 | 8 |
| TIMM10 | -0.2669409 | 1 | 8 |

| | | | |
|---|---|---|---|
| PCLAF | -0.6401106 | 1 | 8 |
| SIVA1 | -0.469397 | 1 | 8 |
| MAD2L1 | -0.2605273 | 1 | 8 |
| PSMB1 | -0.2748617 | 1 | 8 |
| ACTL6A | -0.2665457 | 1 | 8 |
| NOTCH1 | -0.2815208 | 1 | 8 |
| C12orf57 | -0.3601633 | 1 | 8 |
| ARHGAP25 | 0.31648943 | 1 | 8 |
| MBNL1 | 0.31911585 | 1 | 8 |
| RNF167 | -0.2611138 | 1 | 8 |
| C1orf162 | -0.2853939 | 1 | 8 |
| FMNL1 | -0.2568541 | 1 | 8 |
| CAPRIN1 | -0.2702668 | 1 | 8 |
| ESCO1 | 0.28261938 | 1 | 8 |
| CKS2 | -0.3574906 | 1 | 8 |
| M6PR | 0.33915543 | 1 | 8 |
| CHD4 | -0.2791774 | 1 | 8 |
| LBH | 0.25164925 | 1 | 8 |
| FAM111B | -0.2547559 | 1 | 8 |
| COA1 | 0.2801026 | 1 | 8 |
| AATF | 0.26303237 | 1 | 8 |
| AAK1 | 0.32152307 | 1 | 8 |
| HMGA1 | -0.3490843 | 1 | 8 |
| ZNF92 | 0.26078758 | 1 | 8 |
| GTF2A2 | -0.2691668 | 1 | 8 |
| NUDT21 | -0.2838204 | 1 | 8 |
| RPS15 | 1.05756912 | 1.35E-14 | 9 |
| RPL41 | 0.93737603 | 5.62E-14 | 9 |
| RPL18A | 0.99211713 | 9.71E-14 | 9 |
| RPL28 | 1.04224956 | 1.29E-13 | 9 |
| RPL8 | 0.95874603 | 3.44E-13 | 9 |
| RPS19 | 1.03554553 | 3.83E-13 | 9 |
| RPS28 | 0.94315294 | 8.19E-13 | 9 |
| RPS26 | 1.22506296 | 8.46E-13 | 9 |
| RPL35A | 0.97611756 | 1.23E-12 | 9 |
| RPL10 | 0.87631081 | 1.44E-12 | 9 |
| RPL3 | 0.93237119 | 1.55E-12 | 9 |
| MALAT1 | -1.8619301 | 1.68E-12 | 9 |
| RPS9 | 0.95516946 | 2.66E-12 | 9 |
| RPS4X | 0.95527184 | 2.91E-12 | 9 |
| RPL19 | 0.8497144 | 3.51E-12 | 9 |
| RPS15A | 0.84704257 | 4.56E-12 | 9 |
| RPL13 | 0.86211771 | 5.40E-12 | 9 |

| | | | |
|---|---|---|---|
| RPL11 | 0.92211119 | 7.05E-12 | 9 |
| RPS2 | 0.9143841 | 1.54E-11 | 9 |
| RPS18 | 0.86342151 | 1.96E-11 | 9 |
| RPS23 | 0.89645386 | 4.16E-11 | 9 |
| RPS27A | 0.89612869 | 5.55E-11 | 9 |
| RPL18 | 0.87831628 | 7.07E-11 | 9 |
| RPL32 | 0.92282315 | 9.83E-11 | 9 |
| RPS3 | 0.90092822 | 1.22E-10 | 9 |
| RPS5 | 1.11497948 | 1.25E-10 | 9 |
| RPS24 | 0.80975343 | 1.51E-10 | 9 |
| RPL7A | 0.84890108 | 1.56E-10 | 9 |
| RPS13 | 0.97229243 | 1.57E-10 | 9 |
| RPL30 | 0.83095768 | 1.85E-10 | 9 |
| RPLP1 | 0.85320613 | 2.29E-10 | 9 |
| RPS12 | 0.89008735 | 2.35E-10 | 9 |
| RPL31 | 0.85697589 | 3.44E-10 | 9 |
| RPS7 | 1.02773903 | 3.54E-10 | 9 |
| FAU | 0.81994929 | 9.56E-10 | 9 |
| RPL6 | 0.9057232 | 1.09E-09 | 9 |
| RPS21 | 0.85911203 | 1.35E-09 | 9 |
| RPS3A | 0.86036002 | 1.43E-09 | 9 |
| RPL36 | 0.78878726 | 1.44E-09 | 9 |
| RPL35 | 0.86900639 | 2.04E-09 | 9 |
| RPL17 | 0.7855487 | 3.09E-09 | 9 |
| RPS25 | 0.87734709 | 3.27E-09 | 9 |
| RPL29 | 0.82830454 | 4.10E-09 | 9 |
| RPS6 | 0.90004914 | 5.72E-09 | 9 |
| RPL26 | 0.87395704 | 6.73E-09 | 9 |
| NACA | 0.89675197 | 7.15E-09 | 9 |
| RPS14 | 0.86044552 | 1.27E-08 | 9 |
| RPL24 | 0.82163387 | 1.80E-08 | 9 |
| RPL5 | 0.90404653 | 1.80E-08 | 9 |
| PFN1 | 0.98576991 | 2.30E-08 | 9 |
| RPL39 | 0.68825734 | 5.12E-08 | 9 |
| RPL12 | 0.82274926 | 7.38E-08 | 9 |
| TMSB10 | 0.97379366 | 1.92E-07 | 9 |
| RPL21 | 0.75065681 | 2.58E-07 | 9 |
| RPL22 | 0.82425672 | 4.28E-07 | 9 |
| RPL37 | 0.66722374 | 5.53E-07 | 9 |
| RPS27 | 0.6215816 | 6.86E-07 | 9 |
| SH3BGRL3 | 0.9459122 | 7.72E-07 | 9 |
| GAPDH | 1.03218707 | 8.93E-07 | 9 |
| RPL36A | 0.77709682 | 9.85E-07 | 9 |

| | | | |
|---|---|---|---|
| RPS10 | 0.8331319 | 1.00E-06 | 9 |
| RPL37A | 0.55800118 | 1.13E-06 | 9 |
| RPL34 | 0.71720638 | 1.87E-06 | 9 |
| RPS8 | 0.6398648 | 2.04E-06 | 9 |
| EEF1A1 | 0.60058288 | 2.31E-06 | 9 |
| C9orf16 | 0.86509363 | 2.32E-06 | 9 |
| RPLP2 | 0.62800567 | 2.92E-06 | 9 |
| ARHGDIB | 0.66123549 | 5.05E-06 | 9 |
| ATP5MG | 0.70981309 | 7.14E-06 | 9 |
| ACTG1 | 0.58716307 | 1.10E-05 | 9 |
| RPL10A | 0.65467598 | 1.11E-05 | 9 |
| RPSA | 0.64569068 | 1.54E-05 | 9 |
| RPLP0 | 0.70054929 | 2.85E-05 | 9 |
| TECR | 0.96654689 | 2.94E-05 | 9 |
| RPL9 | 0.60950711 | 3.01E-05 | 9 |
| SNHG29 | 0.86256569 | 3.07E-05 | 9 |
| SERF2 | 0.76769398 | 4.46E-05 | 9 |
| CFL1 | 0.6813486 | 5.23E-05 | 9 |
| XIST | -1.2046457 | 5.90E-05 | 9 |
| RPL23A | 0.54701237 | 7.00E-05 | 9 |
| POLR2J3 | -1.0691491 | 0.00010626 | 9 |
| RPL14 | 0.71256815 | 0.00011554 | 9 |
| UBA52 | 0.59324572 | 0.00011566 | 9 |
| MYL6 | 0.63325531 | 0.00013572 | 9 |
| RPS16 | 0.52839551 | 0.00015297 | 9 |
| EEF1B2 | 0.75627137 | 0.00015641 | 9 |
| NDUFB2 | 0.7554405 | 0.00026065 | 9 |
| FXYD2 | 1.08226973 | 0.0002941 | 9 |
| C1orf56 | 0.78197873 | 0.00083383 | 9 |
| TYROBP | 0.33005013 | 0.00084155 | 9 |
| CD3D | 0.59187476 | 0.00101069 | 9 |
| RACK1 | 0.59003723 | 0.00112403 | 9 |
| RPL15 | 0.53957076 | 0.00116994 | 9 |
| ZNF655 | -0.946327 | 0.00131996 | 9 |
| PFDN5 | 0.59912588 | 0.0014983 | 9 |
| IQGAP1 | -0.9485422 | 0.00150417 | 9 |
| ACTB | 0.5113453 | 0.00240381 | 9 |
| ARID1A | -0.9122104 | 0.00281386 | 9 |
| FTL | 0.68812188 | 0.00281517 | 9 |
| COX6B1 | 0.61972666 | 0.00295127 | 9 |
| CD52 | 0.65634173 | 0.0034001 | 9 |
| HINT1 | 0.59475658 | 0.00443533 | 9 |
| SMG1 | -0.9272037 | 0.0046258 | 9 |

| | | | |
|---|---|---|---|
| EEF1G | 0.65172091 | 0.00509945 | 9 |
| IGLL1 | 0.47388726 | 0.00638378 | 9 |
| TTN | -1.2298787 | 0.00712865 | 9 |
| RPL38 | 0.54253947 | 0.0072524 | 9 |
| COX4I1 | 0.6425847 | 0.00746038 | 9 |
| PTPN7 | -0.9411382 | 0.00788023 | 9 |
| PTPRC | -0.801452 | 0.00795177 | 9 |
| MACF1 | -1.0171417 | 0.00852229 | 9 |
| RNF213 | -0.9384753 | 0.00871734 | 9 |
| CAMK4 | -0.9254781 | 0.00926658 | 9 |
| FNBP1 | -0.8496192 | 0.01078872 | 9 |
| FUS | -0.8097077 | 0.01232568 | 9 |
| GSTP1 | 0.75444867 | 0.0123873 | 9 |
| ATP5F1E | 0.58441834 | 0.01260921 | 9 |
| MIF | 0.75679156 | 0.016236 | 9 |
| DDX17 | -0.7026432 | 0.01627656 | 9 |
| BTG2 | -1.0968333 | 0.01678764 | 9 |
| ARPC3 | 0.63406296 | 0.01732032 | 9 |
| UXT | 0.59673327 | 0.021936 | 9 |
| BMP1 | 0.26931788 | 0.02236307 | 9 |
| HIPK1 | -0.852468 | 0.02866737 | 9 |
| USP15 | -0.8358974 | 0.02968338 | 9 |
| HNRNPU | -0.7365808 | 0.03264069 | 9 |
| TRBC2 | 0.52500102 | 0.03301008 | 9 |
| KAT6B | -0.8705001 | 0.03518682 | 9 |
| ZNF292 | -0.8384735 | 0.03574271 | 9 |
| NOP53 | 0.66750528 | 0.04894173 | 9 |
| BTF3 | 0.58780028 | 0.0499459 | 9 |
| KANSL1 | -0.7941504 | 0.05140521 | 9 |
| ATP5MF | 0.68113238 | 0.05400305 | 9 |
| SYNE1 | -0.8565808 | 0.06279632 | 9 |
| ATP5MC2 | 0.62183811 | 0.07298593 | 9 |
| APRT | 0.65673615 | 0.07356435 | 9 |
| HNRNPA2B1 | -0.7200681 | 0.07776439 | 9 |
| KMT2C | -0.8087584 | 0.07946211 | 9 |
| EEF1D | 0.59915213 | 0.0805375 | 9 |
| BCL11B | -0.9517514 | 0.08303278 | 9 |
| COX7C | 0.52993155 | 0.08380216 | 9 |
| ARMH1 | 0.78014057 | 0.09974066 | 9 |
| PPIA | 0.61131235 | 0.10363049 | 9 |
| PRDX2 | 0.71083246 | 0.10432447 | 9 |
| CRIP1 | 0.32400681 | 0.11203062 | 9 |
| LINC00342 | -0.975547 | 0.11240091 | 9 |

| | | | |
|---|---|---|---|
| COMMD6 | 0.5850385 | 0.1125902 | 9 |
| NEAT1 | -1.0341381 | 0.12243152 | 9 |
| RPL36AL | 0.50659122 | 0.13569771 | 9 |
| SLC25A3 | 0.57739002 | 0.13688777 | 9 |
| SON | -0.63563 | 0.15792796 | 9 |
| NPM1 | 0.53498183 | 0.16106254 | 9 |
| ATRX | -0.8291955 | 0.16744547 | 9 |
| GTF2I | -0.6885606 | 0.16881798 | 9 |
| RB1CC1 | -0.7996284 | 0.17392962 | 9 |
| IL32 | 0.52543083 | 0.17425175 | 9 |
| GIMAP7 | 0.47887612 | 0.18089994 | 9 |
| SP100 | -0.824746 | 0.18731 | 9 |
| EIF1 | 0.45679315 | 0.1926555 | 9 |
| ATP5PF | 0.50105385 | 0.20018748 | 9 |
| MT-ND3 | -0.6727653 | 0.22381021 | 9 |
| CCNL1 | -0.7625654 | 0.23094221 | 9 |
| CHD9 | -0.7768527 | 0.23719391 | 9 |
| FXYD5 | 0.64559758 | 0.23953476 | 9 |
| NSD1 | -0.7084643 | 0.25503095 | 9 |
| COX7A2 | 0.59981703 | 0.25930997 | 9 |
| FYB1 | -0.7426595 | 0.28295562 | 9 |
| MTRNR2L12 | -0.7626764 | 0.28386769 | 9 |
| PPDPF | 0.62498108 | 0.29492632 | 9 |
| ZNF217 | -0.7603629 | 0.30057662 | 9 |
| TMEM161B-AS1 | -0.9469663 | 0.31449508 | 9 |
| LINC01222 | -0.9796566 | 0.3177815 | 9 |
| CBL | -0.7312647 | 0.3262645 | 9 |
| ZNF644 | -0.7492624 | 0.33504695 | 9 |
| APBB1IP | -0.6962795 | 0.35025112 | 9 |
| ASH1L | -0.7245251 | 0.35782997 | 9 |
| GNAQ | -0.8094081 | 0.3651666 | 9 |
| DMTF1 | -0.7303059 | 0.37032835 | 9 |
| PCM1 | -0.752412 | 0.37158363 | 9 |
| BDP1 | -0.8015234 | 0.38569315 | 9 |
| CUTA | 0.62115105 | 0.40164557 | 9 |
| ATP5PO | 0.55717287 | 0.40815866 | 9 |
| RPL27 | 0.52683738 | 0.44004727 | 9 |
| KMT2E | -0.6783354 | 0.45258038 | 9 |
| TMEM258 | 0.61337941 | 0.47420638 | 9 |
| TGOLN2 | -0.7133165 | 0.48071394 | 9 |
| STK17B | -0.7956295 | 0.48738637 | 9 |
| STK4 | -0.7695136 | 0.49323986 | 9 |
| PDCD6IP | -0.68461 | 0.50482868 | 9 |

| | | | |
|---|---|---|---|
| MYL12A | 0.50630231 | 0.50704711 | 9 |
| CD27 | 0.59840669 | 0.51692082 | 9 |
| UQCRH | 0.59229837 | 0.53042857 | 9 |
| ME2 | -0.6904816 | 0.53413353 | 9 |
| PNISR | -0.6982716 | 0.54358039 | 9 |
| N4BP2L2 | -0.7418684 | 0.5518 | 9 |
| SMCHD1 | -0.7365416 | 0.59911097 | 9 |
| AC068587.4 | -0.8089354 | 0.60197562 | 9 |
| GCC2 | -0.7968499 | 0.60817753 | 9 |
| TMA7 | 0.51165488 | 0.6233846 | 9 |
| RPL13A | 0.35841135 | 0.62571928 | 9 |
| SEPTIN6 | -0.5569555 | 0.65664969 | 9 |
| SNHG14 | -0.9179631 | 0.67163099 | 9 |
| EIF3K | 0.58132675 | 0.70699861 | 9 |
| TRAPPC6A | 0.42610779 | 0.72838488 | 9 |
| ZMYND8 | -0.6498947 | 0.7359637 | 9 |
| PRDX1 | 0.54632884 | 0.73981144 | 9 |
| SYNRG | -0.7260196 | 0.77052048 | 9 |
| SNRPD2 | 0.61811333 | 0.77067141 | 9 |
| TPT1 | 0.37860998 | 0.77748652 | 9 |
| SHPRH | -0.7743465 | 0.78894554 | 9 |
| SPEN | -0.7361418 | 0.80637092 | 9 |
| LARP1 | -0.6777364 | 0.8298952 | 9 |
| PJA2 | -0.7032036 | 0.86063392 | 9 |
| PRKDC | -0.7197831 | 0.93509816 | 9 |
| ARID1B | -0.7255937 | 0.97955484 | 9 |
| GOLGA4 | -0.7951839 | 1 | 9 |
| AC245060.5 | -0.8093114 | 1 | 9 |
| SLA | -0.9045861 | 1 | 9 |
| RBM25 | -0.6751096 | 1 | 9 |
| ZNF37A | -0.7284558 | 1 | 9 |
| TRIP11 | -0.7109552 | 1 | 9 |
| AAK1 | -0.8146407 | 1 | 9 |
| PPP2R5E | -0.6438035 | 1 | 9 |
| NUTM2A-AS1 | -0.6762753 | 1 | 9 |
| HCST | 0.61639895 | 1 | 9 |
| ID3 | 0.61544419 | 1 | 9 |
| ROCK1 | -0.6175485 | 1 | 9 |
| C12orf57 | 0.58283071 | 1 | 9 |
| ARHGAP26 | -0.6573397 | 1 | 9 |
| NIN | -0.7132948 | 1 | 9 |
| RBM39 | -0.5508049 | 1 | 9 |
| LGALS1 | 0.27724064 | 1 | 9 |

| | | | |
|---|---|---|---|
| BAZ1B | -0.7171446 | 1 | 9 |
| ELOB | 0.60620241 | 1 | 9 |
| NME2 | 0.63289964 | 1 | 9 |
| FTH1 | 0.447896 | 1 | 9 |
| ATXN7L3B | -0.6431884 | 1 | 9 |
| PGLS | 0.61813276 | 1 | 9 |
| HELZ | -0.6452632 | 1 | 9 |
| REST | -0.6562874 | 1 | 9 |
| MT-ND4 | -0.4809564 | 1 | 9 |
| COX6A1 | 0.48205323 | 1 | 9 |
| CHD6 | -0.6275671 | 1 | 9 |
| LSM7 | 0.48503848 | 1 | 9 |
| CD7 | 0.54078785 | 1 | 9 |
| DNAJB14 | -0.6177939 | 1 | 9 |
| MAN1A2 | -0.6611884 | 1 | 9 |
| ACAP2 | -0.6291083 | 1 | 9 |
| H3F3A | 0.3481403 | 1 | 9 |
| SLFN5 | -0.7555901 | 1 | 9 |
| MZT2B | 0.57249104 | 1 | 9 |
| ATP5F1D | 0.6500074 | 1 | 9 |
| ANKRD11 | -0.7446123 | 1 | 9 |
| CYBA | 0.60675445 | 1 | 9 |
| PRRC2C | -0.5777406 | 1 | 9 |
| PIK3R1 | -0.6650331 | 1 | 9 |
| RBL2 | -0.6654058 | 1 | 9 |
| AKNA | -0.6504901 | 1 | 9 |
| RAN | 0.45539337 | 1 | 9 |
| NDUFA13 | 0.52757868 | 1 | 9 |
| TTC14 | -0.7074001 | 1 | 9 |
| RPL4 | 0.35968152 | 1 | 9 |
| HNRNPD | -0.6284497 | 1 | 9 |
| RCSD1 | -0.646363 | 1 | 9 |
| GSTK1 | 0.48062696 | 1 | 9 |
| AL365361.1 | -0.7083708 | 1 | 9 |
| SRSF11 | -0.5656503 | 1 | 9 |
| ATF7IP | -0.6739763 | 1 | 9 |
| SNRNP200 | -0.6246476 | 1 | 9 |
| PRKAR1A | -0.602253 | 1 | 9 |
| BPTF | -0.6642943 | 1 | 9 |
| BOD1L1 | -0.6606514 | 1 | 9 |
| DYNLL1 | 0.48291997 | 1 | 9 |
| TRAF5 | -0.6858569 | 1 | 9 |
| PHIP | -0.7624291 | 1 | 9 |

| | | | |
|---|---|---|---|
| IKZF1 | -0.6257867 | 1 | 9 |
| RUFY2 | -0.6201964 | 1 | 9 |
| ALOX5AP | 0.53761983 | 1 | 9 |
| CEP85L | -0.6589245 | 1 | 9 |
| NDUFA4 | 0.54872454 | 1 | 9 |
| MBNL1 | -0.6545344 | 1 | 9 |
| PCSK7 | -0.7156187 | 1 | 9 |
| TOP2B | -0.5893753 | 1 | 9 |
| LCP2 | -0.6460022 | 1 | 9 |
| CTR9 | -0.566783 | 1 | 9 |
| VPS36 | -0.5901948 | 1 | 9 |
| TRIM24 | -0.5337328 | 1 | 9 |
| UQCRB | 0.45936131 | 1 | 9 |
| ZRANB2 | -0.6420192 | 1 | 9 |
| ERAP2 | -0.6347131 | 1 | 9 |
| ERV3-1 | -0.6049336 | 1 | 9 |
| TPR | -0.6207826 | 1 | 9 |
| CLDND1 | -0.660063 | 1 | 9 |
| NPEPPS | -0.5700514 | 1 | 9 |
| IKZF2 | -0.9301583 | 1 | 9 |
| LIME1 | 0.49922791 | 1 | 9 |
| RPS29 | 0.35120084 | 1 | 9 |
| ABI1 | -0.6051606 | 1 | 9 |
| PSIP1 | -0.6107587 | 1 | 9 |
| BLOC1S1 | 0.47277846 | 1 | 9 |
| CD2AP | -0.5839878 | 1 | 9 |
| EMB | -0.6244533 | 1 | 9 |
| MYCBP2 | -0.6652594 | 1 | 9 |
| SSR2 | 0.49064357 | 1 | 9 |
| TNRC6C | -0.7365263 | 1 | 9 |
| UBL5 | 0.52549566 | 1 | 9 |
| GTF2H5 | 0.4685291 | 1 | 9 |
| R3HDM2 | -0.5713051 | 1 | 9 |
| URI1 | -0.597616 | 1 | 9 |
| TTC39B | -0.5567517 | 1 | 9 |
| RIF1 | -0.6890008 | 1 | 9 |
| SAMD9 | -0.6668793 | 1 | 9 |
| UBXN7 | -0.5464153 | 1 | 9 |
| KAT6A | -0.6237695 | 1 | 9 |
| FAM89B | 0.47918381 | 1 | 9 |
| HLTF | -0.5219339 | 1 | 9 |
| PHF3 | -0.6068752 | 1 | 9 |
| SLC25A6 | 0.46251007 | 1 | 9 |

| | | | |
|---|---|---|---|
| CSNK1A1 | -0.5721658 | 1 | 9 |
| PAG1 | -0.5578514 | 1 | 9 |
| TTC3 | -0.681244 | 1 | 9 |
| BRD2 | -0.4766038 | 1 | 9 |
| GIMAP4 | 0.28695143 | 1 | 9 |
| POLR2B | -0.5133427 | 1 | 9 |
| NUFIP2 | -0.6288491 | 1 | 9 |
| CLINT1 | -0.560131 | 1 | 9 |
| UBE2I | -0.5199097 | 1 | 9 |
| LRRC75A | 0.48613657 | 1 | 9 |
| CD226 | -0.6289572 | 1 | 9 |
| EFCAB14 | -0.5483623 | 1 | 9 |
| CNTRL | -0.6177939 | 1 | 9 |
| TALDO1 | 0.56216106 | 1 | 9 |
| GNG5 | 0.41849396 | 1 | 9 |
| RRP1B | -0.5229993 | 1 | 9 |
| ACTR2 | -0.5537609 | 1 | 9 |
| SUPT20H | -0.5465026 | 1 | 9 |
| PPP4R2 | -0.5761133 | 1 | 9 |
| FAM120A | -0.5557772 | 1 | 9 |
| AFF4 | -0.5300871 | 1 | 9 |
| CASK | -0.5794503 | 1 | 9 |
| SSR4 | 0.48692345 | 1 | 9 |
| TAOK1 | -0.5835082 | 1 | 9 |
| TPM4 | -0.6239938 | 1 | 9 |
| TBC1D10C | 0.57591713 | 1 | 9 |
| NOSIP | 0.51148912 | 1 | 9 |
| EIF3A | -0.6319852 | 1 | 9 |
| OSBPL8 | -0.5841855 | 1 | 9 |
| DLEU2 | -0.6641495 | 1 | 9 |
| TAOK3 | -0.5720211 | 1 | 9 |
| PTBP3 | -0.5685707 | 1 | 9 |
| SENP6 | -0.5686973 | 1 | 9 |
| MYL6B | 0.49991242 | 1 | 9 |
| BAZ2A | -0.5260994 | 1 | 9 |
| TRIP12 | -0.5622508 | 1 | 9 |
| SECISBP2 | -0.5564902 | 1 | 9 |
| ZKSCAN1 | -0.6320763 | 1 | 9 |
| MAP4K4 | -0.5561133 | 1 | 9 |
| POM121 | -0.50478 | 1 | 9 |
| ANKLE2 | -0.4927313 | 1 | 9 |
| NR2C2 | -0.4813548 | 1 | 9 |
| CCDC88C | -0.5604607 | 1 | 9 |

| | | | |
|---|---|---|---|
| USP22 | -0.5596833 | 1 | 9 |
| COX8A | 0.45976285 | 1 | 9 |
| G3BP1 | -0.5479497 | 1 | 9 |
| SLC16A7 | -0.6587066 | 1 | 9 |
| HNRNPA3 | -0.5257265 | 1 | 9 |
| PPIG | -0.6592705 | 1 | 9 |
| OXR1 | -0.5353708 | 1 | 9 |
| CORO1A | 0.43719572 | 1 | 9 |
| PLAC8 | 0.28516614 | 1 | 9 |
| NDUFA11 | 0.74574502 | 1 | 9 |
| TMEM219 | 0.52091238 | 1 | 9 |
| HSPA8 | 0.32722098 | 1 | 9 |
| CD38 | -0.8290155 | 1 | 9 |
| ARFGEF1 | -0.6280622 | 1 | 9 |
| OGA | -0.596808 | 1 | 9 |
| MBD5 | -0.590449 | 1 | 9 |
| SETD2 | -0.6209932 | 1 | 9 |
| USP8 | -0.6084895 | 1 | 9 |
| TRIM38 | -0.6020952 | 1 | 9 |
| PLCG2 | -0.5956795 | 1 | 9 |
| KDM5B | -0.6729184 | 1 | 9 |
| LIMD2 | 0.42116663 | 1 | 9 |
| TCEA1 | -0.5332202 | 1 | 9 |
| ELF1 | -0.6054455 | 1 | 9 |
| NIPBL | -0.6110154 | 1 | 9 |
| NFATC2IP | -0.5627862 | 1 | 9 |
| ZMYM2 | -0.5878266 | 1 | 9 |
| MYO1F | -0.6124801 | 1 | 9 |
| DDHD1 | -0.5335672 | 1 | 9 |
| SCAF11 | -0.5520745 | 1 | 9 |
| CDV3 | -0.5378038 | 1 | 9 |
| TLN1 | -0.5801352 | 1 | 9 |
| GPATCH8 | -0.5376031 | 1 | 9 |
| PDE7A | -0.6826616 | 1 | 9 |
| VAV3 | -0.5750374 | 1 | 9 |
| ZMYM5 | -0.5444711 | 1 | 9 |
| ESCO1 | -0.5385295 | 1 | 9 |
| USP33 | -0.5232682 | 1 | 9 |
| ARHGEF2 | -0.4689101 | 1 | 9 |
| COPS9 | 0.44355545 | 1 | 9 |
| TRIM14 | -0.6282005 | 1 | 9 |
| KMT2A | -0.604498 | 1 | 9 |
| MLXIP | -0.5916347 | 1 | 9 |

| | | | |
|---|---|---|---|
| RNF125 | -0.6486409 | 1 | 9 |
| LPP | -0.6820362 | 1 | 9 |
| AC002454.1 | 0.38515339 | 1 | 9 |
| JPX | -0.6484586 | 1 | 9 |
| CERS6 | -0.5415883 | 1 | 9 |
| ORAI2 | -0.6069056 | 1 | 9 |
| YWHAZ | -0.408641 | 1 | 9 |
| ZBTB37 | -0.5112486 | 1 | 9 |
| KDM4A | -0.4631167 | 1 | 9 |
| CHCHD2 | 0.3718456 | 1 | 9 |
| ZBTB1 | -0.5657844 | 1 | 9 |
| AKAP9 | -0.6167121 | 1 | 9 |
| SACM1L | -0.4902176 | 1 | 9 |
| DRAP1 | 0.44069731 | 1 | 9 |
| GATAD2B | -0.5002623 | 1 | 9 |
| CEP350 | -0.6281934 | 1 | 9 |
| RNASEH2B | -0.6203283 | 1 | 9 |
| STT3B | -0.6196564 | 1 | 9 |
| PTPN22 | -0.5802365 | 1 | 9 |
| KLF13 | -0.514808 | 1 | 9 |
| EBLN3P | -0.5460924 | 1 | 9 |
| ZNF609 | -0.4546798 | 1 | 9 |
| MSL1 | -0.4969395 | 1 | 9 |
| STRN | -0.4583291 | 1 | 9 |
| RABEP1 | -0.4804621 | 1 | 9 |
| LDLRAD4 | -0.5327573 | 1 | 9 |
| KDM5A | -0.5621375 | 1 | 9 |
| LDHB | 0.3713157 | 1 | 9 |
| UQCR11 | 0.3906547 | 1 | 9 |
| DHRS4L2 | 0.29439692 | 1 | 9 |
| GSK3B | -0.4971146 | 1 | 9 |
| ZNF66 | -0.5008561 | 1 | 9 |
| FAM117A | -0.553621 | 1 | 9 |
| CHD2 | -0.5634328 | 1 | 9 |
| ARL4C | -0.5948557 | 1 | 9 |
| YBX1 | 0.31849032 | 1 | 9 |
| CARD8 | -0.5692903 | 1 | 9 |
| TMOD3 | -0.5419367 | 1 | 9 |
| CDC5L | -0.5355569 | 1 | 9 |
| KRIT1 | -0.4508966 | 1 | 9 |
| TRMT112 | 0.51777742 | 1 | 9 |
| ARL6IP4 | 0.39484614 | 1 | 9 |
| JAK1 | -0.6000541 | 1 | 9 |

| | | | |
|---|---|---|---|
| UBR2 | -0.444645 | 1 | 9 |
| CD63 | 0.40008727 | 1 | 9 |
| HERC1 | -0.5931096 | 1 | 9 |
| CHD8 | -0.5553475 | 1 | 9 |
| DCAF7 | -0.4836085 | 1 | 9 |
| BRAF | -0.5344222 | 1 | 9 |
| SMC2 | -0.6903285 | 1 | 9 |
| FTX | -0.677851 | 1 | 9 |
| YIPF4 | -0.6099812 | 1 | 9 |
| SH2D3C | -0.4997381 | 1 | 9 |
| LRP12 | -0.5744812 | 1 | 9 |
| BIRC6 | -0.5550785 | 1 | 9 |
| CUL3 | -0.476871 | 1 | 9 |
| RNF115 | -0.530814 | 1 | 9 |
| SEM1 | 0.42785024 | 1 | 9 |
| FAM214A | -0.4894195 | 1 | 9 |
| HERC4 | -0.533348 | 1 | 9 |
| CAMK1D | -0.5946574 | 1 | 9 |
| ATXN7 | -0.5369588 | 1 | 9 |
| RPS6KA3 | -0.566251 | 1 | 9 |
| NNT | -0.4517682 | 1 | 9 |
| EDF1 | 0.57627942 | 1 | 9 |
| ATP13A3 | -0.4390182 | 1 | 9 |
| AC092683.1 | -0.5603084 | 1 | 9 |
| CASD1 | -0.4935596 | 1 | 9 |
| AFG3L2 | -0.4372674 | 1 | 9 |
| USP34 | -0.5566993 | 1 | 9 |
| MYH9 | -0.5750894 | 1 | 9 |
| MKLN1 | -0.4128889 | 1 | 9 |
| CENPC | -0.5297789 | 1 | 9 |
| DOCK2 | -0.4667923 | 1 | 9 |
| CD164 | -0.5304081 | 1 | 9 |
| NKG7 | 0.43314767 | 1 | 9 |
| THEMIS | -0.5974087 | 1 | 9 |
| KDM2A | -0.5135713 | 1 | 9 |
| ACAP1 | -0.4314814 | 1 | 9 |
| SENP7 | -0.5687883 | 1 | 9 |
| DGKA | -0.5179236 | 1 | 9 |
| ZC3H18 | -0.5136504 | 1 | 9 |
| PHC3 | -0.4962802 | 1 | 9 |
| PHF20 | -0.5102257 | 1 | 9 |
| SIVA1 | 0.4120079 | 1 | 9 |
| TAX1BP1 | -0.4316769 | 1 | 9 |

| | | | |
|---|---|---|---|
| EXOC4 | -0.5372702 | 1 | 9 |
| ZFP36L2 | -0.7048877 | 1 | 9 |
| STAT3 | -0.4893645 | 1 | 9 |
| GIHCG | 0.36140338 | 1 | 9 |
| SMAD2 | -0.5638816 | 1 | 9 |
| MYL12B | 0.45409512 | 1 | 9 |
| PSME1 | 0.42182149 | 1 | 9 |
| CASC4 | -0.4895352 | 1 | 9 |
| FABP5 | 0.32211379 | 1 | 9 |
| GPBP1 | -0.5413152 | 1 | 9 |
| NCKAP1L | -0.4816219 | 1 | 9 |
| PDCD7 | -0.5167239 | 1 | 9 |
| NDUFB11 | 0.53200896 | 1 | 9 |
| RSRC1 | -0.4992297 | 1 | 9 |
| PITPNB | -0.4326239 | 1 | 9 |
| WNK1 | -0.4359792 | 1 | 9 |
| SUZ12 | -0.5654568 | 1 | 9 |
| ITPKB | -0.5684167 | 1 | 9 |
| FBXL20 | -0.4233812 | 1 | 9 |
| STMN1 | 0.3022788 | 1 | 9 |
| INO80D | -0.4574563 | 1 | 9 |
| NDUFA1 | 0.39985184 | 1 | 9 |
| SNW1 | -0.4770378 | 1 | 9 |
| TIMP1 | 0.33735949 | 1 | 9 |
| NKTR | -0.5590924 | 1 | 9 |
| SRP14 | 0.31817479 | 1 | 9 |
| ETS1 | -0.5239797 | 1 | 9 |
| AKT1 | -0.4129635 | 1 | 9 |
| SEC61B | 0.44055835 | 1 | 9 |
| ATP5MD | 0.50194535 | 1 | 9 |
| PHACTR4 | -0.5075118 | 1 | 9 |
| CELF2 | -0.5944194 | 1 | 9 |
| NUDT8 | 0.43895452 | 1 | 9 |
| PHACTR2 | -0.5461368 | 1 | 9 |
| MT-ND5 | -0.4733018 | 1 | 9 |
| SIKE1 | -0.4222967 | 1 | 9 |
| MFSD6 | -0.4511271 | 1 | 9 |
| EIF3G | 0.38054879 | 1 | 9 |
| ATXN2L | -0.4575164 | 1 | 9 |
| PLEKHA3 | -0.4107484 | 1 | 9 |
| CNOT4 | -0.5098824 | 1 | 9 |
| EPB41 | -0.5951083 | 1 | 9 |
| CDCA7 | -0.5394097 | 1 | 9 |

| | | | |
|---|---|---|---|
| ABRACL | 0.36869321 | 1 | 9 |
| DIS3 | -0.4535504 | 1 | 9 |
| GNPTAB | -0.5244731 | 1 | 9 |
| MAPK1 | -0.4995828 | 1 | 9 |
| ARAP2 | -0.5826514 | 1 | 9 |
| ZNF107 | -0.5801187 | 1 | 9 |
| NEMF | -0.5270078 | 1 | 9 |
| HSPA1A | 0.60769855 | 1 | 9 |
| HSPB1 | 0.53664241 | 1 | 9 |
| ZNF207 | -0.5000002 | 1 | 9 |
| DIP2A | -0.556263 | 1 | 9 |
| Mar 07 | -0.5062222 | 1 | 9 |
| OFD1 | -0.4661486 | 1 | 9 |
| INTS6 | -0.5459326 | 1 | 9 |
| SMARCA5 | -0.485635 | 1 | 9 |
| ARPC5 | -0.4772979 | 1 | 9 |
| USP12 | -0.398004 | 1 | 9 |
| EPM2AIP1 | -0.5325835 | 1 | 9 |
| SAP18 | 0.33059634 | 1 | 9 |
| RBM6 | -0.5072906 | 1 | 9 |
| SERINC3 | -0.4595617 | 1 | 9 |
| UPF3A | -0.4808894 | 1 | 9 |
| TMCO1 | -0.4414139 | 1 | 9 |
| ZNF737 | -0.476663 | 1 | 9 |
| GAS5 | 0.37915006 | 1 | 9 |
| WAC | -0.5128503 | 1 | 9 |
| ZMIZ1 | -0.5000576 | 1 | 9 |
| NCR3 | 0.40547152 | 1 | 9 |
| REV3L | -0.5510582 | 1 | 9 |
| DYNC1H1 | -0.5691546 | 1 | 9 |
| TLK2 | -0.4566694 | 1 | 9 |
| MTPN | -0.4248123 | 1 | 9 |
| RBBP6 | -0.5124038 | 1 | 9 |
| ERGIC1 | -0.4000856 | 1 | 9 |
| MT-ND6 | -0.513035 | 1 | 9 |
| LBR | -0.519509 | 1 | 9 |
| MAP3K2 | -0.5075756 | 1 | 9 |
| MT-CYB | -0.4900843 | 1 | 9 |
| BRD7 | -0.4980466 | 1 | 9 |
| TUT7 | -0.479775 | 1 | 9 |
| EIF2S3 | -0.5005663 | 1 | 9 |
| NAP1L4 | -0.5025934 | 1 | 9 |
| PARK7 | 0.38792016 | 1 | 9 |

| | | | |
|---|---|---|---|
| GOLGB1 | -0.6534178 | 1 | 9 |
| ETHE1 | 0.39962316 | 1 | 9 |
| GPR174 | -0.5673041 | 1 | 9 |
| MOB1A | -0.5105168 | 1 | 9 |
| ACSF3 | -0.4893645 | 1 | 9 |
| CHD1 | -0.5067715 | 1 | 9 |
| SLC5A3 | -0.6232953 | 1 | 9 |
| GTF3A | 0.3453816 | 1 | 9 |
| HOOK3 | -0.4944756 | 1 | 9 |
| TRIR | 0.38782958 | 1 | 9 |
| CCDC14 | -0.5739323 | 1 | 9 |
| PTRHD1 | 0.29211013 | 1 | 9 |
| DNMT3A | -0.5024664 | 1 | 9 |
| RBM5 | -0.530952 | 1 | 9 |
| CEPT1 | -0.3885726 | 1 | 9 |
| CAPG | 0.55545693 | 1 | 9 |
| NPIPB5 | -0.4331564 | 1 | 9 |
| DBI | 0.39331393 | 1 | 9 |
| PRKCQ | -0.437608 | 1 | 9 |
| ST8SIA4 | -0.4970951 | 1 | 9 |
| SPN | -0.5512586 | 1 | 9 |
| N4BP2 | -0.571679 | 1 | 9 |
| SMC5 | -0.4806293 | 1 | 9 |
| SEC11A | 0.39142305 | 1 | 9 |
| CHD4 | -0.5412125 | 1 | 9 |
| NDUFB9 | 0.37714179 | 1 | 9 |
| GPX4 | 0.48824256 | 1 | 9 |
| DOCK8 | -0.455808 | 1 | 9 |
| ODF2L | -0.5559797 | 1 | 9 |
| IFT80 | -0.4514069 | 1 | 9 |
| ZBTB7A | -0.5348045 | 1 | 9 |
| PIKFYVE | -0.4295165 | 1 | 9 |
| VEZT | -0.4701214 | 1 | 9 |
| HNRNPUL2 | -0.4888155 | 1 | 9 |
| NUTM2B-AS1 | -0.497318 | 1 | 9 |
| S100A11 | 0.51437147 | 1 | 9 |
| SAFB2 | -0.3959166 | 1 | 9 |
| PPARA | -0.5452991 | 1 | 9 |
| PAK2 | -0.4998283 | 1 | 9 |
| ISY1 | -0.4012131 | 1 | 9 |
| EP300 | -0.490281 | 1 | 9 |
| RAB6A | -0.4440362 | 1 | 9 |
| GYPC | 0.40454083 | 1 | 9 |

| | | | |
|---|---|---|---|
| NDUFS6 | 0.43335479 | 1 | 9 |
| HNRNPK | -0.3684344 | 1 | 9 |
| AIF1 | 0.42092482 | 1 | 9 |
| PPP1R2 | -0.5286952 | 1 | 9 |
| MCMBP | -0.4123655 | 1 | 9 |
| ZNF91 | -0.4668809 | 1 | 9 |
| TMEM165 | -0.4551605 | 1 | 9 |
| HIST1H1E | -0.6018896 | 1 | 9 |
| PTPRA | -0.5255555 | 1 | 9 |
| DGKE | -0.4492602 | 1 | 9 |
| FAM102A | -0.5191107 | 1 | 9 |
| UBE3A | -0.4758426 | 1 | 9 |
| ELF2 | -0.4293954 | 1 | 9 |
| PIH1D1 | 0.33101696 | 1 | 9 |
| PPP2R5C | -0.5364527 | 1 | 9 |
| ST3GAL1 | -0.4802904 | 1 | 9 |
| ANKZF1 | -0.3668684 | 1 | 9 |
| VCP | -0.4030054 | 1 | 9 |
| PLSCR1 | -0.3885986 | 1 | 9 |
| NUDT1 | 0.2543156 | 1 | 9 |
| UPF1 | -0.3868441 | 1 | 9 |
| ATP6V1F | 0.46991828 | 1 | 9 |
| PNRC2 | -0.5029143 | 1 | 9 |
| BCLAF1 | -0.525741 | 1 | 9 |
| PTCD3 | -0.4100639 | 1 | 9 |
| NUMB | -0.3661785 | 1 | 9 |
| EML4 | -0.5619086 | 1 | 9 |
| MAPRE2 | -0.475064 | 1 | 9 |
| WIPF1 | -0.497111 | 1 | 9 |
| SERINC5 | -0.5687109 | 1 | 9 |
| SORL1 | -0.4481817 | 1 | 9 |
| MTR | -0.4304577 | 1 | 9 |
| HIST1H2AC | -0.5568413 | 1 | 9 |
| RAC2 | 0.33103832 | 1 | 9 |
| MED13L | -0.5509582 | 1 | 9 |
| SLK | -0.419069 | 1 | 9 |
| HDAC2 | -0.5028891 | 1 | 9 |
| INO80E | 0.32166456 | 1 | 9 |
| CEP57 | -0.4825665 | 1 | 9 |
| UBB | 0.30657868 | 1 | 9 |
| VIM | 0.27386951 | 1 | 9 |
| QDPR | 0.26124849 | 1 | 9 |
| ACTL6A | -0.3856732 | 1 | 9 |

| | | | |
|---|---|---|---|
| DGKH | -0.4672675 | 1 | 9 |
| CNOT1 | -0.4570177 | 1 | 9 |
| CMAS | -0.3615448 | 1 | 9 |
| ZNF43 | -0.4114069 | 1 | 9 |
| SMAD4 | -0.4425309 | 1 | 9 |
| HECTD1 | -0.4490239 | 1 | 9 |
| DHFR | -0.4856904 | 1 | 9 |
| ZBTB11 | -0.4148547 | 1 | 9 |
| ADA | 0.27041167 | 1 | 9 |
| MZT2A | 0.46887809 | 1 | 9 |
| KLHL36 | -0.3754268 | 1 | 9 |
| CYFIP2 | -0.4854337 | 1 | 9 |
| RBM12B | -0.4144024 | 1 | 9 |
| DLG1 | -0.5047327 | 1 | 9 |
| RBM26 | -0.4883767 | 1 | 9 |
| TCF7 | -0.5082225 | 1 | 9 |
| ARL17B | -0.3889882 | 1 | 9 |
| FAR1 | -0.4315751 | 1 | 9 |
| SRSF6 | -0.437477 | 1 | 9 |
| DHX36 | -0.4512957 | 1 | 9 |
| SINHCAF | -0.4063217 | 1 | 9 |
| MT-ATP6 | -0.4289716 | 1 | 9 |
| GARS-DT | -0.393597 | 1 | 9 |
| SIN3A | -0.4505563 | 1 | 9 |
| SETX | -0.5078982 | 1 | 9 |
| USP16 | -0.4942831 | 1 | 9 |
| MYO9B | -0.4796693 | 1 | 9 |
| TOPBP1 | -0.3760003 | 1 | 9 |
| B2M | 0.28779227 | 1 | 9 |
| MIS18BP1 | -0.4605333 | 1 | 9 |
| RAD9A | -0.3809802 | 1 | 9 |
| DCAF11 | -0.3948378 | 1 | 9 |
| USP25 | -0.3578477 | 1 | 9 |
| TRAPPC10 | -0.406109 | 1 | 9 |
| AC025171.2 | -0.4721955 | 1 | 9 |
| VIPR2 | -0.5728447 | 1 | 9 |
| ATXN2 | -0.3713067 | 1 | 9 |
| ZNF720 | -0.5072838 | 1 | 9 |
| INPP4A | -0.4882291 | 1 | 9 |
| BBX | -0.4761489 | 1 | 9 |
| SYNE2 | -0.7507079 | 1 | 9 |
| ATR | -0.5028891 | 1 | 9 |
| XRN1 | -0.4217097 | 1 | 9 |

| | | | |
|---|---|---|---|
| NCAPG2 | -0.5523962 | 1 | 9 |
| SOX4 | -0.6219973 | 1 | 9 |
| HELB | -0.4939653 | 1 | 9 |
| SCAPER | -0.3950136 | 1 | 9 |
| STK24 | -0.4057172 | 1 | 9 |
| IREB2 | -0.3707834 | 1 | 9 |
| SMG7 | -0.390886 | 1 | 9 |
| MAVS | -0.4198024 | 1 | 9 |
| HNRNPC | -0.4191237 | 1 | 9 |
| SLC25A36 | -0.4926953 | 1 | 9 |
| VAMP5 | 0.3915903 | 1 | 9 |
| ABCE1 | -0.4149922 | 1 | 9 |
| EXOC1 | -0.3988876 | 1 | 9 |
| TPI1 | 0.38802808 | 1 | 9 |
| HBS1L | -0.4284165 | 1 | 9 |
| IK | -0.4443503 | 1 | 9 |
| MDH2 | 0.46788531 | 1 | 9 |
| DOCK10 | -0.5562879 | 1 | 9 |
| CLIP1 | -0.4321246 | 1 | 9 |
| MIR181A1HG | -0.5515602 | 1 | 9 |
| ZNF652 | -0.4225486 | 1 | 9 |
| FBXW7 | -0.3950371 | 1 | 9 |
| KDM4C | -0.4056197 | 1 | 9 |
| PLEKHB2 | -0.389189 | 1 | 9 |
| GSE1 | -0.4848431 | 1 | 9 |
| YWHAB | -0.3515544 | 1 | 9 |
| RBM48 | -0.336299 | 1 | 9 |
| RAD50 | -0.5059054 | 1 | 9 |
| ATP5F1C | 0.46205941 | 1 | 9 |
| NEDD8 | 0.471332 | 1 | 9 |
| CHDH | -0.6823442 | 1 | 9 |
| ZNF708 | -0.4671372 | 1 | 9 |
| MRE11 | -0.3817635 | 1 | 9 |
| TGFBR2 | -0.4395458 | 1 | 9 |
| KCTD20 | -0.3882454 | 1 | 9 |
| MBTPS1 | -0.4480015 | 1 | 9 |
| FLI1 | -0.4248123 | 1 | 9 |
| YY1 | -0.4753687 | 1 | 9 |
| TCOF1 | -0.3825463 | 1 | 9 |
| ZC3HAV1 | -0.5169695 | 1 | 9 |
| ZNF302 | -0.4205019 | 1 | 9 |
| CERT1 | -0.428001 | 1 | 9 |
| DHX38 | -0.3662156 | 1 | 9 |

| | | | |
|---|---|---|---|
| SUPT6H | -0.401763 | 1 | 9 |
| CLEC11A | 0.33357133 | 1 | 9 |
| CNOT6L | -0.4501288 | 1 | 9 |
| DICER1 | -0.4319761 | 1 | 9 |
| SF3B2 | -0.3610934 | 1 | 9 |
| MPC2 | 0.34809902 | 1 | 9 |
| AKIRIN1 | -0.3929573 | 1 | 9 |
| STAG2 | -0.5078393 | 1 | 9 |
| MSI2 | -0.5442105 | 1 | 9 |
| EIF4A1 | 0.36143416 | 1 | 9 |
| ZNF638 | -0.4521437 | 1 | 9 |
| S1PR3 | -0.4503806 | 1 | 9 |
| C12orf75 | 0.41149918 | 1 | 9 |
| PCSK5 | -0.4423436 | 1 | 9 |
| NR3C1 | -0.4568745 | 1 | 9 |
| ERICH1 | -0.5099957 | 1 | 9 |
| IPCEF1 | -0.4546346 | 1 | 9 |
| KAT2B | -0.33421 | 1 | 9 |
| FRYL | -0.3628223 | 1 | 9 |
| DPYSL2 | -0.373912 | 1 | 9 |
| PSMA3-AS1 | -0.4817114 | 1 | 9 |
| NHLRC2 | -0.3687856 | 1 | 9 |
| RSRP1 | -0.4185684 | 1 | 9 |
| LCP1 | -0.4375168 | 1 | 9 |
| PRKD2 | -0.3448862 | 1 | 9 |
| RORB | -0.5503883 | 1 | 9 |
| RGS18 | 0.27711285 | 1 | 9 |
| PRDX6 | 0.4304052 | 1 | 9 |
| PHYKPL | -0.4391819 | 1 | 9 |
| DNAJC10 | -0.4240778 | 1 | 9 |
| TFAM | -0.398661 | 1 | 9 |
| VPS13B | -0.3715244 | 1 | 9 |
| EIF5B | -0.4620098 | 1 | 9 |
| SNRPG | 0.48029268 | 1 | 9 |
| PRMT2 | -0.5541728 | 1 | 9 |
| DCAF6 | -0.3316991 | 1 | 9 |
| RASGRP1 | -0.5382266 | 1 | 9 |
| STK17A | -0.5281542 | 1 | 9 |
| UBA1 | -0.424644 | 1 | 9 |
| SEC16A | -0.3144237 | 1 | 9 |
| RELCH | -0.3369252 | 1 | 9 |
| TRAPPC1 | 0.29488262 | 1 | 9 |
| RIOK3 | -0.4014002 | 1 | 9 |

| | | | |
|---|---|---|---|
| TRIM56 | -0.521333 | 1 | 9 |
| EPC2 | -0.347695 | 1 | 9 |
| UGCG | -0.3306517 | 1 | 9 |
| HUWE1 | -0.4233484 | 1 | 9 |
| RALY | 0.41764837 | 1 | 9 |
| KTN1 | -0.4901729 | 1 | 9 |
| FAM111A | -0.5059379 | 1 | 9 |
| SAMD12 | -0.4727135 | 1 | 9 |
| NORAD | -0.4676979 | 1 | 9 |
| CBX5 | -0.4793692 | 1 | 9 |
| PHF20L1 | -0.4818113 | 1 | 9 |
| NDUFB4 | 0.35058635 | 1 | 9 |
| ZEB1 | -0.4458483 | 1 | 9 |
| DNAJB1 | -0.4056474 | 1 | 9 |
| LATS1 | -0.3598697 | 1 | 9 |
| UQCRFS1 | 0.33456801 | 1 | 9 |
| SMIM26 | 0.38250483 | 1 | 9 |
| ZFP36L1 | -0.6177939 | 1 | 9 |
| RAB27A | -0.4200061 | 1 | 9 |
| DDX42 | -0.4089063 | 1 | 9 |
| USP37 | -0.3283445 | 1 | 9 |
| SNX1 | -0.3805707 | 1 | 9 |
| VPS4B | -0.4386293 | 1 | 9 |
| MECP2 | -0.3568705 | 1 | 9 |
| AGO2 | -0.4224697 | 1 | 9 |
| MRPS18A | 0.25088508 | 1 | 9 |
| BIRC2 | -0.4746373 | 1 | 9 |
| PTPN4 | -0.3656218 | 1 | 9 |
| MCRIP1 | 0.31882017 | 1 | 9 |
| RTF1 | -0.4609483 | 1 | 9 |
| RECQL | -0.4626929 | 1 | 9 |
| U2SURP | -0.3949107 | 1 | 9 |
| SNRPB | 0.33891454 | 1 | 9 |
| PARP14 | -0.442541 | 1 | 9 |
| SERINC1 | -0.4413287 | 1 | 9 |
| MGA | -0.4086381 | 1 | 9 |
| DCLRE1C | -0.4014002 | 1 | 9 |
| LARP4 | -0.3713449 | 1 | 9 |
| PARP4 | -0.3697148 | 1 | 9 |
| BTBD7 | -0.3532774 | 1 | 9 |
| SET | -0.3736251 | 1 | 9 |
| MOSPD2 | -0.318445 | 1 | 9 |
| ANO6 | -0.3672046 | 1 | 9 |

| | | | |
|---|---|---|---|
| TIAM1 | -0.4420782 | 1 | 9 |
| ZMYM4 | -0.4173887 | 1 | 9 |
| MIEN1 | 0.42151131 | 1 | 9 |
| EEA1 | -0.4436159 | 1 | 9 |
| R3HDM1 | -0.421274 | 1 | 9 |
| ZNF22 | -0.3788765 | 1 | 9 |
| ZSCAN30 | -0.3423402 | 1 | 9 |
| CTDSPL2 | -0.4131488 | 1 | 9 |
| INPP5B | -0.3462913 | 1 | 9 |
| ATG10 | -0.3120904 | 1 | 9 |
| GABPB1-IT1 | -0.378328 | 1 | 9 |
| ZFR | -0.3883713 | 1 | 9 |
| RBM33 | -0.4698743 | 1 | 9 |
| ZDHHC21 | -0.3935217 | 1 | 9 |
| NPAT | -0.3536954 | 1 | 9 |
| MFSD14C | -0.3980194 | 1 | 9 |
| PLEKHG1 | -0.5410789 | 1 | 9 |
| TARDBP | -0.4062145 | 1 | 9 |
| TTC17 | -0.4338711 | 1 | 9 |
| SNRPF | 0.2911108 | 1 | 9 |
| HSD17B11 | -0.3438985 | 1 | 9 |
| XIAP | -0.4264215 | 1 | 9 |
| PAN3 | -0.3006423 | 1 | 9 |
| AURKAIP1 | 0.33008836 | 1 | 9 |
| GOLIM4 | -0.3015146 | 1 | 9 |
| TRIM27 | -0.3626786 | 1 | 9 |
| NAP1L1 | -0.4113747 | 1 | 9 |
| CD46 | -0.4303736 | 1 | 9 |
| PTPN1 | -0.387181 | 1 | 9 |
| MRPL27 | 0.31160798 | 1 | 9 |
| ZBED5 | -0.3636239 | 1 | 9 |
| HTATSF1 | -0.3918279 | 1 | 9 |
| POLR2H | 0.29154621 | 1 | 9 |
| SCAI | -0.4559219 | 1 | 9 |
| SNRPN | 0.38808618 | 1 | 9 |
| TAB2 | -0.3630971 | 1 | 9 |
| MFSD1 | -0.3159065 | 1 | 9 |
| ETNK1 | -0.4431546 | 1 | 9 |
| SRPRA | -0.3544374 | 1 | 9 |
| CCSER2 | -0.457878 | 1 | 9 |
| STX16 | -0.4620194 | 1 | 9 |
| REXO4 | -0.3426755 | 1 | 9 |
| MAP3K4 | -0.3480958 | 1 | 9 |

| | | | |
|---|---|---|---|
| ICAM3 | 0.46520031 | 1 | 9 |
| KPNA5 | -0.3635388 | 1 | 9 |
| BRWD1 | -0.4471028 | 1 | 9 |
| STRADA | -0.3370329 | 1 | 9 |
| GNA13 | -0.4006958 | 1 | 9 |
| SLC35E2B | -0.3448862 | 1 | 9 |
| EIF4G1 | -0.4070234 | 1 | 9 |
| MIER1 | -0.3693466 | 1 | 9 |
| CPLANE1 | -0.3965281 | 1 | 9 |
| FBXL5 | -0.372265 | 1 | 9 |
| TMEM131L | -0.4897289 | 1 | 9 |
| MBD4 | -0.3720027 | 1 | 9 |
| CD84 | -0.5778739 | 1 | 9 |
| PRKCA | -0.3999029 | 1 | 9 |
| SLC25A24 | -0.3698161 | 1 | 9 |
| TMEM41B | -0.3688597 | 1 | 9 |
| BCL9L | -0.3463034 | 1 | 9 |
| MRPL52 | 0.37921744 | 1 | 9 |
| ZNF138 | -0.4392126 | 1 | 9 |
| TTC37 | -0.4297215 | 1 | 9 |
| SRGAP3 | -0.3833286 | 1 | 9 |
| RMND5A | -0.3911013 | 1 | 9 |
| COG5 | -0.3609934 | 1 | 9 |
| COX5A | 0.4273222 | 1 | 9 |
| CFLAR | -0.5005767 | 1 | 9 |
| CDK11B | -0.3555965 | 1 | 9 |
| PXK | -0.352697 | 1 | 9 |
| TRIM73 | -0.3815254 | 1 | 9 |
| CLIP2 | -0.3061341 | 1 | 9 |
| ADSS | -0.3557896 | 1 | 9 |
| AL390728.6 | -0.3680066 | 1 | 9 |
| DNAJC13 | -0.299725 | 1 | 9 |
| REX1BD | 0.39016793 | 1 | 9 |
| HELLS | -0.656235 | 1 | 9 |
| TCAF1 | -0.4417373 | 1 | 9 |
| FUBP1 | -0.4603651 | 1 | 9 |
| UFC1 | 0.38559989 | 1 | 9 |
| ATP11B | -0.3783463 | 1 | 9 |
| IARS2 | -0.3680941 | 1 | 9 |
| RUFY3 | -0.4843081 | 1 | 9 |
| PCF11 | -0.4104322 | 1 | 9 |
| KLC1 | -0.406848 | 1 | 9 |
| PWWP2A | -0.3596461 | 1 | 9 |

| | | | |
|---|---|---|---|
| REV1 | -0.3743979 | 1 | 9 |
| HIST1H1C | -0.7152318 | 1 | 9 |
| TUT4 | -0.4572737 | 1 | 9 |
| KIAA1109 | -0.4051581 | 1 | 9 |
| DIDO1 | -0.3290762 | 1 | 9 |
| GMCL1 | -0.4430396 | 1 | 9 |
| PAXBP1 | -0.4401894 | 1 | 9 |
| GALNT1 | -0.3607582 | 1 | 9 |
| PPP1R15B | -0.3315708 | 1 | 9 |
| SFXN1 | -0.4324908 | 1 | 9 |
| CCDC144A | -0.3258234 | 1 | 9 |
| MRPS34 | 0.3537011 | 1 | 9 |
| FBLN2 | -0.3742072 | 1 | 9 |
| KANSL1L | -0.3329551 | 1 | 9 |
| ZNF445 | -0.3484965 | 1 | 9 |
| CYLD | -0.4349825 | 1 | 9 |
| NRDC | -0.3982461 | 1 | 9 |
| SMC1A | -0.5521811 | 1 | 9 |
| NDUFB7 | 0.39943895 | 1 | 9 |
| NUDT3 | -0.3416695 | 1 | 9 |
| HP1BP3 | -0.4279839 | 1 | 9 |
| PPP1R12A | -0.4483164 | 1 | 9 |
| GLTP | -0.2919963 | 1 | 9 |
| TOP3A | -0.2975823 | 1 | 9 |
| AUTS2 | -0.423936 | 1 | 9 |
| NOL8 | -0.3893777 | 1 | 9 |
| CBX4 | -0.379466 | 1 | 9 |
| ZNF680 | -0.4129935 | 1 | 9 |
| SETD5 | -0.40254 | 1 | 9 |
| PPP1R10 | -0.3447369 | 1 | 9 |
| C4orf3 | 0.38466176 | 1 | 9 |
| VPS45 | -0.301223 | 1 | 9 |
| SPOP | -0.3831503 | 1 | 9 |
| AC060780.1 | -0.3754858 | 1 | 9 |
| RDX | -0.3441122 | 1 | 9 |
| TMEM107 | -0.3258845 | 1 | 9 |
| TAF3 | -0.3577542 | 1 | 9 |
| CHORDC1 | -0.359026 | 1 | 9 |
| PPP6R3 | -0.3687066 | 1 | 9 |
| PI4KB | -0.2842259 | 1 | 9 |
| ARPC1B | 0.32413705 | 1 | 9 |
| DDX24 | -0.4125962 | 1 | 9 |
| PCMTD1 | -0.3949616 | 1 | 9 |

| | | | |
|---|---|---|---|
| MDM1 | -0.3480958 | 1 | 9 |
| SUGP2 | -0.4256136 | 1 | 9 |
| MTDH | -0.4175306 | 1 | 9 |
| CSNK2A1 | -0.4304667 | 1 | 9 |
| RIC8B | -0.2883321 | 1 | 9 |
| POGLUT1 | -0.3108161 | 1 | 9 |
| GABPB1-AS1 | -0.6301738 | 1 | 9 |
| PPFIA1 | -0.3093281 | 1 | 9 |
| ANKHD1 | -0.4458253 | 1 | 9 |
| ZNF721 | -0.4374928 | 1 | 9 |
| PBXIP1 | -0.3947143 | 1 | 9 |
| PPM1K | -0.3633525 | 1 | 9 |
| RO60 | -0.4102031 | 1 | 9 |
| TRANK1 | -0.3098863 | 1 | 9 |
| MFSD4B | -0.3165415 | 1 | 9 |
| COX14 | 0.40759942 | 1 | 9 |
| CCNK | -0.3088833 | 1 | 9 |
| UTRN | -0.4628588 | 1 | 9 |
| MIR646HG | -0.3285544 | 1 | 9 |
| DNM1L | -0.3324795 | 1 | 9 |
| JADE1 | -0.3223311 | 1 | 9 |
| RALGAPB | -0.3207884 | 1 | 9 |
| ANAPC1 | -0.3152644 | 1 | 9 |
| TXLNA | -0.3396743 | 1 | 9 |
| CEBPZ | -0.4019132 | 1 | 9 |
| RAB3IP | -0.4647802 | 1 | 9 |
| LPIN2 | -0.390292 | 1 | 9 |
| TGFBR1 | -0.3189494 | 1 | 9 |
| NBR1 | -0.3508747 | 1 | 9 |
| RBMS1 | -0.448384 | 1 | 9 |
| AP005482.1 | -0.5642574 | 1 | 9 |
| OGT | -0.3699941 | 1 | 9 |
| GPATCH2 | -0.3575672 | 1 | 9 |

**ALL P1 scRNA-seq.**
