## Supplementary Table S10 for "Haplotype-aware single-cell multiomics uncovers functional effects of somatic structural variation"

**Table S10. Top20 significant TFs for Cluster3 and Cluster7 of T-ALL P1 scRNA-seq identified using EnrichR analysis**

**Cluster 3 ChEA 2016 top 20 hits**

| Term | Overlap | P-value | Adjusted P-valu | Old P-value | Old Adjusted P- | Odds Ratio | Combined Scor | Genes |
|---|---|---|---|---|---|---|---|---|
| EKLF 21900194 ChIP-Seq ERYTHROCYTE Mouse | 190/1239 | 7.16E-49 | 4.61E-46 | 0 | 0 | 4.22053819 | 467.879546 | RPL4;TCERG1;NDUFA12;NUDT1;PEBP1;YEATS4;IKZF1;ENO1;PARK7;IPO7;ACTG1;GCSH;FBL;CHCHD2;LGALS1;RPS16;TRIM28;SNRPD2;HERC1;SNRPD1;RUVBL2;MYC;STMN1;MYB; |
| MYC 19030024 ChIP-ChIP MESCs Mouse | 352/3868 | 7.86E-38 | 2.53E-35 | 0 | 0 | 2.54748503 | 217.648584 | RPL4;EIF4A1;RPL5;TCERG1;POP7;TFRC;NUDT1;GPATCH4;SPINT2;ENO1;IKZF2;SMC4;PHB2;HS6ST1;SMC2;GCSH;RPS17;PSMD7;RPS16;TRIM28;SNRPD2;RPL18A;FTH1;STMN1;MYB |
| MYB 26560356 Chip-Seq TH2 Human | 209/2000 | 1.98E-28 | 4.25E-26 | 0 | 0 | 2.67281738 | 170.495681 | EIF4A1;TRAF3IP3;IKZF1;ENO1;SMC2;TCL1A;PSMD7;STMN1;MYB;ARL6IP5;B2M;GLUL;CDK5RAP3;MBNL1;CPT1A;PRKCH;GTPBP4;RUNX1;CARHSP1;MTHFD2;TAGLN2;CD226;CDC4 |
| XRN2 22483619 ChIP-Seq HELA Human | 168/1529 | 5.81E-25 | 9.35E-23 | 0 | 0 | 2.74814229 | 153.360226 | RPL4;EIF4A1;RPL5;TRAF3IP3;TFRC;NDUFA11;HNRNPR;PARK7;SMC4;ACTG1;SMC2;FBL;RPS17;CHCHD2;SNRPD2;MYC;FTH1;MAGOH;RPL35;B2M;GLUL;RPS12;TIPIN;ACTN1;RPL23; |
| TAL1 20887958 ChIP-Seq HPC-7 Mouse | 201/2067 | 2.62E-23 | 3.38E-21 | 0 | 0 | 2.43064 | 126.382716 | RPL4;EIF4A1;LPGAT1;TFRC;LST1;NUDT1;IKZF1;SMC3;VPREB1;RPS17;PSMD7;TRIM28;FTH1;STMN1;B2M;GLUL;RPS12;MBNL1;TIPIN;CPT1A;CLEC11A;CIRBP;EWSR1;CSNK2B;VDAC |
| MYC 18555785 ChIP-Seq MESCs Mouse | 135/1200 | 6.62E-21 | 7.11E-19 | 0 | 0 | 2.75485805 | 128.002352 | RPL4;RPL5;TCERG1;NDUFA13;TFRC;NUDT1;NUCKS1;SPINT2;SMC2;EEF1B2;GCSH;RPS17;PSMD7;RPL18A;SNRPD1;RPL35;CDK5RAP3;RPS12;RPL23;RPS5;RPS6;CIRBP;SCRIB;RPSA; |
| UTX 26944678 Chip-Seq JUKART Human | 190/2000 | 8.35E-21 | 7.68E-19 | 0 | 0 | 2.34257006 | 108.301734 | RPL5;SMARCB1;TRAF3IP3;DGKA;LST1;SIRPG;PEBP1;GIMAP4;NUCKS1;ENO1;PARK7;ETS1;ACTG1;SMC2;FADS2;CCND3;ZFP36;LGALS1;RGS3;RPS16;DPYSL2;FTH1;STMN1;CHEK1; |
| MYC 18358816 ChIP-ChIP MESCs Mouse | 276/3413 | 2.90E-20 | 2.34E-18 | 0 | 0 | 2.03936464 | 91.7442625 | RPL4;EIF4A1;RPL5;TCERG1;SMARCB1;TFRC;SPINT2;ENO1;SMC4;PHB2;GCSH;RPS17;PSMD7;SNRPD2;RPL18A;SNRPD1;STMN1;CHEK1;MAGOH;RPL35;RPS11;GLUL;CDK5RAP3;RPS |
| GATA3 27048872 Chip-Seq THYMUS Human | 183/2000 | 2.85E-18 | 2.04E-16 | 0 | 0 | 2.22647115 | 89.9481293 | ITSN2;FAM49B;TFRC;HHIP;DGKA;SIRPG;MSI2;IKZF1;IKZF2;ETS1;ACTG1;FADS2;VPREB1;ZFP36;FTH1;STMN1;MYB;ARL6IP5;BAHCC1;B2M;GLUL;CDK5RAP3;ZNF683;ST6GAL1;TIPIN; |
| KDM2B 26808549 Chip-Seq HPB-ALL Human | 178/2000 | 1.48E-16 | 9.52E-15 | 0 | 0 | 2.14529895 | 78.196524 | DGKA;HNRNPR;IKZF1;ENO1;PARK7;SMC4;HS6ST1;ACTG1;IPO5;ZFP36;DPYSL2;MYC;STMN1;MYB;HMGN2;BAHCC1;GLUL;DDX17;TIPIN;CPT1A;TPM4;IGFBP5;ACTN1;IGFBP2;CIRBP |
| MYC 19079543 ChIP-ChIP MESCs Mouse | 142/1458 | 2.22E-16 | 1.30E-14 | 0 | 0 | 2.33201498 | 84.0546093 | RPL4;EIF4A1;RPL5;ENO1;ACTG1;EEF1B2;FBL;ZFP36;RPS17;PSMD7;RGS3;RPS16;TRIM28;RPL18A;RPL35;RPS11;HADH;RPS12;DDX17;TIPIN;ACTN1;RPL23;RPS5;RPS6;IGFBP2;RPSA; |
| MYB 26560356 Chip-Seq TH1 Human | 177/2000 | 3.19E-16 | 1.71E-14 | 0 | 0 | 2.12923682 | 75.9761765 | NDUFA12;DGKA;GIMAP4;NUCKS1;IKZF1;ENO1;PARK7;SMC3;ETS1;DOCK10;MT2A;CCND3;ZFP36;LGALS1;SNRPD1;DPYSL2;MYC;STMN1;MYB;ARL6IP5;HMGN2;B2M;GLUL;CPT1A; |
| NELFA 20434984 ChIP-Seq ESCs Mouse | 171/2000 | 2.73E-14 | 1.20E-12 | 0 | 0 | 2.03404968 | 63.5270695 | EIF4A1;NDUFA13;TFRC;NDUFA12;PEBP1;YEATS4;SPINT2;ENO1;SMC3;IPO7;GCSH;ZFP36;CHCHD2;PSMD7;TRIM28;SNRPD2;FTH1;STMN1;CHEK1;COTL1;RPS11;GLUL;RPS12;TIPIN; |
| MAF 26560356 Chip-Seq TH1 Human | 171/2000 | 2.73E-14 | 1.20E-12 | 0 | 0 | 2.03404968 | 63.5270695 | DGKA;GIMAP4;GPATCH4;HNRNPR;YEATS4;NUCKS1;ENO1;PARK7;COX6A1;ETS1;ACTG1;DOCK10;FADS2;FBL;MT2A;ZFP36;LGALS1;RGS3;SESN3;STMN1;CHEK1;MAGOH;ARL6IP5; |
| E2F4 17652178 ChIP-ChIP JURKAT Human | 105/1002 | 2.80E-14 | 1.20E-12 | 0 | 0 | 2.47786281 | 77.3250438 | TCERG1;RIF1;HHIP;GMNN;ICAM3;YEATS4;NUCKS1;MT2A;CHCHD2;CHCHD3;MYC;CHEK1;B2M;CDK5RAP3;RPS12;DDX17;MBNL1;SUPT16H;WDR76;RUNX1;MTHFD1;SMS;SNRPG; |
| E2F7 22180533 ChIP-Seq HELA Human | 19/46 | 9.28E-14 | 3.74E-12 | 0 | 0 | 14.1867562 | 425.715111 | SLBP;DUT;PCNA;MCM7;RIF1;PRKDC;ATAD2;CDCA7;NOLC1;H2AFV;SLC1A5;BRCA1;MSH6;MCM3;E2F2;MCM4;MCM5;MCM6;DTL |
| CREM 20920259 ChIP-Seq GC1-SPG Mouse | 380/5776 | 2.80E-13 | 1.06E-11 | 0 | 0 | 1.65069455 | 47.7132161 | ITSN2;EIF4A1;RPL5;TCERG1;SMARCB1;LPGAT1;TFRC;HHIP;HNRNPR;SPINT2;SMC3;SMC4;TXNDC17;PSMD7;RPS16;TRIM28;SNRPD2;RPL18A;FTH1;STMN1;MYB;CHEK1;MAGOH; |
| E2F1 21310950 ChIP-Seq MCF-7 Human | 112/1145 | 3.75E-13 | 1.34E-11 | 0 | 0 | 2.29663459 | 65.7084056 | SMARCB1;GMNN;YEATS4;NUCKS1;MSI2;SMC3;IPO7;FBL;CHCHD3;SNRPD1;DPYSL2;MYC;CHEK1;CDK5RAP3;HIST1H2AC;DDX17;MBNL1;TIPIN;MIF;PGD;HCFC1;MSH6;ILF3;MTHFD |
| MYC 22102868 ChIP-Seq BL Human | 87/797 | 5.85E-13 | 1.98E-11 | 0 | 0 | 2.56665755 | 72.2971046 | EIF4A1;DAZAP1;HSP90AB1;ARPC1B;GDI2;MYL6B;PARK7;YBX1;CHD2;ETS1;IPO7;LMNB1;LIMD2;SYNCRIP;LYL1;PTBP1;TUBA1B;ZFP36;BCL7A;DPYSL2;COTL1;LRRFIP1;RPS12;KLF13; |
| E2F1 18555785 ChIP-Seq MESCs Mouse | 291/4172 | 1.29E-12 | 4.15E-11 | 0 | 0 | 1.693714 | 46.3684798 | RPL4;EIF4A1;TCERG1;SMARCB1;SPINT2;ENO1;SMC4;SMC2;GCSH;RPS17;PSMD7;RPS16;TRIM28;SNRPD2;RPL18A;SNRPD1;MYC;FTH1;STMN1;MAGOH;RPL35;RPS11;B2M;GLUL; |

**Cluster 7 ChEA 2016 top 20 hits**

| Term | Overlap | P-value | Adjusted P-valu | Odds Ratio | Combined Scor | Genes |
|---|---|---|---|---|---|---|
| FOXM1 25889361 ChIP-Seq OE33 AND U2OS Hum | 289/932 | 1.08E-88 | 6.95E-86 | 5.38062653 | 1089.84533 | TCERG1;ARL6IP1;LRR1;CCNF;IKZF2;SMC3;MKI67;RPL9;DCAF7;CDC20;RASSF1;FTH1;FBXO5;SNRPD3;SEPHS1;GTSE1;IER5;SOX4;TMPO;ARL6IP6;CMC2;KNSTRN;DEPDC1B;DDX39A;SARNP;TXNIP;KIF20A;DNA2;KIF2 |
| E2F4 17652178 ChIP-ChIP JURKAT Human | 271/1002 | 2.16E-68 | 6.95E-66 | 4.36249687 | 679.695594 | VPS29;TCERG1;NUP107;ICAM3;MKI67;MT2A;CHEK1;RTTN;FBXO5;SKP2;GTSE1;B2M;CDK5RAP3;RPS13;TMPO;CDK5RAP2;ARL6IP6;RPS12;DDX17;ESCO2;WDR76;RUNX1;DEPDC1B;KIF20A;RPL28;PRR11;ASF1B;SL |
| MYC 19030024 ChIP-ChIP MESCs Mouse | 617/3868 | 4.92E-60 | 1.05E-57 | 2.49117205 | 340.200348 | RPL5;RPL30;POP7;NUP107;RPL3;TFRC;RPL32;RPL31;RPL34;STMN3;SPINT2;ENO1;RPL8;PHB2;RPL9;RPL6;SMC2;RPL7;PSMD7;STMN1;RPL36;RPL38;RPL37;RPL39;CDK5RAP3;CDK5RAP2;DDX17;WDHD1;RPL21;DD |
| FOXM1 23109430 ChIP-Seq U2OS Human | 113/267 | 1.54E-49 | 2.48E-47 | 8.06295861 | 906.224466 | ARL6IP1;ZMYND8;CCNF;HJURP;NUCKS1;TTF2;MKI67;CDC20;PTTG1;NUSAP1;PIM1;HIST1H2AG;NEK2;FBXO5;KPNA2;TMPO;CDC25B;DEPDC1B;PGP;TXNIP;KIF20A;KIF20B;KPNB1;PIF1;CDCA2;TROAP;CDCA8;HMM |
| EKLF 21900194 ChIP-Seq ERYTHROCYTE Mouse | 270/1239 | 1.18E-47 | 1.52E-45 | 3.23212329 | 349.238012 | TCERG1;RPL3;RPL32;NUDT1;IKZF1;ENO1;MKI67;RPL6;RPL7;CDC20;RPS14;TRIM28;SNRPD2;HERC1;SNRPD1;RPS18;STMN1;MAGOH;RCC1;RPS11;MIF;ASRGL1;NUP93;MYL6;DEPDC1B;UQCRC1;PSME2;RPL24;RPL |
| XRN2 22483619 ChIP-Seq HELA Human | 303/1529 | 1.02E-44 | 1.09E-42 | 2.88816989 | 292.562928 | EIF4A1;RPL5;NUP107;RPL3;TRAF3IP3;TFRC;RPL32;ARL6IP1;RPL31;ZNF292;ZMYND8;EIF4A3;HNRNPU;HNRNPR;NUDT5;RPL8;RPL9;DCAF7;RPL6;SMC2;CDC20;RPS14;RASSF1;SNRPD2;RPS19;RPS18;FTH1;MAGOH |
| MYC 18358816 ChIP-ChIP MESCs Mouse | 504/3413 | 4.67E-37 | 4.29E-35 | 2.11661515 | 177.065132 | VPS29;RPL5;RPL30;NUP107;RPL3;TFRC;RPL32;RPL31;STMN3;SPINT2;ENO1;RPL8;PHB2;RPL9;RPL6;RPL7;PSMD7;STMN1;RPL36;RPL38;RPL37;GTSE1;RPL39;CDK5RAP3;CDK5RAP2;DDX17;WDHD1;RPL21;RPL22; |
| NELFA 20434984 ChIP-Seq ESCs Mouse | 343/2000 | 9.26E-37 | 7.44E-35 | 2.424363 | 201.148818 | EIF4A1;NUP107;RPL3;TFRC;CCNF;HNRNPU;SPINT2;ENO1;SMC3;RPL8;RPL6;CDC20;RPS15;PSMD8;RASSF1;PSMD7;TRIM28;SNRPD2;RPS18;FTH1;PSMD2;STMN1;CHEK1;TRIM24;RCC1;FBXO5;RPL37;RPS11;IER5; |
| E2F1 18555785 ChIP-Seq MESCs Mouse | 569/4172 | 2.03E-32 | 1.45E-30 | 1.94259732 | 141.760021 | PI4K2B;RPL30;RPL3;RPL32;RPL31;SPINT2;ENO1;RPL8;RPL6;SMC2;RPL7;PSMD7;STMN1;PSMD1;RPL37;RPL39;IER5;CDK5RAP3;DDX17;RPL21;FNBP1;DDX11;HCFC1;SIGIRR;NUP93;DEPDC1B;MYL6;RNF125;MTHF |
| MYC 19079543 ChIP-ChIP MESCs Mouse | 263/1458 | 2.38E-31 | 1.53E-29 | 2.50771821 | 176.827814 | EIF4A1;RPL5;RPL30;RPL3;RPL32;RPL31;CCNF;ENO1;RPL8;RPL9;RPL6;RPL7;RPS15;RPS14;PSMD7;TRIM28;RPS19;RPL18A;RPS18;RPL36;RCC1;RPL38;RPL37;RPS11;GTSE1;RPL39;RPS10;RPS13;RPS12;DDX17;RPL21 |
| CREM 20920259 ChIP-Seq GC1-SPG Mouse | 721/5776 | 3.18E-30 | 1.86E-28 | 1.81188163 | 123.064732 | PI4K2B;RPL5;RPL30;NUP107;TES;RPL3;TFRC;RPL31;RPL34;HNRNPU;HNRNPR;SPINT2;SMC3;RPL8;DCAF5;TSEN54;RPL6;DCAF7;RPL7;PSMD8;PSMD7;STMN1;RPL36;PSMD1;RPL38;DLEU2;PRKACB;IER5;CDK5RAP3 |
| MYC 18555785 ChIP-Seq MESCs Mouse | 226/1200 | 1.43E-29 | 7.65E-28 | 2.6135103 | 173.586428 | RPL5;TCERG1;RPL30;RPL3;TFRC;RPL32;ARL6IP1;RPL31;EIF4A3;NUDT1;SPINT2;RPL8;RPL6;RPL7;SMC2;RPS15;RPS14;PSMD7;RPS19;RPL18A;SNRPD1;RPS18;RPL36;RCC1;RPL38;RPL37;CDK5RAP3;RPS13;ARL6IP6; |
| E2F4 21247883 ChIP-Seq LYMPHOBLASTOID Hum | 434/2998 | 3.77E-29 | 1.84E-27 | 2.00271333 | 131.075631 | VPS29;NUP107;HNRNPU;HNRNPR;ENO1;SMC3;PSMD7;STMN1;DLEU2;PRKACB;IER5;WDHD1;PRKCH;ACOT7;RPL21;DDX11;WDR76;SIGIRR;NUP93;MYL6;DDIT4;PSME2;RPL26;ZNF276;CDC42SE1;PIF1;CDCA3; |
| TTF2 22483619 ChIP-Seq HELA Human | 264/1512 | 4.01E-29 | 1.84E-27 | 2.4044636 | 157.219385 | EIF4A1;VPS29;RPL5;RPL3;TFRC;RPL32;ARL6IP1;RPL31;ZNF292;RPL34;STMN3;ENO1;NUDT5;RPL9;RPL6;RPL7;RPS15;RPS14;RPS18;STMN1;RPL36;PSMD1;RCC1;RPL38;RTTN;SNRPD3;RPL37;RPS11;PRKACB;RPL39; |
| TAL1 20887958 ChIP-Seq HPC-7 Mouse | 329/2067 | 8.48E-29 | 3.64E-27 | 2.18460533 | 141.206916 | EIF4A1;LPGAT1;RPL3;TFRC;RPL31;LST1;NUDT1;IKZF1;SMC3;RPL8;RPL6;CDC20;RPS15;RPS14;VPREB1;PSMD7;TRIM28;RPS19;RPS18;FTH1;PSMD2;STMN1;RPL36;RCC1;RPS10;IER5;B2M;GLUL;SOX4;RPS12;ARL6IP |
| VDR 23849224 ChIP-Seq CD4+ Human | 346/2231 | 3.35E-28 | 1.35E-26 | 2.12471011 | 134.413373 | VPS29;TCERG1;EIF4A1;RPL5;NUP107;RPL3;TFRC;RPL32;RPL31;ZMYND8;GCC2;DCAF5;DCAF7;SMC2;RPS15;PSMD8;RPS14;HERC1;RPS19;SNRPD1;DPYSL2;CDC27;RPL36;ARL6IP5;PSMD1;RPL38;RTTN;SNRPD3; |
| MYB 26560356 Chip-Seq TH2 Human | 319/2000 | 4.98E-28 | 1.88E-26 | 2.18233195 | 137.195464 | EIF4A1;TES;TRAF3IP3;ZNF292;IKZF1;ENO1;AQP3;SMC2;CCAR1;RPL7;CDC20;RASSF1;TCL1A;PSMD7;STMN1;TBC1D10C;ARL6IP5;DLEU2;IER5;B2M;GLUL;CDK5RAP3;SOX4;TMPO;PTGDR;PRKCH;ATP6V0E1;ACOT7; |
| KDM5B 21448134 ChIP-Seq MESCs Mouse | 504/3724 | 3.70E-27 | 1.32E-25 | 1.87339685 | 114.015385 | VPS29;RPL5;NUP107;TFRC;RPL31;HNRNPU;HNRNPR;SMC3;SMC2;VPREB1;PSMD7;DPYSL2;PSMD2;PSMD1;IER5;CDK5RAP2;DDX17;WDHD1;DDX11;HCFC1;NUP93;EWSR1;PARD3;SARNP;UQCRC1;PRR11;SKAP1; |
| E2F1 21310950 ChIP-Seq MCF-7 Human | 203/1145 | 3.86E-23 | 1.26E-21 | 2.39583299 | 123.646054 | VPS29;SMARCB1;SMC3;SNRPD1;DPYSL2;CHEK1;FBXO5;CDK5RAP3;SOX4;TMPO;ARL6IP6;DDX17;SFMBT1;DDX11;MIF;HCFC1;RNF125;DHX9;PRKDC;CDCA5;CDCA7;ZBTB1;MRPL12;EPB41L2;ZNF704;HIST1H3F; |
| UTX 26944678 Chip-Seq JUKART Human | 304/2000 | 3.92E-23 | 1.26E-21 | 2.03822214 | 105.161247 | RPL5;SMARCB1;TRAF3IP3;LST1;HNRNPU;STMN3;ENO1;TSEN54;SMC2;DPYSL2;FTH1;STMN1;CHEK1;TBC1D10C;RPL36;RCC1;FBXO5;SNRPD3;DLEU2;ZNF720;GLUL;SOX4;ARL6IP6;PTGDR;ACOT7;NUP210;ACTN1; |
